# Effects of polystyrene and polylactide nanoparticles on macrophages under a repeated exposure mode

**DOI:** 10.64898/2026.03.20.713103

**Authors:** Véronique Collin-Faure, Marianne Vitipon, Hélène Diemer, Sarah Cianférani, Daphna Fenel, Elisabeth Darrouzet, Thierry Rabilloud

## Abstract

Micro and nanoplastics are pollutants which concentration in different biotopes increases continuously over time, which poses the question of their potential effects on health. In animals, these micro and nanoplastics are recognized as particulate materials and thus handled by macrophages, which are therefore a key cell type to study. Most studies have used an experimental scheme in which the cells are exposed to a single dose of plastics, with a readout made immediately after exposure. However, this classical experimental scheme does not take into account the impact of biopersistence, nor the potential cellular adaptation that may take place when cells are exposed repeatedly to a low dose of plastics. We thus used a repeated exposure scheme, in order to better take into account these phenomena. Within this frame, we compared the macrophages responses to a persistent nanoplastic, i.e. polystyrene nanoparticles and to a biodegradable nanoplastic, i.e. polylactide, by a combination of proteomic and targeted experiments. Our results show that under this repeated exposure scheme, the proteome changes were of a lesser (for PS) or similar (for PLA) extent than under the acute exposure mode, indicating cell adaptation. However, PLA particles induced mitochondrial dysfunction and depression of response to bacterial molecules perceived as danger signals, such as lipopolysaccharide. Polystyrene nanoparticles also induced a slight alteration of the immune functions of macrophages. This indicates harmful effects even in the repeated exposure scheme.

## 1. Introduction

Our modern societies can be described as plastic-addicts, as the plastic production is steadily increasing ^1^ and is now close to 500Mt/year worldwide ^2^. Indeed, plastic production in the 21st century consumes slightly less than 10% of global oil consumption ^3^, which represents billion of tons/year. Unfortunately, the poor waste handling, at the global scale ^1^ results in an important flow of plastics released in the environment, the sole release into oceans being estimated to 10 Mt/year ^4^ , leading to an estimated accumulation in the oceans of 150 Mt in 2025 ^5^. Although plastic pollution was first documented in marine environments ^6–9^, it has now been documented in marine sediments ^10,11^ , freshwater environments ^12–14^ and terrestrial ones ^15,16^ .

Part of problem lies in plastics’ chemical resistance, which in turn translates into very long persistence in the environment, amounting to decades ^17^ . This does not mean that plastics do not degrade. They fragment first into microplastics (less than 1mm in size) and then into nanoplastics (less than 1µm in size). From what we know from other types of particles, it can be assumed that the smaller the plastic particles, the more easily they will cross the biological barriers and the heavier their interference with biological functions will be. This hypothesis has been documented for at least the intestinal barrier (e.g. in ^18^). In animals, once the barriers have been crossed, a second line of defense comes into play, represented by the phagocytes. This cell types is encountered in invertebrates, from the worms’ coelomocytes to the insects’ hemocytes ^19^. It also occurs in vertebrates in the form of macrophages and neutrophils, and it has been demonstrated that these cells respond to nanoplastics ^20–24^ .

Part of the paradox is our global use of plastic is that close to half of plastic production ends in single use items ^25^. In this case the durability of plastics clearly becomes a negative aspect, especially in the light of the poor waste management worldwide ^1^. Thus, at least for single use applications, the development of biodegradable plastics, which will not release such long-lasting particles, is a promising and highly active avenue of research. Among these biodegradable plastics and beside the ones based on polysaccharides, poly aliphatic hydroxyesters such as polylactide (PLA) or poly hydroxy butyrate/valerate have been extensively proposed. PLA uses a single biosourced monomer, which is polymerized by ring opening, which allows a precise control of polymerization. It has been first introduced for biological sutures ^26,26^ , and has been described as biodegradable ^27^ . However, it must be noted that PLA biodegradation varies widely with the biological conditions. While PLA degrades rapidly within cellular lysosomes ^28^ , which explains its use in drug delivery ^29^, its degradation under composting conditions is highly dependent upon temperature and humidity ^30–32^ , leading to good degradation in industrial, controlled conditions but very slow degradation under home composting conditions . In the same trend, PLA degradation in water, e.g. seawater, is much slower than the one of other aliphatic polyesters ^32^.

However, it has been shown that PLA degradation leads to nanoparticles ^33^, even in conditions reproducing domestic use ^34^ . This further increases the probability that living organisms may encounter PLA nanoparticles, and especially if the use of this plastic increases.

Nevertheless, even if the use of biodegradable plastics increases in the future, we are also facing a very important burden of non-degradable plastics that will be present in the environment for decades, so that their effects are also worth investigating. We thus decided to investigate macrophages responses to PLA and polystyrene (PS) nanoparticles, using a combination of proteomic and targeted approaches. Proteomics allows a wide appraisal of the cellular responses, highlighting phenomena that are further investigated using targeted approaches ^28,35^ . However, in contrast with our previous work and to work published by other teams (e.g. in ^22–24^), we decided to use a repeated exposure scheme. Repeated exposure better mimics environmental or occupational exposures, i.e. low doses over extended periods of time. Furthermore, it has been recently shown that repeated exposure to nanoplastics induces responses that were not detected with a classical, short term and one dose exposure ^36–38^ .

## 2. Materials and Methods

Most of the experiments were carried out as previously described ^39^. However, detailed methods are provided here for the consistency of the paper.

### 2.1. Particles

PS particles (spherical, 200 nm nominal diameter, #AB210806) were purchased from ABCR (Karlsruhe, Germany). Yellowgreen fluorescent particles (spherical, 200 nm nominal diameter, #17151-10) were purchased from Polysciences (Hirschberg an der Bergstrasse, Germany). PLA particles (spherical, 250 nm nominal diameter, #11-00-252) and their green fluorescent labelled counterparts (#51-00-252) were purchased from Micromod (Rostock, Germany).

Infrared (IR) spectra were recorded with a PerkinElmer Spectrum 100 FT-IR spectrometer equipped with a Pike MIRacle attenuated total reflectance (ATR) module, in the 4000-600 cm-1 range. For the measurements a Ge crystal was mounted on the ATR module. In brief, 500µl of PS or PLA nanoparticles dispersion (10 mg/ml) were diluted in 1 ml EtOH and centrifuged at 15000g for 1 hour to pellet the beads. After centrifugation, all but 300 µl of the supernatant was removed and the beads dispersed in the remaining EtOH. 2.5 µL of the resulting suspension was drop-casted on the Ge crystal. The spectra were recorded after complete evaporation of the solvents, under mild pressure (built-in ATR press)

PLA and PS particle primary diameters were assessed from transmission electron microscopy (TEM) images, obtained using either stain on grid or negative staining on a grid techniques. For the stain on grid technique, 10 µL of a 1mg/ml PS or PLA particles were added to a glow discharge grid coated with a carbon supporting film for 5 minutes. The excess solution was soaked off using a filter paper and the grid was air-dried. The images were taken under low dose conditions (<10 e-/Å2) with defocus values comprised between 1.2 and 2.5 mm on a Tecnai 12 LaB6 electron microscope at 120 kV accelerating voltage using a 4k x 4k CEMOS TVIPS F416 camera. For the negative staining technique, samples of 10 mg/mL PLA or PS particles were deposited on the clean side of a carbon film on mica, stained with 1% phosphotungstic acid, before being transferred onto a 400-mesh copper grid. Images were acquired with defocus values comprised between 1.2 and 2.5 mm on a Tecnai 12 LaB6 electron microscope equipped with a Gatan Orius 1000 CCD camera at an accelerating voltage of 120 kV.

### 2.2. Cell culture and exposure to particles

The J774A.1 cell line (female mouse macrophages, RRID CVCL_0358) was purchased from European cell culture collection (Salisbury, UK). Cells were routinely propagated in DMEM supplemented with 10% fetal bovine serum (FBS) in non-adherent flasks (Cellstar flasks for suspension culture, Greiner Bio One, Les Ulis, France). For routine culture, the cells were seeded at 200,000 cells/ml and split two days later, with a cell density ranging from 800,000 to 1,000,000 cells/ml.

For exposure to plastic particles and to limit the effects of cell growth, cells were seeded at 500,000 cells/ml in 6 or 12 wells adherent plates in DMEM supplemented with 1% horse serum ^40^. The cells were let to settle and recover for 24 h, and then exposed to the selected nanoplastics at 10µg/ml and per day, with a medium renewal every two days. This exposure was carried out for 8 days, plus a final 24 hours period. Proteomic experiments were carried out in 6 well plates, and all the other experiments in 12 well plates. Cells were used at passage numbers from 5 to 15 post-reception from the repository. Cell viability was measured by the propidium iodide method ^41^, or with the SytoxGreen probe (Thermofisher S7020) using the protocol provided by the supplier.

The internalized dose was estimated as followed: cells cultured in 12 wells adherent plates, were exposed to fluorescent particles (PS or PLA) for 8 days as described previously (cumulated dose 80µg/ml). To measure the cells associated fluorescence, the medium was removed, the cell layer was then rinsed once with 1ml PBS and the PBS was removed. The cell layer was then lysed by the addition of 400µl of 10mM Hepes pH 7.5, and the cell lysate was collected. Its fluorescence was then measured using a DeNovix QFX instrument. Excitation was set at 470 nm and emission collected in the 515-567 nm window for the green fluorescent particles. Untreated cell cultures of matched age were used to determine the cell autofluorescence background. In order to determine the reference value, the cumulated dose of the particles (i.e. 80µg/ml) was added into 400µl of 10mM Hepes pH 7.5, and the fluorescence measured as for the cell extracts. This protocol is by construction independent of cell division, as each well is considered as a whole.

### 2.3. Proteomics

Proteomics was carried out essentially as described previously ^42^. However, the experimental details are given here for the sake of consistency.

#### 2.3.1. Sample preparation

After exposure to the plastic particles, the cells were harvested by flushing the 6 well plates. They were collected by centrifugation (200g, 5 minutes) and rinsed twice in PBS. The cell pellets were lysed in 100 µl of extraction buffer (4M urea, 2.5% cetyltrimethylammonium chloride, 100mM sodium phosphate buffer pH 3, 150µM methylene blue). The extraction was let to proceed at room temperature for 30 minutes, after which the lysate was centrifuged (15,000g, 15 minutes) to pellet the nucleic acids. The supernatants were then stored at -20°C until use.

#### 2.3.2. Shotgun proteomics

For the shotgun proteomic analysis, the samples were included in polyacrylamide plugs according to Muller et al. ^43^ with some modifications to downscale the process ^44^. To this purpose, the photopolymerization system using methylene blue, toluene sulfinate and diphenyliodonium chloride was used ^45^.

As mentioned above, the methylene blue was included in the cell lysis buffer. The other initiator solutions consisted in a 1 M solution of sodium toluene sulfinate in water and in a saturated water solution of diphenyliodonium chloride. The ready-to-use polyacrylamide solution consisted of 1.2 ml of a commercial 40% acrylamide/bis solution (37.5/1) to which 100 µl of diphenyliodonium chloride solution, 100 µl of sodium toluene sulfinate solution and 100 µl of water were added.

To the protein samples (15 µl), 5 µl of acrylamide solution were added and mixed by pipetting in a 500µl conical polypropylene microtube. 100 µl of water-saturated butanol were then layered on top of the samples, and polymerization was carried out under a 1500 lumen 2700K LED lamp for 2 hours, during which the initially blue gel solution discolored. At the end of the polymerization period, the butanol was removed, and the gel plugs were fixed for 1 hr with 200 µl of 30% ethanol 2 % phosphoric acid, followed by 3x 15 minutes washes in 20% ethanol. The fixed gel plugs were then stored at -20°C until use.

The gel plugs were washed three times with 50 µL of 25 mM ammonium hydrogen carbonate (NH4HCO3) and 50 µL of acetonitrile. The cysteine residues were reduced by 50 µL of 10 mM dithiothreitol at 57°C and alkylated by 50 µL of 55 mM iodoacetamide. After two washes with NH4HCO3 and acetonitrile, the gel plugs were dehydrated by acetonitrile. In-gel protein digestion was performed with 20 µL of 20 ng/µL of trypsin (Promega V5111) in 25 mM NH4HCO3, overnight at room temperature. Peptides were extracted with 50 µL of 60% acetonitrile in 0.1% formic acid. Acetonitrile was evaporated under vacuum and samples were resuspended with 50 µL 1% acetonitrile and 0.1% formic acid to reach a final concentration of 400 ng/µL.

NanoLC-MS/MS analysis was performed using a nanoACQUITY Ultra-Performance-LC (Waters Corporation, Milford, USA) coupled to a Q Exactive HF-X mass spectrometer (Thermo Fisher Scientific, Bremen, Germany).

Samples (1 µL) were first concentrated/desalted onto a NanoEaseTM M/Z Symmetry C18 precolumn (100Å, 5 µm, 180 µm × 20 mm, Waters Corporation, Milford, USA) using 99% of solvent A (0.1% formic acid in water) and 1% of solvent B (0.1% formic acid in acetonitrile) at a flow rate of 5 µl/min for 3 min. A solvent gradient from 1 to 6% of B in 0.5 min then from 6 to 35% of B in 58 min was used for peptide elution, which was performed at a flow rate of 350 nl/min using a NanoEaseTM M/Z BEH C18 column (130Å, 1.7 µm, 75 µm × 250 mm, Waters Corporation, Milford, USA) maintained at 60 °C.

The Q-Exactive HF-X was operated in positive ion mode with source temperature set to 250 °C and spray voltage to 1.8 kV. Full-scan MS spectra (300–1800 m/z) were acquired at a resolution of 60,000 at m/z 200. MS parameters were set as follows: maximum injection time of 50 ms, AGC target value of 3e6 ions, lock-mass option enabled (polysiloxane, 445.12002 m/z), selection of up to 10 most intense precursor ions (doubly charged or more) per full scan for subsequent isolation using a 2 m/z window, fragmentation using higher energy collisional dissociation (HCD, normalised collision energy of 27), dynamic exclusion of already fragmented precursors set to 30 s. MS/MS spectra (200–2000 m/z) were acquired with a resolution of 15,000 at m/z 200. MS/MS parameters were set as follows: maximum injection time of 50 ms, AGC target value of 1e5 ions, peptide match selection option turned on. Raw data were converted into mgf files using the MSConvert tool from ProteomeWizard (v3.0.6090; http://proteowizard.sourceforge.net/)

For protein identification, the MS/MS data were interpreted using a local Mascot server with MASCOT 2.6.2 algorithm (Matrix Science, London, UK) against an in-house database containing all Mus musculus and Rattus norvegicus entries from UniProtKB/SwissProt (version 2025_04, 17,362 sequences) and their corresponding 17,362 reverse entries. Spectra were searched with a mass tolerance of 10 ppm for MS and 0.07 Da for MS/MS data, allowing a maximum of one trypsin missed cleavage. Trypsin was selected as the cutting enzyme and a maximum of one missed cleavage was allowed. Acetylation of protein N-termini, carbamidomethylation of cysteine residues and oxidation of methionine residues were specified as variable modifications. Identification results were imported into Proline software version 2.3 (http://proline.profiproteomics.fr/) for validation ^46^. Peptide Spectrum Matches (PSM) with pretty rank equal to one were retained. False Discovery Rate was then optimized to be below 1% at PSM level using Mascot Adjusted E-value and below 1% at Protein Level using Mascot Mudpit score.

Mass spectrometry data are available via ProteomeXchange with the identifier PXD049997.

### 2.4. Data analysis

For the global analysis of the protein abundances data, missing data were imputed with a low, non-null value. Proteins that were detected less than 3 times out of the 5 replicates in both groups were removed from the analysis. The complete abundance dataset was then analyzed by the PAST software ^47^.

Proteins were considered as significantly different if their p value in the Mann-Whitney U-test against control values was inferior to 0.05. No quantitative change threshold value was applied, as this can lead to the artefactual elimination of relevant changes ^48^. The selected proteins were then submitted to pathway analysis using the DAVID tool ^49^, with a cutoff value set at a FDR of 0.25.

### 2.5. Mitochondrial transmembrane potential assay

The mitochondrial transmembrane potential assay was performed essentially as described previously ^50^. Rhodamine 123 (Rh123) was added to the cultures at an 80 nM final concentration (to avoid quenching ^51^), and the cultures were further incubated at 37°C for 30 minutes. At the end of this period, the cells were collected, washed in cold PBS containing 0.1% glucose, resuspended in PBS glucose and analyzed for the green fluorescence (excitation 488 nm emission 525nm) on a Melody flow cytometer. As a negative control, carbonyl cyanide 4-(trifluoromethoxy)phenylhydrazone (FCCP) was added at 5µM final concentration together with the Rh123 ^52^.

### 2.6. Assay for oxidative stress

For the oxidative stress assay, a protocol based on the oxidation of dihydrorhodamine 123 (DHR123) was used, essentially as described previously ^50^. After exposure to plastic beads as described in section 2.2, the cells were treated in PBS containing 500 ng/ml DHR123 for 20 minutes at 37°C. The cells were then harvested, washed in cold PBS containing 0.1% glucose, resuspended in PBS glucose and analyzed for the green fluorescence (same parameters as rhodamine 123) on a Melody flow cytometer. Menadione (applied on the cells for 2 hours prior to treatment with DHR123) was used as a positive control in a concentration range of 25-50µM.

### 2.7. Lysosomal assay

For the lysosomal function assay, the Lysosensor method was used, as described previously ^50^. After exposure to nanoplastics as described in section 2.2, the medium was removed, the cell layer was rinsed with complete culture medium and incubated with 1µM Lysosensor Green (Molecular Probes) diluted in warm (37°C) complete culture medium for 1 hour at 37°C. At the end of this period, the cells were collected, washed in cold PBS containing 0.1% glucose, resuspended in PBS glucose and analyzed for the green fluorescence (excitation 488 nm emission 540nm) on a Melody flow cytometer.

### 2.8. Phagocytosis assay

For this assay, the cells were first exposed to the PS or PLA nanoparticles as described in section 2.2. Twenty four hours after the final daily exposure to the plastics, the cells were then exposed to 0.5 µm latex beads (carboxylated surface, yellow green-labelled, from Polysciences excitation 488 nm emission 527/32 nm) for 3 hours. After this second exposure, the cells were collected, rinsed twice with PBS, and analyzed for the two fluorescences (green and red) on a Melody flow cytometer.

### 2.9. Cell surface markers

Cells were seeded into 12-well plates at a concentration of 500,000 cells/ml, and exposed to the nanoplastic particles as described in section 2.2. The day following the last addition of nanoplastics, the cells were harvested and washed in DMEM containing 3% FBS. For all labelled antibodies, the cells were treated with the fluorochrome-conjugated antibodies at the adequate dilution for 30 minutes on ice in DMEM-FBS in a final volume of 100µl. For collectin 12, the cells were first treated with the primary antibody at the adequate dilution for 30 minutes on ice in DMEM-FBS in a final volume of 100µl. The cells were then collected by centrifugation, resuspended in 100 µl DMEM.FBS and treated for 30 minutes on ice with the secondary antibody (Rabbit Anti-Goat igG, FITC conjugated, Invitrogen #31509, dilution 1/300)

In all cases, the cells were then washed with 2ml of PBS. The cell pellet was resuspended in 400 µl of PBS containing 1 µM Sytox blue for checking cell viability, and the suspension analyzed by flow cytometry on a Melody flow cytometer. First, live cells (Sytox blue-negative) were selected at 450 nm (excitation at 405 nm). The gated cells were then analyzed for the nanoplastics fluorescence (excitation 561nm emission 695 nm) and for the antibody fluorescence (excitation 488 nm emission 527 nm for FITC or A488 labelled antibodies or excitation 561 nm emission 582 nm for PE-conjugated antibodies). Isotypic control antibodies were used to compensate for non-specific binding.

The following antibodies were used:

Collectin-12 (Invitrogen #PA5-47456, RRID:AB_2606750) dilution 1/6

CD86-FITC (BD-Pharmingen #553-691, RRID:AB_394993) dilution 1/25

CD204-A488 (Invitrogen #53-2046-82, RRID:AB_2802322) dilution 1/25

PD-L1-A488 (BD-Pharmingen #566-864, RRID:AB_2869917) dilution 1/25

TLR2-FITC (Invitrogen# 11-9021-82, RRID:AB_465440) dilution 1/25

TLR7-PE ((BD-Pharmingen #565-557, RRID:AB_2739295) dilution 1/50

In the experiments with TLR7, the cells were collected after exposure to the nanoplastics, washed with PBS, and then fixed and permeabilized with the BD cytofix-cytoperm kit (#554715). The cells were incubated with the anti-TLR7 antibody diluted to 1/50 (respectively) for 30 minutes on ice. The cells were then washed with the permeabilization buffer, resuspended in PBS and analyzed by flow cytometry as described above.

### 2.10. Cytokine release assays

Cells were first exposed to nanoplastics as described in section 2.2. At the end of this exposure period, i.e. 24 hours after the last addition of beads, the culture medium was collected and analyzed for proin-flammatory cytokines. In half of the wells LPS (1 ng/ml) was added 18 hours before medium collection. Tumor necrosis factor (catalog number 558299, RRID:AB_2869144, BD Biosciences, Le Pont de Claix, France), MCP1 (catalog number 558342, RRID:AB_2869167, BD Biosciences, Le Pont de Claix, France)and interleukin 6 (IL-6) (catalog number 558301, RRID:AB_2869146, BD Biosciences, Le Pont de Claix) levels were measured using the Cytometric Bead Array Mouse Inflammation Kit (catalog number 558266, BD Biosciences, Le Pont de Claix), and analyzed with FCAP Array software (RRID:SCR_025927, version 3.0, BD Biosciences) according to the manufacturer’s instructions.

## 3. Results

### 3.1. Particle characterization

First, the chemical nature of the particles was verified by IR spectroscopy. The results, shown on **Figure 1**, confirmed the chemical nature of the polystyrene (PS) and polylactide (PLA) particles, respectively, with characteristic peaks assigned in the IR spectra.

**Figure 1.**
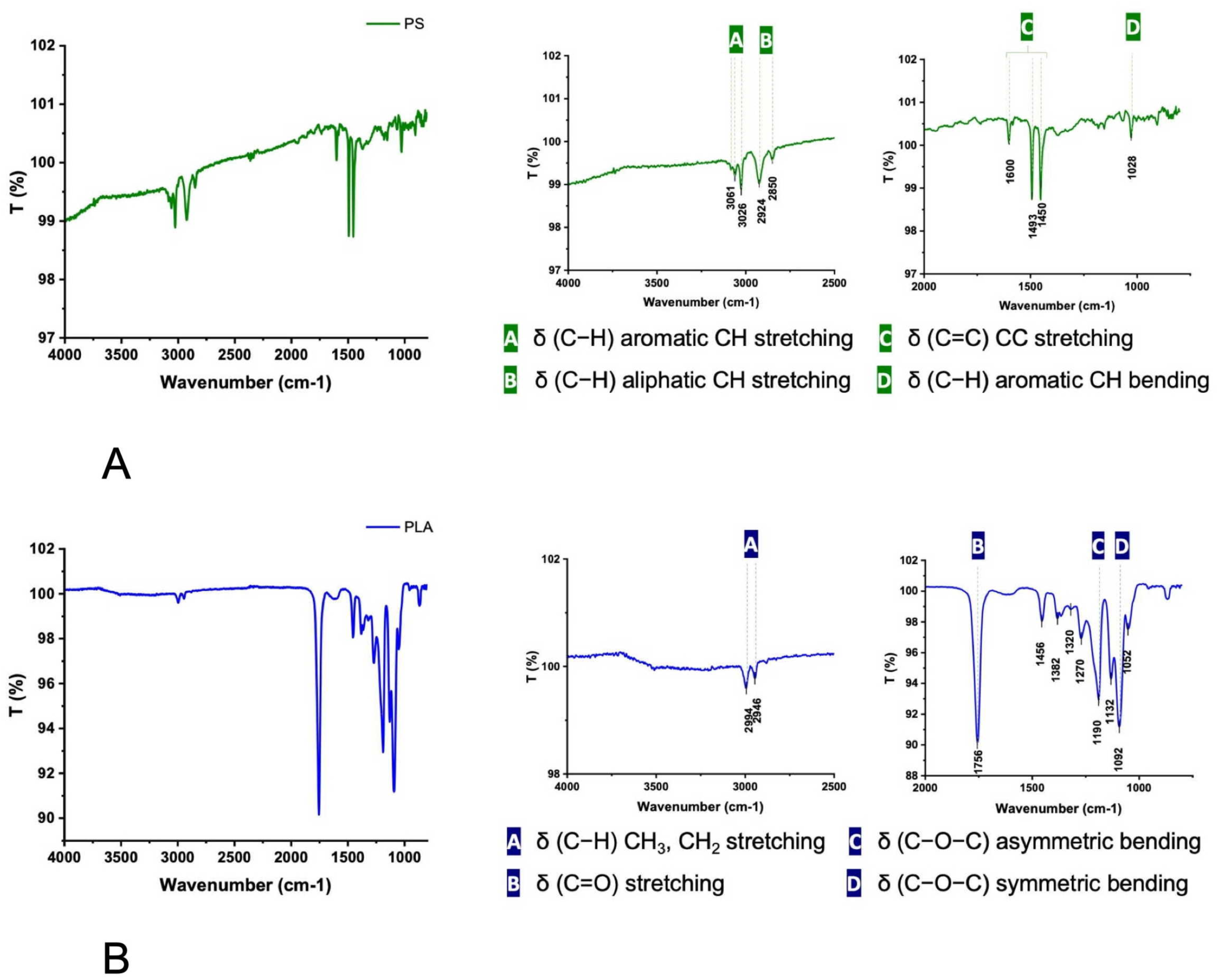
Infrared spectra of the PS and PLA particles Infrared spectroscopy data were recorded on the dry plastic particles. Panel A: data obtained for the PS particles the complete spectrum is shown on the left panel. The two insets describe the main characteristic vibrations that confirm the chemical identity of the PS particles. Panel B: data obtained for the PLA particles the complete spectrum is shown on the left panel. The two insets describe the main characteristic vibrations that confirm the chemical identity of the PLA particles.

The size of the particles was then determined by transmission electron microscopy. The results, displayed on **Figure 2**, showed spherical particles of rather uniform size, in the 200-250 nm size range as expected.

**Figure 2.**
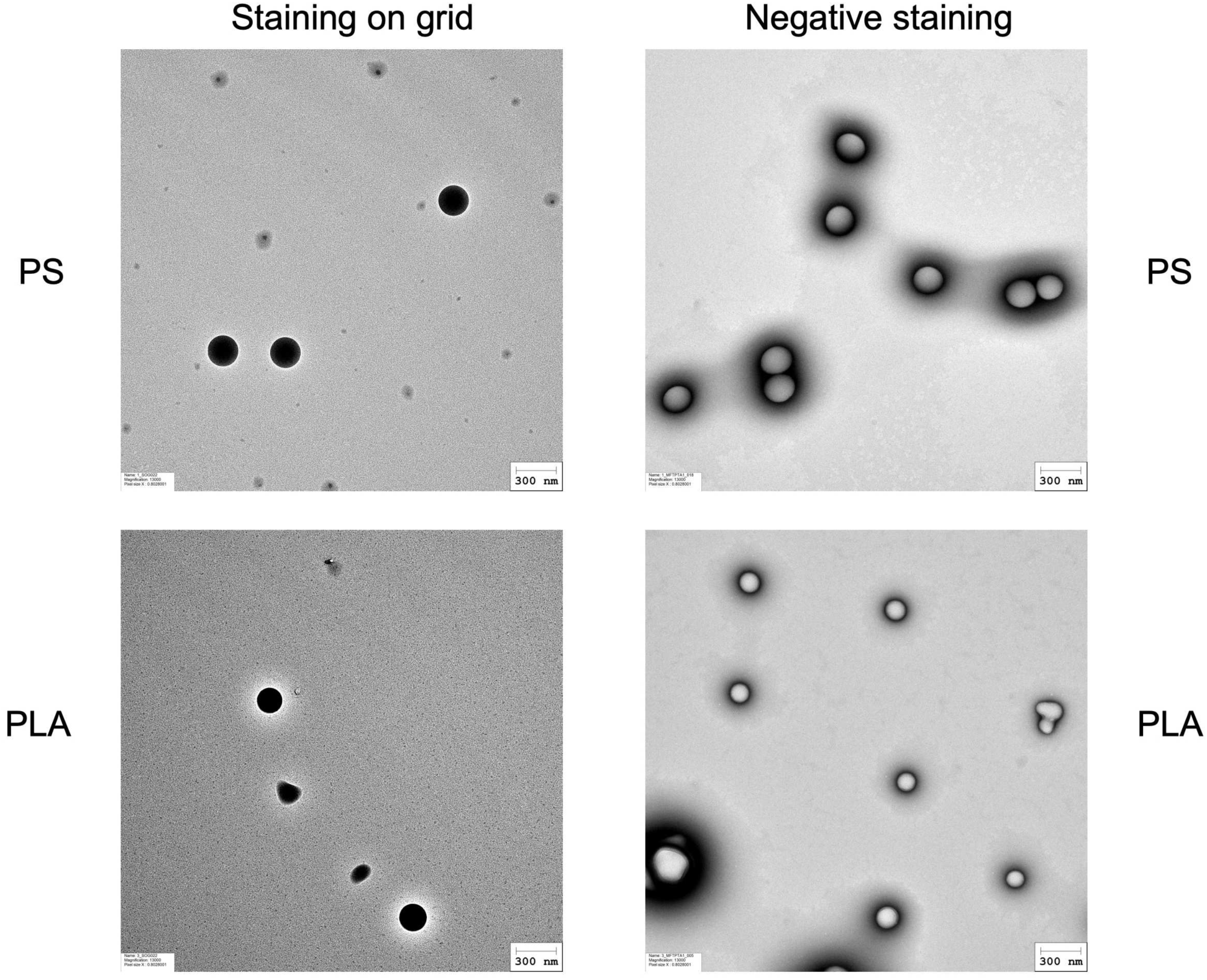
Electron microscopy images of the PS and PLA particles. Particles dispersions were deposited on a carbon grid (staining on grid) or on a mica film which was post-stained with phosphotungstic acid (negative staining). The samples were then observed by transmission electron microscopy First row images of PS particles Second row: images of PLA particles

The in solution parameters of the two plastic particles were characterized using Dynamic Light Scattering (DLS) to assess their size distribution, polydispersity index (PI), and sedimentation behaviour. As summarized in **Table 1**.

**Table 1:**
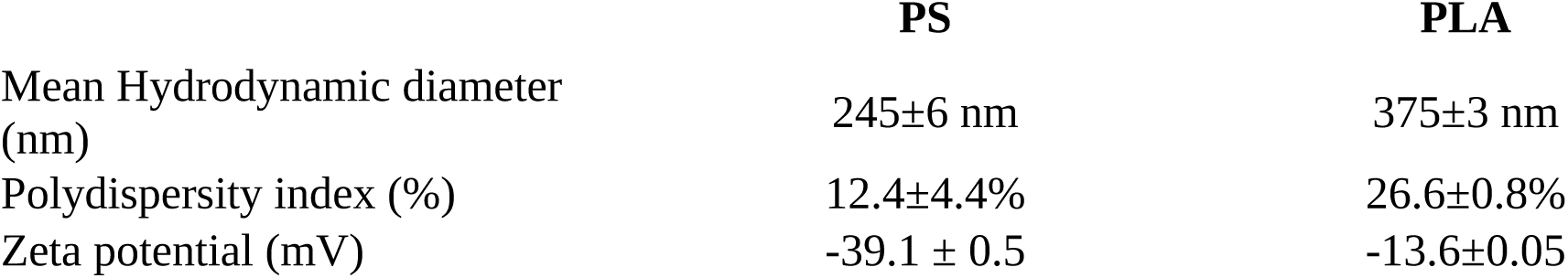
Physicochemical characterization of the PLA and PS particles used in this study.

### 3.2. Particle internalization

We then focused on the total particle internalization during this repeated exposure scheme, using commercially-available labelled particles. The results (N=6) showed a moderate internalization of the PS particles (25.2±1.6%), but a much higher internalization of the PLA particles (63.8±8.5%). By contrast, when the same cumulative dose (80µg/ml) was delivered as a single 24h exposure, 47±3% of the PS beads were internalized and 67±13% of the PLA beads. We also verified the biodegradability of the PLA beads, by exposing the cells to a single 80 µg/ml dose for 24 hours, followed by a recovery period without beads. After 48 hours of recovery, only 54% of the beads-associated fluorescence was still associated with the cells, and 45% after 5 days of recovery.

### 3.3. Global analysis of the proteomic results

The proteomic analysis led to the detection and quantification of 3328 proteins (**Table S1**). In order to evaluate the global extent of changes at the proteome scale, we performed a Principal Coordinate analysis, using the PAST software ^47^ . The results, displayed on **Figure 3**, showed a good separation of the control from the two plastics-treated ones, while the PS and PLA groups did not differ by a large extent. This showed that the processes that are common to the two plastic treated groups, e.g. particle internalization and generic responses to plastics, induced more proteome-wide changes than the specific reactions to each nanoplastic.

**Figure 3.**
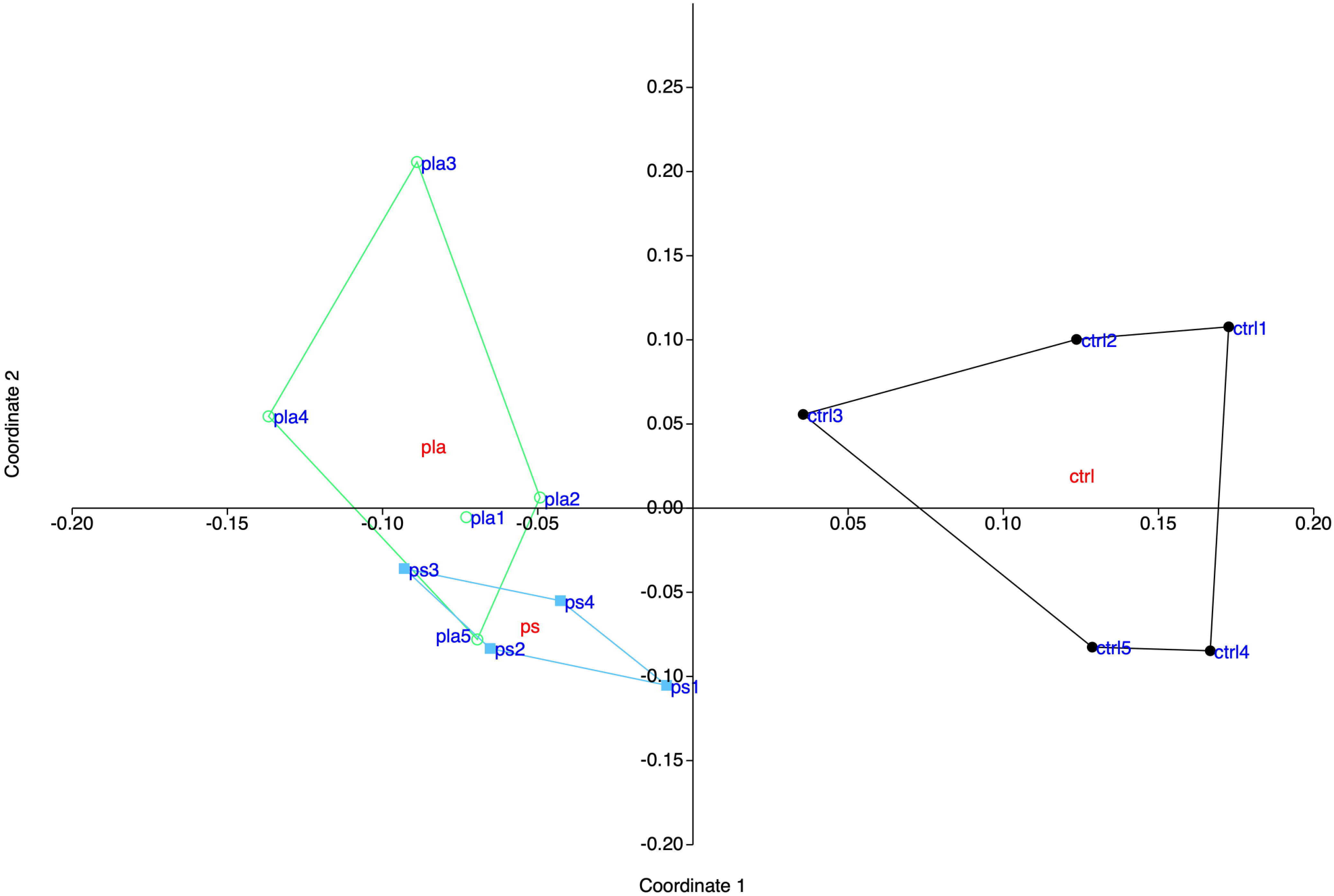
Global analysis of the proteomic data. The complete proteomic data table (3328 proteins) was analyzed by Principal Coordinates Analysis (Gower distance), using the PAST software. The results are represented as the X-Y diagram of the first two axes of the Principal Coordinate Analysis, representing 35% of the total variance. Eigenvalue scale. This representation allows to figure out how, at the global proteome scale, the samples are related to each other. Samples grouped in such a diagram indicate similar proteomes, and the larger the distance between samples are, the more dissimilar their respective proteomes.

We could however select proteins which expression was significantly changed in response to the repeated exposure to each of the particles. To this purpose, we used the Mann-Whitney U test, with a cutoff at p<0.05. This process led to the selection of 361 proteins responding to the treatment with PS particles (**Table S2**) and of 432 proteins responding to the treatment with the PLA particles (**Table S3**). Only 153 proteins were modulated in common by the two nanoplastics **(Table S4)**.. These selected proteins were then submitted to a pathway using the DAVID pathway analysis tool ^53^, and the results are displayed on **Tables S5 and S6**. The pathway analysis pointed out several cellular processes which were modulated upon treatment with plastic particles. Most were generic, e.g. mitochondria, lysosomes or endoplasmic reticulum, but immune response also appeared in the top selected pathways.

### 3.3. Mitochondria and carbon metabolism

Mitochondrial proteins represented an abundant class among the proteins modulated in response to nanoplastics, with 110 proteins (**Table S7**). As a large collection of mitochondrial proteins in the particles-responsive proteins may indicate mitochondrial perturbations ^54^ , we analyzed the transmembrane mitochondrial potential. The results, displayed in **Figure 4A**, did not show any perturbation of the transmembrane mitochondrial potential in response to repeated exposure to PS particles, indicating that the response observed at the proteome level for these particles is homeostatic. However, an increase in the transmembrane mitochondrial potential was observed in response to repeated exposure to PLA particles.

**Figure 4.**
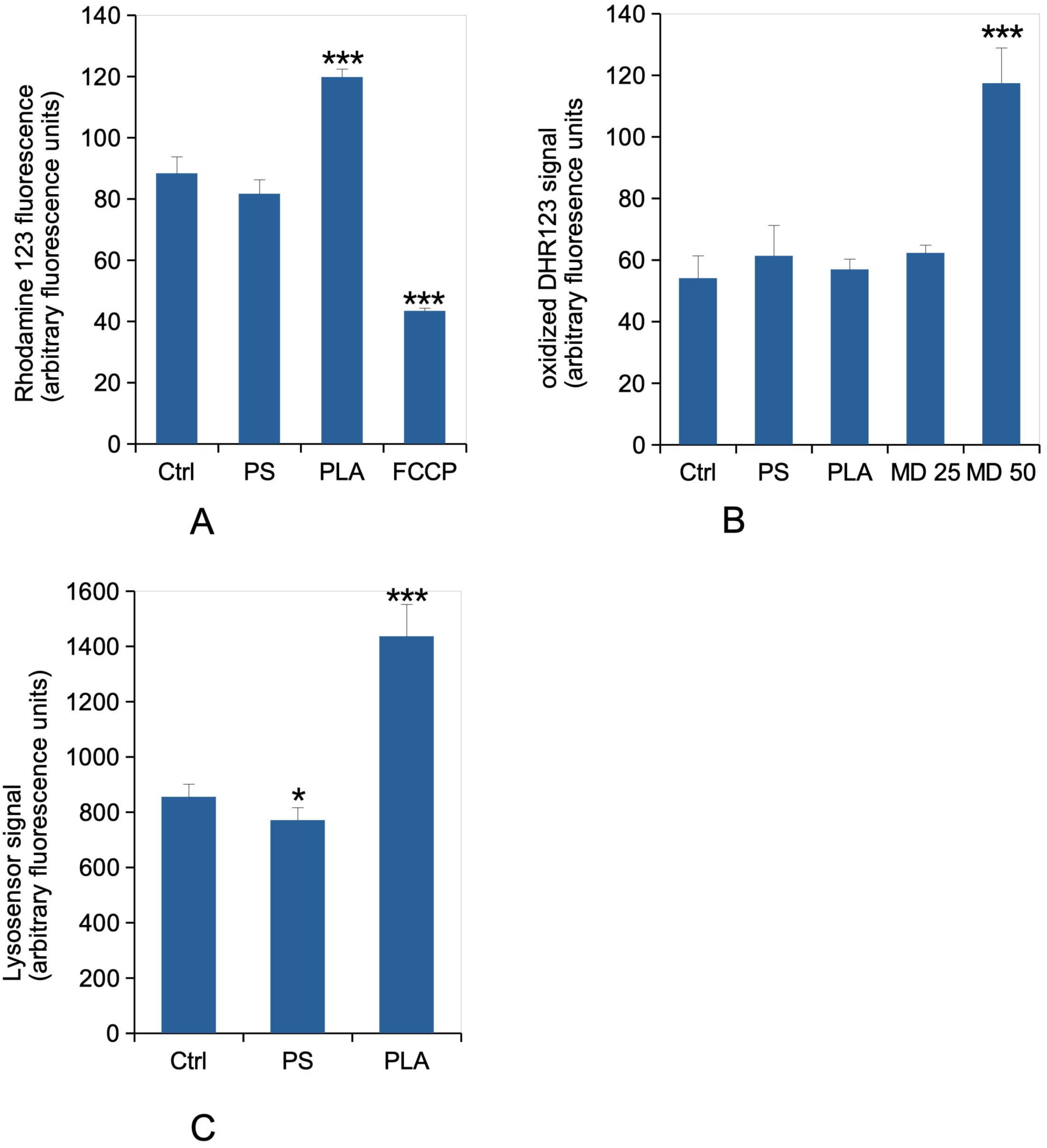
Mitochondria, oxidative stress and lysosomes experiments. Panel A: mitochondrial transmembrane potential (rhodamine 123 method). All cells were positive for rhodamine 123 internalization in mitochondria, and the mean fluorescence is the displayed parameter. Results are displayed as mean± standard deviation (N=4). FCCP: carbonyl cyanide 4-(trifluoromethoxy)phenylhydrazone. Significance marks: *** p<0.001 (Student T test, treated vs control comparison) Panel B: Cellular oxidative stress, measured with the dihydrorhodamine 123 (DHR123) indicator The cells were exposed to nanoplastics as described in section 2.2, and finally for 20 minutes to the DHR 123 probe. Menadione (25 or 50 µg/ml for 2 hours) was used as a positive oxidative stress control. Results are displayed as mean± standard deviation (N=4). Significance marks: *** p≤0.001 (Student T test, treated vs control comparison). MD: Menadione Panel C : lysosomal proton pumping (Lysosensor method). All cells were positive for lysosensor internalization in lysosomes, and the mean fluorescence is the displayed parameter. Results are displayed as mean± standard deviation (N=4). Significance marks: * p≤0.05 ; *** p<0.001 (Student T test, treated vs control comparison)

As lactic acid is a byproduct of glycolysis and also liberated by PLA particles upon their intracellular degradation, we also analysed our proteomic data for the proteins involved in glycolysis and in the pentose phosphate pathway. The results, displayed in **Table S8**, showed that PLA particles (and PS particles to a lesser extent) did impact these pathways. For the pentose phosphate pathway, 6-phosphogluconate dehydrogenase, the protein catalyzing the irreversible degradation of the glucose unit, was induced in response to both nanoplastics. PS particles also induced an increase in the amount of glucose phosphate dehydrogenase and transketolase. This may indicate a higher demand in NADPH in response to PS particles.

Glycolysis was much more impacted. On the 16 proteins implicated in glycolysis that we identified in our proteomic screen, 2 proteins (hexokinase 3 and enolase) were impacted by both nanoparticles, while 7 were increased in response to PLA nanoparticles.

### 3.4. Oxidative stress

In our studies on different particles in a high dose, acute exposure mode, we observed that PS increased the level of oxidative stress, while PLA did not show any effect ^28^. We thus investigated whether these responses also existed in response to a repeated exposure to a lower daily dose. The results, displayed in **Figure 4B**, did not show any significant change in the level of oxidative stress in response to both particles in the repeated exposure scheme.

### 3.5 Lysosomes

The proteomic screen led to a list of 48 lysosomal proteins which abundances were modulated in response to repeated exposure to either PS or PLA nanoplastics (**Table S9**). Furthermore, in our studies on different particles in a high dose, acute exposure mode, we observed that PLA increased the lyso-sensor response ^28^, while PS did not ^42^. We thus investigated whether these responses also existed in response to a repeated exposure to a lower daily dose. The results, displayed in **Figure 4C**, showed a small but significant decrease in response to PS particles, and a significant one in response to PLA particles.

### 3.6 Phagocytosis

In a scheme of acute exposure to a high dose of nanoparticles, PS particles did not change the phagocytic capacities of macrophages ^42^, while PLA particles increased phagocytosis ^28^. The situation was however different when the same particles were given to macrophages in a repeated exposure scheme. In this case, as shown in **Figure 5**, a repeated exposure to PS or PLA nanoparticles slightly depressed phagocytosis, but to a non significant extent in both cases.

**Figure 5.**
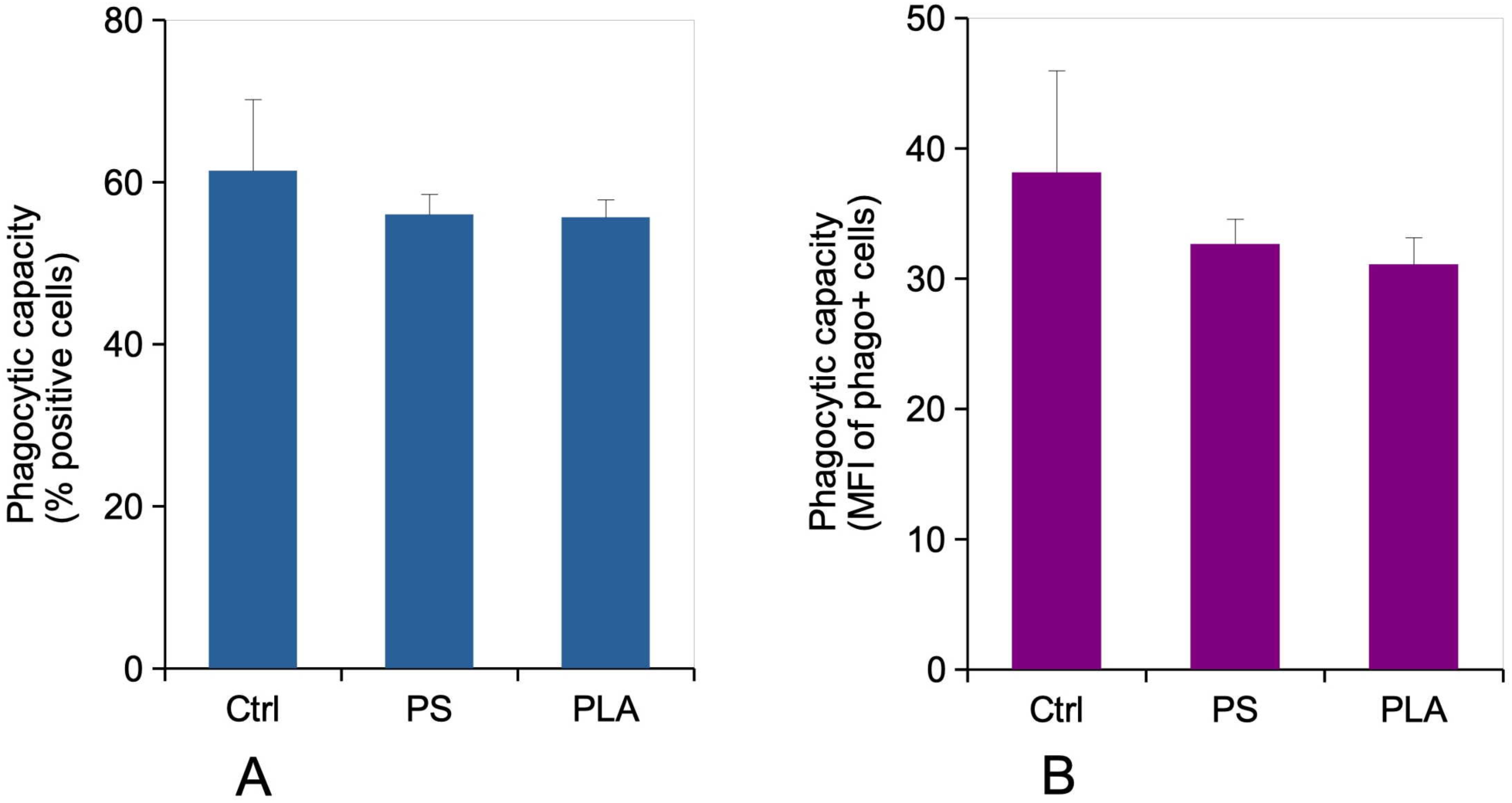
Phagocytic capacity. The cells were exposed to nanoplastics as described in section 2.2. After removal of the plastics-containing cell culture medium, the cells were treated with green fluorophore labelled carboxylated polystyrene beads for 3 hours. Panel A: The percentage of green fluorescence-positive cells, indicating the percentage of cells able to internalize the test beads in 3 hours, is the displayed parameter. Results are displayed as mean± standard deviation (N=4). Significance marks: * p≤0.05 (Student T test, treated vs control comparison) Panel B: The mean fluorescence, indicating the amount of green beads internalized, is the displayed parameter. Results are displayed as mean± standard deviation (N=4).

### 3.7. Surface markers

The pathway analysis highlighted immune response as modulated in response to a repeated exposure to PS or PLA nanoplastics (**Table S10**). We also included in this table proteins that we knew to be modulated in response to acute exposures, but that did not appear as such in the proteomic screen in response to repeated exposure. However, proteomics measures the total protein concentration, and not the one correctly folded and addressed to the relevant cell membrane, which is the useful parameter for membrane receptors. We thus measured this parameter using antibody labelling and flow cytometry, and the results are displayed on **Table 2**.

**Table 2:** Assay of surface antigens by flow cytometry.

|  | Control cells | PS-exposed cells | PLA-exposed cells |
| --- | --- | --- | --- |
| CD86 | 60±0.4 | **49.1±1.5** | **51.7±2.2** |
| Collectin 12 | 20.0±8.8 | 19.7±6.6 | 14.8±3 |
| CD204 (scavenger receptor) | 13.8±0.8 | **10.7±0.8** | **7.4±0.1** |
| PD-L1 | 6.1±0.5 | **7.7±0.5** | **8.5±0.3** |
| TLR2 | 44.9±1.5 | **69±4.5** | **75.5±4.5** |
| TLR7 | 63±3.8 | **47.5±4.3** | **45.6±5.3** |

The results are expressed as mean fluorescence ± standard deviation (N=4)
Bold figures indicate significant changes (Student T test, p<0.05, treated vs control comparison)

Despite not being detected by the proteomic screen, several of these surface markers were indeed modulated in response to repeated exposure to PS or PLA nanoplastics. The variations observed here for TLR7 or CD204 were consistent with those observed in proteomics.

### 3.8. Pro-inflammatory cytokine release

In a scheme of acute exposure to a high dose of nanoparticles, PLA and PS particles increased the release of TNF, while PLA and PS particles depressed the release of both TNF and IL-6 after LPS stimulation ^28^. We thus investigated this parameter in response to a repeated exposure to the same particles. The results, displayed in **Figure 6** showed that PS particles decreased the release of IL-6, MCP-1, and TNF without any additional stimulation (**Figure 6A, C, E**), while PLA particles only decreased the intrinsic release of IL-6 (**Figure 6A**). After an additional stimulation with LPS, PLA particles significantly decreased the release of IL-6, MCP-1 and TNF, while PS particles only decreased the release of IL-6 (**Figure 6B, D, F**).

**Figure 6:**
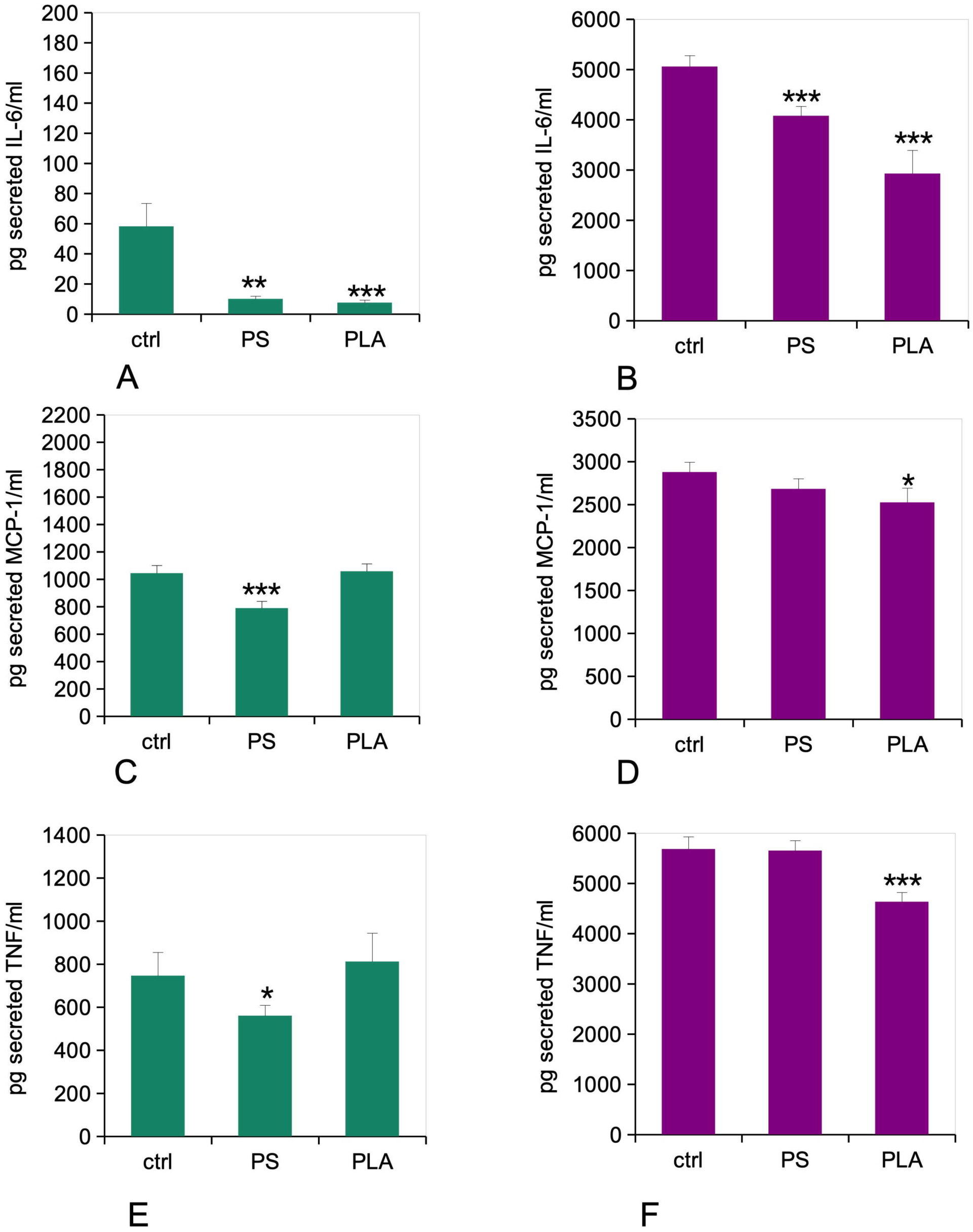
Cytokine release. The cells were exposed to nanoplastics as described in section 2.2. The cells were treated (or not) with 50ng/ml lipopolysaccharide in complete cell culture medium for the last 18 hours. The cell medium was then collected for secreted IL-6, MCP-1 and TNF alpha measurements. Panel A: IL-6 release without LPS stimulation Panel B: IL-6 release with LPS stimulation (50 ng/ml) Panel C: MCP-1 release without LPS stimulation Panel D: MCP-1 release with LPS stimulation (50 ng/ml) Panel E: TNF-alpha release without LPS stimulation Panel F: TNF-alpha release with LPS stimulation (50 ng/ml) *: statistically different from the unexposed control (p≤0.05, Student T test, treated vs control comparison) **: statistically different from the unexposed control (p≤0.01, Student T test, treated vs control comparison) ***: statistically different from the unexposed control (p≤0.001, Student T test, treated vs control comparison)

## 4. Discussion

Many plastics are quite persistent in the environment and also accumulate in the body over time ^55–58^. In this respect, they are similar to some inorganic nanoparticles such as titanium dioxide, which accumulates in gut macrophages to form so-called pigmented cells ^59^ . In addition, biodegradable plastics can release their degradation products (oligomers, monomers or other small molecules) over time, adding another toxicological issue.

Most plastic contaminations are rather chronic and nonetheless, most of the in vitro toxicology studies devoted to the effects of micro and nanoplastics on living cells use an acute exposure mode, i.e. expose the cells to plastics for a short time (24 hours or less) with the readout performed immediately after exposure. Only a few studies used repeated and extended exposure over time ^36–39,60–62^. As these publications on epithelial or fibroblastic cells showed results that were substantially different from those obtained under an acute exposure mode , we decided to perform such repeated exposure experiments on macrophages with a non-biodegradable polymer (PS) and a biodegradable one (PLA). However, such repeated exposure studies must be adapted to the various cell physiologies that can be encountered. The above-cited studies have dealt with epithelial cells, which divide continuously to ensure self renewal and tissue homeostasis. Consequently, these studies have been performed under conditions which ensure cell division. When dealing with macrophages, the context is completely different. Macrophages were long believed to be post-mitotic, although it is now known that they have a slow self-renewal capacity ^63^ . The situation is however rather complex and seems to depend on the various localizations of the macrophages. Alveolar macrophages exhibit a long lifespan ^64^, while dermal or intestinal macrophages show a much shorter lifespan ^65,66^. When dealing with persistent particulate material that has been ingested by macrophages, the key issue becomes the fate of the particles. Here again the alveolar macrophages show a peculiar behavior, as they are eliminated via the mucociliary escalator ^67^. Oppositely, macrophages that are located below the epithelial barriers, such as dermal or intestinal macrophages seem to die on site, liberating their particle cargo that is taken up again by new macrophages replenishing the site. Such a mechanism explains tattoos persistence during life ^66^, and also the progressive appearance of heavily loaded cells when the exposure is continuous ^59^.

When taking all these parameters into account, we devised an exposure system that was based on the peritoneal macrophage J774A.1 ^68^, slowed down by serum decrease in the culture medium to reduce proliferation ^40^, and not on the THP-1 system, which is known to be rather variable in its responses ^69^. Our experiments took place over a total period of 10 days, which is not unreasonable regarding the life-span of dermal and intestinal macrophages ^65,66^ . Furthermore, and opposite to epithelial cells that are present in well-organized tissues, macrophages are usually present in the connective tissues below the epithelia, i.e. in environments where cell-cell contacts are minimal. Indeed, immune cells generally communicate by diffusible signals over rather long distances, instead of cell-cell contacts. Thus, macrophages monocultures represent a rather good approximation of the in vivo situation. We thus reasoned that a split dose over one week may be a good mimic of the final situation in gut macrophages. For biodegradable plastics, such as PLA, this is a good tradeoff between the degradation rate of the plastics and the exposure time. For non degradable plastics, such as PS, the final concentration in macrophages after a lifelong exposure is difficult to evaluate a priori, but the example of pigmented cells observed with titanium dioxide ^59^ shows that the final concentration may be quite high. Thus, we used a non-cytotoxic dose of 10 µg/ml nanoplastics repeated over 8 consecutive days (which is still non cytotoxic at the cumulated level) as a model repeated exposure for macrophages.

We therefore investigated in detail, through the proteomic and targeted experiments, the macrophages responses to a repeated exposure to either PS or PLA nanoparticles.

The first striking result was the low response of macrophages to a repeated exposure to PS nanoparticles in a repeated mode. Only 360 proteins were modulated in response to this exposure, compared to the 900 modulated in response to an acute exposure to a high dose ^35^. This may be linked to the relatively low intake of PS nanoparticles in the repeated mode (18% of the total dose), compared to the acute mode (47%). This low apparent intake may be due to different phenomena. Either the cells having internalized PS beads gets desensitized and internalize less beads over time, or the cells having internalized the most beads in the first exposure detach from the culture support and get lost during the media changes that must be made to extend the culture time to more than one week. In this case, only cells having a low internalization rate would be kept. As we know that the cells in a single culture are highly variable in their particles internalization capacities ^50^ , this hypothesis cannot be ruled out.

Nevertheless, the cells that remain at the end of the exposure show a weak decrease in their lysosomal activity and an insignificant decrease in their phagocytic ability. This may be consistent with a desensitization occurring during the repeated exposure time frame. This desensitization hypothesis is consistent with the decrease observed in CD86, CD204 and TLR7 expression, and also with the decrease in IL-6 observed after a LPS stimulation. The change in CD86, CD204 and TLR7 expression was indeed more pronounced in the repeated exposure scheme than in an acute exposure scheme, while the increase in PD-L1 expression was consistent between both schemes ^50^. The cytokine secretion data obtained after the repeated exposure were in line with the data obtained in response to an acute exposure ^35^. However, the response to a repeated exposure was more significant than the one to an acute exposure.

The situation was quite different for the response to PLA nanoparticles. Here, the proteomic response was slightly higher in the repeated exposure scheme (432 modulated proteins) that in the acute exposure scheme (346 modulated proteins) ^28^. This higher number may be indicative of a higher response in the case of repeated exposure. Indeed, an important mitochondrial response was seen in the repeated exposure scheme, but completely absent in the acute exposure scheme. This may be explained by the progressive release of lactic acid when cells are exposed to PLA for a prolonged period of time. Lactic acid is released from the PLA particles within the lysosomes, where the particles are sequestered. It may then reach and acidify the cytosol, including the outer side of the inner mitochondrial membrane. In addition, lactic acid can be converted back to pyruvate by lactate dehydrogenase, leading to a pyruvate influx into mitochondria and thus an increased mitochondrial metabolism. Both mechanisms may explain the increase in the transmembrane mitochondrial potential that is observed in the repeated exposure scheme. PLA dissolution within cells is relatively rapid, but it still takes a few days ^28^. Thus, this effect may just have had not enough time to appear when the cells are analyzed immediately after exposure to the PLA particles ^28^ . Regarding the lysosomal activity, both the acute exposure ^28^ and the repeated one (this work) induced an increase in activity.

Regarding the cytokine release, the data were consistent between the two exposure modes for IL-6, but divergent for TNF. An increase was observed immediately after exposure to a high dose ^28^ , while no response was observed after a repeated exposure to PLA particle only and a decrease was observed when a further LPS stimulation was applied.

Regarding the expression of surface marker, the cells repeatedly exposed to PLA particles showed the same pattern than those exposed to PS particles, i.e. a decrease in activation markers (CD86, CD204) and in TLR7, but an increase in PD-L1 and TLR2 expression.

## 5. Conclusions

First of all, the detailed analysis of the cellular responses showed how important it is to perform validation experiments to refine the conclusions that can be derived from proteomics. For example, both nanoplastics induce a strong proteomic response for mitochondrial proteins. However, this response is homeostatic for PS particles, while PLA particles induce an alteration of the mitochondrial transmembrane potential.

All in all, the repeated exposure regime induces a macrophage response that is somewhat different from the one observed after acute exposure. Cells repeatedly exposed to PS or PLA particles appears as partially desensitized and are poorly responsive to bacterial stimuli, which indicates a putative lower response to infections. In addition, cells treated with PLA particles show a disturbed metabolism (e.g. in mitochondria), which may result in an altered lifespan of macrophages in vivo, and thus a strain put on the immune homeostasis. This state of fact also shows that everything should be made to limit exposure to nanoplastics, which shows in turn the crucial role of waste management and of a better control of plastics conditions of use (e.g. mechanical or thermal stress during intended use) to limit this exposure.

## Acknowledgments

Thanks are due to the Biostudies database for hosting the non-proteomic data associated with this work, under the doi 10.6019/S-BSST2314 ^70^

Thanks are also due to the ProteomXchange consortium for hosting the proteomic data associated with this work, under the doi 10.6019/PXD049997 ^71^

We thank Guy Schoehn for the establishment of the IBS/ISBG EM facility, and Matthieu Koepf for the IR spectra

## Author Contributions

TR designed and supervised the study. MV and DF characterized the nanoplastic particles. VCF, HD and SC performed the proteomic experiments, which were then analyzed by TR and ED. VCF performed and interpreted the flow cytometry experiments. TR drafted the initial version of the manuscript, which was completed and amended by all co-authors, who approved the final version of the manuscript.

## References

1. Geyer R, Jambeck JR, Law KL. Production, use, and fate of all plastics ever made. Sci Adv. 2017;3(7):e1700782. doi:10.1126/sciadv.1700782

2. Lehner R, Weder C, Petri-Fink A, Rothen-Rutishauser B. Emergence of Nanoplastic in the Environment and Possible Impact on Human Health. Environmental Science & Technology. 2019;53(4):4. doi:10.1021/acs.est.8b05512

3. Thompson RC, Swan SH, Moore CJ, vom Saal FS. Our plastic age. Philos Trans R Soc Lond B Biol Sci. 2009;364(1526):1973–1976. doi:10.1098/rstb.2009.0054

4. Landrigan PJ, Stegeman JJ, Fleming LE, et al. Human Health and Ocean Pollution. Ann Glob Health. 2020;86(1):151. doi:10.5334/aogh.2831

5. Jambeck JR, Geyer R, Wilcox C, et al. Plastic waste inputs from land into the ocean. Science. 2015;347(6223):6223. doi:10.1126/science.1260352

6. Enders K, Lenz R, Stedmon CA, Nielsen TG. Abundance, size and polymer composition of marine microplastics ≥ 10 µm in the Atlantic Ocean and their modelled vertical distribution. Marine Pollution Bulletin. 2015;100(1):70–81. doi:10.1016/j.marpolbul.2015.09.027

7. Ter Halle A, Jeanneau L, Martignac M, et al. Nanoplastic in the North Atlantic Subtropical Gyre. Environ Sci Technol. 2017;51(23):13689–13697. doi:10.1021/acs.est.7b03667

8. Brandon JA, Freibott A, Sala LM. Patterns of suspended and salp-ingested microplastic debris in the North Pacific investigated with epifluorescence microscopy. Limnology and Oceanography Letters. 2020;5(1):1. doi:10.1002/lol2.10127

9. Kedzierski M, Palazot M, Soccalingame L, et al. Chemical composition of microplastics floating on the surface of the Mediterranean Sea. Marine Pollution Bulletin. 2022;174:113284. doi:10.1016/j.marpolbul.2021.113284

10. Jones ES, Ross SW, Robertson CM, Young CM. Distributions of microplastics and larger anthropogenic debris in Norfolk Canyon, Baltimore Canyon, and the adjacent continental slope (Western North Atlantic Margin, U.S.A.). Mar Pollut Bull. 2021;174:113047. doi:10.1016/j.marpolbul.2021.113047

11. Cutroneo L, Capello M, Domi A, et al. Microplastics in the abyss: a first investigation into sediments at 2443-m depth (Toulon, France). Environ Sci Pollut Res Int. 2022;29(6):9375–9385. doi:10.1007/s11356-021-17997-z

12. Mani T, Hauk A, Walter U, Burkhardt-Holm P. Microplastics profile along the Rhine River. Sci Rep. 2015;5:17988. doi:10.1038/srep17988

13. Scherer C, Weber A, Stock F, et al. Comparative assessment of microplastics in water and sediment of a large European river. Sci Total Environ. 2020;738:139866. doi:10.1016/j.scitotenv.2020.139866

14. Weiss L, Ludwig W, Heussner S, et al. The missing ocean plastic sink: Gone with the rivers. Science. 2021;373(6550):107–111. doi:10.1126/science.abe0290

15. Weber CJ, Opp C, Prume JA, Koch M, Andersen TJ, Chifflard P. Deposition and in-situ translocation of microplastics in floodplain soils. Sci Total Environ. 2022;819:152039. doi:10.1016/j.scitotenv.2021.152039

16. Golwala H, Zhang X, Iskander SM, Smith AL. Solid waste: An overlooked source of microplastics to the environment. Sci Total Environ. 2021;769:144581. doi:10.1016/j.scitotenv.2020.144581

17. Chamas A, Moon H, Zheng J, et al. Degradation Rates of Plastics in the Environment. ACS Sustainable Chem Eng. 2020;8(9):3494–3511. doi:10.1021/acssuschemeng.9b06635

18. Stock V, Böhmert L, Lisicki E, et al. Uptake and effects of orally ingested polystyrene microplastic particles in vitro and in vivo. Arch Toxicol. 2019;93(7):1817–1833. doi:10.1007/s00204-019-02478-7

19. Perveen N, Kishore U, Al Aiyan A, Willingham AL, Mohteshamuddin K. Exploring the Innate Immunity in Invertebrates. Adv Exp Med Biol. 2025;1476:411–423. doi:10.1007/978-3-031-85340-1_16

20. Jiang X, Chang Y, Zhang T, Qiao Y, Klobučar G, Li M. Toxicological effects of polystyrene microplastics on earthworm (Eisenia fetida). Environmental Pollution. 2020;259:113896. doi:10.1016/j.envpol.2019.113896

21. Alaraby M, Abass D, Domenech J, Hernández A, Marcos R. Hazard assessment of ingested polystyrene nanoplastics in *Drosophila* larvae. Environ Sci: Nano. 2022;9(5):1845–1857. doi:10.1039/D1EN01199E

22. Prietl B, Meindl C, Roblegg E, Pieber TR, Lanzer G, Fröhlich E. Nano-sized and micro-sized polystyrene particles affect phagocyte function. Cell Biol Toxicol. 2014;30(1):1–16. doi:10.1007/s10565-013-9265-y

23. Florance I, Ramasubbu S, Mukherjee A, Chandrasekaran N. Polystyrene nanoplastics dysregulate lipid metabolism in murine macrophages in vitro. Toxicology. 2021;458:152850. doi:10.1016/j.tox.2021.152850

24. Hu Q, Wang H, He C, Jin Y, Fu Z. Polystyrene nanoparticles trigger the activation of p38 MAPK and apoptosis via inducing oxidative stress in zebrafish and macrophage cells. Environmental Pollution. 2021;269:116075. doi:10.1016/j.envpol.2020.116075

25. Chen Y, Awasthi AK, Wei F, Tan Q, Li J. Single-use plastics: Production, usage, disposal, and adverse impacts. Science of The Total Environment. 2021;752:141772. doi:10.1016/j.scitotenv.2020.141772

26. Cutright DE, Beasley JD, Perez B. Histologic comparison of polylactic and polyglycolic acid sutures. Oral Surg Oral Med Oral Pathol. 1971;32(1):165–173. doi:10.1016/0030-4220(71)90265-9

27. Kulkarni RK, Moore EG, Hegyeli AF, Leonard F. Biodegradable poly(lactic acid) polymers. J Biomed Mater Res. 1971;5(3):169–181. doi:10.1002/jbm.820050305

28. Collin-Faure V, Vitipon M, Diemer H, Cianferani S, Darrouzet E, Rabilloud T. Biobased, Biodegradable but not bio-neutral: about the effects of polylactic acid nanoparticles on macrophages. Environ Sci: Nano. 2024;11:4102–4116. doi:10.1039/D4EN00335G

29. Gollwitzer H. Antibacterial poly(D,L-lactic acid) coating of medical implants using a biodegradable drug delivery technology. Journal of Antimicrobial Chemotherapy. 2003;51(3):585–591. doi:10.1093/jac/dkg105

30. Gil-Castell O, Badia JD, Ingles-Mascaros S, Teruel-Juanes R, Serra A, Ribes-Greus A. Polylactide-based self-reinforced composites biodegradation: Individual and combined influence of temperature, water and compost. Polymer Degradation and Stability. 2018;158:40–51. doi:10.1016/j.polymdegradstab.2018.10.017

31. Puchalski M, Siwek P, Panayotov N, Berova M, Kowalska S, Krucińska I. Influence of Various Climatic Conditions on the Structural Changes of Semicrystalline PLA Spun-Bonded Mulching Non-wovens during Outdoor Composting. Polymers. 2019;11(3):559. doi:10.3390/polym11030559

32. Mouhoubi R, Lasschuijt M, Ramon Carrasco S, Gojzewski H, Wurm FR. End-of-life biodegradation? how to assess the composting of polyesters in the lab and the field. Waste Management. 2022;154:36–48. doi:10.1016/j.wasman.2022.09.025

33. Wolf MH, Gil-Castell O, Cea J, Carrasco JC, Ribes-Greus A. Degradation of Plasticised Poly(lactide) Composites with Nanofibrillated Cellulose in Different Hydrothermal Environments. J Polym Environ. 2023;31(5):2055–2072. doi:10.1007/s10924-022-02711-y

34. Banaei G, García-Rodríguez A, Tavakolpournegari A, et al. The release of polylactic acid nanoplastics (PLA-NPLs) from commercial teabags. Obtention, characterization, and hazard effects of true-to-life PLA-NPLs. Journal of Hazardous Materials. 2023;458:131899. doi:10.1016/j.jhazmat.2023.131899

35. Collin-Faure V, Dalzon B, Devcic J, Diemer H, Cianférani S, Rabilloud T. Does size matter? A proteomics-informed comparison of the effects of polystyrene beads of different sizes on macrophages. Environ Sci: Nano. 2022;9(8):2827–2840. doi:10.1039/D2EN00214K

36. Meindl C, Öhlinger K, Zrim V, Steinkogler T, Fröhlich E. Screening for Effects of Inhaled Nanoparticles in Cell Culture Models for Prolonged Exposure. Nanomaterials (Basel). 2021;11(3):606. doi:10.3390/nano11030606

37. Domenech J, de Britto M, Velázquez A, et al. Long-Term Effects of Polystyrene Nanoplastics in Human Intestinal Caco-2 Cells. Biomolecules. 2021;11(10):1442. doi:10.3390/biom11101442

38. Domenech J, Villacorta A, Ferrer JF, et al. In vitro cell-transforming potential of secondary polyethylene terephthalate and polylactic acid nanoplastics. J Hazard Mater. 2024;469:134030. doi:10.1016/j.jhazmat.2024.134030

39. Collin-Faure V, Villacorta A, Diemer H, et al. Effects of true to life polyethylene terephthalate and polycaprolactone nanoparticles on macrophages under a repeated exposure mode. NanoImpact. 2026;41:100615. doi:10.1016/j.impact.2026.100615

40. Dalzon B, Torres A, Devcic J, Fenel D, Sergent JA, Rabilloud T. A Low-Serum Culture System for Prolonged in Vitro Toxicology Experiments on a Macrophage System. Front Toxicol. 2021;3:780778. doi:10.3389/ftox.2021.780778

41. Torres A, Collin-Faure V, Fenel D, Sergent JA, Rabilloud T. About the Transient Effects of Synthetic Amorphous Silica: An In Vitro Study on Macrophages. Int J Mol Sci. 2022;24(1):220. doi:10.3390/ijms24010220

42. Collin-Faure V, Boulée M, Diemer H, et al. A comparison of the effects of polystyrene and polycaprolactone nanoplastics on macrophages. Environ Sci: Nano. 2025;12:3990–4007. doi:10.1039/D5EN00074B

43. Muller L, Fornecker L, Chion M, et al. Extended investigation of tube-gel sample preparation: a versatile and simple choice for high throughput quantitative proteomics. Sci Rep. 2018;8(1):8260. doi:10.1038/s41598-018-26600-4

44. Cavazza C, Collin-Faure V, Pérard J, et al. Proteomic analysis of Rhodospirillum rubrum after carbon monoxide exposure reveals an important effect on metallic cofactor biosynthesis. J Proteomics. 2022;250:104389. doi:10.1016/j.jprot.2021.104389

45. Lyubimova T, Caglio S, Gelfi C, Righetti PG, Rabilloud T. Photopolymerization of polyacrylamide gels with methylene blue. Electrophoresis. 1993;14(1-2):1–2. doi:10.1002/elps.1150140108

46. Bouyssié D, Hesse AM, Mouton-Barbosa E, et al. Proline: an efficient and user-friendly software suite for large-scale proteomics. Valencia A, ed. Bioinformatics. 2020;36(10):3148–3155. doi:10.1093/bioinformatics/btaa118

47. Hammer O, Harper DAT, Ryan P. D. Paleontological statistics software package for education and data analysis. Palaeontologia Electronica. 2001;4:9pp.

48. Herrmann AG, Searcy JL, Le Bihan T, McCulloch J, Deighton RF. Total variance should drive data handling strategies in third generation proteomic studies. Proteomics. 2013;13:3251–3255. doi:10.1002/pmic.201300056

49. Huang DW, Sherman BT, Lempicki RA. Bioinformatics enrichment tools: paths toward the comprehensive functional analysis of large gene lists. Nucleic Acids Res. 2009;37:1–13.

50. Collin-Faure V, Vitipon M, Torres A, Tanyeres O, Dalzon B, Rabilloud T. The internal dose makes the poison: higher internalization of polystyrene particles induce increased perturbation of macrophages. Front Immunol. 2023;14:1092743. doi:10.3389/fimmu.2023.1092743

51. Perry SW, Norman JP, Barbieri J, Brown EB, Gelbard HA. Mitochondrial membrane potential probes and the proton gradient: a practical usage guide. Biotechniques. 2011;50:98–115.

52. Keij JF, Bell-Prince C, Steinkamp JA. Staining of mitochondrial membranes with 10-nonyl acridine orange, MitoFluor Green, and MitoTracker Green is affected by mitochondrial membrane potential altering drugs. Cytometry. 2000;39(3):203–210. doi:10.1002/(sici)1097-0320(20000301)39:3%3C203::aid-cyto5%3E3.0.co;2-z

53. Huang DW, Sherman BT, Lempicki RA. Systematic and integrative analysis of large gene lists using DAVID bioinformatics resources. Nat Protoc. 2009;4:44–57.

54. Vitipon M, Akingbagbohun E, Devime F, et al. Beyond the ink: cellular and molecular effects of iron-based pigments on macrophages. NanoImpact. 2025;39:100578. doi:10.1016/j.impact.2025.100578

55. Schwarzfischer M, Niechcial A, Lee SS, et al. Ingested nano- and microsized polystyrene particles surpass the intestinal barrier and accumulate in the body. NanoImpact. 2022;25:100374. doi:10.1016/j.impact.2021.100374

56. Meng X, Ge L, Zhang J, et al. Systemic effects of nanoplastics on multi-organ at the environmentally relevant dose: The insights in physiological, histological, and oxidative damages. Sci Total Environ. 2023;892:164687. doi:10.1016/j.scitotenv.2023.164687

57. Shin HS, Lee SH, Moon HJ, et al. Exposure to polystyrene particles causes anxiety-, depression-like behavior and abnormal social behavior in mice. J Hazard Mater. 2023;454:131465. doi:10.1016/j.jhazmat.2023.131465

58. Wang K, Du Y, Li P, et al. Nanoplastics causes heart aging/myocardial cell senescence through the Ca2+/mtDNA/cGAS-STING signaling cascade. J Nanobiotechnology. 2024;22(1):96. doi:10.1186/s12951-024-02375-x

59. Powell JJ, Ainley CC, Harvey RS, et al. Characterisation of inorganic microparticles in pigment cells of human gut associated lymphoid tissue. Gut. 1996;38(3):3. doi:10.1136/gut.38.3.390

60. Barguilla I, Domenech J, Rubio L, Marcos R, Hernández A. Nanoplastics and Arsenic Co-Exposures Exacerbate Oncogenic Biomarkers under an In Vitro Long-Term Exposure Scenario. Int J Mol Sci. 2022;23(6):2958. doi:10.3390/ijms23062958

61. Gutiérrez-García J, Egea R, Barguilla I, et al. Long-Term Exposure to Real-Life Polyethylene Terephthalate Nanoplastics Induces Carcinogenesis In Vitro. Environ Sci Technol. 2025;59(22):10891–10904. doi:10.1021/acs.est.5c01628

62. Morataya-Reyes M, Villacorta A, Gutiérrez-García J, et al. The long-term in vitro co-exposure of polyethylene terephthalate (PET) nanoplastics and cigarette smoke condensate exacerbates the induction of carcinogenic traits. J Hazard Mater. 2025;493:138359. doi:10.1016/j.jhazmat.2025.138359

63. Hashimoto D, Chow A, Noizat C, et al. Tissue-Resident Macrophages Self-Maintain Locally throughout Adult Life with Minimal Contribution from Circulating Monocytes. Immunity. 2013;38(4):792–804. doi:10.1016/j.immuni.2013.04.004

64. Murphy J, Summer R, Wilson AA, Kotton DN, Fine A. The Prolonged Life-Span of Alveolar Macrophages. Am J Respir Cell Mol Biol. 2008;38(4):380–385. doi:10.1165/rcmb.2007-0224RC

65. Bain CC, Bravo-Blas A, Scott CL, et al. Constant replenishment from circulating monocytes maintains the macrophage pool in the intestine of adult mice. Nat Immunol. 2014;15(10):929–937. doi:10.1038/ni.2967

66. Baranska A, Shawket A, Jouve M, et al. Unveiling skin macrophage dynamics explains both tattoo persistence and strenuous removal. J Exp Med. 2018;215(4):1115–1133. doi:10.1084/jem.20171608

67. Bowden DH. The alveolar macrophage. Environ Health Perspect. 1984;55:327–341. doi:10.1289/ehp.8455327

68. Ralph P, Nakoinz I. Phagocytosis and cytolysis by a macrophage tumour and its cloned cell line. Nature. 1975;257(5525):393–394. doi:10.1038/257393a0

69. Ruijter N, Soeteman-Hernández LG, Carrière M, et al. The State of the Art and Challenges of In Vitro Methods for Human Hazard Assessment of Nanomaterials in the Context of Safe-by-Design. Nanomaterials. 2023;13(3):472. doi:10.3390/nano13030472

70. Sarkans U, Gostev M, Athar A, et al. The BioStudies database—one stop shop for all data supporting a life sciences study. Nucleic Acids Research. 2018;46(D1):D1266–D1270. doi:10.1093/nar/gkx965

71. Vizcaino JA, Deutsch EW, Wang R, et al. ProteomeXchange provides globally coordinated proteomics data submission and dissemination. Nat Biotechnol. 2014;32(3):223–226.

