## Supplemental Table 1 for "Effects of polystyrene and polylactide nanoparticles on macrophages under a repeated exposure mode"

Supplementary Table 1: list of proteins detected and quantified by the proteomic screen

| accession | gene_name | description | protein_set_s | coverage | MW | #specific_seq | #peptides | Ctrl1 rep | Ctrl5 rep | Ctrl4 rep | Ctrl3 rep | Ctrl2 rep | PLA1 rep | PLA5 rep | PLA4 rep | PLA3 rep | PLA2 rep | PS1 rep | PS5 rep | PS3 rep | PS2 rep |
|---|---|---|---|---|---|---|---|---|---|---|---|---|---|---|---|---|---|---|---|---|---|
| P06238 | A2m | Alpha-2-macroglobulin Rattus norvegicus | 52.82 | 0.68 | 163785 | 1 | 1 | 2571648.5 | 1 | 1982401.88 | 808577.31 | 1 | 1169493.12 | 940893.44 | 1151829.12 | 1065884.685 | 1001323.06 | 948068.44 | 758197.92 | 877097.44 | 449427.88 |
| P58742 | Aaas | Aladin Mus musculus | 312.2 | 21.61 | 59431 | 7 | 7 | 5700100 | 6476691.5 | 5992078 | 5249330.5 | 6063287 | 5254438.5 | 5348615 | 5538208 | 5726840 | 5212821 | 5246579.5 | 6246079 | 6585944.5 | 5888169 |
| Q9D2R0 | Aacs | Acetoacetyl-CoA synthetase Mus musculus | 628.09 | 23.66 | 75200 | 13 | 13 | 19207743 | 19185077 | 16988625 | 13469001 | 15923373 | 12683798 | 15757592 | 19119417 | 14518748 | 12804036 | 13716428 | 11286769 | 14626861 | 13012759 |
| P0C1X8 | Aak1 | AP2-associated protein kinase 1 Rattus norvegicus | 96.52 | 4.47 | 103761 | 2 | 2 | 410858.03 | 1 | 323195.53 | 1 | 310884.16 | 446601.84 | 152201.83 | 672006.31 | 370865.16 | 324663.03 | 384982.93 | 414235.44 | 364929.41 | 375783.94 |
| Q8BGQ7 | Aars | Alanine--tRNA ligase, cytoplasmic Mus musculus | 2313.64 | 52.58 | 106909 | 31 | 32 | 75010617 | 80832313 | 81785641 | 98410385 | 88170257 | 92599121 | 87249009 | 79900193 | 83856273 | 91521169 | 81641337 | 77735553 | 79221569 | 87134577 |
| Q3THG9 | Aarsd1 | Alanyl-tRNA editing protein Aarsd1 Mus musculus | 299.59 | 27.18 | 44971 | 7 | 7 | 3747308 | 5152381 | 5212448 | 5635507 | 2850704.25 | 5129786 | 6033579 | 5475792.5 | 5176719 | 4895227.5 | 6019863 | 2812482 | 2367337.5 | 6043330 |
| Q9JKX4 | Aatf | Protein AATF Mus musculus | 219.63 | 9.13 | 59482 | 3 | 3 | 221213.81 | 370021.16 | 1 | 1 | 1 | 346790.16 | 523735.12 | 273830.84 | 286792.72 | 436566.72 | 276922.34 | 373468.16 | 1 | 1 |
| P06795 | Abcb1b | Multidrug resistance protein 1B Mus musculus | 196.97 | 4.7 | 140994 | 3 | 3 | 1318353.75 | 907364.62 | 1376753.75 | 947343.5 | 625802.44 | 794830.75 | 211937.36 | 634627.12 | 153047.05 | 684592.56 | 890311.38 | 486194.28 | 900684.25 | 978848 |
| Q61102 | Abcb7 | ATP-binding cassette sub-family B member 7, mitochondrial Mus musculus | 373.17 | 16.62 | 82581 | 2 | 6 | 5123448 | 2944719 | 4799254 | 3505946.25 | 3699019.5 | 5133512 | 4565987 | 4223905.5 | 4739214 | 3930049 | 4463696 | 4272578 | 5153818 | 4862506 |
| O35379 | Abcc1 | Multidrug resistance-associated protein 1 Mus musculus | 205.46 | 3.99 | 171185 | 4 | 4 | 1123700 | 1535258.5 | 1524385 | 1034228.31 | 1184643.88 | 1293299.12 | 584651.5 | 2529027.5 | 1555030.75 | 1329895.75 | 2391209 | 1514230.5 | 862776.5 | 1376532.38 |
| B2RX12 | Abcc3 | Canalicular multispecific organic anion transporter 2 Mus musculus | 264.86 | 5.38 | 169124 | 5 | 5 | 2378467.25 | 2236672.25 | 2379280.25 | 2060722.12 | 2223249.25 | 1781674.5 | 2650931.25 | 2580045 | 2553491.5 | 1966224.75 | 1773697.5 | 2582855 | 2151779.5 | 1697319.25 |
| D3ZHR2 | Abcd1 | ATP-binding cassette sub-family D member 1 Rattus norvegicus | 89.86 | 5.83 | 81938 | 2 | 2 | 1 | 1 | 613164.44 | 1 | 274918.53 | 553399.31 | 314440.41 | 492651.75 | 523429.97 | 579337.31 | 624485.23 | 580768.56 | 551884.94 | 740802.19 |
| P55096 | Abcd3 | ATP-binding cassette sub-family D member 3 Mus musculus | 45.76 | 1.82 | 75475 | 1 | 1 | 969135.44 | 753819.81 | 577534.38 | 775170.56 | 586447.5 | 878386.62 | 857286.69 | 1023502.62 | 606039.12 | 746025.88 | 836338.25 | 544119.06 | 807102.19 | 822504.25 |
| P61222 | Abce1 | ATP-binding cassette sub-family E member 1 Mus musculus | 1117.73 | 41.9 | 67314 | 18 | 18 | 15265587 | 13586266 | 14239910 | 19967685 | 20255783 | 15466849 | 14225522 | 13786508 | 15499469 | 14277301 | 20872541 | 17289219 | 20344087 | 14186561 |
| Q6P542 | Abcf1 | ATP-binding cassette sub-family F member 1 Mus musculus | 464.31 | 19.12 | 94945 | 9 | 9 | 7467059 | 5819978.5 | 6286385.5 | 6161664 | 6544938.5 | 5697914 | 4865801 | 5484054 | 5606019 | 5931955 | 5619560 | 3511276.5 | 6638671 | 5949459 |
| Q99LE6 | Abcf2 | ATP-binding cassette sub-family F member 2 Mus musculus | 352.77 | 14.97 | 71782 | 8 | 8 | 2561787 | 3042396.75 | 3490268.5 | 2401754.25 | 2664650.5 | 2547586.5 | 2240873.75 | 4126751.25 | 2487444 | 3329546.75 | 4321099.5 | 3535197.75 | 4177778.75 | 2658721.25 |
| Q66H39 | Abcf3 | ATP-binding cassette sub-family F member 3 Rattus norvegicus | 49.48 | 2.96 | 79856 | 1 | 1 | 501458.97 | 405761.75 | 536086.56 | 779650.12 | 269166.16 | 344104.53 | 1 | 1 | 303409.12 | 279257.38 | 272382.31 | 443527.44 | 218958.05 | 215270.86 |
| Q6AYT7 | Abhd12 | Monoacylglycerol lipase ABHD12 Rattus norvegicus 2 | 471.79 | 31.41 | 45296 | 9 | 9 | 8299984.5 | 5311613 | 9413133 | 7260250 | 5648225 | 7597999.5 | 7526677 | 9605509 | 10941961 | 11722427 | 6487724 | 8979812 | 12932866 | 7973497.5 |
| Q8VCR7 | Abhd14b | Protein ABHD14B Mus musculus | 102.13 | 15.24 | 22451 | 3 | 3 | 1861727.88 | 3702895 | 1522411.5 | 2582999 | 1558661.25 | 3105555.5 | 1714127.62 | 2027791.62 | 2691876 | 2627837 | 3260470.25 | 1670226.62 | 1722154.75 | 1714735.88 |
| Q8CBW3 | Abi1 | Abl interactor 1 Mus musculus | 85.4 | 4.99 | 52288 | 2 | 2 | 4001681.75 | 4286474.5 | 4502418 | 3655020.5 | 4132653.75 | 4938977.5 | 4619476 | 5560154 | 4280459 | 4232544 | 3703317 | 3945565.5 | 4557025 | 4248989 |
| Q4JIM5 | Abl2 | Tyrosine-protein kinase ABL2 Mus musculus | 43.38 | 1.52 | 128196 | 1 | 1 | 1 | 251990.64 | 195264.5 | 1 | 1 | 1 | 1 | 176132.41 | 180381.3 | 1 | 138011.42 | 1 | 145515.44 | 1 |
| Q5SSL4 | Abr | Active breakpoint cluster region-related protein Mus musculus | 413.1 | 12.81 | 97667 | 8 | 8 | 3828630 | 4065722.5 | 4025891.75 | 1980865 | 4399927.5 | 4683297.5 | 5323579 | 8079637.5 | 7000732 | 3856223.75 | 5715948 | 3816959.75 | 6813539.5 | 7111839.5 |
| Q4KML4 | Abracl | Costars family protein ABRACL Mus musculus | 186.33 | 46.91 | 9030 | 3 | 3 | 29953261 | 21856413 | 18335557 | 21333597 | 19702327 | 20495383 | 18806181 | 23950173 | 19439411 | 18504031 | 18780621 | 15998573 | 16608635 | 18579929 |
| Q3TCJ1 | Abraxas2 | BRISC complex subunit Abraxas 2 Mus musculus | 49.2 | 2.89 | 46943 | 1 | 1 | 2015625 | 2341368 | 2138161.5 | 1699554 | 1894750.38 | 1502298.88 | 1606693.88 | 1780526.12 | 1559051.5 | 1319721.12 | 1701372.75 | 1670824.12 | 1509087.25 | 2086443.75 |
| Q921H8 | Acaa1a | 3-ketoacyl-CoA thiolase A, peroxisomal Mus musculus | 923.52 | 52.12 | 43953 | 13 | 13 | 42564881 | 40742409 | 43531681 | 41126677 | 34604249 | 37010741 | 44085381 | 41522909 | 37632613 | 37608961 | 32063441 | 33557013 | 38465265 | 35878349 |
| Q8BWT1 | Acaa2 | 3-ketoacyl-CoA thiolase, mitochondrial Mus musculus | 346.08 | 26.2 | 41830 | 6 | 6 | 8748761 | 11940587 | 12142917 | 9079196 | 9597698 | 10561389 | 9644347 | 10413639 | 13382588 | 12256088 | 10274625 | 9610003 | 11361908 | 8402716 |
| P11497 | Acaca | Acetyl-CoA carboxylase 1 Rattus norvegicus | 978.75 | 11.81 | 265194 | 17 | 22 | 16053941 | 14432279 | 18560953 | 15344241 | 16481721 | 18526585 | 17263955 | 18979493 | 20570223 | 17986053 | 17928801 | 19140561 | 16984357 | 21050987 |
| E9Q4Z2 | Acacb | Acetyl-CoA carboxylase 2 Mus musculus | 244.8 | 2.98 | 275750 | 1 | 6 | 2287776 | 2000558 | 2973958.5 | 2163551.5 | 1704186.88 | 1875639 | 1470689.25 | 3369095.5 | 2760222 | 2067019.12 | 1841344.25 | 2671208.75 | 1600026.38 | 4810538.5 |
| Q9D7B6 | Acad8 | Isobutyryl-CoA dehydrogenase, mitochondrial Mus musculus | 37.13 | 2.42 | 45020 | 1 | 1 | 1885928.75 | 2224785.5 | 1598112.25 | 1778057.62 | 1712586.12 | 1683622 | 1381464.25 | 1652782.75 | 1424091 | 1764484.62 | 738271.62 | 1670079 | 1874508.5 | 1714635.38 |
| Q8JZN5 | Acad9 | Acyl-CoA dehydrogenase family member 9, mitochondrial Mus musculus | 741.86 | 28 | 68722 | 13 | 13 | 21458219 | 19486827 | 26590593 | 19012471 | 21718827 | 19661759 | 23867921 | 26553827 | 27996773 | 22943279 | 21503433 | 25226857 | 22563911 | 23826937 |
| P51174 | Acadl | Long-chain specific acyl-CoA dehydrogenase, mitochondrial Mus musculus | 970.14 | 50 | 47908 | 4 | 16 | 129661433 | 123162641 | 115800057 | 121847297 | 123761337 | 117184193 | 116301633 | 136743521 | 125887697 | 107687865 | 121864313 | 116315209 | 113887121 | 111874425 |
| P45952 | Acadm | Medium-chain specific acyl-CoA dehydrogenase, mitochondrial Mus musculus | 444.97 | 28.5 | 46481 | 9 | 9 | 42677457 | 50715201 | 46500937 | 45997517 | 47525865 | 40269057 | 44646721 | 39783081 | 44266085 | 38346169 | 45339965 | 44181925 | 44745873 | 49146833 |
| P15651 | Acads | Short-chain specific acyl-CoA dehydrogenase, mitochondrial Rattus norvegicus | 215.77 | 14.81 | 44765 | 5 | 5 | 4089614.5 | 4404529 | 4467067.5 | 4722528 | 3925899 | 3653320 | 3944372.5 | 4341742 | 4233071 | 4086744 | 3842311 | 4013600.5 | 4215407 | 4170792.5 |
| Q9DBL1 | Acadsb | Short/branched chain specific acyl-CoA dehydrogenase, mitochondrial Mus musculus | 542.63 | 33.8 | 47874 | 9 | 9 | 9718412 | 9999263 | 12876759 | 11010008 | 9727089 | 13291001 | 14581746 | 14659703 | 13334390 | 13814395 | 13384975 | 10085830 | 14800241 | 13585437 |
| P50544 | Acadvl | Very long-chain specific acyl-CoA dehydrogenase, mitochondrial Mus musculus | 610.36 | 27.9 | 70875 | 11 | 11 | 7397203.5 | 6852240 | 8378622.5 | 7202088.5 | 7427555 | 7845857 | 8131071 | 6441970 | 7907080.5 | 8097252 | 7757683 | 7495602 | 7829288 | 7724013 |
| Q5FVC7 | Acap2 | Arf-GAP with coiled-coil, ANK repeat and PH domain-containing protein 2 Rattus norvegicu | 88.33 | 2.47 | 87230 | 1 | 1 | 468502.62 | 649814.81 | 542981.44 | 733051.56 | 620654.19 | 1 | 1 | 463504.91 | 1 | 672407.12 | 522415.94 | 414957.62 | 503000.38 | 616751.62 |
| Q8QZT1 | Acat1 | Acetyl-CoA acetyltransferase, mitochondrial Mus musculus | 601.7 | 31.84 | 44816 | 10 | 10 | 21018019 | 20848719 | 21482667 | 24846375 | 24763999 | 21658727 | 25595085 | 22712881 | 25355313 | 27060283 | 26085471 | 27650245 | 27050899 | 25543849 |
| Q8CAY6 | Acat2 | Acetyl-CoA acetyltransferase, cytosolic Mus musculus | 467.61 | 27.46 | 41298 | 6 | 6 | 6614755.5 | 6744047 | 6972167.5 | 7411894 | 7119811.5 | 6794511 | 6812571.5 | 6689108 | 5786914.5 | 8032716.5 | 8547801 | 6406017.5 | 6695768 | 7902625 |
| Q8BMP6 | Acbd3 | Golgi resident protein GCP60 Mus musculus | 311.62 | 13.71 | 60181 | 4 | 4 | 2446929.5 | 2741295 | 4110402.75 | 2876475 | 2201998 | 2679826 | 3059900 | 3112054.5 | 3989313.5 | 3439183 | 2955443 | 3490148.5 | 3262992 | 3659143 |
| Q9D061 | Acbd6 | Acyl-CoA-binding domain-containing protein 6 Mus musculus | 54.8 | 4.26 | 30887 | 1 | 1 | 143866.42 | 238036.62 | 219430.64 | 136224.08 | 251276.25 | 247858.5 | 185758.02 | 213275.91 | 276135.84 | 209627.7 | 162376.89 | 209821.42 | 164585.73 | 208343.84 |
| Q9JIX8 | Acin1 | Apoptotic chromatin condensation inducer in the nucleus Mus musculus | 469.93 | 8.07 | 150719 | 9 | 9 | 16880435 | 17403197 | 18043895 | 15951028 | 18141687 | 18114653 | 17049495 | 18489553 | 17471947 | 18735271 | 17214949 | 17569353 | 17476629 | 16647445 |
| Q91V92 | Acly | ATP-citrate synthase Mus musculus | 2898.74 | 53.99 | 119728 | 4 | 45 | 82236001 | 82837753 | 88651937 | 93847417 | 80978689 | 100457705 | 101209593 | 99180121 | 92174977 | 100754649 | 82173609 | 91873073 | 92448697 | 92403353 |
| P28271 | Aco1 | Cytoplasmic aconitate hydratase Mus musculus | 561.96 | 18.56 | 98126 | 10 | 10 | 14127157 | 14481343 | 16563087 | 16485249 | 16188815 | 14904103 | 14081163 | 13739234 | 14113963 | 14670862 | 14140976 | 14560370 | 14895008 | 15012376 |
| Q99KI0 | Aco2 | Aconitate hydratase, mitochondrial Mus musculus | 1715.58 | 40.38 | 85464 | 27 | 27 | 113500217 | 122642265 | 125108009 | 117830081 | 128722697 | 132302993 | 136039729 | 129805801 | 140362865 | 142224657 | 130724961 | 130182385 | 142546593 | 139967137 |
| P54987 | Acod1 | Cis-aconitate decarboxylase Mus musculus | 98.01 | 6.97 | 53758 | 2 | 2 | 516474.31 | 509054.03 | 870485.12 | 464057.28 | 567807.75 | 245930.42 | 232549.61 | 453477.31 | 220441.86 | 187289.31 | 236652.28 | 371332.25 | 532760.5 | 219037.17 |
| Q9CQR4 | Acot13 | Acyl-coenzyme A thioesterase 13 Mus musculus | 140.56 | 28.57 | 15183 | 3 | 3 | 2040353.5 | 1102507.12 | 3148655.5 | 2605679.25 | 984908 | 3384609.75 | 3623302 | 3133603 | 2026862.25 | 3145810.5 | 3116961.75 | 3338680 | 3112503 | 4192040 |
| O55171 | Acot2 | Acyl-coenzyme A thioesterase 2, mitochondrial Rattus norvegicus | 142.45 | 8.83 | 49701 | 3 | 3 | 550374.44 | 402008.53 | 423205.84 | 959802.75 | 493726.22 | 1203795 | 566856.19 | 449734.28 | 606925.19 | 1256344.88 | 939486.38 | 562261.44 | 679472 | 336698.59 |
| Q91V12 | Acot7 | Cytosolic acyl coenzyme A thioester hydrolase Mus musculus | 440.58 | 23.88 | 42537 | 7 | 7 | 8168401 | 15032150 | 13087661 | 8471117 | 12175181 | 14076377 | 14574925 | 15465647 | 14691127 | 9755983 | 13454787 | 15379133 | 14135539 | 14769985 |
| Q9R0X4 | Acot9 | Acyl-coenzyme A thioesterase 9, mitochondrial Mus musculus | 207.23 | 13.67 | 50560 | 4 | 4 | 3850710 | 1346032 | 2549555.25 | 2736440.5 | 2932182 | 3083578.25 | 2352073.5 | 1653214.38 | 1822222.88 | 2112967.25 | 3411385.5 | 3802070.25 | 4370973.5 | 3491818.5 |
| Q9EPL9 | Acox3 | Peroxisomal acyl-coenzyme A oxidase 3 Mus musculus | 163.39 | 6.43 | 78404 | 2 | 3 | 2464755.25 | 2313931.75 | 2644338.25 | 1952754.62 | 2369746 | 2825278.5 | 2387081.5 | 2392145 | 2414242.75 | 2425126.75 | 2512192.5 | 2683649.25 | 2810664.75 | 3137525.25 |
| P41498 | Acp1 | Low molecular weight phosphotyrosine protein phosphatase Rattus norvegicus | 359.06 | 51.27 | 18152 | 5 | 5 | 20943549 | 22746611 | 20750335 | 23506131 | 23260161 | 22222221 | 22596295 | 23491305 | 23851589 | 24872291 | 22915605 | 21078261 | 21534639 | 22963815 |
| P20611 | Acp2 | Lysosomal acid phosphatase Rattus norvegicus | 136.81 | 6.38 | 48320 | 3 | 3 | 2490458.25 | 4108593.75 | 4197776.5 | 4352387.5 | 4202293.5 | 3381419.75 | 2879859.25 | 4761546 | 3505359.5 | 3013919.5 | 3078982.25 | 2679709 | 3688125.25 | 2027044.12 |
| Q8BP40 | Acp6 | Lysophosphatidic acid phosphatase type 6 Mus musculus | 216.89 | 13.64 | 47625 | 4 | 4 | 5040321 | 4781136.5 | 5236165.5 | 5986873 | 6641041 | 5707575.5 | 5905777 | 5419394 | 5815748.5 | 5798750 | 5855807 | 5553029 | 5435129 | 4810598 |
| Q8VCW8 | Acsf2 | Acyl-CoA synthetase family member 2, mitochondrial Mus musculus | 435.77 | 26.02 | 67951 | 8 | 8 | 6112308.5 | 6232736.5 | 7270555 | 10534029 | 6490749 | 12155309 | 12857558 | 8889328 | 10873004 | 8065915 | 13921524 | 12177727 | 7943379 | 8942086 |
| P41216 | Acsl1 | Long-chain-fatty-acid--CoA ligase 1 Mus musculus | 1095.66 | 40.63 | 77951 | 19 | 21 | 35063401 | 37848509 | 35577673 | 29520365 | 34174989 | 30783355 | 32392091 | 34113621 | 34760153 | 29620123 | 30132581 | 29964471 | 32502633 | 31534959 |
| Q9CZW4 | Acsl3 | Long-chain-fatty-acid--CoA ligase 3 Mus musculus | 602.7 | 22.36 | 80492 | 11 | 12 | 103249249 | 90838929 | 98982273 | 96452401 | 114946073 | 92821481 | 91746625 | 62202541 | 97559153 | 103579377 | 113011801 | 126462985 | 95541649 | 106606713 |
| Q9QUJ7 | Acsl4 | Long-chain-fatty-acid--CoA ligase 4 Mus musculus | 1179.65 | 41.63 | 79077 | 22 | 23 | 77413993 | 74544825 | 73041393 | 67446297 | 74989745 | 62402901 | 57477041 | 59842265 | 57744017 | 56595649 | 76577297 | 82590777 | 63239773 | 76564017 |
| Q8JZR0 | Acsl5 | Long-chain-fatty-acid--CoA ligase 5 Mus musculus | 568.47 | 25.04 | 76206 | 10 | 12 | 16369395 | 17501567 | 17539055 | 17835385 | 16363560 | 15615403 | 16808933 | 14337916 | 16406352 | 14564169 | 17457021 | 17255829 | 17852789 | 16611258 |
| Q9QXG4 | Acss2 | Acetyl-coenzyme A synthetase, cytoplasmic Mus musculus | 491.28 | 18.54 | 78862 | 8 | 8 | 11705761 | 10209376 | 11267837 | 12468933 | 11857124 | 14087967 | 13800367 | 12570557 | 13475720 | 11850761 | 13505003 | 11802905 | 11221238 | 12561764 |
| P60710 | Actb | Actin, cytoplasmic 1 Mus musculus | 4409.75 | 98.4 | 41737 | 2 | 61 | 8796781569 | 10125157377 | 9494925313 | 9269221377 | 9283770369 | 9079844865 | 9468689409 | 9350021121 | 8999975937 | 9134427137 | 8860333057 | 8997057537 | 8713240577 | 9951795201 |
| Q8BFZ3 | Actbl2 | Beta-actin-like protein 2 Mus musculus | 1372.44 | 40.43 | 42004 | 1 | 23 | 3744656897 | 4347484673 | 3986426369 | 3925038081 | 3874814209 | 3885582849 | 4100939521 | 4168554497 | 3917593345 | 3682102017 | 3672310017 | 3746009345 | 3670357761 | 4151612673 |
| P68033 | Actc1 | Actin, alpha cardiac muscle 1 Mus musculus | 1950.93 | 55.97 | 42019 | 4 | 35 | 6090285569 | 7046930433 | 6816580609 | 6644038657 | 6506530305 | 6377374209 | 6635017217 | 6490405377 | 6256486913 | 6396615169 | 6202600961 | 6481373185 | 6102921217 | 6900223489 |
| P63259 | Actg1 | Actin, cytoplasmic 2 Rattus norvegicus | 4409.58 | 98.4 | 41793 | 2 | 61 | 8772901889 | 10096184321 | 9473689601 | 9240883201 | 9255208961 | 9060348929 | 9445854209 | 9330324481 | 8984396801 | 9106027521 | 8836078593 | 8976064513 | 8693986305 | 9928692737 |
| Q9Z2N8 | Actl6a | Actin-like protein 6A Mus musculus | 638.6 | 42.19 | 47448 | 13 | 13 | 20609735 | 27996013 | 27881319 | 25040099 | 22246899 | 26154061 | 23359251 | 21161609 | 24820093 | 22916471 | 20764805 | 26506493 | 25732019 | 27620383 |
| Q7TPR4 | Actn1 | Alpha-actinin-1 Mus musculus | 2014.61 | 43.5 | 103068 | 1 | 34 | 93833825 | 94325033 | 103145209 | 95372721 | 84718417 | 103720737 | 104869953 | 118580505 | 111266985 | 97087129 | 100674945 | 98627897 | 99062257 | 99698417 |
| P57780 | Actn4 | Alpha-actinin-4 Mus musculus | 3053.32 | 63.93 | 104977 | 2 | 49 | 209436129 | 203842033 | 213859377 | 201181649 | 193270273 | 201629009 | 212923185 | 227658513 | 215626017 | 209062993 | 214587745 | 197283649 | 201368625 | 209753921 |
| Q9QZB7 | Actr10 | Actin-related protein 10 Mus musculus | 201.02 | 7.43 | 46208 | 2 | 2 | 1627365.25 | 1443089.5 | 1543676.75 | 1814677.25 | 1030554.06 | 1776229.38 | 1560526 | 1564716.12 | 1663410 | 1579934.12 | 1781899.62 | 1512905.38 | 1747300.75 | 1693206.75 |
| P61164 | Actr1a | Alpha-centractin Mus musculus | 481.85 | 35.64 | 42614 | 4 | 9 | 6828401.5 | 47553585 | 13629455 | 15522523 | 15123009 | 12403303 | 15871631 | 15028503 | 14666267 | 12676868 | 13266028 | 14401003 | 15071499 | 15247256 |
| Q8R5C5 | Actr1b | Beta-centractin Mus musculus | 411.77 | 23.67 | 42281 | 2 | 7 | 4891109.5 | 14206069 | 12028197 | 12383290 | 12416062 | 11075346 | 11897747 | 12554422 | 11617989 | 11747934 | 11197191 | 11434217 | 11935367 | 12562327 |
| P61161 | Actr2 | Actin-related protein 2 Mus musculus | 1276.89 | 57.36 | 44761 | 19 | 19 | 164446689 | 194105441 | 166601889 | 156775073 | 177940385 | 167667041 | 177910529 | 181946945 | 168778881 | 171136097 | 164878561 | 167808657 | 161614817 | 175303457 |
| Q4V7C7 | Actr3 | Actin-related protein 3 Rattus norvegicus | 1283.95 | 61.24 | 47357 | 20 | 20 | 192447953 | 232151825 | 219706193 | 217601153 | 234542769 | 235476337 | 233003137 | 230150961 | 236717393 | 241031249 | 241731025 | 224184657 | 221544657 | 242045313 |
| Q99JW2 | Acy1 | Aminoacylase-1 Mus musculus | 424.76 | 28.92 | 45781 | 7 | 7 | 7514963 | 4298089.5 | 6403944.5 | 6028198 | 5054992.5 | 2668281.5 | 4369942 | 4973562.5 | 4238939 | 3666013.5 | 3885533 | 5359361 | 5295716.5 | 5136124.5 |
| P03958 | Ada | Adenosine deaminase Mus musculus | 161.96 | 9.66 | 39992 | 3 | 3 | 4214858.5 | 4943317 | 4736053 | 3611837.25 | 4193373.5 | 4243115 | 4362883.5 | 4818302 | 4352760.5 | 3762484 | 4505798 | 4466750.5 | 4647510 | 4519759 |
| O88839 | Adam15 | Disintegrin and metalloproteinase domain-containing protein 15 Mus musculus | 102.67 | 2.08 | 92664 | 1 | 1 | 202693.86 | 264608.88 | 222039.39 | 219734.345 | 189595.25 | 416076.72 | 317525.62 | 365111.09 | 336759.44 | 249386.42 | 318597.91 | 522001.75 | 333007.41 | 250278.95 |
| Q9Z0F8 | Adam17 | Disintegrin and metalloproteinase domain-containing protein 17 Mus musculus | 243.27 | 4.47 | 93056 | 3 | 3 | 352922.03 | 434138.34 | 751878.5 | 450703.3825 | 263874.66 | 734034.94 | 871016.44 | 1050635 | 1341085.75 | 495135.84 | 801569.75 | 201937.03 | 405257.72 | 1066279.62 |
| Q05910 | Adam8 | Disintegrin and metalloproteinase domain-containing protein 8 Mus musculus | 110.43 | 5.69 | 90046 | 3 | 3 | 1 | 1 | 1 | 1 | 1 | 1 | 1 | 1 | 232260.86 | 340338.09 | 1 | 448051.69 | 1 | 205252.55 |
| Q61072 | Adam9 | Disintegrin and metalloproteinase domain-containing protein 9 Mus musculus | 45.96 | 2.84 | 92079 | 1 | 1 | 559447.06 | 848215.62 | 617705.295 | 655406.69 | 407751.81 | 895548.56 | 708394.62 | 816062.31 | 969765.38 | 455616.47 | 691102.06 | 530578.25 | 683733.38 | 606880.44 |
| Q8C9W3 | Adamts2 | A disintegrin and metalloproteinase with thrombospondin motifs 2 Mus musculus | 36.67 | 0.58 | 135299 | 1 | 1 | 4741734.5 | 6284738.5 | 7280784.5 | 6663445.5 | 6696178.5 | 7433690 | 7263210.5 | 4651603.5 | 7200794 | 7172799 | 6819474.5 | 7251303.5 | 6708784.5 | 7223080.5 |
| Q99MU3 | Adar | Double-stranded RNA-specific adenosine deaminase Mus musculus | 220.57 | 4.58 | 130447 | 4 | 4 | 236765 | 130874.45 | 1194792 | 874160.94 | 135664.2 | 131269.34 | 739996.88 | 121864.5 | 121719.86 | 561231.25 | 124681.79 | 1045603.5 | 645397.19 | 605227.4933 |
| P51829 | Adcy7 | Adenylate cyclase type 7 Mus musculus | 104.34 | 2.64 | 122708 | 2 | 2 | 879758.88 | 466861.75 | 1108726.38 | 486994.72 | 359808.03 | 775061.75 | 694747.19 | 1392017.88 | 1015061.88 | 542185.69 | 635087.56 | 1211137.88 | 727483.38 | 668725.88 |
| Q63028 | Add1 | Alpha-adducin Rattus norvegicus | 35.3 | 3.67 | 80355 | 1 | 1 | 231378.59 | 1 | 188683.8 | 203929.55 | 1 | 1 | 1 | 155349.23 | 1 | 1 | 179429.7 | 1 | 116483.07 | 1 |
| P28474 | Adh5 | Alcohol dehydrogenase class-3 Mus musculus | 837.02 | 56.15 | 39548 | 14 | 14 | 36729549 | 41145705 | 37752825 | 35846885 | 40345165 | 40323753 | 40901505 | 40189149 | 41103089 | 42113037 | 39420241 | 37067089 | 39359941 | 41977605 |
| P55264 | Adk | Adenosine kinase Mus musculus | 231.14 | 21.05 | 40149 | 2 | 4 | 1433594.25 | 1214920.5 | 2043164.25 | 1937322.12 | 1291554.38 | 1184379.75 | 1841783.62 | 1702019.12 | 2464621.25 | 2144590.25 | 468561.88 | 2734752.25 | 2230824.25 | 1505156.5 |
| Q9JKL8 | Adnp | Activity-dependent neuroprotector homeobox protein Rattus norvegicus | 120.24 | 2.81 | 123626 | 3 | 3 | 1 | 1 | 1 | 996820.62 | 1 | 635765.56 | 1 | 1 | 1 | 1 | 1 | 1 | 1 | 1 |
| Q8VDL4 | Adpgk | ADP-dependent glucokinase Mus musculus | 209.99 | 11.29 | 53902 | 4 | 4 | 8361254.5 | 6984721.5 | 7357322.5 | 4871773 | 6078658 | 7506674.5 | 7888904.5 | 7512767 | 8186415.5 | 7122746.5 | 6671541.5 | 7044421 | 8009570 | 7325986.5 |
| P54923 | Adprh | [Protein ADP-ribosylarginine] hydrolase Mus musculus | 158.22 | 9.67 | 40068 | 3 | 3 | 4881424 | 3971885 | 4591478 | 4401517 | 4606919.5 | 4017540.5 | 3959218.75 | 4230903 | 3902701.5 | 3637002.5 | 4488538 | 4455396 | 4220719 | 4200843.5 |
| Q8CG72 | Adprhl2 | Poly(ADP-ribose) glycohydrolase ARH3 Mus musculus | 177.7 | 15.68 | 39414 | 4 | 4 | 3002238 | 4763269.5 | 4780709.5 | 4387344 | 4546301 | 4816668 | 5367212.5 | 5522041 | 5734347.5 | 5159242 | 4312480 | 4467942.5 | 4820463 | 4118514.75 |

| | | | | | | | | | | | | | | | | | | | | | |
|---|---|---|---|---|---|---|---|---|---|---|---|---|---|---|---|---|---|---|---|---|---|
| Q9JKV1 | Adrm1 | Proteasomal ubiquitin receptor ADRM1 Mus musculus | 226.99 | 16.46 | 42060 | 5 | 5 | 9169358 | 9070299 | 9524340 | 14480429 | 14783161 | 8417409 | 9031911 | 7830195 | 10541937 | 11369733 | 9508183 | 10380946 | 9557592 | 9670674 |
| P54822 | Adsl | Adenylosuccinate lyase Mus musculus | 813.72 | 48.76 | 54866 | 15 | 15 | 17095313 | 18313047 | 15103116 | 16259600 | 13355404 | 13919630 | 17354279 | 15899942 | 12754431 | 15842919 | 17165737 | 13555247 | 16612197 | 16073127 |
| P46664 | Adss | Adenylosuccinate synthetase isozyme 2 Mus musculus | 990.01 | 42.32 | 50021 | 16 | 17 | 42509193 | 49394993 | 54515937 | 40716057 | 39294245 | 41876085 | 44849825 | 46856737 | 33994985 | 43477885 | 37636629 | 42849817 | 37774537 | 36238897 |
| P28650 | Adssl1 | Adenylosuccinate synthetase isozyme 1 Mus musculus | 452.2 | 25.16 | 50254 | 8 | 8 | 24815345 | 19505815 | 7947449 | 16252192 | 5332461.5 | 9005155 | 16472182 | 28287295 | 9778385 | 8379258.5 | 12618939 | 8690363 | 9098495 | 8730673 |
| Q9QZQ1 | Afdn | Afadin Mus musculus | 78.26 | 0.99 | 206499 | 1 | 1 | 997290.12 | 1019074 | 1066013.75 | 898114.5 | 978138.25 | 1167800 | 1200748.75 | 1225231.38 | 1311944.25 | 1150000.25 | 963122.81 | 1358992.62 | 1119267 | 1147127.477 |
| Q920A7 | Afg3l1 | AFG3-like protein 1 Mus musculus | 223.41 | 8.11 | 87047 | 3 | 4 | 3144155 | 5008066.5 | 5348821.5 | 3231860.5 | 3044049.25 | 3210017 | 4003017.25 | 3977637.5 | 2840090.5 | 4397222.5 | 5346757 | 4332210 | 4970186.5 | 5075049 |
| Q8JZQ2 | Afg3l2 | AFG3-like protein 2 Mus musculus | 377.27 | 12.72 | 89519 | 8 | 9 | 4367963.5 | 4278519.5 | 4811836.5 | 5242098 | 4480939 | 5319758 | 5102967 | 4379725 | 5509449 | 4956074 | 5239304 | 4822039.5 | 5484184.5 | 4039066.5 |
| Q4KLH5 | Agfg1 | Arf-GAP domain and FG repeat-containing protein 1 Rattus norvegicus | 160.28 | 9.8 | 58170 | 3 | 3 | 862080.38 | 1442490.75 | 2389525.25 | 1366870.5 | 1249627.88 | 1476406.75 | 1445668.25 | 972610.75 | 2112506.5 | 1959222.5 | 1547691.88 | 1626013.5 | 1243567 | 1333089.75 |
| Q80WC7 | Agfg2 | Arf-GAP domain and FG repeat-containing protein 2 Mus musculus | 105.3 | 5.64 | 48968 | 2 | 2 | 735089.94 | 1670095.5 | 1333213.25 | 1370309 | 1242813.25 | 1517911.75 | 1175260.75 | 1252373.75 | 1314035 | 899117 | 923030.69 | 1801724 | 1336876.75 | 1014145.5 |
| Q9ESW4 | Agk | Acylglycerol kinase, mitochondrial Mus musculus | 114.69 | 6.65 | 46976 | 2 | 2 | 2610002 | 2282347 | 1911229.25 | 2118221.25 | 2412607.75 | 2351986 | 2227272.75 | 2302195.25 | 2048708.38 | 1955539 | 1987889.75 | 2122265 | 1648454.25 | 1659580.88 |
| Q8CJG0 | Ago2 | Protein argonaute-2 Mus musculus | 123.91 | 4.65 | 97304 | 3 | 3 | 376920.59 | 384765.5 | 519341.53 | 353858.5 | 449541.78 | 564732.19 | 548199.94 | 487260.09 | 632594.94 | 393408.53 | 528465.62 | 410613.38 | 356758.03 | 481321 |
| Q8C0I1 | Agps | Alkyldihydroxyacetonephosphate synthase, peroxisomal Mus musculus | 624.79 | 23.26 | 71684 | 11 | 11 | 6235871 | 5000361 | 8535396 | 3113150 | 5289479.5 | 9094774 | 9202690 | 9930530 | 8488279 | 11379509 | 4702857 | 10497420 | 8836275 | 9850253 |
| Q8CJF7 | Ahctf1 | Protein ELYS Mus musculus | 111.96 | 1.52 | 247646 | 2 | 2 | 91999.9 | 180918.52 | 92771.87 | 1 | 1 | 1 | 1 | 1 | 1 | 1 | 1 | 1 | 1 | 1 |
| P50247 | Ahcy | Adenosylhomocysteinase Mus musculus | 1645.95 | 73.15 | 47688 | 26 | 27 | 261898465 | 220012497 | 216327809 | 185914257 | 190846721 | 228664177 | 237433841 | 238370497 | 226486705 | 233903457 | 204937761 | 218081137 | 227094977 | 222611537 |
| B5DFN2 | Ahcyl1 | S-adenosylhomocysteine hydrolase-like protein 1 Rattus norvegicus | 241.82 | 16.15 | 53753 | 3 | 6 | 5914869 | 8225328.5 | 8047324 | 7341394.5 | 8698729 | 9572509 | 8872227 | 8347488.5 | 8380391.5 | 8857981 | 9192662 | 8321410 | 8609571 | 9353807 |
| Q68FL4 | Ahcyl2 | Putative adenosylhomocysteinase 3 Mus musculus | 147.13 | 7.18 | 66899 | 1 | 4 | 5611331 | 5789497 | 6573405.5 | 5489033.5 | 5941449 | 6133041.5 | 5582266 | 4743824.5 | 5190822 | 6214971 | 6131650 | 3942700 | 6277595 | 6413043 |
| Q8BK64 | Ahsa1 | Activator of 90 kDa heat shock protein ATPase homolog 1 Mus musculus | 710.49 | 57.4 | 38117 | 13 | 13 | 34326565 | 33961857 | 38453801 | 32439079 | 36210013 | 35771865 | 32779507 | 35842525 | 35070661 | 34553665 | 34184457 | 40275885 | 38968609 | 37301753 |
| O70200 | Aif1 | Allograft inflammatory factor 1 Mus musculus | 191.28 | 21.77 | 16911 | 2 | 2 | 6277422 | 5196575.5 | 3569797.5 | 3538031 | 3821591.25 | 2853234.5 | 3170443.5 | 4167430.75 | 2955057.75 | 3176482.75 | 3807434.75 | 2846131.5 | 3450931.5 | 3856774.5 |
| Q9Z0X1 | Aifm1 | Apoptosis-inducing factor 1, mitochondrial Mus musculus | 612.78 | 34.8 | 66765 | 13 | 13 | 16463537 | 16582204 | 19528453 | 20219293 | 23095575 | 18233591 | 18972297 | 15009254 | 18525133 | 17835163 | 18596183 | 19621897 | 16595023 | 18575875 |
| Q91VJ1 | Aim2 | Interferon-inducible protein AIM2 Mus musculus | 222.28 | 12.71 | 40155 | 4 | 4 | 3277398 | 2867726 | 3099296.5 | 3292299 | 2809503 | 2882544 | 2594525.25 | 3237006.5 | 3271054.75 | 3294081.25 | 3405462.25 | 3544333 | 3350711.75 | 3847142.5 |
| P31230 | Aimp1 | Aminoacyl tRNA synthase complex-interacting multifunctional protein 1 Mus musculus | 463.32 | 44.19 | 33997 | 8 | 8 | 7778655 | 8279096 | 8006657 | 9662687 | 7513502 | 7108828 | 7960055.5 | 6329900 | 8762497 | 10035231 | 7732882.5 | 8199481 | 8462626 | 7716257 |
| Q8R010 | Aimp2 | Aminoacyl tRNA synthase complex-interacting multifunctional protein 2 Mus musculus | 411.23 | 35.62 | 35378 | 5 | 5 | 2585745.75 | 3658902 | 2180424 | 2601406.5 | 3133163.75 | 2864451 | 3331634.25 | 3958411 | 2812083.75 | 3316007.75 | 2967155 | 3286956.5 | 3262234.5 | 3242433.5 |
| O08915 | Aip | AH receptor-interacting protein Mus musculus | 106.21 | 8.18 | 37605 | 2 | 2 | 3069253 | 4603496 | 4547992.5 | 2998881.25 | 3235235.25 | 4086435.5 | 4634313.5 | 4364326.5 | 4516267 | 4999169 | 4577910.5 | 4497684.5 | 4476793.5 | 4757338 |
| Q9WTP6 | Ak2 | Adenylate kinase 2, mitochondrial Mus musculus | 315.83 | 33.89 | 26469 | 6 | 6 | 20322113 | 21748331 | 21522219 | 22517561 | 21268597 | 24824239 | 24086941 | 25325895 | 23625907 | 26267657 | 24645387 | 24001141 | 23193215 | 24466869 |
| Q9WTP7 | Ak3 | GTP:AMP phosphotransferase AK3, mitochondrial Mus musculus | 211.67 | 25.11 | 25426 | 4 | 4 | 2491269.5 | 2715702 | 2521727.25 | 2693779.75 | 2454971.5 | 2595361.75 | 2755950 | 1790283.62 | 2421259.5 | 2346266.25 | 2704115.75 | 2268941.5 | 2471092.5 | 1329266.25 |
| O08715 | Akap1 | A-kinase anchor protein 1, mitochondrial Mus musculus | 43.88 | 1.52 | 92195 | 1 | 1 | 1 | 1 | 1 | 1 | 1 | 1 | 256094.66 | 220002.33 | 205825.53 | 1 | 201830.61 | 267186.75 | 1 | 1 |
| Q9DBR0 | Akap8 | A-kinase anchor protein 8 Mus musculus | 169.5 | 8.73 | 76294 | 3 | 3 | 1934461.5 | 3442657.5 | 3314414.5 | 4143002.5 | 4506964.5 | 1661564 | 4109248.5 | 3882028 | 3833059.5 | 3920550.5 | 3811264.25 | 1958121.62 | 3245483.5 | 3508860.5 |
| Q9JII6 | Akr1a1 | Alcohol dehydrogenase [NADP(+)] Mus musculus | 451.41 | 30.46 | 36587 | 9 | 9 | 17977653 | 19295517 | 19866239 | 15493695 | 17257193 | 16143472 | 16021025 | 13515514 | 15453446 | 17800587 | 17760123 | 17202335 | 19204343 | 18054359 |
| P45376 | Akr1b1 | Aldose reductase Mus musculus | 841.51 | 52.53 | 35732 | 15 | 15 | 190222113 | 212004353 | 199337377 | 196657073 | 209078065 | 161110577 | 174705681 | 163635473 | 179064801 | 182479457 | 194898833 | 208309905 | 201504657 | 194774241 |
| P21300 | Akr1b7 | Aldose reductase-related protein 1 Mus musculus | 262.98 | 15.82 | 35988 | 0 | 6 | 44295361 | 48995329 | 44048337 | 41327449 | 46377081 | 42969441 | 42042825 | 46063865 | 44739313 | 42475777 | 47922329 | 51357969 | 46366129 | 47523709 |
| Q5RJP0 | Akr1b7 | Aldose reductase-related protein 1 Rattus norvegicus | 348.17 | 17.72 | 36122 | 1 | 6 | 36651325 | 37410433 | 31974077 | 29491129 | 35974133 | 32741885 | 30778577 | 37351617 | 34255721 | 32101461 | 35825281 | 35757025 | 33054613 | 35435897 |
| P45377 | Akr1b8 | Aldose reductase-related protein 2 Mus musculus | 810.43 | 49.05 | 36121 | 6 | 13 | 58479109 | 60477413 | 54098949 | 51269453 | 57613033 | 52305225 | 53052513 | 59805649 | 56439781 | 53584985 | 59528589 | 61924853 | 58374965 | 57667429 |
| Q8CG45 | Akr7a2 | Aflatoxin B1 aldehyde reductase member 2 Rattus norvegicus | 61.95 | 4.09 | 40675 | 1 | 1 | 1 | 458293.44 | 694835.62 | 1 | 777704.19 | 565569.12 | 730556.44 | 344745.47 | 719809.12 | 827677.31 | 780510.12 | 884820.38 | 952128.69 | 684276.94 |
| P31750 | Akt1 | RAC-alpha serine/threonine-protein kinase Mus musculus | 170.55 | 10.62 | 55707 | 2 | 3 | 1051679.38 | 943814.19 | 878722.9375 | 769563.62 | 749834.56 | 458136.59 | 614655.94 | 820263.62 | 1679068.5 | 629130 | 1686174 | 1502030.29 | 1811631.25 | 1008285.62 |
| P47197 | Akt2 | RAC-beta serine/threonine-protein kinase Rattus norvegicus | 71.85 | 6.03 | 55543 | 0 | 2 | 645641.81 | 1 | 1 | 1 | 749834.56 | 458136.59 | 614655.94 | 1 | 746557.44 | 1 | 700310.75 | 1 | 582936.31 | 1 |
| P10518 | Alad | Delta-aminolevulinic acid dehydratase Mus musculus | 418.56 | 31.21 | 36024 | 6 | 7 | 7514570.5 | 8549166 | 8025639.5 | 7233266 | 7346623.5 | 6436315 | 8323216.5 | 8151862.5 | 7510325.5 | 7191854.5 | 7434749 | 8094906 | 7053081.5 | 7713927 |
| Q61490 | Alcam | CD166 antigen Mus musculus | 348.19 | 23.67 | 65092 | 8 | 8 | 3922109.75 | 3693753 | 4025636.5 | 1983142.12 | 5778949.5 | 3354314.5 | 2384545 | 4573160 | 4163940.5 | 3685425 | 4389511 | 5260661 | 4881504 | 3429013.75 |
| Q571I9 | Aldh16a1 | Aldehyde dehydrogenase family 16 member A1 Mus musculus | 328.47 | 11.47 | 84756 | 6 | 6 | 3665910 | 4166355.25 | 4120707 | 4882765 | 4007680 | 5533708.5 | 3025788.5 | 4405451 | 4013739.5 | 5332937 | 4526760 | 4683442 | 3714779 | 3565178.5 |
| Q9Z110 | Aldh18a1 | Delta-1-pyrroline-5-carboxylate synthase Mus musculus | 944.96 | 25.91 | 87266 | 16 | 17 | 27464813 | 31389645 | 29768825 | 29898957 | 27695143 | 30910441 | 23163219 | 27428377 | 27990553 | 25174783 | 25491405 | 33930405 | 25307475 | 28734841 |
| Q66HF8 | Aldh1b1 | Aldehyde dehydrogenase X, mitochondrial Rattus norvegicus | 111.29 | 4.82 | 57625 | 1 | 2 | 11603215 | 11684018 | 13051814 | 12502605 | 11462587 | 13215648 | 12968689 | 14213994 | 14635127 | 13888444 | 13951681 | 16082327 | 14318778 | 14370415 |
| Q8R0Y6 | Aldh1l1 | Cytosolic 10-formyltetrahydrofolate dehydrogenase Mus musculus | 426.66 | 15.08 | 98709 | 9 | 9 | 6059573.5 | 7205715.5 | 8514740 | 7488813 | 7148226 | 5200596 | 5326867 | 6174886.5 | 6405054.5 | 5357527 | 7682267 | 5702402 | 4765501.5 | 5613908 |
| P47738 | Aldh2 | Aldehyde dehydrogenase, mitochondrial Mus musculus | 1506.28 | 68.21 | 56538 | 24 | 25 | 182985393 | 204331585 | 209732513 | 200265169 | 210116641 | 223336913 | 224602097 | 222453585 | 231237777 | 223844209 | 247455457 | 248172561 | 250940065 | 254757489 |
| P47740 | Aldh3a2 | Fatty aldehyde dehydrogenase Mus musculus | 186.2 | 5.58 | 53971 | 2 | 2 | 1676653.38 | 1683196.62 | 1 | 486477.28 | 1 | 2287216.5 | 1570011.75 | 1309452.88 | 506534.38 | 445512.91 | 1987269.12 | 1816892.75 | 995444.19 | 840057.44 |
| Q80VQ0 | Aldh3b1 | Aldehyde dehydrogenase family 3 member B1 Mus musculus | 105.23 | 8.55 | 52292 | 2 | 2 | 2891052.25 | 3489360.75 | 2962967.5 | 2642890.75 | 2990492.75 | 1797691.25 | 1787047.38 | 1641571.12 | 1986077.5 | 1821045.38 | 1948526.25 | 2216548.5 | 2892468.5 | 2791471.75 |
| Q9DBF1 | Aldh7a1 | Alpha-aminoadipic semialdehyde dehydrogenase Mus musculus | 42.14 | 2.6 | 58861 | 1 | 1 | 959145.5 | 1088013.62 | 726082.81 | 815029.56 | 992774.5 | 1195092 | 1049457.88 | 1072075.25 | 844296.56 | 968668.5 | 1208569.75 | 1100472.5 | 1112100.62 | 1123956.38 |
| Q9JLJ2 | Aldh9a1 | 4-trimethylaminobutyraldehyde dehydrogenase Mus musculus | 1173.24 | 58.7 | 53515 | 20 | 21 | 53134621 | 59317857 | 56155481 | 64412857 | 62693641 | 59796585 | 60388769 | 58287577 | 61998197 | 61917117 | 62957341 | 60514917 | 54965397 | 59368177 |
| P05064 | Aldoa | Fructose-bisphosphate aldolase A Mus musculus | 2292.67 | 93.68 | 39356 | 27 | 36 | 1180237697 | 1044185217 | 1001171329 | 1153109889 | 1062884545 | 1194690945 | 1324459905 | 1179178369 | 1419098497 | 1384668801 | 1151078913 | 1103589377 | 1139217921 | 1112596353 |
| P05063 | Aldoc | Fructose-bisphosphate aldolase C Mus musculus | 1570.54 | 70.25 | 39395 | 16 | 23 | 264098593 | 255447873 | 239960209 | 228017905 | 262386721 | 291617729 | 297236609 | 293018529 | 308779137 | 315111137 | 293155009 | 242416257 | 242333217 | 276175521 |
| Q9DB25 | Alg5 | Dolichyl-phosphate beta-glucosyltransferase Mus musculus | 187.22 | 16.67 | 36791 | 4 | 4 | 7048862 | 4045596.5 | 8730081 | 7732013.5 | 4468782 | 7751763 | 7852320 | 6892287 | 6762993.5 | 6331149 | 5240725.5 | 7227564 | 7429627 | 2825804.25 |
| D3ZKD3 | Alkbh5 | RNA demethylase ALKBH5 Rattus norvegicus | 44.83 | 2.78 | 44453 | 1 | 1 | 770056.12 | 951099.19 | 902697.38 | 930684 | 958012.81 | 1075607.38 | 1239157.5 | 900158.06 | 1391791.25 | 1287506.38 | 1041651.44 | 1296742.12 | 1015747.44 | 708619.38 |
| P48999 | Alox5 | Arachidonate 5-lipoxygenase Mus musculus | 772.07 | 31.01 | 77967 | 15 | 15 | 21994769 | 25299785 | 21351103 | 17280719 | 21516281 | 19351801 | 20303479 | 20725145 | 22095675 | 20696275 | 18792425 | 21480737 | 19199609 | 19587775 |
| P20291 | Alox5ap | Arachidonate 5-lipoxygenase-activating protein Rattus norvegicus | 38.16 | 4.97 | 18070 | 1 | 1 | 1 | 273593.47 | 1 | 1529866.12 | 199316.78 | 1510337.38 | 1553615.5 | 1795744.5 | 265876.69 | 272368.31 | 1655682.62 | 1886929.75 | 744044.69 | 1296724.88 |
| O08583 | Alyref | THO complex subunit 4 Mus musculus | 452.53 | 40 | 26940 | 8 | 8 | 10586816 | 10303344 | 15216617 | 17258003 | 13529074 | 11336691 | 13854047 | 7566935 | 7200794.5 | 13274640 | 11146921 | 9554568 | 11643931 | 11189601 |
| Q8JZV7 | Amdhd2 | N-acetylglucosamine-6-phosphate deacetylase Mus musculus | 196.82 | 22.49 | 43501 | 5 | 5 | 6918675 | 8580279 | 6596870 | 8868889 | 8538026 | 7324317 | 7806967.5 | 6763082 | 7019898 | 7973238 | 8701013 | 7249427 | 5473545 | 7647705 |
| Q02356 | Ampd2 | AMP deaminase 2 Rattus norvegicus | 282.16 | 7.89 | 94787 | 6 | 6 | 4205408 | 3954117 | 3249520.5 | 5278274 | 4210056.5 | 3647650.75 | 2886895.5 | 3424186.75 | 4576892 | 6367781.5 | 3620423 | 3958486 | 5169392.5 | 3593774.25 |
| O08739 | Ampd3 | AMP deaminase 3 Mus musculus | 117.49 | 2.35 | 88652 | 2 | 2 | 3647009.5 | 1566312.5 | 1664332.25 | 1268097.25 | 1485964.62 | 2249554 | 1608768.62 | 1457203.75 | 2001025.5 | 1933679.62 | 1591406 | 1914763.25 | 2044820.38 | 1737058.25 |
| P53995 | Anapc1 | Anaphase-promoting complex subunit 1 Mus musculus | 85.35 | 1.7 | 215994 | 2 | 2 | 374076.56 | 272313.72 | 350240.84 | 205304.02 | 191318.83 | 341524.28 | 231829.83 | 426825.62 | 507368.34 | 204881.83 | 279340.31 | 400120.81 | 340253.09 | 228860.84 |
| Q9WVM3 | Anapc7 | Anaphase-promoting complex subunit 7 Mus musculus | 46.32 | 2.65 | 63021 | 1 | 1 | 170881.94 | 225512.88 | 205361.45 | 196778.5375 | 185357.88 | 130679.76 | 150193.03 | 158993.08 | 176718.61 | 178380.92 | 178774.42 | 202512.97 | 154957.02 | 184571.31 |
| Q810B6 | Ankfy1 | Rabankyrin-5 Mus musculus | 680.21 | 15.83 | 128653 | 11 | 11 | 7869792.5 | 13013437 | 8444439 | 12555536 | 9923451 | 16777439 | 5703096 | 10161557 | 8348284.5 | 8649663 | 14917240 | 6977231 | 12491625 | 6899708.5 |
| P58366 | Ankh | Progressive ankylosis protein homolog Rattus norvegicus | 85.84 | 4.27 | 54265 | 1 | 1 | 288447.91 | 143634.0975 | 98545.91 | 99877.84 | 87664.73 | 93527.75 | 1 | 1 | 1 | 63191.25 | 70649.23 | 1 | 136383.8 | 1 |
| Q8BTI7 | Ankrd52 | Serine/threonine-protein phosphatase 6 regulatory ankyrin repeat subunit C Mus musculus | 72.83 | 1.21 | 115032 | 1 | 1 | 191753.88 | 388408.34 | 336735.25 | 275266.265 | 184167.59 | 164778.98 | 182481.05 | 142039.55 | 201289.25 | 275863.84 | 208990.97 | 176886.86 | 245341.22 | 156511.3 |
| Q8K298 | Anln | Anillin Mus musculus | 544.79 | 13.47 | 122794 | 10 | 10 | 5513382 | 4529574 | 5273436 | 6093720.5 | 4500061 | 6563026.5 | 4951654.5 | 4614435 | 4089299 | 4875014 | 6144002.5 | 6118307 | 5706481 | 5409363 |
| O35381 | Anp32a | Acidic leucine-rich nuclear phosphoprotein 32 family member A Mus musculus | 648.42 | 44.53 | 28538 | 9 | 11 | 65718145 | 61853977 | 59668621 | 48358549 | 57908813 | 57155125 | 67671057 | 58620741 | 55423629 | 67439313 | 47381661 | 49648777 | 58090569 | 51173653 |
| Q9EST5 | Anp32b | Acidic leucine-rich nuclear phosphoprotein 32 family member B Mus musculus | 354.38 | 27.21 | 31079 | 5 | 7 | 45747313 | 26284427 | 27068685 | 22621367 | 27982597 | 29294155 | 28922425 | 25397677 | 28317601 | 29674509 | 23784413 | 21650777 | 25840303 | 26661685 |
| P97822 | Anp32e | Acidic leucine-rich nuclear phosphoprotein 32 family member E Mus musculus | 639.73 | 30.38 | 29622 | 7 | 8 | 30915129 | 28389643 | 29572829 | 22445685 | 25878157 | 27752363 | 28577233 | 26014917 | 28171833 | 29646561 | 20567095 | 22472889 | 28284539 | 27078137 |
| P97449 | Anpep | Aminopeptidase N Mus musculus | 868.1 | 21.74 | 109651 | 10 | 14 | 13964333 | 13958208 | 17584101 | 11778133 | 13488800 | 11916638 | 10822664 | 16590036 | 10487807 | 11239396 | 11690896 | 7986223 | 14399492 | 20082131 |
| P10107 | Anxa1 | Annexin A1 Mus musculus | 1826.49 | 66.47 | 38734 | 25 | 27 | 236659969 | 249002929 | 250478305 | 247777201 | 248102913 | 217017617 | 229036337 | 234542833 | 238513441 | 248290433 | 240241201 | 221934097 | 237650417 | 245278865 |
| P97384 | Anxa11 | Annexin A11 Mus musculus | 445.24 | 18.69 | 54079 | 8 | 8 | 13249033 | 14049104 | 13320381 | 12341820 | 13008711 | 10494576 | 13983861 | 13484079 | 13140501 | 11547279 | 13196794 | 13560549 | 13558590 | 12341479 |
| P07356 | Anxa2 | Annexin A2 Mus musculus | 1632.99 | 70.8 | 38676 | 25 | 25 | 413457761 | 402877729 | 423519713 | 427536897 | 441378049 | 471697665 | 521019233 | 505123873 | 568528513 | 556714433 | 470546401 | 450212353 | 440189633 | 446280225 |
| O35639 | Anxa3 | Annexin A3 Mus musculus | 1289.29 | 65.94 | 36384 | 22 | 22 | 79483705 | 73001289 | 74065025 | 79753297 | 70614609 | 79003297 | 76068929 | 76581961 | 79727089 | 85351825 | 78331433 | 77192673 | 73145761 | 77023273 |
| P97429 | Anxa4 | Annexin A4 Mus musculus | 1416.22 | 68.97 | 35916 | 11 | 21 | 283375777 | 255470785 | 235954209 | 264655041 | 254395585 | 276882465 | 256027569 | 297192385 | 284778817 | 274517505 | 265324625 | 251882161 | 259877505 | 268101345 |
| P55260 | Anxa4 | Annexin A4 Rattus norvegicus | 674.85 | 29.47 | 35849 | 1 | 10 | 100536425 | 85344593 | 78703625 | 87442833 | 84842137 | 94267841 | 84634073 | 100877665 | 91730945 | 88142657 | 91952161 | 80199361 | 86361073 | 89368401 |
| P48036 | Anxa5 | Annexin A5 Mus musculus | 2513.52 | 81.19 | 35752 | 31 | 34 | 850601857 | 746683393 | 836200641 | 775538113 | 756466305 | 846890817 | 758451201 | 960231809 | 822459457 | 848555137 | 783807937 | 853777217 | 829436481 | 789288961 |
| P14824 | Anxa6 | Annexin A6 Mus musculus | 1282.66 | 47.99 | 75885 | 25 | 26 | 57874777 | 59606777 | 50453753 | 57954801 | 48983085 | 58794525 | 59524529 | 71684929 | 67952121 | 62019637 | 53976017 | 57076297 | 53751265 | 58010533 |
| Q07076 | Anxa7 | Annexin A7 Mus musculus | 685.25 | 27.21 | 49925 | 12 | 12 | 35527049 | 36895441 | 38392973 | 38771653 | 40532297 | 36345065 | 36901117 | 32933171 | 35987541 | 36309717 | 36192025 | 38404845 | 35712541 | 37557533 |
| O35643 | Ap1b1 | AP-1 complex subunit beta-1 Mus musculus | 2073.93 | 49.63 | 103935 | 20 | 34 | 43716193 | 42529389 | 50980005 | 37876845 | 44609745 | 53359701 | 46317473 | 58898685 | 59589265 | 45151273 | 45600829 | 56944301 | 55968641 | 41221777 |
| P22892 | Ap1g1 | AP-1 complex subunit gamma-1 Mus musculus | 662.83 | 20.92 | 91350 | 14 | 14 | 26613081 | 26423947 | 23144335 | 23785225 | 26812109 | 26668767 | 23651867 | 24876577 | 26051537 | 23386249 | 24421953 | 26709647 | 26579441 | 25964041 |
| O88512 | Ap1g2 | AP-1 complex subunit gamma-like 2 Mus musculus | 65.18 | 2.4 | 87863 | 2 | 2 | 887665.88 | 570473.915 | 515224.12 | 454511.97 | 424493.69 | 312314.88 | 294088.03 | 1 | 364976.81 | 1 | 629513.94 | 475944.62 | 470210.22 | 448755.78 |
| P35585 | Ap1m1 | AP-1 complex subunit mu-1 Mus musculus | 302.74 | 23.17 | 48543 | 7 | 7 | 6224573.5 | 7832745.5 | 8689140 | 5319143.5 | 2913590.5 | 6776742.5 | 9302417 | 7933643.5 | 7140774.5 | 6994427 | 7465457.5 | 8682351 | 8039500 | 8093893 |
| P61967 | Ap1s1 | AP-1 complex subunit sigma-1A Mus musculus | 210.85 | 22.15 | 18733 | 2 | 2 | 2613343.5 | 2705115 | 2649773.5 | 1826789.88 | 2470660.5 | 2680452.5 | 2440622.75 | 2803788.75 | 2441603.25 | 1943095 | 2132503 | 2601829.25 | 2931198.5 | 2654084.5 |
| P17426 | Ap2a1 | AP-2 complex subunit alpha-1 Mus musculus | 1066.92 | 33.27 | 107664 | 14 | 21 | 19275267 | 18982619 | 19978341 | 24179009 | 19944041 | 16245064 | 21697277 | 17294671 | 20491075 | 20495659 | 20390841 | 23494887 | 26177555 | 20312457 |
| P18484 | Ap2a2 | AP-2 complex subunit alpha-2 Rattus norvegicus | 821.51 | 22.6 | 104045 | 9 | 16 | 16922133 | 16935013 | 15711798 | 14436035 | 14473565 | 16071232 | 15925430 | 17602593 | 17017929 | 14789274 | 15824650 | 16761382 | 16924289 | 16553157 |
| P62944 | Ap2b1 | AP-2 complex subunit beta Rattus norvegicus | 1888.98 | 48.56 | 104553 | 21 | 35 | 62308241 | 63961745 | 70913601 | 56382481 | 64318025 | 66827213 | 67064805 | 78495177 | 74444025 | 63376441 | 60108605 | 72371961 | 72260057 | 62523265 |
| P84091 | Ap2m1 | AP-2 complex subunit mu Mus musculus | 464.25 | 28.05 | 49655 | 10 | 10 | 12989671 | 13649889 | 12941551 | 11748584 | 9434192 | 11787104 | 11865069 | 12399760 | 9204751 | 10996847 | 12933143 | 12500523 | 13449590 | 12722041 |
| P62743 | Ap2s1 | AP-2 complex subunit sigma Mus musculus | 148.16 | 21.13 | 17018 | 3 | 3 | 4508175 | 4206873 | 3162647 | 3444398 | 2829109 | 3614864 | 3878249.75 | 4303824 | 4418225.5 | 3235657.5 | 3102008 | 3166741.75 | 3242775.5 | 4097622 |
| Q9Z1T1 | Ap3b1 | AP-3 complex subunit beta-1 Mus musculus | 311.82 | 5.79 | 122740 | 6 | 6 | 10678527 | 9409184 | 8660827 | 8851153 | 10371617 | 8993233 | 8819339 | 7970008.5 | 8311021 | 8148320 | 8984811 | 8143025 | 8579643 | 9018012 |
| O54774 | Ap3d1 | AP-3 complex subunit delta-1 Mus musculus | 512.81 | 10.59 | 135081 | 9 | 9 | 8338149 | 9023508 | 8500252 | 6781809 | 7314837.5 | 8443194 | 7900948 | 8280885 | 8757313 | 7084399 | 7981646.5 | 6014708.5 | 7883758 | 8089833 |
| P53676 | Ap3m1 | AP-3 complex subunit mu-1 Rattus norvegicus | 218.03 | 16.99 | 46981 | 4 | 4 | 1528154.38 | 1988228.25 | 2611411.5 | 1744980.25 | 1936171.25 | 2076224.75 | 1467675.62 | 1301327.5 | 2099081.75 | 1945564 | 1066948.88 | 2278913.5 | 1985734.88 | 2098520.5 |
| Q9DCR2 | Ap3s1 | AP-3 complex subunit sigma-1 Mus musculus | 52.29 | 7.77 | 21732 | 1 | 1 | 1 | 1 | 1 | 1 | 1 | 1 | 1 | 1 | 1 | 1 | 416111.22 | 448480.75 | 1 | 1 |

| | | | | | | | | | | | | | | | | | | | | | |
|---|---|---|---|---|---|---|---|---|---|---|---|---|---|---|---|---|---|---|---|---|---|
| Q80V94 | Ap4e1 | AP-4 complex subunit epsilon-1 Mus musculus | 78.97 | 1.78 | 124845 | 1 | 1 | 681580.38 | 454765.38 | 548853.5175 | 674298.69 | 384769.62 | 1 | 395299.84 | 373866.31 | 1 | 1 | 1 | 331713.78 | 1 | 1 |
| Q3TAP4 | Ap5b1 | AP-5 complex subunit beta-1 Mus musculus | 89.99 | 3.31 | 94679 | 2 | 2 | 495568.12 | 610638.62 | 653603.31 | 594973.88 | 572988.12 | 443462.97 | 505977.75 | 323197.91 | 537224.19 | 440818.47 | 559783.56 | 386920.81 | 510894.88 | 463886.44 |
| O88879 | Apaf1 | Apoptotic protease-activating factor 1 Mus musculus | 59.28 | 1.12 | 141003 | 1 | 1 | 410739.94 | 693278.06 | 611642.69 | 525197.38 | 571885.19 | 669924.5 | 534075.19 | 765750.94 | 577509.81 | 586399.94 | 604485.06 | 872207.19 | 604394 | 621138.38 |
| Q8R5A3 | Apbb1ip | Amyloid beta A4 precursor protein-binding family B member 1-interacting protein Mus mus | 77.88 | 6.12 | 74319 | 2 | 2 | 596750.38 | 539546.88 | 1 | 1 | 1 | 532156.75 | 552728.38 | 1 | 403743.88 | 599431.31 | 398084.25 | 1 | 1 | 1 |
| Q8R146 | Apeh | Acylamino-acid-releasing enzyme Mus musculus | 287.14 | 10.93 | 81581 | 5 | 5 | 7637083 | 8027463 | 7691495.5 | 5069089 | 8057225 | 8607333 | 7768928 | 5862674 | 9012273 | 8195693 | 5789273.5 | 9336231 | 7346988 | 8429543 |
| P28352 | Apex1 | DNA-(apurinic or apyrimidinic site) lyase Mus musculus | 595.7 | 40.69 | 35490 | 8 | 8 | 11758977 | 10707655 | 11214372 | 12466522 | 11296019 | 11175564 | 13123712 | 13366022 | 12616016 | 12894583 | 10512785 | 12262709 | 11564579 | 11666598 |
| O35841 | Api5 | Apoptosis inhibitor 5 Mus musculus | 768.84 | 43.06 | 56785 | 14 | 14 | 23142681 | 28603149 | 25478085 | 22739941 | 27038451 | 22475227 | 24934857 | 28730701 | 24908697 | 25102785 | 23670969 | 26664429 | 24354715 | 25795795 |
| Q9WVQ5 | Apip | Methylthioribulose-1-phosphate dehydratase Mus musculus | 35 | 6.22 | 26949 | 1 | 1 | 1 | 1187221.75 | 782819.5 | 1508583.25 | 1 | 1 | 637379.38 | 1 | 1 | 399477.16 | 505346.09 | 1117047.25 | 1 | 1 |
| P15943 | Aplp2 | Amyloid-like protein 2 Rattus norvegicus | 65.22 | 2.61 | 86883 | 1 | 1 | 281568.31 | 399158.03 | 118031.78 | 171600.78 | 122762.62 | 153210.47 | 265573.53 | 160664.56 | 110526.57 | 203415.08 | 164548.39 | 626421.44 | 102759.45 | 297909.76 |
| Q9D7N9 | Apmap | Adipocyte plasma membrane-associated protein Mus musculus | 331.15 | 23.61 | 46434 | 5 | 5 | 2389094.5 | 2061793.88 | 2761905.75 | 2761008.75 | 2897959.75 | 3085396.5 | 4163013.5 | 2711648.75 | 2410147 | 2844496.5 | 2887134.25 | 2675561 | 3401309.25 | 2811079.5 |
| P51908 | Apobec1 | C->U-editing enzyme APOBEC-1 Mus musculus | 150.25 | 18.78 | 27522 | 1 | 3 | 4181783 | 3128322 | 3258482 | 3074789.75 | 3974909.5 | 2983143.5 | 1326838.38 | 2388297 | 2709799 | 3244947.75 | 3665762.75 | 2977916.5 | 2662353.5 | 3459895 |
| Q8VBT6 | Apobr | Apolipoprotein B receptor Mus musculus | 1003.48 | 23.89 | 102705 | 16 | 16 | 22938505 | 36570777 | 27218489 | 30706401 | 31003593 | 27759441 | 27952053 | 30488389 | 29440633 | 28712053 | 27851391 | 27157259 | 25601209 | 29610879 |
| Q9DCZ4 | Apoo | MICOS complex subunit Mic26 Mus musculus | 109.78 | 20.2 | 22604 | 2 | 2 | 571167.81 | 388203.75 | 801746.06 | 435004.59 | 431366.28 | 918111.31 | 394634.91 | 1261841.12 | 799569.44 | 476930.09 | 629336.94 | 555185.25 | 452173.12 | 521406.84 |
| Q78IK4 | Apool | MICOS complex subunit Mic27 Mus musculus | 389.35 | 33.96 | 29261 | 6 | 6 | 8955794 | 9403101 | 9080211 | 6970131 | 10111369 | 10838823 | 12188252 | 10543495 | 9161167 | 12014285 | 10280277 | 10627857 | 9798840 | 9853381 |
| Q8K3G9 | Appl2 | DCC-interacting protein 13-beta Mus musculus | 37.43 | 3.93 | 73854 | 1 | 1 | 140612.2 | 224415.69 | 157621.38 | 164640.83 | 230919.08 | 165833.88 | 142075.64 | 201390.02 | 135565.36 | 133740.36 | 212335.38 | 173769.44 | 507641.12 | 166358.14 |
| P08030 | Aprt | Adenine phosphoribosyltransferase Mus musculus | 757.74 | 77.78 | 19724 | 10 | 11 | 70375441 | 67236081 | 71763953 | 76840985 | 74646585 | 72398873 | 72881209 | 69376737 | 73652777 | 74962241 | 72176985 | 73867169 | 73032905 | 72226089 |
| Q8CFQ3 | Aqr | RNA helicase aquarius Mus musculus | 96.52 | 2.5 | 170294 | 2 | 2 | 1 | 301020.62 | 305347.97 | 459881.59 | 1 | 343060.5 | 401866.38 | 687799.19 | 369148.47 | 296162.09 | 416875.88 | 388845.3333 | 458281.84 | 291378.28 |
| P04627 | Araf | Serine/threonine-protein kinase A-Raf Mus musculus | 95.75 | 4.64 | 67581 | 1 | 2 | 204934.8 | 156852.69 | 125078.04 | 328889.34 | 281851.28 | 295140.56 | 196447.69 | 153178.7 | 282714.59 | 175343.98 | 275873.03 | 193962.62 | 134409.17 | 183824.31 |
| Q4LDD4 | Arap1 | Arf-GAP with Rho-GAP domain, ANK repeat and PH domain-containing protein 1 Mus mus | 305.56 | 6.34 | 162276 | 6 | 6 | 4658516 | 4736144 | 1639542.5 | 5982300.5 | 5228927.5 | 5259733.5 | 4700928 | 4980438 | 3620461 | 2820946.5 | 4089119.25 | 4759267 | 4779686 | 4696521.5 |
| Q5XJY5 | Arcn1 | Coatomer subunit delta Mus musculus | 877.29 | 35.23 | 57229 | 16 | 16 | 19772421 | 21118951 | 21787121 | 21443305 | 21915279 | 22544067 | 20980317 | 20838267 | 21864999 | 21097653 | 18541639 | 23474327 | 21041875 | 19542593 |
| P84078 | Arf1 | ADP-ribosylation factor 1 Mus musculus | 1015.5 | 80.66 | 20697 | 3 | 16 | 139083505 | 125766337 | 118190753 | 113588457 | 106964153 | 124850249 | 113201801 | 150921073 | 133665321 | 111852321 | 115878753 | 132096697 | 121263745 | 115394665 |
| P84082 | Arf2 | ADP-ribosylation factor 2 Rattus norvegicus | 868.88 | 59.67 | 20746 | 1 | 13 | 113368641 | 100004065 | 92356241 | 84586289 | 82072665 | 99380089 | 88129481 | 122474001 | 105355553 | 88728257 | 87610529 | 102045241 | 93548593 | 89341921 |
| P61750 | Arf4 | ADP-ribosylation factor 4 Mus musculus | 735 | 60 | 20397 | 3 | 10 | 69729241 | 66907981 | 59064017 | 48535037 | 53583585 | 65962189 | 56673429 | 74494841 | 67511905 | 52030405 | 50823377 | 65279525 | 60768517 | 58935245 |
| P84083 | Arf5 | ADP-ribosylation factor 5 Rattus norvegicus | 792.64 | 65.56 | 20530 | 4 | 12 | 101121185 | 101248833 | 91550977 | 83105881 | 82470113 | 97075465 | 88550641 | 110168801 | 101770089 | 80316121 | 84416209 | 101148313 | 93152801 | 90688529 |
| P62331 | Arf6 | ADP-ribosylation factor 6 Mus musculus | 278.71 | 48 | 20082 | 5 | 5 | 9888003 | 7395230 | 6450047 | 6249971.5 | 7713709 | 9021354 | 7383165 | 9427039 | 8988807 | 8150426.5 | 7949840 | 8515543 | 8202797 | 7570287 |
| Q3MID3 | Arfgap2 | ADP-ribosylation factor GTPase-activating protein 2 Rattus norvegicus | 218.78 | 9.62 | 56556 | 4 | 4 | 1611204.88 | 3975532.5 | 2190533.5 | 465425.59 | 988052.31 | 962548.19 | 1204261 | 808087.19 | 1864894.12 | 1787862.25 | 1853396.5 | 666386.31 | 1592879 | 1015094.94 |
| D4A631 | Arfgef1 | Brefeldin A-inhibited guanine nucleotide-exchange protein 1 Rattus norvegicus | 206.15 | 2.65 | 207890 | 1 | 4 | 1543873 | 1527367.62 | 1536154.12 | 1412977 | 1764370.25 | 1554354.12 | 1587834.62 | 1885056.5 | 1696265.5 | 1510763.75 | 1762708.75 | 1419551 | 1503471.25 | 1769946.75 |
| A2A5R2 | Arfgef2 | Brefeldin A-inhibited guanine nucleotide-exchange protein 2 Mus musculus | 312.42 | 4.85 | 202240 | 3 | 6 | 2486530 | 2432411.5 | 2001258.38 | 2357806.25 | 2523400.25 | 2321056 | 2593670 | 2366164 | 2439105 | 2257284.25 | 2712360.25 | 2116048.75 | 2147996.75 | 2356546 |
| Q9JHU5 | Arfip1 | Arfaptin-1 Rattus norvegicus | 217.28 | 12.3 | 40779 | 3 | 3 | 4189904.25 | 2635163 | 3106042.25 | 3730031.75 | 5448298.5 | 3184125.5 | 2600223.5 | 5877065.5 | 2922926.5 | 3254913 | 4845095 | 2476006 | 4850850.5 | 3043858 |
| Q63055 | Arfrp1 | ADP-ribosylation factor-related protein 1 Rattus norvegicus 2 | 74.71 | 8.96 | 22659 | 1 | 1 | 609896.19 | 1 | 775940.44 | 1 | 896574.25 | 893475.31 | 561702.5 | 346706.44 | 696107 | 856068.69 | 1192839.12 | 1004777.69 | 973784.88 | 828399.31 |
| Q3UL36 | Arglu1 | Arginine and glutamate-rich protein 1 Mus musculus | 38.25 | 2.95 | 32887 | 1 | 1 | 1 | 1 | 13492843 | 444879.12 | 370253.31 | 340522.5 | 18266907 | 1 | 13684635 | 1 | 1 | 1 | 1 | 1 |
| Q5FWK3 | Arhgap1 | Rho GTPase-activating protein 1 Mus musculus | 352.45 | 25.28 | 50411 | 8 | 8 | 7618287.5 | 5292056 | 5819092 | 5097715.5 | 5488070 | 5231241.5 | 4866285.5 | 6227612 | 5283328 | 5547908 | 5396994.5 | 5747752.5 | 6037448 | 5633871 |
| Q3UIA2 | Arhgap17 | Rho GTPase-activating protein 17 Mus musculus | 444.6 | 14.89 | 92202 | 10 | 10 | 12156895 | 8757226 | 8017522 | 8870471 | 10846009 | 7692902 | 7969897.5 | 6999208 | 7397576 | 7272889 | 8430568 | 6067848.5 | 7748019 | 5477966 |
| Q8BYW1 | Arhgap25 | Rho GTPase-activating protein 25 Mus musculus | 636.68 | 23.61 | 73383 | 12 | 12 | 16615914 | 18968055 | 18833181 | 20017501 | 16429793 | 18539137 | 20055489 | 20091447 | 19671233 | 18440389 | 17797013 | 15676314 | 18233817 | 19343247 |
| Q640N3 | Arhgap30 | Rho GTPase-activating protein 30 Mus musculus | 311.04 | 5.99 | 120114 | 5 | 5 | 861896.06 | 966804 | 1010682.69 | 953408.6725 | 974251.94 | 1147957.75 | 967622.25 | 1267865.25 | 883143.31 | 812003.31 | 984507.69 | 872786.56 | 691169.25 | 1048298.5 |
| Q3TBD2 | Arhgap45 | Rho GTPase-activating protein 45 Mus musculus | 567.96 | 15.14 | 122902 | 11 | 11 | 9840119 | 10974965 | 11893690 | 12042698 | 12688360 | 12195328 | 12510469 | 10114447 | 12604651 | 12831445 | 12666524 | 12376337 | 12267763 | 12189398 |
| Q5XI73 | Arhgdia | Rho GDP-dissociation inhibitor 1 Rattus norvegicus | 883.41 | 80.39 | 23407 | 12 | 12 | 202342753 | 186281601 | 218119665 | 207268609 | 209968353 | 201379457 | 218415025 | 174440801 | 214995249 | 223634929 | 205715969 | 212228801 | 211417633 | 209361921 |
| Q61599 | Arhgdib | Rho GDP-dissociation inhibitor 2 Mus musculus | 530.49 | 71.5 | 22851 | 11 | 11 | 93330705 | 72605081 | 95821105 | 106086849 | 96515977 | 94879449 | 111075145 | 76527961 | 107577537 | 105055353 | 97351217 | 102169041 | 100233977 | 90804993 |
| Q61210 | Arhgef1 | Rho guanine nucleotide exchange factor 1 Mus musculus | 975.61 | 26.63 | 102805 | 18 | 18 | 14399199 | 18696363 | 17802819 | 19125075 | 15907942 | 14103252 | 18078791 | 21065131 | 17629375 | 17607741 | 17701511 | 16582406 | 17061053 | 15389596 |
| A2AWP8 | Arhgef10l | Rho guanine nucleotide exchange factor 10-like protein Mus musculus | 115.37 | 3.2 | 139987 | 3 | 3 | 1919286 | 1586151.38 | 1712004.5 | 2007021.5 | 1053128.5 | 1820814.62 | 1590486.25 | 683867.44 | 1830558.75 | 1435089 | 1945481.25 | 2138147.75 | 1507994.5 | 2230628 |
| Q60875 | Arhgef2 | Rho guanine nucleotide exchange factor 2 Mus musculus | 950.87 | 22.94 | 111974 | 18 | 18 | 21502987 | 18211187 | 19553369 | 20133037 | 22657689 | 20826867 | 13862235 | 15875779 | 18265075 | 20828739 | 19477807 | 20336897 | 19631343 | 18060955 |
| Q8K4I3 | Arhgef6 | Rho guanine nucleotide exchange factor 6 Mus musculus | 394.07 | 10.64 | 87051 | 6 | 7 | 8605478 | 16651154 | 9383427 | 8057653.5 | 8661356 | 8152410.5 | 7871337 | 8986289 | 8135512 | 7507531 | 8286484 | 7789160.5 | 7615862 | 9456906 |
| O55043 | Arhgef7 | Rho guanine nucleotide exchange factor 7 Rattus norvegicus | 409.57 | 17.18 | 73140 | 6 | 7 | 4854628 | 12482605 | 5101338 | 4127232 | 4401331 | 4290849.5 | 4294639.5 | 4518417 | 3788639.5 | 3960974.25 | 4261760.5 | 3839343.75 | 3752595.5 | 4578836 |
| A2BH40 | Arid1a | AT-rich interactive domain-containing protein 1A Mus musculus | 160.72 | 1.14 | 242092 | 0 | 2 | 27109421 | 27486317 | 30441325 | 29938315 | 29588977 | 24688835 | 24675491 | 25498973 | 25350019 | 27800611 | 27259883 | 25294327 | 26233273 | 29537277 |
| E9Q4N7 | Arid1b | AT-rich interactive domain-containing protein 1B Mus musculus | 145.35 | 1.43 | 237795 | 1 | 2 | 629789.69 | 1076270.62 | 663479.69 | 1036901.88 | 942483.75 | 1028769.69 | 922071.75 | 1134723.38 | 740019.12 | 725402.69 | 944417 | 675269.69 | 724790.38 | 851943.75 |
| E9Q7E2 | Arid2 | AT-rich interactive domain-containing protein 2 Mus musculus | 60.99 | 0.49 | 195988 | 1 | 1 | 1190090.12 | 779792.44 | 634920.94 | 854276.69 | 935487.62 | 612579.69 | 551698.62 | 737782.94 | 644899.88 | 770607.75 | 831674.12 | 735484.88 | 472223.72 | 591959.31 |
| Q9Z1K5 | Arih1 | E3 ubiquitin-protein ligase ARIH1 Mus musculus | 132.77 | 7.39 | 64017 | 2 | 2 | 1302469.5 | 1893415.38 | 1967524.62 | 1277822.88 | 1766508.88 | 1884985.25 | 1275357.38 | 1605773.5 | 1817335.38 | 1109184 | 979437.31 | 1858034.25 | 1559614.75 | 1400658.75 |
| P61211 | Arl1 | ADP-ribosylation factor-like protein 1 Mus musculus | 382.62 | 52.49 | 20412 | 6 | 7 | 14524514 | 15820364 | 15911993 | 10523001 | 13318608 | 14956121 | 15084565 | 17861835 | 15859038 | 15200875 | 11919045 | 14574822 | 16597886 | 14622397 |
| O08697 | Arl2 | ADP-ribosylation factor-like protein 2 Rattus norvegicus | 383.51 | 45.65 | 20836 | 5 | 5 | 2996704 | 3156385.75 | 5255592.5 | 5050121.5 | 3702653.25 | 5368991 | 4628059.5 | 4616334 | 3804540.75 | 4903664.5 | 3013494.25 | 5155463.5 | 5626385.5 | 3438916.25 |
| Q9WUL7 | Arl3 | ADP-ribosylation factor-like protein 3 Mus musculus | 462.95 | 53.3 | 20487 | 8 | 8 | 5054397.5 | 5894785 | 5649855.5 | 4968932.5 | 5805500 | 5679538.5 | 6202498 | 3982004 | 6185416.5 | 5042442 | 7033004 | 7086544.5 | 6270796 | 5761479.5 |
| P61208 | Arl4c | ADP-ribosylation factor-like protein 4C Mus musculus | 41.78 | 7.81 | 21487 | 1 | 1 | 1230139.12 | 2216677.25 | 1874746.12 | 2059655.88 | 1853685.25 | 2941661 | 2277054.75 | 1935899 | 1635682.5 | 1788366.62 | 1521741.88 | 1205209.75 | 1532801.75 | 1438604.5 |
| Q8R5J9 | Arl6ip5 | PRA1 family protein 3 Mus musculus | 77.99 | 10.64 | 21558 | 2 | 2 | 9217537 | 2853742.5 | 2801725.25 | 2138155.5 | 2987978.5 | 3747305.5 | 2282113.5 | 1356661.75 | 2590548.5 | 2817196.5 | 1715526.75 | 3204899.5 | 3484972.5 | 4168009.75 |
| Q8VEH3 | Arl8a | ADP-ribosylation factor-like protein 8A Mus musculus | 362.06 | 44.62 | 21390 | 3 | 7 | 19094271 | 21402879 | 19415433 | 15555749 | 13841333 | 17176025 | 19078585 | 18273241 | 20145805 | 21119145 | 17845333 | 18793927 | 19534789 | 18876185 |
| Q66HA6 | Arl8b | ADP-ribosylation factor-like protein 8B Rattus norvegicus | 540.17 | 61.83 | 21539 | 6 | 10 | 34705193 | 28682611 | 34539589 | 33500075 | 34521117 | 27450573 | 37513605 | 37479233 | 36985337 | 35889145 | 33046965 | 37005477 | 37287949 | 35338325 |
| Q9D7A8 | Armc1 | Armadillo repeat-containing protein 1 Mus musculus | 57.15 | 9.57 | 31247 | 1 | 1 | 409928.34 | 566910.56 | 1 | 1 | 341695.69 | 432694.22 | 409780.03 | 1 | 477262.94 | 1 | 496383.97 | 297931.78 | 445763.91 | 425242.28 |
| Q9D0L7 | Armc10 | Armadillo repeat-containing protein 10 Mus musculus | 172.68 | 15.36 | 33311 | 3 | 3 | 1656080 | 1723679.12 | 1740246.5 | 1079363 | 1922319.38 | 1423307.75 | 1821133.12 | 2176469.5 | 1976578.75 | 1428418 | 2151719.75 | 2111165 | 2096870.75 | 2413043.25 |
| Q8BNU0 | Armc6 | Armadillo repeat-containing protein 6 Mus musculus | 134.17 | 8.33 | 50683 | 2 | 2 | 621959.12 | 564142.25 | 682003.25 | 615147.5 | 650408.81 | 1252661.75 | 669301.62 | 230061.78 | 1372714.75 | 1015539.5 | 737465.62 | 696961.2067 | 647578.94 | 705839.06 |
| Q9DBR3 | Armc8 | Armadillo repeat-containing protein 8 Mus musculus | 104.88 | 4.75 | 75364 | 2 | 2 | 1089267.25 | 789229.56 | 1345845.12 | 875397.12 | 936497.12 | 858203.56 | 959304.25 | 819251.81 | 917430 | 989764.94 | 838419 | 996229 | 1010502.62 | 1267318 |
| Q9WV32 | Arpc1b | Actin-related protein 2/3 complex subunit 1B Mus musculus | 931.68 | 55.65 | 41064 | 16 | 16 | 58040457 | 61409085 | 63811549 | 67494473 | 73469161 | 66690917 | 70015033 | 56656381 | 72151521 | 82448337 | 73903393 | 70634353 | 67816137 | 77939033 |
| Q9CVB6 | Arpc2 | Actin-related protein 2/3 complex subunit 2 Mus musculus | 1177.12 | 70.67 | 34357 | 19 | 19 | 127419329 | 119070625 | 129353473 | 118420425 | 118105409 | 133355161 | 114365353 | 151557953 | 136897377 | 130196513 | 113599105 | 137389697 | 132679697 | 117926513 |
| Q9JM76 | Arpc3 | Actin-related protein 2/3 complex subunit 3 Mus musculus | 481.08 | 53.37 | 20525 | 8 | 8 | 84300433 | 103555009 | 86298033 | 81326905 | 86582385 | 82149529 | 88646665 | 97900409 | 84841113 | 80635457 | 88629113 | 84246601 | 86306185 | 94335585 |
| P59999 | Arpc4 | Actin-related protein 2/3 complex subunit 4 Mus musculus | 400.77 | 54.17 | 19667 | 7 | 7 | 155915953 | 140423329 | 129911689 | 148521505 | 154074081 | 144898465 | 127953897 | 113479513 | 133072273 | 136518241 | 151769601 | 144936977 | 136547265 | 143642353 |
| Q9CPW4 | Arpc5 | Actin-related protein 2/3 complex subunit 5 Mus musculus | 544.13 | 58.94 | 16288 | 8 | 8 | 81309769 | 83109705 | 80807793 | 87037905 | 88914985 | 81629113 | 81735417 | 82704793 | 80621089 | 84645273 | 84506497 | 79142777 | 80508929 | 87017353 |
| P29066 | Arrb1 | Beta-arrestin-1 Rattus norvegicus | 199.58 | 15.31 | 47020 | 4 | 5 | 2347984.25 | 1851920.12 | 3963660.5 | 2899776.5 | 1999813 | 2265942.25 | 1989832.5 | 3504716 | 3749494 | 3376224 | 2062313.38 | 2962713 | 2172285.5 | 2153036.75 |
| P29067 | Arrb2 | Beta-arrestin-2 Rattus norvegicus | 455.84 | 20.73 | 46340 | 8 | 9 | 6769582 | 8025146 | 7476405 | 7378432 | 8299979 | 7657235.5 | 7876933 | 7226334.5 | 8035681.5 | 8630448 | 7624816.5 | 6321922 | 7238294 | 7682043 |
| P50428 | Arsa | Arylsulfatase A Mus musculus | 197.28 | 14.23 | 53748 | 3 | 4 | 12926354 | 3676284.5 | 3297498 | 3617777.5 | 5369760 | 5086908 | 4210050 | 4890346 | 4635105 | 4398067 | 4652566.5 | 5276526 | 5386068 | 3031160.75 |
| P50429 | Arsb | Arylsulfatase B Mus musculus | 676.72 | 24.16 | 59647 | 12 | 12 | 61991101 | 75576577 | 45951285 | 58174565 | 83004793 | 70061593 | 65081929 | 61776641 | 69050369 | 66685641 | 68064201 | 62044977 | 64751101 | 58680121 |
| Q91WU5 | As3mt | Arsenite methyltransferase Mus musculus | 119.12 | 8.51 | 41794 | 2 | 2 | 1717695.88 | 1549232.75 | 2006569.12 | 1131476.38 | 1260063.75 | 1502511.75 | 1525328.38 | 1746514.12 | 1568227.5 | 1468570.62 | 1324287.88 | 1484968.38 | 1481095.62 | 1769243.88 |
| Q9WV54 | Asah1 | Acid ceramidase Mus musculus | 647.15 | 34.01 | 44670 | 10 | 10 | 28765885 | 34030621 | 29372711 | 31656023 | 31882991 | 33320925 | 32397965 | 32849419 | 31598965 | 29658017 | 31204039 | 30153913 | 30250643 | 30770365 |
| Q9JHE3 | Asah2 | Neutral ceramidase Mus musculus | 37.1 | 1.72 | 83509 | 1 | 1 | 454990.69 | 401785 | 349317.94 | 207973.19 | 360640.59 | 380623.25 | 361541.66 | 419769.88 | 471286.25 | 335365.34 | 335264.06 | 392228.38 | 391754.94 | 304421.34 |
| Q1AAU6 | Asap1 | Arf-GAP with SH3 domain, ANK repeat and PH domain-containing protein 1 Rattus norveg | 459.92 | 9.53 | 127088 | 9 | 9 | 8820306 | 9953314 | 9387332 | 9091122 | 9597520 | 9608660 | 10004441 | 10351313 | 11853411 | 11547661 | 10488327 | 10368535 | 10301394 | 11688451 |
| Q9DAP7 | Asf1b | Histone chaperone ASF1B Mus musculus | 164.03 | 31.19 | 22464 | 3 | 3 | 249335.05 | 1 | 183284.09 | 330287.91 | 1 | 206735.52 | 277134.445 | 305315.88 | 363096.69 | 233389.69 | 1 | 1 | 211110.86 | 251780.61 |
| Q91X20 | Ash2l | Set1/Ash2 histone methyltransferase complex subunit ASH2 Mus musculus | 112.32 | 5.3 | 68250 | 2 | 2 | 690015.06 | 1 | 364462.91 | 1 | 613051.56 | 645969.62 | 1638574.62 | 459979.47 | 515459 | 618212.88 | 568268.6967 | 671936.56 | 603895.81 | 428973.72 |
| Q91YI0 | Asl | Argininosuccinate lyase Mus musculus | 528.51 | 28.23 | 51739 | 9 | 9 | 10089356 | 7535517.5 | 7922997.5 | 7927115 | 8766312 | 8660031 | 8744593 | 7501519 | 9506547 | 8836417 | 6472061 | 9928534 | 8745512 | 9019169 |
| O54984 | Asna1 | ATPase Asna1 Mus musculus | 491.22 | 48.56 | 38823 | 11 | 11 | 10520835 | 10712636 | 11106910 | 10759026 | 8052762 | 12026983 | 11247318 | 10786837 | 12850331 | 12713434 | 9912416 | 10081743 | 12311976 | 12029832 |
| Q61024 | Asns | Asparagine synthetase [glutamine-hydrolyzing] Mus musculus | 807.35 | 33.51 | 64283 | 14 | 14 | 22282757 | 16045791 | 22277965 | 22302875 | 22211675 | 34332637 | 24759677 | 23762539 | 22689153 | 25055077 | 23824953 | 28357385 | 23592073 | 24117823 |
| Q8BSY0 | Asph | Aspartyl/asparaginyl beta-hydroxylase Mus musculus | 704.88 | 24.16 | 83042 | 11 | 11 | 8928812 | 9671144 | 9140283 | 10347683 | 10067577 | 9367531 | 10691637 | 11005368 | 10456686 | 10530595 | 8792334 | 9212137 | 9529294 | 9310529 |
| Q8VBT9 | Aspscr1 | Tether containing UBX domain for GLUT4 Mus musculus | 141.82 | 6.73 | 59796 | 2 | 2 | 654962.44 | 539152.06 | 575547.265 | 693931.62 | 414142.94 | 462849.41 | 1 | 702454.75 | 438055.12 | 394186.12 | 446300.66 | 337056.69 | 367681.97 | 350275.06 |
| Q8C0M9 | Asrgl1 | Isoaspartyl peptidase/L-asparaginase Mus musculus | 136.57 | 11.04 | 33950 | 3 | 3 | 5926281 | 9226601 | 9620158 | 3857887.75 | 3708439.25 | 5678126 | 5721453 | 9016008 | 5866665 | 7992680 | 6603791 | 7198446 | 5938247 | 6533847 |
| P16460 | Ass1 | Argininosuccinate synthase Mus musculus | 608.06 | 35.92 | 46584 | 11 | 11 | 13874908 | 10818543 | 12943893 | 12171462 | 10458345 | 11462286 | 10063978 | 10398203 | 12786895 | 10986804 | 11343610 | 12126048 | 12211910 | 10361423 |
| Q505J9 | Atad1 | ATPase family AAA domain-containing protein 1 Rattus norvegicus | 124.53 | 7.76 | 40717 | 2 | 2 | 1973895.75 | 2010580.62 | 2134381.75 | 2198934 | 2198041.25 | 1829851 | 1461598.25 | 2256868.75 | 2302542.5 | 1343975.12 | 2112895 | 2398699.25 | 2052857.5 | 1820675 |
| Q3KRE0 | Atad3 | ATPase family AAA domain-containing protein 3 Rattus norvegicus | 182.8 | 7.28 | 66759 | 3 | 3 | 1 | 1 | 153757.5 | 1 | 196889.88 | 260729.42 | 205999.27 | 294182.22 | 242422.42 | 282221.62 | 531651.44 | 354055.62 | 360888.22 | 169627.2 |
| Q60765 | Atf3 | Cyclic AMP-dependent transcription factor ATF-3 Mus musculus | 37.84 | 5.52 | 20722 | 1 | 1 | 136233.5975 | 121549.55 | 125725.25 | 121069.61 | 176589.98 | 94550.63 | 152725.09 | 129766.71 | 137178.53 | 56622.71 | 32238.48 | 142370.14 | 35274.48 | 219122.92 |
| Q8C0J2 | Atg16l1 | Autophagy-related protein 16-1 Mus musculus | 37.68 | 4.61 | 68172 | 1 | 1 | 283115.12 | 277572.12 | 243052.09 | 295654.28 | 251957.73 | 363904.25 | 209897.11 | 166164.86 | 684148.44 | 252400.59 | 307900.31 | 258506.2 | 236983.38 | 352384.03 |
| Q6AZ50 | Atg3 | Ubiquitin-like-conjugating enzyme ATG3 Rattus norvegicus | 117.78 | 10.51 | 35822 | 3 | 3 | 836774.06 | 1038730.56 | 2332083 | 2298163.25 | 2343778 | 2865089.25 | 2437585.5 | 2873648.25 | 2651260 | 2509774.25 | 2322526.5 | 1251042.62 | 2246637.5 | 1649689.5 |
| Q8BGE6 | Atg4b | Cysteine protease ATG4B Mus musculus | 356.4 | 26.46 | 44375 | 7 | 7 | 5191828.5 | 3700420.25 | 4350060 | 3010067.5 | 3811216.75 | 5192974 | 5702951 | 4884073 | 4691446 | 5086594 | 5640412.5 | 4395238 | 3729435.75 | 5056069.5 |
| Q9D906 | Atg7 | Ubiquitin-like modifier-activating enzyme ATG7 Mus musculus | 879.05 | 30.95 | 77520 | 17 | 17 | 21356659 | 16975681 | 21445151 | 22598491 | 21669667 | 22660869 | 24427023 | 26245317 | 22287843 | 25785223 | 23180135 | 21138159 | 23389657 | 25133443 |
| Q9CWJ9 | Atic | Bifunctional purine biosynthesis protein PURH Mus musculus | 2297.18 | 75.68 | 64217 | 34 | 34 | 100501897 | 103995289 | 114436017 | 101709217 | 103917497 | 91965265 | 92060817 | 90759465 | 92896721 | 97746585 | 109017473 | 106490857 | 107272081 | 106332249 |
| Q6PA06 | Atl2 | Atlastin-2 Mus musculus | 92.44 | 4.12 | 66224 | 2 | 2 | 733836.25 | 899531.75 | 923835.75 | 877421.19 | 803702.88 | 1116392.38 | 1110131 | 1190888.5 | 1285978.75 | 933306.44 | 1116237.38 | 1200585 | 1236324 | 1006278.12 |

| | | | | | | | | | | | | | | | | | | | | | |
|---|---|---|---|---|---|---|---|---|---|---|---|---|---|---|---|---|---|---|---|---|---|
| Q91YH5 | Atl3 | Atlastin-3 Mus musculus | 719.84 | 34.01 | 60575 | 12 | 12 | 12180010 | 10808828 | 10887446 | 9888703 | 10379211 | 11147114 | 11473032 | 13461913 | 12128952 | 10358093 | 9689811 | 8677545 | 9937148 | 10527438 |
| O08997 | Atox1 | Copper transport protein ATOX1 Mus musculus | 342.18 | 88.24 | 7338 | 5 | 5 | 72652377 | 90571233 | 76901185 | 84529169 | 80775057 | 70288777 | 74821937 | 81985665 | 81892353 | 81834865 | 73459473 | 71971729 | 73340145 | 83694313 |
| P98197 | Atp11a | Probable phospholipid-transporting ATPase IH Mus musculus | 35.86 | 1.1 | 135502 | 1 | 1 | 1 | 1 | 216804.14 | 1 | 192745.98 | 1 | 1 | 213541.69 | 1 | 1 | 1 | 269621.47 | 1 | 175343.03 |
| Q9Z1W8 | Atp12a | Potassium-transporting ATPase alpha chain 2 Mus musculus | 170.42 | 2.71 | 114727 | 1 | 4 | 17240277 | 17120835 | 17445753 | 14583029 | 18905811 | 18125081 | 14699113 | 16942239 | 16950801 | 18613145 | 17657465 | 16783579 | 18797349 | 19198197 |
| Q9EPE9 | Atp13a1 | Manganese-transporting ATPase 13A1 Mus musculus | 294.17 | 9 | 132388 | 6 | 6 | 3277617 | 3065685.75 | 4283905 | 3475658.5 | 4203536 | 3481029.75 | 3556447 | 2800960 | 3521475.5 | 4067172.75 | 5029892 | 4656637 | 4609959 | 4443279 |
| Q8VDN2 | Atp1a1 | Sodium/potassium-transporting ATPase subunit alpha-1 Mus musculus | 2261.13 | 38.22 | 112982 | 22 | 35 | 233579809 | 261796497 | 257596817 | 243048401 | 255043793 | 247369313 | 239857025 | 237750049 | 241666961 | 257948321 | 240986017 | 237591473 | 249948065 | 263995377 |
| P06686 | Atp1a2 | Sodium/potassium-transporting ATPase subunit alpha-2 Rattus norvegicus | 950.36 | 15.1 | 112217 | 1 | 13 | 95076665 | 101328361 | 101793449 | 94733745 | 107726785 | 106499537 | 92320953 | 98096385 | 97041137 | 105251937 | 98068801 | 97190329 | 105620897 | 108438025 |
| P97370 | Atp1b3 | Sodium/potassium-transporting ATPase subunit beta-3 Mus musculus | 329.66 | 30.22 | 31776 | 5 | 5 | 14856685 | 11095817 | 12642517 | 9760679 | 10397499 | 12210738 | 12349141 | 12638752 | 10762199 | 10479633 | 12061747 | 10855213 | 11283860 | 11500161 |
| Q64578 | Atp2a1 | Sarcoplasmic/endoplasmic reticulum calcium ATPase 1 Rattus norvegicus | 625.01 | 12.98 | 109409 | 1 | 9 | 4560039.5 | 3859643 | 7743334 | 4583503 | 3654250.75 | 6406847 | 4212267.5 | 2775409.25 | 5743623 | 5398915 | 6430583.5 | 7853750.5 | 7672781.5 | 7825039.5 |
| O55143 | Atp2a2 | Sarcoplasmic/endoplasmic reticulum calcium ATPase 2 Mus musculus | 1778.63 | 35.06 | 114858 | 19 | 30 | 84705337 | 74889233 | 78532473 | 81246145 | 80584153 | 80121665 | 77736505 | 69938097 | 70138433 | 71227265 | 91030433 | 83612001 | 79107209 | 83184073 |
| Q64518 | Atp2a3 | Sarcoplasmic/endoplasmic reticulum calcium ATPase 3 Mus musculus | 740.84 | 15.9 | 113638 | 7 | 12 | 21217385 | 21617743 | 20382085 | 22246409 | 24183677 | 23850321 | 24953457 | 22223931 | 24122077 | 24198801 | 25532695 | 25672589 | 23845941 | 25181271 |
| G5E829 | Atp2b1 | Plasma membrane calcium-transporting ATPase 1 Mus musculus | 1028.14 | 18.03 | 134747 | 11 | 18 | 27450699 | 31212519 | 29367493 | 24706963 | 28031679 | 28519171 | 26604059 | 24810163 | 26907679 | 27185145 | 26091273 | 26947355 | 29095759 | 29839825 |
| Q64568 | Atp2b3 | Plasma membrane calcium-transporting ATPase 3 Rattus norvegicus | 544.51 | 7.15 | 138559 | 2 | 9 | 14151445 | 16483920 | 15763355 | 12696652 | 14428417 | 16069986 | 14419607 | 12962003 | 14114805 | 13069836 | 13282862 | 13825825 | 14611956 | 17075341 |
| Q03265 | Atp5f1a | ATP synthase subunit alpha, mitochondrial Mus musculus | 2133.34 | 60.94 | 59753 | 5 | 34 | 320112897 | 370741889 | 391404577 | 381308225 | 352192897 | 332440993 | 390419073 | 371811713 | 348489217 | 350497921 | 375998945 | 371648577 | 385242561 | 320617121 |
| P15999 | Atp5f1a | ATP synthase subunit alpha, mitochondrial Rattus norvegicus | 1877.48 | 54.25 | 59754 | 1 | 30 | 267228273 | 310851905 | 331039073 | 328174785 | 295759873 | 274769505 | 331466273 | 306400065 | 288894081 | 282669505 | 309344609 | 314383361 | 319037697 | 259881137 |
| P56480 | Atp5f1b | ATP synthase subunit beta, mitochondrial Mus musculus | 2406.32 | 78.26 | 56300 | 29 | 33 | 297902081 | 322207489 | 360714177 | 311538945 | 312757313 | 369774977 | 339196289 | 343579553 | 345979041 | 351858753 | 339415649 | 377274017 | 381537665 | 352649281 |
| Q91VR2 | Atp5f1c | ATP synthase subunit gamma, mitochondrial Mus musculus | 513.01 | 38.59 | 32886 | 10 | 10 | 29189741 | 35319389 | 32932073 | 34564013 | 34362513 | 34312585 | 28572129 | 28117227 | 26355275 | 30012601 | 32416861 | 34295201 | 29743273 | 34028669 |
| Q9D3D9 | Atp5f1d | ATP synthase subunit delta, mitochondrial Mus musculus | 205.66 | 49.4 | 17600 | 4 | 4 | 24280685 | 31734101 | 27633073 | 29550233 | 28133089 | 28717525 | 30466745 | 31732297 | 26237577 | 28169477 | 27340093 | 29434593 | 28344251 | 30534497 |
| O35143 | ATP5IF1 | ATPase inhibitor, mitochondrial Mus musculus | 45.13 | 8.49 | 12159 | 1 | 1 | 3372996.75 | 2822545.75 | 3568362 | 4602494 | 3475411.5 | 3831327 | 4120508.75 | 3442412.25 | 3102892.25 | 4520392 | 4457560.5 | 3403416 | 4388398.5 | 4012072 |
| P56383 | Atp5mc2 | ATP synthase F(0) complex subunit C2, mitochondrial Mus musculus | 111.41 | 21.23 | 15476 | 1 | 1 | 491922.41 | 4113024.5 | 2933016 | 1811611.5 | 2245393.25 | 2711854.75 | 4691664.5 | 3926261.75 | 3767578.75 | 2583846.5 | 2240109.75 | 3227062.75 | 2508488.5 | 2983567.75 |
| Q78IK2 | Atp5md | Up-regulated during skeletal muscle growth protein 5 Mus musculus | 147.05 | 43.1 | 6382 | 2 | 2 | 5997268.5 | 5239435 | 4859546 | 5485673.5 | 6008910 | 5359749 | 5312684 | 4495099 | 5173727.5 | 5675313 | 5639110 | 6339356.5 | 5518029 | 5512556 |
| Q06185 | Atp5me | ATP synthase subunit e, mitochondrial Mus musculus | 106.08 | 42.25 | 8236 | 3 | 3 | 7695626 | 3353336 | 1131103.38 | 1549188.88 | 1145770.88 | 925798.69 | 3578606.25 | 2897337.5 | 2429135.75 | 3674276.5 | 1579706 | 1077858.12 | 1052533.75 | 924340.69 |
| P56135 | Atp5mf | ATP synthase subunit f, mitochondrial Mus musculus | 81.01 | 23.86 | 10344 | 1 | 2 | 17268877 | 15526217 | 13439054 | 11799988 | 15739141 | 15573788 | 14159780 | 14462822 | 13439171 | 13235959 | 14934679 | 13989046 | 16013356 | 14409942 |
| Q9CQQ7 | Atp5pb | ATP synthase F(0) complex subunit B1, mitochondrial Mus musculus | 357.22 | 29.3 | 28949 | 7 | 7 | 18217805 | 20069307 | 17556477 | 16688425 | 16813723 | 19478099 | 16315606 | 19189415 | 13404739 | 14620618 | 16091655 | 16471062 | 17699113 | 17040375 |
| Q9DCX2 | Atp5pd | ATP synthase subunit d, mitochondrial Mus musculus | 502.6 | 63.35 | 18749 | 9 | 9 | 27412769 | 28160661 | 28176317 | 28045433 | 23592771 | 23842847 | 25388689 | 32720377 | 22013595 | 24273539 | 23675529 | 26102247 | 24409897 | 25250151 |
| P97450 | Atp5pf | ATP synthase-coupling factor 6, mitochondrial Mus musculus | 111.26 | 17.59 | 12496 | 1 | 1 | 1423215.25 | 1208755.62 | 1075771.62 | 1035423.06 | 1000578.38 | 1055748.5 | 862907.75 | 905113.56 | 778138.56 | 1078898.62 | 813857.88 | 678042.12 | 811868.88 | 934275.62 |
| Q9DB20 | Atp5po | ATP synthase subunit O, mitochondrial Mus musculus | 573.24 | 48.83 | 23364 | 8 | 9 | 16159396 | 16036384 | 16720265 | 16439538 | 16090287 | 15083501 | 18243861 | 16097272 | 15176505 | 17104011 | 18890243 | 14480927 | 16742232 | 16286389 |
| O54715 | Atp6ap1 | V-type proton ATPase subunit S1 Rattus norvegicus | 33.07 | 1.51 | 51123 | 1 | 1 | 2637324.75 | 2791397.5 | 3484690.5 | 3701668.75 | 3812607.75 | 2374916.25 | 2613749.75 | 2442275.5 | 2349059.5 | 2525420.5 | 2283052 | 2530357.75 | 2638727.75 | 2887804.25 |
| Q9Z1G4 | Atp6v0a1 | V-type proton ATPase 116 kDa subunit a isoform 1 Mus musculus | 262.69 | 9.06 | 96467 | 5 | 5 | 4060833.875 | 3957057 | 3542756 | 4797077 | 3946445.5 | 3861644.5 | 4273693 | 5266857.5 | 4488372.5 | 5605045 | 4108604.75 | 1353263.5 | 4091360 | 3321075 |
| P63081 | Atp6v0c | V-type proton ATPase 16 kDa proteolipid subunit Rattus norvegicus | 493.46 | 76.13 | 15808 | 5 | 8 | 33167599 | 61258881 | 39233713 | 32201433 | 44337745 | 66399977 | 59388005 | 55777529 | 51159741 | 52441885 | 44433793 | 55870013 | 59187773 | 51913465 |
| P51863 | Atp6v0d1 | V-type proton ATPase subunit d 1 Mus musculus | 952.93 | 54.13 | 40301 | 15 | 15 | 77478201 | 91544969 | 85966673 | 75775697 | 81929729 | 86353985 | 85801801 | 89450521 | 84231481 | 80664569 | 82047921 | 84033537 | 86611001 | 88723865 |
| P50516 | Atp6v1a | V-type proton ATPase catalytic subunit A Mus musculus | 2341.2 | 73.91 | 68326 | 34 | 35 | 142434705 | 139243073 | 154388161 | 126802617 | 127971937 | 139649073 | 140930801 | 139369777 | 143628865 | 140880721 | 137940417 | 143311825 | 152183697 | 149080289 |
| P62814 | Atp6v1b2 | V-type proton ATPase subunit B, brain isoform Mus musculus | 1684.59 | 76.13 | 56551 | 25 | 25 | 120825793 | 138188385 | 137582017 | 137719441 | 132128145 | 127680905 | 139031009 | 130704641 | 126655729 | 139154305 | 134412033 | 124117425 | 136060257 | 132119633 |
| Q5FVI6 | Atp6v1c1 | V-type proton ATPase subunit C 1 Rattus norvegicus | 974.91 | 43.98 | 43901 | 17 | 17 | 40574881 | 40408421 | 36351181 | 37687361 | 33457207 | 35273673 | 32090027 | 35615617 | 29545309 | 30895645 | 29103067 | 34799201 | 35215713 | 36383877 |
| P57746 | Atp6v1d | V-type proton ATPase subunit D Mus musculus | 183.41 | 17.41 | 28369 | 3 | 3 | 5983589 | 4213503 | 4272386.5 | 4684691 | 4494796.5 | 3916660.25 | 4494212 | 3920331.75 | 4568220.5 | 3715119 | 4731451 | 4531952.5 | 4669482 | 4584593 |
| P50518 | Atp6v1e1 | V-type proton ATPase subunit E 1 Mus musculus | 426.99 | 33.63 | 26157 | 6 | 6 | 28323443 | 31801433 | 27329297 | 39835665 | 36506273 | 29579317 | 27641379 | 16094747 | 20655525 | 25425685 | 38522765 | 41625453 | 26792491 | 16965189 |
| P50408 | Atp6v1f | V-type proton ATPase subunit F Rattus norvegicus | 255.94 | 45.38 | 13370 | 4 | 4 | 3539334.5 | 3892337.75 | 3070427.5 | 3382132.5 | 3904443 | 2748778.5 | 2688134 | 3827032.5 | 2431745.25 | 2751667.75 | 3421444 | 2253462.5 | 1932349 | 2256133.5 |
| Q9CR51 | Atp6v1g1 | V-type proton ATPase subunit G 1 Mus musculus | 204.02 | 36.44 | 13724 | 3 | 3 | 6386704.5 | 5091156 | 5424084.5 | 6782994 | 6561993 | 5388240.5 | 6925537.5 | 5080297.5 | 5934768.5 | 6903785 | 6679495.5 | 5977726 | 6104392 | 5713213 |
| Q8BVE3 | Atp6v1h | V-type proton ATPase subunit H Mus musculus | 696.59 | 32.3 | 55855 | 13 | 13 | 20321119 | 20710759 | 23600781 | 18563459 | 19201917 | 27175529 | 22681483 | 23013645 | 26928313 | 23830863 | 22637419 | 25989765 | 25466551 | 23831829 |
| Q64430 | Atp7a | Copper-transporting ATPase 1 Mus musculus | 158.3 | 2.88 | 161959 | 4 | 4 | 1918812.62 | 727557.19 | 1614132 | 1341063.62 | 1364779.5 | 1353237.25 | 1536501.25 | 1334503.38 | 1457355.5 | 1416150.62 | 1305571.75 | 1574347.75 | 1400271.25 | 1398265.75 |
| Q91YY4 | Atpaf2 | ATP synthase mitochondrial F1 complex assembly factor 2 Mus musculus | 40.45 | 3.81 | 33289 | 1 | 1 | 1 | 1 | 714317.5 | 1 | 1 | 1 | 1 | 1 | 1 | 647275.5 | 876839.9167 | 986246.06 | 1037077.31 | 607196.38 |
| Q61687 | Atrx | Transcriptional regulator ATRX Mus musculus | 107.1 | 1.05 | 278587 | 2 | 2 | 626782.69 | 553771.25 | 402648.59 | 573780.31 | 663451.62 | 433169.34 | 554291.5 | 503146.41 | 592175.12 | 503073.5 | 682269.44 | 477768.12 | 566836.06 | 593032 |
| P28658 | Atxn10 | Ataxin-10 Mus musculus | 760.79 | 45.47 | 53707 | 15 | 15 | 25570491 | 19160953 | 23684225 | 17806277 | 20947241 | 23018501 | 20492799 | 26757523 | 24838331 | 21594531 | 20384405 | 19366457 | 24741975 | 23622799 |
| Q7TQH0 | Atxn2l | Ataxin-2-like protein Mus musculus | 632.15 | 16.68 | 110649 | 13 | 13 | 9982069 | 8612443 | 12295653 | 11524145 | 10839567 | 12009047 | 12057053 | 10404407 | 12399008 | 13512460 | 9516453 | 10763198 | 13010568 | 9851514 |
| P70295 | Aup1 | Ancient ubiquitous protein 1 Mus musculus | 269.36 | 18.78 | 46121 | 5 | 5 | 2443452 | 1747838.5 | 2808141 | 2333366.25 | 2666960.5 | 1788951.88 | 2665675.75 | 1811699.25 | 2692907 | 2509587.5 | 2612346 | 2232784.75 | 2536824 | 2623848 |
| O55099 | Aurkb | Aurora kinase B Rattus norvegicus | 142.8 | 9.91 | 39234 | 2 | 3 | 291684.5 | 331943.88 | 768632.94 | 1160410.25 | 1127376.25 | 443904.5 | 315479.59 | 1 | 927320.25 | 1 | 624924.94 | 951112.88 | 570416.88 | 466704.56 |
| Q9D9K3 | Aven | Cell death regulator Aven Mus musculus | 50.07 | 6.43 | 37195 | 1 | 1 | 543086.62 | 1 | 417981.97 | 382421.25 | 1 | 1 | 1 | 1 | 484343.34 | 717848.25 | 1 | 1 | 1 | 1 |
| P01887 | B2m | Beta-2-microglobulin Mus musculus | 47.53 | 8.4 | 13779 | 1 | 1 | 4676032.5 | 3371963 | 4980485.5 | 3501672.25 | 3830540.75 | 5126411 | 3488941.75 | 4975547.5 | 5138897.5 | 4336420 | 4693573.5 | 5337498.5 | 4639205 | 4431190.5 |
| Q09200 | B4galnt1 | Beta-1,4 N-acetylgalactosaminyltransferase 1 Mus musculus | 165.3 | 12.95 | 59212 | 4 | 4 | 1801928 | 2613827.25 | 1044855.62 | 1320747.88 | 2539414 | 1643022 | 2606619.75 | 2140355 | 2526586.75 | 2573344.5 | 2447143.75 | 2433712.5 | 3228275.5 | 2554666.5 |
| P15535 | B4galt1 | Beta-1,4-galactosyltransferase 1 Mus musculus | 129.14 | 9.52 | 44411 | 3 | 3 | 595688.04 | 677965.44 | 605968 | 461883.34 | 636935.38 | 1090460.62 | 598736.56 | 726263.88 | 730198.3825 | 505332.47 | 609567.06 | 1084064.75 | 633071.19 | 785683.19 |
| Q6P7Q1 | Babam2 | BRISC and BRCA1-A complex member 2 Rattus norvegicus | 141.76 | 9.92 | 43559 | 3 | 3 | 3014357.5 | 2838094 | 3680446.5 | 2320684 | 2456194.5 | 4393692 | 2615714.5 | 2922149.5 | 3064286 | 2630787.5 | 2831365.5 | 3797890 | 3743891.5 | 3251381 |
| Q6MG49 | Bag6 | Large proline-rich protein BAG6 Rattus norvegicus | 182.14 | 3.4 | 120012 | 3 | 3 | 5516473.5 | 5672206.5 | 6263784.5 | 4499107 | 6266172 | 5573416 | 5354470.5 | 6554594 | 6240818.5 | 4957826 | 6116015 | 6196116.5 | 6264639 | 6091562.5 |
| O08734 | Bak1 | Bcl-2 homologous antagonist/killer Mus musculus | 412.94 | 45.45 | 23295 | 8 | 8 | 24148415 | 20861775 | 21346057 | 19750739 | 19323359 | 23619243 | 19120411 | 25169879 | 22325775 | 19234717 | 18886785 | 18069601 | 19331089 | 20925675 |
| O54962 | Banf1 | Barrier-to-autointegration factor Mus musculus | 234.49 | 52.81 | 10103 | 4 | 4 | 31100997 | 37828185 | 33642749 | 36263769 | 34584713 | 32375681 | 40268021 | 37568117 | 39317989 | 36068625 | 35745689 | 35934417 | 30034237 | 37462341 |
| Q91XV3 | Basp1 | Brain acid soluble protein 1 Mus musculus | 926.1 | 77.43 | 22087 | 7 | 12 | 121687753 | 142960081 | 91223177 | 113236305 | 141292561 | 124743353 | 152490529 | 134322561 | 109870025 | 129857721 | 106542305 | 95058161 | 85663641 | 113408385 |
| Q07813 | Bax | Apoptosis regulator BAX Mus musculus | 635.21 | 63.02 | 21395 | 9 | 9 | 33075229 | 33938641 | 33393683 | 25691449 | 33127145 | 35987489 | 35737021 | 35643941 | 35700893 | 32444009 | 37677029 | 34837793 | 36930033 | 37492449 |
| Q9Z277 | Baz1b | Tyrosine-protein kinase BAZ1B Mus musculus | 327.78 | 6.63 | 170651 | 7 | 7 | 4002570.5 | 4994442 | 3172398.25 | 2495981.75 | 5464130 | 5038227.5 | 4595027.5 | 3781865.75 | 4632838 | 4619897 | 3850210 | 3926551.5 | 4592533 | 2828554.25 |
| Q9D287 | Bcas2 | Pre-mRNA-splicing factor SPF27 Mus musculus | 62.44 | 12.44 | 26131 | 1 | 1 | 1 | 1 | 164248.88 | 182089.86 | 155525.17 | 151165.47 | 134388.78 | 348150.94 | 1 | 1 | 1 | 1 | 1 | 131625.53 |
| P24288 | Bcat1 | Branched-chain-amino-acid aminotransferase, cytosolic Mus musculus | 189.22 | 9.59 | 42791 | 4 | 4 | 2924669 | 4449265.5 | 5201358 | 4306470 | 3904854 | 4006281 | 3421553 | 4225557.5 | 2704946.75 | 3705861 | 3770401.5 | 3655180 | 2713849 | 4068167.75 |
| O35855 | Bcat2 | Branched-chain-amino-acid aminotransferase, mitochondrial Mus musculus | 290.18 | 15.01 | 44127 | 5 | 5 | 23033639 | 21263059 | 21014819 | 22546623 | 23722181 | 23063793 | 20956577 | 19684521 | 22068899 | 21164947 | 22559083 | 22270481 | 20292075 | 22750369 |
| Q9CWI3 | Bccip | BRCA2 and CDKN1A-interacting protein Mus musculus | 151.58 | 15.19 | 35942 | 4 | 4 | 3967588 | 5860369 | 6255044.5 | 5587338.5 | 4560883.5 | 5508632 | 5765540 | 5528246 | 5049445 | 4945419.5 | 5658421 | 6684720 | 6166711 | 6014243.5 |
| P35738 | Bckdhb | 2-oxoisovalerate dehydrogenase subunit beta, mitochondrial Rattus norvegicus | 134.18 | 7.44 | 42823 | 2 | 2 | 957250.94 | 2317624.75 | 2008439.62 | 491190.06 | 352572.53 | 1001312.5 | 1 | 2139925.25 | 1 | 1784916.12 | 1136532.62 | 1308364.5 | 2292087 | 2094612.88 |
| O55028 | Bckdk | [3-methyl-2-oxobutanoate dehydrogenase [lipoamide]] kinase, mitochondrial Mus musculus | 123.32 | 11.41 | 46588 | 2 | 2 | 5335562 | 6367086 | 8666488 | 6777259 | 5026245 | 5429819 | 9566477 | 5840154.5 | 9835070 | 7269164 | 7552943 | 7648839.5 | 7324325.5 | 4865535.5 |
| Q9Z0H7 | Bcl10 | B-cell lymphoma/leukemia 10 Mus musculus | 119.69 | 22.75 | 25948 | 3 | 3 | 2468201.5 | 1522003.5 | 1450064.25 | 1656171.5 | 1788120 | 1548546.25 | 1991269.12 | 1549399.75 | 1729522 | 2259781.75 | 1804997.5 | 1650020.38 | 1483324 | 1595970.12 |
| P10417 | Bcl2 | Apoptosis regulator Bcl-2 Mus musculus | 37.3 | 4.24 | 26407 | 1 | 1 | 593319.06 | 778310.62 | 883003.56 | 1 | 1 | 882704.955 | 596061.88 | 1063698.75 | 1198158.75 | 672900.44 | 597112.88 | 728308 | 744044.69 | 729096.5 |
| P53563 | Bcl2l1 | Bcl-2-like protein 1 Rattus norvegicus | 116.49 | 8.15 | 26158 | 2 | 2 | 5620402.5 | 6564159 | 6171908.5 | 5036935 | 5291808 | 5462271 | 5829856 | 6391595 | 5919540.5 | 5332701.5 | 4906118 | 4928178 | 5708010.5 | 5653290 |
| P59017 | Bcl2l13 | Bcl-2-like protein 13 Mus musculus | 266.18 | 20.97 | 46719 | 5 | 5 | 5133233 | 7492595.5 | 7480295.5 | 8898548 | 7648284 | 9898227 | 9877343 | 6319342 | 9542977 | 6798287 | 10489876 | 8563896 | 9486708 | 10260459 |
| Q8K019 | Bclaf1 | Bcl-2-associated transcription factor 1 Mus musculus | 262.86 | 7.73 | 106002 | 6 | 6 | 5567837.5 | 5830390 | 5469708.5 | 6010429 | 6357267 | 6354879 | 6461912 | 4659881 | 5358369 | 5823422.5 | 5460952 | 5264731 | 5638957.5 | 5693408 |
| Q9CZP5 | Bcs1l | Mitochondrial chaperone BCS1 Mus musculus | 72.75 | 4.55 | 47406 | 1 | 1 | 279154.44 | 219089.17 | 206571.38 | 316904.25 | 264576.97 | 332101.66 | 293105.88 | 336330.41 | 323191.06 | 282350.47 | 299724.09 | 274138.09 | 336352.5 | 229844.08 |
| P70444 | Bid | BH3-interacting domain death agonist Mus musculus | 68.36 | 9.74 | 21952 | 1 | 1 | 463153.19 | 284290.16 | 331945.22 | 315233.22 | 250286.28 | 359350.28 | 239240.81 | 446505.03 | 313042.22 | 383571.38 | 403019.38 | 310196.44 | 415413.94 | 313657.22 |
| O08539 | Bin1 | Myc box-dependent-interacting protein 1 Mus musculus | 172.38 | 8.33 | 64470 | 2 | 3 | 1466632.25 | 1737278.88 | 1662260.75 | 1233827.25 | 1523102 | 1929133.5 | 1888381 | 1993632.88 | 2033967.75 | 2255730.25 | 3599544.75 | 3257452.75 | 1773316.38 | 1957549.25 |
| D3Z6Q9 | Bin2 | Bridging integrator 2 Mus musculus | 608.75 | 23.11 | 52554 | 8 | 9 | 13699195 | 11867651 | 12105952 | 13864309 | 13672661 | 10192439 | 12430583 | 9025065 | 12456111 | 11810048 | 14398497 | 13518427 | 11017589 | 14104471 |
| O88738 | Birc6 | Baculoviral IAP repeat-containing protein 6 Mus musculus | 831.91 | 5.28 | 532170 | 17 | 17 | 8297570.5 | 7470425 | 10081143 | 8263433.5 | 8246352 | 9623354 | 8116167.5 | 9801890 | 10275133 | 9027654 | 10489568 | 9916617 | 9566622 | 9308662 |
| Q8R016 | Blmh | Bleomycin hydrolase Mus musculus | 606.62 | 38.68 | 52511 | 2 | 12 | 17496483 | 15199557 | 15461817 | 15561043 | 17376735 | 17647917 | 17895743 | 14794750 | 14311714 | 15661648 | 17419559 | 13581546 | 13189051 | 13254252 |
| Q32WR5 | Bloc1s2 | Biogenesis of lysosome-related organelles complex-1 subunit 2 Rattus norvegicus | 36.14 | 7.75 | 16067 | 1 | 1 | 363234.62 | 1367884.5 | 340128.53 | 1090036.25 | 1005862.62 | 804638.38 | 911183.31 | 1023208.12 | 724892.56 | 774702.12 | 886097.81 | 945625.25 | 1 | 834349.25 |
| Q9CY64 | Blvra | Biliverdin reductase A Mus musculus | 753.23 | 58.64 | 33525 | 14 | 14 | 48784881 | 54583561 | 49228573 | 50434457 | 51295057 | 44310549 | 44582985 | 47738089 | 44683173 | 44107165 | 44423617 | 40271645 | 43850857 | 43421633 |
| Q923D2 | Blvrb | Flavin reductase (NADPH) Mus musculus | 516.99 | 60.19 | 22197 | 7 | 7 | 5689220 | 4773408 | 5004091 | 6257829 | 6131784 | 4959825 | 5258340.5 | 4532174 | 4845444 | 7765961 | 6198497 | 6815858.5 | 6466006 | 6357858.5 |
| Q8BXK4 | Bmt2 | S-adenosylmethionine sensor upstream of mTORC1 Mus musculus | 54.97 | 6.2 | 45952 | 1 | 1 | 148636.78 | 1 | 131433.78 | 228973.48 | 1 | 139076.48 | 99368.76 | 164159.59 | 172478.22 | 143770.7625 | 1 | 1 | 123695.92 | 134794.91 |
| Q9D8S9 | Bola1 | BolA-like protein 1 Mus musculus | 85.03 | 21.17 | 14379 | 2 | 2 | 1910952.5 | 2213255 | 3035279 | 2462605.75 | 2425786.25 | 2516464 | 2719292.25 | 2060039.12 | 2293587.5 | 1688547.75 | 2374514.5 | 2490222.5 | 2252568 | 2503843 |
| P97452 | Bop1 | Ribosome biogenesis protein BOP1 Mus musculus | 260.95 | 10.79 | 82546 | 6 | 6 | 3946396.25 | 3694792.25 | 3855895 | 5074552 | 4697104 | 6020389.5 | 5052080 | 3282742.25 | 5249487.5 | 5704251.5 | 3373756.5 | 4688288 | 7115534 | 4428486 |
| Q8R164 | Bphl | Valacyclovir hydrolase Mus musculus | 116.8 | 11.68 | 32851 | 2 | 2 | 1100669.62 | 1138351.88 | 1481812.12 | 1607256.25 | 1239352.62 | 1686760 | 1382599.75 | 1742254.5 | 1569561.25 | 1343794.12 | 1157134.62 | 1230496.75 | 1376014.75 | 1436224.75 |
| Q9Z0S1 | Bpnt1 | 3'(2'),5'-bisphosphate nucleotidase 1 Mus musculus | 546.68 | 58.12 | 33196 | 5 | 12 | 13379334 | 13102381 | 11183731 | 12615153 | 13661309 | 11247682 | 13429091 | 12217519 | 12382225 | 12341158 | 13556694 | 13069957 | 11724538 | 12729835 |
| Q9ESU6 | Brd4 | Bromodomain-containing protein 4 Mus musculus | 43.71 | 1.43 | 155895 | 1 | 1 | 1527573 | 2074683.5 | 1545319 | 1 | 1 | 1 | 2163970.25 | 1839124.88 | 1784019.62 | 1 | 1 | 1 | 1668954.12 | 1 |
| O88665 | Brd7 | Bromodomain-containing protein 7 Mus musculus | 65.44 | 4.15 | 74000 | 1 | 1 | 339867.565 | 310889.31 | 243446.92 | 365756.91 | 439377.12 | 289989.38 | 251164.72 | 261145.98 | 264751.605 | 256706.34 | 316395.47 | 317652.72 | 744044.69 | 273383.53 |
| Q8BXV2 | Bri3bp | BRI3-binding protein Mus musculus | 109.82 | 7.51 | 28263 | 1 | 1 | 2298517.25 | 1 | 1171718.75 | 1 | 1322235.62 | 1750025.75 | 1323338.88 | 1190853.75 | 1029613.25 | 1225893.38 | 1382083.38 | 1210212 | 1613035.38 | 1675358.25 |
| Q4QQT6 | Brix1 | Ribosome biogenesis protein BRX1 homolog Rattus norvegicus | 271.25 | 17.9 | 41115 | 5 | 5 | 5540502 | 3599404.5 | 4631790.5 | 4308019 | 3612766.5 | 5316179.5 | 4368629 | 4460523 | 4096792.5 | 4310553.5 | 3445699.75 | 2923515.5 | 4159066.25 | 4236672 |
| Q91VR8 | Brk1 | Protein BRICK1 Mus musculus | 81.48 | 24 | 8761 | 2 | 2 | 1986134 | 1097114.5 | 2865366.5 | 1 | 1 | 2542781 | 1540509.12 | 3411424.75 | 3108680 | 1379221 | 1839176.75 | 2320067 | 1960570.75 | 1941660.5 |
| Q8K2Q7 | Brox | BRO1 domain-containing protein BROX Mus musculus | 362.44 | 25.79 | 46202 | 7 | 7 | 8301968 | 8458051 | 6718905 | 6004193.5 | 7874387.5 | 6884492 | 7183560.5 | 8205023 | 5585821 | 7074829 | 6475049 | 6739797.5 | 6581380.5 | 8136992 |
| Q80Y55 | Bsdc1 | BSD domain-containing protein 1 Mus musculus | 97.89 | 7.96 | 46953 | 2 | 2 | 926742.06 | 675796.88 | 724778.12 | 788749.06 | 798811.62 | 768877.19 | 563023.88 | 1022172.69 | 763038.56 | 618960.19 | 693407.56 | 692605.88 | 926012.56 | 749818.75 |

| | | | | | | | | | | | | | | | | | | | | | |
|---|---|---|---|---|---|---|---|---|---|---|---|---|---|---|---|---|---|---|---|---|---|
| P18572 | Bsg | Basigin Mus musculus | 39.32 | 2.57 | 42445 | 1 | 1 | 1494309.62 | 1713495.38 | 1650802.25 | 1894312.5 | 1921786.12 | 1478428.75 | 1794948.12 | 1911232.75 | 1386155.12 | 1794077.88 | 1851147 | 1776601.75 | 2045947.88 | 1969285.75 |
| Q8R2Q8 | Bst2 | Bone marrow stromal antigen 2 Mus musculus | 110.88 | 11.63 | 19152 | 2 | 2 | 3549342 | 3271166.5 | 2628263 | 2198618.5 | 3423167 | 1919391.5 | 2287608.75 | 1598402.62 | 2082280.88 | 1783525.62 | 1678759.38 | 2267397.75 | 2377634.5 | 1841394 |
| Q64152 | Btf3 | Transcription factor BTF3 Mus musculus | 345.33 | 49.51 | 22031 | 5 | 6 | 8559690 | 3963968 | 4071065.75 | 5078866.5 | 3670445.75 | 3336583.5 | 7835755.5 | 3740741 | 3146954.25 | 4279316 | 3510721.75 | 3547773.25 | 3593206.75 | 3419285.25 |
| P35991 | Btk | Tyrosine-protein kinase BTK Mus musculus | 360.86 | 14.42 | 76437 | 7 | 7 | 2571734 | 1760990.5 | 4117530.5 | 2402241.5 | 2565336 | 2306141.25 | 1674619.75 | 1976427 | 2156884.25 | 1617684.75 | 2346284.75 | 5263701 | 2394040.5 | 2064818.88 |
| Q9WVA3 | Bub3 | Mitotic checkpoint protein BUB3 Mus musculus | 438.52 | 28.83 | 36955 | 7 | 7 | 16847627 | 19373095 | 17989345 | 18040573 | 16922063 | 14512527 | 17121833 | 16098568 | 15281613 | 17037767 | 14124080 | 15379591 | 14832119 | 15379073 |
| Q9CY21 | Bud23 | Probable 18S rRNA (guanine-N(7))-methyltransferase Mus musculus | 139.18 | 18.51 | 31587 | 3 | 3 | 975878.31 | 1687088.25 | 1467269 | 2788204 | 1119058.38 | 2175620.5 | 2317618 | 2083509.5 | 2197771 | 1753914 | 1469047.88 | 2992955.5 | 2363543 | 2477856.5 |
| O70454 | Bud31 | Protein BUD31 homolog Rattus norvegicus | 142.59 | 26.39 | 17000 | 3 | 3 | 3593911.5 | 3148453.5 | 2838716.5 | 1610258 | 2934452.25 | 2571536.5 | 2822109.25 | 3294713 | 2805012.5 | 1770887.62 | 1841646.75 | 1781671.75 | 2539217.75 | 2889267.25 |
| O54825 | Bysl | Bystin Mus musculus | 320.47 | 23.39 | 49784 | 7 | 7 | 6040633 | 3295103.25 | 2628441 | 3875066 | 4431506 | 2739912 | 5283587.5 | 2738138 | 4261319 | 2515156.75 | 4145098.5 | 3988747.5 | 2788601.5 | 2253046.25 |
| Q6P7P5 | Bzw1 | Basic leucine zipper and W2 domain-containing protein 1 Rattus norvegicus | 1012.89 | 61.34 | 48043 | 17 | 19 | 33573389 | 24747669 | 28850221 | 19980147 | 21328787 | 26714983 | 26889715 | 33340433 | 30575429 | 21072459 | 23480143 | 30508311 | 27430947 | 23261887 |
| Q2L4X1 | Bzw2 | Basic leucine zipper and W2 domain-containing protein 2 Mus musculus molossinus (Jap | 388.29 | 30.07 | 48063 | 7 | 9 | 7297814 | 4181495.5 | 6675814 | 3399781 | 4155829.5 | 4640361.5 | 5189722 | 7625703 | 7087156 | 4331829 | 4376958 | 7227306.5 | 6261464.5 | 5325246.5 |
| P98086 | C1qa | Complement C1q subcomponent subunit A Mus musculus | 50.02 | 6.12 | 25974 | 1 | 1 | 476973.56 | 391320.94 | 672807 | 879280.31 | 652380.44 | 382105.22 | 344430.47 | 249674.16 | 315717.97 | 375473.78 | 431360.09 | 385306.88 | 479654.56 | 351893.84 |
| O35658 | C1qbp | Complement component 1 Q subcomponent-binding protein, mitochondrial Mus musculus | 427.26 | 43.53 | 31013 | 1 | 7 | 9895254 | 7132176.5 | 11802255 | 8187858 | 7418910.5 | 12389617 | 11660238 | 14080177 | 13593956 | 10148720 | 11269229 | 11573135 | 12072293 | 8713768 |
| Q80X80 | C2cd2l | Phospholipid transfer protein C2CD2L Mus musculus | 49.59 | 1.42 | 76329 | 1 | 1 | 1 | 1 | 349341.56 | 1 | 1 | 505218.97 | 495357.28 | 569019.19 | 693052.94 | 364224.81 | 446286.72 | 497738.44 | 511167.75 | 485064.3033 |
| P30993 | C5ar1 | C5a anaphylatoxin chemotactic receptor 1 Mus musculus | 90.77 | 6.84 | 39023 | 2 | 2 | 3853467 | 2171287.75 | 1685147 | 1996552.75 | 1890232 | 2015879.38 | 2009257.62 | 1959171.38 | 2052005.62 | 2125921 | 320695.62 | 1392116.62 | 1544688.38 | 1752020.75 |
| Q9D6N1 | Ca13 | Carbonic anhydrase 13 Mus musculus | 57.94 | 3.44 | 29522 | 1 | 1 | 2512639.25 | 3071649 | 2929762.75 | 2469199.5 | 2384382.75 | 2982757.5 | 2600762.5 | 2909995 | 2568370 | 2396833.25 | 2347055.25 | 2332430 | 2273190.75 | 2154185.75 |
| P00920 | Ca2 | Carbonic anhydrase 2 Mus musculus | 72.87 | 6.15 | 29033 | 1 | 1 | 995448.44 | 967557.69 | 745657 | 1006444.56 | 713991.19 | 1199222.5 | 1319754.38 | 1269903.12 | 1257928.88 | 1000709.62 | 1030893.19 | 1065497.62 | 826465.69 | 936044.31 |
| Q06138 | Cab39 | Calcium-binding protein 39 Mus musculus | 538.05 | 40.76 | 39843 | 11 | 12 | 19173705 | 12546711 | 12553784 | 12871691 | 8612880 | 17860829 | 13474044 | 16801849 | 18356579 | 13494572 | 9657478 | 11383983 | 18232027 | 13728920 |
| Q9DB16 | Cab39l | Calcium-binding protein 39-like Mus musculus | 83.15 | 5.34 | 39106 | 1 | 2 | 6080585.5 | 1 | 3140046.25 | 3207864.25 | 1 | 3452073.5 | 3666847.938 | 3663121.75 | 3783982 | 3768214.5 | 1 | 1 | 3709789.5 | 3537789.75 |
| Q9D424 | Cabyr | Calcium-binding tyrosine phosphorylation-regulated protein Mus musculus | 35.11 | 1.55 | 48333 | 1 | 1 | 3739951.5 | 4223158.5 | 3593227.25 | 3195507.25 | 3261948.75 | 3540140.75 | 4231272.5 | 3734857.5 | 3829342 | 3619721.25 | 2909649.25 | 3285202 | 3527276.5 | 3701510.75 |
| Q9CXW3 | Cacybp | Calcyclin-binding protein Mus musculus | 666.09 | 55.46 | 26510 | 11 | 11 | 22912845 | 22584781 | 26027987 | 19970155 | 21790039 | 21316739 | 20983241 | 22000765 | 20938005 | 24189553 | 21031901 | 24125471 | 27603035 | 24597851 |
| B2RQC6 | Cad | CAD protein Mus musculus | 1956.76 | 23.28 | 243238 | 37 | 37 | 42515117 | 45114417 | 50362805 | 46463789 | 44529117 | 41966693 | 39472741 | 39367589 | 43304169 | 41710433 | 43789457 | 45553821 | 43951765 | 51205269 |
| Q6AYP5 | Cadm1 | Cell adhesion molecule 1 Rattus norvegicus | 137 | 7.77 | 51854 | 2 | 2 | 2619958 | 3106014.75 | 2897103 | 2986280 | 2788312.25 | 3269429.75 | 2964670.75 | 3552310.5 | 3456375.25 | 3103610 | 2871987.75 | 3456789.5 | 3386466 | 3392276.25 |
| Q8VEC4 | Calhm2 | Calcium homeostasis modulator protein 2 Mus musculus 2 | 162.2 | 12.69 | 35823 | 3 | 3 | 1478173.12 | 1066539 | 1002386.44 | 524569.94 | 550972.19 | 1231179.88 | 584603.62 | 1312927.75 | 1168291.5 | 539084.25 | 1118974.25 | 1302815.62 | 971582.38 | 611067.5 |
| P0DP26 | Calm1 | Calmodulin-1 Mus musculus | 661.71 | 80.54 | 16838 | 10 | 10 | 72509633 | 55863917 | 78980857 | 56340881 | 50930209 | 74906025 | 45190477 | 99569393 | 67117217 | 51750057 | 57424977 | 71220289 | 66704581 | 52227849 |
| P14211 | Calr | Calreticulin Mus musculus | 1301.77 | 59.62 | 47995 | 20 | 20 | 245965841 | 256566625 | 267784321 | 231734545 | 298205889 | 305657761 | 254900641 | 265679521 | 293298177 | 294845921 | 286706497 | 268442817 | 313402785 | 289483841 |
| O35887 | Calu | Calumenin Mus musculus | 548.33 | 47.94 | 37064 | 6 | 9 | 10071065 | 11226470 | 13939179 | 8559093 | 8033591 | 15623780 | 13041527 | 16754011 | 13940671 | 11137169 | 11685679 | 14567292 | 14008983 | 11136438 |
| Q63450 | Camk1 | Calcium/calmodulin-dependent protein kinase type 1 Rattus norvegicus | 188.81 | 10.16 | 41638 | 3 | 3 | 7956965 | 7125809 | 9297724 | 6216543 | 6378720.5 | 8254010 | 6235470 | 8479715 | 8158396 | 5818220 | 6686711 | 7735044.5 | 8084490 | 8874549 |
| Q6PHZ2 | Camk2d | Calcium/calmodulin-dependent protein kinase type II subunit delta Mus musculus | 490.44 | 26.45 | 56369 | 9 | 9 | 15552478 | 16226495 | 16876161 | 17452865 | 16448511 | 21893837 | 19909357 | 21355475 | 21208995 | 21389053 | 19836729 | 19635099 | 19042353 | 19131423 |
| P97536 | Cand1 | Cullin-associated NEDD8-dissociated protein 1 Rattus norvegicus | 1278.81 | 27.56 | 136362 | 24 | 24 | 60754897 | 57666085 | 56663609 | 50113409 | 60987517 | 55199457 | 52994305 | 61819965 | 55595321 | 54614685 | 55065493 | 57287177 | 54429485 | 58713101 |
| P35564 | Canx | Calnexin Mus musculus | 1226.02 | 49.58 | 67278 | 22 | 22 | 75944481 | 74517137 | 79112257 | 75547153 | 73482825 | 76148985 | 71898929 | 82528801 | 77842393 | 76958265 | 71962137 | 73018681 | 73568129 | 72016993 |
| P40124 | Cap1 | Adenylyl cyclase-associated protein 1 Mus musculus | 2231.67 | 70.25 | 51565 | 33 | 35 | 294169825 | 366957793 | 344019201 | 316606305 | 327385057 | 341739265 | 407582849 | 405840417 | 393304993 | 392298881 | 350190241 | 369820225 | 350521921 | 365544481 |
| P24452 | Capg | Macrophage-capping protein Mus musculus | 892.52 | 42.05 | 39240 | 1 | 13 | 390912129 | 435303329 | 312020865 | 343049249 | 445382593 | 494020577 | 392876961 | 426437185 | 546421697 | 403064833 | 494793857 | 481687617 | 421595553 | 524728193 |
| O35350 | Capn1 | Calpain-1 catalytic subunit Mus musculus | 559.77 | 18.65 | 82106 | 11 | 11 | 11923398 | 14869476 | 14141708 | 9304230 | 10484122 | 13538515 | 10323378 | 15186473 | 14529057 | 10876353 | 11271395 | 10706992 | 13032738 | 11254150 |
| O08529 | Capn2 | Calpain-2 catalytic subunit Mus musculus | 1130.69 | 34 | 79872 | 20 | 20 | 52366493 | 53023433 | 44892601 | 43010409 | 48304345 | 54471217 | 44880845 | 55675521 | 50907121 | 46421989 | 51643121 | 47898801 | 50319253 | 51644265 |
| O88456 | Capns1 | Calpain small subunit 1 Mus musculus | 587.5 | 59.48 | 28463 | 3 | 9 | 30998365 | 34028041 | 29501311 | 25698957 | 30787103 | 35859877 | 35178441 | 38785485 | 35084545 | 32038903 | 35672193 | 33780065 | 35818065 | 37239785 |
| Q60865 | Caprin1 | Caprin-1 Mus musculus | 867.1 | 27.72 | 78169 | 15 | 15 | 24154185 | 14004299 | 20487127 | 14336851 | 13727249 | 19974611 | 15825306 | 22845165 | 24416619 | 16640469 | 15602373 | 21035469 | 20819297 | 15566890 |
| P47753 | Capza1 | F-actin-capping protein subunit alpha-1 Mus musculus | 528 | 45.1 | 32940 | 8 | 10 | 59613385 | 60653597 | 55100237 | 56731757 | 63947313 | 68557569 | 65010953 | 66143533 | 67203049 | 62702825 | 66975649 | 65080177 | 64062505 | 67503489 |
| P47754 | Capza2 | F-actin-capping protein subunit alpha-2 Mus musculus | 760.1 | 64.34 | 32967 | 12 | 14 | 74794169 | 70675769 | 65121949 | 68851193 | 73661457 | 78113313 | 68706857 | 72925649 | 73161097 | 70127649 | 78293633 | 75466857 | 70388849 | 83865745 |
| Q5XI32 | Capzb | F-actin-capping protein subunit beta Rattus norvegicus | 845.61 | 65.44 | 30629 | 17 | 17 | 66935225 | 84005313 | 81562849 | 80085769 | 77732337 | 75300721 | 89907721 | 84804353 | 91546657 | 92166425 | 84180169 | 84955609 | 77227089 | 85048625 |
| Q9D1I2 | Card19 | Caspase recruitment domain-containing protein 19 Mus musculus | 46.92 | 4.37 | 20936 | 1 | 1 | 1020654.75 | 886030.19 | 868942.94 | 1367625.75 | 960020.12 | 1150514.88 | 740120.44 | 1135957.88 | 1000704.38 | 1013113.19 | 1227685.5 | 261344.23 | 964746.44 | 958232.88 |
| A2AIV8 | Card9 | Caspase recruitment domain-containing protein 9 Mus musculus | 126.62 | 3.54 | 62462 | 2 | 2 | 722139.81 | 972047 | 650519.81 | 1374586.12 | 546785.94 | 529304.44 | 679611.25 | 596707.44 | 532104.81 | 542636.12 | 436703.28 | 1310598.88 | 586336.25 | 442106.25 |
| Q4AE70 | Carm1 | Histone-arginine methyltransferase CARM1 Rattus norvegicus | 134.61 | 7.53 | 70341 | 3 | 3 | 2199049.75 | 2187363.25 | 2090592 | 1574659.12 | 914684.31 | 1422664.88 | 987093.12 | 2170193.5 | 1983808 | 930212.38 | 1969643.12 | 2091633.25 | 2009237.25 | 2035114.5 |
| Q9ER72 | Cars | Cysteine--tRNA ligase, cytoplasmic Mus musculus | 955.15 | 23.83 | 94860 | 17 | 17 | 18081165 | 15996308 | 18579007 | 17411201 | 12776451 | 18610473 | 20338941 | 20677853 | 21007881 | 18411105 | 15022956 | 17093307 | 17232021 | 18480583 |
| P29452 | Casp1 | Caspase-1 Mus musculus | 890.3 | 41.29 | 45640 | 16 | 16 | 26517867 | 27996515 | 28144805 | 26615815 | 29227735 | 22356923 | 23036571 | 23089263 | 23713299 | 22791975 | 26056177 | 24088509 | 22483423 | 26143101 |
| P29594 | Casp2 | Caspase-2 Mus musculus | 45.19 | 1.99 | 50661 | 1 | 1 | 778569.94 | 877849.38 | 870675.62 | 927981.62 | 863769.14 | 796943.44 | 1 | 1 | 609012.81 | 1 | 853217.3133 | 794729.75 | 774508.69 | 990413.5 |
| P70677 | Casp3 | Caspase-3 Mus musculus | 315.97 | 29.6 | 31475 | 6 | 6 | 5906910.5 | 9678122 | 8811055 | 7282491 | 7355221.5 | 9663199 | 10159919 | 6736514 | 9906292 | 6934846 | 11242393 | 7794577 | 8044719 | 7248692.5 |
| O08738 | Casp6 | Caspase-6 Mus musculus | 100.48 | 6.88 | 31595 | 2 | 2 | 993009.38 | 1089150 | 1270771.62 | 714426.56 | 856172.25 | 943744.56 | 582711.12 | 1353685 | 1136692.12 | 669868.75 | 914504.56 | 904916.06 | 833895.75 | 888217.75 |
| P97864 | Casp7 | Caspase-7 Mus musculus | 38.54 | 3.3 | 34061 | 1 | 1 | 1 | 1 | 760315.75 | 1 | 640585.56 | 616397.31 | 1 | 1 | 1 | 561458.12 | 706785.81 | 605859.25 | 488161.41 | 489501.12 |
| O89110 | Casp8 | Caspase-8 Mus musculus | 381.44 | 21.25 | 55357 | 8 | 9 | 8308729 | 14283832 | 12380746 | 12562698 | 13577395 | 12535579 | 11580493 | 13401387 | 13002584 | 13805841 | 13181448 | 11826314 | 12891755 | 13517735 |
| Q8C3Q9 | Casp9 | Caspase-9 Mus musculus | 182.09 | 11.67 | 49979 | 3 | 3 | 553626.38 | 1743199.5 | 139710.88 | 1245217.12 | 232351.16 | 1540166.38 | 1069683.62 | 1054557.75 | 951622.62 | 1065528.5 | 1187083.25 | 218832.78 | 168178.47 | 95963.4 |
| P51125 | Cast | Calpastatin Mus musculus | 360.36 | 15.36 | 84922 | 6 | 6 | 3787703.5 | 7911193.5 | 6578158 | 8220052.5 | 8366228 | 6724197 | 8150081 | 7594524.5 | 8302021 | 7763717 | 7397415.5 | 6044353 | 8041788 | 8434093 |
| P24270 | Cat | Catalase Mus musculus | 326.05 | 19.54 | 59795 | 6 | 6 | 4729484 | 5277336 | 7071070 | 5557493 | 5415771 | 5435238 | 7528464 | 6658203 | 8201350.5 | 7857858.5 | 5824343 | 5459779 | 5519879.5 | 6473673.5 |
| Q08024 | Cbfb | Core-binding factor subunit beta Mus musculus | 275.05 | 29.41 | 22030 | 3 | 3 | 4640339 | 6989548 | 4756424 | 4919753.5 | 5366629.5 | 2918851.5 | 5764102 | 5343189 | 5146874.5 | 5062916 | 5322519 | 4987469.5 | 4620929.5 | 5750946 |
| P22682 | Cbl | E3 ubiquitin-protein ligase CBL Mus musculus | 166.92 | 6.24 | 100564 | 4 | 4 | 3931353.5 | 4262439 | 887598.12 | 3733659 | 2774012 | 3396660.5 | 2393085 | 2945851.25 | 2570821 | 1765039 | 3933943.5 | 4689575.5 | 1273245.62 | 4030883.75 |
| Q7TS56 | Cbr4 | Carbonyl reductase family member 4 Rattus norvegicus | 52.99 | 8.05 | 25287 | 1 | 1 | 277254.19 | 499812.47 | 559444.06 | 939096.25 | 859095.06 | 811766.12 | 1003628 | 580208.94 | 830395.44 | 978932.31 | 784682.44 | 727990.5 | 632208.12 | 1017201 |
| Q8VEH6 | Cbwd1 | COBW domain-containing protein 1 Mus musculus | 132.73 | 9.16 | 43772 | 2 | 2 | 1518167.38 | 361844.88 | 1746521.25 | 1475068.88 | 1537275.12 | 1960452.38 | 1745335.5 | 2236414.5 | 2241467.25 | 1859763.25 | 1796521.5 | 2317916.5 | 1876033 | 1950675.62 |
| P83917 | Cbx1 | Chromobox protein homolog 1 Mus musculus | 368.92 | 45.95 | 21418 | 4 | 5 | 12186138 | 15126752 | 13201249 | 14069240 | 18252925 | 17587037 | 18268437 | 16005048 | 17804937 | 18846813 | 18476091 | 18556811 | 16343273 | 18917501 |
| P23198 | Cbx3 | Chromobox protein homolog 3 Mus musculus | 440.56 | 36.61 | 20855 | 5 | 7 | 36266237 | 42719905 | 35782081 | 40401865 | 45056341 | 46545185 | 43567441 | 40244113 | 44586989 | 48252117 | 42534049 | 44824909 | 43279493 | 44795125 |
| Q61686 | Cbx5 | Chromobox protein homolog 5 Mus musculus | 318.98 | 34.55 | 22186 | 5 | 6 | 11663887 | 14353037 | 11469198 | 11127766 | 14267771 | 15773595 | 10965422 | 8443643 | 13049264 | 13886517 | 13440765 | 13981768 | 14500958 | 15374802 |
| Q8K1A6 | Cc2d1a | Coiled-coil and C2 domain-containing protein 1A Mus musculus | 60.43 | 2.12 | 103698 | 1 | 1 | 1 | 1 | 1 | 769127.75 | 1 | 455547.31 | 394483.97 | 313331.91 | 409874.78 | 393309.4925 | 533071.94 | 412420.81 | 447385.81 | 553934.88 |
| Q8BRN9 | Cc2d1b | Coiled-coil and C2 domain-containing protein 1B Mus musculus | 177.26 | 11.32 | 93091 | 4 | 4 | 554515.06 | 907815.56 | 706672 | 1 | 740409.44 | 1351665.5 | 1111377.36 | 1562820.75 | 782064.81 | 748958.38 | 1697086 | 1037577.94 | 794477.75 | 1176380.563 |
| Q8VDP4 | Ccar2 | Cell cycle and apoptosis regulator protein 2 Mus musculus | 257.02 | 8.68 | 103002 | 6 | 6 | 3319470 | 11913621 | 11904172 | 11383513 | 14229631 | 4019527.5 | 14548742 | 2525511.75 | 10940596 | 10844861 | 4214965.5 | 3715054.75 | 10578239 | 6721628 |
| Q8VE99 | Ccdc115 | Coiled-coil domain-containing protein 115 Mus musculus | 153.41 | 28.33 | 19743 | 3 | 3 | 887209.81 | 1163544.75 | 1355171.88 | 128817.3 | 939700.88 | 1000194.88 | 1080565.25 | 946051.75 | 996124.75 | 136740.97 | 1107655.12 | 1210485.88 | 1235707.62 | 1318086.25 |
| Q8R344 | Ccdc12 | Coiled-coil domain-containing protein 12 Mus musculus | 61.88 | 13.86 | 18891 | 1 | 1 | 1 | 1 | 1042977.25 | 877351.94 | 973050.62 | 1125311.88 | 1159305.12 | 1007518.69 | 928112.4925 | 420314.28 | 1 | 1121437.38 | 1 | 1 |
| Q9JIG7 | Ccdc22 | Coiled-coil domain-containing protein 22 Mus musculus | 209.62 | 13.24 | 70844 | 5 | 5 | 1427041.5 | 1870454.5 | 1686675.5 | 1998219.88 | 1917824 | 1150606.5 | 1376797.62 | 1428732.62 | 1529594 | 922738.69 | 1601055.5 | 1691953.75 | 897161.06 | 1533065.38 |
| Q9CR29 | Ccdc43 | Coiled-coil domain-containing protein 43 Mus musculus | 166.47 | 21.62 | 25049 | 3 | 3 | 2385759.25 | 2739219.5 | 2618146 | 2775880.25 | 2504314.75 | 1967098.25 | 2464458.5 | 3011190.5 | 2699677.5 | 1980375.12 | 2504569.25 | 2640884.25 | 2017821 | 2094758.25 |
| Q5U2X6 | Ccdc47 | Coiled-coil domain-containing protein 47 Rattus norvegicus 2 | 236.1 | 11.18 | 55733 | 5 | 5 | 5901036 | 6423451 | 7078342 | 6889855 | 2853142 | 5140765.5 | 5970487 | 6649914 | 2434206 | 3967522.75 | 4195603 | 8983615 | 3733527.25 | 5836013.5 |
| Q3ZAV0 | Ccdc60 | Coiled-coil domain-containing protein 60 Rattus norvegicus 2 | 39.47 | 1.48 | 62503 | 1 | 1 | 5209145 | 4348809 | 4012661 | 4671758 | 4392970.5 | 3891839.5 | 3754229.25 | 4674583 | 4623605 | 3987859.25 | 4414456 | 5232249.5 | 3723803.5 | 4626834.5 |
| Q5XIB5 | Ccdc86 | Coiled-coil domain-containing protein 86 Rattus norvegicus | 94.35 | 5.87 | 38585 | 2 | 2 | 2337335.25 | 1 | 1 | 826935 | 293555.72 | 727382.31 | 1 | 472617.75 | 1 | 279890.88 | 1046844.25 | 2378501.75 | 1 | 1 |
| Q4QRL3 | Ccdc88b | Coiled-coil domain-containing protein 88B Mus musculus | 136.3 | 2.5 | 166608 | 3 | 3 | 1218128.12 | 1376291.75 | 953901 | 1402982.25 | 1371815.5 | 1090789.62 | 1295014.75 | 1121500.5 | 947640.62 | 978185.19 | 1171822.75 | 860390.12 | 958294.81 | 1203234.25 |
| Q5BJT7 | Ccdc93 | Coiled-coil domain-containing protein 93 Rattus norvegicus | 150.33 | 5.25 | 72636 | 3 | 3 | 3673638 | 4200331.5 | 2638253 | 2660034.25 | 2721032.5 | 2262102 | 3807978 | 2329779.25 | 3298580.25 | 3323523.5 | 3766537.75 | 2597605.5 | 3442113.5 | 3449516.25 |
| P51943 | Ccna2 | Cyclin-A2 Mus musculus | 36.96 | 3.32 | 47269 | 1 | 1 | 1 | 1 | 1 | 1 | 1 | 475786.94 | 1 | 1 | 1 | 1 | 1 | 673796.62 | 571199.31 | 1 |
| P24860 | Ccnb1 | G2/mitotic-specific cyclin-B1 Mus musculus | 75.11 | 4.88 | 48052 | 1 | 2 | 1 | 1 | 210056.78 | 1 | 1 | 1 | 1 | 1 | 157967.27 | 218473.39 | 1 | 1 | 287020.5 | 430525.88 |
| P30276 | Ccnb2 | G2/mitotic-specific cyclin-B2 Mus musculus | 62.46 | 5.53 | 45383 | 1 | 2 | 1 | 1 | 210056.78 | 1 | 1 | 1 | 1 | 1 | 157967.27 | 218473.39 | 1 | 1 | 287020.5 | 252020.83 |
| P25322 | Ccnd1 | G1/S-specific cyclin-D1 Mus musculus | 116.65 | 8.14 | 33429 | 2 | 2 | 815802.12 | 537644.81 | 819930.62 | 446154.72 | 909177.38 | 391547.75 | 1202621 | 470958.44 | 1051428.62 | 1037181 | 520777.72 | 923510.06 | 478563.44 | 680934.06 |
| Q8BGU5 | Ccny | Cyclin-Y Mus musculus | 98.72 | 8.21 | 39395 | 2 | 2 | 1668393.38 | 500108.31 | 1841246 | 1178046.38 | 1366125.75 | 987759.56 | 1724228.12 | 1686097.88 | 1958774 | 1898790.5 | 887733.62 | 427332.44 | 1642426.25 | 1198567.12 |
| Q9WU84 | Ccs | Copper chaperone for superoxide dismutase Mus musculus | 129.26 | 18.25 | 28912 | 2 | 2 | 1 | 405529.62 | 485801.06 | 1 | 1 | 1633314.25 | 2479462.75 | 2229614.25 | 1918707.25 | 440717.06 | 1100983.62 | 363290.62 | 1130680.38 | 363333.5 |
| P80314 | Cct2 | T-complex protein 1 subunit beta Mus musculus | 2345.441 | 73.08 | 57477 | 6 | 36 | 94758409 | 77278281 | 87927633 | 78477073 | 75809265 | 79006049 | 81974217 | 93551937 | 80890057 | 84385945 | 81310673 | 79955953 | 86905761 | 83227089 |
| P80318 | Cct3 | T-complex protein 1 subunit gamma Mus musculus | 1900.41 | 63.85 | 60630 | 32 | 33 | 188583281 | 213317041 | 224270465 | 219631633 | 215753985 | 211003969 | 205981073 | 184887201 | 202176673 | 203044161 | 217903121 | 209931745 | 206336769 | 214108801 |
| P80315 | Cct4 | T-complex protein 1 subunit delta Mus musculus | 2634.18 | 79.59 | 58066 | 36 | 39 | 203020705 | 232607457 | 233029761 | 218289569 | 202955649 | 207244353 | 244357377 | 228984497 | 211899073 | 207601921 | 204513537 | 235012641 | 226230001 | 221996881 |
| P80316 | Cct5 | T-complex protein 1 subunit epsilon Mus musculus | 1763.8 | 72.09 | 59624 | 30 | 30 | 126087761 | 135700049 | 141273249 | 132033305 | 129718777 | 134232097 | 144698193 | 137283905 | 137293729 | 138679025 | 139075425 | 139749201 | 145495361 | 142004177 |
| P80317 | Cct6a | T-complex protein 1 subunit zeta Mus musculus | 1564.93 | 57.44 | 58004 | 21 | 25 | 103082353 | 93190521 | 98589273 | 101548561 | 99895985 | 98883553 | 99346577 | 100799841 | 97302561 | 93856609 | 101716089 | 98994993 | 102123841 | 100243409 |
| Q61390 | Cct6b | T-complex protein 1 subunit zeta-2 Mus musculus | 253.78 | 11.49 | 58185 | 1 | 5 | 5120923 | 3767310 | 2864682 | 3987979.25 | 4153049.25 | 2407713.25 | 3716256 | 3170837.75 | 4186694.75 | 3498025 | 5166962 | 2640864 | 3033813.25 | 2742968 |
| P80313 | Cct7 | T-complex protein 1 subunit eta Mus musculus | 1864.45 | 72.24 | 59652 | 29 | 30 | 96133929 | 99549185 | 108503177 | 99808177 | 103498409 | 97505161 | 98285521 | 96832977 | 94284193 | 101845529 | 101616073 | 99764921 | 104186745 | 101798705 |
| P42932 | Cct8 | T-complex protein 1 subunit theta Mus musculus | 1978.12 | 66.24 | 59555 | 30 | 31 | 155005009 | 161322865 | 171043905 | 172423009 | 167592417 | 169758017 | 168037777 | 158987969 | 167151249 | 172604273 | 173989025 | 170984881 | 176050657 | 182076369 |
| P10810 | Cd14 | Monocyte differentiation antigen CD14 Mus musculus | 919.15 | 51.37 | 39204 | 13 | 13 | 46938701 | 51451925 | 49639101 | 46202521 | 44246909 | 37399081 | 39845889 | 41435497 | 33770573 | 39076653 | 33665017 | 33582077 | 38490937 | 30320721 |
| Q62192 | Cd180 | CD180 antigen Mus musculus | 843.69 | 28.59 | 74302 | 14 | 14 | 24539871 | 22443247 | 26022701 | 18453029 | 21320899 | 21692441 | 18966443 | 16394237 | 21803559 | 18374365 | 20962877 | 16745570 | 19761449 | 17882425 |
| Q9JLQ0 | Cd2ap | CD2-associated protein Mus musculus | 209.12 | 8.32 | 70450 | 4 | 4 | 2313159 | 1819108.5 | 2543226 | 1921423.62 | 2484053.5 | 2692327 | 3158640.25 | 3662151.75 | 2628092 | 2563852 | 2924724.25 | 3169045.75 | 2384775.5 | 3175545.25 |
| Q9CWK3 | Cd2bp2 | CD2 antigen cytoplasmic tail-binding protein 2 Mus musculus | 42.89 | 4.09 | 37694 | 1 | 1 | 924238.69 | 937526.25 | 700660.44 | 880414.75 | 959233.62 | 666843.25 | 789463.25 | 701619.11 | 524462.38 | 825707.56 | 635182.56 | 594330.38 | 607225.56 | 716460.5 |

| | | | | | | | | | | | | | | | | | | | | | |
|---|---|---|---|---|---|---|---|---|---|---|---|---|---|---|---|---|---|---|---|---|---|
| P15379 | Cd44 | CD44 antigen Mus musculus | 248.12 | 4.5 | 85617 | 4 | 4 | 21419985 | 27528795 | 25653127 | 22303749 | 26399949 | 29994273 | 28164401 | 30882579 | 28157001 | 27638877 | 24428085 | 23396093 | 25576425 | 27238867 |
| Q61735 | Cd47 | Leukocyte surface antigen CD47 Mus musculus | 204.21 | 11.22 | 33098 | 3 | 3 | 10764981 | 8533328 | 10018729 | 7033180 | 9425581 | 10106561 | 8572951 | 10092247 | 8989319 | 8764565 | 9121427 | 8107942 | 10028924 | 9477439 |
| P13379 | Cd5 | T-cell surface glycoprotein CD5 Mus musculus | 303.12 | 14.98 | 53849 | 5 | 5 | 2857282.5 | 4351480.5 | 2455905 | 2650475 | 2625699.25 | 3878224.25 | 4265424 | 4472959 | 4133004.5 | 4678570 | 3483098.25 | 3104727 | 3640931 | 3884749.75 |
| P31996 | Cd68 | Macrosialin Mus musculus | 141.59 | 13.5 | 34818 | 3 | 3 | 100387481 | 94790417 | 93696321 | 89821873 | 111325049 | 102249753 | 92408129 | 99944465 | 103096065 | 98969457 | 102390057 | 87046081 | 94482225 | 104750681 |
| P21855 | Cd72 | B-cell differentiation antigen CD72 Mus musculus | 200.92 | 13.28 | 40347 | 4 | 4 | 5472007 | 4998917 | 4320119.5 | 4838255 | 4878189.5 | 5775861.5 | 6823931 | 5990614 | 6522884 | 5704567 | 5120527 | 5111056 | 4680059 | 5577287.5 |
| P42082 | Cd86 | T-lymphocyte activation antigen CD86 Mus musculus | 46.91 | 4.53 | 34666 | 1 | 1 | 3228302 | 3338046.5 | 3090599.25 | 3026812.25 | 3151998.25 | 4316440 | 4112833.25 | 4330235.5 | 3371236.5 | 3920199.25 | 3515333 | 3138004.25 | 3088859.75 | 3636546.5 |
| P40240 | Cd9 | CD9 antigen Mus musculus | 82.53 | 11.06 | 25258 | 1 | 1 | 6655790.5 | 1 | 1 | 1 | 993260.69 | 929114.56 | 818824.5 | 1068747 | 741804.06 | 822826.44 | 1374873.98 | 762260.69 | 2067185.75 | 1295175.5 |
| Q8CII2 | Cdc123 | Cell division cycle protein 123 homolog Mus musculus | 166.21 | 9.82 | 38816 | 3 | 3 | 1019178.25 | 1426347.5 | 1078103.25 | 1218294.12 | 1018506.25 | 1111336.62 | 1125850.12 | 1645885.25 | 1312669.25 | 988639.88 | 1262405 | 1273592.62 | 1249785.88 | 1359003.12 |
| Q8BGZ4 | Cdc23 | Cell division cycle protein 23 homolog Mus musculus | 100.45 | 4.02 | 68562 | 2 | 2 | 821238.69 | 1 | 1 | 312803.72 | 1 | 510519.84 | 486181.12 | 604683.38 | 581209.38 | 323043.75 | 519622 | 639545.94 | 555808.88 | 622236.44 |
| Q8CFI2 | Cdc34 | Ubiquitin-conjugating enzyme E2 R1 Mus musculus | 49.06 | 11.91 | 26622 | 1 | 1 | 190175.52 | 354373 | 278904.62 | 179926.92 | 421873.56 | 1 | 259551.36 | 261199.77 | 1 | 1 | 295548.72 | 155485.64 | 221905.72 | 310333.62 |
| Q61081 | Cdc37 | Hsp90 co-chaperone Cdc37 Mus musculus | 512.12 | 25.86 | 44593 | 11 | 11 | 17853611 | 10750638 | 13867500 | 10214349 | 13448626 | 13855706 | 14400365 | 15254864 | 13207591 | 12236786 | 14986703 | 12702154 | 12825406 | 11251374 |
| P60766 | Cdc42 | Cell division control protein 42 homolog Mus musculus | 537.62 | 58.64 | 21259 | 8 | 9 | 41102253 | 34127069 | 39225869 | 42724749 | 36411677 | 41515929 | 46319169 | 42854045 | 42987637 | 45671493 | 42492733 | 40941549 | 38666049 | 42134193 |
| Q6A068 | Cdc5l | Cell division cycle 5-like protein Mus musculus | 774.22 | 25.44 | 90192 | 13 | 13 | 4205941.5 | 5390162.5 | 6357427.5 | 8175404.5 | 6828637 | 6446000 | 5903174 | 2764722.25 | 6628491.5 | 4800660.5 | 8040491 | 7124752 | 5906947.5 | 7172435.5 |
| Q4V8C8 | Cdc73 | Parafibromin Rattus norvegicus | 210.79 | 10.36 | 60603 | 5 | 5 | 1646665.25 | 613052.5 | 676812.5 | 1749981.25 | 758705.88 | 585047.75 | 997735.06 | 856875.25 | 1131947.62 | 819897.75 | 1642948.88 | 1220319.62 | 969984.19 | 606363.94 |
| Q8VDP6 | Cdipt | CDP-diacylglycerol--inositol 3-phosphatidyltransferase Mus musculus | 291.04 | 37.56 | 23599 | 6 | 6 | 2878746.75 | 3684822.25 | 2994393.5 | 3307712.5 | 3192957 | 3905561.25 | 3547569.5 | 3557820.75 | 4082019.5 | 3441427.5 | 3275907.25 | 4079089.75 | 4043612 | 3909570.5 |
| P11440 | Cdk1 | Cyclin-dependent kinase 1 Mus musculus | 766.05 | 56.9 | 34107 | 13 | 14 | 9423399 | 6054676 | 7544436 | 6785699 | 5225055.5 | 3950288.5 | 6474866 | 5301028 | 4065637.5 | 5682500.5 | 6720254.5 | 9244485 | 8566037 | 5673077 |
| Q04735 | Cdk16 | Cyclin-dependent kinase 16 Mus musculus | 78.35 | 4.03 | 55914 | 1 | 2 | 450709.94 | 1 | 1 | 342831.16 | 155028.75 | 545313.31 | 1 | 252032.27 | 563089 | 484396 | 238805.97 | 377147.97 | 523778.94 | 197883.12 |
| Q04899 | Cdk18 | Cyclin-dependent kinase 18 Mus musculus | 295.61 | 17.07 | 51848 | 5 | 6 | 565470.75 | 616130.44 | 1319615.12 | 2423496 | 1317503.75 | 628049.69 | 215380.58 | 1255729.5 | 793675.44 | 778660.12 | 2143422.75 | 2542873.75 | 1171278.25 | 1434384.5 |
| P97377 | Cdk2 | Cyclin-dependent kinase 2 Mus musculus | 199.38 | 15.32 | 38978 | 1 | 4 | 1 | 337874.97 | 1 | 602628.19 | 1 | 1 | 442822.38 | 305784.94 | 482499.25 | 1 | 546422.56 | 561051.05 | 651342.62 | 485387.97 |
| O35207 | Cdk2ap1 | Cyclin-dependent kinase 2-associated protein 1 Mus musculus | 71.07 | 11.4 | 12354 | 1 | 1 | 1 | 1 | 141132.53 | 1 | 1 | 1 | 179434.5 | 182930.23 | 162344.55 | 1 | 1 | 85578.22 | 93567.16 | 1 |
| P30285 | Cdk4 | Cyclin-dependent kinase 4 Mus musculus | 162.21 | 15.84 | 33751 | 3 | 3 | 1 | 357160.56 | 1 | 1937852.62 | 1 | 1 | 1 | 370745.94 | 1 | 762954.25 | 2960195.75 | 451953.06 | 1 | 1 |
| P49615 | Cdk5 | Cyclin-dependent-like kinase 5 Mus musculus | 107.15 | 10.62 | 33288 | 3 | 3 | 1243612.25 | 2242653.25 | 2584870.25 | 1929038.88 | 1953487.5 | 2324289 | 2277975.5 | 2453818.25 | 2743163 | 2070568.62 | 2563500.25 | 2382581 | 2751822 | 2247027.25 |
| Q99LM2 | Cdk5rap3 | CDK5 regulatory subunit-associated protein 3 Mus musculus | 146.32 | 7.55 | 56991 | 3 | 3 | 594501.25 | 551663.5 | 323095.75 | 164270.67 | 355088.12 | 501550.84 | 147669.12 | 647854 | 429846.28 | 136933.91 | 290494.34 | 206236.77 | 469974.94 | 463462.66 |
| Q64261 | Cdk6 | Cyclin-dependent kinase 6 Mus musculus | 130.25 | 13.8 | 37028 | 3 | 3 | 602926.81 | 1661433.5 | 1175220 | 1501017 | 1249175.5 | 1397908.75 | 1407623.62 | 886018 | 637637.44 | 841904.31 | 1747608.38 | 949528.38 | 674198.06 | 1279503.25 |
| Q641Z4 | Cdk9 | Cyclin-dependent kinase 9 Rattus norvegicus | 93.84 | 5.91 | 42762 | 2 | 2 | 294274.34 | 389637.75 | 429396.91 | 227575.97 | 373023.84 | 343241.16 | 184427.45 | 360560 | 264498.41 | 369553.09 | 459910.16 | 282580.09 | 365317.22 | 335133.91 |
| Q5U2X0 | Cdkn2aip | CDKN2A-interacting protein Rattus norvegicus | 34.6 | 1.93 | 60744 | 1 | 1 | 1137211.38 | 1 | 1566931.25 | 392017.84 | 1 | 1 | 1 | 1 | 1 | 1 | 1 | 1 | 1 | 326992.72 |
| Q4VAA2 | Cdv3 | Protein CDV3 Mus musculus | 199.71 | 18.86 | 29729 | 3 | 3 | 7609962 | 8281187 | 6559194 | 7483821 | 9106882 | 7915252.5 | 7077676 | 4803765 | 7519069 | 7663126 | 9255784 | 7822206 | 7905480 | 8994160 |
| P28659 | Celf1 | CUGBP Elav-like family member 1 Mus musculus | 137.04 | 7.2 | 52107 | 2 | 3 | 989301.38 | 1403307.62 | 1312929.62 | 1368515.25 | 1008869 | 1223469.5 | 1476805.25 | 1264740.38 | 1453330.5 | 1442520.25 | 1220095.12 | 1379849.12 | 1354314.5 | 1426366.62 |
| Q792H5 | Celf2 | CUGBP Elav-like family member 2 Rattus norvegicus | 180.57 | 7.68 | 54271 | 2 | 3 | 957990.44 | 317687.5 | 1052966.25 | 816434.69 | 648280.06 | 1099133.5 | 1011289.62 | 963850.69 | 973417.94 | 1046881.25 | 1211629.167 | 1179579.5 | 1379732.62 | 1075575.38 |
| O88278 | Celsr3 | Cadherin EGF LAG seven-pass G-type receptor 3 Rattus norvegicus | 36.49 | 0.24 | 359355 | 1 | 1 | 3292740.75 | 3573454.5 | 4433192.5 | 2411884.5 | 3863846.25 | 4209626 | 2279722.75 | 4552911 | 4791733.5 | 3231159 | 3383461.25 | 3550319.5 | 4326520.5 | 3660083.25 |
| Q6A065 | Cep170 | Centrosomal protein of 170 kDa Mus musculus | 176.73 | 2.9 | 175050 | 4 | 4 | 6052194 | 4725799 | 4774246 | 6748361 | 4813490.5 | 4259247.5 | 3799332.25 | 5103960.5 | 3028309 | 3152480.5 | 4571319.5 | 4632852.5 | 3250293.5 | 4878807 |
| Q8BT07 | Cep55 | Centrosomal protein of 55 kDa Mus musculus | 89.24 | 4.55 | 53930 | 2 | 2 | 551833.94 | 1 | 682923.56 | 548555.31 | 1 | 831548.5 | 649003.75 | 508030.31 | 579707.38 | 435882.03 | 576330.9067 | 600202.19 | 612258.81 | 516531.72 |
| Q8BGS7 | Cept1 | Choline/ethanolaminephosphotransferase 1 Mus musculus | 135.24 | 8.89 | 46434 | 2 | 2 | 1634994 | 960700 | 1007702.62 | 882793.5 | 814427.62 | 1095620 | 566667.75 | 844200.19 | 741375.12 | 796001.06 | 824008.38 | 982540.12 | 1168294.75 | 967219.5 |
| Q924Z4 | Cers2 | Ceramide synthase 2 Mus musculus | 107.02 | 9.47 | 45024 | 2 | 2 | 4171508.5 | 1 | 1 | 1 | 1 | 1 | 1 | 1 | 1 | 294617.81 | 1 | 440388.34 | 1 | 374395.47 |
| Q9D6J1 | Cers4 | Ceramide synthase 4 Mus musculus | 33.15 | 2.04 | 46017 | 1 | 1 | 1075401 | 1 | 1 | 1 | 1 | 296311.59 | 1 | 420590.12 | 1 | 1 | 1 | 406310.72 | 417995.06 | 1 |
| Q9D6K9 | Cers5 | Ceramide synthase 5 Mus musculus | 111.7 | 9.18 | 48167 | 2 | 2 | 834003.69 | 202209.67 | 186993.09 | 234875.84 | 311980.56 | 194372.44 | 268970.72 | 187277.67 | 169072.38 | 224814.64 | 344351.69 | 124715.23 | 260832.33 | 298626.81 |
| Q8BTU1 | Cfap20 | Cilia- and flagella-associated protein 20 Mus musculus | 173.2 | 30.05 | 22748 | 4 | 4 | 7629927 | 8499859 | 8663983 | 7950704 | 6956509.5 | 6192437 | 5840535 | 6917020.5 | 4997719 | 5935685 | 4541339 | 6196100 | 4910781 | 5646966.5 |
| O88271 | Cfdp1 | Craniofacial development protein 1 Mus musculus | 215.9 | 21.02 | 32921 | 4 | 4 | 1691867.12 | 2441219.5 | 1857860.38 | 1772106.75 | 2289406 | 2137445 | 2519050 | 1992428 | 1655807 | 2340329 | 2233082.25 | 1362534.75 | 2771744.25 | 1499763.75 |
| P18760 | Cfl1 | Cofilin-1 Mus musculus | 813.63 | 74.7 | 18560 | 12 | 12 | 337236609 | 319565633 | 351056481 | 363054849 | 393487073 | 415494209 | 392304641 | 329385633 | 406457281 | 452801537 | 442737121 | 434238977 | 460522241 | 457483873 |
| Q9CRB9 | Chchd3 | MICOS complex subunit Mic19 Mus musculus | 238.76 | 18.94 | 26335 | 4 | 4 | 2915369 | 2642379.5 | 3237089.5 | 3782201.75 | 2855165.5 | 2825799 | 3725828.5 | 796654.19 | 2730573 | 3674128.25 | 3130501.5 | 3659010.5 | 2996796.5 | 2583546.25 |
| Q5BJN5 | Chchd4 | Mitochondrial intermembrane space import and assembly protein 40 Rattus norvegicus | 91.49 | 26.62 | 15467 | 2 | 2 | 2378723 | 2261497.75 | 1914399.5 | 1950681.75 | 2899819.5 | 2459515.5 | 1317610.25 | 2028469.5 | 1860177 | 2062091.88 | 2405035.25 | 2527461.5 | 3337418.75 | 2613116.5 |
| Q91VN4 | Chchd6 | MICOS complex subunit Mic25 Mus musculus | 145.23 | 15.02 | 29852 | 3 | 3 | 1233463 | 1436226.88 | 1608062.88 | 1344892.88 | 1678401.88 | 1578437.88 | 1714162.62 | 905464.25 | 1674662 | 1580228.5 | 1430479.5 | 1859128.25 | 1373426.12 | 1986407 |
| Q6PDQ2 | Chd4 | Chromodomain-helicase-DNA-binding protein 4 Mus musculus | 1228.67 | 15.87 | 217751 | 26 | 26 | 23176529 | 30768121 | 24352573 | 28578991 | 32957677 | 28366803 | 31980049 | 29506903 | 31496915 | 29623431 | 29913885 | 29675649 | 29684173 | 28968737 |
| O35280 | Chek1 | Serine/threonine-protein kinase Chk1 Mus musculus | 45.24 | 2.31 | 54381 | 1 | 1 | 1 | 1 | 1 | 1 | 538319.56 | 1 | 1 | 1 | 1 | 1 | 347540.3433 | 498485.75 | 266703.78 | 277431.5 |
| Q8CGZ0 | Cherp | Calcium homeostasis endoplasmic reticulum protein Mus musculus | 102.33 | 5.02 | 106169 | 3 | 3 | 1 | 1 | 1 | 1 | 1 | 632533.905 | 536792 | 640966.12 | 981653.81 | 370723.69 | 509218.72 | 1 | 1 | 795864.56 |
| P37727 | Chm | Rab proteins geranylgeranyltransferase component A 1 Rattus norvegicus | 127.88 | 4.31 | 72480 | 2 | 2 | 1589408.5 | 2299672 | 2029950.25 | 1617285.5 | 2542880.5 | 2621215.75 | 2100106 | 2540877.25 | 2412968.25 | 1818273.5 | 1671959.75 | 2510240.5 | 2463998.5 | 2515750.5 |
| Q8CGS4 | Chmp3 | Charged multivesicular body protein 3 Rattus norvegicus | 88 | 8.52 | 25062 | 2 | 2 | 1 | 1 | 315959.31 | 1 | 278388.94 | 1 | 1 | 1 | 196905.55 | 1 | 1 | 1 | 1 | 347687.28 |
| Q9D8B3 | Chmp4b | Charged multivesicular body protein 4b Mus musculus | 189.84 | 29.46 | 24936 | 5 | 5 | 3567356.25 | 3260036 | 2346466.25 | 3154089 | 2301949.25 | 1996828.12 | 3213589 | 3288482.75 | 2315636 | 3005479 | 2728655 | 3251262.5 | 3093455.5 | 2090782 |
| P0C0A3 | Chmp6 | Charged multivesicular body protein 6 Mus musculus | 47.93 | 6.5 | 23416 | 1 | 1 | 843311.44 | 977564.75 | 696011 | 936585.69 | 910120.06 | 819305.88 | 934772 | 874784.44 | 696232.12 | 1 | 888312.44 | 780695.81 | 838206.62 | 927197.44 |
| Q8R1T1 | Chmp7 | Charged multivesicular body protein 7 Mus musculus | 129.92 | 3.77 | 50633 | 2 | 2 | 587070.62 | 493761 | 444649.38 | 485101.81 | 473849.34 | 429037.25 | 681633 | 561710.62 | 546893.5 | 495777.38 | 414638.62 | 561452.75 | 626061.75 | 487315.53 |
| D4A4T9 | Chordc1 | Cysteine and histidine-rich domain-containing protein 1 Rattus norvegicus | 436.58 | 35.65 | 37362 | 9 | 9 | 8176075 | 9282825 | 8180874.5 | 8925010 | 7551701 | 10693211 | 15437735 | 9469615 | 10953471 | 11913802 | 11772235 | 8170458.5 | 9848022 | 11920077 |
| P61022 | Chp1 | Calcineurin B homologous protein 1 Mus musculus | 69.3 | 6.15 | 22432 | 1 | 1 | 1712638 | 1428324.25 | 1986900.5 | 1183351 | 1516923 | 1747236.88 | 2051698.5 | 2123146.75 | 1764717 | 1401110.25 | 1232417.75 | 1915153.38 | 2057047.38 | 1720372.75 |
| Q9CY57 | Chtop | Chromatin target of PRMT1 protein Mus musculus | 78.43 | 5.22 | 26585 | 1 | 1 | 1 | 1 | 1 | 1 | 1 | 1 | 1 | 1 | 1 | 1 | 1 | 663233.69 | 1 | 713091.38 |
| Q60680 | Chuk | Inhibitor of nuclear factor kappa-B kinase subunit alpha Mus musculus | 68.26 | 3.36 | 84729 | 2 | 2 | 713013.5 | 800140.5 | 876958.75 | 758293.38 | 912225.06 | 874086.75 | 764886.06 | 673511.56 | 1128456.25 | 734128.81 | 672710.88 | 701683 | 1126378.25 | 819159.75 |
| Q9DCL2 | Ciao2a | Cytosolic iron-sulfur assembly component 2A Mus musculus | 43.48 | 5.62 | 18418 | 1 | 1 | 1704677.88 | 1674099.25 | 1883725.62 | 1286717 | 1468428.25 | 1448722 | 1251545.75 | 1714384 | 1224645.75 | 1372792.62 | 1551750.62 | 1723733.62 | 1503416.62 | 1637056.75 |
| Q8WTY4 | Ciapin1 | Anamorsin Mus musculus | 383.78 | 35.28 | 33429 | 7 | 7 | 4608548.5 | 8163353 | 5240400 | 7114874.5 | 7039459.5 | 8085573 | 9440385 | 7324889.5 | 8398237 | 9839864 | 7227465 | 6190520.5 | 8402461 | 7653529 |
| Q8BWY9 | Cip2a | Protein CIP2A Mus musculus | 52.33 | 1.54 | 102043 | 1 | 1 | 1 | 1 | 1 | 1 | 1 | 1 | 1 | 1 | 1 | 1 | 219241.22 | 1 | 274027.12 | 1 |
| B0K020 | Cisd1 | CDGSH iron-sulfur domain-containing protein 1 Rattus norvegicus | 36.33 | 8.33 | 12097 | 1 | 1 | 1284328.875 | 1110574 | 1254126.12 | 1242726.5 | 1529888.88 | 1696193.62 | 1189677.25 | 728460.94 | 1362649.75 | 1890749.5 | 1725112 | 1707670.12 | 1733234.88 | 1801308.25 |
| Q9CQB5 | Cisd2 | CDGSH iron-sulfur domain-containing protein 2 Mus musculus | 172.63 | 31.11 | 15242 | 4 | 4 | 7221406 | 4986387.5 | 5483813 | 8125508 | 5127212 | 8731020 | 8204317 | 5228982 | 7976130 | 8636327 | 5923902 | 9108253 | 8715005 | 8611047 |
| Q8BMK4 | Ckap4 | Cytoskeleton-associated protein 4 Mus musculus | 932.77 | 36.7 | 63692 | 15 | 15 | 21448823 | 17194699 | 19210087 | 15689124 | 16096324 | 20157879 | 20214697 | 19149877 | 19123517 | 20145421 | 19876229 | 20801457 | 19346373 | 19840185 |
| A2AGT5 | Ckap5 | Cytoskeleton-associated protein 5 Mus musculus | 888.94 | 11.71 | 225635 | 20 | 20 | 21181283 | 23680455 | 23635451 | 20457251 | 20914393 | 23192951 | 22385191 | 17000595 | 25066245 | 22049393 | 21900993 | 25819479 | 22006245 | 21879563 |
| Q04447 | Ckb | Creatine kinase B-type Mus musculus | 1693.04 | 67.72 | 42713 | 20 | 23 | 255307153 | 251038737 | 223875297 | 278372481 | 251370417 | 271377857 | 135431441 | 266102673 | 246365249 | 244120577 | 238150417 | 273051873 | 235959041 | 245760577 |
| P00564 | Ckm | Creatine kinase M-type Rattus norvegicus | 47.42 | 8.4 | 43045 | 1 | 1 | 2756509.75 | 1012554.81 | 1432916.62 | 2873567.75 | 3550010.25 | 2595258 | 2379591.5 | 1811409.38 | 2295154.25 | 1503600.38 | 4942392.5 | 2878271.25 | 865102.06 | 2237395.75 |
| Q80TV8 | Clasp1 | CLIP-associating protein 1 Mus musculus | 136.78 | 1.95 | 169227 | 1 | 3 | 739252.69 | 1063765 | 701905.12 | 591517.31 | 617944 | 784381.69 | 1133370.12 | 920357.94 | 1109469.75 | 528210.44 | 623078.81 | 572489.69 | 479337.09 | 885741.5 |
| Q8BRT1 | Clasp2 | CLIP-associating protein 2 Mus musculus | 128.81 | 2.41 | 140739 | 1 | 3 | 739252.69 | 715702 | 453115.66 | 591517.31 | 617944 | 784381.69 | 531339.81 | 792376.81 | 542951.94 | 528210.44 | 623078.81 | 572489.69 | 479337.09 | 885741.5 |
| Q99LI2 | Clcc1 | Chloride channel CLIC-like protein 1 Mus musculus | 183.63 | 8.91 | 60621 | 3 | 3 | 1968337.25 | 1 | 571174.94 | 272969.69 | 1 | 475675.22 | 1 | 274638.72 | 597659.75 | 1 | 772083 | 780437.62 | 613604.5 | 368880.69 |
| P49300 | Clec10a | C-type lectin domain family 10 member A Mus musculus | 187.5 | 15.46 | 34596 | 4 | 4 | 2979940.25 | 2545870.25 | 3094110.5 | 3825382.25 | 2324808 | 3790145.5 | 3498472.25 | 3357119.5 | 2732623.5 | 3931328 | 2891476.5 | 4575340 | 2586123.25 | 3750628.5 |
| Q504P2 | Clec12a | C-type lectin domain family 12 member A Mus musculus | 58.83 | 3.37 | 30757 | 1 | 1 | 833108.38 | 804176.25 | 742673.62 | 603270.94 | 734008.75 | 576056.62 | 481652.47 | 700981.69 | 642220.5 | 635727.38 | 702211.62 | 656591.62 | 655670.62 | 667034.38 |
| Q6MG61 | Clic1 | Chloride intracellular channel protein 1 Rattus norvegicus | 1123.47 | 62.66 | 26981 | 14 | 14 | 202635617 | 232196769 | 213661681 | 198470033 | 209748017 | 235330193 | 247376785 | 239691601 | 249050065 | 246995137 | 221458097 | 228320849 | 229401105 | 240769345 |
| Q9QYB1 | Clic4 | Chloride intracellular channel protein 4 Mus musculus | 1131.61 | 74.31 | 28729 | 17 | 17 | 97964481 | 99478145 | 87956129 | 100850505 | 100354753 | 120676057 | 119147137 | 113144737 | 129646161 | 139642721 | 116781297 | 118015865 | 113190041 | 127598233 |
| Q99KN9 | Clint1 | Clathrin interactor 1 Mus musculus | 269.06 | 10.14 | 68513 | 5 | 5 | 10063867 | 10178283 | 9416835 | 8592088 | 9400185 | 9317638 | 9756241 | 10213264 | 9471971 | 9561411 | 9258390 | 9215317 | 8910421 | 10801455 |
| Q922J3 | Clip1 | CAP-Gly domain-containing linker protein 1 Mus musculus | 279.64 | 5.1 | 155814 | 5 | 5 | 3655231 | 1922632.5 | 3088183 | 4579470 | 4091953.25 | 2380725.75 | 2342367 | 3485804.25 | 2344430 | 2784125.75 | 4326262 | 5447502.5 | 2502129.75 | 3680019.5 |
| Q3UMW8 | Cln5 | Ceroid-lipofuscinosis neuronal protein 5 homolog Mus musculus | 100.62 | 4.69 | 39329 | 2 | 2 | 2946472.75 | 2793477.5 | 2899517.25 | 1730103.12 | 2395201 | 2408217.25 | 1681281.5 | 2294859.5 | 2334099 | 2332409.5 | 2213799.25 | 2033994.75 | 2277659 | 2474815.75 |
| Q8VBZ3 | Clptm1 | Cleft lip and palate transmembrane protein 1 homolog Mus musculus | 260.95 | 11.3 | 75291 | 4 | 4 | 8212460 | 8418120 | 5859314 | 5419005.5 | 6493413.5 | 3923409.5 | 2448907.75 | 5876071.5 | 6251724 | 7070323.5 | 6874873.5 | 9541077 | 11194607 | 8531515 |
| Q5U2U0 | Clpx | ATP-dependent Clp protease ATP-binding subunit clpX-like, mitochondrial Rattus norvegic | 222.6 | 10.43 | 69208 | 5 | 5 | 3114605.75 | 1965507.62 | 24007401 | 3122183.25 | 43430297 | 19265385 | 15251105 | 2872975 | 2281291.75 | 2538111 | 38935265 | 36446213 | 2893644 | 1702267.75 |
| O08585 | Clta | Clathrin light chain A Mus musculus | 123.19 | 10.64 | 25604 | 3 | 3 | 22239505 | 23305617 | 20524421 | 27492887 | 23438983 | 19110209 | 19091705 | 19940957 | 16651577 | 19197859 | 20148757 | 17940843 | 18731121 | 19499133 |
| Q68FD5 | Cltc | Clathrin heavy chain 1 Mus musculus | 5817.88 | 65.13 | 191557 | 85 | 85 | 464104705 | 492453857 | 482883425 | 470813473 | 474982721 | 415601153 | 443680929 | 478636065 | 465366945 | 446243137 | 464605345 | 439327233 | 445865025 | 460420769 |
| Q5SW19 | Cluh | Clustered mitochondria protein homolog Mus musculus | 502.78 | 11.03 | 148068 | 11 | 11 | 15011846 | 13132903 | 10006947 | 8579896 | 11412827 | 8735745 | 10435339 | 10769481 | 10549735 | 8701678 | 10097520 | 10191227 | 11741956 | 11772810 |
| Q8R4N0 | Clybl | Citramalyl-CoA lyase, mitochondrial Mus musculus | 129.86 | 14.79 | 37549 | 3 | 3 | 1999024.75 | 2276726 | 1732721.88 | 2577930.25 | 2283561.5 | 2726415.25 | 2628278.75 | 1777678 | 2627799.5 | 2506425 | 2693341.5 | 2636959.25 | 2170393.5 | 2551797.75 |
| Q99KK2 | Cmas | N-acylneuraminate cytidylyltransferase Mus musculus | 181.28 | 11.11 | 48058 | 4 | 4 | 11565397 | 13408209 | 12874419 | 9874251 | 13388289 | 11534415 | 12542103 | 13146943 | 12730071 | 14084919 | 12019947 | 12069673 | 14923693 | 13165894 |
| Q9DBP5 | Cmpk1 | UMP-CMP kinase Mus musculus | 367.8 | 47.45 | 22165 | 7 | 7 | 13344170 | 12639166 | 15350142 | 12876946 | 14379416 | 16518671 | 14282503 | 12153021 | 15923792 | 17102335 | 15744873 | 17061547 | 16354873 | 14364234 |
| Q9DBC3 | Cmtr1 | Cap-specific mRNA (nucleoside-2'-O-)-methyltransferase 1 Mus musculus | 100.11 | 3.94 | 95676 | 3 | 3 | 1037354.94 | 2110121.5 | 936033.56 | 1068002.62 | 2170214.5 | 2026412.75 | 1909432.5 | 1826958.25 | 2232509.5 | 2137789 | 1708831.12 | 1211519.38 | 2153971.5 | 839732.62 |
| P62634 | Cnbp | Cellular nucleic acid-binding protein Rattus norvegicus | 251.06 | 28.25 | 19463 | 4 | 4 | 7434644.5 | 13141701 | 11193080 | 14584421 | 11707286 | 12889126 | 11893131 | 13871277 | 11734475 | 13658721 | 11067409 | 11123751 | 7670785 | 11554949 |
| Q9D1A2 | Cndp2 | Cytosolic non-specific dipeptidase Mus musculus | 1509.08 | 60.84 | 52767 | 24 | 24 | 35378917 | 43228145 | 46832133 | 35846397 | 39982001 | 41995897 | 41264069 | 41579485 | 39621921 | 42616057 | 42005725 | 45351257 | 46290005 | 40625013 |
| Q9CX13 | Cnih4 | Protein cornichon homolog 4 Mus musculus | 67.89 | 14.39 | 16089 | 1 | 1 | 4690036 | 1326770.5 | 1530951.62 | 1344254.12 | 2211565.25 | 2823673.5 | 1283043.5 | 1115214.75 | 2444365.25 | 2690157.5 | 2211388.25 | 2719572 | 3219945.5 | 3042280.5 |
| Q6ZQ08 | Cnot1 | CCR4-NOT transcription complex subunit 1 Mus musculus | 298.98 | 4.46 | 266808 | 8 | 8 | 3597935.5 | 3165644.5 | 4145893 | 3276575.5 | 2317906.5 | 3972465.5 | 1967631 | 1727759.88 | 2090027.25 | 1757433.25 | 3458114 | 1963443.75 | 1753191.75 | 1209799.5 |
| Q5XIA4 | Cnot10 | CCR4-NOT transcription complex subunit 10 Rattus norvegicus 2 | 60.17 | 1.61 | 81819 | 1 | 1 | 716241.38 | 1 | 325269.28 | 645010.31 | 1 | 1 | 1 | 471293.38 | 533708.75 | 1 | 1 | 1 | 1 | 1 |
| Q8K0V4 | Cnot3 | CCR4-NOT transcription complex subunit 3 Mus musculus | 97.76 | 4.13 | 81946 | 2 | 2 | 575017.19 | 608880.31 | 571949.31 | 805101.12 | 757929 | 756296.88 | 678729.69 | 529119.25 | 627372.94 | 677257.5 | 704720.56 | 118731.84 | 600757.12 | 474736.5067 |
| Q60809 | Cnot7 | CCR4-NOT transcription complex subunit 7 Mus musculus | 78.56 | 10.18 | 32718 | 2 | 2 | 1 | 452534.75 | 1 | 1 | 321391.88 | 544725.06 | 468870.06 | 738707.44 | 565524.69 | 108246.51 | 73928.09 | 423870.47 | 296847.09 | 264881.8833 |

| | | | | | | | | | | | | | | | | | | | | | |
|---|---|---|---|---|---|---|---|---|---|---|---|---|---|---|---|---|---|---|---|---|---|
| Q5PQL2 | Cnot9 | CCR4-NOT transcription complex subunit 9 Rattus norvegicus | 132.57 | 11.71 | 33601 | 3 | 3 | 3470911 | 3178055.75 | 3712563 | 2878176.25 | 2229217 | 2331417.25 | 2866343.5 | 3227667.5 | 2064978.5 | 1992703.75 | 2590431.25 | 2922241.5 | 2262396.5 | 2818674 |
| P13233 | Cnp | 2',3'-cyclic-nucleotide 3'-phosphodiesterase Rattus norvegicus | 99.23 | 5.71 | 47268 | 2 | 2 | 177527.52 | 200313.7825 | 131076.45 | 256356.52 | 236294.64 | 151559.52 | 1 | 178914.59 | 150637.64 | 1 | 1 | 1 | 142196.69 | 193305.5 |
| Q9QXT0 | Cnpy2 | Protein canopy homolog 2 Mus musculus | 147.19 | 15.93 | 20767 | 2 | 2 | 2663602.5 | 2918795.5 | 2321279.5 | 2802940.5 | 2482477.5 | 2203988.5 | 2372301.75 | 2489026.25 | 1994294.12 | 2230383 | 2376435.75 | 2230855.25 | 2095805.5 | 2558675 |
| Q8BQ47 | Cnpy4 | Protein canopy homolog 4 Mus musculus | 35.44 | 4.9 | 28094 | 1 | 1 | 926013.4675 | 1050913.62 | 921976.56 | 817126.19 | 914037.5 | 896065 | 812071.25 | 976427.81 | 786182.62 | 787716.81 | 892688.19 | 1153545.62 | 1053613.38 | 1096158.5 |
| Q5M7A7 | Cnrip1 | CB1 cannabinoid receptor-interacting protein 1 Rattus norvegicus | 57.64 | 9.76 | 18658 | 1 | 1 | 1062242.12 | 911571.5 | 1075807 | 917962.69 | 1316742.88 | 965356 | 818823.38 | 763043.19 | 1072274.88 | 898793.44 | 1153807.5 | 997375.81 | 953448.12 | 1102915.12 |
| Q9DBL7 | Coasy | Bifunctional coenzyme A synthase Mus musculus | 180.57 | 12.79 | 62023 | 4 | 4 | 2029152.75 | 1589304.88 | 1651493.75 | 1497238.62 | 1222557.25 | 1593249.75 | 1714448 | 1927328.5 | 1897090.5 | 1230288.62 | 1268273.38 | 1471678.12 | 1149914.25 | 1172533 |
| Q9Z160 | Cog1 | Conserved oligomeric Golgi complex subunit 1 Mus musculus | 108.17 | 5.2 | 109050 | 3 | 3 | 1624671.5 | 1343315.75 | 1836370.12 | 1212280.12 | 1640043.62 | 1631342.5 | 1405041.62 | 1118134 | 1354409.25 | 1154200.88 | 1612963 | 967780.75 | 1533826.75 | 1228718.25 |
| Q8C0L8 | Cog5 | Conserved oligomeric Golgi complex subunit 5 Mus musculus | 61.13 | 1.69 | 91391 | 1 | 1 | 666407.69 | 692205.75 | 561824.38 | 637569.38 | 589091.31 | 640687.81 | 573383.62 | 596897.06 | 574073.06 | 472994.34 | 778006.31 | 550074.5 | 691751.75 | 622889.44 |
| Q3UM29 | Cog7 | Conserved oligomeric Golgi complex subunit 7 Mus musculus | 39.89 | 1.3 | 86073 | 1 | 1 | 877942.88 | 1048416.69 | 953760.94 | 772125.38 | 767377.81 | 623994.62 | 1106338.5 | 692372.56 | 660758.62 | 799329.5 | 755083.12 | 687895.31 | 700532.94 | 664636.44 |
| Q5SU73 | Coil | Coilin Mus musculus | 68.24 | 1.75 | 61917 | 1 | 1 | 716566.94 | 651259.75 | 686701.88 | 586638.19 | 793094.94 | 629922.06 | 591245.19 | 602385.62 | 471311.16 | 658201.12 | 620393.75 | 612193.94 | 670877.94 | 684768.62 |
| Q9EQG9 | Col4a3bp | Collagen type IV alpha-3-binding protein Mus musculus | 226.77 | 10.26 | 71111 | 4 | 4 | 2020628 | 2719257.5 | 2112373.5 | 2053642.25 | 2452292 | 2120234.25 | 1918969.5 | 2021722.88 | 2269175 | 2366688.25 | 2036609.75 | 1283330.12 | 2297424.5 | 2083307.5 |
| Q8K4Q8 | Colec12 | Collectin-12 Mus musculus | 136.96 | 4.72 | 81304 | 2 | 2 | 1318988.75 | 1573271.25 | 1555674 | 927166.12 | 1725151.75 | 1588737.38 | 1729193.25 | 1288371.815 | 748772.75 | 1086783.88 | 523929.69 | 689163.8333 | 720038.06 | 823523.75 |
| Q8K297 | Colgalt1 | Procollagen galactosyltransferase 1 Mus musculus | 390.19 | 14.26 | 71061 | 8 | 8 | 5236550 | 5642999.5 | 5347797 | 5270898 | 5695256 | 6070436.5 | 5159578 | 6098443.5 | 6172645 | 5136687.5 | 5246438 | 6807876.5 | 5565680 | 7482563.5 |
| Q8BXC6 | Commd2 | COMM domain-containing protein 2 Mus musculus | 74.28 | 9.55 | 22848 | 2 | 2 | 2578403.25 | 2830499.25 | 2162274.75 | 2516614.75 | 2383693 | 2175381.5 | 2183641.25 | 2800303.25 | 2028409.12 | 2099681 | 1967882 | 2133353.25 | 2020462.88 | 2087530.5 |
| Q63829 | Commd3 | COMM domain-containing protein 3 Mus musculus | 218.1 | 22.05 | 22037 | 3 | 3 | 2526894.25 | 2162907.75 | 1817149.75 | 2571232.75 | 2248228.75 | 1681704.38 | 2381098.25 | 2720431 | 1840457.62 | 1978124.88 | 2339141.25 | 1770864.25 | 1912515.5 | 2047477.38 |
| Q9CQ02 | Commd4 | COMM domain-containing protein 4 Mus musculus | 99.52 | 17.59 | 21860 | 2 | 2 | 740026.56 | 614819 | 679925.75 | 539464.88 | 352466.81 | 571773.31 | 502967.12 | 716800.5 | 722073.69 | 527381.25 | 517627.19 | 678309.69 | 648040.06 | 631816.69 |
| O88587 | Comt | Catechol O-methyltransferase Mus musculus | 409.03 | 37.74 | 29486 | 6 | 6 | 5544021 | 1821514.25 | 5037485 | 2181069 | 3859265.25 | 7328726.5 | 4292333.5 | 7694870 | 7431140.5 | 4228091 | 4993448.5 | 5020969.5 | 6315986.5 | 4651063 |
| Q8BIG7 | Comtd1 | Catechol O-methyltransferase domain-containing protein 1 Mus musculus | 557.36 | 41.22 | 28961 | 8 | 8 | 9628291 | 9743043 | 5514307.5 | 7986243 | 8284010 | 8090972.5 | 7046741.5 | 7517129 | 7348340 | 6038652 | 8815913 | 10030180 | 9475046 | 8113273 |
| Q8CIE6 | Copa | Coatomer subunit alpha Mus musculus | 2029.95 | 35.7 | 138432 | 37 | 37 | 80718177 | 78354401 | 77292553 | 71220417 | 76961153 | 76225545 | 94914649 | 74714825 | 76847993 | 75294233 | 74371009 | 73115337 | 69805529 | 73831233 |
| Q9JIF7 | Copb1 | Coatomer subunit beta Mus musculus | 1995.78 | 51.52 | 107066 | 35 | 35 | 56270101 | 53143837 | 54010401 | 48913977 | 52834477 | 52875381 | 53307925 | 56770893 | 52587849 | 49603073 | 47297009 | 53058925 | 51639557 | 52136897 |
| O35142 | Copb2 | Coatomer subunit beta' Rattus norvegicus | 1590.56 | 45.3 | 102551 | 29 | 29 | 65436313 | 74778705 | 72073345 | 71345665 | 73445985 | 71311281 | 66888857 | 74449553 | 77249801 | 67317985 | 67867337 | 67384737 | 70302529 | 73223969 |
| O89079 | Cope | Coatomer subunit epsilon Mus musculus | 679.2 | 45.45 | 34567 | 8 | 8 | 11416745 | 11867459 | 14669194 | 13050814 | 13300484 | 13739992 | 14488589 | 17353559 | 17527815 | 12454122 | 13831607 | 11467524 | 15337310 | 14342049 |
| Q9QZE5 | Copg1 | Coatomer subunit gamma-1 Mus musculus | 1278.92 | 42.33 | 97513 | 21 | 24 | 51265409 | 77387665 | 41395341 | 85211601 | 50153717 | 69680065 | 56873689 | 51660917 | 48596737 | 37765705 | 38323677 | 41299797 | 51419093 | 59829849 |
| Q9QXK3 | Copg2 | Coatomer subunit gamma-2 Mus musculus | 814.5 | 24.91 | 97681 | 12 | 15 | 40872573 | 61695881 | 28478757 | 73457185 | 41367229 | 57687017 | 47531941 | 39211593 | 37441105 | 25926703 | 24035741 | 31827941 | 38928285 | 45820569 |
| P61202 | Cops2 | COP9 signalosome complex subunit 2 Mus musculus | 427.81 | 25.51 | 51597 | 8 | 8 | 8871354 | 6822554 | 6067290 | 6486620.5 | 7420980.5 | 8357043 | 8141436.5 | 8263420 | 7309333 | 2755083.75 | 7158050 | 7439761 | 7633820 | 7263402 |
| O88543 | Cops3 | COP9 signalosome complex subunit 3 Mus musculus | 504.81 | 34.28 | 47832 | 10 | 10 | 13723129 | 12723765 | 11742243 | 12190972 | 12781533 | 13277078 | 9498319 | 12348987 | 11395079 | 12487233 | 13852721 | 12409287 | 11879336 | 12484594 |
| O88544 | Cops4 | COP9 signalosome complex subunit 4 Mus musculus | 650.88 | 33.99 | 46285 | 10 | 10 | 10610030 | 11319453 | 7651990 | 10390833 | 7450855 | 9540383 | 9371411 | 12226973 | 10104827 | 9626457 | 9606605 | 7517886 | 9766280 | 8065464 |
| O35864 | Cops5 | COP9 signalosome complex subunit 5 Mus musculus | 418.43 | 38.02 | 37549 | 9 | 9 | 12679536 | 13750784 | 13470223 | 12646730 | 11731257 | 12452033 | 9107874 | 15119111 | 13521951 | 11036580 | 11066795 | 12008852 | 11378255 | 12763292 |
| O88545 | Cops6 | COP9 signalosome complex subunit 6 Mus musculus | 83.87 | 9.57 | 35880 | 3 | 3 | 2011342.5 | 1936682.5 | 1757356.12 | 3251360 | 1788448.25 | 3068645 | 2302460.25 | 1994257.38 | 2035653.62 | 2917137.75 | 1581466 | 1807321.12 | 1836849.5 | 2729308.5 |
| Q9CZ04 | Cops7a | COP9 signalosome complex subunit 7a Mus musculus | 330.49 | 26.91 | 30224 | 5 | 5 | 1254091.88 | 881626.88 | 919515.62 | 940309.25 | 818783 | 1336141.5 | 1205929.12 | 827096.19 | 1298947.12 | 1039810.62 | 935139.62 | 842781.12 | 1074338.25 | 1275887.75 |
| Q8BV13 | Cops7b | COP9 signalosome complex subunit 7b Mus musculus | 179.7 | 21.97 | 29689 | 4 | 4 | 1596074.75 | 1832160.75 | 2051897 | 1354905 | 1326802.75 | 927606.56 | 1415874.25 | 1601701.75 | 1435927.5 | 1206279.62 | 943214.88 | 1120390.25 | 2364326.5 | 1302848.25 |
| Q8VBV7 | Cops8 | COP9 signalosome complex subunit 8 Mus musculus | 155.15 | 22.97 | 23256 | 3 | 3 | 2836882 | 2869992.25 | 3055049.25 | 3014558.75 | 3912004 | 2849256.5 | 3031056.75 | 3339655 | 3378296.25 | 3183738.25 | 3249900.5 | 3189873 | 3028594.75 | 2880103.25 |
| P61924 | Copz1 | Coatomer subunit zeta-1 Mus musculus | 326.15 | 51.41 | 20198 | 6 | 6 | 16257596 | 18866305 | 17206581 | 14601937 | 16798483 | 11527423 | 17590537 | 15983920 | 15814931 | 15756122 | 17022019 | 16533650 | 17412413 | 17155155 |
| P97478 | Coq7 | 5-demethoxyubiquinone hydroxylase, mitochondrial Mus musculus | 38 | 11.98 | 24042 | 1 | 1 | 170601.22 | 236639.27 | 152764.27 | 219756.44 | 170123.34 | 155928.64 | 178439.88 | 156339.64 | 171033.34 | 149953.09 | 209848.77 | 225781.33 | 171284.61 | 146940.08 |
| Q6AY19 | Coq8b | Atypical kinase COQ8B, mitochondrial Rattus norvegicus | 34.75 | 1.89 | 58905 | 1 | 1 | 3658946.25 | 2610860.5 | 3624262.25 | 2453319 | 2308136 | 2670578.5 | 2206137 | 3805195 | 3090057.25 | 2122850.25 | 2273137.75 | 2297229 | 2243443.25 | 2025130.12 |
| O89053 | Coro1a | Coronin-1A Mus musculus | 1296.8 | 52.49 | 50989 | 20 | 21 | 234062945 | 239528401 | 267821985 | 303063969 | 299673473 | 282927009 | 256563169 | 207672001 | 266206033 | 325302945 | 308543233 | 280389953 | 287637089 | 286338049 |
| O89046 | Coro1b | Coronin-1B Rattus norvegicus | 128.19 | 4.96 | 53845 | 2 | 2 | 4789922 | 5263920 | 4542603 | 5334586 | 5267367 | 4636197 | 4861552.5 | 2107214.25 | 3882279.25 | 4877946.5 | 5791354 | 4388930 | 4555298 | 4487325 |
| Q9WUM4 | Coro1c | Coronin-1C Mus musculus | 900.89 | 43.25 | 53121 | 15 | 15 | 52956961 | 61785905 | 62616461 | 66189629 | 63500001 | 56474657 | 58290029 | 48318501 | 55190049 | 63327849 | 66537665 | 58575749 | 57717061 | 65580065 |
| Q9D2V7 | Coro7 | Coronin-7 Mus musculus | 719.38 | 33.41 | 100812 | 11 | 15 | 4761044 | 6848584.5 | 5650407 | 9400334 | 6425346.5 | 7879861 | 7669433.5 | 6256078 | 6944286 | 7000851.5 | 10415321 | 7396715 | 7700529 | 8620052 |
| Q9CQI6 | Cotl1 | Coactosin-like protein Mus musculus | 560.68 | 72.54 | 15944 | 11 | 11 | 82886673 | 79513065 | 77089809 | 91332793 | 108169353 | 109589441 | 75085633 | 66209325 | 90163481 | 98840305 | 98984233 | 92546001 | 92868561 | 97721345 |
| Q8BJ03 | Cox15 | Cytochrome c oxidase assembly protein COX15 homolog Mus musculus | 40.99 | 4.36 | 45853 | 1 | 1 | 422081.69 | 1 | 1 | 1 | 1 | 1 | 1 | 276809.97 | 75281.68 | 128818.72 | 269988.91 | 1 | 1 | 193581.31 |
| P56394 | Cox17 | Cytochrome c oxidase copper chaperone Mus musculus | 73.75 | 25.4 | 6784 | 1 | 1 | 853887.655 | 675136.25 | 976623.56 | 849466.06 | 914324.75 | 1243552.5 | 1305873.25 | 704204 | 1423785.25 | 1726207.12 | 1119831.88 | 1323507.62 | 1517114 | 1120257.5 |
| Q9D7J4 | Cox20 | Cytochrome c oxidase assembly protein COX20, mitochondrial Mus musculus | 54.77 | 19.66 | 13163 | 1 | 1 | 399966.47 | 385341.97 | 316094.22 | 304034.22 | 344977.81 | 217116.62 | 387090.5 | 206536.62 | 201388.03 | 197661.84 | 419860.03 | 273182.28 | 287732.34 | 259381.95 |
| P19783 | Cox4i1 | Cytochrome c oxidase subunit 4 isoform 1, mitochondrial Mus musculus | 215.7 | 30.77 | 19530 | 4 | 4 | 30285453 | 33025669 | 27630631 | 26359977 | 29911717 | 28602201 | 27487479 | 25874075 | 25248589 | 29015169 | 25853013 | 22462841 | 25196947 | 23308119 |
| P11240 | Cox5a | Cytochrome c oxidase subunit 5A, mitochondrial Rattus norvegicus | 288.69 | 38.36 | 16130 | 6 | 6 | 40878529 | 47620297 | 40817437 | 37513889 | 38771337 | 32348581 | 35937237 | 40773493 | 33111355 | 28772209 | 31308093 | 29085567 | 31354793 | 34658793 |
| P19536 | Cox5b | Cytochrome c oxidase subunit 5B, mitochondrial Mus musculus | 53.61 | 9.38 | 13813 | 1 | 1 | 7728957.5 | 8080394.5 | 9121866 | 9044157 | 8073188.5 | 5919199.5 | 7450418 | 6304691 | 6546995 | 8021108 | 5163765.5 | 6167710.5 | 4713191.5 | 5723266.5 |
| P56391 | Cox6b1 | Cytochrome c oxidase subunit 6B1 Mus musculus | 43.9 | 16.28 | 10071 | 1 | 1 | 2872200.25 | 4056435 | 2684032.25 | 2218423 | 3453565.75 | 2030394.88 | 2620808.75 | 2981121.25 | 2234880.25 | 2236088.25 | 1640230.5 | 1700160.71 | 1498464.25 | 1961787.38 |
| Q9CPQ1 | Cox6c | Cytochrome c oxidase subunit 6C Mus musculus | 67.57 | 19.74 | 8469 | 2 | 2 | 5674408 | 3656424 | 3796538 | 4099267.5 | 3276160.75 | 3148131.5 | 3580100.25 | 3345972.25 | 2841076.75 | 3019356.25 | 2281812.5 | 2360568.5 | 2293236.25 | 2253690 |
| P48771 | Cox7a2 | Cytochrome c oxidase subunit 7A2, mitochondrial Mus musculus | 55.83 | 12.05 | 9291 | 1 | 1 | 2024143.12 | 2778264.75 | 2558337.5 | 2066369.38 | 1934297.38 | 1397679 | 2069152 | 2100055.5 | 1573706.88 | 1520218.75 | 325632.38 | 695491.81 | 1112682 | 711268.73 |
| O89001 | Cpd | Carboxypeptidase D Mus musculus | 285.86 | 6.25 | 152406 | 7 | 7 | 5301483 | 4425193.5 | 3982986.5 | 4140849 | 5125316.5 | 4098473.5 | 4482425 | 3978502.5 | 5490180 | 5267524.5 | 4545021 | 4611463 | 5650253 | 5066801.5 |
| Q8C166 | Cpne1 | Copine-1 Mus musculus | 608.36 | 28.36 | 58887 | 10 | 11 | 17995381 | 17383457 | 16692100 | 21040703 | 19500999 | 29652387 | 22701409 | 25226597 | 24229393 | 25495135 | 19475509 | 18864543 | 18260365 | 18881301 |
| Q8BT60 | Cpne3 | Copine-3 Mus musculus | 333.64 | 19.14 | 59585 | 5 | 7 | 12764879 | 9280576 | 9306238 | 10076015 | 10355059 | 9913400 | 8964607 | 12160198 | 10238117 | 10494270 | 10052956 | 10033543 | 9168273 | 8713955 |
| Q9DC53 | Cpne8 | Copine-8 Mus musculus | 166.02 | 6.93 | 64667 | 2 | 3 | 1 | 1 | 208111.88 | 619525.88 | 1 | 906988.75 | 1 | 217960.78 | 738262 | 1 | 1 | 1 | 633890 | 1 |
| P36552 | Cpox | Oxygen-dependent coproporphyrinogen-III oxidase, mitochondrial Mus musculus | 149.92 | 7.22 | 49715 | 3 | 3 | 1503168 | 1347360.5 | 1329728.25 | 1128550.25 | 1129803.62 | 1787710.75 | 1646714.88 | 1371607.25 | 3656753 | 3044954.5 | 1432165.5 | 3589638.5 | 1740911.75 | 3050852.5 |
| Q9WVJ3 | Cpq | Carboxypeptidase Q Mus musculus | 159.27 | 9.79 | 51813 | 3 | 3 | 4235704.5 | 5433789.5 | 3963128 | 4633763.5 | 4864550 | 2740996.75 | 3432032.25 | 3105766.25 | 3189807.75 | 2950120 | 3112566.25 | 3491155.25 | 2712508.75 | 3580672 |
| Q9EPU4 | Cpsf1 | Cleavage and polyadenylation specificity factor subunit 1 Mus musculus | 280.91 | 4.86 | 160818 | 6 | 6 | 3494121.75 | 2602098.25 | 2750698.5 | 3393683.5 | 3468244.5 | 3329892 | 3086502 | 2315742.5 | 2401737 | 2699688.5 | 3700851.75 | 4226635.5 | 2916957.75 | 3041508.25 |
| O35218 | Cpsf2 | Cleavage and polyadenylation specificity factor subunit 2 Mus musculus | 347.87 | 11.76 | 88383 | 6 | 6 | 2244283.75 | 3929254 | 3872361.25 | 2699875 | 3522009 | 2304963.25 | 2595124.5 | 3252766 | 3146407 | 3261641.5 | 3442908.75 | 3372981.75 | 3191451.25 | 3177276 |
| Q9QXK7 | Cpsf3 | Cleavage and polyadenylation specificity factor subunit 3 Mus musculus | 176.93 | 8.48 | 77505 | 4 | 4 | 2630602.25 | 2300776.5 | 1728431 | 2361033.5 | 2241506 | 2806665.5 | 2839439 | 3075626 | 2360146.75 | 3194808.5 | 2726656.5 | 3271560.25 | 2636069.5 | 1829672.5 |
| Q6NVF9 | Cpsf6 | Cleavage and polyadenylation specificity factor subunit 6 Mus musculus | 453.08 | 17.06 | 59153 | 7 | 7 | 17312125 | 18308605 | 15433369 | 15300309 | 17152403 | 13593899 | 14460246 | 17829391 | 15108296 | 16804881 | 13069781 | 12068177 | 14335607 | 15930365 |
| Q5XI29 | Cpsf7 | Cleavage and polyadenylation specificity factor subunit 7 Rattus norvegicus | 210.07 | 13.42 | 51073 | 5 | 5 | 3155922 | 3872699 | 3390567.5 | 3668620.5 | 3314346.5 | 3158044 | 3179262.5 | 3882158.5 | 2417373.25 | 2974243.5 | 2800432.5 | 2946060.5 | 2737779 | 2988918 |
| P32198 | Cpt1a | Carnitine O-palmitoyltransferase 1, liver isoform Rattus norvegicus | 39.87 | 2.2 | 88125 | 1 | 1 | 220662.52 | 263728.84 | 292934.12 | 1 | 1 | 279076.66 | 260102.25 | 315610.75 | 355859.47 | 322773.5 | 261388.48 | 254520.31 | 303295.5 | 279242.25 |
| P52825 | Cpt2 | Carnitine O-palmitoyltransferase 2, mitochondrial Mus musculus | 194.82 | 10.49 | 73981 | 5 | 5 | 9623966 | 7989849 | 7172153.5 | 8663070 | 8746708 | 8150988.5 | 8309098.5 | 5689783.5 | 7669948 | 7595161.5 | 8519634 | 8097593 | 5478264.5 | 7804111 |
| P15337 | Creb1 | Cyclic AMP-responsive element-binding protein 1 Rattus norvegicus | 88.21 | 10.56 | 36633 | 2 | 2 | 1094556.12 | 1386957.88 | 1462360.62 | 1320651.38 | 1367049.38 | 1046445.56 | 1245007.75 | 1239712.12 | 1442803.38 | 1525690.62 | 1253843.25 | 1369782.88 | 1338368.12 | 1466607.75 |
| O88668 | Creg1 | Protein CREG1 Mus musculus | 112.73 | 13.18 | 24452 | 2 | 2 | 5879620 | 4904086 | 3288679.75 | 5032750 | 5737065 | 6149892.5 | 6169681 | 5546140 | 6243145 | 5166010 | 5134289 | 5000084.5 | 5574432 | 3585188.25 |
| Q4V7F2 | Creld1 | Cysteine-rich with EGF-like domain protein 1 Rattus norvegicus | 85.57 | 7.38 | 45697 | 2 | 2 | 1567646.38 | 1949247.25 | 1451439.5 | 1601468.5 | 2347912.5 | 1718869.25 | 1505372.75 | 2022989.25 | 1700548.5 | 1625508.12 | 914258.75 | 811783 | 1681290.25 | 1673049.25 |
| Q9CYA0 | Creld2 | Cysteine-rich with EGF-like domain protein 2 Mus musculus | 139.69 | 7.43 | 38220 | 2 | 2 | 1209143.38 | 1328185.25 | 1600187 | 1328698.88 | 1718282 | 354444.78 | 283145.53 | 1139288.12 | 1290724.12 | 1451044.75 | 1488502.62 | 2105022.75 | 1795252 | 1754534.38 |
| P63254 | Crip1 | Cysteine-rich protein 1 Mus musculus | 51.61 | 12.99 | 8550 | 1 | 1 | 1382662.62 | 567465.25 | 805599.19 | 1115996.62 | 734611.19 | 679004.56 | 1211261.75 | 742990.75 | 1241579.25 | 1223638 | 1000386.25 | 939140.75 | 724391.75 | 872484.62 |
| Q63768 | Crk | Adapter molecule crk Rattus norvegicus | 200.61 | 14.8 | 33845 | 4 | 4 | 5765189 | 3860508.5 | 3913847.25 | 6410894 | 6449763 | 5381169 | 2499059 | 3879259 | 2596005 | 4328357 | 6899917 | 7275631 | 3984644 | 5105239 |
| Q9Z2L7 | Crlf3 | Cytokine receptor-like factor 3 Mus musculus | 141.76 | 10.86 | 49559 | 3 | 3 | 970408.38 | 1690071.75 | 2738421.25 | 1509731.62 | 1445621.5 | 2103998 | 1098760.88 | 2136706.75 | 2091186.62 | 1988928.12 | 1681226.75 | 657447.38 | 2038196.5 | 1625794.75 |
| P63154 | Crnkl1 | Crooked neck-like protein 1 Mus musculus | 108.38 | 4.49 | 83416 | 3 | 3 | 2482272.75 | 2071717.75 | 2441203.75 | 3190848 | 2019391.75 | 2262778.5 | 2311658 | 2660261.5 | 2230697.5 | 2282258.5 | 2850124.5 | 2682435 | 2462730.25 | 3144775.25 |
| Q99KP3 | Cryl1 | Lambda-crystallin homolog Mus musculus | 305.99 | 25.39 | 35209 | 5 | 5 | 4758469.5 | 4669841 | 5362881.5 | 5612916 | 4733707 | 3760955.75 | 3771054 | 6154298 | 6303065.5 | 3616397 | 3421030.25 | 6694852 | 5218574 | 4188295.5 |
| P47199 | Cryz | Quinone oxidoreductase Mus musculus | 44.42 | 6.65 | 35269 | 1 | 1 | 1067594.25 | 1308112.12 | 1488117.25 | 1360542.88 | 1056024.75 | 1830280.38 | 1295732.88 | 1537155.12 | 1916072.12 | 1893861.75 | 1614763.62 | 1761491.75 | 2126802.25 | 1628377.25 |
| Q921W4 | Cryzl1 | Quinone oxidoreductase-like protein 1 Mus musculus | 83.52 | 10.06 | 38725 | 2 | 2 | 1 | 1 | 1485640.38 | 1012541 | 968118.94 | 1540200.25 | 1 | 1727876 | 1649023.25 | 1 | 1 | 1538163.12 | 1978923.25 | 1 |
| Q9CZU6 | Cs | Citrate synthase, mitochondrial Mus musculus | 782.65 | 42.67 | 51737 | 15 | 15 | 86699297 | 81383289 | 77836641 | 70827313 | 76641065 | 78995449 | 75218225 | 74973777 | 73114857 | 80540769 | 78185329 | 77008185 | 86961073 | 79591793 |
| P18395 | Csde1 | Cold shock domain-containing protein E1 Rattus norvegicus | 860.82 | 25.56 | 88895 | 18 | 18 | 26972421 | 24937707 | 24642163 | 25598829 | 21724057 | 24472405 | 23410607 | 23000067 | 21485895 | 22380707 | 23168107 | 25991005 | 23613615 | 23712455 |
| Q9ERK4 | Cse1l | Exportin-2 Mus musculus | 1738.81 | 37.59 | 110455 | 28 | 28 | 55513865 | 41289213 | 38080361 | 32701209 | 33126635 | 40139693 | 41553521 | 51818473 | 38220797 | 31744985 | 39855417 | 37398713 | 39203697 | 38551817 |
| P09581 | Csf1r | Macrophage colony-stimulating factor 1 receptor Mus musculus | 356.57 | 12.28 | 109179 | 7 | 8 | 10909409 | 11710867 | 10370802 | 7828121 | 10516187 | 9632718 | 9419007 | 10975851 | 11943086 | 11990509 | 10728645 | 10568285 | 12142606 | 11938429 |
| P32577 | Csk | Tyrosine-protein kinase CSK Rattus norvegicus | 339.91 | 15.78 | 50746 | 6 | 6 | 9015732 | 8588454 | 9982495 | 8774781 | 9298991 | 10080958 | 10298362 | 8660045 | 10431245 | 10223942 | 9930851 | 10192406 | 9192737 | 10542107 |
| P48729 | CSNK1A1 | Casein kinase I isoform alpha Homo sapiens | 86.86 | 9.2 | 38915 | 2 | 2 | 536957.25 | 1 | 1 | 1 | 1 | 229944.61 | 1 | 1 | 298371.41 | 212198.98 | 53802.4 | 1 | 1 | 1 |
| Q06486 | Csnk1d | Casein kinase I isoform delta Rattus norvegicus | 162.38 | 9.4 | 47316 | 3 | 3 | 408847.41 | 930260.62 | 637895.75 | 582409.04 | 352632.38 | 852648.75 | 850107.5 | 781898.75 | 905007.5 | 483633.62 | 560487.38 | 347513.22 | 910465.88 | 802639.38 |
| P19139 | Csnk2a1 | Casein kinase II subunit alpha Rattus norvegicus | 464.53 | 30.18 | 45073 | 9 | 9 | 18715861 | 14206223 | 12980470 | 13937643 | 13187150 | 14913258 | 15971410 | 16357001 | 16124517 | 16825893 | 12651507 | 14659136 | 16907691 | 16363610 |
| O54833 | Csnk2a2 | Casein kinase II subunit alpha' Mus musculus | 259.18 | 19.71 | 41215 | 5 | 5 | 4948351 | 3063600.5 | 4773701.5 | 3121328 | 3421565.5 | 4317929 | 2703138.25 | 3672243.5 | 4014049.75 | 3983584.75 | 3019305 | 3059108.75 | 3573774.5 | 4001538.75 |
| P67871 | Csnk2b | Casein kinase II subunit beta Mus musculus | 83.12 | 15.81 | 24942 | 2 | 2 | 3663886.5 | 4403108 | 3072081.25 | 3583049.5 | 4103227 | 3643471.5 | 3420486.25 | 4278240.5 | 3433358.75 | 4119229.75 | 3974655 | 3155274 | 4062436.5 | 4516483.5 |
| Q99388 | Csprs | Component of Sp100-rs Mus musculus | 91.15 | 8.65 | 24061 | 1 | 2 | 2141751.5 | 2036986.12 | 2158798.5 | 1540366.88 | 1584585.62 | 1324223.75 | 1275136.5 | 1619174.88 | 1521761.75 | 1286175.25 | 1327284.38 | 1794179.88 | 1312826 | 1244349 |
| P97315 | Csrp1 | Cysteine and glycine-rich protein 1 Mus musculus | 258.97 | 33.16 | 20583 | 5 | 5 | 1776077.75 | 1699707.62 | 1975467.25 | 2826553.5 | 1726693.5 | 4060239.5 | 2485635.25 | 4567123.5 | 4943919.5 | 2277767.25 | 4142359 | 4486704 | 2507449.25 | 1209048.62 |
| P14841 | Cst3 | Cystatin-C Rattus norvegicus | 37.83 | 7.86 | 15437 | 1 | 1 | 1246088.62 | 894169.31 | 927652.62 | 1088590.5 | 1185652.5 | 1085111.75 | 1092025.12 | 1197694.62 | 726513.88 | 1068976 | 1202227 | 1439122.5 | 942972.12 | 957153.56 |
| Q62426 | Cstb | Cystatin-B Mus musculus | 145.46 | 57.14 | 11046 | 3 | 3 | 7521179 | 7137837.5 | 6904083 | 12055766 | 6465746.5 | 3031240.75 | 5190731 | 2957673 | 4015037.5 | 4353343.5 | 4867321 | 3217625.75 | 5260377 | 2877489 |
| Q5BJQ6 | Cstf1 | Cleavage stimulation factor subunit 1 Rattus norvegicus | 93.52 | 3.71 | 48382 | 1 | 1 | 928786.5625 | 930250.25 | 920181.12 | 1003143.5 | 861571.38 | 1022831 | 1431887.88 | 1066624 | 1073831.5 | 924489.69 | 829787.62 | 998945.69 | 884344.44 | 905757.38 |
| Q8BIQ5 | Cstf2 | Cleavage stimulation factor subunit 2 Mus musculus | 167.32 | 14.31 | 61341 | 4 | 4 | 1416987.5 | 738162.56 | 917080 | 2133209 | 1123908.62 | 1254048.62 | 1046342.75 | 1069448.88 | 1722729.75 | 1020201.94 | 1694630.62 | 1959697.38 | 2240427.75 | 1955137.25 |

| | | | | | | | | | | | | | | | | | | | | | |
|---|---|---|---|---|---|---|---|---|---|---|---|---|---|---|---|---|---|---|---|---|---|
| Q99LI7 | Cstf3 | Cleavage stimulation factor subunit 3 Mus musculus | 372.6 | 13.81 | 82877 | 7 | 7 | 2502103 | 2949152.25 | 3310061.75 | 3082721.25 | 2948490 | 3288276 | 3569139.5 | 3212734 | 3500076.5 | 3676061 | 3380831.75 | 3028354.5 | 3154689.5 | 2977506 |
| Q8R311 | Ctage5 | Endoplasmic reticulum export factor CTAGE5 Mus musculus | 84.57 | 3.47 | 87718 | 2 | 2 | 606834.88 | 705294.5 | 929798.62 | 774351.5 | 855478 | 723228.81 | 714465.12 | 644363.12 | 765426.75 | 772696.19 | 892348.75 | 828276.31 | 830647 | 781088.62 |
| O88712 | Ctbp1 | C-terminal-binding protein 1 Mus musculus | 357.62 | 21.54 | 47745 | 4 | 7 | 6084126.5 | 6867470.5 | 7400646 | 7924728 | 7334426.5 | 9430714 | 7924752 | 7319881 | 9867541 | 9616093 | 8397520 | 7571956 | 8847623 | 7905200.5 |
| P56546 | Ctbp2 | C-terminal-binding protein 2 Mus musculus | 495.05 | 36.63 | 48957 | 9 | 12 | 14222561 | 12440207 | 12064810 | 16314195 | 15451092 | 13990514 | 15174011 | 12710962 | 13620627 | 17159451 | 14601281 | 15532437 | 15076583 | 15152072 |
| Q8R242 | Ctbs | Di-N-acetylchitobiase Mus musculus | 84.47 | 8.2 | 41295 | 2 | 2 | 2802038 | 3222664.25 | 2995471.75 | 958706.62 | 2494720.155 | 2037735.62 | 812468.38 | 969283.44 | 2104498.5 | 2174028.5 | 981597.19 | 2173146 | 1231777 | 2567925 |
| Q5SUQ9 | Ctc1 | CST complex subunit CTC1 Mus musculus | 89.28 | 3.3 | 134028 | 2 | 2 | 1 | 386601.38 | 1 | 315971.44 | 479609.97 | 521541.315 | 332316.88 | 455204.88 | 959720.5 | 338923 | 317311.75 | 417675.34 | 1005266 | 425932.66 |
| Q61164 | Ctcf | Transcriptional repressor CTCF Mus musculus | 123.49 | 3.67 | 83745 | 2 | 2 | 589231.62 | 706246.56 | 1244668.5 | 667464.88 | 1033430.69 | 833635.56 | 1081637.25 | 871246.62 | 1111791.5 | 976852.62 | 975429.81 | 1024833.5 | 1157134.88 | 770180.88 |
| Q8VCN5 | Cth | Cystathionine gamma-lyase Mus musculus | 688.3 | 43.47 | 43567 | 13 | 13 | 41636849 | 40732541 | 49172553 | 51631009 | 46581893 | 48540305 | 42209745 | 44339353 | 42522381 | 43343765 | 35893937 | 36933165 | 36345461 | 37593289 |
| Q9CWL8 | Ctnnbl1 | Beta-catenin-like protein 1 Mus musculus | 289.25 | 13.32 | 64980 | 6 | 6 | 1876150.25 | 2011940.12 | 2942662.5 | 2278631.5 | 3450300 | 2698397.75 | 3338329 | 2708399.25 | 2819634 | 2610201.25 | 3023481.75 | 2215927 | 1924920.25 | 2988842.25 |
| P30999 | Ctnnd1 | Catenin delta-1 Mus musculus | 503.48 | 16.63 | 104925 | 10 | 10 | 3632788.5 | 5114385.5 | 6365331 | 7988858 | 5750182 | 4497701.5 | 5127876.5 | 4029252 | 5560320 | 5101681 | 5056257 | 4413752 | 5573508 | 5348033.5 |
| P70698 | Ctps1 | CTP synthase 1 Mus musculus | 717.68 | 27.41 | 66682 | 11 | 12 | 11293901 | 12770289 | 11429613 | 12607571 | 10775793 | 11147971 | 11544509 | 10293005 | 12308024 | 10984212 | 11113973 | 10472703 | 10176380 | 10660851 |
| P70303 | Ctps2 | CTP synthase 2 Mus musculus | 80.46 | 3.75 | 65514 | 1 | 2 | 881614.19 | 1296648.12 | 1375734.75 | 1357951.38 | 534595.62 | 1445160.88 | 1184633.38 | 1331406.75 | 1479598.75 | 1108698 | 1127967.25 | 1273447.25 | 1196532.25 | 1484463.62 |
| P16675 | Ctsa | Lysosomal protective protein Mus musculus | 1103.64 | 36.71 | 53844 | 14 | 16 | 144273681 | 156101457 | 125186585 | 127415905 | 147836225 | 141878577 | 144054321 | 166781105 | 147314721 | 150106097 | 136156417 | 120880953 | 139485921 | 142960817 |
| P10605 | Ctsb | Cathepsin B Mus musculus | 1317.47 | 67.26 | 37280 | 19 | 20 | 334932353 | 395148609 | 352905249 | 324442689 | 350109057 | 326914977 | 301984129 | 381076705 | 339120993 | 305167521 | 315144577 | 308200609 | 354121889 | 334587905 |
| P97821 | Ctsc | Dipeptidyl peptidase 1 Mus musculus | 595.33 | 16.67 | 52376 | 7 | 7 | 49376273 | 52577253 | 45175817 | 45652129 | 52748553 | 36184393 | 32687601 | 37102473 | 32423235 | 31497689 | 37962265 | 34302401 | 38151173 | 39475241 |
| P18242 | Ctsd | Cathepsin D Mus musculus | 1308.71 | 65.37 | 44954 | 18 | 24 | 406401761 | 489922369 | 425776257 | 393210113 | 412787073 | 566899457 | 570130689 | 552964993 | 524020897 | 533450113 | 536090305 | 506827169 | 585553345 | 525133889 |
| P49935 | Ctsh | Pro-cathepsin H Mus musculus | 121.9 | 8.41 | 37170 | 2 | 2 | 1658226.88 | 4813508.5 | 3965152 | 3522042 | 4001618.5 | 2795950 | 2711203 | 2895378 | 3281238 | 2610760.25 | 3254221.5 | 2790502 | 3319752.25 | 3099382.5 |
| P06797 | Ctsl | Cathepsin L1 Mus musculus | 294.45 | 37.72 | 37547 | 6 | 6 | 13943462 | 17121151 | 17095199 | 10881181 | 14913645 | 15854645 | 15483579 | 20445239 | 17907291 | 16089471 | 14843341 | 14395614 | 17152487 | 17337579 |
| O70370 | Ctss | Cathepsin S Mus musculus | 809.73 | 49.12 | 38475 | 9 | 10 | 238743409 | 309175649 | 274737185 | 216478641 | 242877905 | 188496833 | 181920641 | 208634577 | 183625857 | 167515569 | 163645185 | 144368785 | 225630465 | 196082705 |
| Q9WUU7 | Ctsz | Cathepsin Z Mus musculus | 745.53 | 45.75 | 33996 | 12 | 12 | 342145953 | 436895265 | 362891745 | 375083137 | 406212577 | 388462337 | 355202177 | 417323169 | 363276737 | 378226433 | 376606049 | 350311809 | 394775425 | 404304769 |
| Q99LJ0 | Cttnbp2nl | CTTNBP2 N-terminal-like protein Mus musculus | 157.93 | 6.58 | 69841 | 2 | 2 | 370077.53 | 532828.12 | 236185.23 | 429336.66 | 462252.47 | 628559.12 | 465412.91 | 631763.25 | 443450.53 | 416470.22 | 405681.34 | 506016.53 | 610697.69 | 312364.44 |
| Q3U308 | Ctu2 | Cytoplasmic tRNA 2-thiolation protein 2 Mus musculus | 54.55 | 3.7 | 56105 | 1 | 1 | 1 | 1 | 1 | 1 | 270970.25 | 1 | 1 | 1 | 305560.69 | 267004.19 | 336719.03 | 361082.0833 | 327399.53 | 419127.69 |
| Q9WTX6 | Cul1 | Cullin-1 Mus musculus | 508.56 | 14.69 | 89692 | 10 | 10 | 9356674 | 8197263 | 7475768 | 8358356 | 10032447 | 11403163 | 8733838 | 12266343 | 11508169 | 10748202 | 10499824 | 12952264 | 12832699 | 11439313 |
| Q9D4H8 | Cul2 | Cullin-2 Mus musculus | 222.18 | 7.92 | 86877 | 5 | 5 | 1236850.5 | 1862539.88 | 1368809.25 | 1538803.75 | 1462571.5 | 655457.19 | 2131048.75 | 3279417 | 2955035.25 | 1284577.75 | 2026049.5 | 1597622.25 | 1895738.12 | 2344790 |
| B5DF89 | Cul3 | Cullin-3 Rattus norvegicus | 415.77 | 13.41 | 88932 | 7 | 7 | 4959384.5 | 5196358 | 5075169.5 | 4650876 | 4922087 | 6820248.5 | 4787354.5 | 6839920 | 4197612.5 | 4737988 | 6324329.5 | 4260057.5 | 5881802.5 | 3128280.25 |
| Q3TCH7 | Cul4a | Cullin-4A Mus musculus | 193.37 | 5.67 | 87753 | 2 | 4 | 2419670.5 | 2496802.75 | 2800189 | 2487509 | 2312425.5 | 2703769.25 | 2230956 | 2802537 | 3169368.5 | 2526152 | 2355697.5 | 2022061.25 | 2731464.75 | 2459069 |
| A2A432 | Cul4b | Cullin-4B Mus musculus | 489.3 | 12.06 | 110699 | 7 | 9 | 5683032 | 3562498 | 5907558 | 3791890.75 | 3234073 | 4515149 | 2807788.75 | 6049258 | 4567405.5 | 4793523 | 3843725.5 | 5596016 | 4957681.5 | 4323771 |
| Q9D5V5 | Cul5 | Cullin-5 Mus musculus | 37.15 | 1.28 | 90974 | 1 | 1 | 257088.89 | 1 | 1 | 1 | 455063.47 | 398494.12 | 201253.92 | 1 | 1 | 370552.66 | 380663.53 | 400298 | 437463.41 | 324381.44 |
| Q6MGD0 | Cuta | Protein CutA Rattus norvegicus | 181.04 | 29.94 | 18659 | 4 | 4 | 7819751 | 10969266 | 9721237 | 9869074 | 8752632 | 8600881 | 8412533 | 9060140 | 8975825 | 8136363 | 9267912 | 8619711 | 8849813 | 9333743 |
| Q5BJP2 | Cwc15 | Spliceosome-associated protein CWC15 homolog Rattus norvegicus | 173.56 | 16.16 | 26638 | 3 | 3 | 585857.06 | 3540086 | 2168444.25 | 2685335 | 3972490.25 | 3627474.5 | 3648875.25 | 1884542.88 | 3279349 | 4019204.5 | 2631039.75 | 1858291.25 | 3230294 | 2608121 |
| Q8C5N3 | Cwc22 | Pre-mRNA-splicing factor CWC22 homolog Mus musculus | 319.04 | 9.69 | 104773 | 6 | 6 | 3895319.5 | 4046651.25 | 1822862 | 3242381 | 2941762.5 | 3183897.25 | 3078697.5 | 3642518.5 | 1817117.88 | 4680753.5 | 2485200 | 2662471.75 | 1419262.25 | 1693499.5 |
| Q8CI33 | Cwf19l1 | CWF19-like protein 1 Mus musculus | 37.67 | 2.23 | 60193 | 1 | 1 | 1 | 1 | 1 | 1 | 1 | 1 | 1 | 1 | 1 | 1 | 1 | 1 | 728816.19 | 1 |
| Q9CQX2 | Cyb5b | Cytochrome b5 type B Mus musculus | 328.72 | 58.22 | 16318 | 5 | 5 | 13836063 | 8341799.5 | 10466041 | 13242618 | 11826971 | 9121908 | 8341185.5 | 12099917 | 7257061.5 | 9711717 | 11700505 | 9697557 | 8846196 | 10158302 |
| Q9DB73 | Cyb5r1 | NADH-cytochrome b5 reductase 1 Mus musculus | 274.09 | 25.57 | 34135 | 6 | 6 | 3272358.25 | 3996759 | 3589742.5 | 2439730 | 2806576.75 | 4032830.75 | 5249223 | 3971796.5 | 3135731 | 4786758 | 3529941 | 3057868 | 3781504 | 3490929.75 |
| Q9DCN2 | Cyb5r3 | NADH-cytochrome b5 reductase 3 Mus musculus | 705.95 | 45.18 | 34128 | 11 | 11 | 30700057 | 30305555 | 27908413 | 27498421 | 27467849 | 27060713 | 27824485 | 28032559 | 25440971 | 30115743 | 26765125 | 30250171 | 27780071 | 26819803 |
| Q61462 | Cyba | Cytochrome b-245 light chain Mus musculus | 136.89 | 7.29 | 20748 | 3 | 3 | 10207428 | 9629593 | 6990939 | 6998513.5 | 9168180 | 8418469 | 6150572 | 7804055.5 | 7331240 | 6547116.5 | 7727846.5 | 7081598 | 8961659 | 9184594 |
| Q61093 | Cybb | Cytochrome b-245 heavy chain Mus musculus | 666.11 | 30 | 65305 | 12 | 12 | 17238975 | 9279005 | 8931859 | 7187517 | 11210569 | 10307075 | 7359234 | 11581564 | 7893411 | 8603878 | 9676965 | 7794356 | 12061344 | 9860013 |
| Q3TYS2 | Cybc1 | Cytochrome b-245 chaperone 1 Mus musculus | 113.78 | 14.97 | 20921 | 3 | 3 | 4608574 | 2390832 | 2792405 | 1635330.25 | 2396028.5 | 2834147.75 | 2264166.75 | 2237399 | 2724396.5 | 2216179.25 | 1913315.75 | 2577006 | 2740913.5 | 3142287.5 |
| Q9D0M3 | Cyc1 | Cytochrome c1, heme protein, mitochondrial Mus musculus | 406.16 | 52.62 | 35328 | 9 | 9 | 15720952 | 12139899 | 12626407 | 10599710 | 13279112 | 12918118 | 14413132 | 17864799 | 16719168 | 13398738 | 14932583 | 14426790 | 16593078 | 17330887 |
| P62897 | Cycs | Cytochrome c, somatic Mus musculus | 296.2 | 55.24 | 11605 | 6 | 6 | 24296731 | 23385439 | 24839441 | 24829385 | 23277525 | 18623457 | 22805401 | 21979819 | 21256253 | 26364561 | 21420971 | 22683533 | 27067801 | 22376863 |
| Q7TMB8 | Cyfip1 | Cytoplasmic FMR1-interacting protein 1 Mus musculus | 1945.15 | 33.2 | 145241 | 22 | 37 | 53762173 | 46518805 | 46904561 | 42789213 | 44979477 | 56949453 | 54642505 | 52869509 | 62710625 | 54195625 | 51165733 | 54168297 | 52025097 | 53359213 |
| Q5SQX6 | Cyfip2 | Cytoplasmic FMR1-interacting protein 2 Mus musculus | 846.05 | 16.04 | 145659 | 3 | 17 | 14203615 | 13260923 | 13543792 | 11617989 | 9311673 | 18029937 | 14095068 | 16040019 | 20379449 | 15796638 | 13576555 | 18370377 | 14383378 | 13005055 |
| Q8BKE6 | Cyp20a1 | Cytochrome P450 20A1 Mus musculus | 218.71 | 17.32 | 52149 | 5 | 5 | 4086177.5 | 3415466 | 3399480.25 | 3047147 | 2857976.5 | 3550361 | 2755912.25 | 2771629 | 2660382.75 | 3199472.75 | 3331242.25 | 3086707.75 | 3112701.5 | 3493866 |
| Q9DBG1 | Cyp27a1 | Sterol 26-hydroxylase, mitochondrial Mus musculus | 148.06 | 9.19 | 60720 | 3 | 3 | 1309922.5 | 1143625.12 | 882316.56 | 749493 | 1021339.295 | 975128.12 | 741316.25 | 1019511.5 | 641959.75 | 1079438 | 1340672 | 1237169.5 | 1136221 | 1193099.75 |
| Q64654 | Cyp51a1 | Lanosterol 14-alpha demethylase Rattus norvegicus | 316.36 | 19.09 | 56707 | 7 | 7 | 3869052.75 | 4539243.5 | 4142854 | 2842192 | 3333263.5 | 2977100.5 | 3007905.5 | 2877830.25 | 3007581 | 2464328 | 2757151.5 | 2747478.5 | 3849183.75 | 3096148.5 |
| Q80YW0 | Cyth4 | Cytohesin-4 Mus musculus | 232.17 | 12.21 | 45285 | 4 | 4 | 5024233 | 4180476 | 3568252.5 | 3380212.5 | 4476086 | 4044814.25 | 2516578.5 | 4696851.5 | 3999067.75 | 2291064.25 | 3013928.75 | 3338131.25 | 3605225.5 | 2529773 |
| P98078 | Dab2 | Disabled homolog 2 Mus musculus | 1579.37 | 45.04 | 82312 | 9 | 22 | 33908749 | 31361015 | 43650805 | 33180885 | 30821723 | 32450225 | 30363561 | 28733057 | 35910361 | 30762065 | 31344041 | 38973149 | 34234933 | 32137191 |
| P61804 | Dad1 | Dolichyl-diphosphooligosaccharide--protein glycosyltransferase subunit DAD1 Mus muscul | 166.75 | 28.32 | 12497 | 3 | 3 | 13215215 | 13330451 | 11556073 | 10955833 | 15067723 | 14099778 | 11961920 | 10981207 | 12339314 | 13018405 | 15633830 | 15490589 | 15116859 | 17593443 |
| Q9ER88 | Dap3 | 28S ribosomal protein S29, mitochondrial Mus musculus | 264.87 | 20.46 | 44699 | 5 | 5 | 3156793 | 4620741.5 | 4832820.5 | 2470616 | 4198174.5 | 3692942 | 3831015 | 4202905.5 | 4558979.5 | 3427475.75 | 4069690.5 | 3818907.5 | 3213954.5 | 3633658 |
| Q922B2 | Dars | Aspartate--tRNA ligase, cytoplasmic Mus musculus | 1050.3 | 42.51 | 57147 | 19 | 19 | 66303153 | 75627097 | 70327633 | 64611905 | 65696493 | 64689109 | 69870225 | 74759641 | 66139085 | 66909153 | 62921357 | 67162865 | 64593905 | 75283921 |
| Q8BIP0 | Dars2 | Aspartate--tRNA ligase, mitochondrial Mus musculus | 111.11 | 4.59 | 74102 | 3 | 3 | 2521803.75 | 4048034 | 3894850.25 | 2972283.5 | 4086272.75 | 4269687.5 | 3834908 | 4029764.5 | 4306810 | 3605008.25 | 2787916 | 3600336 | 4151163 | 4040467.5 |
| Q9JII5 | Dazap1 | DAZ-associated protein 1 Mus musculus | 250.56 | 15.27 | 43214 | 4 | 4 | 3752942 | 5246539 | 3905766 | 5174724 | 5566712 | 5490872.5 | 5541856 | 4862856.5 | 5315936 | 5817361 | 5707034 | 5535279 | 5578547 | 5631757 |
| P11030 | Dbi | Acyl-CoA-binding protein Rattus norvegicus | 144.97 | 39.08 | 10027 | 2 | 2 | 8975738 | 7477482 | 6957561 | 8622616 | 7808447.5 | 4107992.25 | 6293613.5 | 7137555 | 4203308 | 7113129 | 7026639.5 | 5648527 | 5563802 | 6502662 |
| Q62418 | Dbnl | Drebrin-like protein Mus musculus | 587.36 | 24.08 | 48700 | 7 | 7 | 8094816.5 | 7775421.5 | 11942636 | 11969308 | 11536735 | 9972487 | 8908499 | 7185464 | 9306166 | 10102071 | 9836754 | 11894790 | 9111891 | 10836874 |
| Q923B1 | Dbr1 | Lariat debranching enzyme Mus musculus | 53.22 | 1.82 | 62289 | 1 | 1 | 1 | 244158.84 | 1 | 1 | 196371.23 | 219173.28 | 264881.75 | 233189.6525 | 196955.94 | 251747.64 | 248104.98 | 156770.67 | 180508.58 | 170506.44 |
| P53395 | Dbt | Lipoamide acyltransferase component of branched-chain alpha-keto acid dehydrogenase c | 201.85 | 10.79 | 53247 | 4 | 4 | 934585.25 | 1303108.5 | 1277103.75 | 1273488.12 | 1362557.25 | 1285597.75 | 1457624.5 | 1274119.5 | 2093373.5 | 1721101.5 | 1531920.25 | 941396.38 | 1709686.25 | 1816122.25 |
| Q80TR8 | Dcaf1 | DDB1- and CUL4-associated factor 1 Mus musculus | 36.68 | 0.93 | 168931 | 1 | 1 | 116573.26 | 298595.81 | 200191.31 | 193877.98 | 140019.59 | 227753.73 | 288159.19 | 280744.69 | 161589.28 | 175565.55 | 202920.31 | 318212.28 | 209042.77 | 172140.2 |
| Q5U2M6 | Dcaf8 | DDB1- and CUL4-associated factor 8 Rattus norvegicus | 36.78 | 1.35 | 66156 | 1 | 1 | 1 | 1 | 724706.62 | 1 | 296630.31 | 465376.72 | 380969.75 | 431134.44 | 558835.12 | 459229.19 | 1 | 441614.53 | 512903.66 | 562800.12 |
| Q8BHC4 | Dcakd | Dephospho-CoA kinase domain-containing protein Mus musculus | 283.83 | 28.57 | 26476 | 6 | 6 | 5708384 | 4926203 | 4425020.5 | 4603862.5 | 3977113.5 | 4259485 | 4620305 | 4237366.5 | 4732710 | 4337645 | 3785582.25 | 4884806 | 4867986 | 4262553.5 |
| Q9DAR7 | Dcps | m7GpppX diphosphatase Mus musculus | 651.43 | 49.41 | 38988 | 13 | 13 | 12611214 | 10769439 | 11305334 | 11576916 | 13026258 | 11194877 | 9626292 | 7366642 | 11974249 | 9855454 | 8130524.5 | 11045285 | 10200207 | 11149930 |
| O08788 | Dctn1 | Dynactin subunit 1 Mus musculus | 395.37 | 9.76 | 141676 | 7 | 7 | 4209534 | 4921418 | 3515108.75 | 6853147 | 3822319.25 | 4664068 | 4790328 | 4251574.5 | 5387035.5 | 4138230.5 | 5513127 | 6303265 | 4269476.5 | 4762295 |
| Q99KJ8 | Dctn2 | Dynactin subunit 2 Mus musculus | 634.94 | 33.58 | 44117 | 11 | 11 | 22103775 | 23893517 | 21376959 | 24522623 | 24590845 | 22141649 | 24643257 | 24132473 | 23093965 | 24204007 | 24026359 | 23873981 | 20998345 | 22401809 |
| Q8CBY8 | Dctn4 | Dynactin subunit 4 Mus musculus | 116.78 | 5.78 | 53057 | 2 | 2 | 871438.31 | 877763.31 | 756759.69 | 956049.19 | 691533.44 | 890174.38 | 1047061.06 | 816058.5 | 1078694.38 | 1064026.62 | 873020.62 | 934084.88 | 840727.56 | 762572.31 |
| Q9QZ73 | Dcun1d1 | DCN1-like protein 1 Mus musculus | 250.03 | 22.78 | 30097 | 4 | 4 | 3379584.25 | 2601892.75 | 2882740.75 | 3204482.25 | 2563609 | 2934865.25 | 2848700.25 | 3300154 | 3026443.5 | 2432227.75 | 3502762.25 | 2484852.5 | 3059450 | 2667876.5 |
| Q91X52 | Dcxr | L-xylulose reductase Mus musculus | 65.94 | 7.38 | 25746 | 2 | 2 | 134480689 | 89065553 | 74447881 | 112225169 | 119284305 | 84487033 | 45989785 | 47740653 | 37871061 | 42012133 | 134733681 | 134293105 | 48283509 | 109253649 |
| Q3U1J4 | Ddb1 | DNA damage-binding protein 1 Mus musculus | 2069.98 | 37.02 | 126853 | 5 | 37 | 74617521 | 78232241 | 78385473 | 74483321 | 76977025 | 76718633 | 73967473 | 78798401 | 79462353 | 79675537 | 74283025 | 77669353 | 79617073 | 82100849 |
| A2ADY9 | Ddi2 | Protein DDI1 homolog 2 Mus musculus | 288.58 | 18.8 | 44591 | 6 | 6 | 4650878 | 6068306 | 3721573 | 7267678 | 6312279.5 | 4694112 | 6476619 | 6488743.5 | 6677014.5 | 8252989 | 7391709 | 6876387.5 | 6468297 | 7318237 |
| O54734 | Ddost | Dolichyl-diphosphooligosaccharide--protein glycosyltransferase 48 kDa subunit Mus muscu | 751.18 | 41.5 | 49028 | 13 | 13 | 519012865 | 34175413 | 129967369 | 173564145 | 191119313 | 130328401 | 35557189 | 33369871 | 37143369 | 38918041 | 176693313 | 48209881 | 43006825 | 168711617 |
| Q80WW9 | Ddrgk1 | DDRGK domain-containing protein 1 Mus musculus | 215.13 | 15.24 | 35977 | 3 | 3 | 2937053.5 | 3908093.75 | 3401647.25 | 2747720.25 | 3207205.75 | 3412633.5 | 3199082.25 | 4837944.5 | 3688148 | 1491919.62 | 2227082.25 | 3277854 | 3047188 | 3157043.25 |
| O35215 | Ddt | D-dopachrome decarboxylase Mus musculus | 52.72 | 9.32 | 13077 | 1 | 1 | 1 | 1 | 1 | 1 | 1 | 1 | 706920.44 | 1 | 1 | 1 | 717825.5 | 1 | 871358.56 | 1 |
| Q91VR5 | Ddx1 | ATP-dependent RNA helicase DDX1 Mus musculus | 1054.07 | 34.32 | 82500 | 19 | 19 | 21494465 | 23311527 | 25019387 | 23063183 | 23753523 | 23760623 | 26720961 | 26121527 | 24885585 | 26870247 | 25130829 | 25252313 | 26245641 | 25898643 |
| Q80Y44 | Ddx10 | Probable ATP-dependent RNA helicase DDX10 Mus musculus | 110.36 | 3.2 | 100739 | 2 | 2 | 197709.77 | 182579.72 | 181611.52 | 224758.7 | 201889.14 | 182456.89 | 236065.44 | 188143.97 | 171036.84 | 162428.61 | 179085.66 | 278889.5 | 230950.73 | 167980.45 |
| Q501J6 | Ddx17 | Probable ATP-dependent RNA helicase DDX17 Mus musculus | 957.64 | 31.54 | 72399 | 11 | 18 | 150077089 | 155918529 | 158525361 | 155182561 | 152922897 | 136444433 | 140492241 | 146613361 | 135731617 | 144161297 | 143302945 | 132117089 | 146279233 | 155489377 |
| Q8K363 | Ddx18 | ATP-dependent RNA helicase DDX18 Mus musculus | 1186.96 | 41.67 | 74181 | 20 | 20 | 17321211 | 17827805 | 16888109 | 16197246 | 11728795 | 14264832 | 13708876 | 19072849 | 13061348 | 13890341 | 13531973 | 17123963 | 18250811 | 14587090 |
| Q61655 | Ddx19a | ATP-dependent RNA helicase DDX19A Mus musculus | 416.28 | 23.43 | 53933 | 8 | 8 | 2878807 | 5348518 | 3112134 | 3867302.25 | 2950477 | 2990728 | 4444176.5 | 3057596 | 3378369.5 | 5374622 | 3684467.25 | 4416218 | 4045603.5 | 3665064 |
| Q9JJY4 | Ddx20 | Probable ATP-dependent RNA helicase DDX20 Mus musculus | 286.83 | 10.18 | 91710 | 6 | 6 | 5664927 | 6365350 | 1952999.12 | 6582219.5 | 5223471.5 | 4538543 | 1741949 | 3346132.25 | 2460349.25 | 2774857.25 | 5594324 | 4665036.5 | 4445907.5 | 4678584.5 |
| Q9JIK5 | Ddx21 | Nucleolar RNA helicase 2 Mus musculus | 2053.9 | 54.76 | 93551 | 35 | 36 | 42154801 | 34022433 | 42058417 | 42927625 | 38194141 | 31975175 | 37257153 | 29146309 | 32689745 | 33366561 | 36178761 | 34434737 | 38966069 | 32566989 |
| Q9ESV0 | Ddx24 | ATP-dependent RNA helicase DDX24 Mus musculus | 345.21 | 10.74 | 96429 | 6 | 6 | 1468147.25 | 1989461.12 | 877275.38 | 1869901.25 | 1698180.5 | 1778299.75 | 1279129.38 | 751556.69 | 532291.75 | 1212506.5 | 556266.56 | 951383.25 | 1485208.12 | 1310087.25 |
| Q8VDW0 | Ddx39a | ATP-dependent RNA helicase DDX39A Mus musculus | 1355.7 | 69.32 | 49067 | 12 | 26 | 165816881 | 171790321 | 166252737 | 161726881 | 158953009 | 149920225 | 163697425 | 163853585 | 154511377 | 155633505 | 151715313 | 157699105 | 150613457 | 163284817 |
| Q63413 | Ddx39b | Spliceosome RNA helicase Ddx39b Rattus norvegicus 2 | 1524.45 | 73.83 | 49035 | 12 | 25 | 190927009 | 204810945 | 194309809 | 190996753 | 184235201 | 176000593 | 195061233 | 191997537 | 180648033 | 183251217 | 178748833 | 181634881 | 170056833 | 189674785 |
| Q62167 | Ddx3x | ATP-dependent RNA helicase DDX3X Mus musculus | 1544.52 | 48.79 | 73101 | 8 | 29 | 87973793 | 79139657 | 82385113 | 69691041 | 81013609 | 69881217 | 72466593 | 65692121 | 67788089 | 72041737 | 63908561 | 65681021 | 70222729 | 68686049 |
| Q62095 | Ddx3y | ATP-dependent RNA helicase DDX3Y Mus musculus | 1104.92 | 32.98 | 73428 | 2 | 23 | 77379209 | 69522481 | 72049065 | 60994129 | 71025609 | 60714649 | 64453085 | 58073865 | 58504413 | 62334405 | 53894289 | 56623925 | 60489421 | 59040217 |
| Q91VN6 | Ddx41 | Probable ATP-dependent RNA helicase DDX41 Mus musculus | 42.28 | 3.22 | 69820 | 1 | 1 | 1 | 1 | 962122.94 | 1 | 1 | 1 | 785518.31 | 1129144.5 | 1211864.75 | 1 | 1 | 910662.69 | 1 | 409081.91 |
| Q569Z5 | Ddx46 | Probable ATP-dependent RNA helicase DDX46 Mus musculus | 802.91 | 19.96 | 117448 | 17 | 17 | 19793049 | 18057853 | 20214551 | 16679657 | 19095113 | 17325125 | 17888469 | 19031819 | 19863953 | 17610369 | 17175241 | 16952003 | 17998313 | 18507387 |
| Q9CWX9 | Ddx47 | Probable ATP-dependent RNA helicase DDX47 Mus musculus | 375.31 | 22.2 | 50639 | 8 | 8 | 6133848 | 7564842 | 7843512 | 5960981 | 5154378.5 | 6811186 | 7571907 | 8254358 | 8012453.5 | 7144217 | 7256071.5 | 8148162 | 7949271.5 | 7904846.5 |
| Q4FZF3 | Ddx49 | Probable ATP-dependent RNA helicase DDX49 Mus musculus | 177.44 | 10.62 | 54094 | 3 | 3 | 804309.06 | 680725.81 | 1172160.75 | 642922.56 | 722912.12 | 701699 | 889818 | 609977.88 | 917736.25 | 401755.88 | 841283.06 | 925203.12 | 732417.62 | 772064.5 |
| Q61656 | Ddx5 | Probable ATP-dependent RNA helicase DDX5 Mus musculus | 1563.06 | 58.63 | 69290 | 22 | 29 | 225700161 | 233248081 | 244238129 | 239267329 | 231313041 | 211316033 | 207093025 | 219695521 | 211457249 | 227407761 | 224661905 | 195167313 | 221794673 | 231716417 |
| Q8K301 | Ddx52 | Probable ATP-dependent RNA helicase DDX52 Mus musculus | 150.01 | 8.19 | 67474 | 3 | 3 | 343005.78 | 337891.59 | 532897.12 | 330994.75 | 334590.16 | 313755.03 | 294793.56 | 274959.28 | 426768.81 | 513032.94 | 287861.53 | 306124 | 337008.25 | 274335.53 |
| Q8K4L0 | Ddx54 | ATP-dependent RNA helicase DDX54 Mus musculus | 179.67 | 7.67 | 97748 | 5 | 5 | 1234306.88 | 602764.81 | 1249032 | 793462.38 | 979783.06 | 966031.94 | 659861.44 | 931775.38 | 821991 | 776946.38 | 744982.38 | 524331.81 | 985302 | 916911.81 |

| | | | | | | | | | | | | | | | | | | | | | |
|---|---|---|---|---|---|---|---|---|---|---|---|---|---|---|---|---|---|---|---|---|---|
| Q9D0R4 | Ddx56 | Probable ATP-dependent RNA helicase DDX56 Mus musculus | 178.1 | 9.34 | 61212 | 3 | 3 | 932268.62 | 817630.69 | 536774.25 | 1242635.12 | 1438026.62 | 750218.38 | 568040.88 | 396626.72 | 536878.05 | 432626.22 | 1307926.25 | 1368493.62 | 824386.62 | 1032922.69 |
| Q66HG7 | Ddx59 | Probable ATP-dependent RNA helicase DDX59 Rattus norvegicus | 37.42 | 1.87 | 65066 | 1 | 1 | 175512.03 | 251675.5 | 279745.94 | 249485.97 | 195653.36 | 137653.25 | 288546.06 | 188128.28 | 248767.59 | 198492.16 | 1 | 1 | 198807.66 | 218606.25 |
| P54823 | Ddx6 | Probable ATP-dependent RNA helicase DDX6 Mus musculus | 907.65 | 44.1 | 54192 | 13 | 13 | 17343113 | 18286627 | 16789805 | 17062831 | 17349095 | 17104823 | 18585343 | 17827401 | 16604509 | 16805119 | 17099197 | 17449469 | 17107579 | 18091145 |
| Q9CQ62 | Decr1 | 2,4-dienoyl-CoA reductase, mitochondrial Mus musculus | 266.22 | 26.57 | 36214 | 5 | 5 | 9110959 | 3718888 | 3367780.5 | 9925201 | 10073742 | 8987500 | 6943719.5 | 3245453.5 | 8728591 | 8674892 | 7791277 | 7867054 | 8993294 | 4709861 |
| Q8C2K1 | Def6 | Differentially expressed in FDCP 6 Mus musculus | 45.02 | 1.59 | 73454 | 1 | 1 | 79740.24 | 1 | 40264.1 | 58635.16 | 1 | 1 | 63138.12 | 1 | 1 | 56326.38 | 81149.95 | 68977.3 | 73952.04 | 51829.91 |
| Q6AXS3 | Dek | Protein DEK Rattus norvegicus | 458.83 | 24.07 | 42892 | 9 | 9 | 24901535 | 25517797 | 23843257 | 21857401 | 26834323 | 24293327 | 24289997 | 23478365 | 23421489 | 23730145 | 20190411 | 21965169 | 24424987 | 26203297 |
| Q3U1Y4 | Dennd4b | DENN domain-containing protein 4B Mus musculus | 37.34 | 0.87 | 164740 | 1 | 1 | 364688.84 | 196452.06 | 1 | 1 | 1 | 297499.66 | 223026.06 | 1 | 1 | 1 | 1 | 224068.88 | 1 | 353926.34 |
| A6H8H2 | Dennd4c | DENN domain-containing protein 4C Mus musculus | 427.89 | 6.56 | 211458 | 8 | 8 | 3449451.25 | 4322360.5 | 5971060.5 | 4559859 | 6918938 | 6749721.5 | 6992494 | 5905397 | 6445326 | 5187324.5 | 6769124 | 7026759.5 | 6446023.5 | 6658921.5 |
| Q9CQJ6 | Denr | Density-regulated protein Mus musculus | 212.91 | 19.19 | 22166 | 3 | 3 | 4609877 | 5230410 | 5181511.5 | 4923461 | 4933535 | 4319952 | 4705343 | 4732829 | 4132269 | 4807452.5 | 4638834 | 5872405.5 | 4524220.5 | 4700444.5 |
| Q91YP3 | Dera | Deoxyribose-phosphate aldolase Mus musculus | 437.65 | 32.08 | 34975 | 7 | 7 | 4085484.5 | 3181956 | 3918847.75 | 3109663.5 | 4620571.5 | 4012048 | 4510107 | 3117761 | 2964168.5 | 4976940 | 4316600.5 | 4895686 | 4418909 | 3832614 |
| O08560 | Dgkz | Diacylglycerol kinase zeta Rattus norvegicus | 71.91 | 4.41 | 104009 | 2 | 2 | 1274328.25 | 1207170.12 | 1073381.62 | 1266015.88 | 1295138.75 | 1424261.38 | 1702466.12 | 1422879.62 | 1570516.88 | 1423209.62 | 1703564.5 | 1523170.38 | 1465322.62 | 1469373.12 |
| Q8BH86 | Dglucy | D-glutamate cyclase, mitochondrial Mus musculus | 46.52 | 3.24 | 66366 | 1 | 1 | 783220.88 | 994559.88 | 892727.81 | 801676.81 | 900332.75 | 1045247.19 | 884992.5475 | 881594.38 | 891730.5 | 721398.12 | 964261.88 | 1133439.75 | 999162.06 | 1169527 |
| O88455 | Dhcr7 | 7-dehydrocholesterol reductase Mus musculus | 210.85 | 11.25 | 53919 | 4 | 4 | 4165302.5 | 652550.19 | 2017097.62 | 731162.75 | 1030888.75 | 1083276.75 | 1064480.38 | 2686411.5 | 1172009.38 | 963705.56 | 908809 | 2355493.5 | 3053416.5 | 957070.5 |
| Q9DBB8 | Dhdh | Trans-1,2-dihydrobenzene-1,2-diol dehydrogenase Mus musculus | 253.19 | 17.42 | 36301 | 4 | 4 | 4518272.5 | 3689742.5 | 4388964 | 3267605 | 3540505.25 | 4346486.5 | 3162936 | 4453474 | 4938681.5 | 4482715 | 3754622.25 | 4745305 | 4867514 | 4541646 |
| P00375 | Dhfr | Dihydrofolate reductase Mus musculus | 279.43 | 32.62 | 21606 | 5 | 5 | 14760133 | 15929109 | 14222924 | 13578961 | 14830744 | 14828341 | 13415088 | 14754172 | 12590337 | 12846213 | 13439015 | 13205320 | 12969294 | 14625293 |
| O35435 | Dhodh | Dihydroorotate dehydrogenase (quinone), mitochondrial Mus musculus | 93.26 | 8.35 | 42700 | 2 | 2 | 858975 | 905831.12 | 701636.5 | 959163.44 | 692961.5 | 696603.56 | 591537.12 | 659897.94 | 461137 | 651447.62 | 689094 | 690762 | 477557 | 804970.25 |
| Q3TXU5 | Dhps | Deoxyhypusine synthase Mus musculus | 118.73 | 8.94 | 40642 | 2 | 2 | 323303.9 | 239570.72 | 328548.91 | 490637.78 | 234458.19 | 302127.59 | 334356.72 | 319743.25 | 331833.72 | 213544.66 | 1 | 1 | 363999.34 | 1 |
| Q99L04 | Dhrs1 | Dehydrogenase/reductase SDR family member 1 Mus musculus | 449.48 | 31.31 | 34005 | 8 | 8 | 9228698 | 9496024 | 10085423 | 8401514 | 7871312 | 7010929 | 10125811 | 7509473.5 | 10110766 | 8532169 | 8155674.5 | 6374579.5 | 9575515 | 10648475 |
| Q99LB2 | Dhrs4 | Dehydrogenase/reductase SDR family member 4 Mus musculus | 158.01 | 26.16 | 29885 | 5 | 5 | 1410262.12 | 1028558.19 | 1760978.75 | 1354389.12 | 1388547.045 | 678901.88 | 1205027 | 557063.31 | 1104334.62 | 1267402.12 | 684962.88 | 968857.56 | 1667560.5 | 1048123.69 |
| Q9CXR1 | Dhrs7 | Dehydrogenase/reductase SDR family member 7 Mus musculus | 173.06 | 13.91 | 38167 | 4 | 4 | 7001172.5 | 3615721.75 | 5859879 | 5189765 | 5816470 | 6344196.5 | 3287581.5 | 4296096 | 5270910 | 4236593 | 7575637 | 7991564 | 5591288 | 7442195 |
| Q99J47 | Dhrs7b | Dehydrogenase/reductase SDR family member 7B Mus musculus | 94.12 | 9.29 | 34987 | 2 | 2 | 1416543.75 | 1785989.88 | 1364105.88 | 1098790 | 1191268.75 | 1204873.62 | 1215560.75 | 829018.75 | 1243786.88 | 1243443.62 | 1225953.38 | 1230353.62 | 1283817.12 | 1204834.25 |
| O35286 | Dhx15 | Pre-mRNA-splicing factor ATP-dependent RNA helicase DHX15 Mus musculus | 1227.15 | 33.08 | 91007 | 21 | 23 | 40992317 | 41301829 | 43735069 | 41636641 | 43165537 | 43822629 | 40796345 | 42361921 | 43302649 | 42538657 | 39679601 | 43520441 | 43929789 | 41419209 |
| Q6PGC1 | Dhx29 | ATP-dependent RNA helicase DHX29 Mus musculus | 81.86 | 1.61 | 153975 | 2 | 2 | 517866.44 | 265201.62 | 577928.75 | 253528.73 | 222110.31 | 454068 | 553906.53 | 816775.75 | 498065.59 | 446716.78 | 379586.97 | 430110.56 | 573125.88 | 370463.47 |
| Q5BJS0 | Dhx30 | ATP-dependent RNA helicase DHX30 Rattus norvegicus | 417.71 | 7.2 | 133997 | 7 | 7 | 1912891 | 1417401.5 | 1832725.5 | 1846246.5 | 1779634.38 | 847820.25 | 4604106 | 1416766.75 | 1846611.25 | 2010823.25 | 4535818 | 800068.94 | 2087003.88 | 4335913 |
| D4A2Z8 | Dhx36 | ATP-dependent DNA/RNA helicase DHX36 Rattus norvegicus | 261.69 | 7.4 | 113843 | 6 | 6 | 1602403.38 | 2228548.25 | 2125017.75 | 1845307 | 3894404.25 | 1977454.38 | 1425083 | 1334479.25 | 1297270.75 | 1280549.5 | 1438098.62 | 1360321.75 | 1207351.75 | 1171439.25 |
| Q99J87 | Dhx58 | Probable ATP-dependent RNA helicase DHX58 Mus musculus | 65.33 | 1.62 | 76709 | 1 | 1 | 189459.12 | 153219.42 | 177991.3725 | 204719.97 | 164566.98 | 1 | 1 | 1 | 1 | 1 | 1 | 1 | 173103.42 | 1 |
| A2A4P0 | Dhx8 | ATP-dependent RNA helicase DHX8 Mus musculus | 221.25 | 5.95 | 142572 | 5 | 5 | 1605208.75 | 1430641 | 1840368.5 | 1847155.38 | 1598596.25 | 1853350.343 | 1780559.62 | 1866162.5 | 2126034 | 1640645.25 | 1687349.38 | 2835835.25 | 1692003.25 | 1309896.12 |
| O70133 | Dhx9 | ATP-dependent RNA helicase A Mus musculus | 2773.33 | 43.12 | 149475 | 48 | 49 | 86166737 | 101898305 | 103340321 | 102476721 | 98230921 | 100241225 | 95527057 | 88709513 | 96047305 | 86779553 | 98921473 | 99417233 | 98238161 | 103620769 |
| F1M775 | Diaph1 | Protein diaphanous homolog 1 Rattus norvegicus | 716.21 | 15.57 | 140409 | 14 | 14 | 8559447 | 9502858 | 9598082 | 8899180 | 9757611 | 11403285 | 10264303 | 10886076 | 10530646 | 11004361 | 10484697 | 7826268.5 | 9353816 | 10188028 |
| Q8R418 | Dicer1 | Endoribonuclease Dicer Mus musculus | 128.52 | 3.08 | 216821 | 3 | 3 | 385942.12 | 259771.64 | 133663.34 | 624994.94 | 668523.62 | 798753.38 | 615792.88 | 614594 | 672918.06 | 499222.41 | 441556.09 | 667436.06 | 260988.81 | 709939.62 |
| Q5M9G7 | Diexf | Digestive organ expansion factor homolog Rattus norvegicus | 160.09 | 4.59 | 87825 | 3 | 3 | 1 | 1 | 1 | 1 | 380879.97 | 1 | 1 | 283632.78 | 469666.59 | 316789.94 | 377079.88 | 569836.8367 | 658585.69 | 673844.94 |
| Q9D0D4 | Dimt1 | Probable dimethyladenosine transferase Mus musculus | 65.46 | 6.39 | 35274 | 2 | 2 | 2119810.25 | 1234545.88 | 3318215.5 | 1956084.72 | 1151767.25 | 1153983.12 | 1894791 | 3629478.5 | 1130185.62 | 2063323.88 | 999654.5 | 1771050.75 | 1325347.5 | 3247924 |
| Q3UH60 | Dip2b | Disco-interacting protein 2 homolog B Mus musculus | 436.25 | 7.24 | 171126 | 8 | 8 | 6769005 | 8648224 | 4753545 | 4431788.5 | 5067007 | 5037080 | 6530347 | 4373674 | 5566877.5 | 5573832.5 | 4712360.5 | 4715220 | 5305928 | 4660716.5 |
| Q9CSH3 | Dis3 | Exosome complex exonuclease RRP44 Mus musculus | 293.29 | 6.58 | 108838 | 5 | 5 | 7311804 | 4772028 | 5428749 | 5526741 | 5278218.5 | 6698805.5 | 5444000 | 4287531 | 4481656.5 | 6051164.5 | 5796377 | 6443862 | 4906086.5 | 9613221 |
| Q8CI75 | Dis3l2 | DIS3-like exonuclease 2 Mus musculus | 134.43 | 4.94 | 97775 | 3 | 3 | 1995034.5 | 1161081.88 | 1202536.38 | 1251003 | 1456505 | 1879837.5 | 1671128.62 | 1912665.88 | 2083408 | 1617647.5 | 1777283.25 | 2252873 | 1838838.38 | 1197430.38 |
| Q9ESX5 | Dkc1 | H/ACA ribonucleoprotein complex subunit DKC1 Mus musculus | 643.3 | 30.84 | 57402 | 11 | 11 | 15481263 | 14396183 | 14286213 | 14283651 | 16759178 | 12066253 | 14389273 | 13993985 | 13170665 | 14370807 | 13546061 | 12185101 | 15053765 | 14479111 |
| Q8BMF4 | Dlat | Dihydrolipoyllysine-residue acetyltransferase component of pyruvate dehydrogenase compl | 709.5 | 31 | 67942 | 13 | 13 | 31919889 | 36952193 | 38732433 | 35680857 | 43382769 | 43247585 | 38816361 | 48691205 | 40780925 | 40879889 | 38362281 | 37807373 | 40952709 | 43503453 |
| O08749 | Dld | Dihydrolipoyl dehydrogenase, mitochondrial Mus musculus | 1010.55 | 45.38 | 54272 | 16 | 16 | 91068521 | 71589561 | 59574589 | 75143865 | 65895429 | 57105733 | 84849641 | 82482873 | 67335329 | 56066889 | 57607657 | 52261449 | 63132309 | 57933181 |
| B1AZP2 | Dlgap4 | Disks large-associated protein 4 Mus musculus | 100.25 | 2.62 | 108037 | 2 | 2 | 8189513.5 | 6492710 | 3329314 | 7200943 | 2492961.5 | 6324504 | 1646540.5 | 4067401.5 | 4301812.5 | 4315111 | 4231862.5 | 6645924 | 4650576 | 7028648 |
| Q01205 | Dlst | Dihydrolipoyllysine-residue succinyltransferase component of 2-oxoglutarate dehydrogenas | 615.58 | 33.48 | 48925 | 11 | 11 | 51557229 | 38689277 | 32494921 | 46522977 | 32387917 | 34972777 | 52629365 | 46242953 | 38224285 | 29105193 | 32633845 | 28501077 | 29887571 | 30392693 |
| P63036 | Dnaja1 | DnaJ homolog subfamily A member 1 Rattus norvegicus | 389.08 | 28.97 | 44868 | 7 | 7 | 5205532.5 | 4361426.5 | 5415119 | 5853004 | 4832383 | 4785149 | 5262764 | 5488631 | 5739502.5 | 6685295 | 5553994 | 4827159.5 | 5654962 | 5472955 |
| Q9QYJ0 | Dnaja2 | DnaJ homolog subfamily A member 2 Mus musculus | 666.3 | 48.79 | 45746 | 12 | 12 | 29684813 | 9392349 | 19706347 | 29303685 | 10326661 | 25861725 | 9295060 | 9095655 | 11610171 | 10603224 | 12378520 | 12698437 | 11309700 | 10207254 |
| Q99M87 | Dnaja3 | DnaJ homolog subfamily A member 3, mitochondrial Mus musculus | 95.77 | 8.12 | 52443 | 2 | 2 | 739663.94 | 749693.12 | 618731.38 | 834531.69 | 774624.56 | 1073251.5 | 1134339.25 | 582344.62 | 906846.94 | 1104711.12 | 765075.19 | 893784.81 | 939155.81 | 1021993.75 |
| Q9QYJ3 | Dnajb1 | DnaJ homolog subfamily B member 1 Mus musculus | 165.74 | 15 | 38167 | 5 | 5 | 4165890 | 3318580 | 3386349.25 | 3622478 | 3842204.5 | 5365432.5 | 5675294 | 4051349.75 | 5571722 | 3914628.5 | 4397374 | 3961700.75 | 5283370 | 5117586 |
| Q6TUG0 | Dnajb11 | DnaJ homolog subfamily B member 11 Rattus norvegicus | 321.83 | 19.55 | 40495 | 5 | 5 | 7270898.5 | 7374524.5 | 9912708 | 8706725 | 7944480 | 8426074 | 9513764 | 6408655 | 8716583 | 9135160 | 8339242 | 9717904 | 9951676 | 9706729 |
| O54946 | Dnajb6 | DnaJ homolog subfamily B member 6 Mus musculus | 48.66 | 3.56 | 39807 | 1 | 1 | 1401260.25 | 534497.81 | 799267.75 | 530233.31 | 1341338.5 | 501134.66 | 532040.56 | 486009.72 | 554511.19 | 743604.5 | 744226.38 | 1490431.5 | 503728.94 | 1190738.75 |
| Q498R3 | Dnajc10 | DnaJ homolog subfamily C member 10 Rattus norvegicus | 77.08 | 2.52 | 90747 | 2 | 2 | 1558368.5 | 1356446.38 | 959210.19 | 1223846.38 | 1740151.12 | 1580706 | 1383554.12 | 2016727.75 | 204572.3 | 1500805 | 1824139.12 | 1 | 1508272.5 | 1 |
| Q5U458 | Dnajc11 | DnaJ homolog subfamily C member 11 Mus musculus | 68.93 | 6.08 | 63233 | 2 | 2 | 1756696.12 | 1877092.38 | 1830327.75 | 1543130.25 | 1021202.5 | 1406979.62 | 1632311.25 | 1905918.12 | 1914440.25 | 1590604.88 | 1273310.62 | 766577.19 | 1526849.25 | 1380862.38 |
| Q9CQV7 | Dnajc19 | Mitochondrial import inner membrane translocase subunit TIM14 Mus musculus | 72.39 | 16.38 | 12437 | 1 | 1 | 2629887.25 | 1677089.62 | 1913949.5 | 2179990.75 | 1895540.75 | 1589805.25 | 1882904.62 | 1775896 | 1273819.5 | 1133331.88 | 2185832 | 2130678.25 | 1962397.62 | 1985468.25 |
| P54103 | Dnajc2 | DnaJ homolog subfamily C member 2 Mus musculus | 218.77 | 10.79 | 71722 | 5 | 5 | 2841132.5 | 2349083.75 | 3566257 | 3432205 | 3433569.5 | 3923222.5 | 4275470.5 | 3619431.5 | 2410579.75 | 3387510 | 4247780 | 4833312 | 4023677.5 | 4711293.5 |
| Q91YW3 | Dnajc3 | DnaJ homolog subfamily C member 3 Mus musculus | 326.8 | 15.67 | 57464 | 5 | 5 | 2656988.75 | 1071912 | 1458179.25 | 816552.88 | 1249762.5 | 644871.69 | 694546.25 | 960467.06 | 816284.155 | 965251.62 | 1193582.5 | 652094.62 | 880727.56 | 981326.06 |
| P60904 | Dnajc5 | DnaJ homolog subfamily C member 5 Mus musculus | 99.44 | 19.7 | 22101 | 2 | 2 | 2099868.25 | 1786901.88 | 1646291.75 | 1272206.5 | 1521797.38 | 1268991.5 | 1543803.75 | 1873298 | 1379744.25 | 712753.44 | 816834.75 | 590675.44 | 1429995.25 | 1428896.75 |
| Q9QYI3 | Dnajc7 | DnaJ homolog subfamily C member 7 Mus musculus | 490.69 | 20.65 | 56476 | 8 | 8 | 12213347 | 12484708 | 14398115 | 11520088 | 11159544 | 12847733 | 12068424 | 14024911 | 13140105 | 11447443 | 11349869 | 12514636 | 11007361 | 12183570 |
| Q642C0 | Dnajc8 | DnaJ homolog subfamily C member 8 Rattus norvegicus 2 | 265.66 | 25.69 | 29813 | 6 | 6 | 11509153 | 7377620 | 6784950.5 | 9237301 | 10761839 | 10296797 | 8291085 | 7290652 | 7311711 | 8891710 | 11238783 | 8561919 | 10422898 | 9671358 |
| P21704 | Dnase1 | Deoxyribonuclease-1 Rattus norvegicus 2 | 35.75 | 2.46 | 32064 | 1 | 1 | 1642476 | 1399387.75 | 1698094.88 | 1988958.5 | 1682229.283 | 525279.88 | 1393843.25 | 1559901 | 1896942 | 1 | 1 | 1 | 2165657.75 | 1 |
| P56542 | Dnase2 | Deoxyribonuclease-2-alpha Mus musculus | 215.95 | 23.8 | 38811 | 5 | 5 | 4207111 | 2816676.5 | 3643136.5 | 2673681.75 | 3293379.25 | 3408443 | 3168977.75 | 3786091.75 | 3367688.75 | 3621635.5 | 3250105.5 | 2658750 | 3964946 | 3206103.25 |
| P21575 | Dnm1 | Dynamin-1 Rattus norvegicus | 204.22 | 5.56 | 97295 | 1 | 5 | 5910187 | 5974341.5 | 4994817 | 5374599.5 | 5425708 | 5404157 | 5384508.5 | 4948333 | 5310773 | 5101730.5 | 5222991 | 4766753 | 3689965.75 | 4639552 |
| O35303 | Dnm1l | Dynamin-1-like protein Rattus norvegicus | 547.71 | 20.93 | 83908 | 11 | 11 | 8967139 | 7922948.5 | 9526005 | 7493573.5 | 6815973 | 6568428 | 6951001 | 8621467 | 9410451 | 7315750 | 6495592 | 7276593 | 6800646 | 6117038 |
| P39052 | Dnm2 | Dynamin-2 Rattus norvegicus | 1083.22 | 34.25 | 98230 | 15 | 22 | 24288561 | 25117165 | 23822793 | 25118757 | 24548741 | 29560125 | 27019033 | 27048409 | 29030973 | 26399053 | 24601069 | 24540613 | 24127935 | 27190765 |
| Q08877 | Dnm3 | Dynamin-3 Rattus norvegicus | 280.16 | 8.63 | 97914 | 2 | 7 | 7508758 | 5974341.5 | 6584546.5 | 6278341.5 | 6236007 | 7041571.5 | 6563686.5 | 7012119 | 7002964 | 6048926.5 | 6323907 | 6577375.5 | 4822192 | 5743764 |
| Q6TXD4 | Dnmbp | Dynamin-binding protein Mus musculus | 52.96 | 0.95 | 177277 | 1 | 1 | 159029.23 | 210863.73 | 187377.3 | 206897.78 | 180768.41 | 230102.17 | 196946 | 243582.34 | 213043.03 | 154412.05 | 188856.61 | 179357.94 | 174358.3 | 200729.64 |
| P13864 | Dnmt1 | DNA (cytosine-5)-methyltransferase 1 Mus musculus | 1939.1 | 31.23 | 183189 | 20 | 36 | 39514585 | 29406143 | 31782447 | 43598445 | 37225265 | 32307311 | 33475359 | 30314753 | 33106721 | 27177299 | 30195139 | 35889513 | 25908249 | 34682597 |
| O88508 | Dnmt3a | DNA (cytosine-5)-methyltransferase 3A Mus musculus | 410.78 | 9.03 | 101672 | 7 | 7 | 7736163 | 7329320.5 | 7615047 | 8064499 | 7192452 | 7349423 | 7914292 | 10346947 | 7135477 | 8258928 | 7781242 | 7372221.5 | 6734877 | 7909178.5 |
| Q9Z2W0 | Dnpep | Aspartyl aminopeptidase Mus musculus | 473.59 | 34.25 | 52207 | 9 | 9 | 10101427 | 10563212 | 10241226 | 12578738 | 13703730 | 12471996 | 11351080 | 11344307 | 11287376 | 10070709 | 13448608 | 10403896 | 12418131 | 11887447 |
| Q8R2M2 | Dnttip2 | Deoxynucleotidyltransferase terminal-interacting protein 2 Mus musculus | 328.7 | 10.29 | 84277 | 6 | 6 | 1357707.25 | 1842853.75 | 756072 | 1547577 | 2191822.5 | 1298457 | 961008.62 | 640480.25 | 605221.38 | 849042.81 | 2173618.5 | 1208912.75 | 1017838.25 | 767445.75 |
| Q8BZN6 | Dock10 | Dedicator of cytokinesis protein 10 Mus musculus | 444.44 | 7.35 | 245758 | 10 | 10 | 5128567.5 | 5359183.5 | 5712585 | 5539528 | 5490950.5 | 6377783.5 | 5121350 | 5376155.5 | 5929483 | 5423164 | 5992834 | 5688210 | 5663250 | 5753689 |
| Q8C3J5 | Dock2 | Dedicator of cytokinesis protein 2 Mus musculus | 1397.04 | 22.26 | 211704 | 30 | 30 | 46099521 | 35438617 | 30045473 | 35338525 | 37959521 | 41848309 | 36924205 | 39700601 | 39682217 | 41518837 | 42324505 | 40268745 | 38768781 | 39276533 |
| Q8R1A4 | Dock7 | Dedicator of cytokinesis protein 7 Mus musculus | 78.76 | 1.46 | 241438 | 2 | 2 | 285566.44 | 297161.34 | 223013.19 | 1 | 1 | 176476.44 | 233978.7225 | 238262.25 | 266267.59 | 254908.61 | 194495.41 | 221422.95 | 245239.17 | 220385.8433 |
| Q8C147 | Dock8 | Dedicator of cytokinesis protein 8 Mus musculus | 692.25 | 8.52 | 238978 | 13 | 13 | 8653259 | 7572209 | 10662542 | 6787040 | 9179315 | 6081105.5 | 8786867 | 6803790.5 | 6348813.5 | 8863552 | 5496445 | 10834220 | 6732298.5 | 7071868.5 |
| Q99LN9 | Dohh | Deoxyhypusine hydroxylase Mus musculus | 35.57 | 6.29 | 32905 | 1 | 1 | 893574.75 | 1 | 1 | 1 | 1 | 1 | 1 | 555380.94 | 1 | 1 | 645517.75 | 1 | 1 | 467182.06 |
| P97465 | Dok1 | Docking protein 1 Mus musculus | 54.25 | 1.87 | 52452 | 1 | 1 | 1090736.38 | 982401.06 | 880917 | 1042535.5 | 1077710.5 | 1013748.5 | 864150 | 728587.19 | 1122047.25 | 1032512.25 | 1129546.75 | 1066574 | 906905.25 | 1148297.88 |
| O70469 | Dok2 | Docking protein 2 Mus musculus | 120.68 | 6.8 | 45522 | 2 | 2 | 238499.89 | 1 | 1 | 328955.28 | 202227.56 | 300348.19 | 185279.11 | 150383.48 | 139165.8 | 95193.5 | 1 | 1 | 87839.27 | 246834.2 |
| Q61103 | Dpf2 | Zinc finger protein ubi-d4 Mus musculus | 145.11 | 6.39 | 44230 | 2 | 2 | 1463768.38 | 2731879.75 | 2694503.5 | 3174368 | 2388616 | 1155697.38 | 2810261 | 2680117.25 | 2495084.25 | 2814779 | 2865080 | 3322811 | 2476509.5 | 2895395.75 |
| Q9CWQ0 | Dph5 | Diphthine methyl ester synthase Mus musculus | 94.73 | 12.81 | 31218 | 2 | 2 | 1 | 444648.53 | 1 | 446650.34 | 803049.88 | 309570.19 | 557145.31 | 194778.66 | 309977.19 | 892668.44 | 400980.97 | 321258.16 | 192482.03 | 692308.75 |
| O70152 | Dpm1 | Dolichol-phosphate mannosyltransferase subunit 1 Mus musculus | 248.54 | 28.85 | 29175 | 5 | 5 | 6037562 | 7843319 | 3448376.5 | 3115702.25 | 5243636.5 | 4887479 | 5235028 | 6886676 | 6277263.5 | 5124418 | 3320262.5 | 5062083.5 | 6432943.5 | 4996181 |
| Q99KK7 | Dpp3 | Dipeptidyl peptidase 3 Mus musculus | 1139.97 | 36.99 | 82898 | 8 | 18 | 32399055 | 34278557 | 35120525 | 27244561 | 30640621 | 33834197 | 31896121 | 35247977 | 35796665 | 31366291 | 34077921 | 33291303 | 34622081 | 34426401 |
| Q9ET22 | Dpp7 | Dipeptidyl peptidase 2 Mus musculus | 242.86 | 13.44 | 56254 | 5 | 5 | 2962367 | 3716168.5 | 3367190.5 | 2330683.25 | 2784412 | 3697525.5 | 3184744 | 2615106.75 | 2907720.5 | 3409950.75 | 3520182.5 | 3220810.5 | 3737444.75 | 3574181 |
| Q8BVG4 | Dpp9 | Dipeptidyl peptidase 9 Mus musculus | 203.67 | 9.05 | 98001 | 5 | 5 | 1704558.75 | 1444616.25 | 1742400.5 | 946576 | 1329590.38 | 1272352 | 1334957 | 2316074.75 | 1129482.88 | 978518.75 | 1481483.5 | 1463699.5 | 1301916.75 | 2496638.25 |
| Q99LT0 | Dpy30 | Protein dpy-30 homolog Mus musculus | 102.07 | 36.36 | 11213 | 2 | 2 | 4752878 | 7616383.5 | 6300841 | 6682824 | 5467161 | 4261067 | 7470912 | 5843031 | 6014257 | 4385720.5 | 5811124 | 6277723 | 4094769 | 4603933 |
| O08553 | Dpysl2 | Dihydropyrimidinase-related protein 2 Mus musculus | 1856.42 | 76.57 | 62278 | 28 | 30 | 191156849 | 225539953 | 227241681 | 212314257 | 229917153 | 215552049 | 217731665 | 232599585 | 230699025 | 240074017 | 202166945 | 203703121 | 192845345 | 210494657 |
| A0JPP1 | Drap1 | Dr1-associated corepressor Rattus norvegicus | 134.49 | 11.22 | 22326 | 3 | 3 | 3956198.25 | 4866696.5 | 3782367.25 | 4190249 | 4447282.5 | 4052880.5 | 4367711.5 | 4170948.5 | 4059633.25 | 3926182.75 | 4115817.5 | 4648239.5 | 3926595.5 | 4515315.5 |
| P32233 | Drg1 | Developmentally-regulated GTP-binding protein 1 Mus musculus | 409.81 | 28.34 | 40512 | 7 | 8 | 5193407.5 | 4370919 | 6439510.5 | 5552117 | 4935575 | 6509186.5 | 4998553.5 | 3950294.5 | 4035522.5 | 3680583.75 | 5793649 | 6090952.5 | 5804571.5 | 5248755 |
| Q9QXB9 | Drg2 | Developmentally-regulated GTP-binding protein 2 Mus musculus | 390.09 | 23.9 | 40718 | 6 | 7 | 5568224 | 3879384.75 | 5953378 | 6886031 | 6015587 | 6517355 | 5607033 | 4533285.5 | 6500848 | 6126306 | 7273772.5 | 5933183 | 7457752 | 6699880 |
| O35075 | Dscr3 | Down syndrome critical region protein 3 homolog Mus musculus | 181.98 | 18.86 | 32970 | 4 | 4 | 3226952.5 | 2074669 | 2602682 | 3277601 | 2343939.5 | 3699260 | 3700280 | 3005929.5 | 3198500.75 | 3557376.5 | 3515577 | 3599492.25 | 3444214.25 | 3758079 |
| O35902 | Dsg3 | Desmoglein-3 Mus musculus | 39.5 | 0.7 | 107889 | 1 | 1 | 1 | 1 | 661985.75 | 550804 | 511728.59 | 744833.38 | 494369.62 | 621809.7975 | 574040.88 | 673995.31 | 606268.94 | 1001943.88 | 610158 | 537401.31 |
| Q9R0P5 | Dstn | Destrin Mus musculus | 489.98 | 52.12 | 18522 | 7 | 7 | 29927785 | 31798485 | 33788209 | 31869721 | 32292155 | 36284041 | 34494133 | 34565989 | 34277933 | 40759389 | 39374597 | 35627793 | 34732637 | 44021397 |
| Q9DD18 | Dtd1 | D-aminoacyl-tRNA deacylase 1 Mus musculus | 52.97 | 7.18 | 23384 | 1 | 1 | 1673924 | 1824673.25 | 1704369.62 | 2156659 | 1926094.25 | 1793847.38 | 2165485.25 | 1825279 | 1955305.62 | 2108060.75 | 1956570.62 | 1827574 | 2138711 | 1477602.75 |
| Q3UIR3 | Dtx3l | E3 ubiquitin-protein ligase DTX3L Mus musculus | 64.55 | 2.14 | 83044 | 1 | 1 | 569658.75 | 527961.69 | 302946.41 | 348627.94 | 294791.12 | 351811.09 | 315014.47 | 460105.72 | 472283.34 | 398799.69 | 311512.6867 | 309742.88 | 323129.62 | 301665.56 |
| P97930 | Dtymk | Thymidylate kinase Mus musculus | 165.99 | 22.64 | 23915 | 3 | 3 | 2683525.5 | 3438320 | 3873143.25 | 3131515 | 3096032 | 2603926.5 | 2825610.25 | 2785701.5 | 3245956.75 | 2138721 | 2786470.5 | 3809269 | 2821864 | 2757985 |

| | | | | | | | | | | | | | | | | | | | | | |
|---|---|---|---|---|---|---|---|---|---|---|---|---|---|---|---|---|---|---|---|---|---|
| Q3KRC5 | Dus3l | tRNA-dihydrouridine(47) synthase [NAD(P)(+)]-like Rattus norvegicus | 142.11 | 6.88 | 71540 | 3 | 3 | 157501.19 | 827886.69 | 1165373.12 | 570771.88 | 584694.12 | 562131.75 | 820449.88 | 637792.25 | 945224.62 | 836160.12 | 785029.19 | 655535.56 | 986398.88 | 632372 |
| Q9D7X3 | Dusp3 | Dual specificity protein phosphatase 3 Mus musculus | 502.78 | 50.81 | 20472 | 7 | 7 | 19808213 | 22795277 | 21355283 | 21772333 | 21776871 | 20919559 | 22307145 | 20855033 | 21220907 | 20972361 | 21450039 | 19088379 | 19146521 | 20982893 |
| P70583 | Dut | Deoxyuridine 5'-triphosphate nucleotidohydrolase Rattus norvegicus | 229.27 | 21.46 | 22003 | 4 | 4 | 2970678.75 | 3781558.5 | 4836467.5 | 4905225 | 5730771 | 4541068.5 | 3977481 | 4908803 | 4511467 | 4829970 | 4256481.5 | 6208005 | 5958897 | 5351240 |
| Q9JHU4 | Dync1h1 | Cytoplasmic dynein 1 heavy chain 1 Mus musculus | 7019.131 | 36.95 | 532045 | 129 | 129 | 286282433 | 273644129 | 270429345 | 240687649 | 255813489 | 264730849 | 261976065 | 277339297 | 275727361 | 270377057 | 259389041 | 256412385 | 272108865 | 260939985 |
| O88487 | Dync1i2 | Cytoplasmic dynein 1 intermediate chain 2 Mus musculus | 631.77 | 21.73 | 68394 | 7 | 7 | 7223076.5 | 7905716.5 | 8210516.5 | 5927793.5 | 8112328 | 9440856 | 9085156 | 7771617 | 7354350 | 9140906 | 7844017 | 7069392.5 | 9177453 | 9583163 |
| Q8R1Q8 | Dync1li1 | Cytoplasmic dynein 1 light intermediate chain 1 Mus musculus | 358.85 | 18.74 | 56614 | 7 | 7 | 14092548 | 11484200 | 11401878 | 11801322 | 12836591 | 14434362 | 10802351 | 15106537 | 14398969 | 12742647 | 13291326 | 14633052 | 13861502 | 14134802 |
| Q62698 | Dync1li2 | Cytoplasmic dynein 1 light intermediate chain 2 Rattus norvegicus | 282.53 | 18.51 | 54745 | 5 | 5 | 1057448 | 1081162.62 | 400488.09 | 996846.9275 | 1448289 | 677694.06 | 381639.59 | 1040039.25 | 1049026.5 | 568405.62 | 650396.88 | 545404 | 369193.69 | 754762.75 |
| P63168 | Dynll1 | Dynein light chain 1, cytoplasmic Mus musculus | 183.83 | 49.44 | 10366 | 1 | 3 | 8300341 | 7898261.5 | 5338728 | 5613036.5 | 7264544 | 6856328 | 6232337.5 | 7770049.5 | 6489973.5 | 6603360 | 6194718.5 | 5368665 | 6034645 | 6813088 |
| Q78P75 | Dynll2 | Dynein light chain 2, cytoplasmic Rattus norvegicus | 121.15 | 49.44 | 10350 | 1 | 3 | 6247390.5 | 5720783.5 | 3357348.5 | 4144981.5 | 5320865 | 4614856 | 4773525 | 5596313.5 | 4431660.5 | 4965437.5 | 4733524 | 3793548.75 | 4521027 | 4878553.5 |
| P51807 | Dynlt1 | Dynein light chain Tctex-type 1 Mus musculus | 216.52 | 44.25 | 12483 | 3 | 3 | 1041992.75 | 870806.56 | 694207.62 | 601861 | 634501.62 | 1051696.25 | 548996.81 | 646958.12 | 749995.56 | 841053.88 | 999107.75 | 616139.75 | 667366.06 | 875219.69 |
| Q9ESD7 | Dysf | Dysferlin Mus musculus | 123.45 | 1 | 237911 | 1 | 2 | 4066897.25 | 4145274.25 | 4969147.5 | 2661205.75 | 3306966.5 | 3920354 | 3594352.75 | 4138949 | 4482063.5 | 3666642.75 | 3476375 | 3642162.25 | 4516820.5 | 4103523 |
| Q9D903 | Ebna1bp2 | Probable rRNA-processing protein EBP2 Mus musculus E | 179.36 | 17.97 | 34703 | 4 | 4 | 1567059.38 | 2113400.5 | 1280972 | 2690933 | 1567930.5 | 2198458.5 | 1892652.25 | 1779422.88 | 1996623.5 | 2265153 | 4261450 | 3059703.5 | 4068897.5 | 5568977.5 |
| Q9CS74 | Ecd | Protein ecdysoneless homolog Mus musculus | 72.85 | 5.77 | 71710 | 2 | 2 | 1 | 161375.7 | 160322.73 | 190806.33 | 1 | 1 | 1 | 1 | 1 | 1 | 1 | 1 | 88618.08 | 1 |
| O35459 | Ech1 | Delta(3,5)-Delta(2,4)-dienoyl-CoA isomerase, mitochondrial Mus musculus | 237.98 | 19.27 | 36118 | 5 | 5 | 4140307.5 | 3485194 | 2630151.5 | 2344078.5 | 5104339 | 2769596.5 | 3012022.5 | 4379939 | 2863489.5 | 3014528.25 | 3051047.5 | 3709461.75 | 2745480.5 | 3021517.5 |
| Q9D9V3 | Echdc1 | Ethylmalonyl-CoA decarboxylase Mus musculus | 151.89 | 17.7 | 35467 | 3 | 4 | 3508413.25 | 3236497 | 2768885.75 | 3074386.75 | 3010218.25 | 3217053.25 | 2410408.5 | 2349392.25 | 3115213 | 1723640.12 | 2743556 | 3196660.5 | 2828329.75 | 2729988 |
| Q8BH95 | Echs1 | Enoyl-CoA hydratase, mitochondrial Mus musculus | 221.95 | 18.28 | 31474 | 4 | 4 | 1069732 | 682670.5 | 2071879.5 | 1335962.62 | 1448323 | 1388431.5 | 1388512.25 | 786034.75 | 1866722.25 | 2547823.75 | 2045750 | 2189380 | 2676057 | 1704333.38 |
| P42125 | Eci1 | Enoyl-CoA delta isomerase 1, mitochondrial Mus musculus | 457.5 | 34.26 | 32250 | 6 | 6 | 7640934 | 8610438 | 9263151 | 7298505 | 8384398 | 9542675 | 8317529 | 10241237 | 8374461.5 | 6105004 | 6133395 | 7561436 | 7828195 | 7861822 |
| Q9WUR2 | Eci2 | Enoyl-CoA delta isomerase 2, mitochondrial Mus musculus | 237.6 | 12.53 | 43268 | 4 | 4 | 8570729 | 2119102.5 | 1910965.75 | 2013180 | 2101377 | 8518567 | 978466.06 | 8711249 | 1664050.75 | 1856673.38 | 7451204.5 | 1734958 | 1867380.25 | 9392752 |
| Q6PDI5 | Ecpas | Proteasome adapter and scaffold protein ECM29 Mus musculus | 1066.03 | 16.47 | 203703 | 19 | 19 | 12597425 | 11245704 | 10650420 | 10602916 | 12927743 | 11729638 | 11770303 | 12662172 | 10719007 | 10904320 | 12310510 | 9991071 | 12381996 | 12139845 |
| Q8K2D3 | Edc3 | Enhancer of mRNA-decapping protein 3 Mus musculus | 81.51 | 7.87 | 55957 | 2 | 2 | 176901.52 | 250268.17 | 273306.28 | 347917.59 | 300374.81 | 522904.25 | 187677.61 | 168956.95 | 260944.17 | 357484.81 | 594012.44 | 1 | 260178.44 | 1 |
| Q3UJB9 | Edc4 | Enhancer of mRNA-decapping protein 4 Mus musculus | 374.94 | 9.39 | 152484 | 1 | 7 | 4353501 | 4966455 | 6571259 | 5920117.5 | 6745177.5 | 4583599 | 4710631 | 6912591 | 5945215 | 4835685 | 5767570.5 | 6465669 | 4705997 | 4792388 |
| Q8BL66 | Eea1 | Early endosome antigen 1 Mus musculus | 405.09 | 8.5 | 160915 | 8 | 8 | 1834322.75 | 1916873.25 | 1880146.88 | 3753542.75 | 2587111 | 2037316.75 | 3460101.25 | 1994031.5 | 2437493.75 | 2369758.75 | 2240230.75 | 3500059 | 1587198.75 | 2163271.5 |
| Q921E6 | Eed | Polycomb protein EED Mus musculus | 113.96 | 7.71 | 50198 | 3 | 3 | 1085072.12 | 1522864.88 | 1627659 | 2256247.5 | 2388143.5 | 1130694.12 | 1519780.12 | 1191034.75 | 2402969.5 | 1192057.12 | 1130155.12 | 2584589.5 | 1247150.62 | 1676137.75 |
| P10126 | Eef1a1 | Elongation factor 1-alpha 1 Mus musculus | 2132.89 | 61.26 | 50114 | 17 | 36 | 1027456385 | 965531201 | 1188962817 | 1044751617 | 929421057 | 953019265 | 1054875649 | 993126273 | 1040565249 | 1072151233 | 999525953 | 1087915777 | 1020407297 | 995564353 |
| P62631 | Eef1a2 | Elongation factor 1-alpha 2 Mus musculus | 988.28 | 29.16 | 50454 | 1 | 17 | 684303553 | 651505345 | 811386049 | 745664257 | 637148673 | 646525505 | 732425281 | 610859073 | 712283777 | 747590529 | 681522817 | 760596417 | 694924353 | 669860609 |
| Q9D853 | Eef1akmt2 | EEF1A lysine methyltransferase 2 Mus musculus | 47.95 | 8.61 | 26864 | 1 | 1 | 1 | 202524.69 | 209280.45 | 1 | 1 | 134764.48 | 185319.41 | 166409.78 | 143046.67 | 155356.19 | 1 | 1 | 1 | 1 |
| O70251 | Eef1b | Elongation factor 1-beta Mus musculus | 753.58 | 66.22 | 24694 | 10 | 12 | 36619877 | 39156001 | 42035241 | 37366705 | 33599133 | 32904041 | 35011453 | 40616445 | 37892005 | 37087821 | 32331483 | 38694101 | 37487469 | 36452289 |
| P57776 | Eef1d | Elongation factor 1-delta Mus musculus | 933.53 | 56.94 | 31293 | 12 | 15 | 102198297 | 121863249 | 108618625 | 112663921 | 113851521 | 103507257 | 110478257 | 107205577 | 101421465 | 115565345 | 109149545 | 108214953 | 110197337 | 118385369 |
| Q9D1M4 | Eef1e1 | Eukaryotic translation elongation factor 1 epsilon-1 Mus musculus | 98.65 | 20.11 | 19859 | 2 | 2 | 1216115.75 | 895468.0925 | 645879.75 | 441789.62 | 1278087.25 | 1388506.75 | 614223.06 | 1310354.89 | 1230678.5 | 2008011.25 | 1 | 1669722.38 | 1 | 1 |
| Q9D8N0 | Eef1g | Elongation factor 1-gamma Mus musculus | 1580.78 | 77.8 | 50061 | 26 | 26 | 137479553 | 131329681 | 126536641 | 116026641 | 130042561 | 130389897 | 126088225 | 131621337 | 119579633 | 127075241 | 131980473 | 120046729 | 131319313 | 135572641 |
| P58252 | Eef2 | Elongation factor 2 Mus musculus | 3344.97 | 61.54 | 95314 | 50 | 52 | 326634849 | 330409569 | 354644641 | 332265505 | 309318529 | 319295617 | 335195137 | 345666145 | 354945985 | 329864833 | 328167297 | 347663777 | 345752833 | 328488833 |
| Q9JHW4 | Eefsec | Selenocysteine-specific elongation factor Mus musculus | 318.03 | 16.64 | 63539 | 7 | 7 | 1463852.38 | 1441641.12 | 1455018.88 | 2027843 | 1809071 | 2708750 | 1147499.25 | 1953776 | 1706999.75 | 1510919.25 | 1665944.75 | 3895084.75 | 2176082 | 2273189.75 |
| Q3TGW2 | Eepd1 | Endonuclease/exonuclease/phosphatase family domain-containing protein 1 Mus musculus | 41.23 | 2.64 | 62953 | 1 | 1 | 884563.31 | 910401.94 | 1 | 646673 | 1 | 1015498.62 | 1052696.62 | 1135702.12 | 1087941.25 | 932121.44 | 1332034.5 | 1077017.25 | 957574.06 | 1058873.5 |
| Q9D8Y0 | Efhd2 | EF-hand domain-containing protein D2 Mus musculus | 850.28 | 68.33 | 26791 | 1 | 16 | 111222113 | 112214137 | 105129297 | 120351553 | 124314817 | 137244401 | 94090753 | 121174321 | 119232873 | 134029009 | 124425345 | 125054225 | 113676233 | 119179513 |
| Q8C0D5 | Efl1 | Elongation factor-like GTPase 1 Mus musculus | 96.08 | 2.13 | 125777 | 2 | 2 | 948018.19 | 1185063 | 1237860.62 | 1375599 | 1284588.88 | 1264720.38 | 1806917.5 | 1425036.88 | 1263848.12 | 1234130.75 | 1322149.75 | 1334086.5 | 1445605.88 | 1460848.75 |
| Q8BG67 | Efr3a | Protein EFR3 homolog A Mus musculus | 102.1 | 3.91 | 92613 | 2 | 2 | 246584.97 | 2561251 | 1414644.75 | 1100180.145 | 178239.86 | 223832.36 | 1831338.75 | 1339877.75 | 242905.39 | 1171172.62 | 192622.06 | 184069.53 | 169342.94 | 1877184.75 |
| O08810 | Eftud2 | 116 kDa U5 small nuclear ribonucleoprotein component Mus musculus | 1673.83 | 45.93 | 109361 | 31 | 31 | 31971007 | 28137881 | 35297185 | 29333603 | 27991733 | 38356533 | 28873071 | 36176125 | 34722373 | 31261567 | 28485921 | 34267409 | 35411773 | 32857303 |
| Q99MS7 | Ehbp1l1 | EH domain-binding protein 1-like protein 1 Mus musculus | 330.96 | 5.19 | 184834 | 6 | 6 | 3124244.5 | 2300516.5 | 3008927.75 | 2818594.25 | 3847507.25 | 2789632.5 | 2288165.25 | 2330174.5 | 2678896.5 | 2351470.5 | 3213875.5 | 3082915 | 2972232.5 | 3182990.5 |
| Q641Z6 | Ehd1 | EH domain-containing protein 1 Rattus norvegicus | 1135.41 | 48.69 | 60603 | 9 | 21 | 37865081 | 28337299 | 30381685 | 28628741 | 27820769 | 34030737 | 37163581 | 34173897 | 35972837 | 38206693 | 32207329 | 32877989 | 31064197 | 31254005 |
| Q8R491 | Ehd3 | EH domain-containing protein 3 Rattus norvegicus | 724.66 | 27.29 | 60791 | 1 | 15 | 19499203 | 14338282 | 16051792 | 15090915 | 13285443 | 17522715 | 18181385 | 16563661 | 18813023 | 20579707 | 16522527 | 18005433 | 17306563 | 16761165 |
| Q9EQP2 | Ehd4 | EH domain-containing protein 4 Mus musculus | 1173.64 | 47.13 | 61481 | 17 | 21 | 45898105 | 35515837 | 35702145 | 34853389 | 35500921 | 33299337 | 34314649 | 30702581 | 33440849 | 35102593 | 31913481 | 34874433 | 35917621 | 32514789 |
| P48024 | Eif1 | Eukaryotic translation initiation factor 1 Mus musculus | 234.65 | 53.1 | 12747 | 4 | 4 | 931350.94 | 396710.88 | 481420.78 | 522596.165 | 280902.06 | 547497.75 | 537898.94 | 650386.25 | 661904.81 | 506597.38 | 417938.59 | 537821.19 | 595451.31 | 292143.94 |
| Q60872 | Eif1a | Eukaryotic translation initiation factor 1A Mus musculus | 297.5 | 29.17 | 16502 | 1 | 4 | 6098777.5 | 3301234.25 | 4047254 | 4697406.5 | 5013342 | 4288429 | 4533033 | 3335546 | 4510665 | 5163793 | 4119202 | 4695035 | 4117465.25 | 4433278 |
| Q8BMJ3 | Eif1ax | Eukaryotic translation initiation factor 1A, X-chromosomal Mus musculus 2 | 343.49 | 36.11 | 16460 | 2 | 5 | 9468905 | 5668160 | 6419049.5 | 7456208 | 7460835 | 7123439.5 | 6960449.5 | 5535719 | 7133845 | 7829767 | 7091995 | 7705174.5 | 6663468 | 7187062 |
| Q8BJW6 | Eif2a | Eukaryotic translation initiation factor 2A Mus musculus | 567.33 | 28.92 | 64403 | 11 | 11 | 5282611 | 4372255 | 6031035 | 5546082.5 | 4430541 | 4158442.75 | 3982337.5 | 4112891 | 5316611.5 | 5018592.5 | 4776460.5 | 4452275.5 | 4849810.5 | 5759045 |
| Q03963 | Eif2ak2 | Interferon-induced, double-stranded RNA-activated protein kinase Mus musculus | 387.12 | 17.09 | 58280 | 6 | 6 | 6568350 | 3811978 | 3449479.75 | 2326365 | 2465058.5 | 3344496.25 | 1535612.38 | 2292577.5 | 1797691.5 | 2961848.75 | 3063750.25 | 4366944 | 3664662.75 | 2626936.5 |
| Q99LC8 | Eif2b1 | Translation initiation factor eIF-2B subunit alpha Mus musculus | 206.58 | 13.11 | 33816 | 3 | 3 | 1076993.88 | 3882521.25 | 3815415 | 2441191 | 2117378 | 4451907 | 3545106 | 6866003.5 | 4853475 | 1056371.75 | 2559697.5 | 5169764 | 3464045.75 | 3108552.75 |
| Q62818 | Eif2b2 | Translation initiation factor eIF-2B subunit beta Rattus norvegicus 2 | 93.04 | 8.55 | 38875 | 2 | 2 | 551711.5 | 161822.95 | 189998.22 | 226018.19 | 151747.2 | 144340.61 | 171876.08 | 1 | 178085.12 | 1 | 147518.83 | 141694.95 | 169306.81 | 131554.73 |
| P70541 | Eif2b3 | Translation initiation factor eIF-2B subunit gamma Rattus norvegicus 2 | 184.38 | 11.06 | 50435 | 4 | 4 | 3885131 | 5173464 | 5457932 | 4769345.5 | 4171754 | 4339871.5 | 6104583.5 | 5596407.5 | 6319935.5 | 5394965.5 | 4752591.5 | 4760056 | 5591033 | 5397565 |
| Q61749 | Eif2b4 | Translation initiation factor eIF-2B subunit delta Mus musculus | 270.95 | 14.31 | 57624 | 5 | 5 | 5332604 | 5539814 | 7310752 | 5345673 | 5296609 | 6460334.5 | 5870835.5 | 4313669 | 5446728 | 5543978.5 | 5265709.5 | 8423859 | 6682002.5 | 5820357 |
| Q8CHW4 | Eif2b5 | Translation initiation factor eIF-2B subunit epsilon Mus musculus | 252.21 | 9.07 | 80086 | 5 | 5 | 6108872 | 4428276.5 | 6308202 | 4816150 | 5639807 | 6573563 | 3805471.25 | 6073813 | 5772690 | 5311493.5 | 5810293.5 | 6059403 | 5717401.5 | 5235013 |
| P68101 | Eif2s1 | Eukaryotic translation initiation factor 2 subunit 1 Rattus norvegicus | 1198.18 | 70.79 | 36108 | 19 | 20 | 74835921 | 83887617 | 85171801 | 80947633 | 85725169 | 93875897 | 84537305 | 88022513 | 86702537 | 89098241 | 81731305 | 88946153 | 94515857 | 91406385 |
| Q99L45 | Eif2s2 | Eukaryotic translation initiation factor 2 subunit 2 Mus musculus | 736.98 | 45.92 | 38092 | 14 | 14 | 13474387 | 6245353 | 8476924 | 8165452.5 | 4590240.5 | 9154604 | 8758123 | 10285436 | 10079600 | 10575643 | 8332889.5 | 8706542 | 8954181 | 9052971 |
| Q9Z0N1 | Eif2s3x | Eukaryotic translation initiation factor 2 subunit 3, X-linked Mus musculus | 1176.57 | 54.66 | 51065 | 19 | 20 | 34462809 | 33809809 | 35290821 | 40723009 | 31704717 | 28845563 | 38391633 | 37316745 | 29933517 | 34476513 | 38929689 | 37762081 | 32497957 | 29891929 |
| P23116 | Eif3a | Eukaryotic translation initiation factor 3 subunit A Mus musculus | 1322.97 | 25.89 | 161936 | 29 | 29 | 57096769 | 44178701 | 37261657 | 41998193 | 48662077 | 46450841 | 40604301 | 38929309 | 46242353 | 48256393 | 45024917 | 41904469 | 41555081 | 47418861 |
| Q8JZQ9 | Eif3b | Eukaryotic translation initiation factor 3 subunit B Mus musculus | 1398.48 | 48.19 | 91370 | 27 | 27 | 26909591 | 23923837 | 28357913 | 21258719 | 20898801 | 23047267 | 23121917 | 22890481 | 29230849 | 24132907 | 19169629 | 23115487 | 23347335 | 20371957 |
| B5DFC8 | Eif3c | Eukaryotic translation initiation factor 3 subunit C Rattus norvegicus | 1050.91 | 23.05 | 105435 | 20 | 20 | 42405489 | 36620485 | 39037785 | 35756045 | 41643897 | 37568249 | 33698801 | 35754621 | 37442825 | 37980537 | 37710389 | 35609917 | 37159777 | 39780125 |
| O70194 | Eif3d | Eukaryotic translation initiation factor 3 subunit D Mus musculus | 736.8 | 40.69 | 63989 | 13 | 13 | 16592148 | 18260059 | 19410627 | 16976993 | 19256145 | 20605241 | 17853237 | 17122629 | 20794139 | 18560397 | 20446249 | 16203981 | 22074855 | 20483763 |
| P60229 | Eif3e | Eukaryotic translation initiation factor 3 subunit E Mus musculus | 1219.69 | 54.16 | 52221 | 20 | 20 | 40375681 | 33781045 | 46782873 | 37196785 | 31846987 | 39211649 | 34523301 | 45985909 | 43415485 | 33731497 | 37235461 | 39763697 | 38830773 | 36678165 |
| Q9DCH4 | Eif3f | Eukaryotic translation initiation factor 3 subunit F Mus musculus | 896.39 | 48.75 | 37984 | 13 | 13 | 35004189 | 38736741 | 47615741 | 29747607 | 28078943 | 36213409 | 32407635 | 43799997 | 42115533 | 32348567 | 31923391 | 42572337 | 44903217 | 35302865 |
| Q9Z1D1 | Eif3g | Eukaryotic translation initiation factor 3 subunit G Mus musculus | 445.95 | 36.88 | 35638 | 9 | 9 | 22536869 | 21529325 | 26223417 | 25183975 | 24896031 | 23475963 | 22396907 | 20574289 | 22513641 | 23791471 | 25966401 | 24424203 | 22906757 | 24315641 |
| Q91WK2 | Eif3h | Eukaryotic translation initiation factor 3 subunit H Mus musculus | 774.68 | 52.27 | 39832 | 13 | 13 | 28118545 | 27135455 | 24421703 | 25477219 | 31439245 | 28777171 | 23199699 | 21817245 | 26631247 | 29268805 | 29026955 | 28557595 | 24280863 | 30131389 |
| Q9QZD9 | Eif3i | Eukaryotic translation initiation factor 3 subunit I Mus musculus | 737.39 | 64.92 | 36461 | 15 | 15 | 13739393 | 9488356 | 13646486 | 11548385 | 12774445 | 12925812 | 11673102 | 13001693 | 12852923 | 13273945 | 14090297 | 13397522 | 13864057 | 13551669 |
| Q3UGC7 | Eif3j1 | Eukaryotic translation initiation factor 3 subunit J-A Mus musculus 2 | 397.51 | 39.85 | 29344 | 8 | 8 | 6638919.5 | 8447682 | 5934762 | 7575206 | 8514462 | 9194735 | 11699943 | 7544610 | 7782351 | 8611373 | 9450860 | 5816444.5 | 8311961 | 10349713 |
| Q9DBZ5 | Eif3k | Eukaryotic translation initiation factor 3 subunit K Mus musculus | 282.58 | 27.98 | 25087 | 4 | 4 | 11773104 | 10850116 | 12716353 | 6303225 | 11711519 | 12330026 | 10291080 | 11037607 | 6989932.5 | 10929177 | 10736804 | 11902208 | 12441889 | 10753558 |
| Q8QZY1 | Eif3l | Eukaryotic translation initiation factor 3 subunit L Mus musculus | 989.73 | 37.06 | 66613 | 16 | 17 | 25309131 | 20410773 | 20383457 | 17605967 | 19905763 | 23720069 | 18794731 | 24437527 | 21857559 | 18227155 | 19497967 | 17153627 | 22415839 | 21673703 |
| Q99JX4 | Eif3m | Eukaryotic translation initiation factor 3 subunit M Mus musculus | 811.31 | 53.21 | 42517 | 12 | 12 | 15678228 | 13660635 | 15983906 | 13005503 | 10542719 | 13411919 | 13412366 | 15867793 | 14678714 | 13769444 | 12338823 | 14144101 | 13112824 | 10714219 |
| P60843 | Eif4a1 | Eukaryotic initiation factor 4A-I Mus musculus | 2206.6 | 83.25 | 46154 | 18 | 32 | 325793025 | 335565217 | 354217793 | 289204417 | 307334721 | 302127105 | 314754497 | 321874273 | 303494145 | 297120545 | 276102337 | 311796993 | 297724193 | 286355553 |
| P10630 | Eif4a2 | Eukaryotic initiation factor 4A-II Mus musculus | 938.42 | 33.42 | 46402 | 1 | 15 | 216110977 | 216853345 | 236527617 | 198028017 | 208413969 | 197293457 | 206115505 | 207606657 | 196323425 | 200602705 | 186702353 | 199770977 | 198326481 | 189803553 |
| Q91VC3 | Eif4a3 | Eukaryotic initiation factor 4A-III Mus musculus | 1315.96 | 59.61 | 46840 | 20 | 23 | 60105225 | 72170793 | 70375913 | 67179833 | 72956833 | 69281689 | 70385945 | 72777001 | 68628649 | 72897353 | 64660221 | 70861209 | 68810609 | 66719049 |
| Q8BGD9 | Eif4b | Eukaryotic translation initiation factor 4B Mus musculus | 1121.46 | 37.97 | 68840 | 19 | 19 | 26457303 | 19061721 | 21141321 | 25850855 | 22335831 | 29444335 | 26995367 | 22274587 | 20799573 | 29753875 | 29161497 | 23755477 | 29712187 | 30535009 |
| P63073 | Eif4e | Eukaryotic translation initiation factor 4E Mus musculus | 221.31 | 21.66 | 25053 | 5 | 5 | 10787272 | 7963477 | 8493354 | 12022545 | 8465217 | 7852479.5 | 8785402 | 9353487 | 8810601 | 11519077 | 7647948.5 | 9098724 | 8927771 | 8753847 |
| Q8BMB3 | Eif4e2 | Eukaryotic translation initiation factor 4E type 2 Mus musculus | 34.37 | 4.49 | 28263 | 1 | 1 | 1081350.5 | 1431085.12 | 1438545.88 | 1158269.62 | 1095246.75 | 1106279.75 | 1317892.75 | 1316892 | 1057513.88 | 1278957.38 | 997289.19 | 1069576.5 | 1038758.69 | 1172681.62 |
| Q6NZJ6 | Eif4g1 | Eukaryotic translation initiation factor 4 gamma 1 Mus musculus | 2441.32 | 30.88 | 176077 | 36 | 39 | 60170577 | 60680209 | 52271645 | 45380969 | 48350205 | 50700453 | 55548033 | 59202193 | 50542129 | 55181961 | 53332777 | 60923777 | 59577193 | 51017541 |
| Q62448 | Eif4g2 | Eukaryotic translation initiation factor 4 gamma 2 Mus musculus | 1138.33 | 36.09 | 102106 | 24 | 24 | 22325795 | 16805373 | 20646151 | 14888943 | 16771886 | 18761781 | 18492253 | 22017887 | 19322223 | 17988821 | 16183223 | 19259731 | 21867999 | 18742339 |
| Q80XI3 | Eif4g3 | Eukaryotic translation initiation factor 4 gamma 3 Mus musculus | 492.37 | 6.84 | 174890 | 7 | 10 | 14518640 | 13511615 | 14310582 | 11453613 | 10705974 | 13701986 | 13212282 | 14525976 | 13673346 | 12742569 | 13906315 | 12619349 | 13841760 | 15248673 |
| Q5XI72 | Eif4h | Eukaryotic translation initiation factor 4H Rattus norvegicus | 652.42 | 49.6 | 27324 | 11 | 11 | 23070013 | 32763969 | 22664043 | 25098147 | 26287501 | 22959489 | 25683345 | 31609673 | 27218095 | 23132511 | 27038465 | 21227621 | 21150165 | 24789737 |
| P59325 | Eif5 | Eukaryotic translation initiation factor 5 Mus musculus | 290.59 | 13.29 | 48968 | 5 | 5 | 10179051 | 8915562 | 9307121 | 9392106 | 10261104 | 10624993 | 10484176 | 10996545 | 10274364 | 10722081 | 10795490 | 10582504 | 11922213 | 11142316 |
| P63242 | Eif5a | Eukaryotic translation initiation factor 5A-1 Mus musculus | 1045.08 | 88.31 | 16832 | 14 | 15 | 125870945 | 127178625 | 152176417 | 127524241 | 88316369 | 135720497 | 139334961 | 165439825 | 160879089 | 144208609 | 142868353 | 152212369 | 156684913 | 157686529 |
| Q05D44 | Eif5b | Eukaryotic translation initiation factor 5B Mus musculus | 1257.04 | 32.24 | 137616 | 25 | 25 | 27146141 | 22656091 | 19270283 | 24558887 | 22207337 | 32076987 | 21573595 | 25275587 | 27133763 | 31281033 | 27341947 | 25834085 | 27195345 | 23094847 |
| O55135 | Eif6 | Eukaryotic translation initiation factor 6 Mus musculus | 504.25 | 46.53 | 26511 | 8 | 8 | 16486477 | 17500049 | 18771293 | 19021457 | 16594559 | 18590255 | 18540673 | 18404879 | 18865585 | 20500099 | 15422814 | 16019709 | 18555553 | 18588083 |
| Q80Y81 | Elac2 | Zinc phosphodiesterase ELAC protein 2 Mus musculus | 254.86 | 8.66 | 92719 | 6 | 6 | 6394434 | 6164484.5 | 5424469 | 6997979 | 6631873 | 5834920 | 6481397 | 5352327 | 5327019.5 | 5887636 | 5625542 | 6485929 | 5770198.5 | 5967164 |
| B5DF91 | Elavl1 | ELAV-like protein 1 Rattus norvegicus | 600.85 | 38.65 | 36169 | 10 | 10 | 24938427 | 24757395 | 27992295 | 29491109 | 24272397 | 24344577 | 26659787 | 25227497 | 23851091 | 27248961 | 25908037 | 24387781 | 24763877 | 25295543 |
| O08856 | Ell | RNA polymerase II elongation factor ELL Mus musculus | 67.85 | 1.83 | 67146 | 1 | 1 | 285321.78 | 228550.61 | 198114.03 | 274333.495 | 385347.56 | 391270.69 | 337763.59 | 1 | 294997.53 | 1 | 1 | 335357.56 | 1 | 326019.91 |
| Q8BPU7 | Elmo1 | Engulfment and cell motility protein 1 Mus musculus | 636.57 | 24.9 | 83936 | 12 | 13 | 12739519 | 14630181 | 15591896 | 15240503 | 16312340 | 13638091 | 15895690 | 16937365 | 15807121 | 14174281 | 14747930 | 13642096 | 10343916 | 13686612 |
| Q8BHL5 | Elmo2 | Engulfment and cell motility protein 2 Mus musculus | 164.3 | 7.38 | 83887 | 3 | 4 | 2126161.25 | 1153252.12 | 1210852.62 | 2955921 | 1700594.5 | 2048333.12 | 2669245.25 | 2493268 | 1898584.38 | 2147026.25 | 2097114.12 | 1117902.12 | 1391491.75 | 1095302 |
| Q8CB77 | Eloa | Elongin-A Mus musculus | 105.93 | 5.3 | 87161 | 2 | 2 | 861514 | 854246.44 | 685421.5 | 931225.75 | 914140.5 | 823284.38 | 773544.38 | 890407.62 | 850358.25 | 848276.38 | 740278.62 | 941042.19 | 876390.12 | 866783.81 |
| P62869 | Elob | Elongin-B Mus musculus | 420.08 | 59.32 | 13170 | 8 | 8 | 40089149 | 39224705 | 40423361 | 46266561 | 39398197 | 37677413 | 37533953 | 38442097 | 39750989 | 41510349 | 38228565 | 39037369 | 37950597 | 40691765 |
| P83940 | Eloc | Elongin-C Mus musculus | 207.44 | 30.36 | 12473 | 2 | 2 | 3720203.5 | 3612083.5 | 3997336 | 4232823 | 4573965 | 5138137.5 | 4283929.5 | 3193797.75 | 5126163 | 6214968 | 5172096 | 4905423 | 5666234 | 4760511.5 |

| | | | | | | | | | | | | | | | | | | | | | |
|---|---|---|---|---|---|---|---|---|---|---|---|---|---|---|---|---|---|---|---|---|---|
| Q7TT37 | Elp1 | Elongator complex protein 1 Mus musculus | 267.45 | 4.65 | 149584 | 5 | 5 | 2316355 | 1619828.5 | 1629935.12 | 1721657.25 | 2597709.5 | 3143377.25 | 1607188.25 | 2924868.75 | 2481901 | 2437639 | 3018784.25 | 2815225.5 | 2577866.25 | 3220420 |
| Q9CZX0 | Elp3 | Elongator complex protein 3 Mus musculus | 259.32 | 11.88 | 62385 | 5 | 5 | 950490.5 | 482626.75 | 1815457.38 | 1871549.75 | 1444335.88 | 2511329.25 | 2058298 | 752426.62 | 1315737 | 741678.88 | 1185380.75 | 2493047.5 | 1671697.25 | 1161068 |
| Q8C7X2 | Emc1 | ER membrane protein complex subunit 1 Mus musculus | 706.36 | 19.36 | 111605 | 12 | 12 | 11590910 | 9152788 | 9775971 | 9215133 | 8609014 | 10015037 | 11805245 | 10916150 | 11314294 | 9715801 | 10864630 | 10851058 | 11189965 | 9778974 |
| Q5U2V8 | Emc3 | ER membrane protein complex subunit 3 Rattus norvegicus | 153.74 | 14.18 | 29980 | 3 | 3 | 3914306 | 5538289 | 4979377.5 | 4787448 | 3768954.75 | 2564742.5 | 4835779 | 4352444.5 | 4722816.5 | 3867850.5 | 4787084.5 | 4865499 | 4657085.5 | 4786192 |
| Q9CZX9 | Emc4 | ER membrane protein complex subunit 4 Mus musculus | 53.73 | 8.74 | 20117 | 1 | 1 | 3121233 | 3703782.5 | 3606992.75 | 4100907.5 | 3399092.25 | 4006762.75 | 3142540.25 | 2590723.25 | 2806799 | 3120217.75 | 3179908.5 | 4219055.5 | 3053554.75 | 3681084.75 |
| O70378 | Emc8 | ER membrane protein complex subunit 8 Mus musculus | 272.81 | 26.09 | 23348 | 4 | 4 | 6220320 | 7126992 | 6707797 | 6752939.5 | 5990222 | 5620938 | 7347929.5 | 5082604 | 6404004.5 | 5856890.5 | 7421213.5 | 6737461.5 | 7055588.5 | 7513481 |
| O08579 | Emd | Emerin Mus musculus | 347.14 | 41.7 | 29436 | 7 | 7 | 4667484.5 | 6607882.5 | 5862455 | 8795026 | 4948861 | 6431103 | 6949030 | 6497713 | 7148007 | 7028363 | 6557111 | 5336212 | 5820905 | 5161019 |
| O35130 | Emg1 | Ribosomal RNA small subunit methyltransferase NEP1 Mus musculus | 399.82 | 33.61 | 26974 | 6 | 6 | 3260497 | 2907007 | 4997991.5 | 2031453.5 | 3379251.5 | 4825036.5 | 5609383 | 5495119 | 3181017.75 | 2962544.75 | 2966473.5 | 2908584 | 3362361.5 | 3672508.25 |
| Q6P6T4 | Eml2 | Echinoderm microtubule-associated protein-like 2 Rattus norvegicus | 151.15 | 7.7 | 70711 | 3 | 3 | 5018700.938 | 6989340 | 7348600 | 2637560.75 | 3099303 | 9606035 | 8795315 | 7381280 | 8336462 | 7777409 | 3748214.75 | 9723933 | 7177481 | 4100047 |
| Q8VC03 | Eml3 | Echinoderm microtubule-associated protein-like 3 Mus musculus | 41.79 | 1.78 | 95695 | 1 | 1 | 1 | 1 | 320697.59 | 1 | 275340.22 | 306950.84 | 311770.94 | 1 | 1 | 342507.41 | 1 | 355695.69 | 396763.53 | 1 |
| Q3UMY5 | Eml4 | Echinoderm microtubule-associated protein-like 4 Mus musculus | 82.13 | 1.62 | 110027 | 1 | 1 | 1 | 1 | 1 | 1 | 1 | 733450.0475 | 834772.44 | 603441.25 | 692215.75 | 803370.75 | 600970.06 | 620397.03 | 799164.06 | 461056.97 |
| Q8BX80 | Engase | Cytosolic endo-beta-N-acetylglucosaminidase Mus musculus | 38.33 | 2.45 | 82945 | 1 | 1 | 630382.69 | 406771.66 | 582075.31 | 577889.12 | 318679.78 | 379506.34 | 406837.78 | 395422 | 479089.28 | 373333.5 | 491443.75 | 543001.81 | 328425.25 | 703515.19 |
| P17182 | Eno1 | Alpha-enolase Mus musculus | 2394.93 | 77.42 | 47141 | 24 | 32 | 483027841 | 544320321 | 569124609 | 442800673 | 497630657 | 576118465 | 683387137 | 669707137 | 704570817 | 678481153 | 568724865 | 618923585 | 630460033 | 581501569 |
| P17183 | Eno2 | Gamma-enolase Mus musculus | 645.04 | 23.73 | 47297 | 2 | 7 | 73958097 | 73393017 | 84455273 | 71131409 | 67966833 | 70267569 | 95670033 | 72616977 | 83336209 | 97182009 | 76567401 | 87854417 | 78859145 | 73257185 |
| P15429 | Eno3 | Beta-enolase Rattus norvegicus | 565.86 | 29.26 | 47014 | 3 | 7 | 47131237 | 45882161 | 56982809 | 44746569 | 45084993 | 45395793 | 61282017 | 45761993 | 51318829 | 61781561 | 50229401 | 58215209 | 51696865 | 46864777 |
| Q8BGB7 | Enoph1 | Enolase-phosphatase E1 Mus musculus | 101.97 | 5.45 | 28600 | 2 | 2 | 1674472 | 514303.97 | 1643080.12 | 1983792.75 | 2291924 | 1669732.75 | 1876117.38 | 1249551.5 | 1604725.5 | 1929748 | 692810.5 | 1565590.5 | 544458.69 | 1717635.88 |
| Q8BTJ4 | Enpp4 | Bis(5'-adenosyl)-triphosphatase enpp4 Mus musculus | 99.91 | 4.82 | 51611 | 2 | 2 | 985076.69 | 1300517.25 | 1556378.12 | 1119300 | 1032830.94 | 802762.88 | 979898.94 | 1244291 | 883281.62 | 971595.62 | 653590.19 | 660817.12 | 971313.44 | 843465.81 |
| Q5NDL0 | Eogt | EGF domain-specific O-linked N-acetylglucosamine transferase Rattus norvegicus | 98.17 | 3.42 | 61524 | 2 | 2 | 1099925 | 2217043 | 2163710.25 | 1962838.38 | 1994900.62 | 2163412.75 | 2102640 | 2346245.5 | 2073211.25 | 2009438.25 | 1897003 | 2220117 | 2187976 | 2141309.75 |
| P97481 | Epas1 | Endothelial PAS domain-containing protein 1 Mus musculus | 34.03 | 0.8 | 96712 | 1 | 1 | 2291132.25 | 1687931.88 | 1177006.25 | 2503481.5 | 2628611.75 | 1560509.25 | 917629.25 | 978144.69 | 618671.19 | 732402.69 | 2386942 | 2456274 | 831776.06 | 1641924.12 |
| P48193 | Epb41 | Protein 4.1 Mus musculus | 115.17 | 5.48 | 95911 | 3 | 3 | 2396828.5 | 1270234.38 | 2874250.5 | 3088874.75 | 3092124 | 3803812.5 | 3551729 | 2980751 | 3441233.5 | 3982010.5 | 3149823 | 3099586 | 3051549.75 | 3054908.5 |
| O70318 | Epb41l2 | Band 4.1-like protein 2 Mus musculus | 413.11 | 12.55 | 109940 | 8 | 8 | 7989059 | 4522140.5 | 5149627 | 7183709 | 9050957 | 10224838 | 5817005 | 7272232.5 | 7540088 | 10056795 | 6044065 | 5274418 | 7207497 | 6798329 |
| P58107 | EPPK1 | Epiplakin Homo sapiens OX=9606 | 213.95 | 2.95 | 555658 | 1 | 4 | 20477909 | 21831199 | 18154501 | 22418725 | 20974463 | 22075539 | 22491365 | 23942865 | 22022103 | 19743397 | 22085619 | 17984329 | 18837849 | 19708705 |
| Q8CGC7 | Eprs | Bifunctional glutamate/proline--tRNA ligase Mus musculus | 3129.3 | 44.97 | 170079 | 53 | 53 | 86093945 | 90823649 | 85946025 | 96304833 | 92856417 | 98712305 | 94255857 | 89438513 | 94415809 | 103358505 | 97153633 | 89921449 | 90498161 | 93605417 |
| P42567 | Eps15 | Epidermal growth factor receptor substrate 15 Mus musculus | 88.53 | 3.34 | 98471 | 2 | 2 | 781225.5 | 753682.5 | 836381.38 | 758778.5 | 593401 | 704979.5 | 913333.62 | 668046.62 | 656420.12 | 1021560.88 | 1132061.75 | 1599505.38 | 1033020.5 | 563309.38 |
| Q60902 | Eps15l1 | Epidermal growth factor receptor substrate 15-like 1 Mus musculus | 416.16 | 10.8 | 99309 | 8 | 8 | 1579996.25 | 631869.94 | 4871119.5 | 6067258.5 | 1889298.5 | 2648998.5 | 2018438.065 | 1259952.88 | 3585589.5 | 579211.38 | 3656547.75 | 13707442 | 982657.06 | 2452154 |
| Q08509 | Eps8 | Epidermal growth factor receptor kinase substrate 8 Mus musculus | 139.73 | 5.24 | 91737 | 2 | 2 | 1 | 1 | 1 | 1 | 1 | 1 | 1 | 326622.75 | 733055.25 | 561163.75 | 1 | 1 | 1 | 1 |
| Q9EQH2 | Erap1 | Endoplasmic reticulum aminopeptidase 1 Mus musculus | 946.18 | 26.34 | 106599 | 20 | 20 | 32018785 | 26761759 | 35973577 | 28325499 | 29424585 | 25553445 | 25330507 | 28156379 | 24026919 | 23602235 | 26816195 | 25693203 | 28954555 | 27258149 |
| Q80TH2 | Erbin | Erbin Mus musculus | 190.2 | 3.99 | 157248 | 3 | 3 | 1450685.75 | 1594659.12 | 1688859.75 | 1646393.25 | 1693545.5 | 1857364.38 | 1139402.75 | 1283240.5 | 1800873 | 1367196.75 | 1596150.62 | 1749348.88 | 1449968.75 | 1610416.75 |
| P07903 | Ercc1 | DNA excision repair protein ERCC-1 Mus musculus | 57.37 | 7.05 | 32910 | 1 | 1 | 1 | 1 | 1 | 101706.07 | 117672.57 | 1 | 96436.88 | 187887.98 | 142161.69 | 1 | 1 | 1 | 1 | 1 |
| Q8BHK9 | Ercc6l | DNA excision repair protein ERCC-6-like Mus musculus | 45.2 | 1.21 | 138854 | 1 | 1 | 221364.56 | 403636.19 | 266241.91 | 396379.38 | 415197.25 | 458503.75 | 311299.56 | 280930 | 321746.47 | 414287.28 | 398474.91 | 453389.31 | 589393 | 358525.53 |
| Q9ERY9 | Erg28 | Probable ergosterol biosynthetic protein 28 Mus musculus | 72.04 | 9.29 | 15806 | 1 | 1 | 666625.88 | 1048970.25 | 532570.44 | 500417.12 | 608880.25 | 600117 | 740018.12 | 565562.94 | 591261.12 | 515569.38 | 644031.94 | 942477.75 | 608361.25 | 653326.69 |
| Q9DC16 | Ergic1 | Endoplasmic reticulum-Golgi intermediate compartment protein 1 Mus musculus | 77.85 | 6.21 | 32562 | 2 | 2 | 3226603 | 3379903.5 | 3210803.25 | 3267271.5 | 3166848 | 3127740.75 | 2970197.5 | 3209533 | 3374236.75 | 3026255 | 2898832.25 | 3544579 | 3199977.25 | 3271404 |
| P84089 | Erh | Enhancer of rudimentary homolog Mus musculus | 247.72 | 45.19 | 12259 | 4 | 4 | 9151626 | 11230285 | 8897056 | 8986972 | 10046686 | 9178282 | 9146434 | 9627002 | 9092208 | 8741290 | 9320055 | 7393286 | 8282305.5 | 8958505 |
| Q7TMF2 | Eri1 | 3'-5' exoribonuclease 1 Mus musculus | 105.39 | 6.96 | 39493 | 2 | 2 | 3691419 | 4465231 | 4722021 | 4387705 | 4198916 | 4215196 | 4622320.5 | 4818858.5 | 4522628 | 4388523 | 4312755 | 4977518 | 4243759 | 4796125 |
| Q8BFZ9 | Erlin2 | Erlin-2 Mus musculus | 206.7 | 17.65 | 37873 | 5 | 5 | 1397666 | 1118907 | 1771453.12 | 1481336.12 | 724465.38 | 795913.06 | 1183217 | 723069.62 | 1640517.5 | 1002377.12 | 972311.69 | 1554360.75 | 1106878.25 | 839299.88 |
| Q3UVK0 | Ermp1 | Endoplasmic reticulum metallopeptidase 1 Mus musculus | 81.45 | 3.67 | 100148 | 2 | 2 | 177502.88 | 426273.2925 | 204563.97 | 918256.38 | 404769.94 | 150623.81 | 320287.34 | 216473.91 | 104187.24 | 290797.25 | 360757.59 | 151223.61 | 71247.48 | 272480.31 |
| Q8R180 | Ero1a | ERO1-like protein alpha Mus musculus | 1208.87 | 53.66 | 54084 | 17 | 19 | 34275897 | 37585565 | 33318769 | 30588371 | 32604637 | 52807873 | 55697817 | 67111121 | 56419889 | 53866457 | 31954029 | 34456857 | 36277085 | 37950649 |
| Q8R2E9 | Ero1b | ERO1-like protein beta Mus musculus | 314.36 | 12.85 | 53518 | 3 | 5 | 6508726 | 6918732 | 7956128 | 5295514.5 | 5025839.5 | 8813798 | 11036895 | 12714955 | 10234134 | 8538276 | 5906597.5 | 6736327.5 | 7119441 | 7604306.5 |
| P57759 | Erp29 | Endoplasmic reticulum resident protein 29 Mus musculus | 421.21 | 42.75 | 28823 | 7 | 7 | 23509187 | 25599647 | 25032001 | 19475835 | 23330189 | 17848159 | 18310825 | 21764247 | 19237601 | 17908189 | 21789521 | 23503477 | 26228381 | 23041821 |
| Q9D1Q6 | Erp44 | Endoplasmic reticulum resident protein 44 Mus musculus | 170.88 | 12.32 | 46853 | 4 | 4 | 759549.19 | 1013525.75 | 4785617.5 | 711205.12 | 1054735.25 | 2993364.83 | 1027213.88 | 5911436 | 758714.94 | 4276094.5 | 3312329.5 | 827444.12 | 1028358.88 | 4774724 |
| Q9R0P3 | Esd | S-formylglutathione hydrolase Mus musculus | 964.57 | 64.89 | 31320 | 13 | 14 | 83114041 | 96321745 | 83802617 | 68926065 | 83589385 | 76685665 | 78710913 | 84664905 | 80836249 | 84889969 | 86825473 | 87450065 | 90954209 | 90397833 |
| Q3V1V3 | Esf1 | ESF1 homolog Mus musculus | 43.06 | 1.42 | 98048 | 1 | 1 | 295701.62 | 221579.8 | 296168.06 | 246615.015 | 173010.58 | 291382.16 | 270064.655 | 295835.38 | 246984.55 | 246056.53 | 164955.7 | 211752.34 | 387348.16 | 222384.28 |
| P60330 | Espl1 | Separin Mus musculus | 109.06 | 1.56 | 233035 | 3 | 3 | 392002.09 | 188101.06 | 4667197.5 | 7043474.5 | 270140.88 | 413096.31 | 2396179.64 | 3456689.5 | 3165768.25 | 2549164.5 | 254261.72 | 471697.19 | 133556.61 | 249839.03 |
| Q3U7R1 | Esyt1 | Extended synaptotagmin-1 Mus musculus | 848.32 | 17.12 | 121554 | 14 | 14 | 10172406 | 12263189 | 11941009 | 13940559 | 12000911 | 13769315 | 10060706 | 11993031 | 13388635 | 10736338 | 10997511 | 12836045 | 12093596 | 11731764 |
| Q3TZZ7 | Esyt2 | Extended synaptotagmin-2 Mus musculus | 119.37 | 4.5 | 94139 | 2 | 2 | 1064953.5 | 1177318.88 | 1586444.38 | 1074387.38 | 1557714.5 | 1733379.88 | 2089133.5 | 1733767.88 | 1960167.12 | 1562902 | 2275910 | 1470853.25 | 1649729.75 | 1518644.25 |
| Q5U2Q7 | Etf1 | Eukaryotic peptide chain release factor subunit 1 Rattus norvegicus | 1018.21 | 59.5 | 49031 | 18 | 18 | 16706434 | 18603379 | 22570753 | 18915315 | 16865905 | 20350831 | 22217527 | 24730837 | 22900761 | 19328585 | 20875921 | 23177161 | 22262429 | 20515099 |
| Q99LC5 | Etfa | Electron transfer flavoprotein subunit alpha, mitochondrial Mus musculus | 782.9 | 54.65 | 35009 | 13 | 13 | 57741413 | 66087065 | 70525881 | 65579277 | 65035497 | 64453841 | 73952921 | 69655345 | 71553889 | 67138097 | 64588113 | 69222497 | 64620221 | 69280969 |
| Q68FU3 | Etfb | Electron transfer flavoprotein subunit beta Rattus norvegicus | 567.95 | 36.86 | 27687 | 1 | 11 | 47981317 | 51292781 | 52961961 | 57274581 | 57576025 | 53512033 | 56423821 | 49054713 | 55791173 | 54468653 | 54013013 | 56676909 | 53942745 | 55213085 |
| Q9DCM0 | Ethe1 | Persulfide dioxygenase ETHE1, mitochondrial Mus musculus | 166.01 | 24.41 | 27739 | 4 | 4 | 8303762.5 | 9284230 | 9074427 | 10790023 | 10753311 | 8386768 | 8354889 | 7501611 | 7327678.5 | 8757601 | 10588234 | 8474962 | 7211660 | 9932822 |
| P97360 | Etv6 | Transcription factor ETV6 Mus musculus | 149.74 | 7.84 | 56406 | 3 | 3 | 1 | 119053.66 | 201921.03 | 1 | 1 | 1 | 106952.92 | 287862.81 | 1 | 1 | 201754.53 | 241476.12 | 1 | 1 |
| Q8VD58 | Evi2b | Protein EVI2B Mus musculus | 39.26 | 2.48 | 47993 | 1 | 1 | 3684902.25 | 2832548.5 | 2309091.5 | 1882541 | 2129549.25 | 1573820.5 | 2069914.88 | 2132688.75 | 1717356.12 | 1615230.62 | 1113547.25 | 1533057.38 | 1955776.75 | 1665572.12 |
| Q61545 | Ewsr1 | RNA-binding protein EWS Mus musculus | 292.15 | 12.21 | 68462 | 6 | 6 | 9790247 | 11386937 | 9336300 | 11701854 | 11464452 | 8151007 | 13421140 | 9046718 | 10774652 | 12048937 | 5276147 | 8332456 | 8746660 | 8851677 |
| Q9D4H1 | Exoc2 | Exocyst complex component 2 Mus musculus | 65.3 | 2.06 | 103959 | 2 | 2 | 1514566.12 | 1450794.75 | 1546134.88 | 1230913.12 | 1554094.12 | 1626518.12 | 1246017.62 | 1551404.62 | 1689082.62 | 1548564.5 | 1097117.38 | 1280030.25 | 1750300.62 | 1532413.62 |
| O35382 | Exoc4 | Exocyst complex component 4 Mus musculus | 290.01 | 6.77 | 110545 | 5 | 5 | 1848533.75 | 2479471 | 2428685 | 2457477 | 1848064.5 | 2152875.25 | 1620175 | 2772357.25 | 2630271 | 2016730.38 | 1754162 | 2230131 | 1835662.25 | 2454435.5 |
| Q3TPX4 | Exoc5 | Exocyst complex component 5 Mus musculus | 39.09 | 1.27 | 81738 | 1 | 1 | 1582412.25 | 1 | 1420503.88 | 415890.53 | 1 | 1316517.25 | 924563 | 1703718.12 | 1596158.75 | 978704.81 | 1025742.81 | 1539988.5 | 1143894.75 | 1195949.88 |
| O54923 | Exoc6 | Exocyst complex component 6 Rattus norvegicus | 100.95 | 2.49 | 93178 | 2 | 2 | 610581.38 | 603179.94 | 577174.06 | 650635.94 | 571622.25 | 720974.94 | 478189.81 | 545035.12 | 615718.69 | 628970.06 | 722428.25 | 452019.12 | 577933.44 | 753653.56 |
| A6H5Z3 | Exoc6b | Exocyst complex component 6B Mus musculus | 104.09 | 3.95 | 94130 | 2 | 2 | 650293.88 | 556235.75 | 483403 | 618612.25 | 364234.78 | 488429.72 | 485162.78 | 693119.5 | 536518.81 | 414602.69 | 465116.91 | 570431.81 | 536888.12 | 404184.25 |
| O54924 | Exoc8 | Exocyst complex component 8 Rattus norvegicus | 39.47 | 2.23 | 81043 | 1 | 1 | 1680718 | 2204113.75 | 1868724.5 | 1831553.12 | 1860770.5 | 1644193.25 | 1709266.38 | 1889605.25 | 1426325.75 | 1644810.62 | 1500142.88 | 1413343.25 | 1371372.75 | 1862619.25 |
| P56960 | Exosc10 | Exosome component 10 Mus musculus | 395.79 | 11.61 | 100942 | 8 | 8 | 3946711 | 5951535.5 | 5056039.5 | 4645242.5 | 5054771.5 | 5204211 | 5714009 | 3738156 | 3590203.75 | 4055049.5 | 4990687 | 3925689 | 4544430 | 4749634.5 |
| Q8VBV3 | Exosc2 | Exosome complex component RRP4 Mus musculus | 199.32 | 21.84 | 32632 | 4 | 4 | 2140369 | 2067023.62 | 1843501.5 | 4046490 | 2557314.25 | 3933297 | 1648760.5 | 4565743 | 1810482.5 | 4105612.5 | 2635769.25 | 1866453.75 | 4155499.5 | 1692696.38 |
| Q7TQK4 | Exosc3 | Exosome complex component RRP40 Mus musculus | 126.95 | 12.41 | 29546 | 2 | 2 | 1306568 | 1349430.5 | 2213209.5 | 1403089.38 | 1568074.345 | 1828918.5 | 1713696.12 | 1933426.5 | 2120763.25 | 1809852.88 | 1370991.38 | 1585925.12 | 1883722.5 | 2363399 |
| Q921I9 | Exosc4 | Exosome complex component RRP41 Mus musculus | 60.6 | 4.9 | 26250 | 1 | 1 | 1 | 1 | 1 | 1 | 354221.69 | 374872.72 | 289572.47 | 1 | 1 | 1 | 1 | 1 | 1 | 374704.25 |
| Q9CRA8 | Exosc5 | Exosome complex component RRP46 Mus musculus | 145.76 | 13.62 | 25194 | 3 | 3 | 4624671.5 | 5615735 | 5979488 | 4829062.5 | 4547397 | 4433066 | 5378759 | 4838787 | 5188973 | 4693448 | 4433340 | 6283818.5 | 5368849 | 4811139 |
| Q9D0M0 | Exosc7 | Exosome complex exonuclease RRP42 Mus musculus | 187.06 | 19.24 | 31825 | 4 | 4 | 2964501.75 | 3029343.5 | 2762999.5 | 2791264 | 1334217 | 2791397.75 | 3016215 | 2759473 | 2911419 | 2609495.5 | 2533583 | 2963200.5 | 2551283.5 | 2745490 |
| Q9D753 | Exosc8 | Exosome complex component RRP43 Mus musculus | 213.44 | 24.28 | 29949 | 4 | 4 | 2567726 | 4718768 | 5453596.5 | 5592656.5 | 5231408.5 | 5238204.5 | 6015746 | 5603923 | 5966166 | 4072091 | 5153995 | 5851679 | 4623779.5 | 5904373.5 |
| Q9JHI7 | Exosc9 | Exosome complex component RRP45 Mus musculus | 282.82 | 13.47 | 48937 | 4 | 4 | 4550806 | 6657117 | 4695681 | 6797694 | 5475161 | 4327291 | 4604538 | 4292619 | 6389767 | 5550483.5 | 4833677 | 6841462 | 5469455 | 6748772 |
| P26040 | Ezr | Ezrin Mus musculus | 1125.14 | 34.13 | 69407 | 9 | 21 | 422243009 | 401172545 | 355826241 | 385307809 | 379793089 | 426534657 | 413574465 | 429468193 | 404784993 | 398752097 | 397178113 | 352466209 | 331874145 | 361883969 |
| P04117 | Fabp4 | Fatty acid-binding protein, adipocyte Mus musculus | 270.09 | 51.52 | 14650 | 6 | 6 | 14000153 | 14500083 | 15950010 | 6045960 | 11049351 | 6664707 | 11067365 | 13879454 | 11125036 | 10443999 | 6024804.5 | 6746711 | 17115957 | 12005427 |
| Q05816 | Fabp5 | Fatty acid-binding protein 5 Mus musculus | 1005.52 | 79.26 | 15137 | 13 | 16 | 367597665 | 362926977 | 390861889 | 341166817 | 309307073 | 348621505 | 401450913 | 399667425 | 376506625 | 378119297 | 395330977 | 398830881 | 434537025 | 457692673 |
| Q61160 | Fadd | FAS-associated death domain protein Mus musculus | 155.35 | 17.56 | 22960 | 2 | 2 | 948964.56 | 794998.44 | 736284 | 765240.405 | 580714.62 | 615911.25 | 716612.81 | 712067.12 | 650559.31 | 536337.31 | 661390 | 571219 | 651086.88 | 584658.81 |
| Q920L1 | Fads1 | Fatty acid desaturase 1 Mus musculus | 88.65 | 6.26 | 52323 | 2 | 2 | 535070.75 | 1 | 1 | 1 | 1 | 1 | 1 | 1 | 168692.39 | 1 | 179718.2667 | 170185.67 | 189785.91 | 179183.22 |
| Q9Z0R9 | Fads2 | Fatty acid desaturase 2 Mus musculus | 123.95 | 6.76 | 52387 | 2 | 3 | 392607.81 | 1 | 371367.59 | 1 | 1 | 1 | 1 | 1 | 1 | 1 | 1 | 1 | 351027.22 | 1 |
| P54731 | Faf1 | FAS-associated factor 1 Mus musculus | 138.61 | 5.7 | 73863 | 3 | 3 | 1324567.75 | 794858.38 | 1910030 | 1 | 661915.69 | 1149229.62 | 1207180.38 | 2080602.5 | 1898244 | 2061012.12 | 1789786.62 | 1335186.88 | 1405671.38 | 1663771 |
| Q3TDN2 | Faf2 | FAS-associated factor 2 Mus musculus | 150.93 | 12.58 | 52471 | 3 | 3 | 1774138.75 | 1 | 1315747.25 | 1 | 1 | 1296483.5 | 1591352.12 | 1146010.62 | 1034661.62 | 1562835 | 1590334.75 | 1379546.25 | 1653225.38 | 1738232.62 |
| P25093 | Fah | Fumarylacetoacetase Rattus norvegicus | 142.45 | 14.8 | 45976 | 3 | 3 | 633424.75 | 1208979.5 | 1571140 | 1002720 | 718923.38 | 639873.25 | 1497314.38 | 1 | 1092950.5 | 1026018.62 | 924111.44 | 947091.75 | 1950566.5 | 634198.25 |
| Q9WUD8 | Faim | Fas apoptotic inhibitory molecule 1 Mus musculus | 108.3 | 13.97 | 20201 | 2 | 2 | 1509374.88 | 1934835.75 | 721436.44 | 948193.38 | 1248292.75 | 2418244.75 | 626177.75 | 2374215.75 | 2200284.25 | 1 | 1114634.75 | 2087034.88 | 1009685.75 | 2235166 |
| Q8VE88 | Fam114a2 | Protein FAM114A2 Mus musculus | 121.26 | 6.04 | 54045 | 2 | 2 | 701795.5 | 651817.5 | 774680.12 | 421943 | 1631702.25 | 1582904 | 1059918 | 1837207.38 | 1889494.12 | 1382197.12 | 1383175.5 | 813203.12 | 1492156.25 | 456807.91 |
| Q6A0A9 | FAM120A | Constitutive coactivator of PPAR-gamma-like protein 1 Mus musculus | 274.76 | 6.65 | 121646 | 6 | 6 | 3361158 | 3630206 | 3036079.75 | 1367072.12 | 3035716 | 1518155 | 3425002.5 | 4933171 | 4631203 | 1908938.75 | 1706037.12 | 1033363.5 | 3538329 | 2766903.5 |
| Q3UW53 | Fam129a | Protein Niban Mus musculus | 1269.04 | 33.26 | 102649 | 23 | 23 | 37346441 | 32697601 | 38217685 | 38209249 | 31493591 | 34300965 | 34329389 | 33928057 | 32412151 | 37348917 | 34571681 | 35827813 | 34872381 | 31817665 |
| Q8R1F1 | Fam129b | Niban-like protein 1 Mus musculus | 513.98 | 19.36 | 84819 | 9 | 9 | 21481605 | 20756937 | 21639297 | 18641639 | 18686853 | 24123223 | 23228745 | 24436909 | 23561257 | 21288829 | 22190621 | 23111867 | 22226109 | 22595651 |
| Q4QQV3 | Fam162a | Protein FAM162A Rattus norvegicus | 35.31 | 4.52 | 17827 | 1 | 1 | 421356.06 | 463096.21 | 430423.62 | 530424.94 | 470180.22 | 1 | 398820.31 | 1 | 402747.31 | 484585.88 | 459039.55 | 512958.28 | 511555.31 | 352605.06 |
| Q91WE2 | Fam192a | PSME3-interacting protein Mus musculus | 49.63 | 8.66 | 28702 | 2 | 2 | 1037927.298 | 1135491.5 | 760348.69 | 1142106.25 | 1113762.75 | 959999 | 815611.12 | 1372714.25 | 753290.19 | 683800.38 | 1097130.75 | 1013449.19 | 906164.56 | 791727.62 |
| Q8CGI1 | Fam193a | Protein FAM193A Mus musculus | 35.38 | 0.81 | 136714 | 1 | 1 | 4412990 | 2779078.25 | 2425787 | 4287065.5 | 4312787 | 1263273.88 | 2120620.75 | 1657398.75 | 1911461.62 | 1650282.5 | 4412990 | 4386696.5 | 762691.62 | 3370027.5 |
| Q91VU0 | Fam3c | Protein FAM3C Mus musculus | 272.59 | 24.23 | 24753 | 4 | 4 | 3161076.75 | 3497232.25 | 3286509.25 | 3563666.75 | 3167611.75 | 3315723.5 | 3166684 | 3555858.25 | 3183598.5 | 3615402.75 | 3611576.75 | 3578932.5 | 3703299.75 | 2757006.25 |
| Q8BHZ0 | Fam49a | Protein FAM49A Mus musculus | 265.66 | 15.79 | 37343 | 2 | 4 | 20813073 | 23976357 | 20775189 | 20002189 | 23508867 | 22837621 | 23624373 | 23128407 | 23061965 | 21181123 | 19346061 | 20163679 | 19929849 | 20930437 |
| Q921M7 | Fam49b | Protein FAM49B Mus musculus | 1135.29 | 73.46 | 36776 | 17 | 19 | 52385545 | 53019641 | 50129689 | 44368365 | 50591877 | 47774805 | 49030085 | 51663705 | 50546941 | 50851881 | 45876473 | 46069557 | 48505465 | 48084657 |
| Q9WV03 | Fam50a | Protein FAM50A Mus musculus | 96.5 | 7.96 | 40252 | 2 | 2 | 1302013.25 | 465316.88 | 1157272.75 | 1587247.62 | 536215.12 | 1330067.62 | 1338158.25 | 1069227.12 | 1410019.25 | 1675246.25 | 1548724.75 | 1409831.25 | 1351041 | 786166.62 |
| Q3TJZ6 | Fam98a | Protein FAM98A Mus musculus | 109.07 | 10.1 | 55055 | 2 | 3 | 1 | 323133.53 | 773351.38 | 482654.88 | 1 | 602756.12 | 407554.97 | 508812.94 | 585000.44 | 203598.17 | 1 | 1 | 178675.27 | 344186.12 |
| Q80VD1 | Fam98b | Protein FAM98B Mus musculus | 231.79 | 15.85 | 45349 | 4 | 5 | 2202856.75 | 6076433 | 6850369 | 2981043 | 4320574 | 3049656 | 4958395 | 6938220 | 3002911.5 | 5061146 | 6718394 | 4540014 | 7013997 | 2835989 |

| | | | | | | | | | | | | | | | | | | | | | |
|---|---|---|---|---|---|---|---|---|---|---|---|---|---|---|---|---|---|---|---|---|---|
| Q922J9 | Far1 | Fatty acyl-CoA reductase 1 Mus musculus | 493.86 | 24.27 | 59435 | 9 | 9 | 18782985 | 12180763 | 12503131 | 11126202 | 14845694 | 13756003 | 10952911 | 13167841 | 12700844 | 11446783 | 13400021 | 13181665 | 13037003 | 13741521 |
| Q8C0C7 | Farsa | Phenylalanine--tRNA ligase alpha subunit Mus musculus | 594.17 | 23.03 | 57599 | 4 | 11 | 7553203 | 9834617 | 6850016 | 8593985 | 9612251 | 7256307 | 6428624.5 | 8554877 | 7658390.5 | 6570739.5 | 7797626 | 7719867 | 6511081.5 | 8144221.5 |
| Q9WUA2 | Farsb | Phenylalanine--tRNA ligase beta subunit Mus musculus | 868.51 | 39.56 | 65697 | 20 | 20 | 38717701 | 36137033 | 39857009 | 33997365 | 34118893 | 37396457 | 34323801 | 41123157 | 38708153 | 35511713 | 36598293 | 40960133 | 39565529 | 37031721 |
| P19096 | Fasn | Fatty acid synthase Mus musculus | 6246.8 | 61.86 | 272428 | 51 | 103 | 346178113 | 337950113 | 356189569 | 357735041 | 323721473 | 334791361 | 342689793 | 341150145 | 336258529 | 328394305 | 341145313 | 333249153 | 326903649 | 341676769 |
| P62862 | Fau | 40S ribosomal protein S30 Mus musculus | 94.78 | 18.64 | 6648 | 2 | 2 | 2374680.75 | 2146666.25 | 4830429 | 3816281.25 | 4727740 | 6375753 | 4694967 | 1848487.12 | 5831644 | 8275803 | 6668771 | 8601862 | 8358926.5 | 7226102 |
| P22509 | Fbl | rRNA 2'-O-methyltransferase fibrillarin Rattus norvegicus | 758.15 | 51.68 | 34222 | 9 | 13 | 11956617 | 12822240 | 16228307 | 13784236 | 12760894 | 12311932 | 16126875 | 12045175 | 13876513 | 16119087 | 13994292 | 12999887 | 15669475 | 14062370 |
| Q80WS3 | Fbll1 | rRNA/tRNA 2'-O-methyltransferase fibrillarin-like protein 1 Mus musculus | 193.75 | 11.46 | 33339 | 1 | 3 | 1879900.25 | 1368595.75 | 2122934.75 | 1937742.38 | 1605991 | 1606010.12 | 1818624.12 | 1264604.12 | 1564863.62 | 2681657 | 1732267 | 1963356.12 | 2219345 | 1995010 |
| Q9CZV8 | Fbxl20 | F-box/LRR-repeat protein 20 Mus musculus | 134.55 | 4.59 | 48396 | 2 | 2 | 1667723.12 | 1688105 | 1393354.75 | 845205.38 | 1029334.38 | 950312.94 | 1096521.88 | 1601200.88 | 1318204 | 1558557.12 | 1576620.88 | 976424.81 | 1096774.38 | 1003757.44 |
| Q78JE5 | Fbxo22 | F-box only protein 22 Mus musculus | 226.06 | 24.38 | 44203 | 5 | 5 | 5727194 | 6290344.5 | 3759569.75 | 5147191.5 | 5312682.5 | 3761430.25 | 3551218.25 | 5278593.5 | 5425252 | 3035843.25 | 3551516.5 | 3724194 | 3564878.75 | 5253938 |
| Q9QZN4 | Fbxo6 | F-box only protein 6 Mus musculus | 34.09 | 3.39 | 34493 | 1 | 1 | 323568.19 | 250572.7 | 270163.19 | 281705.925 | 282519.62 | 245961.03 | 306078.76 | 357342.31 | 416691.25 | 204320.45 | 197213.44 | 303465.59 | 239411.19 | 246696.74 |
| P20491 | Fcer1g | High affinity immunoglobulin epsilon receptor subunit gamma Mus musculus | 102.69 | 15.12 | 9652 | 2 | 2 | 3695508.25 | 2696233.25 | 4694493.5 | 3697796.5 | 5004032.5 | 3881418.25 | 353448.69 | 1504566.88 | 2975626.5 | 5032781 | 3263798.75 | 3551863.75 | 3599762 | 4626755 |
| P26151 | Fcgr1 | High affinity immunoglobulin gamma Fc receptor I Mus musculus | 275.57 | 22.77 | 44888 | 6 | 6 | 10544918 | 9299983 | 11888508 | 9437379 | 9035087 | 7037936 | 6074016 | 6070993.5 | 7317976 | 5974343 | 6363764 | 6071877.5 | 7110427 | 4110293.75 |
| P08101 | Fcgr2 | Low affinity immunoglobulin gamma Fc region receptor II Mus musculus | 176.94 | 11.21 | 36695 | 2 | 3 | 10047039 | 11144037 | 9987515 | 11168614 | 10237842 | 9170796 | 10371663 | 8652388 | 7745995 | 10653992 | 9368193 | 7853520 | 9759479 | 8793518 |
| P08508 | Fcgr3 | Low affinity immunoglobulin gamma Fc region receptor III Mus musculus | 190.29 | 16.09 | 30036 | 2 | 3 | 8633707 | 10154023 | 8342685.5 | 11564095 | 10968568 | 9087691 | 8170459 | 8212834.5 | 8077483.5 | 10018629 | 9749846 | 8987717 | 8910594 | 7707230 |
| A0A0B4 | Fcgr4 | Low affinity immunoglobulin gamma Fc region receptor IV Mus musculus | 45.23 | 4.02 | 28398 | 1 | 1 | 2495273 | 2113805.5 | 2117607.5 | 2473083 | 2382982.25 | 1894300.5 | 1841483.38 | 2000441.75 | 1741781.12 | 2120373.5 | 1889272.12 | 1783561.25 | 1607984.75 | 1968628.62 |
| D3ZYR1 | Fcho2 | F-BAR domain only protein 2 Rattus norvegicus | 115.95 | 3.83 | 88712 | 2 | 2 | 830894.94 | 883108.62 | 884735.75 | 744802.12 | 843027.44 | 1208799.88 | 1217656.62 | 985770.38 | 1326516 | 1219684.62 | 1048347.88 | 729531.06 | 865548 | 930317.62 |
| P53798 | Fdft1 | Squalene synthase Mus musculus | 184 | 9.86 | 48154 | 3 | 3 | 221825.56 | 1 | 1 | 316029.34 | 1 | 1 | 1 | 1 | 1 | 1 | 320486.91 | 305460.03 | 744044.69 | 218683.61 |
| Q920E5 | Fdps | Farnesyl pyrophosphate synthase Mus musculus | 1360.53 | 65.44 | 40582 | 20 | 21 | 184978865 | 157284449 | 182328657 | 169005761 | 156573297 | 174010833 | 170506001 | 184441761 | 179312577 | 167460673 | 183938529 | 163477473 | 179663937 | 176954561 |
| Q61578 | Fdxr | NADPH:adrenodoxin oxidoreductase, mitochondrial Mus musculus | 42.92 | 2.43 | 54202 | 1 | 1 | 1531694.25 | 1348532.75 | 1561898.88 | 1238214 | 2042713 | 1375772.88 | 1520341.5 | 1436242.38 | 1559145.5 | 1527084.88 | 1688709.75 | 1164738.88 | 1367383.5 | 1361577.12 |
| P39749 | Fen1 | Flap endonuclease 1 Mus musculus | 507.06 | 29.1 | 42315 | 9 | 9 | 20376619 | 16223644 | 14952076 | 17508535 | 15983569 | 15778612 | 16351187 | 18301489 | 16101388 | 15245840 | 15378574 | 16238940 | 15837186 | 12123189 |
| Q8K1B8 | Fermt3 | Fermitin family homolog 3 Mus musculus | 1846.33 | 60.15 | 75635 | 29 | 29 | 128422081 | 140474417 | 129359985 | 123999265 | 132227425 | 132468561 | 124299937 | 148843361 | 132593089 | 123791369 | 123178473 | 127113817 | 126923817 | 133930049 |
| P16879 | Fes | Tyrosine-protein kinase Fes/Fps Mus musculus | 35.95 | 1.46 | 93779 | 1 | 1 | 2004771.88 | 1651227.75 | 1695138.62 | 1868781.12 | 1642265.75 | 1624663 | 1466674 | 1680877.12 | 1401896.5 | 1825407.38 | 1835718.38 | 1303219.12 | 1639539.38 | 1633854.5 |
| O88842 | Fgd3 | FYVE, RhoGEF and PH domain-containing protein 3 Mus musculus | 113.28 | 4.09 | 80624 | 2 | 2 | 373792.97 | 382110 | 424591.03 | 406523.38 | 442048.72 | 318586.56 | 291633.72 | 543437.31 | 420460.88 | 420062.44 | 358986.78 | 307629.78 | 300141.69 | 425776.28 |
| P97807 | Fh | Fumarate hydratase, mitochondrial Mus musculus | 1200.89 | 50.3 | 54357 | 17 | 19 | 86103625 | 83282137 | 88677953 | 80313417 | 82207745 | 83393489 | 93351513 | 92565545 | 88980857 | 86502265 | 83627905 | 81766081 | 83369281 | 83069145 |
| Q9R059 | Fhl3 | Four and a half LIM domains protein 3 Mus musculus | 227.87 | 19.03 | 31795 | 4 | 4 | 6045332.5 | 10425108 | 4754440 | 8569628 | 8681190 | 8032475 | 8193082 | 4440852 | 8164092.5 | 9959419 | 8693954 | 9350019 | 7571937 | 8923509 |
| Q6P9Q4 | Fhod1 | FH1/FH2 domain-containing protein 1 Mus musculus | 213.66 | 6.43 | 129600 | 5 | 5 | 3240982 | 2549733 | 2016947.75 | 2310462 | 2904079.5 | 2425104 | 2784137.5 | 3550233.5 | 3265756.75 | 3011047.5 | 2758289 | 841391.19 | 2407205.5 | 3175905.25 |
| Q5U317 | Fip1l1 | Pre-mRNA 3'-end-processing factor FIP1 Rattus norvegicus | 224.01 | 13.62 | 60188 | 5 | 5 | 6427216 | 425029.22 | 277373.94 | 324814.94 | 306673.88 | 1242730.62 | 1523347.75 | 776983.06 | 1418247.75 | 1035314.62 | 1814855.38 | 373343.19 | 1894319.75 | 494424.03 |
| P84817 | Fis1 | Mitochondrial fission 1 protein Rattus norvegicus | 381.05 | 60.53 | 16995 | 6 | 6 | 14938037 | 9948465 | 9113640 | 8535140 | 8707892 | 6092679.5 | 6198266.5 | 6328685 | 7185594.5 | 7633101 | 8499026 | 6001722 | 9117733 | 6506754 |
| Q6P9Q6 | Fkbp15 | FK506-binding protein 15 Mus musculus | 71.65 | 0.9 | 132961 | 1 | 1 | 850937.69 | 875626.44 | 835598.31 | 831313.94 | 1455444.62 | 904057.94 | 790797.25 | 655812.75 | 707830.06 | 876486.25 | 966557.19 | 676452.38 | 671755.75 | 1140923.25 |
| P26883 | Fkbp1a | Peptidyl-prolyl cis-trans isomerase FKBP1A Mus musculus | 89.35 | 12.96 | 11923 | 1 | 1 | 3188589.5 | 2845916.5 | 2171562.25 | 3069237.5 | 3164370 | 2722950.5 | 3630331 | 3018585 | 2669283.75 | 3310789 | 3087272.5 | 3375607.25 | 3358400.75 | 2460506.25 |
| P45878 | Fkbp2 | Peptidyl-prolyl cis-trans isomerase FKBP2 Mus musculus | 125.57 | 19.29 | 15344 | 2 | 2 | 3244866 | 3025755.5 | 3779809.5 | 4243410.5 | 2996151 | 2476533.25 | 2402041.25 | 2739740 | 1806613.25 | 2899214 | 3283477.75 | 2750951 | 2478876.5 | 2652518.5 |
| Q62446 | Fkbp3 | Peptidyl-prolyl cis-trans isomerase FKBP3 Mus musculus | 46.87 | 4.91 | 25148 | 1 | 1 | 1944479.88 | 1016109.56 | 1316950 | 1529155.88 | 1503591.5 | 1288249.12 | 1407481.5 | 1059804.38 | 1144764.62 | 1400069.62 | 1248974.12 | 1333283 | 1336360 | 1066898.62 |
| P30416 | Fkbp4 | Peptidyl-prolyl cis-trans isomerase FKBP4 Mus musculus | 1578 | 71.4 | 51572 | 27 | 27 | 63051041 | 57052389 | 60547837 | 56181509 | 58160157 | 55564913 | 54880121 | 59651421 | 55996949 | 57397429 | 63283181 | 59269093 | 57129337 | 62024329 |
| Q64378 | Fkbp5 | Peptidyl-prolyl cis-trans isomerase FKBP5 Mus musculus | 386.09 | 20.18 | 50966 | 8 | 8 | 6534787 | 8083692 | 9353432 | 8812209 | 9785943 | 9097383 | 9412392 | 7674144 | 9181603 | 10015511 | 9368324 | 9453704 | 9960211 | 9941976 |
| O35465 | Fkbp8 | Peptidyl-prolyl cis-trans isomerase FKBP8 Mus musculus | 522.65 | 29.1 | 43529 | 8 | 8 | 13277937 | 14322264 | 15320881 | 9608378 | 11283122 | 14168371 | 16037043 | 13530283 | 14473254 | 14097243 | 14300108 | 10406040 | 16319420 | 15490078 |
| Q8QZS3 | Flcn | Folliculin Mus musculus | 43.05 | 2.25 | 64327 | 1 | 1 | 235372.88 | 227347.08 | 1 | 1 | 210333.64 | 210600.2225 | 194717.5 | 227220.34 | 211677.02 | 208786.03 | 208366.98 | 194942.31 | 190943.08 | 253639.44 |
| Q9JJ28 | Flii | Protein flightless-1 homolog Mus musculus | 1236.46 | 25.96 | 144803 | 23 | 23 | 33758693 | 35097469 | 24368847 | 33923405 | 33465589 | 35324701 | 30569741 | 34900745 | 29308523 | 37593837 | 32688033 | 35832645 | 33795161 | 31659823 |
| Q8BTM8 | Flna | Filamin-A Mus musculus | 5048.24 | 49.87 | 281222 | 80 | 84 | 369365921 | 351985377 | 418238977 | 354114593 | 340416193 | 299507617 | 332472097 | 274723489 | 279353921 | 326676577 | 278621537 | 289579969 | 383951745 | 319704929 |
| Q80X90 | Flnb | Filamin-B Mus musculus | 556.33 | 5.88 | 277825 | 8 | 11 | 16415002 | 14864654 | 17498361 | 15034736 | 14546203 | 12402659 | 10174523 | 11341972 | 11000152 | 11902811 | 13906003 | 14000077 | 19377567 | 14977597 |
| Q60634 | Flot2 | Flotillin-2 Mus musculus | 245.6 | 10.28 | 47038 | 4 | 4 | 765938.69 | 1848567.12 | 2336999.75 | 1 | 1162812.25 | 1660661.62 | 2537925 | 2745098 | 2613886 | 2558559.25 | 1986047.88 | 2485231.5 | 1786466.38 | 1669692.38 |
| Q9JL26 | Fmnl1 | Formin-like protein 1 Mus musculus | 1753.47 | 41.04 | 122060 | 30 | 35 | 57177841 | 58380137 | 60175849 | 68217953 | 54049305 | 55786417 | 54548093 | 53782877 | 55511413 | 63105941 | 51195693 | 58224305 | 49162477 | 53427037 |
| A2APV2 | Fmnl2 | Formin-like protein 2 Mus musculus | 296.73 | 6.45 | 123101 | 0 | 7 | 8938427 | 9728446 | 8357934 | 9296369 | 7466358.5 | 6589770 | 8536496 | 8120341 | 6439241 | 9245420 | 8237426.5 | 7834973.5 | 6339505.5 | 7543199.5 |
| Q6ZPF4 | Fmnl3 | Formin-like protein 3 Mus musculus | 324.93 | 8.27 | 117169 | 2 | 8 | 4148227 | 4034978.5 | 2735316 | 3048927 | 3106976 | 2294376 | 2911728 | 2923270 | 2187319.25 | 4158158.5 | 3683141.75 | 3057034.5 | 2291198.75 | 3061146 |
| P35922 | Fmr1 | Synaptic functional regulator FMR1 Mus musculus | 128.12 | 3.58 | 68989 | 1 | 2 | 8126656.5 | 8517745 | 8133283 | 8277533 | 8584711 | 8328885 | 8368202 | 9118463 | 8137485 | 8543702 | 8434421 | 9070251 | 8260224 | 8660171 |
| Q80TY0 | Fnbp1 | Formin-binding protein 1 Mus musculus | 797.37 | 28.57 | 71344 | 16 | 16 | 21962303 | 26344899 | 22767065 | 27794309 | 26876757 | 26968981 | 28623121 | 29073977 | 27696497 | 28881731 | 23275953 | 23090863 | 21758103 | 24006653 |
| Q2HWF0 | Fnbp1l | Formin-binding protein 1-like Rattus norvegicus | 101.73 | 3.47 | 69974 | 1 | 1 | 250212.36 | 1 | 169275.83 | 1 | 157250.34 | 1 | 1 | 165152.89 | 1 | 1 | 1 | 248328.72 | 206494.09 | 1 |
| Q6NWW9 | Fndc3b | Fibronectin type III domain-containing protein 3B Mus musculus | 137.45 | 2.57 | 132764 | 2 | 2 | 239133.61 | 324202 | 728307.06 | 279746.22 | 794567 | 249004.48 | 721106.88 | 740445 | 1035397.75 | 695165.56 | 217168.22 | 208674.19 | 271787.34 | 666677.81 |
| Q8K385 | FRRS1 | Ferric-chelate reductase 1 Mus musculus | 280.32 | 16.05 | 66047 | 5 | 5 | 7680971 | 8136465 | 7587437.5 | 6404462 | 7100279.5 | 7477670 | 6735212 | 8264281 | 6976210 | 6009498 | 5919187 | 7494734 | 7988006.5 | 6013706 |
| A2ARZ3 | Fsip2 | Fibrous sheath-interacting protein 2 Mus musculus | 40.85 | 0.1 | 784857 | 1 | 1 | 2031017 | 1698953.5 | 2305389.75 | 2006768.25 | 2115784.25 | 2106571.5 | 1964987.75 | 1952154.75 | 1894555 | 2371351.5 | 2105113 | 2308895 | 2152015 | 2153232.25 |
| P09528 | Fth1 | Ferritin heavy chain Mus musculus | 284.6 | 49.45 | 21067 | 7 | 7 | 30465959 | 35178017 | 27363913 | 23865913 | 23612213 | 23818541 | 27881259 | 35917517 | 26625265 | 29858575 | 31156673 | 26187439 | 29517275 | 30877563 |
| P29391 | Ftl1 | Ferritin light chain 1 Mus musculus | 527.69 | 57.38 | 20802 | 9 | 9 | 79946201 | 83928089 | 69717713 | 67122257 | 64559313 | 81511425 | 82787529 | 102819713 | 81299105 | 77043105 | 80681033 | 74852753 | 85268377 | 91899481 |
| Q8BGW1 | Fto | Alpha-ketoglutarate-dependent dioxygenase FTO Mus musculus | 118.81 | 6.57 | 58007 | 2 | 2 | 1 | 1 | 1 | 225302.11 | 1 | 197345.09 | 1 | 1 | 192606.64 | 179165.58 | 265976.12 | 327578.03 | 239117.73 | 277557.2933 |
| Q9DBE9 | Ftsj3 | pre-rRNA processing protein FTSJ3 Mus musculus | 724.5 | 26.73 | 95532 | 13 | 13 | 4936288 | 2077780 | 1827505.12 | 3731672.25 | 2251585.5 | 4390191.5 | 2012008 | 3710191.75 | 1096129 | 4121466.5 | 2784046.5 | 2555402 | 2561507 | 2118351 |
| Q32PX7 | Fubp1 | Far upstream element-binding protein 1 Rattus norvegicus | 758.75 | 29.26 | 67197 | 14 | 16 | 19452977 | 18649261 | 23757205 | 27023873 | 18323221 | 21248905 | 22855821 | 21343015 | 20289593 | 24043189 | 20810843 | 23929663 | 21800123 | 22877373 |
| Q9D6K8 | Fundc2 | FUN14 domain-containing protein 2 Mus musculus | 190.78 | 21.85 | 16564 | 3 | 3 | 5720753 | 6413602 | 5358784 | 1250962.12 | 5218878 | 6022844 | 4435958 | 5656974.5 | 4100256.5 | 4968351.5 | 3933950.25 | 5304460 | 5139792 | 5067433.5 |
| Q8R2K1 | Fuom | Fucose mutarotase Mus musculus | 126.66 | 17.65 | 16805 | 2 | 2 | 2262723.25 | 1758226.5 | 1265212.5 | 2424451 | 1251164.12 | 1359057 | 2084125.25 | 2033296.38 | 1414952.5 | 925365 | 1229686.75 | 1155378.75 | 859052.25 | 999137.94 |
| P56959 | Fus | RNA-binding protein FUS Mus musculus | 626.84 | 23.55 | 52673 | 10 | 10 | 35919305 | 40310081 | 39968925 | 42177397 | 43356985 | 42149361 | 40124533 | 32134073 | 39799865 | 50041881 | 39491133 | 34925793 | 40026637 | 41332145 |
| D3ZYW7 | Fxn | Frataxin, mitochondrial Rattus norvegicus | 48.28 | 5.77 | 23066 | 1 | 1 | 1470573.38 | 2018505.62 | 1671755.12 | 1573553.88 | 1436761.75 | 1305965 | 1669705.12 | 1574757.5 | 1504039.31 | 1465729.62 | 1517389.38 | 1579516.75 | 1564232.12 | 1435155 |
| Q61584 | Fxr1 | Fragile X mental retardation syndrome-related protein 1 Mus musculus | 897.56 | 29.69 | 76222 | 14 | 15 | 22631109 | 24512787 | 24441133 | 22195725 | 23071475 | 23125903 | 24227719 | 29673489 | 23335015 | 24734069 | 24288169 | 24764769 | 22287907 | 25428255 |
| Q9WVR4 | Fxr2 | Fragile X mental retardation syndrome-related protein 2 Mus musculus | 228.17 | 7.88 | 73743 | 3 | 4 | 8126656.5 | 8517745 | 8133283 | 8277533 | 8584711 | 8328885 | 8368202 | 9118463 | 8137485 | 8543702 | 8434421 | 9070251 | 8260224 | 8660171 |
| P97808 | Fxyd5 | FXYD domain-containing ion transport regulator 5 Mus musculus | 74.01 | 20.79 | 19454 | 2 | 2 | 700896.44 | 1 | 1 | 1 | 1 | 1 | 1 | 1462963.62 | 747003.12 | 602642.81 | 1 | 668148.62 | 687868.19 | 1 |
| O35601 | Fyb1 | FYN-binding protein 1 Mus musculus | 454.81 | 14.53 | 90055 | 9 | 9 | 6926826 | 6698229 | 6649610.5 | 11833141 | 7970181 | 6609765.5 | 6944785.5 | 6371316 | 7659888 | 9486459 | 8737728 | 6748771.5 | 5092670 | 6163651.5 |
| P97855 | G3bp1 | Ras GTPase-activating protein-binding protein 1 Mus musculus | 846.24 | 35.91 | 51829 | 13 | 15 | 15862112 | 18706691 | 18623369 | 16756893 | 19503099 | 18489215 | 17912373 | 17239991 | 18776265 | 18468591 | 18303373 | 17364131 | 18872361 | 17213423 |
| P97379 | G3bp2 | Ras GTPase-activating protein-binding protein 2 Mus musculus | 256.45 | 11 | 54088 | 3 | 5 | 4602877.5 | 6003054.5 | 5904047.5 | 5445513 | 6130925 | 3433733.25 | 5006298 | 5130017.5 | 4254653.5 | 5048372.5 | 5301427.5 | 5160429 | 4970253 | 4389167 |
| Q00612 | G6pdx | Glucose-6-phosphate 1-dehydrogenase X Mus musculus | 1454.99 | 55.73 | 59263 | 27 | 27 | 97138265 | 93448481 | 95200425 | 96327689 | 96271081 | 100245673 | 98799889 | 95811273 | 95528729 | 108365089 | 102213569 | 111241657 | 105843505 | 98231041 |
| P70699 | Gaa | Lysosomal alpha-glucosidase Mus musculus | 690.87 | 23.08 | 106248 | 14 | 14 | 22272737 | 15396345 | 14726017 | 16850859 | 17535857 | 16226853 | 18420603 | 15215264 | 19980349 | 16812677 | 17957735 | 16314774 | 19419419 | 16754811 |
| P60517 | Gabarap | Gamma-aminobutyric acid receptor-associated protein Rattus norvegicus | 148.57 | 20.51 | 13918 | 1 | 2 | 3734132.75 | 3583941 | 5045262 | 4634650.5 | 4044040.75 | 4961763.5 | 5505267 | 3233401.5 | 3970549.75 | 4630802.5 | 4381063 | 5122040 | 4187900.75 | 4945026 |
| P60521 | Gabarapl2 | Gamma-aminobutyric acid receptor-associated protein-like 2 Mus musculus | 127.07 | 27.35 | 13667 | 2 | 3 | 3766432.5 | 3512370 | 5199525 | 4685455.5 | 4001487.25 | 4939595 | 5330890.5 | 3109017 | 4025355.5 | 4667050.5 | 4585217 | 5152809.5 | 4519868.5 | 5234210 |
| Q00422 | Gabpa | GA-binding protein alpha chain Mus musculus | 221.16 | 14.32 | 51344 | 5 | 5 | 4205862 | 4537628.5 | 3042829.75 | 3104090.75 | 2691167.5 | 5209564.5 | 2761917.25 | 4202786.5 | 3789153.5 | 2944303.5 | 3936057.75 | 3309814 | 3747887.75 | 4074088.25 |
| Q99KY4 | Gak | Cyclin-G-associated kinase Mus musculus | 198.88 | 5.21 | 143641 | 3 | 3 | 3257501.5 | 3764614 | 4303446 | 2971749 | 2496956 | 2435808.25 | 2873148.75 | 3692414 | 1460724.62 | 2639480.75 | 2693713.25 | 3625443.25 | 1544908.5 | 3068043.25 |
| P54818 | Galc | Galactocerebrosidase Mus musculus | 84.13 | 5.12 | 77256 | 2 | 2 | 1318176.12 | 1087074.12 | 921299.69 | 1000666.94 | 1183955 | 1157308.62 | 849291.5 | 863285.99 | 396568.72 | 1049975.12 | 1069473.88 | 1151478.5 | 1356934.38 | 1215034.88 |
| P18645 | Gale | UDP-glucose 4-epimerase Rattus norvegicus | 126.2 | 8.65 | 38225 | 2 | 2 | 3551099.5 | 3361979.5 | 3645733.75 | 2802935.5 | 3505035 | 4324152 | 821840.69 | 3941289 | 1160605.5 | 3108605.5 | 3631896.75 | 1221190.75 | 3614922 | 3936490 |
| Q9R0N0 | Galk1 | Galactokinase Mus musculus | 453.75 | 37.5 | 42295 | 9 | 10 | 8279408 | 12743603 | 18287175 | 10402457 | 10066241 | 19555699 | 18757369 | 19496847 | 22014131 | 22097199 | 11510724 | 12738829 | 18303935 | 19423973 |
| Q68FH4 | Galk2 | N-acetylgalactosamine kinase Mus musculus | 220.02 | 12.01 | 50503 | 4 | 4 | 4180592.5 | 5116891 | 4586589 | 5857616 | 5836480.5 | 3977218.5 | 4684769 | 5953130.5 | 4545496 | 3190362.75 | 4959292 | 7426904 | 4579288 | 4592748.5 |
| Q571E4 | Galns | N-acetylgalactosamine-6-sulfatase Mus musculus | 394.08 | 15.96 | 57673 | 6 | 7 | 10118437 | 11670897 | 9301575 | 9488543 | 10695998 | 10728196 | 10377405 | 12217593 | 10235504 | 9559614 | 10737903 | 11769617 | 9990048 | 11172712 |
| Q6PB93 | Galnt2 | Polypeptide N-acetylgalactosaminyltransferase 2 Mus musculus | 160.96 | 8.6 | 64514 | 4 | 4 | 1872842.25 | 1691759.5 | 2217570.5 | 1075653 | 390742.75 | 2259870 | 1478603.5 | 1794737 | 2011590 | 1771109.75 | 454198.5 | 2261460 | 2296308.5 | 2058507.25 |
| Q80VA0 | Galnt7 | N-acetylgalactosaminyltransferase 7 Mus musculus | 365.49 | 18.11 | 75419 | 8 | 8 | 15136037 | 13011606 | 9584421 | 10479327 | 12797417 | 13901345 | 10915679 | 11651599 | 11756203 | 11892221 | 11867669 | 12666589 | 12608339 | 12710270 |
| Q8BHN3 | Ganab | Neutral alpha-glucosidase AB Mus musculus | 1370.82 | 41.74 | 106911 | 29 | 29 | 34234589 | 29828711 | 32994331 | 32246355 | 28628169 | 33507635 | 33437065 | 29725203 | 34007305 | 35127225 | 34838457 | 32876801 | 38733425 | 35223829 |
| Q8BVW0 | Ganc | Neutral alpha-glucosidase C Mus musculus | 41.45 | 1.45 | 102008 | 1 | 1 | 494299.16 | 529243.2125 | 528985.5 | 519233.81 | 574454.38 | 600611.44 | 423773.31 | 425767.44 | 559476.69 | 629191.62 | 487276.78 | 464546.97 | 351868.94 | 566702.5 |
| P16858 | Gapdh | Glyceraldehyde-3-phosphate dehydrogenase Mus musculus | 2270.76 | 87.69 | 35810 | 29 | 36 | 1044268161 | 1000019969 | 1184233217 | 1225996161 | 1173118081 | 1240618881 | 1350280833 | 1054096193 | 1339714817 | 1578373249 | 1263416705 | 1378445185 | 1291220609 | 1057732929 |
| Q6PAR5 | Gapvd1 | GTPase-activating protein and VPS9 domain-containing protein 1 Mus musculus | 286.74 | 5.56 | 162402 | 7 | 7 | 1588914.12 | 3550225 | 3559893.75 | 4762644.5 | 3431529 | 4284296 | 4310535.5 | 4283576 | 5191882.5 | 3953007.25 | 2567820.5 | 3880148.75 | 3567567.75 | 1762469.88 |
| Q6AYA1 | Gar1 | H/ACA ribonucleoprotein complex subunit 1 Rattus norvegicus | 60.84 | 7.96 | 23011 | 2 | 2 | 3150930.5 | 2862889.313 | 1571096.25 | 3286665 | 3442865.5 | 2214999 | 1609378.248 | 1578279.75 | 1118974.62 | 1525259.62 | 4214360.5 | 2287204.21 | 1339832.75 | 1307419.38 |
| Q9CZD3 | Gars | Glycine--tRNA ligase Mus musculus | 1395.78 | 37.59 | 81878 | 3 | 22 | 96797529 | 94607849 | 95651737 | 99651545 | 93605913 | 109506609 | 108360201 | 106143361 | 109224985 | 107692209 | 102365249 | 99154657 | 106202961 | 101447001 |
| Q64737 | Gart | Trifunctional purine biosynthetic protein adenosine-3 Mus musculus | 1922.33 | 47.03 | 107503 | 30 | 31 | 21127445 | 21301797 | 22663411 | 23803385 | 23066641 | 23197469 | 24876195 | 23285383 | 21835641 | 23995009 | 25761401 | 23834795 | 27190243 | 22440505 |
| O55148 | Gas7 | Growth arrest-specific protein 7 Rattus norvegicus | 296.82 | 18.48 | 48275 | 6 | 6 | 8409904 | 8834103 | 6357651.5 | 10475651 | 11646223 | 8320164.5 | 8873946 | 8943824 | 9870402 | 8224000 | 7817282 | 8016844 | 6762795.5 | 9510166 |
| Q99JT1 | Gatb | Glutamyl-tRNA(Gln) amidotransferase subunit B, mitochondrial Mus musculus | 112.23 | 8.98 | 62118 | 2 | 2 | 1 | 240130.56 | 210578.31 | 187402.14 | 1 | 1 | 1 | 261355.92 | 287132.56 | 1 | 160667.83 | 1 | 1 | 1 |
| Q8BFQ8 | Gatd1 | Glutamine amidotransferase-like class 1 domain-containing protein 1 Mus musculus | 232.11 | 30 | 23277 | 4 | 4 | 3404697.5 | 3548624.75 | 3877741.25 | 3405962.75 | 3242598.75 | 4100336.25 | 3487544 | 4228942 | 4156098.5 | 2977433 | 3488624 | 3945584.5 | 3287051.25 | 3679123 |
| P50442 | Gatm | Glycine amidinotransferase, mitochondrial Rattus norvegicus | 650.58 | 30.97 | 48242 | 10 | 10 | 44178961 | 47179373 | 42780881 | 40331505 | 48363341 | 38891361 | 42791233 | 43660833 | 41006389 | 44653665 | 49273661 | 46248577 | 49506997 | 48534545 |
| P17439 | Gba | Glucosylceramidase Mus musculus | 288.44 | 20.39 | 57622 | 6 | 6 | 7411030.5 | 6799296.5 | 6604925 | 8099719.5 | 6469604 | 5052369.5 | 6231565 | 5997354 | 5270157 | 4826628 | 7181936 | 6700892.5 | 7041698 | 6856018.5 |

| | | | | | | | | | | | | | | | | | | | | | |
|---|---|---|---|---|---|---|---|---|---|---|---|---|---|---|---|---|---|---|---|---|---|
| Q9D6Y9 | Gbe1 | 1,4-alpha-glucan-branching enzyme Mus musculus | 38.75 | 1.85 | 80364 | 1 | 1 | 1 | 1 | 1 | 1 | 1 | 290928.62 | 1 | 1519358 | 998059.88 | 1345397.75 | 331358.88 | 832863.38 | 1 | 771792.75 |
| Q9Z0E6 | Gbp2 | Guanylate-binding protein 2 Mus musculus | 328.57 | 16.3 | 66739 | 6 | 6 | 4043399.75 | 4796837 | 4765077 | 1587843.38 | 3894907.25 | 888460 | 622300.69 | 531811.94 | 305732.81 | 1592889.25 | 946624.88 | 1949364.75 | 1239605 | 1473114.75 |
| Q61107 | Gbp4 | Guanylate-binding protein 4 Mus musculus | 569.72 | 26.13 | 70801 | 11 | 11 | 8514483 | 7450088 | 5133596 | 6443512 | 6430460 | 3197550.5 | 4450390.5 | 3241343.75 | 2997324.5 | 2882868.25 | 3562685 | 2570989.5 | 3680811.25 | 3870607.75 |
| O88986 | Gcat | 2-amino-3-ketobutyrate coenzyme A ligase, mitochondrial Mus musculus | 461.88 | 32.45 | 44931 | 7 | 7 | 2071480 | 2891570.25 | 3612637.25 | 2995040 | 3543770.25 | 4552562 | 3529727 | 2241942.75 | 3769953.5 | 4263833 | 3987094.5 | 4190853.5 | 4462710 | 4411221 |
| Q60759 | Gcdh | Glutaryl-CoA dehydrogenase, mitochondrial Mus musculus | 98.2 | 5.94 | 48606 | 2 | 2 | 1620535.62 | 1467741.88 | 1832715 | 1938497.88 | 1648733.38 | 1658709.62 | 1079956.25 | 1594873.62 | 1651758.38 | 1657990.88 | 1453158.88 | 1487872.12 | 1804176.88 | 1861640.62 |
| P97494 | Gclc | Glutamate--cysteine ligase catalytic subunit Mus musculus | 391.57 | 16.01 | 72571 | 6 | 6 | 1696302.5 | 3405893.5 | 3341330.25 | 3184514.5 | 3020486.75 | 3533322.75 | 3039812.5 | 4288761 | 2531285.5 | 3766027 | 4282653.5 | 4255555 | 3807138 | 3437513 |
| O09172 | Gclm | Glutamate--cysteine ligase regulatory subunit Mus musculus | 278.8 | 20.8 | 30535 | 4 | 4 | 4865610.5 | 4399010.5 | 6606841 | 2449319.75 | 4602144 | 2830404.5 | 3609352.5 | 2152036.25 | 3317049.5 | 3299061.5 | 5551141 | 5708062 | 6027968.5 | 4990138.5 |
| E9PVA8 | Gcn1 | eIF-2-alpha kinase activator GCN1 Mus musculus | 1253.66 | 11.08 | 293021 | 22 | 22 | 36018305 | 32823859 | 29905395 | 29609555 | 28048527 | 32067957 | 30585349 | 31899485 | 29943731 | 30778121 | 27961967 | 28414427 | 30802729 | 29244445 |
| Q91WK5 | Gcsh | Glycine cleavage system H protein, mitochondrial Mus musculus | 76.94 | 11.76 | 18637 | 1 | 1 | 567171.12 | 689098.62 | 795584.44 | 518279.56 | 615326.06 | 806362.62 | 632399.44 | 975940.94 | 714293.69 | 1381483.38 | 575788.88 | 882876.62 | 994238 | 857479.38 |
| Q66H63 | Gdap2 | Ganglioside-induced differentiation-associated-protein 2 Rattus norvegicus | 38.01 | 1.61 | 56130 | 1 | 1 | 831176.88 | 720556.62 | 932246.88 | 1131572.75 | 758783.62 | 1002460.25 | 789216.88 | 173307.62 | 650422.31 | 700932.25 | 892228.25 | 863770.75 | 764069.44 | 934595 |
| P50396 | Gdi1 | Rab GDP dissociation inhibitor alpha Mus musculus | 868.4 | 51.68 | 50522 | 9 | 14 | 52040957 | 48112345 | 39622377 | 44421417 | 52836105 | 54773865 | 51716905 | 51556465 | 53003941 | 51255785 | 52589737 | 49481381 | 52232929 | 51537385 |
| P50399 | Gdi2 | Rab GDP dissociation inhibitor beta Rattus norvegicus | 2045.24 | 74.38 | 50537 | 22 | 29 | 167608433 | 167957473 | 152872913 | 156400529 | 174835505 | 174719537 | 173082465 | 193976993 | 177382993 | 169172929 | 166047761 | 170222929 | 169366113 | 165560337 |
| Q9CQQ4 | Gemin2 | Gem-associated protein 2 Mus musculus 2 | 46.47 | 6.32 | 30441 | 1 | 1 | 1 | 1 | 1 | 131221.45 | 120477.13 | 107514.89 | 99570.95 | 102259.91 | 104127.96 | 134379.05 | 184930.16 | 1 | 1 | 136018.59 |
| Q8BX17 | Gemin5 | Gem-associated protein 5 Mus musculus | 193.13 | 4.53 | 166592 | 5 | 5 | 3853023 | 3625824.5 | 3026013.5 | 2197521.5 | 2166582.25 | 1310570.88 | 1604851.12 | 2008812.38 | 1628851.62 | 1231911.75 | 1028084.69 | 710402 | 1677707.25 | 940815.56 |
| Q9D1H7 | Get4 | Golgi to ER traffic protein 4 homolog Mus musculus | 133.38 | 12.23 | 36525 | 3 | 3 | 2727767 | 2513847.5 | 2044945.5 | 2748242.75 | 2697812.5 | 2023680.75 | 1667199.25 | 1509719.38 | 1007143.12 | 1561593 | 1789479.75 | 3348092 | 1421113.75 | 2742497.25 |
| Q8K0D5 | Gfm1 | Elongation factor G, mitochondrial Mus musculus | 547.2 | 18.24 | 83550 | 3 | 10 | 9691852 | 9473548 | 10245069 | 6776132.5 | 8243651 | 8782423 | 8630944 | 10870203 | 8672735 | 7316871 | 9942389 | 11005298 | 10797326 | 9556613 |
| Q5BJP6 | Gfm2 | Ribosome-releasing factor 2, mitochondrial Rattus norvegicus | 47.55 | 1.67 | 85914 | 1 | 1 | 1 | 1 | 807486.94 | 879959.81 | 820097.69 | 728937.56 | 1 | 1 | 1 | 1 | 893669.94 | 921067 | 853079.81 | 889272.25 |
| P47856 | Gfpt1 | Glutamine--fructose-6-phosphate aminotransferase [isomerizing] 1 Mus musculus | 178.28 | 8.32 | 78539 | 4 | 4 | 2224536 | 2953629 | 2589416 | 2316178.25 | 1497132 | 1733019.5 | 1426399.12 | 3176112.5 | 2286460.75 | 2594273.75 | 1374403.75 | 2343649.083 | 3563548.25 | 2092995.25 |
| Q8R0H9 | Gga1 | ADP-ribosylation factor-binding protein GGA1 Mus musculus | 122.02 | 6.61 | 69972 | 2 | 2 | 1854207 | 395725.62 | 1687098.5 | 1813656.25 | 1705823.5 | 1969647.75 | 1726057.5 | 1770974 | 2144115.5 | 2265988 | 1937131.25 | 1350254.12 | 784850.38 | 429663.38 |
| Q6P5E6 | Gga2 | ADP-ribosylation factor-binding protein GGA2 Mus musculus | 51.11 | 4.48 | 66049 | 1 | 1 | 392778.47 | 599482.06 | 234197.67 | 447510.44 | 418492.16 | 332977.615 | 387613.78 | 351796.62 | 290301.53 | 302198.53 | 412262.34 | 262550.89 | 184787.95 | 190602.38 |
| Q91VC9 | Ghitm | Growth hormone-inducible transmembrane protein Mus musculus | 181.47 | 16.76 | 37275 | 3 | 3 | 806760.25 | 3824930.5 | 4245632 | 4440761.5 | 4651806.5 | 2670675 | 4195592 | 1035448.62 | 3658340.25 | 1321048 | 664230.44 | 3011875.25 | 2527966 | 4868522.5 |
| Q9D7M1 | Gid8 | Glucose-induced degradation protein 8 homolog Mus musculus | 290.44 | 39.04 | 26779 | 5 | 5 | 2313184.25 | 3789896.5 | 3269249.5 | 3287851.75 | 3370374.5 | 3719707.5 | 3755467.5 | 3764215 | 2545261.5 | 1786544.5 | 2814087.5 | 2074284.5 | 3196470.75 | 2359339.75 |
| Q6Y7W8 | Gigyf2 | GRB10-interacting GYF protein 2 Mus musculus | 228.63 | 5.65 | 149193 | 4 | 4 | 1181891.62 | 1094963.62 | 1509011.12 | 950411.12 | 983237.94 | 1733878 | 1438916.12 | 574003.75 | 1414048 | 1174552 | 990902.38 | 1591662 | 1362716.5 | 1042647.75 |
| Q68FF6 | Git1 | ARF GTPase-activating protein GIT1 Mus musculus | 82.2 | 1.82 | 85300 | 1 | 1 | 333864.03 | 464910.84 | 461032.47 | 310569.19 | 351643.22 | 309675.44 | 418786.66 | 503195.56 | 458505.19 | 495127.34 | 439829 | 569019.94 | 532316.75 | 462526.22 |
| Q9JLQ2 | Git2 | ARF GTPase-activating protein GIT2 Mus musculus | 138.18 | 8.47 | 78766 | 3 | 3 | 300345.81 | 1919752.5 | 2231588.5 | 1810593.75 | 763491.31 | 2607851.25 | 182271.52 | 1562675.25 | 1505120.62 | 1178312.88 | 1981731.75 | 1943292.75 | 1414512.5 | 1634475.25 |
| Q64516 | Gk | Glycerol kinase Mus musculus | 418.09 | 20.57 | 61227 | 9 | 9 | 10587397 | 8673071 | 11610506 | 10183176 | 8658168 | 10396198 | 8987588 | 9890937 | 11227581 | 10809801 | 7556988 | 10201135 | 12063701 | 11082126 |
| P51569 | Gla | Alpha-galactosidase A Mus musculus | 791.73 | 39.38 | 47643 | 13 | 13 | 43265073 | 44987593 | 41021769 | 42075301 | 41287025 | 45426405 | 44033309 | 39982649 | 47183433 | 38267797 | 41659417 | 39176089 | 46818017 | 40861957 |
| P23780 | Glb1 | Beta-galactosidase Mus musculus | 782.19 | 24.57 | 73121 | 15 | 15 | 74988033 | 85227161 | 72027969 | 63749433 | 80690641 | 75850049 | 69349449 | 82707729 | 87212793 | 71003721 | 72965217 | 73181705 | 82541761 | 73411857 |
| Q61543 | Glg1 | Golgi apparatus protein 1 Mus musculus | 1636.4 | 32.26 | 133734 | 29 | 29 | 55488997 | 55751933 | 47964597 | 49339425 | 53261433 | 45867373 | 46099345 | 49111949 | 42774749 | 50527941 | 51202229 | 42910741 | 46877805 | 49523897 |
| Q9CWG1 | Glipr1 | Glioma pathogenesis-related protein 1 Mus musculus | 61.17 | 5.49 | 29129 | 1 | 1 | 1234218.25 | 1205148.25 | 1037347 | 1332321.75 | 1437550.5 | 1610373.62 | 1616885.88 | 1726157.62 | 1560643.75 | 1267817.25 | 1606324.75 | 1614322.5 | 1579347.5 | 1447292.12 |
| Q8BZM1 | Glmn | Glomulin Mus musculus | 192.62 | 6.21 | 67756 | 3 | 3 | 1630325.62 | 1510563.5 | 1626378.5 | 1526176 | 1512363.75 | 1368659 | 1420061.38 | 1522409.75 | 1621259.88 | 1244536 | 1345730.75 | 1563846.88 | 1495373.25 | 1394731.62 |
| Q9JHJ3 | Glmp | Glycosylated lysosomal membrane protein Mus musculus | 115.5 | 8.17 | 43804 | 2 | 2 | 390392.56 | 375639.5 | 895242.81 | 521804.69 | 836234.56 | 1268463.12 | 620702.69 | 364765.09 | 256151.8 | 1277358.5 | 1246621.5 | 1066663.75 | 1269519.25 | 1135898.12 |
| Q9CPU0 | Glo1 | Lactoylglutathione lyase Mus musculus | 441.84 | 46.74 | 20810 | 8 | 8 | 22031245 | 26920293 | 19755439 | 26164569 | 24276729 | 29083327 | 29703499 | 28965929 | 30514657 | 28386367 | 26571287 | 29813259 | 23432779 | 24923859 |
| Q9CPV4 | Glod4 | Glyoxalase domain-containing protein 4 Mus musculus | 817.04 | 60.4 | 33317 | 7 | 15 | 21218533 | 106618553 | 18382133 | 23281503 | 22404561 | 20812341 | 24269301 | 22565341 | 22732179 | 22480103 | 22917261 | 22319975 | 19781765 | 22897403 |
| Q9QUH0 | Glrx | Glutaredoxin-1 Mus musculus | 296.6 | 57.94 | 11871 | 4 | 4 | 14472304 | 17170877 | 22626497 | 12931024 | 14314939 | 17952957 | 23487779 | 21301867 | 21697245 | 16675105 | 15902309 | 21791083 | 18866429 | 17416145 |
| Q9CQM9 | Glrx3 | Glutaredoxin-3 Mus musculus | 666.96 | 43.62 | 37778 | 10 | 10 | 17375581 | 19209147 | 16288185 | 17734797 | 16590306 | 16355206 | 16828911 | 22039001 | 16970819 | 16317839 | 16953127 | 17155281 | 15929085 | 15936488 |
| Q80Y14 | Glrx5 | Glutaredoxin-related protein 5, mitochondrial Mus musculus | 164.67 | 20.39 | 16292 | 2 | 2 | 1583419.75 | 2376964 | 2317822.25 | 1911952.12 | 2597586.25 | 2662592.75 | 2313371.5 | 2358786.5 | 2506278.5 | 2582399.5 | 2234289.75 | 2283285.75 | 2573934 | 2733465 |
| D3Z7P3 | Gls | Glutaminase kidney isoform, mitochondrial Mus musculus | 705.94 | 29.08 | 73964 | 12 | 12 | 6084751 | 7923345.5 | 7132730 | 7813474 | 7803654.5 | 7447727.5 | 9725034 | 8178582.5 | 9470943 | 8108374 | 8526909 | 6205809 | 9836598 | 6127518 |
| B0BNM9 | GLTP | Glycolipid transfer protein Rattus norvegicus | 82.99 | 12.44 | 23704 | 2 | 2 | 947167.69 | 1288210.75 | 1221304.5 | 960443 | 1010822.81 | 882689.5 | 1060279.5 | 1647905.38 | 1557285.12 | 1036941.69 | 1413363.12 | 1251070 | 998503.81 | 1356582.62 |
| Q8K0R6 | Gltpd2 | Glycolipid transfer protein domain-containing protein 2 Mus musculus | 40.18 | 2.18 | 34453 | 1 | 1 | 2788235.5 | 248642.72 | 1 | 1 | 1 | 1 | 1 | 1 | 1250316.25 | 1 | 2636152.25 | 2335906.75 | 377894.38 | 1783317.793 |
| P26443 | Glud1 | Glutamate dehydrogenase 1, mitochondrial Mus musculus | 1417.03 | 53.76 | 61337 | 23 | 23 | 107670049 | 126157753 | 121263281 | 120967529 | 120317089 | 112692697 | 117573241 | 117584049 | 113179953 | 117103169 | 110977017 | 102713161 | 106702961 | 116947689 |
| Q5RKH0 | Glyr1 | Putative oxidoreductase GLYR1 Rattus norvegicus | 449 | 22.64 | 60421 | 7 | 7 | 5601259 | 7051058 | 7290744 | 7394831 | 9028924 | 8183828 | 7816592 | 6041579.5 | 8217723.5 | 8387823 | 6293011.5 | 7446675 | 7986553 | 7752856 |
| Q8K0C9 | Gmds | GDP-mannose 4,6 dehydratase Mus musculus | 354.99 | 23.12 | 41985 | 6 | 6 | 4954164.5 | 4834941 | 4326470.5 | 4401193.5 | 2836879.5 | 3014695 | 2744977 | 5182063.5 | 4375740 | 2806547 | 2605723.5 | 4808694 | 2729535 | 2494089.75 |
| Q63228 | Gmfb | Glia maturation factor beta Rattus norvegicus | 78.36 | 14.08 | 16736 | 1 | 2 | 4264387.5 | 3684344 | 5403790.5 | 4835011.5 | 5390039 | 4697552.5 | 3778423.25 | 1948636.12 | 5261657.5 | 6029249.5 | 6480667 | 6032505 | 5564252.5 | 4803169.5 |
| Q9ERL7 | Gmfg | Glia maturation factor gamma Mus musculus | 275.54 | 42.96 | 16748 | 5 | 6 | 7088348 | 9215614 | 7856842 | 10454353 | 12165899 | 7668588 | 7252592 | 4192719.5 | 8200499 | 12620921 | 11718839 | 9838497 | 9876161 | 10812357 |
| Q6PGG2 | Gmip | GEM-interacting protein Mus musculus | 250.95 | 8.75 | 107545 | 5 | 5 | 892273.06 | 1753771.12 | 1248583.75 | 1306761.62 | 944341.38 | 1151791.38 | 1557636 | 2010730 | 1479341.12 | 1672287.88 | 1042261.69 | 1100546.75 | 899107.56 | 1834189.75 |
| Q5XIC1 | Gmppa | Mannose-1-phosphate guanyltransferase alpha Rattus norvegicus | 92.41 | 6.43 | 46191 | 2 | 2 | 932103.75 | 38146285 | 693062.75 | 947676.38 | 50922325 | 888022.25 | 874213.25 | 497862.28 | 21046403 | 1100513.88 | 63354433 | 823490.25 | 29847211 | 1023321.94 |
| Q8BTZ7 | Gmppb | Mannose-1-phosphate guanyltransferase beta Mus musculus | 49.39 | 5 | 39917 | 1 | 1 | 4203369.5 | 4833290.5 | 4242727.5 | 4922349.5 | 5575047 | 3916547.25 | 3445408.25 | 2720064.25 | 2685379.5 | 2948363 | 4816802 | 3610994.75 | 3373055.75 | 4345794 |
| Q9DCZ1 | Gmpr | GMP reductase 1 Mus musculus | 192.79 | 8.99 | 37482 | 3 | 3 | 473300.62 | 958105.69 | 422140.38 | 943952 | 788612.88 | 329385.25 | 591895.25 | 530731.06 | 720694.62 | 810233.44 | 811813.94 | 452938.47 | 979452.25 | 781518.25 |
| Q3THK7 | Gmps | GMP synthase [glutamine-hydrolyzing] Mus musculus | 1328.99 | 52.96 | 76723 | 23 | 23 | 29054957 | 30467919 | 32455597 | 32417665 | 32113261 | 30536041 | 33634037 | 33911477 | 32332507 | 33128155 | 31065503 | 30755097 | 33015631 | 32623909 |
| P21278 | Gna11 | Guanine nucleotide-binding protein subunit alpha-11 Mus musculus | 127.59 | 12.53 | 42024 | 1 | 3 | 1113086.62 | 816279.62 | 1206552.12 | 547198.38 | 527918.75 | 1198028.75 | 798447.06 | 1617357.25 | 1360297.5 | 184285.52 | 845831.62 | 1223613.62 | 1082496.62 | 216101.66 |
| P27601 | Gna13 | Guanine nucleotide-binding protein subunit alpha-13 Mus musculus | 378.98 | 22.81 | 44055 | 8 | 8 | 10219065 | 8161165 | 10680955 | 9740980 | 8717010 | 10575096 | 8603372 | 9959616 | 12527249 | 10164701 | 8203712.5 | 11153471 | 10896655 | 9608413 |
| P08752 | Gnai2 | Guanine nucleotide-binding protein G(i) subunit alpha-2 Mus musculus | 1594.85 | 67.89 | 40489 | 14 | 24 | 152412081 | 163942833 | 159485921 | 144590897 | 157207153 | 148012401 | 153471569 | 162173457 | 157375089 | 153814353 | 153350097 | 144715617 | 156157553 | 160956017 |
| P08753 | Gnai3 | Guanine nucleotide-binding protein G(k) subunit alpha Rattus norvegicus | 832.77 | 39.55 | 40522 | 4 | 12 | 30806729 | 33407441 | 33833005 | 25643085 | 31113389 | 27595705 | 29632045 | 34303205 | 32817639 | 31030537 | 28694303 | 30943099 | 33237419 | 32815841 |
| P21279 | Gnaq | Guanine nucleotide-binding protein G(q) subunit alpha Mus musculus | 358.17 | 24.51 | 42158 | 5 | 7 | 8880789 | 7985632 | 7864490.5 | 7197769.5 | 7928388 | 7973394 | 6870315.5 | 7889930 | 7342849.5 | 5660104.5 | 3901811.25 | 7675033 | 7515092 | 6746829 |
| P63094 | Gnas | Guanine nucleotide-binding protein G(s) subunit alpha isoforms short Mus musculus | 581.65 | 32.74 | 45664 | 10 | 11 | 9744239 | 12054281 | 10497814 | 10054858 | 11257588 | 11958830 | 10949654 | 12724107 | 12494934 | 10637821 | 11849607 | 8734258 | 11344956 | 12803589 |
| P54311 | Gnb1 | Guanine nucleotide-binding protein G(I)/G(S)/G(T) subunit beta-1 Rattus norvegicus | 705.63 | 50.59 | 37377 | 6 | 11 | 16494165 | 15670456 | 22188921 | 18765769 | 17749507 | 17807339 | 21520853 | 19154131 | 20509197 | 21696797 | 19391725 | 19253883 | 22130727 | 20182181 |
| P54313 | Gnb2 | Guanine nucleotide-binding protein G(I)/G(S)/G(T) subunit beta-2 Rattus norvegicus | 1056.32 | 63.53 | 37331 | 10 | 16 | 34841069 | 42208965 | 42039529 | 43463409 | 41996961 | 39172537 | 42451449 | 36768093 | 38788169 | 45933025 | 41990077 | 35773977 | 44940545 | 44710393 |
| P63213 | Gng2 | Guanine nucleotide-binding protein G(I)/G(S)/G(O) subunit gamma-2 Mus musculus | 83.93 | 39.44 | 7850 | 2 | 2 | 6892465 | 7611842 | 7428047 | 8924962 | 8298338.5 | 5637436 | 6061224.5 | 6645098 | 5762194 | 5795699 | 6408208 | 5934865 | 5218212 | 6679262.5 |
| P36916 | Gnl1 | Guanine nucleotide-binding protein-like 1 Mus musculus | 343.07 | 13.84 | 68771 | 5 | 5 | 2113483.5 | 3079817 | 3214730.5 | 2743833.25 | 4014801.5 | 3941300 | 3046383 | 3231129.5 | 3610171.5 | 3418060.5 | 3963749.75 | 3848663.25 | 3468428 | 3543236.25 |
| Q8CI11 | Gnl3 | Guanine nucleotide-binding protein-like 3 Mus musculus | 624.34 | 30.86 | 60786 | 12 | 12 | 6893120 | 6732897.5 | 10253097 | 8987618 | 5074815 | 8063088 | 11446593 | 11635340 | 7495265 | 11745466 | 10239552 | 6687225 | 7075603.5 | 10991698 |
| P98192 | Gnpat | Dihydroxyacetone phosphate acyltransferase Mus musculus | 70.04 | 2.36 | 76870 | 1 | 1 | 1 | 1 | 1 | 3075630.75 | 1 | 2275717.5 | 1 | 1 | 1979153.12 | 1564254 | 1 | 1 | 1577890.62 | 2467910.25 |
| O88958 | Gnpda1 | Glucosamine-6-phosphate isomerase 1 Mus musculus | 160 | 16.26 | 32549 | 3 | 3 | 2575737.25 | 2873447.75 | 1285031.5 | 3602059.5 | 2907078 | 3624560.5 | 2728492 | 3503794.75 | 2965915.75 | 2960945.25 | 2763814.5 | 3501990 | 3229915.5 | 3187604.5 |
| Q9JK38 | Gnpnat1 | Glucosamine 6-phosphate N-acetyltransferase Mus musculus | 111.98 | 15.76 | 20791 | 2 | 2 | 3336251.5 | 4078587.5 | 3908066.5 | 3978580.75 | 3855449.5 | 3621814.5 | 3910776.5 | 4090617 | 3768142.5 | 3837273 | 3755392 | 3785172.25 | 3770731.75 | 3792498.5 |
| Q8BFR4 | Gns | N-acetylglucosamine-6-sulfatase Mus musculus | 715.07 | 25.55 | 61175 | 13 | 13 | 53642101 | 56073937 | 50804413 | 59516449 | 58488501 | 54168009 | 57293253 | 40762445 | 54765685 | 47932913 | 49966917 | 48354757 | 51611917 | 47777021 |
| Q62839 | Golga2 | Golgin subfamily A member 2 Rattus norvegicus | 39.4 | 1 | 112872 | 1 | 1 | 347358.12 | 1 | 1 | 433134.72 | 551018.94 | 480830.25 | 536498.38 | 404406.34 | 405087.88 | 482377.75 | 468199.56 | 518845.25 | 445522.94 | 477522.5833 |
| Q91VW5 | Golga4 | Golgin subfamily A member 4 Mus musculus | 59.57 | 0.58 | 257564 | 1 | 1 | 421708.03 | 476552.31 | 481143.72 | 511592.06 | 283701.88 | 503459.19 | 509830.31 | 481181.22 | 445429.25 | 550146.75 | 561573.81 | 582947.25 | 744044.69 | 500434.44 |
| Q9CRA5 | Golph3 | Golgi phosphoprotein 3 Mus musculus | 161.82 | 9.73 | 33752 | 1 | 2 | 1233853.5 | 1361809.75 | 1235296.25 | 1315822.62 | 1022313.88 | 1292429.75 | 1207900.38 | 1400208.25 | 1158890.12 | 1060284.25 | 1300757.25 | 1092566.88 | 983603.88 | 1395493.12 |
| Q66H74 | Golph3l | Golgi phosphoprotein 3-like Rattus norvegicus | 155.56 | 8.77 | 32922 | 1 | 2 | 1976959.75 | 1936485 | 1743979.62 | 1889268.88 | 1744210.38 | 1789250.12 | 1735125.75 | 1400208.25 | 1648827.5 | 1487278.12 | 1899141 | 1607180.62 | 1340708.25 | 1908126.12 |
| Q9CR60 | Golt1b | Vesicle transport protein GOT1B Mus musculus | 130.62 | 10.87 | 15422 | 2 | 2 | 5513042 | 824664.62 | 806548.38 | 417443.72 | 390140.72 | 1411560.25 | 738161.38 | 767750.75 | 881641.62 | 860113.25 | 781310.5 | 956547.5 | 1228541 | 1302028.5 |
| Q99JX3 | Gorasp2 | Golgi reassembly-stacking protein 2 Mus musculus | 329.11 | 17.29 | 47038 | 5 | 5 | 4681539.5 | 4907770 | 5190959 | 6039763 | 5307992.5 | 5237837 | 6423681 | 5135719 | 5090034.5 | 5320339 | 5041966.5 | 4403033 | 4722376 | 5489804 |
| O88630 | Gosr1 | Golgi SNAP receptor complex member 1 Mus musculus | 47.82 | 5.2 | 28489 | 1 | 1 | 1441950.25 | 1730465.5 | 1363458 | 1220558.88 | 1553179.25 | 1473022.12 | 1368655.12 | 1654186 | 1589600.5 | 1396021.75 | 1570097.88 | 1638646 | 1577749.75 | 1568434 |
| P05201 | Got1 | Aspartate aminotransferase, cytoplasmic Mus musculus | 1339.22 | 63.44 | 46248 | 23 | 23 | 183289009 | 128224865 | 113330681 | 157477169 | 121211169 | 122396961 | 160161329 | 143314017 | 136248353 | 118846833 | 131136793 | 113596081 | 109888041 | 106201473 |
| P05202 | Got2 | Aspartate aminotransferase, mitochondrial Mus musculus | 1388.24 | 53.72 | 47411 | 23 | 23 | 312332705 | 298579585 | 324323841 | 323117889 | 290100417 | 288913985 | 327967169 | 388317089 | 324825473 | 308995521 | 290893729 | 290573953 | 288282177 | 282179137 |
| Q3ULJ0 | Gpd1l | Glycerol-3-phosphate dehydrogenase 1-like protein Mus musculus | 280.65 | 27.64 | 38226 | 7 | 7 | 2543968 | 2371350.5 | 3832269.25 | 988359.56 | 161670.52 | 3222803.75 | 764191.06 | 3709063.75 | 3584242.5 | 3437874.5 | 3303133.75 | 523723.03 | 1100350.25 | 3338987.25 |
| Q64521 | Gpd2 | Glycerol-3-phosphate dehydrogenase, mitochondrial Mus musculus | 536.62 | 22.28 | 80954 | 12 | 12 | 12265822 | 14680938 | 14065567 | 13584454 | 12169963 | 15677707 | 15148026 | 13589635 | 16696129 | 13955227 | 16333337 | 14846157 | 16393345 | 15033927 |
| P06745 | Gpi | Glucose-6-phosphate isomerase Mus musculus | 1639.38 | 58.6 | 62767 | 14 | 27 | 122799385 | 132587617 | 124140889 | 107792553 | 137788401 | 142960385 | 132582481 | 130616985 | 135631617 | 153000705 | 132809865 | 139272545 | 143035905 | 127423449 |
| Q99P91 | Gpnmb | Transmembrane glycoprotein NMB Mus musculus | 299.24 | 10.45 | 63676 | 6 | 6 | 15079978 | 18625543 | 14805663 | 15678684 | 18425977 | 28292055 | 26069569 | 21835903 | 21971033 | 23690629 | 24751743 | 21765275 | 28238209 | 24268953 |
| Q8CIM5 | Gpr84 | G-protein coupled receptor 84 Mus musculus | 320.45 | 27.02 | 43717 | 5 | 5 | 7807401 | 6131564 | 5569723 | 5427608.5 | 7374241 | 7488883 | 6987511 | 5552706 | 5684752 | 6492601 | 7351480 | 7512429 | 7116934 | 7253135 |
| P97834 | Gps1 | COP9 signalosome complex subunit 1 Rattus norvegicus | 423.14 | 24.42 | 53428 | 8 | 8 | 8377154.5 | 7295562.5 | 8448687 | 5767675.5 | 6388505.5 | 7127403.5 | 6982472.5 | 9137393 | 8344485 | 7528249.5 | 7721758 | 7560394.5 | 7365825.5 | 7247850 |
| P11352 | Gpx1 | Glutathione peroxidase 1 Mus musculus | 216.24 | 29.85 | 22329 | 4 | 4 | 12420509 | 9940401 | 8472454 | 11677418 | 11073843 | 8316239 | 9854850 | 8782298 | 8203964 | 7889484 | 8033744 | 7628524 | 8737673 | 6710114 |
| O70325 | Gpx4 | Phospholipid hydroperoxide glutathione peroxidase Mus musculus | 178.09 | 22.34 | 22229 | 4 | 4 | 12119297 | 14274557 | 13153134 | 13247991 | 12999605 | 11885990 | 13190193 | 14428155 | 13218379 | 11809083 | 11402723 | 11629559 | 11910745 | 13811313 |
| Q60631 | Grb2 | Growth factor receptor-bound protein 2 Mus musculus | 359.46 | 40.55 | 25238 | 7 | 7 | 19001541 | 20050225 | 18111553 | 11778371 | 17586897 | 20223567 | 20392341 | 23892001 | 21949587 | 16570933 | 17624887 | 18990317 | 18964185 | 18781955 |
| O35127 | Grcc10 | Protein C10 Mus musculus | 116.25 | 28.57 | 13194 | 2 | 2 | 493293.09 | 2113838.5 | 1189737.88 | 1739117.5 | 1862298.5 | 1602291.25 | 2247705.25 | 1901685.25 | 1775683.25 | 1876648 | 1866430.25 | 1736553.75 | 1791326 | 2008986.5 |
| Q91Z53 | Grhpr | Glyoxylate reductase/hydroxypyruvate reductase Mus musculus | 301.1 | 23.17 | 35329 | 6 | 6 | 4643523 | 5200505 | 4224973.5 | 4792153.5 | 4646698.5 | 4288927 | 5519219 | 5483881.5 | 5000092 | 5521907 | 4659018 | 5757194 | 4614993.5 | 4732315 |
| P26817 | Grk2 | Beta-adrenergic receptor kinase 1 Rattus norvegicus | 607.39 | 22.06 | 79785 | 13 | 13 | 6677560 | 5539719 | 6658146 | 4423462.5 | 6012117 | 6890418 | 5496088.5 | 6438050.5 | 6938860.5 | 6413163 | 7361141 | 7357792 | 7052446.5 | 6780686.5 |
| P28798 | Grn | Granulins Mus musculus | 564.23 | 22.07 | 63458 | 11 | 11 | 36123945 | 42283901 | 28707123 | 37969009 | 44100657 | 33868609 | 33300453 | 35215921 | 30989385 | 39370857 | 42014413 | 34623141 | 42847965 | 43860541 |
| P97576 | Grpel1 | GrpE protein homolog 1, mitochondrial Rattus norvegicus | 223.31 | 25.35 | 24297 | 5 | 5 | 266657.03 | 958875.81 | 245341.73 | 1302648 | 955945.62 | 1787370.5 | 1601601.248 | 2152585.75 | 1825410.62 | 641038.12 | 1955658.25 | 1126871.5 | 1439562.12 | 1564608.25 |
| Q8C5Q4 | Grsf1 | G-rich sequence factor 1 Mus musculus | 124.36 | 7.1 | 53076 | 2 | 2 | 1513430.5 | 265656.28 | 1716902 | 1505736 | 1397569.5 | 1411365.12 | 1693866.12 | 1559422.81 | 1417831.38 | 1714628.62 | 1673637.75 | 1647948 | 1817124.12 | 1472050.38 |

| | | | | | | | | | | | | | | | | | | | | | |
|---|---|---|---|---|---|---|---|---|---|---|---|---|---|---|---|---|---|---|---|---|---|
| Q810D6 | Grwd1 | Glutamate-rich WD repeat-containing protein 1 Mus musculus | 220.73 | 12.56 | 49224 | 4 | 4 | 4084838.5 | 3332763 | 4120762 | 4366309 | 3994580.5 | 3952091 | 4107416 | 1473943.25 | 1931761.25 | 2291639 | 4436612 | 6234604 | 2829027.5 | 3536809 |
| Q9D8T2 | Gsdmdc1 | Gasdermin-D Mus musculus | 559.62 | 28.54 | 53238 | 10 | 10 | 30094597 | 28334321 | 28243353 | 24313145 | 26067663 | 28353653 | 24874685 | 24509161 | 24364011 | 25522673 | 25905825 | 27278701 | 28803887 | 27123301 |
| Q9Z2D3 | Gsdme | Gasdermin-E Mus musculus | 44.84 | 1.76 | 56631 | 1 | 1 | 643205.62 | 1 | 1 | 554476.19 | 1 | 717198.25 | 1 | 1 | 686312.56 | 672671.19 | 798345.38 | 677917.94 | 1 | 1 |
| Q2NL51 | Gsk3a | Glycogen synthase kinase-3 alpha Mus musculus | 114.63 | 4.69 | 51661 | 2 | 2 | 964385.38 | 694561.5 | 677590.38 | 743043.72 | 635637.62 | 839601.735 | 757190.06 | 701533.94 | 1237492.88 | 662190.06 | 755771.44 | 879618.06 | 700028.62 | 1012125.19 |
| P18266 | Gsk3b | Glycogen synthase kinase-3 beta Rattus norvegicus | 49.49 | 6.9 | 46742 | 1 | 1 | 1 | 1 | 1 | 1 | 1 | 160553.52 | 1 | 524925.69 | 1 | 1 | 1 | 353668.78 | 363961.97 | 1 |
| P13020 | Gsn | Gelsolin Mus musculus | 2023.14 | 54.87 | 85942 | 32 | 32 | 165182961 | 175577073 | 172119601 | 185502081 | 203728641 | 194625457 | 195215345 | 155902273 | 202464081 | 235813681 | 196114529 | 185443169 | 172692705 | 187874929 |
| Q8R050 | Gspt1 | Eukaryotic peptide chain release factor GTP-binding subunit ERF3A Mus musculus | 500.2 | 23.58 | 68625 | 9 | 9 | 6721178 | 6278470.5 | 6931352.5 | 7236142.5 | 4968875.5 | 5374094.5 | 6467379.5 | 5542193 | 5488033.5 | 7426291 | 6801280.5 | 5312694.5 | 6378752.5 | 6003435 |
| P47791 | Gsr | Glutathione reductase, mitochondrial Mus musculus | 691.09 | 30 | 53663 | 6 | 10 | 17212551 | 19611767 | 20513933 | 22004241 | 14095437 | 15825484 | 18567387 | 21290097 | 16080771 | 14967496 | 17920513 | 20010077 | 16540153 | 18098553 |
| P46413 | Gss | Glutathione synthetase Rattus norvegicus | 181.99 | 11.81 | 52345 | 4 | 4 | 4631200 | 4455283.5 | 5560903 | 5407622.5 | 5190456 | 5518262.5 | 6339487.5 | 6234936 | 5175755.5 | 5234138 | 5751489 | 5696827.5 | 6323873.5 | 3467626.5 |
| Q9DCM2 | Gstk1 | Glutathione S-transferase kappa 1 Mus musculus | 135.54 | 17.7 | 25704 | 3 | 3 | 1352608.5 | 796224.12 | 2325049 | 1762967.5 | 1370246.5 | 949027.94 | 1602979.5 | 2208718.25 | 2258845.5 | 1891459.25 | 1898817.75 | 2001627.25 | 2048410.25 | 2027169.12 |
| P10649 | Gstm1 | Glutathione S-transferase Mu 1 Mus musculus | 515.64 | 46.33 | 25970 | 8 | 9 | 12996580 | 10876059 | 17243077 | 11134448 | 11213759 | 10050049 | 17740525 | 20164789 | 19646377 | 19330783 | 20700775 | 10986923 | 17349591 | 20350641 |
| P48774 | Gstm5 | Glutathione S-transferase Mu 5 Mus musculus | 485.22 | 42.41 | 26635 | 7 | 8 | 14716581 | 17360573 | 24001869 | 14632357 | 17022199 | 15564605 | 20672531 | 22845959 | 25060383 | 23578205 | 24288985 | 18587005 | 25027437 | 25579923 |
| O09131 | Gsto1 | Glutathione S-transferase omega-1 Mus musculus | 225.92 | 20 | 27498 | 4 | 4 | 5040239 | 5463971 | 4953207.5 | 6277726 | 5098597 | 5474537 | 5270814 | 5626814 | 5846466 | 5209385 | 5784273 | 5326780 | 5071233.5 | 4683463.5 |
| P62915 | Gtf2b | Transcription initiation factor IIB Mus musculus | 257.34 | 18.99 | 34819 | 4 | 4 | 4488791 | 4869705.5 | 4240542 | 4620558 | 4386188.5 | 4213214 | 3835092.5 | 4085552.5 | 4324867.5 | 4468918 | 3995049.5 | 4593472 | 4608801 | 3292576 |
| Q9D0D5 | Gtf2e1 | General transcription factor IIE subunit 1 Mus musculus | 48.41 | 2.95 | 49593 | 1 | 1 | 1 | 445364.44 | 705151.12 | 506598.72 | 746931.44 | 801906.94 | 509523.31 | 711580.5 | 760158.94 | 612365.75 | 744133.0833 | 714883.69 | 780880.75 | 736634.81 |
| Q3THK3 | Gtf2f1 | General transcription factor IIF subunit 1 Mus musculus | 116.06 | 6.5 | 57241 | 2 | 2 | 1 | 1 | 590315.31 | 583785.19 | 1 | 1 | 1 | 1 | 1559414 | 1 | 4705009 | 2914986.25 | 1 | 1 |
| Q9DBA9 | Gtf2h1 | General transcription factor IIH subunit 1 Mus musculus | 48.22 | 2.38 | 61851 | 1 | 1 | 295063.1625 | 436338.5 | 267319.59 | 244863.11 | 231731.45 | 1 | 1 | 371412.22 | 114382.96 | 326251.41 | 253511.7 | 1 | 306016.28 | 1 |
| Q9ESZ8 | Gtf2i | General transcription factor II-I Mus musculus | 383.49 | 15.43 | 112265 | 9 | 9 | 5702867 | 6604590.5 | 2522804 | 4596535.5 | 4069722.75 | 5555471 | 5441449.5 | 4152841.5 | 4548177.5 | 3915738.25 | 5409084.5 | 5016258 | 5183929 | 2565745 |
| Q8BMQ2 | Gtf3c4 | General transcription factor 3C polypeptide 4 Mus musculus | 48.54 | 2.33 | 91613 | 1 | 1 | 148779.95 | 95456.84 | 237425.92 | 144857.19 | 140392.62 | 273780.12 | 244024.67 | 272324.19 | 235925.58 | 177443.38 | 171055.56 | 285742.25 | 298756.91 | 203142.75 |
| D2XV59 | Gtpbp1 | GTP-binding protein 1 Rattus norvegicus | 447.47 | 17.94 | 72489 | 8 | 8 | 5781637 | 3632949 | 5440118 | 4444974 | 5251820 | 4017122.75 | 4343743 | 4069677.75 | 4122506 | 4260135.5 | 4933641.5 | 3861705 | 5160529.5 | 3664526.5 |
| Q99ME9 | Gtpbp4 | Nucleolar GTP-binding protein 1 Mus musculus | 640.33 | 24.76 | 74113 | 13 | 13 | 4191265.5 | 1856079.5 | 4486196.5 | 2227758 | 2976197 | 2453703 | 2388705.5 | 3686468 | 4367512.5 | 2560606.75 | 5717133.5 | 5738777 | 3530399 | 2627310.5 |
| P51840 | Gucy2e | Guanylyl cyclase GC-E Rattus norvegicus | 39.88 | 0.63 | 120801 | 1 | 1 | 770574 | 2047297.25 | 2362873.5 | 881445.12 | 876466.25 | 802174.38 | 1080329.62 | 2485625 | 981458.38 | 1237264.12 | 795538.19 | 1112836 | 1067708.5 | 2106802.25 |
| Q64520 | Guk1 | Guanylate kinase Mus musculus | 98.74 | 14.14 | 21918 | 2 | 2 | 1425565.25 | 676793.38 | 1614261.88 | 1302717 | 1146173 | 1525280.38 | 809121.94 | 1885358.75 | 1031260 | 1142847.38 | 1073313.25 | 1576739.88 | 1445729.88 | 1373958.88 |
| P12265 | Gusb | Beta-glucuronidase Mus musculus | 1304.21 | 44.44 | 74195 | 23 | 23 | 175019537 | 175104497 | 143682769 | 165715329 | 203578257 | 184248913 | 164555217 | 169493793 | 178724513 | 163199777 | 178242129 | 164749297 | 174815041 | 145520033 |
| Q3UHH8 | Gxylt1 | Glucoside xylosyltransferase 1 Mus musculus | 38.58 | 1.98 | 46491 | 1 | 1 | 385412.75 | 306356.34 | 428203.12 | 275930.5 | 1 | 270420.28 | 351606.0325 | 442849.38 | 352449.41 | 340705.06 | 1 | 377925.19 | 368252.5 | 1 |
| O08730 | Gyg1 | Glycogenin-1 Rattus norvegicus | 45.97 | 4.8 | 37378 | 1 | 1 | 3535684.75 | 3298503.5 | 3479268.125 | 3422287 | 3660597.25 | 3340132 | 1 | 1929546.5 | 1 | 1664164 | 3925949 | 4476207.5 | 1 | 1 |
| A2RRU1 | Gys1 | Glycogen [starch] synthase, muscle Rattus norvegicus | 64.7 | 1.9 | 84072 | 1 | 1 | 1 | 1617416.25 | 1889975.62 | 1317772.88 | 1 | 1341459.345 | 1327784.88 | 1231076.75 | 1533112.75 | 1273863 | 1118286.62 | 1325713.62 | 1145055.75 | 1449572 |
| P10922 | H1f0 | Histone H1.0 Mus musculus | 264.71 | 27.84 | 20861 | 6 | 6 | 6077676 | 5028575.5 | 5421710 | 8196977 | 6596668 | 5140076 | 5047117.5 | 5822851 | 4840035 | 6893709.5 | 6924676 | 5555862.5 | 6053688.5 | 7289347.5 |
| P01899 | H2-D1 | H-2 class I histocompatibility antigen, D-B alpha chain Mus musculus | 465.83 | 27.62 | 40836 | 0 | 9 | 94416209 | 100057817 | 97557457 | 82101145 | 98633273 | 87461977 | 74047865 | 86729377 | 84666001 | 83465537 | 89896345 | 80960305 | 95651649 | 94520185 |
| P01900 | H2-D1 | H-2 class I histocompatibility antigen, D-D alpha chain Mus musculus | 821.73 | 49.86 | 41110 | 8 | 16 | 141752129 | 156688449 | 144794481 | 127365297 | 153014273 | 133145489 | 112402753 | 129501857 | 126400217 | 123209249 | 133765249 | 119186737 | 136398545 | 136591089 |
| P14426 | H2-D1 | H-2 class I histocompatibility antigen, D-K alpha chain Mus musculus | 228.67 | 9.67 | 40620 | 0 | 4 | 76935561 | 88768785 | 78007345 | 74458721 | 84181505 | 70632345 | 62804249 | 67617417 | 65458549 | 67753905 | 73245777 | 63252033 | 74996817 | 76354569 |
| P01901 | H2-K1 | H-2 class I histocompatibility antigen, K-B alpha chain Mus musculus | 226.74 | 9.21 | 41302 | 0 | 5 | 69263969 | 77587809 | 70712513 | 67576433 | 76025153 | 63816409 | 58201057 | 59927801 | 59760469 | 61931045 | 66310081 | 57656105 | 69091313 | 68949009 |
| P01902 | H2-K1 | H-2 class I histocompatibility antigen, K-D alpha chain Mus musculus | 881.15 | 45.38 | 41490 | 10 | 16 | 96228993 | 98687489 | 100711857 | 77917329 | 95276449 | 78757441 | 74367601 | 81868753 | 77528273 | 75264657 | 79332945 | 73915873 | 86289529 | 77255185 |
| P01897 | H2-L | H-2 class I histocompatibility antigen, L-D alpha chain Mus musculus | 610.29 | 35.08 | 40711 | 1 | 12 | 97031153 | 107556897 | 103497809 | 84638721 | 106488073 | 90470681 | 76345009 | 93851345 | 92225241 | 85695809 | 92115417 | 86609713 | 102593689 | 101438465 |
| P01898 | H2-Q10 | H-2 class I histocompatibility antigen, Q10 alpha chain Mus musculus | 224.71 | 12 | 37251 | 0 | 5 | 31418471 | 41678173 | 36969873 | 29887583 | 37888525 | 30271239 | 25399051 | 31475941 | 32206549 | 27630075 | 30155445 | 31049273 | 36220125 | 33341589 |
| P06339 | H2-T23 | H-2 class I histocompatibility antigen, D-37 alpha chain Mus musculus | 236.87 | 10.64 | 40875 | 0 | 4 | 49192085 | 52357897 | 48947249 | 45976805 | 54361713 | 43278765 | 40954385 | 41889553 | 40893653 | 43155413 | 44860493 | 38484013 | 47614577 | 48557841 |
| P27661 | H2afx | Histone H2AX Mus musculus | 360.81 | 52.45 | 15143 | 0 | 5 | 229847233 | 215684609 | 251903953 | 234172321 | 248695969 | 226502321 | 235058369 | 176458241 | 197791089 | 315601665 | 265790497 | 263473185 | 262352065 | 245582721 |
| Q9QZQ8 | H2afy | Core histone macro-H2A.1 Mus musculus | 565.73 | 37.1 | 39735 | 10 | 10 | 13904306 | 16380042 | 16779923 | 13020004 | 19447685 | 21471109 | 17988427 | 16172781 | 18675089 | 21882153 | 19602819 | 19257051 | 19166083 | 19841315 |
| P0C0S6 | H2afz | Histone H2A.Z Mus musculus | 293.14 | 51.56 | 13553 | 3 | 5 | 235438561 | 223822273 | 266783489 | 242506881 | 251473025 | 239993553 | 248954081 | 182917041 | 207090113 | 328905921 | 272846721 | 281114849 | 281000193 | 255513825 |
| P02301 | H3f3c | Histone H3.3C Mus musculus | 368.36 | 52.21 | 15315 | 1 | 7 | 1011221633 | 887928705 | 971219457 | 1076199681 | 1066173505 | 891644545 | 883897217 | 762019585 | 835788609 | 1020548737 | 1003031041 | 1026841985 | 970579713 | 913473665 |
| Q9D3B1 | Hacd2 | Very-long-chain (3R)-3-hydroxyacyl-CoA dehydratase 2 Mus musculus | 41.48 | 4.72 | 28402 | 1 | 1 | 1500543.62 | 370147.94 | 374076.28 | 305479.94 | 522270.91 | 1 | 1 | 400976.41 | 491819.47 | 344054.09 | 1 | 433548.72 | 1 | 446524.97 |
| Q8K2C9 | Hacd3 | Very-long-chain (3R)-3-hydroxyacyl-CoA dehydratase 3 Mus musculus | 41.54 | 2.76 | 43131 | 1 | 1 | 1488737.12 | 437472.41 | 527302.81 | 264417.53 | 471759.06 | 871191.81 | 346660.56 | 567195.31 | 565371.62 | 587604.825 | 665647.25 | 581872.06 | 849130.56 | 613964.5 |
| Q61425 | Hadh | Hydroxyacyl-coenzyme A dehydrogenase, mitochondrial Mus musculus | 375.62 | 36.94 | 34464 | 7 | 7 | 12153773 | 15731794 | 13339585 | 13572125 | 14204149 | 14106337 | 12412671 | 12662339 | 13468379 | 14472145 | 14114807 | 14270136 | 14478565 | 12760018 |
| Q8BMS1 | Hadha | Trifunctional enzyme subunit alpha, mitochondrial Mus musculus | 1110.22 | 38.66 | 82670 | 19 | 19 | 28916769 | 24112179 | 27524993 | 26538753 | 25760241 | 28232151 | 29585693 | 29986777 | 30998355 | 24772967 | 31427533 | 32221645 | 33350231 | 31124985 |
| Q99JY0 | Hadhb | Trifunctional enzyme subunit beta, mitochondrial Mus musculus | 447.59 | 21.26 | 51386 | 9 | 9 | 18308545 | 19479009 | 20810149 | 20087425 | 20413189 | 18963227 | 22021683 | 18546233 | 21062735 | 22372113 | 27326747 | 25261597 | 23451767 | 23743363 |
| O35952 | Hagh | Hydroxyacylglutathione hydrolase, mitochondrial Rattus norvegicus | 57.78 | 6.15 | 34109 | 2 | 2 | 2686735.5 | 851342.94 | 3501160.25 | 1167546.12 | 1244698.62 | 3014296 | 2624949.25 | 4164661.5 | 3274113.75 | 740106.19 | 2876432.5 | 887661.44 | 3251958.5 | 1100576.62 |
| Q61035 | Hars | Histidine--tRNA ligase, cytoplasmic Mus musculus | 1010.52 | 48.33 | 57432 | 18 | 18 | 31521269 | 29842935 | 31278805 | 29822403 | 29324697 | 31185113 | 27873379 | 31531973 | 30662169 | 30857407 | 30482035 | 30482371 | 29814555 | 31034777 |
| Q8BY71 | Hat1 | Histone acetyltransferase type B catalytic subunit Mus musculus | 340.62 | 24.52 | 49278 | 7 | 7 | 5537363.5 | 4287962 | 3822267.25 | 4729448 | 4515848 | 3592670.5 | 3325248.5 | 3529465 | 3122781 | 2976022 | 4724355.5 | 4819829 | 3491253 | 3809642.5 |
| Q8BFT2 | Haus4 | HAUS augmin-like complex subunit 4 Mus musculus | 44.95 | 7.16 | 42197 | 1 | 1 | 304402.09 | 227842.42 | 302441.635 | 330933.56 | 346588.47 | 323762.72 | 350493.09 | 168091.64 | 309216.25 | 260909.2 | 373263.66 | 490128.94 | 375260.56 | 299100.47 |
| Q9D786 | Haus5 | HAUS augmin-like complex subunit 5 Mus musculus | 107.36 | 6.79 | 69572 | 2 | 2 | 127841.55 | 262327.03 | 127895.41 | 165718.055 | 144808.23 | 229508.61 | 121149.44 | 238561.9575 | 303465.59 | 300124.19 | 166776.25 | 226495.2633 | 384161.25 | 128548.29 |
| Q69ZS7 | Hbs1l | HBS1-like protein Mus musculus | 76.01 | 3.67 | 75100 | 2 | 2 | 1 | 1 | 289044.81 | 283889.75 | 273839.38 | 325917.09 | 372204.06 | 237434.31 | 324246.3425 | 361429.91 | 388208.06 | 438782.38 | 341281.25 | 307390.81 |
| P53702 | Hccs | Cytochrome c-type heme lyase Mus musculus | 71.53 | 7.35 | 30978 | 2 | 2 | 1115353 | 972654.62 | 1113247.25 | 1247881.25 | 1202632.75 | 1087086.88 | 1131189.25 | 989811.75 | 1174971.38 | 1186920.88 | 1162849.12 | 1352464.38 | 1351521.62 | 1381771.5 |
| Q61191 | Hcfc1 | Host cell factor 1 Mus musculus | 644.41 | 11.34 | 210437 | 13 | 13 | 6203405.5 | 7394230.5 | 6269429.5 | 9585636 | 6893013.5 | 9244245 | 10315910 | 8604170 | 8803035 | 10148131 | 6954694.5 | 8376769.5 | 8338137 | 6992164.5 |
| P08103 | Hck | Tyrosine-protein kinase HCK Mus musculus | 202.92 | 13.17 | 59129 | 5 | 5 | 2724987.5 | 2864456 | 4521957 | 3100563.25 | 4244812 | 3448394.5 | 3263131 | 3425672 | 2757609 | 3182085 | 3121371.5 | 3245645.5 | 3414840.5 | 2895579.75 |
| P49710 | Hcls1 | Hematopoietic lineage cell-specific protein Mus musculus | 412.3 | 18.52 | 54240 | 8 | 8 | 11973537 | 10518004 | 12753000 | 12874864 | 16029502 | 17020421 | 11226291 | 10964757 | 12024353 | 16137784 | 14628815 | 15192635 | 19529609 | 13509808 |
| O09106 | Hdac1 | Histone deacetylase 1 Mus musculus | 633.27 | 35.48 | 55075 | 8 | 11 | 32608535 | 33157919 | 33158403 | 27421663 | 30621875 | 32880409 | 32784507 | 34355085 | 35725089 | 30886565 | 32097017 | 31781939 | 32375215 | 29484989 |
| P70288 | Hdac2 | Histone deacetylase 2 Mus musculus | 439.68 | 24.59 | 55302 | 5 | 8 | 20816513 | 19463119 | 18366981 | 16977519 | 21372809 | 20917677 | 18145683 | 18522179 | 20393679 | 19537805 | 19658933 | 19349131 | 19637407 | 19662503 |
| O88895 | Hdac3 | Histone deacetylase 3 Mus musculus | 72.08 | 4.01 | 48364 | 1 | 1 | 1348115.283 | 1613038.25 | 1429792.62 | 1183903.88 | 1165726.38 | 1386062 | 1 | 1 | 1678652 | 1301631.12 | 982481.5 | 1357597.62 | 1350723.75 | 1251556.62 |
| P51859 | Hdgf | Hepatoma-derived growth factor Mus musculus | 448.32 | 48.95 | 26269 | 8 | 9 | 21098055 | 19599647 | 19918309 | 19440521 | 21249851 | 21098271 | 24100527 | 19624253 | 23167539 | 21451037 | 22221985 | 17393649 | 21748791 | 19984683 |
| Q3UMU9 | Hdgfl2 | Hepatoma-derived growth factor-related protein 2 Mus musculus | 88.95 | 3.14 | 74291 | 1 | 2 | 2615269.5 | 2156333 | 1839871.12 | 2171031.75 | 2839698.5 | 1854955.62 | 3048079.75 | 1878729 | 2677314.25 | 2830559.25 | 2538459.75 | 224036.97 | 2151884.75 | 2531987.25 |
| Q3UGR5 | Hdhd2 | Haloacid dehalogenase-like hydrolase domain-containing protein 2 Mus musculus | 106.24 | 11.2 | 28730 | 2 | 2 | 1601858.88 | 1578865.75 | 1937501.25 | 1620819.12 | 1657198.5 | 1907892.62 | 1330143.62 | 1688784.25 | 1654707.88 | 1389900.5 | 1669670.75 | 1631931 | 1845524.75 | 1762162.88 |
| Q91WM2 | Hdhd5 | Haloacid dehalogenase-like hydrolase domain-containing 5 Mus musculus | 44.63 | 3.82 | 46306 | 1 | 1 | 360665.69 | 272300.16 | 435080.22 | 430365.12 | 235908.73 | 258892.59 | 498333.28 | 286657.31 | 301274.25 | 204102.06 | 294954.9767 | 318588.62 | 282919.56 | 283356.75 |
| Q8VDJ3 | Hdlbp | Vigilin Mus musculus | 1592.19 | 29.02 | 141743 | 29 | 29 | 23232933 | 23506171 | 26037487 | 24295201 | 24547027 | 24577607 | 27614497 | 22064999 | 26684741 | 26717391 | 28245225 | 24331689 | 28401115 | 26681447 |
| Q8BQM4 | Heatr3 | HEAT repeat-containing protein 3 Mus musculus | 210.68 | 5.45 | 74305 | 3 | 3 | 2602077.75 | 1358947.62 | 2857003 | 2405509.25 | 1953410.5 | 2175822 | 2614771.5 | 2156530.5 | 2586344.75 | 2669260.5 | 2073216.62 | 2731540 | 2569264.25 | 2077539 |
| Q5PRF0 | Heatr5a | HEAT repeat-containing protein 5A Mus musculus | 55.06 | 0.59 | 219893 | 1 | 1 | 460746.09 | 1 | 373829 | 1 | 428216.5 | 416232.16 | 406993.08 | 366478.66 | 444240.5 | 401021 | 320531.09 | 324188.26 | 313924.22 | 338109.47 |
| Q8C547 | Heatr5b | HEAT repeat-containing protein 5B Mus musculus | 63.18 | 0.82 | 224319 | 1 | 1 | 156976.59 | 135018.84 | 194960.61 | 217384.62 | 220957.33 | 152532.19 | 167774.41 | 125866.74 | 128925.16 | 249098.42 | 258706.38 | 245411.03 | 218303.77 | 259222.94 |
| A1EC95 | Heatr6 | HEAT repeat-containing protein 6 Rattus norvegicus | 82.91 | 2.16 | 136505 | 2 | 2 | 577690.69 | 396189.25 | 377223.16 | 446741.7975 | 435864.09 | 352418.34 | 363479.575 | 422473.09 | 369836.62 | 309190.25 | 312233.9 | 362360.72 | 237844.64 | 336496.34 |
| Q69ZR2 | Hectd1 | E3 ubiquitin-protein ligase HECTD1 Mus musculus | 121 | 0.92 | 290086 | 2 | 2 | 278612.47 | 408957.69 | 258623.62 | 419540.75 | 365158.09 | 331282.16 | 261558.98 | 356152.62 | 282231.62 | 292927.38 | 287809.69 | 281069.25 | 315800.22 | 248898.27 |
| Q6NVF4 | Helb | DNA helicase B Mus musculus | 40.58 | 0.74 | 121471 | 1 | 1 | 1 | 1 | 1 | 1309828.38 | 1 | 1 | 1 | 1 | 1 | 1 | 1221964.12 | 286485.69 | 1 | 1 |
| Q60848 | Hells | Lymphocyte-specific helicase Mus musculus | 625.19 | 15.71 | 95126 | 11 | 11 | 1720837.25 | 1543517.75 | 4692735.5 | 2017514.5 | 3456545 | 2566515 | 3326569.75 | 3020546.5 | 3758815 | 2289546.5 | 2276010 | 3174233 | 3772057 | 3904321.5 |
| E9QAM5 | Helz2 | Helicase with zinc finger domain 2 Mus musculus | 151.11 | 2.07 | 331563 | 4 | 4 | 379286.59 | 418748.47 | 384684.03 | 307553.22 | 292926.03 | 180810.45 | 341476.72 | 227275 | 215808.92 | 241342.7725 | 312501.56 | 320097.28 | 280820.34 | 158677.58 |
| P29416 | Hexa | Beta-hexosaminidase subunit alpha Mus musculus | 1086.12 | 42.23 | 60613 | 6 | 16 | 70220577 | 94596145 | 76982145 | 71568745 | 87501009 | 66215849 | 63692437 | 76452817 | 68969409 | 63487673 | 73454281 | 71298273 | 68323281 | 81361849 |
| P20060 | Hexb | Beta-hexosaminidase subunit beta Mus musculus | 738.72 | 39.37 | 61116 | 16 | 16 | 55997421 | 70052937 | 50176317 | 45176281 | 61938809 | 45185985 | 41383497 | 48473765 | 44602829 | 43844301 | 49839685 | 44752501 | 52119533 | 53569377 |
| Q99L13 | Hibadh | 3-hydroxyisobutyrate dehydrogenase, mitochondrial Mus musculus | 245.21 | 23.28 | 35440 | 5 | 5 | 2553083.25 | 2951650 | 1218244.12 | 2513776 | 3179769.5 | 2978749.5 | 3115297.75 | 3002447.75 | 3304830.5 | 3467776.25 | 3401932 | 3599280 | 3304719 | 2948626.75 |
| Q8QZS1 | Hibch | 3-hydroxyisobutyryl-CoA hydrolase, mitochondrial Mus musculus | 168.89 | 11.43 | 43038 | 3 | 3 | 2631392.25 | 2949682.5 | 3708862.5 | 2917451.5 | 3732863.25 | 3848548.5 | 2381056.25 | 3158293.75 | 4214499.5 | 3886919.5 | 2481279.25 | 3248999.5 | 4125406.5 | 3764821.5 |
| Q8VH49 | Higd1a | HIG1 domain family member 1A, mitochondrial Rattus norvegicus | 147.95 | 43.01 | 10301 | 2 | 2 | 813655.38 | 698692.56 | 782062.25 | 630587.57 | 227940.09 | 264727.19 | 816548.88 | 544879.38 | 808258.88 | 831105.56 | 1162508.75 | 1433592.38 | 1245085.25 | 1104648.75 |
| P62959 | Hint1 | Histidine triad nucleotide-binding protein 1 Rattus norvegicus | 163.67 | 35.71 | 13777 | 3 | 3 | 41841437 | 48005725 | 47729601 | 49812909 | 48736601 | 38098337 | 43058569 | 42895177 | 39513597 | 41674329 | 43209365 | 43843257 | 39779449 | 45466081 |
| Q9D0S9 | Hint2 | Histidine triad nucleotide-binding protein 2, mitochondrial Mus musculus | 42.21 | 7.98 | 17320 | 1 | 1 | 1147041.25 | 1143952.62 | 1336661.75 | 1596866.62 | 1333003.38 | 1353291.5 | 1384808.5 | 1128504.12 | 1541515.25 | 1537812.75 | 1470471.88 | 1301628.5 | 1312059 | 1519850.12 |
| Q8VD75 | Hip1 | Huntingtin-interacting protein 1 Mus musculus | 61.07 | 2.24 | 115202 | 2 | 2 | 2968139.75 | 1731633.38 | 1329137.38 | 3062914 | 3098671 | 2605225.5 | 1402042.88 | 837240.62 | 1449326.88 | 1083401.88 | 2393296.75 | 3540937.75 | 1722002.88 | 2423584.5 |
| Q8BLH7 | Hirip3 | HIRA-interacting protein 3 Mus musculus | 142.01 | 3.66 | 65215 | 2 | 2 | 1 | 1 | 796076.69 | 711181.25 | 668247.88 | 1 | 1 | 1 | 1 | 685636.81 | 1 | 1 | 1 | 1 |
| P43275 | Hist1h1a | Histone H1.1 Mus musculus | 314.39 | 34.27 | 21785 | 4 | 6 | 73412177 | 52340465 | 53922981 | 63098833 | 49039541 | 43580101 | 50579665 | 50396945 | 43592625 | 51534285 | 47865645 | 54792421 | 46160089 | 42322573 |
| P43276 | Hist1h1b | Histone H1.5 Mus musculus | 292.83 | 22.42 | 22576 | 6 | 7 | 10660957 | 10001917 | 10339258 | 11442148 | 11154373 | 9470985 | 10959836 | 8551160 | 8919582 | 14092127 | 10818876 | 13279295 | 11927869 | 11001427 |
| P15864 | Hist1h1c | Histone H1.2 Mus musculus | 549.35 | 33.02 | 21267 | 3 | 10 | 142196337 | 107721441 | 120461889 | 129245409 | 101859425 | 98974913 | 120665793 | 103185209 | 101573985 | 126201761 | 105911521 | 127566361 | 106883889 | 93249721 |
| P43277 | Hist1h1d | Histone H1.3 Mus musculus | 504 | 24.43 | 22100 | 1 | 9 | 146072417 | 110155473 | 120200137 | 130880401 | 102216593 | 99597481 | 122041881 | 106140105 | 101935497 | 127680449 | 106550105 | 127255105 | 105382785 | 93908529 |
| P43274 | Hist1h1e | Histone H1.4 Mus musculus | 498.91 | 24.66 | 21977 | 2 | 8 | 113203889 | 80511689 | 90889553 | 98874025 | 73935537 | 72272057 | 89940497 | 79847713 | 77094625 | 93958289 | 80011841 | 94763697 | 77709489 | 71503713 |
| P15865 | Hist1h1e | Histone H1.4 Rattus norvegicus | 455.21 | 24.66 | 21987 | 0 | 8 | 130091425 | 95335217 | 108519433 | 117127905 | 89543865 | 88441761 | 109106625 | 92637841 | 91738593 | 113448873 | 94004881 | 113480209 | 93705753 | 83229121 |
| C0HKE1 | Hist1h2ab | Histone H2A type 1-B Mus musculus | 507.43 | 63.85 | 14135 | 1 | 7 | 231118113 | 216707329 | 253135185 | 234484961 | 250703665 | 228225521 | 236704129 | 176458241 | 198126097 | 317765441 | 267852641 | 265767617 | 265323969 | 247604353 |
| P10853 | Hist1h2bf | Histone H2B type 1-F/J/L Mus musculus | 1153.51 | 55.56 | 13936 | 2 | 18 | 2311764993 | 2088850177 | 1691605505 | 1780771073 | 1772783873 | 1521681665 | 1719559297 | 1880200961 | 1436857089 | 1671866241 | 1596670337 | 1497583489 | 1423864065 | 1678264833 |
| P10854 | Hist1h2bm | Histone H2B type 1-M Mus musculus | 1109.6 | 55.56 | 13936 | 2 | 17 | 2307947009 | 2087647233 | 1690756609 | 1778428161 | 1768388353 | 1521673217 | 1718052097 | 1880057857 | 1436131329 | 1670938881 | 1597435905 | 1497685249 | 1424740993 | 1677980161 |

| | | | | | | | | | | | | | | | | | | | | | |
|---|---|---|---|---|---|---|---|---|---|---|---|---|---|---|---|---|---|---|---|---|---|
| P68433 | Hist1h3a | Histone H3.1 Mus musculus | 547.58 | 52.21 | 15404 | 1 | 9 | 1012020993 | 888395521 | 972652545 | 1078272513 | 1066560641 | 892698881 | 884519681 | 762588673 | 835778753 | 1021285441 | 1006867009 | 1026898689 | 971209089 | 913959489 |
| P84228 | Hist1h3b | Histone H3.2 Mus musculus | 513.57 | 52.94 | 15388 | 2 | 9 | 1010180225 | 887055361 | 970322305 | 1074411521 | 1064539265 | 890050369 | 882928897 | 761630209 | 834671617 | 1018985217 | 1001479809 | 1025140865 | 969170753 | 912129921 |
| P62804 | Hist1h4b | Histone H4 Rattus norvegicus | 1131.95 | 58.25 | 11367 | 13 | 16 | 4833988609 | 2000589441 | 2137045889 | 1928947969 | 2121537153 | 1806478977 | 1684927361 | 1617567873 | 1833690625 | 2107090305 | 1791839233 | 1792816641 | 1988163585 | 2023052161 |
| P0CC09 | Hist2h2aa3 | Histone H2A type 2-A Rattus norvegicus | 545.2 | 58.46 | 14095 | 0 | 7 | 232415809 | 218299201 | 257881473 | 244310337 | 255039889 | 233198401 | 238218177 | 179464129 | 198844097 | 320439201 | 272293537 | 273284481 | 270955937 | 249004465 |
| Q64522 | Hist2h2ab | Histone H2A type 2-B Mus musculus | 404.79 | 52.31 | 14013 | 0 | 5 | 65926153 | 43979121 | 106511801 | 92464297 | 88569009 | 100197417 | 71374041 | 42014825 | 80647337 | 136808817 | 117431937 | 140340849 | 123040849 | 98975033 |
| P17710 | Hk1 | Hexokinase-1 Mus musculus | 1409.73 | 30.6 | 108303 | 4 | 24 | 51614469 | 58252017 | 56549513 | 52917917 | 53608305 | 59347841 | 66062785 | 61506405 | 65622465 | 66010489 | 54835997 | 56745281 | 54734325 | 58304449 |
| O08528 | Hk2 | Hexokinase-2 Mus musculus | 1259.89 | 33.04 | 102535 | 18 | 23 | 36978961 | 40303117 | 41025301 | 33905165 | 36865553 | 46153885 | 49299353 | 45904545 | 46528041 | 47517737 | 33184919 | 35763957 | 36774897 | 36253249 |
| Q3TRM8 | Hk3 | Hexokinase-3 Mus musculus | 2099.18 | 46.85 | 100101 | 13 | 30 | 111634473 | 134472689 | 119574729 | 111517449 | 118247505 | 113794553 | 107501257 | 111184673 | 103951961 | 107363473 | 108582865 | 94170505 | 105660569 | 109304409 |
| Q91W97 | Hkdc1 | Putative hexokinase HKDC1 Mus musculus | 161.04 | 3.72 | 102259 | 1 | 4 | 11155605 | 13182875 | 13107003 | 10640257 | 11549691 | 12545467 | 16006957 | 12990197 | 14124487 | 16178127 | 11411110 | 12041853 | 11570603 | 15417217 |
| Q9D8V0 | Hm13 | Minor histocompatibility antigen H13 Mus musculus | 278.34 | 16.4 | 41748 | 6 | 6 | 17344729 | 18118161 | 16397292 | 14235864 | 17362193 | 21622459 | 19333771 | 21930545 | 19732055 | 19179163 | 20518239 | 19863279 | 22094347 | 21011995 |
| P22907 | Hmbs | Porphobilinogen deaminase Mus musculus | 319.45 | 25.21 | 39344 | 5 | 5 | 3290961.25 | 4298848 | 4512083 | 5402619.5 | 3872265.75 | 4147303.75 | 5003795 | 4728411.5 | 4817332 | 4623870 | 4998648.5 | 4942670 | 4478314.5 | 2995520 |
| P17095 | Hmga1 | High mobility group protein HMG-I/HMG-Y Mus musculus | 45.7 | 14.95 | 11614 | 1 | 1 | 911566.44 | 658399.25 | 938192.88 | 1175260.38 | 868148.69 | 818665.31 | 1033552.31 | 661255.44 | 900902.38 | 1304006.38 | 1062800.25 | 1075640.25 | 1125129.5 | 907055.19 |
| P63158 | Hmgb1 | High mobility group protein B1 Mus musculus | 423.23 | 31.16 | 24894 | 6 | 7 | 27314087 | 22562343 | 23011071 | 23944357 | 31560505 | 26056929 | 27973265 | 12699752 | 23047601 | 27359769 | 25433669 | 22419935 | 17817363 | 22482635 |
| P30681 | Hmgb2 | High mobility group protein B2 Mus musculus | 509.12 | 29.05 | 24162 | 7 | 8 | 50777745 | 51925705 | 49958501 | 67947057 | 60783049 | 47121305 | 56739953 | 51936301 | 50742557 | 64643793 | 56295209 | 49047521 | 49111573 | 56093765 |
| O54879 | Hmgb3 | High mobility group protein B3 Mus musculus | 107.64 | 13 | 23010 | 2 | 2 | 5262773.5 | 4866769.5 | 4656933 | 5142224 | 4846237 | 4914099 | 4847400.5 | 4529970.5 | 4639146.5 | 5242811 | 5069471 | 5139917.5 | 4605610 | 5772732 |
| P38060 | Hmgcl | Hydroxymethylglutaryl-CoA lyase, mitochondrial Mus musculus | 605.33 | 46.77 | 34239 | 10 | 10 | 20361335 | 25321399 | 25609585 | 17527379 | 23998657 | 24400257 | 28256333 | 28503143 | 28488217 | 27767281 | 28131443 | 21966409 | 29815053 | 28196835 |
| Q8JZK9 | Hmgcs1 | Hydroxymethylglutaryl-CoA synthase, cytoplasmic Mus musculus | 1358.43 | 53.27 | 57569 | 20 | 21 | 7472597 | 6713622 | 4069837.25 | 6253870.5 | 3350192 | 3717322.25 | 4321885 | 5066904.5 | 3058924.5 | 3228456.5 | 5877180 | 5757336.5 | 5943686 | 3781959.5 |
| Q9JL35 | Hmgn5 | High mobility group nucleosome-binding domain-containing protein 5 Mus musculus | 301.6 | 20.2 | 45344 | 6 | 6 | 7709765 | 8175025.5 | 9029629 | 7347658.5 | 10840799 | 10582521 | 9666949 | 6443514 | 9273328 | 10787695 | 8652645 | 9479369 | 10300777 | 12010579 |
| O70252 | Hmox2 | Heme oxygenase 2 Mus musculus | 417.53 | 38.1 | 35739 | 8 | 8 | 16528945 | 14790119 | 13183153 | 13672824 | 13986250 | 13560446 | 15103705 | 15098434 | 12472848 | 11462087 | 14179128 | 13101698 | 13942378 | 13263475 |
| Q9CX86 | Hnrnpa0 | Heterogeneous nuclear ribonucleoprotein A0 Mus musculus | 501.93 | 39.02 | 30530 | 8 | 8 | 18543561 | 23033749 | 22235577 | 24538709 | 22462929 | 23200993 | 19196487 | 21514069 | 24237727 | 23307227 | 20744725 | 15381837 | 18922743 | 20393365 |
| P04256 | Hnrnpa1 | Heterogeneous nuclear ribonucleoprotein A1 Rattus norvegicus | 1393.54 | 63.12 | 34212 | 15 | 19 | 25362331 | 29835313 | 38211317 | 30269097 | 30000265 | 28990019 | 31018687 | 25067601 | 30209053 | 36903293 | 37378741 | 33931981 | 42623065 | 33139911 |
| O88569 | Hnrnpa2b1 | Heterogeneous nuclear ribonucleoproteins A2/B1 Mus musculus | 1699.02 | 82.44 | 37403 | 25 | 25 | 139062753 | 131963097 | 176265057 | 137647809 | 127125321 | 148214625 | 133536713 | 169648529 | 176090257 | 147481425 | 140441617 | 153804961 | 162882209 | 137088705 |
| Q6URK4 | Hnrnpa3 | Heterogeneous nuclear ribonucleoprotein A3 Rattus norvegicus | 1355.72 | 48.55 | 39652 | 19 | 22 | 85827065 | 91923281 | 122147929 | 86692369 | 85959385 | 108092305 | 100458529 | 110027281 | 114058409 | 114531945 | 101202833 | 102456785 | 107699905 | 102917161 |
| Q99020 | Hnrnpab | Heterogeneous nuclear ribonucleoprotein A/B Mus musculus | 498.94 | 30.53 | 30831 | 8 | 9 | 23145561 | 25227909 | 26947213 | 24505421 | 26309121 | 23840677 | 23795749 | 24882661 | 23787067 | 25588241 | 25712285 | 26032819 | 25577095 | 27515735 |
| G3V9R8 | Hnrnpc | Heterogeneous nuclear ribonucleoprotein C Rattus norvegicus | 820.63 | 42.28 | 32857 | 1 | 13 | 91587977 | 96398145 | 97646769 | 95248209 | 110042449 | 103368089 | 100603161 | 84349089 | 103978297 | 103453545 | 101158585 | 100183697 | 105476737 | 105677889 |
| Q9Z204 | Hnrnpc | Heterogeneous nuclear ribonucleoproteins C1/C2 Mus musculus | 783.56 | 44.41 | 34385 | 1 | 13 | 90037665 | 95079089 | 95737849 | 93441361 | 108013793 | 101317553 | 98931673 | 82930073 | 102028105 | 101482785 | 98938321 | 98120537 | 103044225 | 103251457 |
| Q60668 | Hnrnpd | Heterogeneous nuclear ribonucleoprotein D0 Mus musculus | 495.15 | 25.63 | 38354 | 8 | 9 | 8594504 | 3998791.75 | 7767368.5 | 10836933 | 13714794 | 13134239 | 10626843 | 11627727 | 6423377 | 13462923 | 9546326 | 16262345 | 9810471 | 12181420 |
| Q3SWU3 | Hnrnpdl | Heterogeneous nuclear ribonucleoprotein D-like Rattus norvegicus | 135.9 | 16.46 | 35294 | 1 | 3 | 1 | 1 | 1 | 1 | 1 | 301879.19 | 386733.03 | 1 | 1 | 1 | 338917.44 | 1 | 1 | 1 |
| Q794E4 | Hnrnpf | Heterogeneous nuclear ribonucleoprotein F Rattus norvegicus | 935.53 | 54.46 | 45730 | 12 | 14 | 85709761 | 91869393 | 90898321 | 96375137 | 105867153 | 93052113 | 92435633 | 91595393 | 84296849 | 99282193 | 98842009 | 78288873 | 90732257 | 96781217 |
| O35737 | Hnrnph1 | Heterogeneous nuclear ribonucleoprotein H Mus musculus | 740.41 | 37.86 | 49199 | 6 | 12 | 74856865 | 82025825 | 78275569 | 89255617 | 91103553 | 81392993 | 75508193 | 76146649 | 70749913 | 85986937 | 85157001 | 68748177 | 80490545 | 84196521 |
| P70333 | Hnrnph2 | Heterogeneous nuclear ribonucleoprotein H2 Mus musculus | 596.54 | 32.74 | 49280 | 5 | 10 | 36482773 | 41749225 | 36871281 | 45399625 | 43615345 | 41624201 | 40408301 | 39480829 | 37253961 | 47124917 | 41499981 | 34847185 | 41364757 | 43312457 |
| P61979 | Hnrnpk | Heterogeneous nuclear ribonucleoprotein K Mus musculus | 1469.61 | 63.28 | 50976 | 23 | 23 | 144774369 | 158442625 | 157376049 | 163918337 | 162241809 | 162417569 | 165292833 | 148988993 | 156737985 | 180751009 | 163519617 | 157960337 | 160792321 | 167525665 |
| F1LQ48 | Hnrnpl | Heterogeneous nuclear ribonucleoprotein L Rattus norvegicus | 1366.59 | 67.09 | 67903 | 22 | 22 | 71784449 | 80882889 | 91346017 | 94987377 | 99295065 | 95983153 | 97306425 | 71616209 | 94754441 | 102962033 | 97486393 | 94863377 | 96682481 | 84017913 |
| Q921F4 | Hnrnpll | Heterogeneous nuclear ribonucleoprotein L-like Mus musculus | 554.52 | 26.23 | 64125 | 10 | 10 | 9680591 | 9633316 | 9131778 | 11410217 | 11651424 | 11106570 | 10627917 | 11698749 | 10567780 | 11328315 | 10621412 | 9201046 | 9864990 | 12069861 |
| Q9D0E1 | Hnrnpm | Heterogeneous nuclear ribonucleoprotein M Mus musculus | 1663.1 | 47.74 | 77649 | 27 | 28 | 85412833 | 88315073 | 105917417 | 107523809 | 106907113 | 106957641 | 112233681 | 82838609 | 105829081 | 133405825 | 105118153 | 113590441 | 107935721 | 105028745 |
| Q6IMY8 | Hnrnpu | Heterogeneous nuclear ribonucleoprotein U Rattus norvegicus | 2107.2 | 43.36 | 87732 | 31 | 32 | 164142273 | 185724625 | 181767889 | 187177761 | 182851089 | 164649601 | 191990657 | 186404849 | 179226209 | 198744673 | 174067681 | 153309185 | 175300497 | 174789873 |
| Q8VDM6 | Hnrnpul1 | Heterogeneous nuclear ribonucleoprotein U-like protein 1 Mus musculus | 398.51 | 17.46 | 96002 | 10 | 10 | 7594204 | 6369281 | 7597508 | 6940959 | 8571770 | 8101624.5 | 5428014.5 | 8059455 | 7050722 | 8312756.5 | 7362905 | 6058577 | 6903094 | 7674782 |
| Q00PI9 | Hnrnpul2 | Heterogeneous nuclear ribonucleoprotein U-like protein 2 Mus musculus | 568.93 | 17.58 | 84940 | 11 | 11 | 33132101 | 38198973 | 37533853 | 38153349 | 38778877 | 38077053 | 37728417 | 36258373 | 40292757 | 42195929 | 37476961 | 38079237 | 37861417 | 41239557 |
| Q8BUK6 | Hook3 | Protein Hook homolog 3 Mus musculus | 50.94 | 1.95 | 83218 | 1 | 1 | 250239.77 | 1 | 429937.62 | 1 | 1 | 309082.53 | 325163.3175 | 506869.59 | 286235.31 | 198465.84 | 271503.04 | 304517.25 | 280564.59 | 229427.28 |
| Q3TEA8 | Hp1bp3 | Heterochromatin protein 1-binding protein 3 Mus musculus | 499.96 | 22.74 | 60867 | 9 | 9 | 15570894 | 17605471 | 16048955 | 19989929 | 18067201 | 17320645 | 17404155 | 17778505 | 17204513 | 18767909 | 17912385 | 15803711 | 17772153 | 18429689 |
| P62748 | Hpcal1 | Hippocalcin-like protein 1 Mus musculus | 128.53 | 16.06 | 22338 | 3 | 3 | 4420884 | 4650699 | 4425670 | 4190072 | 3879160 | 2455384 | 3967564.25 | 3967655 | 4029262.5 | 3569625.75 | 4431663 | 3926056.25 | 3537659.25 | 4465342.5 |
| Q8CFE2 | Hpf1 | Histone PARylation factor 1 Mus musculus | 195.1 | 15.32 | 39291 | 4 | 4 | 5105827 | 4907595.5 | 4066558 | 4904229 | 3923714.5 | 2967268.5 | 4952774 | 4430104 | 5290509.5 | 4465759 | 5230712.5 | 5534289 | 4727379 | 4990795 |
| P00493 | Hprt1 | Hypoxanthine-guanine phosphoribosyltransferase Mus musculus | 592.38 | 69.72 | 24570 | 12 | 12 | 37369653 | 39280897 | 42360637 | 37940021 | 36330717 | 36037993 | 38332769 | 38624197 | 38121989 | 35355561 | 36777453 | 38099045 | 40002133 | 37376393 |
| P59438 | Hps5 | Hermansky-Pudlak syndrome 5 protein homolog Mus musculus | 54.34 | 0.98 | 126337 | 1 | 1 | 1293994 | 977232.12 | 679642.94 | 1212638.62 | 1405209.38 | 1384881.38 | 607042.75 | 513103.16 | 353394.91 | 374554.47 | 1246655.5 | 1747951 | 674995.94 | 871290.88 |
| Q3TC93 | Hs1bp3 | HCLS1-binding protein 3 Mus musculus | 94.78 | 4.3 | 43691 | 1 | 1 | 1310085.62 | 1358001.75 | 1127502 | 1406968.38 | 1162737.12 | 1167165.88 | 1516085 | 1209508.25 | 1212655.5 | 1278244.75 | 1556925.5 | 1252149.12 | 1325809.12 | 1309121.88 |
| Q8R3H7 | Hs2st1 | Heparan sulfate 2-O-sulfotransferase 1 Mus musculus | 117.24 | 7.3 | 41813 | 2 | 2 | 2687547.75 | 1947724.62 | 1716276.5 | 1655924.12 | 1704163.75 | 1905689.62 | 2146076 | 1769446.12 | 1417437.38 | 1703101.25 | 1515029.5 | 1725671.88 | 1520854.25 | 1722173.88 |
| O08756 | Hsd17b10 | 3-hydroxyacyl-CoA dehydrogenase type-2 Mus musculus | 712.23 | 55.94 | 27419 | 5 | 9 | 23563111 | 25149027 | 29836307 | 30778667 | 26688145 | 26588485 | 32122755 | 27162493 | 30631385 | 29028111 | 27330129 | 32268991 | 29912133 | 29641081 |
| Q9EQ06 | Hsd17b11 | Estradiol 17-beta-dehydrogenase 11 Mus musculus | 68.04 | 11.74 | 32881 | 2 | 2 | 1640220.5 | 380058.09 | 923520 | 395099.84 | 510289 | 325970.34 | 253953.95 | 503947.41 | 492992.53 | 1097527 | 428469.38 | 999009.25 | 513539.78 | 630369.12 |
| O70503 | Hsd17b12 | Very-long-chain 3-oxoacyl-CoA reductase Mus musculus | 1048.91 | 69.55 | 34742 | 17 | 17 | 29122711 | 30014685 | 31073283 | 28494255 | 27393423 | 30289733 | 29603029 | 31449389 | 26708147 | 27515511 | 26402463 | 30095909 | 28442037 | 28027353 |
| P51660 | Hsd17b4 | Peroxisomal multifunctional enzyme type 2 Mus musculus | 696.87 | 21.77 | 79482 | 11 | 11 | 15125930 | 10977580 | 14185545 | 13229414 | 12428735 | 11817173 | 13158129 | 13798612 | 13454169 | 12409687 | 11814502 | 11788424 | 13430504 | 12437004 |
| O88736 | Hsd17b7 | 3-keto-steroid reductase Mus musculus | 381.96 | 26.65 | 37317 | 7 | 7 | 4350245.5 | 3922892.75 | 4406959.5 | 3599696 | 3864512 | 3846372.25 | 3743803 | 4474243.5 | 4432062.5 | 3606522 | 3524424.5 | 4899891.5 | 4559142 | 4545992 |
| P50171 | Hsd17b8 | Estradiol 17-beta-dehydrogenase 8 Mus musculus | 194.79 | 18.15 | 26588 | 4 | 4 | 3826551 | 2408823.25 | 2803414 | 4490665.5 | 4305340 | 4659270.5 | 5484088.5 | 4018437.5 | 4983222 | 5325776 | 4845972 | 4886681 | 4865597 | 3941924.5 |
| Q8BTX9 | Hsdl1 | Inactive hydroxysteroid dehydrogenase-like protein 1 Mus musculus | 138.33 | 10.91 | 36868 | 2 | 2 | 1532226.12 | 1417680.75 | 1644205.25 | 1829637.88 | 1761696.75 | 1051358.75 | 1358895.12 | 1245302.25 | 1625902 | 1276974.12 | 1263502.12 | 1304692.12 | 1262580.62 | 1349341.75 |
| P07901 | Hsp90aa1 | Heat shock protein HSP 90-alpha Mus musculus | 2808.68 | 62.62 | 84788 | 28 | 46 | 416174593 | 466700545 | 429501857 | 418725409 | 478100257 | 485711521 | 447464545 | 465466945 | 439408193 | 479584993 | 485990913 | 498430721 | 507168833 | 517039393 |
| P11499 | Hsp90ab1 | Heat shock protein HSP 90-beta Mus musculus | 4068.121 | 76.38 | 83281 | 38 | 61 | 858858113 | 907080065 | 860898369 | 795965633 | 908185857 | 934317825 | 867120065 | 913728897 | 873618945 | 890316225 | 922130049 | 965213697 | 959466689 | 958405569 |
| P08113 | Hsp90b1 | Endoplasmin Mus musculus | 3099.08 | 61.97 | 92476 | 44 | 46 | 438865441 | 435713089 | 428330785 | 391069985 | 457535489 | 490388065 | 421803585 | 515039329 | 459849313 | 447182273 | 464496897 | 497617281 | 489239169 | 481065057 |
| Q99M31 | Hspa14 | Heat shock 70 kDa protein 14 Mus musculus | 296.23 | 19.84 | 54650 | 7 | 7 | 3661207 | 4327122.5 | 3964697.75 | 2309327.25 | 5929704.5 | 2257359 | 5143785.5 | 3260811 | 4265515.5 | 6843077.5 | 5136178.5 | 4783021.5 | 3616097.25 | 6974864.5 |
| Q61316 | Hspa4 | Heat shock 70 kDa protein 4 Mus musculus | 2190.29 | 54.7 | 94133 | 5 | 33 | 144136369 | 161532769 | 164268225 | 142548609 | 158958481 | 151734097 | 159660097 | 156008817 | 162266769 | 150475793 | 151334881 | 153210753 | 148501793 | 156750081 |
| P48722 | Hspa4l | Heat shock 70 kDa protein 4L Mus musculus | 219.27 | 5.37 | 94382 | 1 | 3 | 5077805.5 | 7223141 | 6410148 | 5993005.5 | 6154040 | 6540946.5 | 6402121 | 5802784.5 | 6167127 | 5974289.5 | 5534131 | 6965436.5 | 6607245.5 | 7102655.5 |
| P06761 | Hspa5 | Endoplasmic reticulum chaperone BiP Rattus norvegicus | 2885.9 | 64.53 | 72347 | 34 | 40 | 943839105 | 1021198273 | 1015736385 | 966251521 | 977109313 | 956006017 | 985471937 | 987955841 | 922050753 | 931761729 | 944878465 | 950579905 | 1008761793 | 986553217 |
| P63017 | Hspa8 | Heat shock cognate 71 kDa protein Mus musculus | 3579.499 | 77.86 | 70871 | 44 | 52 | 1801361665 | 1869525377 | 1957913473 | 1742636801 | 1764108289 | 1649671041 | 1699606401 | 1699964801 | 1645360129 | 1663941761 | 1694374529 | 1615660545 | 1761841537 | 1766261249 |
| P38647 | Hspa9 | Stress-70 protein, mitochondrial Mus musculus | 2182.07 | 55.67 | 73461 | 31 | 32 | 183261889 | 181052913 | 198643089 | 186770193 | 185384353 | 213031393 | 212484529 | 189584337 | 194952609 | 203820193 | 195486641 | 184821457 | 203430913 | 198804001 |
| Q99P31 | Hspbp1 | Hsp70-binding protein 1 Mus musculus | 572.21 | 45.66 | 39167 | 10 | 10 | 14451627 | 14545725 | 16243478 | 15873172 | 14678513 | 14233226 | 12644041 | 13985019 | 13554151 | 12725764 | 14455019 | 14324560 | 14405397 | 14767091 |
| P63038 | Hspd1 | 60 kDa heat shock protein, mitochondrial Mus musculus | 3463.88 | 79.06 | 60955 | 43 | 47 | 544798337 | 529123073 | 613124481 | 569205889 | 537038273 | 583622785 | 573542017 | 584274753 | 566607489 | 583392641 | 585433025 | 600132225 | 647663233 | 570655361 |
| P26772 | Hspe1 | 10 kDa heat shock protein, mitochondrial Rattus norvegicus | 215.66 | 42.16 | 10902 | 4 | 4 | 18476301 | 19934239 | 19126895 | 21050439 | 16966677 | 12317349 | 15474625 | 14810171 | 10708773 | 15614746 | 15044636 | 14590793 | 15932658 | 15271881 |
| Q61699 | Hsph1 | Heat shock protein 105 kDa Mus musculus | 1053.25 | 31.12 | 96407 | 17 | 18 | 13979423 | 14701681 | 15538737 | 12002855 | 13341430 | 11505919 | 12724204 | 11726425 | 9868152 | 10778031 | 13668253 | 16255472 | 15678931 | 14659286 |
| Q9Z2G9 | Htatip2 | Oxidoreductase HTATIP2 Mus musculus | 209.49 | 17.77 | 26870 | 4 | 4 | 2660624.25 | 2843695.25 | 3325615.5 | 1404470 | 1155086.5 | 3179811.5 | 4311161 | 2278153.5 | 4502977 | 1259505.38 | 4305708.5 | 4029175.75 | 2632579.75 | 2252173.25 |
| Q9JIY5 | Htra2 | Serine protease HTRA2, mitochondrial Mus musculus | 220.09 | 17.69 | 49348 | 4 | 5 | 747603.06 | 685581.5 | 941821.56 | 487902.75 | 834827.5 | 997479.31 | 474083 | 794950.75 | 1100611.88 | 607628.81 | 2001827.75 | 1800737.5 | 1692637.5 | 1097997.75 |
| P42859 | Htt | Huntingtin Mus musculus | 156.62 | 1.44 | 344690 | 4 | 4 | 6380583 | 4782427 | 4312055.5 | 4736919.5 | 5056132.5 | 4781647 | 4971312.5 | 3281574.5 | 5193406.5 | 4247296 | 4621722 | 4366892 | 4467006 | 4752972.5 |
| Q7TMY8 | Huwe1 | E3 ubiquitin-protein ligase HUWE1 Mus musculus | 998.72 | 7.86 | 482635 | 22 | 22 | 5177047 | 6848453.5 | 9441940 | 6243555.5 | 7194109 | 8445926 | 11872217 | 7516080.5 | 6044917.5 | 11200916 | 9459361 | 8428893 | 9614492 | 9658723 |
| Q8R1F5 | Hyi | Putative hydroxypyruvate isomerase Mus musculus | 282.88 | 24.55 | 30449 | 6 | 6 | 9398018 | 1449733.62 | 3642734.5 | 9416271 | 8546109 | 6508929 | 8350908 | 4000078 | 7184138 | 5505450 | 7736957 | 6495792 | 4930102 | 2284729.25 |
| Q9JKR6 | Hyou1 | Hypoxia up-regulated protein 1 Mus musculus | 2199.98 | 52.25 | 111181 | 37 | 37 | 68673689 | 74321409 | 74723737 | 70327585 | 73010289 | 81279161 | 74372057 | 81924777 | 73661641 | 67764257 | 74046081 | 78299313 | 78839825 | 74403849 |
| P03975 | Iap | IgE-binding protein Mus musculus V=1 | 590.01 | 23.16 | 62747 | 11 | 11 | 21785451 | 19728165 | 25046163 | 23513657 | 20108189 | 31279901 | 34253705 | 32022245 | 31745995 | 30166807 | 32709733 | 34841109 | 32528817 | 34805005 |
| Q8BU30 | Iars | Isoleucine--tRNA ligase, cytoplasmic Mus musculus | 622.99 | 13.55 | 144271 | 14 | 14 | 49474901 | 37118825 | 34908689 | 42228605 | 41116593 | 40683885 | 31770677 | 31836009 | 31662037 | 34950537 | 42095897 | 39480861 | 30380117 | 36440817 |
| Q8BIJ6 | Iars2 | Isoleucine--tRNA ligase, mitochondrial Mus musculus | 743.05 | 18.08 | 112804 | 13 | 13 | 19898681 | 25412257 | 23707031 | 20511649 | 23922073 | 25874443 | 28077879 | 24911379 | 27941139 | 23808957 | 26065137 | 25083243 | 24275297 | 26453295 |
| P13597 | Icam1 | Intercellular adhesion molecule 1 Mus musculus | 99.99 | 5.21 | 58844 | 2 | 2 | 1823847.25 | 1870921.75 | 1795167 | 1391329.25 | 1687625.75 | 1686406.25 | 2045399.62 | 2070661.12 | 1789631.75 | 1955418.38 | 1752152 | 1758315.5 | 1875641.75 | 1852095 |
| Q9JHR7 | Ide | Insulin-degrading enzyme Mus musculus | 413.9 | 9.72 | 117772 | 8 | 8 | 8222862 | 8821802 | 7243275 | 7681324 | 8548060 | 6692895 | 7949691 | 9294116 | 9283447 | 6829474.5 | 9654265 | 8446112 | 4955661 | 8508812 |
| O88844 | Idh1 | Isocitrate dehydrogenase [NADP] cytoplasmic Mus musculus | 1107.41 | 61.11 | 46674 | 19 | 20 | 63572689 | 55378865 | 71228601 | 62620189 | 52962105 | 54530457 | 65728701 | 59993269 | 60997469 | 59374857 | 58311757 | 59407445 | 68155385 | 60913109 |
| P54071 | Idh2 | Isocitrate dehydrogenase [NADP], mitochondrial Mus musculus | 1170.59 | 48.67 | 50906 | 19 | 21 | 72911833 | 74086593 | 74553665 | 72508329 | 70330505 | 71828065 | 79011729 | 83732601 | 78756425 | 79949545 | 82717289 | 76308801 | 78930425 | 83983257 |
| Q9D6R2 | Idh3a | Isocitrate dehydrogenase [NAD] subunit alpha, mitochondrial Mus musculus | 714.15 | 36.61 | 39639 | 12 | 12 | 108897041 | 108297993 | 103560273 | 101560265 | 102674825 | 97084449 | 107383097 | 117435145 | 105069017 | 111792673 | 102740625 | 95675105 | 106604177 | 107363449 |
| Q68FX0 | Idh3B | Isocitrate dehydrogenase [NAD] subunit beta, mitochondrial Rattus norvegicus | 357.97 | 25.71 | 42354 | 8 | 8 | 10234452 | 8873762 | 12745161 | 10257745 | 10238926 | 9926804 | 11011899 | 10802423 | 11437063 | 11929578 | 11173842 | 12119854 | 12698398 | 10247410 |
| P41565 | Idh3g | Isocitrate dehydrogenase [NAD] subunit gamma 1, mitochondrial Rattus norvegicus | 253.8 | 14.76 | 42851 | 6 | 6 | 16545307 | 13982288 | 16847583 | 16369147 | 13910443 | 18911789 | 15494213 | 15909703 | 17160641 | 18949001 | 19599161 | 17961009 | 21329011 | 17529899 |
| P58044 | Idi1 | Isopentenyl-diphosphate Delta-isomerase 1 Mus musculus | 580.47 | 37.89 | 26289 | 8 | 9 | 20077199 | 17077927 | 16589274 | 13619267 | 15197891 | 16890257 | 14892117 | 21304623 | 21038849 | 14154783 | 17368961 | 17694129 | 17890389 | 16334471 |
| Q9CR20 | Ier3ip1 | Immediate early response 3-interacting protein 1 Mus musculus =3 | 90.03 | 34.15 | 9017 | 2 | 2 | 1776517.62 | 1800477.88 | 1646131.38 | 1235866.38 | 1670221.38 | 2057467.62 | 1553786 | 1562193.62 | 1378950.88 | 1878677.88 | 1468097.88 | 1657696 | 1700484.88 | 1786355.75 |
| Q9R002 | Ifi202 | Interferon-activable protein 202 Mus musculus | 144.04 | 6.29 | 50466 | 2 | 2 | 2326020.25 | 2754549.75 | 2657673.25 | 1744767.5 | 1884176.88 | 1 | 1 | 1 | 1 | 1 | 1 | 1 | 1 | 1 |
| O35368 | Ifi203 | Interferon-activable protein 203 Mus musculus | 141.11 | 12.25 | 46300 | 2 | 3 | 3073621 | 4515976 | 3747676 | 3639146.25 | 3943940.5 | 3896708.25 | 3220888.5 | 3169128.75 | 2820361.5 | 1691877.25 | 3061970 | 2135649.5 | 2236953.25 | 3607709 |
| P0DOV2 | Ifi204 | Interferon-activable protein 204 Mus musculus | 675.02 | 27.46 | 69438 | 8 | 12 | 18012943 | 19144417 | 19639325 | 15222543 | 14087636 | 10929171 | 11615319 | 12017299 | 11915774 | 9637524 | 9370458 | 9949678 | 11449507 | 8969819 |
| Q9ESY9 | Ifi30 | Gamma-interferon-inducible lysosomal thiol reductase Mus musculus | 276.9 | 20.97 | 27784 | 4 | 4 | 9177378 | 12348510 | 9824344 | 9560805 | 11264979 | 11316278 | 13420224 | 10753703 | 9051181 | 9501100 | 12512991 | 9502271 | 14529967 | 15277725 |
| Q9D8C4 | Ifi35 | Interferon-induced 35 kDa protein homolog Mus musculus | 309.27 | 27.62 | 31876 | 6 | 6 | 1691151.5 | 2825326 | 1568373.88 | 2018846 | 2413499.75 | 2463729 | 2021990.75 | 1146408.25 | 2129804.5 | 1084939.75 | 2167048 | 2922008.5 | 2537894.75 | 2500188.5 |

| | | | | | | | | | | | | | | | | | | | | | |
|---|---|---|---|---|---|---|---|---|---|---|---|---|---|---|---|---|---|---|---|---|---|
| Q8BV66 | Ifi44 | Interferon-induced protein 44 Mus musculus | 78.42 | 5.92 | 47852 | 2 | 2 | 6552790.5 | 6363975 | 5733456 | 3897284.25 | 5024364 | 2039624.12 | 2116467.25 | 1792802.5 | 2103463.25 | 2083649.25 | 1789207.75 | 1374940.88 | 461732.31 | 1208626.98 |
| Q9BDB7 | Ifi44l | Interferon-induced protein 44-like Mus musculus 2 | 231.32 | 12.98 | 49680 | 4 | 4 | 2844332.75 | 2146209.25 | 2052692.5 | 1848826.62 | 1618923.62 | 842147.125 | 1044910.31 | 669528.31 | 942336.44 | 711813.44 | 737215.38 | 424957.41 | 802695.25 | 167065.08 |
| Q8R5F7 | Ifih1 | Interferon-induced helicase C domain-containing protein 1 Mus musculus | 51.61 | 1.17 | 115971 | 1 | 1 | 303777.95 | 307292.41 | 403792.19 | 272384.78 | 231642.42 | 300433.19 | 220947.42 | 379357.94 | 275787.91 | 230970.39 | 340671.22 | 305446.66 | 263506.66 | 196191.56 |
| Q99J93 | Ifitm2 | Interferon-induced transmembrane protein 2 Mus musculus | 146.22 | 16.67 | 15743 | 1 | 2 | 53026101 | 38082793 | 41511925 | 25423989 | 37172865 | 28936005 | 24759839 | 20878133 | 20745715 | 22083409 | 18167669 | 21079547 | 32500441 | 20010805 |
| Q9CQW9 | Ifitm3 | Interferon-induced transmembrane protein 3 Mus musculus | 539.03 | 48.91 | 14954 | 4 | 6 | 65032261 | 45017053 | 49339585 | 30601089 | 45552461 | 33990385 | 30262735 | 24008263 | 23870285 | 26900781 | 21719847 | 21401057 | 39564185 | 21880273 |
| P19182 | Ifrd1 | Interferon-related developmental regulator 1 Mus musculus | 141.02 | 5.35 | 49935 | 2 | 2 | 1 | 1 | 1 | 1 | 285720.09 | 339624.62 | 1 | 1 | 1 | 1 | 1 | 1 | 301021.53 | 1 |
| Q61249 | Igbp1 | Immunoglobulin-binding protein 1 Mus musculus | 307.62 | 18.24 | 38971 | 4 | 4 | 2160710 | 1080741.12 | 1455760 | 1554181.88 | 508005 | 1024517.94 | 840984.94 | 626720.62 | 835446.62 | 265473.09 | 384612.62 | 240125.86 | 1682172.38 | 388227.53 |
| Q9CPN8 | Igf2bp3 | Insulin-like growth factor 2 mRNA-binding protein 3 Mus musculus | 714.89 | 26.6 | 63575 | 11 | 11 | 11325107 | 11967692 | 12592103 | 11890570 | 14149274 | 11871218 | 11982451 | 8912142 | 11567337 | 12607661 | 12663937 | 14245362 | 13915909 | 12469808 |
| Q66HG8 | Ik | Protein Red Rattus norvegicus | 222.1 | 12.57 | 65588 | 5 | 5 | 5506375 | 4898198 | 2074125.12 | 4769928.5 | 4593217 | 5328247.5 | 5047525.5 | 4618095 | 5430646 | 5422640.5 | 5200812.5 | 5715690 | 5855394.5 | 5811492.5 |
| O88351 | Ikbkb | Inhibitor of nuclear factor kappa-B kinase subunit beta Mus musculus | 264.42 | 10.7 | 86690 | 5 | 5 | 9063181 | 2698539.25 | 8796161 | 2351809.5 | 2684371 | 9118863 | 2988006 | 12217903 | 10922312 | 9449693 | 8293788 | 10684524 | 9719247 | 9020347 |
| Q9R0T8 | Ikbke | Inhibitor of nuclear factor kappa-B kinase subunit epsilon Mus musculus | 202.04 | 6.83 | 80953 | 4 | 4 | 498817.88 | 1032041.25 | 1727046.75 | 2295590.75 | 1765663.25 | 537208.88 | 2153206.5 | 1475590.25 | 1388792.75 | 569193.94 | 1 | 1 | 1 | 1111947.62 |
| Q9CXY6 | Ilf2 | Interleukin enhancer-binding factor 2 Mus musculus | 732.59 | 53.85 | 43062 | 14 | 14 | 20563567 | 19345431 | 20165027 | 20896637 | 20034229 | 19942845 | 17199653 | 19258859 | 18856737 | 19025955 | 17879289 | 18110727 | 20047185 | 16444238 |
| Q9Z1X4 | Ilf3 | Interleukin enhancer-binding factor 3 Mus musculus | 511.49 | 16.82 | 96021 | 10 | 11 | 6328511.5 | 6775481 | 7170851 | 6997951.5 | 7813943 | 5572864.5 | 6491910.5 | 7228274 | 5794679.5 | 6308578.5 | 5717686 | 5359599 | 7363603.5 | 6447672 |
| O55222 | Ilk | Integrin-linked protein kinase Mus musculus | 80.71 | 6.42 | 51373 | 2 | 2 | 2229950.25 | 2900513 | 2954583 | 2837169.25 | 3166686 | 2242888.75 | 3178199 | 2772220.5 | 3697066.5 | 1416947.62 | 3204413 | 3348711.75 | 3061072.25 | 3109199.25 |
| Q8R0F6 | Ilkap | Integrin-linked kinase-associated serine/threonine phosphatase 2C Mus musculus | 135.03 | 10.97 | 42774 | 3 | 3 | 1029290.25 | 1227866.88 | 1092892.12 | 1115054.88 | 891648.75 | 1064947 | 983881.5 | 1237336.88 | 1181223.62 | 916076.62 | 1139025.88 | 1253732.12 | 1152945.62 | 1421809.5 |
| Q8CAQ8 | Immt | MICOS complex subunit Mic60 Mus musculus | 1666.42 | 45.18 | 83900 | 28 | 28 | 63464085 | 66969681 | 64434625 | 74027753 | 63271897 | 61703577 | 59618921 | 60560085 | 69284425 | 63618925 | 59620713 | 62884857 | 62110133 | 65892033 |
| O55023 | Impa1 | Inositol monophosphatase 1 Mus musculus | 611.74 | 46.93 | 30436 | 8 | 12 | 42729101 | 42525937 | 44979457 | 46504489 | 46628441 | 43510893 | 43486493 | 41005525 | 43398049 | 44725177 | 44491521 | 43417997 | 43094877 | 46166605 |
| Q8CIN7 | Impa2 | Inositol monophosphatase 2 Rattus norvegicus | 138.48 | 13.45 | 31796 | 3 | 3 | 3346350 | 2725572.75 | 1079522.75 | 2314741.75 | 3517454.5 | 2033842.5 | 4099381.75 | 3000745 | 2262335 | 3589267.75 | 1913979.38 | 1958691 | 2773889 | 1452300.75 |
| D3ZLZ7 | Impdh1 | Inosine-5'-monophosphate dehydrogenase 1 Rattus norvegicus 3 | 186.18 | 7.2 | 55293 | 3 | 4 | 10691699 | 7636193.5 | 8596957 | 10345150 | 9403159 | 9972892 | 10259992 | 8473558 | 10312983 | 9655576 | 9808429 | 10780468 | 8264693 | 9007438 |
| P24547 | Impdh2 | Inosine-5'-monophosphate dehydrogenase 2 Mus musculus | 1273.46 | 44.94 | 55815 | 18 | 19 | 76242937 | 59049025 | 62302537 | 75801265 | 75943457 | 69249841 | 78956673 | 61043957 | 73751089 | 71604241 | 73829225 | 74829257 | 67416705 | 65064157 |
| Q0GNC1 | Inf2 | Inverted formin-2 Mus musculus | 432.48 | 8.25 | 138560 | 8 | 8 | 1647709.12 | 1456264.75 | 1339991.88 | 1237981.5 | 1380480 | 1720009.88 | 1324928 | 2113867.75 | 1550908.5 | 2262031.75 | 1630009.75 | 3229328.5 | 2525351.25 | 1948364 |
| P49442 | Inpp1 | Inositol polyphosphate 1-phosphatase Mus musculus | 89.5 | 6.82 | 43346 | 2 | 2 | 3260557 | 2083278.62 | 2319977 | 1820683.38 | 3456455.5 | 1936161.12 | 2065716.62 | 2935704.25 | 3506708.5 | 2285259.5 | 1981008.75 | 3124747.5 | 2229535.5 | 2077793 |
| Q8K337 | Inpp5b | Type II inositol 1,4,5-trisphosphate 5-phosphatase Mus musculus | 49.91 | 2.22 | 112762 | 1 | 1 | 286034.19 | 270461.94 | 1 | 1 | 1 | 194172.66 | 118652.17 | 1 | 203965.06 | 1 | 153996.05 | 253725.3 | 201682.41 | 1 |
| Q9ES52 | Inpp5d | Phosphatidylinositol 3,4,5-trisphosphate 5-phosphatase 1 Mus musculus | 1433.81 | 30.81 | 133542 | 27 | 29 | 33906385 | 34887021 | 39670337 | 38651161 | 41056853 | 38116793 | 38938561 | 31742885 | 38192469 | 41943417 | 37646249 | 39459665 | 37471237 | 38558177 |
| Q6P549 | Inppl1 | Phosphatidylinositol 3,4,5-trisphosphate 5-phosphatase 2 Mus musculus | 580.63 | 13.37 | 138973 | 11 | 13 | 10057740 | 8966307 | 9686605 | 9075967 | 7102800 | 7736722.5 | 8610283 | 7488930 | 8801840 | 9561610 | 10341035 | 8261365.5 | 9045841 | 10904745 |
| Q6P4S8 | Ints1 | Integrator complex subunit 1 Mus musculus | 39.05 | 0.91 | 245168 | 1 | 1 | 181085.66 | 288791.91 | 246775.52 | 1 | 318217.59 | 176969.14 | 199120.195 | 232933.03 | 191170.81 | 195407.8 | 350937.56 | 232560.83 | 275035.5 | 193558.06 |
| Q3MHC2 | Ints11 | Integrator complex subunit 11 Rattus norvegicus 2 | 43.91 | 1.67 | 67841 | 1 | 1 | 974125.75 | 675339.88 | 783002.31 | 834236.25 | 706802.12 | 776426.94 | 752664.19 | 788318.75 | 820622.12 | 767821.56 | 687770.56 | 743813.69 | 772333.19 | 853302.31 |
| Q68FR3 | Ints12 | Integrator complex subunit 12 Rattus norvegicus | 54.09 | 3.04 | 48477 | 1 | 1 | 1 | 1 | 1 | 1 | 1 | 233719.3 | 188584.84 | 199350.3975 | 202375.92 | 172721.53 | 1 | 132676.22 | 124364.33 | 1 |
| Q8QZV7 | IntS13 | Integrator complex subunit 13 Mus musculus | 38.84 | 1.5 | 82780 | 1 | 1 | 1 | 1232785.62 | 1174522.25 | 1 | 1 | 952227.3575 | 1051861.25 | 1197137.12 | 850061.06 | 709850 | 1081880.12 | 1088809.62 | 864635.19 | 866763.69 |
| Q66H58 | Ints14 | Integrator complex subunit 14 Rattus norvegicus 2 | 62.67 | 2.33 | 57144 | 1 | 1 | 247919.2 | 239601.92 | 277329.53 | 306764.06 | 318210.34 | 265793.59 | 266178.19 | 297987.5 | 317637.44 | 229583.34 | 329531.69 | 279825.66 | 261419.98 | 187139.36 |
| Q7TPD0 | Ints3 | Integrator complex subunit 3 Mus musculus | 52.77 | 1.63 | 117938 | 1 | 1 | 278796.5 | 139191.3 | 172355.92 | 147970.53 | 151160.78 | 106623.33 | 1 | 374587.53 | 183484.75 | 1 | 168650.5167 | 255864.88 | 92419.37 | 157667.3 |
| Q8CIM8 | Ints4 | Integrator complex subunit 4 Mus musculus | 226.28 | 7.57 | 108193 | 4 | 4 | 867053.75 | 534346.56 | 1268611.25 | 409629.25 | 935553.06 | 909280.38 | 1060416.25 | 1173503.75 | 900166.12 | 415204.56 | 1008080.5 | 478403.03 | 820090.94 | 1005062.88 |
| Q7TQK1 | Ints7 | Integrator complex subunit 7 Mus musculus | 162.44 | 6 | 106861 | 4 | 4 | 744211.305 | 459033.56 | 468421.53 | 1178950.25 | 870439.88 | 723729.5 | 522926 | 291398.81 | 1126237.75 | 690522.44 | 952260.88 | 419888.06 | 1332664.5 | 431283.91 |
| Q80V86 | Ints8 | Integrator complex subunit 8 Mus musculus | 34.38 | 0.7 | 113353 | 1 | 1 | 931067.31 | 809143.56 | 771522.56 | 846315.19 | 706830.19 | 541543.5 | 729858.5 | 709024.19 | 523866.44 | 640948.31 | 647398.12 | 750522.69 | 719151.62 | 601051.25 |
| Q8K114 | Ints9 | Integrator complex subunit 9 Mus musculus | 39.47 | 3.19 | 74078 | 1 | 1 | 404984 | 477826.62 | 556368.56 | 373949.28 | 446336.06 | 457792.44 | 504021.69 | 537420.19 | 437579.44 | 357556.34 | 628481 | 475898.88 | 496984.72 | 331205.84 |
| Q8K2V6 | Ipo11 | Importin-11 Mus musculus | 136.31 | 5.85 | 112416 | 3 | 3 | 1769204.38 | 847586.31 | 1007741.88 | 182755.36 | 1513397.75 | 409686.75 | 406189.16 | 441848.19 | 1049091.75 | 722837.88 | 882179.56 | 1317114.12 | 439023.44 | 615990.12 |
| Q8VI75 | Ipo4 | Importin-4 Mus musculus | 259.13 | 6.65 | 119275 | 6 | 7 | 2748288.5 | 2278374.25 | 3183473.75 | 2987209.5 | 3082576.75 | 2667036 | 2677922 | 2364929.75 | 2690972.75 | 2700471 | 2532938.25 | 2700247.75 | 2517097.5 | 2536995.25 |
| Q8BKC5 | Ipo5 | Importin-5 Mus musculus | 1521.09 | 37.65 | 123591 | 28 | 28 | 48631161 | 48013341 | 51696005 | 32527311 | 41215461 | 53493613 | 49722969 | 48792409 | 51160501 | 50527349 | 47934185 | 52077377 | 52017329 | 41483597 |
| Q9EPL8 | Ipo7 | Importin-7 Mus musculus | 638.96 | 15.9 | 119486 | 11 | 12 | 21407035 | 12272100 | 17098311 | 10782678 | 9871825 | 15622479 | 16499560 | 23478247 | 15794815 | 12582251 | 11150541 | 14627855 | 16944637 | 10754171 |
| Q7TMY7 | Ipo8 | Importin-8 Mus musculus | 113.41 | 4.16 | 117078 | 2 | 3 | 222764.27 | 164700.03 | 2676387 | 134075.33 | 156044.33 | 99893.2 | 2714296.25 | 3226368.25 | 79862.47 | 76156.7 | 110356.7 | 146948.94 | 2350763.5 | 869356.38 |
| Q91YE6 | Ipo9 | Importin-9 Mus musculus | 1449.63 | 33.72 | 116052 | 22 | 22 | 43797889 | 48878905 | 45943877 | 43513609 | 41303201 | 51128825 | 43634321 | 56560781 | 53736069 | 41501381 | 48191373 | 48717925 | 48325745 | 49347753 |
| Q9JKF1 | Iqgap1 | Ras GTPase-activating-like protein IQGAP1 Mus musculus | 6063.399 | 68.56 | 188742 | 90 | 96 | 519050561 | 488924513 | 508137121 | 496845985 | 463244961 | 576668161 | 560585793 | 599433985 | 578488513 | 558550977 | 499355777 | 497528801 | 520964737 | 541056769 |
| Q3UQ44 | Iqgap2 | Ras GTPase-activating-like protein IQGAP2 Mus musculus | 196.71 | 2.6 | 180528 | 1 | 5 | 22371009 | 24136241 | 22486175 | 24295237 | 22065695 | 27804099 | 25029741 | 25726317 | 25219491 | 28893215 | 23858859 | 27781209 | 24645635 | 24493761 |
| Q8R0S2 | Iqsec1 | IQ motif and SEC7 domain-containing protein 1 Mus musculus | 137.24 | 4.37 | 108015 | 3 | 3 | 1 | 289479.97 | 171411.81 | 1 | 147244.22 | 1 | 1 | 164722.94 | 143829.59 | 119323.54 | 221291.17 | 120683.93 | 107365.76 | 149780.2867 |
| Q8R4K2 | Irak4 | Interleukin-1 receptor-associated kinase 4 Mus musculus | 74.58 | 3.7 | 50872 | 1 | 1 | 1060406.25 | 1101917 | 1002067.94 | 1146826.38 | 1106412.25 | 922905.25 | 875769.12 | 996127 | 1027434 | 850466.44 | 930253.19 | 901446.62 | 803519.38 | 893944.81 |
| E9Q1P8 | Irf2bp2 | Interferon regulatory factor 2-binding protein 2 Mus musculus | 121.44 | 7.19 | 59292 | 1 | 2 | 376504.16 | 340499.75 | 560696.69 | 29403.25 | 326775.9625 | 417958.25 | 381190.59 | 456377.5 | 658671.56 | 511402.16 | 442766.28 | 463452.69 | 550650.06 | 549348.19 |
| Q5EIC4 | Irf2bpl | Interferon regulatory factor 2-binding protein-like Rattus norvegicus | 79.29 | 4.73 | 81496 | 1 | 2 | 886338.5 | 1033053.44 | 1089360.38 | 573580.88 | 509385.41 | 1123089.5 | 1158167.88 | 1381445.75 | 1591920.5 | 1102901.75 | 1146767 | 1037823.94 | 1283478.25 | 1304527.25 |
| P56477 | Irf5 | Interferon regulatory factor 5 Mus musculus | 469.74 | 21.33 | 56005 | 8 | 8 | 6195636 | 4725676.5 | 6614123.5 | 5685785.5 | 5408696 | 7044485 | 5292203 | 6750491.5 | 6633115.5 | 6402820 | 5997214.5 | 6562640 | 6484901 | 6554878 |
| Q60766 | Irgm1 | Immunity-related GTPase family M protein 1 Mus musculus | 382.54 | 19.07 | 46552 | 7 | 8 | 6445768.5 | 4892167.5 | 4781422.5 | 3869175 | 4389789 | 2883047.75 | 2765580 | 2933151.25 | 3284595 | 2833628 | 2599356.5 | 2450496.5 | 3855157.5 | 2993222.25 |
| Q64339 | Isg15 | Ubiquitin-like protein ISG15 Mus musculus | 59.95 | 11.18 | 17898 | 1 | 1 | 239879.86 | 383989.53 | 247970.45 | 363461.97 | 266334.69 | 207630.58 | 274218.12 | 418383.72 | 284436.6 | 237513.98 | 291434.72 | 250020.75 | 260463.84 | 360658.25 |
| Q6I7R3 | Isoc1 | Isochorismatase domain-containing protein 1 Rattus norvegicus | 171.97 | 14.81 | 32002 | 3 | 3 | 3014118 | 2495339.5 | 2088693 | 479990.72 | 3078048.75 | 2773609.5 | 2291453.25 | 2168728.5 | 2466883.25 | 2457650.25 | 2539431.5 | 2685891.75 | 2385459.25 | 2551059 |
| P85094 | Isoc2a | Isochorismatase domain-containing protein 2A Mus musculus | 237.74 | 40.78 | 22417 | 4 | 4 | 3103277.25 | 2942515 | 2963688.25 | 2998752.25 | 3032113.75 | 3567827.75 | 3256465.25 | 2750156 | 4007995 | 3503977.25 | 4070244.75 | 3735360.25 | 4640719 | 4501481 |
| Q568Z6 | Ist1 | IST1 homolog Rattus norvegicus | 147.61 | 9.29 | 39942 | 3 | 3 | 1452078.5 | 824591 | 902055.94 | 1459026 | 1172317.5 | 791035.5 | 544406.88 | 1425762.62 | 504516.47 | 1237185 | 1147390.25 | 1297556.25 | 559168.5 | 799635.75 |
| Q9JHU9 | Isyna1 | Inositol-3-phosphate synthase 1 Mus musculus | 663.52 | 29.26 | 60932 | 11 | 11 | 19801961 | 23093807 | 24332087 | 22423971 | 21288923 | 23101655 | 24394055 | 23515929 | 25244925 | 21761001 | 23163881 | 23560049 | 27403043 | 23245903 |
| Q00651 | Itga4 | Integrin alpha-4 Mus musculus | 78.48 | 2.12 | 115696 | 2 | 2 | 1 | 1 | 758519.5 | 1 | 265363.81 | 199960.12 | 1 | 507060.16 | 1 | 1 | 1 | 405717.25 | 332913.44 | 1 |
| P11688 | Itga5 | Integrin alpha-5 Mus musculus | 466.22 | 15.76 | 115043 | 10 | 10 | 5198483 | 3406540.75 | 7189628 | 4701444 | 4655489.5 | 5542421.5 | 6058128.5 | 5639336 | 5491996 | 5256880.5 | 5303205.5 | 6055553 | 5352772 | 6360909.5 |
| P05555 | Itgam | Integrin alpha-M Mus musculus | 1606.42 | 30.01 | 127481 | 24 | 24 | 68837145 | 72955153 | 78229753 | 61495005 | 73197409 | 56020089 | 55143293 | 60602621 | 60426461 | 57877545 | 59598233 | 60240989 | 68919881 | 61229769 |
| P09055 | Itgb1 | Integrin beta-1 Mus musculus | 902.36 | 27.19 | 88231 | 16 | 16 | 32506807 | 38435029 | 33373171 | 29439243 | 34020877 | 37723109 | 35276129 | 37860321 | 38948945 | 36952865 | 35933413 | 34654269 | 35472177 | 37574993 |
| P11835 | Itgb2 | Integrin beta-2 Mus musculus | 1755.53 | 44.75 | 85026 | 27 | 27 | 89636257 | 102390457 | 115078817 | 77016329 | 99660521 | 82846617 | 75295793 | 78449497 | 79430689 | 83396537 | 86968169 | 81247073 | 83874425 | 94043081 |
| O70309 | Itgb5 | Integrin beta-5 Mus musculus | 72.13 | 2.88 | 87909 | 2 | 2 | 1 | 1 | 1 | 1 | 1 | 68096.88 | 104749.095 | 153777.06 | 127744.94 | 69377.5 | 91404.76 | 131483.73 | 82495.11 | 101794.5333 |
| O89051 | Itm2b | Integral membrane protein 2B Mus musculus | 242.83 | 19.55 | 30260 | 4 | 4 | 3923463.5 | 4806815 | 4322810 | 3662523.25 | 2545112 | 4230018 | 4459268.5 | 6102388 | 5211909 | 4463752.5 | 5147811.5 | 4589960.5 | 3497168 | 4867620 |
| Q5PQL7 | Itm2c | Integral membrane protein 2C Rattus norvegicus | 105.92 | 11.52 | 30480 | 2 | 2 | 1782204.5 | 1484172.5 | 1676717.88 | 1326225.5 | 1185751 | 1866026.75 | 1721062.5 | 2472878.75 | 2119065.5 | 1993988.5 | 1764260.75 | 2158080.75 | 2409142.75 | 2332807.5 |
| Q9D892 | Itpa | Inosine triphosphate pyrophosphatase Mus musculus | 303.95 | 44.44 | 21897 | 6 | 6 | 11202732 | 6558981 | 11430209 | 10363730 | 9197700 | 9747878 | 10360995 | 8982742 | 10568615 | 10244829 | 9776717 | 10514001 | 11285417 | 11332917 |
| P11881 | Itpr1 | Inositol 1,4,5-trisphosphate receptor type 1 Mus musculus | 137.18 | 1.2 | 313167 | 1 | 3 | 2044397 | 2403410 | 2190753.75 | 2525277.75 | 1417940.88 | 2427041 | 1358073.75 | 2361419.5 | 2507992.75 | 2023316.5 | 2288530.75 | 1617326.88 | 2192902 | 2505567.5 |
| P29995 | Itpr2 | Inositol 1,4,5-trisphosphate receptor type 2 Rattus norvegicus | 202.91 | 1.93 | 307058 | 3 | 5 | 2451430.75 | 2791530.5 | 2356905.5 | 2536544.75 | 1545694 | 2620741.5 | 1855344.12 | 2768546.25 | 2612028.5 | 2158270 | 2641540.75 | 2074498.5 | 2475133.5 | 2521843.25 |
| P70227 | Itpr3 | Inositol 1,4,5-trisphosphate receptor type 3 Mus musculus | 131.55 | 1.91 | 304275 | 4 | 4 | 639147.94 | 1556119.5 | 1574866.25 | 1596880.12 | 1718204 | 2475911.75 | 1699047 | 1880152 | 2103020.75 | 2218357.5 | 2138167.5 | 2033202 | 1825938 | 1802469.75 |
| Q9JHI5 | Ivd | Isovaleryl-CoA dehydrogenase, mitochondrial Mus musculus | 286 | 17.22 | 46325 | 7 | 7 | 9670996 | 6238688 | 11463977 | 12403107 | 9045478 | 9626245 | 11865673 | 7819387 | 12187270 | 6971389 | 4898558.5 | 10130486 | 11793393 | 9823841 |
| Q4KM64 | Jagn1 | Protein jagunal homolog 1 Rattus norvegicus | 63.67 | 6.56 | 21102 | 1 | 1 | 1 | 1 | 1 | 1 | 1954519.75 | 1 | 293684.72 | 1 | 292557.88 | 1 | 370578.1867 | 304871.47 | 438947.03 | 367916.06 |
| P52332 | Jak1 | Tyrosine-protein kinase JAK1 Mus musculus | 169.62 | 4.94 | 133367 | 3 | 3 | 1758193.25 | 1617794.75 | 2500729.5 | 1903785.5 | 2008187.25 | 2131179.25 | 1561907 | 1664972.75 | 2991740 | 1491117.25 | 1576795.5 | 1833251.38 | 2010208 | 1850008.25 |
| P97825 | Jpt1 | Jupiter microtubule associated homolog 1 Mus musculus | 106.39 | 27.27 | 16081 | 2 | 2 | 2924789.25 | 4087840.5 | 3059245.5 | 4315181 | 3478096 | 1926713.38 | 1266849.12 | 2953416.75 | 1909610.25 | 2426835.25 | 2296861.25 | 1162097.38 | 1083107.12 | 2803540.5 |
| Q5BK20 | Jpt2 | Jupiter microtubule associated homolog 2 Rattus norvegicus | 105.31 | 18.42 | 20070 | 2 | 2 | 1159687.88 | 1111391.12 | 1737516.38 | 1378039.75 | 1613009.25 | 1073075.25 | 1063749.38 | 1183393 | 979843.94 | 1955527.25 | 1334339.5 | 1575931.5 | 1077850.75 | 1324308.5 |
| Q02257 | Jup | Junction plakoglobin Mus musculus | 66.82 | 2.42 | 81801 | 1 | 1 | 681373.81 | 1 | 1 | 1 | 1 | 1 | 1 | 1 | 1 | 122476.52 | 1 | 131256.72 | 148330.28 | 1 |
| Q99MN1 | Kars | Lysine--tRNA ligase Mus musculus | 801.4 | 33.45 | 67840 | 4 | 16 | 25529641 | 27604621 | 26149621 | 23147309 | 25424887 | 27118193 | 27826457 | 25373905 | 29446619 | 25623251 | 25002227 | 28549269 | 30604197 | 28316661 |
| Q5SVQ0 | Kat7 | Histone acetyltransferase KAT7 Mus musculus | 181.99 | 7.01 | 70641 | 3 | 3 | 2823160.75 | 1656111.88 | 1134877.5 | 950151 | 1963719.5 | 1774442.25 | 1622311.25 | 2042530.62 | 1232929 | 1079748.5 | 1126402.75 | 1417017 | 2714823.25 | 2843579.25 |
| Q80UY2 | Kcmf1 | E3 ubiquitin-protein ligase KCMF1 Mus musculus | 169.46 | 8.14 | 41791 | 2 | 2 | 424839.94 | 440292.12 | 223873.81 | 451242.31 | 465944.56 | 214918.64 | 191453.97 | 395658.06 | 366751.44 | 268202.75 | 439550.31 | 567460.25 | 223322.3 | 422113.69 |
| P62482 | Kcnab2 | Voltage-gated potassium channel subunit beta-2 Mus musculus | 379.23 | 35.97 | 41021 | 7 | 7 | 11593159 | 12929697 | 12474758 | 13290734 | 11773923 | 13282590 | 14060553 | 12990218 | 14285813 | 12341438 | 11852202 | 10747507 | 11878111 | 9421511 |
| Q6WVG3 | Kctd12 | BTB/POZ domain-containing protein KCTD12 Mus musculus | 512.69 | 36.39 | 35892 | 10 | 10 | 14306049 | 15819779 | 15026211 | 17265059 | 23246457 | 18088725 | 15807113 | 14526873 | 14649204 | 17087915 | 17643457 | 15070851 | 14405221 | 16162942 |
| Q9JHP7 | Kdelc1 | KDEL motif-containing protein 1 Mus musculus | 240.67 | 12.35 | 57985 | 5 | 5 | 1223272.62 | 1045887.25 | 1499118.38 | 367626.06 | 736394.88 | 1194721 | 942390.5 | 1887783.88 | 1953648.75 | 907518.44 | 1180377.25 | 1257543.62 | 1150431.88 | 1307532.25 |
| Q569A6 | Kdelr1 | ER lumen protein-retaining receptor 1 Rattus norvegicus 2 | 97.28 | 10.38 | 24546 | 2 | 2 | 1144506.25 | 213300.92 | 162754.42 | 189118.08 | 427419.9175 | 1 | 102192.34 | 158620.48 | 1 | 1 | 1 | 1 | 138202.08 | 147157.95 |
| Q6ZQ88 | Kdm1a | Lysine-specific histone demethylase 1A Mus musculus | 155.68 | 5.16 | 92851 | 2 | 2 | 1145119.88 | 987750.56 | 1188586.62 | 1060773.375 | 921636.44 | 1002121.19 | 974195.31 | 1064346.25 | 1131527.88 | 1063227.62 | 1019830.81 | 918299.06 | 1150267.25 | 1100447.5 |
| P59997 | Kdm2a | Lysine-specific demethylase 2A Mus musculus | 161.2 | 2.76 | 132680 | 3 | 3 | 248617.81 | 1953887 | 2227117 | 1956581.5 | 3216276.75 | 3134206.25 | 3092727.5 | 1991727.12 | 3024348.5 | 3033234.5 | 3163457.75 | 2164621.25 | 2031215 | 2572334.5 |
| Q3UXZ9 | Kdm5a | Lysine-specific demethylase 5A Mus musculus | 47.87 | 0.77 | 192216 | 1 | 1 | 227078.34 | 255711.59 | 206055.94 | 242048.64 | 279348.69 | 174793.67 | 164065.64 | 320100.41 | 194433.2 | 156873.34 | 241210.48 | 249592.33 | 175300.59 | 132548.48 |
| P41230 | Kdm5c | Lysine-specific demethylase 5C Mus musculus | 40.19 | 0.84 | 175313 | 1 | 1 | 105584.94 | 244402.84 | 183996.55 | 201710.8 | 210575.95 | 121499.27 | 124476.85 | 158326.17 | 167603.34 | 196906.83 | 189205.61 | 235637.03 | 744044.69 | 106341.96 |
| Q6GV12 | Kdsr | 3-ketodihydrosphingosine reductase Mus musculus | 65.45 | 4.22 | 35956 | 1 | 1 | 1429564 | 1541790.62 | 1477920.38 | 1612317.12 | 1591729.62 | 1734343.12 | 1274162.62 | 1570917.88 | 1353793.88 | 1483304.375 | 1 | 1327692.12 | 1417061.38 | 1 |
| Q60749 | Khdrbs1 | KH domain-containing, RNA-binding, signal transduction-associated protein 1 Mus musculi | 156.47 | 16.48 | 48371 | 4 | 4 | 1929417.88 | 1149907.12 | 2048198 | 1872508.62 | 2247208.25 | 2106668.75 | 2332169 | 1887153.88 | 2168887.75 | 2466514.75 | 2495453 | 1973671 | 2457235 | 2290427.75 |
| Q3U0V1 | Khsrp | Far upstream element-binding protein 2 Mus musculus | 1332.31 | 48.93 | 76775 | 20 | 22 | 56492473 | 69930305 | 58396933 | 74231513 | 77922577 | 64550333 | 74844457 | 65513129 | 71514393 | 76144665 | 70178265 | 56435681 | 68219609 | 79849769 |
| Q6P9P6 | Kif11 | Kinesin-like protein KIF11 Mus musculus | 575.89 | 14.16 | 118027 | 9 | 9 | 1801478.25 | 1844002.5 | 1682134.75 | 1739286 | 1683549 | 2769442.25 | 2144045.5 | 1744872.88 | 2268083.25 | 2153403.5 | 2794677.5 | 2142315.5 | 1691018.62 | 1437019.75 |
| Q6P9L6 | Kif15 | Kinesin-like protein KIF15 Mus musculus | 215.27 | 3.53 | 160120 | 3 | 3 | 613811.88 | 450786.91 | 756072.19 | 419258.38 | 885964.5 | 1102867.38 | 851302.25 | 629978.94 | 1066865.38 | 522565.88 | 632580.25 | 1608650.5 | 1316431.38 | 973782 |
| O35071 | Kif1c | Kinesin-like protein KIF1C Mus musculus | 63.83 | 1.18 | 122434 | 1 | 1 | 125842.36 | 103534.19 | 129607.06 | 111424.1 | 289267.75 | 158966.16 | 113365.58 | 68163.71 | 172627.94 | 128280.8475 | 119935.85 | 173776.84 | 105449.16 | 242591.06 |

| | | | | | | | | | | | | | | | | | | | | | |
|---|---|---|---|---|---|---|---|---|---|---|---|---|---|---|---|---|---|---|---|---|---|
| P97329 | Kif20a | Kinesin-like protein KIF20A Mus musculus | 57.07 | 3.16 | 99876 | 1 | 1 | 1 | 1 | 1 | 384765.12 | 260453.38 | 253352.72 | 190681.42 | 1 | 157333.42 | 1 | 209755.66 | 143621.72 | 283397.44 | 212258.2733 |
| F1M5N7 | Kif21b | Kinesin-like protein KIF21B Rattus norvegicus | 236.75 | 5.2 | 182563 | 5 | 5 | 1035331.31 | 1890741 | 1367951.38 | 1164323.38 | 1275484.25 | 1784628.5 | 1359197.38 | 1674837.5 | 1576579.12 | 1746144.38 | 1529583.5 | 1344461 | 1138854.62 | 1162457.5 |
| Q3V300 | Kif22 | Kinesin-like protein KIF22 Mus musculus | 78.78 | 3.94 | 73190 | 2 | 2 | 1265744.75 | 1064443.25 | 957997.25 | 1238799.25 | 931019.81 | 1299079.38 | 686134.4625 | 656093.5 | 292359.53 | 497005.44 | 1147785.75 | 1823298.75 | 811664.88 | 1091851.25 |
| E9Q5G3 | Kif23 | Kinesin-like protein KIF23 Mus musculus | 46.24 | 1.26 | 108776 | 1 | 1 | 896212.19 | 1619307 | 1253198.38 | 1764848.75 | 588226.38 | 1061361.25 | 1233739.62 | 1053476.12 | 671267.75 | 574471.19 | 351483.31 | 1104989 | 1349986.62 | 1357498.88 |
| P28740 | Kif2a | Kinesin-like protein KIF2A Mus musculus | 102.84 | 2.7 | 79756 | 2 | 2 | 573731.31 | 1145016 | 1255727.5 | 935530.19 | 599947.31 | 361999.69 | 669003.94 | 1263010.75 | 1158376.38 | 921613.12 | 569511.75 | 1136319.88 | 686502.62 | 733435.56 |
| Q922S8 | Kif2c | Kinesin-like protein KIF2C Mus musculus | 95.48 | 4.72 | 81085 | 2 | 2 | 242709.95 | 1 | 275403.44 | 1 | 402230.97 | 345746.72 | 667487.75 | 865196.25 | 668064.5 | 636623.805 | 1032694.56 | 1090669.5 | 972195.62 | 532133.69 |
| Q61771 | Kif3b | Kinesin-like protein KIF3B Mus musculus | 116.12 | 2.54 | 85288 | 1 | 1 | 2063001 | 2194726 | 1 | 1 | 1314586.12 | 1965726.25 | 1741508.88 | 2703465.5 | 1608145.75 | 2990421.5 | 1956656 | 1518262.62 | 2133437 | 2911599.5 |
| P33174 | Kif4 | Chromosome-associated kinesin KIF4 Mus musculus | 181.14 | 3.66 | 139519 | 4 | 4 | 2610901.75 | 2644056.5 | 2799358.5 | 1277284.25 | 2421715.5 | 2597166.5 | 2022307 | 830647 | 1369147.75 | 1529479.88 | 2727065 | 1652155.12 | 1255043.75 | 2365856.5 |
| P33175 | Kif5a | Kinesin heavy chain isoform 5A Mus musculus | 188.16 | 4.58 | 117019 | 0 | 4 | 3830728.5 | 2069237.5 | 2084377.88 | 1940073.25 | 2864564 | 1756975 | 1619471.5 | 1182498.75 | 1585857.468 | 1784484.62 | 3130023.5 | 1716612.75 | 2907706.5 | 1499499.88 |
| Q2PQA9 | Kif5b | Kinesin-1 heavy chain Rattus norvegicus | 993.88 | 23.36 | 109531 | 13 | 17 | 10117376 | 11674566 | 10884703 | 12310351 | 9799467 | 11803854 | 10391265 | 12570015 | 11803177 | 13342517 | 13781332 | 11008825 | 12601933 | 13123322 |
| P28738 | Kif5c | Kinesin heavy chain isoform 5C Mus musculus | 193.65 | 5.13 | 109275 | 1 | 4 | 1634748.62 | 1525816.5 | 2336881.75 | 1455427.75 | 1495552.12 | 1456539.75 | 2419700 | 2355356.75 | 1303587 | 1784484.62 | 1438625.38 | 1716612.75 | 1531278.38 | 2029047.5 |
| P70188 | Kifap3 | Kinesin-associated protein 3 Mus musculus | 232.68 | 5.8 | 91291 | 3 | 3 | 1491804.12 | 1501712.5 | 1419731.25 | 1250890.5 | 1425476 | 1864644.75 | 1384948.38 | 1721172.38 | 1506714.5 | 1096676.62 | 1181126.38 | 1734581.88 | 1491729.5 | 1552447.75 |
| O88447 | Klc1 | Kinesin light chain 1 Mus musculus | 38.25 | 2.4 | 61450 | 1 | 1 | 804088.31 | 841914.5 | 922713.69 | 1119057.12 | 1045303.44 | 724371.19 | 1173988.25 | 967664.38 | 1103912.25 | 1032298.94 | 1006323.88 | 1002021.25 | 853811.62 | 966977.56 |
| Q921I2 | Klhdc4 | Kelch domain-containing protein 4 Mus musculus 2 | 52.98 | 2.57 | 64860 | 1 | 1 | 1290601.62 | 1177804.25 | 1553057.88 | 1303753.75 | 1331525 | 1237858.88 | 1110034.88 | 1040095.44 | 911102.38 | 998585.06 | 1126783.88 | 1222575.25 | 1166588.62 | 1330169.75 |
| Q8C3Y4 | Kntc1 | Kinetochore-associated protein 1 Mus musculus | 106.39 | 1.63 | 250358 | 3 | 3 | 1642317.75 | 1803738.75 | 1552117.5 | 1696947.75 | 1526644.5 | 1438818.5 | 1387031.5 | 1432149.25 | 1324792 | 1088379.62 | 1622234.5 | 1434481.75 | 1410783.88 | 1251005.5 |
| P83953 | Kpna1 | Importin subunit alpha-5 Rattus norvegicus | 477.59 | 22.68 | 60137 | 3 | 7 | 4530387.5 | 4801344.5 | 4995068.5 | 4916343 | 3771598.75 | 4824911.5 | 4741556.5 | 3937946.75 | 4973683 | 3520077.5 | 4945946.5 | 5526356.5 | 5209703 | 5375926.5 |
| P52293 | Kpna2 | Importin subunit alpha-1 Mus musculus | 1192.77 | 48.77 | 57928 | 17 | 17 | 17248655 | 15881089 | 20675819 | 17715973 | 15642924 | 14031604 | 15633144 | 13114095 | 16035950 | 14784199 | 17153887 | 21234893 | 21426397 | 19724155 |
| O35344 | Kpna3 | Importin subunit alpha-4 Mus musculus | 574.8 | 27.45 | 57773 | 9 | 10 | 4867773 | 11178238 | 5797838.5 | 4495336 | 7245831 | 9267289 | 8289657 | 13406877 | 12377464 | 7977978 | 8918785 | 9097895 | 10165519 | 7840146 |
| O35343 | Kpna4 | Importin subunit alpha-3 Mus musculus | 520.35 | 26.1 | 57923 | 7 | 8 | 7532582 | 9383363 | 8000332 | 6291915 | 9114079 | 9482772 | 8053814 | 8601470 | 9109903 | 6717419.5 | 6564023.5 | 6683475 | 9631345 | 7628518 |
| Q56R16 | Kpna5 | Importin subunit alpha-6 Rattus norvegicus | 233.82 | 9.89 | 60299 | 1 | 5 | 4009986.5 | 4253468.5 | 4582332 | 4451326 | 2915368.25 | 4371649 | 4259116.5 | 3469477.25 | 4299261.5 | 2885765 | 4194815 | 9234573 | 8813116 | 4743917 |
| O35345 | Kpna6 | Importin subunit alpha-7 Mus musculus | 324.87 | 13.99 | 59964 | 2 | 5 | 4916037.5 | 3161914.75 | 3009315.75 | 5540338.5 | 1484844.38 | 4576209 | 5015974 | 1784129 | 6050003.5 | 4343048.5 | 5278684.5 | 5835186.5 | 6279347 | 3519768 |
| P70168 | Kpnb1 | Importin subunit beta-1 Mus musculus | 1892.39 | 55.25 | 97184 | 30 | 31 | 89736977 | 92410529 | 92801505 | 78001705 | 70889225 | 92674329 | 90374081 | 104734057 | 99893097 | 87161753 | 82264729 | 99858873 | 102962977 | 88590537 |
| Q8BGA5 | Krr1 | KRR1 small subunit processome component homolog Mus musculus | 104.57 | 7.89 | 43538 | 3 | 3 | 6586508 | 7482851.5 | 6580950 | 8051483 | 7085955.5 | 7442363 | 6488608.5 | 7578450 | 6810511.5 | 6883826 | 5822115.5 | 6639074.5 | 6526532.5 | 7094742 |
| Q61595 | Ktn1 | Kinectin Mus musculus | 162.01 | 3.47 | 152592 | 3 | 3 | 362564.12 | 347393.81 | 616992.12 | 217837.8 | 567984.19 | 695818 | 627839.69 | 544844 | 576056.62 | 738287.56 | 607491.25 | 563299.5 | 652587.94 | 603573.12 |
| Q8BZT9 | Lacc1 | Laccase domain-containing protein 1 Mus musculus | 54.75 | 3.95 | 47514 | 1 | 1 | 1 | 311313.31 | 280255.16 | 1 | 407837.97 | 210776.0975 | 244956.7 | 177187.66 | 222418.22 | 198541.81 | 253784.34 | 344585.38 | 172593.59 | 194435 |
| Q9EP89 | Lactb | Serine beta-lactamase-like protein LACTB, mitochondrial Mus musculus | 233.63 | 10.89 | 60705 | 5 | 5 | 3384683 | 3194884 | 2470866 | 3067446 | 1985797.75 | 3243407.5 | 2770043.5 | 2778257.5 | 2207083 | 2848978 | 2424302.75 | 2048506.5 | 2655592 | 3000379.25 |
| Q99KR3 | Lactb2 | Endoribonuclease LACTB2 Mus musculus | 137.57 | 11.46 | 32754 | 3 | 3 | 3368817.5 | 2992152.5 | 3056815.25 | 3435738 | 3577917 | 3739374.25 | 3664004.5 | 3710150.25 | 3487669.5 | 3706514.25 | 3435647 | 3399116 | 3602768.5 | 3727644.75 |
| P11438 | Lamp1 | Lysosome-associated membrane glycoprotein 1 Mus musculus | 568.22 | 28.33 | 43865 | 8 | 8 | 43631817 | 55507593 | 51127957 | 45247497 | 51616885 | 58910861 | 54424409 | 69183185 | 59832969 | 52330705 | 47542797 | 55563593 | 55145113 | 44633385 |
| P17047 | Lamp2 | Lysosome-associated membrane glycoprotein 2 Mus musculus | 263.8 | 12.29 | 45681 | 5 | 5 | 142638177 | 104922305 | 105360225 | 135854225 | 149493297 | 156102865 | 141263873 | 126288897 | 144596321 | 128989089 | 122947241 | 121529457 | 130368721 | 99882673 |
| Q6P791 | Lamtor1 | Ragulator complex protein LAMTOR1 Rattus norvegicus | 96.67 | 16.15 | 17721 | 2 | 2 | 3387687.5 | 3668629.5 | 2649623.5 | 3601528.25 | 3677311 | 2439902.5 | 2771505 | 2523526.75 | 1127593.12 | 2774678 | 2861861 | 2781462.5 | 2352882.5 | 2578986.5 |
| Q9JHS3 | Lamtor2 | Ragulator complex protein LAMTOR2 Mus musculus | 327.95 | 75.2 | 13480 | 5 | 5 | 3061190 | 3380003 | 2833678 | 3459445 | 3677292 | 3135139.75 | 2842444.5 | 3054852 | 2685858.25 | 2623011 | 2955658.5 | 3180415.5 | 3554531.5 | 1805479.12 |
| O88653 | Lamtor3 | Ragulator complex protein LAMTOR3 Mus musculus | 54.18 | 8.87 | 13553 | 1 | 1 | 332034.62 | 235556.84 | 417342.41 | 1 | 1 | 377075.75 | 315712.06 | 588976.12 | 473379.72 | 211296.47 | 269096.16 | 346450.81 | 464246.56 | 359931.1767 |
| Q9D1L9 | Lamtor5 | Ragulator complex protein LAMTOR5 Mus musculus | 51.75 | 21.98 | 9642 | 1 | 1 | 2614597.25 | 2651509.75 | 3639202.75 | 2723047 | 3201065.75 | 2272561.25 | 2567972.75 | 2065061.38 | 2366158.75 | 2342385.5 | 3204254.5 | 2217009.25 | 2566614.5 | 3257206 |
| O89112 | Lancl1 | LanC-like protein 1 Mus musculus | 192.41 | 13.28 | 45341 | 4 | 4 | 2849887.5 | 2832787.5 | 2081036 | 2611689.75 | 2867080 | 2505188 | 2808023 | 3763544 | 3026691.5 | 2405063.25 | 2454414.75 | 2514682 | 2729176.5 | 2721831.5 |
| Q9CPY7 | Lap3 | Cytosol aminopeptidase Mus musculus | 787.76 | 39.5 | 56141 | 15 | 15 | 15427193 | 19944985 | 18782535 | 19058895 | 12815697 | 21164185 | 14973483 | 18034805 | 17258759 | 16581409 | 20239917 | 19349123 | 18955045 | 21169639 |
| Q6ZQ58 | Larp1 | La-related protein 1 Mus musculus | 512.21 | 12.5 | 121125 | 10 | 10 | 2039866.5 | 3495226.75 | 3501751.75 | 3309802.25 | 3028093.25 | 2788713.75 | 3139366.25 | 2948463 | 3460307 | 3001768.5 | 3664658.5 | 3957481.75 | 3642988.5 | 2058757.75 |
| Q8BWW4 | Larp4 | La-related protein 4 Mus musculus | 347.74 | 10.29 | 79763 | 6 | 6 | 5036020 | 4904411 | 5728868 | 5108949 | 6015560 | 3549385.5 | 3411596.25 | 5684779 | 4481363 | 2653790.75 | 3125525 | 4126945.75 | 5258634.5 | 4493025 |
| Q6A0A2 | Larp4b | La-related protein 4B Mus musculus | 316.71 | 11.2 | 81627 | 6 | 6 | 832142.81 | 3606668.5 | 2519778.75 | 3920381 | 2112063.25 | 1179564.88 | 3482647.5 | 2362489.25 | 3561082.5 | 3482375.25 | 2453327 | 2167968.75 | 2259181 | 3148418.5 |
| Q05CL8 | Larp7 | La-related protein 7 Mus musculus | 297.48 | 11.4 | 64802 | 5 | 5 | 10513111 | 11670235 | 7440189.5 | 11271201 | 11376124 | 9429243 | 6000653 | 6368686 | 5520512 | 10661035 | 10089153 | 8844383 | 7844631 | 5328739 |
| Q8BMJ2 | Lars | Leucine--tRNA ligase, cytoplasmic Mus musculus | 1924.83 | 41.17 | 134192 | 41 | 41 | 46871145 | 43986337 | 46065385 | 49140165 | 46022913 | 47200281 | 46106457 | 40900377 | 47557709 | 38383713 | 41450177 | 48358873 | 49389049 | 46705141 |
| A2BE28 | Las1l | Ribosomal biogenesis protein LAS1L Mus musculus | 58.04 | 2.19 | 89414 | 1 | 1 | 697141.19 | 724645.56 | 741313.75 | 604094.31 | 491069.44 | 450763.06 | 485417.66 | 775932.62 | 672006.12 | 463982 | 521777.2067 | 457239.81 | 595622.12 | 512469.69 |
| Q61792 | Lasp1 | LIM and SH3 domain protein 1 Mus musculus | 418.55 | 28.9 | 29994 | 8 | 8 | 11757164 | 14238055 | 13827363 | 17997865 | 12548104 | 16043297 | 17041625 | 13314048 | 16789501 | 20093063 | 15520823 | 16271151 | 15856422 | 11591105 |
| Q3U9G9 | Lbr | Lamin-B receptor Mus musculus | 538.78 | 18.21 | 71440 | 8 | 9 | 20571297 | 4351733 | 7152046 | 7971330 | 8114032.5 | 8228797 | 6832436 | 6387961 | 7362729.5 | 7993963.5 | 6430120.5 | 8098242 | 8919548 | 7337112 |
| Q6P4Z6 | Lcmt1 | Leucine carboxyl methyltransferase 1 Rattus norvegicus | 44.66 | 2.71 | 38330 | 1 | 1 | 2115761.75 | 2183631.25 | 1710164.75 | 2253409.25 | 1937010.12 | 1755236.88 | 1529209.62 | 1369825.75 | 1517203.12 | 1654059 | 2062621.88 | 2175203.75 | 1453889.38 | 1705589.5 |
| Q61233 | Lcp1 | Plastin-2 Mus musculus | 4110.059 | 87.88 | 70149 | 48 | 62 | 1483341441 | 1456825729 | 1599929345 | 1449602433 | 1517976193 | 1695252353 | 1459549953 | 1646451457 | 1754942465 | 1675444481 | 1588014593 | 1606976001 | 1637279873 | 1627295233 |
| Q60787 | Lcp2 | Lymphocyte cytosolic protein 2 Mus musculus | 350.12 | 15.95 | 60238 | 7 | 7 | 4554089.5 | 3082293.5 | 3843718.5 | 1894505.88 | 2389610.75 | 4557804 | 3986660 | 2597825.5 | 2306852.5 | 1693727.5 | 2790840.25 | 2999953 | 1800517.62 | 1829555.75 |
| P06151 | Ldha | L-lactate dehydrogenase A chain Mus musculus | 2215.19 | 87.05 | 36499 | 28 | 32 | 712083585 | 734669377 | 670777793 | 591029249 | 691721537 | 678447233 | 753604353 | 773292737 | 765662209 | 786916609 | 693276353 | 711156929 | 671447425 | 675062529 |
| P35951 | Ldlr | Low-density lipoprotein receptor Mus musculus | 247.14 | 7.31 | 94947 | 4 | 5 | 503156.88 | 299449.81 | 499312.5 | 751841.75 | 937581.75 | 561302.94 | 586157.71 | 839636.06 | 580659.12 | 363032.72 | 672347.38 | 1209777.25 | 829231.62 | 995924.69 |
| Q8C142 | Ldlrap1 | Low density lipoprotein receptor adapter protein 1 Mus musculus | 154.1 | 12.34 | 33975 | 3 | 3 | 5834719.5 | 5932507 | 2432658.75 | 5441983 | 3378121.5 | 3232427 | 4572475.5 | 4592041.5 | 3470473.25 | 4686776 | 2909007 | 2322111 | 3378801.5 | 5093773.5 |
| Q6DVA0 | Lemd2 | LEM domain-containing protein 2 Mus musculus | 81.54 | 5.09 | 57507 | 2 | 2 | 1213773.5 | 1456737.88 | 1518883.25 | 1223020.12 | 1783008.38 | 2192399.5 | 1497458.5 | 1929877.62 | 1837782 | 1986906.12 | 1614966.12 | 1490715.5 | 2041762.12 | 2216279.25 |
| Q5XJE5 | Leo1 | RNA polymerase-associated protein LEO1 Mus musculus | 98.94 | 3.45 | 75597 | 2 | 2 | 1584213.25 | 626570.31 | 2074630.12 | 2103116 | 1275064.25 | 512013.22 | 1925142.62 | 558892.62 | 1584263.62 | 1684165.12 | 2093589.25 | 1426254.88 | 1896098.62 | 1724526.25 |
| Q9Z2I0 | Letm1 | Mitochondrial proton/calcium exchanger protein Mus musculus | 228.53 | 15.85 | 82989 | 5 | 5 | 3292706 | 3563616.75 | 3094787.5 | 3588668 | 3040835.5 | 3588820.75 | 3228422.75 | 3624931 | 4342376.5 | 3039779.5 | 3983413 | 3939329.5 | 3501451.5 | 3433051 |
| P16045 | Lgals1 | Galectin-1 Mus musculus | 1008.62 | 92.59 | 14866 | 12 | 14 | 395518401 | 477600097 | 483867105 | 452777025 | 462985857 | 425536929 | 407105249 | 431826913 | 439702401 | 434879329 | 457049121 | 427814049 | 480328513 | 486210817 |
| P16110 | Lgals3 | Galectin-3 Mus musculus | 516.11 | 36.36 | 27515 | 8 | 8 | 66420117 | 61348193 | 74391713 | 82619921 | 79175249 | 76876129 | 77264337 | 55938101 | 77543793 | 104901417 | 86584249 | 70924073 | 87247065 | 85772545 |
| Q07797 | Lgals3bp | Galectin-3-binding protein Mus musculus | 1156.02 | 39.34 | 64491 | 16 | 16 | 84722809 | 85872353 | 84740609 | 60414577 | 75841993 | 48594913 | 47905609 | 47771129 | 44640049 | 45468081 | 48904841 | 45471689 | 70658721 | 50665385 |
| O89017 | Lgmn | Legumain Mus musculus | 339.48 | 14.48 | 49373 | 5 | 5 | 16174174 | 13847101 | 17326819 | 8485222 | 10875123 | 17590017 | 15370876 | 17198165 | 15729986 | 14117877 | 18229133 | 17334623 | 20155363 | 15862572 |
| Q5I0D5 | Lhpp | Phospholysine phosphohistidine inorganic pyrophosphate phosphatase Rattus norvegicus | 149.53 | 11.85 | 29190 | 2 | 2 | 1550441.62 | 1435424.5 | 365567.44 | 2014820.5 | 1277171.75 | 1958688.25 | 1717775.5 | 1862611 | 1935752.5 | 1804088.5 | 1779357.75 | 833488.56 | 1825399 | 954739.94 |
| P37913 | Lig1 | DNA ligase 1 Mus musculus | 1179.01 | 34.83 | 102290 | 23 | 23 | 14698591 | 15500032 | 14990569 | 18926393 | 14219613 | 14135799 | 14556043 | 14874858 | 15585770 | 15938075 | 15408687 | 21520149 | 14595616 | 17564347 |
| P97386 | Lig3 | DNA ligase 3 Mus musculus | 132.74 | 3.45 | 113072 | 3 | 3 | 2243806 | 2093747.75 | 2771492.25 | 2356689.5 | 2377674.5 | 2333669.25 | 2591394 | 2356729.5 | 2440665 | 2450938.5 | 2365112 | 2583741 | 2403936.5 | 2681469.5 |
| Q64281 | Lilrb4 | Leukocyte immunoglobulin-like receptor subfamily B member 4 Mus musculus | 272.17 | 22.09 | 37544 | 4 | 4 | 6278929.5 | 6655734.5 | 7452466.5 | 7018348 | 6715694 | 7567146 | 8171333 | 6839654.5 | 6650095 | 6940729 | 7083005 | 5622785 | 6841767 | 6296285 |
| Q4KM31 | Limd2 | LIM domain-containing protein 2 Rattus norvegicus | 61.9 | 13.28 | 14236 | 1 | 1 | 5087711 | 5415087 | 5407738 | 5114712 | 5454524 | 3992986.5 | 5148498 | 4087321.25 | 4910629.5 | 5214855 | 3680553 | 4044729.25 | 4217600 | 3957633 |
| Q99JW4 | Lims1 | LIM and senescent cell antigen-like-containing domain protein 1 Mus musculus | 64.65 | 4.62 | 37240 | 1 | 1 | 807918.62 | 615479.19 | 607873.88 | 693736.845 | 743675.69 | 809672.38 | 741941.62 | 715139.19 | 923332.19 | 976209.12 | 1013626.94 | 791070 | 912606.94 | 725541.19 |
| Q9Z0M5 | Lipa | Lysosomal acid lipase/cholesteryl ester hydrolase Mus musculus | 683.19 | 22.17 | 45325 | 9 | 9 | 14897863 | 18597939 | 18244205 | 13282737 | 12877767 | 27890647 | 30345103 | 34120141 | 34454009 | 32353231 | 40208761 | 55868197 | 56020901 | 45081977 |
| P54310 | Lipe | Hormone-sensitive lipase Mus musculus | 253.27 | 10.01 | 83348 | 4 | 4 | 1120625 | 1076144.75 | 1234195.38 | 588111.56 | 545402.5 | 958237.38 | 829990.75 | 948834.5 | 859638.75 | 656634.38 | 971813.06 | 861192 | 729057 | 939846.88 |
| Q9D0F3 | Lman1 | Protein ERGIC-53 Mus musculus | 395.53 | 26.5 | 57789 | 8 | 8 | 23873111 | 23560533 | 23336553 | 21160401 | 25816227 | 25166201 | 23478273 | 27150179 | 25446841 | 25532127 | 23328765 | 23878237 | 25084655 | 27300421 |
| Q9DBH5 | Lman2 | Vesicular integral-membrane protein VIP36 Mus musculus | 187.23 | 18.99 | 40430 | 4 | 4 | 3604307 | 3640636.5 | 3198043.5 | 2183465.5 | 2834169.5 | 3254505 | 2430495.5 | 2864373.75 | 3194454.5 | 2777745.25 | 2715193 | 2178250.5 | 3992701.5 | 3045847.25 |
| P48678 | Lmna | Prelamin-A/C Mus musculus | 2521.97 | 56.54 | 74238 | 39 | 42 | 281224001 | 298698145 | 292395201 | 342851937 | 302216961 | 316718401 | 340511585 | 319097633 | 315763201 | 348189089 | 325984193 | 303872641 | 302146529 | 311966817 |
| P14733 | Lmnb1 | Lamin-B1 Mus musculus | 2033.55 | 62.24 | 66786 | 29 | 34 | 87113625 | 89518945 | 92807689 | 108647713 | 102776625 | 97557753 | 99677641 | 89985601 | 91674953 | 107331577 | 97698281 | 93289873 | 92570049 | 100102889 |
| P21619 | Lmnb2 | Lamin-B2 Mus musculus | 713.07 | 25.84 | 67318 | 10 | 15 | 24706799 | 24913981 | 25952291 | 25501765 | 26472293 | 29523549 | 28922769 | 22451395 | 26395947 | 30991325 | 28984125 | 28020161 | 25399105 | 28776439 |
| Q8C129 | Lnpep | Leucyl-cystinyl aminopeptidase Mus musculus | 682.9 | 13.95 | 117304 | 11 | 11 | 12268318 | 9276440 | 12960425 | 9956491 | 8948255 | 17623309 | 15894234 | 16932119 | 20007805 | 15619805 | 10285751 | 12964193 | 16254150 | 14058953 |
| Q8CGK3 | Lonp1 | Lon protease homolog, mitochondrial Mus musculus | 1072.35 | 34.04 | 105843 | 20 | 20 | 28760953 | 28925937 | 29730303 | 31896783 | 30825573 | 39436801 | 39184665 | 39297381 | 35430021 | 34813457 | 37455065 | 32411825 | 35603089 | 31406583 |
| Q3TFD2 | Lpcat1 | Lysophosphatidylcholine acyltransferase 1 Mus musculus | 222.91 | 8.61 | 59744 | 4 | 4 | 1168222 | 1306953.75 | 1139208.25 | 1965804.25 | 1955921.25 | 1562880.75 | 975118.5 | 1775242.5 | 735296.44 | 1031170.44 | 1412343.25 | 2123016.5 | 1855290.75 | 1758101.25 |
| Q8BYI6 | Lpcat2 | Lysophosphatidylcholine acyltransferase 2 Mus musculus | 480.64 | 21.14 | 60254 | 6 | 10 | 2924251.5 | 3248657.25 | 5680712 | 3541806 | 3749135.5 | 5552612 | 6246263 | 6334870 | 6720229 | 5677482 | 4759444 | 5573457 | 7019060.5 | 6105467.5 |
| P11152 | Lpl | Lipoprotein lipase Mus musculus | 177.45 | 11.18 | 53109 | 3 | 3 | 308996.12 | 678906.56 | 317656.5 | 1969860 | 818854.795 | 230586.58 | 288498.69 | 833030.62 | 1342658.25 | 317949.41 | 430611.5 | 294375.88 | 1029638.25 | 821505.44 |
| Q99N69 | Lpxn | Leupaxin Mus musculus | 542.15 | 26.94 | 43478 | 7 | 7 | 6798333.5 | 7566228.5 | 7889414 | 7975122 | 4547158 | 7304910 | 7252421 | 7965298.5 | 9928673 | 7954476 | 5731387 | 7407565 | 7541129 | 5970694 |
| Q8BVU0 | Lrch3 | Leucine-rich repeat and calponin homology domain-containing protein 3 Mus musculus | 35.82 | 2.44 | 86341 | 1 | 1 | 1 | 1 | 1 | 1 | 101635.62 | 1 | 1 | 1 | 1 | 70568.89 | 94790.66667 | 101658.43 | 73287.95 | 109425.62 |
| Q921G6 | Lrch4 | Leucine-rich repeat and calponin homology domain-containing protein 4 Mus musculus | 192.74 | 8.97 | 73163 | 3 | 3 | 1620449 | 1876541.88 | 1666251.5 | 1657480 | 2064907.25 | 1279420.12 | 2099251.5 | 1644682 | 2250551.75 | 2230971.25 | 1896072 | 769949.44 | 2304446.75 | 1906343 |
| Q60664 | Lrmp | Lymphoid-restricted membrane protein Mus musculus | 110.8 | 5.57 | 59588 | 2 | 2 | 2122874.5 | 1019856.94 | 720089.81 | 1669346 | 1802042.38 | 1762830.5 | 1621125.25 | 1612228.5 | 867298.94 | 1466019.12 | 1544940.12 | 2147245.5 | 1489350.75 | 2003574 |
| Q91ZX7 | Lrp1 | Prolow-density lipoprotein receptor-related protein 1 Mus musculus | 3428.96 | 18.13 | 504742 | 59 | 60 | 98539633 | 125103761 | 100120561 | 105283761 | 133641889 | 118375633 | 102409449 | 105671177 | 107297073 | 114920497 | 120053249 | 100440801 | 125445433 | 125024665 |
| Q9JI18 | Lrp1b | Low-density lipoprotein receptor-related protein 1B Mus musculus | 63.99 | 0.37 | 513634 | 1 | 2 | 1296615.25 | 2340284.75 | 1373193 | 1505938.5 | 1582396.62 | 1746386.5 | 1223866.75 | 1739599.12 | 1240024.62 | 1170115.5 | 1517052.88 | 1048227.38 | 1539044.38 | 1516726.88 |
| P55302 | Lrpap1 | Alpha-2-macroglobulin receptor-associated protein Mus musculus | 371.87 | 25.56 | 42215 | 7 | 7 | 25035591 | 17206813 | 19915209 | 19133793 | 15211021 | 19464997 | 23976985 | 22647863 | 19895129 | 23974017 | 20921965 | 18047123 | 21908223 | 19979023 |
| Q6PB66 | Lrpprc | Leucine-rich PPR motif-containing protein, mitochondrial Mus musculus | 1840.37 | 30.17 | 156615 | 24 | 33 | 51690861 | 44126069 | 48019153 | 43048493 | 46688893 | 47828125 | 46427785 | 48814081 | 48702737 | 45659057 | 50447165 | 49696189 | 55459893 | 51356921 |
| Q8K1T1 | Lrrc25 | Leucine-rich repeat-containing protein 25 Mus musculus | 53.64 | 6.06 | 32673 | 1 | 1 | 654861.69 | 776330.31 | 764671.89 | 721633 | 905862.56 | 624420.81 | 1 | 1 | 1 | 1 | 596315.12 | 595834 | 671994.12 | 608548.25 |
| Q9CRC8 | Lrrc40 | Leucine-rich repeat-containing protein 40 Mus musculus | 346.08 | 14.78 | 68076 | 7 | 7 | 6767478 | 4537993 | 5081819.5 | 4743589 | 5067823 | 5382521.5 | 3347321 | 5345219.5 | 5726484.5 | 4621923 | 6160940 | 6929887 | 6867298 | 6217938 |
| Q8K1C9 | Lrrc41 | Leucine-rich repeat-containing protein 41 Mus musculus | 136.54 | 7.56 | 88180 | 3 | 3 | 496309.31 | 682914.62 | 789214 | 456527 | 560264.12 | 450324.66 | 814074.12 | 635692.62 | 724581 | 590136.88 | 816692.94 | 684346.5 | 740446.25 | 652286.44 |
| Q505F5 | Lrrc47 | Leucine-rich repeat-containing protein 47 Mus musculus | 188.49 | 11.88 | 63590 | 4 | 4 | 2665587 | 2880447.25 | 2403833.75 | 2991008 | 3062402.5 | 2181662.5 | 2430071.25 | 2210709.5 | 2539899 | 2393514 | 3145687.75 | 2741638.25 | 2095935.75 | 2574405.25 |
| Q5RJR8 | Lrrc59 | Leucine-rich repeat-containing protein 59 Rattus norvegicus | 560.67 | 38.44 | 34869 | 10 | 10 | 61149985 | 65145753 | 58488453 | 57033969 | 60603913 | 62902317 | 65926569 | 62337193 | 63393881 | 64312753 | 66181505 | 61435385 | 64701225 | 67535953 |
| Q498T9 | Lrrc8c | Volume-regulated anion channel subunit LRRC8C Rattus norvegicus | 181.54 | 3.86 | 92463 | 2 | 2 | 285118.25 | 180492.56 | 295830.62 | 227927.3775 | 150268.08 | 272492.41 | 227617.89 | 253047.97 | 312444.03 | 464188.38 | 215423.53 | 252929.72 | 177604.17 | 175398.95 |
| Q3UZ39 | Lrrfip1 | Leucine-rich repeat flightless-interacting protein 1 Mus musculus | 1055.96 | 27.43 | 79249 | 16 | 16 | 25806603 | 32912419 | 25704533 | 31371381 | 31016885 | 30377317 | 30588865 | 31929741 | 31359669 | 29773481 | 27168693 | 26495901 | 26604175 | 29526207 |

| | | | | | | | | | | | | | | | | | | | | | |
|---|---|---|---|---|---|---|---|---|---|---|---|---|---|---|---|---|---|---|---|---|---|
| Q3UM18 | Lsg1 | Large subunit GTPase 1 homolog Mus musculus | 67.9 | 4.66 | 73157 | 1 | 1 | 676685.31 | 163861.2 | 1 | 1 | 222588.08 | 272614.28 | 122867.09 | 99781.29 | 92686.05 | 95887.99 | 280662.47 | 177658.47 | 144048.66 | 120657.27 |
| Q9D0R8 | Lsm12 | Protein LSM12 homolog Mus musculus | 170.35 | 27.69 | 21701 | 3 | 3 | 1417764.38 | 1503190.063 | 1301414.62 | 1666060.5 | 1627520.75 | 1846937.25 | 2037866.5 | 1277141.38 | 1918455.62 | 1842879.62 | 1656458.12 | 2079249.5 | 1748233.25 | 1478397.75 |
| Q8K2F8 | Lsm14a | Protein LSM14 homolog A Mus musculus | 113.18 | 6.28 | 50546 | 2 | 2 | 507717.91 | 651041.44 | 616954.69 | 1233059.25 | 1107622.38 | 880299.88 | 312389.16 | 314359.72 | 873240.5 | 781324.31 | 927596.3533 | 1107089.62 | 662223.75 | 1013475.69 |
| O35900 | Lsm2 | U6 snRNA-associated Sm-like protein LSm2 Mus musculus | 229.45 | 60 | 10835 | 4 | 4 | 2667687 | 1314288.12 | 682846.94 | 1468977.62 | 1897637.25 | 2407252 | 2061215.88 | 2841446.5 | 2269167.5 | 1728343.25 | 2241752.5 | 2004164.12 | 1858332.25 | 725637.19 |
| P62311 | Lsm3 | U6 snRNA-associated Sm-like protein LSm3 Mus musculus | 139.07 | 32.35 | 11845 | 2 | 2 | 3110765 | 3487201.5 | 3687708 | 4149744 | 5675498.5 | 5410144 | 3086265 | 2336221.25 | 3664570.5 | 5244513.5 | 5538689 | 5762724.5 | 6036211 | 5347757.5 |
| Q9QXA5 | Lsm4 | U6 snRNA-associated Sm-like protein LSm4 Mus musculus | 83.69 | 12.41 | 15076 | 2 | 2 | 1314896.5 | 2189684 | 1335778.5 | 1566682.25 | 1819093.5 | 1454204.62 | 1586266 | 1568759.88 | 1298857.5 | 1354238.75 | 1593969.75 | 1392736.5 | 1436788.62 | 1613311.88 |
| P62322 | Lsm5 | U6 snRNA-associated Sm-like protein LSm5 Mus musculus | 81.46 | 20.88 | 9937 | 1 | 1 | 3710096.25 | 3849165.75 | 3430881.5 | 4940065.5 | 4260182.5 | 3646703 | 3888621.75 | 4051761.25 | 3105697.25 | 3563818.5 | 4042722.5 | 3770828.5 | 3502299.75 | 3702172.25 |
| P62313 | Lsm6 | U6 snRNA-associated Sm-like protein LSm6 Mus musculus | 43.15 | 10 | 9128 | 1 | 1 | 3327701 | 2948293.25 | 2895213.25 | 3063587.25 | 2961613 | 2748458.25 | 2807458.5 | 2351072 | 2245264.75 | 2222299.5 | 2413814 | 3338756.5 | 2971443.75 | 2118903.75 |
| Q9CQQ8 | Lsm7 | U6 snRNA-associated Sm-like protein LSm7 Mus musculus | 125.58 | 33.01 | 11636 | 3 | 3 | 4030209.75 | 3583573.75 | 2910486.75 | 4427214.5 | 3855994.25 | 3057894.75 | 3083905.25 | 3220732.75 | 2512152.5 | 2943542.5 | 3393962 | 2946428.5 | 3241128 | 3643697.75 |
| Q6ZWM4 | Lsm8 | U6 snRNA-associated Sm-like protein LSm8 Mus musculus | 357.2 | 82.29 | 10403 | 5 | 5 | 6014043.5 | 6108049 | 5574844.5 | 6910368 | 7500382 | 5870320.5 | 5834091 | 5416966 | 5524660 | 6544393 | 7040619.5 | 5989835 | 6354515 | 6052931 |
| P19973 | Lsp1 | Lymphocyte-specific protein 1 Mus musculus | 677.48 | 42.12 | 36714 | 9 | 9 | 91496321 | 96268945 | 91651585 | 113484225 | 131125113 | 102667193 | 105248057 | 83274681 | 102107057 | 120220633 | 113582625 | 99195145 | 97263329 | 110756193 |
| Q8BLN5 | Lss | Lanosterol synthase Mus musculus | 321.46 | 8.59 | 83141 | 2 | 5 | 5062840 | 5013338 | 4277994 | 4248439 | 3634731.75 | 4353357 | 4677803.5 | 4964144 | 3946070.75 | 3678749.25 | 3561984.5 | 4310518 | 4931656 | 6373000.5 |
| P24527 | Lta4h | Leukotriene A-4 hydrolase Mus musculus | 476.73 | 19.15 | 69051 | 9 | 9 | 11273067 | 12678380 | 11299983 | 12328483 | 8893505 | 14530157 | 12769497 | 14811241 | 12085676 | 12902393 | 10949869 | 13702669 | 11657485 | 11071309 |
| Q6A009 | Ltn1 | E3 ubiquitin-protein ligase listerin Mus musculus | 141.11 | 1.08 | 198921 | 2 | 2 | 173121.45 | 418949.69 | 139976.73 | 341836.88 | 295224.5 | 125300.26 | 249701.3 | 1 | 1 | 161657.66 | 275862.94 | 244078.45 | 197421.86 | 150750.34 |
| Q68FR7 | Ltv1 | Protein LTV1 homolog Rattus norvegicus | 48.27 | 4.04 | 53562 | 1 | 1 | 1 | 1 | 1 | 483846.47 | 1 | 585971.62 | 455048.16 | 471036.62 | 452096.59 | 491038.2475 | 1 | 1 | 548177.06 | 357973.03 |
| Q9CYI4 | Luc7l | Putative RNA-binding protein Luc7-like 1 Mus musculus | 229.78 | 15.36 | 43934 | 3 | 6 | 10329964 | 8477223 | 9781321 | 10140205 | 10196263 | 11279487 | 12236431 | 9379219 | 11127909 | 11139125 | 9468247 | 10387012 | 9461750 | 9955047 |
| Q7TNC4 | Luc7l2 | Putative RNA-binding protein Luc7-like 2 Mus musculus | 395.75 | 20.15 | 46583 | 5 | 8 | 23241649 | 21309971 | 22017795 | 25316113 | 23186323 | 24192105 | 25695821 | 22717333 | 24786427 | 25735951 | 22069021 | 22631187 | 20934327 | 22793487 |
| Q5SUF2 | Luc7l3 | Luc7-like protein 3 Mus musculus | 640.75 | 26.39 | 51450 | 9 | 10 | 9580275 | 9362646 | 12781082 | 11444175 | 8638739 | 8646819 | 15234064 | 6691071 | 10212407 | 13814043 | 5733710.5 | 9976912 | 10796009 | 7993212.5 |
| P70202 | Lxn | Latexin Mus musculus | 92.6 | 9.46 | 25492 | 2 | 2 | 4335441.5 | 5021072 | 4927896 | 4757538.5 | 4461408.5 | 3813518 | 4037597.75 | 4750108.5 | 4314252.5 | 3536690 | 3864358.75 | 3692705.25 | 3778510 | 3973488.75 |
| O88188 | Ly86 | Lymphocyte antigen 86 Mus musculus | 309.35 | 43.83 | 17811 | 4 | 4 | 5974965.5 | 6362964.5 | 5429174.5 | 3877232.75 | 5239046 | 2333309.25 | 2539961.75 | 4649433.5 | 3992437.5 | 3548827.75 | 3586800.25 | 3616600.5 | 3760531 | 3984031.25 |
| Q08288 | Lyar | Cell growth-regulating nucleolar protein Mus musculus | 79.37 | 3.61 | 43736 | 1 | 1 | 84915.59 | 120043.96 | 108362.73 | 126396.12 | 193602.12 | 89651.73 | 109313.53 | 117996.16 | 139914.08 | 115058.91 | 170829.55 | 134126.39 | 146897.22 | 164263.81 |
| P25911 | Lyn | Tyrosine-protein kinase Lyn Mus musculus | 639.8 | 34.18 | 58812 | 12 | 12 | 19735451 | 21564789 | 19844963 | 14761444 | 15744967 | 20086575 | 17147273 | 16132549 | 19059299 | 17024837 | 14160733 | 16532984 | 18757531 | 15399210 |
| P97823 | Lypla1 | Acyl-protein thioesterase 1 Mus musculus | 226.94 | 26.52 | 24688 | 4 | 4 | 4853141 | 6326202.5 | 5610991.5 | 5826329 | 6182496 | 4784356 | 5914389 | 5436433.5 | 4269883 | 4777823 | 7005920 | 5773841.5 | 5858104.5 | 5495080 |
| Q9QYL8 | Lypla2 | Acyl-protein thioesterase 2 Rattus norvegicus | 40.87 | 10.82 | 24807 | 1 | 1 | 875456.56 | 942451.81 | 740642.75 | 577799.25 | 760094.81 | 987983.12 | 1483220.62 | 2577308.75 | 2390149.25 | 1009287.06 | 2236100.75 | 1277549.75 | 1083784.25 | 1011178.44 |
| P08905 | Lyz2 | Lysozyme C-2 Mus musculus | 403.81 | 49.32 | 16689 | 4 | 4 | 23830993 | 30157265 | 25073081 | 23394253 | 22318589 | 19638397 | 21972163 | 22522759 | 15524209 | 19922069 | 20743953 | 17378173 | 21759069 | 23216407 |
| P24668 | M6pr | Cation-dependent mannose-6-phosphate receptor Mus musculus | 423.18 | 32.37 | 31172 | 6 | 6 | 27936077 | 31874237 | 26059501 | 30118059 | 28856205 | 28829245 | 31497911 | 35144113 | 25739449 | 29946513 | 29532125 | 24204737 | 27500529 | 29504725 |
| Q9QXZ0 | Macf1 | Microtubule-actin cross-linking factor 1 Mus musculus | 1182.11 | 4.98 | 831878 | 26 | 27 | 6852718.5 | 8346064 | 8966120 | 7128024 | 6447014 | 9448019 | 9302008 | 10202712 | 12248702 | 9357708 | 10193320 | 11638077 | 9482751 | 10385722 |
| Q9Z1B5 | Mad2l1 | Mitotic spindle assembly checkpoint protein MAD2A Mus musculus | 341.8 | 37.56 | 23598 | 6 | 6 | 3833004.5 | 7963356 | 6602414 | 5019572.5 | 4244114.5 | 1950531.12 | 6115794.5 | 5736827.5 | 5756840 | 3780349.25 | 3260969.75 | 5235739.5 | 6076085 | 7297557 |
| Q80U28 | Madd | MAP kinase-activating death domain protein Mus musculus | 55.4 | 1.52 | 175180 | 1 | 1 | 351095.78 | 223346.52 | 1 | 1 | 187522.52 | 252006.27 | 311276.66 | 222518.64 | 295145.09 | 223564.06 | 291033 | 292121.44 | 221945.31 | 211473.23 |
| P61327 | Magoh | Protein mago nashi homolog Mus musculus | 278.44 | 56.85 | 17164 | 6 | 6 | 5254852 | 2559180.25 | 2357303.5 | 2722990 | 3714786.5 | 4584322 | 2493556.5 | 5820958.5 | 3368054.5 | 3730514.75 | 3657454 | 4830306 | 3948736.5 | 4084153 |
| Q6AY04 | Maip1 | m-AAA protease-interacting protein 1, mitochondrial Rattus norvegicus | 91.79 | 8.59 | 33090 | 2 | 2 | 1 | 1 | 1380012.25 | 1500286.62 | 1859372.12 | 445459 | 1058742.813 | 1151396.5 | 1411363.5 | 1226752.25 | 1409163.88 | 1177544.5 | 1247737.25 | 1540197.12 |
| Q8BGS0 | Mak16 | Protein MAK16 homolog Mus musculus | 150.13 | 11.49 | 35185 | 3 | 3 | 1338627.62 | 1080735.5 | 1124333.88 | 1422618.5 | 1444412.88 | 1523970.75 | 1004266.5 | 1171150.123 | 1123038.62 | 1033324.62 | 1234404.12 | 1050477 | 1479515.25 | 933844.69 |
| P27046 | Man2a1 | Alpha-mannosidase 2 Mus musculus | 74.78 | 2.96 | 131631 | 2 | 2 | 245755.61 | 444731.75 | 384437.62 | 267502.25 | 468527.28 | 551889.19 | 96172.29 | 87442.55 | 477016.75 | 389855.34 | 454814.25 | 608452.44 | 489702.19 | 434890.03 |
| O09159 | Man2b1 | Lysosomal alpha-mannosidase Mus musculus | 636.26 | 14.61 | 114648 | 10 | 10 | 29760767 | 29134727 | 24821189 | 32504589 | 31066853 | 22524719 | 22186813 | 22868383 | 23590495 | 20176759 | 24421871 | 22808655 | 21914893 | 23430521 |
| O54782 | Man2b2 | Epididymis-specific alpha-mannosidase Mus musculus | 370.99 | 8.74 | 115610 | 6 | 6 | 6902012 | 8631566 | 8239087.5 | 8540969 | 7448676 | 9207460 | 8592020 | 9111038 | 8590994 | 8105749 | 6870336 | 7346116.5 | 7554881 | 8388824 |
| Q91W89 | Man2c1 | Alpha-mannosidase 2C1 Mus musculus | 310.28 | 6.93 | 115688 | 5 | 5 | 3701658.25 | 4677847 | 3995086.5 | 4081363.25 | 4436063 | 5188776.5 | 5349075.5 | 3791053.5 | 5046583.5 | 4573996 | 4252665 | 5258346 | 4240865.5 | 4762512.5 |
| Q8K2I4 | Manba | Beta-mannosidase Mus musculus | 358.22 | 8.99 | 100831 | 6 | 6 | 9049945 | 13534204 | 10444386 | 8421119 | 12710343 | 11597144 | 9785396 | 11367436 | 11458863 | 9136331 | 10904286 | 9509131 | 11104775 | 8781094 |
| P0C5H9 | Manf | Mesencephalic astrocyte-derived neurotrophic factor Rattus norvegicus | 263.78 | 21.23 | 20388 | 5 | 5 | 13466765 | 12135204 | 10767937 | 10744876 | 10496057 | 8718691 | 8108153 | 10755421 | 7393491 | 9374199 | 11167409 | 8767509 | 10698030 | 11831972 |
| P21396 | Maoa | Amine oxidase [flavin-containing] A Rattus norvegicus | 54.51 | 3.8 | 59508 | 1 | 1 | 241152.31 | 184421.59 | 172652.61 | 1 | 1 | 193129.12 | 1 | 220678.8 | 159792.03 | 1 | 218761.09 | 1 | 1 | 191389.45 |
| Q8C052 | Map1s | Microtubule-associated protein 1S Mus musculus | 322.18 | 8.74 | 102939 | 7 | 7 | 3955355.25 | 2762875.25 | 4707181 | 4585850.5 | 5361405.5 | 2999163 | 2752173.25 | 3520419 | 3929263 | 3725320 | 5510982.5 | 3580867.5 | 5216362 | 4718750 |
| P31938 | Map2k1 | Dual specificity mitogen-activated protein kinase kinase 1 Mus musculus | 779.04 | 53.18 | 43474 | 10 | 14 | 19489445 | 34046405 | 26518319 | 17445085 | 18984441 | 26721137 | 32797363 | 34227869 | 33132369 | 34149421 | 23607019 | 24398481 | 32978897 | 17506553 |
| Q63932 | Map2k2 | Dual specificity mitogen-activated protein kinase kinase 2 Mus musculus | 544.34 | 33.92 | 44402 | 6 | 11 | 14525667 | 18130511 | 18250865 | 9743858 | 14086204 | 16234233 | 19362363 | 19185753 | 20138015 | 19338113 | 18610899 | 15415304 | 19699629 | 13266392 |
| O09110 | Map2k3 | Dual specificity mitogen-activated protein kinase kinase 3 Mus musculus | 181.02 | 14.7 | 39296 | 3 | 3 | 3462949.75 | 3640004.5 | 5152590 | 4161518.5 | 3491682 | 4946086.5 | 4042933 | 4290644 | 4419173.5 | 4041022.25 | 4660687 | 4488571 | 4375412 | 4931507.5 |
| P47809 | Map2k4 | Dual specificity mitogen-activated protein kinase kinase 4 Mus musculus | 37.09 | 4.28 | 44114 | 1 | 1 | 1 | 1 | 1 | 132612.8 | 1 | 1 | 1 | 1 | 112754.61 | 1 | 1 | 79266.92 | 100801.73 | 1 |
| Q9ESL4 | Map3k20 | Mitogen-activated protein kinase kinase kinase 20 Mus musculus | 282.33 | 7.48 | 91720 | 5 | 5 | 9457455 | 9591159 | 8883040 | 8947703 | 9302907 | 9875387 | 10254225 | 10923897 | 10057027 | 10172491 | 10556757 | 9972235 | 9422916 | 7723086 |
| P27546 | Map4 | Microtubule-associated protein 4 Mus musculus | 897.26 | 23.91 | 117429 | 19 | 19 | 24230473 | 27092593 | 23207729 | 27445529 | 28212669 | 28903119 | 30368293 | 25507457 | 30614079 | 28833887 | 30152775 | 29561549 | 28140585 | 26934361 |
| P70218 | Map4k1 | Mitogen-activated protein kinase kinase kinase kinase 1 Mus musculus | 156.07 | 5.44 | 91535 | 3 | 3 | 354441.78 | 1 | 330072.12 | 1 | 1 | 1 | 237723.8 | 1 | 278167.09 | 1 | 1 | 245084.22 | 744044.69 | 254552.06 |
| P97820 | Map4k4 | Mitogen-activated protein kinase kinase kinase kinase 4 Mus musculus | 71.94 | 2.68 | 140602 | 1 | 1 | 1 | 1 | 1 | 1 | 1 | 1 | 1 | 1 | 1 | 1 | 193544.75 | 204118.06 | 1 | 1 |
| A2AJI0 | Map7d1 | MAP7 domain-containing protein 1 Mus musculus | 112.83 | 5.79 | 93276 | 3 | 3 | 6545858 | 7890116 | 2292030.5 | 6643053.5 | 6494837.5 | 4992840.5 | 5679536.5 | 1987008 | 2984923.25 | 5908131 | 6313216.5 | 4949306 | 1967318.25 | 5872205.5 |
| P63085 | Mapk1 | Mitogen-activated protein kinase 1 Mus musculus | 341.41 | 25.7 | 41276 | 4 | 7 | 18330447 | 19120885 | 17701097 | 17579577 | 20027901 | 21189127 | 18214603 | 18930849 | 20448803 | 20604117 | 19712285 | 19053873 | 20372973 | 21148571 |
| P47811 | Mapk14 | Mitogen-activated protein kinase 14 Mus musculus | 583.22 | 39.17 | 41287 | 11 | 11 | 13121958 | 13253919 | 12043329 | 12279388 | 13826778 | 12244761 | 13301273 | 15270886 | 13534353 | 12859185 | 13115301 | 12546038 | 12012134 | 13635398 |
| Q8BH93 | Mapk1ip1l | MAPK-interacting and spindle-stabilizing protein-like Mus musculus PE | 36.11 | 10.33 | 23885 | 1 | 1 | 1 | 1 | 1522731.12 | 1 | 1938282.88 | 1 | 1427204.12 | 1427836.75 | 1522226.88 | 1 | 1501487.62 | 1481008.54 | 1400757.62 | 1540780.38 |
| P21708 | Mapk3 | Mitogen-activated protein kinase 3 Rattus norvegicus | 285.39 | 25.53 | 43081 | 3 | 6 | 14046033 | 13884697 | 13244656 | 12927679 | 14114616 | 17948245 | 14339667 | 15353954 | 16576580 | 16775449 | 16691656 | 15289569 | 16162203 | 16747251 |
| P49186 | Mapk9 | Mitogen-activated protein kinase 9 Rattus norvegicus | 83.81 | 8.04 | 48017 | 2 | 2 | 570272.56 | 933954.75 | 656750.62 | 349013.44 | 688350.19 | 755984.31 | 931781.75 | 711369.88 | 713786.25 | 674205.56 | 684356.12 | 380730.09 | 699370.5 | 690171.94 |
| P49138 | Mapkapk2 | MAP kinase-activated protein kinase 2 Mus musculus | 193.8 | 11.66 | 44050 | 5 | 5 | 5646235 | 7711157 | 6409378 | 4354192.5 | 5777333.5 | 4518503.5 | 7576411 | 3571469 | 5029995.5 | 4646068 | 5890217 | 5351306 | 8053307 | 5713878 |
| Q61166 | Mapre1 | Microtubule-associated protein RP/EB family member 1 Mus musculus | 569.78 | 41.42 | 30016 | 9 | 9 | 16125955 | 14248805 | 17313273 | 15565018 | 14982298 | 16313731 | 16824713 | 14834413 | 17621871 | 19727685 | 17574161 | 19524247 | 16654463 | 17687607 |
| Q3B8Q0 | Mapre2 | Microtubule-associated protein RP/EB family member 2 Rattus norvegicus | 65.24 | 3.68 | 36988 | 1 | 1 | 494036.25 | 466571.81 | 1 | 1 | 1 | 477079.88 | 477702.095 | 522277 | 436779.91 | 474671.59 | 479650.84 | 523658.12 | 469630.34 | 490979.7667 |
| Q922Q1 | Marc2 | Mitochondrial amidoxime reducing component 2 Mus musculus | 363.39 | 26.04 | 38194 | 3 | 8 | 7982832 | 7891377.5 | 7739494 | 9074475 | 10240457 | 10593121 | 9214571 | 9266890 | 4989632 | 11515621 | 12031591 | 9771661 | 12126803 | 10905011 |
| Q3KNM2 | March5 | E3 ubiquitin-protein ligase MARCH5 Mus musculus | 89.65 | 13.67 | 31232 | 3 | 3 | 209912.09 | 1351820.25 | 220500.03 | 316599.56 | 313799.12 | 2616609.5 | 1604976.38 | 171616.36 | 1384196.88 | 2232297.25 | 1343158.5 | 135355.39 | 1148209.5 | 2394815.25 |
| P26645 | Marcks | Myristoylated alanine-rich C-kinase substrate Mus musculus | 463.97 | 34.63 | 29661 | 7 | 7 | 12324777 | 14816291 | 15684935 | 17136421 | 13898977 | 13959959 | 14892003 | 13897375 | 14465680 | 16716566 | 18734465 | 14846687 | 16231339 | 19347283 |
| P28667 | Marcksl1 | MARCKS-related protein Mus musculus | 205.08 | 34.5 | 20165 | 4 | 4 | 9196065 | 8834829 | 8280734.5 | 8980619 | 9934519 | 8961878 | 10228555 | 9286730 | 9281941 | 10735024 | 8878136 | 9732336 | 8403939 | 9563279 |
| Q68FL6 | Mars | Methionine--tRNA ligase, cytoplasmic Mus musculus | 1471.44 | 44.79 | 101431 | 28 | 28 | 33976821 | 34399789 | 34109697 | 36394849 | 35676701 | 40249581 | 34487997 | 36927465 | 36080565 | 35942969 | 39800313 | 38507585 | 33054967 | 37249885 |
| Q3THS6 | Mat2a | S-adenosylmethionine synthase isoform type-2 Mus musculus | 436.56 | 25.06 | 43689 | 2 | 9 | 25768187 | 26025637 | 27081829 | 21670393 | 25990649 | 20955279 | 21122445 | 17982077 | 20557909 | 21761937 | 20024881 | 19028405 | 21115993 | 22779985 |
| Q5U2R0 | Mat2b | Methionine adenosyltransferase 2 subunit beta Rattus norvegicus | 217.94 | 20.66 | 37375 | 4 | 4 | 7180173.5 | 7486520 | 7835277.5 | 8525633 | 7883800.5 | 7023584.5 | 6369925 | 7857753.5 | 8033081 | 7112648 | 6855283.5 | 7679022.5 | 7461366 | 6610614 |
| Q8K310 | Matr3 | Matrin-3 Mus musculus | 768.83 | 23.4 | 94630 | 13 | 13 | 13998072 | 13357893 | 12686952 | 13257240 | 14324118 | 11271320 | 12847892 | 13491489 | 13008358 | 14596295 | 13566050 | 12297956 | 12743053 | 13305785 |
| Q9D2X5 | Mau2 | MAU2 chromatid cohesion factor homolog Mus musculus | 45.49 | 1.45 | 69579 | 1 | 1 | 741195.69 | 486170.12 | 475540.56 | 651269.19 | 563693.38 | 533056.69 | 489969.97 | 666078.62 | 469639.19 | 436321.16 | 474913.75 | 408895.25 | 427851.97 | 420013.66 |
| Q8VCF0 | Mavs | Mitochondrial antiviral-signaling protein Mus musculus | 442.01 | 21.27 | 53399 | 7 | 7 | 6068584 | 6987083 | 7016185.5 | 7826510.5 | 7408904.5 | 7451757 | 7821120 | 6250278.5 | 7432838.5 | 7678173.5 | 10640234 | 8653704 | 8332632 | 7701887 |
| Q9Z2D8 | Mbd3 | Methyl-CpG-binding domain protein 3 Mus musculus | 273.07 | 16.84 | 32168 | 5 | 5 | 4428769 | 4878586 | 4117598.5 | 4551651.5 | 4654179 | 4012158 | 4152682.25 | 4630507.5 | 4584897 | 3107936.25 | 5275999.5 | 4342071.5 | 4549249 | 4481379 |
| Q9JKP5 | Mbnl1 | Muscleblind-like protein 1 Mus musculus | 210.35 | 21.99 | 36976 | 4 | 4 | 7373277 | 7182238 | 8920765 | 10352111 | 7573455 | 6451318 | 7674755 | 5334059.5 | 6836716 | 8244195 | 7971089.5 | 6500444.5 | 7819273 | 6768189 |
| Q8CHK3 | Mboat7 | Lysophospholipid acyltransferase 7 Mus musculus | 112.25 | 7.19 | 53436 | 2 | 2 | 1874685.25 | 1911352.25 | 1763730.75 | 1283265.38 | 1490587.75 | 2701595.25 | 1557784.5 | 1926041.75 | 2547342.5 | 1887739.75 | 1500818.75 | 2459292 | 2641210.5 | 2144343 |
| Q8R3F5 | Mcat | Malonyl-CoA-acyl carrier protein transacylase, mitochondrial Mus musculus | 70.75 | 3.15 | 41928 | 1 | 1 | 263039.78 | 643665.5 | 685530.56 | 664288.5 | 560193.69 | 868637.75 | 506822.91 | 631941.7825 | 676848.44 | 475458.03 | 784668.69 | 651014.88 | 561750.81 | 563818.94 |
| Q99MR8 | Mccc1 | Methylcrotonoyl-CoA carboxylase subunit alpha, mitochondrial Mus musculus | 38.88 | 1.12 | 79344 | 1 | 1 | 1006580.12 | 1059265.62 | 795484.06 | 1220062.38 | 1020348.045 | 973099.12 | 829181.94 | 899207.75 | 894923.44 | 1005994.81 | 933480.69 | 980728.12 | 1051987.62 | 887257.75 |
| P97287 | Mcl1 | Induced myeloid leukemia cell differentiation protein Mcl-1 homolog Mus musculus | 133.72 | 10.27 | 35217 | 2 | 2 | 1736595.25 | 1185432.5 | 784035.94 | 1118408.38 | 1162852.75 | 757698 | 930485.38 | 1031097.31 | 882170.38 | 1103018.38 | 623723.69 | 896991.12 | 933646.25 | 526586.06 |
| P97310 | Mcm2 | DNA replication licensing factor MCM2 Mus musculus | 1917.27 | 46.57 | 102078 | 31 | 31 | 51710697 | 47636637 | 44773761 | 45241849 | 44152637 | 43531117 | 46710745 | 39382049 | 38292597 | 43636561 | 42618753 | 45243189 | 39042873 | 40642849 |
| P25206 | Mcm3 | DNA replication licensing factor MCM3 Mus musculus | 2118.5 | 59.11 | 91546 | 35 | 35 | 71992825 | 70460033 | 64243505 | 77191561 | 76612785 | 66122557 | 65562313 | 53989133 | 58321333 | 65732413 | 60856393 | 64366125 | 61021669 | 60529881 |
| P49717 | Mcm4 | DNA replication licensing factor MCM4 Mus musculus | 1408.25 | 42.58 | 96736 | 24 | 25 | 22723659 | 21678979 | 24885785 | 23017115 | 20432631 | 21421683 | 21578161 | 22494335 | 23419087 | 24499043 | 23742303 | 22218757 | 22743407 | 21144359 |
| P49718 | Mcm5 | DNA replication licensing factor MCM5 Mus musculus | 1617.39 | 52.8 | 82343 | 29 | 30 | 45386225 | 52188905 | 47732161 | 51570645 | 48091649 | 43327481 | 40836581 | 40987033 | 44097633 | 43083477 | 40616589 | 49738469 | 43903841 | 41701089 |
| P97311 | Mcm6 | DNA replication licensing factor MCM6 Mus musculus | 1859.34 | 44.7 | 92867 | 32 | 32 | 40374621 | 38405533 | 37149001 | 44161857 | 42873825 | 38716133 | 34479485 | 40604593 | 35473505 | 39324737 | 39354969 | 34491553 | 37369033 | 36216853 |
| Q61881 | Mcm7 | DNA replication licensing factor MCM7 Mus musculus | 1286.23 | 44.37 | 81211 | 22 | 22 | 33899465 | 33035453 | 32675285 | 34860397 | 31904891 | 27325927 | 29044059 | 30638405 | 30152625 | 26885287 | 28408079 | 27961453 | 26078545 | 25678947 |
| Q8R3C0 | Mcmbp | Mini-chromosome maintenance complex-binding protein Mus musculus | 462.89 | 16.98 | 72891 | 8 | 8 | 4546606.5 | 6107745 | 5097431.5 | 7207752.5 | 5396642 | 6312326 | 7810061.5 | 4106685.5 | 5400688 | 5310568 | 5588807 | 6803035 | 4479473.5 | 4811642 |
| Q4G009 | Mcts1 | Malignant T-cell-amplified sequence 1 Rattus norvegicus | 140.67 | 21.43 | 20550 | 2 | 2 | 1085782 | 917313.94 | 2002678.5 | 2162327.5 | 2437907.25 | 1786891 | 2279349.75 | 2157599.25 | 1947060.25 | 2105984.75 | 2000361.25 | 999067.44 | 2082598.75 | 2127111.25 |
| Q3UMR5 | Mcu | Calcium uniporter protein, mitochondrial Mus musculus | 55.29 | 2.86 | 39682 | 1 | 1 | 1657174.38 | 1464395 | 2021857.12 | 1750472.5 | 1484662.75 | 1381859.5 | 2219578.5 | 1435272.75 | 725154.44 | 2303515.5 | 1868737.62 | 1166886.25 | 956863.88 | 1560742.88 |
| Q5PSV9 | Mdc1 | Mediator of DNA damage checkpoint protein 1 Mus musculus | 144.43 | 3.4 | 184670 | 3 | 3 | 1 | 1665963.38 | 1 | 1 | 2225106.5 | 1270198 | 1825947.905 | 2482773 | 2055410.5 | 1495410.12 | 1498101.38 | 2240526.25 | 744044.69 | 2105086.25 |
| P14152 | Mdh1 | Malate dehydrogenase, cytoplasmic Mus musculus | 893.88 | 62.28 | 36511 | 15 | 15 | 66041913 | 69056273 | 74115449 | 65314393 | 67860985 | 90727257 | 84473145 | 80912601 | 105949033 | 89114865 | 80568481 | 97653977 | 91438297 | 64822617 |
| P08249 | Mdh2 | Malate dehydrogenase, mitochondrial Mus musculus | 1155.56 | 63.02 | 35611 | 17 | 18 | 362337793 | 384085889 | 367957345 | 372896545 | 337987393 | 332483617 | 409677313 | 410964737 | 323656833 | 346755617 | 347503265 | 349305825 | 374817057 | 367920097 |
| Q9D967 | Mdp1 | Magnesium-dependent phosphatase 1 Mus musculus | 152.52 | 21.95 | 18582 | 3 | 3 | 1781096.62 | 1341321.88 | 991375.06 | 1564949.12 | 1129446.88 | 1310897.5 | 1528323.12 | 2389993.25 | 2200841 | 1444996.38 | 1165839.38 | 1217744.5 | 1018688.81 | 950664.12 |
| P06801 | Me1 | NADP-dependent malic enzyme Mus musculus | 247.32 | 15.03 | 63954 | 4 | 4 | 589037.31 | 1014610 | 448303.91 | 355041.78 | 263235.97 | 945507.88 | 1103155.5 | 1091711.62 | 179480.42 | 1232759.75 | 844179.62 | 392299.59 | 1227219.75 | 679923.31 |
| Q99KE1 | Me2 | NAD-dependent malic enzyme, mitochondrial Mus musculus | 842.78 | 47.88 | 65799 | 16 | 16 | 6508553 | 10857120 | 9220604 | 8049399 | 11346127 | 10355369 | 7493749.5 | 9230377 | 9077148 | 9797728 | 10210649 | 9211162 | 10064353 | 7305768 |
| Q9DCS3 | Mecr | Enoyl-[acyl-carrier-protein] reductase, mitochondrial Mus musculus | 190.64 | 23.59 | 40343 | 3 | 3 | 200994.44 | 270153.69 | 361610.19 | 359813.31 | 298142.9075 | 201024.41 | 179565.95 | 231832.06 | 486162.03 | 1598337 | 1326938.38 | 277279.91 | 431449.31 | 1343942.75 |

| | | | | | | | | | | | | | | | | | | | | | |
|---|---|---|---|---|---|---|---|---|---|---|---|---|---|---|---|---|---|---|---|---|---|
| Q9CQ39 | Med21 | Mediator of RNA polymerase II transcription subunit 21 Mus musculus | 43.56 | 13.89 | 15588 | 1 | 1 | 146402.66 | 1 | 197672.39 | 1 | 190558.34 | 150487.6825 | 113099.83 | 234065.31 | 124017.28 | 130768.31 | 125188.16 | 164886.06 | 155843.33 | 227716.39 |
| Q4QQR9 | Memo1 | Protein MEMO1 Rattus norvegicus | 200.91 | 18.18 | 33679 | 4 | 4 | 2324965.25 | 3121353.75 | 2687123.5 | 2386020.5 | 2087837.75 | 3686239.5 | 3522288.5 | 2399831 | 2733387.5 | 2395835.5 | 2406844.25 | 3077596 | 2592135 | 2349562 |
| O88559 | Men1 | Menin Mus musculus | 33.31 | 2.29 | 67501 | 1 | 1 | 36176.1 | 378646.75 | 1 | 1 | 1 | 46373.46 | 1 | 1 | 75420.67 | 66289.84 | 39068.25 | 77214.02 | 1 | 1 |
| Q9ERE7 | Mesd | LRP chaperone MESD Mus musculus | 345.63 | 37.5 | 25207 | 6 | 6 | 5726722.5 | 7060420 | 5772009 | 6588132.5 | 5376967 | 4327514.5 | 5422149 | 5000957 | 4327137.5 | 5600284.5 | 5297414 | 4399870.5 | 4885446 | 6070852 |
| Q8BP48 | Metap1 | Methionine aminopeptidase 1 Mus musculus | 271.74 | 17.1 | 43221 | 5 | 5 | 6111351 | 6590463 | 6052901 | 6348026 | 6844733.5 | 7778479.5 | 8297604 | 6086174 | 7142611.5 | 6733561 | 7627016 | 7982255 | 7282722.5 | 7888992 |
| O08663 | Metap2 | Methionine aminopeptidase 2 Mus musculus | 500.59 | 32.01 | 52922 | 10 | 10 | 3759914.25 | 7570577.5 | 4777828 | 7124577 | 2888116.25 | 5519362 | 5320225.5 | 5111956.5 | 4646399 | 6544213.5 | 4081549.75 | 4585518 | 5289418 | 6815290 |
| Q9Z120 | Mettl1 | tRNA (guanine-N(7)-)-methyltransferase Mus musculus | 67.78 | 5.6 | 30603 | 1 | 1 | 251424.14 | 278458.44 | 217846.14 | 404747.72 | 347269.28 | 210418.48 | 240212.11 | 308422.84 | 249746 | 203438.8 | 312426.01 | 247386.44 | 318996.5 | 370895.09 |
| Q91YR5 | Mettl13 | Methyltransferase-like protein 13 Mus musculus | 177.6 | 5.73 | 78757 | 4 | 4 | 2987178 | 2296236.25 | 1540658.75 | 2430628.5 | 2493596.75 | 961580.19 | 1976723.75 | 1690095.12 | 1819286.25 | 850016 | 2046361.88 | 1215939.75 | 1435229 | 1554713.5 |
| Q9CQG2 | Mettl16 | U6 small nuclear RNA (adenine-(43)-N(6))-methyltransferase Mus musculus | 192.29 | 9.22 | 62341 | 4 | 4 | 31932109 | 2204096.5 | 2618429.75 | 24724479 | 35056745 | 2356071 | 1242981.12 | 2174420.5 | 1953284 | 1907661.38 | 40096529 | 2582518.75 | 2495090.75 | 31278831 |
| Q8BMK1 | Mettl2 | Methyltransferase-like protein 2 Mus musculus 2 | 37.87 | 5.14 | 43912 | 1 | 1 | 354146.12 | 277974.16 | 1 | 1 | 396939 | 292638.56 | 1 | 1 | 1 | 1 | 1 | 1 | 1 | 1 |
| Q9DCS2 | Mettl26 | Methyltransferase-like 26 Mus musculus | 98.55 | 8.82 | 22687 | 1 | 1 | 385797.88 | 276300.4525 | 205587.38 | 275702.25 | 238114.3 | 347897.56 | 403964.91 | 537628.25 | 494495.97 | 329711.44 | 417953.97 | 525149.44 | 383055.22 | 285989.78 |
| P21956 | Mfge8 | Lactadherin Mus musculus | 401.72 | 23.11 | 51241 | 8 | 8 | 25970557 | 26130477 | 26266839 | 20592657 | 23680475 | 20323777 | 19623881 | 19144729 | 18653009 | 19247727 | 17701069 | 13561518 | 21956337 | 18465437 |
| Q811U4 | Mfn1 | Mitofusin-1 Mus musculus | 55.91 | 1.89 | 83726 | 1 | 1 | 438864.47 | 544852.44 | 673544.94 | 489100.62 | 551190.5 | 947622.94 | 888932.44 | 495321.75 | 585021.31 | 694564.19 | 732513.62 | 898191.25 | 632499.06 | 831145.75 |
| Q02527 | Mgat3 | Beta-1,4-mannosyl-glycoprotein 4-beta-N-acetylglucosaminyltransferase Rattus norvegicus | 52.47 | 1.3 | 62022 | 1 | 1 | 7478302.5 | 7832926.5 | 4383691.5 | 6138261 | 8969417 | 5199680.5 | 3911749.5 | 4176866.5 | 2998327.5 | 4071656 | 7424262 | 7369179 | 3104444.75 | 4982473.5 |
| Q9CPU4 | Mgst3 | Microsomal glutathione S-transferase 3 Mus musculus | 221.11 | 33.99 | 16958 | 3 | 3 | 9168685 | 7486690 | 8288514 | 6290236 | 7887625 | 8243886.5 | 7819897 | 7861023 | 10824975 | 8012313.5 | 8696937 | 10469529 | 9092562 | 9698527 |
| Q8BI84 | Mia3 | Transport and Golgi organization protein 1 homolog Mus musculus | 39.75 | 0.62 | 213675 | 1 | 1 | 1082089.38 | 883635.25 | 987032.12 | 1040188.593 | 1207997.62 | 1371629.38 | 1006131.31 | 1189062.268 | 1158123.5 | 1220364.88 | 964549.19 | 877841.12 | 947882.94 | 1052335 |
| Q8R404 | Mic13 | MICOS complex subunit MIC13 Mus musculus | 246.72 | 68.91 | 13373 | 4 | 5 | 8285664 | 10155471 | 7339599 | 7039126.5 | 8109818.5 | 10048246 | 9329541 | 10007187 | 7412841.5 | 9511580 | 7597700.5 | 7443008 | 7573107 | 9573061 |
| Q8VDP3 | Mical1 | [F-actin]-monooxygenase MICAL1 Mus musculus | 145.43 | 6.11 | 116785 | 3 | 3 | 1849722 | 1880653.12 | 1756609.62 | 2122856.25 | 2245656.75 | 2033439.5 | 1968014.38 | 2253015.25 | 1841231.75 | 1849520.38 | 2021063.5 | 2195532.5 | 1834118.88 | 2159629.5 |
| Q6P6Q9 | Micu1 | Calcium uptake protein 1, mitochondrial Rattus norvegicus | 66.62 | 3.14 | 54195 | 1 | 1 | 344107.8825 | 471585.47 | 205662.03 | 347466.62 | 351717.41 | 1 | 1 | 544611.38 | 1 | 224663.64 | 462011.1767 | 560082.25 | 419544.94 | 406406.34 |
| P34884 | Mif | Macrophage migration inhibitory factor Mus musculus | 354.63 | 83.48 | 12504 | 5 | 5 | 53402745 | 43973705 | 51153873 | 36193417 | 43227489 | 57576541 | 49177593 | 64047189 | 68629393 | 53185769 | 57395533 | 59597025 | 50172601 | 49847353 |
| Q3TB92 | Milr1 | Allergin-1 Mus musculus | 71.87 | 5.69 | 27366 | 1 | 1 | 221939.36 | 275979.75 | 322224.44 | 265192 | 387710.78 | 364402.66 | 226353.06 | 316360.81 | 501373.16 | 321400.72 | 239910.09 | 316862.81 | 264206.72 | 437379.38 |
| Q76LS9 | Mindy1 | Ubiquitin carboxyl-terminal hydrolase MINDY-1 Mus musculus | 79.69 | 4.49 | 51226 | 1 | 1 | 580975.62 | 1 | 568249.25 | 719290.12 | 1 | 730955.19 | 584400.5 | 542218.94 | 600087.4225 | 542775.06 | 784075.94 | 1 | 592382.12 | 1 |
| Q9Z2L6 | Minpp1 | Multiple inositol polyphosphate phosphatase 1 Mus musculus | 92.8 | 6.44 | 54537 | 2 | 2 | 257190.27 | 1 | 1 | 1 | 1 | 1 | 167540.27 | 1 | 230146.2 | 174523.84 | 1 | 1 | 329528 | 1 |
| E9PVX6 | Mki67 | Proliferation marker protein Ki-67 Mus musculus | 3338.53 | 27.7 | 350864 | 60 | 60 | 194457617 | 64824269 | 67822953 | 70583001 | 81777945 | 53681725 | 31672173 | 36557573 | 32160395 | 52863345 | 83109921 | 47769329 | 50978801 | 75608257 |
| O08605 | Mknk1 | MAP kinase-interacting serine/threonine-protein kinase 1 Mus musculus | 48.96 | 6.32 | 47915 | 1 | 1 | 296722.34 | 372066.1075 | 349093.56 | 498455.59 | 343992.94 | 1 | 1 | 468229.38 | 395738.75 | 372338.41 | 412052.31 | 154827.98 | 744044.69 | 411099.88 |
| Q6ZQI3 | Mlec | Malectin Mus musculus | 238.37 | 16.49 | 32342 | 4 | 4 | 7046681.5 | 6176772.5 | 7601901 | 7534347.5 | 7374707 | 7877989 | 7661747.5 | 7951675.5 | 8297902 | 7359219 | 5761110.5 | 8145035 | 7630384.5 | 8133242 |
| P97679 | Mlh1 | DNA mismatch repair protein Mlh1 Rattus norvegicus | 86.38 | 5.28 | 84540 | 2 | 2 | 1 | 1 | 1 | 1 | 1 | 1 | 83354.81 | 1 | 233349.03 | 1 | 1 | 1 | 1 | 1 |
| P34960 | Mmp12 | Macrophage metalloelastase Mus musculus | 180.84 | 8.67 | 54971 | 4 | 4 | 3640142.5 | 664269.56 | 2515680.75 | 1 | 2652994.75 | 1285814.75 | 2718186 | 4539423.5 | 4647434 | 4680252 | 2571935 | 3562892.75 | 4010076.5 | 1142462.88 |
| Q9D071 | Mms19 | MMS19 nucleotide excision repair protein homolog Mus musculus | 833.62 | 19.98 | 113089 | 16 | 16 | 10553330 | 9221958 | 10853849 | 9209429 | 9126669 | 9938618 | 9285139 | 11871687 | 11040275 | 8212445 | 10088711 | 10115890 | 9658803 | 9173233 |
| P0DOV1 | Mnda | Interferon-activable protein 205-B Mus musculus | 59.12 | 2.82 | 46977 | 1 | 1 | 1493655.88 | 1912190.38 | 1427573.62 | 1756698.25 | 1660475.75 | 1618703.88 | 1436500.38 | 1239946.5 | 925458.44 | 1491002.38 | 1551222.5 | 1044461.94 | 1078398.62 | 1243619.88 |
| D0QMC3 | Mndal | Myeloid cell nuclear differentiation antigen-like protein Mus musculus | 513.11 | 22.68 | 60526 | 6 | 11 | 21679509 | 23776097 | 24524045 | 21039025 | 22592671 | 16122662 | 13932207 | 17640987 | 17385345 | 14221786 | 14060852 | 14942461 | 15294745 | 16304416 |
| Q3T1J9 | Mob1a | MOB kinase activator 1A Rattus norvegicus | 133.11 | 13.89 | 25080 | 3 | 3 | 15805977 | 9417855 | 8342740.5 | 8268693 | 9284386 | 9655179 | 8920247 | 9788105 | 8172502 | 9458717 | 8327933 | 8590804 | 8621169 | 9161568 |
| Q8VI63 | Mob2 | MOB kinase activator 2 Mus musculus | 122.93 | 10.64 | 26851 | 3 | 3 | 1377899.12 | 1714181.12 | 1378204.62 | 1764960.88 | 1905178.12 | 1743566.25 | 1472074.38 | 1570121.62 | 1460861.12 | 1350251 | 1591305.75 | 1805029.25 | 1311786 | 1764175.88 |
| A2BDX3 | Mocs3 | Adenylyltransferase and sulfurtransferase MOCS3 Mus musculus | 96.99 | 6.74 | 49375 | 3 | 3 | 688121.12 | 1 | 1 | 1 | 443013.94 | 334728.47 | 1 | 384461.16 | 1 | 230493.61 | 1 | 299145.75 | 276840.97 | 1 |
| Q80UM7 | Mogs | Mannosyl-oligosaccharide glucosidase Mus musculus | 693.4 | 23.26 | 91831 | 6 | 12 | 7922013 | 5397955 | 5427015 | 6176937.5 | 7854608 | 6081145.5 | 6348673 | 4818009.5 | 4859203.5 | 5896373.5 | 6567069 | 6193618.5 | 6180378 | 5272677 |
| Q80TL7 | Mon2 | Protein MON2 homolog Mus musculus | 269.9 | 4.66 | 189078 | 6 | 6 | 1295220.62 | 1792681.75 | 1658617.25 | 700139.25 | 525037.75 | 1898434 | 593356.75 | 1978818.25 | 1861732 | 1873055.25 | 641779.81 | 1477362.75 | 1763329.5 | 2064523.5 |
| P23249 | Mov10 | Putative helicase MOV-10 Mus musculus | 410.31 | 9.76 | 113583 | 8 | 8 | 6950960 | 6710272 | 6152657.5 | 4463334 | 5575883 | 5938043 | 4467471 | 4641793 | 4540489.5 | 4294482 | 3925563 | 7132080 | 6086712 | 6152628 |
| P63030 | Mpc1 | Mitochondrial pyruvate carrier 1 Mus musculus | 79.86 | 19.27 | 12455 | 1 | 1 | 487731.34 | 372120.62 | 303557.84 | 549422.75 | 332989.03 | 287578.06 | 326512.22 | 512950.47 | 245380.91 | 414876.53 | 359169.06 | 379913.94 | 269230.69 | 367537.41 |
| P38718 | Mpc2 | Mitochondrial pyruvate carrier 2 Rattus norvegicus | 117.89 | 28.35 | 14258 | 3 | 3 | 3960276.25 | 6464934 | 5513718 | 3025832.5 | 6765437 | 5743027.5 | 6089108 | 6111947 | 6069020 | 4695889 | 4671011.5 | 2469002 | 6123127 | 5476269.5 |
| Q9R0Q9 | Mpdu1 | Mannose-P-dolichol utilization defect 1 protein Mus musculus | 159.91 | 12.55 | 26498 | 3 | 3 | 5326098.5 | 4257626.5 | 4327888 | 3015309 | 4579706 | 5175932.5 | 4270083 | 3843708.75 | 3937211.5 | 4110939.5 | 4119772 | 4453835 | 5578305 | 4422784.5 |
| A1L314 | Mpeg1 | Macrophage-expressed gene 1 protein Mus musculus | 1083.6 | 44.04 | 78390 | 18 | 18 | 51348273 | 40907589 | 51192469 | 39580849 | 48383313 | 39353189 | 30205523 | 28185057 | 34183377 | 38079869 | 31211605 | 33129407 | 43881609 | 32021371 |
| Q810V0 | Mphosph10 | U3 small nucleolar ribonucleoprotein protein MPP10 Mus musculus | 118.73 | 10.28 | 78735 | 3 | 3 | 854143.12 | 794846.12 | 663416.75 | 879502.38 | 1050323.12 | 1028327.69 | 654926.5 | 800415.75 | 1011017.38 | 910981.06 | 829489.44 | 618254.31 | 343244.75 | 1101545.62 |
| Q9D1Q1 | Mphosph6 | M-phase phosphoprotein 6 Mus musculus | 39.11 | 6.83 | 19086 | 1 | 1 | 1656604.75 | 1630591 | 1144415.12 | 2229464 | 1933099.5 | 1603939.88 | 1561722.25 | 1597742.62 | 1329018.62 | 1463743.12 | 1742352.75 | 1819427.88 | 1714098.38 | 1516373.25 |
| P70290 | Mpp1 | 55 kDa erythrocyte membrane protein Mus musculus | 329.81 | 19.53 | 52227 | 6 | 6 | 6443307 | 4687410 | 8043545 | 4995342.5 | 4394527.5 | 5718051 | 4470090 | 7037111 | 6412846.5 | 5359899 | 5926417 | 5289640 | 6433798 | 5922609 |
| Q9JLB0 | Mpp6 | MAGUK p55 subfamily member 6 Mus musculus | 317.28 | 15.37 | 62631 | 6 | 6 | 5821362.5 | 6624546.5 | 6686558 | 4830265.5 | 5605394 | 6244530.5 | 6804290 | 7060152.5 | 6990546 | 7361588.5 | 7093408.5 | 4696316 | 7515800.5 | 6493294 |
| Q99J99 | Mpst | 3-mercaptopyruvate sulfurtransferase Mus musculus | 83.11 | 13.13 | 33097 | 2 | 2 | 1035780.94 | 1259697.62 | 1375035.5 | 965799.88 | 2324861.25 | 952104.81 | 2117898.25 | 1855888.25 | 1444309.25 | 1430997.75 | 1114146.25 | 631303.31 | 1495745.38 | 1846765.75 |
| Q61216 | Mre11 | Double-strand break repair protein MRE11 Mus musculus | 259.71 | 9.92 | 80223 | 6 | 6 | 598519.19 | 4096804.5 | 4784435 | 2311033.5 | 1564075.12 | 2995411.75 | 4810776 | 3652116.5 | 3507421.5 | 2991520.5 | 3734317.75 | 4276177 | 3534759.75 | 3344010.75 |
| Q9DAT2 | Mrgbp | MRG/MORF4L-binding protein Mus musculus | 47.29 | 6.37 | 22436 | 1 | 1 | 802670.06 | 979215.6225 | 1053447.12 | 1030878.19 | 1029867.12 | 1021464.19 | 891994.83 | 813563.88 | 771067.5 | 961883.75 | 901776 | 922245.62 | 926512.44 | 939891.38 |
| Q9CQT1 | Mri1 | Methylthioribose-1-phosphate isomerase Mus musculus | 494.63 | 38.75 | 39411 | 9 | 9 | 6321781.5 | 6129971 | 6594098 | 5600923.5 | 6837747 | 7051785.5 | 6845926.5 | 6466178 | 6744318 | 8227161.5 | 6573668 | 6833362 | 6285308 | 6144904 |
| Q99N96 | Mrpl1 | 39S ribosomal protein L1, mitochondrial Mus musculus | 45.19 | 6.25 | 37597 | 1 | 1 | 979944.81 | 1 | 659672 | 1 | 1 | 1 | 899916.81 | 1 | 692552 | 1 | 1 | 1 | 1 | 542350.25 |
| Q3TBW2 | Mrpl10 | 39S ribosomal protein L10, mitochondrial Mus musculus | 145.13 | 16.03 | 29396 | 3 | 3 | 7641554 | 8895426 | 1781394.5 | 9099606 | 2036918.5 | 7416096.5 | 1617284.5 | 1853697.25 | 1681507 | 2566281.75 | 1890753.12 | 9735469 | 2080748.25 | 1900926 |
| Q7M0E7 | Mrpl14 | 39S ribosomal protein L14, mitochondrial Rattus norvegicus | 71.54 | 6.9 | 15913 | 1 | 1 | 632822.62 | 389845.28 | 377724.06 | 928022.75 | 734948.62 | 365889.25 | 268193.97 | 197096.45 | 213796.72 | 261244.0975 | 805939.19 | 865177.75 | 744044.69 | 629384.56 |
| Q9CPR5 | Mrpl15 | 39S ribosomal protein L15, mitochondrial Mus musculus | 45.13 | 3.05 | 33542 | 1 | 1 | 894098.5 | 766796 | 1178934 | 936892.38 | 944180.22 | 964284.5 | 982733.12 | 898454.62 | 891619.81 | 1258827.62 | 1273393 | 1522701.5 | 1056277.38 | 834992.25 |
| Q9D338 | Mrpl19 | 39S ribosomal protein L19, mitochondrial Mus musculus | 107.4 | 10.62 | 33578 | 3 | 3 | 1592018.25 | 2688175.25 | 3359624.5 | 4051172.75 | 3064025 | 3016419.5 | 3331262.25 | 2801310.75 | 3317112.25 | 3423738 | 3406578 | 3364592.25 | 3140006.25 | 3148132.75 |
| Q9CQL4 | Mrpl20 | 39S ribosomal protein L20, mitochondrial Mus musculus | 44.48 | 7.38 | 17595 | 1 | 1 | 784133 | 1138434 | 711862.38 | 820748 | 988449.62 | 864064.56 | 854094 | 423959.44 | 760394 | 836374.56 | 937449 | 894493.88 | 1058552.25 | 963498.3767 |
| Q9D1N9 | Mrpl21 | 39S ribosomal protein L21, mitochondrial Mus musculus | 138.58 | 17.7 | 23366 | 3 | 3 | 1 | 1 | 717395.31 | 1 | 239972.38 | 422293.5 | 358022.34 | 446807.29 | 352345.44 | 654567.88 | 379638.78 | 654854.88 | 706971.88 | 257350.73 |
| Q9D1B9 | Mrpl28 | 39S ribosomal protein L28, mitochondrial Mus musculus | 111.93 | 18.68 | 30170 | 3 | 3 | 1 | 323581.31 | 1 | 1 | 1 | 1 | 215780.27 | 313020.06 | 111601.22 | 1 | 1 | 1 | 158743.91 | 146738.86 |
| Q6AXT0 | Mrpl37 | 39S ribosomal protein L37, mitochondrial Rattus norvegicus 2 | 106.56 | 7.09 | 48366 | 2 | 2 | 1101481.5 | 817284.75 | 1022061.88 | 1101236.25 | 898417.94 | 1077175.25 | 1102326.62 | 914765.62 | 1095964.75 | 1002188.88 | 1106689.25 | 1147189.88 | 1086400 | 719911.25 |
| Q9JKF7 | Mrpl39 | 39S ribosomal protein L39, mitochondrial Mus musculus | 79.52 | 8.04 | 38549 | 2 | 2 | 2685010.063 | 2745866 | 1424881.75 | 3128495.75 | 3440796.75 | 1645021.62 | 1640824.75 | 2664573.25 | 1352060 | 1725144.5 | 3167840.5 | 3317534.5 | 2129127.25 | 3192920 |
| Q9CY73 | Mrpl44 | 39S ribosomal protein L44, mitochondrial Mus musculus | 254.42 | 15.02 | 37527 | 3 | 3 | 1087725.88 | 1012119 | 1242485.88 | 1022207.38 | 918183.12 | 722306.62 | 1122911 | 1096844.75 | 1056445 | 815830.81 | 1039557.25 | 955534.5 | 1101811.25 | 921284.31 |
| Q9D0Q7 | Mrpl45 | 39S ribosomal protein L45, mitochondrial Mus musculus | 110.26 | 15.03 | 35411 | 3 | 3 | 2053782.88 | 1848439.12 | 2141995.25 | 1893388.25 | 2197622.5 | 1598413.75 | 2413353.75 | 1872333 | 1803440 | 2608691.25 | 2787036 | 2037119.88 | 1819551.75 | 2366615 |
| Q9EQI8 | Mrpl46 | 39S ribosomal protein L46, mitochondrial Mus musculus | 139.41 | 14.13 | 32132 | 3 | 3 | 861975.12 | 601137.44 | 545947.44 | 716169.155 | 855616.62 | 895545.5 | 781710.19 | 1 | 858697.69 | 889470.12 | 943940.19 | 941359.06 | 910827.25 | 704899.88 |
| Q9CPW3 | Mrpl54 | 39S ribosomal protein L54, mitochondrial Mus musculus | 56.12 | 13.33 | 15422 | 1 | 1 | 695990 | 584164.38 | 978873.12 | 794012.12 | 825405.12 | 707305.44 | 1 | 644357.19 | 627021.31 | 1 | 1 | 944514.44 | 1 | 651351.06 |
| Q9CPX7 | Mrps16 | 28S ribosomal protein S16, mitochondrial Mus musculus | 38.09 | 5.93 | 15192 | 1 | 1 | 1601795.12 | 1679947.62 | 1821793.75 | 2012624.5 | 1772623.25 | 1362614.88 | 1394538.12 | 1343179.88 | 1723120.25 | 1633714.75 | 1757595.25 | 1810428.62 | 1583394.12 | 1931761 |
| Q9CQE3 | Mrps17 | 28S ribosomal protein S17, mitochondrial Mus musculus | 32.7 | 7.5 | 13382 | 1 | 1 | 596581.88 | 571504.94 | 569459.94 | 384478.53 | 563171 | 440758.59 | 654718.12 | 617976.44 | 566275.56 | 504829.62 | 512854.84 | 497371 | 636747.38 | 560860.19 |
| Q924T2 | Mrps2 | 28S ribosomal protein S2, mitochondrial Mus musculus | 210.62 | 23.02 | 32313 | 4 | 4 | 3232446 | 2209466.5 | 3335009 | 2988228.25 | 3097873.75 | 3537302.75 | 2139357.75 | 3637178.75 | 3918392 | 3320751.5 | 2482769.5 | 3157006 | 3504698.25 | 2852916 |
| P58059 | Mrps21 | 28S ribosomal protein S21, mitochondrial Mus musculus | 91.71 | 14.94 | 10561 | 1 | 1 | 1 | 1 | 1 | 1 | 1 | 1 | 1 | 1 | 4652529 | 3602645.5 | 1 | 1 | 4842453.5 | 1 |
| Q9CXW2 | Mrps22 | 28S ribosomal protein S22, mitochondrial Mus musculus | 97.09 | 6.41 | 41192 | 2 | 2 | 1 | 471504.78 | 1 | 1 | 1741433.75 | 1 | 1 | 1 | 1 | 1 | 706856.38 | 1 | 1 | 564219.62 |
| Q8VE22 | Mrps23 | 28S ribosomal protein S23, mitochondrial Mus musculus | 212.44 | 24.86 | 20348 | 4 | 4 | 5079033 | 5106976.5 | 6020982 | 4458610.5 | 5111542 | 6203772 | 5486471.5 | 4509080.5 | 5123661.5 | 4634664.5 | 4873162 | 6265546 | 6807651 | 5550370.5 |
| Q9CQV5 | Mrps24 | 28S ribosomal protein S24, mitochondrial Mus musculus | 100.12 | 11.98 | 18901 | 2 | 2 | 3103318 | 3143296 | 2894432 | 2772794.25 | 3048753.25 | 2611449.5 | 2711583.5 | 1745325.75 | 3183983 | 1595479.62 | 2986874.5 | 3841231.25 | 3479869.75 | 3112544 |
| Q4QR80 | Mrps25 | 28S ribosomal protein S25, mitochondrial Rattus norvegicus 2 | 48.31 | 7.6 | 19775 | 1 | 1 | 551060.31 | 297166.72 | 379615.66 | 630213.75 | 412936.53 | 452641.28 | 352821.22 | 286898.81 | 404376.41 | 411184.53 | 379780.81 | 407755.09 | 748232 | 236289.03 |
| Q8BK72 | Mrps27 | 28S ribosomal protein S27, mitochondrial Mus musculus | 225.09 | 14.22 | 47779 | 5 | 5 | 4435851 | 4147179.25 | 6525800 | 4556549 | 3804033.25 | 4906396 | 4418495 | 6247535.5 | 5519309 | 4148132 | 5654387 | 6019560 | 6543340 | 5141177.5 |
| Q9CY16 | Mrps28 | 28S ribosomal protein S28, mitochondrial Mus musculus | 201.01 | 23.12 | 20520 | 3 | 3 | 8898625 | 3262783.75 | 3165366.75 | 2974766.75 | 2653114 | 2379017 | 2607358.75 | 6679363 | 2905079.5 | 2190415 | 2799834.75 | 3020188.75 | 2610261.75 | 2632242.5 |
| Q9D0G0 | Mrps30 | 28S ribosomal protein S30, mitochondrial Mus musculus | 214.45 | 16.52 | 49939 | 4 | 4 | 1011809.5 | 96723.46 | 1106397.88 | 1256107.75 | 1277453.75 | 2397141.5 | 1991002.5 | 833092.12 | 1413076 | 2188909.25 | 1494909.12 | 1697879 | 2480267.75 | 1823879.25 |
| Q61733 | Mrps31 | 28S ribosomal protein S31, mitochondrial Mus musculus | 148.48 | 13.54 | 43881 | 3 | 3 | 26679185 | 36153341 | 27488103 | 34474009 | 34572117 | 31242407 | 28097519 | 32179473 | 28617881 | 28799153 | 31432807 | 26541591 | 28855457 | 35423821 |
| Q9JIK9 | Mrps34 | 28S ribosomal protein S34, mitochondrial Mus musculus | 76.26 | 6.88 | 25827 | 2 | 2 | 3645397.25 | 1378870.12 | 1469048.5 | 1533717.12 | 1899317.62 | 1828549.75 | 1309544 | 1167900.12 | 1322603.25 | 1491400.88 | 1680367.62 | 1515963.25 | 1751247.88 | 2035269.38 |
| Q8BJZ4 | Mrps35 | 28S ribosomal protein S35, mitochondrial Mus musculus | 77.65 | 12.5 | 35975 | 2 | 2 | 1252758.38 | 385422.41 | 657607.44 | 394956.56 | 497457.38 | 702479.88 | 437323.09 | 841318.75 | 1037764.56 | 352768.06 | 993639.62 | 1054043.88 | 802897.5 | 842146.62 |
| P58064 | Mrps6 | 28S ribosomal protein S6, mitochondrial Mus musculus | 44.2 | 6.4 | 14309 | 1 | 1 | 925275.12 | 452644.59 | 603942.81 | 514467.69 | 467362.88 | 401379.22 | 637761.44 | 432746.16 | 480611.88 | 488124.675 | 394163.31 | 438208.44 | 546034.62 | 517231.34 |
| Q5I0K8 | Mrps7 | 28S ribosomal protein S7, mitochondrial Rattus norvegicus | 71.51 | 4.96 | 28197 | 1 | 1 | 2196894.25 | 593319.44 | 1789828.5 | 3287881 | 1966980.798 | 1786291.25 | 1439464.62 | 1229803 | 1790264.62 | 1833157.38 | 1 | 1 | 1236639.25 | 1 |
| Q9D7N3 | Mrps9 | 28S ribosomal protein S9, mitochondrial Mus musculus | 133.67 | 10.77 | 44929 | 3 | 3 | 1 | 1 | 191513.38 | 311392.88 | 357242.94 | 1 | 1 | 1 | 1 | 286915.97 | 226452.8 | 307579.44 | 244559.3 | 259530.5133 |
| Q9D0I8 | Mrto4 | mRNA turnover protein 4 homolog Mus musculus | 345.49 | 31.8 | 27546 | 8 | 8 | 10086518 | 9320718 | 10442178 | 9928135 | 9072290 | 10776985 | 9597031 | 9514083 | 8165329.5 | 9662761 | 6694744 | 9605433 | 8853762 | 8847276 |
| P43247 | Msh2 | DNA mismatch repair protein Msh2 Mus musculus | 918.75 | 24.17 | 104151 | 15 | 15 | 10034431 | 10094448 | 12280420 | 10476568 | 9570140 | 10735188 | 10617246 | 11490051 | 12580919 | 11017228 | 11687874 | 14181993 | 11256533 | 13076377 |
| P54276 | Msh6 | DNA mismatch repair protein Msh6 Mus musculus | 430.13 | 7.44 | 151084 | 9 | 9 | 9817449 | 10534568 | 16664263 | 7030455 | 16395957 | 10091629 | 13652669 | 11498538 | 14175403 | 10538319 | 9701401 | 16019451 | 10447562 | 15228954 |
| Q9CRA4 | Msmo1 | Methylsterol monooxygenase 1 Mus musculus | 141.9 | 17.75 | 34772 | 1 | 3 | 3628856.25 | 3325793.5 | 2443600.5 | 1321450.25 | 2322797.75 | 3094121.5 | 3117801 | 3987793 | 3674740.25 | 3036194.25 | 2498757.5 | 3006821.75 | 3533948.75 | 3179997.25 |
| P26041 | Msn | Moesin Mus musculus | 3037.55 | 68.11 | 67767 | 34 | 50 | 691954433 | 676819137 | 608385089 | 702829697 | 649914689 | 713954753 | 729744513 | 735206401 | 692807489 | 702614401 | 662856705 | 622155137 | 571317505 | 619211713 |
| P30204 | Msr1 | Macrophage scavenger receptor types I and II Mus musculus | 227.23 | 9.17 | 50170 | 5 | 5 | 29196489 | 30667877 | 38901625 | 33723585 | 36951481 | 25439385 | 24688635 | 23763053 | 27207317 | 26961627 | 27099437 | 29979159 | 28305943 | 32740393 |
| Q9D6Y7 | Msra | Mitochondrial peptide methionine sulfoxide reductase Mus musculus | 86.64 | 6.87 | 25988 | 1 | 1 | 1688829.12 | 1673941.5 | 1439020.12 | 1893916.5 | 1607827.38 | 1532487.75 | 1769473.88 | 1563389 | 1906581.75 | 1696728 | 1482283.75 | 1682374.75 | 1701873.62 | 1846341.75 |
| Q2YDW2 | Msto1 | Protein misato homolog 1 Mus musculus | 144.39 | 8.45 | 61230 | 3 | 3 | 717175.69 | 994928.44 | 1171781.75 | 1458062.75 | 1224273.62 | 1134969.38 | 1331154 | 299371.66 | 883597.81 | 964570 | 1115595.38 | 1169256 | 1517963.62 | 1229319.12 |
| P02802 | Mt1 | Metallothionein-1 Mus musculus | 193.08 | 68.85 | 6018 | 3 | 3 | 1 | 3184470.75 | 1 | 1963729.25 | 533792.25 | 6911727.5 | 5090203.5 | 7274615 | 8056611 | 9215157 | 1491512 | 2475808.75 | 495071.62 | 1142451.12 |

| | | | | | | | | | | | | | | | | | | | | | |
|---|---|---|---|---|---|---|---|---|---|---|---|---|---|---|---|---|---|---|---|---|---|
| P02798 | Mt2 | Metallothionein-2 Mus musculus | 54.76 | 19.67 | 6115 | 1 | 1 | 1 | 1 | 1 | 1 | 1 | 2719493.5 | 1972055.88 | 3008447.75 | 3353212.75 | 3903325.25 | 1 | 876340.12 | 1 | 1 |
| Q62599 | Mta1 | Metastasis-associated protein MTA1 Rattus norvegicus | 167.11 | 5.69 | 79412 | 1 | 3 | 6018245 | 5941622.5 | 5369196.5 | 6008476 | 7029515 | 7548297 | 5114915 | 5663994 | 5737300 | 5094412.5 | 6558016.5 | 5536009 | 5470983.5 | 5242203.5 |
| Q9R190 | Mta2 | Metastasis-associated protein MTA2 Mus musculus | 489.42 | 19.61 | 75030 | 9 | 11 | 11753179 | 11028814 | 10616998 | 12457280 | 13647856 | 13963038 | 11683730 | 12056455 | 13609668 | 13830695 | 15289526 | 12874397 | 12999347 | 10759306 |
| Q924K8 | Mta3 | Metastasis-associated protein MTA3 Mus musculus | 159.11 | 5.92 | 67077 | 0 | 3 | 8610102 | 7772625 | 7776590 | 9174893 | 9826137 | 9757543 | 7448026.5 | 7438728 | 8120222.5 | 7706254 | 9446932 | 7356513 | 7683380 | 7460241 |
| Q9CQ65 | Mtap | S-methyl-5'-thioadenosine phosphorylase Mus musculus | 475.76 | 42.05 | 31062 | 8 | 8 | 10984400 | 16676826 | 17895295 | 18858059 | 17826605 | 16842041 | 13959749 | 17314115 | 18363127 | 13884589 | 15562043 | 18470873 | 17911107 | 16814949 |
| Q791T5 | Mtch1 | Mitochondrial carrier homolog 1 Mus musculus | 140.74 | 15.42 | 41565 | 4 | 4 | 272478.12 | 1 | 1 | 1 | 1 | 468563.88 | 341604.41 | 1 | 1 | 286233.16 | 539887.38 | 519330.41 | 437482.22 | 280412.81 |
| Q791V5 | Mtch2 | Mitochondrial carrier homolog 2 Mus musculus | 539.84 | 49.17 | 33499 | 11 | 11 | 20859653 | 18690925 | 19395263 | 19571011 | 20258059 | 22602769 | 22605763 | 21323819 | 25902417 | 21592661 | 20651543 | 21899347 | 28518225 | 24182107 |
| P00405 | Mtco2 | Cytochrome c oxidase subunit 2 Mus musculus | 271.03 | 26.43 | 25976 | 5 | 5 | 66056129 | 80171209 | 69072009 | 67014545 | 75373865 | 73883257 | 70129417 | 66272065 | 67682065 | 69854697 | 41636253 | 43376381 | 51372705 | 50640613 |
| Q80WJ7 | Mtdh | Protein LYRIC Mus musculus | 728.98 | 26.08 | 63846 | 11 | 11 | 11561751 | 14092343 | 12164339 | 11791865 | 11385665 | 11422501 | 15940453 | 16759333 | 15058605 | 12527285 | 12288238 | 14431605 | 15826711 | 15495929 |
| Q8BVN4 | Mterf4 | Transcription termination factor 4, mitochondrial Mus musculus 2 | 35.3 | 2.02 | 40249 | 1 | 1 | 4459182.5 | 4166899.75 | 4839473.5 | 4991558.5 | 4571420.5 | 4913611.5 | 4242239.5 | 4348033.5 | 4798275 | 5061476 | 6260129 | 5393287 | 5279134.5 | 5790976 |
| Q5XII9 | Mtfr1l | Mitochondrial fission regulator 1-like Rattus norvegicus | 38.76 | 5.88 | 31730 | 1 | 1 | 267904.22 | 287358.22 | 237853.92 | 502640.09 | 459358.84 | 303031.97 | 463893.75 | 413161.66 | 485750.66 | 431327.03 | 470668.88 | 417930.59 | 586953.88 | 660876.38 |
| Q922D8 | Mthfd1 | C-1-tetrahydrofolate synthase, cytoplasmic Mus musculus | 979.69 | 27.38 | 101200 | 20 | 20 | 32226977 | 32587449 | 33619313 | 35887057 | 31192873 | 29830341 | 34224269 | 36322245 | 32726623 | 30977197 | 30086389 | 40286649 | 36904433 | 35868169 |
| Q3V3R1 | Mthfd1l | Monofunctional C1-tetrahydrofolate synthase, mitochondrial Mus musculus | 1183.16 | 30.3 | 105729 | 21 | 21 | 21410699 | 23157723 | 30254647 | 22551873 | 23307525 | 27432937 | 27225705 | 24544719 | 28756483 | 30676247 | 28042853 | 29152171 | 31643541 | 28773079 |
| P18155 | Mthfd2 | Bifunctional methylenetetrahydrofolate dehydrogenase/cyclohydrolase, mitochondrial Mus | 324.3 | 38.86 | 37863 | 7 | 7 | 6193325.5 | 5235505 | 6414935 | 1609984.25 | 5457327.5 | 1981267.25 | 1917171.75 | 5610306 | 5416547 | 1546046.62 | 4030647.75 | 4381240 | 8024883 | 6475610 |
| Q9D110 | Mthfs | 5-formyltetrahydrofolate cyclo-ligase Mus musculus | 79.25 | 10.34 | 23202 | 1 | 1 | 825248.19 | 586814.81 | 853625.56 | 654623.62 | 573649.5 | 801296.75 | 545301.31 | 527296.81 | 554050.06 | 451064.38 | 628122.88 | 774597.69 | 619781.62 | 341277.25 |
| Q8VEL2 | Mtmr14 | Myotubularin-related protein 14 Mus musculus | 95.82 | 4.32 | 72443 | 2 | 2 | 1152797.75 | 907174.81 | 900426.94 | 1559847 | 1803581.5 | 1660714.5 | 760952.25 | 973505.62 | 1498045.75 | 802028.69 | 659575.88 | 1011280.25 | 1480483 | 904678.44 |
| Q5PQT2 | Mtmr3 | Myotubularin-related protein 3 Rattus norvegicus | 38.67 | 1.42 | 133500 | 1 | 1 | 147283.08 | 1 | 218126.02 | 206626.69 | 1 | 1 | 1 | 160462.77 | 171092.52 | 1 | 113741.96 | 85329.12 | 744044.69 | 130690.21 |
| P03911 | Mtnd4 | NADH-ubiquinone oxidoreductase chain 4 Mus musculus | 73.08 | 6.1 | 51882 | 2 | 2 | 2704138.25 | 1376480.75 | 574861.81 | 1463478.828 | 1198434.5 | 2072572.25 | 477153.28 | 647895.31 | 1460925.25 | 1448986 | 569668.94 | 1335405 | 2380865.75 | 1869835 |
| P03921 | Mtnd5 | NADH-ubiquinone oxidoreductase chain 5 Mus musculus | 87.32 | 5.44 | 68475 | 2 | 2 | 696661.12 | 653871.69 | 411395.28 | 362957.81 | 502774.44 | 515644.75 | 321077.78 | 444791.66 | 485099.41 | 387445.28 | 508272.78 | 542400.94 | 486252.34 | 403960.75 |
| P42346 | Mtor | Serine/threonine-protein kinase mTOR Rattus norvegicus | 173.43 | 1.8 | 288794 | 3 | 3 | 373131.56 | 360144.03 | 170056.17 | 984476.19 | 700564.44 | 804879.88 | 959051 | 513076.25 | 691546.06 | 550283 | 454338.75 | 423781.62 | 691093.88 | 677628.5 |
| P62774 | Mtpn | Myotrophin Mus musculus | 212.54 | 43.22 | 12861 | 3 | 3 | 12414931 | 9254644 | 8558970 | 11747155 | 10568561 | 12925697 | 15846595 | 12813013 | 16022103 | 16317372 | 13558337 | 12699394 | 14288535 | 10384550 |
| Q9CZU3 | Mtrex | Exosome RNA helicase MTR4 Mus musculus | 704.57 | 17.5 | 117636 | 13 | 13 | 11266927 | 13654202 | 12180767 | 12814407 | 12901595 | 12712269 | 11727219 | 13273297 | 13466939 | 11898094 | 13163480 | 12460293 | 13459230 | 14043662 |
| Q8R1S4 | Mtss1 | Metastasis suppressor protein 1 Mus musculus | 496.42 | 18.45 | 82408 | 10 | 10 | 9374076 | 7320747 | 12592615 | 9704686 | 9922731 | 12159103 | 12007060 | 7731531 | 12336889 | 11379501 | 8372231 | 4972459.5 | 8208258.5 | 6426825.5 |
| P47802 | Mtx1 | Metaxin-1 Mus musculus | 233.22 | 16.72 | 35624 | 5 | 5 | 2503474.25 | 1896446.12 | 1906389 | 5213587 | 1850215.25 | 5190706 | 2063815.75 | 1625731 | 2076030.75 | 1761744.75 | 1996997 | 1948960.12 | 2620966.5 | 4590042 |
| O88441 | Mtx2 | Metaxin-2 Mus musculus | 363.42 | 51.71 | 29758 | 6 | 6 | 11953773 | 15446370 | 16026865 | 13889297 | 12873955 | 11433947 | 15876069 | 13474718 | 17106409 | 13141901 | 14563795 | 5969074.5 | 14726044 | 16919611 |
| P16332 | Mut | Methylmalonyl-CoA mutase, mitochondrial Mus musculus | 326.95 | 15.11 | 82844 | 7 | 7 | 7811386 | 8878732 | 7843282 | 8798362 | 8054289.5 | 6970764 | 9825029 | 9681101 | 8973153 | 8838839 | 9963017 | 7088723 | 10404638 | 10376828 |
| Q99JF5 | Mvd | Diphosphomevalonate decarboxylase Mus musculus | 633.68 | 33.42 | 44072 | 11 | 11 | 4716165 | 2736008 | 3696750 | 3064832.5 | 2196102 | 3036713.5 | 3998442.75 | 3587044 | 3034370.75 | 3641597.5 | 2539695.75 | 3581664.5 | 3088744.5 | 3323584 |
| Q9R008 | Mvk | Mevalonate kinase Mus musculus | 98.03 | 7.85 | 41877 | 2 | 2 | 1 | 1 | 159690.31 | 145431.55 | 1 | 1 | 1 | 1 | 1 | 1 | 1 | 1 | 1 | 1 |
| Q9EQK5 | Mvp | Major vault protein Mus musculus | 1618.74 | 49.13 | 95924 | 26 | 26 | 34631369 | 49552661 | 51757525 | 35835677 | 33339521 | 47339501 | 51812405 | 54684693 | 51129813 | 52038389 | 40128417 | 49560329 | 51389161 | 42000249 |
| O35682 | Myadm | Myeloid-associated differentiation marker Mus musculus | 142.12 | 16.25 | 35285 | 3 | 3 | 1 | 400232.06 | 1 | 628709.5 | 532174.56 | 446582.5 | 806921.25 | 678009.9375 | 695979.19 | 762556.81 | 542196.5 | 858932.81 | 981764.5 | 794297.9367 |
| Q7TPV4 | Mybbp1a | Myb-binding protein 1A Mus musculus | 1507.34 | 28.5 | 152037 | 21 | 27 | 54774545 | 29585349 | 29353129 | 34995205 | 34132297 | 37312985 | 34503717 | 28521193 | 28869303 | 37978989 | 34237637 | 40157961 | 43888041 | 32823037 |
| P22366 | Myd88 | Myeloid differentiation primary response protein MyD88 Mus musculus | 127.41 | 15.54 | 33753 | 3 | 3 | 241400.28 | 381308.16 | 334468.31 | 323696.16 | 173338.16 | 700743.31 | 915457.94 | 884638.69 | 715355 | 229963.56 | 501141.38 | 318508.59 | 305476.47 | 615166.62 |
| Q9CPT4 | Mydgf | Myeloid-derived growth factor Mus musculus | 164.89 | 21.69 | 17982 | 4 | 4 | 11454445 | 12826983 | 12146643 | 12124269 | 12093198 | 11743287 | 11204550 | 13121672 | 11606630 | 9643201 | 12501403 | 12580669 | 11336945 | 12577573 |
| Q8C854 | Myef2 | Myelin expression factor 2 Mus musculus | 368.43 | 17.26 | 63295 | 8 | 8 | 12451083 | 10399837 | 13744827 | 13478114 | 12110127 | 12444582 | 12824973 | 11806477 | 10349247 | 18146287 | 12877945 | 16170170 | 16891291 | 17410257 |
| Q9JK81 | Myg1 | UPF0160 protein MYG1, mitochondrial Mus musculus | 541.26 | 36.32 | 42723 | 11 | 11 | 17665041 | 19092641 | 16112001 | 19558717 | 18558771 | 14437871 | 15265776 | 15087716 | 12738415 | 14976250 | 13547084 | 14803579 | 13206613 | 14359070 |
| Q61879 | Myh10 | Myosin-10 Mus musculus | 1341.1 | 16.3 | 228996 | 8 | 24 | 74988881 | 60605669 | 69193473 | 77776465 | 65900389 | 71497145 | 81761073 | 67735193 | 78119785 | 83971361 | 70536937 | 67848313 | 69180041 | 66655853 |
| O08638 | Myh11 | Myosin-11 Mus musculus | 687.13 | 5.93 | 227028 | 0 | 12 | 73974265 | 58429205 | 65075965 | 70564665 | 50668577 | 70975865 | 71461009 | 59483565 | 65172541 | 75858185 | 73651025 | 64093765 | 63286953 | 63066673 |
| Q6URW6 | Myh14 | Myosin-14 Mus musculus | 411.8 | 4.15 | 228586 | 1 | 7 | 43131049 | 32586251 | 38244089 | 42434093 | 36154369 | 39343689 | 40920749 | 34526369 | 38341345 | 43738401 | 39208385 | 36900157 | 35464141 | 35587113 |
| Q8VDD5 | Myh9 | Myosin-9 Mus musculus | 7949.92 | 59.54 | 226372 | 96 | 119 | 543778561 | 529906881 | 549097857 | 587732737 | 528381601 | 607297857 | 684445825 | 542283905 | 611592449 | 641490177 | 587579649 | 552672897 | 552689217 | 548586369 |
| Q3THE2 | Myl12b | Myosin regulatory light chain 12B Mus musculus | 507.79 | 57.56 | 19779 | 1 | 8 | 51397537 | 54624149 | 54150541 | 53190337 | 52433781 | 61876445 | 51802825 | 69420817 | 66393913 | 60025369 | 56725825 | 56143129 | 55255449 | 51995821 |
| P16409 | Myl3 | Myosin light chain 3 Rattus norvegicus | 152.53 | 12 | 22156 | 1 | 2 | 49731337 | 46274817 | 55362829 | 41102137 | 40091953 | 55581985 | 48993581 | 68914905 | 62806777 | 44069197 | 52805641 | 54799937 | 50844953 | 55646165 |
| Q60605 | Myl6 | Myosin light polypeptide 6 Mus musculus | 692.56 | 66.23 | 16930 | 9 | 10 | 247381857 | 287915105 | 259841569 | 272482433 | 250616641 | 267724305 | 282016129 | 323066497 | 277912513 | 268017137 | 270163425 | 247204193 | 241652033 | 273634113 |
| Q9CQ19 | Myl9 | Myosin regulatory light polypeptide 9 Mus musculus | 357.35 | 45.35 | 19854 | 1 | 6 | 26152073 | 28341347 | 28584151 | 26103673 | 24852313 | 31823803 | 27647341 | 35699409 | 33782497 | 32548851 | 29581851 | 29213619 | 30036973 | 26568097 |
| Q9JMH9 | Myo18a | Unconventional myosin-XVIIIa Mus musculus | 490.17 | 6.59 | 232755 | 9 | 9 | 4793741 | 6160377.5 | 3045286.25 | 4735160.5 | 3423998.25 | 4536288 | 4882860 | 3741452.25 | 4143828.5 | 3342137.5 | 6130950.5 | 4490298 | 3017165.25 | 6234149 |
| Q9WTI7 | Myo1c | Unconventional myosin-Ic Mus musculus | 1457.81 | 27.66 | 121944 | 23 | 24 | 35383717 | 19853017 | 14692594 | 18217151 | 19301913 | 17891621 | 19431027 | 21476851 | 17922225 | 16512819 | 17519481 | 17371251 | 17790307 | 15160151 |
| E9Q634 | Myo1e | Unconventional myosin-Ie Mus musculus | 1357.31 | 32.25 | 126818 | 21 | 26 | 33731409 | 33920177 | 35435257 | 31804043 | 30204529 | 32727155 | 36163601 | 31974301 | 35200169 | 35641233 | 33410051 | 37107581 | 36431445 | 33909517 |
| P70248 | Myo1f | Unconventional myosin-If Mus musculus | 612.94 | 12.47 | 125947 | 5 | 10 | 17446385 | 18365493 | 20330939 | 17379969 | 17404041 | 18999235 | 19819955 | 20115285 | 21588253 | 18433567 | 18101615 | 19441709 | 19036053 | 18776571 |
| Q5SUA5 | Myo1g | Unconventional myosin-Ig Mus musculus | 345.99 | 9.47 | 117227 | 7 | 7 | 5781062.5 | 4118383 | 3759806.75 | 2363663.5 | 3330902.5 | 3728698.75 | 3080312 | 3554295.5 | 3893215.75 | 2330355.75 | 3298749.25 | 2592337.5 | 3515610.25 | 3092089.5 |
| Q99104 | Myo5a | Unconventional myosin-Va Mus musculus | 229.53 | 3.72 | 215538 | 5 | 5 | 1932483.25 | 1126682.25 | 1334531.75 | 1824382.25 | 737221.5 | 1425989.25 | 1384577.88 | 1543087.75 | 1583081.25 | 1395116.5 | 2415091.75 | 1492823 | 2066559.25 | 1926800.12 |
| Q63358 | Myo9b | Unconventional myosin-IXb Rattus norvegicus | 384.83 | 5.35 | 225037 | 7 | 7 | 1958592.38 | 2189105.75 | 2147990.75 | 1461775.75 | 2002667.5 | 2262807 | 2255004.75 | 2709591.25 | 2473535 | 2830665.5 | 1658130.25 | 2318432.75 | 2192890.75 | 2105076 |
| Q69ZN7 | Myof | Myoferlin Mus musculus | 889.94 | 12.65 | 233324 | 18 | 19 | 19943977 | 22640817 | 24982759 | 14619662 | 16671501 | 30093631 | 17543915 | 18947945 | 19762373 | 20447297 | 20859325 | 20925043 | 31446979 | 21346037 |
| Q80UM3 | Naa15 | N-alpha-acetyltransferase 15, NatA auxiliary subunit Mus musculus | 805.53 | 21.62 | 100961 | 17 | 17 | 26797527 | 30386665 | 28246201 | 25291909 | 28832999 | 31215095 | 27049415 | 31753255 | 28953379 | 26479989 | 25438347 | 27112099 | 28368061 | 29198697 |
| Q8BWZ3 | Naa25 | N-alpha-acetyltransferase 25, NatB auxiliary subunit Mus musculus | 298.5 | 10.29 | 111708 | 7 | 7 | 7524617.5 | 7141157 | 8268304.5 | 6840852 | 6667260 | 7016593.5 | 6525719 | 7762595 | 7902196 | 6272259 | 6207010.5 | 6804357.5 | 7686207.5 | 6551238.5 |
| Q8VE10 | Naa40 | N-alpha-acetyltransferase 40 Mus musculus | 76.46 | 12.24 | 27229 | 2 | 2 | 1771650.12 | 1859755.5 | 1967480.38 | 1627302.75 | 1980099.25 | 1881331.75 | 1933601.25 | 1780618.968 | 1847746.12 | 1459796.75 | 2024254.75 | 2099732 | 1847657.25 | 2038390.5 |
| Q6PGB6 | Naa50 | N-alpha-acetyltransferase 50 Mus musculus | 413.5 | 46.15 | 19414 | 9 | 9 | 8352918.5 | 7574797.5 | 10420285 | 6676337 | 6330412 | 8543519 | 6694141 | 9475609 | 9763754 | 8272671.5 | 8313723.5 | 8816664 | 8926705 | 7740886 |
| Q9D7V9 | Naaa | N-acylethanolamine-hydrolyzing acid amidase Mus musculus | 120.6 | 6.63 | 40075 | 2 | 2 | 1150586.12 | 1279250.5 | 1134581.25 | 614297.88 | 1006296.25 | 1048185.69 | 918550 | 1120085.38 | 1088881.5 | 875974.5 | 1530869.25 | 1131337.62 | 996751.44 | 1044423.06 |
| Q3SWT1 | Nabp2 | SOSS complex subunit B1 Rattus norvegicus | 155.87 | 17.06 | 22377 | 2 | 2 | 590464.69 | 490507.22 | 419391.28 | 516070.09 | 181658.59 | 376977.47 | 410855.03 | 355731.28 | 543210.75 | 380162.12 | 1 | 1 | 568753.5 | 398037.56 |
| P70670 | Naca | Nascent polypeptide-associated complex subunit alpha, muscle-specific form Mus muscul | 450.4 | 3.2 | 220499 | 6 | 6 | 16618589 | 2569047.5 | 20120981 | 21405269 | 23384415 | 25656853 | 21488095 | 17408961 | 22986857 | 26415465 | 21659267 | 24710441 | 27801965 | 23619005 |
| P58058 | Nadk | NAD kinase Mus musculus | 117.21 | 7.29 | 48597 | 2 | 2 | 1 | 528761.81 | 1 | 859106.5 | 634041.38 | 676636.81 | 681017.78 | 717243.62 | 663544.75 | 666645.94 | 748767.5 | 2229787.5 | 619642.69 | 1584447.5 |
| Q8VBW6 | Nae1 | NEDD8-activating enzyme E1 regulatory subunit Mus musculus | 551.95 | 32.21 | 60274 | 10 | 10 | 1727273 | 6163646 | 5096017.5 | 7515592.5 | 7532802 | 5603706 | 7480095.5 | 7076524 | 8453739 | 6120686.5 | 7313509.5 | 7267900.5 | 7600516 | 6995704.5 |
| Q9QWR8 | Naga | Alpha-N-acetylgalactosaminidase Mus musculus | 613.16 | 36.63 | 47235 | 12 | 12 | 29794001 | 21989381 | 21418603 | 26626067 | 23937645 | 23639115 | 29644369 | 27823455 | 29453821 | 23591503 | 27322447 | 27173035 | 24618809 | 23046301 |
| Q9QZ08 | Nagk | N-acetyl-D-glucosamine kinase Mus musculus | 217.37 | 14.29 | 37268 | 5 | 5 | 8062789 | 6713548 | 6884579 | 6825881.5 | 7194295 | 6607882 | 4962628.5 | 6840466 | 6593326 | 5349132 | 6609201.5 | 5562475 | 5993183.5 | 6841478.5 |
| Q8BJ48 | Nagpa | N-acetylglucosamine-1-phosphodiester alpha-N-acetylglucosaminidase Mus musculus | 98.94 | 3.09 | 56044 | 1 | 1 | 2048881.38 | 2216414.75 | 2235319.5 | 1364360.88 | 1683257.25 | 1555792.5 | 1747755.75 | 1957345.88 | 1763128.5 | 1536927.25 | 1600093.88 | 1925760.5 | 1760216.5 | 1677480.38 |
| Q9JIB3 | Naip7 | Baculoviral IAP repeat-containing protein 1g Mus musculus | 48.44 | 1 | 159664 | 1 | 1 | 238930.83 | 261196.81 | 289199.44 | 216950.89 | 175438.23 | 299522.84 | 326030.94 | 331597.91 | 324241.88 | 194549.48 | 313725.44 | 400236.69 | 276076.38 | 242278.19 |
| Q99KQ4 | Nampt | Nicotinamide phosphoribosyltransferase Mus musculus | 731.17 | 32.38 | 55447 | 13 | 13 | 30820605 | 24699147 | 27517631 | 22974717 | 25876253 | 27646577 | 29177039 | 31736719 | 30398801 | 28983257 | 24044679 | 22803953 | 25327249 | 24938137 |
| Q5M969 | Nanp | N-acylneuraminate-9-phosphatase Rattus norvegicus | 83.18 | 7.26 | 27741 | 1 | 1 | 564400.44 | 1 | 1 | 1 | 388258.62 | 1 | 380659.16 | 1 | 323253.69 | 266489.47 | 1 | 366503.34 | 1 | 1 |
| Q99J77 | Nans | Sialic acid synthase Mus musculus | 469.57 | 40.11 | 40024 | 9 | 9 | 13509025 | 10797360 | 11397695 | 12177869 | 13164238 | 14493721 | 14600310 | 11978542 | 13064987 | 13644092 | 13047870 | 11914465 | 12438173 | 12922142 |
| P28656 | Nap1l1 | Nucleosome assembly protein 1-like 1 Mus musculus | 477.44 | 29.67 | 45345 | 7 | 8 | 19945157 | 16459026 | 16837247 | 16041722 | 14483890 | 19846067 | 15787098 | 20845735 | 19376947 | 17484515 | 17225001 | 18927387 | 20828933 | 16247528 |
| Q78ZA7 | Nap1l4 | Nucleosome assembly protein 1-like 4 Mus musculus | 735.31 | 49.6 | 42679 | 11 | 12 | 20299347 | 22217325 | 19984163 | 19435575 | 21151159 | 19828931 | 21652323 | 22412957 | 21635135 | 21323851 | 21524375 | 19930205 | 19934905 | 19765273 |
| Q5PPG6 | Nap1l5 | Nucleosome assembly protein 1-like 5 Rattus norvegicus 2 | 35.42 | 4.52 | 16858 | 1 | 1 | 2030543 | 1892679.25 | 1967458.75 | 2355257 | 1713276.25 | 1402256.38 | 1443987.75 | 1327458.88 | 1358701.62 | 1509884.25 | 1741801.62 | 1917897.38 | 1365048.5 | 1856196.75 |
| P54921 | Napa | Alpha-soluble NSF attachment protein Rattus norvegicus | 501.16 | 38.31 | 33193 | 9 | 9 | 20515501 | 19961555 | 22465731 | 19491621 | 22112343 | 21078087 | 23011023 | 23755503 | 23145177 | 20406511 | 22582059 | 23699813 | 26875043 | 22398461 |
| Q9CWZ7 | Napg | Gamma-soluble NSF attachment protein Mus musculus | 301.23 | 20.83 | 34732 | 6 | 6 | 14751396 | 14937638 | 14208314 | 11512289 | 12807202 | 11623381 | 13809689 | 11931965 | 12823853 | 13523354 | 14235778 | 11832348 | 13963039 | 14287076 |
| O09043 | Napsa | Napsin-A Mus musculus | 217.39 | 18.62 | 45544 | 4 | 4 | 8752844 | 10914111 | 8979887 | 9367902 | 8667265 | 9321483 | 10836249 | 10200148 | 9967983 | 8936050 | 6740373 | 7521599.5 | 7877479 | 8742607 |
| Q8BP47 | Nars | Asparagine--tRNA ligase, cytoplasmic Mus musculus | 1235.19 | 42.04 | 64279 | 19 | 19 | 57360417 | 58800577 | 63115161 | 62486093 | 61635505 | 68363913 | 64754697 | 66663197 | 69494041 | 69905617 | 59691693 | 60229793 | 63878785 | 63270693 |
| Q99MD9 | Nasp | Nuclear autoantigenic sperm protein Mus musculus | 853.85 | 28.85 | 83954 | 12 | 12 | 17193899 | 22030449 | 23544573 | 18471017 | 22064153 | 21591093 | 21176717 | 22444649 | 22116225 | 21115937 | 19277963 | 20790531 | 19755639 | 19555725 |
| Q8K224 | Nat10 | RNA cytidine acetyltransferase Mus musculus | 588.29 | 16.41 | 115419 | 11 | 11 | 3630593.75 | 2518550 | 3124602.75 | 3726393 | 2927654 | 2950718.5 | 1862551.25 | 3499876 | 2370055.5 | 3929132.5 | 3015760.5 | 3049406 | 4466577 | 4748532 |
| Q9CZ42 | Naxd | ATP-dependent (S)-NAD(P)H-hydrate dehydratase Mus musculus | 95.41 | 8.45 | 36717 | 2 | 2 | 1031333.31 | 1133381.38 | 1491334.25 | 1585236.38 | 1723380.75 | 1424460 | 964699.75 | 1282024 | 1308703.88 | 1415371.62 | 1425985.38 | 1424945.12 | 1363195.38 | 1798684.62 |
| Q9EPN1 | Nbea | Neurobeachin Mus musculus | 40.57 | 0.41 | 326743 | 1 | 1 | 1 | 403126.84 | 412973.84 | 1 | 154410.02 | 198608.97 | 375468.8525 | 481021.16 | 466297.84 | 355947.44 | 1 | 178570.47 | 1 | 1 |
| Q6ZQA0 | Nbeal2 | Neurobeachin-like protein 2 Mus musculus | 204.87 | 2.59 | 301924 | 5 | 5 | 552293.81 | 632431.62 | 866172.62 | 536646.12 | 296409.5 | 343855.25 | 844915.31 | 820257 | 1244882.25 | 679602.56 | 735676 | 583106.19 | 929578.38 | 749453.5233 |
| Q8K2Z4 | Ncapd2 | Condensin complex subunit 1 Mus musculus | 831.87 | 17.17 | 155665 | 17 | 17 | 16534728 | 14165364 | 16248981 | 12173057 | 12275724 | 12984456 | 13105599 | 13799287 | 13502280 | 12321882 | 13862331 | 15292649 | 15783131 | 14966242 |
| Q8C156 | Ncaph | Condensin complex subunit 2 Mus musculus | 335.07 | 11.22 | 82303 | 5 | 5 | 1761884.62 | 2871634 | 1722349.75 | 1184029.75 | 2470834.25 | 1740894.12 | 1778790.75 | 1672322 | 1753001.88 | 1794876.75 | 1835524.38 | 1805700.62 | 1722564.75 | 1994131.75 |
| Q3UYV9 | Ncbp1 | Nuclear cap-binding protein subunit 1 Mus musculus | 480.77 | 13.04 | 91927 | 7 | 7 | 5706985 | 4680056 | 3885938.75 | 3584168 | 3357126 | 5004967 | 3873667 | 5277315.5 | 4581183 | 4171382.5 | 4913906.5 | 4810874 | 4367267.5 | 4304982 |
| Q8BLF1 | Nceh1 | Neutral cholesterol ester hydrolase 1 Mus musculus | 759.72 | 42.16 | 45740 | 11 | 11 | 39716793 | 42137821 | 51028613 | 47627873 | 41299869 | 49714409 | 46364413 | 44404141 | 47719149 | 46509241 | 42611473 | 49256741 | 52646477 | 45952801 |
| Q09014 | Ncf1 | Neutrophil cytosol factor 1 Mus musculus | 735.9 | 43.33 | 44667 | 13 | 13 | 33010849 | 37826905 | 34606425 | 33224639 | 42468509 | 40737505 | 35117361 | 34619181 | 38552889 | 39637401 | 36853581 | 38223617 | 38343801 | 43484697 |
| O70145 | Ncf2 | Neutrophil cytosol factor 2 Mus musculus | 1100.57 | 50.48 | 59485 | 21 | 21 | 69790921 | 61107409 | 54740737 | 56351425 | 58903945 | 53135781 | 53183137 | 57343985 | 49652681 | 52358545 | 52031457 | 46979217 | 49328633 | 51797241 |
| P97369 | Ncf4 | Neutrophil cytosol factor 4 Mus musculus | 1302.8 | 59.88 | 38707 | 19 | 19 | 81977657 | 88707193 | 93146001 | 82076441 | 86537425 | 83068145 | 85735233 | 99390209 | 89060425 | 78406737 | 83681273 | 78695921 | 80817953 | 89944305 |
| Q8K1X4 | Nckap1l | Nck-associated protein 1-like Mus musculus | 1623.37 | 32.72 | 128905 | 25 | 26 | 41108977 | 46357361 | 44054669 | 39256133 | 43858401 | 47610313 | 48682701 | 54800649 | 51818541 | 45465341 | 44643573 | 45631721 | 46334749 | 47776105 |
| P09405 | Ncl | Nucleolin Mus musculus | 2657.22 | 41.58 | 76723 | 41 | 43 | 512448801 | 493881793 | 545296833 | 505632737 | 517477665 | 546814785 | 557909633 | 523227073 | 571253825 | 519261889 | 481219489 | 532755297 | 556250113 | 481294433 |
| Q5XIA1 | Ncln | Nicalin Rattus norvegicus | 196.73 | 7.28 | 62992 | 4 | 4 | 5398326.5 | 5630011.5 | 5604782 | 4915900.5 | 7233276.5 | 5395313 | 5894605.5 | 4233968 | 4957850 | 4875217.5 | 4694031 | 3742528.5 | 4228253.5 | 4378723 |

| | | | | | | | | | | | | | | | | | | | | | |
|---|---|---|---|---|---|---|---|---|---|---|---|---|---|---|---|---|---|---|---|---|---|
| P57716 | Ncstn | Nicastrin Mus musculus | 874.54 | 25.99 | 78492 | 13 | 13 | 66070841 | 61829253 | 59012321 | 56884313 | 61278481 | 63410585 | 57526361 | 59194677 | 50824225 | 51005569 | 49550433 | 53796573 | 55036769 | 55373305 |
| Q8VCB1 | Ndc1 | Nucleoporin NDC1 Mus musculus | 108.12 | 6.24 | 75409 | 2 | 2 | 1 | 569035.88 | 1 | 1142505 | 474074.88 | 1 | 828521.94 | 1 | 575830.69 | 685407.06 | 699124.62 | 1075651.207 | 1659112.5 | 868716.5 |
| Q78PB6 | Ndel1 | Nuclear distribution protein nudE-like 1 Rattus norvegicus | 70.68 | 3.48 | 38366 | 1 | 1 | 413963 | 358324.19 | 535885.44 | 692874.75 | 438339.22 | 368744.41 | 424354.22 | 322893.31 | 449210.19 | 424886.94 | 493590.69 | 765726.56 | 608368.62 | 569718.38 |
| Q62433 | Ndrg1 | Protein NDRG1 Mus musculus | 402.01 | 32.74 | 43009 | 7 | 7 | 12050821 | 17884593 | 12918770 | 12185809 | 17734777 | 23029545 | 23079657 | 26153407 | 27206085 | 28481307 | 19393255 | 15887399 | 16453997 | 18457141 |
| Q6AYR2 | Ndrg3 | Protein NDRG3 Rattus norvegicus | 61.03 | 5.07 | 41507 | 1 | 1 | 1 | 1 | 1 | 1 | 1 | 468500.78 | 1 | 403068 | 1 | 1 | 1 | 538201.38 | 1 | 481301.97 |
| Q99LC3 | Ndufa10 | NADH dehydrogenase [ubiquinone] 1 alpha subcomplex subunit 10, mitochondrial Mus mu | 384.06 | 29.01 | 40603 | 6 | 6 | 7593042.5 | 6444211 | 6788890 | 5793231.5 | 6718288 | 5982342 | 6371891 | 6730492 | 6922600 | 6341350 | 6645300 | 6858966 | 7139362.5 | 7201826.5 |
| Q9D8B4 | Ndufa11 | NADH dehydrogenase [ubiquinone] 1 alpha subcomplex subunit 11 Mus musculus | 59.19 | 7.8 | 14982 | 1 | 1 | 1637960.25 | 1386063.25 | 1465539.38 | 1019381.25 | 1312374.5 | 1410113.5 | 1312291.5 | 1390527.75 | 1738220.75 | 1182789 | 1654438.88 | 1664576.38 | 1582344.12 | 1498887.5 |
| Q7TMF3 | Ndufa12 | NADH dehydrogenase [ubiquinone] 1 alpha subcomplex subunit 12 Mus musculus | 139.85 | 17.93 | 17086 | 3 | 3 | 1921479.5 | 1833003.38 | 851830.31 | 1647268.38 | 1864334.75 | 1773579.25 | 1959347.62 | 1814663 | 1585689.62 | 1683590 | 1262927.75 | 1833383.75 | 2165629.5 | 1155377.75 |
| Q9ERS2 | Ndufa13 | NADH dehydrogenase [ubiquinone] 1 alpha subcomplex subunit 13 Mus musculus | 88.83 | 16.67 | 16860 | 2 | 2 | 4614131 | 4919397 | 3511846 | 3350242.75 | 3415066.5 | 4450796 | 4098168.5 | 4945254 | 4415811 | 3916618.75 | 3842576 | 2923785 | 4786252 | 4700986 |
| Q9CQ75 | Ndufa2 | NADH dehydrogenase [ubiquinone] 1 alpha subcomplex subunit 2 Mus musculus | 84.03 | 14.14 | 10916 | 1 | 1 | 2463771.5 | 3071521.25 | 2938854.5 | 3102648.5 | 3003064.75 | 2054305.12 | 3184983.5 | 2638694.5 | 3057146 | 2824422.75 | 2644195.25 | 2854947 | 2398434.5 | 2336564.25 |
| Q62425 | Ndufa4 | Cytochrome c oxidase subunit NDUFA4 Mus musculus | 180.38 | 68.29 | 9327 | 4 | 4 | 24869155 | 20107309 | 17811985 | 16011653 | 17451605 | 13934563 | 13859357 | 14726675 | 15424429 | 12746563 | 11183773 | 6177599 | 10366876 | 11151371 |
| Q9CPP6 | Ndufa5 | NADH dehydrogenase [ubiquinone] 1 alpha subcomplex subunit 5 Mus musculus | 64.98 | 17.24 | 13360 | 1 | 1 | 703088.19 | 303784.12 | 667029.88 | 1 | 1 | 605270.44 | 411185.47 | 1064695.62 | 745168.69 | 296783.12 | 650613.8767 | 765023.94 | 740815.31 | 446002.38 |
| Q9DCJ5 | Ndufa8 | NADH dehydrogenase [ubiquinone] 1 alpha subcomplex subunit 8 Mus musculus | 115.78 | 24.42 | 19992 | 3 | 3 | 2939968 | 2200855.5 | 2275720 | 2544502.25 | 2041394.75 | 2166904.75 | 2187849.25 | 2112275.75 | 2130579 | 2197107.25 | 2203241.5 | 2843536.5 | 2296145.25 | 1826972.88 |
| Q9DC69 | Ndufa9 | NADH dehydrogenase [ubiquinone] 1 alpha subcomplex subunit 9, mitochondrial Mus mus | 551 | 36.87 | 42525 | 9 | 9 | 17553387 | 7885362 | 8357411 | 12999701 | 12867800 | 13081089 | 12728057 | 9022749 | 9330587 | 9106890 | 14055119 | 9240465 | 13321976 | 9863841 |
| Q9DCS9 | Ndufb10 | NADH dehydrogenase [ubiquinone] 1 beta subcomplex subunit 10 Mus musculus | 237.4 | 31.25 | 21024 | 4 | 4 | 3483242.5 | 3497644.5 | 3301506 | 3235374 | 4771064 | 712077.81 | 3238075.75 | 2344082 | 2693239.75 | 3887845.5 | 5157265.5 | 1943758 | 3482991.5 | 3500455.25 |
| O09111 | Ndufb11 | NADH dehydrogenase [ubiquinone] 1 beta subcomplex subunit 11, mitochondrial Mus mus | 37.18 | 11.26 | 17444 | 1 | 1 | 1 | 1 | 1 | 1 | 835302.44 | 1 | 1 | 586911.38 | 1 | 1123373.62 | 2479955.25 | 1712495.833 | 1463830.25 | 1193702 |
| Q9CQZ6 | Ndufb3 | NADH dehydrogenase [ubiquinone] 1 beta subcomplex subunit 3 Mus musculus | 49.58 | 16.35 | 11692 | 1 | 1 | 478849.84 | 939127.62 | 1228604 | 1 | 1 | 301887.53 | 543350.94 | 1178516 | 948843.94 | 389196.25 | 368821.38 | 352597.22 | 744044.69 | 285321.72 |
| Q9CQH3 | Ndufb5 | NADH dehydrogenase [ubiquinone] 1 beta subcomplex subunit 5, mitochondrial Mus musc | 95.5 | 12.7 | 21710 | 2 | 2 | 1 | 1 | 1 | 1 | 1 | 1 | 352113.62 | 1 | 1 | 1 | 1 | 1 | 325236.75 | 371568.81 |
| Q9CQ54 | Ndufc2 | NADH dehydrogenase [ubiquinone] 1 subunit C2 Mus musculus | 41.04 | 10.83 | 14164 | 1 | 1 | 1530846.62 | 1188782.38 | 1155049.5 | 983285.12 | 1004454.69 | 1101889 | 1424151.62 | 1100653.62 | 810604.19 | 1045581.75 | 1212747.38 | 853282.19 | 1115059.75 | 1366438 |
| Q91VD9 | Ndufs1 | NADH-ubiquinone oxidoreductase 75 kDa subunit, mitochondrial Mus musculus | 822.8 | 32.6 | 79777 | 15 | 15 | 17999677 | 19507047 | 19924165 | 19055877 | 18308039 | 19397315 | 20607681 | 16615241 | 20372451 | 19623889 | 19353883 | 18621959 | 20528713 | 19163313 |
| Q91WD5 | Ndufs2 | NADH dehydrogenase [ubiquinone] iron-sulfur protein 2, mitochondrial Mus musculus | 481.97 | 19.65 | 52626 | 7 | 7 | 12808319 | 15790347 | 14714339 | 14247455 | 13583423 | 14250283 | 14641209 | 17082167 | 15426262 | 13794102 | 12945464 | 15299515 | 15262903 | 14555298 |
| Q9DCT2 | Ndufs3 | NADH dehydrogenase [ubiquinone] iron-sulfur protein 3, mitochondrial Mus musculus | 441.05 | 33.08 | 30149 | 7 | 7 | 11549762 | 10014481 | 10929866 | 10903163 | 10815180 | 10659149 | 9660369 | 11341558 | 11064966 | 9309770 | 11130895 | 10249497 | 10631192 | 11899731 |
| Q9DC70 | Ndufs7 | NADH dehydrogenase [ubiquinone] iron-sulfur protein 7, mitochondrial Mus musculus | 360.71 | 30.8 | 24683 | 6 | 6 | 9863442 | 10096206 | 9947674 | 9556809 | 11060647 | 10410752 | 10491083 | 9492803 | 11016054 | 11690093 | 11537952 | 12172173 | 12314664 | 11484388 |
| Q8K3J1 | Ndufs8 | NADH dehydrogenase [ubiquinone] iron-sulfur protein 8, mitochondrial Mus musculus | 221.95 | 17.92 | 24038 | 3 | 3 | 1898490.5 | 2204512 | 1916932.75 | 1833128.62 | 2253961.75 | 2535078 | 2059565.5 | 2602214.5 | 2425363 | 2143960.25 | 2481266.5 | 2556399.5 | 2826770.75 | 2442564.5 |
| Q91YT0 | Ndufv1 | NADH dehydrogenase [ubiquinone] flavoprotein 1, mitochondrial Mus musculus | 482.5 | 37.5 | 50834 | 10 | 10 | 14304968 | 9603685 | 7987856.5 | 10541915 | 11027295 | 8325026 | 9698043 | 11618025 | 11399937 | 11338268 | 13342847 | 7120621 | 9455609 | 9361891 |
| Q9D6J6 | Ndufv2 | NADH dehydrogenase [ubiquinone] flavoprotein 2, mitochondrial Mus musculus | 83.11 | 8.47 | 27285 | 2 | 2 | 2964811.5 | 3032189.75 | 2951593.938 | 2531769.5 | 3277605 | 2699179.75 | 3407906.5 | 3804912.25 | 3901760 | 3300352.5 | 2829209.5 | 3515123 | 3047977 | 3103952.5 |
| Q9CR95 | Necap1 | Adaptin ear-binding coat-associated protein 1 Mus musculus | 227.34 | 26.55 | 29639 | 5 | 5 | 4157405.25 | 4732588.5 | 5920900 | 4974133.5 | 2267945.25 | 4899086 | 2362826.25 | 6072789 | 6541736.5 | 4436228.5 | 2503657.5 | 7786197 | 4882232.5 | 4242051.5 |
| Q9D1J1 | Necap2 | Adaptin ear-binding coat-associated protein 2 Mus musculus | 270.8 | 31.95 | 28598 | 2 | 5 | 3710688 | 4054495.5 | 3515519.5 | 3933436.5 | 4115200 | 5013994 | 5581684.5 | 4389146.5 | 5045821.5 | 5895955.5 | 4591696 | 3863060 | 4302733 | 4362804.5 |
| P29595 | Nedd8 | NEDD8 Mus musculus | 143.7 | 34.57 | 8972 | 3 | 3 | 2820473.75 | 2314128.5 | 2063765 | 2557613.5 | 2498568.5 | 2147223.75 | 2341645.25 | 1868491.5 | 2124641.25 | 2548357.25 | 2259870 | 908756.31 | 942246.5 | 2005464.25 |
| D3ZBE5 | Nek7 | Serine/threonine-protein kinase Nek7 Rattus norvegicus | 34.07 | 2.65 | 34529 | 1 | 1 | 1883261.25 | 1644848.5 | 1576378 | 1386857.75 | 1375839.62 | 1221107.62 | 1434051.12 | 612162.62 | 1331321.88 | 1487324.25 | 1588064 | 1550290.38 | 1355501.5 | 1220094.88 |
| Q8K1R7 | Nek9 | Serine/threonine-protein kinase Nek9 Mus musculus | 255 | 5.89 | 107143 | 5 | 5 | 12113090 | 7071253 | 9067258 | 10640010 | 10020693 | 8287767 | 7676927 | 5826613 | 6125222.5 | 7207948 | 13187758 | 15137051 | 7613699 | 9491888 |
| Q8C4Y3 | Nelfb | Negative elongation factor B Mus musculus | 75.63 | 5.34 | 65636 | 2 | 2 | 1081040.62 | 867708.44 | 1432506.62 | 402737.62 | 398659.06 | 1587010.5 | 974288.81 | 2248569.25 | 2055497.75 | 889497.38 | 1263496.25 | 1210495.5 | 1998030.62 | 1072818.62 |
| Q922L6 | Nelfcd | Negative elongation factor D Mus musculus | 107.55 | 5.25 | 66276 | 2 | 2 | 1230443.75 | 1123769.88 | 1206099 | 1379372.75 | 1355226.75 | 1506306 | 1772333.62 | 1458071 | 1675131.62 | 1627184 | 1595173 | 1401060.5 | 448711.31 | 1207126.25 |
| Q8CCP0 | Nemf | Nuclear export mediator factor Nemf Mus musculus | 38.68 | 0.66 | 121188 | 1 | 1 | 484501.22 | 500323.765 | 477963.78 | 596490 | 442340.06 | 1 | 519787.53 | 465431.09 | 1 | 1 | 542089.19 | 524060.56 | 520910.88 | 480769.16 |
| Q6P5H2 | Nes | Nestin Mus musculus | 2639.19 | 34.39 | 207124 | 46 | 46 | 79465009 | 89528201 | 88990681 | 93163257 | 81493441 | 96563737 | 107526281 | 101315457 | 102183537 | 110900217 | 87106777 | 76921713 | 81664481 | 85680297 |
| O35657 | Neu1 | Sialidase-1 Mus musculus | 37.63 | 2.2 | 44591 | 1 | 1 | 1 | 1 | 1 | 1 | 1 | 291035.47 | 267082.62 | 1 | 1 | 1 | 257865.5 | 1 | 1 | 1 |
| P97526 | Nf1 | Neurofibromin Rattus norvegicus | 65.62 | 0.78 | 317083 | 1 | 1 | 279746.625 | 297530.44 | 277005.28 | 275340.78 | 269110 | 264020.62 | 266138.66 | 245558.67 | 223492.73 | 249802.67 | 280086.03 | 216161.66 | 207308 | 247418.09 |
| Q60591 | Nfatc2 | Nuclear factor of activated T-cells, cytoplasmic 2 Mus musculus | 39.21 | 0.97 | 100020 | 1 | 1 | 668746.62 | 1 | 763157.81 | 837551.31 | 1001204.44 | 847837.56 | 773356.12 | 763969.75 | 765775.69 | 717239.31 | 783058.19 | 871050.31 | 679799.12 | 762864.19 |
| P25799 | Nfkb1 | Nuclear factor NF-kappa-B p105 subunit Mus musculus | 417.44 | 10.71 | 105615 | 9 | 9 | 7976543 | 6919470.5 | 7398418 | 7659858.5 | 7483320 | 7037566 | 6300524 | 6667246 | 7139988 | 7596915 | 6920653.5 | 7201275 | 6591541 | 6821641 |
| Q9WTK5 | Nfkb2 | Nuclear factor NF-kappa-B p100 subunit Mus musculus | 402.2 | 13.35 | 96832 | 8 | 8 | 1745666.38 | 1787161 | 1787656.12 | 1938950.5 | 2539147.5 | 2173838 | 2230846 | 1720164.5 | 2273414 | 1587467.62 | 2200815.5 | 2455245 | 2600027.75 | 2312894.75 |
| O54910 | Nfkbie | NF-kappa-B inhibitor epsilon Mus musculus | 66.86 | 3.3 | 39200 | 1 | 1 | 547031.81 | 576346.38 | 607238.94 | 826548.88 | 678006.31 | 802566 | 700561.62 | 574969.31 | 665226.25 | 569764.62 | 715242.19 | 640290.38 | 737901.25 | 767503.25 |
| Q9QZ23 | Nfu1 | NFU1 iron-sulfur cluster scaffold homolog, mitochondrial Mus musculus | 172.78 | 17.25 | 28568 | 3 | 3 | 7487796 | 4663643 | 8237416 | 8452325 | 9131911 | 8826859 | 9417228 | 8370998.5 | 8222132 | 3837078.5 | 9104046 | 3840594 | 8808137 | 8126258 |
| P63139 | Nfyb | Nuclear transcription factor Y subunit beta Mus musculus | 53.87 | 10.14 | 22787 | 1 | 1 | 130565.93 | 226263.86 | 158491.7 | 181244.34 | 240914.52 | 150331.55 | 151083.55 | 224969.08 | 188662.44 | 177528.41 | 185037.16 | 245525.2 | 146463.47 | 223255.69 |
| P70353 | Nfyc | Nuclear transcription factor Y subunit gamma Mus musculus | 100.51 | 7.46 | 37254 | 2 | 2 | 483421.32 | 322263.41 | 645538.25 | 392668.31 | 573215.31 | 609991.25 | 452662.34 | 435658.06 | 680002.38 | 730935.38 | 507556.31 | 649288.12 | 665427.31 | 416406 |
| Q9DB96 | Ngdn | Neuroguidin Mus musculus | 67.49 | 5.71 | 35659 | 1 | 1 | 402678.41 | 253262.73 | 370786.69 | 345428.28 | 240595.23 | 207479.25 | 281698.47 | 307920.06 | 270298.81 | 266849.1475 | 296121.7967 | 281486.41 | 349422.56 | 257456.42 |
| Q8BZW8 | Nhlrc2 | NHL repeat-containing protein 2 Mus musculus | 38.12 | 1.66 | 78430 | 1 | 1 | 716599.81 | 273726.16 | 923782.62 | 975982.38 | 940498.81 | 930322.88 | 1026392.19 | 829874.56 | 776205.56 | 811157.94 | 832115 | 1058519 | 894799.5 | 828778.12 |
| Q8CCH2 | Nhlrc3 | NHL repeat-containing protein 3 Mus musculus | 485.16 | 52.74 | 38195 | 11 | 11 | 7638655 | 13080141 | 8901204 | 6809373.5 | 12766574 | 13340505 | 12752131 | 13725154 | 11869678 | 10663453 | 12180060 | 10834109 | 11815272 | 12787853 |
| Q9CRB2 | Nhp2 | H/ACA ribonucleoprotein complex subunit 2 Mus musculus | 136.73 | 16.99 | 17247 | 2 | 2 | 5380567 | 5823086.5 | 6158144 | 6858105 | 7214916 | 5769436 | 6333426 | 6162586 | 6282813 | 6408451 | 6802644.5 | 6034491 | 6402899 | 2009959.62 |
| Q9EQ80 | Nif3l1 | NIF3-like protein 1 Mus musculus | 202.23 | 17.29 | 41746 | 5 | 5 | 6159853.5 | 6868709 | 6970456 | 6814075 | 6238669.5 | 6439283 | 7403442 | 6468800 | 5987383 | 7542702.5 | 7506630 | 7617705.5 | 7625864 | 6658359 |
| Q91VE6 | Nifk | MKI67 FHA domain-interacting nucleolar phosphoprotein Mus musculus | 390.61 | 28.71 | 36265 | 7 | 7 | 3278893.5 | 2901934.5 | 5127383 | 4733415.5 | 4422989 | 4105508.75 | 6258921.5 | 3148597.75 | 4972174 | 4498952 | 4739832.5 | 5515447 | 2738695.5 | 4376764 |
| Q9CXK8 | Nip7 | 60S ribosome subunit biogenesis protein NIP7 homolog Mus musculus | 196.89 | 42.78 | 20452 | 5 | 5 | 6615078 | 5026366.5 | 4666064 | 5472882.5 | 4653561 | 4937485 | 5609427.5 | 6184283.5 | 7036061.5 | 5640202 | 4812681.5 | 4516796 | 6131842 | 4554375 |
| Q6KCD5 | Nipbl | Nipped-B-like protein Mus musculus | 106.75 | 0.93 | 315450 | 2 | 2 | 214160.67 | 1076155.88 | 1031369 | 1035851.31 | 914168.38 | 907483.25 | 972907.12 | 1066515.62 | 981399.62 | 832473.62 | 1268763.62 | 1116732.62 | 967901.12 | 807946.62 |
| O55125 | Nipsnap1 | Protein NipSnap homolog 1 Mus musculus | 49.61 | 3.17 | 33363 | 1 | 1 | 6229544 | 3744931.5 | 4046890.5 | 3863978.75 | 4515974.5 | 4804138.5 | 3847106.5 | 4318099.5 | 4218885.5 | 4487122.5 | 4943370.5 | 3484867.75 | 4600299.5 | 4342845.917 |
| O55126 | Nipsnap2 | Protein NipSnap homolog 2 Mus musculus | 94.84 | 9.96 | 32933 | 2 | 2 | 524071.69 | 492649.66 | 356549.66 | 453230.065 | 439649.25 | 340320.25 | 447107.22 | 493098.78 | 473948.09 | 711019.56 | 442633.41 | 600941 | 679460.31 | 561239 |
| Q9CQE1 | Nipsnap3b | Protein NipSnap homolog 3B Mus musculus | 128.67 | 13.77 | 28308 | 2 | 2 | 2797086 | 2111008.5 | 2202724 | 737958.81 | 2399881.5 | 1816095.25 | 2817949 | 2532825.5 | 3311314 | 1937596 | 1697562.38 | 2123787.25 | 2524358 | 1794579.62 |
| Q80TM9 | Nisch | Nischarin Mus musculus | 422.25 | 5.15 | 175012 | 6 | 6 | 7754023 | 3385398 | 4322361.5 | 3333334.5 | 4967568 | 5752486.5 | 4370999.5 | 5189663 | 5388123.5 | 4586552 | 4339880 | 7112966 | 3900587 | 4810991.5 |
| Q8VDK1 | Nit1 | Deaminated glutathione amidase Mus musculus | 177.25 | 15.17 | 35705 | 3 | 3 | 2300366 | 2724879.5 | 3690259.5 | 3415774 | 2825136.25 | 3218703.25 | 3313335.25 | 3508677.75 | 3786869.25 | 3107660.25 | 2877110.5 | 2014476.75 | 3009346.25 | 2624210.75 |
| Q9JHW2 | Nit2 | Omega-amidase NIT2 Mus musculus | 414.17 | 32.61 | 30502 | 6 | 6 | 6482804 | 8996017 | 8459023 | 7126254 | 7951580 | 7483090 | 7631883 | 11813896 | 7334443 | 6539870 | 9369764 | 7462286 | 7780491 | 9162395 |
| P42676 | Nln | Neurolysin, mitochondrial Rattus norvegicus | 83.57 | 2.41 | 80254 | 2 | 2 | 1119613.5 | 946692.56 | 1929110.25 | 1672242.62 | 2221845 | 1897786 | 980441.62 | 559619 | 1697266.75 | 2144839.25 | 1298514 | 1999994 | 2165455 | 941992.19 |
| Q3TL44 | Nlrx1 | NLR family member X1 Mus musculus | 82.6 | 4.31 | 107831 | 2 | 2 | 673997.12 | 496556.88 | 627119 | 480419.31 | 634754.94 | 664639.5 | 363206.72 | 700599.56 | 487622.12 | 311436.94 | 644352 | 476817.56 | 788712.81 | 431229.22 |
| Q99L48 | Nmd3 | 60S ribosomal export protein NMD3 Mus musculus | 165.05 | 9.54 | 57611 | 4 | 4 | 1698565 | 695013.06 | 1551777 | 491514.81 | 1111073 | 725061.06 | 651329 | 1284058.38 | 538977.31 | 799856.4375 | 702219.69 | 848449.81 | 1063038.75 | 991702.12 |
| P15532 | Nme1 | Nucleoside diphosphate kinase A Mus musculus | 480.54 | 73.03 | 17208 | 4 | 10 | 104322881 | 79151873 | 83748209 | 89986673 | 72114777 | 74421577 | 85996841 | 85371401 | 55565741 | 77345793 | 82290609 | 79972969 | 78264753 | 73063441 |
| Q01768 | Nme2 | Nucleoside diphosphate kinase B Mus musculus | 594.75 | 69.08 | 17363 | 3 | 10 | 146289873 | 108808457 | 121069241 | 123355457 | 99568065 | 99580001 | 117121217 | 112471961 | 86216425 | 106182025 | 111967345 | 105825209 | 107361585 | 98366049 |
| O35309 | Nmi | N-myc-interactor Mus musculus | 100.72 | 9.55 | 35236 | 2 | 2 | 2499665.5 | 2235807.25 | 2187643.75 | 2451689.75 | 2832923.5 | 1924270.12 | 1708475.75 | 2000299.75 | 1985911.62 | 2119268.5 | 1853734.75 | 1856813.88 | 1985596.25 | 2076469.75 |
| Q8K2T1 | Nmral1 | NmrA-like family domain-containing protein 1 Mus musculus | 327.72 | 27.18 | 34376 | 6 | 6 | 3122672.75 | 3768937.75 | 2600572 | 2868376 | 2902405.25 | 2784182.5 | 3380288.25 | 3328940 | 3447933.5 | 3208299.75 | 2861374.5 | 3695901.5 | 3595160.75 | 2278966.25 |
| O70310 | Nmt1 | Glycylpeptide N-tetradecanoyltransferase 1 Mus musculus | 804.42 | 38.51 | 56888 | 13 | 16 | 16814465 | 22363301 | 20927843 | 25617065 | 22983845 | 27136609 | 27996637 | 25131361 | 26008205 | 27772369 | 24512173 | 25642613 | 22160425 | 21705153 |
| O70311 | Nmt2 | Glycylpeptide N-tetradecanoyltransferase 2 Mus musculus | 225.93 | 10.02 | 60484 | 1 | 4 | 3513524.25 | 2774711.5 | 3552664 | 3528102 | 1920932.75 | 2616762.25 | 3341969 | 4432627 | 4137221 | 2936518.25 | 2809168 | 3422018 | 2807941 | 2786955.25 |
| Q61941 | Nnt | NAD(P) transhydrogenase, mitochondrial Mus musculus | 251.93 | 9.3 | 113838 | 7 | 7 | 5226674.5 | 5245070.5 | 7385412.5 | 7033046 | 9416817 | 7020352 | 8028919.5 | 6834275.5 | 7243213 | 7535295 | 6675738.5 | 4676549 | 7274160.5 | 6355252.5 |
| Q8BW10 | Nob1 | RNA-binding protein NOB1 Mus musculus | 112.9 | 7.94 | 45464 | 2 | 2 | 1 | 683194.5 | 1 | 1 | 1 | 399147.19 | 1 | 1 | 701287.88 | 786419.69 | 1 | 989083.94 | 799372.19 | 1 |
| Q8BJW5 | Nol11 | Nucleolar protein 11 Mus musculus | 286.94 | 10.24 | 80818 | 7 | 7 | 924443.62 | 2229335.5 | 2206846.5 | 2595892.5 | 2166788.25 | 2032539.75 | 1597533.5 | 1806600.62 | 1661428.62 | 1918885.5 | 2611832.75 | 2411682.25 | 1472112 | 1807073.12 |
| Q8R5K4 | Nol6 | Nucleolar protein 6 Mus musculus | 141.63 | 4.08 | 129228 | 3 | 3 | 1330696.12 | 620115 | 258588.98 | 663051.7525 | 442806.91 | 232329.73 | 455118.31 | 410166.47 | 940396.38 | 1162880.5 | 980589.25 | 341043.12 | 1175385.12 | 939250.25 |
| Q3TZX8 | Nol9 | Polynucleotide 5'-hydroxyl-kinase NOL9 Mus musculus | 169.82 | 5.6 | 80840 | 3 | 3 | 628201.56 | 809801.6325 | 462534.03 | 1214701.25 | 933769.69 | 623293.31 | 558006.62 | 596378.69 | 1464659.12 | 727271.62 | 1 | 1008842.19 | 665226.25 | 615351.88 |
| E9Q5C9 | Nolc1 | Nucleolar and coiled-body phosphoprotein 1 Mus musculus | 137.62 | 3.85 | 73698 | 3 | 3 | 74356.43 | 100289.9025 | 93275.03 | 72647.51 | 160880.64 | 243495.84 | 103542.52 | 58916.68 | 1 | 1 | 1 | 1 | 1 | 128452.16 |
| Q6GQT9 | Nomo1 | Nodal modulator 1 Mus musculus | 1342.23 | 33.69 | 133420 | 24 | 24 | 23861947 | 19982323 | 21988367 | 22763577 | 23571889 | 23100981 | 21726689 | 21187129 | 21286619 | 21816903 | 22521585 | 20929623 | 23779793 | 23654617 |
| Q99K48 | Nono | Non-POU domain-containing octamer-binding protein Mus musculus | 914.56 | 49.68 | 54541 | 15 | 16 | 23923795 | 25871525 | 28505361 | 29361659 | 24664091 | 25283995 | 25652885 | 21702423 | 28922769 | 30612999 | 29939305 | 27258545 | 31435499 | 29200449 |
| Q8R3N1 | Nop14 | Nucleolar protein 14 Mus musculus | 132.46 | 5.12 | 98769 | 3 | 3 | 1 | 399382.25 | 674298.75 | 271218.31 | 1 | 730307.19 | 599676.5 | 759479.06 | 674728.75 | 402867.47 | 745488.12 | 566358.75 | 588692 | 697021.81 |
| Q9CPT5 | Nop16 | Nucleolar protein 16 Mus musculus | 79.68 | 8.99 | 21139 | 1 | 1 | 1 | 397197.81 | 604668.38 | 1 | 1 | 1 | 493364.12 | 262030.47 | 445541.44 | 1 | 1 | 1 | 1 | 617516.88 |
| Q922K7 | Nop2 | Probable 28S rRNA (cytosine-C(5))-methyltransferase Mus musculus | 669.24 | 21.56 | 86752 | 13 | 13 | 10487542 | 7249033.5 | 16366221 | 8456655 | 8513229 | 11395624 | 13326432 | 11370652 | 11304950 | 10908963 | 8168240 | 10455470 | 12748198 | 9808905 |
| Q9D6Z1 | Nop56 | Nucleolar protein 56 Mus musculus | 1414.75 | 56.9 | 64464 | 24 | 24 | 47372017 | 40001381 | 43229321 | 44106425 | 45624717 | 41749561 | 41148133 | 40565105 | 40820873 | 43295689 | 36609689 | 37039849 | 42972041 | 40682253 |
| Q6DFW4 | Nop58 | Nucleolar protein 58 Mus musculus | 988 | 39.18 | 60343 | 15 | 17 | 29170077 | 31379703 | 29777683 | 27070727 | 30653845 | 27886977 | 29497003 | 29480497 | 28375265 | 28285413 | 26309365 | 25949129 | 26671935 | 27507261 |
| Q8BMC4 | Nop9 | Nucleolar protein 9 Mus musculus | 231.92 | 6.6 | 70046 | 3 | 3 | 639804.75 | 481929.97 | 787109.12 | 1003767.75 | 918819.75 | 833017 | 434852.91 | 504399.06 | 820938 | 967026.06 | 1033216.88 | 849146.31 | 1127192.88 | 862558.5 |
| O35604 | Npc1 | NPC intracellular cholesterol transporter 1 Mus musculus | 240.09 | 5.56 | 142885 | 5 | 5 | 5466884 | 8042086 | 6392956.5 | 5789944.5 | 7449742 | 8357751 | 6803594 | 7030585.5 | 7627408 | 6941569.5 | 7703148 | 7257868.5 | 8333883 | 7459235.5 |
| Q9Z0J0 | Npc2 | NPC intracellular cholesterol transporter 2 Mus musculus | 223.6 | 41.61 | 16442 | 5 | 5 | 44724541 | 60019929 | 47149653 | 40513769 | 46260973 | 43541525 | 41508521 | 50368157 | 37531517 | 39154353 | 39613041 | 35064985 | 48718329 | 42032141 |
| Q6NSR8 | Npepl1 | Probable aminopeptidase NPEPL1 Mus musculus | 394.99 | 16.6 | 55940 | 6 | 6 | 22768453 | 16636672 | 17058027 | 19636999 | 14535225 | 15084541 | 17668589 | 13377728 | 14303742 | 16406094 | 12712750 | 16162779 | 13830483 | 11917200 |
| Q11011 | Npepps | Puromycin-sensitive aminopeptidase Mus musculus | 879.35 | 30.11 | 103325 | 19 | 19 | 29901503 | 28245597 | 28708541 | 28451621 | 28245683 | 30624523 | 32642347 | 36832961 | 33104643 | 31351347 | 23796579 | 30500277 | 28903423 | 29151335 |
| P60670 | Nploc4 | Nuclear protein localization protein 4 homolog Mus musculus | 561.66 | 28.45 | 68017 | 9 | 9 | 3993311 | 3392168 | 6051331.5 | 2780395.5 | 3892635.5 | 5666318 | 3710267 | 4447966 | 5938746 | 6029245 | 3915883.25 | 6158794.5 | 6635508 | 4865594 |
| P13084 | Npm1 | Nucleophosmin Rattus norvegicus | 979.92 | 45.21 | 32560 | 12 | 14 | 136147089 | 112855337 | 132390833 | 127735353 | 137749089 | 128580457 | 115802137 | 84910697 | 105577569 | 144389041 | 114415505 | 120761025 | 141716145 | 132780065 |
| Q9CPP0 | Npm3 | Nucleoplasmin-3 Mus musculus | 214.55 | 25.71 | 19023 | 3 | 3 | 3219045 | 2339045.5 | 4039733.25 | 4012519.25 | 4656113 | 4518106 | 3781551 | 1812436 | 3530968.5 | 3902311.25 | 6276559 | 6426448 | 5180483.5 | 4017750 |
| P97300 | Nptn | Neuroplastin Mus musculus | 130.03 | 6.3 | 44373 | 2 | 2 | 3599201 | 3667696.75 | 3625200 | 4168693 | 4074775 | 3848506 | 4486480 | 4067270 | 3203171 | 4212236 | 4563831 | 4168169 | 4989253 | 3838349 |

| | | | | | | | | | | | | | | | | | | | | | |
|---|---|---|---|---|---|---|---|---|---|---|---|---|---|---|---|---|---|---|---|---|---|
| Q9JI75 | Nqo2 | Ribosyldihydronicotinamide dehydrogenase [quinone] Mus musculus | 120.65 | 14.72 | 26248 | 2 | 2 | 2998852.75 | 3186597 | 3126613.75 | 3019940.5 | 2777386.5 | 2932392 | 2881538.75 | 3869768.5 | 3381857.25 | 2831846.5 | 3095132 | 2868639.5 | 3760020.25 | 3032757 |
| P08556 | Nras | GTPase NRas Mus musculus | 188.98 | 25.4 | 21199 | 3 | 3 | 4121496.5 | 3810453.25 | 5570454 | 5352785 | 6521370 | 4965758 | 6390140 | 5694309 | 6369140.5 | 5000817.5 | 4539525 | 4813464.5 | 6551577 | 6106566 |
| Q8BHG1 | Nrdc | Nardilysin Mus musculus | 374.9 | 10.51 | 132891 | 9 | 9 | 10176996 | 8305207 | 7807701.5 | 7826306.5 | 7628328 | 9184277 | 8397776 | 8553708 | 8506037 | 6806395 | 7148031.5 | 7291352.5 | 7414464 | 5565442.5 |
| P97333 | Nrp1 | Neuropilin-1 Mus musculus | 105.47 | 3.58 | 103000 | 2 | 2 | 2158467 | 1629235.62 | 1624815.75 | 1570022 | 1405104.25 | 1159563.25 | 1206585.75 | 818527.38 | 1518075.5 | 1235982.25 | 846107.31 | 1576009.88 | 1843321 | 1457088.5 |
| Q9R1J0 | Nsdhl | Sterol-4-alpha-carboxylate 3-dehydrogenase, decarboxylating Mus musculus | 330.15 | 25.97 | 40686 | 6 | 6 | 4963761 | 2714816.5 | 4315897 | 3331419.5 | 3430446 | 3702811.5 | 4779635.5 | 5447125.5 | 5465270.5 | 4535292.5 | 3867191.25 | 5999567.5 | 5942568 | 4402627.5 |
| P46460 | Nsf | Vesicle-fusing ATPase Mus musculus | 1487.1 | 46.24 | 82613 | 28 | 28 | 28648385 | 35985257 | 34471449 | 30676035 | 35588877 | 27917119 | 31029069 | 32253119 | 31423795 | 31872981 | 36433893 | 34006465 | 37673045 | 33935913 |
| O35987 | Nsfl1c | NSFL1 cofactor p47 Rattus norvegicus | 667.58 | 40.54 | 40680 | 11 | 11 | 18535299 | 21998747 | 17681253 | 24993347 | 20707339 | 24815205 | 22486945 | 22766843 | 22060637 | 25184213 | 23302969 | 22815001 | 24445089 | 21239425 |
| Q91VT1 | Nsmce2 | E3 SUMO-protein ligase NSE2 Mus musculus 2 | 75.68 | 6.07 | 28231 | 1 | 1 | 1 | 1 | 507294.69 | 1 | 1 | 373920.12 | 496186.4275 | 827743.62 | 389414.81 | 393667.16 | 1 | 1 | 646454.25 | 1 |
| Q4FZU3 | Nsrp1 | Nuclear speckle splicing regulatory protein 1 Rattus norvegicus | 36.96 | 1.83 | 64180 | 1 | 1 | 90277.63 | 2875351.25 | 2368752.5 | 3279576.75 | 3486250 | 116635.16 | 127676.48 | 325041.2075 | 924618.25 | 131234.94 | 3615135 | 134687.25 | 1 | 2205424.25 |
| Q1HFZ0 | Nsun2 | tRNA (cytosine(34)-C(5))-methyltransferase Mus musculus | 549.78 | 19.68 | 85452 | 10 | 10 | 11462024 | 11726053 | 11252619 | 10511617 | 9920910 | 12373602 | 11274469 | 12853241 | 12096182 | 10906245 | 11469071 | 10961164 | 11645194 | 10995175 |
| Q9JM14 | Nt5c | 5'(3')-deoxyribonucleotidase, cytosolic type Mus musculus | 350.22 | 38.5 | 23076 | 4 | 4 | 4898428.5 | 5409124.5 | 4442569 | 6602330.5 | 4849104 | 5711646 | 4712784 | 5665211 | 5571696 | 5542706 | 5176059 | 4394817 | 4564345 | 4155570.5 |
| Q3V1L4 | Nt5c2 | Cytosolic purine 5'-nucleotidase Mus musculus | 785.37 | 36.07 | 64809 | 15 | 15 | 11805436 | 15177108 | 12163181 | 14185695 | 11898906 | 11413090 | 8817471 | 11833355 | 10476306 | 13334923 | 11876799 | 12659233 | 11606854 | 12063915 |
| Q3UFY7 | Nt5c3b | 7-methylguanosine phosphate-specific 5'-nucleotidase Mus musculus | 172.13 | 19.67 | 34425 | 3 | 3 | 1345667.12 | 1863601.25 | 2155446.75 | 689296.62 | 1895672 | 1824904.5 | 1972082.12 | 3107463.75 | 2671884.5 | 1649248.38 | 1503464 | 1584113.88 | 2398119.75 | 2094887.62 |
| Q5BJX0 | Ntmt1 | N-terminal Xaa-Pro-Lys N-methyltransferase 1 Rattus norvegicus | 53.68 | 3.14 | 25464 | 1 | 1 | 826081 | 663944.56 | 877264.38 | 470766.31 | 690390.19 | 729619.38 | 819478.06 | 611749.56 | 600513.31 | 714777.62 | 593977.25 | 657541.75 | 549476.88 | 630548.06 |
| Q9CQA9 | Ntpcr | Cancer-related nucleoside-triphosphatase homolog Mus musculus | 144.1 | 28.95 | 20667 | 3 | 3 | 7992914.5 | 5703867 | 2309010 | 4058435.5 | 3470464.25 | 3532130.75 | 2242638 | 3287174.25 | 2484500.75 | 1878141.5 | 4141190.5 | 11291505 | 1943357.88 | 8304120 |
| P54729 | Nub1 | NEDD8 ultimate buster 1 Mus musculus | 132.86 | 7.98 | 70307 | 3 | 3 | 999581.62 | 1611360.38 | 839202.38 | 806820.19 | 554152.69 | 943152.62 | 1293741.75 | 418387.78 | 470539.62 | 621647.31 | 598639.19 | 1345459.62 | 593042.31 | 485880.81 |
| Q9R061 | Nubp2 | Cytosolic Fe-S cluster assembly factor NUBP2 Mus musculus | 133.66 | 14.91 | 29518 | 3 | 3 | 3554955 | 4566682 | 2580230 | 3886354 | 3901968 | 3684290.5 | 4339375 | 4390924 | 3883403 | 4174767.75 | 4004405 | 2334667.75 | 4357604 | 4553663 |
| Q02819 | Nucb1 | Nucleobindin-1 Mus musculus | 158.35 | 10.68 | 53409 | 3 | 3 | 1698301.12 | 1853747.88 | 1118186.75 | 1884950.25 | 1270880.88 | 1686537.75 | 1501021.62 | 1210231.62 | 906851.69 | 1180110 | 1104514 | 1157213.75 | 1568818.12 | 1361571.25 |
| P81117 | Nucb2 | Nucleobindin-2 Mus musculus | 207.66 | 14.05 | 50304 | 4 | 4 | 4508465 | 4875450.5 | 4607830.5 | 4960994 | 4038016.25 | 5023095 | 4819713 | 5995018.5 | 5246441.5 | 4078044.5 | 4564077 | 3941968.5 | 4429040 | 4601885 |
| Q80XU3 | Nucks1 | Nuclear ubiquitous casein and cyclin-dependent kinase substrate 1 Mus musculus | 140.91 | 10.26 | 26313 | 2 | 2 | 1549337.88 | 5748996 | 4899099 | 5975176 | 8229839.5 | 6144813 | 7093305 | 4121454.5 | 5670334 | 7240687 | 6486060 | 6722346.5 | 6854845 | 7219808 |
| O35685 | Nudc | Nuclear migration protein nudC Mus musculus | 1046.91 | 67.47 | 38358 | 18 | 18 | 52287017 | 53520913 | 51342013 | 55752165 | 56364201 | 55256685 | 53613489 | 55893205 | 51583341 | 59422521 | 56021937 | 55235965 | 55846529 | 57259277 |
| Q6PIP5 | Nudcd1 | NudC domain-containing protein 1 Mus musculus | 391 | 24.4 | 66705 | 9 | 9 | 8369140.5 | 8550534 | 7894536 | 8618849 | 8807447 | 9948377 | 8114257 | 8104016 | 9682271 | 9112804 | 7270102.5 | 8957691 | 9748315 | 10246612 |
| Q5M823 | Nudcd2 | NudC domain-containing protein 2 Rattus norvegicus 2 | 177.9 | 24.2 | 17674 | 3 | 3 | 2781873.25 | 2213616.75 | 1811639.62 | 2710745.5 | 2033663.25 | 2266439.25 | 1862861.12 | 380480.81 | 1733112.62 | 2009280.38 | 2396760 | 2109264.5 | 1834830.88 | 1795667.75 |
| Q8JZU0 | Nudt13 | Nucleoside diphosphate-linked moiety X motif 13 Mus musculus | 50.78 | 5.68 | 39137 | 1 | 1 | 508560.72 | 513306.47 | 533769.44 | 528764.38 | 458402.59 | 1 | 784849.62 | 1 | 604025.19 | 477041.47 | 565302.69 | 529990.94 | 646958.5 | 580750.71 |
| Q8VHN8 | Nudt16l1 | Tudor-interacting repair regulator protein Mus musculus | 111.47 | 19.43 | 23414 | 3 | 3 | 1841913.25 | 1 | 1 | 1 | 254077.17 | 473499.28 | 1 | 1 | 308796.44 | 1 | 263609.69 | 376339.31 | 744044.69 | 302809.06 |
| Q3U2V3 | Nudt18 | 8-oxo-dGDP phosphatase NUDT18 Mus musculus | 59.15 | 4.95 | 35694 | 1 | 1 | 211769.22 | 261989.75 | 238622.64 | 305746.12 | 292658.69 | 295838.31 | 269093.47 | 251387.38 | 289970.91 | 271048.84 | 297852.34 | 346240.81 | 227357.02 | 284822.16 |
| P56380 | Nudt2 | Bis(5'-nucleosyl)-tetraphosphatase [asymmetrical] Mus musculus | 172.09 | 27.89 | 16989 | 3 | 3 | 1725483.5 | 1933229.5 | 3011968 | 2932848.5 | 2588678.25 | 978708.69 | 1643077 | 3896739.5 | 3360089.25 | 2377452 | 850681.81 | 2536435.5 | 1428941.25 | 1666403 |
| Q4KM65 | Nudt21 | Cleavage and polyadenylation specificity factor subunit 5 Rattus norvegicus 2 | 385.6 | 49.78 | 26240 | 8 | 8 | 738469.69 | 646067.06 | 523869.62 | 985249.62 | 627260.62 | 450886.12 | 676489 | 386052.75 | 1274801.62 | 616907.25 | 475224 | 523894.88 | 430083.38 | 364384.31 |
| Q9JKX6 | Nudt5 | ADP-sugar pyrophosphatase Mus musculus | 128.9 | 23.85 | 23984 | 3 | 3 | 2263312.75 | 1435845.38 | 2279520.75 | 2242799 | 1905312.38 | 1622416.5 | 1968444.5 | 1545127.38 | 1797826.25 | 2018167.12 | 1715199.5 | 2686612.5 | 2086853.25 | 2149794.25 |
| E9Q7G0 | Numa1 | Nuclear mitotic apparatus protein 1 Mus musculus | 2416.61 | 29.89 | 235630 | 47 | 47 | 27765579 | 32704075 | 33247279 | 38427785 | 36773177 | 28247333 | 33143867 | 24669315 | 32452051 | 34839833 | 36699961 | 31102257 | 32406731 | 32193059 |
| Q8BH74 | Nup107 | Nuclear pore complex protein Nup107 Mus musculus | 111.85 | 3.02 | 106717 | 2 | 2 | 1156890.25 | 880226.56 | 1242359.88 | 986539.31 | 916754.88 | 1229506.25 | 915353.88 | 1348421.88 | 1221404.75 | 883657.62 | 1032342.38 | 1092385.75 | 1050017.5 | 1132616.5 |
| Q8R0G9 | Nup133 | Nuclear pore complex protein Nup133 Mus musculus | 1105.75 | 22.16 | 128620 | 18 | 18 | 5994157 | 6022322 | 6925766.5 | 7015285.5 | 5036463.5 | 7401675 | 4458878.5 | 6352625.5 | 6554018.5 | 7173141 | 6545229.5 | 7067937.5 | 6836429.5 | 7728178.5 |
| P49791 | Nup153 | Nuclear pore complex protein Nup153 Rattus norvegicus | 155.16 | 3.54 | 152824 | 3 | 3 | 705399.83 | 598635.19 | 886343.19 | 637318.19 | 699302.75 | 542243.19 | 755674.38 | 369400.75 | 521996.44 | 462220.25 | 460223.41 | 699763.88 | 744044.69 | 484108.44 |
| Q99P88 | Nup155 | Nuclear pore complex protein Nup155 Mus musculus | 1193.77 | 24.51 | 155118 | 22 | 22 | 10530650 | 10842067 | 9080624 | 8302901 | 10888363 | 10274555 | 11190274 | 12492964 | 18025745 | 10693184 | 13959500 | 12450801 | 10165123 | 10513355 |
| Q9Z0W3 | Nup160 | Nuclear pore complex protein Nup160 Mus musculus | 297.61 | 6.28 | 158232 | 6 | 6 | 2887493.25 | 2270639 | 2146823 | 1754315 | 2257415.5 | 2388964 | 2603269.5 | 3385009.25 | 2956806 | 2249611 | 2346292 | 2081715.75 | 2546587.75 | 2823699.75 |
| Q6ZQH8 | Nup188 | Nucleoporin NUP188 homolog Mus musculus | 42.66 | 0.97 | 196696 | 1 | 1 | 194343.38 | 175196.77 | 140924.03 | 174023.38 | 229176.86 | 332882.53 | 157218.55 | 143082.16 | 203194.67 | 166850.3 | 149749.92 | 166936.64 | 175667.02 | 192528.45 |
| Q9QY81 | Nup210 | Nuclear pore membrane glycoprotein 210 Mus musculus | 293.1 | 4.88 | 204101 | 7 | 7 | 1168368.62 | 2198284 | 1950679.5 | 1867510 | 787036.12 | 2320107 | 2474352.5 | 3043822 | 1531970.5 | 1347138.62 | 2328416.75 | 1766895.62 | 1169463.25 | 2404587 |
| Q80U93 | Nup214 | Nuclear pore complex protein Nup214 Mus musculus | 531.86 | 9.16 | 212979 | 10 | 10 | 3388357.25 | 7711486.5 | 7344560 | 8551104 | 7801118 | 7986248 | 2964795.25 | 4287624 | 4930800 | 4158120.75 | 3125545.75 | 11782749 | 5879064 | 6526024 |
| Q8R4R6 | Nup35 | Nucleoporin NUP35 Mus musculus | 173.28 | 14.15 | 34786 | 3 | 3 | 1 | 1 | 670254.44 | 1409240.25 | 1 | 369759.44 | 417039.62 | 527652.3725 | 794974.31 | 528836.12 | 439433.38 | 580166.75 | 604524.44 | 1303188 |
| Q9CWU9 | Nup37 | Nucleoporin Nup37 Mus musculus | 165.38 | 13.5 | 36732 | 3 | 3 | 1842334.75 | 2400950.25 | 686927.44 | 2705120 | 1737793 | 2427204 | 698270.25 | 1768264.62 | 1987154.88 | 1726848 | 1573037 | 1655277.75 | 1153346.75 | 1699094.25 |
| P59235 | Nup43 | Nucleoporin Nup43 Mus musculus | 92.14 | 8.16 | 41990 | 2 | 2 | 1128446.5 | 328673.97 | 1697451.62 | 595154.75 | 1537837.5 | 1 | 1784960.62 | 1 | 300642.56 | 1411530.12 | 1345697.12 | 1639469 | 2086417 | 1455811.12 |
| Q9JIH2 | Nup50 | Nuclear pore complex protein Nup50 Mus musculus | 343.54 | 19.96 | 49485 | 7 | 7 | 2637541 | 1693044.5 | 1724740.5 | 2573843.5 | 1836018.38 | 2717343.5 | 1283970.38 | 2088073.5 | 2155530 | 1791553.5 | 3295505 | 2572841.5 | 3118086 | 3245504.5 |
| Q8R332 | Nup58 | Nucleoporin p58/p45 Mus musculus | 42.75 | 2.21 | 59445 | 1 | 1 | 559396.3575 | 383305.03 | 709591.94 | 662437.12 | 482251.34 | 633939.88 | 1 | 1 | 1 | 739483 | 1 | 1 | 585993 | 672210.44 |
| P17955 | Nup62 | Nuclear pore glycoprotein p62 Rattus norvegicus | 55.66 | 5.33 | 53397 | 2 | 2 | 589935.69 | 521881.25 | 1020462.38 | 638039.3225 | 419877.97 | 490925.09 | 379840.34 | 572300.31 | 1599397.88 | 423750.59 | 398476.88 | 562164.56 | 1 | 1 |
| Q8R480 | Nup85 | Nuclear pore complex protein Nup85 Mus musculus | 83.37 | 2.9 | 74776 | 1 | 2 | 1118061.62 | 1210350.38 | 1055577.515 | 1357544.5 | 536353.56 | 1246453.25 | 1237365.12 | 794644.5 | 1052164.38 | 1504367.75 | 1149092.5 | 1176176.12 | 1144966.25 | 1334476.12 |
| O08658 | Nup88 | Nuclear pore complex protein Nup88 Rattus norvegicus | 98.15 | 4.45 | 83585 | 2 | 2 | 839509.62 | 807910.88 | 751638.9375 | 702566.31 | 656568.94 | 698011.06 | 821490.81 | 1024340.12 | 916390 | 633406.38 | 629571.19 | 827591.62 | 675152.44 | 819079.5 |
| Q8BJ71 | Nup93 | Nuclear pore complex protein Nup93 Mus musculus | 1117.96 | 38.83 | 93281 | 22 | 22 | 15696478 | 18930093 | 18347425 | 14754703 | 15892601 | 17926043 | 16125125 | 19651781 | 18332439 | 18205665 | 16419797 | 19433177 | 17565805 | 16804729 |
| Q6PFD9 | Nup98 | Nuclear pore complex protein Nup98-Nup96 Mus musculus | 440.96 | 10.63 | 197241 | 4 | 11 | 4648943.5 | 4615951 | 4380564 | 3515120 | 4242840.5 | 3587882 | 3049986.75 | 2646424.5 | 1838146.5 | 2977052.5 | 4061990.75 | 3270712.75 | 2630142.5 | 3682314.5 |
| P61971 | Nutf2 | Nuclear transport factor 2 Mus musculus | 220.68 | 55.12 | 14478 | 4 | 4 | 3060826 | 4045924.75 | 3370719.75 | 4297897.5 | 3546473.5 | 2113440.5 | 3764887.25 | 4325519.5 | 2490633.5 | 2864943.5 | 3392063.5 | 3058325.25 | 2632412 | 3170072 |
| Q9DBY8 | Nvl | Nuclear valosin-containing protein-like Mus musculus | 211.68 | 7.25 | 94476 | 4 | 4 | 5006893 | 3699745.25 | 4004343 | 4415970 | 4322784.5 | 3261414.75 | 2164080.5 | 1853806.5 | 1463487 | 1505472.75 | 4264351.5 | 5289536 | 3331857 | 3668079 |
| Q99JX7 | Nxf1 | Nuclear RNA export factor 1 Mus musculus | 419.97 | 21.68 | 70300 | 8 | 8 | 4546236.5 | 3721399 | 4519531 | 4359245 | 3964913.25 | 4827388 | 4173215.25 | 4199958.5 | 4402474 | 3779387.25 | 5039329 | 4808097 | 3865545.5 | 3565534.5 |
| P11928 | Oas1a | 2'-5'-oligoadenylate synthase 1A Mus musculus | 477.04 | 27.79 | 42429 | 7 | 7 | 6047583.5 | 6469717 | 5685870 | 2889836.75 | 4769320.5 | 2935058 | 2408509.25 | 1142124 | 2262447.75 | 4146578.5 | 3218126.75 | 2340372.5 | 4985915 | 3930411.5 |
| Q8VI93 | Oas3 | 2'-5'-oligoadenylate synthase 3 Mus musculus | 1016.07 | 27.15 | 126333 | 21 | 21 | 15974175 | 14548404 | 17765663 | 11573377 | 15456911 | 14922496 | 10426809 | 14042148 | 15902040 | 15410571 | 11308939 | 11799536 | 13401145 | 8710309 |
| Q8VI94 | Oasl1 | 2'-5'-oligoadenylate synthase-like protein 1 Mus musculus | 173.91 | 10.37 | 59088 | 4 | 4 | 902312 | 652955.69 | 787475.69 | 421891.06 | 521197.62 | 1 | 1 | 1 | 807015.75 | 1 | 1 | 105680.22 | 90020.33 | 1 |
| P29758 | Oat | Ornithine aminotransferase, mitochondrial Mus musculus | 1084.71 | 50.11 | 48355 | 16 | 16 | 54754557 | 48940553 | 49442789 | 54602601 | 45663557 | 52761209 | 58228573 | 60343269 | 58608221 | 55362069 | 62513061 | 64798829 | 64568749 | 55856461 |
| Q9CRD0 | Ociad1 | OCIA domain-containing protein 1 Mus musculus | 177.45 | 19.43 | 27610 | 1 | 3 | 1188468.12 | 1515241.5 | 1298614.5 | 1118878 | 1385194 | 1393171.75 | 1679475.75 | 1646007.88 | 1415659.5 | 1500017.75 | 1348513.88 | 1515646.5 | 1579048.38 | 1744926.75 |
| Q4PJX1 | Odr4 | Protein odr-4 homolog Mus musculus | 49.47 | 3.13 | 49998 | 1 | 1 | 927952.94 | 1150725.38 | 1484045.62 | 1079645.25 | 1102688.75 | 1207669.62 | 1110242.62 | 1083308 | 1033625.56 | 1388030.75 | 1052438.38 | 1067821.12 | 1210329.25 | 1110196.25 |
| Q9EQQ9 | Oga | Protein O-GlcNAcase Mus musculus | 122.1 | 7.64 | 103162 | 3 | 3 | 1154215.38 | 1077223 | 1109541.12 | 977999.25 | 985560.69 | 1346692.5 | 916000.25 | 1087383.62 | 902381.31 | 923932.38 | 1156488.62 | 1110957.12 | 903614.06 | 967129.5 |
| Q60597 | Ogdh | 2-oxoglutarate dehydrogenase, mitochondrial Mus musculus | 1003.28 | 23.66 | 116449 | 18 | 18 | 27483485 | 22267293 | 26240773 | 22031441 | 23816069 | 22223819 | 22136785 | 24211629 | 25370947 | 22805409 | 26591155 | 24583419 | 26639391 | 25966529 |
| Q3U0K8 | Ogfod1 | Prolyl 3-hydroxylase OGFOD1 Mus musculus | 117.14 | 9.54 | 62734 | 3 | 3 | 247794.14 | 409292.31 | 1993545.75 | 931927.94 | 465156.06 | 1160925.5 | 1490746.25 | 247089.09 | 195753.55 | 419879.31 | 331108.75 | 1439742.38 | 545161.44 | 502738.19 |
| Q99PG2 | Ogfr | Opioid growth factor receptor Mus musculus | 388.28 | 21.48 | 70679 | 7 | 7 | 2979361.75 | 4873649 | 4787515 | 4835608 | 3108700 | 3350059.5 | 2978024 | 2028704.38 | 3186623.75 | 3961944.5 | 2184625 | 3021672.75 | 3674001.5 | 2406048.25 |
| Q8CGY8 | Ogt | UDP-N-acetylglucosamine--peptide N-acetylglucosaminyltransferase 110 kDa subunit Mus | 734.49 | 17.69 | 116952 | 13 | 13 | 6992855.5 | 9327094 | 9089789 | 8556394 | 11275407 | 9635991 | 10924863 | 11679026 | 9252339 | 11602447 | 10585478 | 13942679 | 9673418 | 11641524 |
| Q9CZ30 | Ola1 | Obg-like ATPase 1 Mus musculus | 538.06 | 31.31 | 44730 | 8 | 8 | 24990359 | 24290319 | 25150643 | 25384167 | 26113907 | 23645711 | 22221783 | 28133973 | 25618341 | 24956979 | 23089645 | 29184197 | 25516235 | 23795251 |
| P58281 | Opa1 | Dynamin-like 120 kDa protein, mitochondrial Mus musculus | 176.73 | 5.73 | 111339 | 3 | 3 | 2297043.75 | 2142200.5 | 617528.69 | 1406200.75 | 1987594 | 1954232.38 | 1071433 | 2374934.75 | 2054423 | 1942067.38 | 1483824.25 | 695799.19 | 1411557.5 | 2379504.25 |
| Q60862 | Orc2 | Origin recognition complex subunit 2 Mus musculus | 41.47 | 4.17 | 65893 | 1 | 1 | 356100.53 | 312731.78 | 203458.05 | 250421.69 | 165191.72 | 231958.59 | 208496.92 | 1 | 1 | 132557.94 | 216900.22 | 262633.22 | 208392.55 | 240952.69 |
| Q5RKH6 | Os9 | Protein OS-9 Rattus norvegicus | 35.79 | 1.95 | 75365 | 1 | 1 | 625222.5 | 872828.5 | 781162.19 | 778855.0775 | 836207.12 | 752089.38 | 668432.88 | 629966.12 | 887788 | 823182.81 | 664714.31 | 841456.31 | 678550 | 819447.56 |
| Q3B7Z2 | Osbp | Oxysterol-binding protein 1 Mus musculus | 323 | 13.29 | 88797 | 7 | 7 | 3191671 | 4634675.5 | 6035773 | 4057666.5 | 6499011.5 | 6786330 | 5765942 | 5308943 | 6137013.5 | 5999007 | 6593737.5 | 5545120 | 5592966 | 4481377 |
| Q8CI95 | Osbpl11 | Oxysterol-binding protein-related protein 11 Mus musculus | 125.9 | 5.73 | 83629 | 3 | 3 | 903306.31 | 1133303.38 | 865538.25 | 1084914.75 | 1070425.62 | 987408.19 | 664239 | 1053821.62 | 932175.75 | 834913.75 | 920752 | 962107.12 | 1006001.44 | 937188.38 |
| Q8BX94 | Osbpl2 | Oxysterol-binding protein-related protein 2 Mus musculus | 84.85 | 6.82 | 55384 | 2 | 2 | 1280324.328 | 819243.31 | 1299020.62 | 942533.5 | 2060499.88 | 986355.25 | 1197713.12 | 1454300.75 | 2362420 | 1723252 | 1132197.38 | 1373261.25 | 2207980.75 | 2107237.5 |
| B9EJ86 | Osbpl8 | Oxysterol-binding protein-related protein 8 Mus musculus | 956.37 | 31.05 | 101269 | 18 | 18 | 42724825 | 33936841 | 33793157 | 26560865 | 40318105 | 47991073 | 36408233 | 35172501 | 41749253 | 40953581 | 41094217 | 41668185 | 52092365 | 49301957 |
| Q8BWU5 | Osgep | Probable tRNA N6-adenosine threonylcarbamoyltransferase Mus musculus | 196.87 | 13.43 | 36301 | 3 | 3 | 1212880.38 | 1635722.25 | 1653178.75 | 1559167 | 1533169.75 | 1509493.75 | 2099391 | 1746301.75 | 1596460.75 | 1559993.12 | 1497801 | 1771530.88 | 1640983.75 | 708252.19 |
| Q62422 | Ostf1 | Osteoclast-stimulating factor 1 Mus musculus | 340.08 | 38.6 | 23783 | 6 | 6 | 18533831 | 17938783 | 16404530 | 14253617 | 17516889 | 16275538 | 17658323 | 20619501 | 18271391 | 18299913 | 18837531 | 14642855 | 17682095 | 18688017 |
| B2RYG6 | Otub1 | Ubiquitin thioesterase OTUB1 Rattus norvegicus | 504.51 | 49.82 | 31270 | 9 | 9 | 21379525 | 21825843 | 20062337 | 18928377 | 18055105 | 20459555 | 17889337 | 25050149 | 23596537 | 19122511 | 19221759 | 22505615 | 21189799 | 21506407 |
| B2RRE7 | Otud4 | OTU domain-containing protein 4 Mus musculus | 52.92 | 0.72 | 123055 | 1 | 1 | 577512.62 | 870309.88 | 664846.75 | 937472.62 | 694400.5 | 546035.31 | 684608.19 | 656585.44 | 579327.44 | 596128.38 | 636578.81 | 761610.25 | 680768.38 | 702731.06 |
| Q8K2H2 | Otud6b | Deubiquitinase OTUD6B Mus musculus | 37.55 | 4.42 | 33758 | 1 | 1 | 1063783.563 | 913113.69 | 923101.06 | 1267940.75 | 1150978.75 | 953819 | 1011147.31 | 908278.06 | 1072234.62 | 890832.06 | 1129266.88 | 1290228.5 | 929058 | 1087133.5 |
| Q9D7E3 | Ovca2 | Esterase OVCA2 Mus musculus | 38.44 | 4.89 | 24246 | 1 | 1 | 1623532.033 | 1819773.25 | 1587083.38 | 1572344.25 | 1514927.25 | 1491165.62 | 1771940.5 | 1815157.88 | 1507409.62 | 1376377 | 1305901.12 | 1629916.75 | 1382473.25 | 1439430.373 |
| Q9D0K2 | Oxct1 | Succinyl-CoA:3-ketoacid coenzyme A transferase 1, mitochondrial Mus musculus | 1174.91 | 43.65 | 55989 | 15 | 15 | 38869481 | 50020709 | 50447477 | 51833617 | 52585401 | 50145901 | 57669321 | 50684585 | 56898329 | 61044753 | 61466453 | 64137069 | 67769793 | 56002905 |
| Q6P9R2 | Oxsr1 | Serine/threonine-protein kinase OSR1 Mus musculus | 314.35 | 15.56 | 58214 | 6 | 6 | 4060466.75 | 2940360 | 4116421.5 | 1372253.25 | 2363157.5 | 2415193 | 3713253 | 4304582 | 4061427.25 | 2594516.5 | 2448150.75 | 2525475.5 | 3210366.5 | 3364157.5 |
| P54001 | P4ha1 | Prolyl 4-hydroxylase subunit alpha-1 Rattus norvegicus | 390.79 | 18.54 | 60898 | 8 | 8 | 2903602.75 | 507346.91 | 1275662 | 2642445 | 1580002.75 | 1340572.38 | 1411442.38 | 1591628.815 | 1424436.75 | 2190063.75 | 1882320.75 | 1555707.25 | 929026.38 | 1527475.38 |
| P09103 | P4hb | Protein disulfide-isomerase Mus musculus | 1769.22 | 55.6 | 57058 | 28 | 28 | 136559073 | 139142049 | 125258673 | 127324713 | 149719969 | 156637729 | 165703089 | 171032881 | 160082321 | 169302545 | 150237809 | 138558721 | 150462081 | 148395649 |
| P50580 | Pa2g4 | Proliferation-associated protein 2G4 Mus musculus | 935.58 | 59.64 | 43699 | 18 | 18 | 38409153 | 32099041 | 39336401 | 40108437 | 35228189 | 32987921 | 42909673 | 33122483 | 33350223 | 39964081 | 40758321 | 35882305 | 38017221 | 35464989 |
| P29341 | Pabpc1 | Polyadenylate-binding protein 1 Mus musculus | 1665.87 | 60.69 | 70671 | 30 | 31 | 76494185 | 81910409 | 88746609 | 87175105 | 83188929 | 78413121 | 77602737 | 71740025 | 78670897 | 83481521 | 78455009 | 82834569 | 84055025 | 82088033 |
| Q8CCS6 | Pabpn1 | Polyadenylate-binding protein 2 Mus musculus | 218.4 | 27.15 | 32297 | 4 | 4 | 10433612 | 11457711 | 10696737 | 11764491 | 12181833 | 10674895 | 10735051 | 10170512 | 10438958 | 10139141 | 11254448 | 11113864 | 7045748 | 10794495 |
| Q9WVE8 | Pacsin2 | Protein kinase C and casein kinase substrate in neurons protein 2 Mus musculus | 267.12 | 16.87 | 55833 | 7 | 7 | 8856287 | 9064631 | 9657616 | 8117228 | 8317619 | 10108433 | 8660341 | 10682041 | 9050735 | 10516920 | 8488284 | 8503720 | 9336183 | 9030766 |
| Q4V886 | Paf1 | RNA polymerase II-associated factor 1 homolog Rattus norvegicus | 43.27 | 2.06 | 60546 | 1 | 1 | 287980.97 | 360996.91 | 289336.97 | 1 | 1 | 290797.44 | 248594.8 | 381320.09 | 316527.84 | 245833.38 | 280336.94 | 312942.56 | 309819.84 | 301033.1133 |
| P63004 | Pafah1b1 | Platelet-activating factor acetylhydrolase IB subunit alpha Rattus norvegicus | 532.77 | 29.51 | 46670 | 11 | 11 | 15892587 | 9029093 | 15389069 | 14904659 | 13506808 | 10726711 | 15077545 | 12962993 | 13392445 | 15004884 | 14506597 | 17053501 | 13570629 | 13582161 |
| O35264 | Pafah1b2 | Platelet-activating factor acetylhydrolase IB subunit beta Rattus norvegicus | 136.06 | 12.23 | 25581 | 2 | 2 | 2320830.5 | 1351602 | 1813642.62 | 1926399 | 1 | 2022298.88 | 1638564.38 | 3029142.25 | 2225802 | 1453605.38 | 2205914.75 | 1839028.5 | 1843406.75 | 1694010.75 |
| Q9DCL9 | Paics | Multifunctional protein ADE2 Mus musculus | 1061.37 | 48.71 | 47006 | 18 | 18 | 18406545 | 19401191 | 24465001 | 22549427 | 24582029 | 22667763 | 20918139 | 19451783 | 20334237 | 22140409 | 26830319 | 26148601 | 25532079 | 26142395 |
| Q8VE62 | Paip1 | Polyadenylate-binding protein-interacting protein 1 Mus musculus | 38.81 | 2.75 | 45702 | 1 | 1 | 1277745.5 | 1062720.25 | 1153127.12 | 1012264.56 | 391925.97 | 393025.16 | 1028063.19 | 928388.62 | 1078223.5 | 1195693.12 | 1154474.75 | 1054808.25 | 1320462.88 | 1176581.96 |

| | | | | | | | | | | | | | | | | | | | | | |
|---|---|---|---|---|---|---|---|---|---|---|---|---|---|---|---|---|---|---|---|---|---|
| Q9DCE5 | Pak1ip1 | p21-activated protein kinase-interacting protein 1 Mus musculus | 269.41 | 20.42 | 42116 | 6 | 6 | 1412074.38 | 1034778.94 | 1321317.75 | 1364280.5 | 1623268.38 | 885734.2325 | 775390.81 | 1152813.12 | 849503.69 | 765229.31 | 1047729.75 | 756952 | 788854.88 | 1071687.62 |
| Q8CIN4 | Pak2 | Serine/threonine-protein kinase PAK 2 Mus musculus | 756.6 | 32.82 | 57930 | 9 | 13 | 13375337 | 8896386 | 13045911 | 10489531 | 11003372 | 13199163 | 14499667 | 11627142 | 12303631 | 12416514 | 13348858 | 13207906 | 11706030 | 12164570 |
| Q61036 | Pak3 | Serine/threonine-protein kinase PAK 3 Mus musculus | 343.94 | 15.56 | 62398 | 2 | 6 | 3598529.5 | 3919888.25 | 3638312.25 | 3730203 | 3620463.5 | 3811791.5 | 3193423 | 3155009.75 | 2610567.5 | 2838928.25 | 3397802.5 | 3394999.5 | 3328516.25 | 3149755 |
| Q9ET54 | Palld | Palladin Mus musculus | 79.78 | 3.12 | 152131 | 2 | 2 | 1 | 1 | 210994.59 | 212372.02 | 108996.18 | 222332.17 | 187968.08 | 120773.92 | 243587.61 | 248268.58 | 212616.64 | 197933.48 | 226787.33 | 167814.16 |
| Q8K4K6 | Pank1 | Pantothenate kinase 1 Mus musculus | 50.91 | 1.64 | 60091 | 1 | 1 | 681330.06 | 668368.5 | 897217.38 | 723368.625 | 646558.56 | 687092.62 | 709307.94 | 676458.56 | 565644.12 | 633865.06 | 845546.2733 | 966797.5 | 889123.94 | 680717.38 |
| Q80YV4 | Pank4 | Pantothenate kinase 4 Mus musculus | 83.69 | 5.12 | 91522 | 2 | 2 | 390686.81 | 509115.09 | 599090.44 | 1 | 1 | 663642.25 | 544814.06 | 148922.69 | 792390.12 | 485302.31 | 600946.5567 | 189964.42 | 578472.06 | 1034403.19 |
| Q60967 | Papss1 | Bifunctional 3'-phosphoadenosine 5'-phosphosulfate synthase 1 Mus musculus | 327.46 | 17.31 | 70794 | 7 | 7 | 3402638.5 | 5502268 | 5422769.5 | 4365005.5 | 5463396.5 | 5452476.5 | 4524866 | 5945372 | 6868583 | 3438975.75 | 6227615 | 3111323.75 | 4840378 | 4027532.25 |
| Q99LX0 | Park7 | Protein/nucleic acid deglycase DJ-1 Mus musculus | 623.23 | 76.19 | 20021 | 10 | 10 | 39469889 | 48986005 | 47944301 | 48110437 | 49732665 | 50031037 | 60516577 | 55527521 | 58796837 | 62060417 | 48712385 | 57808505 | 51117441 | 58712413 |
| Q5XJY4 | Parl | Presenilins-associated rhomboid-like protein, mitochondrial Mus musculus | 131.53 | 10.08 | 41964 | 2 | 2 | 1511375.75 | 853155.31 | 1231294.75 | 373937.16 | 1110664.88 | 1469106.5 | 1551110.75 | 887767.88 | 1531794.38 | 1626891.5 | 1155150.88 | 817024 | 2004750.12 | 1659277.25 |
| Q8VDG3 | Parn | Poly(A)-specific ribonuclease PARN Mus musculus | 55.84 | 2.88 | 71559 | 1 | 1 | 317543.69 | 526220.81 | 517402.84 | 359931.91 | 392415.41 | 1186148.5 | 263202.16 | 1824238.25 | 733327.38 | 475524.44 | 466945.59 | 446275.34 | 348429.59 | 308358.94 |
| P11103 | Parp1 | Poly [ADP-ribose] polymerase 1 Mus musculus | 1111.99 | 23 | 113100 | 4 | 17 | 23729741 | 29371337 | 31731749 | 23119535 | 26062539 | 23210453 | 29436919 | 27465751 | 26311997 | 27620203 | 29747363 | 30226277 | 28431843 | 31500735 |
| Q8BZ20 | Parp12 | Poly [ADP-ribose] polymerase 12 Mus musculus | 125.95 | 5.2 | 79917 | 2 | 2 | 1 | 1 | 1 | 1 | 1 | 296647.56 | 1 | 1009868.62 | 2431780 | 1 | 1 | 845868.62 | 1 | 1 |
| Q2EMV9 | Parp14 | Poly [ADP-ribose] polymerase 14 Mus musculus | 321.01 | 6.05 | 203802 | 7 | 7 | 2628801.75 | 1012310.38 | 2278531 | 1326686.62 | 1773543.5 | 993230.12 | 842959.62 | 1425405.5 | 1243292.38 | 820152.25 | 931966.62 | 1400419.25 | 942447.5 | 1191223.38 |
| Q9ES46 | Parvb | Beta-parvin Mus musculus | 161.49 | 15.34 | 41669 | 4 | 4 | 429852 | 248073.16 | 193111.55 | 316781.56 | 451728.94 | 199306.06 | 203186.7 | 946598.12 | 400356 | 120129.2 | 146303.89 | 178032 | 449716.12 | 258017.3367 |
| Q9ERD8 | Parvg | Gamma-parvin Mus musculus | 81.35 | 9.06 | 37603 | 2 | 2 | 2437660.25 | 1708778.12 | 1937751.75 | 1990288.88 | 2465646.5 | 1993277.12 | 2053304.75 | 2451371.5 | 3221693.25 | 2334232.5 | 2251987.5 | 2297537.75 | 2352619 | 1892727.62 |
| Q9D0B6 | Pbdc1 | Protein PBDC1 Mus musculus | 279.99 | 30.3 | 22223 | 5 | 5 | 9506809 | 10912057 | 10215348 | 11892442 | 9902948 | 9426651 | 8767917 | 11630039 | 9934905 | 9708075 | 11399608 | 9090833 | 10373302 | 10671328 |
| Q9JJ78 | Pbk | Lymphokine-activated killer T-cell-originated protein kinase Mus musculus | 44.52 | 3.94 | 36745 | 1 | 1 | 480654.28 | 856266.69 | 746374.75 | 786979.43 | 1064622 | 801450.75 | 729662.8925 | 456087.69 | 813424.38 | 847688.75 | 999383.5 | 790583.12 | 1019542.19 | 1347686.25 |
| Q8BSQ9 | Pbrm1 | Protein polybromo-1 Mus musculus | 69.92 | 1.1 | 187188 | 1 | 1 | 245477.16 | 297227.59 | 250973.81 | 203114.95 | 210653.03 | 237650.02 | 227430.3 | 204086.95 | 301044.19 | 238992.36 | 278787.28 | 346307.94 | 268619.88 | 269346.84 |
| Q3TVI8 | Pbxip1 | Pre-B-cell leukemia transcription factor-interacting protein 1 Mus musculus | 354.97 | 8.8 | 81164 | 5 | 5 | 7381188.5 | 3296212.5 | 4588834.5 | 5065322 | 6231069.5 | 4756281 | 4651384 | 4875728 | 3945602 | 4470101 | 6367160.5 | 2532751.5 | 4506748.5 | 4748029.5 |
| P60335 | Pcbp1 | Poly(rC)-binding protein 1 Mus musculus | 1103.32 | 72.19 | 37498 | 12 | 16 | 126769065 | 134962737 | 139234417 | 141957777 | 141860001 | 134154201 | 130866577 | 118116897 | 129961921 | 148145601 | 146217121 | 141242977 | 137270225 | 144026769 |
| Q61990 | Pcbp2 | Poly(rC)-binding protein 2 Mus musculus | 925.26 | 72.65 | 38222 | 13 | 16 | 81268449 | 84730753 | 87683425 | 85525457 | 86003377 | 85112265 | 83698657 | 80047561 | 83424617 | 89463665 | 85049673 | 85003321 | 84477081 | 83489833 |
| Q91ZA3 | Pcca | Propionyl-CoA carboxylase alpha chain, mitochondrial Mus musculus | 284.25 | 11.05 | 79922 | 5 | 5 | 3053064 | 3140877.5 | 3337323 | 3037756.5 | 3906972.5 | 3050131.75 | 3374248.5 | 3377716.5 | 3893225.25 | 2556108.25 | 3473455.5 | 3004914.25 | 3384318 | 3552020.5 |
| Q99MN9 | Pccb | Propionyl-CoA carboxylase beta chain, mitochondrial Mus musculus | 479.37 | 22.18 | 58409 | 8 | 8 | 10446075 | 9421685 | 8901953 | 10852820 | 9400124 | 10621293 | 8672009 | 9441719 | 11663810 | 12565391 | 9248659 | 8855995 | 9820914 | 8888671 |
| Q8BFV2 | Pcid2 | PCI domain-containing protein 2 Mus musculus | 113.78 | 8.77 | 46132 | 2 | 2 | 474814.62 | 199473.06 | 396730.5275 | 452961.81 | 459672.62 | 1 | 1 | 1 | 1 | 1 | 299287.41 | 278962.2133 | 252704.23 | 284895 |
| P59114 | Pcif1 | Phosphorylated CTD-interacting factor 1 Mus musculus | 85.52 | 4.11 | 80504 | 2 | 2 | 453812.1875 | 478226.53 | 461242.25 | 408870.56 | 466909.41 | 542656.19 | 432438.09 | 540887.56 | 491491.81 | 420749.69 | 527677.44 | 487623.9267 | 385052.72 | 550141.62 |
| Q8BH04 | Pck2 | Phosphoenolpyruvate carboxykinase [GTP], mitochondrial Mus musculus | 1132.38 | 40.47 | 70528 | 19 | 19 | 44999569 | 51270049 | 49064105 | 55250529 | 52849193 | 58124393 | 55342169 | 56984741 | 56601573 | 53124605 | 54328641 | 46346685 | 47772705 | 54078461 |
| P17918 | Pcna | Proliferating cell nuclear antigen Mus musculus | 1091 | 85.06 | 28785 | 16 | 17 | 197016353 | 229116033 | 214982929 | 200755601 | 222280225 | 191110977 | 184421249 | 172244305 | 184241985 | 198398881 | 200986817 | 196086929 | 221462929 | 210013921 |
| Q9CQF9 | Pcyox1 | Prenylcysteine oxidase Mus musculus | 382.06 | 23.76 | 56495 | 7 | 7 | 15368321 | 16023936 | 15011778 | 10549521 | 14543288 | 14830235 | 15360311 | 15673620 | 15067068 | 13879007 | 13445220 | 14010688 | 14904174 | 15517515 |
| Q8C7K6 | Pcyox1l | Prenylcysteine oxidase-like Mus musculus | 121.79 | 6.87 | 54875 | 3 | 3 | 4274521 | 1098993.75 | 7082154 | 5048351 | 3787326 | 4363406 | 5986186 | 5045521.5 | 5011186 | 5025718 | 4971448.5 | 2988064 | 4579401.5 | 3681372.5 |
| Q922E4 | Pcyt2 | Ethanolamine-phosphate cytidylyltransferase Mus musculus | 202.85 | 12.62 | 45235 | 4 | 4 | 13041319 | 10575002 | 12158069 | 11861909 | 9189696 | 11368877 | 9752674 | 11020215 | 11249509 | 11083007 | 10500773 | 11952353 | 12113160 | 11814329 |
| Q3UHX2 | Pdap1 | 28 kDa heat- and acid-stable phosphoprotein Mus musculus | 275.21 | 33.15 | 20605 | 5 | 5 | 5561523.5 | 6001897.5 | 3730673.5 | 6465843.5 | 5753828 | 5319480.5 | 6562209.5 | 5318257.5 | 2993068.5 | 4549466 | 5362915 | 5388062.5 | 3868453 | 5880596 |
| Q6NX65 | Pdcd10 | Programmed cell death protein 10 Rattus norvegicus 2 | 102.98 | 10.48 | 24355 | 2 | 2 | 5014375 | 1446141.75 | 3302033.75 | 1264186.88 | 2664207.75 | 2910460 | 3608838.5 | 3828375 | 3305194.5 | 3601069.5 | 3207218.25 | 3217725.75 | 2641006.5 | 1501748.88 |
| Q6NS46 | Pdcd11 | Protein RRP5 homolog Mus musculus | 1027.78 | 18.37 | 207779 | 21 | 21 | 11391401 | 10210932 | 14812261 | 10521552 | 6393869 | 14416254 | 13107780 | 8677479 | 17502171 | 13274619 | 16876975 | 27220917 | 10655440 | 19133991 |
| P56812 | Pdcd5 | Programmed cell death protein 5 Mus musculus | 122.84 | 19.84 | 14275 | 2 | 2 | 2714823 | 1559986 | 2663629.25 | 3307334.5 | 2622942.5 | 2637647.5 | 3302508.5 | 2576310 | 2099393 | 2380963.5 | 2839351.25 | 2595980.75 | 2556114 | 2837056 |
| P12815 | Pdcd6 | Programmed cell death protein 6 Mus musculus | 331.17 | 47.12 | 21867 | 6 | 6 | 5503205 | 6544410 | 5117478.5 | 4584561 | 7744203 | 4125307.5 | 6789544.5 | 4888707 | 7109188 | 4790463 | 4842126.5 | 4872136 | 5700339 | 5402719.5 |
| Q9WU78 | Pdcd6ip | Programmed cell death 6-interacting protein Mus musculus | 1562.22 | 41.2 | 96024 | 25 | 25 | 45678109 | 52402553 | 50640681 | 49440153 | 49380501 | 52804869 | 54905465 | 56555949 | 53051033 | 54614657 | 51647509 | 52123933 | 50429949 | 54272649 |
| Q8BVF2 | Pdcl3 | Phosducin-like protein 3 Mus musculus | 303.36 | 32.08 | 27581 | 6 | 6 | 4534774 | 4207844.5 | 4094438 | 4482828.5 | 4854951 | 5378127 | 3399928.5 | 3196745 | 4615747.5 | 5216049.5 | 6264950 | 5809101 | 4742143 | 4962010 |
| Q3TIU4 | Pde12 | 2',5'-phosphodiesterase 12 Mus musculus | 249.54 | 9.38 | 67514 | 4 | 4 | 3170236 | 3794616.75 | 3080938.25 | 3403213 | 2253006.5 | 4059441 | 3205665.25 | 3649739.75 | 3225305.75 | 3549109.75 | 3546123 | 3555978 | 3052486 | 3443183.5 |
| O55057 | Pde6d | Retinal rod rhodopsin-sensitive cGMP 3',5'-cyclic phosphodiesterase subunit delta Mus mu | 41.04 | 8 | 17348 | 1 | 1 | 598764.94 | 946548.19 | 794202.75 | 870121.38 | 681159.19 | 814569.88 | 632900.62 | 899282.69 | 758092.88 | 635369.56 | 720482.25 | 911188.12 | 611734.69 | 770106.94 |
| Q9D051 | Pdhb | Pyruvate dehydrogenase E1 component subunit beta, mitochondrial Mus musculus | 459.56 | 37.88 | 38937 | 10 | 10 | 11917652 | 15018408 | 13561663 | 12633189 | 14306156 | 13304927 | 14013339 | 14155785 | 14159487 | 14643458 | 14588635 | 14672301 | 14447765 | 14095821 |
| Q8BKZ9 | Pdhx | Pyruvate dehydrogenase protein X component, mitochondrial Mus musculus | 164.94 | 8.38 | 53999 | 4 | 4 | 5800042.5 | 6961759.5 | 6824678 | 5991619 | 6342486 | 5368451.5 | 5565463 | 6827628.5 | 6669766 | 6989225.5 | 6779359 | 6362488 | 4920058 | 6646719 |
| P27773 | Pdia3 | Protein disulfide-isomerase A3 Mus musculus | 1767.9 | 60.2 | 56678 | 30 | 30 | 254216289 | 217690513 | 247481441 | 249726273 | 236587809 | 246538001 | 266653649 | 251374081 | 263613041 | 273817761 | 255023905 | 256135217 | 266005025 | 255093329 |
| P08003 | Pdia4 | Protein disulfide-isomerase A4 Mus musculus | 1869.14 | 53.61 | 71982 | 32 | 32 | 53638141 | 41823233 | 54634405 | 44436601 | 43733833 | 54383121 | 50608161 | 55512393 | 58090617 | 57235533 | 51464717 | 52088665 | 58965077 | 40865601 |
| Q922R8 | Pdia6 | Protein disulfide-isomerase A6 Mus musculus | 940.22 | 42.05 | 48100 | 15 | 15 | 100966177 | 86687601 | 92713761 | 90096721 | 83267161 | 88499825 | 101802177 | 104221425 | 101426121 | 98075553 | 97797361 | 97391745 | 98842433 | 85811265 |
| Q63065 | Pdk1 | [Pyruvate dehydrogenase (acetyl-transferring)] kinase isozyme 1, mitochondrial Rattus nor | 190.83 | 11.52 | 49081 | 3 | 3 | 2626979.5 | 3395204 | 1657055.38 | 2282470.5 | 1776084.12 | 3311625.5 | 3405527.5 | 4138147.5 | 3783875.5 | 2302031.5 | 1983502.25 | 3500898.5 | 3243433.5 | 3306275 |
| Q922H2 | Pdk3 | [Pyruvate dehydrogenase (acetyl-transferring)] kinase isozyme 3, mitochondrial Mus musc | 309.05 | 13.49 | 47923 | 4 | 4 | 4694207 | 4650144 | 4087162.75 | 3807998.25 | 4096631 | 3367731.75 | 4074672.25 | 4325741 | 4019888 | 2377792.25 | 4450910 | 4214942.5 | 4274187.5 | 2233876.75 |
| Q8CI51 | Pdlim5 | PDZ and LIM domain protein 5 Mus musculus | 353.4 | 16.41 | 63299 | 7 | 7 | 2167191 | 1759710.75 | 2945875.5 | 1207917.75 | 1192635.38 | 2567303 | 2926862 | 1786632.75 | 2671699 | 2910702 | 1496413.5 | 2058604.75 | 1545732.12 | 1599934.88 |
| A4L9P7 | Pds5a | Sister chromatid cohesion protein PDS5 homolog A Rattus norvegicus | 296.95 | 4.95 | 150286 | 4 | 5 | 3646718.5 | 2701575 | 2707299.5 | 2592853 | 3371456.25 | 3785733.75 | 2453242 | 2937550.25 | 3463306.5 | 3232530.25 | 3803409.25 | 3347367.25 | 3298367.25 | 3430849.75 |
| Q4VA53 | Pds5b | Sister chromatid cohesion protein PDS5 homolog B Mus musculus | 121.11 | 1.66 | 164419 | 1 | 2 | 1654523.75 | 1661654.88 | 1050859.38 | 1462088.25 | 2150609.5 | 2072384.25 | 1388708.12 | 1423165.38 | 1264697.88 | 1378517.5 | 1967273.75 | 1425276.5 | 1585256.62 | 1696590.88 |
| Q99K01 | Pdxdc1 | Pyridoxal-dependent decarboxylase domain-containing protein 1 Mus musculus | 546.27 | 19.31 | 87335 | 11 | 11 | 5451270.5 | 4473760.5 | 5188518 | 6987475 | 8213314 | 8818196 | 6469685.5 | 4624108 | 4735962.5 | 7409180 | 6822929.5 | 4297387.5 | 7450319.5 | 6846438.5 |
| Q8K183 | Pdxk | Pyridoxal kinase Mus musculus | 664.85 | 28.53 | 35015 | 9 | 11 | 15056884 | 15631310 | 21250323 | 17656895 | 14409173 | 19347319 | 16786881 | 18716019 | 19373221 | 18313855 | 19143891 | 19173559 | 20205989 | 16454767 |
| Q62048 | Pea15 | Astrocytic phosphoprotein PEA-15 Mus musculus | 285.53 | 56.15 | 15054 | 6 | 6 | 1709723.25 | 27973147 | 20755053 | 16688222 | 19577303 | 24549165 | 20618633 | 26528901 | 20007095 | 17322009 | 16783831 | 1499900.75 | 22832849 | 25311245 |
| Q69Z38 | Peak1 | Inactive tyrosine-protein kinase PEAK1 Mus musculus | 91.36 | 1.56 | 191097 | 2 | 2 | 693472.88 | 723881.19 | 1 | 1 | 534121.5 | 522005.12 | 625132.88 | 1 | 1 | 535928.69 | 651603 | 581945 | 528227.56 | 566004.44 |
| P70296 | Pebp1 | Phosphatidylethanolamine-binding protein 1 Mus musculus | 293.22 | 44.39 | 20830 | 5 | 5 | 9264044 | 9414791 | 8561913 | 9974959 | 11958726 | 9294026 | 10663440 | 9665325 | 9188848 | 17048857 | 10833777 | 10718981 | 9765320 | 11146075 |
| Q99MZ7 | Pecr | Peroxisomal trans-2-enoyl-CoA reductase Mus musculus | 78.69 | 2.97 | 32410 | 1 | 1 | 1798545.25 | 1351021.75 | 1489071.88 | 1 | 1326310.88 | 1311803.12 | 1483415.62 | 1290883.75 | 1092821.5 | 1234741.25 | 1255136.75 | 1446516.38 | 1267069.62 | 1239558.38 |
| Q8BFY6 | Pef1 | Peflin Mus musculus | 177.9 | 17.45 | 29228 | 4 | 4 | 685249.44 | 1989509 | 3559292.5 | 3504781.5 | 3390456 | 4077110 | 4296750 | 4633742.5 | 4642793 | 2693156.25 | 2504171.5 | 3621043.25 | 4018215.75 | 3003892.5 |
| Q5XIP1 | Pelo | Protein pelota homolog Rattus norvegicus | 47.91 | 4.68 | 43407 | 1 | 1 | 472490.75 | 613910.62 | 1 | 1 | 286769.41 | 216182.27 | 483545.66 | 396186.12 | 317663.7625 | 174741 | 1 | 1 | 197628.22 | 418388.25 |
| Q9DBD5 | Pelp1 | Proline-, glutamic acid- and leucine-rich protein 1 Mus musculus | 331.84 | 4.81 | 118069 | 5 | 5 | 3186774 | 4325926 | 4950494 | 5275342.5 | 3836460.5 | 3147237.25 | 4043144.25 | 4085233 | 3160691.75 | 3081350.25 | 3609782 | 3902025 | 3761455.5 | 3394870.25 |
| Q11136 | Pepd | Xaa-Pro dipeptidase Mus musculus | 711.8 | 34.48 | 55029 | 15 | 15 | 33498953 | 32399651 | 30950619 | 29717185 | 25378425 | 24394553 | 28371033 | 30916511 | 25668675 | 22774377 | 23717367 | 24057411 | 26170783 | 28951187 |
| Q9EQ61 | Pes1 | Pescadillo homolog Mus musculus | 723.77 | 29.45 | 67796 | 16 | 16 | 9817133 | 8869289 | 7449325 | 7575934 | 7898886.5 | 11046548 | 9115557 | 8433276 | 7811292 | 8443398 | 9006141 | 9957600 | 8724138 | 9442241 |
| Q8VCI5 | Pex19 | Peroxisomal biogenesis factor 19 Mus musculus | 53.67 | 9.7 | 32733 | 1 | 1 | 170118.98 | 1 | 228473.72 | 1 | 181924.33 | 127222.84 | 192358.89 | 127895.14 | 160630.6225 | 195045.62 | 893419.25 | 186740.52 | 140602.98 | 183359.27 |
| Q5SUR0 | Pfas | Phosphoribosylformylglycinamidine synthase Mus musculus | 1320.05 | 22.51 | 144629 | 19 | 19 | 22435121 | 30871269 | 25488663 | 34036005 | 26040129 | 26187175 | 26693277 | 26560603 | 22629775 | 29075417 | 30019709 | 31270883 | 25521595 | 29741411 |
| Q9CWM4 | Pfdn1 | Prefoldin subunit 1 Mus musculus | 61.53 | 9.02 | 14255 | 1 | 1 | 2782856.25 | 2875558.25 | 2645507 | 3131745.75 | 2722431 | 2375914.25 | 2936895.25 | 2850401.75 | 2175852 | 2912861.75 | 2803241.5 | 2551755.75 | 2177812.75 | 2062346.25 |
| B0BN18 | Pfdn2 | Prefoldin subunit 2 Rattus norvegicus | 233.75 | 29.22 | 16580 | 4 | 4 | 12981779 | 15286921 | 14376549 | 16331211 | 13527823 | 10710282 | 13436655 | 14911134 | 12632037 | 13818798 | 13273148 | 12899699 | 11883926 | 12446080 |
| Q9WU28 | Pfdn5 | Prefoldin subunit 5 Mus musculus | 227.71 | 33.12 | 17356 | 3 | 3 | 2941656 | 1710702.12 | 3679333.5 | 1357294.38 | 1450175 | 3270239 | 1877216.75 | 5091373 | 3449292.75 | 1961493.88 | 2462581.25 | 3546765.5 | 3479719 | 2700549 |
| Q03958 | Pfdn6 | Prefoldin subunit 6 Mus musculus | 58.42 | 9.45 | 14455 | 1 | 1 | 1665957 | 1816728.12 | 1824478.53 | 2297962.75 | 1517266.25 | 2108387.75 | 1604206.22 | 1314906 | 1608052.75 | 1385478.38 | 1798620.5 | 1429020.75 | 1175958.75 | 1641820.75 |
| P12382 | Pfkl | ATP-dependent 6-phosphofructokinase, liver type Mus musculus | 576.84 | 22.82 | 85360 | 10 | 12 | 19641895 | 16978521 | 16034875 | 13188097 | 19513529 | 20625585 | 20757869 | 18619079 | 20318077 | 19687081 | 18411935 | 18390033 | 17670297 | 18232371 |
| P47857 | Pfkm | ATP-dependent 6-phosphofructokinase, muscle type Mus musculus | 106.5 | 3.85 | 85269 | 2 | 3 | 1968725.25 | 1618983.75 | 1741868.75 | 1884924 | 2017922.5 | 1614429.75 | 1805808 | 1634212.25 | 1698414.25 | 1382708.12 | 1854450.25 | 2029411 | 1967192 | 2030240 |
| Q9WUA3 | Pfkp | ATP-dependent 6-phosphofructokinase, platelet type Mus musculus | 1812.23 | 54.97 | 85455 | 20 | 27 | 62324449 | 64117569 | 62208097 | 54761193 | 61724577 | 73057633 | 78252481 | 84109385 | 84470713 | 83289881 | 67478985 | 61805585 | 67647865 | 64898885 |
| P62962 | Pfn1 | Profilin-1 Mus musculus | 1060.25 | 94.29 | 14957 | 13 | 17 | 698839361 | 817273217 | 818383169 | 799125441 | 853652097 | 772295297 | 781287553 | 758317441 | 796042369 | 832268481 | 829524801 | 727057345 | 832046721 | 892432193 |
| P25113 | Pgam1 | Phosphoglycerate mutase 1 Rattus norvegicus | 1137.31 | 74.02 | 28832 | 15 | 16 | 160273809 | 158861713 | 135597505 | 130547473 | 140905953 | 154864753 | 164400177 | 169627249 | 151610225 | 185304705 | 151290689 | 131547873 | 148156353 | 145995905 |
| Q9DCD0 | Pgd | 6-phosphogluconate dehydrogenase, decarboxylating Mus musculus | 1812.8 | 69.57 | 53247 | 26 | 27 | 151253345 | 164736529 | 175545713 | 136175393 | 153375793 | 178058113 | 165630561 | 182952209 | 192429441 | 181907857 | 181808641 | 192496833 | 207364001 | 189192449 |
| Q8BP56 | Pgghg | Protein-glucosylgalactosylhydroxylysine glucosidase Mus musculus | 52.99 | 2.9 | 76488 | 1 | 1 | 323060.22 | 299136.78 | 216324.42 | 289850.87 | 320882.06 | 223651.34 | 223149.4575 | 280906.44 | 188298.55 | 199741.5 | 253926.83 | 299526.19 | 288431.94 | 233332.39 |
| P09411 | Pgk1 | Phosphoglycerate kinase 1 Mus musculus | 2512.98 | 79.14 | 44550 | 26 | 36 | 841446529 | 935717121 | 922474369 | 795992001 | 906236481 | 962210369 | 1057739841 | 1045803009 | 1095603713 | 1082488833 | 908902721 | 891912129 | 997301121 | 917143809 |
| P09041 | Pgk2 | Phosphoglycerate kinase 2 Mus musculus | 505 | 17.03 | 44853 | 1 | 8 | 44837493 | 39106101 | 49576689 | 51804593 | 54221653 | 43042297 | 52786685 | 38371953 | 52618081 | 68422129 | 48575717 | 53193177 | 55042201 | 50009693 |
| Q9CQ60 | Pgls | 6-phosphogluconolactonase Mus musculus | 557.62 | 59.53 | 27254 | 8 | 8 | 49917609 | 63425085 | 49683365 | 18717755 | 50748305 | 37633125 | 45711501 | 52796177 | 53498265 | 53213377 | 59142361 | 56490525 | 48854265 | 61793541 |
| Q9D0F9 | Pgm1 | Phosphoglucomutase-1 Mus musculus | 1225.89 | 48.04 | 61418 | 18 | 18 | 24706457 | 27459615 | 26923959 | 22012243 | 24741713 | 27569583 | 28038787 | 33208803 | 32010021 | 28321793 | 25857931 | 26726955 | 25721051 | 24685767 |
| Q7TSV4 | Pgm2 | Phosphoglucomutase-2 Mus musculus | 85 | 7.74 | 68748 | 2 | 2 | 2201647.75 | 1594923.38 | 2156999.25 | 2122491.75 | 2083694 | 1682543.62 | 1948690.25 | 1 | 1 | 2703991.5 | 1659131.5 | 1415512 | 796951.12 | 846634.75 |
| Q9CYR6 | Pgm3 | Phosphoacetylglucosamine mutase Mus musculus | 58.22 | 2.58 | 59453 | 1 | 1 | 2303533 | 2253810.5 | 1989791.62 | 2480269.25 | 2497793.5 | 1928146.25 | 1921476.75 | 2051431.25 | 2109985.5 | 2539111.75 | 2007583.12 | 2046845.25 | 2228019.75 | 2257952.5 |
| Q8CHP8 | Pgp | Glycerol-3-phosphate phosphatase Mus musculus | 426.81 | 41.74 | 34541 | 9 | 9 | 19739845 | 10070202 | 22113225 | 7777014.5 | 9715953 | 16929357 | 9380716 | 17482605 | 17792749 | 12647743 | 19929063 | 21129741 | 20637713 | 10288825 |
| O55022 | Pgrmc1 | Membrane-associated progesterone receptor component 1 Mus musculus | 61.64 | 7.18 | 21694 | 1 | 1 | 2714451.25 | 2481273 | 3081889.75 | 3019477.25 | 3099087 | 3229726 | 3145999.75 | 3554294.75 | 3111369.5 | 3296737.25 | 3338519.25 | 2951335 | 3363885.5 | 3229840.5 |
| Q80UU9 | Pgrmc2 | Membrane-associated progesterone receptor component 2 Mus musculus | 254.4 | 21.66 | 23334 | 3 | 3 | 3782386.25 | 3627643.5 | 3018508.25 | 3366951 | 3852169.25 | 2891702.5 | 3128325.5 | 3353998 | 3145518.5 | 2981857.5 | 3568281.5 | 3393729 | 3498896.5 | 3482859.75 |
| Q8BHF7 | Pgs1 | CDP-diacylglycerol--glycerol-3-phosphate 3-phosphatidyltransferase, mitochondrial Mus m | 83.58 | 5.61 | 62489 | 2 | 2 | 679683.38 | 826181.75 | 691918.06 | 720012.625 | 682267.31 | 722155.94 | 666672 | 800632.88 | 667628 | 767313.06 | 890632.62 | 628269.31 | 728876.44 | 699819.38 |
| Q63068 | Phax | Phosphorylated adapter RNA export protein Rattus norvegicus | 131.34 | 5.71 | 43166 | 2 | 2 | 2160803.25 | 578286.62 | 1998157 | 710178.44 | 825671 | 1850391.5 | 1777817.75 | 1849843.88 | 1714909 | 2064228.5 | 1702469.5 | 2126036.75 | 2076473.12 | 2192569.5 |
| P67778 | Phb | Prohibitin Mus musculus | 1423.85 | 81.62 | 29820 | 24 | 24 | 189354209 | 196653537 | 199541985 | 203101233 | 193904209 | 206498305 | 212427409 | 212689233 | 194278273 | 194981793 | 195015009 | 196413953 | 200555137 | 202703169 |
| O35129 | Phb2 | Prohibitin-2 Mus musculus | 910.13 | 53.18 | 33296 | 14 | 14 | 28280971 | 23675733 | 30372779 | 34590209 | 32233809 | 30432163 | 23745145 | 18123451 | 28453055 | 32474403 | 30624525 | 34149477 | 33979457 | 28536049 |
| Q8CHP6 | Phc3 | Polyhomeotic-like protein 3 Mus musculus | 41.81 | 1.33 | 105353 | 1 | 1 | 696666.44 | 526710.75 | 724392.88 | 424297.03 | 641194.19 | 629689 | 708184.4375 | 747476.06 | 718683.88 | 736888.81 | 744340.88 | 968242.69 | 549110.44 | 506102.78 |
| A6H5X4 | Phf11 | PHD finger protein 11 Mus musculus | 71.1 | 10.39 | 37728 | 2 | 2 | 1 | 197548.05 | 936138.56 | 288671.03 | 1 | 507909.62 | 1 | 220330.69 | 1 | 1 | 1 | 391623.97 | 654253.44 | 1 |
| P83870 | Phf5a | PHD finger-like domain-containing protein 5A Mus musculus | 325.89 | 47.27 | 12405 | 5 | 5 | 1582985.75 | 1990887.62 | 836719.88 | 1843662.25 | 2040853.38 | 2243830 | 1798147.5 | 1761533.62 | 2205301 | 2612227 | 2649808.75 | 1991662.88 | 2015362.88 | 2320437.75 |

| | | | | | | | | | | | | | | | | | | | | | |
|---|---|---|---|---|---|---|---|---|---|---|---|---|---|---|---|---|---|---|---|---|---|
| Q9DAK9 | Phpt1 | 14 kDa phosphohistidine phosphatase Mus musculus | 42.6 | 15.32 | 13997 | 1 | 1 | 505115.25 | 469146.16 | 1 | 480579.22 | 1 | 1 | 506018.16 | 497320.5 | 1 | 1 | 674571.31 | 497383.12 | 397147.72 | 366156.94 |
| Q2TBE6 | Pi4k2a | Phosphatidylinositol 4-kinase type 2-alpha Mus musculus | 169.01 | 6.26 | 54258 | 3 | 3 | 2433654.75 | 1953824.25 | 2286650 | 2070547.25 | 2256479 | 1755818.75 | 1848662.25 | 1665317.38 | 1375635.12 | 1578174.75 | 1987015 | 2090155.38 | 2354536 | 1739704.5 |
| E9Q3L2 | Pi4ka | Phosphatidylinositol 4-kinase alpha Mus musculus | 125.61 | 1.9 | 237042 | 2 | 2 | 1 | 1 | 134357.41 | 1 | 1 | 136527.52 | 192876.2475 | 256811.02 | 232711.98 | 145454.47 | 206593.7767 | 292065.56 | 146406.19 | 181309.58 |
| Q7M6Y3 | Picalm | Phosphatidylinositol-binding clathrin assembly protein Mus musculus | 827.62 | 35.3 | 71543 | 1 | 16 | 25045929 | 29878061 | 27825441 | 24341813 | 26191467 | 27257385 | 27716175 | 34538861 | 33235267 | 29128393 | 24563093 | 24035095 | 24508317 | 26144337 |
| Q9CXY9 | Pigk | GPI-anchor transamidase Mus musculus | 98.46 | 5.82 | 44895 | 2 | 2 | 2596389.5 | 2604529.5 | 2888618.5 | 2876564 | 2786547.75 | 3576946 | 3151933.75 | 2682178.5 | 3311018 | 3312016 | 3285250 | 2868224 | 3413149.5 | 3042557.75 |
| Q9R1S3 | Pign | GPI ethanolamine phosphate transferase 1 Mus musculus | 162.1 | 4.3 | 105045 | 3 | 3 | 2485611.5 | 1528267.25 | 2648687.5 | 1693881.25 | 1762883.62 | 2960547.25 | 1770386 | 2825514 | 1890465.12 | 2794665.5 | 2632765.75 | 1644443 | 3068368.25 | 1591098.62 |
| Q6PD26 | Pigs | GPI transamidase component PIG-S Mus musculus | 284.3 | 12.97 | 61711 | 5 | 5 | 2806845.25 | 2095252.75 | 1947335.75 | 2793377.5 | 2648203 | 2914957.25 | 2550083.5 | 2715776.5 | 2256158.75 | 2407828.5 | 2725824.25 | 2524368.5 | 2431480 | 2664987.25 |
| Q8BXQ2 | Pigt | GPI transamidase component PIG-T Mus musculus | 90.4 | 5.5 | 65705 | 2 | 2 | 1827111.88 | 2020421.38 | 1231352.25 | 1495157.5 | 1825833.12 | 2335773.5 | 1720722.25 | 1611787.38 | 1607781.5 | 2130011.75 | 2337649.5 | 2296802.5 | 2326108.75 | 2255476.5 |
| Q8CHJ1 | Pigu | Phosphatidylinositol glycan anchor biosynthesis class U protein Rattus norvegicus | 55.11 | 3.91 | 49953 | 1 | 1 | 728036.62 | 1 | 1 | 1 | 1 | 286638.47 | 234765.9725 | 199706.44 | 253971.7 | 198747.28 | 337560.28 | 1 | 305145.75 | 1 |
| Q9EQ32 | Pik3ap1 | Phosphoinositide 3-kinase adapter protein 1 Mus musculus | 769.93 | 23.06 | 90928 | 14 | 14 | 16486035 | 22704965 | 18916209 | 17843853 | 24338759 | 20708419 | 20514391 | 26283085 | 27100673 | 18773319 | 19254341 | 19844293 | 15999013 | 21405321 |
| Q8BTI9 | Pik3cb | Phosphatidylinositol 4,5-bisphosphate 3-kinase catalytic subunit beta isoform Mus musculu | 54.26 | 1.5 | 121711 | 1 | 1 | 528539.31 | 442406.34 | 1 | 1 | 1 | 598015.9075 | 577891.19 | 577632.44 | 613485.31 | 623054.69 | 497625.53 | 571729.76 | 600266.06 | 617297.69 |
| Q9JHG7 | Pik3cg | Phosphatidylinositol 4,5-bisphosphate 3-kinase catalytic subunit gamma isoform Mus musc | 51.14 | 0.82 | 126400 | 1 | 1 | 939349.38 | 814353.44 | 1030546.19 | 1136400.25 | 1166521 | 898422.69 | 987044.38 | 870997.12 | 825100.25 | 850399.88 | 812244.62 | 763833.25 | 828401.44 | 829620.38 |
| P26450 | Pik3r1 | Phosphatidylinositol 3-kinase regulatory subunit alpha Mus musculus | 44.51 | 1.66 | 83517 | 1 | 1 | 983763.25 | 863371.75 | 797035.06 | 946502.44 | 897668.125 | 1115225.25 | 872001.81 | 729259.88 | 945914.81 | 758126.38 | 1162552.25 | 1129352.88 | 679400.94 | 741568.25 |
| P0C0R5 | Pik3r4 | Phosphoinositide 3-kinase regulatory subunit 4 Rattus norvegicus 3 | 197.14 | 3.83 | 152447 | 4 | 4 | 3335696 | 2340216 | 3167534.25 | 2253984.75 | 3034508 | 2915181 | 2977996.5 | 3249247.75 | 3268674.5 | 2979762 | 2722423.5 | 3133627 | 3028587.75 | 2810513 |
| Q9QUR7 | Pin1 | Peptidyl-prolyl cis-trans isomerase NIMA-interacting 1 Mus musculus | 200.2 | 29.7 | 18370 | 3 | 3 | 4130590 | 5457612.5 | 5450382 | 3411387 | 5057563 | 4562297.5 | 3814822.5 | 3090315.5 | 4427448 | 3988596 | 2752817.5 | 4038743 | 4185972.75 | 4249798.5 |
| O70172 | Pip4k2a | Phosphatidylinositol 5-phosphate 4-kinase type-2 alpha Mus musculus | 133.09 | 9.14 | 46152 | 2 | 3 | 1924650.88 | 2897249.5 | 3319645.5 | 2115855.75 | 2787880.5 | 2860255 | 3347037.5 | 3429582.5 | 3438881.5 | 2539166.5 | 2831562 | 3596395 | 2819137.25 | 3110253.25 |
| O88377 | Pip4k2b | Phosphatidylinositol 5-phosphate 4-kinase type-2 beta Rattus norvegicus | 74.26 | 6.25 | 47264 | 1 | 2 | 2861611.25 | 3089132 | 3489054 | 2074659 | 1855538.5 | 2865660.5 | 3532761 | 3826433.75 | 3542013.25 | 1818226 | 1999208.38 | 3588485.5 | 3216945 | 3630974.75 |
| D3ZSI8 | Pip5k1a | Phosphatidylinositol 4-phosphate 5-kinase type-1 alpha Rattus norvegicus | 50.39 | 2.56 | 60605 | 1 | 1 | 1118438.75 | 1 | 1222851.75 | 891078.56 | 1 | 1 | 1 | 421825.25 | 1 | 1 | 1 | 1 | 1 | 918865.75 |
| P16446 | Pitpna | Phosphatidylinositol transfer protein alpha isoform Rattus norvegicus | 535.86 | 45.02 | 31907 | 10 | 10 | 12655302 | 10579984 | 11744957 | 8875902 | 8547021 | 9822397 | 10267691 | 12136296 | 13883247 | 11194318 | 10394940 | 10237026 | 11466740 | 11671718 |
| P53811 | Pitpnb | Phosphatidylinositol transfer protein beta isoform Mus musculus | 267.34 | 33.21 | 31487 | 7 | 7 | 4186602 | 3557856 | 3335846 | 3566810 | 4621709.5 | 3899177.5 | 2991258.25 | 5590419.5 | 4434497.5 | 5661834 | 4903264 | 2905772.5 | 5233114.5 | 4324751 |
| Q8K411 | Pitrm1 | Presequence protease, mitochondrial Mus musculus | 392.16 | 12.64 | 117372 | 8 | 8 | 7504507 | 9185966 | 9678748 | 8046197 | 8562303 | 9594543 | 9190575 | 9705606 | 10058189 | 9817600 | 10208941 | 8099308 | 10191635 | 9261356 |
| P52480 | Pkm | Pyruvate kinase PKM Mus musculus | 3772.789 | 77.02 | 57845 | 13 | 56 | 1353953409 | 1675794817 | 1641204481 | 1688209025 | 1613781761 | 1830508673 | 1775395841 | 1521528449 | 1766776577 | 2066616321 | 1562039041 | 1637012609 | 1659322497 | 1449816193 |
| P70268 | Pkn1 | Serine/threonine-protein kinase N1 Mus musculus | 434.14 | 13.11 | 104411 | 9 | 9 | 3849104.75 | 3042281.75 | 3887365.75 | 4210186 | 2088987.25 | 3144103 | 3660407 | 3120013.5 | 3105630 | 3003106 | 3033096 | 3742396.5 | 3592270.25 | 3299719.75 |
| Q8VEB4 | Pla2g15 | Group XV phospholipase A2 Mus musculus | 229.78 | 17.96 | 47307 | 6 | 6 | 9589357 | 8330239 | 7938845.5 | 8285196 | 8603706 | 6485803.5 | 9248592 | 9776541 | 8076114.5 | 7390138 | 8491637 | 7750953 | 7675695.5 | 7360586 |
| P47713 | Pla2g4a | Cytosolic phospholipase A2 Mus musculus | 977.38 | 35.16 | 85222 | 19 | 19 | 30164811 | 22980913 | 25101831 | 18467209 | 23935063 | 25351359 | 23313605 | 20397213 | 27638697 | 24959153 | 22639877 | 27787659 | 24967373 | 26232251 |
| P27612 | Plaa | Phospholipase A-2-activating protein Mus musculus | 763.86 | 26.45 | 87221 | 14 | 14 | 6908505 | 8042811 | 8340288 | 9604593 | 11049423 | 8980929 | 9551380 | 7991422 | 12785664 | 10467441 | 9295746 | 12181132 | 11886896 | 11000961 |
| Q3TCN2 | Plbd2 | Putative phospholipase B-like 2 Mus musculus | 514.5 | 21.72 | 66289 | 10 | 10 | 10900258 | 13685235 | 8361534 | 13256259 | 15092123 | 14623600 | 13829656 | 12690423 | 14154651 | 13024529 | 13539777 | 12552769 | 14117087 | 11463788 |
| P51432 | Plcb3 | 1-phosphatidylinositol 4,5-bisphosphate phosphodiesterase beta-3 Mus musculus | 235.68 | 7.54 | 139487 | 5 | 5 | 870095.25 | 1536200 | 1617768.75 | 974266.44 | 1494747.25 | 2135581 | 1680163 | 2121126.5 | 2295048 | 1993543 | 2401421 | 2393453.75 | 1902488.5 | 1969742.5 |
| Q8CIH5 | Plcg2 | 1-phosphatidylinositol 4,5-bisphosphate phosphodiesterase gamma-2 Mus musculus | 905.48 | 20.32 | 147592 | 20 | 20 | 19160359 | 20443325 | 19328189 | 19770709 | 21158481 | 24689185 | 21838817 | 21799275 | 22351687 | 23440199 | 19941969 | 17602193 | 20122117 | 19820003 |
| O35405 | Pld3 | Phospholipase D3 Mus musculus | 139.88 | 11.27 | 54389 | 3 | 3 | 9169067 | 7480470.5 | 6721049 | 9324024 | 8638230 | 6298740.5 | 5068346.5 | 4829077 | 3958668.5 | 4416345 | 9512652 | 9294508 | 5930433 | 7938143 |
| Q8BG07 | Pld4 | Phospholipase D4 Mus musculus | 1299.01 | 42.94 | 56154 | 16 | 19 | 217774625 | 283700417 | 237069425 | 208314785 | 226738369 | 151815697 | 173522833 | 199112241 | 179514177 | 161711761 | 189010065 | 166340353 | 174621745 | 194205297 |
| Q9QXS1 | Plec | Plectin Mus musculus | 10489.382 | 46.9 | 534188 | 183 | 188 | 545612481 | 526952289 | 559298369 | 612632001 | 553254721 | 550656897 | 552424321 | 559611969 | 565419521 | 552997441 | 565381761 | 537161985 | 518466657 | 520650369 |
| Q9JHK5 | Plek | Pleckstrin Mus musculus | 743.29 | 43.14 | 39901 | 11 | 11 | 18507219 | 18363019 | 16011034 | 13170645 | 19258467 | 19118811 | 18092299 | 18183051 | 18711277 | 19263265 | 19047031 | 13569854 | 19103745 | 19612317 |
| Q9ERS5 | Plekha2 | Pleckstrin homology domain-containing family A member 2 Mus musculus | 146.21 | 8.94 | 47380 | 3 | 3 | 1054817.12 | 738206.06 | 254992.8 | 786830.62 | 783559.5 | 1564963.38 | 855086.06 | 811666.06 | 2322426.5 | 1036335 | 844541.94 | 802598.44 | 1455185.62 | 1086383.38 |
| Q8K124 | Plekho2 | Pleckstrin homology domain-containing family O member 2 Mus musculus | 766.48 | 46.87 | 53872 | 13 | 13 | 17818069 | 14481283 | 20974085 | 19805351 | 20332395 | 20317659 | 21672741 | 19699145 | 17812453 | 22020317 | 21828149 | 19789301 | 20304237 | 19909337 |
| P43883 | Plin2 | Perilipin-2 Mus musculus | 38.59 | 3.76 | 46646 | 1 | 1 | 1331561.88 | 1921657.12 | 2026767.25 | 1684054.12 | 1694100.12 | 1296307.75 | 1603171.75 | 1338532.5 | 1442220.62 | 1260768.88 | 1371015.38 | 1233610.62 | 1375488.38 | 1439612 |
| Q9DBG5 | Plin3 | Perilipin-3 Mus musculus | 899.51 | 55.38 | 47262 | 13 | 14 | 63572549 | 62403309 | 57810949 | 82654337 | 80980849 | 72709561 | 66346125 | 54648745 | 57179613 | 66450553 | 94316857 | 105207033 | 56012761 | 78047825 |
| Q07832 | Plk1 | Serine/threonine-protein kinase PLK1 Mus musculus | 176.32 | 8.13 | 68301 | 4 | 4 | 661800.31 | 1368742.25 | 1096273.25 | 1421582.62 | 593047.19 | 584507.06 | 1 | 1 | 316370.06 | 1 | 601306.19 | 1612779.75 | 717066.31 | 1133826.75 |
| Q9R0E2 | Plod1 | Procollagen-lysine,2-oxoglutarate 5-dioxygenase 1 Mus musculus | 218.15 | 8.79 | 83595 | 5 | 5 | 1740708.88 | 996182.12 | 2236154.5 | 505680.28 | 1145521.25 | 2774194.5 | 1605163.5 | 2454534 | 2152946.25 | 1753634.38 | 755654.06 | 1312860.25 | 2147555.75 | 732849.94 |
| Q9R0E1 | Plod3 | Multifunctional procollagen lysine hydroxylase and glycosyltransferase LH3 Mus musculus | 424.41 | 14.98 | 84922 | 8 | 8 | 5364478 | 5032839.5 | 5109065.5 | 4433972 | 4825268 | 4635728 | 4899789 | 4705287.5 | 3916408 | 4813765 | 4431840 | 4185398.5 | 4347884.5 | 3604398.5 |
| Q9Z2Y8 | Plpbp | Pyridoxal phosphate homeostasis protein Mus musculus | 97.09 | 8.03 | 30049 | 2 | 2 | 2219025.25 | 2299979.75 | 2153731.75 | 2477237.75 | 2008060.75 | 2182897.75 | 2418064.75 | 1992502.75 | 1942678 | 1984804.88 | 2350087.25 | 2397449 | 2433353.75 | 2303647.25 |
| Q922V4 | Plrg1 | Pleiotropic regulator 1 Mus musculus | 336.99 | 16.37 | 56938 | 6 | 6 | 1673796.25 | 1381166.25 | 1512445.25 | 1609265 | 1577880.62 | 1019731.62 | 1555922 | 2021479.25 | 1746975.62 | 1412922 | 1458643.12 | 1070390.88 | 902995.88 | 1668873.75 |
| Q3V0K9 | Pls1 | Plastin-1 Mus musculus | 239.73 | 7.46 | 70408 | 0 | 4 | 162932801 | 139718385 | 169041841 | 120631785 | 120178753 | 180728609 | 132377809 | 190820097 | 184877521 | 141066065 | 147082017 | 167066337 | 162003249 | 151677793 |
| Q99K51 | Pls3 | Plastin-3 Mus musculus | 1757.78 | 62.38 | 70742 | 23 | 32 | 428140929 | 438180577 | 453587393 | 434619361 | 434747297 | 523168641 | 448838209 | 530365121 | 537169857 | 478631329 | 468837249 | 474582017 | 470946593 | 490720929 |
| P58195 | Plscr1 | Phospholipid scramblase 1 Rattus norvegicus | 58.81 | 5.97 | 36711 | 1 | 1 | 1147784.62 | 1143446 | 1155475.38 | 1194112.88 | 1016785.81 | 1057941.25 | 910286.5 | 1227599.75 | 1158016.62 | 826703.88 | 886059.4567 | 775410.56 | 975481.56 | 907286.25 |
| Q6QBQ4 | Plscr3 | Phospholipid scramblase 3 Rattus norvegicus 2 | 49.15 | 2.7 | 31646 | 1 | 1 | 1329259.75 | 887402.75 | 908221.69 | 739300.56 | 984196.69 | 836961.81 | 811623 | 856646 | 852112.38 | 735298.19 | 705206.62 | 1251676.88 | 975613 | 780467 |
| P70206 | Plxna1 | Plexin-A1 Mus musculus | 409.96 | 6.12 | 211099 | 7 | 10 | 7789652.5 | 11109825 | 8503231 | 8246097.5 | 9001116 | 6290151.5 | 6913009 | 10529014 | 7534522 | 7859419 | 7751602 | 7298075.5 | 7311485.5 | 6265236 |
| Q80UG2 | Plxna4 | Plexin-A4 Mus musculus | 152.08 | 2.06 | 212560 | 1 | 4 | 202236.52 | 669311.75 | 676408.44 | 395640.09 | 662095 | 204641.47 | 157710.62 | 761954.62 | 622346.62 | 813205.56 | 352464.25 | 136492.56 | 620890.81 | 427847.72 |
| B2RXS4 | Plxnb2 | Plexin-B2 Mus musculus | 1544.57 | 23.45 | 206230 | 27 | 28 | 38733833 | 38123869 | 35281401 | 38462425 | 39450841 | 38870833 | 39079437 | 42287937 | 38543121 | 39042321 | 40045549 | 36839229 | 38536093 | 37826569 |
| Q9QZC2 | Plxnc1 | Plexin-C1 Mus musculus | 80.51 | 1.59 | 176474 | 1 | 2 | 2130857.5 | 1942698.88 | 1995050.25 | 1645297.38 | 1675762.75 | 2106170.75 | 1372831.5 | 2298204 | 1930068.25 | 1674566.25 | 1862034.75 | 1565713.62 | 1604098.5 | 1892427.5 |
| Q3UH93 | Plxnd1 | Plexin-D1 Mus musculus | 82.31 | 0.73 | 211608 | 1 | 2 | 2130857.5 | 1942698.88 | 1995050.25 | 1645297.38 | 1675762.75 | 2106170.75 | 1372831.5 | 2298204 | 1930068.25 | 1674566.25 | 1862034.75 | 1565713.62 | 1604098.5 | 1892427.5 |
| Q60953 | Pml | Protein PML Mus musculus | 405.72 | 12.54 | 98242 | 8 | 8 | 3727383 | 4943247.5 | 3920596 | 5544937 | 3840300.75 | 5259101 | 4721861 | 2867216.5 | 4956309 | 3786297.5 | 3552411 | 4462518 | 4119213 | 2960967.75 |
| O35621 | Pmm1 | Phosphomannomutase 1 Mus musculus | 37.88 | 2.67 | 29775 | 1 | 1 | 1099842.12 | 1270457.62 | 1381751 | 1062134.88 | 1327062.62 | 1116528.38 | 1238694.62 | 1234211.62 | 1268262.62 | 1323385.12 | 1 | 1367676.12 | 1216442.75 | 1332745.25 |
| Q9CXT8 | Pmpcb | Mitochondrial-processing peptidase subunit beta Mus musculus | 236.7 | 9.2 | 54614 | 3 | 3 | 1601344.25 | 572129.62 | 954106.81 | 815037.75 | 929149.25 | 720140.44 | 1267929.12 | 1339049.75 | 1071953.75 | 1039268.81 | 499902.47 | 1372414.5 | 1393798.38 | 743150.19 |
| Q9D1G2 | Pmvk | Phosphomevalonate kinase Mus musculus | 364.17 | 40.62 | 21916 | 7 | 7 | 4786072 | 6624294 | 4326060 | 4172114.5 | 3573549.75 | 5038993 | 4851727 | 2399492.25 | 3228199.75 | 2849410.75 | 3437099 | 2755714.5 | 3742224.5 | 4645706.5 |
| Q9JLV6 | Pnkp | Bifunctional polynucleotide phosphatase/kinase Mus musculus | 286.05 | 21.84 | 57223 | 6 | 6 | 3381179.5 | 3531104.75 | 2664787.75 | 3616525 | 3762678 | 3589898 | 3154724 | 3517325 | 3195913.5 | 3319376 | 3945149.5 | 3524120 | 3406840.5 | 3147851 |
| O35691 | Pnn | Pinin Mus musculus | 87.17 | 3.03 | 82436 | 2 | 2 | 1 | 1 | 1 | 1 | 1 | 255579.23 | 1 | 1 | 292478.25 | 294963.84 | 1 | 373783.44 | 1 | 1 |
| Q9CPS7 | Pno1 | RNA-binding protein PNO1 Mus musculus | 360.37 | 35.08 | 27454 | 6 | 6 | 10794129 | 9796901 | 10321737 | 12206626 | 9797798 | 10995880 | 10744866 | 9242350 | 8095196.5 | 11016702 | 10286537 | 10566674 | 8177167 | 9666873 |
| P23492 | Pnp | Purine nucleoside phosphorylase Mus musculus | 568.02 | 53.63 | 32277 | 9 | 9 | 49712993 | 48014693 | 53075109 | 46328725 | 46769449 | 36315229 | 40750337 | 39961977 | 38437057 | 30295669 | 27103267 | 35282353 | 38767565 | 34755837 |
| O88794 | Pnpo | Pyridoxine-5'-phosphate oxidase Rattus norvegicus | 78.54 | 7.28 | 30184 | 1 | 1 | 211857.48 | 289780 | 1 | 1 | 1 | 1 | 1 | 1 | 167390.19 | 1 | 188403 | 1 | 183752.2 | 1 |
| Q8K1R3 | Pnpt1 | Polyribonucleotide nucleotidyltransferase 1, mitochondrial Mus musculus | 362.7 | 10.47 | 85683 | 6 | 6 | 2268576.25 | 3983668 | 4151390.5 | 3165388 | 2837620.5 | 4351492.5 | 3638037.5 | 3284019.75 | 4192402.25 | 3288215 | 3479135.5 | 2837682.75 | 4085415.5 | 4005184.25 |
| Q91ZW2 | Pofut1 | GDP-fucose protein O-fucosyltransferase 1 Mus musculus | 161.58 | 6.87 | 44688 | 2 | 2 | 1080820.38 | 1082729.38 | 1334043.38 | 1367417.38 | 994716.12 | 1104224.25 | 968019.19 | 1554570.5 | 1145729.12 | 1385566.75 | 1418353.25 | 1044893.31 | 1370408.12 | 1592230.5 |
| Q8VHI3 | Pofut2 | GDP-fucose protein O-fucosyltransferase 2 Mus musculus | 123.34 | 8.39 | 49429 | 3 | 3 | 2027957.62 | 2636437 | 2116389.75 | 2418792.5 | 1831347.38 | 1986822.5 | 2124638.5 | 2257516 | 940784.62 | 2155636 | 2128636.5 | 2550350.75 | 2173222.5 | 2173438.75 |
| Q8BYB9 | Poglut1 | Protein O-glucosyltransferase 1 Mus musculus | 63.4 | 3.57 | 46379 | 1 | 1 | 1357154.38 | 1038875.62 | 1063874.25 | 1 | 1 | 1072737.578 | 1015363.44 | 1391201.62 | 954713.19 | 929672.06 | 1058845.62 | 1081952.453 | 1089254.62 | 1097757.12 |
| P33609 | Pola1 | DNA polymerase alpha catalytic subunit Mus musculus | 298.12 | 7.24 | 167340 | 3 | 7 | 2384784.75 | 1858212.75 | 2085828 | 1297634.75 | 2496580.5 | 2309892.75 | 974450.12 | 2132676.5 | 1961241 | 1570096.25 | 1503445.12 | 1454414.25 | 1350367.75 | 2643687.5 |
| P33611 | Pola2 | DNA polymerase alpha subunit B Mus musculus | 373.46 | 19.17 | 66214 | 8 | 8 | 1202134.5 | 3810813.75 | 2276942.5 | 2836926.5 | 4987065.5 | 1560121.75 | 1311696.75 | 1143023.5 | 1225302.75 | 1754015.5 | 2835265.25 | 3513662.5 | 2046400.62 | 1527945.38 |
| P52431 | Pold1 | DNA polymerase delta catalytic subunit Mus musculus | 851.72 | 17.1 | 123790 | 15 | 15 | 5683090 | 6021531.5 | 7041082.5 | 6819180.5 | 6743368.5 | 5696603 | 4390660 | 6341691.5 | 7730625 | 6888514.5 | 8828413 | 7233209.5 | 6674074.5 | 6496381.5 |
| O35654 | Pold2 | DNA polymerase delta subunit 2 Mus musculus | 164.16 | 5.97 | 51355 | 2 | 2 | 2079043.75 | 1584234 | 1449829.75 | 2223484 | 2042590 | 1721378.38 | 2010782.62 | 1477743.62 | 2072848.88 | 1816243.25 | 2032585.25 | 2438752.75 | 2305868 | 1833449.12 |
| Q91VA6 | Poldip2 | Polymerase delta-interacting protein 2 Mus musculus | 97.95 | 6.52 | 41870 | 2 | 2 | 2267955.75 | 2765633.5 | 2291774 | 2530238.5 | 2195065 | 2138795.5 | 2241895.5 | 2298637.25 | 2135066 | 1935947.25 | 2196960.25 | 2395239.25 | 2125467 | 2239262.5 |
| Q8BG81 | Poldip3 | Polymerase delta-interacting protein 3 Mus musculus | 135.29 | 6.43 | 46132 | 2 | 2 | 2322907.75 | 2237060 | 4153206.25 | 2973884.75 | 3496466.25 | 3223439.75 | 3421584 | 2261658.75 | 4189507.25 | 5386223 | 3726600.5 | 4723547.5 | 4501212 | 3920333.25 |
| O35134 | Polr1a | DNA-directed RNA polymerase I subunit RPA1 Mus musculus | 294.49 | 6.52 | 194110 | 7 | 7 | 1208313.25 | 1723323 | 1655271 | 1210774.88 | 1437643.12 | 2046297.75 | 1952410.5 | 1814990.62 | 1658023.25 | 1428109.88 | 927984.44 | 1341224 | 1847980.75 | 1643851.38 |
| O54888 | Polr1b | DNA-directed RNA polymerase I subunit RPA2 Rattus norvegicus | 35.31 | 1.06 | 127719 | 1 | 1 | 301151.81 | 472892.16 | 363816.28 | 373456.12 | 304530.62 | 304871.84 | 289841.06 | 260074.66 | 439335.88 | 192831.5 | 326653.97 | 336488.97 | 333110.19 | 320686.22 |
| P52432 | Polr1c | DNA-directed RNA polymerases I and III subunit RPAC1 Mus musculus | 80.67 | 4.62 | 39107 | 1 | 1 | 649597.62 | 1 | 563999.06 | 1 | 1 | 498979.38 | 584278.54 | 477941.22 | 747186.06 | 613007.5 | 621503.69 | 522851.06 | 428611 | 711918.62 |
| P08775 | Polr2a | DNA-directed RNA polymerase II subunit RPB1 Mus musculus | 859.94 | 10.81 | 217176 | 15 | 15 | 6130495 | 7017554.5 | 4406641 | 8638785 | 8580928 | 8943941 | 8700812 | 7362786 | 7886023.5 | 7020778.5 | 7910046 | 9900446 | 8668624 | 6635718.5 |
| Q8CFI7 | Polr2b | DNA-directed RNA polymerase II subunit RPB2 Mus musculus | 503.31 | 9.54 | 133911 | 10 | 10 | 3887331.5 | 5512474 | 3498497.25 | 5503975.5 | 6842985 | 3707085.75 | 5654148 | 3749715 | 3328187.5 | 6156603.5 | 4212719 | 5172819.5 | 3305193.5 | 5262571.5 |
| P97760 | Polr2c | DNA-directed RNA polymerase II subunit RPB3 Mus musculus | 247.91 | 16 | 31443 | 5 | 5 | 3377675 | 3345010.5 | 3336368.5 | 2728157.25 | 3007289.25 | 3464399 | 3332690.25 | 4230243 | 3381938 | 2784153.75 | 3456503 | 3538848.25 | 3194375 | 3342902.5 |
| B0BNE2 | Polr2e | DNA-directed RNA polymerases I, II, and III subunit RPABC1 Rattus norvegicus 2 | 83.11 | 11.43 | 24570 | 2 | 2 | 824166.38 | 435735 | 690125.94 | 834303.69 | 666298.69 | 578396.985 | 380695.22 | 848554.31 | 510143.66 | 574194.75 | 448591.81 | 1 | 729596.25 | 1 |
| O88828 | Polr2f | DNA-directed RNA polymerases I, II, and III subunit RPABC2 Rattus norvegicus | 52.49 | 5.51 | 14464 | 1 | 1 | 1859274.38 | 1941991.62 | 1347851.62 | 1806857.25 | 1563165.38 | 2060666.38 | 1811296 | 2067044.62 | 1745089 | 1747267.12 | 1544723.12 | 1767324.25 | 1562232.38 | 1854496.25 |
| P62488 | Polr2g | DNA-directed RNA polymerase II subunit RPB7 Mus musculus | 117.01 | 18.6 | 19294 | 3 | 3 | 1126125.38 | 1104469.75 | 1446462.62 | 1017018.19 | 1332020.12 | 1397288.25 | 2101770.25 | 1161861.62 | 951442 | 1822942.88 | 1191564.5 | 1182641.5 | 1418295.12 | 1116112 |
| Q923G2 | Polr2h | DNA-directed RNA polymerases I, II, and III subunit RPABC3 Mus musculus | 267.07 | 30 | 17143 | 4 | 4 | 9638787 | 11448442 | 13317813 | 9702410 | 8062018.5 | 15240274 | 8956453 | 14879252 | 9449022 | 8104991 | 9526490 | 17369163 | 15034153 | 10921077 |
| P60898 | Polr2i | DNA-directed RNA polymerase II subunit RPB9 Mus musculus | 35.06 | 18.4 | 14523 | 1 | 1 | 1 | 561935.12 | 363359.94 | 1 | 424748.72 | 522352.22 | 397795.62 | 299215.53 | 417934.19 | 442038.56 | 289827.34 | 1 | 1 | 447926.94 |
| Q62086 | Pon2 | Serum paraoxonase/arylesterase 2 Mus musculus | 270.17 | 19.49 | 39617 | 4 | 4 | 952454.44 | 618924.5 | 1425951.25 | 957423.25 | 898176.5 | 627853.19 | 1535853.25 | 1542403.25 | 1429529.38 | 1448907.75 | 1240321.25 | 1606522.5 | 1542674.62 | 690154.81 |
| Q62087 | Pon3 | Serum paraoxonase/lactonase 3 Mus musculus | 312.35 | 30.79 | 39351 | 6 | 6 | 4878769 | 3702919.25 | 4530359.5 | 4513536.5 | 6223161.5 | 2860906 | 3260553 | 4103699.25 | 1667644 | 4788597.5 | 5486497 | 5931205.5 | 2742848.5 | 3902739 |
| P37040 | Por | NADPH--cytochrome P450 reductase Mus musculus | 626.12 | 21.98 | 77044 | 10 | 10 | 10977568 | 12653331 | 11925998 | 10276973 | 11639826 | 14579491 | 10984724 | 11941388 | 12691447 | 12197590 | 12126911 | 11020129 | 15030268 | 12080017 |
| P25425 | Pou2f1 | POU domain, class 2, transcription factor 1 Mus musculus | 168.61 | 4.68 | 79547 | 3 | 3 | 2128809.5 | 3422702 | 927902 | 2532669.5 | 3173457.5 | 3212768 | 3253650.75 | 3095617.25 | 2639025.5 | 3393287 | 2801046 | 3484491.75 | 2983665.5 | 3630145 |
| Q9D819 | Ppa1 | Inorganic pyrophosphatase Mus musculus | 459.65 | 31.83 | 32667 | 8 | 8 | 13805321 | 17046017 | 17311405 | 13757168 | 13618210 | 18038331 | 15711325 | 17476529 | 17083037 | 16750092 | 15566933 | 16632088 | 14982642 | 17143749 |
| Q91VM9 | Ppa2 | Inorganic pyrophosphatase 2, mitochondrial Mus musculus | 328 | 26.36 | 38115 | 7 | 7 | 2332932.75 | 2222790 | 4204944 | 4992291.5 | 5508218 | 5252365 | 4840193 | 5867536 | 6346136 | 6412382.5 | 6004738 | 6050781.5 | 4323268.5 | 2816007 |
| Q91YU8 | Ppan | Suppressor of SWI4 1 homolog Mus musculus | 194.17 | 10 | 52756 | 4 | 4 | 1447403.38 | 906248.94 | 1049341.12 | 1373492.62 | 1488146.75 | 2253746.5 | 1327387.12 | 758186.75 | 1995560.75 | 2141249.5 | 1425888.88 | 1819888.88 | 1435911.25 | 1229190.12 |
| O35711 | Ppfibp2 | Liprin-beta-2 Mus musculus | 103.44 | 1.7 | 98753 | 1 | 1 | 1006869.12 | 749892.81 | 1093282.75 | 949830.06 | 933834.31 | 843772.69 | 837142.1725 | 847486.88 | 761990.31 | 895318.81 | 828588.12 | 964555.38 | 1383104 | 1058749.167 |

| | | | | | | | | | | | | | | | | | | | | | |
|---|---|---|---|---|---|---|---|---|---|---|---|---|---|---|---|---|---|---|---|---|---|
| P17742 | Ppia | Peptidyl-prolyl cis-trans isomerase A Mus musculus | 1083.22 | 79.27 | 17971 | 14 | 16 | 258784449 | 231428353 | 278078177 | 284652577 | 258668129 | 232205617 | 249239793 | 199440481 | 241590625 | 296195649 | 270839617 | 288625025 | 256105457 | 259967185 |
| P24369 | Ppib | Peptidyl-prolyl cis-trans isomerase B Mus musculus | 527.2 | 44.44 | 23713 | 8 | 8 | 71991793 | 64528973 | 64954889 | 56645589 | 54968821 | 56848217 | 59789033 | 58367193 | 60673869 | 56140977 | 59340029 | 57893705 | 62003089 | 54980053 |
| Q9CR16 | Ppid | Peptidyl-prolyl cis-trans isomerase D Mus musculus | 600.07 | 36.76 | 40743 | 12 | 12 | 9842316 | 10942507 | 10216312 | 8024192.5 | 8889505 | 8042992.5 | 8232084 | 8747592 | 11202110 | 8713743 | 9684025 | 11791269 | 11409651 | 9000007 |
| A2AR02 | Ppig | Peptidyl-prolyl cis-trans isomerase G Mus musculus | 102.79 | 3.19 | 88325 | 2 | 2 | 3784205 | 657225 | 569312 | 728397.94 | 826942.56 | 4663139 | 3470473.25 | 3626177.5 | 2714378.5 | 1361528.88 | 4027302 | 3413133.75 | 3052910.5 | 676810.94 |
| Q9D868 | Ppih | Peptidyl-prolyl cis-trans isomerase H Mus musculus | 248.13 | 29.26 | 20464 | 5 | 5 | 5652242 | 4064936 | 3795146.5 | 6647986.5 | 6445592 | 5153454.5 | 5289320.5 | 4382825 | 4168296.25 | 5503907 | 5279168 | 5484852.5 | 5104189.5 | 4719075.5 |
| Q9D0W5 | Ppil1 | Peptidyl-prolyl cis-trans isomerase-like 1 Mus musculus | 52.84 | 20.48 | 18237 | 1 | 1 | 288334.03 | 1 | 1 | 1 | 1 | 267309.78 | 1 | 1 | 1 | 1 | 491628.25 | 1 | 1 | 1 |
| Q9CXG3 | Ppil4 | Peptidyl-prolyl cis-trans isomerase-like 4 Mus musculus | 117.62 | 7.72 | 57230 | 2 | 2 | 1875297 | 1 | 1724944 | 1 | 1 | 831517.62 | 1691165.12 | 1 | 1 | 1 | 1550743.62 | 2115337.5 | 2141181 | 1935754.04 |
| Q6ZQB6 | Ppip5k2 | Inositol hexakisphosphate and diphosphoinositol-pentakisphosphate kinase 2 Mus musculu | 141.43 | 4.07 | 128427 | 3 | 3 | 1313955.12 | 1195377.88 | 756056.56 | 1340971.5 | 1055065.38 | 1533340 | 759403.81 | 546862.19 | 346838.16 | 472768.03 | 1730721.5 | 1953245.5 | 664386.69 | 1202642 |
| P20650 | Ppm1a | Protein phosphatase 1A Rattus norvegicus | 46.33 | 2.88 | 42417 | 1 | 1 | 833024.5 | 879418.62 | 771367.88 | 770505.12 | 910806.38 | 704371.38 | 929085.31 | 1113493.5 | 859451.56 | 813758.25 | 910953.62 | 625497.31 | 766957.62 | 1034635.75 |
| P36993 | Ppm1b | Protein phosphatase 1B Mus musculus | 61.6 | 4.62 | 42795 | 1 | 1 | 1816424 | 2651372.5 | 2343631.75 | 1847520.25 | 1682665.88 | 2211452.5 | 2460205.25 | 2733777.5 | 2469327 | 2322785.5 | 1990340.62 | 2075568.12 | 2130634 | 2775602.25 |
| Q61074 | Ppm1g | Protein phosphatase 1G Mus musculus | 525.34 | 27.12 | 58728 | 10 | 10 | 11635980 | 11763391 | 11682340 | 6516474 | 12106905 | 14358687 | 11587165 | 13537872 | 14029165 | 11765048 | 11403611 | 12625146 | 14194663 | 11868783 |
| Q4FZT2 | Ppme1 | Protein phosphatase methylesterase 1 Rattus norvegicus | 146.88 | 6.99 | 42316 | 3 | 3 | 6184713.5 | 4729126 | 4370266 | 5407718 | 6564517 | 5188333 | 4176921.5 | 3535666.75 | 4010244 | 4043384 | 6063492.5 | 5665088 | 3661940 | 5184402 |
| P51175 | Ppox | Protoporphyrinogen oxidase Mus musculus | 93.2 | 6.92 | 50871 | 2 | 2 | 2312565.25 | 1977656.25 | 2023392.38 | 2096344 | 2418702 | 2330388 | 2617051 | 1494407.88 | 1876089.25 | 1688463.75 | 1928030.12 | 2199618.25 | 2253485.75 | 2014295.62 |
| P62137 | Ppp1ca | Serine/threonine-protein phosphatase PP1-alpha catalytic subunit Mus musculus | 955.57 | 61.21 | 37540 | 1 | 16 | 63125605 | 63142109 | 58299197 | 55401437 | 64901245 | 61654889 | 57019985 | 62211969 | 60019377 | 60336389 | 65474025 | 56228117 | 58965425 | 64666017 |
| P62141 | Ppp1cb | Serine/threonine-protein phosphatase PP1-beta catalytic subunit Mus musculus | 849.14 | 51.68 | 37187 | 4 | 14 | 56258513 | 56350577 | 49453221 | 48099637 | 56901533 | 55764405 | 48944353 | 54925949 | 52220933 | 53372925 | 55982085 | 47906941 | 51951173 | 55874461 |
| P63087 | Ppp1cc | Serine/threonine-protein phosphatase PP1-gamma catalytic subunit Mus musculus | 961.19 | 60.06 | 36984 | 2 | 15 | 57093889 | 55505417 | 52111693 | 50224805 | 59528773 | 56486089 | 50884117 | 54885525 | 55394585 | 54702285 | 58993005 | 50252125 | 53274853 | 58459057 |
| O55000 | Ppp1r10 | Serine/threonine-protein phosphatase 1 regulatory subunit 10 Rattus norvegicus | 39.52 | 2.06 | 92828 | 1 | 1 | 1 | 870048.5 | 553815.44 | 1 | 540611.88 | 1 | 1 | 1 | 1 | 1 | 1 | 1 | 327651.03 | 490496.72 |
| Q9DBR7 | Ppp1r12a | Protein phosphatase 1 regulatory subunit 12A Mus musculus | 147.14 | 3.98 | 114996 | 3 | 3 | 276169.25 | 1491922 | 1230581.12 | 2161822.5 | 1647309.25 | 667972.38 | 1275348.12 | 1511748.38 | 1722962.38 | 2309438.75 | 2654759 | 2056714.38 | 1367632.12 | 1798086.38 |
| Q62084 | Ppp1r14b | Protein phosphatase 1 regulatory subunit 14B Mus musculus | 138.95 | 9.52 | 15957 | 2 | 2 | 378832.88 | 1017930.19 | 1128509.12 | 1069769.12 | 832537.12 | 973217.19 | 1002307 | 1730055.5 | 1104816.62 | 736921.31 | 967180.25 | 1051733 | 638583.19 | 870802.5 |
| Q8BQ30 | Ppp1r18 | Phostensin Mus musculus | 211.23 | 8.59 | 65630 | 5 | 5 | 3270208.75 | 1535491.12 | 1069069.25 | 1639436.5 | 877134.12 | 1404104.25 | 1537484.25 | 1831989.75 | 1618915.25 | 1592215 | 587674.19 | 730581.94 | 1548475.75 | 1770285 |
| Q3TDD9 | Ppp1r21 | Protein phosphatase 1 regulatory subunit 21 Mus musculus | 137.28 | 4.1 | 88338 | 2 | 2 | 530425.69 | 357747.75 | 1 | 779400.19 | 1 | 1093154 | 756437.25 | 771334.12 | 191035.67 | 835074.5 | 490537.81 | 437447.28 | 551218.31 | 270585.72 |
| Q3UM45 | Ppp1r7 | Protein phosphatase 1 regulatory subunit 7 Mus musculus | 1010.14 | 52.35 | 41292 | 14 | 14 | 20763125 | 19350099 | 17294293 | 19419697 | 24535133 | 22307391 | 20442139 | 29946021 | 24623191 | 21231401 | 19271597 | 23350705 | 27057523 | 20800175 |
| O35274 | Ppp1r9b | Neurabin-2 Rattus norvegicus | 171.3 | 7.22 | 89646 | 3 | 3 | 1191231 | 1580052.38 | 1734393 | 1338055.75 | 1314881.75 | 1478694.75 | 1223603.38 | 1373591 | 1133706 | 926193 | 1048438.25 | 1415620.75 | 1436132.5 | 1536637.62 |
| P63330 | Ppp2ca | Serine/threonine-protein phosphatase 2A catalytic subunit alpha isoform Mus musculus | 455.31 | 41.75 | 35608 | 1 | 9 | 32261029 | 34238525 | 30054383 | 29710217 | 36659589 | 38126205 | 32476311 | 36103981 | 33467865 | 36307593 | 37997557 | 33589537 | 36084089 | 39441037 |
| P62715 | Ppp2cb | Serine/threonine-protein phosphatase 2A catalytic subunit beta isoform Mus musculus | 444.64 | 40.13 | 35575 | 1 | 9 | 31010917 | 33454095 | 29690579 | 33054857 | 36666785 | 37803513 | 31170203 | 34643277 | 32238395 | 35384957 | 38292029 | 32446509 | 35479265 | 38889201 |
| Q76MZ3 | Ppp2r1a | Serine/threonine-protein phosphatase 2A 65 kDa regulatory subunit A alpha isoform Mus n | 2107.18 | 64.35 | 65323 | 23 | 33 | 114746873 | 128048809 | 126684097 | 118607377 | 116083617 | 141708465 | 126688105 | 144928385 | 137329841 | 127348945 | 129637289 | 132702345 | 134147513 | 136290289 |
| Q4QQT4 | Ppp2r1b | Serine/threonine-protein phosphatase 2A 65 kDa regulatory subunit A beta isoform Rattus | 933.86 | 29.78 | 66006 | 4 | 13 | 39195481 | 41542317 | 44276041 | 40827941 | 41004665 | 47447493 | 40281437 | 47086229 | 45605901 | 41383129 | 44158613 | 42876037 | 43911297 | 44756705 |
| P36876 | Ppp2r2a | Serine/threonine-protein phosphatase 2A 55 kDa regulatory subunit B alpha isoform Rattus | 271.45 | 13.2 | 51678 | 5 | 5 | 8231431 | 8330970 | 11728833 | 8788771 | 6324596 | 10758032 | 10282099 | 8560013 | 11715667 | 7824176.5 | 10134631 | 11159074 | 10859322 | 10529899 |
| Q6PD03 | Ppp2r5a | Serine/threonine-protein phosphatase 2A 56 kDa regulatory subunit alpha isoform Mus mu | 286.95 | 14.81 | 56347 | 5 | 6 | 5057821.5 | 4250047 | 5595199 | 5685191 | 7293131.5 | 4248495.5 | 7321480 | 6591210.5 | 6258503 | 4324095.5 | 1867846.5 | 6661755.5 | 7133225 | 7801223 |
| Q60996 | Ppp2r5c | Serine/threonine-protein phosphatase 2A 56 kDa regulatory subunit gamma isoform Mus n | 205.03 | 9.73 | 60824 | 2 | 3 | 824246.5 | 524290.88 | 517954.97 | 545492.62 | 430880.75 | 591869.69 | 659892.94 | 776394.88 | 418218.62 | 514862.06 | 641202.62 | 384956.69 | 585823.5 | 142552.91 |
| Q61151 | Ppp2r5e | Serine/threonine-protein phosphatase 2A 56 kDa regulatory subunit epsilon isoform Mus m | 110.81 | 5.14 | 54713 | 0 | 2 | 1416328.25 | 1445775.12 | 1328024.88 | 1152912.12 | 1212347.38 | 1397147.25 | 1627820.75 | 1468078.25 | 1427407.5 | 1089463 | 1331589.75 | 1352609.12 | 1247480.88 | 1540161.88 |
| P63328 | Ppp3ca | Serine/threonine-protein phosphatase 2B catalytic subunit alpha isoform Mus musculus | 428.71 | 18.43 | 58644 | 3 | 8 | 7832728.5 | 7283644 | 10544078 | 8440826 | 9449262 | 7005434 | 9010868 | 8555364 | 9326767 | 9299610 | 7321889.5 | 9322199 | 9108919 | 9948883 |
| P20651 | Ppp3cb | Serine/threonine-protein phosphatase 2B catalytic subunit beta isoform Rattus norvegicus | 280.43 | 12.95 | 59113 | 1 | 6 | 7411415.5 | 7283644 | 10116448 | 8440826 | 9009060 | 7005434 | 8562251 | 8555364 | 9326767 | 9299610 | 7321889.5 | 9322199 | 8417067 | 9311708 |
| P63100 | Ppp3r1 | Calcineurin subunit B type 1 Rattus norvegicus | 207.88 | 52.35 | 19300 | 4 | 4 | 2545973.25 | 4231836 | 4503000 | 3813856.5 | 2342452.5 | 3272138 | 4705582 | 4360382 | 3045129 | 4069312 | 4195258 | 2831794.5 | 3453007.75 | 4701937 |
| P97470 | Ppp4c | Serine/threonine-protein phosphatase 4 catalytic subunit Mus musculus | 253.62 | 18.89 | 35080 | 3 | 5 | 8024741 | 9617601 | 7312884 | 9913145 | 8796322 | 8552854 | 8691860 | 9454233 | 8259164 | 8574160 | 9349349 | 7708831 | 8789324 | 9274957 |
| Q8K2V1 | Ppp4r1 | Serine/threonine-protein phosphatase 4 regulatory subunit 1 Mus musculus | 405.36 | 11.25 | 106298 | 7 | 7 | 4969056 | 3926590 | 3827186 | 4019834.5 | 3875424.5 | 3560705 | 4576339 | 4746056 | 3957207.75 | 3461821.5 | 3667844.5 | 3670637 | 4218131.5 | 3938146.75 |
| Q0VGB7 | Ppp4r2 | Serine/threonine-protein phosphatase 4 regulatory subunit 2 Mus musculus | 177.66 | 10.55 | 46479 | 3 | 3 | 1850140 | 2034461.25 | 2903052.75 | 1955833 | 2421902 | 3027132.5 | 3224331 | 3485005.5 | 3319220.25 | 3209334.5 | 3189156 | 1803652 | 3804962 | 3350605 |
| Q6P2K6 | Ppp4r3a | Serine/threonine-protein phosphatase 4 regulatory subunit 3A Mus musculus | 170.73 | 7.2 | 93842 | 3 | 3 | 727461.94 | 249066.88 | 549090.38 | 178199.16 | 939538.94 | 395838.12 | 709535.88 | 474365.66 | 678032.56 | 476326.81 | 313069.62 | 825777.81 | 859398.56 | 599390.69 |
| Q922R5 | Ppp4r3b | Serine/threonine-protein phosphatase 4 regulatory subunit 3B Mus musculus | 152.42 | 4.15 | 93934 | 2 | 2 | 282221.12 | 310789.46 | 234770.5 | 304347.28 | 421818.94 | 1 | 1 | 1 | 314378.31 | 225663.28 | 286825.4267 | 306067.84 | 267651 | 286757.44 |
| Q60676 | Ppp5c | Serine/threonine-protein phosphatase 5 Mus musculus | 317.02 | 13.63 | 56877 | 5 | 5 | 7287939.5 | 7789658 | 7772254 | 6768153.5 | 7353698 | 6527098 | 7598275.5 | 6979646.5 | 7036337 | 6761019 | 7465686 | 7266636.5 | 7338236 | 7210146 |
| Q64620 | Ppp6c | Serine/threonine-protein phosphatase 6 catalytic subunit Rattus norvegicus | 410.62 | 31.48 | 35159 | 7 | 7 | 7714178.5 | 8356844 | 6788874 | 8338708 | 7995899.5 | 8224132 | 7909893.5 | 8198971 | 8560355 | 7484471.5 | 8327255 | 8521837 | 7649502 | 7511152 |
| Q7TSI3 | Ppp6r1 | Serine/threonine-protein phosphatase 6 regulatory subunit 1 Mus musculus | 258.19 | 7.71 | 94527 | 4 | 4 | 3986228.5 | 3452987.5 | 3294930.5 | 3856874.5 | 3947366 | 3711228.25 | 3549707 | 3702940.75 | 3684503 | 4343336 | 3549801 | 3413421.25 | 3702475.5 | 3866093 |
| Q922D4 | Ppp6r3 | Serine/threonine-protein phosphatase 6 regulatory subunit 3 Mus musculus | 372.73 | 10.31 | 94653 | 7 | 7 | 7959086 | 7394944 | 7224796 | 6713954.5 | 7296041 | 6600551.5 | 7507074 | 7655000 | 7689509 | 6380844 | 5766490 | 6682267 | 6132965.5 | 6455693 |
| O88531 | Ppt1 | Palmitoyl-protein thioesterase 1 Mus musculus | 361.35 | 27.45 | 34490 | 6 | 6 | 23402251 | 21343829 | 19147169 | 20901289 | 22389045 | 25206765 | 16059733 | 18755181 | 28282909 | 16205002 | 29076177 | 25610589 | 25337227 | 14781999 |
| Q8CEC6 | Ppwd1 | Peptidylprolyl isomerase domain and WD repeat-containing protein 1 Mus musculus | 52.69 | 2.01 | 73385 | 1 | 1 | 6784737.5 | 14874537 | 46376205 | 30773133 | 69166489 | 83548193 | 20117743 | 3388300 | 37399177 | 90466713 | 77970193 | 86852145 | 87349105 | 92075785 |
| P35700 | Prdx1 | Peroxiredoxin-1 Mus musculus | 1304.28 | 89.45 | 22176 | 8 | 23 | 943166273 | 1082265217 | 1051464513 | 721838849 | 938903553 | 681353281 | 698175745 | 789360257 | 754446209 | 738384449 | 883846273 | 968239233 | 953239041 | 976386369 |
| Q61171 | Prdx2 | Peroxiredoxin-2 Mus musculus | 761.15 | 60.1 | 21779 | 13 | 13 | 193823601 | 219943553 | 203400849 | 203886273 | 214063009 | 224128481 | 200042081 | 217034657 | 208909841 | 208376801 | 209269633 | 208100017 | 207743873 | 225472081 |
| P20108 | Prdx3 | Thioredoxin-dependent peroxide reductase, mitochondrial Mus musculus | 403.04 | 42.8 | 28127 | 3 | 8 | 35121993 | 37198285 | 39623681 | 31474173 | 32779713 | 37331045 | 36232697 | 43078745 | 42407737 | 34146865 | 35763969 | 35989441 | 39149889 | 36109457 |
| O08807 | Prdx4 | Peroxiredoxin-4 Mus musculus | 864.57 | 63.87 | 31053 | 14 | 16 | 217877649 | 262339073 | 235947489 | 166877601 | 216095537 | 159523185 | 180677569 | 210574737 | 187434433 | 173811761 | 178932753 | 210008017 | 211464241 | 230749777 |
| P99029 | Prdx5 | Peroxiredoxin-5, mitochondrial Mus musculus | 774.76 | 57.14 | 21897 | 12 | 12 | 56091017 | 48160737 | 47277485 | 38873749 | 49548785 | 44597389 | 49405089 | 40686829 | 48408037 | 48692697 | 47494617 | 49725777 | 54260325 | 42405449 |
| O08709 | Prdx6 | Peroxiredoxin-6 Mus musculus | 1011.65 | 80.36 | 24871 | 17 | 17 | 76961673 | 86185241 | 90438033 | 75179377 | 87438225 | 78102801 | 74558929 | 83283729 | 85087337 | 85131249 | 87621353 | 95580401 | 88368993 | 94130673 |
| Q9WUQ2 | Preb | Prolactin regulatory element-binding protein Mus musculus | 186.51 | 11.75 | 45437 | 3 | 3 | 1280340.5 | 1837462.88 | 1958280.5 | 1251712.12 | 460748.56 | 407268.53 | 604428.12 | 1453625.75 | 1623236.25 | 1393938.62 | 929789.62 | 1946388.75 | 1335319 | 238860.66 |
| Q9QUR6 | Prep | Prolyl endopeptidase Mus musculus | 1197.57 | 38.59 | 80752 | 19 | 19 | 24552079 | 25021749 | 26155509 | 23544407 | 22921967 | 24381545 | 24172529 | 29420695 | 26796907 | 24634427 | 25491957 | 22964963 | 25430397 | 26695833 |
| Q69ZK0 | Prex1 | Phosphatidylinositol 3,4,5-trisphosphate-dependent Rac exchanger 1 protein Mus musculu | 1388.41 | 22.42 | 184935 | 27 | 27 | 26847303 | 27958801 | 27497355 | 24238315 | 29390133 | 29025799 | 28728313 | 32196119 | 31290789 | 26931377 | 29062003 | 26870949 | 27430805 | 30143961 |
| P20664 | Prim1 | DNA primase small subunit Mus musculus | 79.63 | 6 | 49295 | 2 | 2 | 1491189.25 | 1334787.75 | 1393445.62 | 1285053.62 | 1336268.62 | 1484642 | 1334205.25 | 633480.94 | 1331680.25 | 923888.62 | 549756.69 | 1695279.88 | 1443104.5 | 1487598 |
| P33610 | Prim2 | DNA primase large subunit Mus musculus | 281.84 | 12.48 | 58408 | 5 | 5 | 2131409.25 | 2714766.25 | 3903885 | 2777066.25 | 1657883.38 | 2627025.5 | 3429755.5 | 2853950.5 | 2927672.75 | 3689052 | 3610194.25 | 3784593.5 | 2536028 | 3649831.5 |
| P54645 | Prkaa1 | 5'-AMP-activated protein kinase catalytic subunit alpha-1 Rattus norvegicus | 131.28 | 6.44 | 63973 | 3 | 3 | 1389563.25 | 1004741 | 1438128.75 | 1325978 | 1516875 | 1682883.25 | 1626149.5 | 1217798.5 | 1620742 | 1859502 | 1548625.12 | 1818288.5 | 1738368.12 | 1356494.75 |
| P05132 | Prkaca | cAMP-dependent protein kinase catalytic subunit alpha Mus musculus | 381.87 | 26.5 | 40571 | 2 | 8 | 24790171 | 23126309 | 21086561 | 19925703 | 21379051 | 23454681 | 25869275 | 22591377 | 24098525 | 22977071 | 23035099 | 25603557 | 22432551 | 23198141 |
| P68181 | Prkacb | cAMP-dependent protein kinase catalytic subunit beta Mus musculus | 394.28 | 26.5 | 40708 | 2 | 8 | 23713019 | 22095197 | 19357943 | 18849219 | 21044219 | 22309039 | 24203625 | 21214897 | 22377009 | 21392307 | 21775077 | 23812533 | 20370401 | 21226395 |
| O54950 | Prkag1 | 5'-AMP-activated protein kinase subunit gamma-1 Mus musculus | 591.44 | 43.03 | 37520 | 9 | 9 | 3652924.25 | 3171194.5 | 3296764.75 | 3395168.25 | 1629812 | 3241449.5 | 4789996.5 | 5651160.5 | 3104472 | 3004139.75 | 3123961.25 | 2094249.5 | 4177229.5 | 4854916 |
| Q9DBC7 | Prkar1a | cAMP-dependent protein kinase type I-alpha regulatory subunit Mus musculus | 631.42 | 34.91 | 43185 | 11 | 11 | 20935051 | 23816355 | 21509539 | 23857553 | 24367271 | 20280033 | 20458831 | 20114593 | 19862863 | 21942341 | 22525745 | 19538983 | 20724205 | 22213181 |
| P12368 | Prkar2a | cAMP-dependent protein kinase type II-alpha regulatory subunit Rattus norvegicus | 337.12 | 20.95 | 45480 | 6 | 6 | 5819845.5 | 6378244.5 | 3912883.5 | 5561034 | 8573565 | 9191183 | 8665423 | 6052968 | 8419529 | 6989055 | 8447003 | 9207045 | 8916037 | 5252936 |
| P68403 | Prkcb | Protein kinase C beta type Rattus norvegicus | 192.2 | 6.11 | 76751 | 3 | 4 | 629502.56 | 477210.38 | 1 | 1 | 284359.34 | 403026.97 | 1 | 350326.75 | 1 | 1 | 1 | 1 | 1 | 1 |
| P28867 | Prkcd | Protein kinase C delta type Mus musculus | 396.02 | 13.2 | 77547 | 7 | 8 | 9568542 | 12952222 | 10283728 | 12139951 | 14702367 | 11126005 | 11332786 | 10707586 | 13941058 | 12607472 | 12864308 | 13054753 | 11841435 | 15189415 |
| O08795 | Prkcsh | Glucosidase 2 subunit beta Mus musculus | 423.96 | 19.39 | 58793 | 8 | 8 | 14769977 | 14824747 | 16255006 | 15607839 | 14695123 | 14626441 | 15648041 | 14677563 | 13347025 | 14074395 | 14125877 | 13542592 | 13251337 | 14463818 |
| Q5XIS9 | Prkd2 | Serine/threonine-protein kinase D2 Rattus norvegicus | 73.48 | 1.26 | 96497 | 1 | 1 | 1 | 143415.03 | 1 | 1 | 144991.11 | 129389.72 | 126323.55 | 170978.66 | 177755.91 | 119926.11 | 205412.16 | 181738.61 | 172690.94 | 116241.67 |
| Q63009 | Prmt1 | Protein arginine N-methyltransferase 1 Rattus norvegicus | 1101.34 | 53.54 | 40522 | 20 | 20 | 48532225 | 47985949 | 46430077 | 42264709 | 45721829 | 41912681 | 35487577 | 40275569 | 37532961 | 37666145 | 26250871 | 48974569 | 37189773 | 42121745 |
| Q8CIG8 | Prmt5 | Protein arginine N-methyltransferase 5 Mus musculus | 264.02 | 12.24 | 72680 | 6 | 6 | 5985871 | 4608271 | 5556597 | 2380798 | 3578270 | 3879445.5 | 2864243 | 4388351 | 4409214 | 4587343 | 3330222 | 3840951 | 5109314 | 4604025 |
| Q64695 | Procr | Endothelial protein C receptor Mus musculus | 40.87 | 7.02 | 27189 | 1 | 1 | 793269.44 | 716483.94 | 656806.94 | 634748.1975 | 372432.47 | 751266.8675 | 734241.44 | 874520.62 | 894712 | 501593.41 | 806146.75 | 890274.38 | 729458.94 | 647810.81 |
| Q9D820 | Prorsd1 | Prolyl-tRNA synthetase associated domain-containing protein 1 Mus musculus | 74.67 | 10.06 | 18862 | 2 | 2 | 615482.5 | 1214442.62 | 849631.19 | 1042563.75 | 827395.31 | 754252.12 | 769632.56 | 1160169.5 | 792154 | 799988 | 450607.69 | 2235478.5 | 778331.31 | 934584.19 |
| Q99KP6 | Prpf19 | Pre-mRNA-processing factor 19 Mus musculus | 682.12 | 52.18 | 55239 | 13 | 13 | 22918815 | 21627957 | 27461579 | 24921621 | 24320841 | 24115929 | 25615773 | 23527675 | 26485841 | 27819133 | 21902741 | 24533849 | 25165397 | 22870045 |
| Q922U1 | Prpf3 | U4/U6 small nuclear ribonucleoprotein Prp3 Mus musculus | 117.93 | 4.83 | 77455 | 3 | 3 | 761582 | 645401.25 | 668652.5 | 694886.12 | 692630.4675 | 425397.44 | 695242.75 | 567076.4225 | 442535.5 | 705130 | 618626.19 | 643891.31 | 875648 | 636032.5 |
| Q8CCF0 | Prpf31 | U4/U6 small nuclear ribonucleoprotein Prp31 Mus musculus | 664.09 | 31.66 | 55430 | 11 | 11 | 10524550 | 9749807 | 9692875 | 8772612 | 8067656.5 | 8634876 | 9352214 | 9999866 | 10319928 | 8689089 | 9248835 | 11672306 | 9570526 | 9708497 |
| Q4FK66 | Prpf38a | Pre-mRNA-splicing factor 38A Mus musculus | 220.03 | 12.82 | 37437 | 4 | 4 | 3209023.5 | 3982061 | 3610993.25 | 3744178.25 | 4512680 | 3016640 | 3193747.5 | 4054402.25 | 3654625.75 | 4141366 | 3755531 | 3873593 | 2959702 | 4539450.5 |
| Q6AXY7 | Prpf38b | Pre-mRNA-splicing factor 38B Rattus norvegicus | 94.67 | 5.35 | 63932 | 2 | 2 | 1203714.38 | 1538512.62 | 1099557.25 | 1420370 | 1767775.12 | 1436615.62 | 1480167.25 | 1746809.12 | 1649858.12 | 1711017.5 | 1315481.75 | 1588527 | 1590791.5 | 1595884.88 |
| Q9DAW6 | Prpf4 | U4/U6 small nuclear ribonucleoprotein Prp4 Mus musculus | 103.33 | 4.99 | 58370 | 2 | 2 | 3025687.75 | 505579.22 | 3164263.25 | 3665880.5 | 3719378.5 | 1961805.12 | 2502765.5 | 2782544.25 | 2865354.75 | 2526367.75 | 700654.44 | 3534662.5 | 1271322.25 | 2993061 |
| Q9R1C7 | Prpf40a | Pre-mRNA-processing factor 40 homolog A Mus musculus | 288.99 | 10.18 | 108481 | 6 | 6 | 7514857 | 7282676.5 | 4758313 | 5327646.5 | 7648230 | 7338669.5 | 6759585 | 8331431.5 | 8326241.5 | 7547538 | 4834835 | 8016013.5 | 6727482 | 7957815.5 |
| Q5RKH1 | Prpf4b | Serine/threonine-protein kinase PRP4 homolog Rattus norvegicus | 51.07 | 1.69 | 117007 | 1 | 1 | 1424606.88 | 1689704.62 | 1048682.38 | 1515060.25 | 1387228.25 | 1244764.12 | 1320071.38 | 1421908.25 | 1220538.62 | 1204857.75 | 953380.31 | 923301.12 | 889005 | 1316950.5 |
| A1A5S1 | Prpf6 | Pre-mRNA-processing factor 6 Rattus norvegicus | 217.37 | 4.89 | 106764 | 6 | 6 | 1923015.5 | 4360011.5 | 4236407 | 2191759 | 2376259.5 | 2043599.25 | 3702453.25 | 3927891.5 | 3077242.75 | 3452741.5 | 2461148.25 | 2286594.5 | 3668849.75 | 2379987 |
| Q99PV0 | Prpf8 | Pre-mRNA-processing-splicing factor 8 Mus musculus | 1434.24 | 18.03 | 273616 | 29 | 29 | 31754929 | 29173377 | 25986165 | 24888961 | 31501925 | 32235845 | 26606077 | 23401445 | 28281721 | 28499879 | 30440497 | 28631513 | 32374615 | 30705027 |
| P60892 | Prps1 | Ribose-phosphate pyrophosphokinase 1 Rattus norvegicus | 753.02 | 41.82 | 34834 | 5 | 12 | 13199419 | 12918571 | 13263252 | 13187058 | 13574412 | 15598464 | 11561083 | 12227264 | 12877634 | 14057268 | 14655282 | 13945043 | 12897274 | 12437864 |
| Q9CS42 | Prps2 | Ribose-phosphate pyrophosphokinase 2 Mus musculus | 653.17 | 39.31 | 34786 | 3 | 10 | 10548243 | 10978101 | 10697736 | 11481259 | 13096030 | 12629257 | 9097256 | 7551114.5 | 11029104 | 11726949 | 12172339 | 12090278 | 12340429 | 11906948 |
| Q9D0M1 | Prpsap1 | Phosphoribosyl pyrophosphate synthase-associated protein 1 Mus musculus | 446.08 | 27.25 | 39432 | 6 | 8 | 10095758 | 8078865 | 5478888 | 6610907.5 | 7423608 | 8600723 | 8156839 | 9923553 | 7554720.5 | 7631229.5 | 9539898 | 10250357 | 7364263.5 | 7533634 |
| Q8R574 | Prpsap2 | Phosphoribosyl pyrophosphate synthase-associated protein 2 Mus musculus | 388.39 | 24.39 | 40881 | 5 | 7 | 7194530 | 6435119 | 5972740.5 | 6093794 | 6516345 | 5992907 | 6724877 | 6996434.5 | 6481862 | 6474586 | 6318293.5 | 6964855 | 6177725 | 6193517 |
| Q3T1I4 | Prrc1 | Protein PRRC1 Rattus norvegicus | 295.8 | 27.31 | 46295 | 6 | 6 | 2347264 | 5160084 | 4681518 | 4292462 | 3467514.75 | 4168958.5 | 3705844.5 | 5061324.5 | 3109505 | 3793591.5 | 3515677 | 4339010 | 3176743.75 | 3665599.25 |
| Q3TLH4 | Prrc2c | Protein PRRC2C Mus musculus | 602.04 | 6.01 | 310892 | 11 | 11 | 1514673.25 | 2235882 | 2596029 | 2538311.25 | 1600206.5 | 2271578 | 2007723.12 | 1847377 | 2538546.5 | 4142253.5 | 2976039 | 2526141.5 | 3364323.5 | 2218278 |
| Q61207 | Psap | Prosaposin Mus musculus | 500.34 | 16.34 | 61422 | 8 | 9 | 62608617 | 85619153 | 61576009 | 67607481 | 62274225 | 50240585 | 61469061 | 68667873 | 45177369 | 55014417 | 55335193 | 56597797 | 54787129 | 64248065 |
| Q99K85 | Psat1 | Phosphoserine aminotransferase Mus musculus | 1147.95 | 60 | 40473 | 18 | 18 | 65698977 | 68842881 | 75055409 | 73179913 | 70061217 | 68063641 | 72252569 | 69837697 | 69875145 | 71767585 | 65165625 | 70766657 | 69380977 | 71416273 |
| O88777 | Psen2 | Presenilin-2 Rattus norvegicus | 101.35 | 3.35 | 50051 | 1 | 1 | 1112437.88 | 1977275.62 | 1674955.25 | 1653055.25 | 1574831.88 | 1255809.5 | 1462820.62 | 1733572.38 | 1823083.5 | 1427782.12 | 1644232.75 | 1919254.38 | 1390247.5 | 1507549 |

| | | | | | | | | | | | | | | | | | | | | | |
|---|---|---|---|---|---|---|---|---|---|---|---|---|---|---|---|---|---|---|---|---|---|
| Q99JF8 | Psip1 | PC4 and SFRS1-interacting protein Mus musculus | 460.55 | 19.89 | 59697 | 8 | 9 | 12897072 | 12902274 | 8302025.5 | 13204524 | 13383367 | 12171392 | 13537311 | 11004723 | 13168898 | 12693244 | 11267877 | 12065961 | 12902655 | 14699911 |
| Q9R1P4 | Psma1 | Proteasome subunit alpha type-1 Mus musculus | 521.4 | 60.46 | 29547 | 12 | 12 | 31595289 | 35210129 | 31054659 | 36645581 | 35170449 | 31328893 | 30190913 | 32691079 | 29411711 | 34062585 | 31041197 | 27388313 | 30424051 | 31240233 |
| P17220 | Psma2 | Proteasome subunit alpha type-2 Rattus norvegicus | 631.85 | 46.15 | 25927 | 9 | 9 | 5367493 | 3770205.5 | 4416430 | 4643863.5 | 5352430 | 4090701 | 4674239 | 4813675.5 | 4960198 | 4000707.5 | 5080979 | 6056246.5 | 6216345 | 5680444.5 |
| O70435 | Psma3 | Proteasome subunit alpha type-3 Mus musculus | 525.17 | 34.51 | 28405 | 8 | 8 | 46703641 | 45811433 | 40112433 | 37888277 | 41449493 | 38952433 | 40181201 | 45318289 | 42420813 | 39417257 | 37648857 | 39183593 | 39695641 | 41539089 |
| P21670 | Psma4 | Proteasome subunit alpha type-4 Rattus norvegicus | 623.23 | 52.87 | 29498 | 9 | 9 | 23731827 | 22714291 | 22937429 | 19437545 | 20412123 | 21589293 | 19622753 | 26228217 | 22585727 | 19499539 | 22977007 | 20836001 | 21026601 | 20560241 |
| Q9Z2U1 | Psma5 | Proteasome subunit alpha type-5 Mus musculus | 628.62 | 65.98 | 26411 | 10 | 10 | 48418773 | 54426885 | 61356933 | 50844089 | 53689413 | 52071073 | 53258469 | 53317841 | 55810637 | 53184533 | 53708337 | 57260141 | 54939297 | 53750609 |
| Q9QUM9 | Psma6 | Proteasome subunit alpha type-6 Mus musculus | 601.56 | 59.35 | 27372 | 10 | 10 | 40426513 | 36778433 | 43377193 | 41162473 | 38554765 | 38417077 | 46109721 | 35185037 | 43003853 | 47292265 | 45774233 | 42772165 | 44258745 | 42906169 |
| Q9Z2U0 | Psma7 | Proteasome subunit alpha type-7 Mus musculus | 662.54 | 51.61 | 27855 | 11 | 11 | 48832961 | 37826333 | 51889441 | 36299857 | 35391637 | 46339701 | 41592373 | 44808405 | 48475825 | 41587057 | 45154113 | 51520873 | 52856509 | 47229525 |
| O09061 | Psmb1 | Proteasome subunit beta type-1 Mus musculus | 889.38 | 65.42 | 26372 | 11 | 13 | 44223393 | 44306617 | 46898557 | 46310325 | 47877153 | 42559633 | 45224585 | 41998777 | 44204529 | 45229533 | 43931049 | 42514097 | 45795613 | 48975485 |
| O35955 | Psmb10 | Proteasome subunit beta type-10 Mus musculus | 263.8 | 21.25 | 29063 | 4 | 4 | 4053446 | 4426572 | 5276907.5 | 5294959 | 3176399 | 4030263.75 | 4467116 | 3309078.75 | 4240396 | 4555254.5 | 4338776 | 4059786 | 4295687.5 | 4479535.5 |
| Q9R1P3 | Psmb2 | Proteasome subunit beta type-2 Mus musculus | 564.71 | 64.18 | 22906 | 10 | 10 | 25188259 | 23431355 | 23791241 | 22826801 | 21087685 | 20522263 | 21122711 | 31285935 | 27445387 | 29176923 | 21361967 | 20670025 | 29799179 | 20114605 |
| Q9R1P1 | Psmb3 | Proteasome subunit beta type-3 Mus musculus | 493.69 | 53.17 | 22965 | 8 | 8 | 38088821 | 39792409 | 41585749 | 38853289 | 41475689 | 43095721 | 40983641 | 38006333 | 44259581 | 43750361 | 43412085 | 43693037 | 45077701 | 43424569 |
| P99026 | Psmb4 | Proteasome subunit beta type-4 Mus musculus | 390.14 | 37.12 | 29116 | 6 | 6 | 28099033 | 25629421 | 30363549 | 29061781 | 29095897 | 29506421 | 26440475 | 25705993 | 29967291 | 27653713 | 26061757 | 27761695 | 29207581 | 27435163 |
| Q60692 | Psmb6 | Proteasome subunit beta type-6 Mus musculus | 201.31 | 26.89 | 25379 | 4 | 4 | 10038085 | 11115873 | 7537159 | 8827584 | 10774531 | 8987874 | 8101392 | 11106551 | 7399331 | 9689384 | 9300162 | 6674578.5 | 9881826 | 10885678 |
| P70195 | Psmb7 | Proteasome subunit beta type-7 Mus musculus | 151.7 | 28.16 | 29891 | 4 | 4 | 4540134 | 3425151.5 | 3747979.25 | 3307585.75 | 3136085.75 | 4211839 | 4792129 | 4805163 | 4010537.5 | 3882361.25 | 4130477.5 | 6382910.5 | 4125211 | 4524401 |
| P28063 | Psmb8 | Proteasome subunit beta type-8 Mus musculus | 763.5 | 51.81 | 30260 | 11 | 11 | 27842901 | 29829893 | 29364637 | 24339501 | 27690625 | 27444579 | 28109699 | 26543861 | 27158935 | 28107445 | 27677703 | 27687825 | 28026085 | 28898801 |
| O35522 | Psmb9 | Proteasome subunit beta type-9 Mus musculus bactrianus | 260.75 | 34.25 | 23427 | 5 | 5 | 23776365 | 27161923 | 22946583 | 26063835 | 24925917 | 20062789 | 21509665 | 21172077 | 18464499 | 19879483 | 20457933 | 17895225 | 19753685 | 20238313 |
| P62192 | Psmc1 | 26S proteasome regulatory subunit 4 Mus musculus | 766.86 | 32.5 | 49185 | 10 | 12 | 14150257 | 12966606 | 15130525 | 12603510 | 12033492 | 14438416 | 13439354 | 13380781 | 13270433 | 16444658 | 14264743 | 13351477 | 15734899 | 15577627 |
| P46471 | Psmc2 | 26S proteasome regulatory subunit 7 Mus musculus | 1108.77 | 47.34 | 48648 | 17 | 17 | 27022051 | 24684971 | 23998349 | 28707055 | 25778543 | 25900465 | 27246309 | 28547313 | 24721741 | 28430285 | 29970781 | 26309021 | 30500727 | 29686577 |
| O88685 | Psmc3 | 26S proteasome regulatory subunit 6A Mus musculus | 1011.96 | 48.87 | 49549 | 16 | 16 | 29903755 | 30740665 | 30914379 | 34134981 | 32561789 | 34142569 | 32252621 | 32383761 | 33231109 | 33759477 | 35515105 | 28878661 | 35416317 | 35147605 |
| P54775 | Psmc4 | 26S proteasome regulatory subunit 6B Mus musculus | 1029.62 | 51.44 | 47408 | 16 | 17 | 19170535 | 17510731 | 16672966 | 14593041 | 15335300 | 16436831 | 20407199 | 22035899 | 16799447 | 15041296 | 17796225 | 16458292 | 17541975 | 15574577 |
| P62196 | Psmc5 | 26S proteasome regulatory subunit 8 Mus musculus | 930.07 | 47.78 | 45626 | 15 | 17 | 20966437 | 18491385 | 16955559 | 21492891 | 19581869 | 20297011 | 19648681 | 17481853 | 16829947 | 19046237 | 21320197 | 20908843 | 18114693 | 18904329 |
| P62334 | Psmc6 | 26S proteasome regulatory subunit 10B Mus musculus | 719.43 | 44.47 | 44173 | 15 | 15 | 22500033 | 21118061 | 23835547 | 24038083 | 20199929 | 21936629 | 22596891 | 24436797 | 21877917 | 22735005 | 23364009 | 22252457 | 23580727 | 24497551 |
| Q3TXS7 | Psmd1 | 26S proteasome non-ATPase regulatory subunit 1 Mus musculus | 1808.57 | 42.29 | 105730 | 26 | 28 | 46912189 | 51100169 | 50941269 | 59514185 | 53150733 | 56259265 | 62453453 | 57807113 | 59982053 | 54405073 | 53427149 | 57928801 | 51232345 | 51709521 |
| Q9Z2X2 | Psmd10 | 26S proteasome non-ATPase regulatory subunit 10 Mus musculus | 42.58 | 3.46 | 25084 | 1 | 1 | 1447422.5 | 1131062.5 | 1309101.75 | 1804776.25 | 1462486.5 | 1484352.25 | 1222847.88 | 1397216.25 | 1697062 | 1431510.62 | 1418098.75 | 1473166.88 | 1599244.75 | 1261796.88 |
| F1LMZ8 | Psmd11 | 26S proteasome non-ATPase regulatory subunit 11 Rattus norvegicus | 1109.24 | 57.58 | 47464 | 19 | 19 | 45383285 | 36889545 | 33363937 | 32518591 | 34964637 | 34600877 | 34814909 | 36151809 | 35014733 | 32814121 | 34428465 | 35634345 | 32946569 | 32764601 |
| Q9D8W5 | Psmd12 | 26S proteasome non-ATPase regulatory subunit 12 Mus musculus | 1211.41 | 50.88 | 52895 | 19 | 20 | 52846409 | 55731561 | 53422661 | 46323533 | 49097925 | 54220157 | 56626693 | 67624497 | 58159593 | 47375141 | 49814137 | 58390985 | 57786321 | 53381929 |
| Q9WVJ2 | Psmd13 | 26S proteasome non-ATPase regulatory subunit 13 Mus musculus | 918.77 | 47.87 | 42809 | 17 | 17 | 45466909 | 42412457 | 37190153 | 40671525 | 38362953 | 37739729 | 41024773 | 48483857 | 36960169 | 39492617 | 38427553 | 32994179 | 40405325 | 39062885 |
| O35593 | Psmd14 | 26S proteasome non-ATPase regulatory subunit 14 Mus musculus | 392.99 | 38.06 | 34577 | 6 | 6 | 17339677 | 21239563 | 20101507 | 21653543 | 20161839 | 20196317 | 22049425 | 24433519 | 19528843 | 20325631 | 21325821 | 21788837 | 18891157 | 21266471 |
| Q8VDM4 | Psmd2 | 26S proteasome non-ATPase regulatory subunit 2 Mus musculus | 1800.66 | 47.8 | 100203 | 29 | 29 | 66501597 | 65012813 | 71737081 | 67672377 | 68472497 | 73770321 | 70062689 | 63874921 | 72961921 | 75627369 | 75782417 | 69639537 | 75584089 | 74630673 |
| P14685 | Psmd3 | 26S proteasome non-ATPase regulatory subunit 3 Mus musculus | 444.5 | 22.45 | 60718 | 9 | 9 | 16284630 | 7235685 | 4872775 | 6756865 | 9649383 | 11896048 | 4891788.5 | 6403771.5 | 7205343.5 | 9626881 | 12189598 | 7955783.5 | 11299892 | 9406827 |
| O35226 | Psmd4 | 26S proteasome non-ATPase regulatory subunit 4 Mus musculus | 501.55 | 28.72 | 40704 | 10 | 10 | 14012863 | 12021024 | 13135014 | 11817732 | 13702150 | 13177783 | 11837446 | 12032927 | 12292011 | 13555810 | 13655655 | 13111721 | 12981744 | 12423157 |
| Q8BJY1 | Psmd5 | 26S proteasome non-ATPase regulatory subunit 5 Mus musculus | 674.42 | 36.11 | 55972 | 14 | 14 | 7831379 | 8856142 | 9746756 | 10063931 | 8296208 | 10467280 | 8977993 | 12232026 | 10748926 | 9928393 | 10998091 | 11795082 | 10389344 | 9003396 |
| Q99JI4 | Psmd6 | 26S proteasome non-ATPase regulatory subunit 6 Mus musculus | 1081.49 | 61.95 | 45536 | 20 | 20 | 35138045 | 31516525 | 36492601 | 30299551 | 27870467 | 32611107 | 30319889 | 37362677 | 35397217 | 29763273 | 31905421 | 32674229 | 32554553 | 32514055 |
| P26516 | Psmd7 | 26S proteasome non-ATPase regulatory subunit 7 Mus musculus | 747.02 | 57.63 | 36540 | 13 | 13 | 20427787 | 14682115 | 18734853 | 13634627 | 15109045 | 18343179 | 20085933 | 22311685 | 16875357 | 20814127 | 18688185 | 16107567 | 19779579 | 19809681 |
| Q9CX56 | Psmd8 | 26S proteasome non-ATPase regulatory subunit 8 Mus musculus | 305.13 | 24.93 | 39930 | 7 | 7 | 7137359 | 8078915.5 | 8560070 | 7215239 | 6351252 | 7551747 | 9194350 | 10328450 | 9408406 | 7886103 | 8110501.5 | 7967179.5 | 8116069.5 | 8071255 |
| P97371 | Psme1 | Proteasome activator complex subunit 1 Mus musculus | 975.63 | 57.03 | 28673 | 15 | 15 | 89540657 | 95525945 | 95758017 | 75130945 | 81183945 | 75755689 | 74087425 | 86415321 | 86245657 | 78407361 | 69659497 | 77183457 | 70087921 | 73277425 |
| P97372 | Psme2 | Proteasome activator complex subunit 2 Mus musculus | 1003.36 | 56.07 | 27057 | 13 | 13 | 44316197 | 46731149 | 47717777 | 34232265 | 37201365 | 38278585 | 37660221 | 47421921 | 42163009 | 33968617 | 35788645 | 34942001 | 40652481 | 36011461 |
| P61290 | Psme3 | Proteasome activator complex subunit 3 Mus musculus | 587.29 | 41.73 | 29506 | 10 | 10 | 9824825 | 10620431 | 10491492 | 10125554 | 10774689 | 11197419 | 8082523 | 8108236 | 9461851 | 9197790 | 9644236 | 6914624 | 10340915 | 10347007 |
| Q8BHL8 | Psmf1 | Proteasome inhibitor PI31 subunit Mus musculus | 73.24 | 4.43 | 29664 | 1 | 1 | 1691224.25 | 1649087.38 | 1543821.88 | 1740047.75 | 1757861.75 | 1480819.25 | 1561427.38 | 1600148.5 | 1593886.62 | 1487472.5 | 1778619.12 | 1882632.38 | 1543761.88 | 1708240.75 |
| Q9JK23 | Psmg1 | Proteasome assembly chaperone 1 Mus musculus | 251.7 | 17.65 | 33104 | 4 | 4 | 6532307 | 6510476 | 5634383 | 6581602 | 6723938.5 | 5755333 | 6293061 | 6190214 | 7291173 | 6524887 | 5697538 | 6986973.5 | 6182931 | 5658773 |
| Q9EST4 | Psmg2 | Proteasome assembly chaperone 2 Mus musculus | 103.54 | 3.03 | 29524 | 2 | 2 | 1 | 347121.22 | 1 | 1 | 296036.81 | 1 | 1 | 1 | 1 | 1 | 1 | 1 | 575879.94 | 1 |
| Q9CZH3 | Psmg3 | Proteasome assembly chaperone 3 Mus musculus | 106.03 | 22.95 | 13331 | 2 | 2 | 1786032.75 | 853473.06 | 1653060 | 1845699 | 4105884 | 1757558 | 804751 | 1416937.25 | 1064826.75 | 1766495.75 | 1589863.38 | 1237129.38 | 1809358 | 5442128.5 |
| P0C7N9 | Psmg4 | Proteasome assembly chaperone 4 Mus musculus | 80.71 | 33.33 | 13974 | 2 | 2 | 709616.09 | 422771.12 | 248700.12 | 1575617 | 591376.12 | 469909.25 | 437857.53 | 348173 | 360231.5 | 556403.56 | 552387.31 | 448082.0933 | 432805.41 | 359053.56 |
| Q4KLH4 | Pspc1 | Paraspeckle component 1 Rattus norvegicus | 180.35 | 9 | 58760 | 2 | 3 | 935184 | 1933783 | 1416285.88 | 1579770.065 | 2033827.38 | 2009663.25 | 636827.88 | 1620797.5 | 1375997.62 | 1174230.88 | 607099.62 | 1266919.83 | 1559779.75 | 1633880.12 |
| P97814 | Pstpip1 | Proline-serine-threonine phosphatase-interacting protein 1 Mus musculus | 282.97 | 17.11 | 47590 | 5 | 5 | 9298495 | 9274206 | 8585177 | 9588877 | 8833164 | 7339234 | 8400129 | 8584521 | 7509706 | 8168469.5 | 7430355.5 | 8039140.5 | 7276998.5 | 7285711 |
| P17225 | Ptbp1 | Polypyrimidine tract-binding protein 1 Mus musculus | 1433.44 | 61.86 | 56478 | 3 | 18 | 36129277 | 46671741 | 39793857 | 46020513 | 52002501 | 54407809 | 48004117 | 40416193 | 52178589 | 49467409 | 53436285 | 47260813 | 49284517 | 49247741 |
| Q8BHD7 | Ptbp3 | Polypyrimidine tract-binding protein 3 Mus musculus | 640.14 | 43.98 | 56701 | 8 | 11 | 5983338 | 4179773.25 | 2239316.75 | 5139728 | 7018103 | 6966994.5 | 5024604.5 | 5784415.5 | 8267049 | 5124060.5 | 10336314 | 6116370.5 | 4371208.5 | 5957414.5 |
| Q14C51 | Ptcd3 | Pentatricopeptide repeat domain-containing protein 3, mitochondrial Mus musculus | 41.2 | 1.17 | 77796 | 1 | 1 | 3577212.75 | 1079706.12 | 1003975.44 | 1367289.62 | 1257993.88 | 1322158.62 | 1380316 | 1539433.25 | 1061945 | 1374409.88 | 1340954.88 | 1120238.75 | 1439339.25 | 1263307.62 |
| Q60866 | Pter | Phosphotriesterase-related protein Mus musculus | 148.09 | 11.46 | 39218 | 3 | 3 | 3970088 | 3982317.5 | 4329619 | 4762867 | 3923348.5 | 3845309.25 | 5057915 | 4025296.25 | 4933977.5 | 4633489 | 4634379 | 4601474 | 4685160 | 4792128 |
| P83868 | Ptges3 | Prostaglandin E synthase 3 Rattus norvegicus | 173.34 | 26.25 | 18721 | 3 | 3 | 17011015 | 13231800 | 9545441 | 14601500 | 16514794 | 13801698 | 13004568 | 14068629 | 11805297 | 13599677 | 15502186 | 11989341 | 12709328 | 13621515 |
| Q91YR9 | Ptgr1 | Prostaglandin reductase 1 Mus musculus | 711.79 | 57.75 | 35560 | 13 | 13 | 30446173 | 43041713 | 39012065 | 37214841 | 37579401 | 31931789 | 35661549 | 36504929 | 34365433 | 30807333 | 30831227 | 39425117 | 37839713 | 32484277 |
| Q8VDQ1 | Ptgr2 | Prostaglandin reductase 2 Mus musculus | 218.17 | 15.95 | 38015 | 4 | 4 | 5464668 | 5582771 | 5610796.5 | 5416513 | 6787951 | 5855222 | 6006883 | 5168242.5 | 5274605.5 | 5437702.5 | 6266358 | 5820248 | 6237794 | 6660921.5 |
| P22437 | Ptgs1 | Prostaglandin G/H synthase 1 Mus musculus | 354.16 | 15.12 | 69042 | 6 | 7 | 318926.88 | 3227523.5 | 2729618 | 6842449 | 5433438.5 | 8101430 | 8955387 | 8751370 | 8849903 | 9009687 | 8367141 | 5618519 | 5071180.5 | 5009584.5 |
| Q9QVP9 | Ptk2b | Protein-tyrosine kinase 2-beta Mus musculus | 949.64 | 28.25 | 115794 | 20 | 20 | 28455413 | 22882949 | 22717637 | 23024557 | 20719505 | 25797893 | 21692037 | 35443121 | 32470521 | 21774563 | 20608911 | 25032421 | 27320627 | 23817839 |
| P26350 | Ptma | Prothymosin alpha Mus musculus | 518.26 | 36.04 | 12254 | 6 | 6 | 136869921 | 163779153 | 148286337 | 148546609 | 136402113 | 141144833 | 146987697 | 122566617 | 135885553 | 155953681 | 140134225 | 119242777 | 82625473 | 154343649 |
| P04550 | Ptms | Parathymosin Rattus norvegicus | 208.88 | 22.55 | 11559 | 3 | 3 | 39655745 | 50878625 | 34129413 | 44393737 | 47642841 | 38023689 | 49292801 | 32388369 | 37839533 | 39797345 | 45919633 | 31789953 | 31195477 | 41193769 |
| Q5U2W6 | Ptov1 | Prostate tumor-overexpressed gene 1 protein homolog Rattus norvegicus | 51.38 | 4.57 | 46849 | 1 | 1 | 159473.75 | 243645.39 | 307737.06 | 193803.83 | 183177.56 | 295345.5 | 176646.64 | 238874.2 | 217486.12 | 152736.55 | 220020.78 | 275788.12 | 276762.5 | 181376.97 |
| P58389 | Ptpa | Serine/threonine-protein phosphatase 2A activator Mus musculus | 348.32 | 35.29 | 36710 | 6 | 6 | 10686629 | 10124990 | 9324665 | 7714597.5 | 8110287 | 9739038 | 9561375 | 12107562 | 11207093 | 9346346 | 9726505 | 9907925 | 10131350 | 10725432 |
| P35821 | Ptpn1 | Tyrosine-protein phosphatase non-receptor type 1 Mus musculus | 193.39 | 12.5 | 49593 | 5 | 5 | 5496886 | 6492162 | 4846930.5 | 5256600 | 5298270.5 | 5307264 | 4286403.5 | 5740215 | 5188380.5 | 5046442 | 5864193 | 5400434 | 6514274.5 | 4904228.5 |
| P35831 | Ptpn12 | Tyrosine-protein phosphatase non-receptor type 12 Mus musculus | 122.42 | 5.29 | 86526 | 3 | 3 | 1075914.88 | 1664086.5 | 2275661 | 2266905.5 | 2073114 | 1976845.25 | 2186137 | 1301953.12 | 2156916.75 | 2271921 | 2220447.25 | 2473681.25 | 2393932.5 | 2340738 |
| Q6PB44 | Ptpn23 | Tyrosine-protein phosphatase non-receptor type 23 Mus musculus | 40.18 | 0.71 | 185216 | 1 | 1 | 1 | 1 | 1313953.12 | 1 | 1 | 1152120.5 | 1 | 1023506.88 | 1 | 958061.62 | 1 | 1143302.88 | 939879.56 | 1 |
| P29351 | Ptpn6 | Tyrosine-protein phosphatase non-receptor type 6 Mus musculus | 2297.35 | 72.61 | 67559 | 36 | 38 | 126555201 | 130331697 | 140119553 | 135952289 | 137699537 | 126718769 | 130476849 | 135932145 | 136935169 | 144172673 | 127672689 | 134331649 | 127613337 | 132530913 |
| P49445 | Ptpn7 | Tyrosine-protein phosphatase non-receptor type 7 Rattus norvegicus | 46.59 | 3.62 | 40314 | 1 | 1 | 504673.72 | 502326.03 | 530789.44 | 704985.31 | 605939.31 | 580063.94 | 475219.78 | 514228.56 | 379385.38 | 519743.59 | 481192.69 | 490500.62 | 445578.91 | 445627.5 |
| O35239 | Ptpn9 | Tyrosine-protein phosphatase non-receptor type 9 Mus musculus | 70.43 | 3.71 | 67970 | 2 | 2 | 564572 | 206183.41 | 1 | 1 | 335159.47 | 618421.75 | 414485.97 | 260030.67 | 455510.06 | 294147.12 | 309626.7433 | 339615.38 | 319006.66 | 270258.19 |
| P18052 | Ptpra | Receptor-type tyrosine-protein phosphatase alpha Mus musculus | 153.81 | 4.7 | 93698 | 3 | 3 | 3998869.5 | 4070362.5 | 4259567.5 | 3541227 | 3619625 | 3648993 | 3097311.5 | 3295869.5 | 3436489.5 | 3991696.75 | 1616491.88 | 3087454.5 | 3724702 | 3587260.5 |
| P06800 | Ptprc | Receptor-type tyrosine-protein phosphatase C Mus musculus | 2183.95 | 32.71 | 144837 | 36 | 37 | 137451825 | 142186913 | 141481121 | 140101329 | 153271585 | 157866737 | 140575137 | 160629889 | 161057953 | 154518481 | 143151025 | 137637985 | 147279713 | 144030897 |
| B2GV87 | Ptpre | Receptor-type tyrosine-protein phosphatase epsilon Rattus norvegicus | 155.66 | 5.87 | 80734 | 3 | 3 | 1086241.88 | 1589133.38 | 1746180.5 | 1015236.94 | 1454266.12 | 1446705 | 1496522.75 | 1665263.25 | 1320733.12 | 1316315.38 | 1739174.5 | 1458770 | 2045167.12 | 1563144.5 |
| E9Q612 | Ptpro | Receptor-type tyrosine-protein phosphatase O Mus musculus | 40.97 | 0.9 | 138589 | 1 | 1 | 1243094 | 1322124.5 | 897329 | 1212953 | 1481179.75 | 869947.69 | 1321038 | 1154329.75 | 1059608 | 933220.5 | 1174039.88 | 1188205.88 | 1050841 | 1751061.75 |
| Q8R2Y8 | Ptrh2 | Peptidyl-tRNA hydrolase 2, mitochondrial Mus musculus | 138.26 | 27.62 | 19527 | 3 | 3 | 3005910.5 | 2955586 | 3802728.5 | 3109108.5 | 4159630 | 3233012.5 | 2972310 | 2106009.5 | 3285989 | 4050375.75 | 4444746.5 | 3198542.5 | 4708541 | 4124705 |
| Q8R143 | Pttg1ip | Pituitary tumor-transforming gene 1 protein-interacting protein Mus musculus | 68.09 | 8.62 | 19965 | 2 | 2 | 3739812 | 3049899.5 | 2303842.5 | 1109900.88 | 2465014.75 | 3181392.5 | 3200391 | 2212371.25 | 1949721.5 | 1969225.62 | 1762549.12 | 1941391.62 | 1824762 | 1936751.75 |
| Q3UEB3 | Puf60 | Poly(U)-binding-splicing factor PUF60 Mus musculus | 835.78 | 34.75 | 60249 | 13 | 13 | 21663497 | 19814313 | 20259581 | 17078499 | 16589848 | 19928995 | 23013913 | 15422050 | 17570813 | 19016427 | 16412802 | 21250767 | 16434125 | 21514743 |
| Q8BKS9 | Pum3 | Pumilio homolog 3 Mus musculus | 315.32 | 12.06 | 72800 | 5 | 5 | 2330401.25 | 988184.69 | 4035540.75 | 1167950.75 | 1099484.5 | 5026593 | 1127541.25 | 1683363.38 | 1960400 | 1302328.5 | 2092007.38 | 1251511.75 | 1606548.25 | 2185759.75 |
| P42669 | Pura | Transcriptional activator protein Pur-alpha Mus musculus | 268.63 | 20.87 | 34884 | 4 | 4 | 1150833.62 | 905254.06 | 1 | 1301468.38 | 2129022.25 | 1113389.25 | 2064518.12 | 1312536 | 599543.62 | 827399.25 | 1512563.5 | 1002530.75 | 545349.06 | 949679.69 |
| O35295 | Purb | Transcriptional activator protein Pur-beta Mus musculus | 225.25 | 24.07 | 33901 | 5 | 5 | 3858990.5 | 4825599 | 3841536.75 | 4611350 | 4764654.5 | 5980753 | 4540948 | 5135999 | 5481793.5 | 5580309 | 4349804 | 3714919.5 | 4943638.5 | 6412186 |
| Q91VU7 | Pus7 | Pseudouridylate synthase 7 homolog Mus musculus | 171.23 | 9.7 | 74793 | 4 | 4 | 994688.25 | 618611.925 | 941911.62 | 281371.38 | 256476.45 | 283778.66 | 397832.5 | 517615.84 | 393756.41 | 960903.88 | 862106.44 | 913114.75 | 744044.69 | 674227.5 |
| Q99LL5 | Pwp1 | Periodic tryptophan protein 1 homolog Mus musculus | 159.94 | 11.98 | 55587 | 3 | 3 | 1304749.25 | 1121091.5 | 1550567.5 | 1523970.5 | 1444675.25 | 1692176.12 | 1405399.75 | 932509.94 | 857517.62 | 948768.31 | 1697928.25 | 2455174.5 | 1147582.12 | 2732996 |
| Q8BU03 | Pwp2 | Periodic tryptophan protein 2 homolog Mus musculus | 190.18 | 6.42 | 102910 | 4 | 4 | 966987.56 | 924193.19 | 874700.38 | 771334.69 | 928563.06 | 970059.25 | 761625.94 | 805041.81 | 1250649.88 | 1351042.62 | 1241401.5 | 994448.81 | 1188925 | 923628.88 |
| Q8VI36 | Pxn | Paxillin Mus musculus | 80.45 | 9.98 | 64476 | 2 | 2 | 132730.3 | 1 | 1 | 1 | 1 | 1 | 1 | 1 | 163400.91 | 1 | 1 | 1 | 1 | 1 |
| Q9EPB4 | Pycard | Apoptosis-associated speck-like protein containing a CARD Mus musculus | 455.27 | 56.99 | 21459 | 8 | 8 | 16844279 | 12940189 | 15560411 | 14470844 | 12986655 | 15241557 | 14442267 | 14628203 | 14474019 | 13005193 | 15107874 | 14162110 | 14189469 | 12174662 |
| Q922Q4 | Pycr2 | Pyrroline-5-carboxylate reductase 2 Mus musculus | 704.07 | 35.31 | 33659 | 10 | 10 | 18715853 | 16591343 | 17587775 | 21074971 | 21293397 | 18532171 | 20751665 | 16745819 | 21056305 | 21228303 | 22036191 | 22069561 | 20361415 | 19574401 |
| P53534 | Pygb | Glycogen phosphorylase, brain form Rattus norvegicus | 997.77 | 23.39 | 96174 | 13 | 16 | 38852877 | 40796329 | 42306413 | 36641517 | 39003997 | 44483133 | 41864561 | 43885309 | 46652597 | 42594941 | 43729029 | 43060269 | 42179901 | 44278145 |
| P09811 | Pygl | Glycogen phosphorylase, liver form Rattus norvegicus | 237.34 | 7.53 | 97483 | 1 | 4 | 9630024 | 9077226 | 9885653 | 9628375 | 10200595 | 10990385 | 9743619 | 10130671 | 10637783 | 10480683 | 11086207 | 10506221 | 9696289 | 11027949 |
| Q8BML9 | Qars | Glutamine--tRNA ligase Mus musculus | 641.94 | 21.03 | 87677 | 14 | 14 | 10923693 | 12416317 | 13178927 | 11004910 | 11090575 | 12116599 | 11538592 | 14230106 | 11907779 | 10983114 | 11793074 | 12845381 | 13906799 | 12302105 |
| Q8BVI4 | Qdpr | Dihydropteridine reductase Mus musculus | 342.47 | 33.61 | 25570 | 5 | 5 | 5157324.5 | 5930228.5 | 6655475 | 5754099.5 | 5579340 | 5923568 | 3372950.5 | 6544766 | 6942169 | 6783428 | 6096860.5 | 7802934 | 7056802 | 6839028 |
| Q91XU1 | Qki | Protein quaking Rattus norvegicus | 307.42 | 29.62 | 37643 | 8 | 8 | 12454134 | 12073374 | 11243338 | 12401428 | 11836760 | 12375581 | 13536542 | 13072787 | 13495711 | 13113253 | 12004055 | 9655860 | 9752497 | 12054920 |
| Q91X91 | Qprt | Nicotinate-nucleotide pyrophosphorylase [carboxylating] Mus musculus | 248.55 | 22.41 | 31530 | 5 | 5 | 703643.69 | 3376261.25 | 2422194.5 | 1380101.25 | 676490.81 | 1258884 | 912655.62 | 1425679.62 | 1241590.25 | 1461135.12 | 496618.41 | 1239660.38 | 1741822.12 | 1119817.25 |
| B8ZXI1 | Qtrt2 | Queuine tRNA-ribosyltransferase accessory subunit 2 Mus musculus | 36.87 | 3.86 | 46288 | 1 | 1 | 1786591.12 | 1844135.25 | 664873.75 | 1860417 | 1881407.38 | 1434269 | 1298987.5 | 1418956.62 | 1655172.88 | 285261.44 | 1412163.33 | 1374509.12 | 1383908.75 | 1478072.12 |
| P61027 | Rab10 | Ras-related protein Rab-10 Mus musculus | 490.94 | 42 | 22541 | 7 | 10 | 21850323 | 21533451 | 21389139 | 20575261 | 19195639 | 23333243 | 22967457 | 24281881 | 22086963 | 22118249 | 22161477 | 22816339 | 24300193 | 22347823 |

| | | | | | | | | | | | | | | | | | | | | | |
|---|---|---|---|---|---|---|---|---|---|---|---|---|---|---|---|---|---|---|---|---|---|
| O35509 | Rab11b | Ras-related protein Rab-11B Rattus norvegicus | 556.61 | 50.92 | 24489 | 11 | 11 | 69784809 | 61384121 | 58891449 | 50684601 | 51708893 | 57037977 | 56115741 | 62476445 | 57986201 | 54146361 | 49023705 | 59211001 | 54813185 | 55638561 |
| Q9D620 | Rab11fip1 | Rab11 family-interacting protein 1 Mus musculus | 96.99 | 5.43 | 70684 | 2 | 2 | 1 | 1 | 188517.3 | 1 | 1 | 1 | 1 | 382795.31 | 1 | 127464.23 | 1 | 192605.22 | 1 | 167348.11 |
| P35283 | Rab12 | Ras-related protein Rab-12 Mus musculus | 149.63 | 12.76 | 27329 | 2 | 3 | 1353021.5 | 618446.38 | 1338756.5 | 849985.31 | 917115.62 | 1687018.12 | 356996.53 | 1010009.88 | 1529765.12 | 1011570.25 | 891571.62 | 1369757.5 | 1386641.75 | 1300431.25 |
| Q91V41 | Rab14 | Ras-related protein Rab-14 Mus musculus | 815.37 | 80 | 23897 | 12 | 14 | 25779797 | 25251989 | 26092057 | 26823629 | 27031935 | 29939299 | 24049607 | 29460551 | 26784353 | 29716733 | 29123059 | 28947961 | 31373229 | 26516647 |
| P35293 | Rab18 | Ras-related protein Rab-18 Mus musculus | 320.7 | 41.26 | 23035 | 6 | 6 | 4780978 | 4271044.5 | 4749120 | 3416822.5 | 2344066.5 | 4734946 | 3488657.25 | 5330698 | 4612586.5 | 3105068.5 | 2878194.5 | 4499268 | 4895100 | 4705031 |
| P62821 | Rab1A | Ras-related protein Rab-1A Mus musculus | 874.43 | 68.78 | 22678 | 7 | 15 | 50089177 | 53785033 | 55181625 | 40476249 | 45167837 | 56361281 | 51929677 | 59555137 | 51106009 | 48809841 | 51294133 | 52675461 | 55712721 | 55857689 |
| P10536 | Rab1b | Ras-related protein Rab-1B Rattus norvegicus | 837.06 | 72.14 | 22163 | 6 | 13 | 40798405 | 45746005 | 45516297 | 37083493 | 36079633 | 43648169 | 43061725 | 48167401 | 44384505 | 42241457 | 43096409 | 47195681 | 46106025 | 43223673 |
| P35282 | Rab21 | Ras-related protein Rab-21 Mus musculus | 264.32 | 25.23 | 24106 | 4 | 4 | 2766645.25 | 3741388 | 4337648 | 2008819 | 3630291.25 | 4891254 | 4202029 | 5514778 | 3845160.5 | 3519791.75 | 4040633.5 | 2443717.75 | 4374702.5 | 4278893 |
| P35285 | Rab22a | Ras-related protein Rab-22A Mus musculus | 90.91 | 12.89 | 21802 | 1 | 2 | 1856008.5 | 2247866 | 2327043.5 | 2879868 | 526297.94 | 1 | 2380791.75 | 1962036.12 | 2675884.25 | 1 | 4421631 | 1 | 1 | 1 |
| P35290 | Rab24 | Ras-related protein Rab-24 Mus musculus | 35.96 | 6.9 | 23144 | 1 | 1 | 577564.44 | 582131.75 | 1717788.25 | 934448.88 | 599634.31 | 927576.56 | 578554.94 | 1 | 1 | 431319.5 | 677644.62 | 518951.47 | 565791.25 | 525364 |
| P23640 | Rab27a | Ras-related protein Rab-27A Rattus norvegicus | 40.34 | 9.5 | 25068 | 1 | 1 | 1 | 134758.64 | 1 | 235721.61 | 1 | 176932.94 | 115956.9175 | 114252.4 | 106408.08 | 66234.25 | 64311.07 | 139055.94 | 57906.12 | 87091.04333 |
| P05712 | Rab2a | Ras-related protein Rab-2A Rattus norvegicus | 527.07 | 46.7 | 23536 | 9 | 9 | 11523637 | 10334534 | 11464839 | 8168102.5 | 9611938 | 9970685 | 9201436 | 11952641 | 11459287 | 9164857 | 9805703 | 10705247 | 10798662 | 11026551 |
| Q921E2 | Rab31 | Ras-related protein Rab-31 Mus musculus | 240.24 | 24.74 | 21331 | 3 | 4 | 6154674.5 | 6432402 | 7534732 | 7578820 | 3406910 | 4770880 | 6409984 | 5417167 | 5123526.5 | 4286000.5 | 8369679 | 5144328.5 | 4917423 | 4531569 |
| Q9CZE3 | Rab32 | Ras-related protein Rab-32 Mus musculus | 80.73 | 10.76 | 25068 | 2 | 2 | 4747686 | 2531041 | 2179624.25 | 2626370.75 | 1742248.25 | 1194038.88 | 810264.69 | 2346547 | 1377834.738 | 1160488.38 | 1580524.647 | 680515.44 | 2482523.5 | 1578535 |
| Q5U316 | Rab35 | Ras-related protein Rab-35 Rattus norvegicus | 318.15 | 32.84 | 23025 | 4 | 5 | 8934905 | 7395600 | 6895737 | 6688692 | 7213246 | 6493565 | 7243260 | 8882143 | 7902543 | 7344410 | 7275829 | 6932378 | 7647407 | 8021994.5 |
| P35276 | Rab3d | Ras-related protein Rab-3D Mus musculus | 229.91 | 14.16 | 24416 | 1 | 4 | 19148383 | 13234569 | 13854055 | 15892462 | 13911381 | 13655751 | 9792860 | 14301369 | 11949833 | 14124969 | 14226634 | 10363902 | 14344537 | 13747196 |
| Q80UJ7 | Rab3gap1 | Rab3 GTPase-activating protein catalytic subunit Mus musculus | 267.42 | 8.87 | 110198 | 6 | 6 | 4529544 | 4017258 | 3924811 | 3890960.75 | 3313279.25 | 3953368 | 4103238.5 | 1874881.25 | 3060421 | 3145765 | 3139543.75 | 3064455 | 3872433 | 2804394.5 |
| Q8BMG7 | Rab3gap2 | Rab3 GTPase-activating protein non-catalytic subunit Mus musculus | 516.99 | 10.61 | 152535 | 11 | 11 | 3521419.5 | 2475348.75 | 3809159.5 | 3572298.75 | 4909346 | 3504498 | 3228419 | 2460833 | 3427038.5 | 3624842.5 | 4029483.75 | 3720721.75 | 3257369 | 2789358.75 |
| Q8VDV3 | Rab3il1 | Guanine nucleotide exchange factor for Rab-3A Mus musculus | 42.02 | 5.22 | 42713 | 1 | 1 | 243104.86 | 142138.69 | 184855.16 | 275147.28 | 229194.58 | 144101.17 | 202274.77 | 204438.53 | 219311.48 | 146518.58 | 176832.36 | 117017.09 | 121782.1 | 138543.85 |
| Q53B90 | Rab43 | Ras-related protein Rab-43 Rattus norvegicus | 119.83 | 11.9 | 23229 | 1 | 2 | 723732.56 | 825376.44 | 648969.88 | 710570.33 | 644202.44 | 236149.67 | 685413.69 | 691015.88 | 671416.25 | 507194.72 | 667025.81 | 701074.81 | 670895.75 | 755195.62 |
| P51146 | Rab4b | Ras-related protein Rab-4B Rattus norvegicus | 426.26 | 46.48 | 23629 | 6 | 7 | 3815972 | 3995803.75 | 3681148.5 | 4220570.5 | 4985689 | 4414588.5 | 3780604.5 | 1785728.25 | 5128584 | 3577163.5 | 3741757.5 | 4102743.25 | 4380031 | 5351160 |
| Q9CQD1 | Rab5a | Ras-related protein Rab-5A Mus musculus | 424.65 | 48.84 | 23599 | 5 | 7 | 14677289 | 14337149 | 14606747 | 13195073 | 13316318 | 14661658 | 12770437 | 18371149 | 17043645 | 14214683 | 13457352 | 14719951 | 15443541 | 14482086 |
| P61021 | Rab5b | Ras-related protein Rab-5B Mus musculus | 329.18 | 32.09 | 23707 | 3 | 5 | 14392134 | 12708080 | 13514980 | 10427497 | 12971203 | 13401993 | 11354175 | 16963905 | 15325401 | 10572072 | 11116746 | 13298754 | 12118493 | 12897300 |
| P35278 | Rab5c | Ras-related protein Rab-5C Mus musculus | 630.37 | 67.13 | 23413 | 7 | 9 | 41195041 | 37879833 | 39473465 | 38617997 | 43166841 | 41976853 | 36011437 | 38195041 | 38741149 | 35497709 | 39625289 | 38238021 | 40039521 | 41329769 |
| P35279 | Rab6a | Ras-related protein Rab-6A Mus musculus | 401.88 | 42.31 | 23590 | 7 | 8 | 1844885.25 | 528157 | 3234779.25 | 1951627.75 | 1326213.5 | 1009596.19 | 2113414.25 | 3148678.75 | 1879350.75 | 3030021 | 1653055.12 | 1962323.667 | 2399947 | 1833968.88 |
| P09527 | Rab7a | Ras-related protein Rab-7a Rattus norvegicus | 1025.46 | 68.12 | 23504 | 14 | 15 | 97731393 | 109525561 | 103554041 | 103969705 | 107382417 | 97950625 | 93470105 | 96716025 | 94470273 | 95391009 | 92300625 | 98579705 | 101028865 | 97031873 |
| Q8VEA8 | Rab7b | Ras-related protein Rab-7b Mus musculus | 62.26 | 7.54 | 22502 | 1 | 1 | 1112938.12 | 1039819.06 | 913769.69 | 477229.94 | 848847.12 | 883066.75 | 414661.91 | 1027524.44 | 854516.69 | 719842.12 | 1 | 1 | 456781.06 | 1022169.69 |
| P35280 | Rab8a | Ras-related protein Rab-8A Rattus norvegicus | 596.15 | 51.69 | 23668 | 5 | 11 | 5161799 | 6839662 | 3150111.75 | 7603442 | 4447727 | 7116540 | 6011053.5 | 7115734 | 7253591 | 5395729.5 | 6844491.5 | 6753198 | 5538119 | 5595862.5 |
| P61028 | Rab8b | Ras-related protein Rab-8B Mus musculus | 643.59 | 59.42 | 23603 | 6 | 12 | 35356293 | 35358469 | 28363077 | 31688473 | 30676647 | 32427861 | 31282267 | 34816377 | 31492777 | 33249341 | 31646665 | 26665685 | 29500465 | 32986119 |
| Q99P75 | Rab9a | Ras-related protein Rab-9A Rattus norvegicus | 123.28 | 23.38 | 22896 | 3 | 3 | 1211371.12 | 2999493.5 | 4264398.5 | 3507301.5 | 2999012.5 | 3408992.5 | 3464321.75 | 2549955.5 | 2107797 | 2761812 | 3208534 | 1822902.5 | 3355623.75 | 2984947.75 |
| O35550 | Rabep1 | Rab GTPase-binding effector protein 1 Rattus norvegicus | 57.62 | 2.2 | 99428 | 1 | 1 | 643260.19 | 581520.31 | 724240.25 | 902920.62 | 808063.19 | 847739.69 | 704762.06 | 816847.88 | 636271.56 | 701021.12 | 638967.81 | 531364.06 | 267122.31 | 699658.31 |
| A2AWA9 | Rabgap1 | Rab GTPase-activating protein 1 Mus musculus | 81.95 | 2.54 | 120798 | 2 | 2 | 935754.94 | 1237422.44 | 801623.69 | 920566.38 | 2291744.75 | 1802179.25 | 803121 | 970313.38 | 961705.62 | 1416385.25 | 1600996 | 1 | 1536799.25 | 1 |
| Q9JHK4 | Rabggta | Geranylgeranyl transferase type-2 subunit alpha Mus musculus | 99.73 | 7.41 | 64989 | 2 | 2 | 1014871.25 | 1558044.5 | 930421.31 | 1962574 | 1197761.5 | 1059949.62 | 1367643.38 | 991772.44 | 1000065.88 | 1420288.38 | 1243266 | 766758.12 | 1037066.31 | 1499016.5 |
| Q5U3K5 | Rabl6 | Rab-like protein 6 Mus musculus | 58.35 | 1.52 | 79831 | 1 | 1 | 532287.44 | 572339.12 | 348638.31 | 373884.62 | 453624.84 | 388109.69 | 432893.41 | 499226.69 | 536042.44 | 359213.56 | 461004.84 | 361310.44 | 361519.69 | 379698.62 |
| P63001 | Rac1 | Ras-related C3 botulinum toxin substrate 1 Mus musculus | 502.89 | 51.04 | 21450 | 4 | 8 | 117916473 | 124658545 | 104534513 | 105572913 | 111204041 | 96933737 | 109895873 | 116568033 | 102075617 | 107475577 | 105453833 | 100123505 | 99066161 | 114623313 |
| Q05144 | Rac2 | Ras-related C3 botulinum toxin substrate 2 Mus musculus | 504.12 | 51.04 | 21441 | 4 | 8 | 157320433 | 173500753 | 153848577 | 153689553 | 156357969 | 140826161 | 155543073 | 164822305 | 150324961 | 150706641 | 158518705 | 149084145 | 143950161 | 170259313 |
| Q9WVM1 | Racgap1 | Rac GTPase-activating protein 1 Mus musculus | 125.57 | 6.05 | 70158 | 3 | 3 | 888202.88 | 1045679.06 | 1100534 | 830627.69 | 1006828.88 | 915980.56 | 972833.015 | 1103713.62 | 1021134.94 | 850502.94 | 150301.17 | 1352241.88 | 1219921.25 | 1184574.5 |
| P63245 | Rack1 | Receptor of activated protein C kinase 1 Rattus norvegicus | 1368.58 | 86.75 | 35077 | 20 | 20 | 148829329 | 154126033 | 163496273 | 146850577 | 149176609 | 155000945 | 164161521 | 158051521 | 161792273 | 164118785 | 156720689 | 162680849 | 158131441 | 163378673 |
| Q61550 | Rad21 | Double-strand-break repair protein rad21 homolog Mus musculus | 172.78 | 9.13 | 72083 | 4 | 4 | 2132063.25 | 1589362.88 | 1646797 | 3037397.5 | 2517096 | 2439310 | 2226009.5 | 667846.38 | 1622668.75 | 2364144.25 | 2328071.5 | 2249252 | 2196316.5 | 692855.5 |
| P54726 | Rad23a | UV excision repair protein RAD23 homolog A Mus musculus | 123.61 | 8.26 | 39706 | 1 | 1 | 1758432 | 1334628.5 | 1559897.5 | 1908512 | 1431107.25 | 1618350.12 | 1429703.75 | 1085199.38 | 1363408.38 | 1362017.12 | 1884795.38 | 1459397.25 | 1444838.38 | 1436246.5 |
| P54728 | Rad23b | UV excision repair protein RAD23 homolog B Mus musculus | 179.3 | 11.54 | 43513 | 3 | 3 | 2478165.5 | 1950238.75 | 3278661.5 | 2996743.5 | 2192811.25 | 2272444.25 | 2590020.5 | 1725823 | 2397869 | 2663263.5 | 2554767 | 2150467.75 | 2597490.75 | 2823717.5 |
| P70388 | Rad50 | DNA repair protein RAD50 Mus musculus | 453.79 | 9.15 | 153488 | 9 | 9 | 3588007.25 | 3754498 | 5015912 | 4149389.75 | 4204187 | 4120013.75 | 4305095.5 | 4805534 | 4505090.5 | 3978328.25 | 4849859 | 4767959.5 | 5134719 | 4607815 |
| Q3SWS8 | Rae1 | mRNA export factor Rattus norvegicus | 158.18 | 14.13 | 40919 | 4 | 4 | 2581383.5 | 444001.88 | 1662393.38 | 2962797 | 2986501 | 860390.31 | 2075788.12 | 1947058.25 | 620311.12 | 3073885.5 | 2819787.5 | 3478671.25 | 2351596 | 514923.5 |
| O08602 | Raet1a | Retinoic acid early-inducible protein 1-alpha Mus musculus | 51.58 | 4.74 | 28616 | 1 | 1 | 2361884.5 | 2118501.75 | 1474918.38 | 1908892.25 | 1955331.5 | 1943966.12 | 1703417 | 1478562.62 | 1537715.75 | 1271626.75 | 2189614.25 | 1938704.003 | 1744971.88 | 1881525.88 |
| P11345 | Raf1 | RAF proto-oncogene serine/threonine-protein kinase Rattus norvegicus | 81.69 | 3.55 | 72928 | 1 | 2 | 1449602.75 | 1954012.88 | 1695082.62 | 1398639.25 | 1985536.25 | 1593021.44 | 1307289.38 | 1771232.38 | 1543036.38 | 1750527.62 | 1 | 1872756.75 | 2056207.75 | 1 |
| P63321 | Rala | Ras-related protein Ral-A Mus musculus | 343.61 | 40.29 | 23553 | 1 | 6 | 4510105.5 | 3817739.25 | 4884840 | 4392306.5 | 5976085 | 5684868 | 5480208 | 5789696.5 | 6492933.5 | 6141894 | 5749738 | 5880959 | 6322742 | 5327107.5 |
| Q9JIW9 | Ralb | Ras-related protein Ral-B Mus musculus | 396.86 | 50.97 | 23349 | 3 | 8 | 5540295 | 4588644.5 | 5215243 | 3503016 | 5817572.5 | 5698153 | 6410821 | 5284708 | 6098926.5 | 6945932 | 4794597 | 7527990.5 | 5237660 | 4962882.5 |
| A3KGS3 | Ralgapa2 | Ral GTPase-activating protein subunit alpha-2 Mus musculus | 100.01 | 1.28 | 210288 | 2 | 2 | 750213.25 | 618278.19 | 617803.38 | 1975379.62 | 214315.11 | 1009386.44 | 1242459.38 | 1138079.62 | 1147574.38 | 648871.44 | 926927.81 | 253879.12 | 941530.62 | 1256492.62 |
| Q8BQZ4 | Ralgapb | Ral GTPase-activating protein subunit beta Mus musculus | 219.53 | 4.45 | 165200 | 4 | 4 | 2820274.5 | 1817084.12 | 1522997.75 | 2239717.5 | 1341277.5 | 3129268 | 2549162 | 1710809.12 | 2958559.25 | 2776574.75 | 2020423.12 | 2214818.5 | 2681043 | 2664414.75 |
| Q64012 | Raly | RNA-binding protein Raly Mus musculus | 537.27 | 32.37 | 33188 | 11 | 11 | 13281002 | 14878608 | 14140905 | 15923953 | 16666228 | 14399887 | 15952720 | 14277783 | 12828080 | 16291995 | 18773815 | 13980631 | 20158063 | 11601357 |
| P62827 | Ran | GTP-binding nuclear protein Ran Mus musculus | 787.85 | 57.87 | 24423 | 13 | 13 | 25982069 | 24143423 | 22978157 | 25446659 | 33917089 | 31649953 | 25090959 | 17929871 | 27466481 | 34587717 | 34306845 | 33202431 | 36633933 | 27074795 |
| P34022 | Ranbp1 | Ran-specific GTPase-activating protein Mus musculus | 205.91 | 20.69 | 23596 | 5 | 5 | 6620613.5 | 7181381 | 2928177 | 7263486.5 | 6782964 | 5431049 | 5896757 | 5914569.5 | 4695309.5 | 6245819.5 | 6748401 | 4521783 | 6041180.5 | 6605476 |
| Q6VN19 | Ranbp10 | Ran-binding protein 10 Mus musculus | 57.29 | 2.26 | 67188 | 1 | 1 | 394196.38 | 405803.62 | 443727.66 | 964522.69 | 901931.25 | 1174076.25 | 621231.38 | 518469.41 | 896830.75 | 989880.56 | 756064.62 | 981124.88 | 1022283 | 896754.81 |
| Q9ERU9 | Ranbp2 | E3 SUMO-protein ligase RanBP2 Mus musculus | 1516.77 | 13.72 | 341121 | 27 | 27 | 11768863 | 10494381 | 14115749 | 10526775 | 10692732 | 12648369 | 13065762 | 12662127 | 12396402 | 11684288 | 12188053 | 13127760 | 14854322 | 13392306 |
| Q9CT10 | Ranbp3 | Ran-binding protein 3 Mus musculus | 397.34 | 18.53 | 52573 | 6 | 6 | 8239164.5 | 6848450 | 5475961.5 | 6928614.5 | 10246101 | 6239143 | 8725777 | 5810172 | 6211659 | 6436203.5 | 6729800.5 | 11629662 | 5288496 | 6901165.5 |
| P46061 | Rangap1 | Ran GTPase-activating protein 1 Mus musculus | 1367.74 | 45.16 | 63531 | 25 | 26 | 20220831 | 21796129 | 20268741 | 17210879 | 16143941 | 25560223 | 19038905 | 19511375 | 22229745 | 22030437 | 19152445 | 24596885 | 26512043 | 21576175 |
| P62835 | Rap1a | Ras-related protein Rap-1A Mus musculus | 974.52 | 76.63 | 20987 | 4 | 14 | 121260257 | 124766001 | 110734497 | 116421289 | 112651857 | 110300545 | 117541417 | 121290913 | 107380041 | 113540209 | 109046593 | 100297225 | 112175825 | 108177553 |
| Q99JI6 | Rap1b | Ras-related protein Rap-1b Mus musculus | 1023.91 | 80.43 | 20825 | 5 | 16 | 136927169 | 141910849 | 128245809 | 131854817 | 126793345 | 124197873 | 131392225 | 137265777 | 121309553 | 126138625 | 120258377 | 112723361 | 125830321 | 122650929 |
| P61226 | Rap2b | Ras-related protein Rap-2b Mus musculus | 358.41 | 48.09 | 20504 | 2 | 6 | 10758406 | 11178974 | 12366614 | 11726889 | 9633748 | 12616033 | 11057426 | 11726891 | 11377799 | 10686038 | 8569474 | 11698846 | 13506508 | 12126605 |
| Q8BU31 | Rap2c | Ras-related protein Rap-2c Mus musculus | 277.87 | 38.25 | 20745 | 1 | 5 | 10418436 | 17213223 | 9644153 | 13419739 | 15866324 | 9443428 | 12477081 | 19492157 | 8764909 | 8527868 | 10587180 | 11480187 | 10571028 | 15638559 |
| F1M386 | Rapgef2 | Rap guanine nucleotide exchange factor 2 Rattus norvegicus | 41.99 | 0.87 | 166388 | 1 | 1 | 480017.22 | 612132.38 | 555573.75 | 308647.25 | 472058.34 | 525986.31 | 478407.5 | 557074.62 | 604766.94 | 445686.75 | 502384.12 | 424431.5 | 576436.75 | 501084.1233 |
| Q9D0I9 | Rars | Arginine--tRNA ligase, cytoplasmic Mus musculus | 1781.69 | 47.12 | 75674 | 29 | 31 | 118466857 | 109909561 | 121923481 | 111019025 | 108712305 | 121495609 | 118799993 | 123335137 | 118570065 | 114079201 | 117580297 | 109688153 | 109519273 | 113838849 |
| Q3U186 | Rars2 | Probable arginine--tRNA ligase, mitochondrial Mus musculus | 190.86 | 4.84 | 65336 | 2 | 2 | 1508299.5 | 1395321.88 | 1814915.5 | 1228651.75 | 1364387 | 1553847.5 | 1678469.5 | 1270665.25 | 1632557.25 | 1407947.75 | 1403361 | 1657269.5 | 1633560 | 1671213.5 |
| Q60790 | Rasa3 | Ras GTPase-activating protein 3 Mus musculus | 189.07 | 4.68 | 95987 | 4 | 4 | 4543903 | 5110698.5 | 5025591 | 4775877 | 5900853 | 5467864 | 4200614.5 | 5124293 | 5233600.5 | 4967122.5 | 4642075 | 4810444 | 4853485 | 5553361 |
| Q6PFQ7 | Rasa4 | Ras GTPase-activating protein 4 Mus musculus | 72.49 | 4.49 | 90060 | 2 | 2 | 458248.4375 | 514896.69 | 410078.59 | 465511.75 | 442506.72 | 671430.88 | 271571.12 | 384198 | 506994.69 | 396801.72 | 1071865.25 | 415441.22 | 533491.12 | 535264.19 |
| Q8C2K5 | Rasal3 | RAS protein activator like-3 Mus musculus | 178.56 | 5.67 | 114782 | 4 | 4 | 836467.88 | 783370.44 | 924220 | 756868.38 | 873788.38 | 700514.12 | 1031493 | 940000.88 | 1075732.75 | 830025.75 | 868894.75 | 928542.25 | 772327.5 | 1187813.25 |
| Q8CB96 | Rassf4 | Ras association domain-containing protein 4 Mus musculus | 195.53 | 11.8 | 36934 | 3 | 3 | 1276281.25 | 966730 | 1242547.62 | 1017789.06 | 960419.12 | 973151.62 | 1253976.62 | 1448708.25 | 1220531.88 | 1121127.75 | 957063.75 | 1221374.75 | 1497744.75 | 1358754.75 |
| Q9CW46 | Raver1 | Ribonucleoprotein PTB-binding 1 Mus musculus | 387.71 | 21.66 | 79382 | 8 | 8 | 1 | 541926 | 1 | 1 | 1 | 16877637 | 808436.12 | 699444.5 | 473327.84 | 597908 | 1674871.25 | 524371.38 | 744044.69 | 1242835.5 |
| Q60972 | Rbbp4 | Histone-binding protein RBBP4 Mus musculus | 390.27 | 22.35 | 47656 | 3 | 7 | 26910333 | 26352515 | 26713375 | 27903791 | 29951127 | 25853221 | 24061037 | 25295337 | 24680683 | 26449649 | 26769591 | 26927769 | 26762301 | 25678185 |
| Q8BX09 | Rbbp5 | Retinoblastoma-binding protein 5 Mus musculus | 57.48 | 1.67 | 59098 | 1 | 1 | 1099447.25 | 1230452.38 | 1205071 | 1385181.62 | 1310407.38 | 1007460.56 | 1040931.75 | 1118567.38 | 1217478.62 | 899510.88 | 1101121.12 | 1124187.25 | 1141752 | 989748.69 |
| Q60973 | Rbbp7 | Histone-binding protein RBBP7 Mus musculus | 335.63 | 22.12 | 47790 | 2 | 6 | 21178625 | 20790347 | 20901695 | 22812941 | 24752673 | 20217631 | 19442929 | 18979021 | 18839291 | 22067653 | 21462917 | 21808947 | 21365573 | 20167611 |
| Q64701 | Rbl1 | Retinoblastoma-like protein 1 Mus musculus | 76.44 | 1.69 | 119456 | 2 | 2 | 853589.38 | 760856.25 | 1096405 | 1326140 | 1091439.62 | 724916.75 | 695267.5 | 731037.88 | 720605.31 | 1280884.38 | 1339007.38 | 1465271.25 | 809056.38 | 1380923.75 |
| Q8C2Q3 | Rbm14 | RNA-binding protein 14 Mus musculus | 435.13 | 22.57 | 69449 | 10 | 10 | 6665431.5 | 7423135.5 | 7817220.5 | 10178391 | 8662953 | 6909961 | 6427374 | 4012307 | 8163808.5 | 10630943 | 8375748.5 | 8934716 | 8504303 | 8616585 |
| Q0VBL3 | Rbm15 | RNA-binding protein 15 Mus musculus | 179.31 | 5.2 | 105722 | 4 | 4 | 1143650.88 | 1253638.5 | 1074182.25 | 1482568.12 | 1370062.12 | 1663988.5 | 1320482.88 | 1678503.75 | 566045.25 | 750183 | 1372498.5 | 697754.44 | 655920.38 | 1344504.88 |
| Q8JZX4 | Rbm17 | Splicing factor 45 Mus musculus | 142.07 | 8.15 | 45304 | 3 | 3 | 1852686.5 | 2012469.12 | 872493.62 | 2370725 | 1001566.62 | 779788.12 | 1880339 | 1328676.75 | 1908883.5 | 1912465.38 | 2026835.38 | 2185369 | 1870417 | 1939668.38 |
| Q8R3C6 | Rbm19 | Probable RNA-binding protein 19 Mus musculus | 102.13 | 2.94 | 106083 | 2 | 2 | 769180.94 | 791832.655 | 761084.06 | 633465 | 1003600.62 | 508960 | 570596.38 | 576935.88 | 849682.19 | 489423.47 | 1218492.5 | 1238463.75 | 413294.88 | 1371606.75 |
| Q4V7D7 | Rbm22 | Pre-mRNA-splicing factor RBM22 Rattus norvegicus | 79.83 | 4.52 | 46864 | 1 | 1 | 480776.72 | 1 | 393129.44 | 1 | 1 | 1 | 1 | 1 | 442813.25 | 1 | 1 | 338597.03 | 1 | 1 |
| B2RY56 | Rbm25 | RNA-binding protein 25 Mus musculus | 328.76 | 10.26 | 99552 | 6 | 6 | 3141130.75 | 2363907 | 3301986.25 | 2883145.25 | 3357609 | 3531563.5 | 2837237.5 | 2950668.25 | 3626819 | 3796397.25 | 2736737 | 2977742.5 | 3611710.25 | 3356857.5 |
| Q8CGC6 | Rbm28 | RNA-binding protein 28 Mus musculus | 440.11 | 13.87 | 84214 | 9 | 9 | 3444275 | 3734736.5 | 787139.31 | 3987158 | 445939.22 | 5192945.5 | 4845445 | 4122897 | 4916632 | 4967977 | 4456815.5 | 5441411.5 | 1491287.25 | 4301062 |
| O89086 | Rbm3 | RNA-binding protein 3 Mus musculus | 263.18 | 48.37 | 16605 | 4 | 4 | 16201317 | 14056728 | 17451029 | 17496437 | 18559707 | 11945609 | 12359547 | 9477812 | 12530498 | 14840802 | 15729302 | 10670674 | 14948069 | 14099241 |
| Q8C5L7 | Rbm34 | RNA-binding protein 34 Mus musculus | 36.19 | 2.67 | 41326 | 1 | 1 | 980492.62 | 577497 | 764433.94 | 691917.8125 | 445247.69 | 602839.88 | 1 | 915441.62 | 516310.41 | 327109.28 | 547236.06 | 512266.41 | 554962.94 | 483544.75 |
| Q8VH51 | Rbm39 | RNA-binding protein 39 Mus musculus | 850.65 | 34.91 | 59407 | 13 | 13 | 18583955 | 20376495 | 26137809 | 25612119 | 27527821 | 28558967 | 23279427 | 21786725 | 26185141 | 29039601 | 24683235 | 26591703 | 27292775 | 25996825 |
| Q91V81 | Rbm42 | RNA-binding protein 42 Mus musculus | 71.11 | 4.43 | 49875 | 2 | 2 | 1 | 1 | 360364.38 | 1 | 1 | 285877.91 | 1 | 1 | 1 | 1 | 1 | 1 | 385137.12 | 1 |
| Q91WT8 | Rbm47 | RNA-binding protein 47 Mus musculus | 43.03 | 3.39 | 64062 | 1 | 1 | 1 | 1 | 1 | 521404.94 | 318446.12 | 469429.09 | 294802.03 | 369411.405 | 429842.25 | 283572.25 | 373849.66 | 369137.66 | 531041.56 | 431347 |
| Q9CQT2 | Rbm7 | RNA-binding protein 7 Mus musculus | 51.72 | 6.79 | 30148 | 1 | 1 | 316903.72 | 1 | 1 | 1 | 1 | 1 | 1 | 1 | 1 | 1 | 244984.08 | 1 | 1 | 1 |
| Q27W01 | Rbm8a | RNA-binding protein 8A Rattus norvegicus | 202.12 | 47.7 | 19889 | 4 | 4 | 7136055.5 | 6256920 | 6194536 | 5901224 | 6468162 | 6044464 | 6361971.5 | 8376484 | 7414246.5 | 6380739 | 5590129 | 5377645 | 7287050.5 | 3577502.25 |
| Q5PQP1 | Rbms1 | RNA-binding motif, single-stranded-interacting protein 1 Rattus norvegicus | 48.7 | 3.72 | 44068 | 1 | 1 | 1 | 2622672.75 | 2923624 | 1 | 2065368.88 | 2145969.75 | 2536697.25 | 2493730.75 | 2442074.75 | 2349132.25 | 2212312.25 | 1980450.25 | 744044.69 | 2658458.75 |
| Q9WV02 | Rbmx | RNA-binding motif protein, X chromosome Mus musculus | 463.19 | 28.64 | 42301 | 0 | 11 | 14676479 | 14162876 | 11357965 | 15574562 | 17912887 | 14949673 | 8169956 | 8355552 | 10800339 | 18029281 | 14283444 | 13600159 | 18086159 | 17482703 |
| Q91VM5 | Rbmxl1 | RNA binding motif protein, X-linked-like-1 Mus musculus 2 | 628.15 | 36.08 | 42162 | 3 | 13 | 18824915 | 19790105 | 16963171 | 22123673 | 25133949 | 23610821 | 13313579 | 14844145 | 17118327 | 25673597 | 21483177 | 20805809 | 26182377 | 25058947 |

| | | | | | | | | | | | | | | | | | | | | | |
|---|---|---|---|---|---|---|---|---|---|---|---|---|---|---|---|---|---|---|---|---|---|
| P31266 | Rbpj | Recombining binding protein suppressor of hairless Mus musculus | 391.96 | 26.62 | 58537 | 8 | 8 | 25027323 | 23856861 | 19681669 | 24367335 | 24816885 | 23833755 | 20899685 | 23212105 | 20738229 | 21448057 | 21385609 | 18833831 | 22624391 | 19986417 |
| P62878 | Rbx1 | E3 ubiquitin-protein ligase RBX1 Mus musculus | 86.98 | 16.67 | 12274 | 1 | 1 | 4225062 | 4060234.75 | 3408895.25 | 3481492.75 | 4201550.5 | 4001337.25 | 3453596.75 | 3504902.5 | 3549188 | 3460731.5 | 3779895.25 | 3881681.75 | 3186963.75 | 4029857.5 |
| Q8VE37 | Rcc1 | Regulator of chromosome condensation Mus musculus | 336.85 | 21.62 | 44931 | 6 | 6 | 7333107 | 6421106 | 8114995.5 | 9350778 | 9880416 | 8501231 | 10006331 | 5686220 | 9436429 | 10754710 | 9390290 | 8530440 | 10215839 | 9193329 |
| Q8BK67 | Rcc2 | Protein RCC2 Mus musculus | 1140.43 | 57.31 | 55983 | 20 | 20 | 50144673 | 56313645 | 56713385 | 67145321 | 62236345 | 59612013 | 63095001 | 60331705 | 61171097 | 64095589 | 64433813 | 56839465 | 53675721 | 59873077 |
| Q9JJT0 | Rcl1 | RNA 3'-terminal phosphate cyclase-like protein Mus musculus | 110.28 | 6.17 | 40841 | 2 | 2 | 1951306.25 | 2081479 | 1325929.38 | 1148578.38 | 1117398.88 | 2168904 | 1544052.25 | 2226735.5 | 1626953.62 | 1863881 | 1669607.5 | 1810600.75 | 2338432.75 | 1586215.62 |
| Q8BP92 | Rcn2 | Reticulocalbin-2 Mus musculus | 534.81 | 36.56 | 37271 | 8 | 8 | 8223233.5 | 9635146 | 8256234.5 | 4061390 | 8449753 | 4427759 | 5130736 | 10361669 | 6461655.5 | 6740664.5 | 8756283 | 9774934 | 9570397 | 9438323 |
| Q3UZA1 | Rcsd1 | CapZ-interacting protein Mus musculus | 372.28 | 25.97 | 44114 | 7 | 7 | 22690795 | 6848788.5 | 5966656.5 | 6237873.5 | 8090971 | 5975495 | 6540070.5 | 5698964.5 | 5763556 | 7314162 | 7187590.5 | 6669904.5 | 6586964.5 | 5764597.5 |
| Q9QYF1 | Rdh11 | Retinol dehydrogenase 11 Mus musculus | 215.49 | 19.3 | 35148 | 5 | 5 | 6818532.5 | 5668518 | 6806729 | 6052273.5 | 5862307.5 | 5788113 | 7410207 | 5784552.5 | 6227919 | 6460265 | 7035384 | 6787682 | 7216069 | 6885310 |
| Q8CEE7 | Rdh13 | Retinol dehydrogenase 13 Mus musculus | 46.64 | 5.69 | 36464 | 1 | 1 | 229952.3425 | 182689.22 | 281971.94 | 185605.27 | 269542.94 | 218140.53 | 143963.72 | 145564.16 | 159058.86 | 234970.14 | 272393.47 | 421084.03 | 290959.5 | 249839.64 |
| P26043 | Rdx | Radixin Mus musculus | 1301.23 | 39.97 | 68543 | 10 | 24 | 451788545 | 429860481 | 379326657 | 426065889 | 406720193 | 449825505 | 443610081 | 455141857 | 424508577 | 427182881 | 423162561 | 374412257 | 351156321 | 388374273 |
| Q9Z129 | Recql | ATP-dependent DNA helicase Q1 Mus musculus | 37.86 | 1.7 | 72484 | 1 | 1 | 649946.69 | 684190.44 | 933678 | 810436.69 | 956424.56 | 855321.75 | 694015.56 | 767279.44 | 734663.94 | 872363.88 | 740500 | 939801.12 | 873973.06 | 922108.31 |
| B2RZ37 | Reep5 | Receptor expression-enhancing protein 5 Rattus norvegicus | 61.03 | 5.29 | 21431 | 1 | 1 | 7432805 | 3583975.75 | 3614411 | 3200738.75 | 4175538.75 | 745461 | 3847124.75 | 4512975.5 | 3408630.5 | 4916669.5 | 4596934.5 | 3382864.5 | 4149302 | 4476232 |
| P15307 | Rel | Proto-oncogene c-Rel Mus musculus | 41.15 | 1.87 | 64960 | 1 | 1 | 1 | 490613.88 | 1 | 1 | 1 | 1 | 220641.67 | 533814.31 | 199414.69 | 1 | 1 | 1 | 1 | 403024.94 |
| Q04207 | Rela | Transcription factor p65 Mus musculus | 82.81 | 8.93 | 60212 | 2 | 2 | 211088.44 | 1 | 145839.23 | 194205.16 | 1 | 131083.5 | 1 | 1 | 1 | 1 | 120918.61 | 137576.08 | 1 | 1 |
| Q148V7 | Relch | RAB11-binding protein RELCH Mus musculus | 98.82 | 1.97 | 134587 | 2 | 2 | 423175.78 | 438693.66 | 405497.84 | 502120.75 | 370819.09 | 425751.66 | 390608.44 | 628622 | 453931.75 | 322783.03 | 424424.47 | 304179.19 | 326230.38 | 265227.56 |
| P82343 | Renbp | N-acylglucosamine 2-epimerase Mus musculus | 372.96 | 21.16 | 49771 | 9 | 9 | 27482973 | 22888361 | 22671903 | 21481823 | 21501491 | 22207463 | 21423515 | 21410497 | 20568213 | 19921253 | 21980685 | 18911397 | 22334797 | 23548221 |
| Q5FVM3 | Retreg1 | Reticulophagy regulator 1 Rattus norvegicus | 70.42 | 2.5 | 52782 | 1 | 1 | 1 | 635444.75 | 1 | 477877.72 | 390176.22 | 329958.44 | 380819.38 | 1 | 1 | 322977.16 | 381749.56 | 1 | 1 | 1 |
| Q3MHU5 | Retreg2 | Reticulophagy regulator 2 Rattus norvegicus | 58.32 | 1.48 | 57659 | 1 | 1 | 1 | 1 | 999634.12 | 1 | 862420.62 | 833744.94 | 619573.19 | 1 | 1 | 650487.38 | 704452.62 | 1004293.31 | 1138898.5 | 1055240.75 |
| Q64FW2 | Retsat | All-trans-retinol 13,14-reductase Mus musculus | 284.96 | 14.29 | 67334 | 5 | 5 | 2062565.88 | 16080405 | 1844958.62 | 1249876.62 | 25397993 | 12083215 | 727752.69 | 439154.84 | 1190675.25 | 394738.84 | 22935765 | 19898829 | 9723112 | 1186289 |
| Q5U1X1 | Rexo2 | Oligoribonuclease, mitochondrial Rattus norvegicus | 127.73 | 10.97 | 26752 | 2 | 2 | 4250126 | 4397954 | 4659792 | 4377616 | 4009168.25 | 4005421 | 4893656.5 | 4043452.25 | 4082092.25 | 3809945 | 4522037 | 4574626 | 4346259.5 | 4065159 |
| P35601 | Rfc1 | Replication factor C subunit 1 Mus musculus | 223.36 | 5.48 | 125985 | 4 | 4 | 1754153.25 | 1955838.12 | 1638564.38 | 2245255 | 1607385 | 1327430.38 | 1172661.25 | 1716483.5 | 1406396 | 2037559.5 | 2247734.25 | 1859288.75 | 1571918.75 | 1368622 |
| Q9WUK4 | Rfc2 | Replication factor C subunit 2 Mus musculus | 214.45 | 16.91 | 38725 | 4 | 4 | 1973131.25 | 1699930.5 | 2027881.75 | 4414021 | 4219553 | 3747767.75 | 3594989.5 | 1499403.88 | 3676161.75 | 3671283 | 4170287.75 | 4713720 | 2010284.25 | 4348209 |
| Q8R323 | Rfc3 | Replication factor C subunit 3 Mus musculus | 208.7 | 11.24 | 40526 | 4 | 4 | 3114739.25 | 2834600.5 | 2661804 | 1520486.12 | 3500486 | 2542238.5 | 1635788 | 3234662.25 | 3106420.5 | 2284862 | 3878876 | 2719913.5 | 3171179.75 | 2480173 |
| Q99J62 | Rfc4 | Replication factor C subunit 4 Mus musculus | 248.48 | 19.51 | 39867 | 5 | 5 | 6632325 | 10782153 | 7629577 | 9015264 | 9698691 | 8412445 | 7897178 | 7496632.5 | 5901751.5 | 7005553 | 10301413 | 10776182 | 8064885 | 9415825 |
| Q9D0F6 | Rfc5 | Replication factor C subunit 5 Mus musculus | 257.05 | 27.14 | 38096 | 6 | 6 | 3489320.75 | 3808726.5 | 3205226.5 | 3610095 | 3384961.5 | 3267691.75 | 3179072.25 | 3530496.25 | 3351914.75 | 3485231.25 | 3853191.25 | 2912322.5 | 3948673 | 3533524.5 |
| Q8CFV9 | Rfk | Riboflavin kinase Mus musculus | 168.51 | 13.55 | 17437 | 2 | 2 | 1046277.62 | 1499406.25 | 1439349.75 | 909639.38 | 1100302.12 | 1210135.5 | 1249712.75 | 1425552.25 | 1756330 | 1103329.5 | 1179302.88 | 848715.62 | 1518680 | 1110916.75 |
| Q6A0D4 | Rftn1 | Raftlin Mus musculus | 128.03 | 8.12 | 61537 | 3 | 3 | 2560086.25 | 167247.33 | 1422655.25 | 2004712.62 | 2072418 | 2007123.25 | 1915276.75 | 1981017.75 | 1448974.88 | 2064164.5 | 1960786.5 | 171995.66 | 1745184.38 | 1752015.88 |
| Q9CQE5 | Rgs10 | Regulator of G-protein signaling 10 Mus musculus | 92.63 | 9.39 | 21151 | 1 | 1 | 1260703.12 | 1249538.62 | 880824.56 | 927223.12 | 954264.5 | 1194361.12 | 830873.38 | 1216615.75 | 794764 | 833320.19 | 1173830.25 | 1005756.06 | 971311.75 | 834866.19 |
| O70521 | Rgs19 | Regulator of G-protein signaling 19 Rattus norvegicus | 39.26 | 6.02 | 24738 | 1 | 1 | 1425710 | 1221182 | 1282710.75 | 1493605.38 | 1355802.033 | 1674472.62 | 1040793.31 | 1141803.12 | 1199762.88 | 373861.97 | 730590.94 | 771096.56 | 1325314.62 | 1411150.88 |
| Q4V8F3 | Rhbdd1 | Rhomboid-related protein 4 Rattus norvegicus 2 | 55.79 | 4.43 | 35879 | 1 | 1 | 1 | 1 | 1 | 446310.66 | 277795.25 | 1 | 1 | 361274.25 | 1 | 1 | 468928.41 | 421210.03 | 327254.56 | 467447.12 |
| Q921J2 | Rheb | GTP-binding protein Rheb Mus musculus | 214.44 | 28.26 | 20451 | 5 | 5 | 3202839 | 3763590 | 2978251.25 | 2856487.5 | 1729558.62 | 1826587 | 3346840.5 | 2920390.5 | 2360367.5 | 3031434.75 | 2969828.5 | 2769939 | 3229484.5 | 3378129.5 |
| P61589 | Rhoa | Transforming protein RhoA Rattus norvegicus | 759.83 | 65.8 | 21782 | 3 | 12 | 78086985 | 48889153 | 73282449 | 55225093 | 51837897 | 89901969 | 53881713 | 93197585 | 93387537 | 58282101 | 55266349 | 56710973 | 80893145 | 79369449 |
| P62746 | Rhob | Rho-related GTP-binding protein RhoB Mus musculus | 196.09 | 25.51 | 22123 | 1 | 3 | 30660373 | 5411296.5 | 30118673 | 6870403 | 5860114.5 | 35684453 | 4726419 | 41020785 | 39526321 | 5491591 | 5908808 | 7926000 | 31621375 | 27933861 |
| Q62159 | Rhoc | Rho-related GTP-binding protein RhoC Mus musculus | 747.29 | 65.8 | 22006 | 3 | 12 | 72336065 | 43518089 | 68619497 | 49710129 | 46555773 | 84762873 | 47014469 | 89086345 | 90406553 | 52769393 | 48762885 | 52776137 | 77380177 | 74319897 |
| P84096 | Rhog | Rho-related GTP-binding protein RhoG Mus musculus | 660.33 | 53.93 | 21309 | 8 | 9 | 17318869 | 18855237 | 15860312 | 18122857 | 17035473 | 16161437 | 16233627 | 15829496 | 15434019 | 18079195 | 15473371 | 15595804 | 16825773 | 16405257 |
| Q8BG51 | Rhot1 | Mitochondrial Rho GTPase 1 Mus musculus | 186.76 | 6.02 | 72242 | 3 | 4 | 334383.47 | 1 | 295852.75 | 214383.38 | 1 | 269978.56 | 1 | 361517.31 | 1 | 1 | 174936.28 | 1 | 1 | 326268.97 |
| Q7TSA0 | Rhot2 | Mitochondrial Rho GTPase 2 Rattus norvegicus | 193.54 | 6.27 | 69153 | 2 | 3 | 501147.69 | 595937.88 | 646770.06 | 642362.8775 | 825595.88 | 583478.75 | 818704.62 | 543883.19 | 619611.75 | 672592.62 | 776390.38 | 527982.81 | 860306.75 | 653893.31 |
| P52760 | Rida | 2-iminobutanoate/2-iminopropanoate deaminase Mus musculus | 102.6 | 17.78 | 14255 | 2 | 2 | 1087577.62 | 1302952.12 | 983554.19 | 1607621.75 | 397875.16 | 617955.19 | 935076.25 | 978087.3925 | 1010456.88 | 1348861.25 | 1705665 | 1315891.62 | 1495393.38 | 1005124.12 |
| Q6PR54 | Rif1 | Telomere-associated protein RIF1 Mus musculus | 189.22 | 2.32 | 266228 | 3 | 3 | 1624920.38 | 2046007.75 | 1606687.25 | 2072905.25 | 1487789.88 | 1624002.75 | 1508187.25 | 1167074.75 | 738457.38 | 696476.94 | 1867916.12 | 2190692.5 | 666812.62 | 704889.44 |
| Q9QZL0 | Ripk3 | Receptor-interacting serine/threonine-protein kinase 3 Mus musculus | 404.16 | 23.25 | 53322 | 7 | 7 | 13043554 | 12830318 | 15241674 | 3231674.75 | 4065368.25 | 16570341 | 13207548 | 14094715 | 14488202 | 18136541 | 21205087 | 5071155 | 4953180.5 | 19077547 |
| P13832 | Rlc-a | Myosin regulatory light chain RLC-A Rattus norvegicus | 681.86 | 79.65 | 19895 | 4 | 11 | 52345889 | 55833101 | 55399397 | 55234433 | 52837229 | 63959177 | 53457025 | 70549169 | 67313065 | 62092353 | 59591005 | 57391145 | 56817985 | 52933953 |
| Q9CWY8 | Rnaseh2a | Ribonuclease H2 subunit A Mus musculus | 54.23 | 5.32 | 33513 | 1 | 1 | 453792.31 | 325170.91 | 500514.62 | 436523.62 | 482824.31 | 405257.88 | 463679.75 | 323573.22 | 405772.5 | 531020.31 | 561576.88 | 547638.56 | 625447.44 | 485281.5 |
| Q5DTM8 | Rnf20 | E3 ubiquitin-protein ligase BRE1A Mus musculus | 98.04 | 2.77 | 113520 | 2 | 2 | 1 | 161327.36 | 247613.83 | 1 | 1 | 171936.27 | 1 | 211574.17 | 210762.36 | 1 | 138290.4767 | 176706.44 | 130838.01 | 107326.98 |
| E9Q555 | Rnf213 | E3 ubiquitin-protein ligase RNF213 Mus musculus | 2031.58 | 11.9 | 584784 | 43 | 43 | 45362837 | 41682497 | 32155981 | 35600493 | 38076313 | 29922537 | 28911147 | 32672189 | 31496833 | 33959937 | 33480975 | 34904065 | 35576017 | 33303313 |
| Q91VI7 | Rnh1 | Ribonuclease inhibitor Mus musculus | 2471.85 | 90.79 | 49816 | 29 | 29 | 331530049 | 365746049 | 323439425 | 375143137 | 344204865 | 375934497 | 387887553 | 393568545 | 360855105 | 377228801 | 383652609 | 331784801 | 336977025 | 385898241 |
| Q5U2U7 | Rnmt | mRNA cap guanine-N7 methyltransferase Rattus norvegicus | 147.61 | 8.03 | 52817 | 3 | 3 | 3920096.75 | 3663730.25 | 3442175.5 | 3841588.75 | 4448384.5 | 4762323 | 4325686 | 4186189.5 | 4655614.5 | 5594118 | 2992147.25 | 4340487.5 | 4163963.5 | 4423773 |
| Q8VCT3 | Rnpep | Aminopeptidase B Mus musculus | 1563.82 | 54.46 | 72416 | 26 | 26 | 62735149 | 67485457 | 71716025 | 64039325 | 62866457 | 59147557 | 56775197 | 60981129 | 66397173 | 57192385 | 57437581 | 55898233 | 56846157 | 62337717 |
| P70336 | Rock2 | Rho-associated protein kinase 2 Mus musculus | 116.68 | 2.67 | 160586 | 3 | 3 | 918762.94 | 1955630 | 1393872.88 | 1431697.12 | 2111055.5 | 1768919.75 | 717646.31 | 1527776.38 | 1489864.25 | 1048571 | 1941292.25 | 1841606.75 | 975364.06 | 1225059.38 |
| P60603 | Romo1 | Reactive oxygen species modulator 1 Mus musculus | 113.24 | 51.9 | 8183 | 2 | 2 | 507002.25 | 705462.4375 | 827640.75 | 435317 | 1051889.75 | 1166872 | 1148360.25 | 501032.97 | 1334988 | 998214.88 | 1392542.75 | 1779152.5 | 1073323.62 | 1407164.12 |
| Q63132 | Ros1 | Proto-oncogene tyrosine-protein kinase ROS Rattus norvegicus | 71.5 | 0.64 | 261656 | 1 | 2 | 2297278.75 | 1518774.38 | 1538482.75 | 466991.91 | 1850511.62 | 1467977 | 1576639.75 | 1269041.12 | 1597570.5 | 1973535.12 | 1662670.5 | 1345331.12 | 1844025.12 | 1817708.25 |
| Q9EPK2 | Rp2 | Protein XRP2 Mus musculus | 108.05 | 5.19 | 39377 | 2 | 2 | 1473764.12 | 1049215.62 | 1728861.25 | 1075393 | 1417357.62 | 1881287.25 | 1547189.88 | 1698362 | 1905032.25 | 1485710.38 | 1560972.75 | 1685879.75 | 1925046 | 1611516.5 |
| Q8VEE4 | Rpa1 | Replication protein A 70 kDa DNA-binding subunit Mus musculus | 839.18 | 26.16 | 69037 | 15 | 15 | 17327145 | 21230773 | 24112441 | 18150669 | 19528737 | 18194885 | 21971593 | 19455383 | 22008115 | 26827429 | 16815173 | 17989909 | 23237553 | 24170723 |
| Q62193 | Rpa2 | Replication protein A 32 kDa subunit Mus musculus | 182.28 | 21.11 | 29718 | 4 | 4 | 8180673.5 | 5909737 | 7714308 | 10611604 | 9788269 | 5166029 | 8415718 | 5322136 | 8033280.5 | 7922520 | 5832450 | 9017262 | 8079228 | 9111018 |
| Q9CQ71 | Rpa3 | Replication protein A 14 kDa subunit Mus musculus | 118.83 | 32.23 | 13584 | 3 | 3 | 2110414.75 | 2540167 | 2299456 | 2579842.75 | 2428874.5 | 2542425.75 | 1808940.12 | 2707742.75 | 2255835.5 | 2479132.75 | 2440230.5 | 2642599.25 | 2685059 | 2504029.5 |
| Q80TE0 | Rpap1 | RNA polymerase II-associated protein 1 Mus musculus | 44.06 | 0.92 | 155270 | 1 | 1 | 394524.53 | 429404.41 | 408864.81 | 426645.22 | 414859.7425 | 420624.12 | 313191.75 | 1 | 1 | 297633.09 | 462139.41 | 394690.84 | 367269.12 | 330774.84 |
| Q9D706 | Rpap3 | RNA polymerase II-associated protein 3 Mus musculus | 165.13 | 6.82 | 74096 | 3 | 3 | 1504026.88 | 1642477.12 | 1195072.5 | 1533197.62 | 233922.06 | 271351.66 | 1320487.25 | 1259984.25 | 1133065.38 | 844387 | 1011683.94 | 1203497.75 | 929315.12 | 1052584.38 |
| Q8VEE0 | Rpe | Ribulose-phosphate 3-epimerase Mus musculus | 281.86 | 36.84 | 24945 | 4 | 4 | 2575407.75 | 2132401 | 2770585 | 2216684 | 2219400 | 2620099.5 | 1159043.38 | 2550320 | 2092764.75 | 2249094.25 | 2611857.5 | 2369330.5 | 1132308.75 | 2256393.25 |
| Q9JJ80 | Rpf2 | Ribosome production factor 2 homolog Mus musculus | 333.42 | 27.45 | 35364 | 6 | 6 | 1278550 | 1473422 | 1305544.5 | 2019469.38 | 2440933.5 | 895740.5775 | 1150534.88 | 196185.8 | 817088.38 | 1419153.25 | 767881.38 | 1024291.88 | 1101454.75 | 1302622.88 |
| P47968 | Rpia | Ribose-5-phosphate isomerase Mus musculus | 282.23 | 28.38 | 32451 | 7 | 7 | 9771853 | 5273146.5 | 4866029.5 | 9437076 | 9593231 | 6238491.5 | 8722519 | 4623844 | 8587214 | 6948304 | 9082331 | 7635780.5 | 7638889 | 5729791 |
| Q6PDV7 | Rpl10 | 60S ribosomal protein L10 Rattus norvegicus | 504.9 | 38.79 | 24604 | 8 | 8 | 19296765 | 18216813 | 22525409 | 20898781 | 20913821 | 18357499 | 20872635 | 16537775 | 22874817 | 26265045 | 23473355 | 19895573 | 24381733 | 22515333 |
| P62907 | Rpl10a | 60S ribosomal protein L10a Rattus norvegicus | 740.84 | 58.53 | 24831 | 13 | 13 | 14175484 | 14322817 | 21082419 | 16076777 | 13926860 | 13646501 | 18363437 | 11437047 | 14281019 | 19380569 | 14440603 | 19751861 | 18383619 | 13892302 |
| P62914 | Rpl11 | 60S ribosomal protein L11 Rattus norvegicus | 577.37 | 62.92 | 20252 | 11 | 11 | 32958047 | 26836899 | 41043341 | 41170781 | 33928033 | 32519909 | 34145937 | 26190701 | 35814949 | 41687269 | 38601877 | 42250273 | 39086705 | 37871501 |
| P35979 | Rpl12 | 60S ribosomal protein L12 Mus musculus | 490.93 | 51.52 | 17805 | 7 | 7 | 42948813 | 36512005 | 40066069 | 40067593 | 38795725 | 37956381 | 35114249 | 35012577 | 33777105 | 38570625 | 40836957 | 34807781 | 32567541 | 36495265 |
| P41123 | Rpl13 | 60S ribosomal protein L13 Rattus norvegicus | 265.56 | 25.59 | 24309 | 5 | 5 | 9368862 | 6131949 | 9752551 | 10905688 | 8065040 | 8343442 | 8685224 | 5909537 | 7411223.5 | 11712983 | 11206875 | 10573486 | 11217897 | 9305945 |
| P19253 | Rpl13a | 60S ribosomal protein L13a Mus musculus | 449.35 | 33.99 | 23464 | 8 | 9 | 19012237 | 11100493 | 13220825 | 13650727 | 4404996 | 10323142 | 12812241 | 6199949 | 10260033 | 9108587 | 10300401 | 11540810 | 7432724 | 7936385 |
| Q9CR57 | Rpl14 | 60S ribosomal protein L14 Mus musculus | 266.01 | 25.81 | 23564 | 5 | 5 | 48757397 | 41464161 | 49147209 | 50989849 | 40586785 | 38326293 | 48761021 | 48600781 | 44754981 | 47072129 | 50495561 | 45415893 | 45584653 | 47164221 |
| P61314 | Rpl15 | 60S ribosomal protein L15 Rattus norvegicus | 608.47 | 45.1 | 24146 | 11 | 11 | 18648701 | 13333976 | 11521345 | 9381727 | 11246411 | 10970105 | 11839691 | 13222983 | 13866968 | 11049573 | 12634757 | 13495125 | 13448799 | 10343294 |
| P24049 | Rpl17 | 60S ribosomal protein L17 Rattus norvegicus | 292.67 | 30.98 | 21397 | 5 | 5 | 17367001 | 13188777 | 15337442 | 14684893 | 12363700 | 11750297 | 13129854 | 12689103 | 12675760 | 13875902 | 15320755 | 12671805 | 14121331 | 13604093 |
| P35980 | Rpl18 | 60S ribosomal protein L18 Mus musculus | 527.2 | 49.47 | 21645 | 10 | 10 | 41226201 | 23076019 | 20713081 | 23666485 | 27125053 | 25453359 | 19141195 | 111476409 | 75858425 | 24918877 | 22044389 | 21080801 | 23894619 | 21633829 |
| P62717 | Rpl18a | 60S ribosomal protein L18a Mus musculus | 352.19 | 33.52 | 20732 | 8 | 8 | 27571991 | 17590765 | 18432631 | 25090469 | 25397873 | 20786763 | 21180055 | 11587269 | 20801873 | 19521421 | 26103553 | 21309243 | 16311888 | 20920901 |
| P84099 | Rpl19 | 60S ribosomal protein L19 Mus musculus | 308.5 | 18.37 | 23466 | 5 | 5 | 2841524.75 | 3748496.25 | 4027587.5 | 3882654.5 | 4441050 | 4410996.5 | 4222289 | 4074969 | 4142975.5 | 5389346.5 | 4766608.5 | 4733162.5 | 4929564.5 | 4162395.25 |
| O09167 | Rpl21 | 60S ribosomal protein L21 Mus musculus | 244.37 | 32.5 | 18562 | 5 | 5 | 23035485 | 21861553 | 28818509 | 33698633 | 30063827 | 25515007 | 29182547 | 19122313 | 29058135 | 32859735 | 32841345 | 31798663 | 29412965 | 29586589 |
| P67984 | Rpl22 | 60S ribosomal protein L22 Mus musculus | 387.03 | 56.25 | 14759 | 7 | 7 | 35659789 | 27045523 | 32735147 | 24856975 | 31397327 | 38167421 | 37770341 | 32239589 | 44082929 | 43481093 | 36101401 | 45934985 | 42751321 | 39497965 |
| Q9D7S7 | Rpl22l1 | 60S ribosomal protein L22-like 1 Mus musculus | 252.54 | 45.08 | 14467 | 4 | 4 | 8332675.5 | 4641697 | 4296884 | 4652856 | 5900927 | 3076762 | 5109935 | 4053164 | 4014869 | 5415731 | 5259283.5 | 5328888.5 | 5068624 | 4639584 |
| P62830 | Rpl23 | 60S ribosomal protein L23 Mus musculus | 397.75 | 62.86 | 14865 | 8 | 8 | 78422417 | 81408617 | 92665329 | 89340329 | 84629801 | 79281513 | 81363609 | 71833233 | 80579713 | 97478993 | 84910913 | 85564233 | 88287473 | 86464241 |
| P62751 | Rpl23a | 60S ribosomal protein L23a Mus musculus | 345.1 | 50.64 | 17695 | 8 | 8 | 12156759 | 9817307 | 7802661.5 | 17052715 | 8816283 | 8796988 | 10084652 | 5807919 | 6815126 | 8608746 | 9943242 | 11738982 | 7291911 | 7721673 |
| P83732 | Rpl24 | 60S ribosomal protein L24 Rattus norvegicus | 279.62 | 32.48 | 17779 | 5 | 5 | 7935524.5 | 6400114 | 8622589 | 8188637.5 | 6570155.5 | 7521173.5 | 8329627 | 6313278 | 8354251 | 9841903 | 8755254 | 10570493 | 9929607 | 7331159.5 |
| P61255 | Rpl26 | 60S ribosomal protein L26 Mus musculus | 467.96 | 41.38 | 17258 | 9 | 9 | 16771569 | 15726351 | 22666187 | 5770501 | 6629578 | 19158551 | 8285819 | 19671061 | 8744828 | 18594623 | 9464413 | 9527111 | 23065755 | 9449594 |
| P61354 | Rpl27 | 60S ribosomal protein L27 Rattus norvegicus | 213.29 | 36.03 | 15798 | 4 | 4 | 1205200.12 | 1757168.703 | 2642859.5 | 2145694.5 | 1034920.69 | 1755464.25 | 1444608.38 | 2997683.533 | 2407433.5 | 6383228 | 2539342 | 2258738 | 2682524.5 | 2292432.25 |
| P14115 | Rpl27a | 60S ribosomal protein L27a Mus musculus | 371 | 33.11 | 16605 | 7 | 7 | 40037089 | 42905105 | 45102537 | 44705289 | 43530089 | 42901021 | 41043189 | 40867605 | 38300317 | 41492233 | 45804581 | 42898969 | 42430217 | 44067653 |
| P41105 | Rpl28 | 60S ribosomal protein L28 Mus musculus | 337.58 | 32.85 | 15733 | 6 | 6 | 7195704.5 | 6425701 | 9476703 | 11688524 | 6312725 | 10949361 | 9922475 | 6213435 | 8220374 | 10123376 | 10463190 | 11281607 | 8442668 | 5295299 |
| P47915 | Rpl29 | 60S ribosomal protein L29 Mus musculus | 86.32 | 11.25 | 17587 | 2 | 2 | 21712929 | 26435781 | 36960413 | 33214665 | 36677345 | 29749517 | 32249885 | 14893884 | 32346519 | 38448185 | 38515825 | 39887737 | 35837237 | 42919817 |
| P27659 | Rpl3 | 60S ribosomal protein L3 Mus musculus | 1434.59 | 52.85 | 46110 | 22 | 24 | 23059385 | 17892201 | 22967791 | 31215469 | 29197835 | 19868665 | 20011443 | 12321592 | 18014563 | 27963059 | 28234433 | 27324929 | 23692239 | 27726637 |
| P62889 | Rpl30 | 60S ribosomal protein L30 Mus musculus | 490.07 | 59.13 | 12784 | 7 | 7 | 33088765 | 25803993 | 29098033 | 50881621 | 34721301 | 42038097 | 43337721 | 33311781 | 33524711 | 36011785 | 49254385 | 42480697 | 48393921 | 42459081 |
| P62900 | Rpl31 | 60S ribosomal protein L31 Mus musculus | 186.39 | 30.4 | 14463 | 4 | 4 | 4916891 | 3206677.5 | 4019639.5 | 4328282 | 3507651.75 | 7912629 | 3819074.25 | 4297766 | 5413332 | 4859778.5 | 11445761 | 4115789.75 | 3976763.25 | 3768962.25 |
| P62911 | Rpl32 | 60S ribosomal protein L32 Mus musculus | 393.86 | 40 | 15860 | 7 | 7 | 5341341.5 | 3066487.5 | 6492211.5 | 7782269 | 5765616 | 5948176 | 4217205 | 4303519.5 | 5228459 | 6223996 | 5966632 | 6989915 | 5704111 | 6330589 |
| P11250 | Rpl34 | 60S ribosomal protein L34 Rattus norvegicus | 82.51 | 12.82 | 13507 | 2 | 2 | 575018.31 | 1489885.5 | 1470175.75 | 2556895.25 | 1522993.703 | 1557151.12 | 2449011.25 | 1967081.5 | 2236431 | 1948343.25 | 1918670.12 | 2866290.5 | 1568118.25 | 1541056 |
| P17078 | Rpl35 | 60S ribosomal protein L35 Rattus norvegicus | 190.89 | 21.95 | 14553 | 4 | 4 | 1 | 646843.44 | 667667.94 | 1050045.75 | 1 | 652525.5 | 498645.53 | 607419.1475 | 615461.81 | 663043.75 | 1 | 1 | 240358.97 | 646721.5 |
| O55142 | Rpl35a | 60S ribosomal protein L35a Mus musculus | 114.96 | 20.91 | 12554 | 3 | 3 | 8128167.5 | 4272391 | 4715124.5 | 7664477 | 4816295.5 | 4665048 | 6574308 | 4816208 | 4958555.5 | 5382703.5 | 5985015 | 5110698 | 5199384 | 4421944 |

| | | | | | | | | | | | | | | | | | | | | | |
|---|---|---|---|---|---|---|---|---|---|---|---|---|---|---|---|---|---|---|---|---|---|
| P47964 | Rpl36 | 60S ribosomal protein L36 Mus musculus | 82.39 | 18.1 | 12216 | 1 | 2 | 4759367 | 2821063 | 5226443 | 4375304.5 | 4081034.75 | 5004962 | 4172033 | 2621019 | 5024376 | 6551808 | 5349980 | 5856201 | 6328044 | 5137186 |
| P39032 | Rpl36 | 60S ribosomal protein L36 Rattus norvegicus | 105.28 | 19.05 | 12268 | 1 | 2 | 1348309.25 | 1191118.5 | 2931916.5 | 1890920.88 | 1829570 | 2413493 | 1940014.62 | 831572.5 | 2733209 | 3822650.5 | 2577084.5 | 3087368.75 | 3257983 | 2456311 |
| P61514 | Rpl37a | 60S ribosomal protein L37a Mus musculus | 169.49 | 41.3 | 10275 | 3 | 3 | 632743.88 | 1557243.75 | 1481535.38 | 558657.25 | 1394191.38 | 1340081 | 2167878.25 | 866885.88 | 593854.69 | 2009041.75 | 1966053.88 | 2050467.38 | 2335211.75 | 1423280.75 |
| P62892 | Rpl39 | 60S ribosomal protein L39 Mus musculus | 35.58 | 19.61 | 6407 | 1 | 1 | 2724630.5 | 2267559 | 2507127.75 | 2371130.75 | 2485049.75 | 3037738.75 | 2288886 | 1699666.12 | 2708701.75 | 3465083.75 | 4062534.25 | 3798987 | 3846852.75 | 3811686.25 |
| Q9D8E6 | Rpl4 | 60S ribosomal protein L4 Mus musculus | 1226.3 | 46.54 | 47154 | 20 | 20 | 68622793 | 50576013 | 61833941 | 60178089 | 56715809 | 56216201 | 63157993 | 50197481 | 60003061 | 72401361 | 73724481 | 62216353 | 83259225 | 76912265 |
| P47962 | Rpl5 | 60S ribosomal protein L5 Mus musculus | 654.47 | 50.51 | 34401 | 12 | 12 | 58611145 | 60605537 | 58989681 | 61306309 | 64149569 | 58394697 | 58030329 | 57719881 | 55266201 | 62940493 | 62573301 | 58307649 | 61931361 | 68640377 |
| P47911 | Rpl6 | 60S ribosomal protein L6 Mus musculus | 687.47 | 44.59 | 33510 | 14 | 14 | 26706893 | 18230993 | 25584587 | 25728261 | 23738183 | 18253813 | 20075385 | 17753393 | 20640687 | 24639165 | 23606081 | 23339339 | 23538133 | 22963931 |
| P14148 | Rpl7 | 60S ribosomal protein L7 Mus musculus | 1284.82 | 58.15 | 31420 | 23 | 25 | 82860465 | 55623253 | 44417761 | 54578425 | 63209765 | 61777797 | 58980561 | 47279921 | 45244109 | 68382937 | 67387681 | 60787941 | 64940137 | 64638861 |
| P12970 | Rpl7a | 60S ribosomal protein L7a Mus musculus | 828.14 | 49.62 | 29977 | 15 | 15 | 42082725 | 27784061 | 32498789 | 45993913 | 39162697 | 30351185 | 33519313 | 29548047 | 27356493 | 33905289 | 39964217 | 39412809 | 28413589 | 29571235 |
| Q9D8M4 | Rpl7l1 | 60S ribosomal protein L7-like 1 Mus musculus | 317.61 | 41.06 | 28544 | 8 | 8 | 1269463.12 | 1459463.75 | 1023326.69 | 886850 | 715230.25 | 656685.12 | 1260726.75 | 835337.75 | 793694.75 | 630244.38 | 577287 | 804319.31 | 810441.81 | 721006.88 |
| P62918 | Rpl8 | 60S ribosomal protein L8 Mus musculus | 396.23 | 21.79 | 28025 | 7 | 7 | 7948369 | 6989144 | 12437464 | 14813648 | 12839873 | 14171821 | 11276857 | 8037877 | 12268135 | 17090935 | 14073904 | 16884893 | 12058295 | 12805594 |
| P51410 | Rpl9 | 60S ribosomal protein L9 Mus musculus | 1059.24 | 73.44 | 21881 | 15 | 15 | 46457813 | 29655009 | 42141465 | 32684757 | 33831709 | 36912285 | 31004215 | 28434267 | 37050233 | 43065349 | 36654593 | 41975333 | 42276873 | 36203369 |
| P14869 | Rplp0 | 60S acidic ribosomal protein P0 Mus musculus | 871.61 | 54.57 | 34216 | 13 | 13 | 218328369 | 246103585 | 248938705 | 213388145 | 213936561 | 228115889 | 241813345 | 250953809 | 221770497 | 234878721 | 224239393 | 219696241 | 220902161 | 247026321 |
| P47955 | Rplp1 | 60S acidic ribosomal protein P1 Mus musculus | 340.33 | 80.7 | 11475 | 5 | 5 | 82563521 | 83746257 | 79999273 | 84038497 | 79610681 | 85263017 | 87640593 | 100162977 | 82591833 | 76376265 | 87001601 | 80582377 | 84314505 | 80665201 |
| P99027 | Rplp2 | 60S acidic ribosomal protein P2 Mus musculus | 500.73 | 71.3 | 11651 | 6 | 6 | 5981313 | 4798400 | 7109557.5 | 4406071 | 4282963 | 6758006 | 4949369.5 | 6089794 | 5037934 | 5682939 | 6251004.5 | 7119221 | 5676588 | 4156047.75 |
| Q91YQ5 | Rpn1 | Dolichyl-diphosphooligosaccharide--protein glycosyltransferase subunit 1 Mus musculus | 1447.2 | 51.15 | 68528 | 24 | 24 | 61956117 | 57415889 | 67091837 | 56183701 | 55390873 | 63202921 | 61102873 | 61665709 | 61489753 | 67087325 | 62038637 | 59683601 | 65534817 | 64984201 |
| Q9DBG6 | Rpn2 | Dolichyl-diphosphooligosaccharide--protein glycosyltransferase subunit 2 Mus musculus | 1455.44 | 47.7 | 69063 | 21 | 21 | 33773069 | 32964589 | 43071273 | 40588133 | 38263321 | 39727945 | 31172481 | 41089361 | 43769737 | 31852215 | 38338917 | 39829653 | 45066933 | 43129441 |
| O88796 | Rpp30 | Ribonuclease P protein subunit p30 Mus musculus | 248.8 | 21.64 | 29473 | 4 | 4 | 3400004 | 2658113.25 | 3338017.5 | 2545187 | 3049643.25 | 3153082.5 | 2364173.25 | 2412198.5 | 2954417 | 2845346.25 | 2787481.75 | 2744133 | 3430294.25 | 2863863 |
| Q9CSU0 | Rprd1b | Regulation of nuclear pre-mRNA domain-containing protein 1B Mus musculus | 228.51 | 25.46 | 36884 | 6 | 6 | 2211920.5 | 4216523 | 1878051.5 | 4398867 | 4087899 | 4996502 | 3983533 | 5526581 | 3935644.5 | 2187943.25 | 4427882 | 3840262.5 | 4272323 | 3899013 |
| P63325 | Rps10 | 40S ribosomal protein S10 Mus musculus | 322.71 | 32.73 | 18916 | 5 | 5 | 5408785 | 3677941.5 | 4942438 | 4542353 | 4919256.5 | 3337112 | 3996923.5 | 2954962 | 3794745.5 | 4962527 | 4016354.5 | 4094672.25 | 3868029.25 | 4223237 |
| P62281 | Rps11 | 40S ribosomal protein S11 Mus musculus | 365.91 | 37.34 | 18431 | 8 | 8 | 24565721 | 29397857 | 39459553 | 38108433 | 37294617 | 36984009 | 34504213 | 24323863 | 35775441 | 46687985 | 39572737 | 41059757 | 44456165 | 43636773 |
| P63323 | Rps12 | 40S ribosomal protein S12 Mus musculus | 395.05 | 59.09 | 14525 | 6 | 7 | 32113307 | 32623473 | 36488805 | 37181425 | 36133601 | 27592399 | 30623793 | 30973905 | 33338717 | 33002849 | 38690493 | 29140253 | 34990609 | 38670009 |
| P62278 | Rps13 | 40S ribosomal protein S13 Rattus norvegicus | 488.7 | 47.02 | 17222 | 10 | 10 | 50209561 | 26923841 | 31766423 | 28352915 | 23271535 | 25501325 | 29971673 | 28443075 | 25375293 | 28435823 | 23520063 | 23760177 | 25107873 | 23775687 |
| P13471 | Rps14 | 40S ribosomal protein S14 Rattus norvegicus | 499.57 | 65.56 | 16259 | 8 | 8 | 24958129 | 28438065 | 32976529 | 30938233 | 32210651 | 23963889 | 23938817 | 23095375 | 25913605 | 32166437 | 27028761 | 32907517 | 28382013 | 30163745 |
| P62843 | Rps15 | 40S ribosomal protein S15 Mus musculus | 308.52 | 36.55 | 17040 | 3 | 5 | 5904843 | 3562673.5 | 6260171 | 3366951.75 | 3147139.5 | 3467115.75 | 6199225 | 4943792 | 2653688 | 6557672 | 3258470.5 | 5983780 | 4175878.75 | 3951011.5 |
| P62245 | Rps15a | 40S ribosomal protein S15a Mus musculus | 446.73 | 64.62 | 14840 | 8 | 8 | 80841425 | 61381577 | 69723017 | 67028045 | 58379629 | 56527573 | 66882385 | 59267709 | 60462269 | 66496937 | 60736193 | 61009081 | 59927249 | 56940721 |
| P14131 | Rps16 | 40S ribosomal protein S16 Mus musculus | 608.94 | 62.33 | 16445 | 13 | 13 | 96580201 | 62943681 | 77330849 | 81807889 | 73851537 | 68730409 | 74413265 | 59361857 | 68832793 | 71864257 | 74085633 | 87562041 | 75489249 | 69744049 |
| P63276 | Rps17 | 40S ribosomal protein S17 Mus musculus | 574.73 | 65.93 | 15524 | 9 | 9 | 11185217 | 9556584 | 12078351 | 11516377 | 10479789 | 11973393 | 10872885 | 9580313 | 10516548 | 13787366 | 11793397 | 10542473 | 11870279 | 12319415 |
| P62270 | Rps18 | 40S ribosomal protein S18 Mus musculus | 546.05 | 58.55 | 17719 | 11 | 11 | 21092969 | 16196550 | 24059113 | 21873423 | 18565451 | 15633325 | 19671921 | 17596285 | 16111175 | 23485857 | 18777943 | 20466865 | 18932775 | 17793641 |
| P17074 | Rps19 | 40S ribosomal protein S19 Rattus norvegicus | 377.39 | 35.17 | 16085 | 7 | 7 | 25781753 | 23033861 | 20522993 | 22798165 | 21703593 | 18428705 | 12208375 | 20073651 | 16947313 | 19788967 | 20618567 | 18086761 | 16976985 | 21485765 |
| P25444 | Rps2 | 40S ribosomal protein S2 Mus musculus | 707.16 | 39.25 | 31231 | 13 | 13 | 38287237 | 37984777 | 32127023 | 37984445 | 31504255 | 37412013 | 38632489 | 38622413 | 38414121 | 37596253 | 39171945 | 39306797 | 35591513 | 33803425 |
| P60867 | Rps20 | 40S ribosomal protein S20 Mus musculus | 86.11 | 19.33 | 13373 | 2 | 2 | 4084804.5 | 2527764 | 3224031.75 | 4230608 | 3628275 | 3088463.75 | 4058210 | 3194136 | 2664253.75 | 5687191 | 3401830.5 | 2638313.25 | 4084986.25 | 3260228.25 |
| P05765 | Rps21 | 40S ribosomal protein S21 Rattus norvegicus | 259.31 | 57.83 | 9127 | 4 | 4 | 18322141 | 16647120 | 20788409 | 23679899 | 23835877 | 19692825 | 21536481 | 13799885 | 21762251 | 25776495 | 21893393 | 21610883 | 20375281 | 19469069 |
| P62267 | Rps23 | 40S ribosomal protein S23 Mus musculus | 265.94 | 41.96 | 15808 | 4 | 4 | 2338638.5 | 443509.69 | 2379563.5 | 2421914.5 | 933587.94 | 1545059.25 | 2119195.5 | 413704.44 | 1649495.5 | 2579466 | 2698958.75 | 2659552.75 | 514554.56 | 1396913.75 |
| P62849 | Rps24 | 40S ribosomal protein S24 Mus musculus | 342.18 | 35.34 | 15423 | 5 | 6 | 21076983 | 11508957 | 13300076 | 16061965 | 12982588 | 11940313 | 12907333 | 11065839 | 10999725 | 14399359 | 14481738 | 12684614 | 12011935 | 12829326 |
| P62852 | Rps25 | 40S ribosomal protein S25 Mus musculus | 258.58 | 27.2 | 13742 | 5 | 5 | 122681505 | 131242185 | 122062353 | 114899049 | 110873033 | 104163545 | 123359305 | 123159793 | 111545337 | 108463985 | 108224241 | 112941793 | 108367105 | 118590769 |
| P62855 | Rps26 | 40S ribosomal protein S26 Mus musculus | 134.38 | 20.87 | 13015 | 2 | 2 | 4817003.5 | 3925846.25 | 6285140 | 6238324 | 6075653.5 | 6061371 | 5884579.5 | 3701841.25 | 6521393 | 8143887 | 6364207.5 | 7958699.5 | 6904395 | 5765214 |
| P62982 | Rps27a | Ubiquitin-40S ribosomal protein S27a Rattus norvegicus | 749.14 | 59.62 | 17951 | 12 | 12 | 311169857 | 368934945 | 305579585 | 356052225 | 354830369 | 355956481 | 344759841 | 283426881 | 283629825 | 340558625 | 294045697 | 285526913 | 310916897 | 349564481 |
| P24051 | Rps27l | 40S ribosomal protein S27-like Rattus norvegicus | 163.95 | 26.19 | 9477 | 3 | 3 | 1373994 | 717258.62 | 1610744.88 | 2520221.75 | 1822715 | 2187784.25 | 1355050 | 469071.44 | 1909959.75 | 3233539 | 856323.19 | 1609148.25 | 3020824.5 | 2170615.75 |
| P62858 | Rps28 | 40S ribosomal protein S28 Mus musculus | 111.9 | 30.43 | 7841 | 2 | 2 | 22935545 | 19365419 | 21594133 | 26559035 | 24158677 | 22967261 | 25031587 | 15950723 | 24377699 | 24783297 | 28671353 | 26250101 | 23636305 | 25605005 |
| P62908 | Rps3 | 40S ribosomal protein S3 Mus musculus | 1028.02 | 74.07 | 26674 | 20 | 20 | 197884705 | 163660033 | 167921617 | 181191601 | 179688833 | 160192993 | 167194129 | 165947825 | 164712465 | 169294337 | 162641201 | 166227921 | 162666801 | 163382225 |
| P49242 | Rps3a | 40S ribosomal protein S3a Rattus norvegicus | 740.66 | 45.83 | 29945 | 13 | 13 | 113750105 | 112958609 | 114948289 | 118346137 | 109531193 | 97042241 | 105823233 | 100710585 | 100679497 | 105432593 | 102109929 | 103039873 | 99281617 | 107522273 |
| P62702 | Rps4x | 40S ribosomal protein S4, X isoform Mus musculus | 909.89 | 59.7 | 29598 | 17 | 17 | 111978305 | 91429985 | 114099345 | 111929257 | 95494737 | 96954161 | 109846865 | 100516521 | 107897569 | 109868193 | 110327345 | 109817297 | 105173545 | 104167537 |
| P97461 | Rps5 | 40S ribosomal protein S5 Mus musculus | 837 | 55.88 | 22889 | 13 | 13 | 65677265 | 68578081 | 71479009 | 69414161 | 76406825 | 65859193 | 63720657 | 44518953 | 60103913 | 67808585 | 69797105 | 62037685 | 56744709 | 69129137 |
| P62754 | Rps6 | 40S ribosomal protein S6 Mus musculus | 498.01 | 31.33 | 28681 | 8 | 8 | 31294497 | 32319381 | 26106187 | 33848281 | 29148607 | 22841215 | 28185259 | 29275837 | 25526705 | 26722381 | 27496945 | 27597167 | 26922425 | 30837573 |
| P18654 | Rps6ka3 | Ribosomal protein S6 kinase alpha-3 Mus musculus | 192.51 | 5.27 | 83694 | 2 | 4 | 2134561.5 | 1042357 | 1728925.38 | 765146.88 | 1434409.5 | 1806452.12 | 1505325 | 1788807.38 | 1934354.88 | 2015100.88 | 728953.06 | 1952128.75 | 1737135.88 | 671345.06 |
| P62082 | Rps7 | 40S ribosomal protein S7 Mus musculus | 430.09 | 40.72 | 22127 | 9 | 9 | 30112991 | 31808681 | 27780311 | 29600301 | 27330341 | 27460111 | 28216081 | 27897335 | 22558743 | 26054755 | 27909013 | 26803595 | 26789205 | 26277635 |
| P62242 | Rps8 | 40S ribosomal protein S8 Mus musculus | 677.39 | 50 | 24205 | 11 | 11 | 155997169 | 147219825 | 142502753 | 145336257 | 133317769 | 137815041 | 160755873 | 143044817 | 157288593 | 152950289 | 140931889 | 153140129 | 136486289 | 142622369 |
| P29314 | Rps9 | 40S ribosomal protein S9 Rattus norvegicus | 615.29 | 52.06 | 22591 | 14 | 14 | 177725201 | 71877393 | 87435513 | 51817165 | 70549825 | 66300573 | 77066945 | 71742353 | 67691057 | 75106833 | 71298313 | 77242649 | 69086257 | 69439697 |
| P14206 | Rpsa | 40S ribosomal protein SA Mus musculus | 1142.58 | 61.36 | 32838 | 15 | 17 | 75961185 | 67448417 | 76124585 | 82350089 | 72655481 | 76484601 | 74797905 | 63828141 | 70010433 | 87640377 | 81877129 | 86934121 | 80245825 | 74102681 |
| Q8K4Q0 | Rptor | Regulatory-associated protein of mTOR Mus musculus | 117.3 | 2.17 | 149471 | 2 | 2 | 1317334 | 1095099.75 | 1190953.5 | 589439.56 | 1241394 | 1528155.5 | 1061907.25 | 1362429 | 1340412.5 | 976426.5 | 1007920.38 | 985496.5 | 1315070.88 | 1328477.38 |
| Q63486 | Rraga | Ras-related GTP-binding protein A Rattus norvegicus | 415.81 | 34.5 | 36566 | 7 | 7 | 4424732 | 7776330.5 | 7697051.5 | 6519489 | 6985390 | 5458246 | 7712093 | 5879208.5 | 6135253 | 6634883.5 | 6934926.5 | 7734793 | 5599527 | 7433568 |
| Q99K70 | Rragc | Ras-related GTP-binding protein C Mus musculus | 579.47 | 33.67 | 44121 | 10 | 10 | 30362663 | 37790445 | 35462301 | 25095525 | 24505407 | 35297493 | 28472099 | 43400821 | 35874337 | 28206119 | 27750103 | 33281641 | 31283489 | 31364475 |
| D3Z8L7 | Rras | Ras-related protein R-Ras Rattus norvegicus | 235.96 | 28.9 | 23909 | 5 | 5 | 5490828.5 | 9192823 | 5410132.5 | 5420434.5 | 6203214 | 9938922 | 10849354 | 12619552 | 10807559 | 9097382 | 6464193 | 2961916.5 | 8718141 | 10152219 |
| Q99PL5 | Rrbp1 | Ribosome-binding protein 1 Mus musculus | 1522.1 | 24.05 | 172879 | 29 | 29 | 50452633 | 47263069 | 54767909 | 48342281 | 48508321 | 43475609 | 54210289 | 41473865 | 49621749 | 49621917 | 47471433 | 46100657 | 46324645 | 47358665 |
| P07742 | Rrm1 | Ribonucleoside-diphosphate reductase large subunit Mus musculus | 647.75 | 26.64 | 90210 | 14 | 14 | 18246387 | 12284210 | 5937449 | 16716314 | 16616034 | 11217080 | 8490664 | 5081146 | 5170175 | 9090302 | 19011979 | 18595471 | 7972857.5 | 4428564.5 |
| P11157 | Rrm2 | Ribonucleoside-diphosphate reductase subunit M2 Mus musculus | 298.68 | 14.36 | 45096 | 4 | 4 | 4144637.5 | 3025292.5 | 5096769.5 | 3848701.5 | 3711027.5 | 3489877 | 4140944.75 | 3164510 | 4756618.5 | 3837410.25 | 3506086.25 | 4230147.5 | 3032513 | 3479197.5 |
| P56183 | Rrp1 | Ribosomal RNA processing protein 1 homolog A Mus musculus | 284.55 | 21.26 | 54777 | 5 | 5 | 2734320.75 | 2070033.75 | 3952067.5 | 3494916.75 | 1842751.38 | 2777072.5 | 3667832.5 | 3176385.5 | 4898189 | 1765203.88 | 2186773.75 | 1602726.38 | 2413223.25 | 2432487.25 |
| Q6P5B0 | Rrp12 | RRP12-like protein Mus musculus | 114.21 | 2.63 | 143131 | 2 | 2 | 756523.94 | 144750.52 | 304117.66 | 425822.5375 | 497898.03 | 340315.34 | 207384.28 | 401052.59 | 351334.81 | 315130.16 | 483323.75 | 140157.33 | 725532.25 | 353174.66 |
| Q5M947 | Rrp15 | RRP15-like protein Rattus norvegicus | 68.1 | 6.43 | 30926 | 2 | 2 | 771086.56 | 1 | 843622.56 | 1 | 1 | 1 | 1 | 1 | 1 | 1 | 1335347.75 | 1088045.5 | 1 | 1 |
| Q91YK2 | Rrp1b | Ribosomal RNA processing protein 1 homolog B Mus musculus | 268.17 | 10.22 | 80582 | 5 | 5 | 5406735 | 5591422.5 | 6015206 | 4887104 | 5657125 | 5680460.5 | 5776165 | 4659601.5 | 5671608 | 5779362.5 | 6169642.5 | 5619362.5 | 6177638 | 5611289 |
| Q9DB85 | Rrp8 | Ribosomal RNA-processing protein 8 Mus musculus | 46.41 | 3.28 | 51066 | 1 | 1 | 759697 | 1 | 742256.19 | 1 | 794526.88 | 809297.19 | 661495.81 | 729786.8125 | 739539 | 708815.25 | 869511.81 | 835177.04 | 698630.62 | 937388.69 |
| Q91WM3 | Rrp9 | U3 small nucleolar RNA-interacting protein 2 Mus musculus | 86.69 | 6.95 | 52107 | 2 | 2 | 408973.44 | 324433.41 | 385185.69 | 323987.12 | 362741.88 | 363768.81 | 254776.73 | 345943.28 | 323460.28 | 305244.66 | 354939.38 | 277021.9633 | 337508.03 | 138618.48 |
| Q9CYH6 | Rrs1 | Ribosome biogenesis regulatory protein homolog Mus musculus | 233.57 | 16.16 | 41552 | 5 | 5 | 649089.12 | 891159 | 1558807.5 | 415914.56 | 497779.84 | 1192246.62 | 573035 | 444660.97 | 684062.38 | 1355823.75 | 1103355 | 1039195.88 | 767875.5 | 988087.12 |
| Q8BVY0 | Rsl1d1 | Ribosomal L1 domain-containing protein 1 Mus musculus | 603.76 | 22.79 | 50421 | 9 | 9 | 37531885 | 21401785 | 28499665 | 50338025 | 40785601 | 25286479 | 22207909 | 11306976 | 13842887 | 17548941 | 46628805 | 59814081 | 19116073 | 34891213 |
| Q6P6G7 | Rsl24d1 | Probable ribosome biogenesis protein RLP24 Rattus norvegicus | 72.87 | 11.66 | 19653 | 2 | 2 | 612981.56 | 618317 | 2277953.25 | 1096974 | 815260.75 | 654614.38 | 2290005 | 2253042.75 | 312476.38 | 598047.62 | 1908214.12 | 923315.94 | 1965786.12 | 2110475.25 |
| Q01730 | Rsu1 | Ras suppressor protein 1 Mus musculus | 361.67 | 45.85 | 31550 | 7 | 7 | 9417672 | 10143515 | 9792885 | 10924646 | 8617757 | 10902979 | 9898228 | 12114781 | 10509603 | 8678998 | 10131274 | 10823439 | 10090083 | 10501422 |
| Q9D7H3 | RtcA | RNA 3'-terminal phosphate cyclase Mus musculus | 274.63 | 17.21 | 39254 | 4 | 4 | 2739914.75 | 2188632.5 | 2828345.75 | 2486478.5 | 1789373.12 | 2258879.75 | 2261028.75 | 2560717.5 | 2276850.75 | 2548361 | 2290744.25 | 2354767.5 | 2535308.75 | 1989705.25 |
| Q6AYT3 | Rtcb | tRNA-splicing ligase RtcB homolog Rattus norvegicus | 579.08 | 28.71 | 55249 | 12 | 12 | 15347944 | 16400637 | 20071873 | 16082753 | 16561951 | 17855515 | 20343665 | 19262173 | 22609541 | 20570229 | 13212056 | 15771848 | 19672145 | 17958699 |
| Q3T1J8 | Rtfdc1 | Protein RTF2 homolog Rattus norvegicus 2 | 40.84 | 3.27 | 33812 | 1 | 1 | 663241.5 | 444095.25 | 635196.69 | 287354.75 | 386546.75 | 696939.62 | 662056.62 | 648334.69 | 797921.56 | 732653.44 | 732385.5 | 616243.31 | 723199.69 | 583197.5 |
| Q9ES97 | Rtn3 | Reticulon-3 Mus musculus | 186.54 | 6.54 | 103879 | 3 | 3 | 7813480 | 4670133 | 4226623.5 | 3397479.75 | 4866151 | 5460758 | 4717418 | 5880096.5 | 4951937 | 5562293 | 4875845.5 | 4377161.5 | 4512028 | 4505568 |
| Q99P72 | Rtn4 | Reticulon-4 Mus musculus | 984.33 | 25.13 | 126613 | 15 | 15 | 62349229 | 58925973 | 62964929 | 54204753 | 69060793 | 77515713 | 77575649 | 67254601 | 75303641 | 79078345 | 60750797 | 72058809 | 76561441 | 69924809 |
| Q9CQE8 | RTRAF | RNA transcription, translation and transport factor protein Mus musculus | 513.98 | 40.57 | 28152 | 8 | 8 | 10452018 | 7446543.5 | 8612863 | 5779640.5 | 6114199 | 8419569 | 7892765.5 | 8606508 | 9317207 | 6919245 | 8692324 | 8076177 | 7372744 | 8563543 |
| Q8R4Y8 | Rttn | Rotatin Mus musculus | 37.11 | 0.31 | 248489 | 1 | 1 | 1948464.75 | 1 | 1000575.56 | 658666.75 | 1 | 622075.14 | 1063315.25 | 474824.88 | 405978.62 | 544181.81 | 1 | 555914.44 | 1 | 1 |
| Q8BIJ7 | Rufy1 | RUN and FYVE domain-containing protein 1 Mus musculus | 408.16 | 14.19 | 80376 | 9 | 9 | 7251219 | 8579220 | 7904312.5 | 9282424 | 8046382 | 7465190.5 | 8049068.5 | 6698813 | 7506735 | 7873976 | 8982408 | 7438015 | 7870773 | 8350417 |
| P60122 | Ruvbl1 | RuvB-like 1 Mus musculus | 878.56 | 42.76 | 50214 | 15 | 15 | 34250041 | 33843885 | 35951393 | 33710441 | 32442131 | 32382427 | 32418757 | 31658555 | 32795217 | 34361025 | 34724649 | 32296227 | 34629281 | 34055821 |
| Q9WTM5 | Ruvbl2 | RuvB-like 2 Mus musculus | 1156.47 | 54.64 | 51113 | 21 | 21 | 159727153 | 116194353 | 119336737 | 173795409 | 156698625 | 120122153 | 92422225 | 79221297 | 78944489 | 94008201 | 158645713 | 185310657 | 94165385 | 130346329 |
| Q9CQK7 | Rwdd1 | RWD domain-containing protein 1 Mus musculus | 94.49 | 14.4 | 27785 | 2 | 2 | 419287.75 | 1 | 464638 | 1 | 491980.84 | 1 | 545362.5 | 479848.34 | 1 | 1 | 320070.5233 | 301537.97 | 372884.66 | 285788.94 |
| Q8CCI5 | Rybp | RING1 and YY1-binding protein Mus musculus | 79.92 | 18.86 | 24777 | 2 | 2 | 285203.12 | 581921.25 | 569095.38 | 500089.66 | 575833.75 | 506635.5 | 509766.25 | 489872.28 | 649436.06 | 614346.12 | 560596.75 | 787871.38 | 537960.38 | 560080.12 |
| P56565 | S100a1 | Protein S100-A1 Mus musculus | 101.41 | 19.15 | 10505 | 2 | 2 | 967942.06 | 1518875.62 | 417689.59 | 1440948.75 | 1295038.62 | 923617.25 | 1443401.88 | 743047.44 | 617673.12 | 954656 | 1226134.25 | 720953.69 | 805898.88 | 1143547.25 |
| P08207 | S100a10 | Protein S100-A10 Mus musculus | 143.58 | 27.84 | 11186 | 2 | 2 | 19057881 | 19975681 | 17181357 | 18627387 | 24212561 | 20283257 | 22772837 | 13471245 | 21829737 | 19694877 | 21623449 | 16526499 | 14373387 | 20090125 |
| P50543 | S100a11 | Protein S100-A11 Mus musculus | 300.34 | 54.08 | 11083 | 4 | 5 | 8823502 | 10569310 | 9505049 | 9836684 | 10115149 | 8173393 | 9684372 | 11436662 | 10383303 | 11045240 | 11260471 | 8111805.5 | 8643204 | 10935395 |
| P07091 | S100a4 | Protein S100-A4 Mus musculus | 540 | 84.16 | 11721 | 7 | 9 | 342641921 | 365499169 | 282173601 | 455036193 | 395784161 | 268835713 | 293508449 | 299471361 | 275043617 | 350356801 | 353867841 | 287454017 | 252285617 | 295674209 |
| P05964 | S100a6 | Protein S100-A6 Rattus norvegicus | 373.2 | 64.04 | 10035 | 5 | 5 | 147885137 | 153946993 | 109438433 | 153265377 | 135110529 | 116443825 | 122622409 | 150441217 | 98141889 | 126531505 | 131004505 | 105661081 | 94549401 | 131179385 |
| Q9D2C2 | Saal1 | Protein SAAL1 Mus musculus | 228.47 | 10.55 | 52769 | 4 | 4 | 145683.84 | 853157.88 | 868802.88 | 552912.88 | 495066.44 | 439290.5 | 848563.19 | 385618.19 | 972504.31 | 479796.5 | 1025801.62 | 935400.25 | 900051.75 | 513396.62 |
| Q9EP69 | Sacm1l | Phosphatidylinositide phosphatase SAC1 Mus musculus | 310.02 | 14.48 | 66944 | 6 | 6 | 4200392 | 4747371.5 | 4502654.5 | 5046367 | 4558957 | 5242934 | 4826618 | 4720028 | 5886921 | 3820288.5 | 4704008 | 4717182 | 4840087 | 5507722.5 |
| Q9R1T2 | Sae1 | SUMO-activating enzyme subunit 1 Mus musculus | 764.91 | 47.14 | 38620 | 13 | 13 | 17726315 | 19295161 | 16435100 | 20360735 | 21586009 | 19983213 | 19155439 | 10992209 | 18791621 | 20927213 | 19329247 | 18306223 | 17297817 | 19430945 |
| D3YXK2 | Safb | Scaffold attachment factor B1 Mus musculus | 378.03 | 15.58 | 105104 | 4 | 8 | 18522413 | 19448347 | 18781235 | 17714511 | 19606143 | 20507755 | 19990853 | 22320199 | 20586181 | 18884721 | 18167147 | 16904299 | 18881665 | 18297649 |
| Q80YR5 | Safb2 | Scaffold attachment factor B2 Mus musculus | 411.82 | 10.8 | 111838 | 4 | 7 | 14648402 | 16451889 | 15362969 | 13104848 | 15903931 | 16523070 | 15490727 | 17747965 | 17071947 | 14953437 | 13514055 | 14414525 | 15542565 | 15235882 |

| | | | | | | | | | | | | | | | | | | | | | |
|---|---|---|---|---|---|---|---|---|---|---|---|---|---|---|---|---|---|---|---|---|---|
| Q0VE29 | Samd12 | Sterile alpha motif domain-containing protein 12 Mus musculus 2 | 34.63 | 6.21 | 18245 | 1 | 1 | 1921470.25 | 2018619.25 | 463416.31 | 1535382.733 | 1738025.12 | 1171723.25 | 1497352.75 | 1765120.12 | 959212.69 | 1177238.5 | 1523067.38 | 1225077.88 | 910830.62 | 1414527.62 |
| Q60710 | Samhd1 | Deoxynucleoside triphosphate triphosphohydrolase SAMHD1 Mus musculus | 1087.99 | 36.93 | 75893 | 18 | 19 | 79472193 | 78731289 | 68233169 | 67630641 | 75887353 | 73405737 | 58374857 | 78814761 | 83157713 | 73752569 | 70849593 | 67221257 | 77045033 | 71906161 |
| Q8BGH2 | Samm50 | Sorting and assembly machinery component 50 homolog Mus musculus | 769.28 | 41.58 | 51864 | 15 | 15 | 21734131 | 19847863 | 20459641 | 19857669 | 19958101 | 22509353 | 21406595 | 21761089 | 22735655 | 21234795 | 18830151 | 22308693 | 22266007 | 22129885 |
| O55128 | Sap18 | Histone deacetylase complex subunit SAP18 Mus musculus | 126.72 | 15.03 | 17595 | 2 | 2 | 1 | 1 | 331385.66 | 533466.81 | 1 | 1 | 380910.66 | 500984.25 | 1 | 395778.47 | 529056.88 | 1 | 1 | 1 |
| P36536 | Sar1a | GTP-binding protein SAR1a Mus musculus | 240.6 | 27.27 | 22371 | 2 | 3 | 2159137.5 | 1953319.5 | 1877033.5 | 1812069.75 | 1819013.25 | 846541.12 | 999676.94 | 2010583.88 | 1665897.5 | 1529691.75 | 1644286.25 | 1301579.5 | 1569697.25 | 1992059.25 |
| Q5HZY2 | Sar1b | GTP-binding protein SAR1b Rattus norvegicus | 295.25 | 29.8 | 22410 | 3 | 4 | 3482802 | 3752000.25 | 5363574 | 3161496.75 | 2763001.25 | 4530959 | 4920448.5 | 5744016 | 4791704 | 3931088 | 4426938 | 4307661 | 5302339 | 4720940.5 |
| Q498U4 | Sarnp | SAP domain-containing ribonucleoprotein Rattus norvegicus | 306.32 | 27.62 | 23605 | 5 | 5 | 3018168.75 | 4519868 | 2776125.25 | 5417837.5 | 2201666 | 3298892.5 | 3852880.5 | 2307280.75 | 4058015 | 6864431 | 4194893.5 | 5666257.5 | 3860670 | 4073156.5 |
| P26638 | Sars | Serine--tRNA ligase, cytoplasmic Mus musculus | 1343.12 | 51.56 | 58389 | 22 | 22 | 93568409 | 104996937 | 91261817 | 111538577 | 105797465 | 113404985 | 109825777 | 100149649 | 104939665 | 116163945 | 94934425 | 88368241 | 87634489 | 92781089 |
| Q9JJL8 | Sars2 | Serine--tRNA ligase, mitochondrial Mus musculus | 63.79 | 2.32 | 58316 | 1 | 1 | 1031626.44 | 1129053.38 | 2041283.12 | 1310794.25 | 1255277 | 1062469.62 | 1290307.12 | 1482976.88 | 1066138.25 | 1071088.5 | 1443641.62 | 1291258 | 1192575.5 | 1079425.38 |
| Q5XIW8 | Sart1 | U4/U6.U5 tri-snRNP-associated protein 1 Rattus norvegicus | 126.17 | 4.09 | 91011 | 3 | 3 | 1022317.44 | 1042419.88 | 1167091.5 | 1258104.25 | 1175796.25 | 1289140.88 | 1254510.25 | 1132844.88 | 1312432.75 | 1709854.75 | 1172140.75 | 1458020.12 | 1205748.75 | 1286568.62 |
| Q9JLI8 | Sart3 | Squamous cell carcinoma antigen recognized by T-cells 3 Mus musculus | 375.27 | 8.73 | 109619 | 7 | 7 | 5469114.5 | 5766772.5 | 5798954 | 5408690 | 4815226.5 | 5109508.5 | 4400458 | 6172054.5 | 3581267.5 | 3586910.75 | 4621964 | 4952972.5 | 4004430.5 | 5646617.5 |
| P59808 | Sash1 | SAM and SH3 domain-containing protein 1 Mus musculus | 67.6 | 0.89 | 135591 | 1 | 1 | 1415810 | 1346030.88 | 1071407 | 1438931.88 | 1253963 | 1414517.5 | 1340807.75 | 1334337.25 | 951323 | 1499281.38 | 1180117 | 1279898.12 | 1112482 | 1373255.88 |
| P70122 | Sbds | Ribosome maturation protein SBDS Mus musculus | 52.49 | 4 | 28781 | 1 | 1 | 468094.38 | 261346.23 | 200839.12 | 131399.03 | 657034.5 | 376214.25 | 599316.5 | 1 | 281986.47 | 1 | 496022.06 | 243875.12 | 744044.69 | 144292.72 |
| Q6ZPE2 | Sbf1 | Myotubularin-related protein 5 Mus musculus | 110.26 | 1.55 | 208693 | 2 | 2 | 560052.06 | 577469.31 | 622520.06 | 691935.44 | 894105.12 | 1233396 | 502474.12 | 677905.06 | 762652.75 | 490915.59 | 661914.81 | 582652.81 | 537049.62 | 621268.19 |
| Q63623 | Scaf8 | Protein SCAF8 Rattus norvegicus | 74.17 | 0.95 | 139504 | 1 | 1 | 3052153.75 | 2777303.75 | 2723363 | 2300131 | 2893278.5 | 2533245.75 | 2562560.5 | 3103682.75 | 2701789.25 | 3013295.75 | 2661315.75 | 2115664.25 | 2262021 | 2878393 |
| O35114 | Scarb2 | Lysosome membrane protein 2 Mus musculus | 295.67 | 14.85 | 54044 | 5 | 5 | 3792335.75 | 3064972 | 4783295 | 2932860 | 2375041.5 | 7356850 | 5877957 | 8681323 | 8287700.5 | 5328843 | 5932664 | 10648394 | 8507017 | 7792342 |
| Q8R127 | Sccpdh | Saccharopine dehydrogenase-like oxidoreductase Mus musculus | 159.27 | 10.49 | 47129 | 3 | 3 | 1243900.75 | 2271185.75 | 1727978.5 | 1769493 | 1637801.5 | 1843172.25 | 2015387.75 | 2006724.5 | 1928012.88 | 1665322.12 | 1897724 | 1385150 | 1455709.75 | 1520323.12 |
| Q8BRF7 | Scfd1 | Sec1 family domain-containing protein 1 Mus musculus | 512.72 | 21.91 | 72323 | 9 | 9 | 7423727 | 9705450 | 8513180 | 5661733.5 | 9249032 | 5743905 | 7374019 | 7533460.5 | 6623525 | 5289984 | 6613288 | 7804352 | 6711681.5 | 10602036 |
| Q9JLI6 | Scly | Selenocysteine lyase Mus musculus | 238.14 | 13.43 | 47174 | 5 | 5 | 6722862.5 | 6215261 | 6757017 | 5159280 | 7493572 | 4549049.5 | 6940098 | 5364677.5 | 6086449 | 7024510 | 4431521 | 6142608 | 6677410.5 | 4518528 |
| P32020 | Scp2 | Non-specific lipid-transfer protein Mus musculus | 321.76 | 16.09 | 59126 | 6 | 6 | 12118544 | 14167560 | 10426317 | 9385977 | 12512601 | 9774290 | 11554284 | 11971852 | 11519022 | 11980838 | 11511540 | 13743543 | 11574557 | 12275619 |
| Q920A5 | Scpep1 | Retinoid-inducible serine carboxypeptidase Mus musculus | 619.39 | 28.76 | 50965 | 11 | 13 | 64574505 | 79703097 | 73722761 | 67582617 | 66291773 | 78982225 | 76359473 | 84565441 | 73472249 | 73442817 | 71839409 | 71218345 | 78330489 | 74054497 |
| Q80U72 | Scrib | Protein scribble homolog Mus musculus | 108.41 | 3.47 | 174059 | 3 | 3 | 243420.61 | 164917.41 | 227873.67 | 222660.3 | 177761.56 | 411136.75 | 199203.84 | 253438.47 | 306694.25 | 329076.5 | 295576.72 | 1 | 1 | 193082.22 |
| Q8CFE4 | Scyl2 | SCY1-like protein 2 Mus musculus | 80.93 | 2.69 | 103317 | 2 | 2 | 185693.36 | 540642.88 | 685600.88 | 205727.7 | 206164.47 | 909027.12 | 213282.95 | 1002493.25 | 966004.56 | 529786.12 | 580498.5 | 909815 | 845309 | 544065.81 |
| Q9DBQ7 | Scyl3 | Protein-associating with the carboxyl-terminal domain of ezrin Mus musculus | 100.97 | 6.53 | 81333 | 3 | 3 | 1147515.5 | 1362701 | 1369663.38 | 1772628 | 651327.56 | 1073660.88 | 1268754.12 | 882448.12 | 964852 | 830007.62 | 893089.19 | 905812.12 | 1012147.94 | 867306.5 |
| O08992 | Sdcbp | Syntenin-1 Mus musculus | 423.87 | 53.18 | 32379 | 7 | 7 | 1300222.12 | 1207490.5 | 2344484 | 801476.5 | 1769486.5 | 1875984.25 | 1116153.75 | 931284.38 | 1782370.62 | 2053990.88 | 1786360.38 | 2215532.75 | 2364378.75 | 927036.94 |
| Q9ESP1 | Sdf2l1 | Stromal cell-derived factor 2-like protein 1 Mus musculus | 69.61 | 13.57 | 23648 | 2 | 2 | 685499.3125 | 737664.38 | 493002.62 | 688225.5 | 823104.75 | 945478.25 | 1 | 1 | 630537.25 | 1263878.12 | 615038.44 | 1 | 626083.25 | 1 |
| Q8K2B3 | Sdha | Succinate dehydrogenase [ubiquinone] flavoprotein subunit, mitochondrial Mus musculus | 994.47 | 38.86 | 72585 | 16 | 16 | 52981733 | 55452269 | 54265109 | 67279817 | 65987317 | 64443457 | 74263321 | 66438953 | 75203801 | 71476529 | 67968353 | 60704209 | 63219957 | 60731725 |
| Q9CQA3 | Sdhb | Succinate dehydrogenase [ubiquinone] iron-sulfur subunit, mitochondrial Mus musculus | 438.18 | 30.5 | 31814 | 8 | 8 | 44796633 | 49837093 | 50325585 | 50812549 | 49363261 | 57099637 | 59152501 | 52235157 | 64218137 | 59485769 | 53334785 | 55999333 | 53467569 | 55464701 |
| Q9CZB0 | Sdhc | Succinate dehydrogenase cytochrome b560 subunit, mitochondrial Mus musculus | 167.48 | 21.89 | 18382 | 3 | 3 | 12820827 | 11249965 | 9982293 | 8338896 | 10829868 | 12274096 | 11704585 | 13899473 | 12769384 | 10533235 | 11310572 | 10036823 | 11916172 | 10979183 |
| Q9CXV1 | Sdhd | Succinate dehydrogenase [ubiquinone] cytochrome b small subunit, mitochondrial Mus mu | 82.47 | 16.35 | 17014 | 1 | 1 | 2303064.75 | 1947185.12 | 1287627.62 | 1616036.25 | 1565868.62 | 952996.25 | 1037701.69 | 1567015.25 | 819913.06 | 910888.75 | 1212176.75 | 1040106.31 | 881603.31 | 1251570.62 |
| P42667 | Sec11a | Signal peptidase complex catalytic subunit SEC11A Rattus norvegicus 2 | 126.39 | 10.61 | 20599 | 2 | 2 | 1012932.69 | 1468945.88 | 1810254.12 | 783664.69 | 1064958.88 | 959679.56 | 1133396.62 | 1164239.25 | 684274.75 | 921509.25 | 860617.75 | 1037920.81 | 1282061.75 | 1430189 |
| Q5XFW8 | Sec13 | Protein SEC13 homolog Rattus norvegicus | 301.91 | 23.91 | 35548 | 5 | 5 | 6235783 | 7814998.5 | 8158002 | 5373641 | 7645573 | 8625475 | 5460368 | 3646332.25 | 7341191.5 | 8141656 | 7942263.5 | 8139761 | 8210898 | 7768530 |
| O08547 | Sec22b | Vesicle-trafficking protein SEC22b Mus musculus | 415.75 | 43.72 | 24741 | 8 | 8 | 5180140 | 3469864 | 3923791 | 4471955 | 4959938 | 3485140 | 4015192 | 2821130 | 3856196.5 | 4042485 | 4295024 | 4088552 | 3940209.5 | 1804362.38 |
| Q01405 | Sec23a | Protein transport protein Sec23A Mus musculus | 237.9 | 6.93 | 86162 | 2 | 4 | 5033725 | 2407798.75 | 4261623 | 2926656 | 4742403 | 4272312.5 | 5432406 | 1452974.38 | 4321981.5 | 3594826 | 4081459 | 4125259.5 | 2769362.5 | 3229080 |
| Q9D662 | Sec23b | Protein transport protein Sec23B Mus musculus | 799.74 | 32.46 | 86437 | 12 | 14 | 7470714 | 8951778 | 9549329 | 8320190.5 | 8321831 | 8290246 | 10619059 | 7553449 | 10461038 | 9662469 | 8198959.5 | 9384214 | 8867108 | 10330582 |
| Q6NZC7 | Sec23ip | SEC23-interacting protein Mus musculus | 128.94 | 4.11 | 110780 | 3 | 3 | 1 | 1 | 1 | 636808.69 | 668890.06 | 610549.81 | 1 | 1 | 486168.31 | 596515.75 | 681312.19 | 609869.56 | 972238 | 518661.81 |
| Q3U2P1 | Sec24a | Protein transport protein Sec24A Mus musculus | 45.44 | 1.38 | 118782 | 1 | 1 | 1 | 1 | 1 | 1 | 1 | 1 | 236386.28 | 1 | 277140.5 | 1 | 313900.7 | 307739.38 | 359366.22 | 274596.5 |
| Q3UPL0 | Sec31a | Protein transport protein Sec31A Mus musculus | 938.73 | 21.38 | 133569 | 18 | 18 | 14539925 | 15577787 | 15663304 | 13616549 | 14152855 | 16433328 | 13719417 | 17785469 | 19971687 | 17524039 | 13949001 | 18363271 | 16243615 | 14858655 |
| P61620 | Sec61a1 | Protein transport protein Sec61 subunit alpha isoform 1 Mus musculus | 425.25 | 24.58 | 52265 | 6 | 6 | 15066201 | 10202939 | 5386791 | 9174268 | 5527858 | 10454344 | 5727434.5 | 8117550.5 | 9752174 | 7303581 | 6730256 | 10354123 | 9715091 | 10416825 |
| Q9CQS8 | Sec61b | Protein transport protein Sec61 subunit beta Mus musculus | 122.98 | 26.04 | 9958 | 2 | 2 | 12811087 | 15127483 | 16301788 | 15294817 | 15382770 | 14942069 | 15933591 | 14303207 | 16023137 | 17472033 | 16996077 | 16117519 | 18071473 | 16337454 |
| P60060 | Sec61g | Protein transport protein Sec61 subunit gamma Mus musculus 3 | 49.17 | 17.65 | 7741 | 1 | 1 | 1013166.935 | 940403.62 | 1254208.25 | 832944.81 | 1025111.06 | 1048773.438 | 1119073.62 | 937052.75 | 1065544.38 | 1073423 | 1211994.25 | 1178867.88 | 1354800.62 | 1102314.25 |
| Q8BU14 | Sec62 | Translocation protein SEC62 Mus musculus | 41.11 | 2.51 | 45581 | 1 | 1 | 2574255 | 2461246.25 | 2742003.5 | 2221585.5 | 3078395.25 | 2534673.75 | 2840654.5 | 2257911.75 | 2522394.5 | 2651344.25 | 2463168 | 2804771.25 | 2709800.75 | 3344884 |
| Q6A098 | Secisbp2l | Selenocysteine insertion sequence-binding protein 2-like Mus musculus | 49.94 | 1.93 | 119641 | 1 | 1 | 123743.5275 | 161187.5 | 95700.15 | 109622.68 | 128463.78 | 104966.07 | 1 | 109821.23 | 113476.09 | 1 | 1 | 1 | 140026.73 | 85533.91 |
| Q8R2U0 | Seh1l | Nucleoporin SEH1 Mus musculus | 77.55 | 5.28 | 39775 | 2 | 2 | 795350.62 | 860282.765 | 863834.31 | 920956.69 | 860989.44 | 861379 | 1052641.25 | 1 | 947420.12 | 1 | 775490.5 | 946218.69 | 748454.44 | 804098.69 |
| Q9Z2G6 | Sel1l | Protein sel-1 homolog 1 Mus musculus | 180.16 | 7.22 | 88340 | 3 | 3 | 659071.56 | 842661.31 | 786070.31 | 617422.25 | 558371.38 | 538527.88 | 674490.62 | 839048 | 586141 | 596985.31 | 553807.19 | 610468.56 | 706800.94 | 670534.56 |
| Q923V8 | Selenof | Selenoprotein F Rattus norvegicus | 196.66 | 41.36 | 17822 | 4 | 4 | 706844.62 | 1173329.88 | 792476.62 | 955051 | 1068103.25 | 1084835.75 | 1311926.12 | 1278967.75 | 1119840.75 | 1150489.25 | 1066741 | 1125392.75 | 1164672.12 | 1364589.88 |
| O09126 | Sema4d | Semaphorin-4D Mus musculus | 229.88 | 7.08 | 95640 | 5 | 5 | 4311965 | 3744597.75 | 2528419 | 3849758.5 | 2527764.5 | 2911149 | 3152263 | 2951382 | 3383711 | 2777725 | 3044298.75 | 3290777.5 | 3169127 | 2885720 |
| Q8BH69 | Sephs1 | Selenide, water dikinase 1 Mus musculus | 124.17 | 9.95 | 42907 | 4 | 4 | 4561529.5 | 3079394.75 | 4759814.5 | 3105929.25 | 3595903.75 | 3415412.25 | 3347135.25 | 5642203.5 | 4965417 | 3478730.25 | 3994759.75 | 4080021.5 | 4353256.5 | 3781767 |
| B3GNI6 | Sept11 | Septin-11 Rattus norvegicus | 426.09 | 30.16 | 49695 | 6 | 10 | 2556646.75 | 2712279.5 | 3552162.25 | 4299992.5 | 4791268.5 | 3965569.5 | 4098744.5 | 3800961.75 | 3879300.25 | 4345838 | 3654072.25 | 4277610 | 4478832 | 4229017.5 |
| P42208 | Sept2 | Septin-2 Mus musculus | 642.11 | 43.21 | 41526 | 9 | 10 | 28462507 | 28598655 | 27987833 | 26868779 | 28410479 | 31002607 | 30110541 | 30369759 | 30355221 | 30671835 | 28705013 | 28579201 | 27823747 | 28760313 |
| Q9R1T4 | Sept6 | Septin-6 Mus musculus | 173.51 | 8.53 | 49620 | 1 | 4 | 1726269.25 | 1790439.25 | 1607049.5 | 1807783.62 | 2788831.75 | 1426661 | 2126552 | 1903069.88 | 1955579.25 | 2114407 | 1872205 | 2208637.5 | 2080682 | 2058740.12 |
| O55131 | Sept7 | Septin-7 Mus musculus | 811.74 | 41.06 | 50550 | 15 | 16 | 12041432 | 10079734 | 12569328 | 13203832 | 12218833 | 13630385 | 12267489 | 11119578 | 13119359 | 13199626 | 12730930 | 13378511 | 13629417 | 12566661 |
| Q8CHH9 | Sept8 | Septin-8 Mus musculus | 389.48 | 14.45 | 49812 | 4 | 6 | 1973114.75 | 2295610.5 | 2710563.5 | 2638609.5 | 2549811.25 | 2690864.75 | 2945754.5 | 2552342.25 | 1046030.38 | 2345349.5 | 3161122 | 2899460.5 | 2353662.25 | 2553819.25 |
| Q80UG5 | Sept9 | Septin-9 Mus musculus | 247.66 | 10.81 | 65575 | 5 | 5 | 13375406 | 13316225 | 14692731 | 15236556 | 12774570 | 14959122 | 13904698 | 13710806 | 14760871 | 14848932 | 13032281 | 12609731 | 14279687 | 14060000 |
| Q6AXS5 | Serbp1 | Plasminogen activator inhibitor 1 RNA-binding protein Rattus norvegicus | 661.45 | 36.36 | 44754 | 11 | 11 | 24421401 | 20042011 | 22517981 | 26039509 | 20794507 | 19698909 | 22605957 | 18395947 | 17107879 | 26830441 | 24652123 | 24514465 | 22298877 | 18609773 |
| Q9QZI9 | Serinc3 | Serine incorporator 3 Mus musculus | 59.3 | 2.54 | 52623 | 1 | 1 | 1522499.25 | 1 | 1 | 1 | 1 | 1 | 1 | 1 | 445575.69 | 1 | 547085.81 | 537091.5 | 460704.31 | 514960.54 |
| Q9DCQ7 | Serpina1f | Alpha-1-antitrypsin 1-6 Mus musculus | 33.93 | 1.95 | 46337 | 1 | 1 | 5145926 | 4528943.5 | 5142068 | 3530996.75 | 3343327.5 | 1960305.75 | 2068821.88 | 2967468.5 | 1613439.5 | 1921989.38 | 1524142.25 | 2172384.25 | 3404321.25 | 1969269 |
| Q4G075 | Serpinb1a | Leukocyte elastase inhibitor A Rattus norvegicus | 69.24 | 2.64 | 42727 | 1 | 1 | 1023972.69 | 1064173.5 | 1096896.38 | 1305444.12 | 1008452.94 | 1868474.5 | 1546595.12 | 1100378.75 | 1621885.25 | 1312841.5 | 1235579.75 | 1489651.5 | 1297659.62 | 1510886.75 |
| Q60854 | Serpinb6 | Serpin B6 Mus musculus | 1224.84 | 60.85 | 42599 | 18 | 19 | 83696937 | 106701513 | 88571385 | 97106305 | 93216201 | 89508745 | 88093793 | 96750753 | 92560633 | 90640505 | 93737497 | 86320969 | 80151001 | 83897089 |
| O08800 | Serpinb8 | Serpin B8 Mus musculus | 234.19 | 15.51 | 42150 | 3 | 3 | 2481567 | 2146444 | 1905547 | 2350871.75 | 1693934.5 | 1710205 | 2132007 | 2110688.25 | 1490581.5 | 1416441.25 | 1988005.38 | 2192795 | 1314699.5 | 2422663 |
| P32261 | Serpinc1 | Antithrombin-III Mus musculus | 315.63 | 15.05 | 52004 | 6 | 6 | 29093393 | 14399019 | 18221479 | 21083791 | 18452405 | 15196599 | 14321270 | 14468564 | 13993722 | 13181828 | 12153597 | 10939648 | 18360459 | 8842693 |
| Q63945 | Set | Protein SET Rattus norvegicus | 612.54 | 43.94 | 33406 | 9 | 10 | 34842169 | 30783585 | 37072669 | 26052119 | 28159197 | 35129169 | 32027769 | 37898477 | 41493393 | 34551309 | 28725061 | 35045613 | 39779889 | 29131065 |
| Q91WC0 | Setd3 | Histone-lysine N-methyltransferase setd3 Mus musculus | 208.66 | 11.28 | 67176 | 4 | 4 | 1710877 | 1393691.75 | 1837505.38 | 1575329.5 | 1700180.5 | 2206011.75 | 1775680.75 | 1751678.75 | 1528940.75 | 1585965.5 | 1917854.5 | 1576749 | 1721286.62 | 1433492.62 |
| Q8VHL1 | Setd7 | Histone-lysine N-methyltransferase SETD7 Mus musculus | 88.82 | 4.92 | 40506 | 1 | 1 | 1 | 1 | 1 | 1 | 1980008.75 | 1 | 813285.56 | 1 | 1445148.88 | 1521750.88 | 1 | 1 | 1 | 1984639.38 |
| Q64213 | Sf1 | Splicing factor 1 Mus musculus | 595.69 | 24.81 | 70403 | 11 | 11 | 19301177 | 19982713 | 18181747 | 23425367 | 21831543 | 19475585 | 20044853 | 19238959 | 19758771 | 20278755 | 21451799 | 16448691 | 18054243 | 20098737 |
| Q8K4Z5 | Sf3a1 | Splicing factor 3A subunit 1 Mus musculus | 1022.27 | 32.11 | 88545 | 19 | 19 | 30420623 | 25910533 | 30410029 | 30604865 | 32499881 | 31353905 | 27806345 | 25583575 | 29513207 | 32623877 | 25433939 | 29100329 | 28047137 | 26057303 |
| Q62203 | Sf3a2 | Splicing factor 3A subunit 2 Mus musculus | 165.61 | 16.21 | 49911 | 4 | 4 | 1577524 | 1475952 | 1110295.5 | 1313441.25 | 1468282.25 | 1529002 | 2695629.25 | 445041.47 | 2188663.5 | 2299224 | 1613712.5 | 577277.75 | 1757531.38 | 1406476.75 |
| Q9D554 | Sf3a3 | Splicing factor 3A subunit 3 Mus musculus | 868.71 | 38.72 | 58842 | 14 | 14 | 18947025 | 19965373 | 17937693 | 16690158 | 19102867 | 21308965 | 19024135 | 19742131 | 21233429 | 21158951 | 18660723 | 18272653 | 20479675 | 19425941 |
| Q99NB9 | Sf3b1 | Splicing factor 3B subunit 1 Mus musculus | 985.69 | 23.54 | 145816 | 21 | 21 | 34263065 | 14947607 | 13554555 | 16902905 | 15601250 | 14862865 | 14956012 | 17507259 | 13938548 | 15110369 | 15974332 | 9438698 | 15609598 | 13114733 |
| Q921M3 | Sf3b3 | Splicing factor 3B subunit 3 Mus musculus | 2181.02 | 39.19 | 135550 | 34 | 35 | 37008477 | 37335373 | 42581665 | 40881613 | 40728669 | 44396025 | 40689029 | 41002281 | 42931017 | 40451505 | 40201461 | 40888417 | 40673689 | 39315161 |
| Q6AYL5 | Sf3b4 | Splicing factor 3B subunit 4 Rattus norvegicus | 415.15 | 20.99 | 44356 | 6 | 6 | 8127862 | 8459824 | 6111461.5 | 7344115 | 7829358.5 | 8087920 | 6814517.5 | 8770515 | 4622509.5 | 7437833 | 8646837 | 8578870 | 6795935.5 | 4573995 |
| P59708 | Sf3b6 | Splicing factor 3B subunit 6 Mus musculus | 106.7 | 20 | 14585 | 2 | 2 | 2154996.75 | 2299903.25 | 2438491.25 | 2262488.5 | 1923720 | 1966843.62 | 2105065.25 | 2529418.75 | 2675716.5 | 2061465.5 | 2248657 | 1936946.5 | 1971892.88 | 2607119 |
| O70456 | Sfn | 14-3-3 protein sigma Mus musculus | 284.71 | 16.13 | 27706 | 1 | 6 | 144261889 | 165393505 | 152082865 | 155488065 | 144588449 | 151719169 | 148202369 | 174104657 | 183128961 | 164531121 | 145035185 | 157824993 | 141626865 | 155234433 |
| Q8VIJ6 | Sfpq | Splicing factor, proline- and glutamine-rich Mus musculus | 993.57 | 27.18 | 75442 | 16 | 16 | 14007516 | 11468027 | 12631364 | 16690725 | 12777351 | 11896819 | 11022301 | 11704439 | 12804453 | 14737486 | 13314540 | 12205391 | 14441313 | 12588686 |
| Q99JR1 | Sfxn1 | Sideroflexin-1 Mus musculus | 434.94 | 31.99 | 35649 | 5 | 6 | 8041927 | 7853453 | 7753863 | 8344489 | 7582172.5 | 7479436 | 7853814 | 6919617 | 6900343 | 8052574 | 9328649 | 8110641 | 9092919 | 9357381 |
| Q91V61 | Sfxn3 | Sideroflexin-3 Mus musculus | 202.32 | 18.38 | 35406 | 2 | 3 | 1603607.75 | 1896116 | 1578090.88 | 1786388.12 | 1732143 | 1197221.88 | 1553972.62 | 1900640.5 | 1655666.5 | 1684663.38 | 1856514.62 | 1667865.75 | 1501763.12 | 1495111.88 |
| Q8R4V0 | Sgk3 | Serine/threonine-protein kinase Sgk3 Rattus norvegicus | 43 | 2.42 | 57171 | 1 | 1 | 1 | 512506.91 | 1 | 1 | 1 | 1 | 1 | 1 | 1 | 441098.69 | 1 | 1 | 369607.03 | 1 |
| Q8R0X7 | Sgpl1 | Sphingosine-1-phosphate lyase 1 Mus musculus | 342.72 | 16.55 | 63677 | 7 | 7 | 12433941 | 14669872 | 14132386 | 12871850 | 14855282 | 14384436 | 13004845 | 13495315 | 14145264 | 14439873 | 14482327 | 13638032 | 16274623 | 15948561 |
| O70593 | Sgta | Small glutamine-rich tetratricopeptide repeat-containing protein alpha Rattus norvegicus | 63.96 | 4.14 | 34157 | 1 | 1 | 1241965.5 | 1529314.5 | 1433334.38 | 1016860.31 | 1864432.75 | 883875.38 | 1017203.12 | 1010371.31 | 1280786.25 | 898506.12 | 933536 | 1096998.12 | 965641.62 | 1307647.88 |
| Q9JJU8 | Sh3bgrl | SH3 domain-binding glutamic acid-rich-like protein Mus musculus | 383.94 | 56.14 | 12811 | 7 | 7 | 55023249 | 54861409 | 54446401 | 59335037 | 60166041 | 53728281 | 61036005 | 51776285 | 53912453 | 55398185 | 61283109 | 48660881 | 50334109 | 57088141 |
| Q8BG73 | Sh3bgrl2 | SH3 domain-binding glutamic acid-rich-like protein 2 Mus musculus | 138.63 | 38.32 | 12255 | 3 | 3 | 5735677 | 5209213.5 | 6878262 | 4435783 | 4723495 | 7416827 | 5530544.5 | 7546743 | 6786781 | 5961108 | 4630350.5 | 5747597 | 6390881 | 6182943 |
| Q91VW3 | Sh3bgrl3 | SH3 domain-binding glutamic acid-rich-like protein 3 Mus musculus | 237.33 | 66.67 | 10477 | 4 | 4 | 8359976 | 8215580.5 | 10558641 | 12215968 | 9039162 | 9502601 | 10098030 | 9781682 | 10756673 | 11461535 | 9971382 | 11268500 | 9600642 | 9592281 |
| P55194 | Sh3bp1 | SH3 domain-binding protein 1 Mus musculus | 366.42 | 14.71 | 74172 | 6 | 6 | 4435971.5 | 4594058 | 3631084 | 4682397 | 3457839 | 4283551 | 4419885 | 4966386.5 | 4541030 | 4418873 | 4340379 | 4611871.5 | 5144286.5 | 3408942 |
| O35964 | Sh3gl1 | Endophilin-A2 Rattus norvegicus | 262.24 | 17.93 | 41492 | 6 | 6 | 2962311.5 | 1598904.12 | 3551249 | 2889640 | 1 | 2756065.75 | 1169509 | 2832217 | 3639780 | 3313469.75 | 3595007.5 | 3923269 | 2451888.25 | 2852882.5 |
| Q6AYE2 | Sh3glb1 | Endophilin-B1 Rattus norvegicus | 215.15 | 10.68 | 40787 | 3 | 3 | 6654449 | 7580770 | 5955345.5 | 7714775 | 7210518 | 5255211 | 5804351 | 5699925.5 | 5476036 | 6283219.5 | 6525239.5 | 5816611 | 5211846 | 6372754 |
| A2AAY5 | Sh3pxd2b | SH3 and PX domain-containing protein 2B Mus musculus | 87.58 | 1.87 | 101517 | 2 | 2 | 861233.5 | 1130739.38 | 1630043.75 | 1116311 | 1239085.88 | 1131659.38 | 963769.12 | 850838.31 | 1007704.56 | 913461.38 | 1100009.12 | 1179238 | 990180.5 | 971949.75 |
| P50431 | Shmt1 | Serine hydroxymethyltransferase, cytosolic Mus musculus | 184.35 | 8.16 | 52601 | 2 | 3 | 10176252 | 9464960 | 11348258 | 11411757 | 8762703 | 10619852 | 10582433 | 9730348 | 10882089 | 11266423 | 9495418 | 11887216 | 12014774 | 10704177 |
| Q9CZN7 | Shmt2 | Serine hydroxymethyltransferase, mitochondrial Mus musculus | 1282.55 | 57.54 | 55759 | 23 | 24 | 110442577 | 112967289 | 116171249 | 138613057 | 121700257 | 102444689 | 116646337 | 113575185 | 104615297 | 109584985 | 125714713 | 109786921 | 112909489 | 126930857 |
| Q8K2Q9 | Shtn1 | Shootin-1 Mus musculus | 453.89 | 12.84 | 71343 | 9 | 9 | 18959481 | 21447931 | 19827315 | 19525633 | 19291207 | 19012555 | 17929105 | 19085505 | 18397015 | 18127753 | 18088917 | 18162987 | 16660083 | 20857917 |

| | | | | | | | | | | | | | | | | | | | | | |
|---|---|---|---|---|---|---|---|---|---|---|---|---|---|---|---|---|---|---|---|---|---|
| P70665 | Siae | Sialate O-acetylesterase Mus musculus | 186.47 | 7.76 | 60775 | 3 | 3 | 3026954.5 | 6427991 | 5872064.5 | 4669695 | 5355039 | 4599566 | 4039695 | 5363563 | 4577098 | 3519424.75 | 4673151 | 4892544 | 4528089 | 4525481.5 |
| O55242 | Sigmar1 | Sigma non-opioid intracellular receptor 1 Mus musculus | 126.45 | 13.9 | 25250 | 2 | 2 | 1644043 | 888876.69 | 952398.69 | 788086.75 | 1070364 | 782130 | 942550.5 | 693566.69 | 849490.88 | 442935.09 | 809656.81 | 1233380.75 | 443367.47 | 780649.19 |
| Q6P6S4 | Sil1 | Nucleotide exchange factor SIL1 Rattus norvegicus | 140.87 | 6.45 | 52350 | 3 | 3 | 2155357 | 341537.28 | 1373500.62 | 1334709.12 | 1289616.75 | 1902878.5 | 1773197.25 | 1771166.75 | 1874802.12 | 1451034.88 | 1547131.38 | 2022945.25 | 1864207.88 | 1587361 |
| Q60520 | Sin3a | Paired amphipathic helix protein Sin3a Mus musculus | 186.69 | 4.47 | 145088 | 4 | 4 | 842738.75 | 1095125.75 | 844350.25 | 316112.09 | 396585.84 | 213666.59 | 619759.88 | 388012.03 | 947565.38 | 388826.25 | 737508.25 | 955438.75 | 800720.38 | 495675.81 |
| Q8C8M1 | Sinhcaf | SIN3-HDAC complex-associated factor Mus musculus | 45.19 | 5.88 | 24829 | 1 | 1 | 193652.61 | 151803.62 | 1 | 311316.62 | 145616.91 | 227726.62 | 205427.45 | 221141.61 | 150751.55 | 173928.23 | 142878.86 | 247241.86 | 166763.11 | 207064.22 |
| P46062 | Sipa1 | Signal-induced proliferation-associated protein 1 Mus musculus | 144.33 | 4.53 | 112066 | 2 | 2 | 2053289.38 | 2238001 | 2395500 | 2068664.88 | 2462241.25 | 2002720.5 | 1997478.12 | 1813371.12 | 1866137 | 1663222.38 | 2175346.75 | 2116383 | 540939.88 | 2139342 |
| P97797 | Sirpa | Tyrosine-protein phosphatase non-receptor type substrate 1 Mus musculus | 389.84 | 23 | 56425 | 7 | 7 | 23094845 | 21341977 | 20353763 | 19189805 | 20057461 | 19577595 | 21475925 | 18487875 | 16792299 | 19625617 | 19875303 | 18915523 | 18498729 | 18524879 |
| Q5RJQ4 | Sirt2 | NAD-dependent protein deacetylase sirtuin-2 Rattus norvegicus | 136.95 | 7.14 | 39319 | 2 | 2 | 1776509.38 | 1368802.88 | 1590799.25 | 1262741.88 | 1485338.25 | 1239999.38 | 1558391.283 | 1702029.75 | 1917732.25 | 1373803.75 | 1469158.12 | 1 | 1 | 1 |
| Q8R216 | Sirt4 | NAD-dependent protein lipoamidase sirtuin-4, mitochondrial Mus musculus | 51.98 | 3.6 | 37525 | 1 | 1 | 472245.59 | 448417.03 | 421806.75 | 477248.72 | 454929.5225 | 510196.34 | 413236.25 | 539167.56 | 430466.53 | 440966.81 | 420702.19 | 477937.91 | 477534.59 | 581817.06 |
| Q3UND0 | Skap2 | Src kinase-associated phosphoprotein 2 Mus musculus | 634.12 | 60.34 | 40712 | 11 | 11 | 24933905 | 26948779 | 22605333 | 23218359 | 24837851 | 24360677 | 25044069 | 27099931 | 25486501 | 22919585 | 22411163 | 23212519 | 20200163 | 20815125 |
| Q6PEC4 | Skp1 | S-phase kinase-associated protein 1 Rattus norvegicus | 412.72 | 55.21 | 18672 | 6 | 6 | 12821218 | 15930318 | 12726561 | 13665122 | 12955385 | 13671783 | 11563603 | 19048839 | 14621828 | 11641945 | 12194578 | 11721395 | 13968707 | 14818080 |
| Q8CI08 | Slain2 | SLAIN motif-containing protein 2 Mus musculus | 48.31 | 2.07 | 62378 | 1 | 1 | 1065192.75 | 797507.75 | 527056.31 | 1317345.38 | 1272489.5 | 909175.75 | 1 | 1 | 1 | 1 | 1251404.38 | 1955571.25 | 744044.69 | 843040.62 |
| Q99MR3 | Slc12a9 | Solute carrier family 12 member 9 Mus musculus | 135.45 | 3.61 | 96330 | 3 | 3 | 1759231 | 1167908.75 | 481658.88 | 1132727.5 | 752233.5 | 692173.38 | 553375.31 | 953002.38 | 1167049.75 | 494181.62 | 1166323.75 | 1092996.5 | 1069154 | 932546.5 |
| Q8BPX9 | Slc15a3 | Solute carrier family 15 member 3 Mus musculus | 285.85 | 18.86 | 64051 | 6 | 6 | 9729724 | 9355371 | 5480519.5 | 7941604 | 8179974 | 7936137.5 | 8510173 | 5331278 | 5293870.5 | 9096983 | 7207181 | 5249839.5 | 6822525.5 | 5238233 |
| P53986 | Slc16a1 | Monocarboxylate transporter 1 Mus musculus | 323.79 | 14.81 | 53267 | 5 | 5 | 3190276 | 1698462.5 | 1684076 | 1281738.88 | 1902557.12 | 1876960.88 | 1938398 | 1955682.75 | 2096994.75 | 2258160.75 | 2228990.25 | 1967605.75 | 2571043 | 2057473.25 |
| P51912 | Slc1a5 | Neutral amino acid transporter B(0) Mus musculus | 742.2 | 29.29 | 58483 | 11 | 11 | 18431581 | 5509180.5 | 10296889 | 9211809 | 8687740 | 10722033 | 5020683 | 4237944.5 | 7745656 | 5079170.5 | 7757547 | 6274547.5 | 10893871 | 8321042 |
| P32089 | Slc25a1 | Tricarboxylate transport protein, mitochondrial Rattus norvegicus | 236.06 | 20.58 | 33835 | 6 | 6 | 14630337 | 11117192 | 12152416 | 11667673 | 15221807 | 12993797 | 15271721 | 12496816 | 14519285 | 16214638 | 15315478 | 12950639 | 15661461 | 15299264 |
| Q9QZD8 | Slc25a10 | Mitochondrial dicarboxylate carrier Mus musculus | 376.96 | 31.71 | 31715 | 6 | 6 | 7433726.5 | 6977366.5 | 5758274 | 7106157.5 | 6173945 | 6869705 | 8175206 | 7216580 | 7527457 | 7012616 | 8694246 | 8926883 | 8122220.5 | 5798118 |
| Q9CR62 | Slc25a11 | Mitochondrial 2-oxoglutarate/malate carrier protein Mus musculus | 803.98 | 63.06 | 34155 | 13 | 13 | 29607567 | 22388391 | 21583083 | 28706909 | 28475977 | 26764263 | 29935853 | 30251589 | 25458123 | 26428371 | 24534221 | 28227159 | 26400619 | 23852055 |
| Q8BH59 | Slc25a12 | Calcium-binding mitochondrial carrier protein Aralar1 Mus musculus | 1161.56 | 43.72 | 74570 | 17 | 20 | 36904181 | 39552901 | 41379253 | 35412145 | 39784721 | 43641701 | 40349677 | 48736293 | 41627165 | 40790373 | 45488373 | 42274969 | 54611137 | 42473157 |
| Q9QXX4 | Slc25a13 | Calcium-binding mitochondrial carrier protein Aralar2 Mus musculus | 1659.72 | 63.61 | 74467 | 26 | 29 | 62194497 | 61106117 | 65557537 | 57746685 | 64471141 | 67684505 | 68092353 | 81449977 | 75328153 | 72001529 | 73703249 | 66758481 | 87497329 | 74168241 |
| O70579 | Slc25a17 | Peroxisomal membrane protein PMP34 Mus musculus | 72.36 | 4.56 | 34413 | 1 | 1 | 245448.55 | 232337.67 | 208585.3775 | 152327.23 | 204228.06 | 239832.24 | 223624.25 | 257527.38 | 298430.5 | 179746.83 | 278119.81 | 311607.62 | 374917.25 | 280820.88 |
| Q9D6M3 | Slc25a22 | Mitochondrial glutamate carrier 1 Mus musculus | 110.3 | 7.12 | 34670 | 2 | 2 | 1748356.12 | 2944769.25 | 2173978.25 | 3293245.5 | 2781396.75 | 1828596.75 | 2026943.12 | 2223469.5 | 1824502.5 | 2191053.5 | 2499594.75 | 2458332.5 | 2186007.5 | 2208112 |
| Q8BMD8 | Slc25a24 | Calcium-binding mitochondrial carrier protein SCaMC-1 Mus musculus | 335.4 | 19.16 | 52902 | 7 | 7 | 4205191.5 | 3714077.75 | 3746114 | 4745642 | 3702689.75 | 3213535.75 | 3167947 | 3124622.5 | 2970465.25 | 4409744.5 | 4878225.5 | 4083488 | 4378294.5 | 3892001.25 |
| P16036 | Slc25a3 | Phosphate carrier protein, mitochondrial Rattus norvegicus | 713.67 | 44.66 | 39445 | 12 | 12 | 36062841 | 41107849 | 41221105 | 34584221 | 38753701 | 47766305 | 45828705 | 50553545 | 54328689 | 48474313 | 44965977 | 49965213 | 50002517 | 46465757 |
| P48962 | Slc25a4 | ADP/ATP translocase 1 Mus musculus | 1171.58 | 63.76 | 32904 | 8 | 20 | 324441665 | 356342497 | 317014433 | 262045585 | 319668129 | 307007297 | 337986241 | 367036129 | 327870369 | 310991873 | 313181441 | 311626753 | 335774529 | 342906945 |
| Q8CFJ7 | Slc25a45 | Solute carrier family 25 member 45 Mus musculus | 80.69 | 5.9 | 31690 | 1 | 1 | 555261.12 | 936192 | 254531.72 | 661330.19 | 674990.75 | 916973.69 | 544061.69 | 686264 | 736088.19 | 500566.84 | 569572.19 | 685322.31 | 596268.88 | 663532.19 |
| P51881 | Slc25a5 | ADP/ATP translocase 2 Mus musculus | 1478.8 | 76.85 | 32931 | 13 | 25 | 436639297 | 474269697 | 425312257 | 362385185 | 427604641 | 413629025 | 445045473 | 486678497 | 431457313 | 412497921 | 420322017 | 414729537 | 439041761 | 462399521 |
| P97849 | Slc27a1 | Long-chain fatty acid transport protein 1 Rattus norvegicus | 37.03 | 2.01 | 71283 | 1 | 1 | 426460.14 | 441851.38 | 373752.81 | 447369.78 | 442866.59 | 533024.69 | 483106.06 | 467574.28 | 478536.5 | 440785.12 | 425371.88 | 423321.28 | 465400.47 | 482958.31 |
| Q91VE0 | Slc27a4 | Long-chain fatty acid transport protein 4 Mus musculus | 46.27 | 1.4 | 72319 | 1 | 1 | 544022.06 | 804977.62 | 522594.25 | 785672 | 696072.31 | 710662.69 | 1091607.62 | 566765.88 | 801303.19 | 980607.31 | 894143.62 | 868754.88 | 750379.69 | 815482.75 |
| P11167 | Slc2a1 | Solute carrier family 2, facilitated glucose transporter member 1 Rattus norvegicus | 148.34 | 5.69 | 53963 | 4 | 4 | 2903824.5 | 1 | 326483.44 | 1 | 1 | 276774.59 | 333713.94 | 303789.19 | 227634.86 | 329985.53 | 1 | 1 | 260272.08 | 157256.25 |
| Q9JKN1 | Slc30a7 | Zinc transporter 7 Mus musculus | 51.04 | 3.97 | 41790 | 1 | 1 | 1650122.5 | 1308745.75 | 1441584.25 | 1486542.75 | 1122267 | 1789349 | 1599254.12 | 1342025.12 | 1240750.12 | 1306568 | 1251344.25 | 1162397.38 | 1303240.88 | 1283499 |
| Q6AYY8 | Slc33a1 | Acetyl-coenzyme A transporter 1 Rattus norvegicus | 41.69 | 2.36 | 61101 | 1 | 1 | 473776.47 | 1 | 1 | 1 | 205475.41 | 274355.03 | 1 | 1 | 142706.19 | 115260.82 | 147938.41 | 148892.41 | 1 | 1 |
| P97858 | Slc35b1 | Solute carrier family 35 member B1 Mus musculus | 46.8 | 3.11 | 35883 | 1 | 1 | 1833371.62 | 1034841.94 | 1278477.25 | 897085.38 | 978839.69 | 1595296.62 | 876516 | 1010237.56 | 1221088.5 | 925590.31 | 922290.75 | 1204624.38 | 1491669 | 1472770.25 |
| A2AKQ0 | Slc35d1 | UDP-glucuronic acid/UDP-N-acetylgalactosamine transporter Mus musculus | 55.91 | 5.92 | 39272 | 1 | 1 | 238848.25 | 1 | 171245.02 | 1 | 1 | 225676.17 | 1 | 1 | 127506.38 | 1 | 174998.38 | 260140.73 | 228535.5 | 191416.64 |
| Q5RKH7 | Slc35f6 | Solute carrier family 35 member F6 Rattus norvegicus | 40.68 | 4.03 | 41092 | 1 | 1 | 1099326.5 | 1 | 1 | 1 | 1 | 914800.88 | 542464.515 | 856301.38 | 186872.02 | 211883.78 | 807101.3867 | 1040582.38 | 1056716.5 | 324005.28 |
| Q9WU81 | Slc37a2 | Glucose-6-phosphate exchanger SLC37A2 Mus musculus | 160.7 | 7.78 | 55073 | 3 | 3 | 6496954.5 | 1 | 431650.62 | 267736.12 | 386036.78 | 802360.06 | 498235.915 | 318004.94 | 545444.75 | 327133.91 | 230660.08 | 475388 | 639519.25 | 640287.38 |
| Q8CFE6 | Slc38a2 | Sodium-coupled neutral amino acid transporter 2 Mus musculus | 70.86 | 3.77 | 55503 | 2 | 2 | 3341282.5 | 1337418.12 | 1624271.88 | 886244.25 | 842715.62 | 843997 | 890794.38 | 822069.75 | 777737.81 | 839100.62 | 795052.46 | 580019.38 | 1173708.88 | 631429.12 |
| Q6P6S2 | Slc39a11 | Zinc transporter ZIP11 Rattus norvegicus | 78.44 | 6.57 | 34868 | 1 | 1 | 398985.34 | 461895.41 | 569950.69 | 472844.44 | 411936.06 | 629991.25 | 503693.47 | 572189.5 | 603394.38 | 425174.75 | 562263.69 | 523469.59 | 692345.38 | 526708.69 |
| Q78IQ7 | Slc39a4 | Zinc transporter ZIP4 Mus musculus | 201.38 | 12.73 | 71063 | 4 | 4 | 2840990.5 | 2027986.25 | 4243827 | 1403588.5 | 2391140 | 1991673.62 | 979170.56 | 1433551.88 | 748370.44 | 753986.94 | 2292126.75 | 2163706.25 | 4911040 | 2856759.5 |
| Q31125 | Slc39a7 | Zinc transporter SLC39A7 Mus musculus | 136.99 | 8.19 | 50657 | 2 | 2 | 1483161.62 | 1283374.12 | 1112795.5 | 1289551.25 | 1187442.88 | 1364645.62 | 1155224.5 | 1430699 | 184303.66 | 1451737.88 | 1465651.12 | 970540.56 | 1292539.38 | 1315464.12 |
| P10852 | Slc3a2 | 4F2 cell-surface antigen heavy chain Mus musculus | 986.12 | 53.23 | 58337 | 19 | 19 | 41247057 | 18164911 | 19587141 | 18179335 | 20064393 | 17544019 | 16581012 | 16246881 | 16133649 | 15747751 | 14971227 | 13672847 | 18708373 | 17467807 |
| Q8BTY2 | Slc4a7 | Sodium bicarbonate cotransporter 3 Mus musculus | 198.97 | 3.77 | 116514 | 3 | 3 | 1 | 1 | 1 | 1 | 1 | 1 | 1 | 278411.38 | 1 | 142437.42 | 191332.16 | 256858.83 | 744044.69 | 345649.22 |
| Q9CXK4 | Slc50a1 | Sugar transporter SWEET1 Mus musculus | 46.08 | 9.05 | 24649 | 1 | 1 | 1643345 | 1234161.75 | 1432607.12 | 1383773 | 1731784.12 | 1515281.12 | 782830.06 | 848260.19 | 1382884.5 | 883261.06 | 1352029.12 | 1364503.25 | 1329009.88 | 1482030.5 |
| O35316 | Slc6a6 | Sodium- and chloride-dependent taurine transporter Mus musculus | 81.01 | 3.06 | 69856 | 2 | 2 | 11509354 | 10552174 | 11006932 | 8452274 | 9802037 | 11270522 | 10689359 | 10011242 | 10929266 | 10188281 | 9808561 | 9909455 | 11240518 | 11027739 |
| Q6P9P0 | Slf2 | SMC5-SMC6 complex localization factor protein 2 Mus musculus | 36.1 | 0.55 | 143970 | 1 | 1 | 1371724 | 1059111.62 | 392924.06 | 1025785.38 | 1202280.12 | 748016.88 | 1 | 1 | 197343.52 | 1 | 1163531.75 | 1828177.38 | 189973.88 | 1379244.88 |
| B1ARD6 | Slfn9 | Schlafen family member 9 Mus musculus | 404.65 | 14.62 | 103992 | 8 | 8 | 5376753.5 | 4439211 | 5863270 | 3434523.5 | 6125938.5 | 5147135.5 | 5574310.5 | 3647943.5 | 4338384.5 | 5167391 | 4973569 | 8268255 | 6134436 | 5662571 |
| O54988 | Slk | STE20-like serine/threonine-protein kinase Mus musculus | 646.06 | 19.22 | 141457 | 13 | 13 | 9749107 | 8270267 | 9828851 | 15975512 | 13573023 | 10711121 | 11362948 | 9272901 | 10966123 | 9878948 | 10551732 | 9761061 | 9974230 | 11870764 |
| Q8CH25 | Sltm | SAFB-like transcription modulator Mus musculus | 127.99 | 2.42 | 116919 | 2 | 2 | 1 | 3062424.25 | 983633.5 | 1 | 1 | 1801892.62 | 2727775.5 | 1 | 1 | 1 | 1 | 2667404.25 | 726224.5 | 1 |
| O54835 | Smad9 | Mothers against decapentaplegic homolog 9 Rattus norvegicus | 79.33 | 5.53 | 48992 | 2 | 2 | 1104051.5 | 1915360.25 | 1419825.5 | 1939111.5 | 1950017.12 | 1939997 | 1843245.5 | 1728441.75 | 2283356 | 1216681 | 1984123 | 2189404 | 1760471.75 | 1754621.25 |
| Q9R0P4 | Smap | Small acidic protein Mus musculus | 218.45 | 21.55 | 20046 | 2 | 2 | 1115714.75 | 1380716.62 | 1308635.5 | 1145949.62 | 1572490 | 1159948.38 | 1553540.75 | 1560797 | 1270280.5 | 1200256.38 | 1294608 | 1260869 | 1292440.5 | 1184494.88 |
| Q91VZ6 | Smap1 | Stromal membrane-associated protein 1 Mus musculus | 41.36 | 2.73 | 47660 | 1 | 1 | 818227.31 | 1 | 1 | 1 | 1035083.69 | 1 | 1 | 1 | 1 | 1 | 1 | 1 | 983357.62 | 1 |
| Q3TKT4 | Smarca4 | Transcription activator BRG1 Mus musculus | 440.52 | 6.51 | 181427 | 8 | 8 | 3384356.75 | 6823248.5 | 5306927.5 | 5596839.5 | 3889618.25 | 4821368.5 | 3920743 | 3152683.75 | 7357518 | 4153952.25 | 8328985.5 | 7551244.5 | 4671168.5 | 6080734 |
| Q91ZW3 | Smarca5 | SWI/SNF-related matrix-associated actin-dependent regulator of chromatin subfamily A m | 813.84 | 19.7 | 121627 | 18 | 18 | 20324881 | 20998975 | 21987059 | 23634985 | 17373133 | 19249649 | 20166987 | 22187093 | 21088387 | 21512821 | 20458391 | 22545351 | 23958661 | 19917325 |
| Q04692 | Smarcad1 | SWI/SNF-related matrix-associated actin-dependent regulator of chromatin subfamily A co | 34.57 | 1.57 | 116451 | 1 | 1 | 497805.5 | 992761.12 | 860097.06 | 855697.5 | 975915.81 | 748654.56 | 607268.19 | 748390.31 | 778580.5 | 776880.56 | 615306.81 | 1054125.88 | 916806.75 | 968076 |
| B4F769 | Smarcal1 | | 37.68 | 0.99 | 101212 | 1 | 1 | 401547.94 | 351870.12 | 358495.8575 | 324722.06 | 355843.31 | 1 | 333238.34 | 1 | 374463.81 | 421619.81 | 437933.91 | 423986.75 | 346540.47 | 1 |
| Q9Z0H3 | Smarcb1 | SWI/SNF-related matrix-associated actin-dependent regulator of chromatin subfamily B m | 160.83 | 7.79 | 44141 | 2 | 2 | 3590110.5 | 3535351.5 | 4380336 | 4039877 | 4316787 | 4264880.5 | 4292934 | 3439329 | 4608996 | 4141160.5 | 4480635 | 4320599 | 4347393 | 2262003.5 |
| P97496 | Smarcc1 | SWI/SNF complex subunit SMARCC1 Mus musculus | 421.07 | 9.96 | 122890 | 4 | 7 | 15891359 | 19538943 | 17350991 | 13023304 | 10092841 | 12780523 | 16450242 | 17675387 | 12450844 | 11140528 | 11324331 | 19370415 | 11566712 | 10502663 |
| Q6PDG5 | Smarcc2 | SWI/SNF complex subunit SMARCC2 Mus musculus | 426.14 | 7.34 | 132604 | 4 | 7 | 10748632 | 12745533 | 11604058 | 12007481 | 11015288 | 12261596 | 12720306 | 14145898 | 12616769 | 11877902 | 10828069 | 12514804 | 11244882 | 12167279 |
| O54772 | Smarcd2 | SWI/SNF-related matrix-associated actin-dependent regulator of chromatin subfamily D me | 169.1 | 9.23 | 59128 | 3 | 3 | 1 | 1 | 1 | 1 | 229647.23 | 166091.48 | 180889.39 | 176767.5 | 1 | 1 | 1 | 1 | 1 | 1 |
| O54941 | Smarce1 | SWI/SNF-related matrix-associated actin-dependent regulator of chromatin subfamily E m | 151.4 | 6.81 | 46638 | 2 | 2 | 1753462.12 | 2170892.75 | 1625276.88 | 1852051.75 | 1706193.12 | 1632749.25 | 1944933 | 1559611.5 | 1572306.62 | 1684001 | 1681106 | 1572218.88 | 1690301.62 | 1924088 |
| Q9CU62 | Smc1a | Structural maintenance of chromosomes protein 1A Mus musculus | 939.73 | 18.82 | 143235 | 19 | 19 | 17392211 | 15854419 | 16932417 | 13097605 | 16627759 | 12408650 | 15683387 | 11397364 | 16615067 | 15897290 | 14664540 | 16498469 | 17427393 | 12048785 |
| Q8CG48 | Smc2 | Structural maintenance of chromosomes protein 2 Mus musculus | 1010.75 | 28.3 | 134239 | 26 | 26 | 13232687 | 10911837 | 18553215 | 14979910 | 15309549 | 14005323 | 14233445 | 13971462 | 12667563 | 12078784 | 18118129 | 16361701 | 16965931 | 15220894 |
| Q9CW03 | Smc3 | Structural maintenance of chromosomes protein 3 Mus musculus | 1357.97 | 28.02 | 141556 | 26 | 26 | 20230691 | 25036955 | 22632877 | 21715167 | 26981207 | 23496273 | 28286699 | 18013559 | 23229275 | 24623875 | 23403835 | 27095283 | 24449755 | 26419165 |
| Q8CG47 | Smc4 | Structural maintenance of chromosomes protein 4 Mus musculus | 1030.89 | 23.64 | 146895 | 22 | 22 | 11295095 | 13646517 | 16613988 | 17815375 | 16215852 | 15311292 | 13225909 | 13669151 | 13653915 | 15013550 | 15434331 | 17838873 | 16597755 | 14296971 |
| Q6P5D8 | Smchd1 | Structural maintenance of chromosomes flexible hinge domain-containing protein 1 Mus m | 992.29 | 14.25 | 225648 | 18 | 18 | 7834954 | 11752611 | 10098537 | 10123781 | 8980470 | 9383701 | 8715204 | 8427858 | 9084222 | 9998140 | 9443223 | 9592487 | 9314261 | 10895023 |
| P58242 | Smpdl3b | Acid sphingomyelinase-like phosphodiesterase 3b Mus musculus | 269.42 | 18.64 | 51600 | 6 | 6 | 9478891 | 10678620 | 8368701 | 9966309 | 10576836 | 10422891 | 10260592 | 11727323 | 9879722 | 8965652 | 8582033 | 8413799 | 7862410.5 | 6616807.5 |
| P97355 | Sms | Spermine synthase Mus musculus | 332.14 | 28.96 | 41313 | 6 | 6 | 3529398.5 | 4052233.5 | 3624725.75 | 3494258.5 | 3224708.75 | 3132100.25 | 3266161.5 | 3622169.5 | 4132220.25 | 3450940 | 3535642.5 | 3676906 | 3433310.75 | 3699691 |
| Q3UKJ7 | Smu1 | WD40 repeat-containing protein SMU1 Mus musculus | 420.49 | 24.76 | 57544 | 9 | 9 | 5199344.5 | 3802362.75 | 4249785 | 5334716.5 | 5519688 | 5444902.5 | 5548634.5 | 4951032 | 5269399 | 5348591 | 5807639 | 8477330 | 6160661.5 | 6499396.5 |
| Q3TYX3 | Smyd5 | SET and MYND domain-containing protein 5 Mus musculus | 163.18 | 9.13 | 47095 | 3 | 3 | 280062.72 | 286163.16 | 144167.31 | 302051.06 | 416204.91 | 233772.94 | 216568.84 | 141072.84 | 79595.98 | 151395.28 | 170598.12 | 234739.08 | 204072.48 | 235909.88 |
| O09044 | Snap23 | Synaptosomal-associated protein 23 Mus musculus | 89.9 | 6.67 | 23261 | 1 | 1 | 2179954.25 | 2663049.25 | 2291606.75 | 2343632.25 | 1942696.5 | 1913155.38 | 1948148.75 | 2264530 | 1899173.25 | 1916421.5 | 1927433.5 | 1790290 | 1841691.12 | 2130339.5 |
| Q9ERB0 | Snap29 | Synaptosomal-associated protein 29 Mus musculus | 43.16 | 4.23 | 29572 | 1 | 1 | 579252.31 | 476272.28 | 387013.06 | 668182.69 | 526233.5 | 516966.09 | 450213.56 | 585002.75 | 397219.84 | 589578.62 | 559378.81 | 475065.94 | 548980.44 | 387626.38 |
| Q78PY7 | Snd1 | Staphylococcal nuclease domain-containing protein 1 Mus musculus | 2243.23 | 53.52 | 102088 | 38 | 38 | 97090297 | 95015009 | 104223953 | 101757113 | 98515169 | 93528193 | 98166561 | 96836177 | 98623473 | 97329601 | 99314841 | 98342345 | 96199889 | 99541633 |
| Q5RK19 | Snf8 | Vacuolar-sorting protein SNF8 Rattus norvegicus | 42.99 | 5.43 | 28884 | 1 | 1 | 789109.94 | 790146.25 | 834083.19 | 924989.38 | 875019.69 | 845466.88 | 987617.62 | 803863.69 | 968269.62 | 1062072.38 | 787623.25 | 802476.06 | 817779.31 | 956187.25 |
| F1LNJ2 | Snrnp200 | U5 small nuclear ribonucleoprotein 200 kDa helicase Rattus norvegicus | 2586.93 | 31.09 | 244875 | 48 | 48 | 60487281 | 58840725 | 58822573 | 57602289 | 63683121 | 65421589 | 59854937 | 60327913 | 58050285 | 55146281 | 55864897 | 57200417 | 63094613 | 59126217 |
| Q6PE01 | Snrnp40 | U5 small nuclear ribonucleoprotein 40 kDa protein Mus musculus | 382.16 | 32.4 | 39276 | 5 | 5 | 1 | 1 | 1 | 1032575.38 | 1487945.12 | 1 | 1263140.38 | 207364.67 | 1216035.38 | 1 | 929597.06 | 1614495.88 | 1180000.38 | 1241364.44 |
| Q62376 | Snrnp70 | U1 small nuclear ribonucleoprotein 70 kDa Mus musculus | 408.36 | 17.86 | 51992 | 7 | 7 | 16101511 | 13816689 | 11579233 | 17936449 | 16424201 | 17819133 | 16565286 | 17330511 | 14252582 | 18319725 | 14395667 | 22049619 | 15311319 | 20539465 |
| Q62189 | Snrpa | U1 small nuclear ribonucleoprotein A Mus musculus | 295.14 | 31.71 | 31835 | 5 | 6 | 5356152.5 | 5656992.5 | 5636798.5 | 5084548 | 5502430 | 4676490.5 | 5325697.5 | 3946715.5 | 3584942.25 | 4370346.5 | 5313226 | 4528311.5 | 3996467.25 | 4890053.5 |
| P57784 | Snrpa1 | U2 small nuclear ribonucleoprotein A' Mus musculus | 577.45 | 45.88 | 28357 | 11 | 11 | 11704108 | 11399720 | 13774256 | 15603991 | 11045075 | 8645690 | 10169872 | 6992035.5 | 14930463 | 14816768 | 10884130 | 10504986 | 8214811 | 6768221.5 |
| P17136 | Snrpb | Small nuclear ribonucleoprotein-associated protein B Rattus norvegicus | 181.4 | 12.99 | 23656 | 3 | 3 | 6944501.5 | 3822040 | 5843892.5 | 7536809 | 5309032.5 | 4283071 | 5365199.5 | 3946235 | 5444619.5 | 5694206.5 | 5509747 | 6048242.5 | 5659330.5 | 5223806 |
| Q9CQI7 | Snrpb2 | U2 small nuclear ribonucleoprotein B'' Mus musculus | 187.01 | 20 | 25323 | 3 | 4 | 1173842.485 | 341499.44 | 337097.5 | 2369642 | 1647131 | 1 | 1855014.12 | 1 | 1663826 | 2056145.5 | 752565.31 | 1533280.62 | 1715569.25 | 1615603.38 |
| D3ZCL3 | Snrpc | U1 small nuclear ribonucleoprotein C Rattus norvegicus | 75.42 | 7.55 | 17364 | 1 | 1 | 1886324.38 | 2063909.12 | 1915900.5 | 1998641.38 | 2116162.25 | 1993456.5 | 1992491.75 | 2084176.75 | 1901863 | 2342241.25 | 1790236.12 | 2000477.88 | 1944559.25 | 1954809.12 |
| P62315 | Snrpd1 | Small nuclear ribonucleoprotein Sm D1 Mus musculus | 246.73 | 54.62 | 13282 | 5 | 5 | 2063207 | 1520507.5 | 1185624.62 | 2076748.62 | 2390305.5 | 1533975.38 | 1767368 | 99431.53 | 1283037.633 | 1731375.62 | 2031256.62 | 1253606.62 | 1 | 923692.44 |
| P62317 | Snrpd2 | Small nuclear ribonucleoprotein Sm D2 Mus musculus | 221.84 | 35.59 | 13527 | 5 | 5 | 2880013.25 | 1414651.88 | 1495560 | 1183361.5 | 1269746 | 2393292 | 2161647 | 1739079 | 1189461.75 | 3037729.25 | 2396324.75 | 2232480.75 | 763082.12 | 2984825.75 |
| P62320 | Snrpd3 | Small nuclear ribonucleoprotein Sm D3 Mus musculus | 88.47 | 15.08 | 13916 | 2 | 2 | 25269827 | 26480363 | 28734623 | 31722185 | 32007723 | 29846609 | 29218471 | 26075149 | 30747547 | 31018585 | 29322917 | 32433871 | 32407351 | 32631945 |
| P62305 | Snrpe | Small nuclear ribonucleoprotein E Mus musculus | 181.2 | 52.17 | 10804 | 3 | 4 | 24198429 | 32978761 | 34201445 | 29037191 | 30816033 | 30674593 | 28179145 | 30006379 | 32457265 | 28392413 | 26327205 | 31450059 | 31658517 | 31112185 |
| P62307 | Snrpf | Small nuclear ribonucleoprotein F Mus musculus | 229.71 | 83.72 | 9725 | 4 | 4 | 1863000 | 1003075.06 | 1714080 | 1687725.12 | 2597979.25 | 2818866.25 | 1291797.5 | 1215975.5 | 2022246.5 | 2745812.75 | 2160090.5 | 2909514.5 | 3622878.75 | 2802040 |
| Q61235 | Sntb2 | Beta-2-syntrophin Mus musculus | 71.99 | 1.73 | 56382 | 1 | 1 | 449934.72 | 439862.88 | 453626.09 | 498403.125 | 650188.81 | 572536 | 494738.5 | 504387.12 | 469149.09 | 585222.12 | 480050.78 | 472516.12 | 426191.38 | 556519.44 |
| P55770 | Snu13 | NHP2-like protein 1 Rattus norvegicus | 235.11 | 55.47 | 14174 | 5 | 5 | 13444588 | 11597464 | 13798339 | 15383133 | 16392825 | 13639129 | 12975607 | 10907711 | 13335884 | 15811525 | 14713782 | 13606502 | 14204947 | 13574032 |

| | | | | | | | | | | | | | | | | | | | | | |
|---|---|---|---|---|---|---|---|---|---|---|---|---|---|---|---|---|---|---|---|---|---|
| Q9CSN1 | Snw1 | SNW domain-containing protein 1 Mus musculus | 345.85 | 15.11 | 61475 | 6 | 6 | 3172287.75 | 1872244.62 | 3028839.5 | 1570786.62 | 4597744 | 3223292.5 | 5472952 | 4075663 | 4164899.25 | 3685650 | 2819274.5 | 5141239 | 2899771 | 4628490 |
| Q99N27 | Snx1 | Sorting nexin-1 Rattus norvegicus | 741.77 | 30.65 | 59044 | 13 | 14 | 27401709 | 34500917 | 28031507 | 29407069 | 30222635 | 28073307 | 27331505 | 27771389 | 26482087 | 27086541 | 28986517 | 26716427 | 26679225 | 28245859 |
| O70493 | Snx12 | Sorting nexin-12 Mus musculus | 245.49 | 44.24 | 19116 | 4 | 5 | 7651322 | 6783016 | 6796568.5 | 5799132 | 7286726 | 6579928.5 | 5671096.5 | 6048533.5 | 6780847.5 | 5228042.5 | 3844429 | 6660799.5 | 5211139.5 | 4131628 |
| Q8BVL3 | Snx17 | Sorting nexin-17 Mus musculus | 151.8 | 13.62 | 52797 | 4 | 4 | 1532372 | 157143.75 | 918231.44 | 1128782.25 | 934132.36 | 1 | 692742.06 | 620969.5 | 1 | 1 | 1 | 619672.94 | 1 | 1 |
| Q91ZR2 | Snx18 | Sorting nexin-18 Mus musculus | 131.78 | 6.35 | 67904 | 3 | 3 | 5560716.5 | 3640845.5 | 2257478.5 | 3922595.5 | 2254448 | 1998626.5 | 3521044.5 | 4312154.5 | 3380272.25 | 4454715 | 3530497.25 | 3090289 | 1845515 | 3529107.5 |
| Q9CWK8 | Snx2 | Sorting nexin-2 Mus musculus | 1058.94 | 48.36 | 58471 | 17 | 18 | 42139717 | 39019877 | 42667173 | 42373893 | 37328149 | 38252785 | 38707789 | 42189829 | 41611285 | 40859901 | 39611117 | 37980489 | 33742225 | 38470109 |
| Q5BK61 | Snx20 | Sorting nexin-20 Rattus norvegicus | 45.89 | 3.83 | 35723 | 1 | 1 | 781292.69 | 719573.88 | 749767.56 | 606141.81 | 766262.44 | 616120.06 | 536662.12 | 528662.44 | 693264.19 | 634766.38 | 597233.88 | 509362.72 | 576291.06 | 601287.12 |
| Q3UHD6 | Snx27 | Sorting nexin-27 Mus musculus | 160.06 | 9.28 | 60989 | 3 | 3 | 1525009.533 | 1829758.5 | 187545 | 1828063.88 | 2254670.75 | 316077.78 | 1405659.883 | 1862064.75 | 1894744.88 | 1549752.12 | 224955.27 | 243713.1533 | 258143.86 | 248040.33 |
| Q9D3S3 | Snx29 | Sorting nexin-29 Mus musculus | 36.24 | 1.59 | 91387 | 1 | 1 | 258800.97 | 299483.16 | 236869.2 | 275548.41 | 215580.48 | 306104.16 | 248255.6675 | 211654.98 | 266195.78 | 209067.75 | 237601.9233 | 264220.56 | 254943.77 | 193641.44 |
| O70492 | Snx3 | Sorting nexin-3 Mus musculus | 283.42 | 48.15 | 18757 | 5 | 6 | 10380674 | 7210111.5 | 11032516 | 8297273.5 | 7993428 | 7603458 | 6750667.5 | 7861677.5 | 7952916 | 8200715.5 | 6567756.5 | 7919320.5 | 8856754 | 7523027 |
| Q8CE50 | Snx30 | Sorting nexin-30 Mus musculus | 44.73 | 4.35 | 49520 | 1 | 1 | 210270.38 | 292144.66 | 360198.59 | 512337.06 | 174994.08 | 345869.62 | 445195.09 | 308247.72 | 358119.91 | 337775.25 | 468833.31 | 374779.53 | 416582.31 | 272758.53 |
| Q91YJ2 | Snx4 | Sorting nexin-4 Mus musculus | 69.29 | 2 | 51778 | 1 | 1 | 1887102.38 | 1511936.88 | 1589528.12 | 1633072.62 | 2072632.62 | 1957162.12 | 1791263.75 | 1293141.38 | 1620400.62 | 1813740.5 | 2070487.25 | 2146586.25 | 1979416 | 1865476 |
| Q9D8U8 | Snx5 | Sorting nexin-5 Mus musculus | 926.58 | 45.3 | 46797 | 18 | 18 | 50526221 | 47983113 | 50641529 | 45504009 | 45378405 | 47334905 | 52106665 | 57279193 | 61884097 | 50794889 | 45286033 | 50778269 | 48015729 | 48074653 |
| Q6P8X1 | Snx6 | Sorting nexin-6 Mus musculus | 205.46 | 15.02 | 46649 | 5 | 5 | 2909167.25 | 4147249.5 | 2350855.5 | 1733028.88 | 2891085.5 | 3864976 | 3455306.5 | 3376617 | 3798492 | 3037675 | 4332303 | 2135670 | 4031987 | 2546903 |
| Q9CY18 | Snx7 | Sorting nexin-7 Mus musculus | 41.25 | 3.1 | 45000 | 1 | 1 | 1 | 1220634.25 | 1 | 938071.06 | 1293163.12 | 1268477.12 | 722520.38 | 323900.09 | 430469.56 | 867234.75 | 1028038.31 | 775994.12 | 1 | 1 |
| Q8CFD4 | Snx8 | Sorting nexin-8 Mus musculus | 103.54 | 7.19 | 52059 | 2 | 2 | 301617.22 | 1 | 745870.19 | 1 | 1 | 186769.31 | 1 | 382347.03 | 438909.78 | 1 | 139506.12 | 220899.52 | 206161.77 | 188855.8033 |
| Q91VH2 | Snx9 | Sorting nexin-9 Mus musculus | 103.12 | 7.73 | 66546 | 2 | 2 | 1294086.12 | 870328.25 | 1544029 | 670678.25 | 1034551 | 1042604.62 | 1246798.5 | 1385813.5 | 2242280.5 | 1229000.75 | 461820.06 | 685326.75 | 1539839.62 | 1335905 |
| Q61263 | Soat1 | Sterol O-acyltransferase 1 Mus musculus | 297.96 | 13.7 | 63799 | 5 | 5 | 7986168 | 791642.75 | 2632665.5 | 2175211.5 | 1571731.5 | 1999565.25 | 1075641.75 | 2410815 | 2589225 | 1638663.5 | 1985424.25 | 2531148.25 | 2788167.75 | 2354726 |
| P08228 | Sod1 | Superoxide dismutase [Cu-Zn] Mus musculus | 269.37 | 32.47 | 15943 | 4 | 4 | 34171897 | 36139077 | 31668321 | 38968433 | 37639953 | 25617309 | 33953793 | 37489441 | 28651593 | 37931597 | 40141969 | 26710577 | 30773029 | 41788033 |
| P09671 | Sod2 | Superoxide dismutase [Mn], mitochondrial Mus musculus | 292.17 | 41.89 | 24603 | 5 | 5 | 24467249 | 28106105 | 30527805 | 22500363 | 25449799 | 22629517 | 25327485 | 22465857 | 22749131 | 26749399 | 24019129 | 23885713 | 32733925 | 26219795 |
| Q9QX47 | Son | Protein SON Mus musculus | 301.94 | 3.48 | 265651 | 5 | 5 | 6658958 | 6934572 | 5505999 | 6920907 | 7817023.5 | 8059071 | 7077709 | 6571938.5 | 7908050 | 7143248 | 7218888 | 6854021 | 7051892 | 6960747 |
| O35892 | Sp100 | Nuclear autoantigen Sp-100 Mus musculus | 189.49 | 9.96 | 54727 | 3 | 4 | 3439218.5 | 3207927 | 3216481.25 | 2612204 | 2956948.5 | 1324223.75 | 2217736.75 | 2585625 | 2355409.25 | 2315391.5 | 2404158.75 | 2575712.25 | 2204169 | 1244349 |
| Q8BVK9 | Sp110 | Sp110 nuclear body protein Mus musculus | 472.34 | 32.58 | 50140 | 10 | 10 | 9270450 | 7255536 | 8656968 | 7135867.5 | 8281462.5 | 5700259 | 6962201 | 6817715.5 | 6990179.5 | 6948625.5 | 5176007 | 5951477 | 6941924.5 | 5088065.5 |
| Q7TNE3 | Spag7 | Sperm-associated antigen 7 Mus musculus | 88.28 | 15.86 | 25921 | 2 | 2 | 615179.38 | 506796 | 641619.44 | 668591.81 | 904419.81 | 601676.38 | 779538.5 | 546663.62 | 180243.08 | 1244327.5 | 599050.06 | 1 | 1 | 998339.81 |
| Q58A65 | Spag9 | C-Jun-amino-terminal kinase-interacting protein 4 Mus musculus | 403.93 | 9.01 | 146219 | 7 | 7 | 4373358.5 | 4123626 | 3886400 | 3164691 | 5128186 | 6293938.5 | 5564882.5 | 6968959 | 6788624.5 | 5099077 | 5801966 | 6487378 | 4663699 | 6591536 |
| Q8R1X6 | Spart | Spartin Mus musculus | 64.11 | 1.94 | 72655 | 1 | 1 | 678906.94 | 1005219.44 | 895302.44 | 923381.62 | 997698.25 | 885141.25 | 909600.31 | 775472.12 | 1253543.12 | 1276951.88 | 936625.25 | 913758.96 | 726321.75 | 1078329.88 |
| B2RYN7 | Spast | Spastin Rattus norvegicus | 79.52 | 6.88 | 63022 | 2 | 2 | 533605.88 | 178711.44 | 314964.69 | 300102.91 | 403308.5 | 506577.25 | 513196.56 | 423320.7975 | 411261.41 | 262247.97 | 454910.97 | 346881.81 | 387045.34 | 238664.62 |
| Q0P557 | Spata18 | Mitochondria-eating protein Mus musculus | 40.01 | 1.49 | 60398 | 1 | 1 | 1319964.88 | 1498278.88 | 1609656 | 1620784.62 | 1431329 | 1578343.25 | 1557069.38 | 1207620.88 | 1421358.88 | 1402721.25 | 1211029.12 | 1367916.88 | 1264456.62 | 1698195.25 |
| Q9D083 | Spc24 | Kinetochore protein Spc24 Mus musculus | 155.91 | 21.89 | 23416 | 3 | 3 | 2456834.5 | 2157392 | 1952820.75 | 1715648.5 | 1469050.62 | 1410195.38 | 1874569 | 2875444 | 1792200 | 1630219 | 1216369.75 | 1948831.5 | 1951733 | 1758277.75 |
| Q3UA16 | Spc25 | Kinetochore protein Spc25 Mus musculus | 43.59 | 4.87 | 26457 | 1 | 1 | 871541.06 | 1022696.31 | 983653.56 | 1 | 1 | 812132.38 | 944976 | 706657 | 650729.19 | 642622 | 748906.69 | 747162.6867 | 704672.75 | 787908.62 |
| Q9CYN2 | Spcs2 | Signal peptidase complex subunit 2 Mus musculus | 180.34 | 19.47 | 24978 | 4 | 4 | 2645786.75 | 1444518.75 | 2136541 | 1882667.62 | 1688463.88 | 1606329.75 | 1438681.5 | 2694929 | 2410110 | 1471654.5 | 1336058.5 | 2304954.25 | 3499497 | 197386.58 |
| Q6ZWQ7 | Spcs3 | Signal peptidase complex subunit 3 Mus musculus | 177.23 | 21.67 | 20313 | 4 | 4 | 5424392.5 | 4938548 | 4468089 | 5067779.5 | 4013489.25 | 4247326 | 5212095.5 | 5686997.5 | 4508852.5 | 4656359.5 | 4838942.5 | 4223543.5 | 4512127 | 4298159 |
| Q5SXY1 | Specc1 | Cytospin-B Mus musculus | 51.34 | 0.75 | 118101 | 1 | 1 | 891141.69 | 461935.31 | 915271.12 | 938618.12 | 619028.31 | 484018.22 | 863978 | 397164.22 | 638635.06 | 713719.81 | 756569.25 | 698713.25 | 762819.12 | 597363.19 |
| Q9JIA7 | Sphk2 | Sphingosine kinase 2 Mus musculus | 244.92 | 8.75 | 65618 | 4 | 4 | 3011618 | 3209972.75 | 3824089.25 | 3107739.5 | 2815345.25 | 3713318.25 | 2953592.75 | 1925856.62 | 2451323.75 | 2990944 | 2769623 | 3263840 | 3718436.25 | 2543909 |
| P17433 | Spi1 | Transcription factor PU.1 Mus musculus | 74.39 | 3.68 | 31349 | 1 | 1 | 860899.19 | 1514500.38 | 1234576.88 | 1313593.12 | 975769.31 | 1331016.38 | 1137715.88 | 1277902.62 | 1355711.62 | 1132679.38 | 1461340 | 1607997.25 | 1351214.12 | 1368701.75 |
| Q2YDU8 | Spns1 | Protein spinster homolog 1 Rattus norvegicus | 74.92 | 5.11 | 56796 | 1 | 1 | 1171658.5 | 1 | 739291.38 | 1 | 1 | 819386 | 687770.06 | 482149.09 | 1 | 1 | 224627.8 | 1 | 1 | 559534.06 |
| Q3UHX9 | Spout1 | Putative methyltransferase C9orf114 homolog Mus musculus | 38.88 | 3.12 | 42958 | 1 | 1 | 1 | 1 | 1 | 255840.88 | 198152.83 | 1 | 1 | 1 | 223415.86 | 256685.94 | 1 | 1 | 257109.16 | 261735.67 |
| Q64105 | Spr | Sepiapterin reductase Mus musculus | 732.36 | 58.62 | 27883 | 11 | 11 | 20878269 | 23049057 | 24121913 | 26636331 | 23832011 | 22090557 | 22394231 | 19683771 | 22607535 | 23775433 | 24603033 | 23322133 | 18391023 | 24224185 |
| O88838 | Spsb2 | SPRY domain-containing SOCS box protein 2 Mus musculus | 35.47 | 2.65 | 28938 | 1 | 1 | 9523748 | 11776717 | 10869080 | 11468616 | 11094955 | 11049018 | 9252005 | 10523891 | 8604649 | 8720057 | 10494177 | 9512201 | 8680596 | 10287478 |
| P16086 | Sptan1 | Spectrin alpha chain, non-erythrocytic 1 Rattus norvegicus | 117.82 | 1.5 | 284637 | 2 | 2 | 360823.97 | 1043366.38 | 1750962.62 | 868981.36 | 320772.47 | 1226098 | 436591.47 | 2273786.25 | 662460.94 | 509778.22 | 676115.31 | 476022.72 | 1589390.25 | 557648.38 |
| Q62261 | Sptbn1 | Spectrin beta chain, non-erythrocytic 1 Mus musculus | 195.59 | 2.54 | 274223 | 5 | 5 | 6039017 | 5244930 | 4114364.25 | 5571106.5 | 5208593.5 | 4525477.5 | 5072192 | 4997175.5 | 4585712.5 | 4859376 | 5406274.5 | 6263796.5 | 2841320 | 5400430 |
| O35704 | Sptlc1 | Serine palmitoyltransferase 1 Mus musculus | 160.62 | 6.13 | 52535 | 2 | 2 | 101991.68 | 467272.62 | 497187.44 | 432215.12 | 353916.41 | 181767.88 | 578589.94 | 132957.73 | 290637.5 | 402233.25 | 456314.25 | 206450.14 | 155505.42 | 435008.22 |
| P97363 | Sptlc2 | Serine palmitoyltransferase 2 Mus musculus | 89.65 | 6.79 | 62982 | 2 | 2 | 719878.12 | 1 | 1 | 1 | 1 | 225804.12 | 137711.28 | 1 | 1 | 1 | 244210.47 | 250063.9133 | 328912.44 | 177068.83 |
| Q9R112 | Sqor | Sulfide:quinone oxidoreductase, mitochondrial Mus musculus | 758.22 | 35.11 | 50282 | 12 | 12 | 26959017 | 29870575 | 30165297 | 28299773 | 34691545 | 34952825 | 32642601 | 34754561 | 31669533 | 32321261 | 29832653 | 26671445 | 26918625 | 29418213 |
| Q64337 | Sqstm1 | Sequestosome-1 Mus musculus | 493.06 | 34.62 | 48163 | 8 | 10 | 9116029 | 7441562 | 8011082 | 6849564 | 4125496.5 | 6949940 | 6640018.5 | 7951936.5 | 6819349 | 8285588 | 7666042.5 | 6588617 | 7271688 | 6993648 |
| Q497V5 | Srbd1 | S1 RNA-binding domain-containing protein 1 Mus musculus | 286.78 | 9.67 | 110281 | 7 | 7 | 34034501 | 33414335 | 41727673 | 34932977 | 36857381 | 35845105 | 33420093 | 30797405 | 36904221 | 41665877 | 40572377 | 41018141 | 41726513 | 41244173 |
| D4A208 | Srgap2 | SLIT-ROBO Rho GTPase-activating protein 2 Rattus norvegicus | 536.55 | 12.04 | 120878 | 11 | 11 | 11569112 | 10469069 | 10177344 | 8966176 | 11575758 | 8486698 | 8293533 | 10040669 | 12452510 | 12026647 | 10060167 | 10618472 | 11750848 | 10851407 |
| Q812A2 | Srgap3 | SLIT-ROBO Rho GTPase-activating protein 3 Mus musculus | 77.63 | 1.09 | 124420 | 1 | 1 | 402724.81 | 569853.25 | 428571.34 | 488392.74 | 552421.56 | 450833 | 425662.25 | 637870.12 | 587764 | 492096.56 | 440007.12 | 502295.22 | 412182.41 | 413419.12 |
| Q6P069 | Sri | Sorcin Mus musculus | 230.69 | 26.26 | 21627 | 5 | 5 | 6887902.5 | 8405801 | 7203348 | 8216794 | 8774899 | 7638637 | 8369025 | 7215694.5 | 8286242 | 8340127 | 7904559 | 6311475.5 | 7305117 | 8534839 |
| Q64674 | Srm | Spermidine synthase Mus musculus | 482.09 | 44.04 | 33995 | 10 | 10 | 10481149 | 15168779 | 11882889 | 14265563 | 16794055 | 18197215 | 14969335 | 13817627 | 16567729 | 19041729 | 15874473 | 16273927 | 18256349 | 20045125 |
| P16254 | Srp14 | Signal recognition particle 14 kDa protein Mus musculus | 157.1 | 33.64 | 12510 | 3 | 3 | 2109382.5 | 2303947.75 | 1127622.5 | 1105371.38 | 1661581.033 | 1927260 | 2324075.75 | 1798871.063 | 1381858.25 | 1562290.25 | 1272038.73 | 2568431.75 | 521756.38 | 725928.06 |
| Q9D7A6 | Srp19 | Signal recognition particle 19 kDa protein Mus musculus | 108.91 | 25.69 | 16191 | 2 | 2 | 1186770.88 | 1212089.62 | 1358957.625 | 1583497.5 | 1453472.5 | 754597.88 | 1050707 | 973517.75 | 1079149.5 | 1318759.75 | 1367959 | 710563.19 | 1729218 | 1416048.62 |
| P14576 | Srp54 | Signal recognition particle 54 kDa protein Mus musculus | 252.93 | 15.67 | 55721 | 6 | 6 | 4842025.5 | 4733749 | 4221085.5 | 5124884 | 2779329.5 | 4865895.5 | 5263516.5 | 5245638 | 4834873.5 | 3661544.5 | 4738544.5 | 2365878.25 | 3841788.75 | 3161657 |
| Q8BMA6 | Srp68 | Signal recognition particle subunit SRP68 Mus musculus | 758.24 | 24 | 70574 | 12 | 12 | 8877004 | 9019523 | 9309933 | 9427344 | 8799103 | 9350803 | 8182989 | 8568866 | 9923001 | 7162825 | 8237717 | 7605911.5 | 8263081 | 6750717.5 |
| P49962 | Srp9 | Signal recognition particle 9 kDa protein Mus musculus | 77.08 | 25.58 | 10194 | 2 | 2 | 2654223 | 2354702.25 | 2011528.75 | 2710504.5 | 2749296.5 | 2272481 | 2055546.25 | 2553878.75 | 2419980.25 | 2663879 | 1453997.5 | 2411694 | 2046090.75 | 1588952.88 |
| O70551 | Srpk1 | SRSF protein kinase 1 Mus musculus | 500.77 | 19.44 | 73088 | 9 | 9 | 9392259 | 4168226.25 | 2440486.5 | 3807400.5 | 3431408 | 4758589 | 3311573.5 | 5130711 | 3212916 | 3562780.5 | 3891687.25 | 4715940 | 5751209.5 | 4114063.25 |
| Q9DBG7 | Srpra | Signal recognition particle receptor subunit alpha Mus musculus | 906.16 | 32.55 | 69623 | 14 | 14 | 14421949 | 13493169 | 12227201 | 20366015 | 15431319 | 21736667 | 19977879 | 16778509 | 17088211 | 15637099 | 17546541 | 23053753 | 13319934 | 14624238 |
| P47758 | Srprb | Signal recognition particle receptor subunit beta Mus musculus | 350.86 | 32.34 | 29579 | 7 | 7 | 1671979.75 | 4487028.5 | 3294738.25 | 1963707.12 | 1942231.12 | 2308206.25 | 4480727.5 | 534949.31 | 2878709.25 | 1897107.5 | 4439762.5 | 4739471 | 3775860.5 | 1818012.12 |
| Q52KI8 | Srrm1 | Serine/arginine repetitive matrix protein 1 Mus musculus | 417.81 | 10.25 | 106862 | 7 | 7 | 4782300 | 5934505 | 7111439 | 6608492 | 6399699 | 8489364 | 8690769 | 6491351 | 7436672 | 9134941 | 6004613 | 5607186 | 6368805.5 | 6769152.5 |
| Q8BTI8 | Srrm2 | Serine/arginine repetitive matrix protein 2 Mus musculus | 514.83 | 6.51 | 294840 | 11 | 11 | 4423304.5 | 5107881.5 | 4832537 | 4396774 | 5783462.5 | 5228192 | 4870759.5 | 5077284 | 4407644 | 6233854.5 | 4546439.5 | 2564794 | 5010153.5 | 4532779 |
| Q99MR6 | Srrt | Serrate RNA effector molecule homolog Mus musculus | 568.12 | 19.09 | 100452 | 12 | 12 | 13330740 | 11962635 | 18398007 | 19085989 | 19595179 | 19613069 | 19687889 | 15094325 | 18295077 | 20801569 | 20443617 | 19433533 | 13305717 | 14978082 |
| Q6PDM2 | Srsf1 | Serine/arginine-rich splicing factor 1 Mus musculus | 549.92 | 37.5 | 27745 | 11 | 11 | 32474185 | 35429337 | 29109567 | 40848225 | 45081749 | 38521017 | 34398965 | 31547201 | 33121505 | 39097873 | 41449413 | 35286017 | 37299941 | 40843609 |
| Q9R0U0 | Srsf10 | Serine/arginine-rich splicing factor 10 Mus musculus | 316.32 | 23.28 | 31301 | 5 | 5 | 10842501 | 12178756 | 12920575 | 5026208 | 9880966 | 10558975 | 11237553 | 11920330 | 9776083 | 9784673 | 10450442 | 10813421 | 11098500 | 10192795 |
| Q62093 | Srsf2 | Serine/arginine-rich splicing factor 2 Mus musculus | 361.38 | 25.34 | 25476 | 5 | 5 | 14396855 | 13050013 | 14028079 | 16717478 | 16280948 | 14082640 | 15242735 | 11715017 | 14740163 | 18901017 | 14129121 | 18081117 | 15214867 | 14309774 |
| P84104 | Srsf3 | Serine/arginine-rich splicing factor 3 Mus musculus | 264.33 | 36.59 | 19330 | 5 | 6 | 30546295 | 35529941 | 32581633 | 41087673 | 38271949 | 35735241 | 36339281 | 30975369 | 36331005 | 41544049 | 36529765 | 38176465 | 36328581 | 38505389 |
| Q8VE97 | Srsf4 | Serine/arginine-rich splicing factor 4 Mus musculus | 302.3 | 13.29 | 55979 | 4 | 6 | 33271275 | 35565693 | 32026681 | 37520329 | 38076009 | 35497893 | 34167261 | 31855183 | 34413597 | 36812561 | 32210303 | 34073529 | 33903221 | 36091677 |
| O35326 | Srsf5 | Serine/arginine-rich splicing factor 5 Mus musculus | 351.71 | 25.28 | 30891 | 5 | 6 | 46996529 | 46404385 | 42676233 | 45301057 | 48428285 | 44846393 | 42924545 | 38399329 | 42644145 | 47435765 | 39357057 | 39502393 | 41972405 | 41565229 |
| G3V6S8 | Srsf6 | Serine/arginine-rich splicing factor 6 Rattus norvegicus | 448.16 | 24.78 | 39025 | 6 | 8 | 42628281 | 43954549 | 39842809 | 43961929 | 45972813 | 41959569 | 41276817 | 36122657 | 38700201 | 44303345 | 40271109 | 50164573 | 41888933 | 42660633 |
| Q8BL97 | Srsf7 | Serine/arginine-rich splicing factor 7 Mus musculus | 234.32 | 18.73 | 30818 | 4 | 5 | 11821619 | 11745668 | 12058998 | 17130155 | 15611909 | 14740214 | 15585223 | 11005029 | 15954647 | 18885989 | 15550232 | 18379949 | 18221037 | 17832193 |
| P32067 | Ssb | Lupus La protein homolog Mus musculus | 635.85 | 26.51 | 47756 | 11 | 11 | 12495275 | 10936265 | 11841657 | 11789481 | 11767524 | 11350290 | 11468199 | 12065281 | 10152066 | 11589815 | 10674655 | 10565881 | 12888687 | 11569160 |
| Q9CYR0 | Ssbp1 | Single-stranded DNA-binding protein, mitochondrial Mus musculus | 161.08 | 30.92 | 17319 | 4 | 4 | 12816508 | 15230558 | 12364125 | 13573183 | 14595897 | 13383653 | 12251699 | 14448597 | 13642133 | 13191951 | 13832618 | 11691993 | 14068355 | 14113761 |
| Q7TPJ0 | Ssr1 | Translocon-associated protein subunit alpha Rattus norvegicus | 155.14 | 10.66 | 35629 | 3 | 3 | 13274833 | 16414066 | 16455467 | 14157776 | 14268068 | 18070115 | 14397219 | 18274321 | 14744935 | 14873237 | 15781225 | 17872257 | 18921545 | 16792925 |
| Q08013 | Ssr3 | Translocon-associated protein subunit gamma Rattus norvegicus | 88.41 | 7.57 | 21064 | 1 | 1 | 5342460 | 4176789.5 | 4127466.5 | 3821924.25 | 3767802.25 | 4235953 | 3514747 | 4122671.5 | 3312121.25 | 3523313 | 3998947.25 | 3724067.25 | 4713748 | 4336218 |
| Q07984 | Ssr4 | Translocon-associated protein subunit delta Rattus norvegicus | 205.41 | 24.86 | 18980 | 3 | 3 | 5593567.5 | 3916806.5 | 3909770.5 | 3409753 | 3364499.25 | 3113920.75 | 4016415.25 | 3592383.5 | 2798602.5 | 2700922.25 | 3816575.25 | 2360575.75 | 4111493 | 3716528.25 |
| Q04931 | Ssrp1 | FACT complex subunit SSRP1 Rattus norvegicus | 514.66 | 19.04 | 80914 | 9 | 9 | 8968763 | 8489271 | 8607678 | 9083119 | 9493513 | 8889821 | 7533185.5 | 9590582 | 8814054 | 7891401 | 8659560 | 8861205 | 9339470 | 9331576 |
| Q4KLK9 | Ssu72 | RNA polymerase II subunit A C-terminal domain phosphatase SSU72 Rattus norvegicus | 83.33 | 12.37 | 22544 | 2 | 2 | 1450481.88 | 1394365.25 | 2578613.75 | 1868535 | 1867524.38 | 2250835.5 | 2216127.5 | 1592567 | 2072365 | 2073691.75 | 1973185.62 | 2273947.5 | 1981565.75 | 2178772 |
| Q99L47 | St13 | Hsc70-interacting protein Mus musculus | 558.98 | 23.99 | 41656 | 11 | 11 | 29978735 | 32553649 | 36151301 | 31068037 | 31646495 | 30108457 | 29894943 | 29257873 | 29918037 | 32965165 | 32249677 | 32443751 | 32837137 | 34951865 |
| O88829 | St3gal5 | Lactosylceramide alpha-2,3-sialyltransferase Mus musculus | 64.94 | 5.56 | 47360 | 2 | 2 | 2107032.25 | 2109276.25 | 1447866.62 | 2348310.5 | 2417529.5 | 2630961 | 2028636.38 | 1924562.75 | 2392820 | 2569061.75 | 2553602.5 | 2433463.5 | 2611923 | 2993465 |
| Q9D3E6 | Stag1 | Cohesin subunit SA-1 Mus musculus | 290.82 | 6.04 | 144440 | 4 | 6 | 3265961.5 | 1683671.88 | 2003439.88 | 806558.44 | 1580293 | 3739003.25 | 1334966.62 | 4157064 | 3965190.75 | 1792086.75 | 1702567.5 | 2004979.5 | 2323392.25 | 1702994.25 |
| O35638 | Stag2 | Cohesin subunit SA-2 Mus musculus | 406.31 | 10.24 | 141281 | 7 | 9 | 6644299.5 | 3486620.25 | 3699485.5 | 2715289.75 | 3123931 | 5837437 | 3708419.5 | 6422352 | 6276510 | 3525988.25 | 3567616.5 | 3738404 | 4314860.5 | 3298668.75 |
| Q8R424 | Stambp | STAM-binding protein Rattus norvegicus 2 | 57.59 | 4.72 | 48512 | 1 | 1 | 1653348.88 | 1594908.75 | 1296538.12 | 1474115 | 1617054 | 1356715.12 | 1716474.5 | 1727549.88 | 1546419.5 | 1316349.25 | 1095121.12 | 1260645.12 | 1537783.25 | 1475764.25 |
| Q76N33 | Stambpl1 | AMSH-like protease Mus musculus | 56.9 | 4.59 | 49640 | 1 | 1 | 630870.81 | 1153767.12 | 1032593.88 | 1220138.12 | 1061599.25 | 1154897.75 | 1093311.5 | 893314.25 | 981662.62 | 842586.69 | 885317.44 | 1296808.5 | 1099591.75 | 1191983.5 |
| Q9JM90 | Stap1 | Signal-transducing adaptor protein 1 Mus musculus | 637.43 | 59.93 | 34628 | 13 | 13 | 17729511 | 12382445 | 13236752 | 13402185 | 15024507 | 23414919 | 21383787 | 18052759 | 23343591 | 22613869 | 22383089 | 22426727 | 19523953 | 20117637 |
| P42225 | Stat1 | Signal transducer and activator of transcription 1 Mus musculus | 1044.25 | 29.37 | 87197 | 17 | 17 | 27494325 | 28965477 | 27279907 | 21606003 | 25210429 | 16652458 | 15746811 | 15368089 | 18423641 | 16536492 | 14598458 | 14268871 | 18060927 | 15280153 |
| Q9WVL2 | Stat2 | Signal transducer and activator of transcription 2 Mus musculus | 153.72 | 3.47 | 105417 | 3 | 3 | 2279999.25 | 2238939.75 | 2558215.5 | 1865435.12 | 1575645.25 | 1512023 | 247460.8 | 1372053.88 | 1270407.5 | 1310062.38 | 1185951.38 | 1208175.88 | 1289742.88 | 1175394.5 |
| P42227 | Stat3 | Signal transducer and activator of transcription 3 Mus musculus | 483.59 | 22.08 | 88054 | 10 | 10 | 3515281.25 | 2005150.62 | 2500997.5 | 2524098.25 | 2473764.25 | 3163969.25 | 2653809 | 2309740.5 | 3002923.25 | 2453984.5 | 1930244 | 2853838.25 | 2533031.25 | 2413414.25 |
| Q60864 | Stip1 | Stress-induced-phosphoprotein 1 Mus musculus | 1419.29 | 51.93 | 62582 | 25 | 25 | 92142401 | 98960561 | 114734121 | 106186937 | 98563025 | 111416969 | 113462841 | 109652033 | 115267337 | 108921457 | 103799393 | 113045841 | 99960257 | 106205649 |
| O55098 | Stk10 | Serine/threonine-protein kinase 10 Mus musculus | 380.67 | 8.9 | 111906 | 6 | 6 | 4882280.5 | 6873606 | 6064906.5 | 6973655 | 5283605.5 | 6767422 | 6361329 | 6525685 | 5702277 | 6353048.5 | 5533409 | 5755029.5 | 5120246.5 | 5147617.5 |
| Q3TAA7 | Stk11ip | Serine/threonine-protein kinase 11-interacting protein Mus musculus | 104.3 | 2.71 | 118005 | 2 | 2 | 1087222.38 | 435459.94 | 1472558 | 422537.34 | 1048523.12 | 1254704.88 | 393414.59 | 400168.38 | 1172586 | 1153421.5 | 1126184.75 | 1343671.12 | 1363806 | 1132618.5 |

| | | | | | | | | | | | | | | | | | | | | | |
|---|---|---|---|---|---|---|---|---|---|---|---|---|---|---|---|---|---|---|---|---|---|
| Q99KH8 | Stk24 | Serine/threonine-protein kinase 24 Mus musculus | 404.54 | 25.99 | 47954 | 8 | 8 | 3952477.25 | 5645742.5 | 5255120 | 3935718 | 3924994.5 | 8385537 | 8650342 | 5015629 | 8625960 | 7986652 | 5721885.5 | 5286080.5 | 5779808.5 | 7148376.5 |
| Q9JI10 | Stk3 | Serine/threonine-protein kinase 3 Mus musculus | 241.69 | 15.9 | 56855 | 2 | 6 | 11367116 | 10492509 | 11975225 | 7462452 | 8947741 | 10794319 | 10207067 | 11006247 | 10577046 | 8970985 | 9038129 | 8414677 | 10050839 | 10266179 |
| Q9JI11 | Stk4 | Serine/threonine-protein kinase 4 Mus musculus | 300.6 | 17.04 | 55541 | 2 | 6 | 6237303 | 5766168.5 | 6652365 | 5194270 | 5365338 | 6526493 | 5694783 | 5472045 | 5724014 | 5338271 | 5252240.5 | 5382373.5 | 5696308 | 5581898.5 |
| P13668 | Stmn1 | Stathmin Rattus norvegicus | 123.22 | 23.49 | 17288 | 3 | 3 | 2168176.25 | 1101631.5 | 1538180.38 | 3119678.75 | 1734427.75 | 971770.25 | 1579054.88 | 1331267.62 | 465867.5 | 1498069.25 | 2244790 | 1798271.25 | 641686.94 | 1325773.25 |
| P54116 | Stom | Erythrocyte band 7 integral membrane protein Mus musculus | 532.44 | 40.49 | 31375 | 9 | 9 | 100953953 | 109211873 | 100468361 | 87066729 | 103234697 | 107029921 | 112268009 | 115816433 | 107935521 | 103247433 | 97395945 | 93296849 | 105505481 | 105393833 |
| Q99JB2 | Stoml2 | Stomatin-like protein 2, mitochondrial Mus musculus | 587.54 | 39.94 | 38385 | 10 | 10 | 8086672 | 8972740 | 10520236 | 10774133 | 9481619 | 10693588 | 11397595 | 10145533 | 10581765 | 8662388 | 10150286 | 9561723 | 10226316 | 10364525 |
| Q5XIG8 | Strap | Serine-threonine kinase receptor-associated protein Rattus norvegicus | 777.83 | 55.14 | 38456 | 12 | 12 | 19361671 | 18032113 | 19995617 | 13686660 | 16922465 | 17381081 | 19464347 | 20492489 | 19613997 | 19981891 | 17790887 | 19298071 | 20550107 | 20438369 |
| Q8VIM6 | Strc | Stereocilin Mus musculus | 35.29 | 0.44 | 196347 | 1 | 1 | 722819.81 | 2210075.5 | 1922223.88 | 1858499.88 | 719493.31 | 3080604.5 | 3225625.75 | 3418017 | 4935381 | 2952063.25 | 3495577.5 | 2236525 | 3993204 | 3594814.25 |
| Q8C079 | Strip1 | Striatin-interacting protein 1 Mus musculus | 38.35 | 0.84 | 95585 | 1 | 1 | 1 | 475437 | 1 | 1 | 580265.19 | 594375.38 | 490061.5 | 527678.4925 | 522659.56 | 503617.53 | 1 | 580786 | 1 | 598336.75 |
| O55106 | Strn | Striatin Mus musculus | 43.21 | 1.15 | 85966 | 1 | 1 | 783681.75 | 224602.83 | 983225.56 | 865151.06 | 966120.81 | 706670.38 | 724876.44 | 604588.19 | 720222.88 | 845661.88 | 994590.25 | 1131237.12 | 994727.44 | 790095.75 |
| P15589 | Sts | Steryl-sulfatase Rattus norvegicus | 37.5 | 1.21 | 62679 | 1 | 1 | 18672391 | 18864409 | 19582717 | 18834287 | 19137203 | 20631153 | 17560785 | 17414059 | 17574611 | 18616639 | 19207727 | 19609749 | 20383455 | 20274721 |
| P46978 | Stt3a | Dolichyl-diphosphooligosaccharide--protein glycosyltransferase subunit STT3A Mus muscu | 475.62 | 17.87 | 80598 | 11 | 11 | 21405637 | 17041797 | 20985511 | 19263519 | 18418625 | 18276381 | 15234689 | 20423889 | 20170845 | 13724101 | 20855353 | 19487875 | 20094717 | 17650669 |
| Q3TDQ1 | Stt3b | Dolichyl-diphosphooligosaccharide--protein glycosyltransferase subunit STT3B Mus muscu | 160.21 | 4.25 | 93246 | 4 | 4 | 3941781 | 4491079 | 3968801 | 4955797 | 4669015 | 4561818 | 4559948 | 4575558 | 4897722 | 4535791 | 5103285 | 5272529 | 4313962 | 5174484 |
| Q9WUD1 | Stub1 | STIP1 homology and U box-containing protein 1 Mus musculus | 57.9 | 3.95 | 34909 | 1 | 1 | 3146873.188 | 3086832.25 | 2984497.5 | 3650552 | 2865611 | 2090934.5 | 3187515 | 2745167.25 | 2363677.5 | 2170151.25 | 2437110.25 | 2648986.667 | 2685198.5 | 2824651.25 |
| Q9ER00 | Stx12 | Syntaxin-12 Mus musculus | 159.31 | 14.23 | 31195 | 3 | 3 | 5299605 | 5022711 | 5399471.5 | 4769017 | 5899928 | 4988338 | 4629243 | 4599178.5 | 5675749 | 5188153 | 5654708 | 5169556 | 4565737 | 5213805 |
| Q8BVI5 | Stx16 | Syntaxin-16 Mus musculus | 173.78 | 9.82 | 37080 | 2 | 2 | 672622.56 | 793185.31 | 2133389.5 | 2124225.5 | 642382.19 | 685011.81 | 1578329.75 | 1893051.25 | 2315069.5 | 664885.25 | 608115.62 | 975650.94 | 2403423.75 | 2121201.25 |
| Q08849 | Stx3 | Syntaxin-3 Rattus norvegicus | 65.64 | 4.84 | 33257 | 1 | 1 | 1304755.5 | 1256390.12 | 1570297.62 | 1050472.62 | 1043461.81 | 1733287.62 | 1350518.25 | 2013118.88 | 1849019 | 1344744.5 | 1178522.75 | 1319422.38 | 1270346.5 | 1107202.25 |
| Q8K1E0 | Stx5 | Syntaxin-5 Mus musculus | 146.87 | 13.24 | 39713 | 3 | 3 | 1 | 1 | 1 | 1 | 502111.38 | 1 | 1 | 528458.75 | 587626.31 | 1 | 1158666 | 1 | 581148.25 | 1 |
| O70439 | Stx7 | Syntaxin-7 Mus musculus | 225.56 | 22.22 | 29821 | 4 | 4 | 6188589 | 9226391 | 10026963 | 10626933 | 9494574 | 7142655 | 8575288 | 8976337 | 9201994 | 8984403 | 8930549 | 8418296 | 8476526 | 10038357 |
| O88983 | Stx8 | Syntaxin-8 Mus musculus | 152.78 | 11.86 | 26925 | 2 | 2 | 1879497.563 | 1758616.75 | 2002885.88 | 1978806.12 | 1777681.5 | 1351127.75 | 1448355.62 | 1478653.75 | 1335958.88 | 1441302.38 | 1542985.25 | 1589585.083 | 1527788.5 | 1697981.5 |
| O08599 | Stxbp1 | Syntaxin-binding protein 1 Mus musculus | 145.38 | 6.9 | 67569 | 3 | 3 | 2562417.75 | 2881951 | 2218400 | 2610813.75 | 2852319.5 | 2605449 | 2405264.5 | 1061665.62 | 2419341.5 | 1414871.25 | 3385139.5 | 2974812 | 1525077.75 | 2769985 |
| Q64324 | Stxbp2 | Syntaxin-binding protein 2 Mus musculus | 695.14 | 24.28 | 66357 | 11 | 11 | 5440815 | 5271924 | 5078561 | 5142059.5 | 4795616 | 5475848 | 5154398 | 5771421 | 6452571 | 4807936.5 | 4893275 | 5136536.5 | 4284760 | 4601936.5 |
| Q60770 | Stxbp3 | Syntaxin-binding protein 3 Mus musculus | 239.34 | 14.19 | 67942 | 5 | 5 | 2474597 | 3777330.5 | 3009555 | 2729361 | 2934874.75 | 3329068.75 | 2912872.75 | 3400849 | 3368690 | 2763355.75 | 1317611.75 | 1468198 | 3705188 | 3675237 |
| Q5DQR4 | Stxbp5l | Syntaxin-binding protein 5-like Mus musculus | 36.53 | 0.93 | 131844 | 1 | 1 | 5538802.5 | 6822379.5 | 4978563.5 | 5871094 | 7652958.5 | 9780742 | 9647783 | 6495141.5 | 8665200 | 10572685 | 12446183 | 6568418.5 | 11742469 | 10057399 |
| P11031 | Sub1 | Activated RNA polymerase II transcriptional coactivator p15 Mus musculus | 271.01 | 44.88 | 14427 | 6 | 6 | 20297853 | 17595211 | 17189027 | 15575062 | 19474133 | 19151469 | 18454037 | 21394623 | 18318541 | 18620541 | 17129777 | 7266420 | 17331685 | 17083757 |
| Q9Z2I9 | Sucla2 | Succinate--CoA ligase [ADP-forming] subunit beta, mitochondrial Mus musculus | 456.06 | 29.37 | 50114 | 9 | 9 | 8295629.5 | 10424399 | 10797735 | 8882850 | 9556912 | 11354631 | 9859279 | 13265904 | 12864112 | 11100224 | 12045576 | 12539160 | 12071474 | 10814796 |
| Q9WUM5 | Suclg1 | Succinate--CoA ligase [ADP/GDP-forming] subunit alpha, mitochondrial Mus musculus | 308.19 | 23.12 | 36155 | 5 | 5 | 1708247.25 | 1856638 | 2603641.25 | 2963060.5 | 3241120.5 | 4125882.75 | 2771182.25 | 3578916.688 | 2987495.25 | 4431106.5 | 4014572.5 | 2484613 | 4215248 | 3076285.5 |
| Q9Z2I8 | Suclg2 | Succinate--CoA ligase [GDP-forming] subunit beta, mitochondrial Mus musculus | 832.03 | 43.42 | 46840 | 14 | 14 | 46315337 | 40460253 | 48964709 | 41624685 | 43707373 | 55236441 | 51140345 | 52193109 | 55620869 | 46938513 | 52441313 | 56616301 | 50560709 | 52940785 |
| Q9CX34 | Sugt1 | Protein SGT1 homolog Mus musculus | 291.94 | 23.81 | 38159 | 6 | 6 | 6946320 | 7296044.5 | 11091518 | 8172529.5 | 8410025 | 7885454 | 7555362.5 | 8847104 | 9690465 | 8409132 | 8584480 | 8207677 | 7435955 | 9512564 |
| Q8R0F3 | Sumf1 | Formylglycine-generating enzyme Mus musculus | 57.92 | 4.03 | 40659 | 1 | 1 | 810070.88 | 982812.97 | 1167581 | 927408.75 | 1026191.25 | 762364.12 | 1 | 1 | 940496.44 | 905922.88 | 834354.19 | 1048556.56 | 1031610.06 | 1057401.12 |
| P63166 | Sumo1 | Small ubiquitin-related modifier 1 Mus musculus | 33.75 | 6.93 | 11557 | 1 | 1 | 2866870.75 | 2263323 | 2727641.75 | 2733389.5 | 2330636.75 | 2265883.5 | 2557713.25 | 2649103.25 | 2639608.5 | 2676257.75 | 2326051.75 | 2564042.25 | 2390127 | 2113198.75 |
| P61957 | Sumo2 | Small ubiquitin-related modifier 2 Mus musculus | 77.77 | 12.63 | 10871 | 1 | 1 | 17521405 | 2550764.75 | 3024832.5 | 8302931 | 18981269 | 10005600 | 1920606.75 | 1965097.5 | 2617171.25 | 6569732 | 13547958 | 2750152.75 | 3103931.25 | 11562158 |
| Q9D666 | Sun1 | SUN domain-containing protein 1 Mus musculus | 68.36 | 0.99 | 101976 | 1 | 1 | 1243723.62 | 930462.25 | 1211287.453 | 968380.06 | 1702583.88 | 1331332.12 | 939296.88 | 921252.88 | 1042941.565 | 979884.38 | 938427.56 | 1063841.88 | 1305719.25 | 1047457.56 |
| Q8BJS4 | Sun2 | SUN domain-containing protein 2 Mus musculus | 357.59 | 12.72 | 81605 | 7 | 7 | 7134746 | 7171136 | 6667814 | 8281610.5 | 8802811 | 7591476 | 6536819 | 7077459 | 7599344 | 7705661 | 7994056.5 | 6953072 | 6893870.5 | 7618921 |
| Q920B9 | Supt16h | FACT complex subunit SPT16 Mus musculus | 1362.52 | 30.47 | 119825 | 26 | 26 | 35326781 | 36160489 | 38503145 | 33197889 | 36850221 | 36170393 | 34783581 | 35013269 | 33674937 | 36337129 | 35947809 | 38702885 | 37150429 | 37242753 |
| O55201 | Supt5h | Transcription elongation factor SPT5 Mus musculus | 622.98 | 16.82 | 120664 | 13 | 13 | 6384110 | 5187356.5 | 4203389 | 3230445.25 | 5565550 | 3950430.75 | 4026324.5 | 3920013 | 5266818 | 9202531 | 4490831 | 4989519.5 | 3177548.75 | 5566547.5 |
| Q62383 | Supt6h | Transcription elongation factor SPT6 Mus musculus | 38.59 | 0.87 | 199086 | 1 | 1 | 815871 | 938990 | 839073.06 | 844556.06 | 902020 | 729370.62 | 885138.88 | 793959.44 | 730075.56 | 726533.88 | 824633 | 846597.56 | 803620.56 | 857732 |
| Q64310 | Surf4 | Surfeit locus protein 4 Mus musculus | 274.41 | 21.19 | 30381 | 4 | 4 | 9366277 | 4119156.5 | 6308821 | 3941644.25 | 3997260 | 6288492.5 | 4816417.5 | 7257326.5 | 7081237.5 | 4197193 | 3853571.25 | 5723500 | 6882540 | 4755546.5 |
| P70279 | Surf6 | Surfeit locus protein 6 Mus musculus | 80.44 | 4.79 | 41235 | 2 | 2 | 1 | 1 | 371485.03 | 1 | 1 | 1 | 295876.81 | 1 | 1 | 1 | 429008.91 | 322004.53 | 421007.28 | 390673.5733 |
| Q80U70 | Suz12 | Polycomb protein Suz12 Mus musculus | 154.41 | 5.67 | 83026 | 3 | 3 | 1049436 | 949619.69 | 723083.44 | 868012.5 | 936224.38 | 970783 | 1010098.75 | 754355.69 | 650214.38 | 986137.12 | 781760 | 954116.81 | 747686.5 | 1125159.12 |
| Q8K4L3 | Svil | Supervillin Mus musculus | 70.09 | 1.24 | 243162 | 2 | 2 | 903145.62 | 1424224.25 | 1116957.25 | 793862.38 | 1657934.75 | 1749003.75 | 1014828.88 | 1799460.88 | 1132346.38 | 1676395.75 | 927658.81 | 1280121 | 1859833.75 | 1821219.25 |
| Q6A028 | Swap70 | Switch-associated protein 70 Mus musculus | 826.42 | 35.21 | 68996 | 18 | 18 | 8438880 | 4376241.5 | 9912001 | 6593201 | 9192176 | 12313358 | 9991668 | 10447698 | 10027200 | 12633437 | 8687989 | 12469795 | 13438304 | 9246873 |
| Q80X82 | Sympk | Symplekin Mus musculus | 607.63 | 13.55 | 142284 | 12 | 12 | 6440025.5 | 5867272.5 | 7302222 | 5437608 | 5401326.5 | 6865884.5 | 7468577 | 9655819 | 7936289.5 | 6719849.5 | 6671930.5 | 7268310 | 8251929.5 | 6601514 |
| Q7TMK9 | Syncrip | Heterogeneous nuclear ribonucleoprotein Q Mus musculus | 1155.25 | 39.65 | 69633 | 19 | 19 | 40615889 | 36728261 | 36924769 | 44591941 | 42022185 | 37092033 | 36885853 | 41608689 | 38136369 | 39272561 | 37190569 | 37087245 | 39335857 | 35105581 |
| O55100 | Syngr1 | Synaptogyrin-1 Mus musculus | 38.68 | 5.13 | 25653 | 1 | 1 | 2450956.5 | 2020317 | 2084739.25 | 1717727.38 | 2355973 | 2607202.25 | 1876732.62 | 1747543.25 | 2050206.5 | 2416793 | 2187909.5 | 2610802.5 | 3224642 | 2948778.75 |
| O55101 | Syngr2 | Synaptogyrin-2 Mus musculus 2 | 48.79 | 4.46 | 24778 | 1 | 1 | 6716908 | 5932428.5 | 6593905.5 | 6176990 | 6504500.5 | 7131942.5 | 4942594.5 | 4691146.5 | 7286331 | 6242169.5 | 6592178 | 7657609.5 | 7629605.5 | 7683776.5 |
| Q62910 | Synj1 | Synaptojanin-1 Rattus norvegicus | 57.38 | 0.76 | 172881 | 1 | 1 | 650035.81 | 819361.69 | 762494.56 | 645826.94 | 733179.38 | 666011.06 | 539381.56 | 823892.56 | 679032.81 | 422013.09 | 490265.56 | 630727.69 | 570751.31 | 691429.31 |
| O09117 | Sypl1 | Synaptophysin-like protein 1 Mus musculus | 74.56 | 4.21 | 28899 | 2 | 2 | 7626975 | 7761680 | 7000726 | 6510608 | 7570838 | 8136638 | 8299296 | 7413027 | 7333900.5 | 7813165.5 | 6396364 | 7358967 | 6321016.5 | 7721772 |
| Q6Y685 | Tacc1 | Transforming acidic coiled-coil-containing protein 1 Mus musculus | 87.02 | 2.97 | 83952 | 2 | 2 | 1 | 1 | 1281975 | 209819.17 | 230212.3 | 256977.89 | 568102.06 | 1117627.88 | 1028496.5 | 672154.69 | 370429.66 | 1350844.5 | 744044.69 | 808670.56 |
| Q9JJ11 | Tacc3 | Transforming acidic coiled-coil-containing protein 3 Mus musculus | 41.96 | 2.06 | 70627 | 1 | 1 | 570164.88 | 683694.69 | 984348.31 | 634961.25 | 639077.06 | 661778.69 | 672606.94 | 602113 | 722911.69 | 751927.38 | 664764.38 | 624390.62 | 702022.44 | 800288.5 |
| B2RYT9 | Taco1 | Translational activator of cytochrome c oxidase 1 Rattus norvegicus | 72.6 | 6.1 | 32754 | 1 | 1 | 520645.19 | 917651.88 | 1258428.62 | 1122164.12 | 911669.06 | 1174054.62 | 1258741.38 | 1277837.5 | 1142102.12 | 1142220.62 | 1336790.25 | 845858 | 862144.62 | 1227212.38 |
| Q8C092 | Taf5 | Transcription initiation factor TFIID subunit 5 Mus musculus | 48.69 | 1.37 | 87045 | 1 | 1 | 249290.66 | 221653.39 | 291923.91 | 275313.72 | 263304.69 | 278962.09 | 373201.38 | 238473.06 | 287527.44 | 280542.69 | 350749.56 | 247464.59 | 344678.12 | 252350.7 |
| Q91WQ5 | Taf5l | TAF5-like RNA polymerase II p300/CBP-associated factor-associated factor 65 kDa subuni | 43.84 | 3.74 | 65911 | 1 | 1 | 196988.38 | 279299.28 | 218964.33 | 247291.38 | 293913.53 | 1 | 95993.91 | 1 | 190384.42 | 1 | 276832.91 | 184487.11 | 180785.72 | 236676.84 |
| Q9WVA4 | Tagln2 | Transgelin-2 Mus musculus | 1085.59 | 81.41 | 22395 | 15 | 17 | 360666753 | 431408257 | 389466945 | 432111265 | 416786625 | 406705569 | 437845217 | 449314113 | 424956129 | 456099777 | 438363649 | 387668993 | 382331841 | 446616449 |
| Q93092 | Taldo1 | Transaldolase Mus musculus | 1291.53 | 72.4 | 37387 | 24 | 24 | 313788705 | 345215713 | 352452129 | 280869825 | 309088321 | 290711009 | 300426369 | 328447041 | 327133953 | 283559169 | 316289729 | 334889537 | 320368257 | 297152257 |
| Q0VGY8 | Tanc1 | Protein TANC1 Mus musculus | 276.4 | 5.87 | 200805 | 6 | 6 | 1665739.12 | 1572150.38 | 1584758.25 | 1529621.75 | 1553410 | 2260720.75 | 2593082.75 | 2205979 | 2687452.5 | 2599394.75 | 2192618 | 1965477.12 | 1814102.62 | 1908039.88 |
| Q53UA7 | Taok3 | Serine/threonine-protein kinase TAO3 Rattus norvegicus | 226.72 | 5.57 | 105474 | 4 | 4 | 1636289.5 | 1070588.38 | 1438767.5 | 1499944.12 | 1010315 | 1055130.75 | 1202638.62 | 1188569.62 | 1264880.75 | 1346985.75 | 1076981.25 | 1215391.5 | 1193640.5 | 887975 |
| P21958 | Tap1 | Antigen peptide transporter 1 Mus musculus | 423.56 | 17.68 | 78864 | 9 | 9 | 10058543 | 10971870 | 10419274 | 7036405.5 | 7464756 | 5593281 | 5416298 | 6594180 | 6924227 | 6284634 | 4814017 | 3992453.25 | 6695657 | 5874981.5 |
| P36371 | Tap2 | Antigen peptide transporter 2 Mus musculus | 530.08 | 19.52 | 77445 | 9 | 9 | 14567948 | 13917693 | 10892413 | 10225757 | 12761393 | 5557308 | 7911689 | 6487473 | 9713233 | 5826027 | 8362977.5 | 5555959 | 7068710 | 9032469 |
| Q9R233 | Tapbp | Tapasin Mus musculus | 211.01 | 12.9 | 49736 | 4 | 4 | 11257769 | 11536198 | 11120231 | 6555127 | 10087860 | 7521309 | 7555706.5 | 6742979 | 6884508 | 8001356.5 | 8577923 | 8619918 | 8134434.5 | 4916808.5 |
| Q921F2 | Tardbp | TAR DNA-binding protein 43 Mus musculus | 930.72 | 50 | 44548 | 14 | 14 | 23175409 | 22328111 | 23927153 | 27226111 | 27859395 | 29485829 | 27772005 | 22636725 | 30544047 | 33039991 | 29247515 | 27562755 | 31290785 | 29640989 |
| Q9D0R2 | Tars | Threonine--tRNA ligase, cytoplasmic Mus musculus | 1599.79 | 46.12 | 83356 | 28 | 29 | 65506257 | 62482541 | 61105641 | 61218469 | 61199493 | 67305305 | 59347493 | 64304017 | 64449185 | 65464577 | 58892961 | 55011981 | 55924521 | 62468609 |
| Q3UQ84 | Tars2 | Threonine--tRNA ligase, mitochondrial Mus musculus | 118.35 | 3.6 | 81700 | 2 | 2 | 1062753.38 | 1229340 | 1103442.12 | 1291838.75 | 1461157 | 1318593.25 | 1422597 | 1300732.75 | 1191862.25 | 1137297.62 | 1220040.62 | 1287207.5 | 1133859.25 | 1278822.25 |
| Q8R3D1 | Tbc1d13 | TBC1 domain family member 13 Mus musculus | 45.93 | 2.5 | 46453 | 1 | 1 | 724563.75 | 774910.88 | 598706.19 | 716630.38 | 715621.12 | 685328.44 | 876646.81 | 716203.5 | 688941.06 | 661017.94 | 896891.88 | 647943.31 | 613336.5 | 747683.06 |
| Q9CXF4 | Tbc1d15 | TBC1 domain family member 15 Mus musculus | 470.21 | 14.9 | 76527 | 8 | 8 | 11482830 | 9404947 | 13581257 | 11388000 | 11701886 | 12828785 | 10405990 | 12120642 | 11995235 | 11208133 | 13284983 | 14459568 | 11075206 | 13233262 |
| Q8BYH7 | Tbc1d17 | TBC1 domain family member 17 Mus musculus | 69.78 | 1.55 | 72860 | 1 | 1 | 1826476 | 2320990 | 2133825.75 | 2061661.5 | 2286611.5 | 2438345.75 | 1799839.12 | 1785858.62 | 1609931.25 | 1731611 | 2155966.5 | 1916356.75 | 2014059.25 | 2271653.75 |
| Q5SVR0 | Tbc1d9b | TBC1 domain family member 9B Mus musculus | 165.54 | 3.09 | 141779 | 3 | 3 | 1177040.5 | 1080592.25 | 1238921.62 | 1190100.12 | 983897.62 | 1260318.25 | 1323017.12 | 1416112.25 | 1426650 | 1128203.12 | 1369081.75 | 1265815.12 | 1099766.38 | 1158866.75 |
| P48428 | Tbca | Tubulin-specific chaperone A Mus musculus | 224.12 | 56.48 | 12758 | 5 | 5 | 9131711 | 9397210 | 8753151 | 10509868 | 8785775 | 8490261 | 8795788 | 8568958 | 7416747 | 9391328 | 8958221 | 8242874.5 | 7744038 | 8336860 |
| Q9D1E6 | Tbcb | Tubulin-folding cofactor B Mus musculus | 287.78 | 32.79 | 27386 | 5 | 5 | 1688019.12 | 1699028.62 | 2197702.25 | 1859222.75 | 3468018.25 | 3385020 | 2278396.25 | 1568319.5 | 2693830 | 2941233.25 | 2690633.75 | 2839036.5 | 2849970 | 3038751.5 |
| Q8VCN9 | Tbcc | Tubulin-specific chaperone C Mus musculus | 120.93 | 8.8 | 38125 | 2 | 2 | 1930435.62 | 1688610.62 | 2308558.75 | 1972986.88 | 1508172.5 | 1667218.5 | 1372952.38 | 1179736.12 | 3723606.5 | 1387214.88 | 1520596.25 | 484912.41 | 1257391.25 | 1518504 |
| Q8BYA0 | Tbcd | Tubulin-specific chaperone D Mus musculus | 479.79 | 11.12 | 133321 | 10 | 10 | 9827232 | 10397821 | 10799128 | 11129129 | 9635791 | 10576431 | 10834959 | 11296077 | 11370438 | 9349816 | 10144748 | 10099637 | 9596676 | 10616777 |
| Q5FVQ9 | Tbce | Tubulin-specific chaperone E Rattus norvegicus | 38.82 | 2.1 | 59043 | 1 | 1 | 273605.12 | 1 | 202520.45 | 1 | 264504.56 | 223982.59 | 226514.34 | 209166.41 | 186332.39 | 153071.5 | 217987.78 | 189743.8 | 744044.69 | 279906.81 |
| Q5PQJ7 | Tbcel | Tubulin-specific chaperone cofactor E-like protein Rattus norvegicus | 88.23 | 5.9 | 48046 | 1 | 1 | 602050.81 | 885528.81 | 499498.59 | 948888.75 | 915210.38 | 622449.06 | 464868.38 | 728835.25 | 831436.25 | 722504.5 | 779167.38 | 474881.44 | 678815.94 | 477315.59 |
| Q9QXE7 | Tbl1x | F-box-like/WD repeat-containing protein TBL1X Mus musculus | 149.35 | 5.5 | 56802 | 1 | 2 | 2227288 | 1777417.62 | 1661314.12 | 2389203.5 | 1955666.62 | 2360573.5 | 1450900.75 | 986677.69 | 867534.25 | 916350.69 | 2330055 | 3146667 | 1155512 | 1675347 |
| Q8BHJ5 | Tbl1xr1 | F-box-like/WD repeat-containing protein TBL1XR1 Mus musculus | 213.14 | 9.14 | 55661 | 2 | 3 | 14106546 | 16720502 | 16898611 | 15335139 | 1277937 | 3426218.75 | 523779.81 | 17327333 | 499951.09 | 846032.38 | 662916.5 | 1060689.5 | 821011.88 | 665046.19 |
| Q9R099 | Tbl2 | Transducin beta-like protein 2 Mus musculus | 77.91 | 5.88 | 49584 | 2 | 2 | 1479462 | 1627648.5 | 1688083.25 | 696873 | 1092593 | 1377084.25 | 690251.5 | 1695402.38 | 718343.62 | 1559643 | 1450030.75 | 708137.88 | 1604135.62 | 557038.75 |
| Q8C4J7 | Tbl3 | Transducin beta-like protein 3 Mus musculus | 352.15 | 13.61 | 88266 | 8 | 8 | 3707170.75 | 3182216.25 | 3525159.5 | 3772371 | 2670673.75 | 3378390.25 | 3082072.5 | 3377350.25 | 3306403.25 | 3258183.5 | 3121993.5 | 3465155.25 | 3215630 | 2714294.75 |
| Q91YM4 | Tbrg4 | FAST kinase domain-containing protein 4 Mus musculus | 130.32 | 7.62 | 71513 | 3 | 3 | 2510035 | 2642429.75 | 2381086.75 | 1631869.62 | 1641948 | 487617.59 | 480820.78 | 478945.22 | 677948.38 | 621065 | 282645.81 | 527269.88 | 818783.38 | 2638052.75 |
| P10711 | Tcea1 | Transcription elongation factor A protein 1 Mus musculus | 556.38 | 36.21 | 33880 | 2 | 9 | 8782650 | 10094231 | 9063652 | 10786020 | 8662249 | 9711122 | 10123358 | 8473073 | 8282153 | 9658749 | 9308140 | 7182576 | 10041269 | 7651018 |
| Q8CGF7 | Tcerg1 | Transcription elongation regulator 1 Mus musculus | 152.12 | 3.91 | 123788 | 3 | 3 | 4030092.5 | 1967734.62 | 1834887 | 1163685.75 | 743651.25 | 2027085.25 | 1801178.75 | 2872409.75 | 805597.56 | 1774921.5 | 1953045.88 | 1293525.62 | 851390.38 | 1597863.12 |
| O08784 | Tcof1 | Treacle protein Mus musculus | 455.56 | 13.94 | 135001 | 12 | 12 | 5144935.5 | 5856552 | 5091077.5 | 5333352.5 | 4983309.5 | 5809466.5 | 5717061 | 3716768.25 | 4117759.25 | 4305793 | 3941112 | 4467836 | 5531596.5 | 6443511.5 |
| P11983 | Tcp1 | T-complex protein 1 subunit alpha Mus musculus | 1961.24 | 70.14 | 60449 | 28 | 29 | 103142201 | 98834041 | 128821737 | 113434849 | 119067593 | 127784233 | 121612873 | 112257777 | 124960121 | 128860705 | 128124137 | 130652729 | 135114353 | 130183217 |
| Q4G056 | Tdp1 | Tyrosyl-DNA phosphodiesterase 1 Rattus norvegicus | 46.82 | 3.12 | 68632 | 1 | 1 | 1 | 1 | 1 | 1 | 1 | 1 | 1 | 240876.55 | 1 | 1 | 1 | 1 | 376986.69 | 1 |
| Q64232 | Tecr | Very-long-chain enoyl-CoA reductase Rattus norvegicus | 109.64 | 6.82 | 36123 | 2 | 2 | 14777568 | 765068.69 | 857992 | 548268.62 | 1024155.94 | 1196648.75 | 844531.94 | 901943.69 | 778428.56 | 1129739.88 | 1252731.75 | 1215675.62 | 1464300.5 | 1401022.88 |
| Q5EAN7 | Terf2ip | Telomeric repeat-binding factor 2-interacting protein 1 Rattus norvegicus | 69.76 | 3.56 | 43418 | 1 | 1 | 303358.16 | 1 | 368227.56 | 1 | 1 | 425166.81 | 371567.25 | 439172.94 | 368551.03 | 283467.84 | 346011.94 | 387326.56 | 341573.62 | 447683.03 |
| P47226 | Tes | Testin Mus musculus | 353.71 | 21.04 | 47983 | 8 | 8 | 3696221.75 | 4564285.5 | 4730642 | 6025362.5 | 4329370.5 | 7849374 | 5757800 | 5079002 | 6395534 | 6599892.5 | 6140136 | 5024831.5 | 5679737.5 | 6283765.5 |
| Q3URQ0 | Tex10 | Testis-expressed protein 10 Mus musculus | 40.31 | 1.4 | 105209 | 1 | 1 | 482972.75 | 307244.69 | 168606.17 | 225089.27 | 284859.88 | 203006.31 | 202083.89 | 300447.34 | 211472.84 | 227380.52 | 286050.22 | 280742.81 | 251367.86 | 260135.94 |
| Q921I1 | Tf | Serotransferrin Mus musculus | 40.02 | 2.01 | 76724 | 1 | 1 | 307375.94 | 237959.55 | 224392.09 | 422393.12 | 246643.97 | 385051.09 | 329388.19 | 321554.75 | 441268.34 | 347895.31 | 504680.78 | 469763.22 | 356330.72 | 425325.19 |
| Q811P6 | Tfb1m | Dimethyladenosine transferase 1, mitochondrial Rattus norvegicus | 66.78 | 5.51 | 39181 | 2 | 2 | 800350.69 | 1495634.25 | 1176110.5 | 1251058.25 | 1301972.25 | 1261572.12 | 1249497.88 | 1306348 | 510187.81 | 1079245.25 | 1192485.5 | 1464513.5 | 1222191 | 577404.94 |

| | | | | | | | | | | | | | | | | | | | | | |
|---|---|---|---|---|---|---|---|---|---|---|---|---|---|---|---|---|---|---|---|---|---|
| Q9ERA6 | Tfip11 | Tuftelin-interacting protein 11 Mus musculus | 150.74 | 5.49 | 96305 | 3 | 3 | 272489.78 | 336598.19 | 352474.6175 | 357605.84 | 443204.66 | 360103.62 | 351997.09 | 361118.225 | 296120.47 | 436251.72 | 331124 | 206264.83 | 1 | 1 |
| Q62351 | Tfrc | Transferrin receptor protein 1 Mus musculus | 939.45 | 30.54 | 85731 | 20 | 20 | 11530963 | 11633155 | 10900484 | 7395272.5 | 10115463 | 7675129.5 | 5969507 | 7726780.5 | 7213647.5 | 5669373 | 9960069 | 11926839 | 10309869 | 10780530 |
| Q91YX0 | Themis2 | Protein THEMIS2 Mus musculus | 451.31 | 22.32 | 74378 | 9 | 9 | 6519084.5 | 7443011.5 | 6576726.5 | 7534454 | 7404014.5 | 3663874.5 | 4519365 | 4221583.5 | 4518707.5 | 4335175.5 | 5111128.5 | 5144754.5 | 5504107 | 4491972 |
| P59924 | Thoc1 | THO complex subunit 1 Rattus norvegicus | 36.64 | 2.62 | 40243 | 1 | 1 | 1837730 | 1421757.88 | 2604596.75 | 1773867.813 | 1231386.62 | 2069443.25 | 1687968 | 2301756.5 | 1998416.62 | 1225558.38 | 1446609.38 | 1991398.62 | 2090517.12 | 1796324.88 |
| B1AZI6 | Thoc2 | THO complex subunit 2 Mus musculus | 570.07 | 9.28 | 182773 | 12 | 12 | 8149270.5 | 7828305.5 | 9587872 | 8968936 | 9977361 | 8857419 | 15065211 | 9162973 | 9186955 | 9004481 | 8558104 | 8619074 | 9683449 | 9867468 |
| Q8VE80 | Thoc3 | THO complex subunit 3 Mus musculus | 45.07 | 6.55 | 38738 | 1 | 1 | 210528.8425 | 270131.34 | 162510.39 | 237879.44 | 171594.2 | 171699.59 | 154348.25 | 190092.56 | 200224.78 | 181062.11 | 118958.43 | 195646.2 | 203494.3 | 236717.64 |
| Q68FX7 | Thoc5 | THO complex subunit 5 homolog Rattus norvegicus | 48.69 | 1.03 | 78660 | 1 | 1 | 1197032.38 | 946366.56 | 1253918.25 | 743662.75 | 765300.56 | 1131307.38 | 885328.75 | 1230646 | 890794.88 | 825182.62 | 1029854 | 938315.19 | 1004921.19 | 762567.81 |
| Q5U4D9 | Thoc6 | THO complex subunit 6 homolog Mus musculus | 114.37 | 10.56 | 37315 | 2 | 2 | 1086235.25 | 781938 | 1699729.5 | 1547681.75 | 1151054.75 | 1061377.25 | 985322.06 | 1053412.75 | 1182171.88 | 1170923.5 | 1034302.62 | 1174540.5 | 852664.75 | 978240.62 |
| Q8C1A5 | Thop1 | Thimet oligopeptidase Mus musculus | 366.94 | 13.39 | 78026 | 9 | 9 | 11717284 | 10877380 | 10613722 | 12053381 | 12043315 | 10434605 | 10446616 | 10857849 | 11335751 | 11536244 | 11920502 | 10755021 | 11946237 | 10644867 |
| Q569Z6 | Thrap3 | Thyroid hormone receptor-associated protein 3 Mus musculus | 482.4 | 13.67 | 108178 | 10 | 10 | 3810779 | 3779500.5 | 3694478.75 | 2688826.75 | 3653554 | 2720960.75 | 4338419.5 | 2268214.25 | 3286829.25 | 4624024 | 3170618.5 | 2689086 | 2953527 | 4092733.25 |
| Q99J36 | Thumpd1 | THUMP domain-containing protein 1 Mus musculus | 246.32 | 23.14 | 38885 | 6 | 6 | 5087066 | 3713403.5 | 4923494 | 3482210.5 | 4537704 | 4350388.5 | 4919905.5 | 3365397.5 | 4279763 | 3352929.75 | 3750889.5 | 4467125.5 | 4256782 | 5406794.5 |
| P97770 | Thumpd3 | THUMP domain-containing protein 3 Mus musculus | 227.58 | 11.49 | 56431 | 5 | 5 | 899067.62 | 1041571.62 | 1029383.31 | 1288757.88 | 1037148.62 | 1246746.5 | 979046.88 | 965987.75 | 1035807.88 | 1869101.88 | 1648342.25 | 1458954.5 | 1544222.62 | 1375195.5 |
| P52912 | Tia1 | Nucleolysin TIA-1 Mus musculus | 51.35 | 4.66 | 42800 | 1 | 1 | 329928.09 | 292098.72 | 249242.75 | 298583 | 292463.14 | 222929.34 | 282399.94 | 241197.3 | 275345.5 | 184114.42 | 279784.69 | 291728.4267 | 323873.34 | 271527.25 |
| P70318 | Tial1 | Nucleolysin TIAR Mus musculus | 228.31 | 11.99 | 43389 | 4 | 4 | 6755246 | 3901110.25 | 2767010.5 | 3009831 | 4636751 | 4710984.5 | 4218899 | 3354219 | 4068834.75 | 2693398.5 | 3958384 | 3139595 | 4073087 | 4137481 |
| Q793I8 | Tifa | TRAF-interacting protein with FHA domain-containing protein A Mus musculus | 39.76 | 4.89 | 21560 | 1 | 1 | 559648.69 | 481834.75 | 630341.81 | 799239.31 | 653510.75 | 701851.5 | 722485.25 | 832444.56 | 781586.81 | 768169.94 | 567269.69 | 662303.12 | 628207.12 | 722701.25 |
| Q8BZA9 | Tigar | Fructose-2,6-bisphosphatase TIGAR Mus musculus | 247.69 | 34.2 | 29191 | 5 | 5 | 3817089.5 | 4261817 | 4053701 | 3545411.5 | 4825893.5 | 4302078 | 3274367.75 | 3891345.5 | 3579051.5 | 3320066.25 | 3936232.75 | 4669935 | 4493222 | 3625316.5 |
| P62075 | Timm13 | Mitochondrial import inner membrane translocase subunit Tim13 Mus musculus | 36.2 | 14.74 | 10458 | 1 | 1 | 2119784.5 | 2271527.25 | 2447546.5 | 2781436.25 | 2072312.75 | 1404924 | 1992912 | 1928502.38 | 1382621 | 1689689.62 | 2156696.75 | 1935577 | 1747974.12 | 1923287.5 |
| O35092 | Timm17a | Mitochondrial import inner membrane translocase subunit Tim17-A Rattus norvegicus | 225.05 | 43.27 | 18038 | 3 | 3 | 1488920.75 | 1750135.12 | 1635881.25 | 5872844.5 | 1660234.25 | 1607643.62 | 1759371.25 | 1936693 | 1874907.12 | 1560411.5 | 2040129.25 | 2185548.5 | 1997192.38 | 2209612.75 |
| Q9Z0V7 | Timm17b | Mitochondrial import inner membrane translocase subunit Tim17-B Mus musculus | 97.86 | 29.65 | 18352 | 2 | 2 | 571825.06 | 364785.34 | 1 | 1 | 1 | 1 | 1 | 1 | 1 | 1 | 1 | 4235789.5 | 1 | 1767531 |
| Q9WTQ8 | Timm23 | Mitochondrial import inner membrane translocase subunit Tim23 Mus musculus | 208.48 | 37.32 | 21978 | 3 | 3 | 1655220 | 2294022.75 | 2154669.5 | 2537059 | 2370898.5 | 2291069 | 2282806.5 | 2724911 | 2469080.75 | 1782882 | 2302044.75 | 2499509 | 2785956.25 | 766804.75 |
| O35857 | Timm44 | Mitochondrial import inner membrane translocase subunit TIM44 Mus musculus | 644.41 | 25.44 | 51091 | 11 | 11 | 11188471 | 12907923 | 13961999 | 10798965 | 11944740 | 12411624 | 11655814 | 12557621 | 13614326 | 13585994 | 13574402 | 13336830 | 13520685 | 13669614 |
| Q9D880 | Timm50 | Mitochondrial import inner membrane translocase subunit TIM50 Mus musculus | 500.41 | 28.33 | 39776 | 8 | 8 | 21850333 | 21642433 | 23470217 | 21432963 | 19927959 | 21913043 | 22113833 | 22830365 | 21629803 | 22395937 | 21896041 | 22804215 | 24948729 | 21864987 |
| A2VCX1 | Tiprl | TIP41-like protein Rattus norvegicus | 88.43 | 6.64 | 31225 | 2 | 2 | 1010077.5 | 1185112.38 | 1003873.19 | 1027271.5 | 951998.25 | 1107547.12 | 979274.44 | 1240700.25 | 1171214.88 | 813651.25 | 710119.75 | 1019627.19 | 1110389.62 | 1043947.75 |
| Q9Z0U1 | Tjp2 | Tight junction protein ZO-2 Mus musculus | 551.53 | 11.14 | 131280 | 10 | 10 | 4912769 | 5892577 | 5076197 | 6428088.5 | 5285090 | 5611082 | 6050415 | 5094482 | 5915032.5 | 5492947 | 6132314.5 | 5874466.5 | 6680318.5 | 6155205.5 |
| P04184 | Tk1 | Thymidine kinase, cytosolic Mus musculus | 211.99 | 19.74 | 25776 | 4 | 4 | 1585144.5 | 1742422.25 | 1060971.75 | 2080591.25 | 3288614.25 | 928598.25 | 1062107.62 | 1380058 | 1343685 | 959315.31 | 2107370 | 1739533.5 | 1893525.75 | 1032170.88 |
| P40142 | Tkt | Transketolase Mus musculus | 2163.61 | 67.74 | 67630 | 33 | 34 | 278338977 | 278360673 | 292927297 | 297235425 | 310469057 | 319966209 | 285376705 | 298199585 | 327872737 | 315072865 | 307718305 | 325894145 | 323921857 | 314851041 |
| P26039 | Tln1 | Talin-1 Mus musculus | 7638.56 | 62.42 | 269821 | 106 | 106 | 416044193 | 454960129 | 469737793 | 454685089 | 458625025 | 433816129 | 470248225 | 461345057 | 457277825 | 481420321 | 452208289 | 429256769 | 429695233 | 467948225 |
| Q6R5N8 | Tlr13 | Toll-like receptor 13 Mus musculus | 647.17 | 13.62 | 114443 | 9 | 9 | 12779083 | 9427389 | 12441244 | 11319293 | 11369952 | 8719041 | 8643784 | 9305679 | 9360403 | 6470506.5 | 8340983.5 | 8267912 | 11075327 | 7969123 |
| Q9QUN7 | Tlr2 | Toll-like receptor 2 Mus musculus | 571.43 | 19.01 | 89449 | 10 | 10 | 4176299.25 | 2433391 | 3841125 | 3986160.25 | 4094781.25 | 3248038.75 | 3052094.5 | 2773228.5 | 2493626.5 | 2745768 | 2980772.25 | 1918254.38 | 2599399 | 2190405.25 |
| Q99MB1 | Tlr3 | Toll-like receptor 3 Mus musculus | 130.23 | 4.64 | 103671 | 3 | 3 | 1 | 289401.53 | 257530.61 | 1 | 1 | 1 | 1 | 631899.69 | 291006.75 | 1 | 1 | 1 | 214161.78 | 1 |
| P58681 | Tlr7 | Toll-like receptor 7 Mus musculus | 670.44 | 16.38 | 121837 | 14 | 14 | 9379617 | 8604142 | 9166527 | 10330100 | 8842673 | 8929297 | 8443729 | 9944700 | 6979926.5 | 7646626 | 8494588 | 8120686.5 | 6905860.5 | 7610974.5 |
| P58021 | Tm9sf2 | Transmembrane 9 superfamily member 2 Mus musculus | 278.51 | 9.37 | 75330 | 5 | 5 | 13855640 | 12156082 | 12090649 | 9590521 | 11324615 | 13090470 | 10758483 | 12456675 | 11898104 | 9394779 | 10426590 | 11520649 | 13385623 | 10224459 |
| Q9ET30 | Tm9sf3 | Transmembrane 9 superfamily member 3 Mus musculus | 260.78 | 14.65 | 67545 | 6 | 6 | 7030180 | 6193510.5 | 4196395.5 | 4461002.5 | 7014155 | 5821991.5 | 8225485.5 | 5519439 | 6012270 | 8964196 | 6465938 | 4715021 | 9162519 | 6460609.5 |
| Q9CR02 | Tma16 | Translation machinery-associated protein 16 Mus musculus | 97.78 | 17.19 | 25782 | 2 | 2 | 2082810.12 | 1679686.38 | 1689868.88 | 2088082.88 | 2318952.25 | 163200 | 1099331.66 | 1320676.88 | 1354477.38 | 1558972.38 | 2616166 | 2732925.75 | 1900565.603 | 2198541.75 |
| Q7TQ69 | Tmc3 | Transmembrane channel-like protein 3 Mus musculus | 36.33 | 0.62 | 128691 | 1 | 1 | 1 | 236465.83 | 695763.5 | 361556 | 1 | 535011.38 | 478055.29 | 275227.75 | 497117.22 | 604864.81 | 615009.06 | 719421.81 | 1 | 1 |
| Q5I0H4 | Tmco1 | Calcium load-activated calcium channel Rattus norvegicus | 180.96 | 19.68 | 21175 | 3 | 3 | 3399643.25 | 2650346.5 | 3606003 | 3552383 | 5420269 | 3912021 | 3425907 | 3395506.5 | 3115239.75 | 6329713 | 4478566.5 | 4806150.5 | 6186867 | 5179596 |
| Q63584 | Tmed10 | Transmembrane emp24 domain-containing protein 10 Rattus norvegicus | 529.49 | 31.96 | 24858 | 6 | 6 | 40057721 | 39002757 | 39721865 | 38627953 | 39894673 | 46204029 | 35342013 | 43571181 | 38375237 | 39571245 | 38750605 | 34929749 | 39570569 | 43963921 |
| Q63524 | Tmed2 | Transmembrane emp24 domain-containing protein 2 Rattus norvegicus | 189.89 | 16.92 | 22733 | 3 | 3 | 3422113.75 | 3281659.5 | 3057820 | 2860088 | 4130505 | 3648242 | 3384196.5 | 3421078.25 | 4020099.75 | 4893308.5 | 3510925.5 | 2661094.25 | 3156207.5 | 3850377 |
| Q8R1V4 | Tmed4 | Transmembrane emp24 domain-containing protein 4 Mus musculus | 140.56 | 16.3 | 26022 | 2 | 3 | 2765311.5 | 5435591 | 6167950 | 5655069.5 | 2494016.5 | 2559196 | 2634765.5 | 4011368 | 2298675.5 | 4342469 | 6146217 | 2616833 | 3125061 | 2620967.5 |
| Q6AXN3 | Tmed5 | Transmembrane emp24 domain-containing protein 5 Rattus norvegicus | 163.24 | 10.04 | 26136 | 2 | 3 | 1766584.25 | 3235119.75 | 1965206.5 | 1636073.38 | 1891252 | 2521843.75 | 1757006.5 | 4477422.5 | 2644917.75 | 2237753.5 | 1914838.12 | 2585536.5 | 3115258.5 | 1476065.5 |
| D3ZTX0 | Tmed7 | Transmembrane emp24 domain-containing protein 7 Rattus norvegicus | 219.34 | 24.34 | 25479 | 4 | 4 | 3980779.5 | 4075977 | 4367660 | 3285284.25 | 4314586 | 4954215 | 3667313.5 | 5599888.5 | 5447866.5 | 3261455.75 | 3928301 | 4113767 | 3495878.75 | 4326724 |
| Q5I0E7 | Tmed9 | Transmembrane emp24 domain-containing protein 9 Rattus norvegicus | 225.67 | 20 | 27028 | 4 | 5 | 27894041 | 27263373 | 25066293 | 24434101 | 24617893 | 25373113 | 24290801 | 27665369 | 22929421 | 22618409 | 25063123 | 20497319 | 23675195 | 23094175 |
| Q8VC04 | Tmem106a | Transmembrane protein 106A Mus musculus | 138.31 | 14.56 | 29109 | 3 | 3 | 8038553.5 | 10927428 | 7606400.5 | 10944657 | 11518702 | 10045150 | 9934427 | 6263983 | 6128397 | 9194521 | 9925698 | 5595022.5 | 9895054 | 9578600 |
| Q6AYA5 | Tmem106b | Transmembrane protein 106B Rattus norvegicus | 52.41 | 5.82 | 31152 | 1 | 1 | 1 | 808104.06 | 1153426 | 1 | 1 | 1358662.375 | 1316674 | 1427608.5 | 1189302 | 1501065 | 1351118.75 | 1330084.5 | 1385718 | 1374090.88 |
| Q3UBX0 | Tmem109 | Transmembrane protein 109 Mus musculus | 208.92 | 11.93 | 26306 | 3 | 3 | 11520050 | 8938495 | 7579169.5 | 7653542 | 8806654 | 9032588 | 8605221 | 9219168 | 8722821 | 8238002.5 | 9627412 | 11004770 | 8653037 | 9869750 |
| B0BN86 | Tmem11 | Transmembrane protein 11, mitochondrial Rattus norvegicus 2 | 39.37 | 4.21 | 21342 | 1 | 1 | 1 | 841602.56 | 1191889.12 | 755534.25 | 881622.19 | 1281214.62 | 963581.31 | 1102635.25 | 1525328.38 | 1095606.12 | 990600.75 | 1291257 | 1248161.75 | 1078966.75 |
| Q9D1R1 | Tmem126b | Complex I assembly factor TMEM126B, mitochondrial Mus musculus | 48.71 | 7.83 | 25444 | 1 | 1 | 452009.94 | 505386.88 | 705776.31 | 524030.79 | 432950.03 | 416505.16 | 262116.75 | 610044.75 | 629120.12 | 441107.56 | 366651.41 | 419497.25 | 489247.59 | 342182.09 |
| Q9CQN6 | Tmem14c | Transmembrane protein 14C Mus musculus | 340.79 | 92.11 | 11642 | 5 | 5 | 8133633 | 8748256 | 8877349 | 8262232 | 6547783.5 | 8742546 | 9345987 | 10589470 | 9507474 | 7645523 | 8404493 | 9418079 | 9691133 | 9599866 |
| Q9D938 | Tmem160 | Transmembrane protein 160 Mus musculus | 45.37 | 3.72 | 19587 | 1 | 1 | 1820692.62 | 2003869.12 | 1692980 | 1684022.88 | 1704514.62 | 1746157.12 | 1339688.88 | 1417623.5 | 1588838.75 | 1599083.5 | 1784629.88 | 1587043.75 | 1653014.75 | 1519378.5 |
| P52875 | Tmem165 | Transmembrane protein 165 Mus musculus | 128.09 | 13.31 | 34791 | 2 | 2 | 1156994.62 | 1776877 | 2666680.5 | 1616710.88 | 1752277.88 | 1845163.75 | 1377738.88 | 2226832 | 2154705 | 2013992.25 | 1652382.5 | 1733552 | 2220621.75 | 1476207.62 |
| Q9CR64 | Tmem167a | Protein kish-A Mus musculus | 45.37 | 12.5 | 8074 | 1 | 1 | 1144087.38 | 278626.41 | 697241.62 | 661018.47 | 524118.47 | 455784.38 | 318931.19 | 441181.4875 | 465530.69 | 524479.69 | 374078.38 | 497507.72 | 903078.56 | 661856.56 |
| Q5PQM0 | Tmem168 | Transmembrane protein 168 Rattus norvegicus | 168.83 | 7.32 | 79772 | 4 | 4 | 2888776 | 435973.72 | 1 | 1 | 387919.75 | 1196557.5 | 1539182.25 | 1877630 | 1532507.62 | 1536469.343 | 662427.81 | 1682127.88 | 1318304.12 | 1220953.27 |
| Q3TBT3 | Tmem173 | Stimulator of interferon genes protein Mus musculus | 106.37 | 11.9 | 42830 | 3 | 3 | 3873826.25 | 2567173 | 2623640.25 | 2622259.75 | 2671752.25 | 1795765 | 1862165.12 | 1654236.62 | 2591512.5 | 2079581 | 1997885.5 | 2156260.5 | 2350312 | 2926366 |
| Q9R1Q6 | Tmem176b | Transmembrane protein 176B Mus musculus | 199.52 | 10.65 | 28367 | 4 | 4 | 2148956.75 | 2091327.5 | 2594122 | 1331876.12 | 2838658 | 3657955 | 2483858 | 2656355.75 | 2613483 | 2044621.75 | 3258462.25 | 1229810.75 | 3253721.5 | 2385391 |
| Q9D771 | Tmem206 | Transmembrane protein 206 Mus musculus | 190.62 | 17.43 | 40178 | 4 | 4 | 224886.72 | 895148.5 | 1072906.62 | 520046.97 | 596969.31 | 527287.5 | 772867.44 | 807784.12 | 1017649 | 593458.44 | 557030.31 | 632324.69 | 1185840.5 | 607215.75 |
| A1L1L2 | Tmem214 | Transmembrane protein 214 Rattus norvegicus | 58.63 | 1.9 | 76566 | 1 | 1 | 145575.83 | 1 | 166379.84 | 1 | 1 | 228695.64 | 229969.2475 | 338946 | 230705.56 | 121529.79 | 134574.75 | 194711.33 | 195662.78 | 184779.02 |
| Q8VEK0 | Tmem30a | Cell cycle control protein 50A Mus musculus | 42.17 | 4.67 | 41061 | 1 | 1 | 1 | 811879.38 | 1072021.5 | 1 | 523757.34 | 749024.44 | 653259.69 | 926331.12 | 939684.5 | 605821.5 | 561706.94 | 698800.81 | 754504.25 | 683629.88 |
| Q9CR67 | Tmem33 | Transmembrane protein 33 Mus musculus | 222.6 | 19.84 | 28031 | 5 | 5 | 13066707 | 10189787 | 9696099 | 9721362 | 11396923 | 11334893 | 10498830 | 10047239 | 10408967 | 8952559 | 12634705 | 12262248 | 12177312 | 9577509 |
| Q9DAV9 | Tmem38b | Trimeric intracellular cation channel type B Mus musculus | 100.84 | 9.93 | 32640 | 2 | 2 | 2071083.38 | 1757264.38 | 2027588.38 | 1166918 | 1924872.12 | 2013819.25 | 2085608.5 | 2084700.12 | 2149906.75 | 1926007.88 | 1791344.75 | 2386318.25 | 2411892.75 | 2121528.25 |
| Q9DBS1 | Tmem43 | Transmembrane protein 43 Mus musculus | 609.13 | 33.5 | 44783 | 11 | 11 | 9646623 | 8312537 | 7369328.5 | 7296842.5 | 10582217 | 8148528 | 9591098 | 5016997 | 8528342 | 11195031 | 11221543 | 9097162 | 11322794 | 10811057 |
| Q4VAE3 | Tmem65 | Transmembrane protein 65 Mus musculus | 36.81 | 4.7 | 24918 | 1 | 1 | 605327.5 | 993269 | 657702.12 | 560369.31 | 616304.94 | 1003408.75 | 721794.19 | 728543.06 | 641973.31 | 655633.31 | 698864.19 | 726299 | 817662.88 | 893758.19 |
| B9EKI3 | Tmf1 | TATA element modulatory factor Mus musculus | 49.99 | 1.37 | 121803 | 1 | 1 | 971443.31 | 1054957.62 | 913891.81 | 1043313.94 | 967069.44 | 1310620.88 | 961955.31 | 1104701.88 | 1140436.38 | 922498.94 | 929209.62 | 855882.75 | 735158.38 | 913820.69 |
| P49813 | Tmod1 | Tropomodulin-1 Mus musculus | 76.86 | 7.52 | 40466 | 2 | 2 | 1126163.25 | 1262612.5 | 1341541.12 | 1705706.75 | 1437078.62 | 1610057.88 | 1377572.5 | 1692447.88 | 1611344.12 | 1576615.88 | 1877508 | 1984288.62 | 1590483.12 | 2002115.38 |
| Q9JHJ0 | Tmod3 | Tropomodulin-3 Mus musculus | 473.86 | 28.41 | 39503 | 8 | 8 | 9038000 | 9776202 | 8172756 | 9641553 | 9345298 | 8136427.5 | 11804543 | 10339673 | 10490787 | 8826404 | 9651977 | 9044906 | 8308225 | 8624543 |
| Q61033 | Tmpo | Lamina-associated polypeptide 2, isoforms alpha/zeta Mus musculus | 726.2 | 27.85 | 75168 | 6 | 12 | 10506329 | 10680562 | 11460652 | 12180816 | 10627226 | 11433041 | 9259722 | 9149543 | 9467989 | 12519884 | 8628876 | 10067712 | 9138476 | 9306061 |
| Q61029 | Tmpo | Lamina-associated polypeptide 2, isoforms beta/delta/epsilon/gamma Mus musculus | 870.19 | 43.58 | 50373 | 8 | 14 | 22853437 | 25097985 | 24252831 | 27683307 | 23396299 | 20453555 | 22798725 | 22609409 | 21745565 | 28616523 | 20907375 | 21273769 | 25675427 | 25066241 |
| Q8VBT0 | Tmx1 | Thioredoxin-related transmembrane protein 1 Mus musculus | 230.44 | 17.27 | 31396 | 5 | 5 | 12106276 | 12540248 | 11158372 | 10139955 | 10436282 | 12448879 | 12212757 | 14560317 | 12530816 | 12165171 | 12336584 | 12729205 | 12539227 | 12920627 |
| Q5XIK2 | Tmx2 | Thioredoxin-related transmembrane protein 2 Rattus norvegicus | 67.31 | 5.08 | 33866 | 1 | 1 | 290727.25 | 284959.56 | 230181.39 | 201141.98 | 298347.53 | 196443.27 | 196502.42 | 209914.42 | 166982.14 | 147189.75 | 226062.47 | 360765.41 | 215834.38 | 155584.45 |
| Q8BXZ1 | Tmx3 | Protein disulfide-isomerase TMX3 Mus musculus | 404.66 | 19.3 | 51848 | 6 | 6 | 5363058 | 5698235 | 5681977.5 | 6452302.5 | 5437719.5 | 5301959 | 6278907 | 8033944 | 5775556.5 | 6279803 | 5183127.5 | 5782136.5 | 5560681 | 6103899 |
| Q8C0L0 | Tmx4 | Thioredoxin-related transmembrane protein 4 Mus musculus | 133.29 | 11.34 | 37131 | 3 | 3 | 2123360.25 | 2472592.25 | 2666886.75 | 2784210 | 2305559.25 | 2969005 | 2072241.75 | 3161273 | 3395063.25 | 1865315.75 | 2643053.75 | 2874262.75 | 3026597.75 | 3153503 |
| Q61333 | Tnfaip2 | Tumor necrosis factor alpha-induced protein 2 Mus musculus | 61.25 | 1.74 | 78102 | 1 | 1 | 404323.91 | 502148.03 | 584102.56 | 275638.88 | 300789.28 | 580202.62 | 414292.94 | 617727.88 | 425619.16 | 412455.81 | 706366.38 | 1122818.25 | 686408.75 | 546814.75 |
| Q921Z5 | Tnfaip8 | Tumor necrosis factor alpha-induced protein 8 Mus musculus | 48.55 | 10.1 | 22960 | 1 | 1 | 1 | 1 | 200955.91 | 1 | 1 | 1 | 1 | 204444.11 | 275461.44 | 193460.98 | 1 | 263590.09 | 1 | 1 |
| Q9D8Y7 | Tnfaip8l2 | Tumor necrosis factor alpha-induced protein 8-like protein 2 Mus musculus | 111.73 | 26.09 | 20615 | 3 | 3 | 474344.91 | 1228828.75 | 325598.09 | 1 | 1 | 965599.31 | 1 | 358868.06 | 925335.31 | 1 | 1 | 1254878.25 | 467649.66 | 1 |
| Q8BFY9 | Tnpo1 | Transportin-1 Mus musculus | 916.78 | 28.73 | 102357 | 13 | 18 | 39307329 | 40928837 | 39648933 | 39648505 | 41273549 | 43799013 | 40104089 | 44907905 | 42044061 | 39123097 | 44049557 | 43053501 | 44068709 | 44637841 |
| Q99LG2 | Tnpo2 | Transportin-2 Mus musculus | 428.93 | 10.6 | 100456 | 3 | 8 | 18682063 | 19357955 | 19835205 | 20231513 | 20493809 | 20322747 | 20159983 | 19982701 | 20093897 | 20062357 | 21680597 | 19667539 | 20477153 | 23304395 |
| Q6P2B1 | Tnpo3 | Transportin-3 Mus musculus | 427.89 | 17.12 | 104170 | 10 | 10 | 9316738 | 6364965.5 | 9056856 | 8481058 | 9776535 | 10390683 | 8686003 | 6371390 | 9599014 | 6789632 | 8760732 | 6872916 | 7344087 | 6312034 |
| Q5SSZ5 | Tns3 | Tensin-3 Mus musculus | 44.72 | 0.97 | 155589 | 1 | 1 | 596497.81 | 404810.47 | 723231.25 | 484870.19 | 789873 | 1 | 1 | 772518.88 | 1 | 753306.56 | 1 | 664874.69 | 924459.94 | 1 |
| Q9D2E2 | Toe1 | Target of EGR1 protein 1 Mus musculus | 113.89 | 6.26 | 56871 | 3 | 3 | 289474.69 | 192869.33 | 215007.33 | 164737.58 | 1 | 979996.5 | 625284.12 | 121787.96 | 550880 | 538430.31 | 84002.27 | 472074.47 | 120107.38 | 225394.7067 |
| A2RUW1 | Tollip | Toll-interacting protein Rattus norvegicus | 127.83 | 8.39 | 30315 | 2 | 2 | 1371221.5 | 351292.06 | 1948126.5 | 483997.88 | 1203978.62 | 1418644.88 | 1297829 | 980173.25 | 1276369.75 | 1396201.5 | 1196023.75 | 867223.94 | 1584949.38 | 525430.62 |
| O88746 | Tom1 | Target of Myb protein 1 Mus musculus | 362.27 | 20.12 | 54325 | 6 | 6 | 5696313 | 5496928 | 5123093 | 6027098.5 | 5160333.5 | 5821696 | 5673804.5 | 5196539 | 4892461 | 5273377 | 4739190 | 6352727.5 | 5522103.5 | 5131028 |
| Q62760 | Tomm20 | Mitochondrial import receptor subunit TOM20 homolog Rattus norvegicus | 122.93 | 44.14 | 16284 | 3 | 3 | 288771.03 | 299915.03 | 654164.56 | 291334.91 | 182377.92 | 288552.78 | 224265.53 | 659523.06 | 721602.31 | 300250.47 | 291376.31 | 540107.44 | 436521.69 | 194766.66 |
| Q9CPQ3 | Tomm22 | Mitochondrial import receptor subunit TOM22 homolog Mus musculus | 247.17 | 42.25 | 15537 | 4 | 4 | 7626078 | 8556125 | 8900279 | 8178021 | 8377904.5 | 11178517 | 8774223 | 9839948 | 10990508 | 7490971 | 8882940 | 10574362 | 9146619 | 9753334 |
| Q9CYG7 | Tomm34 | Mitochondrial import receptor subunit TOM34 Mus musculus | 659.61 | 45.95 | 34278 | 11 | 11 | 12720540 | 14699377 | 15130840 | 12742153 | 14684081 | 16434287 | 16587682 | 16365119 | 18186055 | 16624583 | 16003520 | 16805789 | 17180667 | 17118991 |
| Q9QYA2 | Tomm40 | Mitochondrial import receptor subunit TOM40 homolog Mus musculus | 481.47 | 29.92 | 37895 | 7 | 7 | 7149887.5 | 8180150 | 8028069 | 8924805 | 9096925 | 9159905 | 8393055 | 8562687 | 8200053 | 8663128 | 8541611 | 8685763 | 9141113 | 6318908 |
| B1AXP6 | Tomm5 | Mitochondrial import receptor subunit TOM5 homolog Mus musculus | 38.98 | 19.61 | 6037 | 1 | 1 | 598000.94 | 803049.94 | 523047.84 | 456146.28 | 518794.09 | 562601.62 | 680288.88 | 745427.19 | 520960 | 586734.44 | 599994.88 | 696660 | 627355.56 | 493748.28 |
| Q9CZW5 | Tomm70 | Mitochondrial import receptor subunit TOM70 Mus musculus | 521.79 | 17.18 | 67590 | 10 | 10 | 26322325 | 23164519 | 18914515 | 23519605 | 21850621 | 20034171 | 18624847 | 19168111 | 18571787 | 18758403 | 22865805 | 26596017 | 20172785 | 22452845 |
| Q04750 | Top1 | DNA topoisomerase 1 Mus musculus | 448.38 | 16.3 | 90876 | 9 | 9 | 11616658 | 13068545 | 12492185 | 10531881 | 10801429 | 12869825 | 12588271 | 12812462 | 12015454 | 13115847 | 11019033 | 11171627 | 10881792 | 13004955 |
| Q01320 | Top2a | DNA topoisomerase 2-alpha Mus musculus | 2130.18 | 33.77 | 172790 | 34 | 41 | 33302683 | 28116785 | 34092885 | 32116889 | 35976841 | 28414301 | 26225959 | 22319379 | 29137521 | 28831121 | 33755761 | 37065873 | 38218009 | 35626761 |
| Q64511 | Top2b | DNA topoisomerase 2-beta Mus musculus | 581.87 | 9.49 | 181909 | 6 | 13 | 6199150.5 | 5327997 | 6893181 | 5923109 | 7973737 | 7997010 | 7174866 | 5347953.5 | 8188947 | 6164808 | 7784697 | 7713884 | 7506552 | 9160439 |

| | | | | | | | | | | | | | | | | | | | | | |
|---|---|---|---|---|---|---|---|---|---|---|---|---|---|---|---|---|---|---|---|---|---|
| Q68G38 | Tor1a | Torsin-1A Rattus norvegicus | 149.4 | 11.41 | 37949 | 3 | 3 | 2457221.25 | 1550796.75 | 1719114.25 | 1065484.25 | 1840050.75 | 2247728.5 | 2534569.5 | 1957562 | 2499332.5 | 2030875.88 | 1865744.75 | 1758577 | 2527396.25 | 2315429.75 |
| Q921T2 | Tor1aip1 | Torsin-1A-interacting protein 1 Mus musculus | 570.9 | 23.7 | 66781 | 11 | 11 | 14347833 | 14843691 | 18382027 | 12886996 | 13369572 | 12578905 | 10830982 | 13021696 | 14351020 | 14691007 | 13700707 | 10446359 | 14423423 | 12966592 |
| Q9ER41 | Tor1b | Torsin-1B Mus musculus | 159.29 | 12.8 | 37818 | 3 | 3 | 1876690.88 | 2223194 | 1662219.12 | 2164013.5 | 2329247.75 | 1548252.62 | 2129986.5 | 2818397.5 | 2497405.5 | 2237168.75 | 2213272.25 | 2141495.5 | 2865307.5 | 1371360.5 |
| Q9ER38 | Tor3a | Torsin-3A Mus musculus | 189.48 | 15.06 | 43814 | 4 | 4 | 5560227.5 | 4338360 | 3494521.5 | 4100928 | 3502685 | 3594878 | 3734765 | 4384059 | 3875051.25 | 3307942.25 | 1764952 | 3256605.5 | 3556890.5 | 2791821 |
| Q8BU11 | Tox4 | TOX high mobility group box family member 4 Mus musculus | 36.05 | 1.62 | 65961 | 1 | 1 | 1 | 365459.31 | 1 | 1 | 233071.77 | 340403.62 | 281942.38 | 698805.94 | 1 | 1 | 1 | 1 | 1 | 1 |
| Q62393 | Tpd52 | Tumor protein D52 Mus musculus | 484.65 | 43.3 | 24313 | 7 | 7 | 31624619 | 34161057 | 30067463 | 34338641 | 28192053 | 25940747 | 34083297 | 41563769 | 32652609 | 32885201 | 28492837 | 24756509 | 23475313 | 27336077 |
| Q6PCT3 | Tpd52l2 | Tumor protein D54 Rattus norvegicus | 368.03 | 35.45 | 23992 | 6 | 6 | 6379085 | 6059255.5 | 8893107 | 8007671 | 7538736 | 7903376.5 | 7445262 | 6768175.5 | 9625298 | 8765475 | 8506221 | 9383963 | 8747937 | 7128878.5 |
| P17751 | Tpi1 | Triosephosphate isomerase Mus musculus | 1395.28 | 72.91 | 32192 | 18 | 19 | 491184737 | 665689857 | 585128065 | 657117441 | 671526529 | 650650433 | 695287617 | 736615169 | 689315777 | 771902529 | 505927585 | 564298113 | 587008065 | 661814657 |
| P04692 | Tpm1 | Tropomyosin alpha-1 chain Rattus norvegicus | 483.89 | 17.96 | 32681 | 4 | 9 | 181939681 | 212132929 | 173293713 | 206825809 | 185180545 | 185013681 | 181433041 | 195000513 | 165206497 | 181729761 | 171899809 | 166830193 | 147172321 | 180091777 |
| P58774 | Tpm2 | Tropomyosin beta chain Mus musculus | 253.82 | 12.68 | 32837 | 0 | 5 | 186449921 | 213143153 | 175054945 | 208202273 | 187314593 | 186966097 | 182343873 | 196719025 | 168164801 | 184638769 | 174001953 | 167863729 | 148637569 | 182446385 |
| P21107 | Tpm3 | Tropomyosin alpha-3 chain Mus musculus | 643.1 | 32.63 | 32994 | 3 | 12 | 274021505 | 324433089 | 267948113 | 309771105 | 276667489 | 270503553 | 271210177 | 297951265 | 251516897 | 269315521 | 249717713 | 252121457 | 222881105 | 270019073 |
| Q6IRU2 | Tpm4 | Tropomyosin alpha-4 chain Mus musculus | 1222.06 | 58.87 | 28468 | 9 | 20 | 171244289 | 181799265 | 161074145 | 175204321 | 158968001 | 192287377 | 220924417 | 209447985 | 182344801 | 186597441 | 174388577 | 172576593 | 165195169 | 188302001 |
| O89023 | Tpp1 | Tripeptidyl-peptidase 1 Mus musculus | 494.53 | 28.29 | 61342 | 3 | 8 | 33809873 | 47168833 | 35362789 | 36958925 | 34351465 | 47698097 | 42514225 | 48979121 | 47771941 | 40657149 | 43374429 | 44452413 | 46082881 | 38542913 |
| Q64514 | Tpp2 | Tripeptidyl-peptidase 2 Mus musculus | 1755.26 | 37.56 | 139879 | 4 | 37 | 51876317 | 46468493 | 61087669 | 54042949 | 48974405 | 52853445 | 49722053 | 60016117 | 58549153 | 57479025 | 50132529 | 59836705 | 66626305 | 59717477 |
| F6ZDS4 | Tpr | Nucleoprotein TPR Mus musculus | 1444.52 | 16.58 | 273990 | 10 | 29 | 19221561 | 24636911 | 26501733 | 24298715 | 23207349 | 20541037 | 22515769 | 21111085 | 22910715 | 21658021 | 18315995 | 20139563 | 23730711 | 21667105 |
| A8WCF8 | Tprg1l | Tumor protein p63-regulated gene 1-like protein Rattus norvegicus | 38.68 | 6.77 | 29844 | 1 | 1 | 1653772.32 | 423464.59 | 650947.94 | 2443804.25 | 3096872.5 | 1 | 3337642.25 | 881302.38 | 1 | 3642289.5 | 2928011.75 | 1985466.207 | 631298.12 | 2397088.75 |
| P63028 | Tpt1 | Translationally-controlled tumor protein Mus musculus | 499.76 | 75.58 | 19462 | 8 | 8 | 26796833 | 23246987 | 23427929 | 17855349 | 18320633 | 23813583 | 23704593 | 33982545 | 29713469 | 19832017 | 20917865 | 20373197 | 27247165 | 23470373 |
| A2APB8 | Tpx2 | Targeting protein for Xklp2 Mus musculus | 145.9 | 6.44 | 85894 | 3 | 3 | 882806.94 | 1429659.75 | 1757683.75 | 1648329.12 | 1584731.88 | 934501.08 | 1143170.25 | 446226.44 | 1065802.25 | 1082805.38 | 1231902.25 | 1750879.75 | 1354553.12 | 1546631.62 |
| Q6PFR5 | Tra2a | Transformer-2 protein homolog alpha Mus musculus | 181.14 | 14.59 | 32316 | 3 | 4 | 5919604 | 5434528 | 3995547.75 | 6059926 | 6816700.5 | 5741717.5 | 5432981 | 4959786 | 4646382.5 | 5300329 | 5757979 | 5822069 | 5475229.5 | 5380131.5 |
| P62996 | Tra2b | Transformer-2 protein homolog beta Mus musculus | 413.94 | 23.96 | 33666 | 5 | 6 | 12482151 | 14761592 | 10882459 | 14648524 | 18349399 | 17489315 | 14840339 | 13705348 | 17226749 | 17072159 | 18845013 | 18032283 | 16895429 | 16319610 |
| Q3U0V2 | Tradd | Tumor necrosis factor receptor type 1-associated DEATH domain protein Mus musculus | 216.01 | 15.16 | 34577 | 3 | 3 | 1922995.5 | 3142418 | 1180384.12 | 1779870.62 | 1701181.12 | 2028545.25 | 1231041.62 | 2335729 | 1781037.75 | 2166155.25 | 819649.31 | 583542.81 | 2073510.5 | 2517245.5 |
| Q5XI41 | Tram1 | Translocating chain-associated membrane protein 1 Rattus norvegicus | 186.08 | 7.75 | 43030 | 3 | 3 | 4125778.25 | 825884.38 | 1388401 | 732517.75 | 1169592.12 | 1489383.62 | 756084.44 | 522455.5 | 1819890.62 | 1602147.62 | 1049307.38 | 1879986 | 2140322 | 1762248.25 |
| Q9CQN1 | Trap1 | Heat shock protein 75 kDa, mitochondrial Mus musculus | 993.01 | 31.16 | 80209 | 6 | 16 | 16313866 | 14580007 | 20722707 | 15664015 | 18790687 | 20664479 | 18313503 | 17061605 | 17616405 | 16191769 | 18402517 | 19760605 | 21240793 | 19479571 |
| B2RXC1 | Trappc11 | Trafficking protein particle complex subunit 11 Mus musculus | 83.56 | 2.82 | 128396 | 2 | 2 | 935231.8125 | 401060.81 | 1186073.38 | 1523631.5 | 630161.56 | 1095743.25 | 721928.735 | 698386.31 | 807955.88 | 285629.5 | 358594.91 | 1667954.25 | 339996.66 | 268172.22 |
| Q8K2L8 | Trappc12 | Trafficking protein particle complex subunit 12 Mus musculus | 44.54 | 1.13 | 87694 | 1 | 1 | 927506.655 | 889102.12 | 1309256.88 | 725802.62 | 785865 | 707273.12 | 1220832.5 | 956035.94 | 1053293.62 | 786948.56 | 819813.75 | 951360.5433 | 1053314.88 | 980953 |
| Q9CQA1 | Trappc5 | Trafficking protein particle complex subunit 5 Mus musculus | 84.5 | 16.49 | 20795 | 2 | 2 | 1020425.75 | 847211.19 | 917518.75 | 952143.19 | 691442.5 | 801275.19 | 857793.38 | 1132649.88 | 810274.25 | 1107915.62 | 1064753.38 | 888131.56 | 1072938.25 | 1129584.38 |
| Q9D289 | Trappc6b | Trafficking protein particle complex subunit 6B Mus musculus | 88.62 | 18.99 | 17936 | 2 | 2 | 1019057.25 | 429156.5 | 80090.02 | 604001.06 | 674858.69 | 654994.12 | 452892.31 | 504226.59 | 456810.06 | 456756.5 | 541904 | 431124.4467 | 407565.12 | 343904.22 |
| Q99NH8 | Trem2 | Triggering receptor expressed on myeloid cells 2 Mus musculus | 177.52 | 12.33 | 24527 | 2 | 2 | 1202371.345 | 1312040.38 | 979611.88 | 1306919.62 | 1210913.5 | 1141367 | 1390604.5 | 1306873.88 | 1421430 | 1632899.5 | 1588167.38 | 1270380 | 1673322.25 | 1749030.38 |
| Q91XB0 | Trex1 | Three-prime repair exonuclease 1 Mus musculus | 159.45 | 14.01 | 33675 | 3 | 3 | 3858302.5 | 6591903 | 3002202.5 | 5408457.5 | 5267453.5 | 2709213 | 1787539.12 | 2425848 | 2479948.25 | 3220306 | 2229514 | 2075390.62 | 2429171.5 | 2635683 |
| Q8BVW3 | Trim14 | Tripartite motif-containing protein 14 Mus musculus | 71.01 | 7.27 | 49641 | 2 | 2 | 1259331.88 | 1518784.5 | 1706086.38 | 852161.62 | 1193682.75 | 1258415.38 | 1006774.62 | 983320 | 1204913.88 | 778918.25 | 1115762.25 | 954924.25 | 1035475.25 | 117567.09 |
| Q61510 | Trim25 | E3 ubiquitin/ISG15 ligase TRIM25 Mus musculus | 42.96 | 1.89 | 71726 | 1 | 1 | 1870693.75 | 1544405.88 | 1901792.25 | 1853437.5 | 1749071.25 | 1385453 | 1568614 | 1712436.75 | 1528736.5 | 1536857.12 | 1493609.25 | 2162860.25 | 1389090.5 | 1927899.62 |
| P62603 | Trim26 | Tripartite motif-containing protein 26 Rattus norvegicus 3 | 48.08 | 3.51 | 62631 | 1 | 1 | 205442 | 153655.53 | 1 | 1 | 1 | 1 | 1 | 1 | 1 | 1 | 1 | 1 | 1 | 1 |
| Q62318 | Trim28 | Transcription intermediary factor 1-beta Mus musculus | 1250.12 | 35.13 | 88847 | 21 | 21 | 46268641 | 52740009 | 54350965 | 65436005 | 52466577 | 52208633 | 67542537 | 54991789 | 60648685 | 69035649 | 55884185 | 59298009 | 50144265 | 52522705 |
| Q8CH72 | Trim32 | E3 ubiquitin-protein ligase TRIM32 Mus musculus | 54 | 2.29 | 72057 | 1 | 1 | 622830.06 | 621233.69 | 698187.44 | 567710.69 | 2221752 | 595915.25 | 826562.56 | 989920.69 | 786647.12 | 646383 | 662676.25 | 598024 | 521344.81 | 460870.69 |
| Q99PP7 | Trim33 | E3 ubiquitin-protein ligase TRIM33 Mus musculus | 54.25 | 0.96 | 123843 | 1 | 1 | 1 | 1 | 902001.06 | 630287.94 | 1 | 777687.81 | 437122.5 | 360043.44 | 1 | 1 | 280374.97 | 869150.88 | 303788.5 | 822228.12 |
| Q80VI1 | Trim56 | E3 ubiquitin-protein ligase TRIM56 Mus musculus | 269.54 | 12.4 | 79513 | 5 | 5 | 1026202.38 | 905165 | 1186880.5 | 710034.62 | 1074505 | 794328.81 | 1135140.25 | 474495.41 | 622185.38 | 757804.12 | 901815.19 | 1256027.12 | 924221.25 | 897881.12 |
| Q99KW3 | Triobp | TRIO and F-actin-binding protein Mus musculus | 36.94 | 0.5 | 223368 | 1 | 1 | 1 | 500874.97 | 849372.25 | 1 | 634677.62 | 553160.12 | 365799.22 | 694710.62 | 313856.88 | 555860.25 | 430799.1867 | 428328.81 | 285786.69 | 578282.06 |
| P97531 | Trip10 | Cdc42-interacting protein 4 Rattus norvegicus | 131.09 | 9.51 | 62799 | 3 | 3 | 648270.38 | 886000.44 | 1134428.38 | 1 | 247482.66 | 809388.31 | 1185064.62 | 766393.75 | 739984.56 | 835714.69 | 990254.94 | 629495.81 | 957530 | 948088 |
| F1LP64 | Trip12 | E3 ubiquitin-protein ligase TRIP12 Rattus norvegicus | 719.57 | 10.32 | 223928 | 14 | 14 | 8402313 | 6882629.5 | 8042867.5 | 6013898 | 7340522.5 | 8662781 | 8516010 | 6910832 | 7804795 | 7809107.5 | 7193116.5 | 7179424.5 | 9415508 | 7024805.5 |
| Q3UA06 | Trip13 | Pachytene checkpoint protein 2 homolog Mus musculus | 95.9 | 5.32 | 48377 | 2 | 2 | 2079211.5 | 3025766.5 | 2844043 | 2085358.38 | 2357648.5 | 2479522.5 | 1025333 | 2499008.5 | 2169613.25 | 2019200.5 | 2589970 | 2460652.25 | 1321102.88 | 2502287.25 |
| Q3TX08 | Trmt1 | tRNA (guanine(26)-N(2))-dimethyltransferase Mus musculus | 105.58 | 4.52 | 72350 | 2 | 2 | 1127776.625 | 1147970.75 | 996056.25 | 1157810.12 | 1209269.38 | 940761.12 | 1074495.5 | 1073776.88 | 995710.81 | 915424.12 | 1163824.5 | 968688.38 | 1016140.81 | 1149427.5 |
| A2RSY6 | Trmt1l | TRMT1-like protein Mus musculus | 359.46 | 13.05 | 80861 | 6 | 6 | 2862737 | 2160277 | 1737797.25 | 1551891.12 | 1599637.25 | 2735926 | 3292781.75 | 2145009 | 2037076.25 | 2280881.5 | 3218521 | 2932769.5 | 2095661.25 | 1977402.38 |
| Q8BNV1 | Trmt2a | tRNA (uracil-5-)-methyltransferase homolog A Mus musculus | 32.54 | 1.74 | 63301 | 1 | 1 | 1 | 1 | 1 | 1 | 1 | 1 | 1 | 1 | 1 | 1 | 1 | 98444.1 | 126935.88 | 1 |
| Q9D0C4 | Trmt5 | tRNA (guanine(37)-N1)-methyltransferase Mus musculus | 42.14 | 1.8 | 56794 | 1 | 1 | 710043.5 | 916123.88 | 730682.31 | 818078.31 | 1 | 850658.06 | 745509.06 | 639919.62 | 665118.81 | 670635.5 | 807324.88 | 1 | 625349.75 | 838772.69 |
| Q8CE96 | Trmt6 | tRNA (adenine(58)-N(1))-methyltransferase non-catalytic subunit TRM6 Mus musculus | 243.1 | 10.87 | 55518 | 4 | 4 | 957405.56 | 714771.25 | 1303577.5 | 853337 | 957272.8275 | 218011.22 | 1194555.38 | 1130268.12 | 1252777 | 984866.12 | 1283039 | 1078599.88 | 1239046.62 | 1200228.5 |
| Q8K1J6 | Trnt1 | CCA tRNA nucleotidyltransferase 1, mitochondrial Mus musculus | 380.86 | 26.04 | 49895 | 9 | 9 | 10779536 | 9099076 | 9943824 | 8867748 | 11954073 | 9274870 | 7692941.5 | 8855822 | 7187872.5 | 8567276 | 10541735 | 7056790.5 | 11693053 | 9402080 |
| Q9WTR1 | Trpv2 | Transient receptor potential cation channel subfamily V member 2 Mus musculus | 274.13 | 8.86 | 85965 | 4 | 4 | 3648372.5 | 3049682.25 | 2692374.5 | 2306182 | 2552853 | 3079282 | 2700655 | 3610521.5 | 2901450.25 | 2757201 | 2955519 | 2932410.25 | 3099093 | 2746789 |
| Q9CZR8 | Tsfm | Elongation factor Ts, mitochondrial Mus musculus | 443.4 | 48.77 | 35334 | 7 | 7 | 8348572 | 9932174 | 8412015 | 8252476.5 | 8658664 | 6962736.5 | 7805462 | 8298605 | 9187687 | 8738711 | 8745494 | 5545814 | 7448281.5 | 7820907 |
| Q61187 | Tsg101 | Tumor susceptibility gene 101 protein Mus musculus | 168 | 12.28 | 44124 | 4 | 4 | 3959817.5 | 3640241.25 | 4122413.25 | 3878653.75 | 3800367.5 | 3938967 | 3694352 | 3114051 | 3540673.75 | 3704972 | 4200035.5 | 4237683.5 | 4020016.25 | 3394111.5 |
| Q62348 | Tsn | Translin Mus musculus | 734.46 | 60.96 | 26201 | 11 | 11 | 48678913 | 39909309 | 40311953 | 36538065 | 36860081 | 35599765 | 35729581 | 41773429 | 37005817 | 35335977 | 33801517 | 32918493 | 33818909 | 40247641 |
| Q9QZE7 | Tsnax | Translin-associated protein X Mus musculus | 385.33 | 42.41 | 32926 | 5 | 8 | 2525334 | 1737998.12 | 2549686 | 2309082 | 2140633.75 | 2128030 | 2402524.25 | 1900052.12 | 2038049.5 | 1995341 | 2197545.75 | 2604338.25 | 2034269.25 | 2838944 |
| Q9CQ88 | Tspan31 | Tetraspanin-31 Mus musculus | 45.3 | 5.71 | 22694 | 1 | 1 | 1429178.5 | 1445725.88 | 1218858.12 | 964262.25 | 1071089.12 | 1379232.25 | 1072913 | 1693323.88 | 1152923.62 | 1225666.25 | 1936776.25 | 1542836.75 | 1257330.38 | 1655167.5 |
| P50637 | Tspo | Translocator protein Mus musculus | 236.83 | 46.75 | 18841 | 4 | 4 | 26044009 | 6385051 | 5716660 | 5612279 | 11752173 | 9924216 | 6291304 | 3143281.5 | 6603584 | 8127208 | 8867401 | 7669372 | 9379265 | 8618802 |
| Q5SWD9 | Tsr1 | Pre-rRNA-processing protein TSR1 homolog Mus musculus | 513.92 | 15.69 | 92105 | 9 | 9 | 3070351 | 3919106.75 | 1639943 | 1836093.25 | 4685954 | 2027505.25 | 2594197.5 | 2582644 | 1519848.62 | 3276506 | 3108649.5 | 3201776 | 3384054.25 | 1602490 |
| Q5XIB1 | Tssc4 | Protein TSSC4 Rattus norvegicus | 39.93 | 3.15 | 33939 | 1 | 1 | 927600.94 | 1171855.62 | 1108860.62 | 1141664 | 1066645.38 | 956150.5 | 969503.88 | 1230436.5 | 847259.88 | 978466.12 | 952198.88 | 1048436 | 1093655.62 | 1186690.5 |
| P23591 | Tsta3 | GDP-L-fucose synthase Mus musculus | 230.02 | 20.25 | 35878 | 5 | 5 | 7730363.5 | 8365780 | 7979294 | 8123320 | 8205173.5 | 6914703 | 6895175 | 7710219 | 6726803.5 | 6897820.5 | 6750698 | 7032807 | 6528529.5 | 7176565 |
| Q9D6K7 | Ttc33 | Tetratricopeptide repeat protein 33 Mus musculus | 78.75 | 4.96 | 29371 | 1 | 1 | 2230320.75 | 2738433.5 | 2508648.25 | 2295292.25 | 2051364 | 2207965.75 | 2516706.5 | 2269535 | 2244324 | 2069343.88 | 2047474.62 | 2138017 | 2142574.5 | 2288533.5 |
| A3KMP2 | Ttc38 | Tetratricopeptide repeat protein 38 Mus musculus | 56.29 | 1.94 | 52224 | 1 | 1 | 476321.41 | 291528.38 | 521739.12 | 191043.45 | 318513.09 | 288087.78 | 310554.785 | 298658.88 | 399027.75 | 256444.73 | 412066.06 | 444073.75 | 423996.91 | 348023.81 |
| Q5BK48 | Ttc5 | Tetratricopeptide repeat protein 5 Rattus norvegicus | 72.23 | 6.36 | 48793 | 2 | 2 | 1 | 412489.59 | 538074.75 | 284557.75 | 1 | 570080.38 | 358013.75 | 1 | 1 | 220953.5 | 419248.78 | 244854.45 | 274071.78 | 279482.69 |
| Q5NC05 | Ttf2 | Transcription termination factor 2 Mus musculus | 100.85 | 2.9 | 125530 | 3 | 3 | 1898142.75 | 1383248 | 1174040.62 | 1850015.38 | 1632152.38 | 1321264.38 | 585877.25 | 415239.88 | 235110.8 | 648483.5 | 1504525.5 | 1336907.25 | 506270.47 | 1107042.38 |
| Q3UDE2 | Ttll12 | Tubulin--tyrosine ligase-like protein 12 Mus musculus | 493.99 | 25.67 | 74043 | 11 | 11 | 7946159.5 | 7170459.5 | 9636674 | 8998819 | 6736971 | 9606616 | 8141468.5 | 6049843.5 | 9677451 | 9564427 | 8149738 | 9731208 | 10477882 | 9080328 |
| P68369 | Tuba1a | Tubulin alpha-1A chain Mus musculus | 2138.28 | 75.39 | 50136 | 1 | 31 | 1277217537 | 1397301377 | 1323573889 | 1206727169 | 1333526913 | 1305008257 | 1312006785 | 1381117441 | 1375105281 | 1253303809 | 1402826753 | 1370941569 | 1346163713 | 1316314369 |
| P05213 | Tuba1b | Tubulin alpha-1B chain Mus musculus | 2208.86 | 75.39 | 50152 | 0 | 32 | 1412480385 | 1560608897 | 1517606529 | 1353083393 | 1480593025 | 1464311041 | 1479759745 | 1569487873 | 1550869249 | 1393855489 | 1573285249 | 1576691201 | 1530101249 | 1482702081 |
| P68373 | Tuba1c | Tubulin alpha-1C chain Mus musculus | 2169.27 | 71.05 | 49909 | 1 | 31 | 1361132033 | 1494732033 | 1480577537 | 1323368193 | 1421069185 | 1429105921 | 1439021953 | 1513045761 | 1498437889 | 1335882497 | 1513825153 | 1523515777 | 1472810113 | 1446271489 |
| P68368 | Tuba4a | Tubulin alpha-4A chain Mus musculus | 1677.86 | 64.96 | 49924 | 4 | 24 | 900939393 | 1041748737 | 1001077889 | 889242113 | 965179777 | 951503937 | 982387713 | 1052490497 | 1026873601 | 900758657 | 1022987009 | 1053322881 | 1006003329 | 918523585 |
| Q3UX10 | Tubal3 | Tubulin alpha chain-like 3 Mus musculus 2 | 144.57 | 11.66 | 49988 | 1 | 3 | 18683813 | 24827005 | 17155457 | 15835367 | 20305229 | 26396885 | 27993255 | 27852099 | 25230437 | 23117065 | 30590573 | 18002667 | 32408537 | 28968605 |
| P85108 | Tubb2a | Tubulin beta-2A chain Rattus norvegicus | 2026.68 | 58.2 | 49907 | 1 | 29 | 824646721 | 1311703297 | 1105619969 | 1120288129 | 1217874817 | 1139842049 | 957325249 | 990190657 | 1141614977 | 1167937281 | 1183348865 | 1166275329 | 1125334145 | 1256991233 |
| Q3KRE8 | Tubb2b | Tubulin beta-2B chain Rattus norvegicus | 2106.08 | 66.97 | 49953 | 1 | 30 | 827297537 | 1315153665 | 1108002561 | 1123333121 | 1220174081 | 1141882881 | 956873089 | 992509057 | 1143967489 | 1170400513 | 1185960577 | 1166011905 | 1125354881 | 1256991233 |
| Q4QRB4 | Tubb3 | Tubulin beta-3 chain Rattus norvegicus | 1327.39 | 38.89 | 50419 | 2 | 20 | 581574337 | 1012644865 | 854428033 | 848915585 | 927739137 | 853800769 | 671223233 | 697592705 | 865079745 | 876445825 | 883568897 | 894411329 | 847602241 | 948922689 |
| Q9D6F9 | Tubb4a | Tubulin beta-4A chain Mus musculus | 1632.81 | 70.95 | 49586 | 1 | 24 | 580736193 | 981451585 | 832888961 | 809792129 | 898003841 | 846021825 | 664602625 | 688424577 | 854360257 | 865500929 | 870651201 | 870636481 | 839448705 | 934485057 |
| P68372 | Tubb4b | Tubulin beta-4B chain Mus musculus | 2411.1 | 82.92 | 49831 | 1 | 34 | 707777793 | 1169986817 | 995606273 | 1004987777 | 1098371969 | 1030574337 | 844440065 | 860197889 | 1032313921 | 1066017025 | 1065478657 | 1059129537 | 1021420929 | 1133122817 |
| P69897 | Tubb5 | Tubulin beta-5 chain Rattus norvegicus | 2650.3 | 83.78 | 49671 | 6 | 36 | 1062513793 | 1632415745 | 1373111809 | 1368667521 | 1503917057 | 1451681281 | 1252703745 | 1287233281 | 1413058305 | 1461101953 | 1480413825 | 1460982017 | 1419812481 | 1571317249 |
| Q922F4 | Tubb6 | Tubulin beta-6 chain Mus musculus | 1448.46 | 65.32 | 50090 | 9 | 23 | 417234241 | 575988545 | 517882305 | 509164769 | 566127745 | 504519745 | 513567457 | 521575201 | 530690433 | 536847873 | 535705889 | 535518049 | 521761537 | 555376321 |
| P83887 | Tubg1 | Tubulin gamma-1 chain Mus musculus | 151.76 | 15.3 | 51101 | 4 | 4 | 3125687.75 | 3614018 | 8423060 | 3376430 | 3839941 | 3826689.5 | 11803887 | 9327195 | 9875611 | 3623616.5 | 2886114.25 | 8398760 | 6419595 | 4017401.25 |
| Q921G8 | Tubgcp2 | Gamma-tubulin complex component 2 Mus musculus | 39.08 | 1.44 | 103223 | 1 | 1 | 1 | 577559.94 | 937959.19 | 1 | 1 | 965030.75 | 749093.62 | 521842.03 | 630769.5 | 591365.69 | 710081.25 | 694554.94 | 710234.5 | 725454.31 |
| Q8BFR5 | Tufm | Elongation factor Tu, mitochondrial Mus musculus | 527.02 | 37.61 | 49508 | 11 | 11 | 17981583 | 18943269 | 14903769 | 19271393 | 19213115 | 19347441 | 20557705 | 19483163 | 21563411 | 16200442 | 19497465 | 16313425 | 18904509 | 19266577 |
| Q91YR1 | Twf1 | Twinfilin-1 Mus musculus | 266 | 21.71 | 40079 | 5 | 5 | 10499071 | 10866777 | 11384323 | 9589999 | 11015919 | 9002594 | 10923225 | 9511898 | 9773675 | 9219404 | 8580519 | 8403734 | 9452270 | 10555929 |
| Q9Z0P5 | Twf2 | Twinfilin-2 Mus musculus | 574.02 | 44.13 | 39471 | 8 | 8 | 7669736 | 5994880.5 | 8146624 | 8317787.5 | 8789805 | 6970179 | 5773652 | 5557620 | 7549960 | 8272131.5 | 6852277 | 7704051.5 | 8649494 | 7098383.5 |
| Q6PAM1 | Txlna | Alpha-taxilin Mus musculus | 130 | 6.14 | 62369 | 2 | 2 | 1280284.75 | 1384573.938 | 1239255.75 | 1673974.75 | 1344780.5 | 1286422.843 | 1188025.62 | 1049875.62 | 1649319.25 | 1258470.88 | 964751.31 | 1186727.75 | 852930.81 | 1151665.5 |
| P10639 | Txn | Thioredoxin Mus musculus | 439.25 | 46.67 | 11675 | 5 | 5 | 103845633 | 105214497 | 117619593 | 56874173 | 76428097 | 117400009 | 101248721 | 145315809 | 136745265 | 104514577 | 111716569 | 128556545 | 119138121 | 121567337 |
| P97493 | Txn2 | Thioredoxin, mitochondrial Mus musculus | 73.23 | 9.04 | 18255 | 1 | 1 | 1 | 339997.91 | 244386.84 | 1 | 298630.62 | 527099.5 | 1 | 1 | 1 | 499319.31 | 293626.53 | 303651.97 | 1 | 1 |
| Q498E0 | Txndc12 | Thioredoxin domain-containing protein 12 Rattus norvegicus | 126.59 | 24.71 | 19019 | 3 | 3 | 1352617.62 | 1492037.12 | 1199635.62 | 1423276.12 | 1626131.5 | 1271107.5 | 1491105.12 | 1561759.25 | 1270011.75 | 1362488.25 | 1769055.75 | 1290509 | 1310887.38 | 1533034.25 |
| Q9CQM5 | Txndc17 | Thioredoxin domain-containing protein 17 Mus musculus | 231.16 | 39.84 | 14015 | 4 | 4 | 10687913 | 11595468 | 12056245 | 11825727 | 11382406 | 11669695 | 11813478 | 11713321 | 12136583 | 12893105 | 12632143 | 12233735 | 12153640 | 12660701 |
| Q91W90 | Txndc5 | Thioredoxin domain-containing protein 5 Mus musculus | 531.49 | 30.22 | 46415 | 9 | 9 | 9467288 | 7339417.5 | 11260229 | 9304043 | 9124493 | 10301322 | 10807179 | 8694723 | 10011357 | 8876035 | 10536149 | 11198726 | 7834649.5 | 10105811 |
| Q8K581 | Txndc9 | Thioredoxin domain-containing protein 9 Rattus norvegicus 2 | 39.36 | 3.98 | 26272 | 1 | 1 | 314342.66 | 1 | 321456.78 | 432841.44 | 1 | 1 | 1 | 1 | 1 | 332622.47 | 1 | 1 | 1 | 1 |
| Q5M7W1 | Txnip | Thioredoxin-interacting protein Rattus norvegicus | 47.22 | 3.05 | 44018 | 1 | 1 | 1007481.5 | 1452248.25 | 1143807.88 | 740502.25 | 934415.31 | 914280 | 1104790.75 | 1029686.5 | 1045420.44 | 1283433 | 880691.75 | 843482.44 | 1038457.62 | 942437.69 |
| Q8CDN6 | Txnl1 | Thioredoxin-like protein 1 Mus musculus | 487.49 | 48.79 | 32237 | 10 | 10 | 8772278 | 10316230 | 7470956 | 9537057 | 11095900 | 9654209 | 8782822 | 10137358 | 9010271 | 9863067 | 8822223 | 9142849 | 9414841 | 10212674 |
| Q9JMH6 | Txnrd1 | Thioredoxin reductase 1, cytoplasmic Mus musculus | 814.75 | 38.17 | 67084 | 15 | 15 | 34148489 | 29293857 | 37767077 | 30790565 | 24352639 | 35039229 | 35218733 | 39995177 | 40289197 | 32210917 | 33625645 | 38290589 | 36724477 | 33540205 |
| P07607 | Tyms | Thymidylate synthase Mus musculus | 152.05 | 11.73 | 34958 | 3 | 3 | 1808373.75 | 1584692.88 | 1165552.25 | 1332723 | 1349304 | 949202.38 | 1386821.25 | 1298956.62 | 1205475.75 | 871274.12 | 490754.31 | 1898373 | 1831699.25 | 1757925.38 |

| | | | | | | | | | | | | | | | | | | | | | |
|---|---|---|---|---|---|---|---|---|---|---|---|---|---|---|---|---|---|---|---|---|---|
| Q9D883 | U2af1 | Splicing factor U2AF 35 kDa subunit Mus musculus | 309.82 | 22.59 | 27815 | 5 | 5 | 11448645 | 9639451 | 12226973 | 11376870 | 11514768 | 13254155 | 11905207 | 8186481 | 12109025 | 13132112 | 14635611 | 11606117 | 13221250 | 10744753 |
| P26369 | U2af2 | Splicing factor U2AF 65 kDa subunit Mus musculus | 826.54 | 42.74 | 53517 | 12 | 12 | 25919089 | 30357559 | 35058081 | 34916045 | 29659301 | 35371321 | 32002923 | 39485465 | 38426841 | 37827961 | 32380563 | 36104461 | 36808109 | 33075245 |
| Q6NV83 | U2surp | U2 snRNP-associated SURP motif-containing protein Mus musculus | 254.9 | 7.77 | 118261 | 6 | 6 | 1312861.75 | 345402.53 | 1885097.88 | 1140783.75 | 522111.66 | 1285734.25 | 1550221 | 918170.38 | 1655540.25 | 930763.75 | 170487.66 | 730691 | 1131757.25 | 179118.73 |
| Q91YN5 | Uap1 | UDP-N-acetylhexosamine pyrophosphorylase Mus musculus | 129.05 | 4.41 | 58609 | 1 | 2 | 6990582.5 | 7103780 | 7469785.5 | 8474543 | 8029916.5 | 6603780.5 | 7916273 | 5660285.5 | 7019534 | 7680168 | 7908706.5 | 8572946 | 7786934.5 | 8685193 |
| Q3TW96 | Uap1l1 | UDP-N-acetylhexosamine pyrophosphorylase-like protein 1 Mus musculus | 793.49 | 36.09 | 56614 | 13 | 14 | 52067277 | 56779461 | 59052273 | 56250797 | 61191093 | 57157121 | 61271297 | 50817661 | 58959793 | 56980593 | 63924037 | 60597257 | 64148873 | 62003045 |
| Q02053 | Uba1 | Ubiquitin-like modifier-activating enzyme 1 Mus musculus | 2035.76 | 41.68 | 117809 | 29 | 29 | 87735785 | 96284081 | 84422105 | 98210033 | 97630369 | 96491937 | 96518825 | 95098697 | 94515097 | 96815825 | 100513401 | 95330745 | 91539257 | 94484353 |
| Q9Z1F9 | Uba2 | SUMO-activating enzyme subunit 2 Mus musculus | 1202.24 | 44.2 | 70569 | 18 | 18 | 33409885 | 41087249 | 28453777 | 27419885 | 30028393 | 30765785 | 34218337 | 38105961 | 36875629 | 37866881 | 31931661 | 37774233 | 37407929 | 33808229 |
| Q8C878 | Uba3 | NEDD8-activating enzyme E1 catalytic subunit Mus musculus | 520.74 | 36.58 | 51737 | 10 | 10 | 7718246 | 9037193 | 11423203 | 12034932 | 11194233 | 11562415 | 10641751 | 12764645 | 11938135 | 12014803 | 10161168 | 11914076 | 10656156 | 12264459 |
| Q8VE47 | Uba5 | Ubiquitin-like modifier-activating enzyme 5 Mus musculus | 207.24 | 16.13 | 44790 | 4 | 4 | 2713038.5 | 2294605.25 | 2945991.75 | 2004612.5 | 3471097.5 | 3733676.25 | 2232783.5 | 2074804.88 | 3204027.5 | 4022232.25 | 2249176 | 2218144.75 | 1778463.12 | 3213514.5 |
| Q8C7R4 | Uba6 | Ubiquitin-like modifier-activating enzyme 6 Mus musculus | 499.16 | 15.19 | 117966 | 11 | 11 | 8222249.5 | 3920951.25 | 5513448.5 | 4615173 | 5408250 | 6578616.5 | 7529287 | 7015569 | 8325254 | 6783479.5 | 6446053.5 | 5980500 | 7125090 | 6227329 |
| Q91VX2 | Ubap2 | Ubiquitin-associated protein 2 Mus musculus | 250.7 | 6.18 | 117966 | 5 | 5 | 1 | 1 | 1018405.44 | 1134142.88 | 1689859.38 | 1425853 | 1373862.75 | 1221093.393 | 1322627.88 | 762029.94 | 1741814.5 | 1017619.75 | 1395769.88 | 1854867.12 |
| Q80X50 | Ubap2l | Ubiquitin-associated protein 2-like Mus musculus | 618.97 | 16.35 | 116799 | 11 | 11 | 21484491 | 20054767 | 21571233 | 20913437 | 22209131 | 20808061 | 19541083 | 20032955 | 21225367 | 22421459 | 21202377 | 19856521 | 20107001 | 21469633 |
| Q9D1C1 | Ube2c | Ubiquitin-conjugating enzyme E2 C Mus musculus | 111.75 | 17.32 | 19606 | 2 | 2 | 2829670.25 | 3431147.5 | 39140193 | 2226269.75 | 64193329 | 2559393 | 60500545 | 1698620.25 | 52990593 | 45766841 | 53180701 | 2979349 | 2671102 | 2680301.75 |
| P61078 | Ube2d3 | Ubiquitin-conjugating enzyme E2 D3 Rattus norvegicus 2 | 177.74 | 36.05 | 16687 | 3 | 3 | 11385062 | 11975149 | 10739178 | 10172582 | 11948288 | 11957667 | 11800608 | 12594949 | 10784067 | 13240657 | 12053140 | 12329890 | 11277965 | 10850190 |
| P63280 | Ube2i | SUMO-conjugating enzyme UBC9 Mus musculus | 286.27 | 53.8 | 18007 | 6 | 6 | 11402778 | 14158973 | 13540894 | 13210278 | 15421310 | 13762391 | 15624529 | 12826937 | 15550940 | 16858439 | 14273253 | 13601271 | 15464353 | 16284199 |
| Q9JJZ4 | Ube2j1 | Ubiquitin-conjugating enzyme E2 J1 Mus musculus | 128.33 | 11.01 | 34990 | 3 | 3 | 2193777.5 | 2154871.5 | 646798 | 3101341 | 3452234.25 | 2120294.75 | 4100785.5 | 2305806.25 | 2408471 | 2105530 | 2351277.5 | 2398574.75 | 2419540.5 | 2540433.5 |
| P61087 | Ube2k | Ubiquitin-conjugating enzyme E2 K Mus musculus | 292.54 | 38.5 | 22407 | 5 | 5 | 12695037 | 12623145 | 12040264 | 11796697 | 11388548 | 5910043.5 | 13052859 | 11546717 | 11436075 | 12594447 | 5660901.5 | 12365291 | 12282625 | 5383071 |
| P68037 | Ube2l3 | Ubiquitin-conjugating enzyme E2 L3 Mus musculus | 392.22 | 59.09 | 17862 | 6 | 6 | 18667483 | 24371525 | 25664853 | 18493715 | 24950803 | 19858265 | 27038209 | 21045047 | 20937217 | 26380821 | 20044885 | 23867087 | 22126995 | 22822883 |
| P61082 | Ube2m | NEDD8-conjugating enzyme Ubc12 Mus musculus | 304.42 | 35.52 | 20900 | 6 | 6 | 15813014 | 15193225 | 18217401 | 15580575 | 21075753 | 14617315 | 21549573 | 22616699 | 20609833 | 19945421 | 18239155 | 16339577 | 23037847 | 11159715 |
| P61089 | Ube2n | Ubiquitin-conjugating enzyme E2 N Mus musculus | 301.24 | 56.58 | 17138 | 6 | 7 | 12204837 | 13243491 | 12766043 | 11743315 | 13561907 | 10560760 | 12005223 | 12963096 | 10357571 | 12460656 | 11730907 | 11458081 | 12050034 | 10723505 |
| Q6ZPJ3 | Ube2o | (E3-independent) E2 ubiquitin-conjugating enzyme UBE2O Mus musculus | 94.23 | 2.56 | 140834 | 2 | 2 | 855169.88 | 1104413.5 | 1068413.38 | 895552.6425 | 554213.81 | 537698.19 | 702599.75 | 729718 | 709437.12 | 1201316.5 | 926204.5 | 608103.94 | 1084108.25 | 601336.56 |
| Q7TSS2 | Ube2q1 | Ubiquitin-conjugating enzyme E2 Q1 Mus musculus | 159.62 | 8.77 | 46173 | 3 | 3 | 852529.38 | 358739.12 | 1268407.12 | 1 | 1 | 439008.31 | 430919.03 | 1074328.5 | 912544.12 | 406217.19 | 598258.56 | 1008950 | 584387.56 | 560445.25 |
| Q9CQ37 | Ube2t | Ubiquitin-conjugating enzyme E2 T Mus musculus | 110.46 | 9.31 | 22975 | 1 | 2 | 2891851 | 3155219.25 | 3087698.25 | 2853902.25 | 2370678 | 2712414.75 | 2321146.5 | 2719548.5 | 2601693.25 | 3536485.5 | 2898664.25 | 2322406.25 | 2977250.25 | 2496097.5 |
| Q9CZY3 | Ube2v1 | Ubiquitin-conjugating enzyme E2 variant 1 Mus musculus | 250.93 | 29.25 | 16355 | 4 | 4 | 18891277 | 19778333 | 22533873 | 21941629 | 18727319 | 19498093 | 20875111 | 18281221 | 20697459 | 22167749 | 19436833 | 21880579 | 21502909 | 20226117 |
| Q3B7D1 | Ube2z | Ubiquitin-conjugating enzyme E2 Z Rattus norvegicus | 89.35 | 8.71 | 38352 | 2 | 2 | 1019622.62 | 1025016.25 | 1030110.81 | 835277.12 | 1065174.25 | 1038477.06 | 1051350.88 | 1280733.25 | 1001464.25 | 1364240.25 | 1068000.5 | 1036512.563 | 1024792.81 | 1016744.38 |
| O08759 | Ube3a | Ubiquitin-protein ligase E3A Mus musculus | 266.1 | 7.47 | 99819 | 5 | 5 | 27386849 | 20405337 | 25101607 | 27857019 | 29841137 | 31929091 | 31161861 | 18861973 | 33304221 | 33831869 | 29316013 | 34615381 | 29164987 | 25045347 |
| Q9ES00 | Ube4b | Ubiquitin conjugation factor E4 B Mus musculus | 102.79 | 4.01 | 133318 | 3 | 3 | 439971 | 962346 | 1530228.12 | 872385.12 | 728183.56 | 1416984.5 | 1064598 | 1781321.25 | 2358598 | 1071610 | 1085315.5 | 1528794.25 | 1398812 | 1086320 |
| Q9EPV8 | Ubl5 | Ubiquitin-like protein 5 Mus musculus | 38.38 | 12.33 | 8547 | 1 | 1 | 3824312.25 | 3188218.25 | 2980332.5 | 2408776.25 | 3759906.25 | 3473935.25 | 376057.62 | 2665005.25 | 2363800.155 | 2940202.5 | 2336059 | 2563763 | 534965.56 | 2854829.5 |
| Q99NB8 | Ubqln4 | Ubiquilin-4 Mus musculus | 147.67 | 11.58 | 63506 | 3 | 3 | 933609.69 | 1449453.75 | 1192275.25 | 987305.8925 | 373884.88 | 332234.09 | 1591570 | 995879.19 | 1194127.75 | 1120239.5 | 454807.59 | 1774508.62 | 1450023 | 1072120.12 |
| Q6WKZ8 | Ubr2 | E3 ubiquitin-protein ligase UBR2 Mus musculus | 84.28 | 0.74 | 199155 | 1 | 1 | 387065.75 | 498156.81 | 519618.09 | 286536.16 | 201287.05 | 545233.25 | 289660.06 | 533465.56 | 555270.75 | 401492.41 | 405610.44 | 571123.94 | 523980.34 | 342735.53 |
| A2AN08 | Ubr4 | E3 ubiquitin-protein ligase UBR4 Mus musculus | 1210.58 | 6.12 | 572290 | 22 | 22 | 13464246 | 15326411 | 13744404 | 14036837 | 11869789 | 17011859 | 14981061 | 17499269 | 17161905 | 15428996 | 14048884 | 15509513 | 16232494 | 14690013 |
| Q62671 | Ubr5 | E3 ubiquitin-protein ligase UBR5 Rattus norvegicus | 126.4 | 1.97 | 308027 | 2 | 2 | 1 | 184980.03 | 406640.84 | 525496.25 | 1 | 624391.19 | 1 | 412724.06 | 1 | 1 | 525036.75 | 1 | 1 | 1 |
| P25976 | Ubtf | Nucleolar transcription factor 1 Mus musculus | 94.05 | 3.27 | 89509 | 2 | 2 | 1509451.88 | 1644149 | 1590948.5 | 1521410.88 | 2415347 | 528094.81 | 535609.31 | 1381853.62 | 1451999 | 1501012.88 | 2249148.5 | 1146579.75 | 1312194.38 | 1339223 |
| Q922Y1 | Ubxn1 | UBX domain-containing protein 1 Mus musculus | 390.34 | 34.34 | 33573 | 5 | 6 | 6506754.5 | 9303493 | 4879799 | 10758774 | 5061277.5 | 6837156.5 | 4849077.5 | 7197456.5 | 3961593 | 5505180.5 | 7345695.5 | 10699069 | 7324001.5 | 6106101.5 |
| Q8VCH8 | Ubxn4 | UBX domain-containing protein 4 Mus musculus | 238.39 | 12.85 | 56472 | 4 | 4 | 3544424.25 | 4576210.5 | 3864571 | 3838892.75 | 4312834.5 | 3904445.25 | 3543010.25 | 3288161.75 | 3604221.75 | 3745262 | 3495833 | 3542748.25 | 3743194 | 1994579.25 |
| Q99PL6 | Ubxn6 | UBX domain-containing protein 6 Mus musculus | 84.89 | 6.33 | 49796 | 2 | 2 | 2164644 | 2293407.75 | 2197554 | 2322861 | 724293.75 | 2256024.5 | 2535539.25 | 2471313.25 | 2489127.75 | 2196002 | 1467967.25 | 1601173.88 | 2291250.5 | 2428906.5 |
| Q9JKB1 | Uchl3 | Ubiquitin carboxyl-terminal hydrolase isozyme L3 Mus musculus | 234.9 | 23.04 | 26152 | 4 | 4 | 5858486 | 5441184 | 5181906 | 5494765 | 5473786.5 | 5661324.5 | 5327440 | 5731085 | 5139306 | 6028757 | 5461353 | 6011416.5 | 5620408 | 5831801 |
| Q9WUP7 | Uchl5 | Ubiquitin carboxyl-terminal hydrolase isozyme L5 Mus musculus | 816.93 | 66.26 | 37617 | 16 | 16 | 18753543 | 16516076 | 16795345 | 18152443 | 18655609 | 19461481 | 17815527 | 21781965 | 20985781 | 18200319 | 21014761 | 21398901 | 19105791 | 17970113 |
| Q3U1V6 | Uevld | Ubiquitin-conjugating enzyme E2 variant 3 Mus musculus | 110.15 | 14.01 | 51681 | 3 | 3 | 1330330 | 1469679 | 2752659.25 | 1350396.62 | 2624959.5 | 1584245 | 1094319.25 | 1654101 | 1316289.88 | 2207276.5 | 1565456.38 | 2143654.5 | 1476874.75 | 2261458 |
| Q9CR09 | Ufc1 | Ubiquitin-fold modifier-conjugating enzyme 1 Mus musculus | 173.85 | 31.14 | 19481 | 4 | 4 | 7331790 | 6625408.5 | 6249144.5 | 5932308 | 6208152.5 | 4703334 | 4986133.5 | 5569932 | 5297996.5 | 4591163 | 5214540.5 | 4845140 | 5559540 | 3579490.25 |
| P70362 | Ufd1 | Ubiquitin recognition factor in ER-associated degradation protein 1 Mus musculus | 206.76 | 19.54 | 34481 | 5 | 5 | 5358554 | 5680012 | 5781273.5 | 5793175 | 5958451 | 5615487 | 5314407.5 | 5580491 | 5112414.5 | 5951618.5 | 5631511 | 5933647 | 6069112 | 5661828 |
| Q8CCJ3 | Ufl1 | E3 UFM1-protein ligase 1 Mus musculus | 407.08 | 13.49 | 89520 | 6 | 6 | 3361500 | 3305349 | 3690832.75 | 3432519.25 | 1910361.5 | 2488742.5 | 3058311.25 | 3941415.5 | 3158208 | 2803838 | 3555218.75 | 2617377 | 3194888.25 | 4196707.5 |
| Q99K23 | Ufsp2 | Ufm1-specific protease 2 Mus musculus | 263.03 | 17.14 | 52515 | 6 | 6 | 3958302.5 | 4498152 | 4389915.5 | 5304419 | 4965032 | 4432660 | 2965417.25 | 2864233 | 3166457 | 4688020.5 | 5160139 | 5668126.5 | 5633820 | 5559861.5 |
| O70199 | Ugdh | UDP-glucose 6-dehydrogenase Rattus norvegicus | 46.3 | 1.83 | 54892 | 1 | 1 | 1 | 1 | 1 | 1 | 1 | 1 | 1 | 134066.55 | 1 | 1 | 1 | 1 | 191973.52 | 190666.55 |
| Q6P5E4 | Uggt1 | UDP-glucose:glycoprotein glucosyltransferase 1 Mus musculus | 2310.6 | 40.04 | 176434 | 41 | 42 | 55591629 | 46955769 | 47187621 | 46302633 | 51739361 | 58756277 | 41854617 | 48844337 | 51505837 | 56241797 | 58260673 | 62031585 | 62840517 | 53944741 |
| Q91ZJ5 | Ugp2 | UTP--glucose-1-phosphate uridylyltransferase Mus musculus | 278.07 | 13.78 | 56979 | 6 | 6 | 1285799.88 | 5131077 | 4584418 | 6727989.5 | 7102984 | 6606651 | 5044091.5 | 4283211 | 2743090.5 | 5283748.5 | 5236974.5 | 7644362 | 4643455.5 | 5191613.5 |
| Q6ZQM8 | Ugt1a7c | UDP-glucuronosyltransferase 1-7C Mus musculus | 1044.94 | 45.01 | 59758 | 18 | 19 | 78058561 | 83750217 | 64516753 | 77695905 | 77941233 | 86669297 | 84737585 | 84050017 | 83624129 | 87147425 | 85040953 | 78555441 | 81012153 | 88514065 |
| Q8VDF2 | Uhrf1 | E3 ubiquitin-protein ligase UHRF1 Mus musculus | 849.41 | 25.45 | 88304 | 11 | 16 | 6293807.5 | 2417324 | 6369521.5 | 7107202 | 6277823 | 7188613.5 | 7208659 | 6399633 | 6025060.5 | 7411285.5 | 6708959 | 7836394 | 7355068.5 | 7598416.5 |
| P13439 | Umps | Uridine 5'-monophosphate synthase Mus musculus | 809.7 | 44.07 | 52292 | 15 | 15 | 22133903 | 23610109 | 22258945 | 21785751 | 18322199 | 21525029 | 19069559 | 17998035 | 17285439 | 18592689 | 21257127 | 18681123 | 20652555 | 19932577 |
| Q62885 | Unc119 | Protein unc-119 homolog A Rattus norvegicus | 61.98 | 5.42 | 27048 | 1 | 1 | 1 | 1 | 1 | 1 | 1 | 1 | 1 | 1 | 606005.44 | 859388.88 | 1 | 1 | 582684.88 | 1 |
| B2RUP2 | Unc13d | Protein unc-13 homolog D Mus musculus | 36.8 | 1.01 | 123119 | 1 | 1 | 247152.02 | 184416.53 | 91451.2 | 211817.7 | 86256.26 | 186871.22 | 235833.48 | 175030.01 | 87108.17 | 190307.17 | 263540.12 | 134711.97 | 155460.42 | 62996.73 |
| Q99KD5 | Unc45a | Protein unc-45 homolog A Mus musculus | 119.85 | 4.24 | 103447 | 2 | 2 | 1 | 1 | 271007.47 | 93428.48 | 1 | 296903 | 1 | 406476.56 | 263799.78 | 1 | 62887.86 | 141898.32 | 192356.66 | 170450.44 |
| Q8VCW4 | Unc93b1 | Protein unc-93 homolog B1 Mus musculus | 358.12 | 13.21 | 66981 | 6 | 6 | 10856783 | 4375958.5 | 4493601 | 3998579 | 3573923 | 4574182 | 3153528.75 | 3670316.5 | 4024221.25 | 2485370.5 | 3351883 | 3924257.5 | 5018807.5 | 3076578 |
| Q9EPU0 | Upf1 | Regulator of nonsense transcripts 1 Mus musculus | 769.85 | 16.64 | 123967 | 15 | 15 | 4513688.5 | 4762802 | 4437975 | 5356068 | 5269039.5 | 6113806.5 | 5674726 | 4905793.5 | 6596554.5 | 6618581.5 | 5310971 | 8054423.5 | 5380112.5 | 6454206 |
| Q9CWU6 | Uqcc1 | Ubiquinol-cytochrome-c reductase complex assembly factor 1 Mus musculus | 139.84 | 10.51 | 34300 | 3 | 3 | 1099861.5 | 3819063.5 | 3298992.5 | 2966367.5 | 2181181.5 | 3014725 | 3382141 | 4102760.5 | 2477155.5 | 3101642.25 | 2065909.75 | 3314080.25 | 3996497 | 3253617.5 |
| Q8R1I1 | Uqcr10 | Cytochrome b-c1 complex subunit 9 Mus musculus | 74.73 | 37.5 | 7446 | 2 | 2 | 5445867 | 1762729.62 | 2465869.75 | 2581725.5 | 3023601 | 2526979.75 | 2382890.5 | 3317095.5 | 2564909.75 | 2280291.5 | 2973806.5 | 2383359.75 | 2661291 | 2457954 |
| Q9CZ13 | Uqcrc1 | Cytochrome b-c1 complex subunit 1, mitochondrial Mus musculus | 833 | 44.38 | 52852 | 13 | 13 | 18654481 | 21889973 | 21706489 | 20146351 | 20867265 | 21523859 | 23278403 | 22013623 | 20275139 | 23378021 | 25586275 | 21354603 | 23129957 | 25110145 |
| Q9DB77 | Uqcrc2 | Cytochrome b-c1 complex subunit 2, mitochondrial Mus musculus | 1118.98 | 68.65 | 48235 | 18 | 18 | 20296187 | 25128709 | 18761921 | 18604255 | 21652477 | 22630673 | 23371405 | 22630111 | 19103593 | 18738701 | 23841179 | 20640319 | 20881753 | 20068369 |
| Q9CR68 | Uqcrfs1 | Cytochrome b-c1 complex subunit Rieske, mitochondrial Mus musculus | 358.37 | 41.97 | 29368 | 7 | 7 | 8335855 | 9108856 | 12914313 | 9917850 | 10165819 | 10832690 | 10761925 | 9001791 | 9803859 | 12266985 | 10850898 | 13131041 | 12565305 | 12427729 |
| P99028 | Uqcrh | Cytochrome b-c1 complex subunit 6, mitochondrial Mus musculus | 131.91 | 28.09 | 10435 | 2 | 2 | 4904808 | 4955743.5 | 4000095 | 5226052 | 5368117 | 4912943 | 4295990 | 5340537 | 4646376.5 | 4783739.5 | 5576356 | 4484605.5 | 4982266 | 5853043 |
| Q7TQ16 | Uqcrq | Cytochrome b-c1 complex subunit 8 Rattus norvegicus | 94.57 | 24.39 | 9849 | 2 | 2 | 1 | 716689.88 | 1 | 224516.77 | 1 | 1 | 1 | 1 | 808951.75 | 706289.56 | 610480.3533 | 418667.72 | 219534.22 | 1193239.12 |
| P70697 | Urod | Uroporphyrinogen decarboxylase Mus musculus | 321.18 | 23.43 | 40692 | 5 | 5 | 5958787.5 | 5213655.5 | 4495623.5 | 5482164 | 5296505 | 4857380.5 | 5246858.5 | 4788502 | 5086060.5 | 5677795 | 4958597.5 | 4938660.5 | 5131658 | 4728020 |
| Q9Z1Z0 | Uso1 | General vesicular transport factor p115 Mus musculus | 717.45 | 20.44 | 106983 | 15 | 15 | 20782991 | 23835381 | 23481537 | 25961857 | 23067939 | 24377347 | 23196913 | 24771263 | 24647993 | 24169501 | 24853731 | 23022323 | 23925319 | 24606753 |
| P52479 | Usp10 | Ubiquitin carboxyl-terminal hydrolase 10 Mus musculus | 283.26 | 12.75 | 87022 | 7 | 7 | 4203533 | 2332308.75 | 3097966.25 | 4501043 | 2947893.5 | 3025427.5 | 2525839.5 | 2671065.5 | 2997817 | 2332892.5 | 3499622 | 2547670 | 2854706 | 2734919.5 |
| Q9JMA1 | Usp14 | Ubiquitin carboxyl-terminal hydrolase 14 Mus musculus | 1417.72 | 68.56 | 56002 | 25 | 25 | 40458277 | 38805877 | 41205489 | 44496385 | 43180869 | 37854729 | 41464689 | 44219341 | 43612057 | 42377981 | 38106457 | 42600541 | 41947481 | 43723097 |
| Q8R5H1 | Usp15 | Ubiquitin carboxyl-terminal hydrolase 15 Mus musculus | 480.37 | 15.7 | 112325 | 10 | 11 | 6104938.5 | 5670110 | 4913352.5 | 5859005.5 | 6201124 | 6122261.5 | 5595544.5 | 5194894 | 5782400 | 5350468 | 4613441.5 | 6195439 | 4643786 | 5911562.5 |
| Q9WTV6 | Usp18 | Ubl carboxyl-terminal hydrolase 18 Mus musculus | 76.99 | 6.25 | 42472 | 2 | 2 | 1 | 1 | 178631.36 | 182691.03 | 1 | 1 | 1 | 1 | 1 | 71848.65 | 180561.36 | 273666 | 109868.52 | 158149.56 |
| Q3UJD6 | Usp19 | Ubiquitin carboxyl-terminal hydrolase 19 Mus musculus | 226.22 | 4.34 | 150549 | 5 | 5 | 1141519.88 | 1723307.75 | 1526260.88 | 1577298.75 | 1645881.88 | 1688510.75 | 1475033 | 1496859.75 | 1823634 | 1433149.75 | 1733014.5 | 1600440.75 | 1359478.25 | 1508824.88 |
| B1AY13 | Usp24 | Ubiquitin carboxyl-terminal hydrolase 24 Mus musculus | 77.01 | 1.15 | 294001 | 2 | 2 | 663189.25 | 624075 | 769901.38 | 528914.75 | 505905.31 | 687056.12 | 616868.12 | 851699.31 | 771447.5 | 700927.62 | 567499.81 | 662880.44 | 730513.12 | 710140.75 |
| P57080 | Usp25 | Ubiquitin carboxyl-terminal hydrolase 25 Mus musculus | 61.6 | 0.95 | 121420 | 1 | 1 | 1 | 242203.67 | 1 | 1 | 1 | 1 | 1 | 181997.88 | 137784.75 | 149873.08 | 163148.48 | 197116.9267 | 168118.77 | 260083.53 |
| Q3TIX9 | Usp39 | U4/U6.U5 tri-snRNP-associated protein 2 Mus musculus | 553.77 | 26.77 | 65146 | 11 | 11 | 8693761 | 10850620 | 9177515 | 7901195.5 | 11353966 | 12422682 | 8750624 | 9977782 | 11694070 | 8238734.5 | 8322220 | 12868541 | 12740237 | 12326037 |
| B2GUZ1 | Usp4 | Ubiquitin carboxyl-terminal hydrolase 4 Rattus norvegicus | 421.77 | 11.24 | 108373 | 7 | 8 | 8456999 | 7108444 | 6530091.5 | 5582978 | 6051841.5 | 7742342 | 6164108 | 8056498 | 7669033.5 | 7479353 | 8050709.5 | 7779185.5 | 7055301 | 7137254.5 |
| Q8BY87 | Usp47 | Ubiquitin carboxyl-terminal hydrolase 47 Mus musculus | 140.46 | 2.69 | 157455 | 3 | 3 | 602432.31 | 2089618.62 | 1939205.5 | 1105125.12 | 1603429 | 1922655.5 | 1652161.25 | 2386808.75 | 2006353.88 | 1554632.12 | 1935029.38 | 2412662 | 2174536.25 | 517943.91 |
| P56399 | Usp5 | Ubiquitin carboxyl-terminal hydrolase 5 Mus musculus | 1665.19 | 45.8 | 95833 | 26 | 26 | 84498225 | 93753113 | 91397561 | 92476553 | 89759745 | 93168073 | 90257329 | 96995057 | 93060593 | 91252377 | 86292369 | 88905241 | 91758337 | 90596025 |
| Q4VSI4 | Usp7 | Ubiquitin carboxyl-terminal hydrolase 7 Rattus norvegicus | 219.57 | 4.81 | 128431 | 5 | 5 | 3975493.25 | 2977166 | 2808469.75 | 2560292.25 | 2880270 | 3385502.5 | 2649717.75 | 3372905.25 | 3864696.75 | 2855063 | 1322608.12 | 2519939 | 2964181.25 | 2582478.75 |
| P70398 | Usp9x | Probable ubiquitin carboxyl-terminal hydrolase FAF-X Mus musculus | 412.87 | 4.34 | 290711 | 8 | 8 | 2325340.5 | 3451192.75 | 3006313.5 | 3166280.75 | 2296593.5 | 3801517.25 | 3528626.75 | 3714584.25 | 3436606.5 | 2162318.5 | 2117481.25 | 3373625 | 3159438.75 | 3017765 |
| Q640M1 | Utp14a | U3 small nucleolar RNA-associated protein 14 homolog A Mus musculus | 67.13 | 4.3 | 87265 | 2 | 2 | 1 | 463575.97 | 1 | 1 | 594368.88 | 1 | 1 | 1 | 1 | 1 | 506896.53 | 415613.69 | 301937.94 | 408149.3867 |
| Q8C7V3 | Utp15 | U3 small nucleolar RNA-associated protein 15 homolog Mus musculus | 345.09 | 18.94 | 59375 | 7 | 7 | 2518481.5 | 2785008 | 2513311.5 | 2797119 | 2384890.75 | 2224997 | 3524819.5 | 2780430.5 | 2189631.25 | 2404993.5 | 2489312.5 | 2270642 | 2648961 | 2924197.5 |
| Q5SSI6 | Utp18 | U3 small nucleolar RNA-associated protein 18 homolog Mus musculus | 502.03 | 30.98 | 61218 | 9 | 10 | 7200306.5 | 6874788.5 | 6430860 | 7406476 | 8152441 | 6361275 | 7112043 | 5903111 | 6727726 | 6322473.5 | 7110709.5 | 6285612.5 | 5525501.5 | 6553286 |
| Q9JI13 | Utp3 | Something about silencing protein 10 Mus musculus | 271.13 | 13.43 | 53399 | 5 | 5 | 3654444.75 | 4159856 | 2575663 | 4289463 | 4867771 | 4238020 | 1562859.38 | 2058519.75 | 1478170.88 | 2930099.25 | 2998254.5 | 3655944.75 | 2510851 | 1687413.5 |
| Q8R2N2 | Utp4 | U3 small nucleolar RNA-associated protein 4 homolog Mus musculus | 271.32 | 11.95 | 76909 | 5 | 5 | 1 | 1 | 806678.25 | 891446 | 1 | 1 | 686317.94 | 583981.75 | 1034494.5 | 1 | 1 | 1327889.38 | 1277346.38 | 1 |
| Q8K245 | Uvrag | UV radiation resistance associated protein Mus musculus | 45.36 | 1.43 | 77525 | 1 | 1 | 1 | 1 | 1 | 1 | 1 | 1 | 1 | 1 | 1 | 187039.83 | 1 | 1 | 166265.88 | 394849.28 |
| Q9WTZ0 | Uxt | Protein UXT Mus musculus | 45.66 | 7.01 | 18187 | 1 | 1 | 840198.69 | 989501.88 | 870619.56 | 887313.31 | 897253.56 | 1135164.38 | 1100218.62 | 977668.56 | 1029321.56 | 855861 | 1047703.69 | 780358.44 | 946210.88 | 872660.75 |
| Q80WQ2 | Vac14 | Protein VAC14 homolog Mus musculus | 768.49 | 25.96 | 88048 | 14 | 14 | 13528564 | 10920689 | 10440508 | 9211076 | 10567934 | 14573422 | 11720757 | 14294620 | 14138150 | 11784666 | 13081669 | 12649819 | 13982791 | 14690806 |
| P63024 | Vamp3 | Vesicle-associated membrane protein 3 Mus musculus | 90.95 | 32.04 | 11480 | 2 | 2 | 1 | 1 | 1 | 1 | 411596.5 | 667802.56 | 346006.72 | 485681.155 | 422271.72 | 506643.62 | 1 | 605893.94 | 1 | 1 |
| O70404 | Vamp8 | Vesicle-associated membrane protein 8 Mus musculus | 63.54 | 11.88 | 11451 | 1 | 1 | 391275.59 | 425782.22 | 429650.78 | 268917.22 | 469063.53 | 301782.06 | 404570.06 | 259131.28 | 297087.4175 | 222866.27 | 353154.5967 | 415894.53 | 223240.45 | 420328.81 |
| Q9WV55 | Vapa | Vesicle-associated membrane protein-associated protein A Mus musculus | 329.96 | 31.73 | 27855 | 6 | 7 | 10996689 | 11994211 | 14204713 | 13406575 | 13966539 | 14221193 | 13735304 | 12784598 | 13922437 | 18068739 | 15357873 | 13316421 | 17025021 | 15547157 |
| Q9QY76 | Vapb | Vesicle-associated membrane protein-associated protein B Mus musculus | 395.28 | 28.81 | 26946 | 7 | 8 | 7922319 | 4367373 | 9264227 | 8058532.5 | 7800590.5 | 8494146 | 7724784 | 6686160.5 | 9549265 | 10734207 | 8546912 | 9518185 | 10804054 | 10221844 |
| Q9Z1Q9 | Vars | Valine--tRNA ligase Mus musculus | 2525.76 | 51.46 | 140215 | 44 | 44 | 99511017 | 107113025 | 104057321 | 105556265 | 107404969 | 105193617 | 109168705 | 114276233 | 111017353 | 116902961 | 105508913 | 106772745 | 108946809 | 110761169 |

| | | | | | | | | | | | | | | | | | | | | | |
|---|---|---|---|---|---|---|---|---|---|---|---|---|---|---|---|---|---|---|---|---|---|
| Q9CZT5 | Vasn | Vasorin Mus musculus | 54.53 | 1.04 | 72261 | 1 | 1 | 3280807.5 | 3123268.5 | 3746866.5 | 3503898 | 3298635.5 | 3284849 | 3381837.25 | 2920945.25 | 3348123.5 | 3463898.25 | 3407533.25 | 3280307.25 | 2974641.25 | 3141709 |
| P70460 | Vasp | Vasodilator-stimulated phosphoprotein Mus musculus | 398.93 | 26.4 | 39667 | 8 | 8 | 7519228 | 6332024 | 9423862 | 10308521 | 9601369 | 10449936 | 10257508 | 7091594 | 9453908 | 12531467 | 10498012 | 10570444 | 9856680 | 9816809 |
| Q62465 | Vat1 | Synaptic vesicle membrane protein VAT-1 homolog Mus musculus | 635.9 | 35.47 | 43097 | 10 | 10 | 13013952 | 12600807 | 11770273 | 9058213 | 10924196 | 15030296 | 22454865 | 16856367 | 19143909 | 19103937 | 17374861 | 18009465 | 19733481 | 17329637 |
| P27870 | Vav1 | Proto-oncogene vav Mus musculus | 1530.61 | 33.96 | 98137 | 22 | 23 | 51367053 | 43368361 | 55340193 | 44861849 | 49595133 | 50096065 | 44261809 | 50862541 | 54029121 | 49860293 | 50323145 | 53105001 | 51524877 | 50474161 |
| Q9R0C8 | Vav3 | Guanine nucleotide exchange factor VAV3 Mus musculus | 464.12 | 12.28 | 97968 | 8 | 9 | 6905495.5 | 5524216 | 6153152.5 | 5147491 | 5953059.5 | 6258924.5 | 5279247.5 | 6920806 | 6850885 | 5264901 | 5077142 | 6250728 | 5999430.5 | 7218218.5 |
| P61759 | Vbp1 | Prefoldin subunit 3 Mus musculus | 364.86 | 38.27 | 22436 | 7 | 7 | 5948499.5 | 6263599 | 6214578.5 | 4969961 | 4569902 | 7385223 | 6022266 | 8264419 | 7723952.5 | 5685436.5 | 5389260 | 8466902 | 5812721.5 | 6048433.5 |
| Q64727 | Vcl | Vinculin Mus musculus | 958.92 | 20.17 | 116717 | 17 | 17 | 16008462 | 27164945 | 24175101 | 29153369 | 28499287 | 28650565 | 29445117 | 18472587 | 18586951 | 29576553 | 29336329 | 26132791 | 24344601 | 29884083 |
| P46462 | Vcp | Transitional endoplasmic reticulum ATPase Rattus norvegicus | 2628.17 | 61.41 | 89349 | 39 | 41 | 111624025 | 112881953 | 111558361 | 120316785 | 111253761 | 113236177 | 112428121 | 119571329 | 106024137 | 122021633 | 109981473 | 124154833 | 112021993 | 108377849 |
| Q8CDG3 | Vcpip1 | Deubiquitinating protein VCIP135 Mus musculus | 117.73 | 2.79 | 134503 | 2 | 2 | 1332798 | 1507742.5 | 1314535.25 | 1718037.88 | 1391673.38 | 1 | 1609224.25 | 1350859.88 | 1 | 1339784.12 | 1244579.75 | 1332891.38 | 1 | 442961.56 |
| Q9Z2L0 | Vdac1 | Voltage-dependent anion-selective channel protein 1 Rattus norvegicus | 1323.92 | 78.8 | 30756 | 17 | 17 | 90162177 | 100447313 | 85169633 | 88416833 | 102944129 | 107703945 | 104512193 | 100083985 | 107296049 | 104988993 | 92626481 | 104174233 | 104296585 | 106404737 |
| Q60930 | Vdac2 | Voltage-dependent anion-selective channel protein 2 Mus musculus | 1007.83 | 56.61 | 31733 | 13 | 14 | 155728449 | 168903905 | 177213537 | 146716865 | 154621441 | 175781217 | 180012129 | 184517217 | 184701425 | 170132913 | 153915905 | 164444097 | 179458209 | 176108481 |
| Q60931 | Vdac3 | Voltage-dependent anion-selective channel protein 3 Mus musculus | 600 | 38.87 | 30753 | 7 | 9 | 42390313 | 40488137 | 52823513 | 36550153 | 37206093 | 50436737 | 49743029 | 48819677 | 56829201 | 43844413 | 40324005 | 51983345 | 50131249 | 45134753 |
| P20152 | Vim | Vimentin Mus musculus | 3269.56 | 86.48 | 53688 | 43 | 51 | 2434394369 | 2796715521 | 2589878017 | 2831042049 | 2816222721 | 3398969601 | 3163222273 | 2930145793 | 3181000961 | 3520160769 | 2958590465 | 2826093825 | 2985940737 | 3154718721 |
| Q91ZQ0 | Vmp1 | Vacuole membrane protein 1 Rattus norvegicus | 102.3 | 3.94 | 45901 | 2 | 2 | 1524172.88 | 1283855.75 | 1099408.88 | 1219152.3 | 969171.69 | 1380313.25 | 1207382.125 | 534966.5 | 1355243.5 | 1559005.25 | 1135989.75 | 1565218.12 | 1129400.25 | 713350.88 |
| Q91W86 | Vps11 | Vacuolar protein sorting-associated protein 11 homolog Mus musculus | 95.53 | 3.4 | 107719 | 2 | 2 | 682377.81 | 683623.4125 | 506056.03 | 893279.75 | 652780.06 | 1 | 1 | 1147448.25 | 1151959.62 | 1068924.38 | 1 | 701804.44 | 1 | 1 |
| Q8BX70 | Vps13c | Vacuolar protein sorting-associated protein 13C Mus musculus | 3355.131 | 24.09 | 420089 | 63 | 63 | 95545865 | 88624833 | 98818273 | 82741225 | 90174377 | 93392393 | 87466241 | 91216241 | 83861977 | 80464153 | 92013025 | 94544065 | 96459505 | 92988337 |
| Q920Q4 | Vps16 | Vacuolar protein sorting-associated protein 16 homolog Mus musculus | 296.13 | 12.28 | 94928 | 6 | 6 | 3724955.25 | 4112053 | 4163690 | 3637693.5 | 3905642.25 | 3273562.5 | 3363030 | 3969108 | 4091005 | 4111639.5 | 3385273.25 | 3612767 | 3814964.5 | 3371887.25 |
| Q8R307 | Vps18 | Vacuolar protein sorting-associated protein 18 homolog Mus musculus | 117.84 | 2.88 | 110219 | 2 | 2 | 887620.56 | 373677.84 | 234487.78 | 376040.69 | 380077.09 | 420651.91 | 514158.28 | 379227.19 | 463324.5 | 300456.16 | 500708.84 | 342573.03 | 281039.34 | 243384.38 |
| P0C0A1 | Vps25 | Vacuolar protein-sorting-associated protein 25 Rattus norvegicus | 167.23 | 21.02 | 20762 | 3 | 3 | 4261923 | 3802196 | 3971357 | 2041096.75 | 3977827.5 | 2154623 | 3341959.5 | 3212699 | 3204648 | 3254453.5 | 1867378.75 | 3866190.5 | 3789499.25 | 1687400.25 |
| P40336 | Vps26a | Vacuolar protein sorting-associated protein 26A Mus musculus | 519.69 | 39.14 | 38114 | 9 | 10 | 11493229 | 8537860 | 10521782 | 8121111.5 | 10240659 | 10460316 | 6252024 | 11096116 | 9552007 | 7899621 | 11557119 | 11045589 | 8446483 | 9170024 |
| Q8C0E2 | Vps26b | Vacuolar protein sorting-associated protein 26B Mus musculus | 117.51 | 9.23 | 39125 | 2 | 3 | 1498711.75 | 8649769 | 8341549 | 5296583.5 | 5104544.5 | 8806583 | 6095570.5 | 5807676 | 5591298.5 | 6454477 | 7528263 | 4257617 | 6248651 | 6606598 |
| B5DEN9 | Vps28 | Vacuolar protein sorting-associated protein 28 homolog Rattus norvegicus | 182.21 | 17.54 | 26110 | 3 | 3 | 1357463.88 | 1802175 | 1675546.88 | 2291588 | 2502938 | 2466213 | 2258172 | 2630574.25 | 2745090.5 | 2278885.75 | 2054939.75 | 2005801 | 2108404.75 | 2526789.5 |
| Q9QZ88 | Vps29 | Vacuolar protein sorting-associated protein 29 Mus musculus | 150.31 | 29.12 | 20496 | 4 | 4 | 5598654 | 5453934 | 4981585.5 | 5349165 | 5998336 | 5020969 | 5866305 | 4815387 | 5513347 | 7515364.5 | 6041502 | 5002860 | 6376829 | 6122622 |
| Q63615 | Vps33a | Vacuolar protein sorting-associated protein 33A Rattus norvegicus | 219.52 | 11.56 | 67514 | 3 | 3 | 1307783.5 | 1303169.62 | 1088316.25 | 1534171 | 1176797.75 | 1197269.25 | 1017876.25 | 1266752.62 | 1242758.25 | 1105970.62 | 1203415.75 | 1299385.38 | 1086371.12 | 909262.5 |
| Q9EQH3 | Vps35 | Vacuolar protein sorting-associated protein 35 Mus musculus | 1651.39 | 42.96 | 91713 | 27 | 27 | 77016625 | 67792897 | 79955761 | 60206765 | 65895489 | 72192585 | 61573857 | 78486057 | 78231793 | 60419245 | 66368925 | 69599385 | 71941265 | 63510157 |
| P0C0A2 | Vps36 | Vacuolar protein-sorting-associated protein 36 Rattus norvegicus | 251.35 | 15.8 | 43718 | 5 | 5 | 7320975.5 | 9481144 | 23444781 | 37198309 | 29594149 | 7140913 | 9445587 | 9004387 | 9316576 | 14500253 | 6017309 | 34983785 | 15717621 | 29379947 |
| Q8R105 | Vps37c | Vacuolar protein sorting-associated protein 37C Mus musculus | 92.67 | 9.09 | 38453 | 2 | 2 | 2901238.5 | 3828342.25 | 2644181.5 | 1177149.75 | 3265096 | 2577944 | 2216310.25 | 2673762 | 2198775.75 | 2574067 | 816855.81 | 1916799.5 | 2514676.25 | 2953169 |
| Q8R5L3 | Vps39 | Vam6/Vps39-like protein Mus musculus | 146.49 | 4.29 | 101693 | 3 | 3 | 1434395.62 | 1657410.75 | 991179.44 | 1260462.12 | 1247210.38 | 2273649.5 | 1657768.5 | 1535092 | 1735517.25 | 1650531.5 | 1185969.38 | 1014356.69 | 1971271.25 | 1928844 |
| Q5KU39 | Vps41 | Vacuolar protein sorting-associated protein 41 homolog Mus musculus | 114.77 | 5.28 | 98602 | 3 | 3 | 1764089.75 | 2877492 | 1975183 | 2186869 | 2298270.5 | 1895617.38 | 1867850 | 2461153.75 | 1888577.75 | 2088514.25 | 2022492.75 | 1702751.62 | 1872859.25 | 2016692.38 |
| P97390 | Vps45 | Vacuolar protein sorting-associated protein 45 Mus musculus | 201.05 | 11.05 | 65053 | 4 | 4 | 2015103 | 2130352.5 | 2321523 | 1991782.25 | 2260159 | 2057833.62 | 1920088.5 | 2510278 | 2111219.25 | 1927935.5 | 2090864.75 | 2092515 | 2105421.5 | 1787821.62 |
| Q793F9 | Vps4a | Vacuolar protein sorting-associated protein 4A Rattus norvegicus | 152.46 | 10.07 | 48907 | 1 | 4 | 3217811.5 | 2766266.25 | 2672727.5 | 2452150.75 | 3413551.5 | 2773369.5 | 2860691.25 | 2697065.25 | 3233291 | 3989467.5 | 2898587.25 | 3427220.75 | 3387395.25 | 3386904.25 |
| P46467 | Vps4b | Vacuolar protein sorting-associated protein 4B Mus musculus | 495.86 | 32.66 | 49419 | 6 | 9 | 7534549 | 8680039 | 6019704.5 | 6241002 | 7659982 | 6001018.5 | 8288249.5 | 8679353 | 7878314 | 9083419 | 7384723.5 | 6892010 | 8587820 | 7480720.5 |
| F1LSG8 | Vps50 | Syndetin Rattus norvegicus | 143.18 | 4.88 | 111188 | 3 | 3 | 1385889.5 | 1878717.5 | 1521972.62 | 1271391.25 | 1867205.25 | 1540538.5 | 1634738.75 | 1854682.75 | 1669943.38 | 1457997.5 | 1464295.38 | 1419002.25 | 1281874.38 | 1743676.25 |
| Q3UVL4 | Vps51 | Vacuolar protein sorting-associated protein 51 homolog Mus musculus | 329.73 | 12.02 | 86187 | 6 | 6 | 6871667 | 7778310.5 | 6939899 | 6797230 | 6580675 | 6280603 | 6998185.5 | 7282023.5 | 6867556 | 5801770 | 4702605.5 | 6611993 | 6193673 | 6547823 |
| O55166 | Vps52 | Vacuolar protein sorting-associated protein 52 homolog Rattus norvegicus | 370.9 | 15.35 | 82103 | 8 | 8 | 68504841 | 72353409 | 70649513 | 86977121 | 70330769 | 78680953 | 72687449 | 77376761 | 78353569 | 83182065 | 77080705 | 72253441 | 71965041 | 72772865 |
| Q8CCB4 | Vps53 | Vacuolar protein sorting-associated protein 53 homolog Mus musculus | 100.89 | 3.85 | 94423 | 2 | 2 | 999012 | 794355.5 | 1 | 1 | 1 | 1 | 662852.12 | 1 | 1 | 900605.25 | 1 | 916217.12 | 1 | 1 |
| Q80X41 | Vrk1 | Serine/threonine-protein kinase VRK1 Mus musculus | 289.35 | 19.32 | 49741 | 7 | 7 | 9471291 | 8664119 | 7830619 | 6107584 | 6991410.5 | 9353552 | 9666751 | 9974227 | 9211645 | 8511387 | 8646288 | 8250020.5 | 7641847 | 8640409 |
| Q8BN21 | Vrk2 | Serine/threonine-protein kinase VRK2 Mus musculus | 280.93 | 14.91 | 58119 | 5 | 5 | 5893602 | 4538886 | 3376686.5 | 4370146.5 | 4444076 | 2088243.5 | 3752075.75 | 4597515.5 | 2384800.5 | 4322700 | 4518232 | 2000677.88 | 2798519 | 3927058.25 |
| P29788 | Vtn | Vitronectin Mus musculus | 80.38 | 3.77 | 54849 | 1 | 1 | 690520.81 | 656204.25 | 617229.56 | 475170.97 | 465189.84 | 680352.81 | 1 | 934377.19 | 716505.69 | 1 | 482538.06 | 556626.5 | 424541.81 | 633243.06 |
| Q99KC8 | Vwa5a | von Willebrand factor A domain-containing protein 5A Mus musculus | 1849.75 | 49.81 | 87143 | 29 | 29 | 141498321 | 159484497 | 143030481 | 122763457 | 156414961 | 161655633 | 124029129 | 137212801 | 127880377 | 150506849 | 153155281 | 146056449 | 147374177 | 128949713 |
| Q8CC88 | Vwa8 | von Willebrand factor A domain-containing protein 8 Mus musculus | 137.71 | 1.84 | 213421 | 3 | 3 | 1 | 1 | 226946.95 | 169567.12 | 137882.53 | 147797.44 | 173544.89 | 163047.94 | 180604.41 | 180730.2 | 143763.27 | 179682.11 | 156338.83 | 130037.8 |
| Q65Z40 | Wapl | Wings apart-like protein homolog Mus musculus | 286.43 | 7.58 | 134070 | 5 | 5 | 1373054.38 | 1563977.25 | 1570771 | 1787935.88 | 1526152.38 | 1543065.62 | 873228.12 | 1357312.62 | 1250861.5 | 1030300.38 | 1788077.38 | 826465 | 2008068.25 | 1905443.88 |
| P32921 | Wars | Tryptophan--tRNA ligase, cytoplasmic Mus musculus | 620.24 | 36.8 | 54358 | 12 | 12 | 20430443 | 20311007 | 19161015 | 23268679 | 18740545 | 20339619 | 18370371 | 20959087 | 21158487 | 17985153 | 16221972 | 16010727 | 19139139 | 20133701 |
| P70315 | Was | Wiskott-Aldrich syndrome protein homolog Mus musculus | 336.07 | 13.65 | 54192 | 7 | 7 | 5165638.5 | 5253825 | 6108551.5 | 5458462.5 | 6272689 | 5992061 | 4690000 | 4232222 | 5046516.5 | 6158731 | 4732399.5 | 5344382 | 5038653.5 | 5737145.5 |
| Q8BH43 | Wasf2 | Wiskott-Aldrich syndrome protein family member 2 Mus musculus | 520.27 | 21.33 | 54074 | 9 | 9 | 7733303.5 | 8904244 | 9569390 | 7331849 | 11301141 | 11083330 | 9736062 | 11897482 | 11779608 | 11686500 | 8882168 | 10968333 | 12126567 | 12147282 |
| B2RYF7 | Washc1 | WASH complex subunit 1 Rattus norvegicus 2 | 232.58 | 15.37 | 51327 | 3 | 3 | 346658.91 | 438681.59 | 1 | 1 | 427266.47 | 300382.91 | 205592.72 | 312808.97 | 471364.19 | 571095.94 | 436459.72 | 431829.47 | 540074.25 | 316667.5 |
| Q6PGL7 | Washc2 | WASH complex subunit 2 Mus musculus | 490.49 | 11.92 | 145311 | 7 | 7 | 4654708.5 | 4389613 | 3335635.25 | 4487746.5 | 4130611.5 | 4431188 | 3472839.5 | 3820809 | 4245200.5 | 3660586 | 3361441 | 3470533 | 4329425.5 | 4134221.75 |
| Q9CR27 | Washc3 | WASH complex subunit 3 Mus musculus | 78.7 | 7.22 | 21093 | 2 | 2 | 1253019.5 | 1583527.12 | 1490122.38 | 1421575.38 | 1757485.62 | 1458151 | 1515257.88 | 965427.44 | 1437794.38 | 1212080.75 | 1604346.75 | 1311863.5 | 790912.56 | 1221490.12 |
| Q3UMB9 | Washc4 | WASH complex subunit 4 Mus musculus | 444.22 | 8.44 | 136370 | 7 | 7 | 6317588 | 4460619 | 6048707.5 | 3869675.75 | 5690689 | 6357270 | 4801745.5 | 5961374.5 | 6307241.5 | 3818266.5 | 3775924.5 | 6376078.5 | 5949272 | 5333964.5 |
| Q8C2E7 | Washc5 | WASH complex subunit 5 Mus musculus | 753.17 | 16.22 | 134110 | 13 | 13 | 15136523 | 15837017 | 13416941 | 14184977 | 13099698 | 14332101 | 14477274 | 15094037 | 13889816 | 14280867 | 13931172 | 13964516 | 14203856 | 15626388 |
| O08816 | Wasl | Neural Wiskott-Aldrich syndrome protein Rattus norvegicus | 38.63 | 4.59 | 54325 | 1 | 1 | 182711.09 | 1 | 1 | 1 | 221595.27 | 1 | 126786.23 | 181316.14 | 1 | 278909.78 | 222473.73 | 1 | 1 | 164800.41 |
| P59328 | Wdhd1 | WD repeat and HMG-box DNA-binding protein 1 Mus musculus | 158.15 | 4.03 | 124254 | 4 | 4 | 514364.81 | 1815379.12 | 523639 | 405847.94 | 554513.62 | 626113.44 | 536918.5 | 921900.06 | 443253.91 | 418593.06 | 1539088.88 | 1932931.88 | 915604.56 | 706939.44 |
| O88342 | Wdr1 | WD repeat-containing protein 1 Mus musculus | 1398.86 | 54.46 | 66407 | 22 | 22 | 92064257 | 82679289 | 89392897 | 99880233 | 90978177 | 87534065 | 81725249 | 83759441 | 92515697 | 101043945 | 90131033 | 94685873 | 86512473 | 94362265 |
| Q8K1X1 | Wdr11 | WD repeat-containing protein 11 Mus musculus | 375.65 | 9.32 | 135937 | 7 | 7 | 1746721.25 | 2654190.5 | 1646001.75 | 2812441 | 2961961.5 | 2684525 | 2436109.5 | 1481890.38 | 2809778 | 2805518.5 | 2254426 | 2734251 | 2666562 | 2512554 |
| Q9JJA4 | Wdr12 | Ribosome biogenesis protein WDR12 Mus musculus | 431.24 | 21.75 | 47347 | 6 | 6 | 3496997.75 | 1817594.12 | 2151153.25 | 2161436.5 | 1697828.5 | 2296013.5 | 2229907.75 | 3184277.75 | 3449342.25 | 1423258.5 | 2026698.25 | 2417991.5 | 2988987.75 | 2298498.25 |
| Q4VBE8 | Wdr18 | WD repeat-containing protein 18 Mus musculus | 108.02 | 9.05 | 47211 | 3 | 3 | 1 | 453416.16 | 309514.03 | 1 | 325002.22 | 1 | 1 | 1 | 1 | 1 | 245278.53 | 1 | 1 | 1 |
| Q8C6G8 | Wdr26 | WD repeat-containing protein 26 Mus musculus | 366.31 | 14.04 | 70543 | 6 | 6 | 3310922.25 | 2065627.88 | 3672164.5 | 3922655.5 | 4275283 | 3854480.75 | 4136032.75 | 2678405 | 2991205.5 | 4404277.5 | 4572262.5 | 2425315.5 | 3380961 | 3923656 |
| Q8BHB4 | Wdr3 | WD repeat-containing protein 3 Mus musculus | 417.79 | 10.62 | 105775 | 8 | 8 | 1887802.75 | 2171924.25 | 2491827.75 | 3237982 | 2894142.5 | 2329654.75 | 2722819 | 1907169.5 | 1955460.62 | 1138855.88 | 3183337.75 | 1486281.25 | 2229804.25 | 3305244.25 |
| Q6ZQL4 | Wdr43 | WD repeat-containing protein 43 Mus musculus | 536.53 | 21.57 | 75381 | 9 | 9 | 3795969 | 3502650.5 | 4467145 | 4847759.5 | 1830786.75 | 3907758 | 3215700.25 | 2786311.25 | 3671122 | 2134906.25 | 2858413 | 3484904.75 | 2627131.5 | 3531097.5 |
| Q6NVE8 | Wdr44 | WD repeat-containing protein 44 Mus musculus | 41.3 | 0.77 | 101555 | 1 | 1 | 3851419.75 | 3086079.75 | 2049291.12 | 3511173.75 | 4193344 | 2539551 | 2228665 | 2261155 | 1817998.38 | 1491871.88 | 4046317.5 | 3925296.25 | 2014360.88 | 3239171.5 |
| Q9Z0H1 | Wdr46 | WD repeat-containing protein 46 Mus musculus | 233.7 | 11.09 | 69048 | 4 | 4 | 6926345.5 | 3481912 | 699087.38 | 5402370 | 4405157 | 5209373 | 4342118 | 1733272.88 | 1169458.12 | 3584778 | 5886722 | 4461777.5 | 5152263 | 795889.94 |
| Q9ERF3 | Wdr61 | WD repeat-containing protein 61 Mus musculus | 345.45 | 28.52 | 33773 | 5 | 5 | 5545834.5 | 5051768 | 5856887.5 | 4464965 | 5519179 | 5482600 | 4738949 | 6915873.5 | 6980923 | 5552535 | 5775577.5 | 6494910.5 | 5002649 | 5499978 |
| Q920I9 | Wdr7 | WD repeat-containing protein 7 Mus musculus | 94.65 | 1.61 | 163450 | 2 | 2 | 1002719.735 | 1159235.5 | 904080.81 | 981449.44 | 966113.19 | 702574.1375 | 737201.62 | 725148.62 | 778086.69 | 569859.62 | 990913 | 956931.44 | 820038.81 | 880263.38 |
| Q8VCG3 | Wdr74 | WD repeat-containing protein 74 Mus musculus | 171.13 | 15.36 | 42636 | 4 | 4 | 1 | 1 | 1 | 914682.19 | 551223.12 | 479457.28 | 1 | 1 | 1 | 1 | 902347.25 | 1 | 1 | 1 |
| Q3U821 | Wdr75 | WD repeat-containing protein 75 Mus musculus | 276.24 | 7.83 | 94037 | 5 | 5 | 1112476.62 | 1787282.12 | 2255014 | 1392409.75 | 1587204.88 | 1334574.5 | 1445656.12 | 1481356.38 | 1161233 | 1941364.25 | 1573063.75 | 1575452 | 1977129.5 | 1841214.62 |
| Q99J09 | Wdr77 | Methylosome protein 50 Mus musculus | 227.72 | 19.88 | 36943 | 4 | 4 | 482799.38 | 1696376.75 | 1950942.75 | 1685725.88 | 1953306.25 | 1676533.75 | 1606704.12 | 456633.84 | 1566021.5 | 1560870 | 638799.75 | 1482777.5 | 626328.62 | 2344026 |
| Q8BFQ4 | Wdr82 | WD repeat-containing protein 82 Mus musculus | 122.95 | 10.22 | 35079 | 2 | 2 | 1241876.62 | 1899788.25 | 2435892.25 | 1747820.75 | 1870588.5 | 1800593.75 | 1837435.12 | 2333477 | 2165443.5 | 2384845.25 | 1855916.12 | 2167757.5 | 1778787.62 | 1816680.88 |
| B2RYI0 | Wdr91 | WD repeat-containing protein 91 Rattus norvegicus | 81.86 | 3.21 | 83146 | 2 | 2 | 943704.44 | 1 | 1218338.5 | 1 | 1 | 1 | 1 | 555281 | 1113445.88 | 973178.19 | 957052.23 | 951019.44 | 1023904.75 | 896232.5 |
| O88286 | Wiz | Protein Wiz Mus musculus | 43.91 | 0.89 | 184291 | 1 | 1 | 1078049.38 | 1476510.38 | 1311159 | 844011.31 | 1269574.75 | 508044.5 | 527435.56 | 1186802.25 | 1 | 1 | 686847.94 | 945573.25 | 918914.31 | 1230957.5 |
| P83741 | Wnk1 | Serine/threonine-protein kinase WNK1 Mus musculus | 553.13 | 6.52 | 250934 | 10 | 12 | 110027521 | 83453697 | 72778305 | 98426177 | 83955393 | 70849145 | 58174965 | 55978133 | 38741957 | 40960961 | 88183441 | 107508809 | 39366733 | 93689369 |
| Q3UH66 | Wnk2 | Serine/threonine-protein kinase WNK2 Mus musculus | 130.28 | 1.44 | 227527 | 1 | 3 | 4534244.5 | 5505072 | 4700283 | 5202611 | 5731587 | 5851336 | 5379042.5 | 5560287.5 | 5517743 | 5248860 | 5136000 | 5016362 | 5153581 | 5578270.5 |
| Q8BZZ3 | Wwp1 | NEDD4-like E3 ubiquitin-protein ligase WWP1 Mus musculus | 51.43 | 1.31 | 104694 | 1 | 1 | 1038385.44 | 968847.12 | 736336.5 | 636559.19 | 642218.94 | 1024110.75 | 967812.81 | 975853.75 | 781173.44 | 885953.5 | 916033.06 | 775920.06 | 737944 | 916865.25 |
| Q9DBH0 | Wwp2 | NEDD4-like E3 ubiquitin-protein ligase WWP2 Mus musculus | 138.69 | 5.17 | 98761 | 3 | 3 | 794372.06 | 844796.31 | 1055113.12 | 1053161.5 | 897406.12 | 794179.12 | 810663.19 | 1323056.38 | 1296288.25 | 744356.44 | 795751.94 | 835967.38 | 863044.12 | 1086827.25 |
| Q00519 | Xdh | Xanthine dehydrogenase/oxidase Mus musculus | 1074.64 | 22.32 | 146562 | 20 | 20 | 33818797 | 31264845 | 26359433 | 38434429 | 29072485 | 30386217 | 35948773 | 42730317 | 32915553 | 32444477 | 30033715 | 28637441 | 28074069 | 28153709 |
| Q6P1B1 | Xpnpep1 | Xaa-Pro aminopeptidase 1 Mus musculus | 787.32 | 36.92 | 69591 | 3 | 15 | 22986119 | 19389019 | 20236287 | 19305361 | 21173427 | 27443181 | 25499117 | 26730447 | 27163809 | 23611945 | 22811497 | 24545289 | 28329321 | 21817261 |
| B7ZMP1 | Xpnpep3 | Xaa-Pro aminopeptidase 3 Mus musculus | 67.36 | 3.36 | 56677 | 1 | 1 | 887883.5 | 1 | 836949.19 | 555046.56 | 692962.06 | 1 | 829332.06 | 825815.94 | 628556.38 | 1 | 1 | 1 | 770950.5 | 939258.88 |
| Q6P5F9 | Xpo1 | Exportin-1 Mus musculus | 1634.78 | 36.04 | 123093 | 31 | 31 | 67067733 | 63259401 | 52247017 | 53474477 | 65256521 | 73953825 | 64048061 | 73270657 | 74356089 | 64719885 | 66832165 | 62560857 | 69468897 | 68927481 |
| Q924C1 | Xpo5 | Exportin-5 Mus musculus | 817.62 | 14.62 | 136973 | 13 | 13 | 13229798 | 15525363 | 13483556 | 10659500 | 10972389 | 11702088 | 11716799 | 13263349 | 11978990 | 11926752 | 15281495 | 13227934 | 13589202 | 14637171 |
| Q9EPK7 | Xpo7 | Exportin-7 Mus musculus | 77.36 | 2.48 | 123810 | 2 | 2 | 763644.25 | 548928.75 | 446127.56 | 175356.64 | 436515.34 | 317504.94 | 451408.81 | 650142.12 | 581583.25 | 627997.94 | 509238.44 | 558073.25 | 554259.19 | 479923.06 |
| Q9CRT8 | Xpot | Exportin-T Mus musculus | 443.92 | 11.53 | 109734 | 9 | 9 | 3612543 | 2686021 | 5183394 | 3536804.5 | 3199093.75 | 7173642.5 | 4895072.5 | 6115287.5 | 5107600.5 | 3356514.25 | 2984050.5 | 5026308 | 4664757 | 2369214.5 |
| Q60596 | Xrcc1 | DNA repair protein XRCC1 Mus musculus | 100.73 | 4.75 | 68971 | 2 | 2 | 1 | 1 | 1 | 1 | 326521.94 | 1 | 365126.97 | 1 | 265994.31 | 1 | 230882.59 | 306806.59 | 306416.22 | 281368.4667 |
| P27641 | Xrcc5 | X-ray repair cross-complementing protein 5 Mus musculus | 265.8 | 8.47 | 83057 | 5 | 5 | 4528747 | 4249620 | 5734083 | 3648882.75 | 3654808 | 5142563 | 5043125 | 5901646 | 5969686 | 4885178 | 5067075.5 | 5800810.5 | 5426034 | 5086937.5 |
| P23475 | Xrcc6 | X-ray repair cross-complementing protein 6 Mus musculus | 82.76 | 4.77 | 69484 | 2 | 2 | 1266195 | 1761616.38 | 1318388.88 | 1495142 | 1732070.88 | 1564701.88 | 1328296.25 | 1756135.88 | 1531491.5 | 1428723.88 | 1787596.88 | 1607733.12 | 1666044.88 | 1646553.75 |
| P97789 | Xrn1 | 5'-3' exoribonuclease 1 Mus musculus | 63.96 | 0.99 | 194307 | 1 | 1 | 1 | 1 | 240862.66 | 1 | 207752.36 | 329733.06 | 246815.38 | 338300.59 | 264674.41 | 200326.83 | 242354.92 | 216120.2 | 223111.81 | 210425.17 |
| Q9DBR1 | Xrn2 | 5'-3' exoribonuclease 2 Mus musculus | 1106.68 | 27.87 | 108687 | 21 | 21 | 15612037 | 14278676 | 19510667 | 14982429 | 17092421 | 21686991 | 19275043 | 20932755 | 24596929 | 19800279 | 20272161 | 18728573 | 22759661 | 18884761 |
| Q91WQ3 | Yars | Tyrosine--tRNA ligase, cytoplasmic Mus musculus | 720.92 | 38.07 | 59105 | 16 | 16 | 28210869 | 21748569 | 27875767 | 28025875 | 24279579 | 28555731 | 28074267 | 30390861 | 30791929 | 28821605 | 28280917 | 31469807 | 26727817 | 27579871 |
| Q8BYL4 | Yars2 | Tyrosine--tRNA ligase, mitochondrial Mus musculus | 102.7 | 6.99 | 52597 | 2 | 2 | 449523.19 | 521184.19 | 537727.75 | 480602.28 | 523785.03 | 466235.38 | 950733.88 | 587034.44 | 820436.88 | 506500.22 | 646178.25 | 605433.94 | 973630.75 | 515705.25 |
| P62960 | Ybx1 | Nuclease-sensitive element-binding protein 1 Mus musculus | 806.76 | 49.07 | 35730 | 7 | 10 | 12411681 | 8943158 | 10193305 | 12765116 | 12786760 | 7080497 | 7872198.5 | 5933162.5 | 8493797 | 11040674 | 10806845 | 9333197 | 10449989 | 8061960.5 |
| Q9JKB3 | Ybx3 | Y-box-binding protein 3 Mus musculus | 414.99 | 26.87 | 38814 | 3 | 6 | 2490044.5 | 1521259.38 | 2400492.22 | 2960451.75 | 2630213.25 | 383667.94 | 453809 | 1 | 1 | 1 | 2128008.75 | 1014960.813 | 409303.44 | 507570.25 |

| | | | | | | | | | | | | | | | | | | | | | |
|---|---|---|---|---|---|---|---|---|---|---|---|---|---|---|---|---|---|---|---|---|---|
| Q5M7T4 | Yipf4 | Protein YIPF4 Rattus norvegicus | 38.77 | 4.47 | 27283 | 1 | 1 | 1563630.25 | 1 | 392467.5 | 438158.66 | 1 | 1 | 410081.94 | 1 | 1 | 1 | 1 | 1 | 1 | 1 |
| Q9CQW1 | Ykt6 | Synaptobrevin homolog YKT6 Mus musculus | 249.38 | 38.38 | 22314 | 6 | 6 | 4701246.5 | 5363432.5 | 5751318 | 3564797.5 | 4207009.5 | 5922344.5 | 4777925 | 4871907.5 | 5705460.5 | 4146314 | 3527412 | 6407400 | 5681614 | 4983951.5 |
| O88967 | Yme1l1 | ATP-dependent zinc metalloprotease YME1L1 Mus musculus | 165.06 | 8.81 | 80028 | 4 | 4 | 1 | 420507.09 | 493427 | 1 | 1 | 1020437 | 354886.28 | 264750.09 | 254567.92 | 240112.98 | 1 | 1 | 297667.03 | 1 |
| Q91YT7 | Ythdf2 | YTH domain-containing family protein 2 Mus musculus | 116.33 | 5.87 | 62280 | 2 | 3 | 1386122.75 | 836985.88 | 794061.81 | 935737.44 | 1361119.88 | 736798.56 | 1076041.75 | 634261.75 | 743322.25 | 796678.69 | 987328.81 | 565266.06 | 938025.19 | 870383.94 |
| Q8BYK6 | Ythdf3 | YTH domain-containing family protein 3 Mus musculus | 215.77 | 8.72 | 63961 | 3 | 4 | 1038374.69 | 836985.88 | 794061.81 | 935737.44 | 1263717.88 | 736798.56 | 782098.88 | 634261.75 | 631624.19 | 796678.69 | 987328.81 | 565266.06 | 938025.19 | 870383.94 |
| Q9CQV8 | Ywhab | 14-3-3 protein beta/alpha Mus musculus | 1360.37 | 73.17 | 28086 | 9 | 20 | 188957809 | 208764881 | 198768225 | 197832049 | 182322161 | 201382241 | 198530609 | 222865569 | 236501649 | 210273745 | 189015937 | 201554305 | 185968801 | 200691457 |
| P62259 | Ywhae | 14-3-3 protein epsilon Mus musculus | 1780.16 | 85.88 | 29174 | 26 | 30 | 469202977 | 506398337 | 465715905 | 483936257 | 451317441 | 493007169 | 499033729 | 589706689 | 537708161 | 505434337 | 466634817 | 472328225 | 439568833 | 485614977 |
| P61982 | Ywhag | 14-3-3 protein gamma Mus musculus | 1202.65 | 71.26 | 28303 | 11 | 19 | 212918113 | 242106865 | 219071857 | 219759761 | 206556465 | 214297249 | 223041249 | 250325617 | 258614113 | 232486545 | 215063057 | 230196673 | 208979521 | 221622881 |
| P68510 | Ywhah | 14-3-3 protein eta Mus musculus | 1279.82 | 60.98 | 28212 | 14 | 23 | 232024433 | 251203857 | 241545553 | 238275393 | 221811153 | 237288321 | 234267521 | 267853121 | 281964097 | 250635281 | 224578241 | 246065793 | 224226481 | 238988993 |
| P68254 | Ywhaq | 14-3-3 protein theta Mus musculus | 1359.98 | 73.47 | 27778 | 14 | 21 | 197620593 | 216977153 | 212601121 | 204379681 | 192531041 | 216283633 | 209891729 | 257932737 | 263455793 | 226779041 | 201703473 | 223162609 | 198159825 | 221101905 |
| P63101 | Ywhaz | 14-3-3 protein zeta/delta Mus musculus | 1487.54 | 78.37 | 27771 | 15 | 24 | 370861025 | 412231649 | 403401985 | 414648225 | 381436033 | 401318145 | 423837633 | 404556353 | 458584417 | 416333569 | 396035201 | 410867745 | 383921345 | 402008417 |
| P0C6P6 | Yy2 | Transcription factor YY2 Rattus norvegicus V=1 | 40.73 | 2.13 | 41805 | 1 | 1 | 670397.06 | 767132.5 | 757949.31 | 1078833.25 | 853574.81 | 1033083.69 | 1257061.38 | 832792.19 | 1167040.12 | 1133266.38 | 811917.75 | 1289149.62 | 729792.44 | 1059099.75 |
| Q8BGC4 | Zadh2 | Prostaglandin reductase-3 Mus musculus | 63.86 | 7.69 | 40529 | 1 | 1 | 227485.3 | 342538.455 | 262807.34 | 575417.56 | 304443.62 | 1 | 536424.38 | 368986.84 | 253385.66 | 1 | 351581.72 | 537223.94 | 466078.97 | 409343.78 |
| Q505B7 | Zbtb8os | Protein archease Mus musculus | 47.66 | 5.95 | 19727 | 1 | 1 | 1287060.12 | 1329972.12 | 1224755.25 | 1403056.38 | 1149620.25 | 1315067.12 | 1288932.38 | 1282312 | 1668301 | 1088225.62 | 1239144.583 | 1162206.88 | 1263885.12 | 1291341.75 |
| Q6NZF1 | Zc3h11a | Zinc finger CCCH domain-containing protein 11A Mus musculus | 217.47 | 8.33 | 86492 | 5 | 5 | 1478486.5 | 1571028 | 1751738.25 | 1529590 | 1550828.5 | 471800 | 1433966.25 | 1472847.5 | 1087586.38 | 1296602.88 | 981408.69 | 1301186.88 | 1336700.62 | 256232.75 |
| Q3TIV5 | Zc3h15 | Zinc finger CCCH domain-containing protein 15 Mus musculus | 202.4 | 9.62 | 48327 | 3 | 3 | 379547.72 | 478608.06 | 597786.88 | 451137.66 | 644188.06 | 1119383.5 | 1215531.38 | 569483.94 | 586552.94 | 742612.69 | 528832.56 | 1011990.75 | 675203.56 | 957459.62 |
| Q6ZPZ3 | Zc3h4 | Zinc finger CCCH domain-containing protein 4 Mus musculus | 56.5 | 1.23 | 140967 | 1 | 1 | 1 | 1 | 1 | 1 | 208738.22 | 196532.59 | 1 | 1 | 1 | 182484.17 | 206285.31 | 1 | 1 | 1 |
| Q3UPF5 | Zc3hav1 | Zinc finger CCCH-type antiviral protein 1 Mus musculus | 33.78 | 1.48 | 106688 | 1 | 1 | 899400.81 | 874464.06 | 819844.75 | 934696.69 | 805721.5 | 818975.12 | 856341.31 | 819374.69 | 816819.94 | 789172.5 | 845376.12 | 993172.06 | 843347.5 | 828135.69 |
| Q80YV2 | Zc3hc1 | Nuclear-interacting partner of ALK Mus musculus | 80.12 | 3.79 | 55196 | 1 | 1 | 192793.98 | 1 | 213104.2 | 964038.62 | 65215.49 | 156019.67 | 1 | 1 | 235977.25 | 1 | 143630.59 | 190355.27 | 1 | 1 |
| Q9CYA6 | Zcchc8 | Zinc finger CCHC domain-containing protein 8 Mus musculus | 171.88 | 7.48 | 78026 | 3 | 3 | 1134459.75 | 1824514.62 | 2568045 | 2077077.5 | 2022760.75 | 2068997.75 | 2093801.12 | 2318274.5 | 1445069.88 | 2061121.5 | 2224793.75 | 2676368.5 | 1672643 | 2130232.5 |
| Q9D270 | Zdhhc21 | Probable palmitoyltransferase ZDHHC21 Mus musculus | 44.93 | 3.02 | 31331 | 1 | 1 | 429161.62 | 221667.58 | 216614.39 | 305444.67 | 354335.09 | 379572.62 | 304936.97 | 290166.94 | 247421.23 | 302587.09 | 365863.06 | 323724.66 | 443922.84 | 399627.88 |
| Q9Z2U2 | Zfp292 | Zinc finger protein 292 Mus musculus | 34.66 | 0.41 | 301048 | 1 | 1 | 923951.88 | 2765312 | 2547395 | 1599058 | 1505235.12 | 1436604.5 | 1 | 1 | 954173.75 | 1 | 983321.62 | 1409616.75 | 1261138.62 | 1218025.663 |
| Q9DB43 | Zfpl1 | Zinc finger protein-like 1 Mus musculus | 43.99 | 3.87 | 34155 | 1 | 1 | 368699.84 | 422595.25 | 443284.66 | 287951.34 | 471469.69 | 452503.72 | 370249 | 482995.09 | 488256.59 | 349336.75 | 351652.44 | 311819.56 | 369955.47 | 383835.22 |
| O88532 | Zfr | Zinc finger RNA-binding protein Mus musculus | 109.18 | 2.79 | 116859 | 2 | 2 | 806900.19 | 1133684.25 | 1 | 1 | 1342708.88 | 1224180.12 | 1483970 | 1205507.12 | 1323913.88 | 1469574.38 | 984236.88 | 1171109.88 | 1340196.88 | 1001358.38 |
| Q3TXX3 | Zfyve27 | Protrudin Mus musculus | 53.67 | 3.37 | 46202 | 1 | 1 | 278513.91 | 1 | 330742.41 | 278902.81 | 1 | 315809.91 | 294692.72 | 325206.22 | 275835.53 | 302886.095 | 292024.8767 | 297525.72 | 289387.66 | 289161.25 |
| Q80W54 | Zmpste24 | CAAX prenyl protease 1 homolog Mus musculus | 40.98 | 2.32 | 54735 | 1 | 1 | 2383957.25 | 1678499.5 | 1610838.38 | 497427.59 | 1428656.25 | 1815056.62 | 1562086.5 | 2076714.62 | 1743502.215 | 1520151.12 | 1344661.38 | 1886670.12 | 1647894.88 | 2096466.12 |
| O88466 | Znf106 | Zinc finger protein 106 Mus musculus | 52.08 | 0.74 | 208964 | 1 | 1 | 1 | 168882.31 | 150078.77 | 1 | 204795.86 | 1 | 196285.12 | 122041.45 | 205635.75 | 234411.8 | 1 | 310877.72 | 1 | 237378.88 |
| Q9JMD0 | Znf207 | BUB3-interacting and GLEBS motif-containing protein ZNF207 Mus musculus | 167.13 | 6.67 | 52793 | 3 | 3 | 1021271.69 | 1589116.12 | 1309227 | 1383830.12 | 988423.06 | 703331.62 | 1554436.62 | 1342154.12 | 730863.5 | 1166550.12 | 1510744.25 | 1092045.75 | 879169.12 | 790897.31 |
| O88291 | Znf326 | DBIRD complex subunit ZNF326 Mus musculus | 87.14 | 2.76 | 65225 | 1 | 1 | 671484.12 | 1 | 434279.41 | 1 | 1 | 1 | 717675.56 | 1 | 553485.06 | 1 | 527380.75 | 1 | 581090.69 | 1 |
| Q80YR4 | Znf598 | E3 ubiquitin-protein ligase ZNF598 Mus musculus | 36.77 | 1.54 | 99192 | 1 | 1 | 669236.88 | 730088.88 | 643516.69 | 1071744.75 | 744059.88 | 1243052.38 | 698939.88 | 777561.56 | 624169.25 | 508385.47 | 758161.62 | 1136037.88 | 564538.94 | 468140.31 |
| Q91VY9 | Znf622 | Zinc finger protein 622 Mus musculus | 208.9 | 11.97 | 53451 | 4 | 4 | 1 | 293586.12 | 963740.31 | 971338.75 | 1 | 760980.62 | 778172.56 | 452410.06 | 407794.25 | 599839.3725 | 1 | 1 | 830780.38 | 1096387.12 |
| Q62384 | Zpr1 | Zinc finger protein ZPR1 Mus musculus | 87.28 | 4.58 | 50715 | 2 | 2 | 1537296.62 | 715885 | 1433982.25 | 958626.5 | 816974.56 | 1620875 | 1151471 | 1734091 | 1802881.5 | 1317984.75 | 1188093.12 | 1510182 | 1367824.75 | 1176009.25 |
| O35986 | Zranb2 | Zinc finger Ran-binding domain-containing protein 2 Rattus norvegicus | 55.42 | 5.45 | 37350 | 2 | 2 | 2275240 | 1912361.75 | 2048002.38 | 2818413.75 | 396779.72 | 2465725.25 | 2891881.25 | 2404159.75 | 2859773.25 | 3414188.75 | 2771167.75 | 3795330.5 | 3466864.25 | 3270741.75 |
| O54692 | Zw10 | Centromere/kinetochore protein zw10 homolog Mus musculus | 209.48 | 7.57 | 88063 | 4 | 4 | 1426157.75 | 666612 | 925775.31 | 897703.06 | 928143.38 | 1180961.12 | 716350.56 | 1549726 | 1474630.5 | 911446.88 | 566161.56 | 1008768.75 | 1344506 | 644933.44 |
| Q9CQU5 | Zwint | ZW10 interactor Mus musculus | 42.04 | 9.52 | 28713 | 1 | 1 | 315264.34 | 276110.19 | 346374.72 | 1 | 291008.81 | 669192.75 | 1 | 165337.89 | 486576.25 | 1 | 372074.53 | 237751.48 | 355467.94 | 297007.25 |
| Q62523 | Zyx | Zyxin Mus musculus | 239.13 | 14.36 | 60546 | 5 | 5 | 1166992.75 | 729891.44 | 1693928.548 | 2839388.5 | 2039441.5 | 2016547.62 | 2626640 | 2007039.12 | 3097314.75 | 3417549 | 2970702 | 3447523 | 1570061.25 | 2971150.25 |
| Q5SSH7 | Zzef1 | Zinc finger ZZ-type and EF-hand domain-containing protein 1 Mus musculus | 95.71 | 1.54 | 328312 | 2 | 2 | 178799.81 | 368272.22 | 280991.78 | 1 | 1 | 1 | 1 | 344286.59 | 116604.92 | 242810.55 | 446564.81 | 290450.5 | 143896.5 | 293637.27 |
