## Supplemental Table 2 for "Effects of polystyrene and polylactide nanoparticles on macrophages under a repeated exposure mode"

Supplementary table 2: List of modulated proteins in response to PS particles, selected by the Mann Whitney U test (p<0.05)

| accession | gene_name | description | otein_set_sco | coverage | MW | pecific_peptid | Ctrl1 rep | Ctrl5 rep | Ctrl4 rep | Ctrl3 rep | Ctrl2 rep | PLA1 rep | PLA5 rep | PLA4 rep | PLA3 rep | PLA2 rep | PS1 rep | PS5 rep | PS3 rep | PS2 rep | U-PLA | U-PS | ratio PLA | ratio PS |
|---|---|---|---|---|---|---|---|---|---|---|---|---|---|---|---|---|---|---|---|---|---|---|---|---|
| Q8QZT1 | Acat1 | Acetyl-CoA acetyltransferase, mitochondrial | 601.7 | 31.84 | 44816 | 10 | 21018019 | 20848719 | 21482667 | 24846375 | 24763999 | 21658727 | 25595085 | 22712881 | 25355313 | 27060283 | 26085471 | 27650245 | 27050899 | 25543849 | 4 | 0 | 1.08 | 1.18 |
| Q99KI0 | Aco2 | Aconitate hydratase, mitochondrial | 1715.58 | 40.38 | 85464 | 27 | 113500217 | 122642265 | 125108009 | 117830081 | 128722697 | 132302993 | 136039729 | 129805801 | 140362865 | 142224657 | 130724961 | 130182385 | 142546593 | 139967137 | 0 | 0 | 1.12 | 1.12 |
| Q91VJ1 | Aim2 | Interferon-inducible protein AIM2 | 222.28 | 12.71 | 40155 | 4 | 3277398 | 2867726 | 3099296.5 | 3292299 | 2809503 | 2882544 | 2594525.25 | 3237006.5 | 3271054.75 | 3294081.25 | 3405462.25 | 3544333 | 3350711.75 | 3847142.5 | 12 | 0 | 1.00 | 1.15 |
| Q9WTP6 | Ak2 | Adenylate kinase 2, mitochondrial | 315.83 | 33.89 | 26469 | 6 | 20322113 | 21748331 | 21522219 | 22517561 | 21268597 | 24824239 | 24086941 | 25325895 | 23625907 | 26267657 | 24645387 | 24001141 | 23193215 | 24466869 | 0 | 0 | 1.16 | 1.12 |
| Q66HF8 | Aldh1b1 | Aldehyde dehydrogenase X, mitochondrial | 111.29 | 4.82 | 57625 | 1 | 11603215 | 11684018 | 13051814 | 12502605 | 11462587 | 13215648 | 12968689 | 14213994 | 14635127 | 13888444 | 13951681 | 16082327 | 14318778 | 14370415 | 1 | 0 | 1.14 | 1.22 |
| P47738 | Aldh2 | Aldehyde dehydrogenase, mitochondrial | 1506.28 | 68.21 | 56538 | 24 | 182985393 | 204331585 | 209732513 | 200265169 | 210116641 | 223336913 | 224602097 | 222453585 | 231237777 | 223844209 | 247455457 | 248172561 | 250940065 | 254757489 | 0 | 0 | 1.12 | 1.24 |
| Q9DBF1 | Aldh7a1 | Alpha-aminoadipic semialdehyde dehydrogenase | 42.14 | 2.6 | 58861 | 1 | 959145.5 | 1088013.62 | 726082.81 | 815029.56 | 992774.5 | 1195092 | 1049457.88 | 1072075.25 | 844296.56 | 968668.5 | 1208569.75 | 1100472.5 | 1112100.62 | 1123956.38 | 7 | 0 | 1.12 | 1.24 |
| Q80V94 | Ap4e1 | AP-4 complex subunit epsilon-1 | 78.97 | 1.78 | 124845 | 1 | 681580.38 | 454765.38 | 548853.5175 | 674298.69 | 384769.62 | 1 | 395299.84 | 373866.31 | 1 | 1 | 1 | 331713.78 | 1 | 1 | 1 | 0 | 0.28 | 0.15 |
| Q9D0L7 | Armc10 | Armadillo repeat-containing protein 10 | 172.68 | 15.36 | 33311 | 3 | 1656080 | 1723679.12 | 1740246.5 | 1079363 | 1922319.38 | 1423307.75 | 1821133.12 | 2176469.5 | 1976578.75 | 1428418 | 2151719.75 | 2111165 | 2096870.75 | 2413043.25 | 9 | 0 | 1.09 | 1.35 |
| Q1AAU6 | Asap1 | Arf-GAP with SH3 domain, ANK repeat and PH domain-containing protein 1 | 459.92 | 9.53 | 127088 | 9 | 8820306 | 9953314 | 9387332 | 9091122 | 9597520 | 9608660 | 10004441 | 10351313 | 11853411 | 11547661 | 10488327 | 10368535 | 10301394 | 11688451 | 1 | 0 | 1.14 | 1.14 |
| Q61024 | Asns | Asparagine synthetase [glutamine-hydrolyzing] | 807.35 | 33.51 | 64283 | 14 | 22282757 | 16045791 | 22277965 | 22302875 | 22211675 | 34332637 | 24759677 | 23762539 | 22689153 | 25055077 | 23824953 | 28357385 | 23592073 | 24117823 | 0 | 0 | 1.24 | 1.19 |
| Q6PA06 | Atl2 | Atlastin-2 | 92.44 | 4.12 | 66224 | 2 | 733836.25 | 899531.75 | 923835.75 | 877421.19 | 803702.88 | 1116392.38 | 1110131 | 1190888.5 | 1285978.75 | 933306.44 | 1116237.38 | 1200585 | 1236324 | 1006278.12 | 0 | 0 | 1.33 | 1.34 |
| Q9EPE9 | Atp13a1 | Manganese-transporting ATPase 13A1 | 294.17 | 9 | 132388 | 6 | 3277617 | 3065685.75 | 4283905 | 3475658.5 | 4203536 | 3481029.75 | 3556447 | 2800960 | 3521475.5 | 4067172.75 | 5029892 | 4656637 | 4609959 | 4443279 | 12 | 0 | 0.95 | 1.28 |
| P97450 | Atp5pf | ATP synthase-coupling factor 6, mitochondrial | 111.26 | 17.59 | 12496 | 1 | 1423215.25 | 1208755.62 | 1075771.62 | 1035423.06 | 1000578.38 | 1055748.5 | 862907.75 | 905113.56 | 778138.56 | 1078898.62 | 813857.88 | 678042.12 | 811868.88 | 934275.62 | 5 | 0 | 0.81 | 0.70 |
| Q07813 | Bax | Apoptosis regulator BAX | 635.21 | 63.02 | 21395 | 9 | 33075229 | 33938641 | 33393683 | 25691449 | 33127145 | 35987489 | 35737021 | 35643941 | 35700893 | 32444009 | 37677029 | 34837793 | 36930033 | 37492449 | 4 | 0 | 1.10 | 1.15 |
| O08539 | Bin1 | Myc box-dependent-interacting protein 1 | 172.38 | 8.33 | 64470 | 2 | 1466632.25 | 1737278.88 | 1662260.75 | 1233827.25 | 1523102 | 1929133.5 | 1888381 | 1993632.88 | 2033967.75 | 2255730.25 | 3599544.75 | 3257452.75 | 1773316.38 | 1957549.25 | 0 | 0 | 1.33 | 1.74 |
| Q9CY64 | Blvra | Biliverdin reductase A | 753.23 | 58.64 | 33525 | 14 | 48784881 | 54583561 | 49228573 | 50434457 | 51295057 | 44310549 | 44582985 | 47738089 | 44683173 | 44107165 | 44423617 | 40271645 | 43850857 | 43421633 | 0 | 0 | 0.89 | 0.85 |
| Q64152 | Btf3 | Transcription factor BTF3 | 345.33 | 49.51 | 22031 | 6 | 8559690 | 3963968 | 4071065.75 | 5078866.5 | 3670445.75 | 3336583.5 | 7835755.5 | 3740741 | 3146954.25 | 4279316 | 3510721.75 | 3547773.25 | 3593206.75 | 3419285.25 | 8 | 0 | 0.88 | 0.69 |
| Q9WVA3 | Bub3 | Mitotic checkpoint protein BUB3 | 438.52 | 28.83 | 36955 | 7 | 16847627 | 19373095 | 17989345 | 18040573 | 16922063 | 14512527 | 17121833 | 16098568 | 15281613 | 17037767 | 14124080 | 15379591 | 14832119 | 15379073 | 4 | 0 | 0.90 | 0.84 |
| Q80X80 | C2cd2l | Phospholipid transfer protein C2CD2L | 49.59 | 1.42 | 76329 | 1 | 1 | 1 | 349341.56 | 1 | 1 | 505218.97 | 495357.28 | 569019.19 | 693052.94 | 364224.81 | 446286.72 | 497738.44 | 511167.75 | 485064.3033 | 0 | 0 | 7.52 | 6.94 |
| Q9D6N1 | Ca13 | Carbonic anhydrase 13 | 57.94 | 3.44 | 29522 | 1 | 2512639.25 | 3071649 | 2929762.75 | 2469199.5 | 2384382.75 | 2982757.5 | 2600762.5 | 2909995 | 2568370 | 2396833.25 | 2347055.25 | 2332430 | 2273190.75 | 2154185.75 | 11 | 0 | 1.01 | 0.85 |
| Q6PHZ2 | Camk2d | Calcium/calmodulin-dependent protein kinase type II subunit delta | 490.44 | 26.45 | 56369 | 9 | 15552478 | 16226495 | 16876161 | 17452865 | 16448511 | 21893837 | 19909357 | 21355475 | 21208995 | 21389053 | 19836729 | 19635099 | 19042353 | 19131423 | 0 | 0 | 1.28 | 1.18 |
| P47753 | Capza1 | F-actin-capping protein subunit alpha-1 | 528 | 45.1 | 32940 | 8 | 59613385 | 60653597 | 55100237 | 56731757 | 63947313 | 68557569 | 65010953 | 66143533 | 67203049 | 62702825 | 66975649 | 65080177 | 64062505 | 67503489 | 1 | 0 | 1.11 | 1.11 |
| P29452 | Casp1 | Caspase-1 | 890.3 | 41.29 | 45640 | 16 | 26517867 | 27996515 | 28144805 | 26615815 | 29227735 | 22356923 | 23036571 | 23089263 | 23713299 | 22791975 | 26056177 | 24088509 | 22483423 | 26143101 | 0 | 0 | 0.83 | 0.89 |
| Q8VEH6 | Cbwd1 | COBW domain-containing protein 1 | 132.73 | 9.16 | 43772 | 2 | 1518167.38 | 361844.88 | 1746521.25 | 1475068.88 | 1537275.12 | 1960452.38 | 1745335.5 | 2236414.5 | 2241467.25 | 1859763.25 | 1796521.5 | 2317916.5 | 1876033 | 1950675.62 | 1 | 0 | 1.51 | 1.50 |
| P10810 | Cd14 | Monocyte differentiation antigen CD14 | 919.15 | 51.37 | 39204 | 13 | 46938701 | 51451925 | 49639101 | 46202521 | 44246909 | 37399081 | 39845889 | 41435497 | 33770573 | 39076653 | 33665017 | 33582077 | 38490937 | 30320721 | 0 | 0 | 0.80 | 0.71 |
| Q792H5 | Celf2 | CUGBP Elav-like family member 2 | 180.57 | 7.68 | 54271 | 2 | 957990.44 | 317687.5 | 1052966.25 | 816434.69 | 648280.06 | 1099133.5 | 1011289.62 | 963850.69 | 973417.94 | 1046881.25 | 1211629.167 | 1179579.5 | 1379732.62 | 1075575.38 | 4 | 0 | 1.34 | 1.60 |
| Q8BTU1 | Cfap20 | Cilia- and flagella-associated protein 20 | 173.2 | 30.05 | 22748 | 4 | 7629927 | 8499859 | 8663983 | 7950704 | 6956509.5 | 6192437 | 5840535 | 6917020.5 | 4997719 | 5935685 | 4541339 | 6196100 | 4910781 | 5646966.5 | 0 | 0 | 0.75 | 0.67 |
| P18760 | Cfl1 | Cofilin-1 | 813.63 | 74.7 | 18560 | 12 | 337236609 | 319565633 | 351056481 | 363054849 | 393487073 | 415494209 | 392304641 | 329385633 | 406457281 | 452801537 | 442737121 | 434238977 | 460522241 | 457483873 | 5 | 0 | 1.13 | 1.27 |
| B0K020 | Cisd1 | CDGSH iron-sulfur domain-containing protein 1 | 36.33 | 8.33 | 12097 | 1 | 1284328.875 | 1110574 | 1254126.12 | 1242726.5 | 1529888.88 | 1696193.62 | 1189677.25 | 728460.94 | 1362649.75 | 1890749.5 | 1725112 | 1707670.12 | 1733234.88 | 1801308.25 | 10 | 0 | 1.07 | 1.36 |
| Q9QYB1 | Clic4 | Chloride intracellular channel protein 4 | 1131.61 | 74.31 | 28729 | 17 | 97964481 | 99478145 | 87956129 | 100850505 | 100354753 | 120676057 | 119147137 | 113144737 | 129646161 | 139642721 | 116781297 | 118015865 | 113190041 | 127598233 | 0 | 0 | 1.28 | 1.22 |
| O08585 | Clta | Clathrin light chain A | 123.19 | 10.64 | 25604 | 3 | 22239505 | 23305617 | 20524421 | 27492887 | 23438983 | 19110209 | 19091705 | 19940957 | 16651577 | 19197859 | 20148757 | 17940843 | 18731121 | 19499133 | 0 | 0 | 0.80 | 0.82 |
| Q3UM29 | Cog7 | Conserved oligomeric Golgi complex subunit 7 | 39.89 | 1.3 | 86073 | 1 | 877942.88 | 1048416.69 | 953760.94 | 772125.38 | 767377.81 | 623994.62 | 1106338.5 | 692372.56 | 660758.62 | 799329.5 | 755083.12 | 687895.31 | 700532.94 | 664636.44 | 7 | 0 | 0.88 | 0.79 |
| Q8K4Q8 | Colec12 | Collectin-12 | 136.96 | 4.72 | 81304 | 2 | 1318988.75 | 1573271.25 | 1555674 | 927166.12 | 1725151.75 | 1588737.38 | 1729193.25 | 1288371.815 | 748772.75 | 1086783.88 | 523929.69 | 689163.8333 | 720038.06 | 823523.75 | 11 | 0 | 0.91 | 0.49 |
| Q8BXC6 | Commd2 | COMM domain-containing protein 2 | 74.28 | 9.55 | 22848 | 2 | 2578403.25 | 2830499.25 | 2162274.75 | 2516614.75 | 2383693 | 2175381.5 | 2183641.25 | 2800303.25 | 2028409.12 | 2099681 | 1967882 | 2133353.25 | 2020462.88 | 2087530.5 | 6 | 0 | 0.91 | 0.82 |
| Q6AY19 | Coq8b | Atypical kinase COQ8B, mitochondrial | 34.75 | 1.89 | 58905 | 1 | 3658946.25 | 2610860.5 | 3624262.25 | 2453319 | 2308136 | 2670578.5 | 2206137 | 3805195 | 3090057.25 | 2122850.25 | 2273137.75 | 2297229 | 2243443.25 | 2025130.12 | 11 | 0 | 0.95 | 0.75 |
| P56394 | Cox17 | Cytochrome c oxidase copper chaperone | 73.75 | 25.4 | 6784 | 1 | 853887.655 | 675136.25 | 976623.56 | 849466.06 | 914324.75 | 1243552.5 | 1305873.25 | 704204 | 1423785.25 | 1726207.12 | 1119831.88 | 1323507.62 | 1517114 | 1120257.5 | 4 | 0 | 1.50 | 1.49 |

| | | | | | | | | | | | | | | | | | | | | | | | | |
|---|---|---|---|---|---|---|---|---|---|---|---|---|---|---|---|---|---|---|---|---|---|---|---|---|
| P19783 | Cox4i1 | Cytochrome c oxidase subunit 4 isoform 1, mitochondrial | 215.7 | 30.77 | 19530 | 4 | 30285453 | 33025669 | 27630631 | 26359977 | 29911717 | 28602201 | 27487479 | 25874075 | 25248589 | 29015169 | 25853013 | 22462841 | 25196947 | 23308119 | 5 | 0 | 0.93 | 0.82 |
| P11240 | Cox5a | Cytochrome c oxidase subunit 5A, mitochondrial | 288.69 | 38.36 | 16130 | 6 | 40878529 | 47620297 | 40817437 | 37513889 | 38771337 | 32348581 | 35937237 | 40773493 | 33111355 | 28772209 | 31308093 | 29085567 | 31354793 | 34658793 | 2 | 0 | 0.83 | 0.77 |
| P19536 | Cox5b | Cytochrome c oxidase subunit 5B, mitochondrial | 53.61 | 9.38 | 13813 | 1 | 7728957.5 | 8080394.5 | 9121866 | 9044157 | 8073188.5 | 5919199.5 | 7450418 | 6304691 | 6546995 | 8021108 | 5163765.5 | 6167710.5 | 4713191.5 | 5723266.5 | 1 | 0 | 0.81 | 0.65 |
| P56391 | Cox6b1 | Cytochrome c oxidase subunit 6B1 | 43.9 | 16.28 | 10071 | 1 | 2872200.25 | 4056435 | 2684032.25 | 2218423 | 3453565.75 | 2030394.88 | 2620808.75 | 2981121.25 | 2234880.25 | 2236088.25 | 1640230.5 | 1700160.71 | 1498464.25 | 1961787.38 | 6 | 0 | 0.79 | 0.56 |
| Q9CPQ1 | Cox6c | Cytochrome c oxidase subunit 6C | 67.57 | 19.74 | 8469 | 2 | 5674408 | 3656424 | 3796538 | 4099267.5 | 3276160.75 | 3148131.5 | 3580100.25 | 3345972.25 | 2841076.75 | 3019356.25 | 2281812.5 | 2360568.5 | 2293236.25 | 2253690 | 2 | 0 | 0.78 | 0.56 |
| P48771 | Cox7a2 | Cytochrome c oxidase subunit 7A2, mitochondrial | 55.83 | 12.05 | 9291 | 1 | 2024143.12 | 2778264.75 | 2558337.5 | 2066369.38 | 1934297.38 | 1397679 | 2069152 | 2100055.5 | 1573706.88 | 1520218.75 | 325632.38 | 695491.81 | 1112682 | 711268.73 | 6 | 0 | 0.76 | 0.31 |
| Q9WVJ3 | Cpq | Carboxypeptidase Q | 159.27 | 9.79 | 51813 | 3 | 4235704.5 | 5433789.5 | 3963128 | 4633763.5 | 4864550 | 2740996.75 | 3432032.25 | 3105766.25 | 3189807.75 | 2950120 | 3112566.25 | 3491155.25 | 2712508.75 | 3580672 | 0 | 0 | 0.67 | 0.70 |
| Q5XI29 | Cpsf7 | Cleavage and polyadenylation specificity factor subunit 7 | 210.07 | 13.42 | 51073 | 5 | 3155922 | 3872699 | 3390567.5 | 3668620.5 | 3314346.5 | 3158044 | 3179262.5 | 3882158.5 | 2417373.25 | 2974243.5 | 2800432.5 | 2946060.5 | 2737779 | 2988918 | 7 | 0 | 0.90 | 0.82 |
| P47199 | Cryz | Quinone oxidoreductase | 44.42 | 6.65 | 35269 | 1 | 1067594.25 | 1308112.12 | 1488117.25 | 1360542.88 | 1056024.75 | 1830280.38 | 1295732.88 | 1537155.12 | 1916072.12 | 1893861.75 | 1614763.62 | 1761491.75 | 2126802.25 | 1628377.25 | 3 | 0 | 1.35 | 1.42 |
| Q62426 | Cstb | Cystatin-B | 145.46 | 57.14 | 11046 | 3 | 7521179 | 7137837.5 | 6904083 | 12055766 | 6465746.5 | 3031240.75 | 5190731 | 2957673 | 4015037.5 | 4353343.5 | 4867321 | 3217625.75 | 5260377 | 2877489 | 0 | 0 | 0.49 | 0.51 |
| Q8VCN5 | Cth | Cystathionine gamma-lyase | 688.3 | 43.47 | 43567 | 13 | 41636849 | 40732541 | 49172553 | 51631009 | 46581893 | 48540305 | 42209745 | 44339353 | 42522381 | 43343765 | 35893937 | 36933165 | 36345461 | 37593289 | 11 | 0 | 0.96 | 0.80 |
| P97821 | Ctsc | Dipeptidyl peptidase 1 | 595.33 | 16.67 | 52376 | 7 | 49376273 | 52577253 | 45175817 | 45652129 | 52748553 | 36184393 | 32687601 | 37102473 | 32423235 | 31497689 | 37962265 | 34302401 | 38151173 | 39475241 | 0 | 0 | 0.69 | 0.76 |
| P18242 | Ctsd | Cathepsin D | 1308.71 | 65.37 | 44954 | 21 | 406401761 | 489922369 | 425776257 | 393210113 | 412787073 | 566899457 | 570130689 | 552964993 | 524020897 | 533450113 | 536090305 | 506827169 | 585553345 | 525133889 | 0 | 0 | 1.29 | 1.26 |
| Q3U308 | Ctu2 | Cytoplasmic tRNA 2-thiolation protein 2 | 54.55 | 3.7 | 56105 | 1 | 1 | 1 | 1 | 1 | 270970.25 | 1 | 1 | 1 | 305560.69 | 267004.19 | 336719.03 | 361082.0833 | 327399.53 | 419127.69 | 10 | 0 | 2.11 | 6.66 |
| Q9WTX6 | Cul1 | Cullin-1 | 508.56 | 14.69 | 89692 | 10 | 9356674 | 8197263 | 7475768 | 8358356 | 10032447 | 11403163 | 8733838 | 12266343 | 11508169 | 10748202 | 10499824 | 12952264 | 12832699 | 11439313 | 2 | 0 | 1.26 | 1.37 |
| P61804 | Dad1 | Dolichyl-diphosphooligosaccharide--protein glycosyltransferase subunit DAD1 | 166.75 | 28.32 | 12497 | 3 | 13215215 | 13330451 | 11556073 | 10955833 | 15067723 | 14099778 | 11961920 | 10981207 | 12339314 | 13018405 | 15633830 | 15490589 | 15116859 | 17593443 | 11 | 0 | 0.97 | 1.24 |
| Q91VR5 | Ddx1 | ATP-dependent RNA helicase DDX1 | 1054.07 | 34.32 | 82500 | 19 | 21494465 | 23311527 | 25019387 | 23063183 | 23753523 | 23760623 | 26720961 | 26121527 | 24885585 | 26870247 | 25130829 | 25252313 | 26245641 | 25898643 | 2 | 0 | 1.10 | 1.10 |
| Q62095 | Ddx3y | ATP-dependent RNA helicase DDX3Y | 1104.92 | 32.98 | 73428 | 2 | 77379209 | 69522481 | 72049065 | 60994129 | 71025609 | 60714649 | 64453085 | 58073865 | 58504413 | 62334405 | 53894289 | 56623925 | 60489421 | 59040217 | 2 | 0 | 0.87 | 0.82 |
| O08560 | Dgkz | Diacylglycerol kinase zeta | 71.91 | 4.41 | 104009 | 2 | 1274328.25 | 1207170.12 | 1073381.62 | 1266015.88 | 1295138.75 | 1424261.38 | 1702466.12 | 1422879.62 | 1570516.88 | 1423209.62 | 1703564.5 | 1523170.38 | 1465322.62 | 1469373.12 | 0 | 0 | 1.23 | 1.26 |
| D4A2Z8 | Dhx36 | ATP-dependent DNA/RNA helicase DHX36 | 261.69 | 7.4 | 113843 | 6 | 1602403.38 | 2228548.25 | 2125017.75 | 1845307 | 3894404.25 | 1977454.38 | 1425083 | 1334479.25 | 1297270.75 | 1280549.5 | 1438098.62 | 1360321.75 | 1207351.75 | 1171439.25 | 2 | 0 | 0.63 | 0.55 |
| P54103 | Dnajc2 | DnaJ homolog subfamily C member 2 | 218.77 | 10.79 | 71722 | 5 | 2841132.5 | 2349083.75 | 3566257 | 3432205 | 3433569.5 | 3923222.5 | 4275470.5 | 3619431.5 | 2410579.75 | 3387510 | 4247780 | 4833312 | 4023677.5 | 4711293.5 | 7 | 0 | 1.13 | 1.43 |
| O35075 | Dscr3 | Down syndrome critical region protein 3 homolog | 181.98 | 18.86 | 32970 | 4 | 3226952.5 | 2074669 | 2602682 | 3277601 | 2343939.5 | 3699260 | 3700280 | 3005929.5 | 3198500.75 | 3557376.5 | 3515577 | 3599492.25 | 3444214.25 | 3758079 | 4 | 0 | 1.27 | 1.32 |
| Q9R0P5 | Dstn | Destrin | 489.98 | 52.12 | 18522 | 7 | 29927785 | 31798485 | 33788209 | 31869721 | 32292155 | 36284041 | 34494133 | 34565989 | 34277933 | 40759389 | 39374597 | 35627793 | 34732637 | 44021397 | 0 | 0 | 1.13 | 1.20 |
| Q9D903 | Ebna1bp2 | Probable rRNA-processing protein EBP2E | 179.36 | 17.97 | 34703 | 4 | 1567059.38 | 2113400.5 | 1280972 | 2690933 | 1567930.5 | 2198458.5 | 1892652.25 | 1779422.88 | 1996623.5 | 2265153 | 4261450 | 3059703.5 | 4068897.5 | 5568977.5 | 8 | 0 | 1.10 | 2.30 |
| Q3TGW2 | Eepd1 | Endonuclease/ exonuclease/ phosphatase family domain-containing protein 1 | 41.23 | 2.64 | 62953 | 1 | 884563.31 | 910401.94 | 1 | 646673 | 1 | 1015498.62 | 1052696.62 | 1135702.12 | 1087941.25 | 932121.44 | 1332034.5 | 1077017.25 | 957574.06 | 1058873.5 | 0 | 0 | 2.14 | 2.27 |
| P59325 | Eif5 | Eukaryotic translation initiation factor 5 | 290.59 | 13.29 | 48968 | 5 | 10179051 | 8915562 | 9307121 | 9392106 | 10261104 | 10624993 | 10484176 | 10996545 | 10274364 | 10722081 | 10795490 | 10582504 | 11922213 | 11142316 | 0 | 0 | 1.11 | 1.16 |
| P83940 | Eloc | Elongin-C | 207.44 | 30.36 | 12473 | 2 | 3720203.5 | 3612083.5 | 3997336 | 4232823 | 4573965 | 5138137.5 | 4283929.5 | 3193797.75 | 5126163 | 6214968 | 5172096 | 4905423 | 5666234 | 4760511.5 | 6 | 0 | 1.19 | 1.27 |
| Q3UMY5 | Eml4 | Echinoderm microtubule-associated protein-like 4 | 82.13 | 1.62 | 110027 | 1 | 1 | 1 | 1 | 1 | 1 | 733450.0475 | 834772.44 | 603441.25 | 692215.75 | 803370.75 | 600970.06 | 620397.03 | 799164.06 | 461056.97 | 0 | 0 | 733450.05 | 620397.03 |
| Q8BTJ4 | Enpp4 | Bis(5'-adenosyl)-triphosphatase enpp4 | 99.91 | 4.82 | 51611 | 2 | 985076.69 | 1300517.25 | 1556378.12 | 1119300 | 1032830.94 | 802762.88 | 979898.94 | 1244291 | 883281.62 | 971595.62 | 653590.19 | 660817.12 | 971313.44 | 843465.81 | 3 | 0 | 0.81 | 0.65 |
| Q61545 | Ewsr1 | RNA-binding protein EWS | 292.15 | 12.21 | 68462 | 6 | 9790247 | 11386937 | 9336300 | 11701854 | 11464452 | 8151007 | 13421140 | 9046718 | 10774652 | 12048937 | 5276147 | 8332456 | 8746660 | 8851677 | 12 | 0 | 1.00 | 0.73 |
| Q05816 | Fabp5 | Fatty acid-binding protein 5 | 1005.52 | 79.26 | 15137 | 16 | 367597665 | 362926977 | 390861889 | 341166817 | 309307073 | 348621505 | 401450913 | 399667425 | 376506625 | 378119297 | 395330977 | 398830881 | 434537025 | 457692673 | 5 | 0 | 1.07 | 1.19 |
| Q8R1F1 | Fam129b | Niban-like protein 1 | 513.98 | 19.36 | 84819 | 9 | 21481605 | 20756937 | 21639297 | 18641639 | 18686853 | 24123223 | 23228745 | 24436909 | 23561257 | 21288829 | 22190621 | 23111867 | 22226109 | 22595651 | 2 | 0 | 1.15 | 1.11 |
| P62862 | Fau | 40S ribosomal protein S30 | 94.78 | 18.64 | 6648 | 2 | 2374680.75 | 2146666.25 | 4830429 | 3816281.25 | 4727740 | 6375753 | 4694967 | 1848487.12 | 5831644 | 8275803 | 6668771 | 8601862 | 8358926.5 | 7226102 | 7 | 0 | 1.51 | 2.16 |
| P26151 | Fcgr1 | High affinity immunoglobulin gamma Fc receptor I | 275.57 | 22.77 | 44888 | 6 | 10544918 | 9299983 | 11888508 | 9437379 | 9035087 | 7037936 | 6074016 | 6070993.5 | 7317976 | 5974343 | 6363764 | 6071877.5 | 7110427 | 4110293.75 | 0 | 0 | 0.65 | 0.59 |
| P08101 | Fcgr2 | Low affinity immunoglobulin gamma Fc region receptor II | 176.94 | 11.21 | 36695 | 2 | 10047039 | 11144037 | 9987515 | 11168614 | 10237842 | 9170796 | 10371663 | 8652388 | 7745995 | 10653992 | 9368193 | 7853520 | 9759479 | 8793518 | 6 | 0 | 0.89 | 0.85 |
| A0A0B4 | Fcgr4 | Low affinity immunoglobulin gamma Fc region receptor IV | 45.23 | 4.02 | 28398 | 1 | 2495273 | 2113805.5 | 2117607.5 | 2473083 | 2382982.25 | 1894300.5 | 1841483.38 | 2000441.75 | 1741781.12 | 2120373.5 | 1889272.12 | 1783561.25 | 1607984.75 | 1968628.62 | 2 | 0 | 0.83 | 0.78 |

| | | | | | | | | | | | | | | | | | | | | | | | | |
|---|---|---|---|---|---|---|---|---|---|---|---|---|---|---|---|---|---|---|---|---|---|---|---|---|
| Q8BGW1 | Fto | Alpha-ketoglutarate-dependent dioxygenase FTO | 118.81 | 6.57 | 58007 | 2 | 1 | 1 | 1 | 225302.11 | 1 | 197345.09 | 1 | 1 | 192606.64 | 179165.58 | 265976.12 | 327578.03 | 239117.73 | 277557.2933 | 9 | 0 | 2.53 | 6.16 |
| Q8R2K1 | Fuom | Fucose mutarotase | 126.66 | 17.65 | 16805 | 2 | 2262723.25 | 1758226.5 | 1265212.5 | 2424451 | 1251164.12 | 1359057 | 2084125.25 | 2033296.38 | 1414952.5 | 925365 | 1229686.75 | 1155378.75 | 859052.25 | 999137.94 | 10 | 0 | 0.87 | 0.59 |
| Q00612 | G6pdx | Glucose-6-phosphate 1-dehydrogenase X | 1454.99 | 55.73 | 59263 | 27 | 97138265 | 93448481 | 95200425 | 96327689 | 96271081 | 100245673 | 98799889 | 95811273 | 95528729 | 108365089 | 102213569 | 111241657 | 105843505 | 98231041 | 6 | 0 | 1.04 | 1.09 |
| Q61107 | Gbp4 | Guanylate-binding protein 4 | 569.72 | 26.13 | 70801 | 11 | 8514483 | 7450088 | 5133596 | 6443512 | 6430460 | 3197550.5 | 4450390.5 | 3241343.75 | 2997324.5 | 2882868.25 | 3562685 | 2570989.5 | 3680811.25 | 3870607.75 | 0 | 0 | 0.49 | 0.50 |
| O88986 | Gcat | 2-amino-3-ketobutyrate coenzyme A ligase, mitochondrial | 461.88 | 32.45 | 44931 | 7 | 2071480 | 2891570.25 | 3612637.25 | 2995040 | 3543770.25 | 4552562 | 3529727 | 2241942.75 | 3769953.5 | 4263833 | 3987094.5 | 4190853.5 | 4462710 | 4411221 | 6 | 0 | 1.21 | 1.41 |
| P97494 | Gclc | Glutamate--cysteine ligase catalytic subunit | 391.57 | 16.01 | 72571 | 6 | 1696302.5 | 3405893.5 | 3341330.25 | 3184514.5 | 3020486.75 | 3533322.75 | 3039812.5 | 4288761 | 2531285.5 | 3766027 | 4282653.5 | 4255555 | 3807138 | 3437513 | 7 | 0 | 1.17 | 1.35 |
| Q8BX17 | Gemin5 | Gem-associated protein 5 | 193.13 | 4.53 | 166592 | 5 | 3853023 | 3625824.5 | 3026013.5 | 2197521.5 | 2166582.25 | 1310570.88 | 1604851.12 | 2008812.38 | 1628851.62 | 1231911.75 | 1028084.69 | 710402 | 1677707.25 | 940815.56 | 0 | 0 | 0.52 | 0.37 |
| Q9CWG1 | Glipr1 | Glioma pathogenesis-related protein 1 Mus musculus | 61.17 | 5.49 | 29129 | 1 | 1234218.25 | 1205148.25 | 1037347 | 1332321.75 | 1437550.5 | 1610373.62 | 1616885.88 | 1726157.62 | 1560643.75 | 1267817.25 | 1606324.75 | 1614322.5 | 1579347.5 | 1447292.12 | 2 | 0 | 1.25 | 1.25 |
| Q9JHJ3 | Glmp | Glycosylated lysosomal membrane protein | 115.5 | 8.17 | 43804 | 2 | 390392.56 | 375639.5 | 895242.81 | 521804.69 | 836234.56 | 1268463.12 | 620702.69 | 364765.09 | 256151.8 | 1277358.5 | 1246621.5 | 1066663.75 | 1269519.25 | 1135898.12 | 12 | 0 | 1.25 | 1.95 |
| P63213 | Gng2 | Guanine nucleotide-binding protein G(I)/G(S)/G(O) subunit gamma-2 | 83.93 | 39.44 | 7850 | 2 | 6892465 | 7611842 | 7428047 | 8924962 | 8298338.5 | 5637436 | 6061224.5 | 6645098 | 5762194 | 5795699 | 6408208 | 5934865 | 5218212 | 6679262.5 | 0 | 0 | 0.76 | 0.77 |
| Q64521 | Gpd2 | Glycerol-3-phosphate dehydrogenase, mitochondrial | 536.62 | 22.28 | 80954 | 12 | 12265822 | 14680938 | 14065567 | 13584454 | 12169963 | 15677707 | 15148026 | 13589635 | 16696129 | 13955227 | 16333337 | 14846157 | 16393345 | 15033927 | 4 | 0 | 1.12 | 1.17 |
| Q99P91 | Gpnmb | Transmembrane glycoprotein NMB | 299.24 | 10.45 | 63676 | 6 | 15079978 | 18625543 | 14805663 | 15678684 | 18425977 | 28292055 | 26069569 | 21835903 | 21971033 | 23690629 | 24751743 | 21765275 | 28238209 | 24268953 | 0 | 0 | 1.48 | 1.50 |
| P26817 | Grk2 | Beta-adrenergic receptor kinase 1 | 607.39 | 22.06 | 79785 | 13 | 6677560 | 5539719 | 6658146 | 4423462.5 | 6012117 | 6890418 | 5496088.5 | 6438050.5 | 6938860.5 | 6413163 | 7361141 | 7357792 | 7052446.5 | 6780686.5 | 8 | 0 | 1.10 | 1.22 |
| Q8BMS1 | Hadha | Trifunctional enzyme subunit alpha, mitochondrial | 1110.22 | 38.66 | 82670 | 19 | 28916769 | 24112179 | 27524993 | 26538753 | 25760241 | 28232151 | 29585693 | 29986777 | 30998355 | 24772967 | 31427533 | 32221645 | 33350231 | 31124985 | 5 | 0 | 1.08 | 1.21 |
| Q99JY0 | Hadhb | Trifunctional enzyme subunit beta, mitochondrial | 447.59 | 21.26 | 51386 | 9 | 18308545 | 19479009 | 20810149 | 20087425 | 20413189 | 18963227 | 22021683 | 18546233 | 21062735 | 22372113 | 27326747 | 25261597 | 23451767 | 23743363 | 8 | 0 | 1.04 | 1.26 |
| Q69ZS7 | Hbs1l | HBS1-like protein | 76.01 | 3.67 | 75100 | 2 | 1 | 1 | 289044.81 | 283889.75 | 273839.38 | 325917.09 | 372204.06 | 237434.31 | 324246.3425 | 361429.91 | 388208.06 | 438782.38 | 341281.25 | 307390.81 | 3 | 0 | 1.91 | 2.18 |
| A1EC95 | Heatr6 | HEAT repeat-containing protein 6 | 82.91 | 2.16 | 136505 | 2 | 577690.69 | 396189.25 | 377223.16 | 446741.7975 | 435864.09 | 352418.34 | 363479.575 | 422473.09 | 369836.62 | 309190.25 | 312233.9 | 362360.72 | 237844.64 | 336496.34 | 2 | 0 | 0.81 | 0.70 |
| Q8VH49 | Higd1a | HIG1 domain family member 1A, mitochondrial | 147.95 | 43.01 | 10301 | 2 | 813655.38 | 698692.56 | 782062.25 | 630587.57 | 227940.09 | 264727.19 | 816548.88 | 544879.38 | 808258.88 | 831105.56 | 1162508.75 | 1433592.38 | 1245085.25 | 1104648.75 | 9 | 0 | 1.04 | 1.96 |
| P10853 | Hist1h2bf | Histone H2B type 1-F/J/L | 1153.51 | 55.56 | 13936 | 3 | 2311764993 | 2088850177 | 1691605505 | 1780771073 | 1772783873 | 1521681665 | 1719559297 | 1880200961 | 1436857089 | 1671866241 | 1596670337 | 1497583489 | 1423864065 | 1678264833 | 4 | 0 | 0.85 | 0.80 |
| P10854 | Hist1h2bm | Histone H2B type 1-M | 1109.6 | 55.56 | 13936 | 2 | 2307947009 | 2087647233 | 1690756609 | 1778428161 | 1768388353 | 1521673217 | 1718052097 | 1880057857 | 1436131329 | 1670938881 | 1597435905 | 1497685249 | 1424740993 | 1677980161 | 4 | 0 | 0.85 | 0.80 |
| Q3TRM8 | Hk3 | Hexokinase-3 | 2099.18 | 46.85 | 100101 | 14 | 111634473 | 134472689 | 119574729 | 111517449 | 118247505 | 113794553 | 107501257 | 111184673 | 103951961 | 107363473 | 108582865 | 94170505 | 105660569 | 109304409 | 2 | 0 | 0.91 | 0.88 |
| Q9D8V0 | Hm13 | Minor histocompatibility antigen H13 | 278.34 | 16.4 | 41748 | 6 | 17344729 | 18118161 | 16397292 | 14235864 | 17362193 | 21622459 | 19333771 | 21930545 | 19732055 | 19179163 | 20518239 | 19863279 | 22094347 | 21011995 | 0 | 0 | 1.22 | 1.25 |
| Q8BTX9 | Hsdl1 | Inactive hydroxysteroid dehydrogenase-like protein 1 | 138.33 | 10.91 | 36868 | 2 | 1532226.12 | 1417680.75 | 1644205.25 | 1829637.88 | 1761696.75 | 1051358.75 | 1358895.12 | 1245302.25 | 1625902 | 1276974.12 | 1263502.12 | 1304692.12 | 1262580.62 | 1349341.75 | 2 | 0 | 0.80 | 0.79 |
| P07901 | Hsp90aa1 | Heat shock protein HSP 90-alpha | 2808.68 | 62.62 | 84788 | 30 | 416174593 | 466700545 | 429501857 | 418725409 | 478100257 | 485711521 | 447464545 | 465466945 | 439408193 | 479584993 | 485990913 | 498430721 | 507168833 | 517039393 | 6 | 0 | 1.05 | 1.14 |
| P11499 | Hsp90ab1 | Heat shock protein HSP 90-beta | 4068.121 | 76.38 | 83281 | 43 | 858858113 | 907080065 | 860898369 | 795965633 | 908185857 | 934317825 | 867120065 | 913728897 | 873618945 | 890316225 | 922130049 | 965213697 | 959466689 | 958405569 | 6 | 0 | 1.03 | 1.10 |
| P08113 | Hsp90b1 | Endoplasmin | 3099.08 | 61.97 | 92476 | 44 | 438865441 | 435713089 | 428330785 | 391069985 | 457535489 | 490388065 | 421803585 | 515039329 | 459849313 | 447182273 | 464496897 | 497617281 | 489239169 | 481065057 | 5 | 0 | 1.08 | 1.12 |
| P26772 | Hspe1 | 10 kDa heat shock protein, mitochondrial | 215.66 | 42.16 | 10902 | 4 | 18476301 | 19934239 | 19126895 | 21050439 | 16966677 | 12317349 | 15474625 | 14810171 | 10708773 | 15614746 | 15044636 | 14590793 | 15932658 | 15271881 | 0 | 0 | 0.72 | 0.80 |
| Q9JIY5 | Htra2 | Serine protease HTRA2, mitochondrial | 220.09 | 17.69 | 49348 | 4 | 747603.06 | 685581.5 | 941821.56 | 487902.75 | 834827.5 | 997479.31 | 474083 | 794950.75 | 1100611.88 | 607628.81 | 2001827.75 | 1800737.5 | 1692637.5 | 1097997.75 | 11 | 0 | 1.07 | 2.23 |
| P03975 | Iap | IgE-binding proteinV=1 | 590.01 | 23.16 | 62747 | 11 | 21785451 | 19728165 | 25046163 | 23513657 | 20108189 | 31279901 | 34253705 | 32022245 | 31745995 | 30166807 | 32709733 | 34841109 | 32528817 | 34805005 | 0 | 0 | 1.45 | 1.53 |
| P54071 | Idh2 | Isocitrate dehydrogenase [NADP], mitochondrial | 1170.59 | 48.67 | 50906 | 20 | 72911833 | 74086593 | 74553665 | 72508329 | 70330505 | 71828065 | 79011729 | 83732601 | 78756425 | 79949545 | 82717289 | 76308801 | 78930425 | 83983257 | 4 | 0 | 1.08 | 1.10 |
| P41565 | Idh3g | Isocitrate dehydrogenase [NAD] subunit gamma 1, mitochondrial | 253.8 | 14.76 | 42851 | 6 | 16545307 | 13982288 | 16847583 | 16369147 | 13910443 | 18911789 | 15494213 | 15909703 | 17160641 | 18949001 | 19599161 | 17961009 | 21329011 | 17529899 | 6 | 0 | 1.11 | 1.23 |
| Q9R002 | Ifi202 | Interferon-activable protein 202 | 144.04 | 6.29 | 50466 | 2 | 2326020.25 | 2754549.75 | 2657673.25 | 1744767.5 | 1884176.88 | 1 | 1 | 1 | 1 | 1 | 1 | 1 | 1 | 1 | 0 | 0 | 0.00 | 0.00 |
| P0DOV2 | Ifi204 | Interferon-activable protein 204 | 675.02 | 27.46 | 69438 | 8 | 18012943 | 19144417 | 19639325 | 15222543 | 14087636 | 10929171 | 11615319 | 12017299 | 11915774 | 9637524 | 9370458 | 9949678 | 11449507 | 8969819 | 0 | 0 | 0.65 | 0.58 |
| Q8BV66 | Ifi44 | Interferon-induced protein 44 | 78.42 | 5.92 | 47852 | 2 | 6552790.5 | 6363975 | 5733456 | 3897284.25 | 5024364 | 2039624.12 | 2116467.25 | 1792802.5 | 2103463.25 | 2083649.25 | 1789207.75 | 1374940.88 | 461732.31 | 1208626.98 | 0 | 0 | 0.37 | 0.22 |
| Q9BDB7 | Ifi44l | Interferon-induced protein 44-like2 | 231.32 | 12.98 | 49680 | 4 | 2844332.75 | 2146209.25 | 2052692.5 | 1848826.62 | 1618923.62 | 842147.125 | 1044910.31 | 669528.31 | 942336.44 | 711813.44 | 737215.38 | 424957.41 | 802695.25 | 167065.08 | 0 | 0 | 0.40 | 0.25 |

| | | | | | | | | | | | | | | | | | | | | | | | | |
|---|---|---|---|---|---|---|---|---|---|---|---|---|---|---|---|---|---|---|---|---|---|---|---|---|
| Q8R4K2 | Irak4 | Interleukin-1 receptor-associated kinase 4 | 74.58 | 3.7 | 50872 | 1 | 1060406.25 | 1101917 | 1002067.94 | 1146826.38 | 1106412.25 | 922905.25 | 875769.12 | 996127 | 1027434 | 850466.44 | 930253.19 | 901446.62 | 803519.38 | 893944.81 | 1 | 0 | 0.86 | 0.81 |
| Q60766 | Irgm1 | Immunity-related GTPase family M protein 1 | 382.54 | 19.07 | 46552 | 7 | 6445768.5 | 4892167.5 | 4781422.5 | 3869175 | 4389789 | 2883047.75 | 2765580 | 2933151.25 | 3284595 | 2833628 | 2599356.5 | 2450496.5 | 3855157.5 | 2993222.25 | 0 | 0 | 0.60 | 0.61 |
| P85094 | Isoc2a | Isochorismatase domain-containing protein 2A | 237.74 | 40.78 | 22417 | 4 | 3103277.25 | 2942515 | 2963688.25 | 2998752.25 | 3032113.75 | 3567827.75 | 3256465.25 | 2750156 | 4007995 | 3503977.25 | 4070244.75 | 3735360.25 | 4640719 | 4501481 | 5 | 0 | 1.14 | 1.41 |
| O70309 | Itgb5 | Integrin beta-5 | 72.13 | 2.88 | 87909 | 2 | 1 | 1 | 1 | 1 | 1 | 68096.88 | 104749.095 | 153777.06 | 127744.94 | 69377.5 | 91404.76 | 131483.73 | 82495.11 | 101794.5333 | 0 | 0 | 104749.10 | 101794.53 |
| P70227 | Itpr3 | Inositol 1,4,5-trisphosphate receptor type 3 | 131.55 | 1.91 | 304275 | 4 | 639147.94 | 1556119.5 | 1574866.25 | 1596880.12 | 1718204 | 2475911.75 | 1699047 | 1880152 | 2103020.75 | 2218357.5 | 2138167.5 | 2033202 | 1825938 | 1802469.75 | 1 | 0 | 1.46 | 1.38 |
| P97825 | Jpt1 | Jupiter microtubule associated homolog 1 | 106.39 | 27.27 | 16081 | 2 | 2924789.25 | 4087840.5 | 3059245.5 | 4315181 | 3478096 | 1926713.38 | 1266849.12 | 2953416.75 | 1909610.25 | 2426835.25 | 2296861.25 | 1162097.38 | 1083107.12 | 2803540.5 | 1 | 0 | 0.59 | 0.51 |
| Q569A6 | Kdelr1 | ER lumen protein-retaining receptor 1 2 | 97.28 | 10.38 | 24546 | 2 | 1144506.25 | 213300.92 | 162754.42 | 189118.08 | 427419.9175 | 1 | 102192.34 | 158620.48 | 1 | 1 | 1 | 1 | 138202.08 | 147157.95 | 0 | 0 | 0.12 | 0.17 |
| Q6GV12 | Kdsr | 3-ketodihydrosphingosine reductase | 65.45 | 4.22 | 35956 | 1 | 1429564 | 1541790.62 | 1477920.38 | 1612317.12 | 1591729.62 | 1734343.12 | 1274162.62 | 1570917.88 | 1353793.88 | 1483304.375 | 1 | 1327692.12 | 1417061.38 | 1 | 10 | 0 | 0.97 | 0.45 |
| Q922S8 | Kif2c | Kinesin-like protein KIF2C | 95.48 | 4.72 | 81085 | 2 | 242709.95 | 1 | 275403.44 | 1 | 402230.97 | 345746.72 | 667487.75 | 865196.25 | 668064.5 | 636623.805 | 1032694.56 | 1090669.5 | 972195.62 | 532133.69 | 1 | 0 | 3.46 | 4.93 |
| Q4KM31 | Limd2 | LIM domain-containing protein 2 | 61.9 | 13.28 | 14236 | 1 | 5087711 | 5415087 | 5407738 | 5114712 | 5454524 | 3992986.5 | 5148498 | 4087321.25 | 4910629.5 | 5214855 | 3680553 | 4044729.25 | 4217600 | 3957633 | 4 | 0 | 0.88 | 0.75 |
| Q9Z0M5 | Lipa | Lysosomal acid lipase/cholesteryl ester hydrolase | 683.19 | 22.17 | 45325 | 9 | 14897863 | 18597939 | 18244205 | 13282737 | 12877767 | 27890647 | 30345103 | 34120141 | 34454009 | 32353231 | 40208761 | 55868197 | 56020901 | 45081977 | 0 | 0 | 2.04 | 3.16 |
| P70202 | Lxn | Latexin | 92.6 | 9.46 | 25492 | 2 | 4335441.5 | 5021072 | 4927896 | 4757538.5 | 4461408.5 | 3813518 | 4037597.75 | 4750108.5 | 4314252.5 | 3536690 | 3864358.75 | 3692705.25 | 3778510 | 3973488.75 | 2 | 0 | 0.87 | 0.81 |
| Q9QYL8 | Lypla2 | Acyl-protein thioesterase 2 | 40.87 | 10.82 | 24807 | 1 | 875456.56 | 942451.81 | 740642.75 | 577799.25 | 760094.81 | 987983.12 | 1483220.62 | 2577308.75 | 2390149.25 | 1009287.06 | 2236100.75 | 1277549.75 | 1083784.25 | 1011178.44 | 0 | 0 | 2.17 | 1.80 |
| Q9QXZ0 | Macf1 | Microtubule-actin cross-linking factor 1 | 1182.11 | 4.98 | 831878 | 26 | 6852718.5 | 8346064 | 8966120 | 7128024 | 6447014 | 9448019 | 9302008 | 10202712 | 12248702 | 9357708 | 10193320 | 11638077 | 9482751 | 10385722 | 0 | 0 | 1.34 | 1.38 |
| O09159 | Man2b1 | Lysosomal alpha-mannosidase | 636.26 | 14.61 | 114648 | 10 | 29760767 | 29134727 | 24821189 | 32504589 | 31066853 | 22524719 | 22186813 | 22868383 | 23590495 | 20176759 | 24421871 | 22808655 | 21914893 | 23430521 | 0 | 0 | 0.76 | 0.79 |
| P21708 | Mapk3 | Mitogen-activated protein kinase 3 | 285.39 | 25.53 | 43081 | 3 | 14046033 | 13884697 | 13244656 | 12927679 | 14114616 | 17948245 | 14339667 | 15353954 | 16576580 | 16775449 | 16691656 | 15289569 | 16162203 | 16747251 | 0 | 0 | 1.19 | 1.19 |
| Q9D2X5 | Mau2 | MAU2 chromatid cohesion factor homolog | 45.49 | 1.45 | 69579 | 1 | 741195.69 | 486170.12 | 475540.56 | 651269.19 | 563693.38 | 533056.69 | 489969.97 | 666078.62 | 469639.19 | 436321.16 | 474913.75 | 408895.25 | 427851.97 | 420013.66 | 8 | 0 | 0.89 | 0.74 |
| Q61881 | Mcm7 | DNA replication licensing factor MCM7 | 1286.23 | 44.37 | 81211 | 22 | 33899465 | 33035453 | 32675285 | 34860397 | 31904891 | 27325927 | 29044059 | 30638405 | 30152625 | 26885287 | 28408079 | 27961453 | 26078545 | 25678947 | 0 | 0 | 0.87 | 0.81 |
| Q8BP48 | Metap1 | Methionine aminopeptidase 1 | 271.74 | 17.1 | 43221 | 5 | 6111351 | 6590463 | 6052901 | 6348026 | 6844733.5 | 7778479.5 | 8297604 | 6086174 | 7142611.5 | 6733561 | 7627016 | 7982255 | 7282722.5 | 7888992 | 5 | 0 | 1.13 | 1.20 |
| D0QMC3 | Mndal | Myeloid cell nuclear differentiation antigen-like protein | 513.11 | 22.68 | 60526 | 6 | 21679509 | 23776097 | 24524045 | 21039025 | 22592671 | 16122662 | 13932207 | 17640987 | 17385345 | 14221786 | 14060852 | 14942461 | 15294745 | 16304416 | 0 | 0 | 0.70 | 0.67 |
| Q9D0G0 | Mrps30 | 28S ribosomal protein S30, mitochondrial | 214.45 | 16.52 | 49939 | 4 | 1011809.5 | 96723.46 | 1106397.88 | 1256107.75 | 1277453.75 | 2397141.5 | 1991002.5 | 833092.12 | 1413076 | 2188909.25 | 1494909.12 | 1697879 | 2480267.75 | 1823879.25 | 4 | 0 | 1.86 | 1.97 |
| Q791T5 | Mtch1 | Mitochondrial carrier homolog 1 | 140.74 | 15.42 | 41565 | 4 | 272478.12 | 1 | 1 | 1 | 1 | 468563.88 | 341604.41 | 1 | 1 | 286233.16 | 539887.38 | 519330.41 | 437482.22 | 280412.81 | 6 | 0 | 4.02 | 8.15 |
| P00405 | Mtco2 | Cytochrome c oxidase subunit 2 | 271.03 | 26.43 | 25976 | 5 | 66056129 | 80171209 | 69072009 | 67014545 | 75373865 | 73883257 | 70129417 | 66272065 | 67682065 | 69854697 | 41636253 | 43376381 | 51372705 | 50640613 | 12 | 0 | 0.97 | 0.65 |
| Q8BVN4 | Mterf4 | Transcription termination factor 4, mitochondrial2 | 35.3 | 2.02 | 40249 | 1 | 4459182.5 | 4166899.75 | 4839473.5 | 4991558.5 | 4571420.5 | 4913611.5 | 4242239.5 | 4348033.5 | 4798275 | 5061476 | 6260129 | 5393287 | 5279134.5 | 5790976 | 11 | 0 | 1.01 | 1.23 |
| Q9JK81 | Myg1 | UPF0160 protein MYG1, mitochondrial | 541.26 | 36.32 | 42723 | 11 | 17665041 | 19092641 | 16112001 | 19558717 | 18558771 | 14437871 | 15265776 | 15087716 | 12738415 | 14976250 | 13547084 | 14803579 | 13206613 | 14359070 | 0 | 0 | 0.80 | 0.77 |
| O70145 | Ncf2 | Neutrophil cytosol factor 2 | 1100.57 | 50.48 | 59485 | 21 | 69790921 | 61107409 | 54740737 | 56351425 | 58903945 | 53135781 | 53183137 | 57343985 | 49652681 | 52358545 | 52031457 | 46979217 | 49328633 | 51797241 | 2 | 0 | 0.88 | 0.83 |
| Q5XIA1 | Ncln | Nicalin | 196.73 | 7.28 | 62992 | 4 | 5398326.5 | 5630011.5 | 5604782 | 4915900.5 | 7233276.5 | 5395313 | 5894605.5 | 4233968 | 4957850 | 4875217.5 | 4694031 | 3742528.5 | 4228253.5 | 4378723 | 6 | 0 | 0.88 | 0.74 |
| P57716 | Ncstn | Nicastrin | 874.54 | 25.99 | 78492 | 13 | 66070841 | 61829253 | 59012321 | 56884313 | 61278481 | 63410585 | 57526361 | 59194677 | 50824225 | 51005569 | 49550433 | 53796573 | 55036769 | 55373305 | 7 | 0 | 0.92 | 0.88 |
| Q62425 | Ndufa4 | Cytochrome c oxidase subunit NDUFA4 | 180.38 | 68.29 | 9327 | 4 | 24869155 | 20107309 | 17811985 | 16011653 | 17451605 | 13934563 | 13859357 | 14726675 | 15424429 | 12746563 | 11183773 | 6177599 | 10366876 | 11151371 | 0 | 0 | 0.73 | 0.50 |
| O09111 | Ndufb11 | NADH dehydrogenase [ubiquinone] 1 beta subcomplex subunit 11, mitochondrial | 37.18 | 11.26 | 17444 | 1 | 1 | 1 | 1 | 1 | 835302.44 | 1 | 1 | 586911.38 | 1 | 1123373.62 | 2479955.25 | 1712495.833 | 1463830.25 | 1193702 | 10 | 0 | 2.05 | 10.25 |
| Q9DC70 | Ndufs7 | NADH dehydrogenase [ubiquinone] iron-sulfur protein 7, mitochondrial | 360.71 | 30.8 | 24683 | 6 | 9863442 | 10096206 | 9947674 | 9556809 | 11060647 | 10410752 | 10491083 | 9492803 | 11016054 | 11690093 | 11537952 | 12172173 | 12314664 | 11484388 | 8 | 0 | 1.05 | 1.18 |
| Q8K3J1 | Ndufs8 | NADH dehydrogenase [ubiquinone] iron-sulfur protein 8, mitochondrial | 221.95 | 17.92 | 24038 | 3 | 1898490.5 | 2204512 | 1916932.75 | 1833128.62 | 2253961.75 | 2535078 | 2059565.5 | 2602214.5 | 2425363 | 2143960.25 | 2481266.5 | 2556399.5 | 2826770.75 | 2442564.5 | 4 | 0 | 1.16 | 1.27 |
| O35309 | Nmi | N-myc-interactor | 100.72 | 9.55 | 35236 | 2 | 2499665.5 | 2235807.25 | 2187643.75 | 2451689.75 | 2832923.5 | 1924270.12 | 1708475.75 | 2000299.75 | 1985911.62 | 2119268.5 | 1853734.75 | 1856813.88 | 1985596.25 | 2076469.75 | 0 | 0 | 0.80 | 0.80 |
| Q8BHG1 | Nrdc | Nardilysin | 374.9 | 10.51 | 132891 | 9 | 10176996 | 8305207 | 7807701.5 | 7826306.5 | 7628328 | 9184277 | 8397776 | 8553708 | 8506037 | 6806395 | 7148031.5 | 7291352.5 | 7414464 | 5565442.5 | 9 | 0 | 0.99 | 0.82 |
| Q8VI94 | Oasl1 | 2'-5'-oligoadenylate synthase-like protein 1 | 173.91 | 10.37 | 59088 | 4 | 902312 | 652955.69 | 787475.69 | 421891.06 | 521197.62 | 1 | 1 | 1 | 807015.75 | 1 | 1 | 105680.22 | 90020.33 | 1 | 4 | 0 | 0.25 | 0.07 |
| P29758 | Oat | Ornithine aminotransferase, mitochondrial | 1084.71 | 50.11 | 48355 | 16 | 54754557 | 48940553 | 49442789 | 54602601 | 45663557 | 52761209 | 58228573 | 60343269 | 58608221 | 55362069 | 62513061 | 64798829 | 64568749 | 55856461 | 2 | 0 | 1.13 | 1.22 |

| | | | | | | | | | | | | | | | | | | | | | | | | |
|---|---|---|---|---|---|---|---|---|---|---|---|---|---|---|---|---|---|---|---|---|---|---|---|---|
| Q9D0K2 | Oxct1 | Succinyl-CoA:3-ketoacid coenzyme A transferase 1, mitochondrial | 1174.91 | 43.65 | 55989 | 15 | 38869481 | 50020709 | 50447477 | 51833617 | 52585401 | 50145901 | 57669321 | 50684585 | 56898329 | 61044753 | 61466453 | 64137069 | 67769793 | 56002905 | 5 | 0 | 1.13 | 1.28 |
| Q9DCL9 | Paics | Multifunctional protein ADE2 | 1061.37 | 48.71 | 47006 | 18 | 18406545 | 19401191 | 24465001 | 22549427 | 24582029 | 22667763 | 20918139 | 19451783 | 20334237 | 22140409 | 26830319 | 26148601 | 25532079 | 26142395 | 11 | 0 | 0.96 | 1.20 |
| Q61036 | Pak3 | Serine/threonine-protein kinase PAK 3 | 343.94 | 15.56 | 62398 | 2 | 3598529.5 | 3919888.25 | 3638312.25 | 3730203 | 3620463.5 | 3811791.5 | 3193423 | 3155009.75 | 2610567.5 | 2838928.25 | 3397802.5 | 3394999.5 | 3328516.25 | 3149755 | 4 | 0 | 0.84 | 0.90 |
| P27773 | Pdia3 | Protein disulfide-isomerase A3 | 1767.9 | 60.2 | 56678 | 30 | 254216289 | 217690513 | 247481441 | 249726273 | 236587809 | 246538001 | 266653649 | 251374081 | 263613041 | 273817761 | 255023905 | 256135217 | 266005025 | 255093329 | 4 | 0 | 1.08 | 1.07 |
| Q9DCD0 | Pgd | 6-phosphogluconate dehydrogenase, decarboxylating | 1812.8 | 69.57 | 53247 | 27 | 151253345 | 164736529 | 175545713 | 136175393 | 153375793 | 178058113 | 165630561 | 182952209 | 192429441 | 181907857 | 181808641 | 192496833 | 207364001 | 189192449 | 1 | 0 | 1.15 | 1.23 |
| E9Q3L2 | Pi4ka | Phosphatidylinositol 4-kinase alpha | 125.61 | 1.9 | 237042 | 2 | 1 | 1 | 134357.41 | 1 | 1 | 136527.52 | 192876.2475 | 256811.02 | 232711.98 | 145454.47 | 206593.7767 | 292065.56 | 146406.19 | 181309.58 | 0 | 0 | 7.18 | 7.69 |
| Q8BXQ2 | Pigt | GPI transamidase component PIG-T | 90.4 | 5.5 | 65705 | 2 | 1827111.88 | 2020421.38 | 1231352.25 | 1495157.5 | 1825833.12 | 2335773.5 | 1720722.25 | 1611787.38 | 1607781.5 | 2130011.75 | 2337649.5 | 2296802.5 | 2326108.75 | 2255476.5 | 9 | 0 | 1.12 | 1.37 |
| P51432 | Plcb3 | 1-phosphatidylinositol 4,5-bisphosphate phosphodiesterase beta-3 | 235.68 | 7.54 | 139487 | 5 | 870095.25 | 1536200 | 1617768.75 | 974266.44 | 1494747.25 | 2135581 | 1680163 | 2121126.5 | 2295048 | 1993543 | 2401421 | 2393453.75 | 1902488.5 | 1969742.5 | 0 | 0 | 1.57 | 1.67 |
| Q8BG07 | Pld4 | Phospholipase D4 | 1299.01 | 42.94 | 56154 | 19 | 217774625 | 283700417 | 237069425 | 208314785 | 226738369 | 151815697 | 173522833 | 199112241 | 179514177 | 161711761 | 189010065 | 166340353 | 174621745 | 194205297 | 0 | 0 | 0.74 | 0.77 |
| Q9R0E1 | Plod3 | Multifunctional procollagen lysine hydroxylase and glycosyltransferase LH3 | 424.41 | 14.98 | 84922 | 8 | 5364478 | 5032839.5 | 5109065.5 | 4433972 | 4825268 | 4635728 | 4899789 | 4705287.5 | 3916408 | 4813765 | 4431840 | 4185398.5 | 4347884.5 | 3604398.5 | 5 | 0 | 0.93 | 0.84 |
| Q99K51 | Pls3 | Plastin-3 | 1757.78 | 62.38 | 70742 | 23 | 428140929 | 438180577 | 453587393 | 434619361 | 434747297 | 523168641 | 448838209 | 530365121 | 537169857 | 478631329 | 468837249 | 474582017 | 470946593 | 490720929 | 1 | 0 | 1.15 | 1.09 |
| P58195 | Plscr1 | Phospholipid scramblase 1 | 58.81 | 5.97 | 36711 | 1 | 1147784.62 | 1143446 | 1155475.38 | 1194112.88 | 1016785.81 | 1057941.25 | 910286.5 | 1227599.75 | 1158016.62 | 826703.88 | 886059.4567 | 775410.56 | 975481.56 | 907286.25 | 10 | 0 | 0.92 | 0.78 |
| P70206 | Plxna1 | Plexin-A1 | 409.96 | 6.12 | 211099 | 7 | 7789652.5 | 11109825 | 8503231 | 8246097.5 | 9001116 | 6290151.5 | 6913009 | 10529014 | 7534522 | 7859419 | 7751602 | 7298075.5 | 7311485.5 | 6265236 | 5 | 0 | 0.88 | 0.80 |
| P23492 | Pnp | Purine nucleoside phosphorylase | 568.02 | 53.63 | 32277 | 9 | 49712993 | 48014693 | 53075109 | 46328725 | 46769449 | 36315229 | 40750337 | 39961977 | 38437057 | 30295669 | 27103267 | 35282353 | 38767565 | 34755837 | 0 | 0 | 0.76 | 0.70 |
| Q76MZ3 | Ppp2r1a | Serine/threonine-protein phosphatase 2A 65 kDa regulatory subunit A alpha isoform | 2107.18 | 64.35 | 65323 | 24 | 114746873 | 128048809 | 126684097 | 118607377 | 116083617 | 141708465 | 126688105 | 144928385 | 137329841 | 127348945 | 129637289 | 132702345 | 134147513 | 136290289 | 2 | 0 | 1.12 | 1.10 |
| Q922D4 | Ppp6r3 | Serine/threonine-protein phosphatase 6 regulatory subunit 3 | 372.73 | 10.31 | 94653 | 7 | 7959086 | 7394944 | 7224796 | 6713954.5 | 7296041 | 6600551.5 | 7507074 | 7655000 | 7689509 | 6380844 | 5766490 | 6682267 | 6132965.5 | 6455693 | 12 | 0 | 0.98 | 0.86 |
| Q8CEC6 | Ppwd1 | Peptidylprolyl isomerase domain and WD repeat-containing protein 1 | 52.69 | 2.01 | 73385 | 1 | 6784737.5 | 14874537 | 46376205 | 30773133 | 69166489 | 83548193 | 20117743 | 3388300 | 37399177 | 90466713 | 77970193 | 86852145 | 87349105 | 92075785 | 10 | 0 | 1.40 | 2.56 |
| O08795 | Prkcsh | Glucosidase 2 subunit beta | 423.96 | 19.39 | 58793 | 8 | 14769977 | 14824747 | 16255006 | 15607839 | 14695123 | 14626441 | 15648041 | 14677563 | 13347025 | 14074395 | 14125877 | 13542592 | 13251337 | 14463818 | 4 | 0 | 0.95 | 0.91 |
| Q9R1P1 | Psmb3 | Proteasome subunit beta type-3 | 493.69 | 53.17 | 22965 | 8 | 38088821 | 39792409 | 41585749 | 38853289 | 41475689 | 43095721 | 40983641 | 38006333 | 44259581 | 43750361 | 43412085 | 43693037 | 45077701 | 43424569 | 7 | 0 | 1.05 | 1.10 |
| O35522 | Psmb9 | Proteasome subunit beta type-9 Mus musculus bactrianus | 260.75 | 34.25 | 23427 | 5 | 23776365 | 27161923 | 22946583 | 26063835 | 24925917 | 20062789 | 21509665 | 21172077 | 18464499 | 19879483 | 20457933 | 17895225 | 19753685 | 20238313 | 0 | 0 | 0.81 | 0.78 |
| P97814 | Pstpip1 | Proline-serine-threonine phosphatase-interacting protein 1 | 282.97 | 17.11 | 47590 | 5 | 9298495 | 9274206 | 8585177 | 9588877 | 8833164 | 7339234 | 8400129 | 8584521 | 7509706 | 8168469.5 | 7430355.5 | 8039140.5 | 7276998.5 | 7285711 | 0 | 0 | 0.88 | 0.82 |
| P49445 | Ptpn7 | Tyrosine-protein phosphatase non-receptor type 7 | 46.59 | 3.62 | 40314 | 1 | 504673.72 | 502326.03 | 530789.44 | 704985.31 | 605939.31 | 580063.94 | 475219.78 | 514228.56 | 379385.38 | 519743.59 | 481192.69 | 490500.62 | 445578.91 | 445627.5 | 7 | 0 | 0.87 | 0.82 |
| P61027 | Rab10 | Ras-related protein Rab-10 | 490.94 | 42 | 22541 | 7 | 21850323 | 21533451 | 21389139 | 20575261 | 19195639 | 23333243 | 22967457 | 24281881 | 22086963 | 22118249 | 22161477 | 22816339 | 24300193 | 22347823 | 0 | 0 | 1.10 | 1.10 |
| Q99JI6 | Rap1b | Ras-related protein Rap-1b | 1023.91 | 80.43 | 20825 | 6 | 136927169 | 141910849 | 128245809 | 131854817 | 126793345 | 124197873 | 131392225 | 137265777 | 121309553 | 126138625 | 120258377 | 112723361 | 125830321 | 122650929 | 6 | 0 | 0.96 | 0.90 |
| Q8VCT3 | Rnpep | Aminopeptidase B | 1563.82 | 54.46 | 72416 | 26 | 62735149 | 67485457 | 71716025 | 64039325 | 62866457 | 59147557 | 56775197 | 60981129 | 66397173 | 57192385 | 57437581 | 55898233 | 56846157 | 62337717 | 3 | 0 | 0.91 | 0.88 |
| P60603 | Romo1 | Reactive oxygen species modulator 1 | 113.24 | 51.9 | 8183 | 2 | 507002.25 | 705462.4375 | 827640.75 | 435317 | 1051889.75 | 1166872 | 1148360.25 | 501032.97 | 1334988 | 998214.88 | 1392542.75 | 1779152.5 | 1073323.62 | 1407164.12 | 5 | 0 | 1.46 | 2.00 |
| P67984 | Rpl22 | 60S ribosomal protein L22 | 387.03 | 56.25 | 14759 | 7 | 35659789 | 27045523 | 32735147 | 24856975 | 31397327 | 38167421 | 37770341 | 32239589 | 44082929 | 43481093 | 36101401 | 45934985 | 42751321 | 39497965 | 2 | 0 | 1.29 | 1.35 |
| P62892 | Rpl39 | 60S ribosomal protein L39 | 35.58 | 19.61 | 6407 | 1 | 2724630.5 | 2267559 | 2507127.75 | 2371130.75 | 2485049.75 | 3037738.75 | 2288886 | 1699666.12 | 2708701.75 | 3465083.75 | 4062534.25 | 3798987 | 3846852.75 | 3811686.25 | 10 | 0 | 1.07 | 1.57 |
| P62281 | Rps11 | 40S ribosomal protein S11 | 365.91 | 37.34 | 18431 | 8 | 24565721 | 29397857 | 39459553 | 38108433 | 37294617 | 36984009 | 34504213 | 24323863 | 35775441 | 46687985 | 39572737 | 41059757 | 44456165 | 43636773 | 11 | 0 | 1.06 | 1.25 |
| P49242 | Rps3a | 40S ribosomal protein S3a | 740.66 | 45.83 | 29945 | 13 | 113750105 | 112958609 | 114948289 | 118346137 | 109531193 | 97042241 | 105823233 | 100710585 | 100679497 | 105432593 | 102109929 | 103039873 | 99281617 | 107522273 | 0 | 0 | 0.89 | 0.90 |
| O35114 | Scarb2 | Lysosome membrane protein 2 | 295.67 | 14.85 | 54044 | 5 | 3792335.75 | 3064972 | 4783295 | 2932860 | 2375041.5 | 7356850 | 5877957 | 8681323 | 8287700.5 | 5328843 | 5932664 | 10648394 | 8507017 | 7792342 | 0 | 0 | 2.10 | 2.43 |
| Q9CQA3 | Sdhb | Succinate dehydrogenase [ubiquinone] iron-sulfur subunit, mitochondrial | 438.18 | 30.5 | 31814 | 8 | 44796633 | 49837093 | 50325585 | 50812549 | 49363261 | 57099637 | 59152501 | 52235157 | 64218137 | 59485769 | 53334785 | 55999333 | 53467569 | 55464701 | 0 | 0 | 1.19 | 1.11 |
| Q9CXV1 | Sdhd | Succinate dehydrogenase [ubiquinone] cytochrome b small subunit, mitochondrial | 82.47 | 16.35 | 17014 | 1 | 2303064.75 | 1947185.12 | 1287627.62 | 1616036.25 | 1565868.62 | 952996.25 | 1037701.69 | 1567015.25 | 819913.06 | 910888.75 | 1212176.75 | 1040106.31 | 881603.31 | 1251570.62 | 2 | 0 | 0.61 | 0.63 |
| Q3U2P1 | Sec24a | Protein transport protein Sec24A | 45.44 | 1.38 | 118782 | 1 | 1 | 1 | 1 | 1 | 1 | 1 | 236386.28 | 1 | 277140.5 | 1 | 313900.7 | 307739.38 | 359366.22 | 274596.5 | 7.5 | 0 | 102705.96 | 313900.70 |

| | | | | | | | | | | | | | | | | | | | | | | | | |
|---|---|---|---|---|---|---|---|---|---|---|---|---|---|---|---|---|---|---|---|---|---|---|---|---|
| Q8BH59 | Slc25a12 | Calcium-binding mitochondrial carrier protein Aralar1 | 1161.56 | 43.72 | 74570 | 17 | 36904181 | 39552901 | 41379253 | 35412145 | 39784721 | 43641701 | 40349677 | 48736293 | 41627165 | 40790373 | 45488373 | 42274969 | 54611137 | 42473157 | 2 | 0 | 1.11 | 1.20 |
| Q9QXX4 | Slc25a13 | Calcium-binding mitochondrial carrier protein Aralar2 | 1659.72 | 63.61 | 74467 | 26 | 62194497 | 61106117 | 65557537 | 57746685 | 64471141 | 67684505 | 68092353 | 81449977 | 75328153 | 72001529 | 73703249 | 66758481 | 87497329 | 74168241 | 0 | 0 | 1.17 | 1.21 |
| O70579 | Slc25a17 | Peroxisomal membrane protein PMP34 | 72.36 | 4.56 | 34413 | 1 | 245448.55 | 232337.67 | 208585.3775 | 152327.23 | 204228.06 | 239832.24 | 223624.25 | 257527.38 | 298430.5 | 179746.83 | 278119.81 | 311607.62 | 374917.25 | 280820.88 | 7 | 0 | 1.15 | 1.49 |
| P16036 | Slc25a3 | Phosphate carrier protein, mitochondrial | 713.67 | 44.66 | 39445 | 12 | 36062841 | 41107849 | 41221105 | 34584221 | 38753701 | 47766305 | 45828705 | 50553545 | 54328689 | 48474313 | 44965977 | 49965213 | 50002517 | 46465757 | 0 | 0 | 1.29 | 1.25 |
| Q8BTY2 | Slc4a7 | Sodium bicarbonate cotransporter 3 | 198.97 | 3.77 | 116514 | 3 | 1 | 1 | 1 | 1 | 1 | 1 | 1 | 278411.38 | 1 | 142437.42 | 191332.16 | 256858.83 | 744044.69 | 345649.22 | 7.5 | 0 | 84170.36 | 384471.23 |
| Q3UKJ7 | Smu1 | WD40 repeat-containing protein SMU1 | 420.49 | 24.76 | 57544 | 9 | 5199344.5 | 3802362.75 | 4249785 | 5334716.5 | 5519688 | 5444902.5 | 5548634.5 | 4951032 | 5269399 | 5348591 | 5807639 | 8477330 | 6160661.5 | 6499396.5 | 7 | 0 | 1.10 | 1.40 |
| P57784 | Snrpa1 | U2 small nuclear ribonucleoprotein A' | 577.45 | 45.88 | 28357 | 11 | 11704108 | 11399720 | 13774256 | 15603991 | 11045075 | 8645690 | 10169872 | 6992035.5 | 14930463 | 14816768 | 10884130 | 10504986 | 8214811 | 6768221.5 | 8 | 0 | 0.87 | 0.72 |
| Q5BK61 | Snx20 | Sorting nexin-20 | 45.89 | 3.83 | 35723 | 1 | 781292.69 | 719573.88 | 749767.56 | 606141.81 | 766262.44 | 616120.06 | 536662.12 | 528662.44 | 693264.19 | 634766.38 | 597233.88 | 509362.72 | 576291.06 | 601287.12 | 3 | 0 | 0.83 | 0.79 |
| O35892 | Sp100 | Nuclear autoantigen Sp-100 | 189.49 | 9.96 | 54727 | 3 | 3439218.5 | 3207927 | 3216481.25 | 2612204 | 2956948.5 | 1324223.75 | 2217736.75 | 2585625 | 2355409.25 | 2315391.5 | 2404158.75 | 2575712.25 | 2204169 | 1244349 | 0 | 0 | 0.70 | 0.68 |
| Q8BVK9 | Sp110 | Sp110 nuclear body protein | 472.34 | 32.58 | 50140 | 10 | 9270450 | 7255536 | 8656968 | 7135867.5 | 8281462.5 | 5700259 | 6962201 | 6817715.5 | 6990179.5 | 6948625.5 | 5176007 | 5951477 | 6941924.5 | 5088065.5 | 0 | 0 | 0.82 | 0.71 |
| Q8BMA6 | Srp68 | Signal recognition particle subunit SRP68 | 758.24 | 24 | 70574 | 12 | 8877004 | 9019523 | 9309933 | 9427344 | 8799103 | 9350803 | 8182989 | 8568866 | 9923001 | 7162825 | 8237717 | 7605911.5 | 8263081 | 6750717.5 | 9 | 0 | 0.95 | 0.85 |
| O35326 | Srsf5 | Serine/arginine-rich splicing factor 5 | 351.71 | 25.28 | 30891 | 5 | 46996529 | 46404385 | 42676233 | 45301057 | 48428285 | 44846393 | 42924545 | 38399329 | 42644145 | 47435765 | 39357057 | 39502393 | 41972405 | 41565229 | 6 | 0 | 0.94 | 0.88 |
| O88829 | St3gal5 | Lactosylceramide alpha-2,3-sialyltransferase | 64.94 | 5.56 | 47360 | 2 | 2107032.25 | 2109276.25 | 1447866.62 | 2348310.5 | 2417529.5 | 2630961 | 2028636.38 | 1924562.75 | 2392820 | 2569061.75 | 2553602.5 | 2433463.5 | 2611923 | 2993465 | 9 | 0 | 1.11 | 1.27 |
| Q9JM90 | Stap1 | Signal-transducing adaptor protein 1 | 637.43 | 59.93 | 34628 | 13 | 17729511 | 12382445 | 13236752 | 13402185 | 15024507 | 23414919 | 21383787 | 18052759 | 23343591 | 22613869 | 22383089 | 22426727 | 19523953 | 20117637 | 0 | 0 | 1.52 | 1.47 |
| P42225 | Stat1 | Signal transducer and activator of transcription 1 | 1044.25 | 29.37 | 87197 | 17 | 27494325 | 28965477 | 27279907 | 21606003 | 25210429 | 16652458 | 15746811 | 15368089 | 18423641 | 16536492 | 14598458 | 14268871 | 18060927 | 15280153 | 0 | 0 | 0.63 | 0.60 |
| Q9WVL2 | Stat2 | Signal transducer and activator of transcription 2 | 153.72 | 3.47 | 105417 | 3 | 2279999.25 | 2238939.75 | 2558215.5 | 1865435.12 | 1575645.25 | 1512023 | 247460.8 | 1372053.88 | 1270407.5 | 1310062.38 | 1185951.38 | 1208175.88 | 1289742.88 | 1175394.5 | 0 | 0 | 0.54 | 0.58 |
| Q8VIM6 | Strc | Stereocilin | 35.29 | 0.44 | 196347 | 1 | 722819.81 | 2210075.5 | 1922223.88 | 1858499.88 | 719493.31 | 3080604.5 | 3225625.75 | 3418017 | 4935381 | 2952063.25 | 3495577.5 | 2236525 | 3993204 | 3594814.25 | 0 | 0 | 2.37 | 2.24 |
| Q9WUD1 | Stub1 | STIP1 homology and U box-containing protein 1 | 57.9 | 3.95 | 34909 | 1 | 3146873.188 | 3086832.25 | 2984497.5 | 3650552 | 2865611 | 2090934.5 | 3187515 | 2745167.25 | 2363677.5 | 2170151.25 | 2437110.25 | 2648986.667 | 2685198.5 | 2824651.25 | 4 | 0 | 0.80 | 0.84 |
| O88983 | Stx8 | Syntaxin-8 | 152.78 | 11.86 | 26925 | 2 | 1879497.563 | 1758616.75 | 2002885.88 | 1978806.12 | 1777681.5 | 1351127.75 | 1448355.62 | 1478653.75 | 1335958.88 | 1441302.38 | 1542985.25 | 1589585.083 | 1527788.5 | 1697981.5 | 0 | 0 | 0.75 | 0.85 |
| Q9Z2I9 | Sucla2 | Succinate--CoA ligase [ADP-forming] subunit beta, mitochondrial | 456.06 | 29.37 | 50114 | 9 | 8295629.5 | 10424399 | 10797735 | 8882850 | 9556912 | 11354631 | 9859279 | 13265904 | 12864112 | 11100224 | 12045576 | 12539160 | 12071474 | 10814796 | 2 | 0 | 1.22 | 1.24 |
| Q9Z2I8 | Suclg2 | Succinate--CoA ligase [GDP-forming] subunit beta, mitochondrial | 832.03 | 43.42 | 46840 | 14 | 46315337 | 40460253 | 48964709 | 41624685 | 43707373 | 55236441 | 51140345 | 52193109 | 55620869 | 46938513 | 52441313 | 56616301 | 50560709 | 52940785 | 1 | 0 | 1.18 | 1.20 |
| Q0VGY8 | Tanc1 | Protein TANC1 | 276.4 | 5.87 | 200805 | 6 | 1665739.12 | 1572150.38 | 1584758.25 | 1529621.75 | 1553410 | 2260720.75 | 2593082.75 | 2205979 | 2687452.5 | 2599394.75 | 2192618 | 1965477.12 | 1814102.62 | 1908039.88 | 0 | 0 | 1.56 | 1.25 |
| P21958 | Tap1 | Antigen peptide transporter 1 | 423.56 | 17.68 | 78864 | 9 | 10058543 | 10971870 | 10419274 | 7036405.5 | 7464756 | 5593281 | 5416298 | 6594180 | 6924227 | 6284634 | 4814017 | 3992453.25 | 6695657 | 5874981.5 | 0 | 0 | 0.67 | 0.58 |
| P36371 | Tap2 | Antigen peptide transporter 2 | 530.08 | 19.52 | 77445 | 9 | 14567948 | 13917693 | 10892413 | 10225757 | 12761393 | 5557308 | 7911689 | 6487473 | 9713233 | 5826027 | 8362977.5 | 5555959 | 7068710 | 9032469 | 0 | 0 | 0.57 | 0.60 |
| Q8BHJ5 | Tbl1xr1 | F-box-like/WD repeat-containing protein TBL1XR1 | 213.14 | 9.14 | 55661 | 2 | 14106546 | 16720502 | 16898611 | 15335139 | 1277937 | 3426218.75 | 523779.81 | 17327333 | 499951.09 | 846032.38 | 662916.5 | 1060689.5 | 821011.88 | 665046.19 | 6 | 0 | 0.35 | 0.06 |
| Q91YX0 | Themis2 | Protein THEMIS2 | 451.31 | 22.32 | 74378 | 9 | 6519084.5 | 7443011.5 | 6576726.5 | 7534454 | 7404014.5 | 3663874.5 | 4519365 | 4221583.5 | 4518707.5 | 4335175.5 | 5111128.5 | 5144754.5 | 5504107 | 4491972 | 0 | 0 | 0.60 | 0.71 |
| P97770 | Thumpd3 | THUMP domain-containing protein 3 | 227.58 | 11.49 | 56431 | 5 | 899067.62 | 1041571.62 | 1029383.31 | 1288757.88 | 1037148.62 | 1246746.5 | 979046.88 | 965987.75 | 1035807.88 | 1869101.88 | 1648342.25 | 1458954.5 | 1544222.62 | 1375195.5 | 12 | 0 | 1.15 | 1.42 |
| P58681 | Tlr7 | Toll-like receptor 7 | 670.44 | 16.38 | 121837 | 14 | 9379617 | 8604142 | 9166527 | 10330100 | 8842673 | 8929297 | 8443729 | 9944700 | 6979926.5 | 7646626 | 8494588 | 8120686.5 | 6905860.5 | 7610974.5 | 6 | 0 | 0.91 | 0.84 |
| Q6AYA5 | Tmem106b | Transmembrane protein 106B | 52.41 | 5.82 | 31152 | 1 | 1 | 808104.06 | 1153426 | 1 | 1 | 1358662.375 | 1316674 | 1427608.5 | 1189302 | 1501065 | 1351118.75 | 1330084.5 | 1385718 | 1374090.88 | 0 | 0 | 3.46 | 3.47 |
| Q61033 | Tmpo | Lamina-associated polypeptide 2, isoforms alpha/zeta | 726.2 | 27.85 | 75168 | 6 | 10506329 | 10680562 | 11460652 | 12180816 | 10627226 | 11433041 | 9259722 | 9149543 | 9467989 | 12519884 | 8628876 | 10067712 | 9138476 | 9306061 | 8 | 0 | 0.93 | 0.84 |
| Q8BFY9 | Tnpo1 | Transportin-1 | 916.78 | 28.73 | 102357 | 13 | 39307329 | 40928837 | 39648933 | 39648505 | 41273549 | 43799013 | 40104089 | 44907905 | 42044061 | 39123097 | 44049557 | 43053501 | 44068709 | 44637841 | 7 | 0 | 1.05 | 1.09 |
| Q9CYG7 | Tomm34 | Mitochondrial import receptor subunit TOM34 | 659.61 | 45.95 | 34278 | 11 | 12720540 | 14699377 | 15130840 | 12742153 | 14684081 | 16434287 | 16587682 | 16365119 | 18186055 | 16624583 | 16003520 | 16805789 | 17180667 | 17118991 | 0 | 0 | 1.20 | 1.20 |
| Q91XB0 | Trex1 | Three-prime repair exonuclease 1 | 159.45 | 14.01 | 33675 | 3 | 3858302.5 | 6591903 | 3002202.5 | 5408457.5 | 5267453.5 | 2709213 | 1787539.12 | 2425848 | 2479948.25 | 3220306 | 2229514 | 2075390.62 | 2429171.5 | 2635683 | 1 | 0 | 0.52 | 0.49 |
| P23591 | Tsta3 | GDP-L-fucose synthase | 230.02 | 20.25 | 35878 | 5 | 7730363.5 | 8365780 | 7979294 | 8123320 | 8205173.5 | 6914703 | 6895175 | 7710219 | 6726803.5 | 6897820.5 | 6750698 | 7032807 | 6528529.5 | 7176565 | 0 | 0 | 0.87 | 0.85 |
| Q6PAM1 | Txlna | Alpha-taxilin | 130 | 6.14 | 62369 | 2 | 1280284.75 | 1384573.938 | 1239255.75 | 1673974.75 | 1344780.5 | 1286422.843 | 1188025.62 | 1049875.62 | 1649319.25 | 1258470.88 | 964751.31 | 1186727.75 | 852930.81 | 1151665.5 | 7 | 0 | 0.93 | 0.75 |
| Q9CQM5 | Txndc17 | Thioredoxin domain-containing protein 17 | 231.16 | 39.84 | 14015 | 4 | 10687913 | 11595468 | 12056245 | 11825727 | 11382406 | 11669695 | 11813478 | 11713321 | 12136583 | 12893105 | 12632143 | 12233735 | 12153640 | 12660701 | 6 | 0 | 1.05 | 1.08 |
| Q9CR09 | Ufc1 | Ubiquitin-fold modifier-conjugating enzyme 1 | 173.85 | 31.14 | 19481 | 4 | 7331790 | 6625408.5 | 6249144.5 | 5932308 | 6208152.5 | 4703334 | 4986133.5 | 5569932 | 5297996.5 | 4591163 | 5214540.5 | 4845140 | 5559540 | 3579490.25 | 0 | 0 | 0.78 | 0.74 |
| Q62465 | Vat1 | Synaptic vesicle membrane protein VAT-1 homolog | 635.9 | 35.47 | 43097 | 10 | 13013952 | 12600807 | 11770273 | 9058213 | 10924196 | 15030296 | 22454865 | 16856367 | 19143909 | 19103937 | 17374861 | 18009465 | 19733481 | 17329637 | 0 | 0 | 1.61 | 1.58 |
| Q6NZF1 | Zc3h11a | Zinc finger CCCH domain-containing protein 11A | 217.47 | 8.33 | 86492 | 5 | 1478486.5 | 1571028 | 1751738.25 | 1529590 | 1550828.5 | 471800 | 1433966.25 | 1472847.5 | 1087586.38 | 1296602.88 | 981408.69 | 1301186.88 | 1336700.62 | 256232.75 | 0 | 0 | 0.73 | 0.61 |

| | | | | | | | | | | | | | | | | | | | | | | | |
|---|---|---|---|---|---|---|---|---|---|---|---|---|---|---|---|---|---|---|---|---|---|---|---|
| Q9EPL9 | Acox3 | Peroxisomal acyl-coenzyme A oxidase 3 | 163.39 | 6.43 | 78404 | 2 | 2464755.25 | 2313931.75 | 2644338.25 | 1952754.62 | 2369746 | 2825278.5 | 2387081.5 | 2392145 | 2414242.75 | 2425126.75 | 2512192.5 | 2683649.25 | 2810664.75 | 3137525.25 | 8 | 1 | 1.06 | 1.19 |
| O88839 | Adam15 | Disintegrin and metalloproteinase domain-containing protein 15 | 102.67 | 2.08 | 92664 | 1 | 202693.86 | 264608.88 | 222039.39 | 219734.345 | 189595.25 | 416076.72 | 317525.62 | 365111.09 | 336759.44 | 249386.42 | 318597.91 | 522001.75 | 333007.41 | 250278.95 | 1 | 1 | 1.53 | 1.62 |
| P31750 | Akt1 | RAC-alpha serine/threonine-protein kinase | 170.55 | 10.62 | 55707 | 2 | 1051679.38 | 943814.19 | 878722.9375 | 769563.62 | 749834.56 | 458136.59 | 614655.94 | 820263.62 | 1679068.5 | 629130 | 1686174 | 1502030.29 | 1811631.25 | 1008285.62 | 7 | 1 | 0.96 | 1.71 |
| P07356 | Anxa2 | Annexin A2 | 1632.99 | 70.8 | 38676 | 25 | 413457761 | 402877729 | 423519713 | 427536897 | 441378049 | 471697665 | 521019233 | 505123873 | 568528513 | 556714433 | 470546401 | 450212353 | 440189633 | 446280225 | 0 | 1 | 1.24 | 1.07 |
| Q63055 | Arfrp1 | ADP-ribosylation factor-related protein 1 2 | 74.71 | 8.96 | 22659 | 1 | 609896.19 | 1 | 775940.44 | 1 | 896574.25 | 893475.31 | 561702.5 | 346706.44 | 696107 | 856068.69 | 1192839.12 | 1004777.69 | 973784.88 | 828399.31 | 10 | 1 | 1.47 | 2.19 |
| Q3UIA2 | Arhgap17 | Rho GTPase-activating protein 17 | 444.6 | 14.89 | 92202 | 10 | 12156895 | 8757226 | 8017522 | 8870471 | 10846009 | 7692902 | 7969897.5 | 6999208 | 7397576 | 7272889 | 8430568 | 6067848.5 | 7748019 | 5477966 | 0 | 1 | 0.77 | 0.71 |
| Q8VBT9 | Aspscr1 | Tether containing UBX domain for GLUT4 | 141.82 | 6.73 | 59796 | 2 | 654962.44 | 539152.06 | 575547.265 | 693931.62 | 414142.94 | 462849.41 | 1 | 702454.75 | 438055.12 | 394186.12 | 446300.66 | 337056.69 | 367681.97 | 350275.06 | 7 | 1 | 0.69 | 0.65 |
| Q3KRE0 | Atad3 | ATPase family AAA domain-containing protein 3 | 182.8 | 7.28 | 66759 | 3 | 1 | 1 | 153757.5 | 1 | 196889.88 | 260729.42 | 205999.27 | 294182.22 | 242422.42 | 282221.62 | 531651.44 | 354055.62 | 360888.22 | 169627.2 | 0 | 1 | 3.67 | 5.05 |
| Q64518 | Atp2a3 | Sarcoplasmic/ endoplasmic reticulum calcium ATPase 3 | 740.84 | 15.9 | 113638 | 7 | 21217385 | 21617743 | 20382085 | 22246409 | 24183677 | 23850321 | 24953457 | 22223931 | 24122077 | 24198801 | 25532695 | 25672589 | 23845941 | 25181271 | 4 | 1 | 1.09 | 1.14 |
| Q8BVE3 | Atp6v1h | V-type proton ATPase subunit H | 696.59 | 32.3 | 55855 | 13 | 20321119 | 20710759 | 23600781 | 18563459 | 19201917 | 27175529 | 22681483 | 23013645 | 26928313 | 23830863 | 22637419 | 25989765 | 25466551 | 23831829 | 2 | 1 | 1.21 | 1.20 |
| Q91YY4 | Atpaf2 | ATP synthase mitochondrial F1 complex assembly factor 2 | 40.45 | 3.81 | 33289 | 1 | 1 | 1 | 714317.5 | 1 | 1 | 1 | 1 | 1 | 1 | 647275.5 | 876839.9167 | 986246.06 | 1037077.31 | 607196.38 | 12 | 1 | 0.91 | 6.14 |
| P59017 | Bcl2l13 | Bcl-2-like protein 13 | 266.18 | 20.97 | 46719 | 5 | 5133233 | 7492595.5 | 7480295.5 | 8898548 | 7648284 | 9898227 | 9877343 | 6319342 | 9542977 | 6798287 | 10489876 | 8563896 | 9486708 | 10260459 | 8 | 1 | 1.16 | 1.32 |
| Q923D2 | Blvrb | Flavin reductase (NADPH) | 516.99 | 60.19 | 22197 | 7 | 5689220 | 4773408 | 5004091 | 6257829 | 6131784 | 4959825 | 5258340.5 | 4532174 | 4845444 | 7765961 | 6198497 | 6815858.5 | 6466006 | 6357858.5 | 9 | 1 | 0.98 | 1.16 |
| P30993 | C5ar1 | C5a anaphylatoxin chemotactic receptor 1 | 90.77 | 6.84 | 39023 | 2 | 3853467 | 2171287.75 | 1685147 | 1996552.75 | 1890232 | 2015879.38 | 2009257.62 | 1959171.38 | 2052005.62 | 2125921 | 320695.62 | 1392116.62 | 1544688.38 | 1752020.75 | 11 | 1 | 0.88 | 0.54 |
| P35564 | Canx | Calnexin | 1226.02 | 49.58 | 67278 | 22 | 75944481 | 74517137 | 79112257 | 75547153 | 73482825 | 76148985 | 71898929 | 82528801 | 77842393 | 76958265 | 71962137 | 73018681 | 73568129 | 72016993 | 8 | 1 | 1.02 | 0.96 |
| O88456 | Capns1 | Calpain small subunit 1 | 587.5 | 59.48 | 28463 | 3 | 30998365 | 34028041 | 29501311 | 25698957 | 30787103 | 35859877 | 35178441 | 38785485 | 35084545 | 32038903 | 35672193 | 33780065 | 35818065 | 37239785 | 1 | 1 | 1.17 | 1.18 |
| P83917 | Cbx1 | Chromobox protein homolog 1 | 368.92 | 45.95 | 21418 | 4 | 12186138 | 15126752 | 13201249 | 14069240 | 18252925 | 17587037 | 18268437 | 16005048 | 17804937 | 18846813 | 18476091 | 18556811 | 16343273 | 18917501 | 3 | 1 | 1.22 | 1.24 |
| Q8BRN9 | Cc2d1b | Coiled-coil and C2 domain-containing protein 1B | 177.26 | 11.32 | 93091 | 4 | 554515.06 | 907815.56 | 706672 | 1 | 740409.44 | 1351665.5 | 1111377.36 | 1562820.75 | 782064.81 | 748958.38 | 1697086 | 1037577.94 | 794477.75 | 1176380.563 | 2 | 1 | 1.91 | 2.02 |
| Q9CWK3 | Cd2bp2 | CD2 antigen cytoplasmic tail-binding protein 2 | 42.89 | 4.09 | 37694 | 1 | 924238.69 | 937526.25 | 700660.44 | 880414.75 | 959233.62 | 666843.25 | 789463.25 | 701619.11 | 524462.38 | 825707.56 | 635182.56 | 594330.38 | 607225.56 | 716460.5 | 3 | 1 | 0.80 | 0.72 |
| Q68FD5 | Cltc | Clathrin heavy chain 1 | 5817.88 | 65.13 | 191557 | 85 | 464104705 | 492453857 | 482883425 | 470813473 | 474982721 | 415601153 | 443680929 | 478636065 | 465366945 | 446243137 | 464605345 | 439327233 | 445865025 | 460420769 | 4 | 1 | 0.94 | 0.95 |
| P36552 | Cpox | Oxygen-dependent coproporphyrinogen-III oxidase, mitochondrial | 149.92 | 7.22 | 49715 | 3 | 1503168 | 1347360.5 | 1329728.25 | 1128550.25 | 1129803.62 | 1787710.75 | 1646714.88 | 1371607.25 | 3656753 | 3044954.5 | 1432165.5 | 3589638.5 | 1740911.75 | 3050852.5 | 1 | 1 | 1.79 | 1.91 |
| P70698 | Ctps1 | CTP synthase 1 | 717.68 | 27.41 | 66682 | 11 | 11293901 | 12770289 | 11429613 | 12607571 | 10775793 | 11147971 | 11544509 | 10293005 | 12308024 | 10984212 | 11113973 | 10472703 | 10176380 | 10660851 | 8 | 1 | 0.96 | 0.90 |
| O70370 | Ctss | Cathepsin S | 809.73 | 49.12 | 38475 | 10 | 238743409 | 309175649 | 274737185 | 216478641 | 242877905 | 188496833 | 181920641 | 208634577 | 183625857 | 167515569 | 163645185 | 144368785 | 225630465 | 196082705 | 0 | 1 | 0.73 | 0.71 |
| Q9D4H8 | Cul2 | Cullin-2 | 222.18 | 7.92 | 86877 | 5 | 1236850.5 | 1862539.88 | 1368809.25 | 1538803.75 | 1462571.5 | 655457.19 | 2131048.75 | 3279417 | 2955035.25 | 1284577.75 | 2026049.5 | 1597622.25 | 1895738.12 | 2344790 | 9 | 1 | 1.38 | 1.32 |
| Q9JII5 | Dazap1 | DAZ-associated protein 1 | 250.56 | 15.27 | 43214 | 4 | 3752942 | 5246539 | 3905766 | 5174724 | 5566712 | 5490872.5 | 5541856 | 4862856.5 | 5315936 | 5817361 | 5707034 | 5535279 | 5578547 | 5631757 | 6 | 1 | 1.14 | 1.19 |
| P11030 | Dbi | Acyl-CoA-binding protein | 144.97 | 39.08 | 10027 | 2 | 8975738 | 7477482 | 6957561 | 8622616 | 7808447.5 | 4107992.25 | 6293613.5 | 7137555 | 4203308 | 7113129 | 7026639.5 | 5648527 | 5563802 | 6502662 | 2 | 1 | 0.72 | 0.78 |
| Q63413 | Ddx39b | Spliceosome RNA helicase Ddx39b 2 | 1524.45 | 73.83 | 49035 | 12 | 190927009 | 204810945 | 194309809 | 190996753 | 184235201 | 176000593 | 195061233 | 191997537 | 180648033 | 183251217 | 178748833 | 181634881 | 177056833 | 189674785 | 7 | 1 | 0.96 | 0.94 |
| Q62167 | Ddx3x | ATP-dependent RNA helicase DDX3X | 1544.52 | 48.79 | 73101 | 8 | 87973793 | 79139657 | 82385113 | 69691041 | 81013609 | 69881217 | 72466593 | 65692121 | 67788089 | 72041737 | 63908561 | 65681021 | 70222729 | 68686049 | 3 | 1 | 0.87 | 0.84 |
| Q8BH86 | Dglucy | D-glutamate cyclase, mitochondrial | 46.52 | 3.24 | 66366 | 1 | 783220.88 | 994559.88 | 892727.81 | 801676.81 | 900332.75 | 1045247.19 | 884992.5475 | 881594.38 | 891730.5 | 721398.12 | 964261.88 | 1133439.75 | 999162.06 | 1169527 | 11 | 1 | 1.01 | 1.22 |
| Q5M9G7 | Diexf | Digestive organ expansion factor homolog | 160.09 | 4.59 | 87825 | 3 | 1 | 1 | 1 | 1 | 380879.97 | 1 | 1 | 283632.78 | 469666.59 | 316789.94 | 377079.88 | 569836.8367 | 658585.69 | 673844.94 | 8 | 1 | 2.81 | 7.48 |
| O08749 | Dld | Dihydrolipoyl dehydrogenase, mitochondrial | 1010.55 | 45.38 | 54272 | 16 | 91068521 | 71589561 | 59574589 | 75143865 | 65895429 | 57105733 | 84849641 | 82482873 | 67335329 | 56066889 | 57607657 | 52261449 | 63132309 | 57933181 | 10 | 1 | 0.96 | 0.79 |
| Q9QYJ3 | Dnajb1 | DnaJ homolog subfamily B member 1 | 165.74 | 15 | 38167 | 5 | 4165890 | 3318580 | 3386349.25 | 3622478 | 3842204.5 | 5365432.5 | 5675294 | 4051349.75 | 5571722 | 3914628.5 | 4397374 | 3961700.75 | 5283370 | 5117586 | 2 | 1 | 1.34 | 1.28 |
| P21575 | Dnm1 | Dynamin-1 | 204.22 | 5.56 | 97295 | 1 | 5910187 | 5974341.5 | 4994817 | 5374599.5 | 5425708 | 5404157 | 5384508.5 | 4948333 | 5310773 | 5101730.5 | 5222991 | 4766753 | 3689965.75 | 4639552 | 6 | 1 | 0.94 | 0.83 |
| O35303 | Dnm1l | Dynamin-1-like protein | 547.71 | 20.93 | 83908 | 11 | 8967139 | 7922948.5 | 9526005 | 7493573.5 | 6815973 | 6568428 | 6951001 | 8621467 | 9410451 | 7315750 | 6495592 | 7276593 | 6800646 | 6117038 | 9 | 1 | 0.95 | 0.82 |
| P63242 | Eif5a | Eukaryotic translation initiation factor 5A-1 | 1045.08 | 88.31 | 16832 | 15 | 125870945 | 127178625 | 152176417 | 127524241 | 88316369 | 135720497 | 139334961 | 165439825 | 160879089 | 144208609 | 142868353 | 152212369 | 156684913 | 157686529 | 3 | 1 | 1.20 | 1.23 |
| Q7TT37 | Elp1 | Elongator complex protein 1 | 267.45 | 4.65 | 149584 | 5 | 2316355 | 1619828.5 | 1629935.12 | 1721657.25 | 2597709.5 | 3143377.25 | 1607188.25 | 2924868.75 | 2481901 | 2437639 | 3018784.25 | 2815225.5 | 2577866.25 | 3220420 | 7 | 1 | 1.27 | 1.47 |
| P17182 | Eno1 | Alpha-enolase | 2394.93 | 77.42 | 47141 | 26 | 483027841 | 544320321 | 569124609 | 442800673 | 497630657 | 576118465 | 683387137 | 669707137 | 704570817 | 678481153 | 568724865 | 618923585 | 630460033 | 581501569 | 0 | 1 | 1.31 | 1.18 |
| Q8VD58 | Evi2b | Protein EVI2B | 39.26 | 2.48 | 47993 | 1 | 3684902.25 | 2832548.5 | 2309091.5 | 1882541 | 2129549.25 | 1573820.5 | 2069914.88 | 2132688.75 | 1717356.12 | 1615230.62 | 1113547.25 | 1533057.38 | 1955776.75 | 1665572.12 | 3 | 1 | 0.71 | 0.61 |
| Q9CZD3 | Gars | Glycine--tRNA ligase | 1395.78 | 37.59 | 81878 | 3 | 96797529 | 94607849 | 95651737 | 99651545 | 93605913 | 109506609 | 108360201 | 106143361 | 109224985 | 107692209 | 102365249 | 99154657 | 106202961 | 101447001 | 0 | 1 | 1.13 | 1.06 |
| Q9Z0E6 | Gbp2 | Guanylate-binding protein 2 | 328.57 | 16.3 | 66739 | 6 | 4043399.75 | 4796837 | 4765077 | 1587843.38 | 3894907.25 | 888460 | 622300.69 | 531811.94 | 305732.81 | 1592889.25 | 946624.88 | 1949364.75 | 1239605 | 1473114.75 | 1 | 1 | 0.21 | 0.37 |

| | | | | | | | | | | | | | | | | | | | | | | | |
|---|---|---|---|---|---|---|---|---|---|---|---|---|---|---|---|---|---|---|---|---|---|---|---|
| Q5BJP6 | Gfm2 | Ribosome-releasing factor 2, mitochondrial | 47.55 | 1.67 | 85914 | 1 | 1 | 1 | 807486.94 | 879959.81 | 820097.69 | 728937.56 | 1 | 1 | 1 | 1 | 893669.94 | 921067 | 853079.81 | 889272.25 | 6 | 1 | 0.29 | 1.77 |
| Q8BFR4 | Gns | N-acetylglucosamine-6-sulfatase | 715.07 | 25.55 | 61175 | 13 | 53642101 | 56073937 | 50804413 | 59516449 | 58488501 | 54168009 | 57293253 | 40762445 | 54765685 | 47932913 | 49966917 | 48354757 | 51611917 | 47777021 | 7 | 1 | 0.92 | 0.89 |
| Q91VW5 | Golga4 | Golgin subfamily A member 4 | 59.57 | 0.58 | 257564 | 1 | 421708.03 | 476552.31 | 481143.72 | 511592.06 | 283701.88 | 503459.19 | 509830.31 | 481181.22 | 445429.25 | 550146.75 | 561573.81 | 582947.25 | 744044.69 | 500434.44 | 6 | 1 | 1.15 | 1.37 |
| P11352 | Gpx1 | Glutathione peroxidase 1 | 216.24 | 29.85 | 22329 | 4 | 12420509 | 9940401 | 8472454 | 11677418 | 11073843 | 8316239 | 9854850 | 8782298 | 8203964 | 7889484 | 8033744 | 7628524 | 8737673 | 6710114 | 2 | 1 | 0.80 | 0.73 |
| P97576 | Grpel1 | GrpE protein homolog 1, mitochondrial | 223.31 | 25.35 | 24297 | 5 | 266657.03 | 958875.81 | 245341.73 | 1302648 | 955945.62 | 1787370.5 | 1601601.248 | 2152585.75 | 1825410.62 | 641038.12 | 1955658.25 | 1126871.5 | 1439562.12 | 1564608.25 | 3 | 1 | 2.15 | 2.04 |
| P48774 | Gstm5 | Glutathione S-transferase Mu 5 | 485.22 | 42.41 | 26635 | 7 | 14716581 | 17360573 | 24001869 | 14632357 | 17022199 | 15564605 | 20672531 | 22845959 | 25060383 | 23578205 | 24288985 | 18587005 | 25027437 | 25579923 | 6 | 1 | 1.23 | 1.33 |
| P0C0S6 | H2afz | Histone H2A.Z | 293.14 | 51.56 | 13553 | 3 | 235438561 | 223822273 | 266783489 | 242506881 | 251473025 | 239993553 | 248954081 | 182917041 | 207090113 | 328905921 | 272846721 | 281114849 | 281000193 | 255513825 | 10 | 1 | 0.99 | 1.12 |
| Q8C547 | Heatr5b | HEAT repeat-containing protein 5B | 63.18 | 0.82 | 224319 | 1 | 156976.59 | 135018.84 | 194960.61 | 217384.62 | 220957.33 | 152532.19 | 167774.41 | 125866.74 | 128925.16 | 249098.42 | 258706.38 | 245411.03 | 218303.77 | 259222.94 | 8 | 1 | 0.89 | 1.33 |
| Q64522 | Hist2h2ab | Histone H2A type 2-B | 404.79 | 52.31 | 14013 | 0 | 65926153 | 43979121 | 106511801 | 92464297 | 88569009 | 100197417 | 71374041 | 42014825 | 80647337 | 136808817 | 117431937 | 140340849 | 123040849 | 98975033 | 12 | 1 | 1.08 | 1.51 |
| Q7TMY8 | Huwe1 | E3 ubiquitin-protein ligase HUWE1 | 998.72 | 7.86 | 482635 | 22 | 5177047 | 6848453.5 | 9441940 | 6243555.5 | 7194109 | 8445926 | 11872217 | 7516080.5 | 6044917.5 | 11200916 | 9459361 | 8428893 | 9614492 | 9658723 | 6 | 1 | 1.29 | 1.33 |
| O35368 | Ifi203 | Interferon-activable protein 203 | 141.11 | 12.25 | 46300 | 2 | 3073621 | 4515976 | 3747676 | 3639146.25 | 3943940.5 | 3896708.25 | 3220888.5 | 3169128.75 | 2820361.5 | 1691877.25 | 3061970 | 2135649.5 | 2236953.25 | 3607709 | 5 | 1 | 0.78 | 0.73 |
| Q99J93 | Ifitm2 | Interferon-induced transmembrane protein 2 | 146.22 | 16.67 | 15743 | 1 | 53026101 | 38082793 | 41511925 | 25423989 | 37172865 | 28936005 | 24759839 | 20878133 | 20745715 | 22083409 | 18167669 | 21079547 | 32500441 | 20010805 | 1 | 1 | 0.60 | 0.59 |
| Q9CQW9 | Ifitm3 | Interferon-induced transmembrane protein 3 | 539.03 | 48.91 | 14954 | 5 | 65032261 | 45017053 | 49339585 | 30601089 | 45552461 | 33990385 | 30262735 | 24008263 | 23870285 | 26900781 | 21719847 | 21401057 | 39564185 | 21880273 | 1 | 1 | 0.59 | 0.55 |
| Q66HG8 | Ik | Protein Red | 222.1 | 12.57 | 65588 | 5 | 5506375 | 4898198 | 2074125.12 | 4769928.5 | 4593217 | 5328247.5 | 5047525.5 | 4618095 | 5430646 | 5422640.5 | 5200812.5 | 5715690 | 5855394.5 | 5811492.5 | 7 | 1 | 1.18 | 1.29 |
| Q0GNC1 | Inf2 | Inverted formin-2 | 432.48 | 8.25 | 138560 | 8 | 1647709.12 | 1456264.75 | 1339991.88 | 1237981.5 | 1380480 | 1720009.88 | 1324928 | 2113867.75 | 1550908.5 | 2262031.75 | 1630009.75 | 3229328.5 | 2525351.25 | 1948364 | 5 | 1 | 1.27 | 1.65 |
| Q5EIC4 | Irf2bpl | Interferon regulatory factor 2-binding protein-like | 79.29 | 4.73 | 81496 | 1 | 886338.5 | 1033053.44 | 1089360.38 | 573580.88 | 509385.41 | 1123089.5 | 1158167.88 | 1381445.75 | 1591920.5 | 1102901.75 | 1146767 | 1037823.94 | 1283478.25 | 1304527.25 | 0 | 1 | 1.55 | 1.46 |
| Q5PQL7 | Itm2c | Integral membrane protein 2C | 105.92 | 11.52 | 30480 | 2 | 1782204.5 | 1484172.5 | 1676717.88 | 1326225.5 | 1185751 | 1866026.75 | 1721062.5 | 2472878.75 | 2119065.5 | 1993988.5 | 1764260.75 | 2158080.75 | 2409142.75 | 2332807.5 | 1 | 1 | 1.36 | 1.45 |
| P83953 | Kpna1 | Importin subunit alpha-5 | 477.59 | 22.68 | 60137 | 3 | 4530387.5 | 4801344.5 | 4995068.5 | 4916343 | 3771598.75 | 4824911.5 | 4741556.5 | 3937946.75 | 4973683 | 3520077.5 | 4945946.5 | 5526356.5 | 5209703 | 5375926.5 | 10 | 1 | 0.96 | 1.14 |
| Q61233 | Lcp1 | Plastin-2 | 4110.059 | 87.88 | 70149 | 54 | 1483341441 | 1456825729 | 1599929345 | 1449602433 | 1517976193 | 1695252353 | 1459549953 | 1646451457 | 1754942465 | 1675444481 | 1588014593 | 1606976001 | 1637279873 | 1627295233 | 3 | 1 | 1.10 | 1.08 |
| Q07797 | Lgals3bp | Galectin-3-binding protein | 1156.02 | 39.34 | 64491 | 16 | 84722809 | 85872353 | 84740609 | 60414577 | 75841993 | 48594913 | 47905609 | 47771129 | 44640049 | 45468081 | 48904841 | 45471689 | 70658721 | 50665385 | 0 | 1 | 0.60 | 0.69 |
| Q8CGK3 | Lonp1 | Lon protease homolog, mitochondrial | 1072.35 | 34.04 | 105843 | 20 | 28760953 | 28925937 | 29730303 | 31896783 | 30825573 | 39436801 | 39184665 | 39297381 | 35430021 | 34813457 | 37455065 | 32411825 | 35603089 | 31406583 | 0 | 1 | 1.25 | 1.14 |
| Q8K1T1 | Lrrc25 | Leucine-rich repeat-containing protein 25 | 53.64 | 6.06 | 32673 | 1 | 654861.69 | 776330.31 | 764671.89 | 721633 | 905862.56 | 624420.81 | 1 | 1 | 1 | 1 | 596315.12 | 595834 | 671994.12 | 608548.25 | 0 | 1 | 0.16 | 0.81 |
| O88188 | Ly86 | Lymphocyte antigen 86 | 309.35 | 43.83 | 17811 | 4 | 5974965.5 | 6362964.5 | 5429174.5 | 3877232.75 | 5239046 | 2333309.25 | 2539961.75 | 4649433.5 | 3992437.5 | 3548827.75 | 3586800.25 | 3616600.5 | 3760531 | 3984031.25 | 2 | 1 | 0.63 | 0.70 |
| P08905 | Lyz2 | Lysozyme C-2 | 403.81 | 49.32 | 16689 | 4 | 23830993 | 30157265 | 25073081 | 23394253 | 22318589 | 19638397 | 21972163 | 22522759 | 15524209 | 19922069 | 20743953 | 17378173 | 21759069 | 23216407 | 1 | 1 | 0.80 | 0.83 |
| Q61166 | Mapre1 | Microtubule-associated protein RP/EB family member 1 | 569.78 | 41.42 | 30016 | 9 | 16125955 | 14248805 | 17313273 | 15565018 | 14982298 | 16313731 | 16824713 | 14834413 | 17621871 | 19727685 | 17574161 | 19524247 | 16654463 | 17687607 | 6 | 1 | 1.09 | 1.14 |
| Q922Q1 | Marc2 | Mitochondrial amidoxime reducing component 2 | 363.39 | 26.04 | 38194 | 3 | 7982832 | 7891377.5 | 7739494 | 9074475 | 10240457 | 10593121 | 9214571 | 9266890 | 4989632 | 11515621 | 12031591 | 9771661 | 12126803 | 10905011 | 7 | 1 | 1.06 | 1.31 |
| Q3THS6 | Mat2a | S-adenosylmethionine synthase isoform type-2 | 436.56 | 25.06 | 43689 | 2 | 25768187 | 26025637 | 27081829 | 21670393 | 25990649 | 20955279 | 21122445 | 17982077 | 20557909 | 21761937 | 20024881 | 19028405 | 21115993 | 22779985 | 1 | 1 | 0.81 | 0.82 |
| Q8VCF0 | Mavs | Mitochondrial antiviral-signaling protein | 442.01 | 21.27 | 53399 | 7 | 6068584 | 6987083 | 7016185.5 | 7826510.5 | 7408904.5 | 7451757 | 7821120 | 6250278.5 | 7432838.5 | 7678173.5 | 10640234 | 8653704 | 8332632 | 7701887 | 8 | 1 | 1.04 | 1.25 |
| P25206 | Mcm3 | DNA replication licensing factor MCM3 | 2118.5 | 59.11 | 91546 | 35 | 71992825 | 70460033 | 64243505 | 77191561 | 76612785 | 66122557 | 65562313 | 53989133 | 58321333 | 65732413 | 60856393 | 64366125 | 61021669 | 60529881 | 3 | 1 | 0.86 | 0.86 |
| P21956 | Mfge8 | Lactadherin | 401.72 | 23.11 | 51241 | 8 | 25970557 | 26130477 | 26266839 | 20592657 | 23680475 | 20323777 | 19623881 | 19144729 | 18653009 | 19247727 | 17701069 | 13561518 | 21956337 | 18465437 | 0 | 1 | 0.79 | 0.73 |
| Q811U4 | Mfn1 | Mitofusin-1 | 55.91 | 1.89 | 83726 | 1 | 438864.47 | 544852.44 | 673544.94 | 489100.62 | 551190.5 | 947622.94 | 888932.44 | 495321.75 | 585021.31 | 694564.19 | 732513.62 | 898191.25 | 632499.06 | 831145.75 | 4 | 1 | 1.34 | 1.43 |
| Q5I0K8 | Mrps7 | 28S ribosomal protein S7, mitochondrial | 71.51 | 4.96 | 28197 | 1 | 2196894.25 | 593319.44 | 1789828.5 | 3287881 | 1966980.798 | 1786291.25 | 1439464.62 | 1229803 | 1790264.62 | 1833157.38 | 1 | 1 | 1236639.25 | 1 | 7 | 1 | 0.82 | 0.16 |
| Q791V5 | Mtch2 | Mitochondrial carrier homolog 2 | 539.84 | 49.17 | 33499 | 11 | 20859653 | 18690925 | 19395263 | 19571011 | 20258059 | 22602769 | 22605763 | 21323819 | 25902417 | 21592661 | 20651543 | 21899347 | 28518225 | 24182107 | 0 | 1 | 1.15 | 1.21 |
| Q80WJ7 | Mtdh | Protein LYRIC | 728.98 | 26.08 | 63846 | 11 | 11561751 | 14092343 | 12164339 | 11791865 | 11385665 | 11422501 | 15940453 | 16759333 | 15058605 | 12527285 | 12288238 | 14431605 | 15826711 | 15495929 | 5 | 1 | 1.18 | 1.19 |
| O35682 | Myadm | Myeloid-associated differentiation marker | 142.12 | 16.25 | 35285 | 3 | 1 | 400232.06 | 1 | 628709.5 | 532174.56 | 446582.5 | 806921.25 | 678009.9375 | 695979.19 | 762556.81 | 542196.5 | 858932.81 | 981764.5 | 794297.9367 | 2 | 1 | 2.17 | 2.54 |
| P70670 | Naca | Nascent polypeptide-associated complex subunit alpha, muscle-specific form | 450.4 | 3.2 | 220499 | 6 | 16618589 | 2569047.5 | 20120981 | 21405269 | 23384415 | 25656853 | 21488095 | 17408961 | 22986857 | 26415465 | 21659267 | 24710441 | 27801965 | 23619005 | 5 | 1 | 1.36 | 1.45 |
| P54921 | Napa | Alpha-soluble NSF attachment protein | 501.16 | 38.31 | 33193 | 9 | 20515501 | 19961555 | 22465731 | 19491621 | 22112343 | 21078087 | 23011023 | 23755503 | 23145177 | 20406511 | 22582059 | 23699813 | 26875043 | 22398461 | 5 | 1 | 1.07 | 1.14 |
| O09043 | Napsa | Napsin-A | 217.39 | 18.62 | 45544 | 4 | 8752844 | 10914111 | 8979887 | 9367902 | 8667265 | 9321483 | 10836249 | 10200148 | 9967983 | 8936050 | 6740373 | 7521599.5 | 7877479 | 8742607 | 8 | 1 | 1.06 | 0.83 |
| P29595 | Nedd8 | NEDD8 | 143.7 | 34.57 | 8972 | 3 | 2820473.75 | 2314128.5 | 2063765 | 2557613.5 | 2498568.5 | 2147223.75 | 2341645.25 | 1868491.5 | 2124641.25 | 2548357.25 | 2259870 | 908756.31 | 942246.5 | 2005464.25 | 7 | 1 | 0.90 | 0.62 |
| Q6DFW4 | Nop58 | Nucleolar protein 58 | 988 | 39.18 | 60343 | 17 | 29170077 | 31379703 | 29777683 | 27070727 | 30653845 | 27886977 | 29497003 | 29480497 | 28375265 | 28285413 | 26309365 | 25949129 | 26671935 | 27507261 | 7 | 1 | 0.97 | 0.90 |
| Q6NSR8 | Npepl1 | Probable aminopeptidase NPEPL1 | 394.99 | 16.6 | 55940 | 6 | 22768453 | 16636672 | 17058027 | 19636999 | 14535225 | 15084541 | 17668589 | 13377728 | 14303742 | 16406094 | 12712750 | 16162779 | 13830483 | 11917200 | 5 | 1 | 0.85 | 0.75 |

| | | | | | | | | | | | | | | | | | | | | | | | | |
|---|---|---|---|---|---|---|---|---|---|---|---|---|---|---|---|---|---|---|---|---|---|---|---|---|
| Q8JZU0 | Nudt13 | Nucleoside diphosphate-linked moiety X motif 13 | 50.78 | 5.68 | 39137 | 1 | 508560.72 | 513306.47 | 533769.44 | 528764.38 | 458402.59 | 1 | 784849.62 | 1 | 604025.19 | 477041.47 | 565302.69 | 529990.94 | 646958.5 | 580750.71 | 11 | 1 | 0.73 | 1.14 |
| Q4KM65 | Nudt21 | Cleavage and polyadenylation specificity factor subunit 5 2 | 385.6 | 49.78 | 26240 | 8 | 738469.69 | 646067.06 | 523869.62 | 985249.62 | 627260.62 | 450886.12 | 676489 | 386052.75 | 1274801.62 | 616907.25 | 475224 | 523894.88 | 430083.38 | 364384.31 | 9 | 1 | 0.97 | 0.64 |
| Q8BVF2 | Pdcl3 | Phosducin-like protein 3 | 303.36 | 32.08 | 27581 | 6 | 4534774 | 4207844.5 | 4094438 | 4482828.5 | 4854951 | 5378127 | 3399928.5 | 3196745 | 4615747.5 | 5216049.5 | 6264950 | 5809101 | 4742143 | 4962010 | 11 | 1 | 0.98 | 1.23 |
| B0BN18 | Pfdn2 | Prefoldin subunit 2 | 233.75 | 29.22 | 16580 | 4 | 12981779 | 15286921 | 14376549 | 16331211 | 13527823 | 10710282 | 13436655 | 14911134 | 12632037 | 13818798 | 13273148 | 12899699 | 11883926 | 12446080 | 6 | 1 | 0.90 | 0.87 |
| Q7TSV4 | Pgm2 | Phosphoglucomutase-2 | 85 | 7.74 | 68748 | 2 | 2201647.75 | 1594923.38 | 2156999.25 | 2122491.75 | 2083694 | 1682543.62 | 1948690.25 | 1 | 1 | 2703991.5 | 1659131.5 | 1415512 | 796951.12 | 846634.75 | 7 | 1 | 0.62 | 0.58 |
| Q8BTI9 | Pik3cb | Phosphatidylinositol 4,5-bisphosphate 3-kinase catalytic subunit beta isoform | 54.26 | 1.5 | 121711 | 1 | 528539.31 | 442406.34 | 1 | 1 | 1 | 598015.9075 | 577891.19 | 577632.44 | 613485.31 | 623054.69 | 497625.53 | 571729.76 | 600266.06 | 617297.69 | 0 | 1 | 3.08 | 2.94 |
| Q5RKH1 | Prpf4b | Serine/threonine-protein kinase PRP4 homolog | 51.07 | 1.69 | 117007 | 1 | 1424606.88 | 1689704.62 | 1048682.38 | 1515060.25 | 1387228.25 | 1244764.12 | 1320071.38 | 1421908.25 | 1220538.62 | 1204857.75 | 953380.31 | 923301.12 | 889005 | 1316950.5 | 6 | 1 | 0.91 | 0.72 |
| Q9R1P4 | Psma1 | Proteasome subunit alpha type-1 | 521.4 | 60.46 | 29547 | 12 | 31595289 | 35210129 | 31054659 | 36645581 | 35170449 | 31328893 | 30190913 | 32691079 | 29411711 | 34062585 | 31041197 | 27388313 | 30424051 | 31240233 | 5 | 1 | 0.93 | 0.88 |
| Q8VDM4 | Psmd2 | 26S proteasome non-ATPase regulatory subunit 2 | 1800.66 | 47.8 | 100203 | 29 | 66501597 | 65012813 | 71737081 | 67672377 | 68472497 | 73770321 | 70062689 | 63874921 | 72961921 | 75627369 | 75782417 | 69639537 | 75584089 | 74630673 | 6 | 1 | 1.05 | 1.09 |
| P97371 | Psme1 | Proteasome activator complex subunit 1 | 975.63 | 57.03 | 28673 | 15 | 89540657 | 95525945 | 95758017 | 75130945 | 81183945 | 75755689 | 74087425 | 86415321 | 86245657 | 78407361 | 69659497 | 77183457 | 70087921 | 73277425 | 6 | 1 | 0.92 | 0.83 |
| P53534 | Pygb | Glycogen phosphorylase, brain form | 997.77 | 23.39 | 96174 | 13 | 38852877 | 40796329 | 42306413 | 36641517 | 39003997 | 44483133 | 41864561 | 43885309 | 46652597 | 42594941 | 43729029 | 43060269 | 42179901 | 44278145 | 1 | 1 | 1.11 | 1.10 |
| Q8BVI4 | Qdpr | Dihydropteridine reductase | 342.47 | 33.61 | 25570 | 5 | 5157324.5 | 5930228.5 | 6655475 | 5754099.5 | 5579340 | 5923568 | 3372950.5 | 6544766 | 6942169 | 6783428 | 6096860.5 | 7802934 | 7056802 | 6839028 | 8 | 1 | 1.02 | 1.19 |
| Q80UJ7 | Rab3gap1 | Rab3 GTPase-activating protein catalytic subunit | 267.42 | 8.87 | 110198 | 6 | 4529544 | 4017258 | 3924811 | 3890960.75 | 3313279.25 | 3953368 | 4103238.5 | 1874881.25 | 3060421 | 3145765 | 3139543.75 | 3064455 | 3872433 | 2804394.5 | 7 | 1 | 0.82 | 0.82 |
| Q8VDV3 | Rab3il1 | Guanine nucleotide exchange factor for Rab-3A | 42.02 | 5.22 | 42713 | 1 | 243104.86 | 142138.69 | 184855.16 | 275147.28 | 229194.58 | 144101.17 | 202274.77 | 204438.53 | 219311.48 | 146518.58 | 176832.36 | 117017.09 | 121782.1 | 138543.85 | 8 | 1 | 0.85 | 0.64 |
| P62835 | Rap1a | Ras-related protein Rap-1A | 974.52 | 76.63 | 20987 | 4 | 121260257 | 124766001 | 110734497 | 116421289 | 112651857 | 110300545 | 117541417 | 121290913 | 107380041 | 113540209 | 109046593 | 100297225 | 112175825 | 108177553 | 9 | 1 | 0.97 | 0.92 |
| Q9CW46 | Raver1 | Ribonucleoprotein PTB-binding 1 | 387.71 | 21.66 | 79382 | 8 | 1 | 541926 | 1 | 1 | 1 | 16877637 | 808436.12 | 699444.5 | 473327.84 | 597908 | 1674871.25 | 524371.38 | 744044.69 | 1242835.5 | 1 | 1 | 35.90 | 9.66 |
| Q9CWY8 | Rnaseh2a | Ribonuclease H2 subunit A | 54.23 | 5.32 | 33513 | 1 | 453792.31 | 325170.91 | 500514.62 | 436523.62 | 482824.31 | 405257.88 | 463679.75 | 323573.22 | 405772.5 | 531020.31 | 561576.88 | 547638.56 | 625447.44 | 485281.5 | 10 | 1 | 0.97 | 1.26 |
| Q9JJ80 | Rpf2 | Ribosome production factor 2 homolog | 333.42 | 27.45 | 35364 | 6 | 1278550 | 1473422 | 1305544.5 | 2019469.38 | 2440933.5 | 895740.5775 | 1150534.88 | 196185.8 | 817088.38 | 1419153.25 | 767881.38 | 1024291.88 | 1101454.75 | 1302622.88 | 2 | 1 | 0.53 | 0.62 |
| P84099 | Rpl19 | 60S ribosomal protein L19 | 308.5 | 18.37 | 23466 | 5 | 2841524.75 | 3748496.25 | 4027587.5 | 3882654.5 | 4441050 | 4410996.5 | 4222289 | 4074969 | 4142975.5 | 5389346.5 | 4766608.5 | 4733162.5 | 4929564.5 | 4162395.25 | 4 | 1 | 1.17 | 1.23 |
| P47964 | Rpl36 | 60S ribosomal protein L36 | 82.39 | 18.1 | 12216 | 1 | 4759367 | 2821063 | 5226443 | 4375304.5 | 4081034.75 | 5004962 | 4172033 | 2621019 | 5024376 | 6551808 | 5349980 | 5856201 | 6328044 | 5137186 | 10 | 1 | 1.10 | 1.33 |
| Q9D8E6 | Rpl4 | 60S ribosomal protein L4 | 1226.3 | 46.54 | 47154 | 20 | 68622793 | 50576013 | 61833941 | 60178089 | 56715809 | 56216201 | 63157993 | 50197481 | 60003061 | 72401361 | 73724481 | 62216353 | 83259225 | 76912265 | 12 | 1 | 1.01 | 1.24 |
| P62908 | Rps3 | 40S ribosomal protein S3 | 1028.02 | 74.07 | 26674 | 20 | 197884705 | 163660033 | 167921617 | 181191601 | 179688833 | 160192993 | 167194129 | 165947825 | 164712465 | 169294337 | 162641201 | 166227921 | 162666801 | 163382225 | 5 | 1 | 0.93 | 0.92 |
| Q9CQS8 | Sec61b | Protein transport protein Sec61 subunit beta | 122.98 | 26.04 | 9958 | 2 | 12811087 | 15127483 | 16301788 | 15294817 | 15382770 | 14942069 | 15933591 | 14303207 | 16023137 | 17472033 | 16996077 | 16117519 | 18071473 | 16337454 | 10 | 1 | 1.05 | 1.13 |
| Q9DCQ7 | Serpina1f | Alpha-1-antitrypsin 1-6 Mus musculus | 33.93 | 1.95 | 46337 | 1 | 5145926 | 4528943.5 | 5142068 | 3530996.75 | 3343327.5 | 1960305.75 | 2068821.88 | 2967468.5 | 1613439.5 | 1921989.38 | 1524142.25 | 2172384.25 | 3404321.25 | 1969269 | 0 | 1 | 0.49 | 0.52 |
| Q99JR1 | Sfxn1 | Sideroflexin-1 | 434.94 | 31.99 | 35649 | 5 | 8041927 | 7853453 | 7753863 | 8344489 | 7582172.5 | 7479436 | 7853814 | 6919617 | 6900343 | 8052574 | 9328649 | 8110641 | 9092919 | 9357381 | 7 | 1 | 0.94 | 1.13 |
| P97797 | Sirpa | Tyrosine-protein phosphatase non-receptor type substrate 1 | 389.84 | 23 | 56425 | 7 | 23094845 | 21341977 | 20353763 | 19189805 | 20057461 | 19577595 | 21475925 | 18487875 | 16792299 | 19625617 | 19875303 | 18915523 | 18498729 | 18524879 | 6 | 1 | 0.92 | 0.91 |
| Q3UND0 | Skap2 | Src kinase-associated phosphoprotein 2 | 634.12 | 60.34 | 40712 | 11 | 24933905 | 26948779 | 22605333 | 23218359 | 24837851 | 24360677 | 25044069 | 27099931 | 25486501 | 22919585 | 22411163 | 23212519 | 20200163 | 20815125 | 9 | 1 | 1.02 | 0.88 |
| O09044 | Snap23 | Synaptosomal-associated protein 23 | 89.9 | 6.67 | 23261 | 1 | 2179954.25 | 2663049.25 | 2291606.75 | 2343632.25 | 1942696.5 | 1913155.38 | 1948148.75 | 2264530 | 1899173.25 | 1916421.5 | 1927433.5 | 1790290 | 1841691.12 | 2130339.5 | 3 | 1 | 0.87 | 0.84 |
| Q62189 | Snrpa | U1 small nuclear ribonucleoprotein A | 295.14 | 31.71 | 31835 | 5 | 5356152.5 | 5656992.5 | 5636798.5 | 5084548 | 5502430 | 4676490.5 | 5325697.5 | 3946715.5 | 3584942.25 | 4370346.5 | 5313226 | 4528311.5 | 3996467.25 | 4890053.5 | 1 | 1 | 0.80 | 0.86 |
| P62307 | Snrpf | Small nuclear ribonucleoprotein F | 229.71 | 83.72 | 9725 | 4 | 1863000 | 1003075.06 | 1714080 | 1687725.12 | 2597979.25 | 2818866.25 | 1291797.5 | 1215975.5 | 2022246.5 | 2745812.75 | 2160090.5 | 2909514.5 | 3622878.75 | 2802040 | 9 | 1 | 1.14 | 1.62 |
| O70493 | Snx12 | Sorting nexin-12 | 245.49 | 44.24 | 19116 | 4 | 7651322 | 6783016 | 6796568.5 | 5799132 | 7286726 | 6579928.5 | 5671096.5 | 6048533.5 | 6780847.5 | 5228042.5 | 3844429 | 6660799.5 | 5211139.5 | 4131628 | 3 | 1 | 0.88 | 0.72 |
| Q8BVL3 | Snx17 | Sorting nexin-17 | 151.8 | 13.62 | 52797 | 4 | 1532372 | 157143.75 | 918231.44 | 1128782.25 | 934132.36 | 1 | 692742.06 | 620969.5 | 1 | 1 | 1 | 619672.94 | 1 | 1 | 2 | 1 | 0.28 | 0.17 |
| Q58A65 | Spag9 | C-Jun-amino-terminal kinase-interacting protein 4 | 403.93 | 9.01 | 146219 | 7 | 4373358.5 | 4123626 | 3886400 | 3164691 | 5128186 | 6293938.5 | 5564882.5 | 6968959 | 6788624.5 | 5099077 | 5801966 | 6487378 | 4663699 | 6591536 | 1 | 1 | 1.49 | 1.42 |
| Q99KH8 | Stk24 | Serine/threonine-protein kinase 24 | 404.54 | 25.99 | 47954 | 8 | 3952477.25 | 5645742.5 | 5255120 | 3935718 | 3924994.5 | 8385537 | 8650342 | 5015629 | 8625960 | 7986652 | 5721885.5 | 5286080.5 | 5779808.5 | 7148376.5 | 2 | 1 | 1.70 | 1.32 |
| P15589 | Sts | Steryl-sulfatase | 37.5 | 1.21 | 62679 | 1 | 18672391 | 18864409 | 19582717 | 18834287 | 19137203 | 20631153 | 17560785 | 17414059 | 17574611 | 18616639 | 19207727 | 19609749 | 20383455 | 20274721 | 5 | 1 | 0.97 | 1.04 |
| P70279 | Surf6 | Surfeit locus protein 6 | 80.44 | 4.79 | 41235 | 2 | 1 | 1 | 371485.03 | 1 | 1 | 1 | 295876.81 | 1 | 1 | 1 | 429008.91 | 322004.53 | 421007.28 | 390673.5733 | 12 | 1 | 0.80 | 5.26 |
| Q921F2 | Tardbp | TAR DNA-binding protein 43 | 930.72 | 50 | 44548 | 14 | 23175409 | 22328111 | 23927153 | 27226111 | 27859395 | 29485829 | 27772005 | 22636725 | 30544047 | 33039991 | 29247515 | 27562755 | 31290785 | 29640989 | 5 | 1 | 1.15 | 1.18 |
| P11983 | Tcp1 | T-complex protein 1 subunit alpha | 1961.24 | 70.14 | 60449 | 29 | 103142201 | 98834041 | 128821737 | 113434849 | 119067593 | 127784233 | 121612873 | 112257777 | 124960121 | 128860705 | 128124137 | 130652729 | 135114353 | 130183217 | 6 | 1 | 1.09 | 1.16 |
| Q921I1 | Tf | Serotransferrin | 40.02 | 2.01 | 76724 | 1 | 307375.94 | 237959.55 | 224392.09 | 422393.12 | 246643.97 | 385051.09 | 329388.19 | 321554.75 | 441268.34 | 347895.31 | 504680.78 | 469763.22 | 356330.72 | 425325.19 | 4 | 1 | 1.27 | 1.53 |
| Q9ERA6 | Tfip11 | Tuftelin-interacting protein 11 | 150.74 | 5.49 | 96305 | 3 | 272489.78 | 336598.19 | 352474.6175 | 357605.84 | 443204.66 | 360103.62 | 351997.09 | 361118.225 | 296120.47 | 436251.72 | 331124 | 206264.83 | 1 | 1 | 10 | 1 | 1.02 | 0.38 |
| P40142 | Tkt | Transketolase | 2163.61 | 67.74 | 67630 | 34 | 278338977 | 278360673 | 292927297 | 297235425 | 310469057 | 319966209 | 285376705 | 298199585 | 327872737 | 315072865 | 307718305 | 325894145 | 323921857 | 314851041 | 4 | 1 | 1.06 | 1.09 |

| | | | | | | | | | | | | | | | | | | | | | | | | |
|---|---|---|---|---|---|---|---|---|---|---|---|---|---|---|---|---|---|---|---|---|---|---|---|---|
| Q6R5N8 | Tlr13 | Toll-like receptor 13 | 647.17 | 13.62 | 114443 | 9 | 12779083 | 9427389 | 12441244 | 11319293 | 11369952 | 8719041 | 8643784 | 9305679 | 9360403 | 6470506.5 | 8340983.5 | 8267912 | 11075327 | 7969123 | 0 | 1 | 0.74 | 0.78 |
| B9EKI3 | Tmf1 | TATA element modulatory factor | 49.99 | 1.37 | 121803 | 1 | 971443.31 | 1054957.62 | 913891.81 | 1043313.94 | 967069.44 | 1310620.88 | 961955.31 | 1104701.88 | 1140436.38 | 922498.94 | 929209.62 | 855882.75 | 735158.38 | 913820.69 | 8 | 1 | 1.10 | 0.87 |
| P49813 | Tmod1 | Tropomodulin-1 | 76.86 | 7.52 | 40466 | 2 | 1126163.25 | 1262612.5 | 1341541.12 | 1705706.75 | 1437078.62 | 1610057.88 | 1377572.5 | 1692447.88 | 1611344.12 | 1576615.88 | 1877508 | 1984288.62 | 1590483.12 | 2002115.38 | 6 | 1 | 1.14 | 1.36 |
| Q61333 | Tnfaip2 | Tumor necrosis factor alpha-induced protein 2 | 61.25 | 1.74 | 78102 | 1 | 404323.91 | 502148.03 | 584102.56 | 275638.88 | 300789.28 | 580202.62 | 414292.94 | 617727.88 | 425619.16 | 412455.81 | 706366.38 | 1122818.25 | 686408.75 | 546814.75 | 7 | 1 | 1.19 | 1.85 |
| Q9CPQ3 | Tomm22 | Mitochondrial import receptor subunit TOM22 homolog | 247.17 | 42.25 | 15537 | 4 | 7626078 | 8556125 | 8900279 | 8178021 | 8377904.5 | 11178517 | 8774223 | 9839948 | 10990508 | 7490971 | 8882940 | 10574362 | 9146619 | 9753334 | 6 | 1 | 1.16 | 1.15 |
| Q62393 | Tpd52 | Tumor protein D52 | 484.65 | 43.3 | 24313 | 7 | 31624619 | 34161057 | 30067463 | 34338641 | 28192053 | 25940747 | 34083297 | 41563769 | 32652609 | 32885201 | 28492837 | 24756509 | 23475313 | 27336077 | 11 | 1 | 1.06 | 0.82 |
| P04692 | Tpm1 | Tropomyosin alpha-1 chain | 483.89 | 17.96 | 32681 | 5 | 181939681 | 212132929 | 173293713 | 206825809 | 185180545 | 185013681 | 181433041 | 195000513 | 165206497 | 181729761 | 171899809 | 166830193 | 147172321 | 180091777 | 7 | 1 | 0.95 | 0.87 |
| P58774 | Tpm2 | Tropomyosin beta chain | 253.82 | 12.68 | 32837 | 0 | 186449921 | 213143153 | 175054945 | 208202273 | 187314593 | 186966097 | 182343873 | 196719025 | 168164801 | 184638769 | 174001953 | 167863729 | 148637569 | 182446385 | 7 | 1 | 0.95 | 0.87 |
| P21107 | Tpm3 | Tropomyosin alpha-3 chain | 643.1 | 32.63 | 32994 | 3 | 274021505 | 324433089 | 267948113 | 309771105 | 276667489 | 270503553 | 271210177 | 297951265 | 251516897 | 269315521 | 249717713 | 252121457 | 222881105 | 270019073 | 6 | 1 | 0.94 | 0.86 |
| P10639 | Txn | Thioredoxin | 439.25 | 46.67 | 11675 | 5 | 103845633 | 105214497 | 117619593 | 56874173 | 76428097 | 117400009 | 101248721 | 145315809 | 136745265 | 104514577 | 111716569 | 128556545 | 119138121 | 121567337 | 6 | 1 | 1.32 | 1.31 |
| Q3TW96 | Uap1l1 | UDP-N-acetylhexosamine pyrophosphorylase-like protein 1 | 793.49 | 36.09 | 56614 | 13 | 52067277 | 56779461 | 59052273 | 56250797 | 61191093 | 57157121 | 61271297 | 50817661 | 58959793 | 56980593 | 63924037 | 60597257 | 64148873 | 62003045 | 11 | 1 | 1.00 | 1.10 |
| P61089 | Ube2n | Ubiquitin-conjugating enzyme E2 N | 301.24 | 56.58 | 17138 | 6 | 12204837 | 13243491 | 12766043 | 11743315 | 13561907 | 10560760 | 12005223 | 12963096 | 10357571 | 12460656 | 11730907 | 11458081 | 12050034 | 10723505 | 6 | 1 | 0.92 | 0.90 |
| Q8VCH8 | Ubxn4 | UBX domain-containing protein 4 | 238.39 | 12.85 | 56472 | 4 | 3544424.25 | 4576210.5 | 3864571 | 3838892.75 | 4312834.5 | 3904445.25 | 3543010.25 | 3288161.75 | 3604221.75 | 3745262 | 3495833 | 3542748.25 | 3743194 | 1994579.25 | 5 | 1 | 0.90 | 0.79 |
| Q99K23 | Ufsp2 | Ufm1-specific protease 2 | 263.03 | 17.14 | 52515 | 6 | 3958302.5 | 4498152 | 4389915.5 | 5304419 | 4965032 | 4432660 | 2965417.25 | 2864233 | 3166457 | 4688020.5 | 5160139 | 5668126.5 | 5633820 | 5559861.5 | 5 | 1 | 0.78 | 1.19 |
| Q6P5E4 | Uggt1 | UDP-glucose:glycoprotein glucosyltransferase 1 | 2310.6 | 40.04 | 176434 | 42 | 55591629 | 46955769 | 47187621 | 46302633 | 51739361 | 58756277 | 41854617 | 48844337 | 51505837 | 56241797 | 58260673 | 62031585 | 62840517 | 53944741 | 9 | 1 | 1.04 | 1.20 |
| Q8VDF2 | Uhrf1 | E3 ubiquitin-protein ligase UHRF1 | 849.41 | 25.45 | 88304 | 11 | 6293807.5 | 2417324 | 6369521.5 | 7107202 | 6277823 | 7188613.5 | 7208659 | 6399633 | 6025060.5 | 7411285.5 | 6708959 | 7836394 | 7355068.5 | 7598416.5 | 5 | 1 | 1.20 | 1.30 |
| Q9EPU0 | Upf1 | Regulator of nonsense transcripts 1 | 769.85 | 16.64 | 123967 | 15 | 4513688.5 | 4762802 | 4437975 | 5356068 | 5269039.5 | 6113806.5 | 5674726 | 4905793.5 | 6596554.5 | 6618581.5 | 5310971 | 8054423.5 | 5380112.5 | 6454206 | 2 | 1 | 1.23 | 1.29 |
| Q9QY76 | Vapb | Vesicle-associated membrane protein-associated protein B | 395.28 | 28.81 | 26946 | 7 | 7922319 | 4367373 | 9264227 | 8058532.5 | 7800590.5 | 8494146 | 7724784 | 6686160.5 | 9549265 | 10734207 | 8546912 | 9518185 | 10804054 | 10221844 | 9 | 1 | 1.15 | 1.31 |
| P20152 | Vim | Vimentin | 3269.56 | 86.48 | 53688 | 48 | 2434394369 | 2796715521 | 2589878017 | 2831042049 | 2816222721 | 3398969601 | 3163222273 | 2930145793 | 3181000961 | 3520160769 | 2958590465 | 2826093825 | 2985940737 | 3154718721 | 0 | 1 | 1.20 | 1.11 |
| Q3UVL4 | Vps51 | Vacuolar protein sorting-associated protein 51 homolog | 329.73 | 12.02 | 86187 | 6 | 6871667 | 7778310.5 | 6939899 | 6797230 | 6580675 | 6280603 | 6998185.5 | 7282023.5 | 6867556 | 5801770 | 4702605.5 | 6611993 | 6193673 | 6547823 | 10 | 1 | 0.95 | 0.86 |
| Q9JKB3 | Ybx3 | Y-box-binding protein 3 | 414.99 | 26.87 | 38814 | 3 | 2490044.5 | 1521259.38 | 2400492.22 | 2960451.75 | 2630213.25 | 383667.94 | 453809 | 1 | 1 | 1 | 2128008.75 | 1014960.813 | 409303.44 | 507570.25 | 0 | 1 | 0.07 | 0.42 |
| O35986 | Zranb2 | Zinc finger Ran-binding domain-containing protein 2 | 55.42 | 5.45 | 37350 | 2 | 2275240 | 1912361.75 | 2048002.38 | 2818413.75 | 396779.72 | 2465725.25 | 2891881.25 | 2404159.75 | 2859773.25 | 3414188.75 | 2771167.75 | 3795330.5 | 3466864.25 | 3270741.75 | 2 | 1 | 1.49 | 1.76 |
