## Supplemental Table 3 for "Effects of polystyrene and polylactide nanoparticles on macrophages under a repeated exposure mode"

Supplementary table 3: List of modulated proteins in response to PLA  particles, selected by the Mann Whitney U test (p<0.05)

| accession | gene_name | description | protein_set_score | coverage | MW | pecific_peptid | Ctrl1 rep | Ctrl5 rep | Ctrl4 rep | Ctrl3 rep | Ctrl2 rep | PLA1 rep | PLA5 rep | PLA4 rep | PLA3 rep | PLA2 rep | PS1 rep | PS5 rep | PS3 rep | PS2 rep | U-PLA | U-PS | ratio PLA | ratio PS |
|---|---|---|---|---|---|---|---|---|---|---|---|---|---|---|---|---|---|---|---|---|---|---|---|---|
| Q9DBL1 | Acadsb | Short/branched chain specific acyl-CoA dehydrogenase, mitochondrial | 542.63 | 33.8 | 47874 | 9 | 9718412 | 9999263 | 12876759 | 11010008 | 9727089 | 13291001 | 14581746 | 14659703 | 13334390 | 13814395 | 13384975 | 10085830 | 14800241 | 13585437 | 0 | 2 | 1.30656731 | 1.215427394 |
| Q99KI0 | Aco2 | Aconitate hydratase, mitochondrial | 1715.58 | 40.38 | 85464 | 27 | 113500217 | 122642265 | 125108009 | 117830081 | 128722697 | 132302993 | 136039729 | 129805801 | 140362865 | 142224657 | 130724961 | 130182385 | 142546593 | 139967137 | 0 | 0 | 1.11999405 | 1.117592451 |
| P54987 | Acod1 | Cis-aconitate decarboxylase | 98.01 | 6.97 | 53758 | 2 | 516474.31 | 509054.03 | 870485.12 | 464057.28 | 567807.75 | 245930.42 | 232549.61 | 453477.31 | 220441.86 | 187289.31 | 236652.28 | 371332.25 | 532760.5 | 219037.17 | 0 | 3 | 0.457562879 | 0.580532203 |
| Q9QUJ7 | Acsl4 | Long-chain-fatty-acid--CoA ligase 4 | 1179.65 | 41.63 | 79077 | 22 | 77413993 | 74544825 | 73041393 | 67446297 | 74989745 | 62402901 | 57477041 | 59842265 | 57744017 | 56595649 | 76577297 | 82590777 | 63239773 | 76564017 | 0 | 7 | 0.800307184 | 1.017087527 |
| Q8CG72 | Adprhl2 | Poly(ADP-ribose) glycohydrolase ARH3 | 177.7 | 15.68 | 39414 | 4 | 3002238 | 4763269.5 | 4780709.5 | 4387344 | 4546301 | 4816668 | 5367212.5 | 5522041 | 5734347.5 | 5159242 | 4312480 | 4467942.5 | 4820463 | 4118514.75 | 0 | 9 | 1.238346457 | 1.031163529 |
| Q9QZQ1 | Afdn | Afadin | 78.26 | 0.99 | 206499 | 1 | 997290.12 | 1019074 | 1066013.75 | 898114.5 | 978138.25 | 1167800 | 1200748.75 | 1225231.38 | 1311944.25 | 1150000.25 | 963122.81 | 1358992.62 | 1119267 | 1147127.477 | 0 | 4 | 1.221249392 | 1.156697851 |
| Q9WTP6 | Ak2 | Adenylate kinase 2, mitochondrial | 315.83 | 33.89 | 26469 | 6 | 20322113 | 21748331 | 21522219 | 22517561 | 21268597 | 24824239 | 24086941 | 25325895 | 23625907 | 26267657 | 24645387 | 24001141 | 23193215 | 24466869 | 0 | 0 | 1.156006723 | 1.121108091 |
| P45376 | Akr1b1 | Aldose reductase | 841.51 | 52.53 | 35732 | 15 | 190222113 | 212004353 | 199337377 | 196657073 | 209078065 | 161110577 | 174705681 | 163635473 | 179064801 | 182479457 | 194898833 | 208309905 | 201504657 | 194774241 | 0 | 8 | 0.854757133 | 0.992118094 |
| P47738 | Aldh2 | Aldehyde dehydrogenase, mitochondrial | 1506.28 | 68.21 | 56538 | 24 | 182985393 | 204331585 | 209732513 | 200265169 | 210116641 | 223336913 | 224602097 | 222453585 | 231237777 | 223844209 | 247455457 | 248172561 | 250940065 | 254757489 | 0 | 0 | 1.117172536 | 1.242424137 |
| Q80VQ0 | Aldh3b1 | Aldehyde dehydrogenase family 3 member B1 | 105.23 | 8.55 | 52292 | 2 | 2891052.25 | 3489360.75 | 2962967.5 | 2642890.75 | 2990492.75 | 1797691.25 | 1787047.38 | 1641571.12 | 1986077.5 | 1821045.38 | 1948526.25 | 2216548.5 | 2892468.5 | 2791471.75 | 0 | 3 | 0.603163182 | 0.822024621 |
| P05063 | Aldoc | Fructose-bisphosphate aldolase C | 1570.54 | 70.25 | 39395 | 16 | 264098593 | 255447873 | 239960209 | 228017905 | 262386721 | 291617729 | 297236609 | 293018529 | 308779137 | 315111137 | 293155009 | 242416257 | 242333217 | 276175521 | 0 | 6 | 1.204695997 | 1.054154806 |
| P07356 | Anxa2 | Annexin A2 | 1632.99 | 70.8 | 38676 | 25 | 413457761 | 402877729 | 423519713 | 427536897 | 441378049 | 471697665 | 521019233 | 505123873 | 568528513 | 556714433 | 470546401 | 450212353 | 440189633 | 446280225 | 0 | 1 | 1.243892663 | 1.071257465 |
| O88512 | Ap1g2 | AP-1 complex subunit gamma-like 2 | 65.18 | 2.4 | 87863 | 2 | 887665.88 | 570473.915 | 515224.12 | 454511.97 | 424493.69 | 312314.88 | 294088.03 | 1 | 364976.81 | 1 | 629513.94 | 475944.62 | 470210.22 | 448755.78 | 0 | 9 | 0.340552546 | 0.887167891 |
| Q3UIA2 | Arhgap17 | Rho GTPase-activating protein 17 | 444.6 | 14.89 | 92202 | 10 | 12156895 | 8757226 | 8017522 | 8870471 | 10846009 | 7692902 | 7969897.5 | 6999208 | 7397576 | 7272889 | 8430568 | 6067848.5 | 7748019 | 5477966 | 0 | 1 | 0.767398004 | 0.712370791 |
| Q61024 | Asns | Asparagine synthetase [glutamine-hydrolyzing] | 807.35 | 33.51 | 64283 | 14 | 22282757 | 16045791 | 22277965 | 22302875 | 22211675 | 34332637 | 24759677 | 23762539 | 22689153 | 25055077 | 23824953 | 28357385 | 23592073 | 24117823 | 0 | 0 | 1.242368363 | 1.187823724 |
| Q3KRE0 | Atad3 | ATPase family AAA domain-containing protein 3 | 182.8 | 7.28 | 66759 | 3 | 1 | 1 | 153757.5 | 1 | 196889.88 | 260729.42 | 205999.27 | 294182.22 | 242422.42 | 282221.62 | 531651.44 | 354055.62 | 360888.22 | 169627.2 | 0 | 1 | 3.666201502 | 5.048556057 |
| Q6AZ50 | Atg3 | Ubiquitin-like-conjugating enzyme ATG3 | 117.78 | 10.51 | 35822 | 3 | 836774.06 | 1038730.56 | 2332083 | 2298163.25 | 2343778 | 2865089.25 | 2437585.5 | 2873648.25 | 2651260 | 2509774.25 | 2322526.5 | 1251042.62 | 2246637.5 | 1649689.5 | 0 | 9 | 1.507126249 | 1.055126243 |
| Q9CWJ9 | Atic | Bifunctional purine biosynthesis protein PURH | 2297.18 | 75.68 | 64217 | 34 | 100501897 | 103995289 | 114436017 | 101709217 | 103917497 | 91965265 | 92060817 | 90759465 | 92896721 | 97746585 | 109017473 | 106490857 | 107272081 | 106332249 | 0 | 4 | 0.88727491 | 1.022553969 |
| Q6PA06 | Atl2 | Atlastin-2 | 92.44 | 4.12 | 66224 | 2 | 733836.25 | 899531.75 | 923835.75 | 877421.19 | 803702.88 | 1116392.38 | 1110131 | 1190888.5 | 1285978.75 | 933306.44 | 1116237.38 | 1200585 | 1236324 | 1006278.12 | 0 | 0 | 1.329934189 | 1.344700284 |
| O54715 | Atp6ap1 | V-type proton ATPase subunit S1 Rattus norvegicus | 33.07 | 1.51 | 51123 | 1 | 2637324.75 | 2791397.5 | 3484690.5 | 3701668.75 | 3812607.75 | 2374916.25 | 2613749.75 | 2442275.5 | 2349059.5 | 2525420.5 | 2283052 | 2530357.75 | 2638727.75 | 2887804.25 | 0 | 3 | 0.74906588 | 0.786776947 |
| O08539 | Bin1 | Myc box-dependent-interacting protein 1 | 172.38 | 8.33 | 64470 | 2 | 1466632.25 | 1737278.88 | 1662260.75 | 1233827.25 | 1523102 | 1929133.5 | 1888381 | 1993632.88 | 2033967.75 | 2255730.25 | 3599544.75 | 3257452.75 | 1773316.38 | 1957549.25 | 0 | 0 | 1.325031009 | 1.736147624 |
| Q9CY64 | Blvra | Biliverdin reductase A | 753.23 | 58.64 | 33525 | 14 | 48784881 | 54583561 | 49228573 | 50434457 | 51295057 | 44310549 | 44582985 | 47738089 | 44683173 | 44107165 | 44423617 | 40271645 | 43850857 | 43421633 | 0 | 0 | 0.886348592 | 0.845211433 |
| P98086 | C1qa | Complement C1q subcomponent subunit A | 50.02 | 6.12 | 25974 | 1 | 476973.56 | 391320.94 | 672807 | 879280.31 | 652380.44 | 382105.22 | 344430.47 | 249674.16 | 315717.97 | 375473.78 | 431360.09 | 385306.88 | 479654.56 | 351893.84 | 0 | 3 | 0.542639314 | 0.670494182 |
| Q80X80 | C2cd2l | Phospholipid transfer protein C2CD2L | 49.59 | 1.42 | 76329 | 1 | 1 | 1 | 349341.56 | 1 | 1 | 505218.97 | 495357.28 | 569019.19 | 693052.94 | 364224.81 | 446286.72 | 497738.44 | 511167.75 | 485064.3033 | 0 | 0 | 7.519411983 | 6.942471279 |
| Q6PHZ2 | Camk2d | Calcium/calmodulin-dependent protein kinase type II subunit delta | 490.44 | 26.45 | 56369 | 9 | 15552478 | 16226495 | 16876161 | 17452865 | 16448511 | 21893837 | 19909357 | 21355475 | 21208995 | 21389053 | 19836729 | 19635099 | 19042353 | 19131423 | 0 | 0 | 1.281022139 | 1.175643265 |
| P29452 | Casp1 | Caspase-1 | 890.3 | 41.29 | 45640 | 16 | 26517867 | 27996515 | 28144805 | 26615815 | 29227735 | 22356923 | 23036571 | 23089263 | 23713299 | 22791975 | 26056177 | 24088509 | 22483423 | 26143101 | 0 | 0 | 0.830222084 | 0.891419297 |
| P10810 | Cd14 | Monocyte differentiation antigen CD14 | 919.15 | 51.37 | 39204 | 13 | 46938701 | 51451925 | 49639101 | 46202521 | 44246909 | 37399081 | 39845889 | 41435497 | 33770573 | 39076653 | 33665017 | 33582077 | 38490937 | 30320721 | 0 | 0 | 0.80312131 | 0.713158509 |
| Q9JLQ0 | Cd2ap | CD2-associated protein | 209.12 | 8.32 | 70450 | 4 | 2313159 | 1819108.5 | 2543226 | 1921423.62 | 2484053.5 | 2692327 | 3158640.25 | 3662151.75 | 2628092 | 2563852 | 2924724.25 | 3169045.75 | 2384775.5 | 3175545.25 | 0 | 2 | 1.327055499 | 1.314651391 |
| P15379 | Cd44 | CD44 antigen | 248.12 | 4.5 | 85617 | 4 | 21419985 | 27528795 | 25653127 | 22303749 | 26399949 | 29994273 | 28164401 | 30882579 | 28157001 | 27638877 | 24428085 | 23396093 | 25576425 | 27238867 | 0 | 10 | 1.174619199 | 1.020224 |
| P21855 | Cd72 | B-cell differentiation antigen CD72 | 200.92 | 13.28 | 40347 | 4 | 5472007 | 4998917 | 4320119.5 | 4838255 | 4878189.5 | 5775861.5 | 6823931 | 5990614 | 6522884 | 5704567 | 5120527 | 5111056 | 4680059 | 5577287.5 | 0 | 6 | 1.257487405 | 1.045034149 |
| P42082 | Cd86 | T-lymphocyte activation antigen CD86 | 46.91 | 4.53 | 34666 | 1 | 3228302 | 3338046.5 | 3090599.25 | 3026812.25 | 3151998.25 | 4316440 | 4112833.25 | 4330235.5 | 3371236.5 | 3920199.25 | 3515333 | 3138004.25 | 3088859.75 | 3636546.5 | 0 | 7 | 1.266181523 | 1.056054855 |
| Q8BTU1 | Cfap20 | Cilia- and flagella-associated protein 20 | 173.2 | 30.05 | 22748 | 4 | 7629927 | 8499859 | 8663983 | 7950704 | 6956509.5 | 6192437 | 5840535 | 6917020.5 | 4997719 | 5935685 | 4541339 | 6196100 | 4910781 | 5646966.5 | 0 | 0 | 0.752711762 | 0.67048676 |
| Q8CGZ0 | Cherp | Calcium homeostasis endoplasmic reticulum protein | 102.33 | 5.02 | 106169 | 3 | 1 | 1 | 1 | 1 | 1 | 632533.905 | 536792 | 640966.12 | 981653.81 | 370723.69 | 509218.72 | 1 | 1 | 795864.56 | 0 | 5 | 632533.905 | 326271.32 |
| D4A4T9 | Chordc1 | Cysteine and histidine-rich domain-containing protein 1 Rattus norvegicus | 436.58 | 35.65 | 37362 | 9 | 8176075 | 9282825 | 8180874.5 | 8925010 | 7551701 | 10693211 | 15437735 | 9469615 | 10953471 | 11913802 | 11772235 | 8170458.5 | 9848022 | 11920077 | 0 | 4 | 1.388241049 | 1.237959198 |
| Q6MG61 | Clic1 | Chloride intracellular channel protein 1 | 1123.47 | 62.66 | 26981 | 14 | 202635617 | 232196769 | 213661681 | 198470033 | 209748017 | 235330193 | 247376785 | 239691601 | 249050065 | 246995137 | 221458097 | 228320849 | 229401105 | 240769345 | 0 | 3 | 1.153051774 | 1.088221405 |
| Q9QYB1 | Clic4 | Chloride intracellular channel protein 4 | 1131.61 | 74.31 | 28729 | 17 | 97964481 | 99478145 | 87956129 | 100850505 | 100354753 | 120676057 | 119147137 | 113144737 | 129646161 | 139642721 | 116781297 | 118015865 | 113190041 | 127598233 | 0 | 0 | 1.27877452 | 1.221695216 |
| O08585 | Clta | Clathrin light chain A | 123.19 | 10.64 | 25604 | 3 | 22239505 | 23305617 | 20524421 | 27492887 | 23438983 | 19110209 | 19091705 | 19940957 | 16651577 | 19197859 | 20148757 | 17940843 | 18731121 | 19499133 | 0 | 0 | 0.803343349 | 0.815373208 |
| Q8C166 | Cpne1 | Copine-1 | 608.36 | 28.36 | 58887 | 10 | 17995381 | 17383457 | 16692100 | 21040703 | 19500999 | 29652387 | 22701409 | 25226597 | 24229393 | 25495135 | 19475509 | 18864543 | 18260365 | 18881301 | 0 | 8 | 1.37459553 | 1.018782614 |
| Q9WVJ3 | Cpq | Carboxypeptidase Q | 159.27 | 9.79 | 51813 | 3 | 4235704.5 | 5433789.5 | 3963128 | 4633763.5 | 4864550 | 2740996.75 | 3432032.25 | 3105766.25 | 3189807.75 | 2950120 | 3112566.25 | 3491155.25 | 2712508.75 | 3580672 | 0 | 0 | 0.666584497 | 0.696950965 |
| Q62426 | Cstb | Cystatin-B | 145.46 | 57.14 | 11046 | 3 | 7521179 | 7137837.5 | 6904083 | 12055766 | 6465746.5 | 3031240.75 | 5190731 | 2957673 | 4015037.5 | 4353343.5 | 4867321 | 3217625.75 | 5260377 | 2877489 | 0 | 0 | 0.487669077 | 0.505892783 |
| P97821 | Ctsc | Dipeptidyl peptidase 1 | 595.33 | 16.67 | 52376 | 7 | 49376273 | 52577253 | 45175817 | 45652129 | 52748553 | 36184393 | 32687601 | 37102473 | 32423235 | 31497689 | 37962265 | 34302401 | 38151173 | 39475241 | 0 | 0 | 0.691953626 | 0.763099544 |
| P18242 | Ctsd | Cathepsin D | 1308.71 | 65.37 | 44954 | 21 | 406401761 | 489922369 | 425776257 | 393210113 | 412787073 | 566899457 | 570130689 | 552964993 | 524020897 | 533450113 | 536090305 | 506827169 | 585553345 | 525133889 | 0 | 0 | 1.291043317 | 1.264982358 |
| O70370 | Ctss | Cathepsin S | 809.73 | 49.12 | 38475 | 10 | 238743409 | 309175649 | 274737185 | 216478641 | 242877905 | 188496833 | 181920641 | 208634577 | 183625857 | 167515569 | 163645185 | 144368785 | 225630465 | 196082705 | 0 | 1 | 0.725572697 | 0.711505324 |
| Q501J6 | Ddx17 | Probable ATP-dependent RNA helicase DDX17 | 957.64 | 31.54 | 72399 | 11 | 150077089 | 155918529 | 158525361 | 155182561 | 152922897 | 136444433 | 140492241 | 146613361 | 135731617 | 144161297 | 143302945 | 132117089 | 146279233 | 155489377 | 0 | 3 | 0.910456742 | 0.933809368 |

| | | | | | | | | | | | | | | | | | | | | | | | | |
|---|---|---|---|---|---|---|---|---|---|---|---|---|---|---|---|---|---|---|---|---|---|---|---|---|
| O08560 | Dgkz | Diacylglycerol kinase zeta | 71.91 | 4.41 | 104009 | 2 | 1274328.25 | 1207170.12 | 1073381.62 | 1266015.88 | 1295138.75 | 1424261.38 | 1702466.12 | 1422879.62 | 1570516.88 | 1423209.62 | 1703564.5 | 1523170.38 | 1465322.62 | 1469373.12 | 0 | 0 | 1.23337 | 1.25927807 |
| Q99J87 | Dhx58 | Probable ATP-dependent RNA helicase DHX58 | 65.33 | 1.62 | 76709 | 1 | 189459.12 | 153219.42 | 177991.3725 | 204719.97 | 164566.98 | 1 | 1 | 1 | 1 | 1 | 1 | 1 | 173103.42 | 1 | 0 | 2 | 5.61825E-06 | 0.243138779 |
| F1M775 | Diaph1 | Protein diaphanous homolog 1 | 716.21 | 15.57 | 140409 | 14 | 8559447 | 9502858 | 9598082 | 8899180 | 9757611 | 11403285 | 10264303 | 10886076 | 10530646 | 11004361 | 10484697 | 7826268.5 | 9353816 | 10188028 | 0 | 8 | 1.167788569 | 1.021565085 |
| P39052 | Dnm2 | Dynamin-2 | 1083.22 | 34.25 | 98230 | 15 | 24288561 | 25117165 | 23822793 | 25118757 | 24548741 | 29560125 | 27019033 | 27048409 | 29030973 | 26399053 | 24601069 | 24540613 | 24127935 | 27190765 | 0 | 9 | 1.131506101 | 1.021802663 |
| Q9R0P5 | Dstn | Destrin | 489.98 | 52.12 | 18522 | 7 | 29927785 | 31798485 | 33788209 | 31869721 | 32292155 | 36284041 | 34494133 | 34565989 | 34277933 | 40759389 | 39374597 | 35627793 | 34732637 | 44021397 | 0 | 0 | 1.129669355 | 1.203656797 |
| Q3TGW2 | Eepd1 | Endonuclease/ exonuclease/ phosphatase family domain-containing protein 1 | 41.23 | 2.64 | 62953 | 1 | 884563.31 | 910401.94 | 1 | 646673 | 1 | 1015498.62 | 1052696.62 | 1135702.12 | 1087941.25 | 932121.44 | 1332034.5 | 1077017.25 | 957574.06 | 1058873.5 | 0 | 0 | 2.13952897 | 2.265638493 |
| P59325 | Eif5 | Eukaryotic translation initiation factor 5 | 290.59 | 13.29 | 48968 | 5 | 10179051 | 8915562 | 9307121 | 9392106 | 10261104 | 10624993 | 10484176 | 10996545 | 10274364 | 10722081 | 10795490 | 10582504 | 11922213 | 11142316 | 0 | 0 | 1.105030088 | 1.156034096 |
| Q6P6T4 | Eml2 | Echinoderm microtubule-associated protein-like 2 | 151.15 | 7.7 | 70711 | 3 | 5018700.938 | 6989340 | 7348600 | 2637560.75 | 3099303 | 9606035 | 8795315 | 7381280 | 8336462 | 7777409 | 3748214.75 | 9723933 | 7177481 | 4100047 | 0 | 7 | 1.669615365 | 1.232872613 |
| Q3UMY5 | Eml4 | Echinoderm microtubule-associated protein-like 4 | 82.13 | 1.62 | 110027 | 1 | 1 | 1 | 1 | 1 | 1 | 733450.0475 | 834772.44 | 603441.25 | 692215.75 | 803370.75 | 600970.06 | 620397.03 | 799164.06 | 461056.97 | 0 | 0 | 733450.0475 | 620397.03 |
| P17182 | Eno1 | Alpha-enolase | 2394.93 | 77.42 | 47141 | 26 | 483027841 | 544320321 | 569124609 | 442800673 | 497630657 | 576118465 | 683387137 | 669707137 | 704570817 | 678481153 | 568724865 | 618923585 | 630460033 | 581501569 | 0 | 1 | 1.305632605 | 1.182351577 |
| Q8R180 | Ero1a | ERO1-like protein alpha | 1208.87 | 53.66 | 54084 | 17 | 34275897 | 37585565 | 33318769 | 30588371 | 32604637 | 52807873 | 55697817 | 67111121 | 56419889 | 53866457 | 31954029 | 34456857 | 36277085 | 37950649 | 0 | 6 | 1.698032055 | 1.044098671 |
| Q8R2E9 | Ero1b | ERO1-like protein beta | 314.36 | 12.85 | 53518 | 3 | 6508726 | 6918732 | 7956128 | 5295514.5 | 5025839.5 | 8813798 | 11036895 | 12714955 | 10234134 | 8538276 | 5906597.5 | 6736327.5 | 7119441 | 7604306.5 | 0 | 7 | 1.619244761 | 1.078959324 |
| Q8CGI1 | Fam193a | Protein FAM193A | 35.38 | 0.81 | 136714 | 1 | 4412990 | 2779078.25 | 2425787 | 4287065.5 | 4312787 | 1263273.88 | 2120620.75 | 1657398.75 | 1911461.62 | 1650282.5 | 4412990 | 4386696.5 | 762691.62 | 3370027.5 | 0 | 9.5 | 0.472234906 | 0.88735132 |
| P26151 | Fcgr1 | High affinity immunoglobulin gamma Fc receptor I | 275.57 | 22.77 | 44888 | 6 | 10544918 | 9299983 | 11888508 | 9437379 | 9035087 | 7037936 | 6074016 | 6070993.5 | 7317976 | 5974343 | 6363764 | 6071877.5 | 7110427 | 4110293.75 | 0 | 0 | 0.646841918 | 0.588983915 |
| D3ZYR1 | Fcho2 | F-BAR domain only protein 2 | 115.95 | 3.83 | 88712 | 2 | 830894.94 | 883108.62 | 884735.75 | 744802.12 | 843027.44 | 1208799.88 | 1217656.62 | 985770.38 | 1326516 | 1219684.62 | 1048347.88 | 729531.06 | 865548 | 930317.62 | 0 | 7 | 1.423224527 | 1.067026684 |
| P84817 | Fis1 | Mitochondrial fission 1 protein | 381.05 | 60.53 | 16995 | 6 | 14938037 | 9948465 | 9113640 | 8535140 | 8707892 | 6092679.5 | 6198266.5 | 6328685 | 7185594.5 | 7633101 | 8499026 | 6001722 | 9117733 | 6506754 | 0 | 3 | 0.652542063 | 0.734859705 |
| P45878 | Fkbp2 | Peptidyl-prolyl cis-trans isomerase FKBP2 | 125.57 | 19.29 | 15344 | 2 | 3244866 | 3025755.5 | 3779809.5 | 4243410.5 | 2996151 | 2476533.25 | 2402041.25 | 2739740 | 1806613.25 | 2899214 | 3283477.75 | 2750951 | 2478876.5 | 2652518.5 | 0 | 3 | 0.712790462 | 0.807246139 |
| Q8BTM8 | Flna | Filamin-A | 5048.24 | 49.87 | 281222 | 81 | 369365921 | 351985377 | 418238977 | 354114593 | 340416193 | 299507617 | 332472097 | 274723489 | 279353921 | 326676577 | 278621537 | 289579969 | 383951745 | 319704929 | 0 | 4 | 0.824773093 | 0.866803593 |
| Q80X90 | Flnb | Filamin-B | 556.33 | 5.88 | 277825 | 8 | 16415002 | 14864654 | 17498361 | 15034736 | 14546203 | 12402659 | 10174523 | 11341972 | 11000152 | 11902811 | 13906003 | 14000077 | 19377567 | 14977597 | 0 | 7 | 0.725151532 | 0.993205614 |
| Q9R0N0 | Galk1 | Galactokinase | 453.75 | 37.5 | 42295 | 10 | 8279408 | 12743603 | 18287175 | 10402457 | 10066241 | 19555699 | 18757369 | 19496847 | 22014131 | 22097199 | 11510724 | 12738829 | 18303935 | 19423973 | 0 | 4 | 1.704970688 | 1.295973111 |
| Q9CZD3 | Gars | Glycine--tRNA ligase | 1395.78 | 37.59 | 81878 | 3 | 96797529 | 94607849 | 95651737 | 99651545 | 93605913 | 109506609 | 108360201 | 106143361 | 109224985 | 107692209 | 102365249 | 99154657 | 106202961 | 101447001 | 0 | 1 | 1.126193947 | 1.064848672 |
| P17439 | Gba | Glucosylceramidase | 288.44 | 20.39 | 57622 | 6 | 7411030.5 | 6799296.5 | 6604925 | 8099719.5 | 6469604 | 5052369.5 | 6231565 | 5997354 | 5270157 | 4826628 | 7181936 | 6700892.5 | 7041698 | 6856018.5 | 0 | 9 | 0.773729036 | 0.981379055 |
| Q61107 | Gbp4 | Guanylate-binding protein 4 | 569.72 | 26.13 | 70801 | 11 | 8514483 | 7450088 | 5133596 | 6443512 | 6430460 | 3197550.5 | 4450390.5 | 3241343.75 | 2997324.5 | 2882868.25 | 3562685 | 2570989.5 | 3680811.25 | 3870607.75 | 0 | 0 | 0.493624423 | 0.50354106 |
| Q8BX17 | Gemin5 | Gem-associated protein 5 | 193.13 | 4.53 | 166592 | 5 | 3853023 | 3625824.5 | 3026013.5 | 2197521.5 | 2166582.25 | 1310570.88 | 1604851.12 | 2008812.38 | 1628851.62 | 1231911.75 | 1028084.69 | 710402 | 1677707.25 | 940815.56 | 0 | 0 | 0.523573623 | 0.366283865 |
| Q9D1H7 | Get4 | Golgi to ER traffic protein 4 homolog | 133.38 | 12.23 | 36525 | 3 | 2727767 | 2513847.5 | 2044945.5 | 2748242.75 | 2697812.5 | 2023680.75 | 1667199.25 | 1509719.38 | 1007143.12 | 1561593 | 1789479.75 | 3348092 | 1421113.75 | 2742497.25 | 0 | 9 | 0.610191649 | 0.913125718 |
| Q9CPU0 | Glo1 | Lactoylglutathione lyase | 441.84 | 46.74 | 20810 | 8 | 22031245 | 26920293 | 19755439 | 26164569 | 24276729 | 29083327 | 29703499 | 28965929 | 30514657 | 28386367 | 26571287 | 29813259 | 23432779 | 24923859 | 0 | 6 | 1.230851047 | 1.098853341 |
| Q8BTZ7 | Gmppb | Mannose-1-phosphate guanyltransferase beta | 49.39 | 5 | 39917 | 1 | 4203369.5 | 4833290.5 | 4242727.5 | 4922349.5 | 5575047 | 3916547.25 | 3445408.25 | 2720064.25 | 2685379.5 | 2948363 | 4816802 | 3610994.75 | 3373055.75 | 4345794 | 0 | 4 | 0.660970897 | 0.848866193 |
| P63213 | Gng2 | Guanine nucleotide-binding protein G(I)/G(S)/G(O) subunit gamma-2 | 83.93 | 39.44 | 7850 | 2 | 6892465 | 7611842 | 7428047 | 8924962 | 8298338.5 | 5637436 | 6061224.5 | 6645098 | 5762194 | 5795699 | 6408208 | 5934865 | 5218212 | 6679262.5 | 0 | 0 | 0.763661134 | 0.773852072 |
| Q99P91 | Gpnmb | Transmembrane glycoprotein NMB | 299.24 | 10.45 | 63676 | 6 | 15079978 | 18625543 | 14805663 | 15678684 | 18425977 | 28292055 | 26069569 | 21835903 | 21971033 | 23690629 | 24751743 | 21765275 | 28238209 | 24268953 | 0 | 0 | 1.475009897 | 1.498262531 |
| P14426 | H2-D1 | H-2 class I histocompatibility antigen, D-K alpha chain | 228.67 | 9.67 | 40620 | 0 | 76935561 | 88768785 | 78007345 | 74458721 | 84181505 | 70632345 | 62804249 | 67617417 | 65458549 | 67753905 | 73245777 | 63252033 | 74996817 | 76354569 | 0 | 2 | 0.830781341 | 0.894270612 |
| P01901 | H2-K1 | H-2 class I histocompatibility antigen, K-B alpha chain | 226.74 | 9.21 | 41302 | 0 | 69263969 | 77587809 | 70712513 | 67576433 | 76025153 | 63816409 | 58201057 | 59927801 | 59760469 | 61931045 | 66310081 | 57656105 | 69091313 | 68949009 | 0 | 2 | 0.840712815 | 0.906808079 |
| P06339 | H2-T23 | H-2 class I histocompatibility antigen, D-37 alpha chain | 236.87 | 10.64 | 40875 | 0 | 49192085 | 52357897 | 48947249 | 45976805 | 54361713 | 43278765 | 40954385 | 41889553 | 40893653 | 43155413 | 44860493 | 38484013 | 47614577 | 48557841 | 0 | 2 | 0.837886026 | 0.894593996 |
| Q8BY71 | Hat1 | Histone acetyltransferase type B catalytic subunit | 340.62 | 24.52 | 49278 | 7 | 5537363.5 | 4287962 | 3822267.25 | 4729448 | 4515848 | 3592670.5 | 3325248.5 | 3529465 | 3122781 | 2976022 | 4724355.5 | 4819829 | 3491253 | 3809642.5 | 0 | 7 | 0.722765361 | 0.919776889 |
| P17710 | Hk1 | Hexokinase-1 | 1409.73 | 30.6 | 108303 | 4 | 51614469 | 58252017 | 56549513 | 52917917 | 53608305 | 59347841 | 66062785 | 61506405 | 65622465 | 66010489 | 54835997 | 56745281 | 54734325 | 58304449 | 0 | 5 | 1.167096772 | 1.02869781 |
| O08528 | Hk2 | Hexokinase-2 | 1259.89 | 33.04 | 102535 | 18 | 36978961 | 40303117 | 41025301 | 33905165 | 36865553 | 46153885 | 49299353 | 45904545 | 46528041 | 47517737 | 33184919 | 35763957 | 36774897 | 36253249 | 0 | 3 | 1.245007035 | 0.938613622 |
| Q9D8V0 | Hm13 | Minor histocompatibility antigen H13 | 278.34 | 16.4 | 41748 | 6 | 17344729 | 18118161 | 16397292 | 14235864 | 17362193 | 21622459 | 19333771 | 21930545 | 19732055 | 19179163 | 20518239 | 19863279 | 22094347 | 21011995 | 0 | 0 | 1.219747675 | 1.25044365 |
| P63017 | Hspa8 | Heat shock cognate 71 kDa protein | 3579.499 | 77.86 | 70871 | 48 | 1801361665 | 1869525377 | 1957913473 | 1742636801 | 1764108289 | 1649671041 | 1699606401 | 1699964801 | 1645360129 | 1663941761 | 1694374529 | 1615660545 | 1761841537 | 1766261249 | 0 | 3 | 0.914947448 | 0.935649899 |
| Q99P31 | Hspbp1 | Hsp70-binding protein 1 | 572.21 | 45.66 | 39167 | 10 | 14451627 | 14545725 | 16243478 | 15873172 | 14678513 | 14233226 | 12644041 | 13985019 | 13554151 | 12725764 | 14455019 | 14324560 | 14405397 | 14767091 | 0 | 4 | 0.885868492 | 0.955768307 |
| P26772 | Hspe1 | 10 kDa heat shock protein, mitochondrial | 215.66 | 42.16 | 10902 | 4 | 18476301 | 19934239 | 19126895 | 21050439 | 16966677 | 12317349 | 15474625 | 14810171 | 10708773 | 15614746 | 15044636 | 14590793 | 15932658 | 15271881 | 0 | 0 | 0.721322671 | 0.795880041 |
| P03975 | Iap | IgE-binding proteinV=1 | 590.01 | 23.16 | 62747 | 11 | 21785451 | 19728165 | 25046163 | 23513657 | 20108189 | 31279901 | 34253705 | 32022245 | 31745995 | 30166807 | 32709733 | 34841109 | 32528817 | 34805005 | 0 | 0 | 1.447325296 | 1.530253615 |
| Q9R002 | Ifi202 | Interferon-activable protein 202 | 144.04 | 6.29 | 50466 | 2 | 2326020.25 | 2754549.75 | 2657673.25 | 1744767.5 | 1884176.88 | 1 | 1 | 1 | 1 | 1 | 1 | 1 | 1 | 1 | 0 | 0 | 4.398625E-07 | 4.398625E-07 |
| P0DOV2 | Ifi204 | Interferon-activable protein 204 | 675.02 | 27.46 | 69438 | 8 | 18012943 | 19144417 | 19639325 | 15222543 | 14087636 | 10929171 | 11615319 | 12017299 | 11915774 | 9637524 | 9370458 | 9949678 | 11449507 | 8969819 | 0 | 0 | 0.651691217 | 0.576891611 |
| Q8BV66 | Ifi44 | Interferon-induced protein 44 | 78.42 | 5.92 | 47852 | 2 | 6552790.5 | 6363975 | 5733456 | 3897284.25 | 5024364 | 2039624.12 | 2116467.25 | 1792802.5 | 2103463.25 | 2083649.25 | 1789207.75 | 1374940.88 | 461732.31 | 1208626.98 | 0 | 0 | 0.367621292 | 0.219177551 |
| Q9BDB7 | Ifi44l | Interferon-induced protein 44-like2 | 231.32 | 12.98 | 49680 | 4 | 2844332.75 | 2146209.25 | 2052692.5 | 1848826.62 | 1618923.62 | 842147.125 | 1044910.31 | 669528.31 | 942336.44 | 711813.44 | 737215.38 | 424957.41 | 802695.25 | 167065.08 | 0 | 0 | 0.400603343 | 0.253536321 |
| Q68FR3 | Ints12 | Integrator complex subunit 12 | 54.09 | 3.04 | 48477 | 1 | 1 | 1 | 1 | 1 | 1 | 233719.3 | 188584.84 | 199350.3975 | 202375.92 | 172721.53 | 1 | 132676.22 | 124364.33 | 1 | 0 | 5 | 199350.3975 | 64260.6375 |
| Q9JKF1 | Iqgap1 | Ras GTPase-activating-like protein IQGAP1 | 6063.399 | 68.56 | 188742 | 92 | 519050561 | 488924513 | 508137121 | 496845985 | 463244961 | 576668161 | 560585793 | 599433985 | 578488513 | 558550977 | 499355777 | 497528801 | 520964737 | 541056769 | 0 | 4 | 1.160537834 | 1.039346313 |

| | | | | | | | | | | | | | | | | | | | | | | | | |
|---|---|---|---|---|---|---|---|---|---|---|---|---|---|---|---|---|---|---|---|---|---|---|---|---|
| Q3UQ44 | Iqgap2 | Ras GTPase-activating-like protein IQGAP2 | 196.71 | 2.6 | 180528 | 1 | 22371009 | 24136241 | 22486175 | 24295237 | 22065695 | 27804099 | 25029741 | 25726317 | 25219491 | 28893215 | 23858859 | 27781209 | 24645635 | 24493761 | 0 | 2 | 1.150133089 | 1.092063909 |
| Q5EIC4 | Irf2bpl | Interferon regulatory factor 2-binding protein-like | 79.29 | 4.73 | 81496 | 1 | 886338.5 | 1033053.44 | 1089360.38 | 573580.88 | 509385.41 | 1123089.5 | 1158167.88 | 1381445.75 | 1591920.5 | 1102901.75 | 1146767 | 1037823.94 | 1283478.25 | 1304527.25 | 0 | 1 | 1.553754299 | 1.458004843 |
| Q60766 | Irgm1 | Immunity-related GTPase family M protein 1 | 382.54 | 19.07 | 46552 | 7 | 6445768.5 | 4892167.5 | 4781422.5 | 3869175 | 4389789 | 2883047.75 | 2765580 | 2933151.25 | 3284595 | 2833628 | 2599356.5 | 2450496.5 | 3855157.5 | 2993222.25 | 0 | 0 | 0.602994812 | 0.610082623 |
| P05555 | Itgam | Integrin alpha-M | 1606.42 | 30.01 | 127481 | 24 | 68837145 | 72955153 | 78229753 | 61495005 | 73197409 | 56020089 | 55143293 | 60602621 | 60426461 | 57877545 | 59598233 | 60240989 | 68919881 | 61229769 | 0 | 2 | 0.817756358 | 0.880951077 |
| O70309 | Itgb5 | Integrin beta-5 | 72.13 | 2.88 | 87909 | 2 | 1 | 1 | 1 | 1 | 1 | 68096.88 | 104749.095 | 153777.06 | 127744.94 | 69377.5 | 91404.76 | 131483.73 | 82495.11 | 101794.5333 | 0 | 0 | 104749.095 | 101794.5333 |
| Q569A6 | Kdelr1 | ER lumen protein-retaining receptor 1 2 | 97.28 | 10.38 | 24546 | 2 | 1144506.25 | 213300.92 | 162754.42 | 189118.08 | 427419.9175 | 1 | 102192.34 | 158620.48 | 1 | 1 | 1 | 1 | 138202.08 | 147157.95 | 0 | 0 | 0.122041959 | 0.166909647 |
| P33175 | Kif5a | Kinesin heavy chain isoform 5A | 188.16 | 4.58 | 117019 | 0 | 3830728.5 | 2069237.5 | 2084377.88 | 1940073.25 | 2864564 | 1756975 | 1619471.5 | 1182498.75 | 1585857.468 | 1784484.62 | 3130023.5 | 1716612.75 | 2907706.5 | 1499499.88 | 0 | 8 | 0.620009308 | 0.904474185 |
| Q8C3Y4 | Kntc1 | Kinetochore-associated protein 1 | 106.39 | 1.63 | 250358 | 3 | 1642317.75 | 1803738.75 | 1552117.5 | 1696947.75 | 1526644.5 | 1438818.5 | 1387031.5 | 1432149.25 | 1324792 | 1088379.62 | 1622234.5 | 1434481.75 | 1410783.88 | 1251005.5 | 0 | 2 | 0.811403617 | 0.869415624 |
| Q9D1L9 | Lamtor5 | Ragulator complex protein LAMTOR5 | 51.75 | 21.98 | 9642 | 1 | 2614597.25 | 2651509.75 | 3639202.75 | 2723047 | 3201065.75 | 2272561.25 | 2567972.75 | 2065061.38 | 2366158.75 | 2342385.5 | 3204254.5 | 2217009.25 | 2566614.5 | 3257206 | 0 | 8 | 0.783182193 | 0.947869367 |
| Q07797 | Lgals3bp | Galectin-3-binding protein | 1156.02 | 39.34 | 64491 | 16 | 84722809 | 85872353 | 84740609 | 60414577 | 75841993 | 48594913 | 47905609 | 47771129 | 44640049 | 45468081 | 48904841 | 45471689 | 70658721 | 50665385 | 0 | 1 | 0.598530044 | 0.688536947 |
| Q9Z0M5 | Lipa | Lysosomal acid lipase/cholesteryl ester hydrolase | 683.19 | 22.17 | 45325 | 9 | 14897863 | 18597939 | 18244205 | 13282737 | 12877767 | 27890647 | 30345103 | 34120141 | 34454009 | 32353231 | 40208761 | 55868197 | 56020901 | 45081977 | 0 | 0 | 2.043159011 | 3.163968912 |
| Q8C129 | Lnpep | Leucyl-cystinyl aminopeptidase | 682.9 | 13.95 | 117304 | 11 | 12268318 | 9276440 | 12960425 | 9956491 | 8948255 | 17623309 | 15894234 | 16932119 | 20007805 | 15619805 | 10285751 | 12964193 | 16254150 | 14058953 | 0 | 2 | 1.611634271 | 1.253583557 |
| Q8CGK3 | Lonp1 | Lon protease homolog, mitochondrial | 1072.35 | 34.04 | 105843 | 20 | 28760953 | 28925937 | 29730303 | 31896783 | 30825573 | 39436801 | 39184665 | 39297381 | 35430021 | 34813457 | 37455065 | 32411825 | 35603089 | 31406583 | 0 | 1 | 1.253249569 | 1.139577837 |
| Q8K1T1 | Lrrc25 | Leucine-rich repeat-containing protein 25 | 53.64 | 6.06 | 32673 | 1 | 654861.69 | 776330.31 | 764671.89 | 721633 | 905862.56 | 624420.81 | 1 | 1 | 1 | 1 | 596315.12 | 595834 | 671994.12 | 608548.25 | 0 | 1 | 0.163318364 | 0.808415846 |
| P62313 | Lsm6 | U6 snRNA-associated Sm-like protein LSm6 | 43.15 | 10 | 9128 | 1 | 3327701 | 2948293.25 | 2895213.25 | 3063587.25 | 2961613 | 2748458.25 | 2807458.5 | 2351072 | 2245264.75 | 2222299.5 | 2413814 | 3338756.5 | 2971443.75 | 2118903.75 | 0 | 8 | 0.814307776 | 0.89189812 |
| Q9QYL8 | Lypla2 | Acyl-protein thioesterase 2 | 40.87 | 10.82 | 24807 | 1 | 875456.56 | 942451.81 | 740642.75 | 577799.25 | 760094.81 | 987983.12 | 1483220.62 | 2577308.75 | 2390149.25 | 1009287.06 | 2236100.75 | 1277549.75 | 1083784.25 | 1011178.44 | 0 | 0 | 2.16811694 | 1.799272456 |
| Q9QXZ0 | Macf1 | Microtubule-actin cross-linking factor 1 | 1182.11 | 4.98 | 831878 | 26 | 6852718.5 | 8346064 | 8966120 | 7128024 | 6447014 | 9448019 | 9302008 | 10202712 | 12248702 | 9357708 | 10193320 | 11638077 | 9482751 | 10385722 | 0 | 0 | 1.339672197 | 1.381158444 |
| O09159 | Man2b1 | Lysosomal alpha-mannosidase | 636.26 | 14.61 | 114648 | 10 | 29760767 | 29134727 | 24821189 | 32504589 | 31066853 | 22524719 | 22186813 | 22868383 | 23590495 | 20176759 | 24421871 | 22808655 | 21914893 | 23430521 | 0 | 0 | 0.755981984 | 0.785670433 |
| Q9ESL4 | Map3k20 | Mitogen-activated protein kinase kinase kinase 20 | 282.33 | 7.48 | 91720 | 5 | 9457455 | 9591159 | 8883040 | 8947703 | 9302907 | 9875387 | 10254225 | 10923897 | 10057027 | 10172491 | 10556757 | 9972235 | 9422916 | 7723086 | 0 | 7 | 1.110448526 | 1.019736549 |
| P21708 | Mapk3 | Mitogen-activated protein kinase 3 | 285.39 | 25.53 | 43081 | 3 | 14046033 | 13884697 | 13244656 | 12927679 | 14114616 | 17948245 | 14339667 | 15353954 | 16576580 | 16775449 | 16691656 | 15289569 | 16162203 | 16747251 | 0 | 0 | 1.187285962 | 1.189037029 |
| P49718 | Mcm5 | DNA replication licensing factor MCM5 | 1617.39 | 52.8 | 82343 | 30 | 45386225 | 52188905 | 47732161 | 51570645 | 48091649 | 43327481 | 40836581 | 40987033 | 44097633 | 43083477 | 40616589 | 49738469 | 43903841 | 41701089 | 0 | 3 | 0.866769664 | 0.897866505 |
| Q61881 | Mcm7 | DNA replication licensing factor MCM7 | 1286.23 | 44.37 | 81211 | 22 | 33899465 | 33035453 | 32675285 | 34860397 | 31904891 | 27325927 | 29044059 | 30638405 | 30152625 | 26885287 | 28408079 | 27961453 | 26078545 | 25678947 | 0 | 0 | 0.865790401 | 0.812371938 |
| P14152 | Mdh1 | Malate dehydrogenase, cytoplasmic | 893.88 | 62.28 | 36511 | 15 | 66041913 | 69056273 | 74115449 | 65314393 | 67860985 | 90727257 | 84473145 | 80912601 | 105949033 | 89114865 | 80568481 | 97653977 | 91438297 | 64822617 | 0 | 5 | 1.317731831 | 1.221137943 |
| P21956 | Mfge8 | Lactadherin | 401.72 | 23.11 | 51241 | 8 | 25970557 | 26130477 | 26266839 | 20592657 | 23680475 | 20323777 | 19623881 | 19144729 | 18653009 | 19247727 | 17701069 | 13561518 | 21956337 | 18465437 | 0 | 1 | 0.790870256 | 0.730632069 |
| E9PVX6 | Mki67 | Proliferation marker protein Ki-67 | 3338.53 | 27.7 | 350864 | 60 | 194457617 | 64824269 | 67822953 | 70583001 | 81777945 | 53681725 | 31672173 | 36557573 | 32160395 | 52863345 | 83109921 | 47769329 | 50978801 | 75608257 | 0 | 7 | 0.431595366 | 0.671232224 |
| D0QMC3 | Mndal | Myeloid cell nuclear differentiation antigen-like protein | 513.11 | 22.68 | 60526 | 6 | 21679509 | 23776097 | 24524045 | 21039025 | 22592671 | 16122662 | 13932207 | 17640987 | 17385345 | 14221786 | 14060852 | 14942461 | 15294745 | 16304416 | 0 | 0 | 0.698019952 | 0.666774002 |
| A1L314 | Mpeg1 | Macrophage-expressed gene 1 protein | 1083.6 | 44.04 | 78390 | 18 | 51348273 | 40907589 | 51192469 | 39580849 | 48383313 | 39353189 | 30205523 | 28185057 | 34183377 | 38079869 | 31211605 | 33129407 | 43881609 | 32021371 | 0 | 2 | 0.734649252 | 0.757543328 |
| Q7M0E7 | Mrpl14 | 39S ribosomal protein L14, mitochondrial | 71.54 | 6.9 | 15913 | 1 | 632822.62 | 389845.28 | 377724.06 | 928022.75 | 734948.62 | 365889.25 | 268193.97 | 197096.45 | 213796.72 | 261244.0975 | 805939.19 | 865177.75 | 744044.69 | 629384.56 | 0 | 6 | 0.426400772 | 1.242321699 |
| P30204 | Msr1 | Macrophage scavenger receptor types I and II | 227.23 | 9.17 | 50170 | 5 | 29196489 | 30667877 | 38901625 | 33723585 | 36951481 | 25439385 | 24688635 | 23763053 | 27207317 | 26961627 | 27099437 | 29979159 | 28305943 | 32740393 | 0 | 3 | 0.755779144 | 0.871430854 |
| P02802 | Mt1 | Metallothionein-1 | 193.08 | 68.85 | 6018 | 3 | 1 | 3184470.75 | 1 | 1963729.25 | 533792.25 | 6911727.5 | 5090203.5 | 7274615 | 8056611 | 9215157 | 1491512 | 2475808.75 | 495071.62 | 1142451.12 | 0 | 8 | 6.43230394 | 1.233027359 |
| P02798 | Mt2 | Metallothionein-2 | 54.76 | 19.67 | 6115 | 1 | 1 | 1 | 1 | 1 | 1 | 2719493.5 | 1972055.88 | 3008447.75 | 3353212.75 | 3903325.25 | 1 | 876340.12 | 1 | 1 | 0 | 7.5 | 2991307.026 | 219085.78 |
| Q791V5 | Mtch2 | Mitochondrial carrier homolog 2 | 539.84 | 49.17 | 33499 | 11 | 20859653 | 18690925 | 19395263 | 19571011 | 20258059 | 22602769 | 22605763 | 21323819 | 25902417 | 21592661 | 20651543 | 21899347 | 28518225 | 24182107 | 0 | 1 | 1.154416925 | 1.205407591 |
| P62774 | Mtpn | Myotrophin | 212.54 | 43.22 | 12861 | 3 | 12414931 | 9254644 | 8558970 | 11747155 | 10568561 | 12925697 | 15846595 | 12813013 | 16022103 | 16317372 | 13558337 | 12699394 | 14288535 | 10384550 | 0 | 3 | 1.406904933 | 1.211617002 |
| Q9JK81 | Myg1 | UPF0160 protein MYG1, mitochondrial | 541.26 | 36.32 | 42723 | 11 | 17665041 | 19092641 | 16112001 | 19558717 | 18558771 | 14437871 | 15265776 | 15087716 | 12738415 | 14976250 | 13547084 | 14803579 | 13206613 | 14359070 | 0 | 0 | 0.796881881 | 0.768189974 |
| Q63358 | Myo9b | Unconventional myosin-IXb | 384.83 | 5.35 | 225037 | 7 | 1958592.38 | 2189105.75 | 2147990.75 | 1461775.75 | 2002667.5 | 2262807 | 2255004.75 | 2709591.25 | 2473535 | 2830665.5 | 1658130.25 | 2318432.75 | 2192890.75 | 2105076 | 0 | 6 | 1.283958386 | 1.059735878 |
| Q5PPG6 | Nap1l5 | Nucleosome assembly protein 1-like 5 2 | 35.42 | 4.52 | 16858 | 1 | 2030543 | 1892679.25 | 1967458.75 | 2355257 | 1713276.25 | 1402256.38 | 1443987.75 | 1327458.88 | 1358701.62 | 1509884.25 | 1741801.62 | 1917897.38 | 1365048.5 | 1856196.75 | 0 | 4 | 0.707112901 | 0.863640454 |
| Q8BP47 | Nars | Asparagine--tRNA ligase, cytoplasmic | 1235.19 | 42.04 | 64279 | 19 | 57360417 | 58800577 | 63115161 | 62486093 | 61635505 | 68363913 | 64754697 | 66663197 | 69494041 | 69905617 | 59691693 | 60229793 | 63878785 | 63270693 | 0 | 6 | 1.117943233 | 1.017933396 |
| Q62433 | Ndrg1 | Protein NDRG1 | 402.01 | 32.74 | 43009 | 7 | 12050821 | 17884593 | 12918770 | 12185809 | 17734777 | 23029545 | 23079657 | 26153407 | 27206085 | 28481307 | 19393255 | 15887399 | 16453997 | 18457141 | 0 | 4 | 1.758164279 | 1.20563404 |
| Q62425 | Ndufa4 | Cytochrome c oxidase subunit NDUFA4 | 180.38 | 68.29 | 9327 | 4 | 24869155 | 20107309 | 17811985 | 16011653 | 17451605 | 13934563 | 13859357 | 14726675 | 15424429 | 12746563 | 11183773 | 6177599 | 10366876 | 11151371 | 0 | 0 | 0.734445021 | 0.504921162 |
| Q9D1J1 | Necap2 | Adaptin ear-binding coat-associated protein 2 | 270.8 | 31.95 | 28598 | 2 | 3710688 | 4054495.5 | 3515519.5 | 3933436.5 | 4115200 | 5013994 | 5581684.5 | 4389146.5 | 5045821.5 | 5895955.5 | 4591696 | 3863060 | 4302733 | 4362804.5 | 0 | 3 | 1.341308222 | 1.107144239 |
| Q922L6 | Nelfcd | Negative elongation factor D | 107.55 | 5.25 | 66276 | 2 | 1230443.75 | 1123769.88 | 1206099 | 1379372.75 | 1355226.75 | 1506306 | 1772333.62 | 1458071 | 1675131.62 | 1627184 | 1595173 | 1401060.5 | 448711.31 | 1207126.25 | 0 | 8 | 1.277067269 | 0.923776012 |
| Q6P5H2 | Nes | Nestin | 2639.19 | 34.39 | 207124 | 46 | 79465009 | 89528201 | 88990681 | 93163257 | 81493441 | 96563737 | 107526281 | 101315457 | 102183537 | 110900217 | 87106777 | 76921713 | 81664481 | 85680297 | 0 | 6 | 1.198429464 | 0.95741499 |
| P97526 | Nf1 | Neurofibromin | 65.62 | 0.78 | 317083 | 1 | 279746.625 | 297530.44 | 277005.28 | 275340.78 | 269110 | 264020.62 | 266138.66 | 245558.67 | 223492.73 | 249802.67 | 280086.03 | 216161.66 | 207308 | 247418.09 | 0 | 4 | 0.892960442 | 0.849852773 |
| O35309 | Nmi | N-myc-interactor | 100.72 | 9.55 | 35236 | 2 | 2499665.5 | 2235807.25 | 2187643.75 | 2451689.75 | 2832923.5 | 1924270.12 | 1708475.75 | 2000299.75 | 1985911.62 | 2119268.5 | 1853734.75 | 1856813.88 | 1985596.25 | 2076469.75 | 0 | 0 | 0.797709807 | 0.795870198 |
| Q11011 | Npepps | Puromycin-sensitive aminopeptidase | 879.35 | 30.11 | 103325 | 19 | 29901503 | 28245597 | 28708541 | 28451621 | 28245683 | 30624523 | 32642347 | 36832961 | 33104643 | 31351347 | 23796579 | 30500277 | 28903423 | 29151335 | 0 | 7 | 1.146307524 | 0.978311643 |
| Q9DBY8 | Nvl | Nuclear valosin-containing protein-like | 211.68 | 7.25 | 94476 | 4 | 5006893 | 3699745.25 | 4004343 | 4415970 | 4322784.5 | 3261414.75 | 2164080.5 | 1853806.5 | 1463487 | 1505472.75 | 4264351.5 | 5289536 | 3331857 | 3668079 | 0 | 7 | 0.477780315 | 0.964686913 |
| P09103 | P4hb | Protein disulfide-isomerase | 1769.22 | 55.6 | 57058 | 28 | 136559073 | 139142049 | 125258673 | 127324713 | 149719969 | 156637729 | 165703089 | 171032881 | 160082321 | 169302545 | 150237809 | 138558721 | 150462081 | 148395649 | 0 | 3 | 1.213500195 | 1.083426216 |
| Q99LX0 | Park7 | Protein/nucleic acid deglycase DJ-1 | 623.23 | 76.19 | 20021 | 10 | 39469889 | 48986005 | 47944301 | 48110437 | 49732665 | 50031037 | 60516577 | 55527521 | 58796837 | 62060417 | 48712385 | 57808505 | 51117441 | 58712413 | 0 | 2 | 1.22493319 | 1.154519397 |

| | | | | | | | | | | | | | | | | | | | | | | | | |
|---|---|---|---|---|---|---|---|---|---|---|---|---|---|---|---|---|---|---|---|---|---|---|---|---|
| Q8BFV2 | Pcid2 | PCI domain-containing protein 2 | 113.78 | 8.77 | 46132 | 2 | 474814.62 | 199473.06 | 396730.5275 | 452961.81 | 459672.62 | 1 | 1 | 1 | 1 | 1 | 299287.41 | 278962.2133 | 252704.23 | 284895 | 0 | 4 | 2.520603E-06 | 0.703152881 |
| Q9WU78 | Pdcd6ip | Programmed cell death 6-interacting protein | 1562.22 | 41.2 | 96024 | 25 | 45678109 | 52402553 | 50640681 | 49440153 | 49380501 | 52804869 | 54905465 | 56555949 | 53051033 | 54614657 | 51647509 | 52123933 | 50429949 | 54272649 | 0 | 4 | 1.098528639 | 1.052720561 |
| Q9WUA3 | Pfkp | ATP-dependent 6-phosphofructokinase, platelet type | 1812.23 | 54.97 | 85455 | 20 | 62324449 | 64117569 | 62208097 | 54761193 | 61724577 | 73057633 | 78252481 | 84109385 | 84470713 | 83289881 | 67478985 | 61805585 | 67647865 | 64898885 | 0 | 3 | 1.321313267 | 1.07260131 |
| P09411 | Pgk1 | Phosphoglycerate kinase 1 | 2512.98 | 79.14 | 44550 | 29 | 841446529 | 935717121 | 922474369 | 795992001 | 906236481 | 962210369 | 1057739841 | 1045803009 | 1095603713 | 1082488833 | 908902721 | 891912129 | 997301121 | 917143809 | 0 | 7 | 1.191277783 | 1.055023982 |
| Q9D0F9 | Pgm1 | Phosphoglucomutase-1 | 1225.89 | 48.04 | 61418 | 18 | 24706457 | 27459615 | 26923959 | 22012243 | 24741713 | 27569583 | 28038787 | 33208803 | 32010021 | 28321793 | 25857931 | 26726955 | 25721051 | 24685767 | 0 | 10 | 1.185189619 | 1.023009784 |
| O55022 | Pgrmc1 | Membrane-associated progesterone receptor component 1 | 61.64 | 7.18 | 21694 | 1 | 2714451.25 | 2481273 | 3081889.75 | 3019477.25 | 3099087 | 3229726 | 3145999.75 | 3554294.75 | 3111369.5 | 3296737.25 | 3338519.25 | 2951335 | 3363885.5 | 3229840.5 | 0 | 3 | 1.13489337 | 1.118663234 |
| Q2TBE6 | Pi4k2a | Phosphatidylinositol 4-kinase type 2-alpha | 169.01 | 6.26 | 54258 | 3 | 2433654.75 | 1953824.25 | 2286650 | 2070547.25 | 2256479 | 1755818.75 | 1848662.25 | 1665317.38 | 1375635.12 | 1578174.75 | 1987015 | 2090155.38 | 2354536 | 1739704.5 | 0 | 7 | 0.747522243 | 0.928471908 |
| E9Q3L2 | Pi4ka | Phosphatidylinositol 4-kinase alpha | 125.61 | 1.9 | 237042 | 2 | 1 | 1 | 134357.41 | 1 | 1 | 136527.52 | 192876.2475 | 256811.02 | 232711.98 | 145454.47 | 206593.7767 | 292065.56 | 146406.19 | 181309.58 | 0 | 0 | 7.177516502 | 7.687987818 |
| Q8BTI9 | Pik3cb | Phosphatidylinositol 4,5-bisphosphate 3-kinase catalytic subunit beta isoform | 54.26 | 1.5 | 121711 | 1 | 528539.31 | 442406.34 | 1 | 1 | 1 | 598015.9075 | 577891.19 | 577632.44 | 613485.31 | 623054.69 | 497625.53 | 571729.76 | 600266.06 | 617297.69 | 0 | 1 | 3.079544461 | 2.94418124 |
| P51432 | Plcb3 | 1-phosphatidylinositol 4,5-bisphosphate phosphodiesterase beta-3 | 235.68 | 7.54 | 139487 | 5 | 870095.25 | 1536200 | 1617768.75 | 974266.44 | 1494747.25 | 2135581 | 1680163 | 2121126.5 | 2295048 | 1993543 | 2401421 | 2393453.75 | 1902488.5 | 1969742.5 | 0 | 0 | 1.574825066 | 1.668528039 |
| Q8CIH5 | Plcg2 | 1-phosphatidylinositol 4,5-bisphosphate phosphodiesterase gamma-2 | 905.48 | 20.32 | 147592 | 20 | 19160359 | 20443325 | 19328189 | 19770709 | 21158481 | 24689185 | 21838817 | 21799275 | 22351687 | 23440199 | 19941969 | 17602193 | 20122117 | 19820003 | 0 | 9 | 1.142779373 | 0.969926111 |
| O35405 | Pld3 | Phospholipase D3 | 139.88 | 11.27 | 54389 | 3 | 9169067 | 7480470.5 | 6721049 | 9324024 | 8638230 | 6298740.5 | 5068346.5 | 4829077 | 3958668.5 | 4416345 | 9512652 | 9294508 | 5930433 | 7938143 | 0 | 9 | 0.59447106 | 0.988189283 |
| Q8BG07 | Pld4 | Phospholipase D4 | 1299.01 | 42.94 | 56154 | 19 | 217774625 | 283700417 | 237069425 | 208314785 | 226738369 | 151815697 | 173522833 | 199112241 | 179514177 | 161711761 | 189010065 | 166340353 | 174621745 | 194205297 | 0 | 0 | 0.737626503 | 0.771322137 |
| Q07832 | Plk1 | Serine/threonine-protein kinase PLK1 | 176.32 | 8.13 | 68301 | 4 | 661800.31 | 1368742.25 | 1096273.25 | 1421582.62 | 593047.19 | 584507.06 | 1 | 1 | 316370.06 | 1 | 601306.19 | 1612779.75 | 717066.31 | 1133826.75 | 0 | 9 | 0.175219226 | 0.988286977 |
| P23492 | Pnp | Purine nucleoside phosphorylase | 568.02 | 53.63 | 32277 | 9 | 49712993 | 48014693 | 53075109 | 46328725 | 46769449 | 36315229 | 40750337 | 39961977 | 38437057 | 30295669 | 27103267 | 35282353 | 38767565 | 34755837 | 0 | 0 | 0.761621693 | 0.696537936 |
| Q0VGB7 | Ppp4r2 | Serine/threonine-protein phosphatase 4 regulatory subunit 2 | 177.66 | 10.55 | 46479 | 3 | 1850140 | 2034461.25 | 2903052.75 | 1955833 | 2421902 | 3027132.5 | 3224331 | 3485005.5 | 3319220.25 | 3209334.5 | 3189156 | 1803652 | 3804962 | 3350605 | 0 | 5 | 1.456735968 | 1.360048338 |
| Q63009 | Prmt1 | Protein arginine N-methyltransferase 1 | 1101.34 | 53.54 | 40522 | 20 | 48532225 | 47985949 | 46430077 | 42264709 | 45721829 | 41912681 | 35487577 | 40275569 | 37532961 | 37666145 | 26250871 | 48974569 | 37189773 | 42121745 | 0 | 5 | 0.835192194 | 0.836475086 |
| O35522 | Psmb9 | Proteasome subunit beta type-9 Mus musculus bactrianus | 260.75 | 34.25 | 23427 | 5 | 23776365 | 27161923 | 22946583 | 26063835 | 24925917 | 20062789 | 21509665 | 21172077 | 18464499 | 19879483 | 20457933 | 17895225 | 19753685 | 20238313 | 0 | 0 | 0.809520066 | 0.784238163 |
| P97814 | Pstpip1 | Proline-serine-threonine phosphatase-interacting protein 1 | 282.97 | 17.11 | 47590 | 5 | 9298495 | 9274206 | 8585177 | 9588877 | 8833164 | 7339234 | 8400129 | 8584521 | 7509706 | 8168469.5 | 7430355.5 | 8039140.5 | 7276998.5 | 7285711 | 0 | 0 | 0.877624629 | 0.823613944 |
| P22437 | Ptgs1 | Prostaglandin G/H synthase 1 | 354.16 | 15.12 | 69042 | 6 | 318926.88 | 3227523.5 | 2729618 | 6842449 | 5433438.5 | 8101430 | 8955387 | 8751370 | 8849903 | 9009687 | 8367141 | 5618519 | 5071180.5 | 5009584.5 | 0 | 5 | 2.353809878 | 1.621555778 |
| P61027 | Rab10 | Ras-related protein Rab-10 | 490.94 | 42 | 22541 | 7 | 21850323 | 21533451 | 21389139 | 20575261 | 19195639 | 23333243 | 22967457 | 24281881 | 22086963 | 22118249 | 22161477 | 22816339 | 24300193 | 22347823 | 0 | 0 | 1.097987434 | 1.095543454 |
| Q8CGC6 | Rbm28 | RNA-binding protein 28 | 440.11 | 13.87 | 84214 | 9 | 3444275 | 3734736.5 | 787139.31 | 3987158 | 445939.22 | 5192945.5 | 4845445 | 4122897 | 4916632 | 4967977 | 4456815.5 | 5441411.5 | 1491287.25 | 4301062 | 0 | 3 | 1.939302806 | 1.581807241 |
| P17074 | Rps19 | 40S ribosomal protein S19 | 377.39 | 35.17 | 16085 | 7 | 25781753 | 23033861 | 20522993 | 22798165 | 21703593 | 18428705 | 12208375 | 20073651 | 16947313 | 19788967 | 20618567 | 18086761 | 16976985 | 21485765 | 0 | 2 | 0.768154696 | 0.847327725 |
| P49242 | Rps3a | 40S ribosomal protein S3a | 740.66 | 45.83 | 29945 | 13 | 113750105 | 112958609 | 114948289 | 118346137 | 109531193 | 97042241 | 105823233 | 100710585 | 100679497 | 105432593 | 102109929 | 103039873 | 99281617 | 107522273 | 0 | 0 | 0.894920849 | 0.90414587 |
| O35114 | Scarb2 | Lysosome membrane protein 2 | 295.67 | 14.85 | 54044 | 5 | 3792335.75 | 3064972 | 4783295 | 2932860 | 2375041.5 | 7356850 | 5877957 | 8681323 | 8287700.5 | 5328843 | 5932664 | 10648394 | 8507017 | 7792342 | 0 | 0 | 2.096507926 | 2.425023509 |
| Q9CQA3 | Sdhb | Succinate dehydrogenase [ubiquinone] iron-sulfur subunit, mitochondrial | 438.18 | 30.5 | 31814 | 8 | 44796633 | 49837093 | 50325585 | 50812549 | 49363261 | 57099637 | 59152501 | 52235157 | 64218137 | 59485769 | 53334785 | 55999333 | 53467569 | 55464701 | 0 | 0 | 1.191959764 | 1.11299019 |
| P42208 | Sept2 | Septin-2 | 642.11 | 43.21 | 41526 | 9 | 28462507 | 28598655 | 27987833 | 26868779 | 28410479 | 31002607 | 30110541 | 30369759 | 30355221 | 30671835 | 28705013 | 28579201 | 27823747 | 28760313 | 0 | 5 | 1.086808677 | 1.014302818 |
| Q9DCQ7 | Serpina1f | Alpha-1-antitrypsin 1-6 Mus musculus | 33.93 | 1.95 | 46337 | 1 | 5145926 | 4528943.5 | 5142068 | 3530996.75 | 3343327.5 | 1960305.75 | 2068821.88 | 2967468.5 | 1613439.5 | 1921989.38 | 1524142.25 | 2172384.25 | 3404321.25 | 1969269 | 0 | 1 | 0.485542295 | 0.522682639 |
| P46062 | Sipa1 | Signal-induced proliferation-associated protein 1 | 144.33 | 4.53 | 112066 | 2 | 2053289.38 | 2238001 | 2395500 | 2068664.88 | 2462241.25 | 2002720.5 | 1997478.12 | 1813371.12 | 1866137 | 1663222.38 | 2175346.75 | 2116383 | 540939.88 | 2139342 | 0 | 6 | 0.832874121 | 0.776898763 |
| Q9QXX4 | Slc25a13 | Calcium-binding mitochondrial carrier protein Aralar2 | 1659.72 | 63.61 | 74467 | 26 | 62194497 | 61106117 | 65557537 | 57746685 | 64471141 | 67684505 | 68092353 | 81449977 | 75328153 | 72001529 | 73703249 | 66758481 | 87497329 | 74168241 | 0 | 0 | 1.171921151 | 1.214041433 |
| P16036 | Slc25a3 | Phosphate carrier protein, mitochondrial | 713.67 | 44.66 | 39445 | 12 | 36062841 | 41107849 | 41221105 | 34584221 | 38753701 | 47766305 | 45828705 | 50553545 | 54328689 | 48474313 | 44965977 | 49965213 | 50002517 | 46465757 | 0 | 0 | 1.288019202 | 1.247846884 |
| P10852 | Slc3a2 | 4F2 cell-surface antigen heavy chain | 986.12 | 53.23 | 58337 | 19 | 41247057 | 18164911 | 19587141 | 18179335 | 20064393 | 17544019 | 16581012 | 16246881 | 16133649 | 15747751 | 14971227 | 13672847 | 18708373 | 17467807 | 0 | 2 | 0.701563644 | 0.691089704 |
| O35892 | Sp100 | Nuclear autoantigen Sp-100 | 189.49 | 9.96 | 54727 | 3 | 3439218.5 | 3207927 | 3216481.25 | 2612204 | 2956948.5 | 1324223.75 | 2217736.75 | 2585625 | 2355409.25 | 2315391.5 | 2404158.75 | 2575712.25 | 2204169 | 1244349 | 0 | 0 | 0.699704575 | 0.682669406 |
| Q8BVK9 | Sp110 | Sp110 nuclear body protein | 472.34 | 32.58 | 50140 | 10 | 9270450 | 7255536 | 8656968 | 7135867.5 | 8281462.5 | 5700259 | 6962201 | 6817715.5 | 6990179.5 | 6948625.5 | 5176007 | 5951477 | 6941924.5 | 5088065.5 | 0 | 0 | 0.823121841 | 0.712971429 |
| Q9JM90 | Stap1 | Signal-transducing adaptor protein 1 | 637.43 | 59.93 | 34628 | 13 | 17729511 | 12382445 | 13236752 | 13402185 | 15024507 | 23414919 | 21383787 | 18052759 | 23343591 | 22613869 | 22383089 | 22426727 | 19523953 | 20117637 | 0 | 0 | 1.515964035 | 1.470758192 |
| P42225 | Stat1 | Signal transducer and activator of transcription 1 | 1044.25 | 29.37 | 87197 | 17 | 27494325 | 28965477 | 27279907 | 21606003 | 25210429 | 16652458 | 15746811 | 15368089 | 18423641 | 16536492 | 14598458 | 14268871 | 18060927 | 15280153 | 0 | 0 | 0.633654536 | 0.595609756 |
| Q9WVL2 | Stat2 | Signal transducer and activator of transcription 2 | 153.72 | 3.47 | 105417 | 3 | 2279999.25 | 2238939.75 | 2558215.5 | 1865435.12 | 1575645.25 | 1512023 | 247460.8 | 1372053.88 | 1270407.5 | 1310062.38 | 1185951.38 | 1208175.88 | 1289742.88 | 1175394.5 | 0 | 0 | 0.543057617 | 0.577481001 |
| Q8VIM6 | Strc | Stereocilin | 35.29 | 0.44 | 196347 | 1 | 722819.81 | 2210075.5 | 1922223.88 | 1858499.88 | 719493.31 | 3080604.5 | 3225625.75 | 3418017 | 4935381 | 2952063.25 | 3495577.5 | 2236525 | 3993204 | 3594814.25 | 0 | 0 | 2.36935628 | 2.239997203 |
| O88983 | Stx8 | Syntaxin-8 | 152.78 | 11.86 | 26925 | 2 | 1879497.563 | 1758616.75 | 2002885.88 | 1978806.12 | 1777681.5 | 1351127.75 | 1448355.62 | 1478653.75 | 1335958.88 | 1441302.38 | 1542985.25 | 1589585.083 | 1527788.5 | 1697981.5 | 0 | 0 | 0.750774943 | 0.845750011 |
| Q6A028 | Swap70 | Switch-associated protein 70 | 826.42 | 35.21 | 68996 | 18 | 8438880 | 4376241.5 | 9912001 | 6593201 | 9192176 | 12313358 | 9991668 | 10447698 | 10027200 | 12633437 | 8687989 | 12469795 | 13438304 | 9246873 | 0 | 3 | 1.43884094 | 1.423010762 |

| | | | | | | | | | | | | | | | | | | | | | | | | |
|---|---|---|---|---|---|---|---|---|---|---|---|---|---|---|---|---|---|---|---|---|---|---|---|---|
| Q91WQ5 | Taf5l | TAF5-like RNA polymerase II p300/CBP-associated factor-associated factor 65 kDa subunit 5L | 43.84 | 3.74 | 65911 | 1 | 196988.38 | 279299.28 | 218964.33 | 247291.38 | 293913.53 | 1 | 95993.91 | 1 | 190384.42 | 1 | 276832.91 | 184487.11 | 180785.72 | 236676.84 | 0 | 5 | 0.231614487 | 0.888408019 |
| Q0VGY8 | Tanc1 | Protein TANC1 | 276.4 | 5.87 | 200805 | 6 | 1665739.12 | 1572150.38 | 1584758.25 | 1529621.75 | 1553410 | 2260720.75 | 2593082.75 | 2205979 | 2687452.5 | 2599394.75 | 2192618 | 1965477.12 | 1814102.62 | 1908039.88 | 0 | 0 | 1.561741752 | 1.245977278 |
| P21958 | Tap1 | Antigen peptide transporter 1 | 423.56 | 17.68 | 78864 | 9 | 10058543 | 10971870 | 10419274 | 7036405.5 | 7464756 | 5593281 | 5416298 | 6594180 | 6924227 | 6284634 | 4814017 | 3992453.25 | 6695657 | 5874981.5 | 0 | 0 | 0.670556062 | 0.581521056 |
| P36371 | Tap2 | Antigen peptide transporter 2 | 530.08 | 19.52 | 77445 | 9 | 14567948 | 13917693 | 10892413 | 10225757 | 12761393 | 5557308 | 7911689 | 6487473 | 9713233 | 5826027 | 8362977.5 | 5555959 | 7068710 | 9032469 | 0 | 0 | 0.569159206 | 0.601700018 |
| Q91YM4 | Tbrg4 | FAST kinase domain-containing protein 4 | 130.32 | 7.62 | 71513 | 3 | 2510035 | 2642429.75 | 2381086.75 | 1631869.62 | 1641948 | 487617.59 | 480820.78 | 478945.22 | 677948.38 | 621065 | 282645.81 | 527269.88 | 818783.38 | 2638052.75 | 0 | 4 | 0.254122621 | 0.493500288 |
| Q91YX0 | Themis2 | Protein THEMIS2 | 451.31 | 22.32 | 74378 | 9 | 6519084.5 | 7443011.5 | 6576726.5 | 7534454 | 7404014.5 | 3663874.5 | 4519365 | 4221583.5 | 4518707.5 | 4335175.5 | 5111128.5 | 5144754.5 | 5504107 | 4491972 | 0 | 0 | 0.599220104 | 0.713553707 |
| P62075 | Timm13 | Mitochondrial import inner membrane translocase subunit Tim13 | 36.2 | 14.74 | 10458 | 1 | 2119784.5 | 2271527.25 | 2447546.5 | 2781436.25 | 2072312.75 | 1404924 | 1992912 | 1928502.38 | 1382621 | 1689689.62 | 2156696.75 | 1935577 | 1747974.12 | 1923287.5 | 0 | 2 | 0.718287104 | 0.829961958 |
| Q6R5N8 | Tlr13 | Toll-like receptor 13 | 647.17 | 13.62 | 114443 | 9 | 12779083 | 9427389 | 12441244 | 11319293 | 11369952 | 8719041 | 8643784 | 9305679 | 9360403 | 6470506.5 | 8340983.5 | 8267912 | 11075327 | 7969123 | 0 | 1 | 0.741221941 | 0.777276666 |
| Q9CR02 | Tma16 | Translation machinery-associated protein 16 | 97.78 | 17.19 | 25782 | 2 | 2082810.12 | 1679686.38 | 1689868.88 | 2088082.88 | 2318952.25 | 163200 | 1099331.66 | 1320676.88 | 1354477.38 | 1558972.38 | 2616166 | 2732925.75 | #RÉF ! | 2198541.75 | 0 | 4 | 0.557504312 | #RÉF ! |
| Q6AYA5 | Tmem106b | Transmembrane protein 106B | 52.41 | 5.82 | 31152 | 1 | 1 | 808104.06 | 1153426 | 1 | 1 | 1358662.375 | 1316674 | 1427608.5 | 1189302 | 1501065 | 1351118.75 | 1330084.5 | 1385718 | 1374090.88 | 0 | 0 | 3.463266571 | 3.4673212 |
| Q9CYG7 | Tomm34 | Mitochondrial import receptor subunit TOM34 | 659.61 | 45.95 | 34278 | 11 | 12720540 | 14699377 | 15130840 | 12742153 | 14684081 | 16434287 | 16587682 | 16365119 | 18186055 | 16624583 | 16003520 | 16805789 | 17180667 | 17118991 | 0 | 0 | 1.203220156 | 1.198768446 |
| Q6IRU2 | Tpm4 | Tropomyosin alpha-4 chain | 1222.06 | 58.87 | 28468 | 9 | 171244289 | 181799265 | 161074145 | 175204321 | 158968001 | 192287377 | 220924417 | 209447985 | 182344801 | 186597441 | 174388577 | 172576593 | 165195169 | 188302001 | 0 | 7 | 1.168942221 | 1.03216813 |
| P23591 | Tsta3 | GDP-L-fucose synthase | 230.02 | 20.25 | 35878 | 5 | 7730363.5 | 8365780 | 7979294 | 8123320 | 8205173.5 | 6914703 | 6895175 | 7710219 | 6726803.5 | 6897820.5 | 6750698 | 7032807 | 6528529.5 | 7176565 | 0 | 0 | 0.8698342 | 0.85043085 |
| P25976 | Ubtf | Nucleolar transcription factor 1 | 94.05 | 3.27 | 89509 | 2 | 1509451.88 | 1644149 | 1590948.5 | 1521410.88 | 2415347 | 528094.81 | 535609.31 | 1381853.62 | 1451999 | 1501012.88 | 2249148.5 | 1146579.75 | 1312194.38 | 1339223 | 0 | 4 | 0.621861369 | 0.87071357 |
| Q9CR09 | Ufc1 | Ubiquitin-fold modifier-conjugating enzyme 1 | 173.85 | 31.14 | 19481 | 4 | 7331790 | 6625408.5 | 6249144.5 | 5932308 | 6208152.5 | 4703334 | 4986133.5 | 5569932 | 5297996.5 | 4591163 | 5214540.5 | 4845140 | 5559540 | 3579490.25 | 0 | 0 | 0.77746659 | 0.741909117 |
| Q62465 | Vat1 | Synaptic vesicle membrane protein VAT-1 homolog | 635.9 | 35.47 | 43097 | 10 | 13013952 | 12600807 | 11770273 | 9058213 | 10924196 | 15030296 | 22454865 | 16856367 | 19143909 | 19103937 | 17374861 | 18009465 | 19733481 | 17329637 | 0 | 0 | 1.613970789 | 1.57858366 |
| P20152 | Vim | Vimentin | 3269.56 | 86.48 | 53688 | 48 | 2434394369 | 2796715521 | 2589878017 | 2831042049 | 2816222721 | 3398969601 | 3163222273 | 2930145793 | 3181000961 | 3520160769 | 2958590465 | 2826093825 | 2985940737 | 3154718721 | 0 | 1 | 1.202345975 | 1.106801308 |
| Q920I9 | Wdr7 | WD repeat-containing protein 7 | 94.65 | 1.61 | 163450 | 2 | 1002719.735 | 1159235.5 | 904080.81 | 981449.44 | 966113.19 | 702574.1375 | 737201.62 | 725148.62 | 778086.69 | 569859.62 | 990913 | 956931.44 | 820038.81 | 880263.38 | 0 | 4 | 0.700668505 | 0.909562887 |
| P83741 | Wnk1 | Serine/threonine-protein kinase WNK1 | 553.13 | 6.52 | 250934 | 10 | 110027521 | 83453697 | 72778305 | 98426177 | 83955393 | 70849145 | 58174965 | 55978133 | 38741957 | 40960961 | 88183441 | 107508809 | 39366733 | 93689369 | 0 | 10 | 0.590015416 | 0.915955864 |
| Q6P1B1 | Xpnpep1 | Xaa-Pro aminopeptidase 1 | 787.32 | 36.92 | 69591 | 3 | 22986119 | 19389019 | 20236287 | 19305361 | 21173427 | 27443181 | 25499117 | 26730447 | 27163809 | 23611945 | 22811497 | 24545289 | 28329321 | 21817261 | 0 | 2 | 1.265381991 | 1.182257815 |
| Q9JKB3 | Ybx3 | Y-box-binding protein 3 | 414.99 | 26.87 | 38814 | 3 | 2490044.5 | 1521259.38 | 2400492.22 | 2960451.75 | 2630213.25 | 383667.94 | 453809 | 1 | 1 | 1 | 2128008.75 | 1014960.813 | 409303.44 | 507570.25 | 0 | 1 | 0.069775685 | 0.422813623 |
| Q6NZF1 | Zc3h11a | Zinc finger CCCH domain-containing protein 11A | 217.47 | 8.33 | 86492 | 5 | 1478486.5 | 1571028 | 1751738.25 | 1529590 | 1550828.5 | 471800 | 1433966.25 | 1472847.5 | 1087586.38 | 1296602.88 | 981408.69 | 1301186.88 | 1336700.62 | 256232.75 | 0 | 0 | 0.731165108 | 0.614642634 |
| Q6P542 | Abcf1 | ATP-binding cassette sub-family F member 1 | 464.31 | 19.12 | 94945 | 9 | 7467059 | 5819978.5 | 6286385.5 | 6161664 | 6544938.5 | 5697914 | 4865801 | 5484054 | 5606019 | 5931955 | 5619560 | 3511276.5 | 6638671 | 5949459 | 1 | 5 | 0.854576246 | 0.841037382 |
| Q3TCJ1 | Abraxas2 | BRISC complex subunit Abraxas 2 | 49.2 | 2.89 | 46943 | 1 | 2015625 | 2341368 | 2138161.5 | 1699554 | 1894750.38 | 1502298.88 | 1606693.88 | 1780526.12 | 1559051.5 | 1319721.12 | 1701372.75 | 1670824.12 | 1509087.25 | 2086443.75 | 1 | 4 | 0.769941341 | 0.863243504 |
| Q91V92 | Acly | ATP-citrate synthase | 2898.74 | 53.99 | 119728 | 4 | 82236001 | 82837753 | 88651937 | 93847417 | 80978689 | 100457705 | 101209593 | 99180121 | 92174977 | 100754649 | 82173609 | 91873073 | 92448697 | 92403353 | 1 | 7 | 1.152199217 | 1.046835921 |
| Q7TPR4 | Actn1 | Alpha-actinin-1 | 2014.61 | 43.5 | 103068 | 1 | 93833825 | 94325033 | 103145209 | 95372721 | 84718417 | 103720737 | 104869953 | 118580505 | 111266985 | 97087129 | 100674945 | 98627897 | 99062257 | 99698417 | 1 | 4 | 1.136043183 | 1.055546153 |
| O88839 | Adam15 | Disintegrin and metalloproteinase domain-containing protein 15 | 102.67 | 2.08 | 92664 | 1 | 202693.86 | 264608.88 | 222039.39 | 219734.345 | 189595.25 | 416076.72 | 317525.62 | 365111.09 | 336759.44 | 249386.42 | 318597.91 | 522001.75 | 333007.41 | 250278.95 | 1 | 1 | 1.533542051 | 1.620008493 |
| Q8C0I1 | Agps | Alkyldihydroxyacetonephosphate synthase, peroxisomal | 624.79 | 23.26 | 71684 | 11 | 6235871 | 5000361 | 8535396 | 3113150 | 5289479.5 | 9094774 | 9202690 | 9930530 | 8488279 | 11379509 | 4702857 | 10497420 | 8836275 | 9850253 | 1 | 4 | 1.707082503 | 1.503447118 |
| Q66HF8 | Aldh1b1 | Aldehyde dehydrogenase X, mitochondrial | 111.29 | 4.82 | 57625 | 1 | 11603215 | 11684018 | 13051814 | 12502605 | 11462587 | 13215648 | 12968689 | 14213994 | 14635127 | 13888444 | 13951681 | 16082327 | 14318778 | 14370415 | 1 | 0 | 1.142903105 | 1.217227884 |
| P05064 | Aldoa | Fructose-bisphosphate aldolase A | 2292.67 | 93.68 | 39356 | 29 | 1180237697 | 1044185217 | 1001171329 | 1153109889 | 1062884545 | 1194690945 | 1324459905 | 1179178369 | 1419098497 | 1384668801 | 1151078913 | 1103589377 | 1139217921 | 1112596353 | 1 | 8 | 1.194889379 | 1.035194598 |
| P58366 | Ankh | Progressive ankylosis protein homolog | 85.84 | 4.27 | 54265 | 1 | 288447.91 | 143634.0975 | 98545.91 | 99877.84 | 87664.73 | 93527.75 | 1 | 1 | 1 | 63191.25 | 70649.23 | 1 | 136383.8 | 1 | 1 | 3 | 0.218223949 | 0.360351465 |
| Q80V94 | Ap4e1 | AP-4 complex subunit epsilon-1 | 78.97 | 1.78 | 124845 | 1 | 681580.38 | 454765.38 | 548853.5175 | 674298.69 | 384769.62 | 1 | 395299.84 | 373866.31 | 1 | 1 | 1 | 331713.78 | 1 | 1 | 1 | 0 | 0.280282125 | 0.151095315 |
| Q1AAU6 | Asap1 | Arf-GAP with SH3 domain, ANK repeat and PH domain-containing protein 1 | 459.92 | 9.53 | 127088 | 9 | 8820306 | 9953314 | 9387332 | 9091122 | 9597520 | 9608660 | 10004441 | 10351313 | 11853411 | 11547661 | 10488327 | 10368535 | 10301394 | 11688451 | 1 | 0 | 1.139081077 | 1.143198461 |
| O54984 | Asna1 | ATPase Asna1 | 491.22 | 48.56 | 38823 | 11 | 10520835 | 10712636 | 11106910 | 10759026 | 8052762 | 12026983 | 11247318 | 10786837 | 12850331 | 12713434 | 9912416 | 10081743 | 12311976 | 12029832 | 1 | 8 | 1.165637825 | 1.083433212 |
| Q9D906 | Atg7 | Ubiquitin-like modifier-activating enzyme ATG7 | 879.05 | 30.95 | 77520 | 17 | 21356659 | 16975681 | 21445151 | 22598491 | 21669667 | 22660869 | 24427023 | 26245317 | 22287843 | 25785223 | 23180135 | 21138159 | 23389657 | 25133443 | 1 | 4 | 1.166855858 | 1.115392557 |
| Q8R2Q8 | Bst2 | Bone marrow stromal antigen 2 | 110.88 | 11.63 | 19152 | 2 | 3549342 | 3271166.5 | 2628263 | 2198618.5 | 3423167 | 1919391.5 | 2287608.75 | 1598402.62 | 2082280.88 | 1783525.62 | 1678759.38 | 2267397.75 | 2377634.5 | 1841394 | 1 | 2 | 0.641728728 | 0.677246504 |
| P00920 | Ca2 | Carbonic anhydrase 2 | 72.87 | 6.15 | 29033 | 1 | 995448.44 | 967557.69 | 745657 | 1006444.56 | 713991.19 | 1199222.5 | 1319754.38 | 1269903.12 | 1257928.88 | 1000709.62 | 1030893.19 | 1065497.62 | 826465.69 | 936044.31 | 1 | 6 | 1.365406071 | 1.089076163 |
| B2RQC6 | Cad | CAD protein | 1956.76 | 23.28 | 243238 | 37 | 42515117 | 45114417 | 50362805 | 46463789 | 44529117 | 41966693 | 39472741 | 39367589 | 43304169 | 41710433 | 43789457 | 45553821 | 43951765 | 51205269 | 1 | 10 | 0.898842303 | 1.007162667 |
| O88456 | Capns1 | Calpain small subunit 1 | 587.5 | 59.48 | 28463 | 3 | 30998365 | 34028041 | 29501311 | 25698957 | 30787103 | 35859877 | 35178441 | 38785485 | 35084545 | 32038903 | 35672193 | 33780065 | 35818065 | 37239785 | 1 | 1 | 1.171729193 | 1.179611811 |
| P47753 | Capza1 | F-actin-capping protein subunit alpha-1 | 528 | 45.1 | 32940 | 8 | 59613385 | 60653597 | 55100237 | 56731757 | 63947313 | 68557569 | 65010953 | 66143533 | 67203049 | 62702825 | 66975649 | 65080177 | 64062505 | 67503489 | 1 | 0 | 1.113399969 | 1.113093753 |
| Q9ER72 | Cars | Cysteine--tRNA ligase, cytoplasmic | 955.15 | 23.83 | 94860 | 17 | 18081165 | 15996308 | 18579007 | 17411201 | 12776451 | 18610473 | 20338941 | 20677853 | 21007881 | 18411105 | 15022956 | 17093307 | 17232021 | 18480583 | 1 | 9 | 1.19557355 | 1.02344103 |
| P29594 | Casp2 | Caspase-2 | 45.19 | 1.99 | 50661 | 1 | 778569.94 | 877849.38 | 870675.62 | 927981.62 | 863769.14 | 796943.44 | 1 | 1 | 609012.81 | 1 | 853217.3133 | 794729.75 | 774508.69 | 990413.5 | 1 | 7 | 0.325540514 | 0.987783974 |
| Q8VEH6 | Cbwd1 | COBW domain-containing protein 1 | 132.73 | 9.16 | 43772 | 2 | 1518167.38 | 361844.88 | 1746521.25 | 1475068.88 | 1537275.12 | 1960452.38 | 1745335.5 | 2236414.5 | 2241467.25 | 1859763.25 | 1796521.5 | 2317916.5 | 1876033 | 1950675.62 | 1 | 0 | 1.512820935 | 1.495197533 |
| Q9WU84 | Ccs | Copper chaperone for superoxide dismutase | 129.26 | 18.25 | 28912 | 2 | 1 | 405529.62 | 485801.06 | 1 | 1 | 1633314.25 | 2479462.75 | 2229614.25 | 1918707.25 | 440717.06 | 1100983.62 | 363290.62 | 1130680.38 | 363333.5 | 1 | 4 | 9.762691296 | 4.148682175 |
| P60766 | Cdc42 | Cell division control protein 42 homolog | 537.62 | 58.64 | 21259 | 8 | 41102253 | 34127069 | 39225869 | 42724749 | 36411677 | 41515929 | 46319169 | 42854045 | 42987637 | 45671493 | 42492733 | 40941549 | 38666049 | 42134193 | 1 | 7 | 1.13304634 | 1.060444446 |

| | | | | | | | | | | | | | | | | | | | | | | | | |
|---|---|---|---|---|---|---|---|---|---|---|---|---|---|---|---|---|---|---|---|---|---|---|---|---|
| P19536 | Cox5b | Cytochrome c oxidase subunit 5B, mitochondrial | 53.61 | 9.38 | 13813 | 1 | 7728957.5 | 8080394.5 | 9121866 | 9044157 | 8073188.5 | 5919199.5 | 7450418 | 6304691 | 6546995 | 8021108 | 5163765.5 | 6167710.5 | 4713191.5 | 5723266.5 | 1 | 0 | 0.814353896 | 0.647106946 |
| P36552 | Cpox | Oxygen-dependent coproporphyrinogen-III oxidase, mitochondrial | 149.92 | 7.22 | 49715 | 3 | 1503168 | 1347360.5 | 1329728.25 | 1128550.25 | 1129803.62 | 1787710.75 | 1646714.88 | 1371607.25 | 3656753 | 3044954.5 | 1432165.5 | 3589638.5 | 1740911.75 | 3050852.5 | 1 | 1 | 1.787301805 | 1.905218538 |
| Q7TMB8 | Cyfip1 | Cytoplasmic FMR1-interacting protein 1 | 1945.15 | 33.2 | 145241 | 23 | 53762173 | 46518805 | 46904561 | 42789213 | 44979477 | 56949453 | 54642505 | 52869509 | 62710625 | 54195625 | 51165733 | 54168297 | 52025097 | 53359213 | 1 | 3 | 1.19754268 | 1.121060583 |
| Q5SQX6 | Cyfip2 | Cytoplasmic FMR1-interacting protein 2 | 846.05 | 16.04 | 145659 | 3 | 14203615 | 13260923 | 13543792 | 11617989 | 9311673 | 18029937 | 14095068 | 16040019 | 20379449 | 15796638 | 13576555 | 18370377 | 14383378 | 13005055 | 1 | 4 | 1.361702378 | 1.19747515 |
| Q91X52 | Dcxr | L-xylulose reductase | 65.94 | 7.38 | 25746 | 2 | 134480689 | 89065553 | 74447881 | 112225169 | 119284305 | 84487033 | 45989785 | 47740653 | 37871061 | 42012133 | 134733681 | 134293105 | 48283509 | 109253649 | 1 | 9 | 0.487438927 | 1.006990194 |
| Q9JIK5 | Ddx21 | Nucleolar RNA helicase 2 | 2053.9 | 54.76 | 93551 | 36 | 42154801 | 34022433 | 42058417 | 42927625 | 38194141 | 31975175 | 37257153 | 29146309 | 32689745 | 33366561 | 36178761 | 34434737 | 38966069 | 32566989 | 1 | 4 | 0.824824807 | 0.891279581 |
| Q61656 | Ddx5 | Probable ATP-dependent RNA helicase DDX5 | 1563.06 | 58.63 | 69290 | 22 | 225700161 | 233248081 | 244238129 | 239267329 | 231313041 | 211316033 | 207093025 | 219695521 | 211457249 | 227407761 | 224661905 | 195167313 | 221794673 | 231716417 | 1 | 2 | 0.917532889 | 0.93006161 |
| O35435 | Dhodh | Dihydroorotate dehydrogenase (quinone), mitochondrial | 93.26 | 8.35 | 42700 | 2 | 858975 | 905831.12 | 701636.5 | 959163.44 | 692961.5 | 696603.56 | 591537.12 | 659897.94 | 461137 | 651447.62 | 689094 | 690762 | 477557 | 804970.25 | 1 | 2 | 0.743128089 | 0.808042848 |
| Q9EQP2 | Ehd4 | EH domain-containing protein 4 | 1173.64 | 47.13 | 61481 | 17 | 45898105 | 35515837 | 35702145 | 34853389 | 35500921 | 33299337 | 34314649 | 30702581 | 33440849 | 35102593 | 31913481 | 34874433 | 35917621 | 32514789 | 1 | 5 | 0.890060573 | 0.901611175 |
| P97481 | Epas1 | Endothelial PAS domain-containing protein 1 | 34.03 | 0.8 | 96712 | 1 | 2291132.25 | 1687931.88 | 1177006.25 | 2503481.5 | 2628611.75 | 1560509.25 | 917629.25 | 978144.69 | 618671.19 | 732402.69 | 2386942 | 2456274 | 831776.06 | 1641924.12 | 1 | 7 | 0.467270666 | 0.888996866 |
| Q9EQH2 | Erap1 | Endoplasmic reticulum aminopeptidase 1 | 946.18 | 26.34 | 106599 | 20 | 32018785 | 26761759 | 35973577 | 28325499 | 29424585 | 25553445 | 25330507 | 28156379 | 24026919 | 23602235 | 26816195 | 25693203 | 28954555 | 27258149 | 1 | 4 | 0.830596671 | 0.891140198 |
| P57759 | Erp29 | Endoplasmic reticulum resident protein 29 | 421.21 | 42.75 | 28823 | 7 | 23509187 | 25599647 | 25032001 | 19475835 | 23330189 | 17848159 | 18310825 | 21764247 | 19237601 | 17908189 | 21789521 | 23503477 | 26228381 | 23041821 | 1 | 9 | 0.812924963 | 1.010749677 |
| Q3TZZ7 | Esyt2 | Extended synaptotagmin-2 | 119.37 | 4.5 | 94139 | 2 | 1064953.5 | 1177318.88 | 1586444.38 | 1074387.38 | 1557714.5 | 1733379.88 | 2089133.5 | 1733767.88 | 1960167.12 | 1562902 | 2275910 | 1470853.25 | 1649729.75 | 1518644.25 | 1 | 4 | 1.40529411 | 1.337898809 |
| Q9Z0E6 | Gbp2 | Guanylate-binding protein 2 | 328.57 | 16.3 | 66739 | 6 | 4043399.75 | 4796837 | 4765077 | 1587843.38 | 3894907.25 | 888460 | 622300.69 | 531811.94 | 305732.81 | 1592889.25 | 946624.88 | 1949364.75 | 1239605 | 1473114.75 | 1 | 1 | 0.206474298 | 0.367291653 |
| O08730 | Gyg1 | Glycogenin-1 | 45.97 | 4.8 | 37378 | 1 | 3535684.75 | 3298503.5 | 3479268.125 | 3422287 | 3660597.25 | 3340132 | 1 | 1929546.5 | 1 | 1664164 | 3925949 | 4476207.5 | 1 | 1 | 1 | 10 | 0.398580635 | 0.60373031 |
| Q61699 | Hsph1 | Heat shock protein 105 kDa | 1053.25 | 31.12 | 96407 | 17 | 13979423 | 14701681 | 15538737 | 12002855 | 13341430 | 11505919 | 12724204 | 11726425 | 9868152 | 10778031 | 13668253 | 16255472 | 15678931 | 14659286 | 1 | 5 | 0.813677024 | 1.082848759 |
| Q99J93 | Ifitm2 | Interferon-induced transmembrane protein 2 | 146.22 | 16.67 | 15743 | 1 | 53026101 | 38082793 | 41511925 | 25423989 | 37172865 | 28936005 | 24759839 | 20878133 | 20745715 | 22083409 | 18167669 | 21079547 | 32500441 | 20010805 | 1 | 1 | 0.601395863 | 0.587539416 |
| Q9CQW9 | Ifitm3 | Interferon-induced transmembrane protein 3 | 539.03 | 48.91 | 14954 | 5 | 65032261 | 45017053 | 49339585 | 30601089 | 45552461 | 33990385 | 30262735 | 24008263 | 23870285 | 26900781 | 21719847 | 21401057 | 39564185 | 21880273 | 1 | 1 | 0.590264938 | 0.554917821 |
| Q9CXY6 | Ilf2 | Interleukin enhancer-binding factor 2 | 732.59 | 53.85 | 43062 | 14 | 20563567 | 19345431 | 20165027 | 20896637 | 20034229 | 19942845 | 17199653 | 19258859 | 18856737 | 19025955 | 17879289 | 18110727 | 20047185 | 16444238 | 1 | 2 | 0.933460232 | 0.897004074 |
| Q8R4K2 | Irak4 | Interleukin-1 receptor-associated kinase 4 | 74.58 | 3.7 | 50872 | 1 | 1060406.25 | 1101917 | 1002067.94 | 1146826.38 | 1106412.25 | 922905.25 | 875769.12 | 996127 | 1027434 | 850466.44 | 930253.19 | 901446.62 | 803519.38 | 893944.81 | 1 | 0 | 0.862499278 | 0.814277672 |
| Q5PQL7 | Itm2c | Integral membrane protein 2C | 105.92 | 11.52 | 30480 | 2 | 1782204.5 | 1484172.5 | 1676717.88 | 1326225.5 | 1185751 | 1866026.75 | 1721062.5 | 2472878.75 | 2119065.5 | 1993988.5 | 1764260.75 | 2158080.75 | 2409142.75 | 2332807.5 | 1 | 1 | 1.364577411 | 1.452751307 |
| P70227 | Itpr3 | Inositol 1,4,5-trisphosphate receptor type 3 | 131.55 | 1.91 | 304275 | 4 | 639147.94 | 1556119.5 | 1574866.25 | 1596880.12 | 1718204 | 2475911.75 | 1699047 | 1880152 | 2103020.75 | 2218357.5 | 2138167.5 | 2033202 | 1825938 | 1802469.75 | 1 | 0 | 1.464526466 | 1.376065186 |
| P97825 | Jpt1 | Jupiter microtubule associated homolog 1 | 106.39 | 27.27 | 16081 | 2 | 2924789.25 | 4087840.5 | 3059245.5 | 4315181 | 3478096 | 1926713.38 | 1266849.12 | 2953416.75 | 1909610.25 | 2426835.25 | 2296861.25 | 1162097.38 | 1083107.12 | 2803540.5 | 1 | 0 | 0.586808587 | 0.513961912 |
| Q922S8 | Kif2c | Kinesin-like protein KIF2C | 95.48 | 4.72 | 81085 | 2 | 242709.95 | 1 | 275403.44 | 1 | 402230.97 | 345746.72 | 667487.75 | 865196.25 | 668064.5 | 636623.805 | 1032694.56 | 1090669.5 | 972195.62 | 532133.69 | 1 | 0 | 3.458609892 | 4.927076272 |
| Q921I2 | Klhdc4 | Kelch domain-containing protein 42 | 52.98 | 2.57 | 64860 | 1 | 1290601.62 | 1177804.25 | 1553057.88 | 1303753.75 | 1331525 | 1237858.88 | 1110034.88 | 1040095.44 | 911102.38 | 998585.06 | 1126783.88 | 1222575.25 | 1166588.62 | 1330169.75 | 1 | 4 | 0.795836198 | 0.910001683 |
| Q99KR3 | Lactb2 | Endoribonuclease LACTB2 | 137.57 | 11.46 | 32754 | 3 | 3368817.5 | 2992152.5 | 3056815.25 | 3435738 | 3577917 | 3739374.25 | 3664004.5 | 3710150.25 | 3487669.5 | 3706514.25 | 3435647 | 3399116 | 3602768.5 | 3727644.75 | 1 | 4 | 1.114187951 | 1.077596975 |
| Q6P4Z6 | Lcmt1 | Leucine carboxyl methyltransferase 1 | 44.66 | 2.71 | 38330 | 1 | 2115761.75 | 2183631.25 | 1710164.75 | 2253409.25 | 1937010.12 | 1755236.88 | 1529209.62 | 1369825.75 | 1517203.12 | 1654059 | 2062621.88 | 2175203.75 | 1453889.38 | 1705589.5 | 1 | 5 | 0.767210973 | 0.906534449 |
| P08905 | Lyz2 | Lysozyme C-2 | 403.81 | 49.32 | 16689 | 4 | 23830993 | 30157265 | 25073081 | 23394253 | 22318589 | 19638397 | 21972163 | 22522759 | 15524209 | 19922069 | 20743953 | 17378173 | 21759069 | 23216407 | 1 | 1 | 0.798078546 | 0.832479938 |
| Q3THS6 | Mat2a | S-adenosylmethionine synthase isoform type-2 | 436.56 | 25.06 | 43689 | 2 | 25768187 | 26025637 | 27081829 | 21670393 | 25990649 | 20955279 | 21122445 | 17982077 | 20557909 | 21761937 | 20024881 | 19028405 | 21115993 | 22779985 | 1 | 1 | 0.809090573 | 0.819419063 |
| Q9CQG2 | Mettl16 | U6 small nuclear RNA (adenine-(43)-N(6))-methyltransferase | 192.29 | 9.22 | 62341 | 4 | 31932109 | 2204096.5 | 2618429.75 | 24724479 | 35056745 | 2356071 | 1242981.12 | 2174420.5 | 1953284 | 1907661.38 | 40096529 | 2582518.75 | 2495090.75 | 31278831 | 1 | 10 | 0.099801442 | 0.989955573 |
| Q02527 | Mgat3 | Beta-1,4-mannosyl-glycoprotein 4-beta-N-acetylglucosaminyltransferase | 52.47 | 1.3 | 62022 | 1 | 7478302.5 | 7832926.5 | 4383691.5 | 6138261 | 8969417 | 5199680.5 | 3911749.5 | 4176866.5 | 2998327.5 | 4071656 | 7424262 | 7369179 | 3104444.75 | 4982473.5 | 1 | 5 | 0.584964367 | 0.821790622 |
| Q8K1X4 | Nckap1l | Nck-associated protein 1-like | 1623.37 | 32.72 | 128905 | 26 | 41108977 | 46357361 | 44054669 | 39256133 | 43858401 | 47610313 | 48682701 | 54800649 | 51818541 | 45465341 | 44643573 | 45631721 | 46334749 | 47776105 | 1 | 3 | 1.157206043 | 1.073832805 |
| O70310 | Nmt1 | Glycylpeptide N-tetradecanoyltransferase 1 | 804.42 | 38.51 | 56888 | 13 | 16814465 | 22363301 | 20927843 | 25617065 | 22983845 | 27136609 | 27996637 | 25131361 | 26008205 | 27772369 | 24512173 | 25642613 | 22160425 | 21705153 | 1 | 7 | 1.233092387 | 1.081126101 |
| P97333 | Nrp1 | Neuropilin-1 | 105.47 | 3.58 | 103000 | 2 | 2158467 | 1629235.62 | 1624815.75 | 1570022 | 1405104.25 | 1159563.25 | 1206585.75 | 818527.38 | 1518075.5 | 1235982.25 | 846107.31 | 1576009.88 | 1843321 | 1457088.5 | 1 | 7 | 0.70803359 | 0.852820868 |
| Q6PFD9 | Nup98 | Nuclear pore complex protein Nup98-Nup96 | 440.96 | 10.63 | 197241 | 4 | 4648943.5 | 4615951 | 4380564 | 3515120 | 4242840.5 | 3587882 | 3049986.75 | 2646424.5 | 1838146.5 | 2977052.5 | 4061990.75 | 3270712.75 | 2630142.5 | 3682314.5 | 1 | 2 | 0.658749532 | 0.796903085 |
| Q9DCE5 | Pak1ip1 | p21-activated protein kinase-interacting protein 1 | 269.41 | 20.42 | 42116 | 6 | 1412074.38 | 1034778.94 | 1321317.75 | 1364280.5 | 1623268.38 | 885734.2325 | 775390.81 | 1152813.12 | 849503.69 | 765229.31 | 1047729.75 | 756952 | 788854.88 | 1071687.62 | 1 | 2 | 0.655543924 | 0.67817055 |
| Q8BH04 | Pck2 | Phosphoenolpyruvate carboxykinase [GTP], mitochondrial | 1132.38 | 40.47 | 70528 | 19 | 44999569 | 51270049 | 49064105 | 55250529 | 52849193 | 58124393 | 55342169 | 56984741 | 56601573 | 53124605 | 54328641 | 46346685 | 47772705 | 54078461 | 1 | 10 | 1.105526861 | 0.9989136 |
| P17918 | Pcna | Proliferating cell nuclear antigen | 1091 | 85.06 | 28785 | 17 | 197016353 | 229116033 | 214982929 | 200755601 | 222280225 | 191110977 | 184421249 | 172244305 | 184241985 | 198398881 | 200986817 | 196086929 | 221462929 | 210013921 | 1 | 7 | 0.874328243 | 0.973252957 |
| Q9DCD0 | Pgd | 6-phosphogluconate dehydrogenase, decarboxylating | 1812.8 | 69.57 | 53247 | 27 | 151253345 | 164736529 | 175545713 | 136175393 | 153375793 | 178058113 | 165630561 | 182952209 | 192429441 | 181907857 | 181808641 | 192496833 | 207364001 | 189192449 | 1 | 0 | 1.153493072 | 1.233636823 |
| Q99K51 | Pls3 | Plastin-3 | 1757.78 | 62.38 | 70742 | 23 | 428140929 | 438180577 | 453587393 | 434619361 | 434747297 | 523168641 | 448838209 | 530365121 | 537169857 | 478631329 | 468837249 | 474582017 | 470946593 | 490720929 | 1 | 0 | 1.150231248 | 1.087738123 |

| | | | | | | | | | | | | | | | | | | | | | | | | |
|---|---|---|---|---|---|---|---|---|---|---|---|---|---|---|---|---|---|---|---|---|---|---|---|---|
| P53534 | Pygb | Glycogen phosphorylase, brain form | 997.77 | 23.39 | 96174 | 13 | 38852877 | 40796329 | 42306413 | 36641517 | 39003997 | 44483133 | 41864561 | 43885309 | 46652597 | 42594941 | 43729029 | 43060269 | 42179901 | 44278145 | 1 | 1 | 1.110725114 | 1.095940983 |
| P09527 | Rab7a | Ras-related protein Rab-7a | 1025.46 | 68.12 | 23504 | 15 | 97731393 | 109525561 | 103554041 | 103969705 | 107382417 | 97950625 | 93470105 | 96716025 | 94470273 | 95391009 | 92300625 | 98579705 | 101028865 | 97031873 | 1 | 2 | 0.915418997 | 0.931081341 |
| Q9CW46 | Raver1 | Ribonucleoprotein PTB-binding 1 | 387.71 | 21.66 | 79382 | 8 | 1 | 541926 | 1 | 1 | 1 | 16877637 | 808436.12 | 699444.5 | 473327.84 | 597908 | 1674871.25 | 524371.38 | 744044.69 | 1242835.5 | 1 | 1 | 35.902706 | 9.655589329 |
| Q60972 | Rbbp4 | Histone-binding protein RBBP4 | 390.27 | 22.35 | 47656 | 3 | 26910333 | 26352515 | 26713375 | 27903791 | 29951127 | 25853221 | 24061037 | 25295337 | 24680683 | 26449649 | 26769591 | 26927769 | 26762301 | 25678185 | 1 | 7 | 0.916628318 | 0.96257135 |
| O89086 | Rbm3 | RNA-binding protein 3 | 263.18 | 48.37 | 16605 | 4 | 16201317 | 14056728 | 17451029 | 17496437 | 18559707 | 11945609 | 12359547 | 9477812 | 12530498 | 14840802 | 15729302 | 10670674 | 14948069 | 14099241 | 1 | 3 | 0.730067556 | 0.827421084 |
| D3Z8L7 | Rras | Ras-related protein R-Ras | 235.96 | 28.9 | 23909 | 5 | 5490828.5 | 9192823 | 5410132.5 | 5420434.5 | 6203214 | 9938922 | 10849354 | 12619552 | 10807559 | 9097382 | 6464193 | 2961916.5 | 8718141 | 10152219 | 1 | 7 | 1.680866476 | 1.115178124 |
| Q99P72 | Rtn4 | Reticulon-4 | 984.33 | 25.13 | 126613 | 15 | 62349229 | 58925973 | 62964929 | 54204753 | 69060793 | 77515713 | 77575649 | 67254601 | 75303641 | 79078345 | 60750797 | 72058809 | 76561441 | 69924809 | 1 | 3 | 1.225108924 | 1.135328048 |
| Q4G075 | Serpinb1a | Leukocyte elastase inhibitor A | 69.24 | 2.64 | 42727 | 1 | 1023972.69 | 1064173.5 | 1096896.38 | 1305444.12 | 1008452.94 | 1868474.5 | 1546595.12 | 1100378.75 | 1621885.25 | 1312841.5 | 1235579.75 | 1489651.5 | 1297659.62 | 1510886.75 | 1 | 2 | 1.3548385 | 1.257919252 |
| Q6AYE2 | Sh3glb1 | Endophilin-B1 | 215.15 | 10.68 | 40787 | 3 | 6654449 | 7580770 | 5955345.5 | 7714775 | 7210518 | 5255211 | 5804351 | 5699925.5 | 5476036 | 6283219.5 | 6525239.5 | 5816611 | 5211846 | 6372754 | 1 | 2 | 0.812132895 | 0.85169679 |
| Q6P9P0 | Slf2 | SMC5-SMC6 complex localization factor protein 2 | 36.1 | 0.55 | 143970 | 1 | 1371724 | 1059111.62 | 392924.06 | 1025785.38 | 1202280.12 | 748016.88 | 1 | 1 | 197343.52 | 1 | 1163531.75 | 1828177.38 | 189973.88 | 1379244.88 | 1 | 7 | 0.187133039 | 1.12853467 |
| Q62189 | Snrpa | U1 small nuclear ribonucleoprotein A | 295.14 | 31.71 | 31835 | 5 | 5356152.5 | 5656992.5 | 5636798.5 | 5084548 | 5502430 | 4676490.5 | 5325697.5 | 3946715.5 | 3584942.25 | 4370346.5 | 5313226 | 4528311.5 | 3996467.25 | 4890053.5 | 1 | 1 | 0.804209545 | 0.859497752 |
| Q58A65 | Spag9 | C-Jun-amino-terminal kinase-interacting protein 4 | 403.93 | 9.01 | 146219 | 7 | 4373358.5 | 4123626 | 3886400 | 3164691 | 5128186 | 6293938.5 | 5564882.5 | 6968959 | 6788624.5 | 5099077 | 5801966 | 6487378 | 4663699 | 6591536 | 1 | 1 | 1.485543288 | 1.423406439 |
| Q9Z2I8 | Suclg2 | Succinate--CoA ligase [GDP-forming] subunit beta, mitochondrial | 832.03 | 43.42 | 46840 | 14 | 46315337 | 40460253 | 48964709 | 41624685 | 43707373 | 55236441 | 51140345 | 52193109 | 55620869 | 46938513 | 52441313 | 56616301 | 50560709 | 52940785 | 1 | 0 | 1.181193708 | 1.201863899 |
| Q3TBT3 | Tmem173 | Stimulator of interferon genes protein | 106.37 | 11.9 | 42830 | 3 | 3873826.25 | 2567173 | 2623640.25 | 2622259.75 | 2671752.25 | 1795765 | 1862165.12 | 1654236.62 | 2591512.5 | 2079581 | 1997885.5 | 2156260.5 | 2350312 | 2926366 | 1 | 4 | 0.6952784 | 0.821005371 |
| Q5XIK2 | Tmx2 | Thioredoxin-related transmembrane protein 2 Rattus norvegicus | 67.31 | 5.08 | 33866 | 1 | 290727.25 | 284959.56 | 230181.39 | 201141.98 | 298347.53 | 196443.27 | 196502.42 | 209914.42 | 166982.14 | 147189.75 | 226062.47 | 360765.41 | 215834.38 | 155584.45 | 1 | 7 | 0.702513949 | 0.91760931 |
| Q91XB0 | Trex1 | Three-prime repair exonuclease 1 | 159.45 | 14.01 | 33675 | 3 | 3858302.5 | 6591903 | 3002202.5 | 5408457.5 | 5267453.5 | 2709213 | 1787539.12 | 2425848 | 2479948.25 | 3220306 | 2229514 | 2075390.62 | 2429171.5 | 2635683 | 1 | 0 | 0.523155151 | 0.485412966 |
| Q5NC05 | Ttf2 | Transcription termination factor 2 | 100.85 | 2.9 | 125530 | 3 | 1898142.75 | 1383248 | 1174040.62 | 1850015.38 | 1632152.38 | 1321264.38 | 585877.25 | 415239.88 | 235110.8 | 648483.5 | 1504525.5 | 1336907.25 | 506270.47 | 1107042.38 | 1 | 3 | 0.40389742 | 0.701525979 |
| Q3UX10 | Tubal3 | Tubulin alpha chain-like 32 | 144.57 | 11.66 | 49988 | 1 | 18683813 | 24827005 | 17155457 | 15835367 | 20305229 | 26396885 | 27993255 | 27852099 | 25230437 | 23117065 | 30590573 | 18002667 | 32408537 | 28968605 | 1 | 3 | 1.34897182 | 1.41997129 |
| A2AN08 | Ubr4 | E3 ubiquitin-protein ligase UBR4 | 1210.58 | 6.12 | 572290 | 22 | 13464246 | 15326411 | 13744404 | 14036837 | 11869789 | 17011859 | 14981061 | 17499269 | 17161905 | 15428996 | 14048884 | 15509513 | 16232494 | 14690013 | 1 | 2 | 1.199314243 | 1.104606466 |
| Q6ZQM8 | Ugt1a7c | UDP-glucuronosyltransferase 1-7C | 1044.94 | 45.01 | 59758 | 19 | 78058561 | 83750217 | 64516753 | 77695905 | 77941233 | 86669297 | 84737585 | 84050017 | 83624129 | 87147425 | 85040953 | 78555441 | 81012153 | 88514065 | 1 | 2 | 1.115890341 | 1.09016744 |
| P63024 | Vamp3 | Vesicle-associated membrane protein 3 | 90.95 | 32.04 | 11480 | 2 | 1 | 1 | 1 | 1 | 411596.5 | 667802.56 | 346006.72 | 485681.155 | 422271.72 | 506643.62 | 1 | 605893.94 | 1 | 1 | 1 | 9 | 5.899909682 | 1.840063788 |
| Q9DBR1 | Xrn2 | 5'-3' exoribonuclease 2 | 1106.68 | 27.87 | 108687 | 21 | 15612037 | 14278676 | 19510667 | 14982429 | 17092421 | 21686991 | 19275043 | 20932755 | 24596929 | 19800279 | 20272161 | 18728573 | 22759661 | 18884761 | 1 | 2 | 1.30457677 | 1.237249748 |
| Q91WQ3 | Yars | Tyrosine--tRNA ligase, cytoplasmic | 720.92 | 38.07 | 59105 | 16 | 28210869 | 21748569 | 27875767 | 28025875 | 24279579 | 28555731 | 28074267 | 30390861 | 30791929 | 28821605 | 28280917 | 31469807 | 26727817 | 27579871 | 1 | 6 | 1.126737748 | 1.095530145 |
| Q8BYK6 | Ythdf3 | YTH domain-containing family protein 3 | 215.77 | 8.72 | 63961 | 3 | 1038374.69 | 836985.88 | 794061.81 | 935737.44 | 1263717.88 | 736798.56 | 782098.88 | 634261.75 | 631624.19 | 796678.69 | 987328.81 | 565266.06 | 938025.19 | 870383.94 | 1 | 8 | 0.735582672 | 0.86287955 |
| P45952 | Acadm | Medium-chain specific acyl-CoA dehydrogenase, mitochondrial | 444.97 | 28.5 | 46481 | 9 | 42677457 | 50715201 | 46500937 | 45997517 | 47525865 | 40269057 | 44646721 | 39783081 | 44266085 | 38346169 | 45339965 | 44181925 | 44745873 | 49146833 | 2 | 7 | 0.888157818 | 0.982226091 |
| Q8VCW8 | Acsf2 | Acyl-CoA synthetase family member 2, mitochondrial | 435.77 | 26.02 | 67951 | 8 | 6112308.5 | 6232736.5 | 7270555 | 10534029 | 6490749 | 12155309 | 12857558 | 8889328 | 10873004 | 8065915 | 13921524 | 12177727 | 7943379 | 8942086 | 2 | 2 | 1.442155264 | 1.466439429 |
| Q9QXG4 | Acss2 | Acetyl-coenzyme A synthetase, cytoplasmic | 491.28 | 18.54 | 78862 | 8 | 11705761 | 10209376 | 11267837 | 12468933 | 11857124 | 14087967 | 13800367 | 12570557 | 13475720 | 11850761 | 13505003 | 11802905 | 11221238 | 12561764 | 2 | 6 | 1.143913762 | 1.067026108 |
| Q99JW2 | Acy1 | Aminoacylase-1 | 424.76 | 28.92 | 45781 | 7 | 7514963 | 4298089.5 | 6403944.5 | 6028198 | 5054992.5 | 2668281.5 | 4369942 | 4973562.5 | 4238939 | 3666013.5 | 3885533 | 5359361 | 5295716.5 | 5136124.5 | 2 | 6 | 0.679747817 | 0.83944578 |
| Q9Z0F8 | Adam17 | Disintegrin and metalloproteinase domain-containing protein 17 | 243.27 | 4.47 | 93056 | 3 | 352922.03 | 434138.34 | 751878.5 | 450703.3825 | 263874.66 | 734034.94 | 871016.44 | 1050635 | 1341085.75 | 495135.84 | 801569.75 | 201937.03 | 405257.72 | 1066279.62 | 2 | 8 | 1.993287889 | 1.372878603 |
| P54923 | Adprh | [Protein ADP-ribosylarginine] hydrolase | 158.22 | 9.67 | 40068 | 3 | 4881424 | 3971885 | 4591478 | 4401517 | 4606919.5 | 4017540.5 | 3959218.75 | 4230903 | 3902701.5 | 3637002.5 | 4488538 | 4455396 | 4220719 | 4200843.5 | 2 | 6 | 0.879489141 | 0.966759656 |
| B5DFN2 | Ahcyl1 | S-adenosylhomocysteine hydrolase-like protein 1 | 241.82 | 16.15 | 53753 | 3 | 5914869 | 8225328.5 | 8047324 | 7341394.5 | 8698729 | 9572509 | 8872227 | 8347488.5 | 8380391.5 | 8857981 | 9192662 | 8321410 | 8609571 | 9353807 | 2 | 2 | 1.151799882 | 1.160071788 |
| Q8R0Y6 | Aldh1l1 | Cytosolic 10-formyltetrahydrofolate dehydrogenase | 426.66 | 15.08 | 98709 | 9 | 6059573.5 | 7205715.5 | 8514740 | 7488813 | 7148226 | 5200596 | 5326867 | 6174886.5 | 6405054.5 | 5357527 | 7682267 | 5702402 | 4765501.5 | 5613908 | 2 | 4 | 0.781637088 | 0.815691646 |
| Q9WVM3 | Anapc7 | Anaphase-promoting complex subunit 7 | 46.32 | 2.65 | 63021 | 1 | 170881.94 | 225512.88 | 205361.45 | 196778.5375 | 185357.88 | 130679.76 | 150193.03 | 158993.08 | 176718.61 | 178380.92 | 178774.42 | 202512.97 | 154957.02 | 184571.31 | 2 | 5 | 0.807979783 | 0.915770248 |
| P14824 | Anxa6 | Annexin A6 | 1282.66 | 47.99 | 75885 | 25 | 57874777 | 59606777 | 50453753 | 57954801 | 48983085 | 58794525 | 59524529 | 71684929 | 67952121 | 62019637 | 53976017 | 57076297 | 53751265 | 58010533 | 2 | 10 | 1.164084928 | 1.013258648 |
| O35643 | Ap1b1 | AP-1 complex subunit beta-1 | 2073.93 | 49.63 | 103935 | 20 | 43716193 | 42529389 | 50980005 | 37876845 | 44609745 | 53359701 | 46317473 | 58898685 | 59589265 | 45151273 | 45600829 | 56944301 | 55968641 | 41221777 | 2 | 5 | 1.198460643 | 1.136347736 |
| Q3TAP4 | Ap5b1 | AP-5 complex subunit beta-1 | 89.99 | 3.31 | 94679 | 2 | 495568.12 | 610638.62 | 653603.31 | 594973.88 | 572988.12 | 443462.97 | 505977.75 | 323197.91 | 537224.19 | 440818.47 | 559783.56 | 386920.81 | 510894.88 | 463886.44 | 2 | 2 | 0.768735151 | 0.820370258 |
| P51908 | Apobec1 | C->U-editing enzyme APOBEC-1 | 150.25 | 18.78 | 27522 | 1 | 4181783 | 3128322 | 3258482 | 3074789.75 | 3974909.5 | 2983143.5 | 1326838.38 | 2388297 | 2709799 | 3244947.75 | 3665762.75 | 2977916.5 | 2662353.5 | 3459895 | 2 | 6 | 0.718175732 | 0.905729959 |
| Q78IK4 | Apool | MICOS complex subunit Mic27 | 389.35 | 33.96 | 29261 | 6 | 8955794 | 9403101 | 9080211 | 6970131 | 10111369 | 10838823 | 12188252 | 10543495 | 9161167 | 12014285 | 10280277 | 10627857 | 9798840 | 9853381 | 2 | 2 | 1.229678275 | 1.138808482 |
| A2BH40 | Arid1a | AT-rich interactive domain-containing protein 1A | 160.72 | 1.14 | 242092 | 0 | 27109421 | 27486317 | 30441325 | 29938315 | 29588977 | 24688835 | 24675491 | 25498973 | 25350019 | 27800611 | 27259883 | 25294327 | 26233273 | 29537277 | 2 | 3 | 0.885515167 | 0.936648249 |
| Q5FVI6 | Atp6v1c1 | V-type proton ATPase subunit C 1 Rattus norvegicus | 974.91 | 43.98 | 43901 | 17 | 40574881 | 40408421 | 36351181 | 37687361 | 33457207 | 35273673 | 32090027 | 35615617 | 29545309 | 30895645 | 29103067 | 34799201 | 35215713 | 36383877 | 2 | 4 | 0.867047399 | 0.898653307 |
| Q8BVE3 | Atp6v1h | V-type proton ATPase subunit H | 696.59 | 32.3 | 55855 | 13 | 20321119 | 20710759 | 23600781 | 18563459 | 19201917 | 27175529 | 22681483 | 23013645 | 26928313 | 23830863 | 22637419 | 25989765 | 25466551 | 23831829 | 2 | 1 | 1.207345756 | 1.195403359 |

| | | | | | | | | | | | | | | | | | | | | | | | | |
|---|---|---|---|---|---|---|---|---|---|---|---|---|---|---|---|---|---|---|---|---|---|---|---|---|
| Q9CZP5 | Bcs1l | Mitochondrial chaperone BCS1 | 72.75 | 4.55 | 47406 | 1 | 279154.44 | 219089.17 | 206571.38 | 316904.25 | 264576.97 | 332101.66 | 293105.88 | 336330.41 | 323191.06 | 282350.47 | 299724.09 | 274138.09 | 336352.5 | 229844.08 | 2 | 6 | 1.218288189 | 1.107889022 |
| O35658 | C1qbp | Complement component 1 Q subcomponent-binding protein, mitochondrial | 427.26 | 43.53 | 31013 | 1 | 9895254 | 7132176.5 | 11802255 | 8187858 | 7418910.5 | 12389617 | 11660238 | 14080177 | 13593956 | 10148720 | 11269229 | 11573135 | 12072293 | 8713768 | 2 | 4 | 1.39238626 | 1.227270098 |
| P40124 | Cap1 | Adenylyl cyclase-associated protein 1 | 2231.67 | 70.25 | 51565 | 35 | 294169825 | 366957793 | 344019201 | 316606305 | 327385057 | 341739265 | 407582849 | 405840417 | 393304993 | 392298881 | 350190241 | 369820225 | 350521921 | 365544481 | 2 | 3 | 1.176836743 | 1.088505564 |
| Q8BRN9 | Cc2d1b | Coiled-coil and C2 domain-containing protein 1B | 177.26 | 11.32 | 93091 | 4 | 554515.06 | 907815.56 | 706672 | 1 | 740409.44 | 1351665.5 | 1111377.36 | 1562820.75 | 782064.81 | 748958.38 | 1697086 | 1037577.94 | 794477.75 | 1176380.563 | 2 | 1 | 1.909968329 | 2.021680214 |
| Q9JIG7 | Ccdc22 | Coiled-coil domain-containing protein 22 | 209.62 | 13.24 | 70844 | 5 | 1427041.5 | 1870454.5 | 1686675.5 | 1998219.88 | 1917824 | 1150606.5 | 1376797.62 | 1428732.62 | 1529594 | 922738.69 | 1601055.5 | 1691953.75 | 897161.06 | 1533065.38 | 2 | 4 | 0.72003532 | 0.803805785 |
| Q8WTY4 | Ciapin1 | Anamorsin | 383.78 | 35.28 | 33429 | 7 | 4608548.5 | 8163353 | 5240400 | 7114874.5 | 7039459.5 | 8085573 | 9440385 | 7324889.5 | 8398237 | 9839864 | 7227465 | 6190520.5 | 8402461 | 7653529 | 2 | 5 | 1.339554101 | 1.145362852 |
| Q60809 | Cnot7 | CCR4-NOT transcription complex subunit 7 | 78.56 | 10.18 | 32718 | 2 | 1 | 452534.75 | 1 | 1 | 321391.88 | 544725.06 | 468870.06 | 738707.44 | 565524.69 | 108246.51 | 73928.09 | 423870.47 | 296847.09 | 264881.8833 | 2 | 7 | 3.134747225 | 1.711278862 |
| P11240 | Cox5a | Cytochrome c oxidase subunit 5A, mitochondrial | 288.69 | 38.36 | 16130 | 6 | 40878529 | 47620297 | 40817437 | 37513889 | 38771337 | 32348581 | 35937237 | 40773493 | 33111355 | 28772209 | 31308093 | 29085567 | 31354793 | 34658793 | 2 | 0 | 0.831428196 | 0.768520978 |
| Q9CPQ1 | Cox6c | Cytochrome c oxidase subunit 6C | 67.57 | 19.74 | 8469 | 2 | 5674408 | 3656424 | 3796538 | 4099267.5 | 3276160.75 | 3148131.5 | 3580100.25 | 3345972.25 | 2841076.75 | 3019356.25 | 2281812.5 | 2360568.5 | 2293236.25 | 2253690 | 2 | 0 | 0.777193279 | 0.560247139 |
| Q9QXK7 | Cpsf3 | Cleavage and polyadenylation specificity factor subunit 3 | 176.93 | 8.48 | 77505 | 4 | 2630602.25 | 2300776.5 | 1728431 | 2361033.5 | 2241506 | 2806665.5 | 2839439 | 3075626 | 2360146.75 | 3194808.5 | 2726656.5 | 3271560.25 | 2636069.5 | 1829672.5 | 2 | 4 | 1.267647223 | 1.161387216 |
| Q99388 | Csprs | Component of Sp100-rs Mus musculus | 91.15 | 8.65 | 24061 | 1 | 2141751.5 | 2036986.12 | 2158798.5 | 1540366.88 | 1584585.62 | 1324223.75 | 1275136.5 | 1619174.88 | 1521761.75 | 1286175.25 | 1327284.38 | 1794179.88 | 1312826 | 1244349 | 2 | 2 | 0.742560695 | 0.750151399 |
| P97315 | Csrp1 | Cysteine and glycine-rich protein 1 | 258.97 | 33.16 | 20583 | 5 | 1776077.75 | 1699707.62 | 1975467.25 | 2826553.5 | 1726693.5 | 4060239.5 | 2485635.25 | 4567123.5 | 4943919.5 | 2277767.25 | 4142359 | 4486704 | 2507449.25 | 1209048.62 | 2 | 6 | 1.83264388 | 1.542501042 |
| Q99LI7 | Cstf3 | Cleavage stimulation factor subunit 3 | 372.6 | 13.81 | 82877 | 7 | 2502103 | 2949152.25 | 3310061.75 | 3082721.25 | 2948490 | 3288276 | 3569139.5 | 3212734 | 3500076.5 | 3676061 | 3380831.75 | 3028354.5 | 3154689.5 | 2977506 | 2 | 5 | 1.165878253 | 1.059773348 |
| Q9WTX6 | Cul1 | Cullin-1 | 508.56 | 14.69 | 89692 | 10 | 9356674 | 8197263 | 7475768 | 8358356 | 10032447 | 11403163 | 8733838 | 12266343 | 11508169 | 10748202 | 10499824 | 12952264 | 12832699 | 11439313 | 2 | 0 | 1.25884559 | 1.373892839 |
| P11030 | Dbi | Acyl-CoA-binding protein | 144.97 | 39.08 | 10027 | 2 | 8975738 | 7477482 | 6957561 | 8622616 | 7808447.5 | 4107992.25 | 6293613.5 | 7137555 | 4203308 | 7113129 | 7026639.5 | 5648527 | 5563802 | 6502662 | 2 | 1 | 0.724253561 | 0.776245139 |
| Q91VR5 | Ddx1 | ATP-dependent RNA helicase DDX1 | 1054.07 | 34.32 | 82500 | 19 | 21494465 | 23311527 | 25019387 | 23063183 | 23753523 | 23760623 | 26720961 | 26121527 | 24885585 | 26870247 | 25130829 | 25252313 | 26245641 | 25898643 | 2 | 0 | 1.100451377 | 1.09873964 |
| Q62095 | Ddx3y | ATP-dependent RNA helicase DDX3Y | 1104.92 | 32.98 | 73428 | 2 | 77379209 | 69522481 | 72049065 | 60994129 | 71025609 | 60714649 | 64453085 | 58073865 | 58504413 | 62334405 | 53894289 | 56623925 | 60489421 | 59040217 | 2 | 0 | 0.866398809 | 0.819327609 |
| D4A2Z8 | Dhx36 | ATP-dependent DNA/RNA helicase DHX36 | 261.69 | 7.4 | 113843 | 6 | 1602403.38 | 2228548.25 | 2125017.75 | 1845307 | 3894404.25 | 1977454.38 | 1425083 | 1334479.25 | 1297270.75 | 1280549.5 | 1438098.62 | 1360321.75 | 1207351.75 | 1171439.25 | 2 | 0 | 0.625430628 | 0.553325148 |
| Q9QYJ3 | Dnajb1 | DnaJ homolog subfamily B member 1 | 165.74 | 15 | 38167 | 5 | 4165890 | 3318580 | 3386349.25 | 3622478 | 3842204.5 | 5365432.5 | 5675294 | 4051349.75 | 5571722 | 3914628.5 | 4397374 | 3961700.75 | 5283370 | 5117586 | 2 | 1 | 1.34048291 | 1.278941736 |
| Q9CQV7 | Dnajc19 | Mitochondrial import inner membrane translocase subunit TIM14 | 72.39 | 16.38 | 12437 | 1 | 2629887.25 | 1677089.62 | 1913949.5 | 2179990.75 | 1895540.75 | 1589805.25 | 1882904.62 | 1775896 | 1273819.5 | 1133331.88 | 2185832 | 2130678.25 | 1962397.62 | 1985468.25 | 2 | 7 | 0.743533101 | 1.003303299 |
| Q91YW3 | Dnajc3 | DnaJ homolog subfamily C member 3 | 326.8 | 15.67 | 57464 | 5 | 2656988.75 | 1071912 | 1458179.25 | 816552.88 | 1249762.5 | 644871.69 | 694546.25 | 960467.06 | 816284.155 | 965251.62 | 1193582.5 | 652094.62 | 880727.56 | 981326.06 | 2 | 4 | 0.562691066 | 0.638964675 |
| P68101 | Eif2s1 | Eukaryotic translation initiation factor 2 subunit 1 | 1198.18 | 70.79 | 36108 | 20 | 74835921 | 83887617 | 85171801 | 80947633 | 85725169 | 93875897 | 84537305 | 88022513 | 86702537 | 89098241 | 81731305 | 88946153 | 94515857 | 91406385 | 2 | 3 | 1.077132999 | 1.085689756 |
| P48193 | Epb41 | Protein 4.1 | 115.17 | 5.48 | 95911 | 3 | 2396828.5 | 1270234.38 | 2874250.5 | 3088874.75 | 3092124 | 3803812.5 | 3551729 | 2980751 | 3441233.5 | 3982010.5 | 3149823 | 3099586 | 3051549.75 | 3054908.5 | 2 | 4 | 1.395936235 | 1.213995845 |
| Q8R1F1 | Fam129b | Niban-like protein 1 | 513.98 | 19.36 | 84819 | 9 | 21481605 | 20756937 | 21639297 | 18641639 | 18686853 | 24123223 | 23228745 | 24436909 | 23561257 | 21288829 | 22190621 | 23111867 | 22226109 | 22595651 | 2 | 0 | 1.152486824 | 1.113125127 |
| A0A0B4 | Fcgr4 | Low affinity immunoglobulin gamma Fc region receptor IV | 45.23 | 4.02 | 28398 | 1 | 2495273 | 2113805.5 | 2117607.5 | 2473083 | 2382982.25 | 1894300.5 | 1841483.38 | 2000441.75 | 1741781.12 | 2120373.5 | 1889272.12 | 1783561.25 | 1607984.75 | 1968628.62 | 2 | 0 | 0.828678786 | 0.782353711 |
| Q60634 | Flot2 | Flotillin-2 | 245.6 | 10.28 | 47038 | 4 | 765938.69 | 1848567.12 | 2336999.75 | 1 | 1162812.25 | 1660661.62 | 2537925 | 2745098 | 2613886 | 2558559.25 | 1986047.88 | 2485231.5 | 1786466.38 | 1669692.38 | 2 | 5 | 1.981599299 | 1.620670754 |
| Q80TY0 | Fnbp1 | Formin-binding protein 1 | 797.37 | 28.57 | 71344 | 16 | 21962303 | 26344899 | 22767065 | 27794309 | 26876757 | 26968981 | 28623121 | 29073977 | 27696497 | 28881731 | 23275953 | 23090863 | 21758103 | 24006653 | 2 | 6 | 1.123256853 | 0.915854786 |
| Q9CWG1 | Glipr1 | Glioma pathogenesis-related protein 1 Mus musculus | 61.17 | 5.49 | 29129 | 1 | 1234218.25 | 1205148.25 | 1037347 | 1332321.75 | 1437550.5 | 1610373.62 | 1616885.88 | 1726157.62 | 1560643.75 | 1267817.25 | 1606324.75 | 1614322.5 | 1579347.5 | 1447292.12 | 2 | 0 | 1.245781044 | 1.250140301 |
| Q66H74 | Golph3l | Golgi phosphoprotein 3-like | 155.56 | 8.77 | 32922 | 1 | 1976959.75 | 1936485 | 1743979.62 | 1889268.88 | 1744210.38 | 1789250.12 | 1735125.75 | 1400208.25 | 1648827.5 | 1487278.12 | 1899141 | 1607180.62 | 1340708.25 | 1908126.12 | 2 | 6 | 0.867589425 | 0.908840014 |
| P11352 | Gpx1 | Glutathione peroxidase 1 | 216.24 | 29.85 | 22329 | 4 | 12420509 | 9940401 | 8472454 | 11677418 | 11073843 | 8316239 | 9854850 | 8782298 | 8203964 | 7889484 | 8033744 | 7628524 | 8737673 | 6710114 | 2 | 1 | 0.80334303 | 0.725722514 |
| Q8BMQ2 | Gtf3c4 | General transcription factor 3C polypeptide 4 | 48.54 | 2.33 | 91613 | 1 | 148779.95 | 95456.84 | 237425.92 | 144857.19 | 140392.62 | 273780.12 | 244024.67 | 272324.19 | 235925.58 | 177443.38 | 171055.56 | 285742.25 | 298756.91 | 203142.75 | 2 | 2 | 1.569276689 | 1.562592612 |
| P01900 | H2-D1 | H-2 class I histocompatibility antigen, D-D alpha chain | 821.73 | 49.86 | 41110 | 8 | 141752129 | 156688449 | 144794481 | 127365297 | 153014273 | 133145489 | 112402753 | 129501857 | 126400217 | 123209249 | 133765249 | 119186737 | 136398545 | 136591089 | 2 | 3 | 0.863248945 | 0.908531971 |
| P01902 | H2-K1 | H-2 class I histocompatibility antigen, K-D alpha chain | 881.15 | 45.38 | 41490 | 10 | 96228993 | 98687489 | 100711857 | 77917329 | 95276449 | 78757441 | 74367601 | 81868753 | 77528273 | 75264657 | 79332945 | 73915873 | 86289529 | 77255185 | 2 | 2 | 0.82715109 | 0.844652803 |
| A1EC95 | Heatr6 | HEAT repeat-containing protein 6 | 82.91 | 2.16 | 136505 | 2 | 577690.69 | 396189.25 | 377223.16 | 446741.7975 | 435864.09 | 352418.34 | 363479.575 | 422473.09 | 369836.62 | 309190.25 | 312233.9 | 362360.72 | 237844.64 | 336496.34 | 2 | 0 | 0.813623388 | 0.69891356 |
| E9QAM5 | Helz2 | Helicase with zinc finger domain 2 | 151.11 | 2.07 | 331563 | 4 | 379286.59 | 418748.47 | 384684.03 | 307553.22 | 292926.03 | 180810.45 | 341476.72 | 227275 | 215808.92 | 241342.7725 | 312501.56 | 320097.28 | 280820.34 | 158677.58 | 2 | 4 | 0.676713204 | 0.75152658 |
| P29416 | Hexa | Beta-hexosaminidase subunit alpha | 1086.12 | 42.23 | 60613 | 6 | 70220577 | 94596145 | 76982145 | 71568745 | 87501009 | 66215849 | 63692437 | 76452817 | 68969409 | 63487673 | 73454281 | 71298273 | 68323281 | 81361849 | 2 | 6 | 0.845210045 | 0.918124008 |
| P20060 | Hexb | Beta-hexosaminidase subunit beta | 738.72 | 39.37 | 61116 | 16 | 55997421 | 70052937 | 50176317 | 45176281 | 61938809 | 45185985 | 41383497 | 48473765 | 44602829 | 43844301 | 49839685 | 44752501 | 52119533 | 53569377 | 2 | 5 | 0.788766093 | 0.883566777 |
| P62959 | Hint1 | Histidine triad nucleotide-binding protein 1 | 163.67 | 35.71 | 13777 | 3 | 41841437 | 48005725 | 47729601 | 49812909 | 48736601 | 38098337 | 43058569 | 42895177 | 39513597 | 41674329 | 43209365 | 43843257 | 39779449 | 45466081 | 2 | 3 | 0.869195987 | 0.912108116 |
| P62804 | Hist1h4b | Histone H4 | 1131.95 | 58.25 | 11367 | 16 | 4833988609 | 2000589441 | 2137045889 | 1928947969 | 2121537153 | 1806478977 | 1684927361 | 1617567873 | 1833690625 | 2107090305 | 1791839233 | 1792816641 | 1988163585 | 2023052161 | 2 | 3 | 0.694953106 | 0.729132238 |
| Q3TRM8 | Hk3 | Hexokinase-3 | 2099.18 | 46.85 | 100101 | 14 | 111634473 | 134472689 | 119574729 | 111517449 | 118247505 | 113794553 | 107501257 | 111184673 | 103951961 | 107363473 | 108582865 | 94170505 | 105660569 | 109304409 | 2 | 0 | 0.913256862 | 0.876901002 |

| | | | | | | | | | | | | | | | | | | | | | | | | |
|---|---|---|---|---|---|---|---|---|---|---|---|---|---|---|---|---|---|---|---|---|---|---|---|---|
| P38060 | Hmgcl | Hydroxymethylglutaryl-CoA lyase, mitochondrial | 605.33 | 46.77 | 34239 | 10 | 20361335 | 25321399 | 25609585 | 17527379 | 23998657 | 24400257 | 28256333 | 28503143 | 28488217 | 27767281 | 28131443 | 21966409 | 29815053 | 28196835 | 2 | 3 | 1.218021934 | 1.197829688 |
| P50171 | Hsd17b8 | Estradiol 17-beta-dehydrogenase 8 | 194.79 | 18.15 | 26588 | 4 | 3826551 | 2408823.25 | 2803414 | 4490665.5 | 4305340 | 4659270.5 | 5484088.5 | 4018437.5 | 4983222 | 5325776 | 4845972 | 4886681 | 4865597 | 3941924.5 | 2 | 2 | 1.372081721 | 1.299438527 |
| Q8BTX9 | Hsdl1 | Inactive hydroxysteroid dehydrogenase-like protein 1 | 138.33 | 10.91 | 36868 | 2 | 1532226.12 | 1417680.75 | 1644205.25 | 1829637.88 | 1761696.75 | 1051358.75 | 1358895.12 | 1245302.25 | 1625902 | 1276974.12 | 1263502.12 | 1304692.12 | 1262580.62 | 1349341.75 | 2 | 0 | 0.801230823 | 0.791055878 |
| P38647 | Hspa9 | Stress-70 protein, mitochondrial | 2182.07 | 55.67 | 73461 | 31 | 183261889 | 181052913 | 198643089 | 186770193 | 185384353 | 213031393 | 212484529 | 189584337 | 194952609 | 203820193 | 195486641 | 184821457 | 203430913 | 198804001 | 2 | 4 | 1.084225833 | 1.046054706 |
| O88351 | Ikbkb | Inhibitor of nuclear factor kappa-B kinase subunit beta | 264.42 | 10.7 | 86690 | 5 | 9063181 | 2698539.25 | 8796161 | 2351809.5 | 2684371 | 9118863 | 2988006 | 12217903 | 10922312 | 9449693 | 8293788 | 10684524 | 9719247 | 9020347 | 2 | 3 | 1.746372945 | 1.842121933 |
| Q80V86 | Ints8 | Integrator complex subunit 8 | 34.38 | 0.7 | 113353 | 1 | 931067.31 | 809143.56 | 771522.56 | 846315.19 | 706830.19 | 541543.5 | 729858.5 | 709024.19 | 523866.44 | 640948.31 | 647398.12 | 750522.69 | 719151.62 | 601051.25 | 2 | 2 | 0.773760077 | 0.835856309 |
| Q6P9P6 | Kif11 | Kinesin-like protein KIF11 | 575.89 | 14.16 | 118027 | 9 | 1801478.25 | 1844002.5 | 1682134.75 | 1739286 | 1683549 | 2769442.25 | 2144045.5 | 1744872.88 | 2268083.25 | 2153403.5 | 2794677.5 | 2142315.5 | 1691018.62 | 1437019.75 | 2 | 8 | 1.266203081 | 1.152088022 |
| P52293 | Kpna2 | Importin subunit alpha-1 | 1192.77 | 48.77 | 57928 | 17 | 17248655 | 15881089 | 20675819 | 17715973 | 15642924 | 14031604 | 15633144 | 13114095 | 16035950 | 14784199 | 17153887 | 21234893 | 21426397 | 19724155 | 2 | 4 | 0.844369276 | 1.140650272 |
| P11438 | Lamp1 | Lysosome-associated membrane glycoprotein 1 | 568.22 | 28.33 | 43865 | 8 | 43631817 | 55507593 | 51127957 | 45247497 | 51616885 | 58910861 | 54424409 | 69183185 | 59832969 | 52330705 | 47542797 | 55563593 | 55145113 | 44633385 | 2 | 8 | 1.19240903 | 1.026198014 |
| Q6P791 | Lamtor1 | Ragulator complex protein LAMTOR1 | 96.67 | 16.15 | 17721 | 2 | 3387687.5 | 3668629.5 | 2649623.5 | 3601528.25 | 3677311 | 2439902.5 | 2771505 | 2523526.75 | 1127593.12 | 2774678 | 2861861 | 2781462.5 | 2352882.5 | 2578986.5 | 2 | 2 | 0.685154918 | 0.778284489 |
| Q6DVA0 | Lemd2 | LEM domain-containing protein 2 | 81.54 | 5.09 | 57507 | 2 | 1213773.5 | 1456737.88 | 1518883.25 | 1223020.12 | 1783008.38 | 2192399.5 | 1497458.5 | 1929877.62 | 1837782 | 1986906.12 | 1614966.12 | 1490715.5 | 2041762.12 | 2216279.25 | 2 | 3 | 1.312559883 | 1.279237311 |
| Q8BYI6 | Lpcat2 | Lysophosphatidylcholine acyltransferase 2 | 480.64 | 21.14 | 60254 | 6 | 2924251.5 | 3248657.25 | 5680712 | 3541806 | 3749135.5 | 5552612 | 6246263 | 6334870 | 6720229 | 5677482 | 4759444 | 5573457 | 7019060.5 | 6105467.5 | 2 | 2 | 1.594784754 | 1.531598679 |
| Q505F5 | Lrrc47 | Leucine-rich repeat-containing protein 47 | 188.49 | 11.88 | 63590 | 4 | 2665587 | 2880447.25 | 2403833.75 | 2991008 | 3062402.5 | 2181662.5 | 2430071.25 | 2210709.5 | 2539899 | 2393514 | 3145687.75 | 2741638.25 | 2095935.75 | 2574405.25 | 2 | 8 | 0.839507423 | 0.942428143 |
| P24527 | Lta4h | Leukotriene A-4 hydrolase | 476.73 | 19.15 | 69051 | 9 | 11273067 | 12678380 | 11299983 | 12328483 | 8893505 | 14530157 | 12769497 | 14811241 | 12085676 | 12902393 | 10949869 | 13702669 | 11657485 | 11071309 | 2 | 10 | 1.188151282 | 1.048752973 |
| P70202 | Lxn | Latexin | 92.6 | 9.46 | 25492 | 2 | 4335441.5 | 5021072 | 4927896 | 4757538.5 | 4461408.5 | 3813518 | 4037597.75 | 4750108.5 | 4314252.5 | 3536690 | 3864358.75 | 3692705.25 | 3778510 | 3973488.75 | 2 | 0 | 0.87018068 | 0.814195557 |
| O88188 | Ly86 | Lymphocyte antigen 86 | 309.35 | 43.83 | 17811 | 4 | 5974965.5 | 6362964.5 | 5429174.5 | 3877232.75 | 5239046 | 2333309.25 | 2539961.75 | 4649433.5 | 3992437.5 | 3548827.75 | 3586800.25 | 3616600.5 | 3760531 | 3984031.25 | 2 | 1 | 0.634740412 | 0.695037287 |
| P0C5H9 | Manf | Mesencephalic astrocyte-derived neurotrophic factor | 263.78 | 21.23 | 20388 | 5 | 13466765 | 12135204 | 10767937 | 10744876 | 10496057 | 8718691 | 8108153 | 10755421 | 7393491 | 9374199 | 11167409 | 8767509 | 10698030 | 11831972 | 2 | 7 | 0.769819634 | 0.921374362 |
| Q63932 | Map2k2 | Dual specificity mitogen-activated protein kinase kinase 2 | 544.34 | 33.92 | 44402 | 7 | 14525667 | 18130511 | 18250865 | 9743858 | 14086204 | 16234233 | 19362363 | 19185753 | 20138015 | 19338113 | 18610899 | 15415304 | 19699629 | 13266392 | 2 | 6 | 1.261200538 | 1.120464594 |
| Q9ERE7 | Mesd | LRP chaperone MESD | 345.63 | 37.5 | 25207 | 6 | 5726722.5 | 7060420 | 5772009 | 6588132.5 | 5376967 | 4327514.5 | 5422149 | 5000957 | 4327137.5 | 5600284.5 | 5297414 | 4399870.5 | 4885446 | 6070852 | 2 | 3 | 0.808473318 | 0.845785802 |
| Q9DCS2 | Mettl26 | Methyltransferase-like 26 | 98.55 | 8.82 | 22687 | 1 | 385797.88 | 276300.4525 | 205587.38 | 275702.25 | 238114.3 | 347897.56 | 403964.91 | 537628.25 | 494495.97 | 329711.44 | 417953.97 | 525149.44 | 383055.22 | 285989.78 | 2 | 2 | 1.52999976 | 1.458691431 |
| Q9JLB0 | Mpp6 | MAGUK p55 subfamily member 6 | 317.28 | 15.37 | 62631 | 6 | 5821362.5 | 6624546.5 | 6686558 | 4830265.5 | 5605394 | 6244530.5 | 6804290 | 7060152.5 | 6990546 | 7361588.5 | 7093408.5 | 4696316 | 7515800.5 | 6493294 | 2 | 7 | 1.165481604 | 1.090651575 |
| P26041 | Msn | Moesin | 3037.55 | 68.11 | 67767 | 37 | 691954433 | 676819137 | 608385089 | 702829697 | 649914689 | 713954753 | 729744513 | 735206401 | 692807489 | 702614401 | 662856705 | 622155137 | 571317505 | 619211713 | 2 | 4 | 1.073402892 | 0.929284211 |
| O35682 | Myadm | Myeloid-associated differentiation marker | 142.12 | 16.25 | 35285 | 3 | 1 | 400232.06 | 1 | 628709.5 | 532174.56 | 446582.5 | 806921.25 | 678009.9375 | 695979.19 | 762556.81 | 542196.5 | 858932.81 | 981764.5 | 794297.9367 | 2 | 1 | 2.171552328 | 2.544003322 |
| Q9CQ19 | Myl9 | Myosin regulatory light polypeptide 9 | 357.35 | 45.35 | 19854 | 1 | 26152073 | 28341347 | 28584151 | 26103673 | 24852313 | 31823803 | 27647341 | 35699409 | 33782497 | 32548851 | 29581851 | 29213619 | 30036973 | 26568097 | 2 | 2 | 1.204936321 | 1.07622806 |
| Q9QZ08 | Nagk | N-acetyl-D-glucosamine kinase | 217.37 | 14.29 | 37268 | 5 | 8062789 | 6713548 | 6884579 | 6825881.5 | 7194295 | 6607882 | 4962628.5 | 6840466 | 6593326 | 5349132 | 6609201.5 | 5562475 | 5993183.5 | 6841478.5 | 2 | 2 | 0.850686803 | 0.876036044 |
| O70145 | Ncf2 | Neutrophil cytosol factor 2 | 1100.57 | 50.48 | 59485 | 21 | 69790921 | 61107409 | 54740737 | 56351425 | 58903945 | 53135781 | 53183137 | 57343985 | 49652681 | 52358545 | 52031457 | 46979217 | 49328633 | 51797241 | 2 | 0 | 0.882947959 | 0.83142343 |
| P09405 | Ncl | Nucleolin | 2657.22 | 41.58 | 76723 | 43 | 512448801 | 493881793 | 545296833 | 505632737 | 517477665 | 546814785 | 557909633 | 523227073 | 571253825 | 519261889 | 481219489 | 532755297 | 556250113 | 481294433 | 2 | 9 | 1.055822917 | 0.995984576 |
| Q8R3N1 | Nop14 | Nucleolar protein 14 | 132.46 | 5.12 | 98769 | 3 | 1 | 399382.25 | 674298.75 | 271218.31 | 1 | 730307.19 | 599676.5 | 759479.06 | 674728.75 | 402867.47 | 745488.12 | 566358.75 | 588692 | 697021.81 | 2 | 2 | 2.354863473 | 2.414266999 |
| P11928 | Oas1a | 2'-5'-oligoadenylate synthase 1A | 477.04 | 27.79 | 42429 | 7 | 6047583.5 | 6469717 | 5685870 | 2889836.75 | 4769320.5 | 2935058 | 2408509.25 | 1142124 | 2262447.75 | 4146578.5 | 3218126.75 | 2340372.5 | 4985915 | 3930411.5 | 2 | 4 | 0.498590754 | 0.699609577 |
| P29758 | Oat | Ornithine aminotransferase, mitochondrial | 1084.71 | 50.11 | 48355 | 16 | 54754557 | 48940553 | 49442789 | 54602601 | 45663557 | 52761209 | 58228573 | 60343269 | 58608221 | 55362069 | 62513061 | 64798829 | 64568749 | 55856461 | 2 | 0 | 1.125883083 | 1.222045845 |
| P12382 | Pfkl | ATP-dependent 6-phosphofructokinase, liver type | 576.84 | 22.82 | 85360 | 10 | 19641895 | 16978521 | 16034875 | 13188097 | 19513529 | 20625585 | 20757869 | 18619079 | 20318077 | 19687081 | 18411935 | 18390033 | 17670297 | 18232371 | 2 | 8 | 1.171641321 | 1.06471506 |
| Q8K411 | Pitrm1 | Presequence protease, mitochondrial | 392.16 | 12.64 | 117372 | 8 | 7504507 | 9185966 | 9678748 | 8046197 | 8562303 | 9594543 | 9190575 | 9705606 | 10058189 | 9817600 | 10208941 | 8099308 | 10191635 | 9261356 | 2 | 4 | 1.125385709 | 1.098279502 |
| Q4FZT2 | Ppme1 | Protein phosphatase methylesterase 1 | 146.88 | 6.99 | 42316 | 3 | 6184713.5 | 4729126 | 4370266 | 5407718 | 6564517 | 5188333 | 4176921.5 | 3535666.75 | 4010244 | 4043384 | 6063492.5 | 5665088 | 3661940 | 5184402 | 2 | 8 | 0.768795402 | 0.943584232 |
| Q76MZ3 | Ppp2r1a | Serine/threonine-protein phosphatase 2A 65 kDa regulatory subunit A alpha isoform | 2107.18 | 64.35 | 65323 | 24 | 114746873 | 128048809 | 126684097 | 118607377 | 116083617 | 141708465 | 126688105 | 144928385 | 137329841 | 127348945 | 129637289 | 132702345 | 134147513 | 136290289 | 2 | 0 | 1.122205461 | 1.102290652 |
| Q9DBC7 | Prkar1a | cAMP-dependent protein kinase type I-alpha regulatory subunit | 631.42 | 34.91 | 43185 | 11 | 20935051 | 23816355 | 21509539 | 23857553 | 24367271 | 20280033 | 20458831 | 20114593 | 19862863 | 21942341 | 22525745 | 19538983 | 20724205 | 22213181 | 2 | 4 | 0.896693641 | 0.928086027 |
| Q91XU1 | Qki | Protein quaking | 307.42 | 29.62 | 37643 | 8 | 12454134 | 12073374 | 11243338 | 12401428 | 11836760 | 12375581 | 13536542 | 13072787 | 13495711 | 13113253 | 12004055 | 9655860 | 9752497 | 12054920 | 2 | 4 | 1.093066654 | 0.905433089 |
| Q9CZE3 | Rab32 | Ras-related protein Rab-32 | 80.73 | 10.76 | 25068 | 2 | 4747686 | 2531041 | 2179624.25 | 2626370.75 | 1742248.25 | 1194038.88 | 810264.69 | 2346547 | 1377834.738 | 1160488.38 | 1580524.647 | 680515.44 | 2482523.5 | 1578535 | 2 | 2 | 0.498241738 | 0.571536865 |
| P82343 | Renbp | N-acylglucosamine 2-epimerase | 372.96 | 21.16 | 49771 | 9 | 27482973 | 22888361 | 22671903 | 21481823 | 21501491 | 22207463 | 21423515 | 21410497 | 20568213 | 19921253 | 21980685 | 18911397 | 22334797 | 23548221 | 2 | 8 | 0.909541308 | 0.93486253 |
| E9Q555 | Rnf213 | E3 ubiquitin-protein ligase RNF213 | 2031.58 | 11.9 | 584784 | 43 | 45362837 | 41682497 | 32155981 | 35600493 | 38076313 | 29922537 | 28911147 | 32672189 | 31496833 | 33959937 | 33480975 | 34904065 | 35576017 | 33303313 | 2 | 4 | 0.81379185 | 0.889579708 |
| Q91VI7 | Rnh1 | Ribonuclease inhibitor | 2471.85 | 90.79 | 49816 | 29 | 331530049 | 365746049 | 323439425 | 375143137 | 344204865 | 375934497 | 387887553 | 393568545 | 360855105 | 377228801 | 383652609 | 331784801 | 336977025 | 385898241 | 2 | 6 | 1.089313392 | 1.033232879 |
| Q5U2U7 | Rnmt | mRNA cap guanine-N7 methyltransferase | 147.61 | 8.03 | 52817 | 3 | 3920096.75 | 3663730.25 | 3442175.5 | 3841588.75 | 4448384.5 | 4762323 | 4325686 | 4186189.5 | 4655614.5 | 5594118 | 2992147.25 | 4340487.5 | 4163963.5 | 4423773 | 2 | 8 | 1.217848443 | 1.030259321 |
| Q9JJ80 | Rpf2 | Ribosome production factor 2 homolog | 333.42 | 27.45 | 35364 | 6 | 1278550 | 1473422 | 1305544.5 | 2019469.38 | 2440933.5 | 895740.5775 | 1150534.88 | 196185.8 | 817088.38 | 1419153.25 | 767881.38 | 1024291.88 | 1101454.75 | 1302622.88 | 2 | 1 | 0.525797755 | 0.615797518 |
| P35979 | Rpl12 | 60S ribosomal protein L12 | 490.93 | 51.52 | 17805 | 7 | 42948813 | 36512005 | 40066069 | 40067593 | 38795725 | 37956381 | 35114249 | 35012577 | 33777105 | 38570625 | 40836957 | 34807781 | 32567541 | 36495265 | 2 | 4 | 0.909475027 | 0.911760891 |
| P67984 | Rpl22 | 60S ribosomal protein L22 | 387.03 | 56.25 | 14759 | 7 | 35659789 | 27045523 | 32735147 | 24856975 | 31397327 | 38167421 | 37770341 | 32239589 | 44082929 | 43481093 | 36101401 | 45934985 | 42751321 | 39497965 | 2 | 0 | 1.290363436 | 1.353752026 |

| | | | | | | | | | | | | | | | | | | | | | | | | |
|---|---|---|---|---|---|---|---|---|---|---|---|---|---|---|---|---|---|---|---|---|---|---|---|---|
| P97461 | Rps5 | 40S ribosomal protein S5 | 837 | 55.88 | 22889 | 13 | 65677265 | 68578081 | 71479009 | 69414161 | 76406825 | 65859193 | 63720657 | 44518953 | 60103913 | 67808585 | 69797105 | 62037685 | 56744709 | 69129137 | 2 | 5 | 0.859071861 | 0.916316032 |
| Q8BVY0 | Rsl1d1 | Ribosomal L1 domain-containing protein 1 | 603.76 | 22.79 | 50421 | 9 | 37531885 | 21401785 | 28499665 | 50338025 | 40785601 | 25286479 | 22207909 | 11306976 | 13842887 | 17548941 | 46628805 | 59814081 | 19116073 | 34891213 | 2 | 9 | 0.505122799 | 1.123242207 |
| Q6AYT3 | Rtcb | tRNA-splicing ligase RtcB homolog | 579.08 | 28.71 | 55249 | 12 | 15347944 | 16400637 | 20071873 | 16082753 | 16561951 | 17855515 | 20343665 | 19262173 | 22609541 | 20570229 | 13212056 | 15771848 | 19672145 | 17958699 | 2 | 9 | 1.191510504 | 0.985831756 |
| Q3T1J8 | Rtfdc1 | Protein RTF2 homolog 2 | 40.84 | 3.27 | 33812 | 1 | 663241.5 | 444095.25 | 635196.69 | 287354.75 | 386546.75 | 696939.62 | 662056.62 | 648334.69 | 797921.56 | 732653.44 | 732385.5 | 616243.31 | 723199.69 | 583197.5 | 2 | 4 | 1.464101463 | 1.373421003 |
| Q9WTM5 | Ruvbl2 | RuvB-like 2 | 1156.47 | 54.64 | 51113 | 21 | 159727153 | 116194353 | 119336737 | 173795409 | 156698625 | 120122153 | 92422225 | 79221297 | 78944489 | 94008201 | 158645713 | 185310657 | 94165385 | 130346329 | 2 | 10 | 0.640326431 | 0.979101448 |
| D3YXK2 | Safb | Scaffold attachment factor B1 | 378.03 | 15.58 | 105104 | 4 | 18522413 | 19448347 | 18781235 | 17714511 | 19606143 | 20507755 | 19990853 | 22320199 | 20586181 | 18884721 | 18167147 | 16904299 | 18881665 | 18297649 | 2 | 5 | 1.087348024 | 0.960039405 |
| Q8BGH2 | Samm50 | Sorting and assembly machinery component 50 homolog | 769.28 | 41.58 | 51864 | 15 | 21734131 | 19847863 | 20459641 | 19857669 | 19958101 | 22509353 | 21406595 | 21761089 | 22735655 | 21234795 | 18830151 | 22308693 | 22266007 | 22129885 | 2 | 5 | 1.076480272 | 1.049687256 |
| Q9CXV1 | Sdhd | Succinate dehydrogenase [ubiquinone] cytochrome b small subunit, mitochondrial | 82.47 | 16.35 | 17014 | 1 | 2303064.75 | 1947185.12 | 1287627.62 | 1616036.25 | 1565868.62 | 952996.25 | 1037701.69 | 1567015.25 | 819913.06 | 910888.75 | 1212176.75 | 1040106.31 | 881603.31 | 1251570.62 | 2 | 0 | 0.606496215 | 0.628664915 |
| P32261 | Serpinc1 | Antithrombin-III | 315.63 | 15.05 | 52004 | 6 | 29093393 | 14399019 | 18221479 | 21083791 | 18452405 | 15196599 | 14321270 | 14468564 | 13993722 | 13181828 | 12153597 | 10939648 | 18360459 | 8842693 | 2 | 2 | 0.702833796 | 0.620942639 |
| Q8K2Q9 | Shtn1 | Shootin-1 | 453.89 | 12.84 | 71343 | 9 | 18959481 | 21447931 | 19827315 | 19525633 | 19291207 | 19012555 | 17929105 | 19085505 | 18397015 | 18127753 | 18088917 | 18162987 | 16660083 | 20857917 | 2 | 4 | 0.934381311 | 0.930953268 |
| Q8CI08 | Slain2 | SLAIN motif-containing protein 2 | 48.31 | 2.07 | 62378 | 1 | 1065192.75 | 797507.75 | 527056.31 | 1317345.38 | 1272489.5 | 909175.75 | 1 | 1 | 1 | 1 | 1251404.38 | 1955571.25 | 744044.69 | 843040.62 | 2 | 9 | 0.182581185 | 1.203427218 |
| Q8BH59 | Slc25a12 | Calcium-binding mitochondrial carrier protein Aralar1 | 1161.56 | 43.72 | 74570 | 17 | 36904181 | 39552901 | 41379253 | 35412145 | 39784721 | 43641701 | 40349677 | 48736293 | 41627165 | 40790373 | 45488373 | 42274969 | 54611137 | 42473157 | 2 | 0 | 1.114550284 | 1.196993801 |
| Q78IQ7 | Slc39a4 | Zinc transporter ZIP4 | 201.38 | 12.73 | 71063 | 4 | 2840990.5 | 2027986.25 | 4243827 | 1403588.5 | 2391140 | 1991673.62 | 979170.56 | 1433551.88 | 748370.44 | 753986.94 | 2292126.75 | 2163706.25 | 4911040 | 2856759.5 | 2 | 7 | 0.457620661 | 1.183769316 |
| Q8BVL3 | Snx17 | Sorting nexin-17 | 151.8 | 13.62 | 52797 | 4 | 1532372 | 157143.75 | 918231.44 | 1128782.25 | 934132.36 | 1 | 692742.06 | 620969.5 | 1 | 1 | 1 | 619672.94 | 1 | 1 | 2 | 1 | 0.281269468 | 0.165842649 |
| Q9D7A6 | Srp19 | Signal recognition particle 19 kDa protein | 108.91 | 25.69 | 16191 | 2 | 1186770.88 | 1212089.62 | 1358957.625 | 1583497.5 | 1453472.5 | 754597.88 | 1050707 | 973517.75 | 1079149.5 | 1318759.75 | 1367959 | 710563.19 | 1729218 | 1416048.62 | 2 | 9 | 0.761868035 | 0.96099185 |
| Q52KI8 | Srrm1 | Serine/arginine repetitive matrix protein 1 | 417.81 | 10.25 | 106862 | 7 | 4782300 | 5934505 | 7111439 | 6608492 | 6399699 | 8489364 | 8690769 | 6491351 | 7436672 | 9134941 | 6004613 | 5607186 | 6368805.5 | 6769152.5 | 2 | 9 | 1.305050243 | 1.003267604 |
| Q99KH8 | Stk24 | Serine/threonine-protein kinase 24 | 404.54 | 25.99 | 47954 | 8 | 3952477.25 | 5645742.5 | 5255120 | 3935718 | 3924994.5 | 8385537 | 8650342 | 5015629 | 8625960 | 7986652 | 5721885.5 | 5286080.5 | 5779808.5 | 7148376.5 | 2 | 1 | 1.702211458 | 1.317254553 |
| Q08849 | Stx3 | Syntaxin-3 | 65.64 | 4.84 | 33257 | 1 | 1304755.5 | 1256390.12 | 1570297.62 | 1050472.62 | 1043461.81 | 1733287.62 | 1350518.25 | 2013118.88 | 1849019 | 1344744.5 | 1178522.75 | 1319422.38 | 1270346.5 | 1107202.25 | 2 | 9 | 1.331756672 | 0.978955442 |
| Q5DQR4 | Stxbp5l | Syntaxin-binding protein 5-like | 36.53 | 0.93 | 131844 | 1 | 5538802.5 | 6822379.5 | 4978563.5 | 5871094 | 7652958.5 | 9780742 | 9647783 | 6495141.5 | 8665200 | 10572685 | 12446183 | 6568418.5 | 11742469 | 10057399 | 2 | 2 | 1.46325321 | 1.653007413 |
| Q9Z2I9 | Sucla2 | Succinate--CoA ligase [ADP-forming] subunit beta, mitochondrial | 456.06 | 29.37 | 50114 | 9 | 8295629.5 | 10424399 | 10797735 | 8882850 | 9556912 | 11354631 | 9859279 | 13265904 | 12864112 | 11100224 | 12045576 | 12539160 | 12071474 | 10814796 | 2 | 0 | 1.218664837 | 1.237319 |
| Q80X82 | Sympk | Symplekin | 607.63 | 13.55 | 142284 | 12 | 6440025.5 | 5867272.5 | 7302222 | 5437608 | 5401326.5 | 6865884.5 | 7468577 | 9655819 | 7936289.5 | 6719849.5 | 6671930.5 | 7268310 | 8251929.5 | 6601514 | 2 | 3 | 1.269240759 | 1.182066729 |
| Q5EAN7 | Terf2ip | Telomeric repeat-binding factor 2-interacting protein 1 | 69.76 | 3.56 | 43418 | 1 | 303358.16 | 1 | 368227.56 | 1 | 1 | 425166.81 | 371567.25 | 439172.94 | 368551.03 | 283467.84 | 346011.94 | 387326.56 | 341573.62 | 447683.03 | 2 | 2 | 2.811133978 | 2.833942681 |
| P47226 | Tes | Testin | 353.71 | 21.04 | 47983 | 8 | 3696221.75 | 4564285.5 | 4730642 | 6025362.5 | 4329370.5 | 7849374 | 5757800 | 5079002 | 6395534 | 6599892.5 | 6140136 | 5024831.5 | 5679737.5 | 6283765.5 | 2 | 2 | 1.357053127 | 1.238359203 |
| Q62351 | Tfrc | Transferrin receptor protein 1 | 939.45 | 30.54 | 85731 | 20 | 11530963 | 11633155 | 10900484 | 7395272.5 | 10115463 | 7675129.5 | 5969507 | 7726780.5 | 7213647.5 | 5669373 | 9960069 | 11926839 | 10309869 | 10780530 | 2 | 10 | 0.664163128 | 1.041614779 |
| P52912 | Tia1 | Nucleolysin TIA-1 | 51.35 | 4.66 | 42800 | 1 | 329928.09 | 292098.72 | 249242.75 | 298583 | 292463.14 | 222929.34 | 282399.94 | 241197.3 | 275345.5 | 184114.42 | 279784.69 | 291728.4267 | 323873.34 | 271527.25 | 2 | 7 | 0.824710081 | 0.997487843 |
| A1L1L2 | Tmem214 | Transmembrane protein 214 | 58.63 | 1.9 | 76566 | 1 | 145575.83 | 1 | 166379.84 | 1 | 1 | 228695.64 | 229969.2475 | 338946 | 230705.56 | 121529.79 | 134574.75 | 194711.33 | 195662.78 | 184779.02 | 2 | 2 | 3.68589287 | 2.843837775 |
| Q9CZW5 | Tomm70 | Mitochondrial import receptor subunit TOM70 | 521.79 | 17.18 | 67590 | 10 | 26322325 | 23164519 | 18914515 | 23519605 | 21850621 | 20034171 | 18624847 | 19168111 | 18571787 | 18758403 | 22865805 | 26596017 | 20172785 | 22452845 | 2 | 10 | 0.836389148 | 1.011758033 |
| O89023 | Tpp1 | Tripeptidyl-peptidase 1 | 494.53 | 28.29 | 61342 | 3 | 33809873 | 47168833 | 35362789 | 36958925 | 34351465 | 47698097 | 42514225 | 48979121 | 47771941 | 40657149 | 43374429 | 44452413 | 46082881 | 38542913 | 2 | 4 | 1.212993587 | 1.148753688 |
| Q61510 | Trim25 | E3 ubiquitin/ISG15 ligase TRIM25 | 42.96 | 1.89 | 71726 | 1 | 1870693.75 | 1544405.88 | 1901792.25 | 1853437.5 | 1749071.25 | 1385453 | 1568614 | 1712436.75 | 1528736.5 | 1536857.12 | 1493609.25 | 2162860.25 | 1389090.5 | 1927899.62 | 2 | 10 | 0.866885309 | 0.977288148 |
| Q3TX08 | Trmt1 | tRNA (guanine(26)-N(2))-dimethyltransferase | 105.58 | 4.52 | 72350 | 2 | 1127776.625 | 1147970.75 | 996056.25 | 1157810.12 | 1209269.38 | 940761.12 | 1074495.5 | 1073776.88 | 995710.81 | 915424.12 | 1163824.5 | 968688.38 | 1016140.81 | 1149427.5 | 2 | 8 | 0.886730283 | 0.952777592 |
| Q8K1J6 | Trnt1 | CCA tRNA nucleotidyltransferase 1, mitochondrial | 380.86 | 26.04 | 49895 | 9 | 10779536 | 9099076 | 9943824 | 8867748 | 11954073 | 9274870 | 7692941.5 | 8855822 | 7117278.5 | 8567276 | 10541735 | 7056790.5 | 11693053 | 9402080 | 2 | 9 | 0.81960306 | 0.955035694 |
| P26369 | U2af2 | Splicing factor U2AF 65 kDa subunit | 826.54 | 42.74 | 53517 | 12 | 25919089 | 30357559 | 35058081 | 34916045 | 29659301 | 35371321 | 32002923 | 39485465 | 38426841 | 37827961 | 32380563 | 36104461 | 36808109 | 33075245 | 2 | 4 | 1.174487993 | 1.10936046 |
| Q9EPU0 | Upf1 | Regulator of nonsense transcripts 1 | 769.85 | 16.64 | 123967 | 15 | 4513688.5 | 4762802 | 4437975 | 5356068 | 5269039.5 | 6113806.5 | 5674726 | 4905793.5 | 6596554.5 | 6618581.5 | 5310971 | 8054423.5 | 5380112.5 | 6454206 | 2 | 1 | 1.228840868 | 1.294173947 |
| Q80WQ2 | Vac14 | Protein VAC14 homolog | 768.49 | 25.96 | 88048 | 14 | 13528564 | 10920689 | 10440508 | 9211076 | 10567934 | 14573422 | 11720757 | 14294620 | 14138150 | 11784666 | 13081669 | 12649819 | 13982791 | 14690806 | 2 | 2 | 1.216629051 | 1.243970827 |
| Q9Z2L0 | Vdac1 | Voltage-dependent anion-selective channel protein 1 | 1323.92 | 78.8 | 30756 | 17 | 90162177 | 100447313 | 85169633 | 88416833 | 102944129 | 107703945 | 104512193 | 100083985 | 107296049 | 104988993 | 92626481 | 104174233 | 104296585 | 106404737 | 2 | 2 | 1.122971849 | 1.090417118 |
| Q60930 | Vdac2 | Voltage-dependent anion-selective channel protein 2 | 1007.83 | 56.61 | 31733 | 13 | 155728449 | 168903905 | 177213537 | 146716865 | 154621441 | 175781217 | 180012129 | 184517217 | 184701425 | 170132913 | 153915905 | 164444097 | 179458209 | 176108481 | 2 | 7 | 1.114495161 | 1.048835831 |
| Q8R5L3 | Vps39 | Vam6/Vps39-like protein | 146.49 | 4.29 | 101693 | 3 | 1434395.62 | 1657410.75 | 991179.44 | 1260462.12 | 1247210.38 | 2273649.5 | 1657768.5 | 1535092 | 1735517.25 | 1650531.5 | 1185969.38 | 1014356.69 | 1971271.25 | 1928844 | 2 | 8 | 1.343197953 | 1.157024275 |
| Q65Z40 | Wapl | Wings apart-like protein homolog | 286.43 | 7.58 | 134070 | 5 | 1373054.38 | 1563977.25 | 1570771 | 1787935.88 | 1526152.38 | 1543065.62 | 873228.12 | 1357312.62 | 1250861.5 | 1030300.38 | 1788077.38 | 826465 | 2008068.25 | 1905443.88 | 2 | 5 | 0.774079864 | 1.043234718 |
| Q8BH43 | Wasf2 | Wiskott-Aldrich syndrome protein family member 2 | 520.27 | 21.33 | 54074 | 9 | 7733303.5 | 8904244 | 9569390 | 7331849 | 11301141 | 11083330 | 9736062 | 11897482 | 11779608 | 11686500 | 8882168 | 10968333 | 12126567 | 12147282 | 2 | 4 | 1.252967726 | 1.230051888 |
| O88286 | Wiz | Protein Wiz | 43.91 | 0.89 | 184291 | 1 | 1078049.38 | 1476510.38 | 1311159 | 844011.31 | 1269574.75 | 508044.5 | 527435.56 | 1186802.25 | 1 | 1 | 686847.94 | 945573.25 | 918914.31 | 1230957.5 | 2 | 4 | 0.371662656 | 0.790705005 |
| P97789 | Xrn1 | 5'-3' exoribonuclease 1 | 63.96 | 0.99 | 194307 | 1 | 1 | 1 | 240862.66 | 1 | 207752.36 | 329733.06 | 246815.38 | 338300.59 | 264674.41 | 200326.83 | 242354.92 | 216120.2 | 223111.81 | 210425.17 | 2 | 3 | 3.075779858 | 2.485444354 |
| P62960 | Ybx1 | Nuclease-sensitive element-binding protein 1 | 806.76 | 49.07 | 35730 | 7 | 12411681 | 8943158 | 10193305 | 12765116 | 12786760 | 7080497 | 7872198.5 | 5933162.5 | 8493797 | 11040674 | 10806845 | 9333197 | 10449989 | 8061960.5 | 2 | 5 | 0.707886425 | 0.846146628 |
| Q9Z2U2 | Zfp292 | Zinc finger protein 292 | 34.66 | 0.41 | 301048 | 1 | 923951.88 | 2765312 | 2547395 | 1599058 | 1505235.12 | 1436604.5 | 1 | 1 | 954173.75 | 1 | 983321.62 | 1409616.75 | 1261138.62 | 1218025.663 | 2 | 4 | 0.255946209 | 0.651981545 |
| O35986 | Zranb2 | Zinc finger Ran-binding domain-containing protein 2 | 55.42 | 5.45 | 37350 | 2 | 2275240 | 1912361.75 | 2048002.38 | 2818413.75 | 396779.72 | 2465725.25 | 2891881.25 | 2404159.75 | 2859773.25 | 3414188.75 | 2771167.75 | 3795330.5 | 3466864.25 | 3270741.75 | 2 | 1 | 1.4851369 | 1.759653631 |
