## Supplemental Table 5 for "Effects of polystyrene and polylactide nanoparticles on macrophages under a repeated exposure mode"

Supplementary Table 5: Results of the pathway analysis by the David tool on the proteins modulated in response to PS particles

Annotation Cluster 1 — Enrichment Score: 16.08622370427957

| Category | Term | Count | % | PValue | Genes | Fold Enrichment | FDR |
|---|---|---|---|---|---|---|---|
| GOTERM_CC_DIRECT | GO:0005739~mitochondrion | 79 | 21.47 | 4.98691978E-27 | P10639, P07901, O70579, P29452, Q99JY0, Q8K3J1, P48771, O08749, P18242, Q8BVI4, Q9WTP6, P36552, O09044, Q9EPL9, Q8BH59, Q8JZU0, Q9CXV1, Q9D0L7, Q791T5, Q3TRM8, Q8BH86, P11499, Q9Z2I8, P19783, P19536, Q07813, Q9Z2I9, Q9CYG7, Q99KI0, Q91YY4, Q922Q1, Q8BMS1, P54071, Q8QZT1, Q9D0K2, Q921F2, Q811U4, Q91VR5, Q62425, Q60766, Q9D0G0, Q91VJ1, Q68FD5, Q9DBF1, Q9QXX4, Q9CPQ3, Q64521, P31750, Q9CPQ1, P47738, Q9CZD3, Q8BTX9, P00405, Q9WUD1, Q7TMY8, Q9DC70, Q9JM90, Q8CGK3, Q9QYB1, P97494, P97450, Q791V5, P59017, Q8VCF0, P11352, Q8BVN4, Q99JR1, P85094, Q9CQA3, O09111, O88986, P21958, P56394, Q8BHG1, Q9JK81, P29758, P60603, Q9JIY5, P56391 | 4.09 | 1.07966813E-24 |
| UP_KW_CELLULAR_COMPONENT | KW-0496~Mitochondrion | 68 | 18.48 | 4.87573117E-18 | P07901, Q99JY0, Q8K3J1, P48771, O08749, Q9WTP6, P36552, Q8BH59, Q8JZU0, Q9CXV1, Q9D0L7, Q791T5, Q8BH86, P35564, Q9Z2I8, P19783, P19536, Q07813, Q9Z2I9, Q9CYG7, Q99KI0, Q91YY4, Q922Q1, Q8BMS1, P54071, Q8QZT1, Q9D0K2, Q921F2, Q811U4, Q91VR5, Q62425, Q60766, Q9D0G0, Q9DBF1, Q9QXX4, Q9CPQ3, Q64521, P31750, Q9CPQ1, Q62465, P47738, Q9CZD3, Q8BTX9, P00405, Q9WUD1, Q7TMY8, Q9DC70, Q9JM90, Q8CGK3, Q9QYB1, P62908, P97450, Q791V5, P59017, Q8VCF0, P11352, Q8BVN4, Q99JR1, Q9CQA3, O09111, O88986, P56394, Q8BHG1, Q9JK81, P29758, P60603, Q9JIY5, P56391 | 3.25 | 1.56023397E-16 |
| GOTERM_CC_DIRECT | GO:0005743~mitochondrial inner membrane | 32 | 8.70 | 3.38398856E-14 | Q811U4, Q62425, Q8K3J1, Q99JY0, Q9D0G0, P48771, Q9QXX4, Q9CPQ3, Q64521, Q9CPQ1, Q9WTP6, P36552, P00405, Q9DC70, Q8BH59, P62908, P97450, Q9CXV1, Q791V5, P19783, Q99JR1, P19536, Q9CQA3, O09111, O88986, Q91YY4, Q922Q1, P60603, Q8BMS1, P56391, P54071, Q8QZT1 | 5.54 | 3.66316761E-12 |
| UP_KW_DOMAIN | KW-0809~Transit peptide | 33 | 8.97 | 9.52181524E-13 | Q9D0K2, Q8K3J1, Q99JY0, P48771, O08749, Q9DBF1, Q64521, P47738, P36552, Q9CZD3, Q9DC70, Q8CGK3, P97450, Q8JZU0, Q9CXV1, Q8BH86, Q9Z2I8, P19783, Q8BVN4, P19536, Q9CQA3, O09111, Q9Z2I9, O88986, Q99KI0, Q91YY4, Q922Q1, P29758, Q9JK81, Q8BMS1, Q9JIY5, P54071, Q8QZT1 | 4.57 | 1.9995812E-11 |
| UP_SEQ_FEATURE | TRANSIT:Mitochondrion | 33 | 8.97 | 4.72999529E-12 | Q9D0K2, Q8K3J1, Q99JY0, P48771, O08749, Q9DBF1, Q64521, P47738, P36552, Q9CZD3, Q9DC70, Q8CGK3, P97450, Q8JZU0, Q9CXV1, Q8BH86, Q9Z2I8, P19783, Q8BVN4, P19536, Q9CQA3, O09111, Q9Z2I9, O88986, Q99KI0, Q91YY4, Q922Q1, P29758, Q9JK81, Q8BMS1, Q9JIY5, P54071, Q8QZT1 | 4.42 | 5.21245481E-09 |

Annotation Cluster 2 — Enrichment Score: 5.858365434023681

| Category | Term | Count | % | PValue | Genes | Fold Enrichment | FDR |
|---|---|---|---|---|---|---|---|
| GOTERM_CC_DIRECT | GO:0005743~mitochondrial inner membrane | 32 | 8.70 | 3.38398856E-14 | Q811U4, Q62425, Q8K3J1, Q99JY0, Q9D0G0, P48771, Q9QXX4, Q9CPQ3, Q64521, Q9CPQ1, Q9WTP6, P36552, P00405, Q9DC70, Q8BH59, P62908, P97450, Q9CXV1, Q791V5, P19783, Q99JR1, P19536, Q9CQA3, O09111, O88986, Q91YY4, Q922Q1, P60603, Q8BMS1, P56391, P54071, Q8QZT1 | 5.54 | 3.66316761E-12 |
| GOTERM_CC_DIRECT | GO:0031966~mitochondrial membrane | 16 | 4.35 | 1.47302523E-08 | Q62425, Q60766, P48771, P35564, P18760, P19783, Q8VCF0, P59017, Q9CPQ1, Q99JR1, P19536, Q9CQA3, Q07813, P00405, Q9JIY5, P56391 | 6.87 | 8.08387613E-07 |
| KEGG_PATHWAY | mmu05415:Diabetic cardiomyopathy | 21 | 5.71 | 2.86272939E-08 | P97450, P51432, Q62425, Q9CXV1, Q6PHZ2, Q64518, Q8BTI9, O70145, Q8K3J1, P48771, P18242, P19783, P31750, Q9CPQ1, P19536, Q9CQA3, O09111, Q00612, P00405, Q9DC70, P56391 | 4.56 | 6.75610177E-06 |
| UP_KW_CELLULAR_COMPONENT | KW-0999~Mitochondrion inner membrane | 22 | 5.98 | 1.59003837E-07 | P62908, P97450, Q62425, Q9CXV1, Q99JY0, Q8K3J1, P48771, Q9QXX4, P19783, Q64521, Q9CPQ1, Q99JR1, P19536, Q9CQA3, O09111, P00405, Q91YY4, Q9DC70, P60603, Q8BMS1, Q8BH59, P56391 | 4.01 | 1.69604093E-06 |
| KEGG_PATHWAY | mmu05012:Parkinson disease | 22 | 5.98 | 3.02838909E-07 | P97450, P10639, Q811U4, Q62425, Q9CXV1, Q6PHZ2, Q8K3J1, P48771, Q8VDM4, Q9R1P1, Q9R1P4, P19783, Q9CPQ1, P70227, P19536, Q9CQA3, O09111, Q07813, P00405, Q9DC70, Q9JIY5, P56391 | 3.78 | 1.9154561E-05 |
| GOTERM_CC_DIRECT | GO:0045277~respiratory chain complex IV | 7 | 1.90 | 3.31587228E-07 | P19536, Q62425, P48771, P00405, P19783, P56391, Q9CPQ1 | 24.39 | 1.30524791E-05 |
| KEGG_PATHWAY | mmu00190:Oxidative phosphorylation | 15 | 4.08 | 1.5318532E-06 | P97450, Q62425, Q9CXV1, Q8BVE3, Q8K3J1, P48771, P19783, Q9CPQ1, P19536, Q9CQA3, O09111, P00405, P56394, Q9DC70, P56391 | 5.01 | 7.7511772E-05 |
| KEGG_PATHWAY | mmu05020:Prion disease | 20 | 5.43 | 4.80951156E-06 | P97450, Q62425, Q9CXV1, Q8BTI9, O70145, Q8K3J1, P48771, Q8VDM4, Q9R1P1, Q9R1P4, P19783, Q9CPQ1, P70227, P19536, Q9CQA3, O09111, Q07813, P00405, Q9DC70, P56391 | 3.45 | 0.000202801071 |
| KEGG_PATHWAY | mmu05016:Huntington disease | 21 | 5.71 | 7.63040351E-06 | P97450, P51432, Q62425, Q9CXV1, Q8K3J1, P48771, Q68FD5, Q8VDM4, Q9R1P1, Q9R1P4, P19783, P11352, Q9CPQ1, P19536, Q9CQA3, O09111, Q07813, O08585, P00405, Q9DC70, P56391 | 3.21 | 0.000250593273 |
| KEGG_PATHWAY | mmu04932:Non-alcoholic fatty liver disease | 15 | 4.08 | 7.92389797E-06 | Q62425, Q9CXV1, Q8BTI9, Q8K3J1, P48771, P19783, P31750, Q9CPQ1, P19536, Q9CQA3, O09111, Q07813, P00405, Q9DC70, P56391 | 4.36 | 0.000250593273 |
| KEGG_PATHWAY | mmu05208:Chemical carcinogenesis - reactive oxygen species | 17 | 4.62 | 2.65080731E-05 | P97450, Q62425, Q9CXV1, Q8BTI9, O70145, Q8K3J1, P48771, P19783, P31750, Q9CPQ1, P19536, Q9CQA3, O09111, P00405, P48774, Q9DC70, P56391 | 3.50 | 0.000560856683 |
| KEGG_PATHWAY | mmu05022:Pathways of neurodegeneration - multiple diseases | 26 | 7.07 | 2.6601898E-05 | Q921F2, Q811U4, Q62425, Q6PHZ2, Q8K3J1, P48771, Q8VDM4, Q9R1P1, Q9R1P4, Q9CPQ1, Q9QY76, P70227, P00405, Q9DC70, P97450, P51432, Q9CXV1, Q64518, P19783, P11352, P19536, Q9CQA3, O09111, Q07813, Q9JIY5, P56391 | 2.55 | 0.000560856683 |
| KEGG_PATHWAY | mmu05014:Amyotrophic lateral sclerosis | 22 | 5.98 | 4.20172022E-05 | Q921F2, P97450, Q62425, Q9CXV1, P29452, Q8K3J1, P48771, Q8VDM4, Q9R1P1, Q9R1P4, P19783, P11352, Q9CPQ1, Q9QY76, P70227, P19536, Q9CQA3, O09111, Q07813, P00405, Q9DC70, P56391 | 2.76 | 0.000817719396 |

| | | | | | | | |
|---|---|---|---|---|---|---|---|
| KEGG_PATHWAY | mmu05010:Alzheimer disease | 22 | 5.98 | 7.14234601E-05 | P97450, P51432, P57716, Q62425, Q9CXV1, Q64518, Q8BTI9, Q8K3J1, P48771, Q8VDM4, Q9R1P1, Q9R1P4, P19783, P31750, Q9CPQ1, P70227, P19536, Q9CQA3, O09111, P00405, Q9DC70, P56391 | 2.66 | 0.001204675694 |
| KEGG_PATHWAY | mmu04260:Cardiac muscle contraction | 9 | 2.45 | 0.000562523275 | P19536, P21107, Q64518, P48771, P00405, P58774, P19783, P56391, Q9CPQ1 | 4.76 | 0.007906577145 |
| KEGG_PATHWAY | mmu04714:Thermogenesis | 14 | 3.80 | 0.001527611149 | P97450, Q62425, Q9CXV1, Q8K3J1, P48771, P19783, Q9CPQ1, P19536, Q9CQA3, O09111, P00405, P56394, Q9DC70, P56391 | 2.78 | 0.017567528212 |
| Annotation Cluster 3 | Enrichment Score: 4.56527399686028 | | | | | | |
| Category | Term | Count | % | PValue | Genes | Fold Enrichment | FDR |
| GOTERM_CC_DIRECT | GO:0005783~endoplasmic reticulum | 39 | 10.60 | 7.97298866E-09 | Q9CQS8, P27773, Q9CQW9, Q60766, B9EKI3, Q6GV12, Q99JY0, Q8BMA6, Q99K23, Q91XB0, Q9QY76, P70227, O08795, Q8VCH8, P36371, Q9WVJ3, P58681, P63242, Q9WUD1, Q8K1T1, P62908, P57716, Q64518, Q9D0L7, Q62393, P35564, Q6PA06, Q8BVF2, Q9D8V0, Q6P5E4, Q9R0E1, Q07813, P21958, Q9DCQ7, P08113, Q80UJ7, Q8QZT1, Q80WJ7, Q8BG07 | 2.91 | 5.75384015E-07 |
| GOTERM_CC_DIRECT | GO:0005789~endoplasmic reticulum membrane | 26 | 7.07 | 0.000360878917 | Q9CQS8, P61804, Q80X80, Q60766, Q6GV12, Q91XB0, Q9QY76, Q3U2P1, P70227, Q8VCH8, P36371, P58681, P40142, P63242, P61027, P57716, Q9D0L7, P35564, Q9D8V0, Q9R0E1, Q07813, P21958, P08113, Q9EPE9, Q80WJ7, Q8BG07 | 2.20 | 0.00744097958 |
| UP_KW_CELLULAR_COMPONENT | KW-0256~Endoplasmic reticulum | 34 | 9.24 | 0.007001568661 | Q9CQS8, P27773, P61804, Q80X80, Q6GV12, Q99JY0, Q8BMA6, Q99K23, Q91XB0, Q9QY76, Q3U2P1, P70227, O08795, Q8VCH8, P36371, Q9WVJ3, P58681, P63242, P61027, Q64518, Q9D0L7, P35564, Q6PA06, Q8BVF2, Q9D8V0, Q6P5E4, Q9R0E1, Q8BXQ2, P21958, P08113, Q9EPE9, Q80UJ7, Q80WJ7, Q8BG07 | 1.61 | 0.032007171022 |
| Annotation Cluster 4 | Enrichment Score: 4.480100896619738 | | | | | | |
| Category | Term | Count | % | PValue | Genes | Fold Enrichment | FDR |
| KEGG_PATHWAY | mmu01200:Carbon metabolism | 15 | 4.08 | 2.57814736E-07 | Q9CXV1, Q3TRM8, Q9DCD0, O08749, Q9Z2I8, Q9CQA3, P40142, Q00612, Q9Z2I9, Q99KI0, P17182, Q9EPL9, P54071, Q8QZT1 | 5.79 | 1.9154561E-05 |
| GOTERM_BP_DIRECT | GO:0006099~tricarboxylic acid cycle | 6 | 1.63 | 2.37716773E-05 | Q9CQA3, Q9CXV1, Q9Z2I9, Q99KI0, Q9Z2I8, P54071 | 17.15 | 0.005390227838 |
| KEGG_PATHWAY | mmu00020:Citrate cycle (TCA cycle) | 7 | 1.90 | 4.71929063E-05 | Q9CQA3, Q9CXV1, Q9Z2I9, Q99KI0, O08749, Q9Z2I8, P54071 | 10.31 | 0.000852843235 |
| UP_KW_BIOLOGICAL_PROCESS | KW-0816~Tricarboxylic acid cycle | 6 | 1.63 | 6.09027378E-05 | Q9CQA3, Q9CXV1, Q9Z2I9, Q99KI0, Q9Z2I8, P54071 | 13.78 | 0.002679720464 |
| GOTERM_BP_DIRECT | GO:0006105~succinate metabolic process | 3 | 0.82 | 0.002257430633 | Q9CQA3, Q9Z2I9, Q9Z2I8 | 40.42 | 0.116999404814 |
| Annotation Cluster 5 | Enrichment Score: 4.326622223231248 | | | | | | |
| Category | Term | Count | % | PValue | Genes | Fold Enrichment | FDR |
| GOTERM_BP_DIRECT | GO:0045087~innate immune response | 21 | 5.71 | 6.08516202E-07 | Q8VI94, Q8R4K2, Q91VR5, Q60766, P26151, P10810, Q8BVK9, Q99J93, Q6R5N8, Q91VJ1, Q9R002, Q8VCF0, P97814, Q62167, Q91XB0, P0DOV2, O88839, P58681, O88188, O35309, Q8BG07 | 3.89 | 0.000275962098 |
| UP_KW_BIOLOGICAL_PROCESS | KW-0399~Innate immunity | 22 | 5.98 | 1.28155643E-06 | Q8VI94, Q8R4K2, Q91VR5, Q9CQW9, Q60766, P26151, P10810, Q61107, Q8BVK9, Q99J93, Q6R5N8, Q91VJ1, Q9R002, Q9Z0E6, Q8VCF0, P97814, Q62167, P0DOV2, P58681, O88188, O35309, Q8BG07 | 3.47 | 0.000112776966 |
| UP_KW_BIOLOGICAL_PROCESS | KW-0395~Inflammatory response | 11 | 2.99 | 0.001434436882 | P58681, P10810, Q6R5N8, Q91VJ1, Q9R002, P70202, Q91YX0, O88188, P97814, Q62167, Q8BG07 | 3.41 | 0.042076815202 |
| UP_KW_BIOLOGICAL_PROCESS | KW-0391~Immunity | 28 | 7.61 | 0.004413872506 | P27773, Q8R4K2, Q91VR5, Q9CQW9, Q60766, P10810, Q6R5N8, Q8BVK9, Q91VJ1, Q9R1P4, Q62167, P0DOV2, P36371, P58681, Q91YX0, O35309, Q8VI94, P26151, Q61107, Q99J93, Q9R002, Q9Z0E6, Q8VCF0, P97814, P70698, P21958, O88188, Q8BG07 | 1.75 | 0.077684156097 |
| Annotation Cluster 6 | Enrichment Score: 3.69561232025453 | | | | | | |
| Category | Term | Count | % | PValue | Genes | Fold Enrichment | FDR |
| GOTERM_BP_DIRECT | GO:0035458~cellular response to interferon-beta | 13 | 3.53 | 1.15442395E-11 | Q60766, Q61107, Q99J93, Q91VJ1, Q9R002, Q9Z0E6, D0QMC3, Q8VCF0, Q91XB0, P0DOV2, P42225, O35368, Q9JIY5 | 17.27 | 2.09412504E-08 |
| GOTERM_BP_DIRECT | GO:0002218~activation of innate immune response | 10 | 2.72 | 8.45447717E-09 | O35368, P07901, P58681, Q61107, Q91VJ1, Q9R002, Q9Z0E6, D0QMC3, Q8VCF0, P0DOV2 | 16.55 | 7.66821079E-06 |
| UP_SEQ_FEATURE | DOMAIN:HIN-200 | 5 | 1.36 | 1.10006066E-05 | O35368, Q91VJ1, Q9R002, D0QMC3, P0DOV2 | 32.73 | 0.004040889489 |
| INTERPRO | IPR004021:HIN200/IF120x | 5 | 1.36 | 1.2436874E-05 | O35368, Q91VJ1, Q9R002, D0QMC3, P0DOV2 | 31.73 | 0.011158049484 |
| INTERPRO | IPR040205:HIN-200 | 5 | 1.36 | 2.63440006E-05 | O35368, Q91VJ1, Q9R002, D0QMC3, P0DOV2 | 26.85 | 0.011158049484 |
| INTERPRO | IPR012340:NA-bd_OB-fold | 10 | 2.72 | 2.89318483E-05 | O35368, Q9JKB3, Q91VJ1, P63242, P25206, Q9R002, P62281, D0QMC3, Q61881, P0DOV2 | 6.35 | 0.011158049484 |
| GOTERM_MF_DIRECT | GO:0003690~double-stranded DNA binding | 9 | 2.45 | 0.000486744937 | P42225, Q921F2, O35368, Q91VJ1, Q9R002, D0QMC3, Q8BVN4, Q91XB0, P0DOV2 | 4.97 | 0.018467675556 |
| GOTERM_BP_DIRECT | GO:0045824~negative regulation of innate immune response | 5 | 1.36 | 0.001783218323 | O35368, Q9R002, D0QMC3, O35309, Q91XB0 | 9.62 | 0.104347033482 |
| INTERPRO | IPR011029:DEATH-like_dom_sf | 7 | 1.90 | 0.003293608048 | O35368, Q8R4K2, P29452, Q91VJ1, D0QMC3, Q8VCF0, P0DOV2 | 4.84 | 0.273454970332 |
| GOTERM_BP_DIRECT | GO:0140972~negative regulation of AIM2 inflammasome complex assembly | 3 | 0.82 | 0.005749386362 | O35368, Q9R002, D0QMC3 | 25.72 | 0.204497781575 |
| UP_SEQ_FEATURE | DOMAIN:Pyrin | 4 | 1.09 | 0.006658134275 | O35368, Q91VJ1, D0QMC3, P0DOV2 | 10.29 | 0.458578998203 |
| INTERPRO | IPR004020:DAPIN | 4 | 1.09 | 0.008005593011 | O35368, Q91VJ1, D0QMC3, P0DOV2 | 9.63 | 0.463123555669 |
| SMART | SM01289:PYRIN | 4 | 1.09 | 0.008053279593 | O35368, Q91VJ1, D0QMC3, P0DOV2 | 9.55 | 0.421454965357 |
| KEGG_PATHWAY | mmu04623:Cytosolic DNA-sensing pathway | 6 | 1.63 | 0.029448654392 | P29452, Q91VJ1, Q9R002, Q8VCF0, Q91XB0, P0DOV2 | 3.45 | 0.160135945349 |
| GOTERM_BP_DIRECT | GO:0032731~positive regulation of interleukin-1 beta production | 4 | 1.09 | 0.03540589416 | O35368, P29452, Q91VJ1, D0QMC3 | 5.55 | 0.553674931082 |
| GOTERM_MF_DIRECT | GO:0008134~transcription factor binding | 4 | 1.09 | 0.099061702986 | O35368, D0QMC3, Q62167, P0DOV2 | 3.61 | 0.591618503943 |
| Annotation Cluster 7 | Enrichment Score: 3.431654880190493 | | | | | | |
| Category | Term | Count | % | PValue | Genes | Fold Enrichment | FDR |

| | | | | | | | |
|---|---|---|---|---|---|---|---|
| GOTERM_MF_DIRECT | GO:0016887~ATP hydrolysis activity | 18 | 4.89 | 3.33615576E-06 | P97450, P07901, Q91VR5, Q64518, Q9QXZ0, P11983, P11499, Q61881, Q62167, P36371, Q9EPU0, P21958, Q62095, P25206, P08113, Q9EPE9, Q922S8, Q8CGK3 | 4.00 | 0.000391267015 |
| GOTERM_MF_DIRECT | GO:0005524~ATP binding | 36 | 9.78 | 2.06308765E-05 | P07901, Q8R4K2, Q91VR5, Q6PHZ2, Q8BTI9, P11983, P31750, Q62167, Q7TT37, Q9DCL9, Q61036, Q9WTP6, P70227, P36371, Q9EPU0, P61089, Q9CZD3, Q3THS6, Q62095, P25206, Q99KH8, Q8CGK3, P97494, Q64518, Q3TRM8, P11499, Q9Z2I8, Q61881, P70698, Q61024, Q9Z2I9, P21958, P08113, Q9EPE9, E9Q3L2, Q922S8 | 2.19 | 0.001478546148 |
| UP_KW_LIGAND | KW-0547~Nucleotide-binding | 48 | 13.04 | 0.005771336226 | Q811U4, P07901, Q8R4K2, Q91VR5, Q6PHZ2, Q60766, Q8BTI9, Q6GV12, Q69ZS7, O08749, P11983, P59325, P31750, Q62167, Q9DCL9, Q61036, Q9WTP6, P70227, P36371, Q8VEH6, Q9EPU0, P61089, Q9CZD3, Q3THS6, Q62095, P25206, P61027, P62835, Q99KH8, Q8CGK3, P97494, Q64518, Q61107, Q3TRM8, P11499, Q9Z0E6, Q6PA06, Q9Z2I8, Q61881, P70698, Q61024, Q99JI6, Q9Z2I9, P21958, P08113, Q9EPE9, E9Q3L2, Q922S8 | 1.41 | 0.080798707157 |
| UP_KW_LIGAND | KW-0067~ATP-binding | 35 | 9.51 | 0.047243286837 | P07901, Q8R4K2, Q91VR5, Q6PHZ2, Q8BTI9, P11983, P31750, Q62167, Q9DCL9, Q61036, Q9WTP6, P70227, P36371, Q9EPU0, P61089, Q9CZD3, Q3THS6, Q62095, P25206, Q99KH8, Q8CGK3, P97494, Q64518, Q3TRM8, P11499, Q9Z2I8, Q61881, P70698, Q61024, Q9Z2I9, P21958, P08113, Q9EPE9, E9Q3L2, Q922S8 | 1.34 | 0.220468671908 |
| Annotation Cluster 8 | Enrichment Score: 2.59562802363803 | | | | | | |
| Category | Term | Count | % | PValue | Genes | Fold Enrichment | FDR |
| GOTERM_BP_DIRECT | GO:0060337~type I interferon-mediated signaling pathway | 6 | 1.63 | 0.000340956799 | Q8VI94, P42225, Q9CQW9, Q99J93, Q9WVL2, Q91XB0 | 9.93 | 0.030924781697 |
| GOTERM_BP_DIRECT | GO:0034341~response to type II interferon | 4 | 1.09 | 0.005958204655 | P42225, Q9CQW9, Q99J93, O35892 | 10.78 | 0.207849677775 |
| GOTERM_BP_DIRECT | GO:0035456~response to interferon-beta | 3 | 0.82 | 0.008040835125 | P42225, Q9CQW9, Q99J93 | 21.77 | 0.247990509761 |
| Annotation Cluster 9 | Enrichment Score: 2.5950966577153136 | | | | | | |
| Category | Term | Count | % | PValue | Genes | Fold Enrichment | FDR |
| GOTERM_MF_DIRECT | GO:0005525~GTP binding | 14 | 3.80 | 0.000303251795 | Q811U4, P07901, Q60766, Q61107, Q69ZS7, P11499, Q9Z0E6, Q6PA06, Q9Z2I8, P59325, Q99JI6, Q8VEH6, P61027, P62835 | 3.37 | 0.013039827171 |
| GOTERM_MF_DIRECT | GO:0003924~GTPase activity | 12 | 3.26 | 0.00057352057 | Q99JI6, Q8VEH6, Q811U4, P63213, Q60766, Q61107, Q69ZS7, P61027, P62835, Q9Z0E6, Q6PA06, Q62167 | 3.59 | 0.019469514072 |
| GOTERM_MF_DIRECT | GO:1990606~membrane scission GTPase motor activity | 5 | 1.36 | 0.000924971286 | Q811U4, Q60766, Q61107, P61027, Q9Z0E6 | 11.45 | 0.028409832344 |
| GOTERM_MF_DIRECT | GO:0003925~G protein activity | 5 | 1.36 | 0.003159753631 | Q99JI6, Q60766, P61027, P62835, Q9Z0E6 | 8.24 | 0.070277279028 |
| INTERPRO | IPR027417:P-loop_NTPase | 24 | 6.52 | 0.007785028116 | Q811U4, Q91VR5, Q60766, Q61107, Q9BDB7, Q69ZS7, Q9Z0E6, Q6PA06, Q61881, P70698, Q62167, Q9WTP6, Q99JI6, P36371, Q8VEH6, Q9EPU0, P21958, Q62095, Q8BV66, P25206, P61027, P62835, Q922S8, Q8CGK3 | 1.80 | 0.463123555669 |
| UP_KW_LIGAND | KW-0342~GTP-binding | 12 | 3.26 | 0.067924432319 | Q99JI6, Q8VEH6, Q811U4, Q60766, Q61107, Q69ZS7, P61027, P62835, Q9Z0E6, Q9Z2I8, Q6PA06, P59325 | 1.79 | 0.271697729276 |
| Annotation Cluster 10 | Enrichment Score: 2.5524559681877137 | | | | | | |
| Category | Term | Count | % | PValue | Genes | Fold Enrichment | FDR |
| KEGG_PATHWAY | mmu00190:Oxidative phosphorylation | 15 | 4.08 | 1.5318532E-06 | P97450, Q62425, Q9CXV1, Q8BVE3, Q8K3J1, P48771, P19783, Q9CPQ1, P19536, Q9CQA3, O09111, P00405, P56394, Q9DC70, P56391 | 5.01 | 7.7511772E-05 |
| KEGG_PATHWAY | mmu05208:Chemical carcinogenesis - reactive oxygen species | 17 | 4.62 | 2.65080731E-05 | P97450, Q62425, Q9CXV1, Q8BTI9, O70145, Q8K3J1, P48771, P19783, P31750, Q9CPQ1, P19536, Q9CQA3, O09111, P00405, P48774, Q9DC70, P56391 | 3.50 | 0.000560856683 |
| GOTERM_BP_DIRECT | GO:0042776~proton motive force-driven mitochondrial ATP synthesis | 6 | 1.63 | 0.000584619988 | P97450, Q9CQA3, Q9CXV1, O09111, Q8K3J1, Q9DC70 | 8.84 | 0.044681200169 |
| KEGG_PATHWAY | mmu04714:Thermogenesis | 14 | 3.80 | 0.001527611149 | P97450, Q62425, Q9CXV1, Q8K3J1, P48771, P19783, Q9CPQ1, P19536, Q9CQA3, O09111, P00405, P56394, Q9DC70, P56391 | 2.78 | 0.017567528212 |
| UP_KW_BIOLOGICAL_PROCESS | KW-0249~Electron transport | 8 | 2.17 | 0.00225290947 | P10639, Q9CQA3, Q62425, Q9CXV1, O09111, Q8K3J1, P00405, Q9DC70 | 4.39 | 0.049564008339 |
| GOTERM_CC_DIRECT | GO:0045271~respiratory chain complex I | 4 | 1.09 | 0.021622276512 | Q62425, O09111, Q8K3J1, Q9DC70 | 6.72 | 0.208054349552 |
| UP_KW_BIOLOGICAL_PROCESS | KW-0679~Respiratory chain | 5 | 1.36 | 0.023531728224 | Q62425, O09111, Q8K3J1, P00405, Q9DC70 | 4.56 | 0.258849010465 |
| GOTERM_BP_DIRECT | GO:0009060~aerobic respiration | 4 | 1.09 | 0.042301915072 | Q9CQA3, O09111, Q8K3J1, Q9DC70 | 5.17 | 0.60421790505 |
| KEGG_PATHWAY | mmu04723:Retrograde endocannabinoid signaling | 7 | 1.90 | 0.105123070331 | P51432, Q62425, P63213, O09111, Q8K3J1, Q9DC70, P70227 | 2.16 | 0.385451257879 |
| GOTERM_BP_DIRECT | GO:0032981~mitochondrial respiratory chain complex I assembly | 3 | 0.82 | 0.161659594065 | O09111, Q8K3J1, Q9DC70 | 4.16 | 0.982665222102 |
| Annotation Cluster 11 | Enrichment Score: 2.4529220434577357 | | | | | | |
| Category | Term | Count | % | PValue | Genes | Fold Enrichment | FDR |
| GOTERM_CC_DIRECT | GO:0022626~cytosolic ribosome | 10 | 2.72 | 8.92911384E-06 | P62892, P62908, P67984, P84099, Q8BP48, Q9D8E6, Q69ZS7, P62281, P62862, P47964 | 7.41 | 0.000276164735 |
| GOTERM_BP_DIRECT | GO:0002181~cytoplasmic translation | 8 | 2.17 | 0.000172975558 | P62892, P62908, P67984, P84099, Q9D8E6, P62281, P62862, P47964 | 6.86 | 0.019611103911 |
| GOTERM_BP_DIRECT | GO:0006412~translation | 11 | 2.99 | 0.000203059608 | P62892, P62908, P67984, P84099, Q9D8E6, Q69ZS7, P62281, P62862, P31750, P47964, Q8BX17 | 4.45 | 0.02083072179 |
| UP_KW_MOLECULAR_FUNCTION | KW-0687~Ribonucleoprotein | 15 | 4.08 | 0.000889741962 | P62908, Q9CW46, Q9D0G0, P57784, P62281, Q8BMA6, Q62189, P62892, P67984, P84099, Q6DFW4, Q9D8E6, P62307, P62862, P47964 | 2.82 | 0.015792919821 |
| GOTERM_MF_DIRECT | GO:0003735~structural constituent of ribosome | 9 | 2.45 | 0.003685770167 | P62892, P62908, P67984, P84099, Q9D8E6, Q9D0G0, P62281, P62862, P47964 | 3.61 | 0.07924405858 |
| GOTERM_CC_DIRECT | GO:0098793~presynapse | 11 | 2.99 | 0.003847264082 | P67984, Q9D8E6, O08539, P62835, P35564, Q9R0P5, P62281, P47964, Q9QY76, Q922S8, P70227 | 3.02 | 0.055528844911 |
| GOTERM_CC_DIRECT | GO:0005840~ribosome | 8 | 2.17 | 0.004186898129 | P62892, P62908, P67984, Q9D8E6, P35564, P62281, P62862, P47964 | 3.98 | 0.058481512577 |
| KEGG_PATHWAY | mmu05171:Coronavirus disease - COVID-19 | 16 | 4.35 | 0.004696329575 | P62908, Q8R4K2, P29452, Q8BTI9, Q9WVL2, P62281, Q8VCF0, P30993, P42225, P62892, P67984, P58681, P84099, Q9D8E6, P62862, P47964 | 2.26 | 0.038196095529 |
| GOTERM_CC_DIRECT | GO:1990904~ribonucleoprotein complex | 10 | 2.72 | 0.007865506323 | P62892, P62908, P67984, Q91VR5, Q9CW46, Q9D8E6, P62281, P62862, Q9JII5, P47964 | 2.91 | 0.094604562158 |

| | | | | | | | |
|---|---|---|---|---|---|---|---|
| GOTERM_CC_DIRECT | GO:0022625~cytosolic large ribosomal subunit | 5 | 1.36 | 0.012303345095 | P62892, P67984, P84099, Q9D8E6, P47964 | 5.60 | 0.136598677589 |
| UP_KW_MOLECULAR_FUNCTION | KW-0689~Ribosomal protein | 9 | 2.45 | 0.019185014428 | P62892, P62908, P67984, P84099, Q9D8E6, Q9D0G0, P62281, P62862, P47964 | 2.68 | 0.123830547669 |
| GOTERM_BP_DIRECT | GO:0140236~translation at presynapse | 4 | 1.09 | 0.030345642223 | P67984, Q9D8E6, P62281, P47964 | 5.90 | 0.520655726567 |
| GOTERM_BP_DIRECT | GO:0140242~translation at postsynapse | 4 | 1.09 | 0.031572287719 | P67984, Q9D8E6, P62281, P47964 | 5.80 | 0.520655726567 |
| KEGG_PATHWAY | mmu03010:Ribosome | 8 | 2.17 | 0.329583194336 | P62892, P62908, P67984, P84099, Q9D8E6, P62281, P62862, P47964 | 1.42 | 0.715611671076 |

Annotation Cluster 12 — Enrichment Score: 2.4176655868880896

| Category | Term | Count | % | PValue | Genes | Fold Enrichment | FDR | |
|---|---|---|---|---|---|---|---|---|
| KEGG_PATHWAY | mmu04613:Neutrophil extracellular trap formation | 15 | 4.08 | 0.000132779932 | P51432, P29452, P26151, P58681, P10854, Q8BTI9, O70145, P0C0S6, P10853, P31750, Q64522, P30993 | 3.38 | 0.002054538774 | |
| INTERPRO | IPR055333:HISTONE_H2B_site | 5 | 1.36 | 0.000133459078 | P10854, P10853 | 18.37 | 0.035467145397 | |
| INTERPRO | IPR000558:Histone_H2B | 5 | 1.36 | 0.000201485003 | P10854, P10853 | 16.62 | 0.035467145397 | |
| SMART | SM00427:H2B | 5 | 1.36 | 0.000201755621 | P10854, P10853 | 16.49 | 0.031675632541 | |
| UP_SEQ_FEATURE | CROSSLNK:Glycyl lysine isopeptide (Lys-Gly) (interchain with G-Cter in ubiq | alternate | 8.00 | 2.173913043478 | 0.000484916321681 | 22253.00 | 0.089066416992 | 0.088664129652624 |
| UP_SEQ_FEATURE | DOMAIN:Histone H2A/H2B/H3 | 7 | 1.90 | 0.000563202275 | P10854, P0C0S6, P10853, Q64522 | 6.81 | 0.088664129653 | |
| INTERPRO | IPR007125:Histone_H2A/H2B/H3 | 7 | 1.90 | 0.000876461061 | P10854, P0C0S6, P10853, Q64522 | 6.27 | 0.112673938611 | |
| GOTERM_BP_DIRECT | GO:0061644~protein localization to CENP-A containing chromatin | 4 | 1.09 | 0.000999167355 | P10853 | 19.86 | 0.064731770772 | |
| GOTERM_MF_DIRECT | GO:0030527~structural constituent of chromatin | 7 | 1.90 | 0.001392079469 | P10854, P0C0S6, P10853, Q64522 | 5.77 | 0.040813238967 | |
| UP_SEQ_FEATURE | CARBOHYD:O-linked (GlcNAc) serine | 6 | 1.63 | 0.001923901643 | P10854, P11499, P10853 | 6.75 | 0.235571067814 | |
| GOTERM_CC_DIRECT | GO:0043505~CENP-A containing nucleosome | 4 | 1.09 | 0.002856350174 | P10853 | 13.93 | 0.047569216365 | |
| UP_KW_CELLULAR_COMPONENT | KW-0544~Nucleosome core | 7 | 1.90 | 0.003774610813 | P10854, P0C0S6, P10853, Q64522 | 4.69 | 0.020131257669 | |
| GOTERM_CC_DIRECT | GO:0000786~nucleosome | 7 | 1.90 | 0.005239030906 | P10854, P0C0S6, P10853, Q64522 | 4.42 | 0.068742435827 | |
| GOTERM_MF_DIRECT | GO:0046982~protein heterodimerization activity | 11 | 2.99 | 0.008328252514 | P10854, Q07813, P0C0S6, P58774, P10853, Q9QY76, Q76MZ3, Q64522 | 2.69 | 0.149214524215 | |
| INTERPRO | IPR009072:Histone-fold | 7 | 1.90 | 0.008525216124 | P10854, P0C0S6, P10853, Q64522 | 3.97 | 0.469698812142 | |
| UP_KW_PTM | KW-0013~ADP-ribosylation | 8 | 2.17 | 0.011010896435 | Q99JI6, P42225, P10854, P10853, Q9WVA3 | 3.28 | 0.027527241087 | |
| UP_KW_PTM | KW-0379~Hydroxylation | 9 | 2.45 | 0.034735217889 | P10854, P08113, P17182, P10853, Q64522 | 2.39 | 0.074432609761 | |
| KEGG_PATHWAY | mmu05322:Systemic lupus erythematosus | 8 | 2.17 | 0.039618495087 | P26151, P10854, P0C0S6, P10853, Q64522 | 2.51 | 0.196538808961 | |
| GOTERM_BP_DIRECT | GO:0006325~chromatin organization | 7 | 1.90 | 0.07311260309 | P54103, Q8BHJ5, P0C0S6, P10853 | 2.40 | 0.740928838019 | |
| KEGG_PATHWAY | mmu05203:Viral carcinogenesis | 9 | 2.45 | 0.110073297162 | P10854, Q07813, Q8BTI9, O35892, P10853, Q62167 | 1.86 | 0.393672676914 | |
| KEGG_PATHWAY | mmu05034:Alcoholism | 8 | 2.17 | 0.148585655505 | P63213, P10854, P0C0S6, P10853, Q64522 | 1.82 | 0.452917720997 | |
| UP_KW_CELLULAR_COMPONENT | KW-0158~Chromosome | 14 | 3.80 | 0.193162821683 | P10854, P0C0S6, Q9D2X5, P25206, P10853, Q61033, Q61881, Q9WVA3, Q922S8, Q64522, Q76MZ3 | 1.41 | 0.474542801696 | |

Annotation Cluster 13 — Enrichment Score: 2.3876161542366323

| Category | Term | Count | % | PValue | Genes | Fold Enrichment | FDR |
|---|---|---|---|---|---|---|---|
| GOTERM_MF_DIRECT | GO:0008233~peptidase activity | 11 | 2.99 | 6.63466165E-06 | P27773, P57716, P29452, Q6NSR8, O70370, P18242, O09043, Q8BHG1, Q9JIY5, Q9D8V0, Q99K23 | 6.66 | 0.000611336681 |
| GOTERM_BP_DIRECT | GO:0006508~proteolysis | 16 | 4.35 | 0.002138740189 | P29452, Q6NSR8, Q8VCT3, O08749, O70370, P97821, P18242, Q99K23, O88839, Q9WVJ3, Q8BP48, O88456, O09043, Q8BHG1, Q9JIY5, Q8CGK3 | 2.49 | 0.114108079511 |
| UP_KW_MOLECULAR_FUNCTION | KW-0645~Protease | 15 | 4.08 | 0.087807701159 | P29452, Q6NSR8, Q8VCT3, O70370, P97821, P18242, Q9D8V0, Q99K23, O88839, Q9WVJ3, Q8BP48, O09043, Q8BHG1, Q9JIY5, Q8CGK3 | 1.59 | 0.470679821462 |
| UP_KW_PTM | KW-0865~Zymogen | 8 | 2.17 | 0.225957086067 | O88839, P29452, Q9WVJ3, O70370, P97821, P18242, O09043, Q9JIY5 | 1.62 | 0.3389356291 |

Annotation Cluster 14 — Enrichment Score: 2.247975454712471

| Category | Term | Count | % | PValue | Genes | Fold Enrichment | FDR |
|---|---|---|---|---|---|---|---|
| GOTERM_BP_DIRECT | GO:0006457~protein folding | 10 | 2.72 | 9.90837916E-06 | P27773, Q8CEC6, P07901, Q9WUD1, Q9QYJ3, P08113, P11499, P11983, P35564, Q8BVF2 | 7.31 | 0.003594759961 |
| KEGG_PATHWAY | mmu04141:Protein processing in endoplasmic reticulum | 15 | 4.08 | 1.92377972E-05 | Q9CQS8, P27773, P61804, P07901, Q9WTX6, P11499, P35564, Q3U2P1, Q6P5E4, Q8VCH8, O08795, Q07813, Q9WUD1, Q9QYJ3, P08113 | 4.04 | 0.00048671627 |
| GOTERM_MF_DIRECT | GO:0051082~unfolded protein binding | 8 | 2.17 | 5.74361089E-05 | P07901, P70670, Q9QYJ3, P08113, P11499, P11983, P35564, Q6P5E4 | 8.17 | 0.003704629022 |
| UP_KW_MOLECULAR_FUNCTION | KW-0143~Chaperone | 14 | 3.80 | 7.47376856E-05 | P07901, P54103, P70670, P11983, P11499, P35564, Q8BVF2, Q8VEH6, Q9CYG7, Q64152, Q9QYJ3, Q91YY4, P08113, P56394 | 3.83 | 0.002653187838 |
| GOTERM_CC_DIRECT | GO:0042470~melanosome | 8 | 2.17 | 9.68766733E-05 | P27773, P57716, P07901, Q68FD5, P18242, P08113, P11499, P07356 | 7.52 | 0.002330422197 |
| GOTERM_MF_DIRECT | GO:0044183~protein folding chaperone | 6 | 1.63 | 0.000347549682 | P07901, Q9QYJ3, P08113, P11499, P11983, Q8BVF2 | 9.89 | 0.014010596565 |
| INTERPRO | IPR019805:Heat_shock_protein_90_CS | 3 | 0.82 | 0.000603976001 | P07901, P08113, P11499 | 69.82 | 0.087350029078 |
| KEGG_PATHWAY | mmu05417:Lipid and atherosclerosis | 14 | 3.80 | 0.000687472811 | P51432, P07901, Q8R4K2, Q6PHZ2, P29452, P10810, Q8BTI9, O70145, P11499, P31750, Q99JI6, Q07813, P08113, P62835 | 3.04 | 0.009154243222 |
| INTERPRO | IPR037196:HSP90_C | 3 | 0.82 | 0.00119653793 | P07901, P08113, P11499 | 52.36 | 0.115366198715 |
| INTERPRO | IPR020575:Hsp90_N | 3 | 0.82 | 0.00119653793 | P07901, P08113, P11499 | 52.36 | 0.115366198715 |
| INTERPRO | IPR001404:Hsp90_fam | 3 | 0.82 | 0.00119653793 | P07901, P08113, P11499 | 52.36 | 0.115366198715 |
| GOTERM_MF_DIRECT | GO:0030235~nitric-oxide synthase regulator activity | 3 | 0.82 | 0.002276583699 | P07901, P11499, P31750 | 40.25 | 0.063843325461 |
| GOTERM_BP_DIRECT | GO:0034605~cellular response to heat | 5 | 1.36 | 0.002384505876 | P07901, Q9QYJ3, P11499, Q8BVF2, Q9JIY5 | 8.90 | 0.120152601653 |
| PIR_SUPERFAMILY | PIRSF002583:Hsp90 | 3 | 0.82 | 0.003759175818 | P07901, P08113, P11499 | 28.64 | 0.184199615096 |
| UP_SEQ_FEATURE | DOMAIN:Histidine kinase/HSP90-like ATPase | 3 | 0.82 | 0.003829974051 | P07901, P08113, P11499 | 30.86 | 0.383693764051 |
| GOTERM_MF_DIRECT | GO:0140662~ATP-dependent protein folding chaperone | 4 | 1.09 | 0.003888352884 | P07901, P08113, P11499, P11983 | 12.52 | 0.080902826141 |
| SMART | SM00387:HATPase_c | 3 | 0.82 | 0.006891965971 | P07901, P08113, P11499 | 23.09 | 0.421454965357 |
| KEGG_PATHWAY | mmu05132:Salmonella infection | 13 | 3.53 | 0.007145847856 | P62908, P10639, P07901, Q8R4K2, P29452, P10810, Q8BTI9, P11499, P31750, Q61036, Q07813, P08113, P07356 | 2.43 | 0.051654271643 |
| INTERPRO | IPR003594:HATPase_dom | 3 | 0.82 | 0.010266231398 | P07901, P08113, P11499 | 19.04 | 0.539910442173 |
| INTERPRO | IPR020568:Ribosomal_Su5_D2-typ_SF | 4 | 1.09 | 0.01803866878 | P07901, P08113, P11499, Q8CGK3 | 7.16 | 0.840020202176 |
| GOTERM_BP_DIRECT | GO:0045429~positive regulation of nitric oxide biosynthetic process | 4 | 1.09 | 0.018560569543 | P07901, Q8BTI9, P11499, P31750 | 7.12 | 0.41956161909 |

| | | | | | | | |
|---|---|---|---|---|---|---|---|
| KEGG_PATHWAY | mmu04915:Estrogen signaling pathway | 8 | 2.17 | 0.023333242736 | P51432, P07901, Q8BTI9, P18242, P08113, P11499, P31750, P70227 | 2.81 | 0.139064262189 |
| KEGG_PATHWAY | mmu05418:Fluid shear stress and atherosclerosis | 8 | 2.17 | 0.037261697509 | P10639, P07901, Q8BTI9, O70145, P08113, P11499, P48774, P31750 | 2.55 | 0.188544189395 |
| UP_KW_BIOLOGICAL_PROCESS | KW-0346~Stress response | 5 | 1.36 | 0.04103613543 | P07901, Q9QYJ3, P08113, P11499, P54071 | 3.83 | 0.328289083439 |
| INTERPRO | IPR036890:HATPase_C_sf | 3 | 0.82 | 0.042216514503 | P07901, P08113, P11499 | 9.11 | 0.993986254296 |
| KEGG_PATHWAY | mmu05215:Prostate cancer | 5 | 1.36 | 0.160873799632 | P07901, Q8BTI9, P08113, P11499, P31750 | 2.36 | 0.47326827101 |
| GOTERM_CC_DIRECT | GO:0031526~brush border membrane | 3 | 0.82 | 0.223670854175 | P07901, Q8BTI9, P11499 | 3.36 | 0.678709607587 |
| KEGG_PATHWAY | mmu04914:Progesterone-mediated oocyte maturation | 4 | 1.09 | 0.31339668917 | P07901, Q8BTI9, P11499, P31750 | 2.03 | 0.714717770895 |
| GOTERM_BP_DIRECT | GO:0032880~regulation of protein localization | 3 | 0.82 | 0.328057545063 | P07901, P11499, P31750 | 2.55 | 0.982665222102 |
| KEGG_PATHWAY | mmu05207:Chemical carcinogenesis - receptor activation | 6 | 1.63 | 0.530573241603 | P07901, Q8BTI9, P08113, P11499, P48774, P31750 | 1.24 | 0.865215634447 |
| KEGG_PATHWAY | mmu04657:IL-17 signaling pathway | 3 | 0.82 | 0.588290302001 | P07901, P08113, P11499 | 1.52 | 0.884220219993 |
| KEGG_PATHWAY | mmu04151:PI3K-Akt signaling pathway | 8 | 2.17 | 0.662078482444 | P07901, P63213, Q8BTI9, O70309, P08113, P11499, P31750, Q76MZ3 | 1.03 | 0.884615384615 |
| Annotation Cluster 15 | Enrichment Score: 1.8379287258566117 | | | | | | |
| Category | Term | Count | % | PValue | Genes | Fold Enrichment | FDR |
| GOTERM_CC_DIRECT | GO:0042825~TAP complex | 3 | 0.82 | 0.000333549301 | P27773, P36371, P21958 | 94.06 | 0.007221342377 |
| KEGG_PATHWAY | mmu04612:Antigen processing and presentation | 8 | 2.17 | 0.002368137244 | P27773, P36371, P07901, P21958, O70370, P11499, P35564, P97371 | 4.33 | 0.023965548907 |
| GOTERM_CC_DIRECT | GO:0042824~MHC class I peptide loading complex | 3 | 0.82 | 0.010732854475 | P27773, P36371, P21958 | 18.81 | 0.125603405075 |
| GOTERM_MF_DIRECT | GO:0042288~MHC class I protein binding | 3 | 0.82 | 0.076318394607 | P27773, P36371, P21958 | 6.55 | 0.512764213763 |
| UP_KW_BIOLOGICAL_PROCESS | KW-1064~Adaptive immunity | 3 | 0.82 | 0.998722881815 | P27773, P36371, P21958 | 0.34 | 0.998722881815 |
| Annotation Cluster 16 | Enrichment Score: 1.7927510650748262 | | | | | | |
| Category | Term | Count | % | PValue | Genes | Fold Enrichment | FDR |
| GOTERM_BP_DIRECT | GO:0006749~glutathione metabolic process | 5 | 1.36 | 0.00273184602 | P97494, Q00612, P48774, P11352, Q8VCN5 | 8.57 | 0.133934288634 |
| KEGG_PATHWAY | mmu00480:Glutathione metabolism | 6 | 1.63 | 0.018842394278 | P97494, Q00612, Q9DCD0, P48774, P11352, P54071 | 3.87 | 0.119178143805 |
| GOTERM_BP_DIRECT | GO:0006979~response to oxidative stress | 5 | 1.36 | 0.081311973326 | P97494, Q00612, Q8K3J1, P31750, P11352 | 3.06 | 0.765942778886 |
| Annotation Cluster 17 | Enrichment Score: 1.7904301455351663 | | | | | | |
| Category | Term | Count | % | PValue | Genes | Fold Enrichment | FDR |
| GOTERM_MF_DIRECT | GO:0051015~actin filament binding | 9 | 2.45 | 0.002985977173 | Q99K51, P21107, Q9QXZ0, P47753, O08539, P58774, P18760, Q9R0P5, Q61233 | 3.74 | 0.069785309762 |
| UP_KW_MOLECULAR_FUNCTION | KW-0009~Actin-binding | 11 | 2.99 | 0.014768856816 | Q99K51, P21107, Q9QXZ0, P47753, P58774, Q0GNC1, P18760, Q9R0P5, Q61233, P49813 | 2.44 | 0.104858883391 |
| GOTERM_MF_DIRECT | GO:0003779~actin binding | 10 | 2.72 | 0.016395694559 | P21107, Q9QXZ0, P47753, P58774, Q0GNC1, Q9Z0E6, P18760, P97814, P49813 | 2.58 | 0.225004744476 |
| GOTERM_CC_DIRECT | GO:0005884~actin filament | 5 | 1.36 | 0.021242104215 | Q99K51, P21107, P58774, Q61233, P49813 | 4.75 | 0.208054349552 |
| GOTERM_CC_DIRECT | GO:0015629~actin cytoskeleton | 7 | 1.90 | 0.072692324172 | P21107, Q9QXZ0, P58774, Q9Z0E6, P18760, Q9R0P5, Q61233 | 2.40 | 0.386204888657 |
| Annotation Cluster 18 | Enrichment Score: 1.7756964164519138 | | | | | | |
| Category | Term | Count | % | PValue | Genes | Fold Enrichment | FDR |
| UP_SEQ_FEATURE | SITE:Contributes to redox potential value | 3 | 0.82 | 0.00276089392 | P27773, P10639, Q9CQM5 | 36.01 | 0.304250509957 |
| INTERPRO | IPR036249:Thioredoxin-like_sf | 8 | 2.17 | 0.003308876046 | P27773, Q8VCH8, P10639, Q9CQM5, P48774, Q8BVF2, P11352, Q9QYB1 | 4.14 | 0.273454970332 |
| UP_KW_DOMAIN | KW-0676~Redox-active center | 4 | 1.09 | 0.030840615629 | P27773, P10639, Q9CQM5, O08749 | 5.83 | 0.295597081216 |
| UP_SEQ_FEATURE | DISULFID:Redox-active | 4 | 1.09 | 0.035270163032 | P27773, P10639, Q9CQM5, O08749 | 5.54 | 0.920745393113 |
| UP_SEQ_FEATURE | DOMAIN:Thioredoxin | 3 | 0.82 | 0.133124255909 | P27773, P10639, Q9CQM5 | 4.70 | 0.996383363472 |
| Annotation Cluster 19 | Enrichment Score: 1.7729264915076417 | | | | | | |
| Category | Term | Count | % | PValue | Genes | Fold Enrichment | FDR |
| KEGG_PATHWAY | mmu00010:Glycolysis / Gluconeogenesis | 7 | 1.90 | 0.002817860485 | P47738, Q7TSV4, Q3TRM8, O08749, Q9DBF1, P17182 | 4.92 | 0.027419950103 |
| GOTERM_BP_DIRECT | GO:0006096~glycolytic process | 4 | 1.09 | 0.015053161295 | Q9Z0M5, Q3TRM8, P17182 | 7.70 | 0.400499705889 |
| UP_KW_BIOLOGICAL_PROCESS | KW-0324~Glycolysis | 3 | 0.82 | 0.113154889123 | Q3TRM8, P17182 | 5.17 | 0.609993145103 |
| Annotation Cluster 20 | Enrichment Score: 1.7667541210743511 | | | | | | |
| Category | Term | Count | % | PValue | Genes | Fold Enrichment | FDR |
| KEGG_PATHWAY | mmu00280:Valine, leucine and isoleucine degradation | 7 | 1.90 | 0.001219526186 | Q9D0K2, P47738, Q99JY0, O08749, Q9DBF1, Q8BMS1, Q8QZT1 | 5.79 | 0.015427006258 |
| GOTERM_BP_DIRECT | GO:0006635~fatty acid beta-oxidation | 5 | 1.36 | 0.002068807874 | O70579, Q99JY0, Q8BMS1, Q9EPL9, Q8QZT1 | 9.25 | 0.113721741922 |
| GOTERM_MF_DIRECT | GO:0003988~acetyl-CoA C-acyltransferase activity | 3 | 0.82 | 0.003014196492 | Q99JY0, Q8BMS1, Q8QZT1 | 35.22 | 0.069785309762 |
| KEGG_PATHWAY | mmu00071:Fatty acid degradation | 6 | 1.63 | 0.004644874979 | P47738, Q99JY0, Q9DBF1, Q8BMS1, Q9EPL9, Q8QZT1 | 5.44 | 0.038196095529 |
| KEGG_PATHWAY | mmu00310:Lysine degradation | 6 | 1.63 | 0.011125765214 | Q9R0E1, P47738, O08749, Q9DBF1, Q8BMS1, Q8QZT1 | 4.42 | 0.075755227557 |
| GOTERM_MF_DIRECT | GO:0051287~NAD binding | 4 | 1.09 | 0.01607162758 | P47738, O08749, Q8BMS1, P54071 | 7.51 | 0.225004744476 |
| KEGG_PATHWAY | mmu00380:Tryptophan metabolism | 5 | 1.36 | 0.02375071359 | P47738, O08749, Q9DBF1, Q8BMS1, Q8QZT1 | 4.53 | 0.139064262189 |
| KEGG_PATHWAY | mmu00410:beta-Alanine metabolism | 4 | 1.09 | 0.029365171076 | P47738, Q9DBF1, Q8BMS1, Q9EPL9 | 5.89 | 0.160135945349 |
| UP_KW_LIGAND | KW-0520~NAD | 9 | 2.45 | 0.032118505089 | Q8JZU0, P47738, Q6R5N8, Q8K3J1, Q9CY64, O08749, Q9DBF1, Q9DC70, Q8BMS1 | 2.40 | 0.184790203221 |
| KEGG_PATHWAY | mmu00620:Pyruvate metabolism | 4 | 1.09 | 0.065367038034 | P47738, O08749, Q9DBF1, Q8QZT1 | 4.28 | 0.266739687463 |
| KEGG_PATHWAY | mmu01212:Fatty acid metabolism | 4 | 1.09 | 0.14278297645 | Q99JY0, Q8BMS1, Q9EPL9, Q8QZT1 | 3.04 | 0.452218878413 |
| UP_KW_BIOLOGICAL_PROCESS | KW-0276~Fatty acid metabolism | 4 | 1.09 | 0.476655005093 | Q99JY0, Q8BMS1, Q9EPL9, Q8QZT1 | 1.55 | 0.967032967033 |
| Annotation Cluster 21 | Enrichment Score: 1.7358139923659024 | | | | | | |
| Category | Term | Count | % | PValue | Genes | Fold Enrichment | FDR |
| GOTERM_BP_DIRECT | GO:0031663~lipopolysaccharide-mediated signaling pathway | 6 | 1.63 | 5.47531018E-05 | P42225, Q8R4K2, P10810, O88188, P31750, Q80WJ7 | 14.51 | 0.009029284236 |
| KEGG_PATHWAY | mmu04620:Toll-like receptor signaling pathway | 7 | 1.90 | 0.02263360854 | P42225, Q8R4K2, P58681, P10810, Q8BTI9, Q9WVL2, P31750 | 3.17 | 0.139064262189 |

| | | | | | | | |
|---|---|---|---|---|---|---|---|
| GOTERM_BP_DIRECT | GO:0048661~positive regulation of smooth muscle cell proliferation | 3 | 0.82 | 0.226630621257 | P42225, Q8R4K2, P31750 | 3.33 | 0.982665222102 |
| KEGG_PATHWAY | mmu05145:Toxoplasmosis | 4 | 1.09 | 0.405754097261 | P42225, Q8R4K2, Q60766, P31750 | 1.73 | 0.814728465135 |
| Annotation Cluster 22 | Enrichment Score: 1.6967800615576325 | | | | | | |
| Category | Term | Count | % | PValue | Genes | Fold Enrichment | FDR |
| INTERPRO | IPR036543:Guanylate-bd_C_sf | 3 | 0.82 | 0.018876858476 | Q61107, Q9Z0E6, Q6PA06 | 13.96 | 0.840020202176 |
| INTERPRO | IPR003191:Guanylate-bd/ATL_C | 3 | 0.82 | 0.018876858476 | Q61107, Q9Z0E6, Q6PA06 | 13.96 | 0.840020202176 |
| UP_SEQ_FEATURE | DOMAIN:GB1/RHD3-type G | 3 | 0.82 | 0.020161659729 | Q61107, Q9Z0E6, Q6PA06 | 13.50 | 0.802456799282 |
| INTERPRO | IPR015894:Guanylate-bd_N | 3 | 0.82 | 0.021372581697 | Q61107, Q9Z0E6, Q6PA06 | 13.09 | 0.883145607996 |
| INTERPRO | IPR030386:G_GB1_RHD3_dom | 3 | 0.82 | 0.021372581697 | Q61107, Q9Z0E6, Q6PA06 | 13.09 | 0.883145607996 |
| Annotation Cluster 23 | Enrichment Score: 1.6828198314742815 | | | | | | |
| Category | Term | Count | % | PValue | Genes | Fold Enrichment | FDR |
| INTERPRO | IPR020568:Ribosomal_Su5_D2-typ_SF | 4 | 1.09 | 0.01803866878 | P07901, P08113, P11499, Q8CGK3 | 7.16 | 0.840020202176 |
| GOTERM_BP_DIRECT | GO:0051131~chaperone-mediated protein complex assembly | 3 | 0.82 | 0.018657838633 | P07901, P11499, Q8CGK3 | 14.15 | 0.41956161909 |
| GOTERM_MF_DIRECT | GO:0070182~DNA polymerase binding | 3 | 0.82 | 0.026575086613 | P07901, P11499, Q8CGK3 | 11.74 | 0.263706628701 |
| Annotation Cluster 24 | Enrichment Score: 1.6009437270262472 | | | | | | |
| Category | Term | Count | % | PValue | Genes | Fold Enrichment | FDR |
| GOTERM_BP_DIRECT | GO:0006006~glucose metabolic process | 6 | 1.63 | 0.001519656223 | Q05816, Q9Z0M5, Q7TSV4, Q3TRM8, Q00612, P31750 | 7.16 | 0.092903022639 |
| UP_KW_BIOLOGICAL_PROCESS | KW-0313~Glucose metabolism | 3 | 0.82 | 0.026486348371 | Q7TSV4, Q00612, P31750 | 11.63 | 0.258977628515 |
| UP_KW_BIOLOGICAL_PROCESS | KW-0119~Carbohydrate metabolism | 3 | 0.82 | 0.391202281713 | Q7TSV4, Q00612, P31750 | 2.21 | 0.967032967033 |
| Annotation Cluster 25 | Enrichment Score: 1.523512737577447 | | | | | | |
| Category | Term | Count | % | PValue | Genes | Fold Enrichment | FDR |
| GOTERM_BP_DIRECT | GO:0042776~proton motive force-driven mitochondrial ATP synthesis | 6 | 1.63 | 0.000584619988 | P97450, Q9CQA3, Q9CXV1, O09111, Q8K3J1, Q9DC70 | 8.84 | 0.044681200169 |
| GOTERM_MF_DIRECT | GO:0051539~4 iron, 4 sulfur cluster binding | 4 | 1.09 | 0.011375630947 | Q9CQA3, Q8K3J1, Q99KI0, Q9DC70 | 8.54 | 0.178039001773 |
| UP_KW_LIGAND | KW-0004~4Fe-4S | 4 | 1.09 | 0.032998250575 | Q9CQA3, Q8K3J1, Q99KI0, Q9DC70 | 5.62 | 0.184790203221 |
| GOTERM_BP_DIRECT | GO:0009060~aerobic respiration | 4 | 1.09 | 0.042301915072 | Q9CQA3, O09111, Q8K3J1, Q9DC70 | 5.17 | 0.60421790505 |
| UP_KW_LIGAND | KW-0411~Iron-sulfur | 4 | 1.09 | 0.133095199108 | Q9CQA3, Q8K3J1, Q99KI0, Q9DC70 | 3.13 | 0.465833196878 |
| UP_KW_LIGAND | KW-0408~Iron | 8 | 2.17 | 0.584872343168 | Q9CQA3, Q9R0E1, Q9CXV1, Q8K3J1, Q99KI0, Q8BGW1, Q9DC70, Q921I1 | 1.10 | 0.933333333333 |
| Annotation Cluster 26 | Enrichment Score: 1.4806995235494202 | | | | | | |
| Category | Term | Count | % | PValue | Genes | Fold Enrichment | FDR |
| GOTERM_BP_DIRECT | GO:0006098~pentose-phosphate shunt | 3 | 0.82 | 0.008040835125 | P40142, Q00612, Q9DCD0 | 21.77 | 0.247990509761 |
| KEGG_PATHWAY | mmu00030:Pentose phosphate pathway | 4 | 1.09 | 0.034363513493 | Q7TSV4, P40142, Q00612, Q9DCD0 | 5.54 | 0.177427937015 |
| GOTERM_MF_DIRECT | GO:0030246~carbohydrate binding | 6 | 1.63 | 0.130768370014 | O09159, P40142, Q00612, Q8K4Q8, Q9DCD0, P35564 | 2.24 | 0.714793208975 |
| Annotation Cluster 27 | Enrichment Score: 1.2723385049942417 | | | | | | |
| Category | Term | Count | % | PValue | Genes | Fold Enrichment | FDR |
| GOTERM_MF_DIRECT | GO:0043531~ADP binding | 4 | 1.09 | 0.01068247785 | P97494, P36371, P21958, Q8CGK3 | 8.74 | 0.172254955332 |
| GOTERM_MF_DIRECT | GO:0003697~single-stranded DNA binding | 5 | 1.36 | 0.030800340871 | Q9JKB3, P25206, Q61881, Q91XB0, Q8CGK3 | 4.23 | 0.278378199537 |
| INTERPRO | IPR003593:AAA+_ATPase | 5 | 1.36 | 0.156651823903 | P36371, P21958, P25206, Q61881, Q8CGK3 | 2.39 | 0.993986254296 |
| SMART | SM00382:AAA | 5 | 1.36 | 0.157936678523 | P36371, P21958, P25206, Q61881, Q8CGK3 | 2.37 | 1 |
| Annotation Cluster 28 | Enrichment Score: 1.27079871683916 | | | | | | |
| Category | Term | Count | % | PValue | Genes | Fold Enrichment | FDR |
| GOTERM_BP_DIRECT | GO:0010756~positive regulation of plasminogen activation | 3 | 0.82 | 0.003816138879 | P17182, P07356 | 31.44 | 0.164820855391 |
| GOTERM_CC_DIRECT | GO:0045121~membrane raft | 6 | 1.63 | 0.111525192286 | P10810, O35682, P18242, P17182, P07356 | 2.36 | 0.459908650093 |
| GOTERM_CC_DIRECT | GO:0005938~cell cortex | 4 | 1.09 | 0.361915095043 | Q9QXZ0, P17182, P07356 | 1.86 | 0.932793072343 |
| Annotation Cluster 29 | Enrichment Score: 1.222952797814185 | | | | | | |
| Category | Term | Count | % | PValue | Genes | Fold Enrichment | FDR |
| UP_KW_BIOLOGICAL_PROCESS | KW-0159~Chromosome partition | 5 | 1.36 | 0.007189579616 | Q9D2X5, Q62167, Q9WVA3, Q922S8, Q76MZ3 | 6.46 | 0.105447167694 |
| GOTERM_CC_DIRECT | GO:0000775~chromosome, centromeric region | 4 | 1.09 | 0.044063751901 | P83917, Q9WVA3, Q922S8, Q76MZ3 | 5.08 | 0.303831217539 |
| UP_KW_CELLULAR_COMPONENT | KW-0137~Centromere | 3 | 0.82 | 0.676638392297 | Q9WVA3, Q922S8, Q76MZ3 | 1.29 | 0.864864864865 |
| Annotation Cluster 30 | Enrichment Score: 1.2075933647509072 | | | | | | |
| Category | Term | Count | % | PValue | Genes | Fold Enrichment | FDR |
| INTERPRO | IPR019956:Ubiquitin_dom | 3 | 0.82 | 0.023996887091 | Q8VDF2, P29595, P62862 | 12.32 | 0.95739304705 |
| SMART | SM00213:UBQ | 4 | 1.09 | 0.036550079404 | Q8VI94, Q8VDF2, P29595, P62862 | 5.43 | 1 |
| INTERPRO | IPR000626:Ubiquitin-like_dom | 4 | 1.09 | 0.063577106814 | Q8VI94, Q8VDF2, P29595, P62862 | 4.36 | 0.993986254296 |
| INTERPRO | IPR029071:Ubiquitin-like_domsf | 7 | 1.90 | 0.067784336671 | Q8VI94, Q8VCH8, Q8VDF2, Q8BTI9, P29595, P62862, Q8VBT9 | 2.44 | 0.993986254296 |
| UP_SEQ_FEATURE | DOMAIN:Ubiquitin-like | 3 | 0.82 | 0.242415191388 | Q8VDF2, P29595, P62862 | 3.18 | 0.996383363472 |
| Annotation Cluster 31 | Enrichment Score: 1.1686687977724204 | | | | | | |
| Category | Term | Count | % | PValue | Genes | Fold Enrichment | FDR |
| UP_KW_DOMAIN | KW-0729~SH3-binding | 4 | 1.09 | 0.042228154459 | O88839, P97797, Q3UIA2, Q61036 | 5.15 | 0.295597081216 |
| GOTERM_MF_DIRECT | GO:0017124~SH3 domain binding | 5 | 1.36 | 0.050122447664 | O88839, P97797, Q3UIA2, P11352, Q61036 | 3.61 | 0.375918357477 |
| UP_SEQ_FEATURE | MOTIF:SH3-binding | 3 | 0.82 | 0.147353085511 | O88839, P97797, Q3UIA2 | 4.41 | 0.996383363472 |

| Annotation Cluster 32 | Enrichment Score: 1.1486852620451533 | | | | | | |
|---|---|---|---|---|---|---|---|
| Category | Term | Count | % | PValue | Genes | Fold Enrichment | FDR |
| UP_SEQ_FEATURE | DOMAIN:EF-hand 2 | 8 | 2.17 | 0.008532571961 | Q99K51, O08795, Q9QXZ0, O88456, Q9QXX4, Q64521, Q8BH59, Q61233 | 3.47 | 0.52238301675 |
| GOTERM_MF_DIRECT | GO:0005509~calcium ion binding | 15 | 4.08 | 0.02371216406 | Q99K51, P51432, Q64518, Q9QXZ0, Q62393, P35564, Q9QXX4, Q64521, Q61233, P70227, O08795, P40142, O88456, P07356, Q8BH59 | 1.94 | 0.246682997079 |
| UP_SEQ_FEATURE | DOMAIN:EF-hand 1 | 7 | 1.90 | 0.027670924113 | Q99K51, O08795, Q9QXZ0, Q9QXX4, Q64521, Q8BH59, Q61233 | 3.06 | 0.802456799282 |
| UP_SEQ_FEATURE | DOMAIN:Calponin-homology (CH) 1 | 3 | 0.82 | 0.039903511903 | Q99K51, Q9QXZ0, Q61233 | 9.39 | 0.921007785899 |
| UP_SEQ_FEATURE | DOMAIN:Calponin-homology (CH) 2 | 3 | 0.82 | 0.039903511903 | Q99K51, Q9QXZ0, Q61233 | 9.39 | 0.921007785899 |
| INTERPRO | IPR011992:EF-hand-dom_pair | 9 | 2.45 | 0.040763367358 | Q99K51, O08795, P51432, Q9QXZ0, O88456, Q9QXX4, Q64521, Q8BH59, Q61233 | 2.33 | 0.993986254296 |
| INTERPRO | IPR001589:Actinin_actin-bd_CS | 3 | 0.82 | 0.042216514503 | Q99K51, Q9QXZ0, Q61233 | 9.11 | 0.993986254296 |
| INTERPRO | IPR002048:EF_hand_dom | 8 | 2.17 | 0.043303656161 | Q99K51, O08795, Q9QXZ0, O88456, Q9QXX4, Q64521, Q8BH59, Q61233 | 2.48 | 0.993986254296 |
| INTERPRO | IPR018247:EF_Hand_1_Ca_BS | 7 | 1.90 | 0.043924653524 | Q99K51, O08795, Q9QXZ0, O88456, Q64521, Q8BH59, Q61233 | 2.73 | 0.993986254296 |
| UP_SEQ_FEATURE | DOMAIN:EF-hand | 7 | 1.90 | 0.058016422666 | Q99K51, O08795, Q9QXZ0, O88456, Q9QXX4, Q64521, Q61233 | 2.55 | 0.996383363472 |
| UP_SEQ_FEATURE | DOMAIN:Calponin-homology (CH) | 4 | 1.09 | 0.091180997233 | Q99K51, Q9QXZ0, Q61166, Q61233 | 3.74 | 0.996383363472 |
| INTERPRO | IPR001715:CH_dom | 4 | 1.09 | 0.1126466653 | Q99K51, Q9QXZ0, Q61166, Q61233 | 3.41 | 0.993986254296 |
| GOTERM_BP_DIRECT | GO:0016477~cell migration | 7 | 1.90 | 0.115943784228 | Q9QXZ0, Q8BTI9, P97797, O70309, Q61166, Q61233, Q61036 | 2.11 | 0.897410629403 |
| INTERPRO | IPR036872:CH_dom_sf | 4 | 1.09 | 0.118793274355 | Q99K51, Q9QXZ0, Q61166, Q61233 | 3.32 | 0.993986254296 |
| SMART | SM00054:EFh | 5 | 1.36 | 0.160788590445 | Q99K51, Q9QXZ0, Q64521, Q8BH59, Q61233 | 2.36 | 1 |
| UP_SEQ_FEATURE | DOMAIN:EF-hand 4 | 3 | 0.82 | 0.206721776339 | O88456, Q9QXX4, Q8BH59 | 3.54 | 0.996383363472 |
| UP_KW_LIGAND | KW-0106~Calcium | 21 | 5.71 | 0.213778220365 | Q99K51, Q8BFR4, P51432, P03975, Q64518, Q9QXZ0, Q8K4Q8, P35564, Q9QXX4, Q64521, Q61233, P70227, O08795, P40142, O08585, O70309, O88456, P08113, P07356, Q8BH59 | 1.26 | 0.66508779669 |
| SMART | SM00033:CH | 3 | 0.82 | 0.244493699144 | Q99K51, Q9QXZ0, Q61233 | 3.15 | 1 |
| UP_SEQ_FEATURE | DOMAIN:EF-hand 3 | 3 | 0.82 | 0.359593719501 | O88456, Q9QXX4, Q8BH59 | 2.37 | 0.996383363472 |

| Annotation Cluster 33 | Enrichment Score: 1.1174468039642005 | | | | | | |
|---|---|---|---|---|---|---|---|
| Category | Term | Count | % | PValue | Genes | Fold Enrichment | FDR |
| UP_KW_CELLULAR_COMPONENT | KW-0647~Proteasome | 4 | 1.09 | 0.050304625612 | Q8VDM4, Q9R1P1, Q9R1P4, P97371 | 4.80 | 0.169236178785 |
| KEGG_PATHWAY | mmu03050:Proteasome | 4 | 1.09 | 0.076502972407 | Q8VDM4, Q9R1P1, Q9R1P4, P97371 | 4.01 | 0.302425812797 |
| GOTERM_CC_DIRECT | GO:0000502~proteasome complex | 3 | 0.82 | 0.115444529375 | Q8VDM4, Q9R1P1, Q9R1P4 | 5.13 | 0.471580011504 |

| Annotation Cluster 34 | Enrichment Score: 1.111304318066078 | | | | | | |
|---|---|---|---|---|---|---|---|
| Category | Term | Count | % | PValue | Genes | Fold Enrichment | FDR |
| KEGG_PATHWAY | mmu04380:Osteoclast differentiation | 10 | 2.72 | 0.002321119045 | P42225, Q64518, P26151, Q8BTI9, O70145, P97797, P08101, Q9WVL2, P31750, P70227 | 3.46 | 0.023965548907 |
| KEGG_PATHWAY | mmu05170:Human immunodeficiency virus 1 infection | 13 | 3.53 | 0.005032985935 | P27773, Q8R4K2, Q8BTI9, Q9WTX6, P18760, P31750, Q61036, P70227, P36371, P63213, Q07813, P21958, P83940 | 2.54 | 0.038586225504 |
| KEGG_PATHWAY | mmu05168:Herpes simplex virus 1 infection | 12 | 3.26 | 0.005505336336 | O35326, P27773, P42225, P36371, Q8R4K2, Q07813, Q8BTI9, P21958, O35892, Q9WVL2, Q8VCF0, P31750 | 2.65 | 0.040966179205 |
| KEGG_PATHWAY | mmu05169:Epstein-Barr virus infection | 12 | 3.26 | 0.008975288239 | P27773, P42225, P36371, Q8R4K2, Q07813, Q8BTI9, P21958, Q8VDM4, Q9WVL2, P20152, Q8VCF0, P31750 | 2.48 | 0.063076331238 |
| KEGG_PATHWAY | mmu05164:Influenza A | 10 | 2.72 | 0.01149675058 | P42225, Q8R4K2, P58681, P29452, Q07813, Q8BTI9, Q9QYJ3, Q9WVL2, Q8VCF0, P31750 | 2.71 | 0.075755227557 |
| KEGG_PATHWAY | mmu05162:Measles | 9 | 2.45 | 0.011677683299 | P42225, Q8R4K2, P58681, Q07813, Q8BTI9, P08101, Q9WVL2, Q8VCF0, P31750 | 2.92 | 0.075755227557 |
| KEGG_PATHWAY | mmu04620:Toll-like receptor signaling pathway | 7 | 1.90 | 0.02263360854 | P42225, Q8R4K2, P58681, P10810, Q8BTI9, Q9WVL2, P31750 | 3.17 | 0.139064262189 |
| KEGG_PATHWAY | mmu05161:Hepatitis B | 8 | 2.17 | 0.058846971556 | P42225, Q8R4K2, Q07813, Q8BTI9, Q9WVL2, Q8VCF0, P31750, Q62167 | 2.30 | 0.24407022629 |
| KEGG_PATHWAY | mmu04625:C-type lectin receptor signaling pathway | 6 | 1.63 | 0.088234427002 | P42225, P29452, Q8BTI9, Q9WVL2, P31750, P70227 | 2.52 | 0.333183731813 |
| KEGG_PATHWAY | mmu04062:Chemokine signaling pathway | 8 | 2.17 | 0.115126902426 | Q99JI6, P42225, P51432, P63213, Q8BTI9, Q9WVL2, P62835, P31750 | 1.95 | 0.398264441946 |
| KEGG_PATHWAY | mmu05160:Hepatitis C | 7 | 1.90 | 0.142994111751 | P42225, Q07813, Q8BTI9, Q9WVL2, Q8VCF0, P31750, Q76MZ3 | 1.98 | 0.452218878413 |
| KEGG_PATHWAY | mmu04933:AGE-RAGE signaling pathway in diabetic complications | 5 | 1.36 | 0.16493454168 | P42225, P51432, Q07813, Q8BTI9, P31750 | 2.33 | 0.479637230404 |
| KEGG_PATHWAY | mmu05163:Human cytomegalovirus infection | 9 | 2.45 | 0.169148585659 | P27773, P36371, P51432, P63213, Q07813, Q8BTI9, P21958, P31750, P70227 | 1.67 | 0.486302183771 |
| KEGG_PATHWAY | mmu05212:Pancreatic cancer | 4 | 1.09 | 0.216395529417 | P42225, Q07813, Q8BTI9, P31750 | 2.48 | 0.564413081881 |
| KEGG_PATHWAY | mmu04935:Growth hormone synthesis, secretion and action | 5 | 1.36 | 0.234272693732 | P42225, P51432, Q8BTI9, P31750, P70227 | 2.01 | 0.586841500141 |
| KEGG_PATHWAY | mmu04919:Thyroid hormone signaling pathway | 5 | 1.36 | 0.248010835913 | P42225, P51432, Q64518, Q8BTI9, P31750 | 1.96 | 0.60333405275 |
| KEGG_PATHWAY | mmu05235:PD-L1 expression and PD-1 checkpoint pathway in cancer | 4 | 1.09 | 0.284447450744 | P42225, Q8BTI9, Q3UMY5, P31750 | 2.14 | 0.660231238882 |
| KEGG_PATHWAY | mmu05167:Kaposi sarcoma-associated herpesvirus infection | 7 | 1.90 | 0.330935041565 | P42225, P63213, Q07813, Q8BTI9, Q9WVL2, P31750, P70227 | 1.48 | 0.715611671076 |
| GOTERM_BP_DIRECT | GO:0019221~cytokine-mediated signaling pathway | 3 | 0.82 | 0.447240237661 | P42225, Q8R4K2, Q9WVL2 | 1.98 | 0.982665222102 |
| KEGG_PATHWAY | mmu04917:Prolactin signaling pathway | 3 | 0.82 | 0.465626276942 | P42225, Q8BTI9, P31750 | 1.91 | 0.864534806435 |
| KEGG_PATHWAY | mmu05165:Human papillomavirus infection | 9 | 2.45 | 0.489761395962 | Q8VI94, P42225, Q07813, Q8BTI9, Q8BVE3, O70309, Q9WVL2, P31750, Q76MZ3 | 1.18 | 0.864534806435 |
| KEGG_PATHWAY | mmu04630:JAK-STAT signaling pathway | 4 | 1.09 | 0.704820525244 | P42225, Q8BTI9, Q9WVL2, P31750 | 1.10 | 0.884615384615 |

| Annotation Cluster 35 | Enrichment Score: 1.0662376119630501 | | | | | | |
|---|---|---|---|---|---|---|---|
| Category | Term | Count | % | PValue | Genes | Fold Enrichment | FDR |
| GOTERM_CC_DIRECT | GO:0016607~nuclear speck | 12 | 3.26 | 0.011501893332 | O35326, Q8C547, Q921F2, P36371, O88986, Q9ERA6, Q91YY4, P57784, Q8BGW1, Q3UKJ7, Q9CWK3, P0DOV2 | 2.42 | 0.131061047703 |
| GOTERM_BP_DIRECT | GO:0008380~RNA splicing | 7 | 1.90 | 0.020455683193 | O35326, Q921F2, Q8CEC6, P62307, Q3UKJ7, Q62189, Q9CWK3 | 3.28 | 0.41956161909 |
| UP_KW_BIOLOGICAL_PROCESS | KW-0508~mRNA splicing | 10 | 2.72 | 0.051659623763 | O35326, Q921F2, Q8CEC6, Q9ERA6, P57784, P62307, Q3UKJ7, Q62189, Q9CWK3, Q8BX17 | 2.07 | 0.378837240929 |
| GOTERM_BP_DIRECT | GO:0006397~mRNA processing | 7 | 1.90 | 0.056402248463 | O35326, Q921F2, Q8CEC6, Q91VR5, Q9JK81, Q62189, Q9CWK3 | 2.57 | 0.69130863995 |

| | | | | | | | | | |
|---|---|---|---|---|---|---|---|---|---|
| UP_KW_BIOLOGICAL_PROCESS | KW-0507~mRNA processing | 11 | 2.99 | 0.081992092287 | O35326, Q921F2, Q8CEC6, Q91VR5, Q9ERA6, P57784, P62307, Q3UKJ7, Q62189, Q9CWK3, Q8BX17 | | 1.81 | 0.515378865807 | |
| UP_KW_CELLULAR_COMPONENT | KW-0747~Spliceosome | 6 | 1.63 | 0.097073887101 | Q8CEC6, Q9ERA6, P57784, P62307, Q3UKJ7, Q62189 | | 2.46 | 0.282396762475 | |
| GOTERM_BP_DIRECT | GO:0000398~mRNA splicing, via spliceosome | 5 | 1.36 | 0.103515552906 | O35326, P57784, P62307, Q3UKJ7, Q62189 | | 2.81 | 0.859295334347 | |
| GOTERM_CC_DIRECT | GO:0071013~catalytic step 2 spliceosome | 4 | 1.09 | 0.128054083003 | Q8CEC6, Q9ERA6, P57784, P62307 | | 3.22 | 0.495734033201 | |
| GOTERM_CC_DIRECT | GO:0071005~U2-type precatalytic spliceosome | 3 | 0.82 | 0.133068272647 | P57784, P62307, Q3UKJ7 | | 4.70 | 0.50542598295 | |
| GOTERM_CC_DIRECT | GO:0005681~spliceosomal complex | 4 | 1.09 | 0.210401818477 | Q9ERA6, P57784, P62307, Q62189 | | 2.53 | 0.646127570215 | |
| KEGG_PATHWAY | mmu03040:Spliceosome | 4 | 1.09 | 0.922481899147 | O35326, P57784, P62307, Q62189 | | 0.71 | 0.922481899147 | |
| Annotation Cluster 36 | Enrichment Score: 1.0648090201147342 | | | | | | | | |
| Category | Term | Count | % | PValue | Genes | | Fold Enrichment | FDR | |
| GOTERM_MF_DIRECT | GO:0061775~cohesin loader activity | 7 | 1.90 | 0.004940672445 | Q9EPU0, Q91VR5, Q62095, Q9D2X5, P25206, Q61881, Q62167 | | 4.47 | 0.096567688689 | |
| GOTERM_MF_DIRECT | GO:0140584~chromatin extrusion motor activity | 6 | 1.63 | 0.019514378609 | Q9EPU0, Q91VR5, Q62095, P25206, Q61881, Q62167 | | 3.89 | 0.226994328708 | |
| GOTERM_MF_DIRECT | GO:0140665~ATP-dependent H3-H4 histone complex chaperone activity | 6 | 1.63 | 0.019514378609 | Q9EPU0, Q91VR5, Q62095, P25206, Q61881, Q62167 | | 3.89 | 0.226994328708 | |
| GOTERM_MF_DIRECT | GO:0140849~ATP-dependent H2AZ histone chaperone activity | 6 | 1.63 | 0.019514378609 | Q9EPU0, Q91VR5, Q62095, P25206, Q61881, Q62167 | | 3.89 | 0.226994328708 | |
| GOTERM_BP_DIRECT | GO:0006260~DNA replication | 5 | 1.36 | 0.023932859874 | Q9EPU0, P54103, P25206, Q61881, Q91XB0 | | 4.58 | 0.452231331367 | |
| GOTERM_MF_DIRECT | GO:0003689~DNA clamp loader activity | 6 | 1.63 | 0.024465852495 | Q9EPU0, Q91VR5, Q62095, P25206, Q61881, Q62167 | | 3.66 | 0.247214296558 | |
| GOTERM_MF_DIRECT | GO:0003724~RNA helicase activity | 4 | 1.09 | 0.038497251692 | Q9EPU0, Q91VR5, Q62095, Q62167 | | 5.37 | 0.310384091764 | |
| GOTERM_MF_DIRECT | GO:0016787~hydrolase activity | 8 | 2.17 | 0.045615274962 | Q8BFR4, Q8JZU0, Q91VR5, Q60766, Q62095, P18242, Q61881, Q62167 | | 2.46 | 0.346139439414 | |
| UP_SEQ_FEATURE | DOMAIN:DEAD-box RNA helicase Q | 3 | 0.82 | 0.057004959614 | Q91VR5, Q62095, Q62167 | | 7.72 | 0.996383363472 | |
| UP_SEQ_FEATURE | MOTIF:DEAD box | 3 | 0.82 | 0.07217916651 | Q91VR5, Q62095, Q62167 | | 6.75 | 0.996383363472 | |
| INTERPRO | IPR014014:RNA_helicase_DEAD_Q_motif | 3 | 0.82 | 0.088902360642 | Q91VR5, Q62095, Q62167 | | 5.98 | 0.993986254296 | |
| UP_KW_MOLECULAR_FUNCTION | KW-0347~Helicase | 6 | 1.63 | 0.09281010564 | Q9EPU0, Q91VR5, Q62095, P25206, Q61881, Q62167 | | 2.49 | 0.470679821462 | |
| GOTERM_MF_DIRECT | GO:0036121~double-stranded DNA helicase activity | 3 | 0.82 | 0.09873459618 | Q9EPU0, P25206, Q61881 | | 5.63 | 0.591618503943 | |
| UP_SEQ_FEATURE | MOTIF:Q motif | 3 | 0.82 | 0.105748151504 | Q91VR5, Q62095, Q62167 | | 5.40 | 0.996383363472 | |
| INTERPRO | IPR011545:DEAD/DEAH_box_helicase_dom | 3 | 0.82 | 0.27993632973 | Q91VR5, Q62095, Q62167 | | 2.87 | 0.993986254296 | |
| UP_SEQ_FEATURE | DOMAIN:Helicase C-terminal | 3 | 0.82 | 0.455930660484 | Q91VR5, Q62095, Q62167 | | 1.95 | 0.996383363472 | |
| INTERPRO | IPR001650:Helicase_C-like | 3 | 0.82 | 0.472020592188 | Q91VR5, Q62095, Q62167 | | 1.89 | 0.993986254296 | |
| SMART | SM00490:HELICc | 3 | 0.82 | 0.475000194859 | Q91VR5, Q62095, Q62167 | | 1.87 | 1 | |
| UP_SEQ_FEATURE | DOMAIN:Helicase ATP-binding | 3 | 0.82 | 0.483205586703 | Q91VR5, Q62095, Q62167 | | 1.85 | 0.996383363472 | |
| SMART | SM00487:DEXDc | 3 | 0.82 | 0.484351972424 | Q91VR5, Q62095, Q62167 | | 1.84 | 1 | |
| INTERPRO | IPR014001:Helicase_ATP-bd | 3 | 0.82 | 0.485941893074 | Q91VR5, Q62095, Q62167 | | 1.84 | 0.993986254296 | |
| Annotation Cluster 37 | Enrichment Score: 1.018612617097401 | | | | | | | | |
| Category | Term | Count | % | PValue | Genes | | Fold Enrichment | FDR | |
| GOTERM_MF_DIRECT | GO:0030170~pyridoxal phosphate binding | 4 | 1.09 | 0.019720970949 | O88986, Q922Q1, P29758, Q8VCN5 | | 6.96 | 0.226994328708 | |
| INTERPRO | IPR015422:PyrdxlP-dep_Trfase_small | 3 | 0.82 | 0.093276618287 | O88986, P29758, Q8VCN5 | | 5.82 | 0.993986254296 | |
| INTERPRO | IPR015421:PyrdxlP-dep_Trfase_major | 3 | 0.82 | 0.120714327752 | O88986, P29758, Q8VCN5 | | 4.99 | 0.993986254296 | |
| INTERPRO | IPR015424:PyrdxlP-dep_Trfase | 3 | 0.82 | 0.125462920353 | O88986, P29758, Q8VCN5 | | 4.87 | 0.993986254296 | |
| UP_KW_LIGAND | KW-0663~Pyridoxal phosphate | 3 | 0.82 | 0.289708752669 | O88986, P29758, Q8VCN5 | | 2.79 | 0.811184507474 | |
| Annotation Cluster 38 | Enrichment Score: 0.9615304266270306 | | | | | | | | |
| Category | Term | Count | % | PValue | Genes | | Fold Enrichment | FDR | |
| UP_SEQ_FEATURE | REPEAT:Solcar 1 | 5 | 1.36 | 0.004673545786 | O70579, Q791T5, Q791V5, Q9QXX4, Q8BH59 | | 7.35 | 0.39617288123 | |
| UP_SEQ_FEATURE | REPEAT:Solcar 2 | 5 | 1.36 | 0.004673545786 | O70579, Q791T5, Q791V5, Q9QXX4, Q8BH59 | | 7.35 | 0.39617288123 | |
| INTERPRO | IPR018108:Mitochondrial_sb/sol_carrier | 5 | 1.36 | 0.006444963252 | O70579, Q791T5, Q791V5, Q9QXX4, Q8BH59 | | 6.71 | 0.463123555669 | |
| INTERPRO | IPR023395:Mt_carrier_dom_sf | 5 | 1.36 | 0.006444963252 | O70579, Q791T5, Q791V5, Q9QXX4, Q8BH59 | | 6.71 | 0.463123555669 | |
| UP_SEQ_FEATURE | REPEAT:Solcar | 4 | 1.09 | 0.018996015132 | O70579, Q791T5, Q791V5, Q9QXX4 | | 7.03 | 0.802456799282 | |
| INTERPRO | IPR002067:Mit_carrier | 3 | 0.82 | 0.072070360092 | O70579, Q9QXX4, Q8BH59 | | 6.76 | 0.993986254296 | |
| UP_SEQ_FEATURE | REPEAT:Solcar 3 | 3 | 0.82 | 0.133124255909 | O70579, Q9QXX4, Q8BH59 | | 4.70 | 0.996383363472 | |
| UP_SEQ_FEATURE | TRANSMEM:Helical | Name=1 | 10.00 | 2.717391304348 | | 0.722986719507322 | 22253.00 | 1 | 0.996383363471971 |
| UP_SEQ_FEATURE | TRANSMEM:Helical | Name=2 | 10.00 | 2.717391304348 | | 0.730069533719899 | 22253.00 | 1 | 0.996383363471971 |
| UP_SEQ_FEATURE | TRANSMEM:Helical | Name=6 | 9.00 | 2.445652173913 | | 0.775239404621747 | 22253.00 | 1 | 0.996383363471971 |
| UP_SEQ_FEATURE | TRANSMEM:Helical | Name=5 | 9.00 | 2.445652173913 | | 0.782951242147234 | 22253.00 | 1 | 0.996383363471971 |
| UP_SEQ_FEATURE | TRANSMEM:Helical | Name=3 | 9.00 | 2.445652173913 | | 0.807258802922292 | 22253.00 | 1 | 0.996383363471971 |
| UP_SEQ_FEATURE | TRANSMEM:Helical | Name=4 | 9.00 | 2.445652173913 | | 0.813003732951358 | 22253.00 | 1 | 0.996383363471971 |
| UP_SEQ_FEATURE | TRANSMEM:Helical | Name=7 | 4.00 | 1.086956521739 | | 0.994015849852411 | 22253.00 | 1 | 0.996383363471971 |
| Annotation Cluster 39 | Enrichment Score: 0.8911220552374226 | | | | | | | | |
| Category | Term | Count | % | PValue | Genes | | Fold Enrichment | FDR | |
| GOTERM_CC_DIRECT | GO:0005819~spindle | 7 | 1.90 | 0.00692882397 | P62908, P83917, O08585, Q68FD5, P31750, Q61166, Q922S8 | | 4.17 | 0.085719450832 | |
| GOTERM_BP_DIRECT | GO:0051301~cell division | 8 | 2.17 | 0.106580953067 | P62908, O08585, Q9D2X5, Q68FD5, Q3UMY5, Q61166, Q9WVA3, Q922S8 | | 2.00 | 0.870891211095 | |
| UP_KW_BIOLOGICAL_PROCESS | KW-0498~Mitosis | 8 | 2.17 | 0.172192423829 | P62908, O08585, Q9D2X5, Q68FD5, Q3UMY5, Q61166, Q9WVA3, Q922S8 | | 1.75 | 0.757646664848 | |
| UP_KW_BIOLOGICAL_PROCESS | KW-0132~Cell division | 8 | 2.17 | 0.448641440199 | P62908, O08585, Q9D2X5, Q68FD5, Q3UMY5, Q61166, Q9WVA3, Q922S8 | | 1.26 | 0.967032967033 | |
| UP_KW_BIOLOGICAL_PROCESS | KW-0131~Cell cycle | 11 | 2.99 | 0.613955891067 | P62908, Q8VDF2, O08585, Q9D2X5, Q68FD5, P25206, Q3UMY5, Q61881, Q61166, Q9WVA3, Q922S8 | | 1.04 | 0.967032967033 | |
| Annotation Cluster 40 | Enrichment Score: 0.8851560133714548 | | | | | | | | |
| Category | Term | Count | % | PValue | Genes | | Fold Enrichment | FDR | |
| UP_SEQ_FEATURE | DOMAIN:RRM 1 | 5 | 1.36 | 0.036408704543 | O35326, Q921F2, Q9CW46, Q62189, Q9JII5 | | 4.00 | 0.920745393113 | |
| UP_SEQ_FEATURE | DOMAIN:RRM 2 | 5 | 1.36 | 0.036408704543 | O35326, Q921F2, Q9CW46, Q62189, Q9JII5 | | 4.00 | 0.920745393113 | |

| | | | | | | | |
|---|---|---|---|---|---|---|---|
| INTERPRO | IPR035979:RBD_domain_sf | 7 | 1.90 | 0.185964793226 | O35326, Q921F2, Q61545, Q9CW46, Q62189, Q9JII5, O35309 | 1.82 | 0.993986254296 |
| INTERPRO | IPR012677:Nucleotide-bd_a/b_plait_sf | 7 | 1.90 | 0.201200933581 | O35326, Q921F2, Q61545, Q9CW46, Q62189, Q9JII5, O35309 | 1.78 | 0.993986254296 |
| SMART | SM00360:RRM | 6 | 1.63 | 0.221915223585 | O35326, Q921F2, Q61545, Q9CW46, Q62189, Q9JII5 | 1.85 | 1 |
| UP_SEQ_FEATURE | DOMAIN:RRM | 6 | 1.63 | 0.227737664475 | O35326, Q921F2, Q61545, Q9CW46, Q62189, Q9JII5 | 1.84 | 0.996383363472 |
| INTERPRO | IPR000504:RRM_dom | 6 | 1.63 | 0.253988919995 | O35326, Q921F2, Q61545, Q9CW46, Q62189, Q9JII5 | 1.76 | 0.993986254296 |

Annotation Cluster 41 Enrichment Score: 0.868212897158529

| Category | Term | Count | % | PValue | Genes | Fold Enrichment | FDR |
|---|---|---|---|---|---|---|---|
| GOTERM_BP_DIRECT | GO:0006511~ubiquitin-dependent protein catabolic process | 8 | 2.17 | 0.013173000473 | Q9D4H8, Q8VDF2, P61089, Q9WTX6, Q9WUD1, Q7TMY8, P83940, Q9R1P4 | 3.20 | 0.373372232168 |
| GOTERM_BP_DIRECT | GO:0006513~protein monoubiquitination | 4 | 1.09 | 0.017644943297 | P61089, Q9WTX6, Q9WUD1, Q7TMY8 | 7.26 | 0.416308572192 |
| GOTERM_MF_DIRECT | GO:0030674~protein-macromolecule adaptor activity | 7 | 1.90 | 0.021614995638 | P57716, Q9D4H8, Q60766, Q9WTX6, Q9WUD1, P83940, Q9JM90 | 3.24 | 0.228552003058 |
| GOTERM_BP_DIRECT | GO:0043161~proteasome-mediated ubiquitin-dependent protein catabolic pro | 9 | 2.45 | 0.02755249938 | Q8BHJ5, P61089, Q9WTX6, Q9WUD1, Q7TMY8, Q8VDM4, Q9R1P1, Q9R1P4, P31750 | 2.52 | 0.504850847231 |
| GOTERM_BP_DIRECT | GO:0006281~DNA repair | 7 | 1.90 | 0.090597162501 | P23492, P62908, Q9EPU0, Q8VDF2, Q9WUD1, Q7TMY8, Q3TGW2 | 2.27 | 0.799227132233 |
| GOTERM_BP_DIRECT | GO:0016567~protein ubiquitination | 9 | 2.45 | 0.098960910386 | Q9D4H8, Q8VDF2, P61089, Q9WTX6, Q9WUD1, Q7TMY8, P83940, P62862, P31750 | 1.92 | 0.851070984578 |
| KEGG_PATHWAY | mmu04120:Ubiquitin mediated proteolysis | 6 | 1.63 | 0.226160866216 | Q9D4H8, P61089, Q9WTX6, Q9WUD1, Q7TMY8, P83940 | 1.84 | 0.572186991526 |
| GOTERM_MF_DIRECT | GO:0004842~ubiquitin-protein transferase activity | 4 | 1.09 | 0.468789381008 | Q8VDF2, P61089, Q9WUD1, Q7TMY8 | 1.56 | 0.959821428571 |
| UP_KW_BIOLOGICAL_PROCESS | KW-0833~Ubl conjugation pathway | 12 | 3.26 | 0.6217488049 | Q9D4H8, Q8VDF2, Q8BHJ5, P29595, P61089, Q9WTX6, Q9WUD1, Q7TMY8, P83940, Q9CR09, Q8BXC6, Q99K23 | 1.03 | 0.967032967033 |
| GOTERM_MF_DIRECT | GO:0061630~ubiquitin protein ligase activity | 4 | 1.09 | 0.711290056338 | Q8VDF2, Q9WTX6, Q9WUD1, Q7TMY8 | 1.09 | 0.959821428571 |
| UP_KW_BIOLOGICAL_PROCESS | KW-0227~DNA damage | 6 | 1.63 | 0.808923953705 | P62908, Q8VDF2, P61089, Q91VJ1, Q9WUD1, Q7TMY8 | 0.89 | 0.967032967033 |
| UP_KW_BIOLOGICAL_PROCESS | KW-0234~DNA repair | 5 | 1.36 | 0.810366000439 | P62908, Q8VDF2, P61089, Q9WUD1, Q7TMY8 | 0.90 | 0.967032967033 |

Annotation Cluster 42 Enrichment Score: 0.8515861112174429

| Category | Term | Count | % | PValue | Genes | Fold Enrichment | FDR |
|---|---|---|---|---|---|---|---|
| SMART | SM00028:TPR | 5 | 1.36 | 0.096147638925 | Q9CYG7, O70145, Q9WUD1, Q9D2X5, Q0VGY8 | 2.86 | 1 |
| UP_SEQ_FEATURE | REPEAT:TPR 3 | 5 | 1.36 | 0.097958211168 | Q9CYG7, O70145, Q9WUD1, Q9D2X5, Q0VGY8 | 2.86 | 0.996383363472 |
| INTERPRO | IPR019734:TPR_rpt | 5 | 1.36 | 0.111249608372 | Q9CYG7, O70145, Q9WUD1, Q9D2X5, Q0VGY8 | 2.73 | 0.993986254296 |
| UP_SEQ_FEATURE | REPEAT:TPR 1 | 5 | 1.36 | 0.12273943216 | Q9CYG7, O70145, Q9WUD1, Q9D2X5, Q0VGY8 | 2.63 | 0.996383363472 |
| UP_SEQ_FEATURE | REPEAT:TPR 2 | 5 | 1.36 | 0.12273943216 | Q9CYG7, O70145, Q9WUD1, Q9D2X5, Q0VGY8 | 2.63 | 0.996383363472 |
| UP_KW_DOMAIN | KW-0802~TPR repeat | 5 | 1.36 | 0.140344184885 | Q9CYG7, O70145, Q9WUD1, Q9D2X5, Q0VGY8 | 2.49 | 0.437315592645 |
| INTERPRO | IPR011990:TPR-like_helical_dom_sf | 6 | 1.63 | 0.207737194635 | Q9CYG7, O70145, Q9WUD1, Q9D2X5, Q68FD5, Q0VGY8 | 1.90 | 0.993986254296 |
| UP_SEQ_FEATURE | REPEAT:TPR | 3 | 0.82 | 0.334469287775 | O70145, Q9WUD1, Q0VGY8 | 2.51 | 0.996383363472 |

Annotation Cluster 43 Enrichment Score: 0.8253457959700067

| Category | Term | Count | % | PValue | Genes | Fold Enrichment | FDR |
|---|---|---|---|---|---|---|---|
| GOTERM_BP_DIRECT | GO:0048015~phosphatidylinositol-mediated signaling | 3 | 0.82 | 0.022372556943 | P51432, Q8BTI9, E9Q3L2 | 12.86 | 0.429299855006 |
| KEGG_PATHWAY | mmu04070:Phosphatidylinositol signaling system | 4 | 1.09 | 0.330813088568 | P51432, Q8BTI9, E9Q3L2, P70227 | 1.96 | 0.715611671076 |
| KEGG_PATHWAY | mmu00562:Inositol phosphate metabolism | 3 | 0.82 | 0.451506810893 | P51432, Q8BTI9, E9Q3L2 | 1.96 | 0.864534806435 |

Annotation Cluster 44 Enrichment Score: 0.8252467973718555

| Category | Term | Count | % | PValue | Genes | Fold Enrichment | FDR |
|---|---|---|---|---|---|---|---|
| GOTERM_CC_DIRECT | GO:0005794~Golgi apparatus | 20 | 5.43 | 0.038017021484 | P57716, Q9QXZ0, Q60766, Q3UVL4, P10810, Q61107, B9EKI3, P11983, Q9Z0E6, Q61166, Q3UM29, Q9QY76, Q8BVL3, Q8C547, Q9R0E1, Q9WVJ3, Q91VW5, P61027, Q99KH8, Q80UJ7 | 1.64 | 0.29395304112 |
| GOTERM_CC_DIRECT | GO:0000139~Golgi membrane | 11 | 2.99 | 0.119103589999 | P57716, Q60766, Q61107, B9EKI3, Q7TMY8, Q91VW5, P61027, Q9Z0E6, Q99KH8, Q3UM29, O88829 | 1.69 | 0.477517171015 |
| UP_KW_CELLULAR_COMPONENT | KW-0333~Golgi apparatus | 14 | 3.80 | 0.738510488324 | Q9QXZ0, Q60766, Q3UVL4, P10810, Q61107, B9EKI3, Q9Z0E6, Q80V94, Q3UM29, O88829, Q9WVJ3, Q91VW5, P61027, Q8BG07 | 0.93 | 0.864864864865 |

Annotation Cluster 45 Enrichment Score: 0.8031770743245792

| Category | Term | Count | % | PValue | Genes | Fold Enrichment | FDR |
|---|---|---|---|---|---|---|---|
| KEGG_PATHWAY | mmu05230:Central carbon metabolism in cancer | 5 | 1.36 | 0.057732930171 | Q8BTI9, Q3TRM8, Q00612, P31750, P54071 | 3.41 | 0.24344052222 |
| KEGG_PATHWAY | mmu04973:Carbohydrate digestion and absorption | 4 | 1.09 | 0.084359243548 | P51432, Q8BTI9, Q3TRM8, P31750 | 3.85 | 0.323377100267 |
| KEGG_PATHWAY | mmu04910:Insulin signaling pathway | 3 | 0.82 | 0.799673123828 | Q8BTI9, Q3TRM8, P31750 | 1.01 | 0.884615384615 |

Annotation Cluster 46 Enrichment Score: 0.7872446197446867

| Category | Term | Count | % | PValue | Genes | Fold Enrichment | FDR |
|---|---|---|---|---|---|---|---|
| UP_SEQ_FEATURE | REPEAT:LRR 23 | 3 | 0.82 | 0.00506007779 | Q9QXZ0, P58681, Q6R5N8 | 27.01 | 0.398300408905 |
| UP_SEQ_FEATURE | REPEAT:LRR 22 | 3 | 0.82 | 0.01149950844 | Q9QXZ0, P58681, Q6R5N8 | 18.00 | 0.633622915039 |
| UP_SEQ_FEATURE | REPEAT:LRR 21 | 3 | 0.82 | 0.020161659729 | Q9QXZ0, P58681, Q6R5N8 | 13.50 | 0.802456799282 |
| UP_SEQ_FEATURE | REPEAT:LRR 20 | 3 | 0.82 | 0.036762973954 | Q9QXZ0, P58681, Q6R5N8 | 9.82 | 0.920745393113 |
| UP_SEQ_FEATURE | REPEAT:LRR 18 | 3 | 0.82 | 0.060683006741 | Q9QXZ0, P58681, Q6R5N8 | 7.45 | 0.996383363472 |
| UP_SEQ_FEATURE | REPEAT:LRR 19 | 3 | 0.82 | 0.060683006741 | Q9QXZ0, P58681, Q6R5N8 | 7.45 | 0.996383363472 |
| UP_SEQ_FEATURE | REPEAT:LRR 17 | 3 | 0.82 | 0.08848228759 | Q9QXZ0, P58681, Q6R5N8 | 6.00 | 0.996383363472 |
| UP_SEQ_FEATURE | REPEAT:LRR 16 | 3 | 0.82 | 0.110196979009 | Q9QXZ0, P58681, Q6R5N8 | 5.27 | 0.996383363472 |
| UP_SEQ_FEATURE | REPEAT:LRR 11 | 4 | 1.09 | 0.113875528281 | Q9QXZ0, P58681, P10810, Q6R5N8 | 3.39 | 0.996383363472 |
| UP_SEQ_FEATURE | REPEAT:LRR 15 | 3 | 0.82 | 0.147353085511 | Q9QXZ0, P58681, Q6R5N8 | 4.41 | 0.996383363472 |
| UP_SEQ_FEATURE | REPEAT:LRR 14 | 3 | 0.82 | 0.157003727333 | Q9QXZ0, P58681, Q6R5N8 | 4.24 | 0.996383363472 |
| UP_SEQ_FEATURE | REPEAT:LRR 10 | 4 | 1.09 | 0.18864244136 | Q9QXZ0, P58681, P10810, Q6R5N8 | 2.67 | 0.996383363472 |
| UP_SEQ_FEATURE | REPEAT:LRR 13 | 3 | 0.82 | 0.191599932697 | Q9QXZ0, P58681, Q6R5N8 | 3.72 | 0.996383363472 |

| | | | | | | | | | |
|---|---|---|---|---|---|---|---|---|---|
| UP_SEQ_FEATURE | REPEAT:LRR 12 | 3 | 0.82 | 0.25780742438 | Q9QXZ0, P58681, Q6R5N8 | | 3.04 | 0.996383363472 | |
| UP_SEQ_FEATURE | REPEAT:LRR 9 | 4 | 1.09 | 0.283649579126 | Q9QXZ0, P58681, P10810, Q6R5N8 | | 2.15 | 0.996383363472 | |
| UP_SEQ_FEATURE | REPEAT:LRR 1 | 6 | 1.63 | 0.319412252086 | Q9QXZ0, P58681, P10810, Q6R5N8, P57784, Q8K1T1 | | 1.61 | 0.996383363472 | |
| UP_SEQ_FEATURE | REPEAT:LRR 8 | 4 | 1.09 | 0.336551028904 | Q9QXZ0, P58681, P10810, Q6R5N8 | | 1.95 | 0.996383363472 | |
| UP_SEQ_FEATURE | REPEAT:LRR 7 | 4 | 1.09 | 0.420621323265 | Q9QXZ0, P58681, P10810, Q6R5N8 | | 1.69 | 0.996383363472 | |
| UP_SEQ_FEATURE | REPEAT:LRR 4 | 5 | 1.36 | 0.431458705444 | Q9QXZ0, P58681, P10810, Q6R5N8, P57784 | | 1.49 | 0.996383363472 | |
| UP_SEQ_FEATURE | REPEAT:LRR 3 | 5 | 1.36 | 0.478944144315 | Q9QXZ0, P58681, P10810, Q6R5N8, P57784 | | 1.40 | 0.996383363472 | |
| UP_SEQ_FEATURE | REPEAT:LRR 2 | 5 | 1.36 | 0.511484232348 | P58681, P10810, Q6R5N8, P57784, Q8K1T1 | | 1.34 | 0.996383363472 | |
| UP_SEQ_FEATURE | REPEAT:LRR 6 | 4 | 1.09 | 0.520912535068 | Q9QXZ0, P58681, P10810, Q6R5N8 | | 1.45 | 0.996383363472 | |
| INTERPRO | IPR001611:Leu-rich_rpt | 5 | 1.36 | 0.521501063192 | P58681, P10810, Q6R5N8, P57784, Q8K1T1 | | 1.33 | 0.993986254296 | |
| UP_KW_DOMAIN | KW-0433~Leucine-rich repeat | 6 | 1.63 | 0.570022779983 | Q9QXZ0, P58681, P10810, Q6R5N8, P57784, Q8K1T1 | | 1.18 | 1 | |
| UP_SEQ_FEATURE | REPEAT:LRR 5 | 4 | 1.09 | 0.607785821212 | Q9QXZ0, P58681, P10810, Q6R5N8 | | 1.27 | 0.996383363472 | |
| INTERPRO | IPR032675:LRR_dom_sf | 6 | 1.63 | 0.701263838712 | P58681, P10810, Q6R5N8, P57784, Q8K1T1, P49813 | | 1.02 | 0.993986254296 | |
| Annotation Cluster 47 | Enrichment Score: 0.7544036815550133 | | | | | | | | |
| Category | Term | Count | % | PValue | Genes | | Fold Enrichment | FDR | |
| KEGG_PATHWAY | mmu05211:Renal cell carcinoma | 7 | 1.90 | 0.003514262597 | Q99JI6, Q9D4H8, Q8BTI9, P62835, P83940, P31750, Q61036 | | 4.71 | 0.031753872751 | |
| KEGG_PATHWAY | mmu04722:Neurotrophin signaling pathway | 7 | 1.90 | 0.04290978953 | Q99JI6, Q6PHZ2, Q8R4K2, Q07813, Q8BTI9, P62835, P31750 | | 2.73 | 0.20302491404 | |
| KEGG_PATHWAY | mmu04611:Platelet activation | 7 | 1.90 | 0.050520290011 | Q99JI6, P51432, Q8BTI9, O09044, P62835, P31750, P70227 | | 2.62 | 0.224239181978 | |
| KEGG_PATHWAY | mmu04062:Chemokine signaling pathway | 8 | 2.17 | 0.115126902426 | Q99JI6, P42225, P51432, P63213, Q8BTI9, Q9WVL2, P62835, P31750 | | 1.95 | 0.398264441946 | |
| KEGG_PATHWAY | mmu04012:ErbB signaling pathway | 4 | 1.09 | 0.261463354402 | Q6PHZ2, Q8BTI9, P31750, Q61036 | | 2.24 | 0.624058760979 | |
| KEGG_PATHWAY | mmu04024:cAMP signaling pathway | 7 | 1.90 | 0.336521491121 | Q99JI6, Q6PHZ2, Q64518, Q8BTI9, P62835, Q9EPL9, P31750 | | 1.47 | 0.72152489198 | |
| KEGG_PATHWAY | mmu04510:Focal adhesion | 6 | 1.63 | 0.428553315038 | Q99JI6, Q8BTI9, O70309, P62835, P31750, Q61036 | | 1.40 | 0.84049603647 | |
| KEGG_PATHWAY | mmu04670:Leukocyte transendothelial migration | 4 | 1.09 | 0.461744319807 | Q99JI6, Q8BTI9, O70145, P62835 | | 1.58 | 0.864534806435 | |
| KEGG_PATHWAY | mmu04014:Ras signaling pathway | 6 | 1.63 | 0.560688812504 | Q99JI6, P63213, Q8BTI9, P62835, P31750, Q61036 | | 1.20 | 0.874566943666 | |
| KEGG_PATHWAY | mmu04015:Rap1 signaling pathway | 5 | 1.36 | 0.666552899701 | Q99JI6, P51432, Q8BTI9, P62835, P31750 | | 1.10 | 0.884615384615 | |
| KEGG_PATHWAY | mmu04010:MAPK signaling pathway | 5 | 1.36 | 0.881332300422 | Q99JI6, Q8R4K2, P10810, P62835, P31750 | | 0.79 | 0.884615384615 | |
| Annotation Cluster 48 | Enrichment Score: 0.7464001750031353 | | | | | | | | |
| Category | Term | Count | % | PValue | Genes | | Fold Enrichment | FDR | |
| UP_SEQ_FEATURE | LIPID:GPI-anchor amidated asparagine | 3 | 0.82 | 0.03078806628 | P03975, P10810 | | 10.80 | 0.869960231807 | |
| UP_SEQ_FEATURE | PROPEP:Removed in mature form | 8 | 2.17 | 0.101430927932 | Q99JI6, Q9Z0M5, P63213, P03975, P10810, P62835, Q9Z0E6 | | 2.03 | 0.996383363472 | |
| GOTERM_CC_DIRECT | GO:0098552~side of membrane | 3 | 0.82 | 0.39400805338 | P03975, P10810 | | 2.20 | 0.935205183585 | |
| UP_KW_PTM | KW-0336~GPI-anchor | 3 | 0.82 | 0.840120127893 | P03975, P10810 | | 0.92 | 0.840120127893 | |
| Annotation Cluster 49 | Enrichment Score: 0.7434712306134643 | | | | | | | | |
| Category | Term | Count | % | PValue | Genes | | Fold Enrichment | FDR | |
| UP_SEQ_FEATURE | REPEAT:HEAT 3 | 3 | 0.82 | 0.152163146554 | Q8C547, Q8BFY9, Q76MZ3 | | 4.32 | 0.996383363472 | |
| UP_SEQ_FEATURE | REPEAT:HEAT 1 | 3 | 0.82 | 0.196624854332 | Q8C547, Q8BFY9, Q76MZ3 | | 3.66 | 0.996383363472 | |
| UP_SEQ_FEATURE | REPEAT:HEAT 2 | 3 | 0.82 | 0.196624854332 | Q8C547, Q8BFY9, Q76MZ3 | | 3.66 | 0.996383363472 | |
| Annotation Cluster 50 | Enrichment Score: 0.7313648136393137 | | | | | | | | |
| Category | Term | Count | % | PValue | Genes | | Fold Enrichment | FDR | |
| GOTERM_MF_DIRECT | GO:0003925~G protein activity | 5 | 1.36 | 0.003159753631 | Q99JI6, Q60766, P61027, P62835, Q9Z0E6 | | 8.24 | 0.070277279028 | |
| GOTERM_MF_DIRECT | GO:0019003~GDP binding | 5 | 1.36 | 0.011869266785 | Q99JI6, P61027, P62835, Q9Z0E6, Q9Z2I8 | | 5.66 | 0.178039001773 | |
| UP_SEQ_FEATURE | LIPID:S-geranylgeranyl cysteine | 5 | 1.36 | 0.05656061412 | Q99JI6, P63213, P61027, P62835, Q9Z0E6 | | 3.46 | 0.996383363472 | |
| SMART | SM00176:RAN | 3 | 0.82 | 0.33478708765 | Q99JI6, P61027, P62835 | | 2.50 | 1 | |
| UP_SEQ_FEATURE | MOTIF:Effector region | 3 | 0.82 | 0.364578253385 | Q99JI6, P61027, P62835 | | 2.35 | 0.996383363472 | |
| UP_KW_PTM | KW-0636~Prenylation | 5 | 1.36 | 0.442275422539 | Q99JI6, P63213, P61027, P62835, Q9Z0E6 | | 1.47 | 0.603102848917 | |
| SMART | SM00174:RHO | 3 | 0.82 | 0.538248673684 | Q99JI6, P61027, P62835 | | 1.66 | 1 | |
| SMART | SM00173:RAS | 3 | 0.82 | 0.571982147635 | Q99JI6, P61027, P62835 | | 1.56 | 1 | |
| INTERPRO | IPR001806:Small_GTPase | 3 | 0.82 | 0.576703799235 | Q99JI6, P61027, P62835 | | 1.55 | 0.993986254296 | |
| SMART | SM00175:RAB | 3 | 0.82 | 0.64125723676 | Q99JI6, P61027, P62835 | | 1.38 | 1 | |
| INTERPRO | IPR005225:Small_GTP-bd | 3 | 0.82 | 0.691501331892 | Q99JI6, P61027, P62835 | | 1.25 | 0.993986254296 | |
| Annotation Cluster 51 | Enrichment Score: 0.7243664365531233 | | | | | | | | |
| Category | Term | Count | % | PValue | Genes | | Fold Enrichment | FDR | |
| UP_KW_MOLECULAR_FUNCTION | KW-1278~Translocase | 7 | 1.90 | 0.00721300463 | P36371, Q64518, P21958, Q8K3J1, P00405, Q9EPE9, Q9DC70 | | 4.09 | 0.082091012822 | |
| UP_SEQ_FEATURE | TRANSMEM:Helical | Name=9 | 3.00 | 0.815217391304 | | 0.403877893181954 | 22253.00 | 1 | 0.996383363471971 |
| UP_SEQ_FEATURE | TRANSMEM:Helical | Name=8 | 3.00 | 0.815217391304 | | 0.437295140613582 | 22253.00 | 1 | 0.996383363471971 |
| UP_SEQ_FEATURE | TRANSMEM:Helical | Name=7 | 4.00 | 1.086956521739 | | 0.994015849852411 | 22253.00 | 1 | 0.996383363471971 |
| Annotation Cluster 52 | Enrichment Score: 0.6828994134334242 | | | | | | | | |
| Category | Term | Count | % | PValue | Genes | | Fold Enrichment | FDR | |
| GOTERM_BP_DIRECT | GO:0006910~phagocytosis, recognition | 4 | 1.09 | 0.000848011258 | P21956, P26151, Q8K4Q8, P97797 | | 20.96 | 0.059165093188 | |
| GOTERM_BP_DIRECT | GO:0050766~positive regulation of phagocytosis | 5 | 1.36 | 0.006492787995 | P21956, P26151, P97797, P08101, P62835 | | 6.74 | 0.218109581913 | |
| GOTERM_BP_DIRECT | GO:0006911~phagocytosis, engulfment | 4 | 1.09 | 0.008065843482 | P21956, P26151, P97797, P08101 | | 9.67 | 0.247990509761 | |
| GOTERM_CC_DIRECT | GO:0009897~external side of plasma membrane | 6 | 1.63 | 0.69670709701 | P21956, P03975, P26151, P10810, Q8VCT3, P08101 | | 1.02 | 0.935205183585 | |
| UP_KW_DOMAIN | KW-0393~Immunoglobulin domain | 3 | 0.82 | 0.997120223895 | P26151, P97797, P08101 | | 0.38 | 1 | |
| INTERPRO | IPR003599:Ig_sub | 3 | 0.82 | 0.998636555493 | P26151, P97797, P08101 | | 0.34 | 0.998636555493 | |
| SMART | SM00409:IG | 3 | 0.82 | 0.998820602485 | P26151, P97797, P08101 | | 0.34 | 1 | |

| | | | | | | | |
|---|---|---|---|---|---|---|---|
| UP_SEQ_FEATURE | DOMAIN:Ig-like | 3 | 0.82 | 0.999883526129 | P26151, P97797, P08101 | 0.26 | 0.999883526129 |
| INTERPRO | IPR013783:Ig-like_fold | 6 | 1.63 | 0.999967448769 | Q99P91, P26151, P97797, P08101, Q9QY76, P70206 | 0.32 | 0.999967448769 |
| INTERPRO | IPR007110:Ig-like_dom | 3 | 0.82 | 0.999993430443 | P26151, P97797, P08101 | 0.21 | 0.999993430443 |
| INTERPRO | IPR036179:Ig-like_dom_sf | 3 | 0.82 | 0.999996978609 | P26151, P97797, P08101 | 0.20 | 0.999996978609 |
| Annotation Cluster 53 | Enrichment Score: 0.6635536561916783 | | | | | | |
| Category | Term | Count | % | PValue | Genes | Fold Enrichment | FDR |
| KEGG_PATHWAY | mmu05164:Influenza A | 10 | 2.72 | 0.01149675058 | P42225, Q8R4K2, P58681, P29452, Q07813, Q8BTI9, Q9QYJ3, Q9WVL2, Q8VCF0, P31750 | 2.71 | 0.075755227557 |
| KEGG_PATHWAY | mmu05162:Measles | 9 | 2.45 | 0.011677683299 | P42225, Q8R4K2, P58681, Q07813, Q8BTI9, P08101, Q9WVL2, Q8VCF0, P31750 | 2.92 | 0.075755227557 |
| KEGG_PATHWAY | mmu04915:Estrogen signaling pathway | 8 | 2.17 | 0.023333242736 | P51432, P07901, Q8BTI9, P18242, P08113, P11499, P31750, P70227 | 2.81 | 0.139064262189 |
| KEGG_PATHWAY | mmu04210:Apoptosis | 8 | 2.17 | 0.024185089076 | Q07813, Q8BTI9, O70370, P97821, P18242, Q9JIY5, P31750, P70227 | 2.79 | 0.139064262189 |
| KEGG_PATHWAY | mmu04722:Neurotrophin signaling pathway | 7 | 1.90 | 0.04290978953 | Q99JI6, Q6PHZ2, Q8R4K2, Q07813, Q8BTI9, P62835, P31750 | 2.73 | 0.20302491404 |
| KEGG_PATHWAY | mmu05161:Hepatitis B | 8 | 2.17 | 0.058846971556 | P42225, Q8R4K2, Q07813, Q8BTI9, Q9WVL2, Q8VCF0, P31750, Q62167 | 2.30 | 0.24407022629 |
| KEGG_PATHWAY | mmu04625:C-type lectin receptor signaling pathway | 6 | 1.63 | 0.088234427002 | P42225, P29452, Q8BTI9, Q9WVL2, P31750, P70227 | 2.52 | 0.333183731813 |
| BIOCARTA | m_chemicalPathway:Apoptotic Signaling in Response to DNA Damage | 3 | 0.82 | 0.107216457394 | P42225, Q07813, P31750 | 5.16 | 1 |
| KEGG_PATHWAY | mmu04071:Sphingolipid signaling pathway | 6 | 1.63 | 0.127917439656 | P51432, Q07813, Q8BTI9, P18242, P31750, Q76MZ3 | 2.24 | 0.425830424118 |
| KEGG_PATHWAY | mmu05160:Hepatitis C | 7 | 1.90 | 0.142994111751 | P42225, Q07813, Q8BTI9, Q9WVL2, Q8VCF0, P31750, Q76MZ3 | 1.98 | 0.452218878413 |
| KEGG_PATHWAY | mmu04933:AGE-RAGE signaling pathway in diabetic complications | 5 | 1.36 | 0.16493454168 | P42225, P51432, Q07813, Q8BTI9, P31750 | 2.33 | 0.479637230404 |
| KEGG_PATHWAY | mmu05223:Non-small cell lung cancer | 4 | 1.09 | 0.194532209215 | Q07813, Q8BTI9, Q3UMY5, P31750 | 2.62 | 0.523581371611 |
| KEGG_PATHWAY | mmu05214:Glioma | 4 | 1.09 | 0.210879703672 | Q6PHZ2, Q07813, Q8BTI9, P31750 | 2.51 | 0.555755885718 |
| KEGG_PATHWAY | mmu05212:Pancreatic cancer | 4 | 1.09 | 0.216395529417 | P42225, Q07813, Q8BTI9, P31750 | 2.48 | 0.564413081881 |
| KEGG_PATHWAY | mmu01524:Platinum drug resistance | 4 | 1.09 | 0.244393035355 | Q07813, Q8BTI9, P48774, P31750 | 2.33 | 0.600305222764 |
| GOTERM_BP_DIRECT | GO:0018105~peptidyl-serine phosphorylation | 3 | 0.82 | 0.308670153828 | Q6PHZ2, Q07813, P31750 | 2.67 | 0.982665222102 |
| KEGG_PATHWAY | mmu05167:Kaposi sarcoma-associated herpesvirus infection | 7 | 1.90 | 0.330935041565 | P42225, P63213, Q07813, Q8BTI9, Q9WVL2, P31750, P70227 | 1.48 | 0.715611671076 |
| KEGG_PATHWAY | mmu05213:Endometrial cancer | 3 | 0.82 | 0.347428643198 | Q07813, Q8BTI9, P31750 | 2.44 | 0.734163642045 |
| KEGG_PATHWAY | mmu05218:Melanoma | 3 | 0.82 | 0.451506810893 | Q07813, Q8BTI9, P31750 | 1.96 | 0.864534806435 |
| KEGG_PATHWAY | mmu05220:Chronic myeloid leukemia | 3 | 0.82 | 0.479530232567 | Q07813, Q8BTI9, P31750 | 1.86 | 0.864534806435 |
| KEGG_PATHWAY | mmu05165:Human papillomavirus infection | 9 | 2.45 | 0.489761395962 | Q8VI94, P42225, Q07813, Q8BTI9, Q8BVE3, O70309, Q9WVL2, P31750, Q76MZ3 | 1.18 | 0.864534806435 |
| KEGG_PATHWAY | mmu01521:EGFR tyrosine kinase inhibitor resistance | 3 | 0.82 | 0.506665727749 | Q07813, Q8BTI9, P31750 | 1.77 | 0.865215634447 |
| KEGG_PATHWAY | mmu05210:Colorectal cancer | 3 | 0.82 | 0.558114368452 | Q07813, Q8BTI9, P31750 | 1.61 | 0.874566943666 |
| KEGG_PATHWAY | mmu04211:Longevity regulating pathway | 3 | 0.82 | 0.570369745869 | Q07813, Q8BTI9, P31750 | 1.57 | 0.874566943666 |
| KEGG_PATHWAY | mmu01522:Endocrine resistance | 3 | 0.82 | 0.588290302001 | Q07813, Q8BTI9, P31750 | 1.52 | 0.884220219993 |
| KEGG_PATHWAY | mmu05222:Small cell lung cancer | 3 | 0.82 | 0.59414006877 | Q07813, Q8BTI9, P31750 | 1.50 | 0.884220219993 |
| KEGG_PATHWAY | mmu04630:JAK-STAT signaling pathway | 4 | 1.09 | 0.704820525244 | P42225, Q8BTI9, Q9WVL2, P31750 | 1.10 | 0.884615384615 |
| KEGG_PATHWAY | mmu05225:Hepatocellular carcinoma | 4 | 1.09 | 0.719554916527 | Q07813, Q8BTI9, P48774, P31750 | 1.08 | 0.884615384615 |
| KEGG_PATHWAY | mmu05166:Human T-cell leukemia virus 1 infection | 5 | 1.36 | 0.770767653973 | Q07813, Q8BTI9, P35564, P31750, Q9WVA3 | 0.95 | 0.884615384615 |
| KEGG_PATHWAY | mmu05224:Breast cancer | 3 | 0.82 | 0.82127781192 | Q07813, Q8BTI9, P31750 | 0.96 | 0.884615384615 |
| KEGG_PATHWAY | mmu05226:Gastric cancer | 3 | 0.82 | 0.829883373078 | Q07813, Q8BTI9, P31750 | 0.94 | 0.884615384615 |
| KEGG_PATHWAY | mmu04140:Autophagy - animal | 3 | 0.82 | 0.871972348501 | Q8BTI9, P18242, P31750 | 0.85 | 0.884615384615 |
| Annotation Cluster 54 | Enrichment Score: 0.6233174870222405 | | | | | | |
| Category | Term | Count | % | PValue | Genes | Fold Enrichment | FDR |
| KEGG_PATHWAY | mmu04071:Sphingolipid signaling pathway | 6 | 1.63 | 0.127917439656 | P51432, Q07813, Q8BTI9, P18242, P31750, Q76MZ3 | 2.24 | 0.425830424118 |
| KEGG_PATHWAY | mmu05160:Hepatitis C | 7 | 1.90 | 0.142994111751 | P42225, Q07813, Q8BTI9, Q9WVL2, Q8VCF0, P31750, Q76MZ3 | 1.98 | 0.452218878413 |
| KEGG_PATHWAY | mmu05142:Chagas disease | 5 | 1.36 | 0.173165362687 | P51432, Q8R4K2, Q8BTI9, P31750, Q76MZ3 | 2.29 | 0.492256592806 |
| KEGG_PATHWAY | mmu04660:T cell receptor signaling pathway | 4 | 1.09 | 0.478111082298 | Q8BTI9, P31750, Q76MZ3, Q61036 | 1.54 | 0.864534806435 |
| KEGG_PATHWAY | mmu04152:AMPK signaling pathway | 4 | 1.09 | 0.504864478047 | Q8BTI9, P61027, P31750, Q76MZ3 | 1.48 | 0.865215634447 |
| Annotation Cluster 55 | Enrichment Score: 0.609922370255613 | | | | | | |
| Category | Term | Count | % | PValue | Genes | Fold Enrichment | FDR |
| KEGG_PATHWAY | mmu04261:Adrenergic signaling in cardiomyocytes | 7 | 1.90 | 0.112569876202 | P21107, P51432, Q6PHZ2, Q64518, P58774, P31750, Q76MZ3 | 2.11 | 0.395558037209 |
| KEGG_PATHWAY | mmu05410:Hypertrophic cardiomyopathy | 4 | 1.09 | 0.354014118847 | P21107, Q64518, O70309, P58774 | 1.88 | 0.740211339407 |
| KEGG_PATHWAY | mmu05414:Dilated cardiomyopathy | 4 | 1.09 | 0.37135572003 | P21107, Q64518, O70309, P58774 | 1.83 | 0.757685461029 |
| Annotation Cluster 56 | Enrichment Score: 0.5496508547180026 | | | | | | |
| Category | Term | Count | % | PValue | Genes | Fold Enrichment | FDR |
| KEGG_PATHWAY | mmu05017:Spinocerebellar ataxia | 8 | 2.17 | 0.029748574828 | P51432, Q64518, Q8BTI9, Q8VDM4, Q9R1P1, Q9R1P4, P31750, P70227 | 2.67 | 0.160135945349 |
| KEGG_PATHWAY | mmu04611:Platelet activation | 7 | 1.90 | 0.050520290011 | Q99JI6, P51432, Q8BTI9, O09044, P62835, P31750, P70227 | 2.62 | 0.224239181978 |
| KEGG_PATHWAY | mmu04720:Long-term potentiation | 5 | 1.36 | 0.052846047724 | Q99JI6, P51432, Q6PHZ2, P62835, P70227 | 3.52 | 0.226611018207 |
| KEGG_PATHWAY | mmu04973:Carbohydrate digestion and absorption | 4 | 1.09 | 0.084359243548 | P51432, Q8BTI9, Q3TRM8, P31750 | 3.85 | 0.323377100267 |
| KEGG_PATHWAY | mmu04625:C-type lectin receptor signaling pathway | 6 | 1.63 | 0.088234427002 | P42225, P29452, Q8BTI9, Q9WVL2, P31750, P70227 | 2.52 | 0.333183731813 |
| KEGG_PATHWAY | mmu04725:Cholinergic synapse | 6 | 1.63 | 0.090823111321 | P51432, Q6PHZ2, P63213, Q8BTI9, P31750, P70227 | 2.50 | 0.337915399475 |
| KEGG_PATHWAY | mmu04723:Retrograde endocannabinoid signaling | 7 | 1.90 | 0.105123070331 | P51432, Q62425, P63213, O09111, Q8K3J1, Q9DC70, P70227 | 2.16 | 0.385451257879 |
| KEGG_PATHWAY | mmu04261:Adrenergic signaling in cardiomyocytes | 7 | 1.90 | 0.112569876202 | P21107, P51432, Q6PHZ2, Q64518, P58774, P31750, Q76MZ3 | 2.11 | 0.395558037209 |
| KEGG_PATHWAY | mmu04062:Chemokine signaling pathway | 8 | 2.17 | 0.115126902426 | Q99JI6, P42225, P51432, P63213, Q8BTI9, Q9WVL2, P62835, P31750 | 1.95 | 0.398264441946 |
| KEGG_PATHWAY | mmu04071:Sphingolipid signaling pathway | 6 | 1.63 | 0.127917439656 | P51432, Q07813, Q8BTI9, P18242, P31750, Q76MZ3 | 2.24 | 0.425830424118 |
| GOTERM_BP_DIRECT | GO:0071320~cellular response to cAMP | 3 | 0.82 | 0.136040573137 | Q99JI6, P62835, P70227 | 4.64 | 0.934574458772 |
| KEGG_PATHWAY | mmu04929:GnRH secretion | 4 | 1.09 | 0.147733047956 | P51432, Q8BTI9, P31750, P70227 | 2.99 | 0.452917720997 |

| | | | | | | | |
|---|---|---|---|---|---|---|---|
| KEGG_PATHWAY | mmu04728:Dopaminergic synapse | 6 | 1.63 | 0.160453902544 | P51432, Q6PHZ2, P63213, P31750, Q76MZ3, P70227 | 2.08 | 0.47326827101 |
| KEGG_PATHWAY | mmu04933:AGE-RAGE signaling pathway in diabetic complications | 5 | 1.36 | 0.16493454168 | P42225, P51432, Q07813, Q8BTI9, P31750 | 2.33 | 0.479637230404 |
| KEGG_PATHWAY | mmu05163:Human cytomegalovirus infection | 9 | 2.45 | 0.169148585659 | P27773, P36371, P51432, P63213, Q07813, Q8BTI9, P21958, P31750, P70227 | 1.67 | 0.486302183771 |
| KEGG_PATHWAY | mmu05142:Chagas disease | 5 | 1.36 | 0.173165362687 | P51432, Q8R4K2, Q8BTI9, P31750, Q76MZ3 | 2.29 | 0.492256592806 |
| GOTERM_MF_DIRECT | GO:0005516~calmodulin binding | 5 | 1.36 | 0.176649898765 | P51432, Q61545, Q6PHZ2, P31750, Q8VCN5 | 2.26 | 0.849684625611 |
| KEGG_PATHWAY | mmu04972:Pancreatic secretion | 5 | 1.36 | 0.225221450832 | Q99JI6, P51432, Q64518, P62835, P70227 | 2.05 | 0.572186991526 |
| GOTERM_BP_DIRECT | GO:0070374~positive regulation of ERK1 and ERK2 cascade | 5 | 1.36 | 0.229682451623 | Q99JI6, Q99P91, Q6PHZ2, P62835, P30993 | 2.04 | 0.982665222102 |
| KEGG_PATHWAY | mmu04935:Growth hormone synthesis, secretion and action | 5 | 1.36 | 0.234272693732 | P42225, P51432, Q8BTI9, P31750, P70227 | 2.01 | 0.586841500141 |
| KEGG_PATHWAY | mmu04919:Thyroid hormone signaling pathway | 5 | 1.36 | 0.248010835913 | P42225, P51432, Q64518, Q8BTI9, P31750 | 1.96 | 0.60333405275 |
| KEGG_PATHWAY | mmu04070:Phosphatidylinositol signaling system | 4 | 1.09 | 0.330813088568 | P51432, Q8BTI9, E9Q3L2, P70227 | 1.96 | 0.715611671076 |
| KEGG_PATHWAY | mmu05167:Kaposi sarcoma-associated herpesvirus infection | 7 | 1.90 | 0.330935041565 | P42225, P63213, Q07813, Q8BTI9, Q9WVL2, P31750, P70227 | 1.48 | 0.715611671076 |
| KEGG_PATHWAY | mmu04713:Circadian entrainment | 4 | 1.09 | 0.348219909271 | P51432, Q6PHZ2, P63213, P70227 | 1.90 | 0.734163642045 |
| KEGG_PATHWAY | mmu04922:Glucagon signaling pathway | 4 | 1.09 | 0.377118753232 | P51432, Q6PHZ2, P31750, P70227 | 1.81 | 0.763288356541 |
| KEGG_PATHWAY | mmu04934:Cushing syndrome | 5 | 1.36 | 0.448006762468 | Q99JI6, P51432, Q6PHZ2, P62835, P70227 | 1.45 | 0.864534806435 |
| KEGG_PATHWAY | mmu04971:Gastric acid secretion | 3 | 0.82 | 0.472605635113 | P51432, Q6PHZ2, P70227 | 1.88 | 0.864534806435 |
| KEGG_PATHWAY | mmu04926:Relaxin signaling pathway | 4 | 1.09 | 0.520576499684 | P51432, P63213, Q8BTI9, P31750 | 1.45 | 0.865215634447 |
| KEGG_PATHWAY | mmu04750:Inflammatory mediator regulation of TRP channels | 4 | 1.09 | 0.520576499684 | P51432, Q6PHZ2, Q8BTI9, P70227 | 1.45 | 0.865215634447 |
| KEGG_PATHWAY | mmu04911:Insulin secretion | 3 | 0.82 | 0.545613920807 | P51432, Q6PHZ2, P70227 | 1.64 | 0.869952760515 |
| KEGG_PATHWAY | mmu04371:Apelin signaling pathway | 4 | 1.09 | 0.56603815467 | P51432, P63213, P31750, P70227 | 1.36 | 0.874566943666 |
| KEGG_PATHWAY | mmu04912:GnRH signaling pathway | 3 | 0.82 | 0.570369745869 | P51432, Q6PHZ2, P70227 | 1.57 | 0.874566943666 |
| KEGG_PATHWAY | mmu05205:Proteoglycans in cancer | 5 | 1.36 | 0.629340459658 | Q6PHZ2, Q8BTI9, O70309, P31750, P70227 | 1.15 | 0.884615384615 |
| KEGG_PATHWAY | mmu04925:Aldosterone synthesis and secretion | 3 | 0.82 | 0.649237881182 | P51432, Q6PHZ2, P70227 | 1.36 | 0.884615384615 |
| KEGG_PATHWAY | mmu04015:Rap1 signaling pathway | 5 | 1.36 | 0.666552899701 | Q99JI6, P51432, Q8BTI9, P62835, P31750 | 1.10 | 0.884615384615 |
| KEGG_PATHWAY | mmu04724:Glutamatergic synapse | 3 | 0.82 | 0.707360786002 | P51432, P63213, P70227 | 1.22 | 0.884615384615 |
| KEGG_PATHWAY | mmu04022:cGMP-PKG signaling pathway | 4 | 1.09 | 0.708558248128 | P51432, Q64518, P31750, P70227 | 1.10 | 0.884615384615 |
| KEGG_PATHWAY | mmu04726:Serotonergic synapse | 3 | 0.82 | 0.786318996969 | P51432, P63213, P70227 | 1.04 | 0.884615384615 |
| KEGG_PATHWAY | mmu04072:Phospholipase D signaling pathway | 3 | 0.82 | 0.827056969827 | P51432, Q8BTI9, P31750 | 0.95 | 0.884615384615 |
| KEGG_PATHWAY | mmu04921:Oxytocin signaling pathway | 3 | 0.82 | 0.840779038169 | P51432, Q6PHZ2, P70227 | 0.92 | 0.884615384615 |
| KEGG_PATHWAY | mmu04218:Cellular senescence | 3 | 0.82 | 0.902652040792 | Q8BTI9, P31750, P70227 | 0.77 | 0.902652040792 |
| KEGG_PATHWAY | mmu04020:Calcium signaling pathway | 4 | 1.09 | 0.909625646209 | P51432, Q6PHZ2, Q64518, P70227 | 0.74 | 0.909625646209 |
| KEGG_PATHWAY | mmu04081:Hormone signaling | 3 | 0.82 | 0.948435551697 | P51432, P63213, Q8BTI9 | 0.65 | 0.948435551697 |
| GOTERM_BP_DIRECT | GO:0007186~G protein-coupled receptor signaling pathway | 4 | 1.09 | 0.999995976406 | P51432, P63213, P30993, P70227 | 0.23 | 0.999995976406 |

Annotation Cluster 57 — Enrichment Score: 0.5367355916410483

| Category | Term | Count | % | PValue | Genes | Fold Enrichment | FDR |
|---|---|---|---|---|---|---|---|
| INTERPRO | IPR020472:G-protein_beta_WD-40_rep | 4 | 1.09 | 0.144547610593 | Q8BHJ5, Q3UKJ7, Q9WVA3, Q8BX17 | 3.04 | 0.993986254296 |
| UP_SEQ_FEATURE | REPEAT:WD 8 | 3 | 0.82 | 0.166769458852 | Q8BHJ5, Q3UMY5, Q8BX17 | 4.08 | 0.996383363472 |
| UP_SEQ_FEATURE | REPEAT:WD 4 | 6 | 1.63 | 0.241382970265 | Q8CEC6, Q8BHJ5, Q3UMY5, Q3UKJ7, Q9WVA3, Q8BX17 | 1.79 | 0.996383363472 |
| INTERPRO | IPR015943:WD40/YVTN_repeat-like_dom_sf | 8 | 2.17 | 0.246823849689 | Q58A65, Q8CEC6, Q8BHJ5, Q3UMY5, Q3UKJ7, Q9WVA3, P70206, Q8BX17 | 1.58 | 0.993986254296 |
| UP_SEQ_FEATURE | REPEAT:WD | 5 | 1.36 | 0.251857135933 | Q8CEC6, Q8BHJ5, Q3UMY5, Q9WVA3, Q8BX17 | 1.96 | 0.996383363472 |
| UP_KW_DOMAIN | KW-0853~WD repeat | 6 | 1.63 | 0.264231928135 | Q8CEC6, Q8BHJ5, Q3UMY5, Q3UKJ7, Q9WVA3, Q8BX17 | 1.73 | 0.675000892156 |
| UP_SEQ_FEATURE | REPEAT:WD 3 | 6 | 1.63 | 0.274463736188 | Q8CEC6, Q8BHJ5, Q3UMY5, Q3UKJ7, Q9WVA3, Q8BX17 | 1.71 | 0.996383363472 |
| INTERPRO | IPR036322:WD40_repeat_dom_sf | 7 | 1.90 | 0.280213276244 | Q58A65, Q8CEC6, Q8BHJ5, Q3UMY5, Q3UKJ7, Q9WVA3, Q8BX17 | 1.58 | 0.993986254296 |
| UP_SEQ_FEATURE | REPEAT:WD 1 | 6 | 1.63 | 0.28520631739 | Q8CEC6, Q8BHJ5, Q3UMY5, Q3UKJ7, Q9WVA3, Q8BX17 | 1.68 | 0.996383363472 |
| UP_SEQ_FEATURE | REPEAT:WD 2 | 6 | 1.63 | 0.28520631739 | Q8CEC6, Q8BHJ5, Q3UMY5, Q3UKJ7, Q9WVA3, Q8BX17 | 1.68 | 0.996383363472 |
| INTERPRO | IPR001680:WD40_rpt | 6 | 1.63 | 0.336677560361 | Q8CEC6, Q8BHJ5, Q3UMY5, Q3UKJ7, Q9WVA3, Q8BX17 | 1.56 | 0.993986254296 |
| SMART | SM00320:WD40 | 6 | 1.63 | 0.337421510604 | Q8CEC6, Q8BHJ5, Q3UMY5, Q3UKJ7, Q9WVA3, Q8BX17 | 1.56 | 1 |
| UP_SEQ_FEATURE | REPEAT:WD 7 | 4 | 1.09 | 0.340333550784 | Q8BHJ5, Q3UMY5, Q3UKJ7, Q8BX17 | 1.93 | 0.996383363472 |
| UP_SEQ_FEATURE | REPEAT:WD 5 | 5 | 1.36 | 0.386043595193 | Q8BHJ5, Q3UMY5, Q3UKJ7, Q9WVA3, Q8BX17 | 1.59 | 0.996383363472 |
| UP_SEQ_FEATURE | REPEAT:WD 6 | 4 | 1.09 | 0.48531701881 | Q8BHJ5, Q3UMY5, Q3UKJ7, Q8BX17 | 1.53 | 0.996383363472 |
| INTERPRO | IPR019775:WD40_repeat_CS | 3 | 0.82 | 0.596518969361 | Q8BHJ5, Q3UKJ7, Q8BX17 | 1.50 | 0.993986254296 |

Annotation Cluster 58 — Enrichment Score: 0.4577038497438003

| Category | Term | Count | % | PValue | Genes | Fold Enrichment | FDR |
|---|---|---|---|---|---|---|---|
| GOTERM_CC_DIRECT | GO:0016529~sarcoplasmic reticulum | 3 | 0.82 | 0.140279582474 | Q6PHZ2, Q64518, P70227 | 4.55 | 0.519154352231 |
| GOTERM_BP_DIRECT | GO:0006816~calcium ion transport | 3 | 0.82 | 0.331918370501 | Q6PHZ2, Q64518, P70227 | 2.53 | 0.982665222102 |
| KEGG_PATHWAY | mmu04020:Calcium signaling pathway | 4 | 1.09 | 0.909625646209 | P51432, Q6PHZ2, Q64518, P70227 | 0.74 | 0.909625646209 |

Annotation Cluster 59 — Enrichment Score: 0.38239255535430594

| Category | Term | Count | % | PValue | Genes | Fold Enrichment | FDR |
|---|---|---|---|---|---|---|---|
| UP_KW_DOMAIN | KW-0728~SH3 domain | 5 | 1.36 | 0.289286096638 | Q9QXZ0, O70145, Q3UND0, O08539, P97814 | 1.83 | 0.675000892156 |
| UP_SEQ_FEATURE | DOMAIN:SH3 | 5 | 1.36 | 0.338850063872 | Q9QXZ0, O70145, Q3UND0, O08539, P97814 | 1.70 | 0.996383363472 |
| INTERPRO | IPR001452:SH3_domain | 5 | 1.36 | 0.388059569205 | Q9QXZ0, O70145, Q3UND0, O08539, P97814 | 1.58 | 0.993986254296 |
| SMART | SM00326:SH3 | 4 | 1.09 | 0.546842486637 | O70145, Q3UND0, O08539, P97814 | 1.39 | 1 |
| INTERPRO | IPR036028:SH3-like_dom_sf | 4 | 1.09 | 0.588764265641 | O70145, Q3UND0, O08539, P97814 | 1.31 | 0.993986254296 |

Annotation Cluster 60 — Enrichment Score: 0.37087529785295403

| Category | Term | Count | % | PValue | Genes | Fold Enrichment | FDR |
|---|---|---|---|---|---|---|---|
| GOTERM_MF_DIRECT | GO:0005178~integrin binding | 5 | 1.36 | 0.0911332314 | P21956, O88839, Q99P91, O70309, Q61233 | 2.93 | 0.565201290897 |
| GOTERM_BP_DIRECT | GO:0007155~cell adhesion | 7 | 1.90 | 0.475947643016 | P21956, O88839, Q99P91, O70309, Q07797, P07356, P97814 | 1.26 | 0.982665222102 |

| | | | | | | | |
|---|---|---|---|---|---|---|---|
| UP_KW_DOMAIN | KW-0245~EGF-like domain | 3 | 0.82 | 0.842421508616 | P21956, O88839, O70309 | 0.91 | 1 |
| UP_KW_BIOLOGICAL_PROCESS | KW-0130~Cell adhesion | 6 | 1.63 | 0.898945601356 | P21956, O88839, Q8BTI9, O70309, Q07797, P97814 | 0.76 | 0.967032967033 |
| Annotation Cluster 61 | Enrichment Score: 0.3557309238633206 | | | | | | |
| Category | Term | Count | % | PValue | Genes | Fold Enrichment | FDR |
| UP_SEQ_FEATURE | LIPID:S-palmitoyl cysteine | 7 | 1.90 | 0.226343231679 | Q9CQS8, Q9CQW9, O09044, Q99J93, P35564, P62281, Q8VCF0 | 1.70 | 0.996383363472 |
| UP_KW_PTM | KW-0564~Palmitate | 9 | 2.45 | 0.597570019419 | Q9CQS8, Q9CQW9, Q60766, O09044, O35114, Q99J93, P35564, P62281, Q8VCF0 | 1.08 | 0.730800477358 |
| UP_KW_PTM | KW-0449~Lipoprotein | 19 | 5.16 | 0.63336041371 | Q9CQS8, Q8R1F1, Q9CQW9, P03975, Q60766, P10810, Q99J93, Q9Z0E6, P35564, P62281, Q8VCF0, Q99JI6, P63213, O09044, O35114, P61027, P62835, Q3TGW2 | 0.99 | 0.730800477358 |
| Annotation Cluster 62 | Enrichment Score: 0.35549147337435677 | | | | | | |
| Category | Term | Count | % | PValue | Genes | Fold Enrichment | FDR |
| KEGG_PATHWAY | mmu04012:ErbB signaling pathway | 4 | 1.09 | 0.261463354402 | Q6PHZ2, Q8BTI9, P31750, Q61036 | 2.24 | 0.624058760979 |
| GOTERM_MF_DIRECT | GO:0106310~protein serine kinase activity | 6 | 1.63 | 0.350556775497 | Q6PHZ2, Q8R4K2, Q8BTI9, Q99KH8, P31750, Q61036 | 1.54 | 0.959821428571 |
| KEGG_PATHWAY | mmu04510:Focal adhesion | 6 | 1.63 | 0.428553315038 | Q99JI6, Q8BTI9, O70309, P62835, P31750, Q61036 | 1.40 | 0.84049603647 |
| KEGG_PATHWAY | mmu04660:T cell receptor signaling pathway | 4 | 1.09 | 0.478111082298 | Q8BTI9, P31750, Q76MZ3, Q61036 | 1.54 | 0.864534806435 |
| KEGG_PATHWAY | mmu04360:Axon guidance | 5 | 1.36 | 0.53477966367 | Q6PHZ2, Q8BTI9, P18760, P70206, Q61036 | 1.30 | 0.865215634447 |
| KEGG_PATHWAY | mmu04810:Regulation of actin cytoskeleton | 5 | 1.36 | 0.733117206276 | Q8BTI9, O70309, P18760, P31750, Q61036 | 1.01 | 0.884615384615 |
| Annotation Cluster 63 | Enrichment Score: 0.349515564657348 | | | | | | |
| Category | Term | Count | % | PValue | Genes | Fold Enrichment | FDR |
| UP_KW_DOMAIN | KW-0727~SH2 domain | 3 | 0.82 | 0.406779423282 | P42225, Q9WVL2, Q9JM90 | 2.14 | 0.854236788893 |
| UP_SEQ_FEATURE | DOMAIN:SH2 | 3 | 0.82 | 0.437295140614 | P42225, Q9WVL2, Q9JM90 | 2.02 | 0.996383363472 |
| INTERPRO | IPR000980:SH2 | 3 | 0.82 | 0.462615066229 | P42225, Q9WVL2, Q9JM90 | 1.92 | 0.993986254296 |
| INTERPRO | IPR036860:SH2_dom_sf | 3 | 0.82 | 0.485941893074 | P42225, Q9WVL2, Q9JM90 | 1.84 | 0.993986254296 |
| Annotation Cluster 64 | Enrichment Score: 0.29111328393460056 | | | | | | |
| Category | Term | Count | % | PValue | Genes | Fold Enrichment | FDR |
| GOTERM_CC_DIRECT | GO:0016020~membrane | 74 | 20.11 | 0.044206389619 | Q9QXZ0, Q8BVE3, Q8K3J1, Q6GV12, O70370, O70493, Q07797, Q61033, Q9QY76, P70227, Q9JHJ3, P36552, O88456, P47753, P61027, P62835, P07356, P49813, P51432, P26151, Q64518, Q791T5, P97797, O35682, P35564, Q8VD58, Q8BVL3, Q58A65, Q07813, O35114, Q3UMY5, Q9CWG1, Q8BTY2, Q9CQS8, Q811U4, Q80X80, Q60766, Q68FD5, Q9CPQ3, P31750, P70206, O88839, P36371, P58681, Q00612, Q7TMY8, P08101, Q8K1T1, O35309, Q9QYB1, Q8VI94, Q99P91, P57716, Q3UVL4, O70145, Q791V5, Q8K4Q8, Q62393, O08539, Q6PA06, P59017, Q8VCF0, O88829, O09159, Q99JI6, P21958, P17182, Q9EPE9, O88983, Q921I1, Q9JIY5, Q80WJ7 | 1.22 | 0.303831217539 |
| UP_SEQ_FEATURE | TOPO_DOM:Cytoplasmic | 46 | 12.50 | 0.616400503237 | Q9CQS8, P61804, Q811U4, Q80X80, Q9CQW9, O70579, Q6R5N8, Q6GV12, Q9CPQ3, P70206, Q9QY76, P70227, O88839, Q8VCH8, P36371, Q9JHJ3, P58681, P08101, Q8K1T1, Q99P91, P57716, Q64518, P26151, Q791T5, Q99J93, Q791V5, Q8K4Q8, P97797, P35564, Q6PA06, Q8VCF0, Q8VD58, Q9D8V0, O88829, Q8BTJ4, P30993, Q8BXQ2, O35114, P21958, O70309, Q9EPE9, O88983, Q8BTY2, Q80WJ7, Q8BG07 | 0.99 | 0.996383363472 |
| UP_KW_DOMAIN | KW-1133~Transmembrane helix | 70 | 19.02 | 0.721746376041 | O70579, Q6GV12, P48771, Q61033, Q9QY76, P70227, Q9JHJ3, Q8BH59, Q9CXV1, Q9D0L7, P26151, Q64518, Q791T5, P97797, O35682, P35564, P19783, Q8VD58, Q8BVL3, P30993, Q58A65, Q8BXQ2, Q07813, O35114, Q3UMY5, Q9CWG1, Q8BTY2, Q9CQS8, P61804, Q811U4, Q62425, Q80X80, Q9CQW9, Q6R5N8, Q9QXX4, Q9CPQ3, P70206, Q9CPQ1, O88839, P36371, P58681, P00405, Q7TMY8, P08101, Q8K1T1, Q9QYB1, Q99P91, P57716, Q99J93, Q791V5, Q8K4Q8, Q62393, Q6PA06, P59017, Q8VCF0, Q9D8V0, O88829, Q99JR1, Q8BTJ4, O09159, O09111, P21958, O70309, Q9EPE9, O88983, P60603, Q9JIY5, Q8BG07, Q80WJ7 | 0.96 | 1 |
| UP_KW_DOMAIN | KW-0812~Transmembrane | 70 | 19.02 | 0.938882565295 | O70579, Q6GV12, P48771, Q61033, Q9QY76, P70227, Q9JHJ3, Q8BH59, Q9CXV1, Q9D0L7, P26151, Q64518, Q791T5, P97797, O35682, P35564, P19783, Q8VD58, Q8BVL3, P30993, Q58A65, Q8BXQ2, Q07813, O35114, Q3UMY5, Q9CWG1, Q8BTY2, Q9CQS8, P61804, Q811U4, Q62425, Q80X80, Q9CQW9, Q6R5N8, Q9QXX4, Q9CPQ3, P70206, Q9CPQ1, O88839, P36371, P58681, P00405, Q7TMY8, P08101, Q8K1T1, Q9QYB1, Q99P91, P57716, Q99J93, Q791V5, Q8K4Q8, Q62393, Q6PA06, P59017, Q8VCF0, Q9D8V0, O88829, Q99JR1, Q8BTJ4, O09159, O09111, P21958, O70309, Q9EPE9, O88983, P60603, Q9JIY5, Q8BG07, Q80WJ7 | 0.88 | 1 |

| | | | | | | | |
|---|---|---|---|---|---|---|---|
| UP_KW_CELLULAR_COMPONENT | KW-0472~Membrane | 133 | 36.14 | 0.970488188636 | P07901, B9EKI3, Q9QXZ0, Q8BVE3, P18760, O70493, Q61033, Q3U2P1, O88456, P61027, P20152, P26151, Q791T5, Q80V94, Q8VD58, Q8BVL3, P30993, Q8BXQ2, P63213, O35114, Q91YY4, Q922Q1, Q8BMS1, Q9CWG1, Q8BTY2, Q9CQS8, Q811U4, Q9CPQ3, Q64521, Q9CPQ1, P70206, Q62465, O88839, P58681, Q7TMY8, P97450, Q61545, Q99J93, Q6PA06, P59017, Q9D8V0, O88829, Q99JR1, Q76MZ3, O09159, Q99JI6, Q9CQA3, O08585, E9Q3L2, P56391, Q80WJ7, O70579, P29452, Q8K3J1, Q6GV12, Q99JY0, P48771, Q3UIA2, Q62167, Q61233, Q9QY76, P70227, Q9JHJ3, Q8VCH8, O09044, P63242, P62835, Q8BH59, P51432, Q9CXV1, Q64518, Q9D0L7, P97797, P11499, O35682, P35564, P19783, Q9Z0E6, P19536, Q58A65, Q9R0E1, Q07813, Q9CYG7, Q91VW5, Q3UMY5, P61804, Q6PHZ2, Q62425, Q80X80, Q9CQW9, Q60766, P10810, Q6R5N8, Q68FD5, Q9QXX4, P31750, Q91XB0, Q3UM29, P36371, Q00612, P00405, P08101, Q8K1T1, Q9DC70, Q99KH8, Q9QYB1, P62908, Q99P91, P57716, Q8R1F1, Q61107, P03975, Q791V5, Q8K4Q8, Q62393, O08539, Q8VCF0, P97814, Q8VBT9, Q8BTJ4, P21956, O09111, P21958, O70309, P17182, Q9EPE9, O88983, P60603, Q9JIY5, Q8BG07 | 0.90 | 0.970488188636 |
| UP_SEQ_FEATURE | TRANSMEM:Helical | 63 | 17.12 | 0.999968050248 | O70579, Q6GV12, P48771, Q61033, Q9QY76, P70227, Q9JHJ3, Q9CXV1, Q9D0L7, P26151, Q64518, Q791T5, P97797, O35682, P35564, P19783, Q8VD58, Q8BVL3, P30993, Q58A65, Q8BXQ2, Q07813, O35114, Q3UMY5, Q9CWG1, Q8BTY2, Q9CQS8, P61804, Q62425, Q80X80, Q9CQW9, Q6R5N8, Q9CPQ3, P70206, Q9CPQ1, O88839, P36371, P58681, P00405, Q7TMY8, P08101, Q8K1T1, Q99P91, P57716, Q99J93, Q8K4Q8, Q62393, Q6PA06, P59017, Q8VCF0, Q9D8V0, O88829, Q99JR1, Q8BTJ4, O09159, O09111, P21958, O70309, Q9EPE9, P60603, Q9JIY5, Q80WJ7 | 0.68 | 0.999968050248 |

Annotation Cluster 65 — Enrichment Score: 0.2882646271857498

| Category | Term | Count | % | PValue | Genes | Fold Enrichment | FDR |
|---|---|---|---|---|---|---|---|
| GOTERM_BP_DIRECT | GO:0035556~intracellular signal transduction | 9 | 2.45 | 0.089365785081 | P51432, Q8R4K2, Q07813, O35682, Q99KH8, Q9JIY5, P31750, Q62167, Q61036 | 1.97 | 0.799227132233 |
| KEGG_PATHWAY | mmu05142:Chagas disease | 5 | 1.36 | 0.173165362687 | P51432, Q8R4K2, Q8BTI9, P31750, Q76MZ3 | 2.29 | 0.492256592806 |
| GOTERM_MF_DIRECT | GO:0016301~kinase activity | 4 | 1.09 | 0.254976191155 | Q8R4K2, Q8BTI9, E9Q3L2, P31750 | 2.28 | 0.959821428571 |
| KEGG_PATHWAY | mmu04012:ErbB signaling pathway | 4 | 1.09 | 0.261463354402 | Q6PHZ2, Q8BTI9, P31750, Q61036 | 2.24 | 0.624058760979 |
| KEGG_PATHWAY | mmu05135:Yersinia infection | 5 | 1.36 | 0.323587391577 | Q8R4K2, P29452, Q8BTI9, Q3UND0, P31750 | 1.73 | 0.715611671076 |
| GOTERM_MF_DIRECT | GO:0106310~protein serine kinase activity | 6 | 1.63 | 0.350556775497 | Q6PHZ2, Q8R4K2, Q8BTI9, Q99KH8, P31750, Q61036 | 1.54 | 0.959821428571 |
| GOTERM_MF_DIRECT | GO:0044022~histone H3S28 kinase activity | 4 | 1.09 | 0.500728406018 | Q8R4K2, Q99KH8, P31750, Q61036 | 1.48 | 0.959821428571 |
| GOTERM_MF_DIRECT | GO:0044023~histone H4S1 kinase activity | 4 | 1.09 | 0.500728406018 | Q8R4K2, Q99KH8, P31750, Q61036 | 1.48 | 0.959821428571 |
| GOTERM_MF_DIRECT | GO:0004676~3-phosphoinositide-dependent protein kinase activity | 4 | 1.09 | 0.500728406018 | Q8R4K2, Q99KH8, P31750, Q61036 | 1.48 | 0.959821428571 |
| GOTERM_MF_DIRECT | GO:0044024~histone H2AS1 kinase activity | 4 | 1.09 | 0.500728406018 | Q8R4K2, Q99KH8, P31750, Q61036 | 1.48 | 0.959821428571 |
| GOTERM_MF_DIRECT | GO:0004677~DNA-dependent protein kinase activity | 4 | 1.09 | 0.500728406018 | Q8R4K2, Q99KH8, P31750, Q61036 | 1.48 | 0.959821428571 |
| GOTERM_MF_DIRECT | GO:0044025~histone H2BS14 kinase activity | 4 | 1.09 | 0.500728406018 | Q8R4K2, Q99KH8, P31750, Q61036 | 1.48 | 0.959821428571 |
| GOTERM_MF_DIRECT | GO:0004694~eukaryotic translation initiation factor 2alpha kinase activity | 4 | 1.09 | 0.500728406018 | Q8R4K2, Q99KH8, P31750, Q61036 | 1.48 | 0.959821428571 |
| GOTERM_MF_DIRECT | GO:0004711~ribosomal protein S6 kinase activity | 4 | 1.09 | 0.500728406018 | Q8R4K2, Q99KH8, P31750, Q61036 | 1.48 | 0.959821428571 |
| GOTERM_MF_DIRECT | GO:0072354~histone H3T3 kinase activity | 4 | 1.09 | 0.500728406018 | Q8R4K2, Q99KH8, P31750, Q61036 | 1.48 | 0.959821428571 |
| GOTERM_MF_DIRECT | GO:0072371~histone H2AS121 kinase activity | 4 | 1.09 | 0.500728406018 | Q8R4K2, Q99KH8, P31750, Q61036 | 1.48 | 0.959821428571 |
| GOTERM_MF_DIRECT | GO:0072518~Rho-dependent protein serine/threonine kinase activity | 4 | 1.09 | 0.500728406018 | Q8R4K2, Q99KH8, P31750, Q61036 | 1.48 | 0.959821428571 |
| GOTERM_MF_DIRECT | GO:0140823~histone H2BS36 kinase activity | 4 | 1.09 | 0.500728406018 | Q8R4K2, Q99KH8, P31750, Q61036 | 1.48 | 0.959821428571 |
| GOTERM_MF_DIRECT | GO:0140855~histone H3S57 kinase activity | 4 | 1.09 | 0.500728406018 | Q8R4K2, Q99KH8, P31750, Q61036 | 1.48 | 0.959821428571 |
| GOTERM_MF_DIRECT | GO:1990244~histone H2AT120 kinase activity | 4 | 1.09 | 0.500728406018 | Q8R4K2, Q99KH8, P31750, Q61036 | 1.48 | 0.959821428571 |
| GOTERM_MF_DIRECT | GO:0035175~histone H3S10 kinase activity | 4 | 1.09 | 0.500728406018 | Q8R4K2, Q99KH8, P31750, Q61036 | 1.48 | 0.959821428571 |
| GOTERM_MF_DIRECT | GO:0035979~histone H2AXS139 kinase activity | 4 | 1.09 | 0.500728406018 | Q8R4K2, Q99KH8, P31750, Q61036 | 1.48 | 0.959821428571 |
| GOTERM_MF_DIRECT | GO:0035402~histone H3T11 kinase activity | 4 | 1.09 | 0.505348889328 | Q8R4K2, Q99KH8, P31750, Q61036 | 1.47 | 0.959821428571 |
| GOTERM_MF_DIRECT | GO:0035403~histone H3T6 kinase activity | 4 | 1.09 | 0.505348889328 | Q8R4K2, Q99KH8, P31750, Q61036 | 1.47 | 0.959821428571 |
| GOTERM_MF_DIRECT | GO:0140857~histone H3T45 kinase activity | 4 | 1.09 | 0.506900423794 | Q8R4K2, Q99KH8, P31750, Q61036 | 1.48 | 0.959821428571 |
| GOTERM_MF_DIRECT | GO:0004679~AMP-activated protein kinase activity | 4 | 1.09 | 0.513614782324 | Q8R4K2, Q99KH8, P31750, Q61036 | 1.46 | 0.959821428571 |
| GOTERM_MF_DIRECT | GO:0004672~protein kinase activity | 4 | 1.09 | 0.5656656643 | Q6PHZ2, P31750, Q7TT37, Q61036 | 1.35 | 0.959821428571 |
| GOTERM_MF_DIRECT | GO:0004674~protein serine/threonine kinase activity | 5 | 1.36 | 0.601920388605 | Q6PHZ2, Q8R4K2, Q99KH8, P31750, Q61036 | 1.19 | 0.959821428571 |
| GOTERM_BP_DIRECT | GO:0006468~protein phosphorylation | 4 | 1.09 | 0.702923956595 | Q6PHZ2, P18760, Q99KH8, P31750 | 1.10 | 0.982665222102 |
| SMART | SM00220:S_TKc | 5 | 1.36 | 0.803138288506 | Q6PHZ2, Q8R4K2, Q99KH8, P31750, Q61036 | 0.91 | 1 |
| INTERPRO | IPR011009:Kinase-like_dom_sf | 7 | 1.90 | 0.813855905432 | Q6PHZ2, Q8R4K2, Q8BTI9, E9Q3L2, Q99KH8, P31750, Q61036 | 0.87 | 0.993986254296 |
| UP_KW_MOLECULAR_FUNCTION | KW-0418~Kinase | 9 | 2.45 | 0.904023289001 | Q6PHZ2, Q8R4K2, Q8BTI9, Q3TRM8, E9Q3L2, Q99KH8, P31750, Q61036, Q9WTP6 | 0.77 | 0.946666666667 |
| UP_KW_MOLECULAR_FUNCTION | KW-0723~Serine/threonine-protein kinase | 5 | 1.36 | 0.906728866993 | Q6PHZ2, Q8R4K2, Q99KH8, P31750, Q61036 | 0.74 | 0.946666666667 |
| INTERPRO | IPR017441:Protein_kinase_ATP_BS | 4 | 1.09 | 0.922388476457 | Q6PHZ2, Q99KH8, P31750, Q61036 | 0.71 | 0.993986254296 |
| GOTERM_BP_DIRECT | GO:0006338~chromatin remodeling | 5 | 1.36 | 0.926212045478 | Q8R4K2, Q7TMY8, Q99KH8, P31750, Q61036 | 0.70 | 0.982665222102 |
| UP_SEQ_FEATURE | DOMAIN:Protein kinase | 5 | 1.36 | 0.932820989105 | Q6PHZ2, Q8R4K2, Q99KH8, P31750, Q61036 | 0.69 | 0.996383363472 |
| INTERPRO | IPR000719:Prot_kinase_dom | 5 | 1.36 | 0.939509874897 | Q6PHZ2, Q8R4K2, Q99KH8, P31750, Q61036 | 0.68 | 0.993986254296 |
| INTERPRO | IPR008271:Ser/Thr_kinase_AS | 3 | 0.82 | 0.952679676665 | Q6PHZ2, P31750, Q61036 | 0.63 | 0.993986254296 |

Annotation Cluster 66 — Enrichment Score: 0.2784334872637739

| Category | Term | Count | % | PValue | Genes | Fold Enrichment | FDR |
|---|---|---|---|---|---|---|---|
| INTERPRO | IPR035892:C2_domain_sf | 4 | 1.09 | 0.435599462009 | P51432, Q80X80, Q8BTI9, Q8BRN9 | 1.64 | 0.993986254296 |
| UP_SEQ_FEATURE | DOMAIN:C2 | 3 | 0.82 | 0.551642540528 | P51432, Q80X80, Q8BRN9 | 1.62 | 0.996383363472 |
| INTERPRO | IPR000008:C2_dom | 3 | 0.82 | 0.608071226801 | P51432, Q80X80, Q8BRN9 | 1.46 | 0.993986254296 |

Annotation Cluster 67 — Enrichment Score: 0.1426126899876609

| Category | Term | Count | % | PValue | Genes | Fold Enrichment | FDR |
|---|---|---|---|---|---|---|---|
| GOTERM_CC_DIRECT | GO:0031514~motile cilium | 3 | 0.82 | 0.466292778966 | O88839, O08749, Q8BTU1 | 1.91 | 0.935205183585 |
| UP_KW_CELLULAR_COMPONENT | KW-0282~Flagellum | 3 | 0.82 | 0.853658827956 | O88839, O08749, Q8BTU1 | 0.89 | 0.864864864865 |
| UP_KW_CELLULAR_COMPONENT | KW-0969~Cilium | 4 | 1.09 | 0.938034699943 | O88839, Q8VIM6, O08749, Q8BTU1 | 0.67 | 0.938034699943 |

Annotation Cluster 68 — Enrichment Score: 0.10086009743083

| Category | Term | Count | % | PValue | Genes | Fold Enrichment | FDR |
|---|---|---|---|---|---|---|---|
| UP_SEQ_FEATURE | DOMAIN:PH | 4 | 1.09 | 0.715168173479 | Q8R1F1, Q3UND0, Q9JM90, P31750 | 1.08 | 0.996383363472 |
| INTERPRO | IPR001849:PH_domain | 4 | 1.09 | 0.727598192556 | Q8R1F1, Q3UND0, Q9JM90, P31750 | 1.06 | 0.993986254296 |
| INTERPRO | IPR011993:PH-like_dom_sf | 5 | 1.36 | 0.857991111738 | Q8R1F1, Q3UND0, Q9JM90, P31750, Q8BVL3 | 0.82 | 0.993986254296 |
| SMART | SM00233:PH | 3 | 0.82 | 0.884661302982 | Q3UND0, Q9JM90, P31750 | 0.81 | 1 |

Annotation Cluster 69 — Enrichment Score: 0.01289806081616931

| Category | Term | Count | % | PValue | Genes | Fold Enrichment | FDR |
|---|---|---|---|---|---|---|---|
| UP_SEQ_FEATURE | CARBOHYD:N-linked (GlcNAc...) asparagine | 45 | 12.23 | 0.897072815104 | Q8BFR4, P10810, Q6R5N8, O70370, P97821, P18242, Q07797, P70206, O08795, O88839, Q9JHJ3, Q9WVJ3, P58681, P08101, O09043, Q8K1T1, Q99P91, P57716, Q9Z0M5, P03975, P26151, Q8K4Q8, P97797, Q8VD58, Q9D8V0, O88829, Q6P5E4, Q8BTJ4, P30993, O09159, P21956, Q9R0E1, Q8BXQ2, O35114, Q9DCQ7, Q8VIM6, O70309, P08113, Q9EPE9, Q921I1, O88188, Q8BTY2, Q8BG07 | 0.87 | 0.996383363472 |
| UP_KW_DOMAIN | KW-0732~Signal | 48 | 13.04 | 0.989881129923 | P27773, Q8BFR4, Q6PHZ2, P10810, Q6R5N8, Q6GV12, O70370, P97821, P18242, Q07797, P70206, O08795, O88839, P36371, Q9JHJ3, Q9WVJ3, P58681, P08101, O09043, Q8K1T1, Q8CGK3, Q99P91, P57716, Q9Z0M5, P03975, Q64518, P08905, P26151, P97797, P35564, Q8VD58, Q6P5E4, Q8BTJ4, O09159, P21956, Q9R0E1, Q8BXQ2, Q9DCQ7, Q8VIM6, O70309, P08113, Q8BHG1, Q921I1, Q9JK81, O88188, Q9CWG1 | 0.78 | 1 |
| UP_KW_PTM | KW-0325~Glycoprotein | 56 | 15.22 | 0.999995987075 | Q8BFR4, O70370, P97821, P18242, Q07797, O08795, Q9JHJ3, P25206, O09043, P20152, Q9Z0M5, P26151, P97797, P11499, Q8VD58, Q61881, P30993, Q9R0E1, Q8BXQ2, O35114, Q9DCQ7, Q8BTY2, P10810, P10854, Q6R5N8, Q9R1P4, P10853, P31750, P70206, O88839, Q9WVJ3, P58681, P08101, Q8K1T1, Q99P91, P57716, P03975, Q8K4Q8, Q9D8V0, O88829, Q6P5E4, Q8BTJ4, O09159, P21956, Q8VIM6, O70309, P08113, Q9EPE9, Q921I1, O88188, Q8BG07 | 0.64 | 0.999995987075 |
| UP_KW_PTM | KW-1015~Disulfide bond | 42 | 11.41 | 0.999997140489 | P27773, P10639, P10810, Q9CQM5, O08749, O70370, P97821, P18242, Q07797, P31750, P70206, Q9WTP6, P70227, O08795, O88839, P08101, O09043, P57716, P03975, P08905, P26151, Q8K4Q8, P97797, P35564, O88829, Q8BTJ4, P30993, O09159, P21956, Q05816, Q9R0E1, Q8BXQ2, O35114, O70309, P08113, P56394, Q921I1, O88188, P56391, Q8BTY2, Q8BG07 | 0.59 | 0.999997140489 |

Annotation Cluster 70 — Enrichment Score: 0.009504556357113437

| Category | Term | Count | % | PValue | Genes | Fold Enrichment | FDR |
|---|---|---|---|---|---|---|---|
| UP_KW_MOLECULAR_FUNCTION | KW-0010~Activator | 8 | 2.17 | 0.941773752681 | P42225, P10639, Q91VR5, P54103, Q8BHJ5, P70670, Q9WVL2, P0DOV2 | 0.70 | 0.946666666667 |
| UP_KW_BIOLOGICAL_PROCESS | KW-0805~Transcription regulation | 22 | 5.98 | 0.996484657569 | Q921F2, P62908, P10639, Q61545, P54103, Q91VR5, Q9JKB3, B9EKI3, Q8BVK9, P70670, Q9WVL2, Q62167, Q8BRN9, P0DOV2, O35326, P42225, Q8BHJ5, Q8VDF2, O35892, Q64152, P83940, Q8BXC6 | 0.65 | 0.996484657569 |
| UP_KW_BIOLOGICAL_PROCESS | KW-0804~Transcription | 22 | 5.98 | 0.997858892737 | Q921F2, P62908, P10639, Q61545, P54103, Q91VR5, Q9JKB3, B9EKI3, Q8BVK9, P70670, Q9WVL2, Q62167, Q8BRN9, P0DOV2, O35326, P42225, Q8BHJ5, Q8VDF2, O35892, Q64152, P83940, Q8BXC6 | 0.63 | 0.997858892737 |
