## Supplemental Table 6 for "Effects of polystyrene and polylactide nanoparticles on macrophages under a repeated exposure mode"

Supplementary Table 6: Results of the pathway analysis by the David tool on the proteins modulated in response to PLA particles

Annotation Cluster 1 — Enrichment Score: 10.293582570566864

| Category | Term | Count | PValue | Genes | Fold Enrichment | FDR |
|---|---|---|---|---|---|---|
| GOTERM_MF_DIRECT | GO:0005524~ATP binding | 68 | 3.905E-19 | P11928, Q8R4K2, Q63932, B2RQC6, Q8BTI9, P38647, Q07832, Q8K1J6, Q99J87, P70227, Q9WTP6, P12382, Q9EPU0, Q62095, Q3THS6, P09411, Q2TBE6, Q91WQ3, Q3TRM8, Q9WTM5, Q9Z2I8, E9QAM5, Q9ER72, Q61881, Q9JIK5, P17710, Q8BP47, Q9Z2I9, Q5NC05, Q9QXG4, P63017, E9Q555, Q8VCW8, Q61699, Q6P9P6, Q91VR5, Q61656, Q6PHZ2, Q501J6, Q9R0N0, O08528, P36371, P49718, Q9CZD3, O54984, Q91V92, Q6A028, Q9QUJ7, P82343, Q99KH8, O88351, Q8CGK3, Q9CZP5, Q9ESL4, Q60930, Q9EQP2, Q61024, P33175, P21958, Q9QZ08, P83741, E9Q3L2, Q9WUA3, Q9DBY8, Q922S8, Q6P542, E9PVX6 | 3.48 | 9.9585E-17 |
| UP_KW_LIGAND | KW-0547~Nucleotide-binding | 81 | 3.139E-07 | P11928, Q8R4K2, Q63932, B2RQC6, Q8BTI9, P38647, Q07832, Q8K1J6, Q99J87, P70227, Q9WTP6, P12382, Q8VEH6, Q9EPU0, Q62095, Q3THS6, P09411, P61027, Q8BTZ7, Q2TBE6, Q9DBC7, Q91WQ3, Q3TRM8, Q9WTM5, Q9Z0E6, Q9Z2I8, E9QAM5, Q3UX10, Q9ER72, Q61881, Q9JIK5, P17710, Q8BP47, Q9Z2I9, Q9CZE3, Q5NC05, Q8BH04, Q9QXG4, P63017, E9Q555, Q8VCW8, Q61699, Q6P9P6, Q91VR5, Q61656, Q6PHZ2, Q60766, Q501J6, Q9R0N0, O08528, P59325, P36371, P49718, Q9CZD3, O54984, Q91V92, Q9QUJ7, P42208, Q99KH8, O88351, Q8CGK3, Q9CZP5, Q3TBT3, Q9ESL4, Q60930, Q61107, Q9EQP2, Q6PA06, Q61024, P33175, P21958, Q9QZ08, P83741, E9Q3L2, Q9WUA3, Q9DBY8, Q922S8, Q6P542, P60766, E9PVX6 | 1.65 | 1.0045E-05 |
| UP_KW_LIGAND | KW-0067~ATP-binding | 66 | 1.073E-06 | P11928, Q8R4K2, Q63932, B2RQC6, Q8BTI9, P38647, Q07832, Q8K1J6, Q99J87, P70227, Q9WTP6, P12382, Q9EPU0, Q62095, Q3THS6, P09411, Q2TBE6, Q91WQ3, Q3TRM8, Q9WTM5, Q9Z2I8, E9QAM5, Q9ER72, Q61881, Q9JIK5, P17710, Q8BP47, Q9Z2I9, Q5NC05, Q9QXG4, P63017, E9Q555, Q8VCW8, Q61699, Q6P9P6, Q91VR5, Q61656, Q6PHZ2, Q501J6, Q9R0N0, O08528, P36371, P49718, Q9CZD3, O54984, Q91V92, Q9QUJ7, Q99KH8, O88351, Q8CGK3, Q9CZP5, Q9ESL4, Q60930, Q9EQP2, Q61024, P33175, P21958, Q9QZ08, P83741, E9Q3L2, Q9WUA3, Q9DBY8, Q922S8, Q6P542, E9PVX6 | 1.75 | 1.7176E-05 |

Annotation Cluster 2 — Enrichment Score: 7.793905474589093

| Category | Term | Count | PValue | Genes | Fold Enrichment | FDR |
|---|---|---|---|---|---|---|
| GOTERM_BP_DIRECT | GO:0045087~innate immune response | 29 | 6.217E-11 | Q61510, P11928, Q8R4K2, Q91VR5, Q60766, P10810, Q6R5N8, Q8BVK9, Q91XB0, Q99J87, P0DOV2, O88839, A1L314, O35309, Q8R2Q8, P98086, O35405, Q3TBT3, P26151, Q99J93, Q9R002, P97814, Q9JIK5, P17710, Q9D906, Q9QZ08, O88188, O35658, Q8BG07 | 4.59 | 7.5077E-08 |
| UP_KW_BIOLOGICAL_PROCESS | KW-0399~Innate immunity | 30 | 2.362E-09 | Q61510, P11928, Q8R4K2, Q91VR5, Q9CQW9, Q60766, P10810, Q6R5N8, Q8BVK9, Q99J87, P0DOV2, A1L314, O35309, Q8R2Q8, P98086, O35405, Q3TBT3, P26151, Q61107, Q99J93, Q9R002, Q9Z0E6, P97814, P54987, Q9JIK5, P17710, Q9QZ08, O88188, O35658, Q8BG07 | 3.68 | 2.0311E-07 |
| UP_KW_BIOLOGICAL_PROCESS | KW-0391~Immunity | 41 | 2.828E-05 | Q61510, P06339, P11928, Q8R4K2, Q91VR5, Q9CQW9, Q60766, P10810, Q6R5N8, Q8BVK9, Q501J6, Q99J87, P0DOV2, P36371, A1L314, Q91YX0, O35309, Q8R2Q8, P98086, O35405, Q3TBT3, Q9EQH2, P26151, Q61107, Q99J93, Q9R002, Q9Z0E6, P42082, P14426, P97814, P54987, Q9JIK5, P17710, P01902, P01901, P21958, P01900, Q9QZ08, O88188, O35658, E9Q555, Q8BG07 | 1.99 | 0.00081068 |

Annotation Cluster 3 — Enrichment Score: 6.771661314476257

| Category | Term | Count | PValue | Genes | Fold Enrichment | FDR |
|---|---|---|---|---|---|---|
| UP_KW_CELLULAR_COMPONENT | KW-0496~Mitochondrion | 62 | 1.371E-11 | P11928, P50171, Q9CZW5, P38647, Q8K1J6, Q9WTP6, Q9DBL1, P36552, Q8BY71, P62075, P09411, Q8BGH2, Q8BH59, Q2TBE6, Q9CXV1, Q91YM4, Q99LX0, Q9Z2I8, Q8CG72, Q99KR3, P54987, Q9JIK5, P17710, P19536, O35435, O55022, Q9Z2I9, Q9CYG7, Q9CZE3, Q99KI0, Q8BH04, Q8VCW8, Q8K411, Q91VR5, Q62425, Q60766, O08528, Q9QXX4, Q9CPQ1, Q62465, P47738, Q9CZD3, Q8BTX9, Q9QUJ7, Q9JM90, P45952, Q8CGK3, Q9QYB1, Q9CZP5, Q3TBT3, Q60930, P97333, Q791V5, Q8WTY4, Q9CQV7, P11352, Q9CQA3, P38060, Q78IK4, Q9JK81, P29758, O35658 | 2.57 | 2.2629E-10 |
| GOTERM_CC_DIRECT | GO:0005759~mitochondrial matrix | 18 | 8.987E-08 | Q8K411, Q91YM4, P50171, Q99LX0, P38647, Q8CG72, Q99KR3, Q9DBL1, P47738, Q99KI0, P38060, P09411, P29758, P45952, Q9JK81, Q8BH04, O35658, Q8CGK3 | 5.19 | 3.017E-06 |
| UP_KW_DOMAIN | KW-0809~Transit peptide | 25 | 7.143E-06 | Q8K411, Q8VCW8, P38647, Q8K1J6, Q9DBL1, P47738, P36552, Q9CZD3, P45952, Q8CGK3, Q9CXV1, Q91YM4, Q9Z2I8, Q8C0I1, P19536, Q9CQA3, O35435, Q9Z2I9, Q99KI0, P38060, Q78IK4, P29758, Q9JK81, Q8BH04, O35658 | 2.86 | 0.00013572 |
| UP_SEQ_FEATURE | TRANSIT:Mitochondrion | 23 | 9.304E-05 | Q8K411, Q8VCW8, Q9CXV1, Q91YM4, P38647, Q8K1J6, Q9Z2I8, Q9DBL1, P19536, Q9CQA3, P47738, P36552, Q9CZD3, Q9Z2I9, Q99KI0, P38060, Q78IK4, P29758, P45952, Q9JK81, Q8BH04, O35658, Q8CGK3 | 2.59 | 0.02122155 |

Annotation Cluster 4 — Enrichment Score: 6.2541465229420625

| Category | Term | Count | PValue | Genes | Fold Enrichment | FDR |
|---|---|---|---|---|---|---|
| KEGG_PATHWAY | mmu04142:Lysosome | 17 | 2.261E-07 | Q9Z0M5, P11438, Q920I9, Q8BVE3, P20060, O70370, P97821, P18242, P17439, Q80V94, O09159, O35643, O35114, O08585, O88512, P29416, O89023 | 5.02 | 1.2391E-05 |
| GOTERM_CC_DIRECT | GO:0005764~lysosome | 18 | 8.721E-07 | Q9Z0M5, Q60766, P11438, P20060, Q80TY0, O70370, P97821, P18242, P17439, O09159, Q9WVJ3, O35114, P02802, Q9D1L9, P61027, P29416, O89023, Q8BG07 | 4.42 | 2.2771E-05 |
| UP_KW_CELLULAR_COMPONENT | KW-0458~Lysosome | 25 | 8.764E-07 | Q9CQW9, Q60766, P20060, Q80TY0, O70370, P97821, P18242, P10852, Q9WVJ3, Q9D1L9, P61027, O35405, O89023, Q3TBT3, Q8R5L3, Q9Z0M5, P11438, Q99J93, P17439, O09159, Q58A65, O35114, P29416, P63017, Q8BG07 | 3.26 | 9.6399E-06 |

Annotation Cluster 5 — Enrichment Score: 6.123497641883696

| Category | Term | Count | PValue | Genes | Fold Enrichment | FDR |
|---|---|---|---|---|---|---|
| GOTERM_BP_DIRECT | GO:0006096~glycolytic process | 11 | 4.024E-10 | P17710, P12382, Q9Z0M5, Q3TRM8, P05063, P05064, O08528, P09411, P17182, Q9WUA3 | 18.08 | 3.1411E-07 |
| KEGG_PATHWAY | mmu00010:Glycolysis / Gluconeogenesis | 15 | 6.832E-10 | Q3TRM8, Q80VQ0, O08528, P17710, P12382, P47738, P05063, P05064, P09411, P17182, Q9WUA3, Q9D0F9, Q9QXG4, Q8BH04 | 9.00 | 1.8721E-07 |
| GOTERM_BP_DIRECT | GO:0061621~canonical glycolysis | 8 | 1.873E-09 | P17710, P12382, P05064, O08528, P09411, P17182, Q9WUA3 | 33.90 | 8.7712E-07 |
| KEGG_PATHWAY | mmu01200:Carbon metabolism | 18 | 7.161E-09 | Q9CXV1, Q3TRM8, Q9DCD0, O08528, Q9Z2I8, P17710, P12382, Q9CQA3, P14152, Q9Z2I9, Q99KI0, P05063, P05064, P09411, P17182, Q9WUA3, Q9QXG4 | 5.93 | 9.8104E-07 |
| UP_KW_BIOLOGICAL_PROCESS | KW-0324~Glycolysis | 10 | 3.408E-08 | P17710, P12382, Q3TRM8, P05063, P05064, O08528, P09411, P17182, Q9WUA3 | 13.39 | 1.4656E-06 |

| | | | | | | |
|---|---|---|---|---|---|---|
| KEGG_PATHWAY | mmu00051:Fructose and mannose metabolism | 10 | 1.589E-07 | P23591, P17710, P12382, Q3TRM8, P05063, P45376, P05064, O08528, Q8BTZ7, Q9WUA3 | 11.16 | 1.1971E-05 |
| KEGG_PATHWAY | mmu04066:HIF-1 signaling pathway | 16 | 1.748E-07 | Q6PHZ2, Q63932, Q3TRM8, Q8BTI9, O08528, Q62351, P17710, P12382, Q8CIH5, P05063, P05064, P09411, P17182, Q9WUA3, P97481 | 5.50 | 1.1971E-05 |
| KEGG_PATHWAY | mmu00052:Galactose metabolism | 8 | 1.041E-05 | P17710, P12382, Q3TRM8, P45376, Q9R0N0, O08528, Q9WUA3, Q9D0F9 | 10.05 | 0.00028516 |
| KEGG_PATHWAY | mmu01230:Biosynthesis of amino acids | 11 | 2.321E-05 | P12382, Q99JW2, Q3THS6, Q99KI0, P05063, P05064, P09411, P17182, Q9WUA3, Q61024 | 5.60 | 0.00057818 |
| GOTERM_BP_DIRECT | GO:0006002~fructose 6-phosphate metabolic process | 5 | 2.837E-05 | P17710, P12382, Q3TRM8, O08528, Q9WUA3 | 26.84 | 0.00390798 |
| GOTERM_BP_DIRECT | GO:0030388~fructose 1,6-bisphosphate metabolic proc | 4 | 0.000212 | P12382, P05063, P05064, Q9WUA3 | 32.21 | 0.01600228 |
| UP_KW_MOLECULAR_FUNCTION | KW-0021~Allosteric enzyme | 7 | 0.000691 | P17710, P12382, B2RQC6, Q3TRM8, O08528, Q9WUA3 | 6.47 | 0.00659819 |
| KEGG_PATHWAY | mmu05230:Central carbon metabolism in cancer | 7 | 0.007086 | P17710, P12382, Q63932, Q8BTI9, Q3TRM8, O08528, Q9WUA3 | 4.08 | 0.077667 |

Annotation Cluster 6 — Enrichment Score: 4.755612806896643

| Category | Term | Count | PValue | Genes | Fold Enrichment | FDR |
|---|---|---|---|---|---|---|
| GOTERM_MF_DIRECT | GO:0016887~ATP hydrolysis activity | 24 | 5.797E-09 | Q91VR5, Q61656, Q9QXZ0, Q501J6, P38647, Q9WTM5, E9QAM5, Q61881, Q99J87, Q9JIK5, P36371, P49718, Q9EPU0, O54984, P33175, P21958, Q62095, P63017, Q9DBY8, Q6P542, Q922S8, Q8CGK3, Q9CZP5, E9Q555 | 4.48 | 6.3358E-07 |
| INTERPRO | IPR003593:AAA+_ATPase | 10 | 0.000783 | P36371, P21958, Q9WTM5, E9QAM5, Q61881, Q9DBY8, Q6P542, Q8CGK3, Q9CZP5, E9Q555 | 4.10 | 0.10194182 |
| SMART | SM00382:AAA | 10 | 0.001192 | P36371, P21958, Q9WTM5, E9QAM5, Q61881, Q9DBY8, Q6P542, Q8CGK3, Q9CZP5, E9Q555 | 3.82 | 0.10789218 |

Annotation Cluster 7 — Enrichment Score: 3.871700081741407

| Category | Term | Count | PValue | Genes | Fold Enrichment | FDR |
|---|---|---|---|---|---|---|
| KEGG_PATHWAY | mmu00020:Citrate cycle (TCA cycle) | 8 | 1.041E-05 | Q9CQA3, Q9CXV1, Q91V92, Q9Z2I9, P14152, Q99KI0, Q9Z2I8, Q8BH04 | 10.05 | 0.00028516 |
| GOTERM_BP_DIRECT | GO:0006099~tricarboxylic acid cycle | 6 | 5.061E-05 | Q9CQA3, Q9CXV1, Q9Z2I9, P14152, Q99KI0, Q9Z2I8 | 14.64 | 0.00592644 |
| UP_KW_BIOLOGICAL_PROCESS | KW-0816~Tricarboxylic acid cycle | 6 | 0.000201 | Q9CQA3, Q9CXV1, Q9Z2I9, P14152, Q99KI0, Q9Z2I8 | 10.71 | 0.00431575 |
| GOTERM_BP_DIRECT | GO:0006105~succinate metabolic process | 3 | 0.003083 | Q9CQA3, Q9Z2I9, Q9Z2I8 | 34.51 | 0.10087921 |

Annotation Cluster 8 — Enrichment Score: 3.5195346632137845

| Category | Term | Count | PValue | Genes | Fold Enrichment | FDR |
|---|---|---|---|---|---|---|
| KEGG_PATHWAY | mmu00020:Citrate cycle (TCA cycle) | 8 | 1.041E-05 | Q9CQA3, Q9CXV1, Q91V92, Q9Z2I9, P14152, Q99KI0, Q9Z2I8, Q8BH04 | 10.05 | 0.00028516 |
| UP_SEQ_FEATURE | DOMAIN:ATP-grasp | 5 | 0.000112 | B2RQC6, Q91V92, Q9Z2I9, Q9Z2I8 | 18.95 | 0.02122155 |
| INTERPRO | IPR017866:Succ-CoA_synthase_bsu_CS | 3 | 0.000821 | Q91V92, Q9Z2I9, Q9Z2I8 | 59.87 | 0.10194182 |
| INTERPRO | IPR005811:SUCC_ACL_C | 3 | 0.001624 | Q91V92, Q9Z2I9, Q9Z2I8 | 44.90 | 0.13764638 |
| INTERPRO | IPR016102:Succinyl-CoA_synth-like | 3 | 0.001624 | Q91V92, Q9Z2I9, Q9Z2I8 | 44.90 | 0.13764638 |

Annotation Cluster 9 — Enrichment Score: 3.3154188039525976

| Category | Term | Count | PValue | Genes | Fold Enrichment | FDR |
|---|---|---|---|---|---|---|
| GOTERM_MF_DIRECT | GO:0016887~ATP hydrolysis activity | 24 | 5.797E-09 | Q91VR5, Q61656, Q9QXZ0, Q501J6, P38647, Q9WTM5, E9QAM5, Q61881, Q99J87, Q9JIK5, P36371, P49718, Q9EPU0, O54984, P33175, P21958, Q62095, P63017, Q9DBY8, Q6P542, Q922S8, Q8CGK3, Q9CZP5, E9Q555 | 4.48 | 6.3358E-07 |
| GOTERM_MF_DIRECT | GO:0016787~hydrolase activity | 17 | 2.064E-06 | Q99JW2, Q91VR5, Q61656, Q60766, P20060, Q501J6, P18242, P17439, Q3UX10, Q61881, Q99J87, Q9JIK5, P51908, P49718, Q62095, Q5NC05, Q9QXK7 | 4.39 | 0.00012713 |
| GOTERM_MF_DIRECT | GO:0140584~chromatin extrusion motor activity | 12 | 2.493E-06 | P49718, Q9EPU0, Q91VR5, Q61656, Q62095, Q501J6, Q9WTM5, E9QAM5, Q61881, Q99J87, E9Q555, Q9JIK5 | 6.54 | 0.00012713 |
| GOTERM_MF_DIRECT | GO:0140665~ATP-dependent H3-H4 histone complex c | 12 | 2.493E-06 | P49718, Q9EPU0, Q91VR5, Q61656, Q62095, Q501J6, Q9WTM5, E9QAM5, Q61881, Q99J87, E9Q555, Q9JIK5 | 6.54 | 0.00012713 |
| GOTERM_MF_DIRECT | GO:0140849~ATP-dependent H2AZ histone chaperone | 12 | 2.493E-06 | P49718, Q9EPU0, Q91VR5, Q61656, Q62095, Q501J6, Q9WTM5, E9QAM5, Q61881, Q99J87, E9Q555, Q9JIK5 | 6.54 | 0.00012713 |
| GOTERM_MF_DIRECT | GO:0061775~cohesin loader activity | 12 | 2.849E-06 | P49718, Q9EPU0, Q91VR5, Q61656, Q62095, Q501J6, Q9WTM5, E9QAM5, Q61881, Q99J87, E9Q555, Q9JIK5 | 6.45 | 0.00013196 |
| GOTERM_MF_DIRECT | GO:0003689~DNA clamp loader activity | 12 | 4.474E-06 | P49718, Q9EPU0, Q91VR5, Q61656, Q62095, Q501J6, Q9WTM5, E9QAM5, Q61881, Q99J87, E9Q555, Q9JIK5 | 6.16 | 0.00019014 |
| GOTERM_MF_DIRECT | GO:0003724~RNA helicase activity | 8 | 2.914E-05 | Q9EPU0, Q91VR5, Q61656, Q62095, Q501J6, E9QAM5, Q99J87, Q9JIK5 | 9.03 | 0.00101342 |
| UP_KW_MOLECULAR_FUNCTION | KW-0347~Helicase | 12 | 0.000131 | P49718, Q9EPU0, Q91VR5, Q61656, Q62095, Q501J6, Q5NC05, Q9WTM5, E9QAM5, Q61881, Q99J87, Q9JIK5 | 4.21 | 0.00178137 |
| UP_SEQ_FEATURE | MOTIF:DEAD box | 5 | 0.001799 | Q91VR5, Q61656, Q62095, Q501J6, Q9JIK5 | 9.47 | 0.09540855 |
| UP_SEQ_FEATURE | MOTIF:Q motif | 5 | 0.004124 | Q91VR5, Q61656, Q62095, Q501J6, Q9JIK5 | 7.58 | 0.17088833 |
| UP_KW_BIOLOGICAL_PROCESS | KW-0051~Antiviral defense | 8 | 0.004877 | Q61510, P11928, Q91VR5, Q9CQW9, Q99J93, Q501J6, Q99J87, Q9JIK5 | 3.82 | 0.05243 |
| UP_SEQ_FEATURE | DOMAIN:Helicase C-terminal | 7 | 0.010149 | Q91VR5, Q61656, Q62095, Q501J6, Q5NC05, Q99J87, Q9JIK5 | 3.82 | 0.32042055 |
| UP_SEQ_FEATURE | DOMAIN:DEAD-box RNA helicase Q | 4 | 0.01066 | Q91VR5, Q61656, Q62095, Q501J6 | 8.66 | 0.32871706 |
| INTERPRO | IPR001650:Helicase_C-like | 7 | 0.01076 | Q91VR5, Q61656, Q62095, Q501J6, Q5NC05, Q99J87, Q9JIK5 | 3.78 | 0.45596284 |
| INTERPRO | IPR014001:Helicase_ATP-bd | 7 | 0.012164 | Q91VR5, Q61656, Q62095, Q501J6, Q5NC05, Q99J87, Q9JIK5 | 3.68 | 0.49982552 |
| UP_SEQ_FEATURE | DOMAIN:Helicase ATP-binding | 7 | 0.012927 | Q91VR5, Q61656, Q62095, Q501J6, Q5NC05, Q99J87, Q9JIK5 | 3.63 | 0.38092308 |
| SMART | SM00490:HELICc | 7 | 0.014332 | Q91VR5, Q61656, Q62095, Q501J6, Q5NC05, Q99J87, Q9JIK5 | 3.52 | 0.56243068 |
| SMART | SM00487:DEXDc | 7 | 0.015537 | Q91VR5, Q61656, Q62095, Q501J6, Q5NC05, Q99J87, Q9JIK5 | 3.46 | 0.56243068 |
| INTERPRO | IPR014014:RNA_helicase_DEAD_Q_motif | 4 | 0.020242 | Q91VR5, Q61656, Q62095, Q501J6 | 6.84 | 0.64083499 |
| INTERPRO | IPR011545:DEAD/DEAH_box_helicase_dom | 5 | 0.033594 | Q91VR5, Q61656, Q62095, Q501J6, Q9JIK5 | 4.10 | 0.64083499 |
| GOTERM_MF_DIRECT | GO:0004386~helicase activity | 4 | 0.069312 | Q62095, Q501J6, Q5NC05, E9QAM5 | 4.21 | 0.42760857 |
| INTERPRO | IPR000629:RNA-helicase_DEAD-box_CS | 3 | 0.088731 | Q61656, Q62095, Q501J6 | 5.99 | 0.86560724 |

Annotation Cluster 10 — Enrichment Score: 3.250211517970625

| Category | Term | Count | PValue | Genes | Fold Enrichment | FDR |
|---|---|---|---|---|---|---|

| | | | | | | |
|---|---|---|---|---|---|---|
| UP_KW_PTM | KW-1017~Isopeptide bond | 64 | 0.000217 | Q61510, P97461, P09405, P35979, Q9CZW5, Q9WTX6, Q8BTM8, Q07832, P40124, P42225, D3YXK2, Q62095, Q3THS6, Q52KI8, P61027, P20152, P07356, Q80X90, Q99LX0, Q6PFD9, Q9WTM5, P97315, Q6NZF1, Q61881, P62960, Q9CYG7, Q9JLQ0, Q9D906, P05064, Q9QXK7, P17918, Q9CR09, P63017, Q8BVY0, O88286, E9Q555, Q9CXY6, Q6P9P6, Q91VR5, Q61656, Q9CQW9, Q60766, Q501J6, P10852, P59325, Q80X82, P63024, Q8CGC6, Q91V92, O35309, O88351, Q60972, Q3TBT3, Q60930, Q9WU84, Q8CGZ0, Q8BX17, O35892, P17182, Q9DBY8, P26369, Q8BMQ2, E9PVX6 | 1.57 | 0.00173271 |
| UP_SEQ_FEATURE | CROSSLNK:Glycyl lysine isopeptide (Lys-Gly) (intercha | 34 | 0.000295 | Q9CXY6, P35979, Q6P9P6, P09405, Q61656, Q9CZW5, Q501J6, Q07832, P10852, P59325, Q80X82, Q8CGC6, D3YXK2, Q62095, Q52KI8, Q3THS6, P20152, Q6PFD9, Q9WTM5, P97315, Q6NZF1, Q8CGZ0, Q61881, Q8BX17, P62960, Q9CYG7, Q9JLQ0, P17918, Q9DBY8, Q8BVY0, Q8BMQ2, E9PVX6, O88286, E9Q555 | 1.97 | 0.03906538 |
| UP_KW_PTM | KW-0832~Ubl conjugation | 83 | 0.002783 | Q61510, P97461, P09405, P35979, P29452, Q9CZW5, Q9WTX6, Q78IQ7, Q8BTM8, Q07832, Q9WVL2, Q9R0P5, P40124, P42225, D3YXK2, Q62095, Q3THS6, Q52KI8, P61027, A1L314, P20152, P07356, O35405, Q2TBE6, Q80X90, Q99LX0, Q6PFD9, Q9WTM5, P97315, Q6NZF1, Q61881, Q5DQR4, P62960, Q9CYG7, Q9JLQ0, P01900, Q9D906, P52293, P05064, Q9QXK7, P17918, Q9CR09, P63017, Q8BVY0, O88286, E9Q555, P97481, Q9CXY6, Q6P9P6, Q91VR5, Q61656, Q9CQW9, Q60766, Q501J6, P10852, P59325, Q80X82, Q91XB0, P63024, P47738, Q8CGC6, Q91V92, O35309, O88351, Q9D1H7, Q60972, Q3TBT3, Q60930, Q9WU84, P42082, P14426, Q8CGZ0, Q8BX17, P14152, O35892, P83741, P17182, O88983, Q9DBY8, Q922S8, P26369, Q8BMQ2, E9PVX6 | 1.35 | 0.01669679 |

Annotation Cluster 11 — Enrichment Score: 3.1289877065953515

| Category | Term | Count | PValue | Genes | Fold Enrichment | FDR |
|---|---|---|---|---|---|---|
| UP_SEQ_FEATURE | TOPO_DOM:Mitochondrial intermembrane | 11 | 8.692E-05 | Q62425, Q9CXV1, O35435, Q9CZW5, Q791V5, Q78IK4, Q9QXX4, Q9CQV7, Q8BH59, Q9CPQ1, Q9CZP5 | 4.90 | 0.02122155 |
| UP_SEQ_FEATURE | TOPO_DOM:Mitochondrial matrix | 9 | 0.000139 | Q62425, Q9CXV1, O35435, Q78IK4, Q9QXX4, Q9CQV7, Q8BH59, Q9CPQ1, Q9CZP5 | 5.93 | 0.02300892 |
| GOTERM_CC_DIRECT | GO:0005743~mitochondrial inner membrane | 18 | 0.000475 | Q62425, Q9CXV1, Q60930, Q791V5, Q9QXX4, Q9CQV7, Q9CPQ1, Q9WTP6, P19536, Q9CQA3, O35435, P36552, P62075, P38060, Q8BGH2, Q78IK4, Q8BH59, Q9CZP5 | 2.68 | 0.00558385 |
| UP_KW_CELLULAR_COMPONENT | KW-0999~Mitochondrion inner membrane | 12 | 0.053161 | P19536, Q9CQA3, Q62425, Q9CXV1, O35435, P62075, Q78IK4, Q9QXX4, Q9CQV7, Q8BH59, Q9CPQ1, Q9CZP5 | 1.90 | 0.17210394 |

Annotation Cluster 12 — Enrichment Score: 3.1194543270543855

| Category | Term | Count | PValue | Genes | Fold Enrichment | FDR |
|---|---|---|---|---|---|---|
| GOTERM_BP_DIRECT | GO:0030388~fructose 1,6-bisphosphate metabolic proc | 4 | 0.000212 | P12382, P05063, P05064, Q9WUA3 | 32.21 | 0.01600228 |
| KEGG_PATHWAY | mmu00030:Pentose phosphate pathway | 6 | 0.001385 | P12382, Q9DCD0, P05063, P05064, Q9WUA3, Q9D0F9 | 7.09 | 0.02107905 |
| GOTERM_BP_DIRECT | GO:0061615~glycolytic process through fructose-6-pho | 3 | 0.001493 | P12382, P05064, Q9WUA3 | 48.31 | 0.06132581 |

Annotation Cluster 13 — Enrichment Score: 3.1187877035317317

| Category | Term | Count | PValue | Genes | Fold Enrichment | FDR |
|---|---|---|---|---|---|---|
| UP_SEQ_FEATURE | DOMAIN:ATP-grasp | 5 | 0.000112 | B2RQC6, Q91V92, Q9Z2I9, Q9Z2I8 | 18.95 | 0.02122155 |
| INTERPRO | IPR011761:ATP-grasp | 4 | 0.001798 | B2RQC6, Q9Z2I9, Q9Z2I8 | 15.96 | 0.1434025 |
| INTERPRO | IPR013815:ATP_grasp_subdomain_1 | 4 | 0.002186 | B2RQC6, Q9Z2I9, Q9Z2I8 | 14.97 | 0.15783187 |

Annotation Cluster 14 — Enrichment Score: 2.9654478039293157

| Category | Term | Count | PValue | Genes | Fold Enrichment | FDR |
|---|---|---|---|---|---|---|
| GOTERM_BP_DIRECT | GO:0016192~vesicle-mediated transport | 14 | 1.326E-05 | Q8R5L3, Q9D1J1, Q07797, Q80V94, P63024, Q58A65, O35643, P33175, Q9CZE3, O08585, O88512, P61027, O88983, Q3TAP4 | 4.60 | 0.00238794 |
| GOTERM_CC_DIRECT | GO:0005802~trans-Golgi network | 9 | 0.006264 | Q2TBE6, O35643, Q9CZE3, Q8BTM8, O88512, P61027, P17439, O88983, Q80V94 | 3.30 | 0.04460866 |
| GOTERM_BP_DIRECT | GO:0006886~intracellular protein transport | 9 | 0.01529 | O35643, Q8R5L3, Q9CZE3, O08585, O88512, P61027, O88983, Q80V94, Q8BVL3 | 2.82 | 0.310715 |

Annotation Cluster 15 — Enrichment Score: 2.8618174795368487

| Category | Term | Count | PValue | Genes | Fold Enrichment | FDR |
|---|---|---|---|---|---|---|
| GOTERM_BP_DIRECT | GO:0043171~peptide catabolic process | 6 | 1.553E-06 | Q8C129, Q9EQH2, P24527, Q9WVJ3, Q11011, O89023 | 28.42 | 0.00036921 |
| GOTERM_MF_DIRECT | GO:0004177~aminopeptidase activity | 6 | 2.882E-05 | Q8C129, Q9EQH2, Q6P1B1, P24527, Q9WVJ3, Q11011 | 16.35 | 0.00101342 |
| GOTERM_MF_DIRECT | GO:0070006~metalloaminopeptidase activity | 5 | 0.000126 | Q8C129, Q9EQH2, Q6P1B1, P24527, Q11011 | 18.82 | 0.00418988 |
| INTERPRO | IPR001930:Peptidase_M1 | 4 | 0.000504 | Q8C129, Q9EQH2, P24527, Q11011 | 23.95 | 0.10194182 |
| INTERPRO | IPR045357:Aminopeptidase_N-like_N | 4 | 0.000504 | Q8C129, Q9EQH2, P24527, Q11011 | 23.95 | 0.10194182 |
| INTERPRO | IPR014782:Peptidase_M1_dom | 4 | 0.000685 | Q8C129, Q9EQH2, P24527, Q11011 | 21.77 | 0.10194182 |
| INTERPRO | IPR042097:Aminopeptidase_N-like_N_sf | 4 | 0.000902 | Q8C129, Q9EQH2, P24527, Q11011 | 19.96 | 0.10194182 |
| INTERPRO | IPR027268:Peptidase_M4/M1_CTD_sf | 4 | 0.000902 | Q8C129, Q9EQH2, P24527, Q11011 | 19.96 | 0.10194182 |
| UP_KW_MOLECULAR_FUNCTION | KW-0031~Aminopeptidase | 5 | 0.002033 | Q8C129, Q9EQH2, Q6P1B1, Q9WVJ3, Q11011 | 9.08 | 0.01536146 |
| GOTERM_MF_DIRECT | GO:0008237~metallopeptidase activity | 6 | 0.002765 | Q8K411, Q8C129, O88839, Q9EQH2, Q9Z0F8, Q11011 | 6.24 | 0.04597768 |
| UP_SEQ_FEATURE | SITE:Transition state stabilizer | 6 | 0.00377 | Q9CWJ9, Q8C129, Q9EQH2, P47738, Q9R0N0, Q11011 | 5.77 | 0.17088833 |
| UP_SEQ_FEATURE | DOMAIN:ERAP1-like C-terminal | 3 | 0.005364 | Q8C129, Q9EQH2, Q11011 | 25.99 | 0.19757378 |
| UP_SEQ_FEATURE | DOMAIN:Peptidase M1 membrane alanine aminopeptid | 3 | 0.005364 | Q8C129, Q9EQH2, Q11011 | 25.99 | 0.19757378 |
| INTERPRO | IPR024571:ERAP1-like_C_dom | 3 | 0.005499 | Q8C129, Q9EQH2, Q11011 | 25.66 | 0.26630648 |
| INTERPRO | IPR050344:Peptidase_M1_aminopeptidases | 3 | 0.005499 | Q8C129, Q9EQH2, Q11011 | 25.66 | 0.26630648 |
| INTERPRO | IPR034016:M1_APN-typ | 3 | 0.005499 | Q8C129, Q9EQH2, Q11011 | 25.66 | 0.26630648 |
| UP_SEQ_FEATURE | DOMAIN:Aminopeptidase N-like N-terminal | 3 | 0.007075 | Q8C129, Q9EQH2, Q11011 | 22.74 | 0.25353551 |
| UP_KW_MOLECULAR_FUNCTION | KW-0482~Metalloprotease | 9 | 0.011732 | Q8K411, Q8C129, O88839, Q9EQH2, Q6P1B1, P24527, Q9WVJ3, Q9Z0F8, Q11011 | 2.93 | 0.07252298 |
| COG_ONTOLOGY | Amino acid transport and metabolism | 5 | 0.062721 | Q8C129, Q9EQH2, Q99JW2, Q6P1B1, Q11011 | 3.23 | 0.39723439 |

| Category | Term | Count | PValue | Genes | Fold Enrichment | FDR |
|---|---|---|---|---|---|---|
| Annotation Cluster 16 | Enrichment Score: 2.777702616597501 | | | | | |
| Category | Term | Count | PValue | Genes | Fold Enrichment | FDR |
| KEGG_PATHWAY | mmu00520:Amino sugar and nucleotide sugar metaboli | 10 | 3.352E-07 | P17710, Q3TRM8, P20060, P82343, Q9QZ08, Q9R0N0, O08528, Q8BTZ7, P29416, Q9D0F9 | 10.30 | 1.5308E-05 |
| KEGG_PATHWAY | mmu00052:Galactose metabolism | 8 | 1.041E-05 | P17710, P12382, Q3TRM8, P45376, Q9R0N0, O08528, Q9WUA3, Q9D0F9 | 10.05 | 0.00028516 |
| GOTERM_BP_DIRECT | GO:0006002~fructose 6-phosphate metabolic process | 5 | 2.837E-05 | P17710, P12382, Q3TRM8, O08528, Q9WUA3 | 26.84 | 0.00390798 |
| KEGG_PATHWAY | mmu01250:Biosynthesis of nucleotide sugars | 8 | 3.438E-05 | P23591, P17710, Q3TRM8, Q9QZ08, Q9R0N0, O08528, Q8BTZ7, Q9D0F9 | 8.46 | 0.00078494 |
| GOTERM_BP_DIRECT | GO:0046835~carbohydrate phosphorylation | 5 | 0.00017 | P17710, Q3TRM8, Q9QZ08, Q9R0N0, O08528 | 17.50 | 0.01423214 |
| GOTERM_BP_DIRECT | GO:0006006~glucose metabolic process | 7 | 0.00045 | P17710, P11928, Q9Z0M5, Q3TRM8, O08528, Q9D0F9, Q91X52 | 7.13 | 0.02703786 |
| INTERPRO | IPR043129:ATPase_NBD | 7 | 0.000627 | P17710, Q61699, Q3TRM8, P38647, Q9QZ08, O08528, P63017 | 6.65 | 0.10194182 |
| UP_KW_MOLECULAR_FUNCTION | KW-0021~Allosteric enzyme | 7 | 0.000691 | P17710, P12382, B2RQC6, Q3TRM8, O08528, Q9WUA3 | 6.47 | 0.00659819 |
| GOTERM_MF_DIRECT | GO:0004396~hexokinase activity | 3 | 0.000937 | P17710, Q3TRM8, O08528 | 59.27 | 0.0204806 |
| GOTERM_BP_DIRECT | GO:0019318~hexose metabolic process | 3 | 0.001493 | P17710, Q3TRM8, O08528 | 48.31 | 0.06132581 |
| GOTERM_MF_DIRECT | GO:0004340~glucokinase activity | 3 | 0.001549 | P17710, Q3TRM8, O08528 | 47.42 | 0.02961814 |
| GOTERM_MF_DIRECT | GO:0008865~fructokinase activity | 3 | 0.001549 | P17710, Q3TRM8, O08528 | 47.42 | 0.02961814 |
| GOTERM_BP_DIRECT | GO:0002931~response to ischemia | 6 | 0.001561 | P05555, P17710, Q63932, Q8BTI9, Q3TCJ1, O08528 | 7.10 | 0.06301892 |
| UP_SEQ_FEATURE | REGION:Hexokinase large subdomain 1 | 3 | 0.001584 | P17710, Q3TRM8, O08528 | 45.48 | 0.08749466 |
| UP_SEQ_FEATURE | REGION:Hexokinase large subdomain 2 | 3 | 0.001584 | P17710, Q3TRM8, O08528 | 45.48 | 0.08749466 |
| UP_SEQ_FEATURE | REGION:Hexokinase small subdomain 1 | 3 | 0.001584 | P17710, Q3TRM8, O08528 | 45.48 | 0.08749466 |
| UP_SEQ_FEATURE | REGION:Hexokinase small subdomain 2 | 3 | 0.001584 | P17710, Q3TRM8, O08528 | 45.48 | 0.08749466 |
| UP_SEQ_FEATURE | DOMAIN:Hexokinase 1 | 3 | 0.001584 | P17710, Q3TRM8, O08528 | 45.48 | 0.08749466 |
| UP_SEQ_FEATURE | DOMAIN:Hexokinase 2 | 3 | 0.001584 | P17710, Q3TRM8, O08528 | 45.48 | 0.08749466 |
| UP_SEQ_FEATURE | DOMAIN:Hexokinase C-terminal | 3 | 0.001584 | P17710, Q3TRM8, O08528 | 45.48 | 0.08749466 |
| UP_SEQ_FEATURE | DOMAIN:Hexokinase N-terminal | 3 | 0.001584 | P17710, Q3TRM8, O08528 | 45.48 | 0.08749466 |
| INTERPRO | IPR001312:Hexokinase | 3 | 0.002677 | P17710, Q3TRM8, O08528 | 35.92 | 0.15783187 |
| INTERPRO | IPR019807:Hexokinase_BS | 3 | 0.002677 | P17710, Q3TRM8, O08528 | 35.92 | 0.15783187 |
| INTERPRO | IPR022672:Hexokinase_N | 3 | 0.002677 | P17710, Q3TRM8, O08528 | 35.92 | 0.15783187 |
| INTERPRO | IPR022673:Hexokinase_C | 3 | 0.002677 | P17710, Q3TRM8, O08528 | 35.92 | 0.15783187 |
| KEGG_PATHWAY | mmu00524:Neomycin, kanamycin and gentamicin biosy | 3 | 0.005819 | P17710, Q3TRM8, O08528 | 24.11 | 0.06643506 |
| KEGG_PATHWAY | mmu05230:Central carbon metabolism in cancer | 7 | 0.007086 | P17710, P12382, Q63932, Q8BTI9, Q3TRM8, O08528, Q9WUA3 | 4.08 | 0.077667 |
| GOTERM_BP_DIRECT | GO:0051156~glucose 6-phosphate metabolic process | 3 | 0.009302 | P17710, Q3TRM8, O08528 | 20.13 | 0.22004419 |
| GOTERM_MF_DIRECT | GO:0005536~D-glucose binding | 3 | 0.013074 | P17710, Q3TRM8, O08528 | 16.93 | 0.16668802 |
| KEGG_PATHWAY | mmu04910:Insulin signaling pathway | 9 | 0.022996 | P17710, Q9DBC7, Q63932, Q60634, Q8BTI9, Q3TRM8, O08528, Q8BH04, O88351 | 2.58 | 0.17502369 |
| KEGG_PATHWAY | mmu04930:Type II diabetes mellitus | 5 | 0.03054 | P17710, Q8BTI9, Q3TRM8, O08528, O88351 | 4.19 | 0.20919559 |
| KEGG_PATHWAY | mmu04973:Carbohydrate digestion and absorption | 5 | 0.032629 | P17710, P51432, Q8BTI9, Q3TRM8, O08528 | 4.10 | 0.21806006 |
| GOTERM_BP_DIRECT | GO:0001678~intracellular glucose homeostasis | 3 | 0.046799 | P17710, Q3TRM8, O08528 | 8.63 | 0.59631298 |
| KEGG_PATHWAY | mmu00500:Starch and sucrose metabolism | 4 | 0.054978 | P17710, Q3TRM8, O08528, Q9D0F9 | 4.59 | 0.28969199 |
| COG_ONTOLOGY | Carbohydrate transport and metabolism | 3 | 0.170058 | P17710, Q3TRM8, O08528 | 3.95 | 0.80777766 |
| Annotation Cluster 17 | Enrichment Score: 2.772267485785425 | | | | | |
| Category | Term | Count | PValue | Genes | Fold Enrichment | FDR |
| GOTERM_BP_DIRECT | GO:0035456~response to interferon-beta | 4 | 0.000491 | P42225, Q9CQW9, Q99J93, Q8R2Q8 | 24.78 | 0.02742429 |
| GOTERM_BP_DIRECT | GO:0060337~type I interferon-mediated signaling pathw | 6 | 0.0007 | P42225, P11928, Q9CQW9, Q99J93, Q9WVL2, Q91XB0 | 8.48 | 0.03727522 |
| GOTERM_BP_DIRECT | GO:0034341~response to type II interferon | 5 | 0.000895 | P42225, Q9CQW9, Q99J93, O35892, Q8R2Q8 | 11.50 | 0.04380303 |
| GOTERM_BP_DIRECT | GO:0045071~negative regulation of viral genome replica | 5 | 0.004803 | P11928, Q9CQW9, Q99J93, D0QMC3, Q8R2Q8 | 7.32 | 0.13717029 |
| GOTERM_BP_DIRECT | GO:0035455~response to interferon-alpha | 3 | 0.009302 | Q9CQW9, Q99J93, Q8R2Q8 | 20.13 | 0.22004419 |
| Annotation Cluster 18 | Enrichment Score: 2.756556253801063 | | | | | |
| Category | Term | Count | PValue | Genes | Fold Enrichment | FDR |
| GOTERM_MF_DIRECT | GO:0051015~actin filament binding | 15 | 1.25E-06 | Q99K51, Q7TPR4, Q80X90, Q9QXZ0, Q8K2Q9, Q8BTM8, O08539, Q9R0P5, Q9JKF1, Q9QZQ1, Q3UQ44, Q7TMB8, Q9JLQ0, P47753, Q6IRU2 | 5.25 | 8.6921E-05 |
| UP_SEQ_FEATURE | DOMAIN:Calponin-homology (CH) 1 | 5 | 0.000498 | Q99K51, Q80X90, Q7TPR4, Q9QXZ0, Q8BTM8 | 13.18 | 0.05499155 |
| UP_SEQ_FEATURE | DOMAIN:Calponin-homology (CH) 2 | 5 | 0.000498 | Q99K51, Q80X90, Q7TPR4, Q9QXZ0, Q8BTM8 | 13.18 | 0.05499155 |
| INTERPRO | IPR001589:Actinin_actin-bd_CS | 5 | 0.000522 | Q99K51, Q80X90, Q7TPR4, Q9QXZ0, Q8BTM8 | 13.01 | 0.10194182 |
| SMART | SM00033:CH | 7 | 0.001108 | Q99K51, Q3UQ44, Q80X90, Q7TPR4, Q9QXZ0, Q8BTM8, Q9JKF1 | 5.92 | 0.10789218 |
| INTERPRO | IPR001715:CH_dom | 7 | 0.002481 | Q99K51, Q3UQ44, Q80X90, Q7TPR4, Q9QXZ0, Q8BTM8, Q9JKF1 | 5.11 | 0.15783187 |
| INTERPRO | IPR036872:CH_dom_sf | 7 | 0.002801 | Q99K51, Q3UQ44, Q80X90, Q7TPR4, Q9QXZ0, Q8BTM8, Q9JKF1 | 4.99 | 0.15826162 |
| UP_SEQ_FEATURE | DOMAIN:Calponin-homology (CH) | 6 | 0.008802 | Q99K51, Q3UQ44, Q7TPR4, Q9QXZ0, Q8BTM8, Q9JKF1 | 4.72 | 0.2909897 |
| UP_KW_MOLECULAR_FUNCTION | KW-0009~Actin-binding | 12 | 0.017937 | Q99K51, P40124, Q8BH43, Q80X90, Q7TPR4, Q7TMB8, Q9QXZ0, Q8BTM8, P47753, Q6IRU2, Q9R0P5, P48193 | 2.25 | 0.09382271 |
| GOTERM_CC_DIRECT | GO:0005884~actin filament | 5 | 0.034495 | Q99K51, Q7TPR4, Q9JLQ0, Q8BTM8, Q6IRU2 | 4.08 | 0.17247537 |
| UP_SEQ_FEATURE | REGION:Actin-binding | 4 | 0.070256 | Q80X90, Q7TPR4, Q9QXZ0, Q8BTM8 | 4.18 | 0.67021426 |
| Annotation Cluster 19 | Enrichment Score: 2.6989388802939955 | | | | | |
| Category | Term | Count | PValue | Genes | Fold Enrichment | FDR |

| | | | | | | |
|---|---|---|---|---|---|---|
| GOTERM_MF_DIRECT | GO:0004151~dihydroorotase activity | 3 | 0.000472 | O35435, B2RQC6 | 79.03 | 0.01129474 |
| GOTERM_BP_DIRECT | GO:0044205~'de novo' UMP biosynthetic process | 3 | 0.000903 | O35435, B2RQC6 | 60.39 | 0.04380303 |
| GOTERM_BP_DIRECT | GO:0006225~UDP biosynthetic process | 3 | 0.002221 | O35435, B2RQC6 | 40.26 | 0.08525287 |
| GOTERM_BP_DIRECT | GO:0006207~'de novo' pyrimidine nucleobase biosynthe | 3 | 0.003083 | O35435, B2RQC6 | 34.51 | 0.10087921 |
| UP_KW_BIOLOGICAL_PROCESS | KW-0665~Pyrimidine biosynthesis | 3 | 0.01096 | O35435, B2RQC6 | 18.08 | 0.09425789 |

Annotation Cluster 20 Enrichment Score: 2.6713247544333476

| Category | Term | Count | PValue | Genes | Fold Enrichment | FDR |
|---|---|---|---|---|---|---|
| GOTERM_BP_DIRECT | GO:0006397~mRNA processing | 17 | 1.598E-07 | Q0VGB7, Q91VR5, Q61656, P52912, Q62189, Q80X82, P51908, P62960, Q8CGC6, Q52KI8, Q5NC05, Q9CQG2, Q9QXK7, Q9JK81, Q9DBR1, O35658, P26369 | 5.33 | 5.3465E-05 |
| GOTERM_BP_DIRECT | GO:0008380~RNA splicing | 11 | 0.00022 | Q0VGB7, P62960, Q8CGC6, Q61656, Q52KI8, Q5NC05, P52912, Q62189, P63017, O35658, P26369 | 4.41 | 0.01600228 |
| UP_KW_BIOLOGICAL_PROCESS | KW-0507~mRNA processing | 20 | 0.000306 | Q0VGB7, Q91VR5, Q61656, Q501J6, P52912, Q62189, Q80X82, Q8BX17, P51908, P62960, Q8CGC6, Q52KI8, Q5NC05, Q9QXK7, P63017, P62313, Q9DBR1, O35658, P26369, Q99LI7 | 2.56 | 0.00526663 |
| GOTERM_CC_DIRECT | GO:0005681~spliceosomal complex | 9 | 0.000551 | O89086, P09405, Q8CGC6, Q61656, Q52KI8, Q5NC05, Q62189, P63017, P26369 | 4.88 | 0.00604247 |
| UP_KW_BIOLOGICAL_PROCESS | KW-0508~mRNA splicing | 14 | 0.009254 | Q0VGB7, Q61656, Q501J6, P52912, Q62189, Q8BX17, P62960, Q8CGC6, Q52KI8, Q5NC05, P63017, P62313, O35658, P26369 | 2.26 | 0.08842293 |
| UP_KW_CELLULAR_COMPONENT | KW-0747~Spliceosome | 7 | 0.062872 | Q8CGC6, Q61656, Q52KI8, Q5NC05, Q62189, P63017, P62313 | 2.49 | 0.17210394 |
| GOTERM_BP_DIRECT | GO:0000398~mRNA splicing, via spliceosome | 5 | 0.156228 | Q52KI8, Q62189, P63017, P62313, P26369 | 2.40 | 0.97339983 |
| KEGG_PATHWAY | mmu03040:Spliceosome | 6 | 0.791033 | Q61656, Q8CGZ0, Q62189, P63017, P62313, P26369 | 0.91 | 0.91333333 |

Annotation Cluster 21 Enrichment Score: 2.490731221181822

| Category | Term | Count | PValue | Genes | Fold Enrichment | FDR |
|---|---|---|---|---|---|---|
| GOTERM_CC_DIRECT | GO:0030863~cortical cytoskeleton | 5 | 0.000791 | Q7TPR4, Q8BTM8, P47753, Q6IRU2, P48193 | 11.88 | 0.00759103 |
| GOTERM_MF_DIRECT | GO:0003779~actin binding | 13 | 0.002463 | Q8BH43, Q7TPR4, Q9QXZ0, P26041, Q8BTM8, Q9Z0E6, P97814, P48193, P40124, Q9JLQ0, P47753, Q6IRU2, P09103 | 2.82 | 0.0428302 |
| GOTERM_BP_DIRECT | GO:0030036~actin cytoskeleton organization | 10 | 0.003115 | P40124, Q8BH43, Q80X90, Q7TPR4, Q8BTM8, P47753, P97315, E9Q3L2, P48193, P60766 | 3.37 | 0.10087921 |
| UP_KW_MOLECULAR_FUNCTION | KW-0009~Actin-binding | 12 | 0.017937 | Q99K51, P40124, Q8BH43, Q80X90, Q7TPR4, Q7TMB8, Q9QXZ0, Q8BTM8, P47753, Q6IRU2, Q9R0P5, P48193 | 2.25 | 0.09382271 |

Annotation Cluster 22 Enrichment Score: 2.4065581066122923

| Category | Term | Count | PValue | Genes | Fold Enrichment | FDR |
|---|---|---|---|---|---|---|
| GOTERM_CC_DIRECT | GO:0005874~microtubule | 17 | 6.731E-06 | Q6P9P6, Q61699, Q63932, Q9QXZ0, P29452, Q8K2Q9, O08539, Q9JKF1, Q3UX10, Q62433, Q3UQ44, P33175, Q3TCJ1, Q8BTU1, Q3UMY5, P63017, Q922S8 | 4.00 | 0.00015065 |
| GOTERM_MF_DIRECT | GO:0008017~microtubule binding | 8 | 0.056371 | Q6P9P6, Q9QXZ0, P33175, Q3TCJ1, Q07832, Q3UMY5, Q62433, Q922S8 | 2.34 | 0.3858111 |
| UP_KW_CELLULAR_COMPONENT | KW-0493~Microtubule | 9 | 0.158923 | Q6P9P6, Q9QXZ0, P33175, Q3TCJ1, Q8BTU1, Q3UMY5, Q3UX10, Q62433, Q922S8 | 1.71 | 0.34963127 |

Annotation Cluster 23 Enrichment Score: 2.3369290982974738

| Category | Term | Count | PValue | Genes | Fold Enrichment | FDR |
|---|---|---|---|---|---|---|
| GOTERM_BP_DIRECT | GO:0006508~proteolysis | 20 | 0.000225 | Q8K411, Q9EQH2, Q6P1B1, P29452, Q99LX0, Q6PFD9, P29594, O70370, P97821, P18242, Q8C129, O88839, P24527, Q9WVJ3, Q9Z0F8, O88456, Q8C166, Q11011, Q8CGK3, O89023 | 2.66 | 0.01600228 |
| GOTERM_MF_DIRECT | GO:0008233~peptidase activity | 9 | 0.000821 | Q9EQH2, P29452, Q9Z0F8, Q99LX0, P29594, O70370, P18242, Q9D8V0, O89023 | 4.59 | 0.01903167 |
| UP_KW_MOLECULAR_FUNCTION | KW-0645~Protease | 19 | 0.030323 | Q8K411, Q9EQH2, Q6P1B1, P29452, Q99LX0, Q6PFD9, P29594, O70370, P97821, P18242, Q9D8V0, Q8C129, O88839, P24527, Q9WVJ3, Q9Z0F8, Q11011, Q8CGK3, O89023 | 1.70 | 0.14728295 |
| UP_KW_PTM | KW-0865~Zymogen | 11 | 0.079997 | O88839, P29452, Q9WVJ3, Q9Z0F8, Q99LX0, P29594, O70370, P97821, P18242, P29416, O89023 | 1.83 | 0.30378459 |

Annotation Cluster 24 Enrichment Score: 2.273176521239321

| Category | Term | Count | PValue | Genes | Fold Enrichment | FDR |
|---|---|---|---|---|---|---|
| GOTERM_MF_DIRECT | GO:0005525~GTP binding | 16 | 0.000148 | Q60766, Q61107, P14824, Q9Z0E6, Q6PA06, Q9Z2I8, Q9EQP2, P59325, Q3UX10, Q8VEH6, Q9CZE3, P61027, Q8BTZ7, P42208, Q8BH04, P60766 | 3.24 | 0.00471044 |
| GOTERM_MF_DIRECT | GO:1990606~membrane scission GTPase motor activit | 5 | 0.001748 | Q60766, Q61107, P61027, Q9Z0E6, P60766 | 9.64 | 0.0326082 |
| GOTERM_MF_DIRECT | GO:0003924~GTPase activity | 10 | 0.018952 | Q8VEH6, P63213, Q60766, Q61107, Q9CZE3, P61027, Q9Z0E6, Q6PA06, P42208, P60766 | 2.52 | 0.22430602 |
| UP_KW_LIGAND | KW-0342~GTP-binding | 14 | 0.165032 | Q60766, Q61107, Q9Z0E6, Q6PA06, Q9Z2I8, P59325, Q3UX10, Q8VEH6, Q9CZE3, P61027, Q8BTZ7, P42208, Q8BH04, P60766 | 1.45 | 1 |

Annotation Cluster 25 Enrichment Score: 2.265289871760981

| Category | Term | Count | PValue | Genes | Fold Enrichment | FDR |
|---|---|---|---|---|---|---|
| KEGG_PATHWAY | mmu05170:Human immunodeficiency virus 1 infection | 18 | 9.938E-05 | Q3TBT3, P06339, Q8R4K2, Q63932, Q8BTI9, Q9WTX6, P14426, P70227, P36371, O35643, P63213, P01902, P21958, P01901, P01900, Q8CIH5, O88512, Q8R2Q8, O88351 | 3.00 | 0.00209464 |
| GOTERM_MF_DIRECT | GO:0042288~MHC class I protein binding | 6 | 0.000202 | P36371, P06339, P01902, P21958, P01901, P01900, P14426 | 11.03 | 0.00572681 |
| KEGG_PATHWAY | mmu05168:Herpes simplex virus 1 infection | 15 | 0.000828 | Q3TBT3, P06339, P11928, Q8R4K2, Q8BTI9, Q9WVL2, P14426, P42225, P36371, P01902, P21958, O35892, P01901, P01900, Q8R2Q8, O88351 | 2.83 | 0.01620678 |
| KEGG_PATHWAY | mmu05169:Epstein-Barr virus infection | 15 | 0.001602 | P06339, P11928, Q8R4K2, Q8BTI9, Q9WVL2, P14426, P15379, P42225, P36371, P01902, P21958, P01901, P01900, Q8CIH5, P20152, O88351 | 2.64 | 0.02194378 |
| GOTERM_MF_DIRECT | GO:0042605~peptide antigen binding | 6 | 0.004258 | P36371, P06339, P01902, P21958, P01901, P01900, P14426 | 5.64 | 0.06250767 |
| KEGG_PATHWAY | mmu04612:Antigen processing and presentation | 8 | 0.005707 | P36371, P06339, P01902, P21958, P01901, P01900, O70370, P14426, P63017 | 3.70 | 0.06643506 |
| KEGG_PATHWAY | mmu05167:Kaposi sarcoma-associated herpesvirus infe | 13 | 0.009496 | P06339, Q63932, Q8BTI9, Q9WVL2, P42082, P14426, P70227, P42225, P63213, P01902, P01901, P01900, Q8CIH5, O88351 | 2.34 | 0.09455644 |

| | | | | | | |
|---|---|---|---|---|---|---|
| KEGG_PATHWAY | mmu05163:Human cytomegalovirus infection | 13 | 0.024165 | Q3TBT3, P06339, P51432, Q63932, Q8BTI9, P14426, P70227, P36371, P63213, P01902, P21958, P01901, P01900, O88351 | 2.06 | 0.17895215 |
| KEGG_PATHWAY | mmu04218:Cellular senescence | 9 | 0.084762 | Q60972, P06339, Q63932, Q60930, Q8BTI9, P01902, P01901, P01900, P14426, P70227 | 1.98 | 0.36288578 |
| KEGG_PATHWAY | mmu05166:Human T-cell leukemia virus 1 infection | 11 | 0.087031 | P06339, Q63932, Q60930, Q8BTI9, P97333, P01902, P01901, P01900, Q9WVM3, P14426, Q80WQ2, O88351 | 1.79 | 0.36687111 |
| KEGG_PATHWAY | mmu05165:Human papillomavirus infection | 14 | 0.110164 | P06339, Q63932, Q8BTI9, Q8BVE3, Q9WVL2, P14426, A2AN08, Q76MZ3, P42225, P01902, P01901, P01900, O70309, O88351, P60766 | 1.57 | 0.40134219 |
| Annotation Cluster 26 | Enrichment Score: 2.192205670318874 | | | | | |
| Category | Term | Count | PValue | Genes | Fold Enrichment | FDR |
| GOTERM_BP_DIRECT | GO:0034975~protein folding in endoplasmic reticulum | 4 | 3.666E-05 | Q8R2E9, P09103, Q8R180, Q91YW3 | 53.68 | 0.00477017 |
| GOTERM_MF_DIRECT | GO:0016972~thiol oxidase activity | 3 | 0.000472 | Q8R2E9, P09103, Q8R180 | 79.03 | 0.01129474 |
| GOTERM_BP_DIRECT | GO:0018401~peptidyl-proline hydroxylation to 4-hydrox | 3 | 0.003083 | Q8R2E9, P09103, Q8R180 | 34.51 | 0.10087921 |
| GOTERM_MF_DIRECT | GO:0015035~protein-disulfide reductase activity | 3 | 0.031063 | Q8R2E9, P09103, Q8R180 | 10.78 | 0.28630474 |
| UP_KW_DOMAIN | KW-0676~Redox-active center | 3 | 0.201053 | Q8R2E9, P09103, Q8R180 | 3.60 | 0.347273 |
| UP_SEQ_FEATURE | DISULFID:Redox-active | 3 | 0.210675 | Q8R2E9, P09103, Q8R180 | 3.50 | 0.99251497 |
| Annotation Cluster 27 | Enrichment Score: 2.191992773734665 | | | | | |
| Category | Term | Count | PValue | Genes | Fold Enrichment | FDR |
| UP_SEQ_FEATURE | DOMAIN:MGS-like | 3 | 0.001584 | Q9CWJ9, B2RQC6 | 45.48 | 0.08749466 |
| INTERPRO | IPR011607:MGS-like_dom | 3 | 0.001624 | Q9CWJ9, B2RQC6 | 44.90 | 0.13764638 |
| INTERPRO | IPR036914:MGS-like_dom_sf | 3 | 0.001624 | Q9CWJ9, B2RQC6 | 44.90 | 0.13764638 |
| SMART | SM00851:MGS | 3 | 0.001851 | Q9CWJ9, B2RQC6 | 41.87 | 0.11169763 |
| UP_KW_MOLECULAR_FUNCTION | KW-0511~Multifunctional enzyme | 5 | 0.069277 | Q9CWJ9, P23591, B2RQC6, E9Q555 | 3.22 | 0.27710689 |
| GOTERM_BP_DIRECT | GO:0031100~animal organ regeneration | 3 | 0.131544 | Q9CWJ9, B2RQC6 | 4.74 | 0.93639901 |
| Annotation Cluster 28 | Enrichment Score: 2.15386763692565 | | | | | |
| Category | Term | Count | PValue | Genes | Fold Enrichment | FDR |
| GOTERM_BP_DIRECT | GO:0002218~activation of innate immune response | 7 | 7.384E-05 | Q3TBT3, Q61107, Q9CZW5, Q9R002, Q9Z0E6, D0QMC3, P0DOV2 | 9.89 | 0.00729694 |
| UP_SEQ_FEATURE | DOMAIN:HIN-200 | 3 | 0.013451 | Q9R002, D0QMC3, P0DOV2 | 16.54 | 0.38773625 |
| INTERPRO | IPR004021:HIN200/IF120x | 3 | 0.013782 | Q9R002, D0QMC3, P0DOV2 | 16.33 | 0.54964256 |
| INTERPRO | IPR040205:HIN-200 | 3 | 0.019121 | Q9R002, D0QMC3, P0DOV2 | 13.82 | 0.63239108 |
| GOTERM_MF_DIRECT | GO:0003690~double-stranded DNA binding | 6 | 0.064986 | P42225, Q9R002, Q8BFV2, D0QMC3, Q91XB0, P0DOV2 | 2.79 | 0.40418262 |
| Annotation Cluster 29 | Enrichment Score: 1.9646486115716821 | | | | | |
| Category | Term | Count | PValue | Genes | Fold Enrichment | FDR |
| GOTERM_CC_DIRECT | GO:0043202~lysosomal lumen | 6 | 2.173E-05 | O09159, O35114, P20060, P17439, P29416, O35405 | 17.31 | 0.00042554 |
| GOTERM_BP_DIRECT | GO:0007040~lysosome organization | 5 | 0.004497 | Q9Z0M5, P20060, P17439, P29416, O89023 | 7.46 | 0.13001299 |
| KEGG_PATHWAY | mmu00511:Other glycan degradation | 4 | 0.009308 | O09159, P20060, P17439, P29416 | 8.93 | 0.09455644 |
| GOTERM_BP_DIRECT | GO:0019915~lipid storage | 4 | 0.009944 | Q9Z0M5, P20060, P17439, P29416 | 8.95 | 0.22832468 |
| INTERPRO | IPR017853:Glycoside_hydrolase_SF | 4 | 0.06379 | P20060, P10852, P17439, P29416 | 4.35 | 0.74057427 |
| UP_KW_MOLECULAR_FUNCTION | KW-0326~Glycosidase | 5 | 0.077158 | O09159, P08905, P20060, P17439, P29416 | 3.10 | 0.29148701 |
| KEGG_PATHWAY | mmu00600:Sphingolipid metabolism | 3 | 0.397178 | P20060, P17439, P29416 | 2.19 | 0.76376094 |
| Annotation Cluster 30 | Enrichment Score: 1.8387567028804936 | | | | | |
| Category | Term | Count | PValue | Genes | Fold Enrichment | FDR |
| INTERPRO | IPR043129:ATPase_NBD | 7 | 0.000627 | P17710, Q61699, Q3TRM8, P38647, Q9QZ08, O08528, P63017 | 6.65 | 0.10194182 |
| GOTERM_BP_DIRECT | GO:0051085~chaperone cofactor-dependent protein ref | 5 | 0.0008 | Q61699, Q9QYJ3, P38647, P63017, Q8R180 | 11.84 | 0.04165925 |
| GOTERM_BP_DIRECT | GO:0006457~protein folding | 7 | 0.005511 | Q61699, Q9QYJ3, P38647, P09103, Q9ERE7, P63017, Q8R180 | 4.37 | 0.14835326 |
| INTERPRO | IPR029047:HSP70_peptide-bd_sf | 3 | 0.016357 | Q61699, P38647, P63017 | 14.97 | 0.5994799 |
| INTERPRO | IPR018181:Heat_shock_70_CS | 3 | 0.019121 | Q61699, P38647, P63017 | 13.82 | 0.63239108 |
| INTERPRO | IPR029048:HSP70_C_sf | 3 | 0.019121 | Q61699, P38647, P63017 | 13.82 | 0.63239108 |
| INTERPRO | IPR013126:Hsp_70_fam | 3 | 0.019121 | Q61699, P38647, P63017 | 13.82 | 0.63239108 |
| KEGG_PATHWAY | mmu04141:Protein processing in endoplasmic reticulum | 10 | 0.029974 | P57759, Q61699, Q9WTX6, Q9QYJ3, Q8R2E9, P09103, Q99P31, P63017, Q8R180, Q91YW3 | 2.30 | 0.20919559 |
| GOTERM_MF_DIRECT | GO:0044183~protein folding chaperone | 4 | 0.035347 | Q9QYJ3, P38647, Q9ERE7, P63017 | 5.55 | 0.3010346 |
| GOTERM_MF_DIRECT | GO:0140662~ATP-dependent protein folding chaperone | 3 | 0.054784 | Q61699, P38647, P63017 | 7.90 | 0.38449196 |
| GOTERM_MF_DIRECT | GO:0051082~unfolded protein binding | 3 | 0.323679 | Q9QYJ3, P38647, P63017 | 2.58 | 0.94211823 |
| Annotation Cluster 31 | Enrichment Score: 1.8329855974408014 | | | | | |
| Category | Term | Count | PValue | Genes | Fold Enrichment | FDR |
| GOTERM_BP_DIRECT | GO:0001913~T cell mediated cytotoxicity | 7 | 2.928E-07 | P23591, P36371, P06339, P01902, P01901, P01900, P97821, P14426 | 24.51 | 8.5726E-05 |
| GOTERM_CC_DIRECT | GO:0030670~phagocytic vesicle membrane | 9 | 3.711E-06 | P06339, Q60766, P01902, Q9CZE3, P21958, P01901, P01900, P61027, A1L314, P14426 | 9.82 | 8.7218E-05 |
| GOTERM_BP_DIRECT | GO:0002485~antigen processing and presentation of er | 4 | 7.469E-06 | P36371, P01902, P01901, P01900, P14426 | 80.52 | 0.00159029 |
| GOTERM_BP_DIRECT | GO:0001916~positive regulation of T cell mediated cyto | 7 | 7.384E-05 | P23492, P36371, P06339, P01902, P01901, P01900, P14426, P63017 | 9.89 | 0.00729694 |
| GOTERM_MF_DIRECT | GO:0042288~MHC class I protein binding | 6 | 0.000202 | P36371, P06339, P01902, P21958, P01901, P01900, P14426 | 11.03 | 0.00572681 |

| | | | | | | |
|---|---|---|---|---|---|---|
| GOTERM_BP_DIRECT | GO:0042590~antigen processing and presentation of ex | 4 | 0.000382 | P26151, P01902, P01901, A1L314 | 26.84 | 0.0241521 |
| GOTERM_CC_DIRECT | GO:0098553~lumenal side of endoplasmic reticulum m | 5 | 0.000553 | P06339, P01902, P01901, P01900, P14426, Q9D8V0 | 13.03 | 0.00604247 |
| GOTERM_CC_DIRECT | GO:0042824~MHC class I peptide loading complex | 4 | 0.000761 | P36371, P01902, P21958, P01901, P01900, P14426 | 21.54 | 0.00745043 |
| GOTERM_BP_DIRECT | GO:0010977~negative regulation of neuron projection | 7 | 0.000954 | Q9Z0F8, P01902, P01901, P01900, Q8BTM8, P20152, P14426, Q99P72 | 6.19 | 0.04380303 |
| UP_SEQ_FEATURE | REGION:Alpha-3 | 4 | 0.001404 | P06339, P01902, P01901, P01900, P14426 | 17.32 | 0.08749466 |
| GOTERM_MF_DIRECT | GO:0046978~TAP1 binding | 4 | 0.001413 | P36371, P01902, P21958, P01901, P01900, P14426 | 17.56 | 0.02914593 |
| GOTERM_MF_DIRECT | GO:0046979~TAP2 binding | 4 | 0.001413 | P36371, P01902, P21958, P01901, P01900, P14426 | 17.56 | 0.02914593 |
| UP_SEQ_FEATURE | DOMAIN:MHC class I alpha chain C-terminal | 3 | 0.001584 | P01902, P01901, P01900, P14426 | 45.48 | 0.08749466 |
| UP_SEQ_FEATURE | REGION:Alpha-1 | 4 | 0.002997 | P06339, P01902, P01901, P01900, P14426 | 13.47 | 0.14193653 |
| UP_SEQ_FEATURE | REGION:Alpha-2 | 4 | 0.002997 | P06339, P01902, P01901, P01900, P14426 | 13.47 | 0.14193653 |
| GOTERM_MF_DIRECT | GO:0042605~peptide antigen binding | 6 | 0.004258 | P36371, P06339, P01902, P21958, P01901, P01900, P14426 | 5.64 | 0.06250767 |
| GOTERM_BP_DIRECT | GO:0002474~antigen processing and presentation of pe | 4 | 0.004421 | P06339, P01902, P01901, P01900, P14426 | 11.93 | 0.12943476 |
| UP_SEQ_FEATURE | DOMAIN:MHC class I-like antigen recognition-like | 4 | 0.00471 | P06339, P01902, P01901, P01900, P14426 | 11.55 | 0.18926402 |
| INTERPRO | IPR010579:MHC_I_a_C | 3 | 0.005499 | P01902, P01901, P01900, P14426 | 25.66 | 0.26630648 |
| KEGG_PATHWAY | mmu04612:Antigen processing and presentation | 8 | 0.005707 | P36371, P06339, P01902, P21958, P01901, P01900, O70370, P14426, P63017 | 3.70 | 0.06643506 |
| UP_KW_CELLULAR_COMPONENT | KW-0490~MHC I | 4 | 0.010593 | P06339, P01902, P01901, P01900, P14426 | 8.65 | 0.0436957 |
| GOTERM_CC_DIRECT | GO:0042612~MHC class I protein complex | 4 | 0.012286 | P06339, P01902, P01901, P01900, P14426 | 8.28 | 0.07597792 |
| GOTERM_BP_DIRECT | GO:0042270~protection from natural killer cell mediated | 3 | 0.01444 | P36371, P06339, P21958 | 16.10 | 0.30195969 |
| GOTERM_CC_DIRECT | GO:0009897~external side of plasma membrane | 14 | 0.020243 | P05555, P06339, P03975, P11438, P26151, P10810, P42082, P14426, Q62351, P15379, P21956, P01902, P01901, P01900, P09103 | 2.05 | 0.11599387 |
| UP_SEQ_FEATURE | DOMAIN:Ig-like C1-type | 4 | 0.021103 | P06339, P01902, P01901, P01900, P14426 | 6.74 | 0.50281011 |
| UP_SEQ_FEATURE | REGION:Connecting peptide | 4 | 0.024343 | P06339, P01902, P01901, P01900, P14426 | 6.38 | 0.50281011 |
| GOTERM_CC_DIRECT | GO:0032398~MHC class Ib protein complex | 3 | 0.040664 | P06339, P01902, P01901, P01900, P14426 | 9.32 | 0.19502014 |
| INTERPRO | IPR001039:MHC_I_a_a1/a2 | 4 | 0.041107 | P06339, P01902, P01901, P01900, P14426 | 5.21 | 0.64083499 |
| INTERPRO | IPR011161:MHC_I-like_Ag-recog | 4 | 0.048127 | P06339, P01902, P01901, P01900, P14426 | 4.89 | 0.64083499 |
| GOTERM_MF_DIRECT | GO:0030881~beta-2-microglobulin binding | 3 | 0.048389 | P06339, P01902, P01901, P01900, P14426 | 8.47 | 0.35747272 |
| GOTERM_BP_DIRECT | GO:0048839~inner ear development | 4 | 0.060157 | P06339, P01902, P01901, P0DOV2 | 4.47 | 0.64484194 |
| KEGG_PATHWAY | mmu05330:Allograft rejection | 5 | 0.061032 | P06339, P01902, P01901, P01900, P14426, P42082 | 3.35 | 0.29862132 |
| KEGG_PATHWAY | mmu05332:Graft-versus-host disease | 5 | 0.061032 | P06339, P01902, P01901, P01900, P14426, P42082 | 3.35 | 0.29862132 |
| INTERPRO | IPR050208:MHC_class-I_related | 4 | 0.06379 | P06339, P01902, P01901, P01900, P14426 | 4.35 | 0.74057427 |
| INTERPRO | IPR037055:MHC_I-like_Ag-recog_sf | 4 | 0.075373 | P06339, P01902, P01901, P01900, P14426 | 4.06 | 0.81532007 |
| GOTERM_MF_DIRECT | GO:0042608~T cell receptor binding | 3 | 0.075564 | P06339, P01902, P01901, P01900, P14426 | 6.59 | 0.45517081 |
| GOTERM_MF_DIRECT | GO:0005102~signaling receptor binding | 10 | 0.079115 | P06339, P09405, P26041, Q99LX0, P01902, P01901, P01900, O70309, P17439, P14426, P42082, P63017 | 1.91 | 0.46916909 |
| KEGG_PATHWAY | mmu04940:Type I diabetes mellitus | 5 | 0.084173 | P06339, P01902, P01901, P01900, P14426, P42082 | 3.00 | 0.36288578 |
| KEGG_PATHWAY | mmu04218:Cellular senescence | 9 | 0.084762 | Q60972, P06339, Q63932, Q60930, Q8BTI9, P01902, P01901, P01900, P14426, P70227 | 1.98 | 0.36288578 |
| KEGG_PATHWAY | mmu05166:Human T-cell leukemia virus 1 infection | 11 | 0.087031 | P06339, Q63932, Q60930, Q8BTI9, P97333, P01902, P01901, P01900, Q9WVM3, P14426, Q80WQ2, O88351 | 1.79 | 0.36687111 |
| GOTERM_BP_DIRECT | GO:0002476~antigen processing and presentation of er | 3 | 0.087755 | P06339, P01902, P01901, P01900, P14426 | 6.04 | 0.76687129 |
| GOTERM_BP_DIRECT | GO:0002486~antigen processing and presentation of er | 3 | 0.087755 | P06339, P01902, P01901, P01900, P14426 | 6.04 | 0.76687129 |
| GOTERM_MF_DIRECT | GO:0046703~natural killer cell lectin-like receptor bindir | 3 | 0.102359 | P06339, P01902, P01901, P01900, P14426 | 5.51 | 0.55535357 |
| KEGG_PATHWAY | mmu05203:Viral carcinogenesis | 10 | 0.114182 | P06339, Q7TPR4, Q8BTI9, P01902, O35892, P01901, P01900, P14426, Q80WQ2, A2AN08, P60766 | 1.76 | 0.40134219 |
| KEGG_PATHWAY | mmu05320:Autoimmune thyroid disease | 5 | 0.119261 | P06339, P01902, P01901, P01900, P14426, P42082 | 2.64 | 0.40846752 |
| INTERPRO | IPR011162:MHC_I/II-like_Ag-recog | 4 | 0.122272 | P06339, P01902, P01901, P01900, P14426 | 3.28 | 0.98905908 |
| GOTERM_BP_DIRECT | GO:0006955~immune response | 10 | 0.132875 | P23492, P06339, Q9BDB7, P01902, P01901, P01900, Q8BV66, P14426, P42082, O88188, Q91XB0 | 1.71 | 0.94300648 |
| KEGG_PATHWAY | mmu05416:Viral myocarditis | 5 | 0.204136 | P06339, P01902, P01901, P01900, P14426, P42082 | 2.14 | 0.56677534 |
| INTERPRO | IPR003006:Ig/MHC_CS | 4 | 0.219673 | P06339, P01902, P01901, P01900, P14426 | 2.47 | 0.98905908 |
| INTERPRO | IPR003597:Ig_C1-set | 4 | 0.304806 | P06339, P01902, P01901, P01900, P14426 | 2.06 | 0.98905908 |
| SMART | SM00407:IGc1 | 4 | 0.331918 | P06339, P01902, P01901, P01900, P14426 | 1.96 | 1 |
| KEGG_PATHWAY | mmu04514:Cell adhesion molecules | 6 | 0.456224 | P05555, P06339, P01902, P01901, P01900, P14426, P42082 | 1.35 | 0.82785038 |
| UP_SEQ_FEATURE | DOMAIN:Ig-like | 6 | 0.997771 | P06339, P26151, P01902, P01901, P01900, P14426, P42082 | 0.44 | 0.99777086 |
| INTERPRO | IPR007110:Ig-like_dom | 6 | 0.999824 | P06339, P26151, P01902, P01901, P01900, P14426, P42082 | 0.36 | 0.99982407 |
| INTERPRO | IPR013783:Ig-like_fold | 9 | 0.999842 | Q99P91, P06339, Q80X90, P26151, P01902, P01901, P01900, Q8BTM8, P14426, P42082 | 0.41 | 0.99984238 |
| INTERPRO | IPR036179:Ig-like_dom_sf | 6 | 0.999918 | P06339, P26151, P01902, P01901, P01900, P14426, P42082 | 0.34 | 0.99991754 |

Annotation Cluster 32 Enrichment Score: 1.8139470590844817

| Category | Term | Count | PValue | Genes | Fold Enrichment | FDR |
|---|---|---|---|---|---|---|
| GOTERM_CC_DIRECT | GO:0005681~spliceosomal complex | 9 | 0.000551 | O89086, P09405, Q8CGC6, Q61656, Q52KI8, Q5NC05, Q62189, P63017, P26369 | 4.88 | 0.00604247 |
| UP_SEQ_FEATURE | DOMAIN:RRM 3 | 5 | 0.005359 | P09405, Q8CGC6, Q9CW46, P52912, P26369 | 7.05 | 0.19757378 |
| INTERPRO | IPR035979:RBD_domain_sf | 11 | 0.014981 | O89086, P09405, Q8CGC6, Q9CW46, D3YXK2, Q8C166, P52912, Q62189, O35309, Q9JIK5, P26369 | 2.46 | 0.57203633 |
| UP_SEQ_FEATURE | DOMAIN:RRM 1 | 6 | 0.016495 | P09405, Q8CGC6, Q9CW46, P52912, Q62189, P26369 | 4.04 | 0.45566531 |
| UP_SEQ_FEATURE | DOMAIN:RRM 2 | 6 | 0.016495 | P09405, Q8CGC6, Q9CW46, P52912, Q62189, P26369 | 4.04 | 0.45566531 |
| UP_SEQ_FEATURE | DOMAIN:RRM | 9 | 0.040973 | O89086, P09405, Q8CGC6, Q9CW46, D3YXK2, Q8C166, P52912, Q62189, P26369 | 2.32 | 0.50281011 |
| INTERPRO | IPR012677:Nucleotide-bd_a/b_plait_sf | 10 | 0.042128 | O89086, P09405, Q8CGC6, Q9CW46, D3YXK2, Q8C166, P52912, Q62189, O35309, P26369 | 2.18 | 0.64083499 |
| INTERPRO | IPR000504:RRM_dom | 9 | 0.047059 | O89086, P09405, Q8CGC6, Q9CW46, D3YXK2, Q8C166, P52912, Q62189, P26369 | 2.26 | 0.64083499 |

| | | | | | | |
|---|---|---|---|---|---|---|
| SMART | SM00360:RRM | 9 | 0.04836 | O89086, P09405, Q8CGC6, Q9CW46, D3YXK2, Q8C166, P52912, Q62189, P26369 | 2.23 | 1 |
| Annotation Cluster 33 | Enrichment Score: 1.6664946416821684 | | | | | |
| Category | Term | Count | PValue | Genes | Fold Enrichment | FDR |
| GOTERM_BP_DIRECT | GO:0051301~cell division | 13 | 0.002824 | Q6P9P6, Q07832, Q9WVM3, Q65Z40, P48193, Q3TCJ1, O08585, Q9JLQ0, Q3UMY5, P42208, Q922S8, Q8C3Y4, P60766 | 2.78 | 0.10019392 |
| GOTERM_CC_DIRECT | GO:0072686~mitotic spindle | 7 | 0.00856 | Q6P9P6, P83741, Q9WVM3, Q3UMY5, Q65Z40, P48193, P60766 | 3.98 | 0.05747198 |
| GOTERM_CC_DIRECT | GO:0005819~spindle | 7 | 0.013966 | Q6P9P6, O08585, Q6DVA0, Q07832, Q9WVM3, P42208, Q922S8 | 3.58 | 0.08524555 |
| UP_KW_BIOLOGICAL_PROCESS | KW-0498~Mitosis | 12 | 0.033253 | Q6P9P6, O08585, Q3TCJ1, Q9JLQ0, Q07832, Q9WVM3, Q3UMY5, P42208, Q65Z40, P48193, Q8C3Y4, Q922S8 | 2.04 | 0.21998296 |
| UP_KW_BIOLOGICAL_PROCESS | KW-0132~Cell division | 14 | 0.066247 | Q6P9P6, Q9WU78, Q07832, Q9WVM3, Q65Z40, P48193, Q3TCJ1, O08585, Q9JLQ0, Q3UMY5, P42208, Q922S8, Q8C3Y4, P60766 | 1.71 | 0.35607738 |
| UP_KW_BIOLOGICAL_PROCESS | KW-0131~Cell cycle | 19 | 0.134807 | Q60972, Q6P9P6, Q9ESL4, Q9WU78, Q07832, Q9WVM3, Q65Z40, Q61881, P48193, P49718, Q3TCJ1, O08585, Q9JLQ0, Q3UMY5, P42208, Q922S8, Q8C3Y4, P60766, E9PVX6 | 1.39 | 0.64407862 |
| Annotation Cluster 34 | Enrichment Score: 1.6554661838066584 | | | | | |
| Category | Term | Count | PValue | Genes | Fold Enrichment | FDR |
| GOTERM_BP_DIRECT | GO:0007229~integrin-mediated signaling pathway | 7 | 0.002521 | P05555, O88839, Q9Z0F8, P97333, O70309, O88351, P60766 | 5.12 | 0.09224571 |
| GOTERM_MF_DIRECT | GO:0005178~integrin binding | 8 | 0.004331 | P21956, P05555, O88839, Q99P91, Q7TPR4, Q9Z0F8, O70309, P09103 | 3.95 | 0.06250767 |
| GOTERM_BP_DIRECT | GO:0033627~cell adhesion mediated by integrin | 3 | 0.087755 | P05555, Q9Z0F8, O70309 | 6.04 | 0.76687129 |
| UP_KW_MOLECULAR_FUNCTION | KW-0401~Integrin | 3 | 0.249332 | P05555, Q9Z0F8, O70309 | 3.11 | 0.76331662 |
| Annotation Cluster 35 | Enrichment Score: 1.5689868309672443 | | | | | |
| Category | Term | Count | PValue | Genes | Fold Enrichment | FDR |
| INTERPRO | IPR003191:Guanylate-bd/ATL_C | 3 | 0.025183 | Q61107, Q9Z0E6, Q6PA06 | 11.97 | 0.64083499 |
| INTERPRO | IPR036543:Guanylate-bd_C_sf | 3 | 0.025183 | Q61107, Q9Z0E6, Q6PA06 | 11.97 | 0.64083499 |
| UP_SEQ_FEATURE | DOMAIN:GB1/RHD3-type G | 3 | 0.027805 | Q61107, Q9Z0E6, Q6PA06 | 11.37 | 0.50281011 |
| INTERPRO | IPR015894:Guanylate-bd_N | 3 | 0.028469 | Q61107, Q9Z0E6, Q6PA06 | 11.23 | 0.64083499 |
| INTERPRO | IPR030386:G_GB1_RHD3_dom | 3 | 0.028469 | Q61107, Q9Z0E6, Q6PA06 | 11.23 | 0.64083499 |
| Annotation Cluster 36 | Enrichment Score: 1.5687415894296417 | | | | | |
| Category | Term | Count | PValue | Genes | Fold Enrichment | FDR |
| GOTERM_MF_DIRECT | GO:0017124~SH3 domain binding | 7 | 0.006245 | Q8BH43, O88839, Q9WU78, Q9Z0F8, Q9JLQ0, Q3UIA2, P11352 | 4.26 | 0.08846872 |
| UP_SEQ_FEATURE | MOTIF:SH3-binding | 4 | 0.046644 | O88839, Q9Z0F8, Q9JLQ0, Q3UIA2 | 4.95 | 0.50281011 |
| UP_KW_DOMAIN | KW-0729~SH3-binding | 4 | 0.067524 | O88839, Q9Z0F8, Q9JLQ0, Q3UIA2 | 4.24 | 0.1755183 |
| Annotation Cluster 37 | Enrichment Score: 1.4913328175631615 | | | | | |
| Category | Term | Count | PValue | Genes | Fold Enrichment | FDR |
| GOTERM_BP_DIRECT | GO:0006418~tRNA aminoacylation for protein translatio | 4 | 0.000936 | Q91WQ3, Q8BP47, Q9CZD3, Q9ER72 | 20.13 | 0.04380303 |
| GOTERM_MF_DIRECT | GO:0004812~aminoacyl-tRNA ligase activity | 3 | 0.013074 | Q91WQ3, Q8BP47, Q9ER72 | 16.93 | 0.16668802 |
| UP_KW_MOLECULAR_FUNCTION | KW-0030~Aminoacyl-tRNA synthetase | 4 | 0.040469 | Q91WQ3, Q8BP47, Q9CZD3, Q9ER72 | 5.21 | 0.18345783 |
| COG_ONTOLOGY | Translation, ribosomal structure and biogenesis | 6 | 0.061915 | Q8R0Y6, Q8BP47, Q9CZD3, Q8K1J6, Q9QXK7, Q9ER72 | 2.73 | 0.39723439 |
| UP_KW_BIOLOGICAL_PROCESS | KW-0648~Protein biosynthesis | 6 | 0.163917 | Q91WQ3, Q8BP47, Q9CZD3, P59325, Q9ER72, Q8BX17 | 2.07 | 0.7016195 |
| KEGG_PATHWAY | mmu00970:Aminoacyl-tRNA biosynthesis | 4 | 0.22424 | Q91WQ3, Q8BP47, Q9CZD3, Q9ER72 | 2.44 | 0.59078623 |
| Annotation Cluster 38 | Enrichment Score: 1.4465657667315286 | | | | | |
| Category | Term | Count | PValue | Genes | Fold Enrichment | FDR |
| GOTERM_MF_DIRECT | GO:0004029~aldehyde dehydrogenase (NAD+) activity | 3 | 0.019059 | Q8R0Y6, P47738, Q80VQ0 | 13.95 | 0.22430602 |
| UP_SEQ_FEATURE | DOMAIN:Aldehyde dehydrogenase | 3 | 0.027805 | Q8R0Y6, P47738, Q80VQ0 | 11.37 | 0.50281011 |
| INTERPRO | IPR029510:Ald_DH_CS_GLU | 3 | 0.031915 | Q8R0Y6, P47738, Q80VQ0 | 10.56 | 0.64083499 |
| INTERPRO | IPR016160:Ald_DH_CS_CYS | 3 | 0.031915 | Q8R0Y6, P47738, Q80VQ0 | 10.56 | 0.64083499 |
| INTERPRO | IPR016163:Ald_DH_C | 3 | 0.047187 | Q8R0Y6, P47738, Q80VQ0 | 8.55 | 0.64083499 |
| INTERPRO | IPR015590:Aldehyde_DH_dom | 3 | 0.047187 | Q8R0Y6, P47738, Q80VQ0 | 8.55 | 0.64083499 |
| INTERPRO | IPR016161:Ald_DH/histidinol_DH | 3 | 0.047187 | Q8R0Y6, P47738, Q80VQ0 | 8.55 | 0.64083499 |
| INTERPRO | IPR016162:Ald_DH_N | 3 | 0.047187 | Q8R0Y6, P47738, Q80VQ0 | 8.55 | 0.64083499 |
| Annotation Cluster 39 | Enrichment Score: 1.2823221788480705 | | | | | |
| Category | Term | Count | PValue | Genes | Fold Enrichment | FDR |
| GOTERM_CC_DIRECT | GO:0005789~endoplasmic reticulum membrane | 24 | 0.011409 | Q3TBT3, Q9EQH2, Q80X80, Q60766, Q8BYI6, Q8R2E9, Q3TZZ7, Q99P72, Q9D8V0, Q8R180, Q91XB0, P70227, P36371, P45878, O55022, P22437, O54984, P21958, Q9QUJ7, P61027, Q6ZQM8, O35405, Q8BG07 | 1.75 | 0.0714962 |
| UP_KW_CELLULAR_COMPONENT | KW-0256~Endoplasmic reticulum | 33 | 0.067799 | P11928, Q80X80, Q8BYI6, Q8R2E9, Q3TZZ7, Q80WQ2, Q91XB0, P70227, P45878, P36371, P22437, Q9WVJ3, O54984, Q9QUJ7, P61027, Q6ZQM8, O35405, Q3TBT3, P57759, Q9EQH2, Q99LX0, Q6PA06, Q99P72, Q8CGZ0, Q8R180, Q9D8V0, O55022, P21958, P09103, Q9ERE7, Q8BG07, Q91YW3 | 1.35 | 0.17210394 |
| UP_SEQ_FEATURE | TOPO_DOM:Lumenal | 15 | 0.183894 | Q3TBT3, Q9EQH2, Q80X80, P11438, Q8BYI6, Q6PA06, Q3TZZ7, Q99P72, Q9D8V0, P36371, O55022, O35114, P21958, O35405, Q8BG07 | 1.40 | 0.99251497 |

| Annotation Cluster 40 | Enrichment Score: 1.2300661269079218 | | | | | |
|---|---|---|---|---|---|---|
| Category | Term | Count | PValue | Genes | Fold Enrichment | FDR |
| INTERPRO | IPR011992:EF-hand-dom_pair | 12 | 0.005894 | Q99K51, P51432, Q7TPR4, Q9QXZ0, Q6A028, Q9CQ19, Q8BYI6, Q8CIH5, O88456, Q9EQP2, Q9QXX4, Q8BH59 | 2.66 | 0.27557567 |
| UP_SEQ_FEATURE | DOMAIN:EF-hand 2 | 8 | 0.020231 | Q99K51, Q7TPR4, Q9QXZ0, Q9CQ19, Q8BYI6, O88456, Q9QXX4, Q8BH59 | 2.92 | 0.50281011 |
| GOTERM_MF_DIRECT | GO:0005509~calcium ion binding | 17 | 0.022765 | Q99K51, P51432, Q7TPR4, P09405, Q9QXZ0, Q9CQ19, Q8BYI6, P14824, Q9QXX4, Q9JKF1, Q9EQP2, Q3TZZ7, P70227, O88456, Q8C166, P07356, Q8BH59 | 1.85 | 0.23837534 |
| INTERPRO | IPR018247:EF_Hand_1_Ca_BS | 8 | 0.030972 | Q99K51, Q7TPR4, Q9QXZ0, Q9CQ19, Q8BYI6, O88456, Q9EQP2, Q8BH59 | 2.68 | 0.64083499 |
| INTERPRO | IPR002048:EF_hand_dom | 9 | 0.035729 | Q99K51, Q7TPR4, Q9QXZ0, Q9CQ19, Q8BYI6, O88456, Q9EQP2, Q9QXX4, Q8BH59 | 2.39 | 0.64083499 |
| UP_SEQ_FEATURE | DOMAIN:EF-hand | 8 | 0.046384 | Q99K51, Q7TPR4, Q9QXZ0, Q9CQ19, Q8BYI6, O88456, Q9EQP2, Q9QXX4 | 2.45 | 0.50281011 |
| UP_SEQ_FEATURE | DOMAIN:EF-hand 1 | 7 | 0.05562 | Q99K51, Q7TPR4, Q9QXZ0, Q9CQ19, Q8BYI6, Q9QXX4, Q8BH59 | 2.57 | 0.56732304 |
| SMART | SM00054:EFh | 6 | 0.12221 | Q99K51, Q7TPR4, Q9QXZ0, Q9CQ19, Q8BYI6, Q8BH59 | 2.28 | 1 |
| UP_SEQ_FEATURE | DOMAIN:EF-hand 3 | 4 | 0.188997 | Q9CQ19, O88456, Q9QXX4, Q8BH59 | 2.67 | 0.99251497 |
| UP_SEQ_FEATURE | DOMAIN:EF-hand 4 | 3 | 0.265443 | O88456, Q9QXX4, Q8BH59 | 2.98 | 0.99251497 |
| UP_KW_LIGAND | KW-0106~Calcium | 24 | 0.620131 | Q99K51, P05555, P51432, Q7TPR4, P03975, Q9QXZ0, P97333, Q9CQ19, Q8BYI6, P14824, Q9QXX4, Q9EQP2, Q3TZZ7, P70227, O08585, Q8CIH5, O70309, O88456, Q6IRU2, Q8C166, P07356, Q8BH59, O89023 | 1.00 | 1 |

| Annotation Cluster 41 | Enrichment Score: 1.214808474280807 | | | | | |
|---|---|---|---|---|---|---|
| Category | Term | Count | PValue | Genes | Fold Enrichment | FDR |
| INTERPRO | IPR043563:Sp110/Sp140/Sp140L-like | 3 | 0.055633 | Q99388, Q8BVK9, O35892 | 7.81 | 0.71168881 |
| UP_SEQ_FEATURE | DOMAIN:HSR | 3 | 0.063133 | Q99388, Q8BVK9, O35892 | 7.28 | 0.61676064 |
| INTERPRO | IPR004865:HSR_dom | 3 | 0.064562 | Q99388, Q8BVK9, O35892 | 7.18 | 0.74057427 |

| Annotation Cluster 42 | Enrichment Score: 1.2011844101432607 | | | | | |
|---|---|---|---|---|---|---|
| Category | Term | Count | PValue | Genes | Fold Enrichment | FDR |
| UP_SEQ_FEATURE | REPEAT:TPR 3 | 7 | 0.018032 | Q9CYG7, O70145, Q9CZW5, Q9WVM3, Q0VGY8, Q80V86, Q91YW3 | 3.37 | 0.48797782 |
| UP_SEQ_FEATURE | REPEAT:TPR 1 | 7 | 0.02596 | Q9CYG7, O70145, Q9CZW5, Q9WVM3, Q0VGY8, Q80V86, Q91YW3 | 3.10 | 0.50281011 |
| UP_SEQ_FEATURE | REPEAT:TPR 2 | 7 | 0.02596 | Q9CYG7, O70145, Q9CZW5, Q9WVM3, Q0VGY8, Q80V86, Q91YW3 | 3.10 | 0.50281011 |
| UP_KW_DOMAIN | KW-0802~TPR repeat | 7 | 0.035098 | Q9CYG7, O70145, Q9CZW5, Q9WVM3, Q0VGY8, Q80V86, Q91YW3 | 2.87 | 0.15343521 |
| UP_SEQ_FEATURE | REPEAT:TPR | 5 | 0.053509 | O70145, Q9CZW5, Q9WVM3, Q0VGY8, Q91YW3 | 3.53 | 0.5500275 |
| UP_SEQ_FEATURE | REPEAT:TPR 4 | 5 | 0.061256 | Q9CYG7, Q9CZW5, Q9WVM3, Q80V86, Q91YW3 | 3.37 | 0.61533998 |
| INTERPRO | IPR019734:TPR_rpt | 6 | 0.063224 | Q9CYG7, O70145, Q9CZW5, Q9WVM3, Q0VGY8, Q91YW3 | 2.81 | 0.74057427 |
| SMART | SM00028:TPR | 6 | 0.065045 | Q9CYG7, O70145, Q9CZW5, Q9WVM3, Q0VGY8, Q91YW3 | 2.77 | 1 |
| INTERPRO | IPR011990:TPR-like_helical_dom_sf | 8 | 0.076705 | Q9CYG7, O70145, Q9CZW5, Q9WVM3, Q0VGY8, Q9D1H7, Q99LI7, Q91YW3 | 2.18 | 0.81532007 |
| UP_SEQ_FEATURE | REPEAT:TPR 9 | 3 | 0.081896 | Q9CZW5, Q9WVM3, Q91YW3 | 6.27 | 0.73873424 |
| UP_SEQ_FEATURE | REPEAT:TPR 6 | 4 | 0.09489 | Q9CYG7, Q9CZW5, Q9WVM3, Q91YW3 | 3.67 | 0.8065641 |
| UP_SEQ_FEATURE | REPEAT:TPR 5 | 4 | 0.108356 | Q9CYG7, Q9CZW5, Q9WVM3, Q91YW3 | 3.46 | 0.89723585 |
| UP_SEQ_FEATURE | REPEAT:TPR 8 | 3 | 0.174863 | Q9CZW5, Q9WVM3, Q91YW3 | 3.95 | 0.99251497 |
| UP_SEQ_FEATURE | REPEAT:TPR 7 | 3 | 0.234932 | Q9CZW5, Q9WVM3, Q91YW3 | 3.25 | 0.99251497 |

| Annotation Cluster 43 | Enrichment Score: 1.201165646267508 | | | | | |
|---|---|---|---|---|---|---|
| Category | Term | Count | PValue | Genes | Fold Enrichment | FDR |
| GOTERM_BP_DIRECT | GO:0002753~cytoplasmic pattern recognition receptor s | 3 | 0.01637 | Q61510, Q3TBT3, Q99J87 | 15.10 | 0.31683987 |
| GOTERM_BP_DIRECT | GO:0140374~antiviral innate immune response | 4 | 0.056201 | Q61510, Q3TBT3, P11928, Q99J87 | 4.60 | 0.64484194 |
| KEGG_PATHWAY | mmu04622:RIG-I-like receptor signaling pathway | 4 | 0.270843 | Q61510, Q3TBT3, O88351, Q99J87 | 2.20 | 0.63967199 |

| Annotation Cluster 44 | Enrichment Score: 1.1584993109731498 | | | | | |
|---|---|---|---|---|---|---|
| Category | Term | Count | PValue | Genes | Fold Enrichment | FDR |
| INTERPRO | IPR045851:AMP-bd_C_sf | 3 | 0.051347 | Q8VCW8, Q9QUJ7, Q9QXG4 | 8.16 | 0.66311253 |
| INTERPRO | IPR020845:AMP-binding_CS | 3 | 0.064562 | Q8VCW8, Q9QUJ7, Q9QXG4 | 7.18 | 0.74057427 |
| INTERPRO | IPR042099:ANL_N_sf | 3 | 0.083705 | Q8VCW8, Q9QUJ7, Q9QXG4 | 6.19 | 0.8345914 |
| INTERPRO | IPR000873:AMP-dep_synth/lig_dom | 3 | 0.083705 | Q8VCW8, Q9QUJ7, Q9QXG4 | 6.19 | 0.8345914 |

| Annotation Cluster 45 | Enrichment Score: 1.0751567752097964 | | | | | |
|---|---|---|---|---|---|---|
| Category | Term | Count | PValue | Genes | Fold Enrichment | FDR |
| GOTERM_MF_DIRECT | GO:0036121~double-stranded DNA helicase activity | 4 | 0.02524 | P49718, Q9EPU0, Q9WTM5, Q61881 | 6.32 | 0.23837534 |
| GOTERM_MF_DIRECT | GO:0061749~forked DNA-dependent helicase activity | 3 | 0.122841 | P49718, Q9WTM5, Q61881 | 4.94 | 0.59102718 |
| GOTERM_MF_DIRECT | GO:1990518~single-stranded 3'-5' DNA helicase activity | 3 | 0.127049 | P49718, Q9WTM5, Q61881 | 4.84 | 0.59995523 |
| GOTERM_MF_DIRECT | GO:0009378~four-way junction helicase activity | 3 | 0.127049 | P49718, Q9WTM5, Q61881 | 4.84 | 0.59995523 |

| Annotation Cluster 46 | Enrichment Score: 1.0737798779316425 | | | | | |
|---|---|---|---|---|---|---|
| Category | Term | Count | PValue | Genes | Fold Enrichment | FDR |
| GOTERM_BP_DIRECT | GO:0006260~DNA replication | 6 | 0.009238 | Q60972, P49718, Q9EPU0, P17918, Q61881, Q91XB0 | 4.69 | 0.22004419 |
| KEGG_PATHWAY | mmu03030:DNA replication | 3 | 0.223979 | P49718, P17918, Q61881 | 3.35 | 0.59078623 |
| UP_KW_BIOLOGICAL_PROCESS | KW-0235~DNA replication | 4 | 0.290326 | Q60972, P49718, P17918, Q61881 | 2.12 | 0.91489362 |

Annotation Cluster 47 — Enrichment Score: 1.0354186669405285

| Category | Term | Count | PValue | Genes | Fold Enrichment | FDR |
|---|---|---|---|---|---|---|
| GOTERM_BP_DIRECT | GO:0016485~protein processing | 5 | 0.035934 | Q8K411, P29452, Q9Z0F8, P29594, O70370 | 4.03 | 0.52091214 |
| GOTERM_MF_DIRECT | GO:0004197~cysteine-type endopeptidase activity | 4 | 0.045426 | P29452, P29594, O70370, P97821 | 5.02 | 0.34751016 |
| UP_KW_PTM | KW-0865~Zymogen | 11 | 0.079997 | O88839, P29452, Q9WVJ3, Q9Z0F8, Q99LX0, P29594, O70370, P97821, P18242, P29416, O89023 | 1.83 | 0.30378459 |
| UP_KW_MOLECULAR_FUNCTION | KW-0788~Thiol protease | 4 | 0.552634 | P29452, P29594, O70370, P97821 | 1.38 | 0.88311688 |

Annotation Cluster 48 — Enrichment Score: 1.0268360325744028

| Category | Term | Count | PValue | Genes | Fold Enrichment | FDR |
|---|---|---|---|---|---|---|
| UP_KW_DOMAIN | KW-0728~SH3 domain | 8 | 0.048453 | Q9QXZ0, O70145, Q9JLB0, Q80TY0, Q9JLQ0, Q8CIH5, O08539, P97814 | 2.41 | 0.15343521 |
| UP_SEQ_FEATURE | DOMAIN:SH3 | 8 | 0.06183 | Q9QXZ0, O70145, Q9JLB0, Q80TY0, Q9JLQ0, Q8CIH5, O08539, P97814 | 2.29 | 0.61644106 |
| INTERPRO | IPR001452:SH3_domain | 8 | 0.078214 | Q9QXZ0, O70145, Q9JLB0, Q80TY0, Q9JLQ0, Q8CIH5, O08539, P97814 | 2.17 | 0.81532007 |
| INTERPRO | IPR027267:AH/BAR_dom_sf | 4 | 0.137244 | Q80TY0, Q3UIA2, O08539, P97814 | 3.11 | 0.98905908 |
| SMART | SM00326:SH3 | 7 | 0.145181 | O70145, Q9JLB0, Q80TY0, Q9JLQ0, Q8CIH5, O08539, P97814 | 1.96 | 1 |
| INTERPRO | IPR036028:SH3-like_dom_sf | 7 | 0.147834 | O70145, Q9JLB0, Q80TY0, Q9JLQ0, Q8CIH5, O08539, P97814 | 1.97 | 0.98905908 |

Annotation Cluster 49 — Enrichment Score: 0.9685172613595215

| Category | Term | Count | PValue | Genes | Fold Enrichment | FDR |
|---|---|---|---|---|---|---|
| UP_SEQ_FEATURE | PROPEP:Removed in mature form | 11 | 0.019713 | Q9Z0M5, P63213, P03975, P10810, Q99LX0, Q80VQ0, Q9Z0E6, Q8R2Q8, P60766, O89023 | 2.35 | 0.50281011 |
| UP_SEQ_FEATURE | LIPID:GPI-anchor amidated asparagine | 3 | 0.042174 | P03975, P10810 | 9.10 | 0.50281011 |
| GOTERM_CC_DIRECT | GO:0098552~side of membrane | 4 | 0.2107 | P03975, P10810, Q8R2Q8 | 2.52 | 0.56266576 |
| UP_KW_PTM | KW-0336~GPI-anchor | 4 | 0.762897 | P03975, P10810, Q8R2Q8 | 1.00 | 0.96 |

Annotation Cluster 50 — Enrichment Score: 0.9341336683754576

| Category | Term | Count | PValue | Genes | Fold Enrichment | FDR |
|---|---|---|---|---|---|---|
| GOTERM_BP_DIRECT | GO:0048015~phosphatidylinositol-mediated signaling | 4 | 0.002437 | P51432, Q8BTI9, Q8CIH5, E9Q3L2 | 14.64 | 0.09058148 |
| UP_SEQ_FEATURE | REGION:Activation loop | 4 | 0.004087 | Q2TBE6, Q8BTI9, Q07832, E9Q3L2 | 12.13 | 0.17088833 |
| UP_SEQ_FEATURE | REGION:Catalytic loop | 3 | 0.031175 | Q2TBE6, Q8BTI9, E9Q3L2 | 10.70 | 0.50281011 |
| UP_SEQ_FEATURE | REGION:G-loop | 3 | 0.031175 | Q2TBE6, Q8BTI9, E9Q3L2 | 10.70 | 0.50281011 |
| UP_SEQ_FEATURE | DOMAIN:PI3K/PI4K catalytic | 3 | 0.034697 | Q2TBE6, Q8BTI9, E9Q3L2 | 10.11 | 0.50281011 |
| INTERPRO | IPR000403:PI3/4_kinase_cat_dom | 3 | 0.035516 | Q2TBE6, Q8BTI9, E9Q3L2 | 9.98 | 0.64083499 |
| GOTERM_BP_DIRECT | GO:0046854~phosphatidylinositol phosphate biosynthet | 3 | 0.076734 | Q2TBE6, Q8BTI9, E9Q3L2 | 6.53 | 0.73221871 |
| KEGG_PATHWAY | mmu04070:Phosphatidylinositol signaling system | 6 | 0.089668 | Q2TBE6, P51432, Q8BTI9, Q8CIH5, E9Q3L2, P70227 | 2.51 | 0.37226005 |
| KEGG_PATHWAY | mmu00562:Inositol phosphate metabolism | 5 | 0.102966 | Q2TBE6, P51432, Q8BTI9, Q8CIH5, E9Q3L2 | 2.79 | 0.39736084 |
| KEGG_PATHWAY | mmu04935:Growth hormone synthesis, secretion and a | 6 | 0.164671 | P42225, P51432, Q63932, Q8BTI9, Q8CIH5, P70227 | 2.06 | 0.53082134 |
| KEGG_PATHWAY | mmu04929:GnRH secretion | 4 | 0.204784 | P51432, Q63932, Q8BTI9, P70227 | 2.55 | 0.56677534 |
| KEGG_PATHWAY | mmu04915:Estrogen signaling pathway | 6 | 0.238599 | P51432, Q63932, Q8BTI9, P18242, P63017, P70227 | 1.80 | 0.59978209 |
| KEGG_PATHWAY | mmu04933:AGE-RAGE signaling pathway in diabetic cc | 5 | 0.241013 | P42225, P51432, Q8BTI9, Q8CIH5, P60766 | 1.99 | 0.6003427 |
| KEGG_PATHWAY | mmu04725:Cholinergic synapse | 5 | 0.30716 | P51432, Q6PHZ2, P63213, Q8BTI9, P70227 | 1.78 | 0.67329568 |
| KEGG_PATHWAY | mmu04611:Platelet activation | 5 | 0.380557 | P51432, P22437, Q8BTI9, Q8CIH5, P70227 | 1.59 | 0.75016283 |
| KEGG_PATHWAY | mmu04750:Inflammatory mediator regulation of TRP ch | 5 | 0.403088 | P51432, Q6PHZ2, Q8BTI9, Q8CIH5, P70227 | 1.55 | 0.76401458 |
| KEGG_PATHWAY | mmu04022:cGMP-PKG signaling pathway | 5 | 0.620884 | P51432, Q63932, Q60930, Q9CQ19, P70227 | 1.17 | 0.91333333 |
| KEGG_PATHWAY | mmu05017:Spinocerebellar ataxia | 4 | 0.683498 | P51432, Q60930, Q8BTI9, P70227 | 1.14 | 0.91333333 |
| KEGG_PATHWAY | mmu04072:Phospholipase D signaling pathway | 4 | 0.719036 | P51432, Q63932, Q8BTI9, Q8CIH5 | 1.08 | 0.91333333 |
| KEGG_PATHWAY | mmu04020:Calcium signaling pathway | 5 | 0.881498 | P51432, Q6PHZ2, Q60930, Q8CIH5, P70227 | 0.79 | 0.91333333 |

Annotation Cluster 51 — Enrichment Score: 0.922980168363563

| Category | Term | Count | PValue | Genes | Fold Enrichment | FDR |
|---|---|---|---|---|---|---|
| INTERPRO | IPR035892:C2_domain_sf | 7 | 0.066103 | P51432, Q80X80, Q8BTI9, Q8CIH5, Q8C166, Q3TZZ7, Q8BRN9 | 2.47 | 0.74695972 |
| INTERPRO | IPR000008:C2_dom | 6 | 0.091306 | P51432, Q80X80, Q8CIH5, Q8C166, Q3TZZ7, Q8BRN9 | 2.51 | 0.88436286 |
| UP_SEQ_FEATURE | DOMAIN:C2 | 5 | 0.176373 | P51432, Q80X80, Q8CIH5, Q8C166, Q8BRN9 | 2.28 | 0.99251497 |
| SMART | SM00239:C2 | 5 | 0.190954 | P51432, Q8CIH5, Q8C166, Q3TZZ7, Q8BRN9 | 2.20 | 1 |

Annotation Cluster 52 — Enrichment Score: 0.9018293541275922

| Category | Term | Count | PValue | Genes | Fold Enrichment | FDR |
|---|---|---|---|---|---|---|
| KEGG_PATHWAY | mmu05020:Prion disease | 13 | 0.039104 | Q62425, Q9CXV1, Q60930, Q8BTI9, O70145, Q3UX10, Q9CPQ1, P70227, P19536, Q9CQA3, P33175, P63017, P98086 | 1.91 | 0.23810074 |
| KEGG_PATHWAY | mmu05415:Diabetic cardiomyopathy | 11 | 0.043458 | P19536, P51432, Q9CQA3, Q62425, Q6PHZ2, Q9CXV1, Q60930, Q8BTI9, O70145, P18242, Q9CPQ1 | 2.04 | 0.24807477 |
| GOTERM_CC_DIRECT | GO:0045277~respiratory chain complex IV | 3 | 0.043566 | P19536, Q62425, Q9CPQ1 | 8.97 | 0.20682928 |
| KEGG_PATHWAY | mmu05010:Alzheimer disease | 16 | 0.060839 | P51432, Q62425, Q9CXV1, Q63932, Q60930, Q8BTI9, Q78IQ7, Q3UX10, Q99P72, Q9CPQ1, P70227, P19536, Q9CQA3, Q9Z0F8, P33175, O88351 | 1.65 | 0.29862132 |
| KEGG_PATHWAY | mmu05012:Parkinson disease | 12 | 0.078618 | P19536, Q9CQA3, Q62425, Q6PHZ2, Q9CXV1, Q60930, Q99LX0, P33175, Q78IQ7, Q3UX10, Q9CPQ1, P70227 | 1.76 | 0.35893768 |
| KEGG_PATHWAY | mmu04932:Non-alcoholic fatty liver disease | 8 | 0.107719 | P19536, Q9CQA3, Q62425, Q9CXV1, Q8BTI9, Q9CPQ1, O88351, P60766 | 1.98 | 0.40134219 |

| | | | | | | |
|---|---|---|---|---|---|---|
| KEGG_PATHWAY | mmu05208:Chemical carcinogenesis - reactive oxygen | 10 | 0.115277 | P19536, Q9CQA3, Q62425, Q9CXV1, Q63932, Q60930, Q8BTI9, O70145, Q9CPQ1, O88351 | 1.75 | 0.40134219 |
| KEGG_PATHWAY | mmu05016:Huntington disease | 11 | 0.236723 | P19536, P51432, Q9CQA3, Q62425, Q9CXV1, Q60930, P33175, O08585, Q3UX10, P11352, Q9CPQ1 | 1.44 | 0.59978209 |
| KEGG_PATHWAY | mmu00190:Oxidative phosphorylation | 6 | 0.271387 | P19536, Q9CQA3, Q62425, Q9CXV1, Q8BVE3, Q9CPQ1 | 1.71 | 0.63967199 |
| KEGG_PATHWAY | mmu05022:Pathways of neurodegeneration - multiple d | 15 | 0.303626 | P51432, Q62425, Q9CXV1, Q6PHZ2, Q63932, Q60930, Q99LX0, Q9WU84, Q3UX10, P11352, Q9CPQ1, P70227, P19536, Q9CQA3, P33175 | 1.25 | 0.67091555 |
| KEGG_PATHWAY | mmu05014:Amyotrophic lateral sclerosis | 12 | 0.327975 | P19536, Q9CQA3, Q62425, Q9CXV1, P29452, P33175, Q6PFD9, Q9WU84, Q3UX10, P11352, Q9CPQ1, P70227 | 1.29 | 0.69662851 |
| KEGG_PATHWAY | mmu04714:Thermogenesis | 7 | 0.535622 | P19536, Q9CQA3, Q62425, Q9CXV1, Q9QUJ7, A2BH40, Q9CPQ1 | 1.19 | 0.91254945 |

Annotation Cluster 53 — Enrichment Score: 0.8873237387204946

| Category | Term | Count | PValue | Genes | Fold Enrichment | FDR |
|---|---|---|---|---|---|---|
| KEGG_PATHWAY | mmu05164:Influenza A | 13 | 0.001282 | Q61510, P11928, Q8R4K2, Q63932, P29452, Q8BTI9, Q6PFD9, Q9WVL2, P42225, Q9QYJ3, P52293, O88351, Q91YW3 | 3.00 | 0.02065855 |
| KEGG_PATHWAY | mmu04620:Toll-like receptor signaling pathway | 8 | 0.014597 | P42225, Q8R4K2, Q63932, P10810, Q8BTI9, Q9WVL2, P42082, O88351 | 3.09 | 0.12902039 |
| KEGG_PATHWAY | mmu05171:Coronavirus disease - COVID-19 | 15 | 0.037616 | Q3TBT3, P97461, P11928, P35979, Q8R4K2, P29452, P97333, Q8BTI9, Q9WVL2, P42225, P67984, Q9Z0F8, Q8CIH5, P98086, O88351 | 1.80 | 0.23424671 |
| KEGG_PATHWAY | mmu04380:Osteoclast differentiation | 8 | 0.052116 | P42225, P26151, Q8BTI9, O70145, Q8CIH5, Q9WVL2, O88351, P70227 | 2.36 | 0.28199135 |
| KEGG_PATHWAY | mmu04625:C-type lectin receptor signaling pathway | 7 | 0.059465 | P42225, P29452, Q8BTI9, Q8CIH5, Q9WVL2, O88351, P70227 | 2.51 | 0.29862132 |
| KEGG_PATHWAY | mmu05162:Measles | 8 | 0.068694 | P42225, P11928, Q8R4K2, P26041, Q8BTI9, Q9WVL2, P63017, O88351 | 2.22 | 0.33021308 |
| KEGG_PATHWAY | mmu05235:PD-L1 expression and PD-1 checkpoint patl | 5 | 0.174018 | P42225, Q63932, Q8BTI9, Q3UMY5, O88351 | 2.28 | 0.54182755 |
| KEGG_PATHWAY | mmu04062:Chemokine signaling pathway | 8 | 0.202855 | P42225, P51432, P63213, Q8BTI9, Q8CIH5, Q9WVL2, O88351, P60766 | 1.67 | 0.56677534 |
| KEGG_PATHWAY | mmu05161:Hepatitis B | 7 | 0.221659 | P42225, Q8R4K2, Q63932, Q8BTI9, Q9WVL2, P17918, O88351 | 1.72 | 0.59078623 |
| KEGG_PATHWAY | mmu05160:Hepatitis C | 7 | 0.233733 | P42225, P11928, Q63932, Q8BTI9, Q9WVL2, O88351, Q76MZ3 | 1.68 | 0.59853086 |
| KEGG_PATHWAY | mmu04917:Prolactin signaling pathway | 3 | 0.551072 | P42225, Q63932, Q8BTI9 | 1.63 | 0.91333333 |
| KEGG_PATHWAY | mmu05220:Chronic myeloid leukemia | 3 | 0.565602 | Q63932, Q8BTI9, O88351 | 1.59 | 0.91333333 |
| KEGG_PATHWAY | mmu05215:Prostate cancer | 3 | 0.713563 | Q63932, Q8BTI9, O88351 | 1.21 | 0.91333333 |
| BIOCARTA | m_mapkPathway:MAPKinase Signaling Pathway | 3 | 0.865805 | P42225, Q63932, O88351 | 0.87 | 1 |
| KEGG_PATHWAY | mmu04630:JAK-STAT signaling pathway | 3 | 0.928524 | P42225, Q8BTI9, Q9WVL2 | 0.71 | 0.92852377 |

Annotation Cluster 54 — Enrichment Score: 0.855712337638276

| Category | Term | Count | PValue | Genes | Fold Enrichment | FDR |
|---|---|---|---|---|---|---|
| INTERPRO | IPR036322:WD40_repeat_dom_sf | 12 | 0.015187 | Q5DQR4, Q58A65, Q91WQ5, Q60972, Q8R5L3, Q920I9, Q3UMY5, Q9DCE5, A2AN08, Q8C3Y4, Q8BMQ2, Q8BX17 | 2.32 | 0.57203633 |
| UP_SEQ_FEATURE | REPEAT:WD 9 | 4 | 0.022691 | Q5DQR4, Q920I9, Q3UMY5, Q8BX17 | 6.56 | 0.50281011 |
| UP_SEQ_FEATURE | REPEAT:WD 8 | 4 | 0.05657 | Q5DQR4, Q920I9, Q3UMY5, Q8BX17 | 4.58 | 0.57260998 |
| UP_SEQ_FEATURE | REPEAT:WD 13 | 3 | 0.063133 | Q5DQR4, Q3UMY5, Q8BX17 | 7.28 | 0.61676064 |
| UP_SEQ_FEATURE | REPEAT:WD 12 | 3 | 0.081896 | Q5DQR4, Q3UMY5, Q8BX17 | 6.27 | 0.73873424 |
| INTERPRO | IPR019775:WD40_repeat_CS | 6 | 0.085245 | Q5DQR4, Q91WQ5, Q60972, Q920I9, Q9DCE5, Q8BX17 | 2.57 | 0.84373474 |
| UP_SEQ_FEATURE | REPEAT:WD 11 | 3 | 0.096937 | Q5DQR4, Q3UMY5, Q8BX17 | 5.68 | 0.8187137 |
| UP_SEQ_FEATURE | REPEAT:WD 10 | 3 | 0.102112 | Q5DQR4, Q3UMY5, Q8BX17 | 5.51 | 0.85696694 |
| INTERPRO | IPR011047:Quinoprotein_ADH-like_sf | 3 | 0.166439 | Q920I9, Q3UMY5, Q8BX17 | 4.08 | 0.98905908 |
| UP_SEQ_FEATURE | REPEAT:WD 5 | 7 | 0.172378 | Q5DQR4, Q91WQ5, Q60972, Q920I9, Q3UMY5, Q9DCE5, Q8BX17 | 1.87 | 0.99251497 |
| UP_SEQ_FEATURE | REPEAT:WD | 6 | 0.187222 | Q5DQR4, Q91WQ5, Q920I9, Q3UMY5, Q9DCE5, Q8BX17 | 1.98 | 0.99251497 |
| INTERPRO | IPR020472:G-protein_beta_WD-40_rep | 4 | 0.19811 | Q91WQ5, Q60972, Q9DCE5, Q8BX17 | 2.60 | 0.98905908 |
| UP_SEQ_FEATURE | REPEAT:WD 6 | 6 | 0.198644 | Q5DQR4, Q91WQ5, Q60972, Q920I9, Q3UMY5, Q8BX17 | 1.94 | 0.99251497 |
| UP_SEQ_FEATURE | REPEAT:WD 4 | 7 | 0.207204 | Q5DQR4, Q91WQ5, Q60972, Q920I9, Q3UMY5, Q9DCE5, Q8BX17 | 1.76 | 0.99251497 |
| UP_SEQ_FEATURE | REPEAT:WD 3 | 7 | 0.238799 | Q5DQR4, Q91WQ5, Q60972, Q920I9, Q3UMY5, Q9DCE5, Q8BX17 | 1.68 | 0.99251497 |
| UP_KW_DOMAIN | KW-0853~WD repeat | 7 | 0.243952 | Q5DQR4, Q91WQ5, Q60972, Q920I9, Q3UMY5, Q9DCE5, Q8BX17 | 1.66 | 0.35654465 |
| INTERPRO | IPR015943:WD40/YVTN_repeat-like_dom_sf | 9 | 0.248489 | Q5DQR4, Q58A65, Q91WQ5, Q60972, Q920I9, Q3UMY5, Q9DCE5, Q8BMQ2, Q8BX17 | 1.52 | 0.98905908 |
| UP_SEQ_FEATURE | REPEAT:WD 1 | 7 | 0.249621 | Q5DQR4, Q91WQ5, Q60972, Q920I9, Q3UMY5, Q9DCE5, Q8BX17 | 1.65 | 0.99251497 |
| UP_SEQ_FEATURE | REPEAT:WD 2 | 7 | 0.249621 | Q5DQR4, Q91WQ5, Q60972, Q920I9, Q3UMY5, Q9DCE5, Q8BX17 | 1.65 | 0.99251497 |
| INTERPRO | IPR001680:WD40_rpt | 7 | 0.293274 | Q5DQR4, Q91WQ5, Q60972, Q920I9, Q3UMY5, Q9DCE5, Q8BX17 | 1.56 | 0.98905908 |
| SMART | SM00320:WD40 | 7 | 0.340829 | Q5DQR4, Q91WQ5, Q60972, Q920I9, Q3UMY5, Q9DCE5, Q8BX17 | 1.46 | 1 |
| UP_SEQ_FEATURE | REPEAT:WD 7 | 4 | 0.44481 | Q5DQR4, Q920I9, Q3UMY5, Q8BX17 | 1.63 | 0.99251497 |

Annotation Cluster 55 — Enrichment Score: 0.8188008733173322

| Category | Term | Count | PValue | Genes | Fold Enrichment | FDR |
|---|---|---|---|---|---|---|
| KEGG_PATHWAY | mmu04666:Fc gamma R-mediated phagocytosis | 6 | 0.083688 | Q8BH43, P26151, Q8BTI9, Q8CIH5, O08539, P60766 | 2.57 | 0.36288578 |
| KEGG_PATHWAY | mmu04370:VEGF signaling pathway | 4 | 0.173338 | Q63932, Q8BTI9, Q8CIH5, P60766 | 2.77 | 0.54182755 |
| KEGG_PATHWAY | mmu04933:AGE-RAGE signaling pathway in diabetic cc | 5 | 0.241013 | P42225, P51432, Q8BTI9, Q8CIH5, P60766 | 1.99 | 0.6003427 |

Annotation Cluster 56 — Enrichment Score: 0.7960781375602481

| Category | Term | Count | PValue | Genes | Fold Enrichment | FDR |
|---|---|---|---|---|---|---|
| GOTERM_CC_DIRECT | GO:0005819~spindle | 7 | 0.013966 | Q6P9P6, O08585, Q6DVA0, Q07832, Q9WVM3, P42208, Q922S8 | 3.58 | 0.08524555 |
| GOTERM_CC_DIRECT | GO:0000775~chromosome, centromeric region | 4 | 0.063775 | Q07832, Q922S8, Q76MZ3, E9PVX6 | 4.37 | 0.26762817 |
| UP_KW_CELLULAR_COMPONENT | KW-0137~Centromere | 5 | 0.279357 | Q07832, P42208, Q8C3Y4, Q922S8, Q76MZ3 | 1.86 | 0.57441868 |
| UP_KW_CELLULAR_COMPONENT | KW-0995~Kinetochore | 4 | 0.295913 | Q07832, P42208, Q8C3Y4, Q922S8 | 2.10 | 0.57441868 |

| | | | | | | |
|---|---|---|---|---|---|---|
| GOTERM_CC_DIRECT | GO:0000776~kinetochore | 4 | 0.320321 | Q6PFD9, Q07832, P42208, Q922S8 | 2.01 | 0.72729882 |
| UP_KW_CELLULAR_COMPONENT | KW-0158~Chromosome | 11 | 0.709423 | Q60972, P49718, Q07832, P42208, Q8CG72, Q65Z40, Q61881, Q8C3Y4, Q922S8, Q76MZ3, E9PVX6 | 0.96 | 0.86842105 |
| Annotation Cluster 57 | Enrichment Score: 0.7697830964214164 | | | | | |
| Category | Term | Count | PValue | Genes | Fold Enrichment | FDR |
| KEGG_PATHWAY | mmu05221:Acute myeloid leukemia | 6 | 0.029439 | P05555, Q63932, P26151, P10810, Q8BTI9, O88351 | 3.44 | 0.20919559 |
| KEGG_PATHWAY | mmu04640:Hematopoietic cell lineage | 5 | 0.204136 | P05555, P26151, P10810, Q62351, P15379 | 2.14 | 0.56677534 |
| KEGG_PATHWAY | mmu05202:Transcriptional misregulation in cancer | 5 | 0.816207 | P05555, Q61656, P26151, P10810, P42082 | 0.89 | 0.91333333 |
| Annotation Cluster 58 | Enrichment Score: 0.7636937029389453 | | | | | |
| Category | Term | Count | PValue | Genes | Fold Enrichment | FDR |
| GOTERM_BP_DIRECT | GO:0006631~fatty acid metabolic process | 5 | 0.059376 | Q9DBL1, Q8VCW8, Q9Z0M5, Q9QUJ7, P45952 | 3.41 | 0.64484194 |
| KEGG_PATHWAY | mmu00071:Fatty acid degradation | 4 | 0.137703 | Q9DBL1, P47738, Q9QUJ7, P45952 | 3.09 | 0.45458603 |
| KEGG_PATHWAY | mmu01212:Fatty acid metabolism | 4 | 0.198388 | Q9DBL1, P50171, Q9QUJ7, P45952 | 2.59 | 0.56677534 |
| UP_KW_BIOLOGICAL_PROCESS | KW-0276~Fatty acid metabolism | 6 | 0.236082 | Q9DBL1, Q8VCW8, P22437, P50171, Q9QUJ7, P45952 | 1.81 | 0.84596112 |
| COG_ONTOLOGY | Lipid metabolism | 4 | 0.396641 | Q9DBL1, O70310, Q9QUJ7, P45952 | 1.73 | 0.92474306 |
| Annotation Cluster 59 | Enrichment Score: 0.7311058851613546 | | | | | |
| Category | Term | Count | PValue | Genes | Fold Enrichment | FDR |
| GOTERM_CC_DIRECT | GO:0000139~Golgi membrane | 13 | 0.080209 | Q2TBE6, Q3TBT3, Q60766, Q61107, Q8BYI6, Q9Z0E6, P14426, P01902, P01901, P01900, O88512, P61027, Q99KH8, O35405, P60766 | 1.71 | 0.31942085 |
| GOTERM_CC_DIRECT | GO:0005794~Golgi apparatus | 20 | 0.120877 | Q2TBE6, Q3TBT3, Q63932, Q9QXZ0, Q60766, P10810, Q61107, Q9Z0E6, P17439, P14426, P15379, Q8BVL3, O35643, P22437, Q9WVJ3, P01902, P01901, P01900, P61027, Q99KH8, Q8R2Q8, O89023 | 1.41 | 0.41129154 |
| UP_KW_CELLULAR_COMPONENT | KW-0333~Golgi apparatus | 17 | 0.660871 | Q2TBE6, Q3TBT3, Q9QXZ0, Q60766, P10810, Q61107, Q8BYI6, Q9Z0E6, Q80V94, Q8R180, O35643, Q9WVJ3, O88512, P61027, Q8R2Q8, O35405, Q8BG07 | 0.98 | 0.86842105 |
| Annotation Cluster 60 | Enrichment Score: 0.7120953386999649 | | | | | |
| Category | Term | Count | PValue | Genes | Fold Enrichment | FDR |
| UP_SEQ_FEATURE | REPEAT:HEAT 5 | 3 | 0.12352 | Q80WQ2, Q80X82, Q76MZ3 | 4.92 | 0.98668679 |
| UP_SEQ_FEATURE | REPEAT:HEAT 4 | 3 | 0.174863 | Q80WQ2, Q80X82, Q76MZ3 | 3.95 | 0.99251497 |
| UP_SEQ_FEATURE | REPEAT:HEAT 3 | 3 | 0.198643 | Q80WQ2, Q80X82, Q76MZ3 | 3.64 | 0.99251497 |
| UP_SEQ_FEATURE | REPEAT:HEAT 1 | 3 | 0.253226 | Q80WQ2, Q80X82, Q76MZ3 | 3.08 | 0.99251497 |
| UP_SEQ_FEATURE | REPEAT:HEAT 2 | 3 | 0.253226 | Q80WQ2, Q80X82, Q76MZ3 | 3.08 | 0.99251497 |
| Annotation Cluster 61 | Enrichment Score: 0.7112166012491469 | | | | | |
| Category | Term | Count | PValue | Genes | Fold Enrichment | FDR |
| UP_SEQ_FEATURE | DOMAIN:ABC transporter | 3 | 0.174863 | P36371, P21958, Q6P542 | 3.95 | 0.99251497 |
| INTERPRO | IPR017871:ABC_transporter-like_CS | 3 | 0.190363 | P36371, P21958, Q6P542 | 3.74 | 0.98905908 |
| INTERPRO | IPR003439:ABC_transporter-like_ATP-bd | 3 | 0.220836 | P36371, P21958, Q6P542 | 3.39 | 0.98905908 |
| Annotation Cluster 62 | Enrichment Score: 0.703734495806643 | | | | | |
| Category | Term | Count | PValue | Genes | Fold Enrichment | FDR |
| UP_SEQ_FEATURE | LIPID:S-geranylgeranyl cysteine | 7 | 0.007478 | P11928, P63213, Q9CZE3, Q80VQ0, P61027, Q9Z0E6, P60766 | 4.08 | 0.26093386 |
| GOTERM_MF_DIRECT | GO:0003924~GTPase activity | 10 | 0.018952 | Q8VEH6, P63213, Q60766, Q61107, Q9CZE3, P61027, Q9Z0E6, Q6PA06, P42208, P60766 | 2.52 | 0.22430602 |
| GOTERM_MF_DIRECT | GO:0003925~G protein activity | 4 | 0.035347 | Q60766, P61027, Q9Z0E6, P60766 | 5.55 | 0.3010346 |
| UP_KW_PTM | KW-0636~Prenylation | 7 | 0.235339 | P11928, P63213, Q9CZE3, Q80VQ0, P61027, Q9Z0E6, P60766 | 1.68 | 0.70601673 |
| UP_SEQ_FEATURE | MOTIF:Effector region | 3 | 0.448327 | Q9CZE3, P61027, P60766 | 1.98 | 0.99251497 |
| SMART | SM00174:RHO | 3 | 0.656401 | Q9CZE3, P61027, P60766 | 1.34 | 1 |
| INTERPRO | IPR001806:Small_GTPase | 3 | 0.660572 | Q9CZE3, P61027, P60766 | 1.33 | 0.98905908 |
| SMART | SM00173:RAS | 3 | 0.689542 | Q9CZE3, P61027, P60766 | 1.26 | 1 |
| SMART | SM00175:RAB | 3 | 0.754556 | Q9CZE3, P61027, P60766 | 1.11 | 1 |
| INTERPRO | IPR005225:Small_GTP-bd | 3 | 0.769553 | Q9CZE3, P61027, P60766 | 1.08 | 0.98905908 |
| Annotation Cluster 63 | Enrichment Score: 0.6991743945624926 | | | | | |
| Category | Term | Count | PValue | Genes | Fold Enrichment | FDR |
| GOTERM_BP_DIRECT | GO:0006281~DNA repair | 10 | 0.010835 | P23492, Q60972, Q9EPU0, Q99LX0, Q6P9P0, Q5NC05, Q9WTM5, Q8CG72, Q9DBR1, Q3TGW2 | 2.77 | 0.24090457 |
| UP_KW_BIOLOGICAL_PROCESS | KW-0234~DNA repair | 6 | 0.846913 | Q60972, Q99LX0, Q6P9P0, Q9WTM5, P17918, Q8CG72 | 0.84 | 0.91489362 |
| UP_KW_BIOLOGICAL_PROCESS | KW-0227~DNA damage | 7 | 0.870564 | Q60972, Q99LX0, Q6P9P0, Q9WTM5, P17918, Q8CG72, O35658 | 0.80 | 0.91489362 |
| Annotation Cluster 64 | Enrichment Score: 0.6821228974800709 | | | | | |
| Category | Term | Count | PValue | Genes | Fold Enrichment | FDR |
| GOTERM_MF_DIRECT | GO:0008017~microtubule binding | 8 | 0.056371 | Q6P9P6, Q9QXZ0, P33175, Q3TCJ1, Q07832, Q3UMY5, Q62433, Q922S8 | 2.34 | 0.3858111 |
| GOTERM_MF_DIRECT | GO:0003777~microtubule motor activity | 3 | 0.144205 | Q6P9P6, P33175, Q922S8 | 4.47 | 0.65276065 |
| INTERPRO | IPR019821:Kinesin_motor_CS | 3 | 0.160544 | Q6P9P6, P33175, Q922S8 | 4.18 | 0.98905908 |

| | | | | | | |
|---|---|---|---|---|---|---|
| UP_SEQ_FEATURE | DOMAIN:Kinesin motor | 3 | 0.174863 | Q6P9P6, P33175, Q922S8 | 3.95 | 0.99251497 |
| INTERPRO | IPR001752:Kinesin_motor_dom | 3 | 0.178338 | Q6P9P6, P33175, Q922S8 | 3.90 | 0.98905908 |
| SMART | SM00129:KISc | 3 | 0.197554 | Q6P9P6, P33175, Q922S8 | 3.64 | 1 |
| GOTERM_BP_DIRECT | GO:0007018~microtubule-based movement | 3 | 0.256246 | Q6P9P6, P33175, Q922S8 | 3.06 | 0.97339983 |
| KEGG_PATHWAY | mmu04814:Motor proteins | 7 | 0.366194 | Q6P9P6, Q9CQ19, P33175, P47753, Q6IRU2, Q3UX10, Q922S8 | 1.42 | 0.73777235 |
| INTERPRO | IPR036961:Kinesin_motor_dom_sf | 3 | 0.420896 | Q6P9P6, P33175, Q922S8 | 2.09 | 0.98905908 |
| UP_KW_MOLECULAR_FUNCTION | KW-0505~Motor protein | 4 | 0.475301 | Q6P9P6, Q9CQ19, P33175, Q922S8 | 1.55 | 0.88311688 |
| Annotation Cluster 65 | Enrichment Score: 0.6818180664748091 | | | | | |
| Category | Term | Count | PValue | Genes | Fold Enrichment | FDR |
| INTERPRO | IPR001623:DnaJ_domain | 3 | 0.202491 | Q9QYJ3, Q9CQV7, Q91YW3 | 3.59 | 0.98905908 |
| UP_SEQ_FEATURE | DOMAIN:J | 3 | 0.204649 | Q9QYJ3, Q9CQV7, Q91YW3 | 3.57 | 0.99251497 |
| SMART | SM00271:DnaJ | 3 | 0.210606 | Q9QYJ3, Q9CQV7, Q91YW3 | 3.49 | 1 |
| INTERPRO | IPR036869:J_dom_sf | 3 | 0.214704 | Q9QYJ3, Q9CQV7, Q91YW3 | 3.45 | 0.98905908 |
| Annotation Cluster 66 | Enrichment Score: 0.6417027051871216 | | | | | |
| Category | Term | Count | PValue | Genes | Fold Enrichment | FDR |
| GOTERM_MF_DIRECT | GO:0071949~FAD binding | 3 | 0.114531 | Q8C0I1, Q8R2E9, Q8R180 | 5.15 | 0.57752717 |
| GOTERM_MF_DIRECT | GO:0050660~flavin adenine dinucleotide binding | 3 | 0.230395 | Q9DBL1, Q8C0I1, P45952 | 3.29 | 0.84100256 |
| UP_KW_LIGAND | KW-0285~Flavoprotein | 6 | 0.254488 | Q9DBL1, Q8C0I1, O35435, Q8R2E9, P45952, Q8R180 | 1.75 | 1 |
| UP_KW_LIGAND | KW-0274~FAD | 5 | 0.403762 | Q9DBL1, Q8C0I1, Q8R2E9, P45952, Q8R180 | 1.54 | 1 |
| Annotation Cluster 67 | Enrichment Score: 0.6051549307888381 | | | | | |
| Category | Term | Count | PValue | Genes | Fold Enrichment | FDR |
| KEGG_PATHWAY | mmu05142:Chagas disease | 6 | 0.112279 | P51432, Q8R4K2, Q8BTI9, P98086, O88351, Q76MZ3 | 2.34 | 0.40134219 |
| KEGG_PATHWAY | mmu04660:T cell receptor signaling pathway | 5 | 0.357945 | Q63932, Q8BTI9, O88351, P60766, Q76MZ3 | 1.65 | 0.72917566 |
| KEGG_PATHWAY | mmu04071:Sphingolipid signaling pathway | 5 | 0.380557 | P51432, Q63932, Q8BTI9, P18242, Q76MZ3 | 1.59 | 0.75016283 |
| Annotation Cluster 68 | Enrichment Score: 0.5787901654817043 | | | | | |
| Category | Term | Count | PValue | Genes | Fold Enrichment | FDR |
| KEGG_PATHWAY | mmu04625:C-type lectin receptor signaling pathway | 7 | 0.059465 | P42225, P29452, Q8BTI9, Q8CIH5, Q9WVL2, O88351, P70227 | 2.51 | 0.29862132 |
| UP_KW_DOMAIN | KW-0727~SH2 domain | 4 | 0.238987 | P42225, Q8CIH5, Q9WVL2, Q9JM90 | 2.36 | 0.35654465 |
| INTERPRO | IPR000980:SH2 | 4 | 0.273048 | P42225, Q8CIH5, Q9WVL2, Q9JM90 | 2.20 | 0.98905908 |
| INTERPRO | IPR036860:SH2_dom_sf | 4 | 0.295707 | P42225, Q8CIH5, Q9WVL2, Q9JM90 | 2.10 | 0.98905908 |
| UP_SEQ_FEATURE | DOMAIN:SH2 | 3 | 0.527567 | P42225, Q9WVL2, Q9JM90 | 1.70 | 0.99251497 |
| SMART | SM00252:SH2 | 3 | 0.556214 | Q8CIH5, Q9WVL2, Q9JM90 | 1.61 | 1 |
| Annotation Cluster 69 | Enrichment Score: 0.5574129432716092 | | | | | |
| Category | Term | Count | PValue | Genes | Fold Enrichment | FDR |
| INTERPRO | IPR000299:FERM_domain | 3 | 0.190363 | P26041, P48193, Q8BVL3 | 3.74 | 0.98905908 |
| UP_SEQ_FEATURE | DOMAIN:FERM | 3 | 0.198643 | P26041, P48193, Q8BVL3 | 3.64 | 0.99251497 |
| INTERPRO | IPR011993:PH-like_dom_sf | 8 | 0.562478 | Q8R1F1, P26041, Q6A028, Q9D1J1, Q8CIH5, Q9JM90, P48193, Q8BVL3 | 1.13 | 0.98905908 |
| Annotation Cluster 70 | Enrichment Score: 0.5350529887951055 | | | | | |
| Category | Term | Count | PValue | Genes | Fold Enrichment | FDR |
| KEGG_PATHWAY | mmu04620:Toll-like receptor signaling pathway | 8 | 0.014597 | P42225, Q8R4K2, Q63932, P10810, Q8BTI9, Q9WVL2, P42082, O88351 | 3.09 | 0.12902039 |
| KEGG_PATHWAY | mmu05221:Acute myeloid leukemia | 6 | 0.029439 | P05555, Q63932, P26151, P10810, Q8BTI9, O88351 | 3.44 | 0.20919559 |
| KEGG_PATHWAY | mmu05417:Lipid and atherosclerosis | 11 | 0.043458 | P51432, Q6PHZ2, Q8R4K2, P29452, P10810, Q8BTI9, O70145, P63017, Q8R180, O88351, P60766 | 2.04 | 0.24807477 |
| KEGG_PATHWAY | mmu04919:Thyroid hormone signaling pathway | 7 | 0.077465 | P42225, P12382, P51432, Q63932, Q8BTI9, Q8CIH5, Q9WUA3 | 2.34 | 0.35893768 |
| KEGG_PATHWAY | mmu04722:Neurotrophin signaling pathway | 7 | 0.079909 | Q6PHZ2, Q8R4K2, Q63932, Q8BTI9, Q8CIH5, O88351, P60766 | 2.32 | 0.35893768 |
| KEGG_PATHWAY | mmu05223:Non-small cell lung cancer | 5 | 0.102966 | Q63932, Q8BTI9, P33175, Q8CIH5, Q3UMY5 | 2.79 | 0.39736084 |
| KEGG_PATHWAY | mmu05135:Yersinia infection | 7 | 0.121629 | Q8BH43, Q8R4K2, Q63932, P29452, Q8BTI9, O88351, P60766 | 2.07 | 0.41143683 |
| KEGG_PATHWAY | mmu04662:B cell receptor signaling pathway | 5 | 0.154875 | P21855, Q63932, Q8BTI9, Q8CIH5, O88351 | 2.39 | 0.50518702 |
| KEGG_PATHWAY | mmu04935:Growth hormone synthesis, secretion and a | 6 | 0.164671 | P42225, P51432, Q63932, Q8BTI9, Q8CIH5, P70227 | 2.06 | 0.53082134 |
| KEGG_PATHWAY | mmu04370:VEGF signaling pathway | 4 | 0.173338 | Q63932, Q8BTI9, Q8CIH5, P60766 | 2.77 | 0.54182755 |
| KEGG_PATHWAY | mmu05235:PD-L1 expression and PD-1 checkpoint path | 5 | 0.174018 | P42225, Q63932, Q8BTI9, Q3UMY5, O88351 | 2.28 | 0.54182755 |
| KEGG_PATHWAY | mmu04062:Chemokine signaling pathway | 8 | 0.202855 | P42225, P51432, P63213, Q8BTI9, Q8CIH5, Q9WVL2, O88351, P60766 | 1.67 | 0.56677534 |
| KEGG_PATHWAY | mmu05161:Hepatitis B | 7 | 0.221659 | P42225, Q8R4K2, Q63932, Q8BTI9, Q9WVL2, P17918, O88351 | 1.72 | 0.59078623 |
| KEGG_PATHWAY | mmu05160:Hepatitis C | 7 | 0.233733 | P42225, P11928, Q63932, Q8BTI9, Q9WVL2, O88351, Q76MZ3 | 1.68 | 0.59853086 |
| KEGG_PATHWAY | mmu05211:Renal cell carcinoma | 4 | 0.250703 | Q63932, Q8BTI9, P60766, P97481 | 2.30 | 0.61332699 |
| KEGG_PATHWAY | mmu05214:Glioma | 4 | 0.284369 | Q6PHZ2, Q63932, Q8BTI9, Q8CIH5 | 2.14 | 0.63967199 |
| KEGG_PATHWAY | mmu05212:Pancreatic cancer | 4 | 0.291154 | P42225, Q8BTI9, O88351, P60766 | 2.12 | 0.64858802 |
| KEGG_PATHWAY | mmu04012:ErbB signaling pathway | 4 | 0.345658 | Q6PHZ2, Q63932, Q8BTI9, Q8CIH5 | 1.91 | 0.72675984 |

| | | | | | | |
|---|---|---|---|---|---|---|
| KEGG_PATHWAY | mmu04150:mTOR signaling pathway | 6 | 0.354136 | Q63932, Q8BTI9, Q8BVE3, Q9D1L9, P10852, O88351 | 1.53 | 0.72917566 |
| KEGG_PATHWAY | mmu04660:T cell receptor signaling pathway | 5 | 0.357945 | Q63932, Q8BTI9, O88351, P60766, Q76MZ3 | 1.65 | 0.72917566 |
| KEGG_PATHWAY | mmu04750:Inflammatory mediator regulation of TRP ch | 5 | 0.403088 | P51432, Q6PHZ2, Q8BTI9, Q8CIH5, P70227 | 1.55 | 0.76401458 |
| KEGG_PATHWAY | mmu04068:FoxO signaling pathway | 5 | 0.419893 | Q63932, Q8BTI9, Q07832, Q8BH04, O88351 | 1.51 | 0.77736956 |
| KEGG_PATHWAY | mmu04015:Rap1 signaling pathway | 7 | 0.438493 | P05555, Q9QZQ1, P51432, P46062, Q63932, Q8BTI9, P60766 | 1.31 | 0.80635679 |
| KEGG_PATHWAY | mmu04360:Axon guidance | 6 | 0.465781 | Q6PHZ2, Q8BTI9, Q9CQ19, P97333, Q8CIH5, P60766 | 1.33 | 0.83414334 |
| KEGG_PATHWAY | mmu04664:Fc epsilon RI signaling pathway | 3 | 0.489631 | Q63932, Q8BTI9, Q8CIH5 | 1.83 | 0.8599934 |
| KEGG_PATHWAY | mmu04014:Ras signaling pathway | 7 | 0.534099 | Q9QZQ1, Q63932, P63213, Q8BTI9, Q8CIH5, O88351, P60766 | 1.19 | 0.91254945 |
| KEGG_PATHWAY | mmu05220:Chronic myeloid leukemia | 3 | 0.565602 | Q63932, Q8BTI9, O88351 | 1.59 | 0.91333333 |
| KEGG_PATHWAY | mmu01521:EGFR tyrosine kinase inhibitor resistance | 3 | 0.593639 | Q63932, Q8BTI9, Q8CIH5 | 1.51 | 0.91333333 |
| KEGG_PATHWAY | mmu04010:MAPK signaling pathway | 8 | 0.614797 | Q9ESL4, Q8R4K2, Q63932, P10810, Q8BTM8, P63017, O88351, P60766 | 1.08 | 0.91333333 |
| KEGG_PATHWAY | mmu04926:Relaxin signaling pathway | 4 | 0.629374 | P51432, Q63932, P63213, Q8BTI9 | 1.24 | 0.91333333 |
| KEGG_PATHWAY | mmu05200:Pathways in cancer | 13 | 0.71277 | P51432, Q6PHZ2, Q63932, Q8BTI9, Q9WTX6, Q9WVL2, P42225, P63213, Q8CIH5, Q3UMY5, O88351, P60766, P97481 | 0.95 | 0.91333333 |
| KEGG_PATHWAY | mmu05215:Prostate cancer | 3 | 0.713563 | Q63932, Q8BTI9, O88351 | 1.21 | 0.91333333 |
| KEGG_PATHWAY | mmu04072:Phospholipase D signaling pathway | 4 | 0.719036 | P51432, Q63932, Q8BTI9, Q8CIH5 | 1.08 | 0.91333333 |
| KEGG_PATHWAY | mmu04650:Natural killer cell mediated cytotoxicity | 3 | 0.752553 | Q63932, Q8BTI9, Q8CIH5 | 1.12 | 0.91333333 |
| KEGG_PATHWAY | mmu04931:Insulin resistance | 3 | 0.761561 | Q8BTI9, Q8BH04, O88351 | 1.10 | 0.91333333 |
| KEGG_PATHWAY | mmu04024:cAMP signaling pathway | 5 | 0.810778 | Q9QZQ1, Q6PHZ2, Q63932, Q8BTI9, Q9CQ19 | 0.90 | 0.91333333 |
| KEGG_PATHWAY | mmu05225:Hepatocellular carcinoma | 4 | 0.813353 | Q63932, Q8BTI9, Q8CIH5, A2BH40 | 0.92 | 0.91333333 |
| KEGG_PATHWAY | mmu05206:MicroRNAs in cancer | 6 | 0.87571 | Q63932, Q8BTI9, Q8CIH5, P20152, O88351, P15379 | 0.80 | 0.91333333 |
| KEGG_PATHWAY | mmu04151:PI3K-Akt signaling pathway | 7 | 0.897401 | Q63932, P63213, Q8BTI9, O70309, Q8BH04, O88351, Q76MZ3 | 0.77 | 0.91333333 |

| Annotation Cluster 71 | Enrichment Score: 0.47904790856763374 | | | | | |
|---|---|---|---|---|---|---|
| Category | Term | Count | PValue | Genes | Fold Enrichment | FDR |
| KEGG_PATHWAY | mmu00565:Ether lipid metabolism | 4 | 0.115539 | Q8C0I1, Q8BYI6, O35405, Q8BG07 | 3.35 | 0.40134219 |
| UP_KW_BIOLOGICAL_PROCESS | KW-1208~Phospholipid metabolism | 3 | 0.440128 | Q8BYI6, O35405, Q8BG07 | 2.01 | 0.91489362 |
| KEGG_PATHWAY | mmu00564:Glycerophospholipid metabolism | 3 | 0.718703 | Q8BYI6, O35405, Q8BG07 | 1.19 | 0.91333333 |

| Annotation Cluster 72 | Enrichment Score: 0.4767349451524851 | | | | | |
|---|---|---|---|---|---|---|
| Category | Term | Count | PValue | Genes | Fold Enrichment | FDR |
| GOTERM_CC_DIRECT | GO:0005777~peroxisome | 5 | 0.090279 | Q8C0I1, Q3TBT3, Q9QUJ7, P38060, P20152 | 2.95 | 0.34218723 |
| KEGG_PATHWAY | mmu04146:Peroxisome | 3 | 0.639416 | Q8C0I1, Q9QUJ7, P38060 | 1.39 | 0.91333333 |
| UP_KW_CELLULAR_COMPONENT | KW-0576~Peroxisome | 3 | 0.643315 | Q8C0I1, Q9QUJ7, P38060 | 1.37 | 0.86842105 |

| Annotation Cluster 73 | Enrichment Score: 0.46146070387408056 | | | | | |
|---|---|---|---|---|---|---|
| Category | Term | Count | PValue | Genes | Fold Enrichment | FDR |
| GOTERM_BP_DIRECT | GO:0006412~translation | 7 | 0.070716 | O89086, P97461, P35979, P67984, Q8BP47, Q9ER72, Q8BX17 | 2.42 | 0.70314547 |
| GOTERM_BP_DIRECT | GO:0140236~translation at presynapse | 3 | 0.188021 | P97461, P35979, P67984 | 3.77 | 0.97339983 |
| GOTERM_BP_DIRECT | GO:0140242~translation at postsynapse | 3 | 0.192502 | P97461, P35979, P67984 | 3.72 | 0.97339983 |
| GOTERM_CC_DIRECT | GO:0005840~ribosome | 5 | 0.207126 | P97461, P11928, P35979, P67984, Q6P542 | 2.14 | 0.56266576 |
| GOTERM_BP_DIRECT | GO:0002181~cytoplasmic translation | 3 | 0.396137 | P97461, P35979, P67984 | 2.20 | 0.97339983 |
| GOTERM_CC_DIRECT | GO:0022626~cytosolic ribosome | 3 | 0.466269 | P97461, P35979, P67984 | 1.91 | 0.87037037 |
| GOTERM_MF_DIRECT | GO:0003735~structural constituent of ribosome | 3 | 0.79683 | P97461, P35979, P67984 | 1.01 | 0.94211823 |
| UP_KW_MOLECULAR_FUNCTION | KW-0689~Ribosomal protein | 3 | 0.909259 | P97461, P35979, P67984 | 0.75 | 0.90925904 |
| KEGG_PATHWAY | mmu03010:Ribosome | 3 | 0.990656 | P97461, P35979, P67984 | 0.45 | 0.99065578 |

| Annotation Cluster 74 | Enrichment Score: 0.45105522242746776 | | | | | |
|---|---|---|---|---|---|---|
| Category | Term | Count | PValue | Genes | Fold Enrichment | FDR |
| GOTERM_MF_DIRECT | GO:0051539~4 iron, 4 sulfur cluster binding | 3 | 0.106375 | Q9CQA3, Q99KI0, Q8WTY4 | 5.39 | 0.56827358 |
| UP_KW_LIGAND | KW-0004~4Fe-4S | 3 | 0.272447 | Q9CQA3, Q99KI0, Q8WTY4 | 2.92 | 1 |
| UP_KW_LIGAND | KW-0411~Iron-sulfur | 3 | 0.551844 | Q9CQA3, Q99KI0, Q8WTY4 | 1.63 | 1 |
| UP_KW_LIGAND | KW-0408~Iron | 6 | 0.981385 | O55022, Q9CQA3, P22437, Q9CXV1, Q99KI0, Q8WTY4 | 0.57 | 1 |

| Annotation Cluster 75 | Enrichment Score: 0.4409331349114943 | | | | | |
|---|---|---|---|---|---|---|
| Category | Term | Count | PValue | Genes | Fold Enrichment | FDR |
| GOTERM_BP_DIRECT | GO:0051209~release of sequestered calcium ion into cy | 6 | 0.000886 | P51432, Q8CIH5, Q8BTM8, Q8CGZ0, Q8R180, P70227 | 8.05 | 0.04380303 |
| KEGG_PATHWAY | mmu04922:Glucagon signaling pathway | 6 | 0.115715 | P12382, P51432, Q6PHZ2, Q9WUA3, Q8BH04, P70227 | 2.32 | 0.40134219 |
| KEGG_PATHWAY | mmu04935:Growth hormone synthesis, secretion and a | 6 | 0.164671 | P42225, P51432, Q63932, Q8BTI9, Q8CIH5, P70227 | 2.06 | 0.53082134 |
| KEGG_PATHWAY | mmu04912:GnRH signaling pathway | 5 | 0.183882 | P51432, Q6PHZ2, Q63932, P60766, P70227 | 2.23 | 0.55257081 |
| KEGG_PATHWAY | mmu04730:Long-term depression | 4 | 0.185751 | P51432, Q63932, Q76MZ3, P70227 | 2.68 | 0.55257081 |
| KEGG_PATHWAY | mmu04929:GnRH secretion | 4 | 0.204784 | P51432, Q63932, Q8BTI9, P70227 | 2.55 | 0.56677534 |
| KEGG_PATHWAY | mmu04720:Long-term potentiation | 4 | 0.230806 | P51432, Q6PHZ2, Q63932, P70227 | 2.40 | 0.59853086 |
| KEGG_PATHWAY | mmu04915:Estrogen signaling pathway | 6 | 0.238599 | P51432, Q63932, Q8BTI9, P18242, P63017, P70227 | 1.80 | 0.59978209 |

| | | | | | | |
|---|---|---|---|---|---|---|
| KEGG_PATHWAY | mmu04728:Dopaminergic synapse | 6 | 0.247857 | P51432, Q6PHZ2, P63213, P33175, Q76MZ3, P70227 | 1.77 | 0.61182716 |
| KEGG_PATHWAY | mmu04971:Gastric acid secretion | 4 | 0.284369 | P51432, Q6PHZ2, P00920, P70227 | 2.14 | 0.63967199 |
| KEGG_PATHWAY | mmu04725:Cholinergic synapse | 5 | 0.30716 | P51432, Q6PHZ2, P63213, Q8BTI9, P70227 | 1.78 | 0.67329568 |
| KEGG_PATHWAY | mmu04540:Gap junction | 4 | 0.359265 | P51432, Q63932, Q3UX10, P70227 | 1.87 | 0.72917566 |
| KEGG_PATHWAY | mmu04071:Sphingolipid signaling pathway | 5 | 0.380557 | P51432, Q63932, Q8BTI9, P18242, Q76MZ3 | 1.59 | 0.75016283 |
| KEGG_PATHWAY | mmu04611:Platelet activation | 5 | 0.380557 | P51432, P22437, Q8BTI9, Q8CIH5, P70227 | 1.59 | 0.75016283 |
| KEGG_PATHWAY | mmu04750:Inflammatory mediator regulation of TRP ch | 5 | 0.403088 | P51432, Q6PHZ2, Q8BTI9, Q8CIH5, P70227 | 1.55 | 0.76401458 |
| KEGG_PATHWAY | mmu04713:Circadian entrainment | 4 | 0.446129 | P51432, Q6PHZ2, P63213, P70227 | 1.62 | 0.81492845 |
| KEGG_PATHWAY | mmu04921:Oxytocin signaling pathway | 5 | 0.533155 | P51432, Q6PHZ2, Q63932, Q9CQ19, P70227 | 1.30 | 0.91254945 |
| KEGG_PATHWAY | mmu04022:cGMP-PKG signaling pathway | 5 | 0.620884 | P51432, Q63932, Q60930, Q9CQ19, P70227 | 1.17 | 0.91333333 |
| KEGG_PATHWAY | mmu04926:Relaxin signaling pathway | 4 | 0.629374 | P51432, Q63932, P63213, Q8BTI9 | 1.24 | 0.91333333 |
| KEGG_PATHWAY | mmu04911:Insulin secretion | 3 | 0.633132 | P51432, Q6PHZ2, P70227 | 1.40 | 0.91333333 |
| KEGG_PATHWAY | mmu04726:Serotonergic synapse | 4 | 0.659653 | P51432, P22437, P63213, P70227 | 1.18 | 0.91333333 |
| KEGG_PATHWAY | mmu04371:Apelin signaling pathway | 4 | 0.674112 | P51432, Q63932, P63213, P70227 | 1.16 | 0.91333333 |
| KEGG_PATHWAY | mmu04270:Vascular smooth muscle contraction | 4 | 0.6972 | P51432, Q63932, Q9CQ19, P70227 | 1.12 | 0.91333333 |
| KEGG_PATHWAY | mmu04916:Melanogenesis | 3 | 0.713563 | P51432, Q6PHZ2, Q63932 | 1.21 | 0.91333333 |
| KEGG_PATHWAY | mmu04072:Phospholipase D signaling pathway | 4 | 0.719036 | P51432, Q63932, Q8BTI9, Q8CIH5 | 1.08 | 0.91333333 |
| KEGG_PATHWAY | mmu04925:Aldosterone synthesis and secretion | 3 | 0.733662 | P51432, Q6PHZ2, P70227 | 1.16 | 0.91333333 |
| KEGG_PATHWAY | mmu04723:Retrograde endocannabinoid signaling | 4 | 0.735614 | P51432, Q62425, P63213, P70227 | 1.05 | 0.91333333 |
| KEGG_PATHWAY | mmu04934:Cushing syndrome | 4 | 0.770094 | P51432, Q6PHZ2, Q63932, P70227 | 0.99 | 0.91333333 |
| KEGG_PATHWAY | mmu04724:Glutamatergic synapse | 3 | 0.786902 | P51432, P63213, P70227 | 1.04 | 0.91333333 |
| KEGG_PATHWAY | mmu04020:Calcium signaling pathway | 5 | 0.881498 | P51432, Q6PHZ2, Q60930, Q8CIH5, P70227 | 0.79 | 0.91333333 |
| KEGG_PATHWAY | mmu04081:Hormone signaling | 3 | 0.9742 | P51432, P63213, Q8BTI9 | 0.55 | 0.97420032 |
| GOTERM_BP_DIRECT | GO:0007186~G protein-coupled receptor signaling path | 3 | 1 | P51432, P63213, P70227 | 0.15 | 0.99999998 |

| Annotation Cluster 76 | Enrichment Score: 0.3992300540593858 | | | | | |
|---|---|---|---|---|---|---|
| Category | Term | Count | PValue | Genes | Fold Enrichment | FDR |
| UP_SEQ_FEATURE | REPEAT:LRR 11 | 4 | 0.164518 | Q9QXZ0, P10810, Q6R5N8, Q91VI7 | 2.85 | 0.99251497 |
| UP_SEQ_FEATURE | REPEAT:LRR 15 | 3 | 0.192659 | Q9QXZ0, Q6R5N8, Q91VI7 | 3.71 | 0.99251497 |
| UP_SEQ_FEATURE | REPEAT:LRR 14 | 3 | 0.204649 | Q9QXZ0, Q6R5N8, Q91VI7 | 3.57 | 0.99251497 |
| UP_SEQ_FEATURE | REPEAT:LRR 13 | 3 | 0.247122 | Q9QXZ0, Q6R5N8, Q91VI7 | 3.14 | 0.99251497 |
| UP_SEQ_FEATURE | REPEAT:LRR 10 | 4 | 0.262379 | Q9QXZ0, P10810, Q6R5N8, Q91VI7 | 2.25 | 0.99251497 |
| UP_SEQ_FEATURE | REPEAT:LRR 7 | 5 | 0.306828 | Q505F5, Q9QXZ0, P10810, Q6R5N8, Q91VI7 | 1.78 | 0.99251497 |
| UP_SEQ_FEATURE | REPEAT:LRR 12 | 3 | 0.326276 | Q9QXZ0, Q6R5N8, Q91VI7 | 2.56 | 0.99251497 |
| UP_SEQ_FEATURE | REPEAT:LRR 9 | 4 | 0.378941 | Q9QXZ0, P10810, Q6R5N8, Q91VI7 | 1.81 | 0.99251497 |
| UP_SEQ_FEATURE | REPEAT:LRR 6 | 5 | 0.415135 | Q505F5, Q9QXZ0, P10810, Q6R5N8, Q91VI7 | 1.52 | 0.99251497 |
| UP_SEQ_FEATURE | REPEAT:LRR 8 | 4 | 0.440496 | Q9QXZ0, P10810, Q6R5N8, Q91VI7 | 1.64 | 0.99251497 |
| UP_SEQ_FEATURE | REPEAT:LRR 1 | 6 | 0.454832 | Q505F5, Q9QXZ0, P10810, Q6R5N8, Q91VI7, Q8K1T1 | 1.35 | 0.99251497 |
| UP_SEQ_FEATURE | REPEAT:LRR 5 | 5 | 0.512 | Q505F5, Q9QXZ0, P10810, Q6R5N8, Q91VI7 | 1.34 | 0.99251497 |
| UP_SEQ_FEATURE | REPEAT:LRR 4 | 5 | 0.565298 | Q505F5, Q9QXZ0, P10810, Q6R5N8, Q91VI7 | 1.25 | 0.99251497 |
| UP_SEQ_FEATURE | REPEAT:LRR 3 | 5 | 0.612738 | Q505F5, Q9QXZ0, P10810, Q6R5N8, Q91VI7 | 1.18 | 0.99251497 |
| INTERPRO | IPR001611:Leu-rich_rpt | 5 | 0.637769 | Q505F5, P10810, Q6R5N8, Q91VI7, Q8K1T1 | 1.14 | 0.98905908 |
| UP_SEQ_FEATURE | REPEAT:LRR 2 | 5 | 0.645427 | Q505F5, P10810, Q6R5N8, Q91VI7, Q8K1T1 | 1.13 | 0.99251497 |
| UP_KW_DOMAIN | KW-0433~Leucine-rich repeat | 6 | 0.735919 | Q505F5, Q9QXZ0, P10810, Q6R5N8, Q91VI7, Q8K1T1 | 0.98 | 0.82604209 |
| INTERPRO | IPR032675:LRR_dom_sf | 5 | 0.913711 | Q505F5, P10810, Q6R5N8, Q91VI7, Q8K1T1 | 0.73 | 0.98905908 |

| Annotation Cluster 77 | Enrichment Score: 0.3873132688711247 | | | | | |
|---|---|---|---|---|---|---|
| Category | Term | Count | PValue | Genes | Fold Enrichment | FDR |
| GOTERM_BP_DIRECT | GO:0070936~protein K48-linked ubiquitination | 5 | 0.034824 | Q61510, Q9WTX6, Q9WVM3, A2AN08, O35309 | 4.07 | 0.52091214 |
| GOTERM_BP_DIRECT | GO:0006511~ubiquitin-dependent protein catabolic proc | 5 | 0.335488 | Q61510, Q9WTX6, A2AN08, Q9D1H7, E9Q555 | 1.71 | 0.97339983 |
| GOTERM_MF_DIRECT | GO:0061630~ubiquitin protein ligase activity | 5 | 0.635367 | Q61510, Q9WTX6, A2AN08, P60766, E9Q555 | 1.15 | 0.94211823 |
| GOTERM_BP_DIRECT | GO:0016567~protein ubiquitination | 5 | 0.800685 | Q9WTX6, Q07832, Q9WVM3, A2AN08, E9Q555 | 0.91 | 0.97339983 |
| GOTERM_MF_DIRECT | GO:0004842~ubiquitin-protein transferase activity | 3 | 0.810041 | Q61510, A2AN08, E9Q555 | 0.98 | 0.94211823 |
| UP_KW_BIOLOGICAL_PROCESS | KW-0833~Ubl conjugation pathway | 9 | 0.985301 | Q61510, Q3TCJ1, Q9WTX6, Q9D906, Q9WVM3, Q9JIG7, Q9CR09, A2AN08, E9Q555 | 0.60 | 0.98530095 |

| Annotation Cluster 78 | Enrichment Score: 0.36639845764385204 | | | | | |
|---|---|---|---|---|---|---|
| Category | Term | Count | PValue | Genes | Fold Enrichment | FDR |
| SMART | SM00327:VWA | 3 | 0.381141 | P05555, O70309, Q8C166 | 2.26 | 1 |
| INTERPRO | IPR002035:VWF_A | 3 | 0.391791 | P05555, O70309, Q8C166 | 2.22 | 0.98905908 |
| UP_SEQ_FEATURE | DOMAIN:VWFA | 3 | 0.437203 | P05555, O70309, Q8C166 | 2.02 | 0.99251497 |
| INTERPRO | IPR036465:vWFA_dom_sf | 3 | 0.524305 | P05555, O70309, Q8C166 | 1.71 | 0.98905908 |

Annotation Cluster 79 Enrichment Score: 0.3576985098803392

| Category | Term | Count | PValue | Genes | Fold Enrichment | FDR |
|---|---|---|---|---|---|---|
| GOTERM_BP_DIRECT | GO:0006468~protein phosphorylation | 10 | 0.027636 | Q9DBC7, Q9ESL4, Q6PHZ2, Q63932, Q9JLQ0, Q07832, P83741, Q99KH8, O88351, P60766 | 2.35 | 0.43732096 |
| GOTERM_MF_DIRECT | GO:0106310~protein serine kinase activity | 10 | 0.044053 | Q9ESL4, Q6PHZ2, Q8R4K2, Q63932, Q8BTI9, Q07832, P83741, P09411, Q99KH8, O88351 | 2.16 | 0.34388047 |
| UP_KW_MOLECULAR_FUNCTION | KW-0418~Kinase | 21 | 0.062162 | Q2TBE6, Q9DBC7, Q9ESL4, Q8R4K2, Q6PHZ2, Q63932, Q3TRM8, Q8BTI9, Q07832, Q9R0N0, O08528, Q9WTP6, P17710, P12382, Q9QZ08, P83741, P09411, E9Q3L2, Q9WUA3, Q99KH8, O88351 | 1.51 | 0.26418661 |
| GOTERM_MF_DIRECT | GO:0004674~protein serine/threonine kinase activity | 9 | 0.126305 | Q9ESL4, Q6PHZ2, Q8R4K2, Q63932, Q07832, P83741, P09411, Q99KH8, O88351 | 1.81 | 0.59995523 |
| GOTERM_MF_DIRECT | GO:0004672~protein kinase activity | 7 | 0.143108 | P17710, Q6PHZ2, B2RQC6, Q07832, P83741, O88351 | 1.98 | 0.65165334 |
| GOTERM_BP_DIRECT | GO:0018105~peptidyl-serine phosphorylation | 4 | 0.144571 | Q6PHZ2, Q07832, P83741, O88351 | 3.04 | 0.97019648 |
| SMART | SM00220:S_TKc | 8 | 0.524443 | Q9ESL4, Q6PHZ2, Q8R4K2, Q63932, Q07832, P83741, Q99KH8, O88351 | 1.17 | 1 |
| INTERPRO | IPR011009:Kinase-like_dom_sf | 10 | 0.590349 | Q9ESL4, Q6PHZ2, Q8R4K2, Q63932, Q8BTI9, Q07832, P83741, E9Q3L2, Q99KH8, O88351 | 1.07 | 0.98905908 |
| GOTERM_BP_DIRECT | GO:0006338~chromatin remodeling | 9 | 0.592595 | Q60972, Q8R4K2, A2BH40, P83741, Q9WTM5, Q99KH8, O88351, Q8BMQ2, Q9JIK5 | 1.08 | 0.97339983 |
| GOTERM_MF_DIRECT | GO:0072354~histone H3T3 kinase activity | 4 | 0.621243 | Q8R4K2, P83741, Q99KH8, O88351 | 1.25 | 0.94211823 |
| GOTERM_MF_DIRECT | GO:0072371~histone H2AS121 kinase activity | 4 | 0.621243 | Q8R4K2, P83741, Q99KH8, O88351 | 1.25 | 0.94211823 |
| GOTERM_MF_DIRECT | GO:0072518~Rho-dependent protein serine/threonine k | 4 | 0.621243 | Q8R4K2, P83741, Q99KH8, O88351 | 1.25 | 0.94211823 |
| GOTERM_MF_DIRECT | GO:0004676~3-phosphoinositide-dependent protein kin | 4 | 0.621243 | Q8R4K2, P83741, Q99KH8, O88351 | 1.25 | 0.94211823 |
| GOTERM_MF_DIRECT | GO:0004677~DNA-dependent protein kinase activity | 4 | 0.621243 | Q8R4K2, P83741, Q99KH8, O88351 | 1.25 | 0.94211823 |
| GOTERM_MF_DIRECT | GO:0140823~histone H2BS36 kinase activity | 4 | 0.621243 | Q8R4K2, P83741, Q99KH8, O88351 | 1.25 | 0.94211823 |
| GOTERM_MF_DIRECT | GO:0004694~eukaryotic translation initiation factor 2alp | 4 | 0.621243 | Q8R4K2, P83741, Q99KH8, O88351 | 1.25 | 0.94211823 |
| GOTERM_MF_DIRECT | GO:0004711~ribosomal protein S6 kinase activity | 4 | 0.621243 | Q8R4K2, P83741, Q99KH8, O88351 | 1.25 | 0.94211823 |
| GOTERM_MF_DIRECT | GO:0140855~histone H3S57 kinase activity | 4 | 0.621243 | Q8R4K2, P83741, Q99KH8, O88351 | 1.25 | 0.94211823 |
| GOTERM_MF_DIRECT | GO:0035175~histone H3S10 kinase activity | 4 | 0.621243 | Q8R4K2, P83741, Q99KH8, O88351 | 1.25 | 0.94211823 |
| GOTERM_MF_DIRECT | GO:1990244~histone H2AT120 kinase activity | 4 | 0.621243 | Q8R4K2, P83741, Q99KH8, O88351 | 1.25 | 0.94211823 |
| GOTERM_MF_DIRECT | GO:0035979~histone H2AXS139 kinase activity | 4 | 0.621243 | Q8R4K2, P83741, Q99KH8, O88351 | 1.25 | 0.94211823 |
| GOTERM_MF_DIRECT | GO:0044022~histone H3S28 kinase activity | 4 | 0.621243 | Q8R4K2, P83741, Q99KH8, O88351 | 1.25 | 0.94211823 |
| GOTERM_MF_DIRECT | GO:0044023~histone H4S1 kinase activity | 4 | 0.621243 | Q8R4K2, P83741, Q99KH8, O88351 | 1.25 | 0.94211823 |
| GOTERM_MF_DIRECT | GO:0044024~histone H2AS1 kinase activity | 4 | 0.621243 | Q8R4K2, P83741, Q99KH8, O88351 | 1.25 | 0.94211823 |
| GOTERM_MF_DIRECT | GO:0044025~histone H2BS14 kinase activity | 4 | 0.621243 | Q8R4K2, P83741, Q99KH8, O88351 | 1.25 | 0.94211823 |
| GOTERM_MF_DIRECT | GO:0140857~histone H3T45 kinase activity | 4 | 0.623897 | Q8R4K2, P83741, Q99KH8, O88351 | 1.24 | 0.94211823 |
| GOTERM_MF_DIRECT | GO:0035402~histone H3T11 kinase activity | 4 | 0.626539 | Q8R4K2, P83741, Q99KH8, O88351 | 1.24 | 0.94211823 |
| GOTERM_MF_DIRECT | GO:0035403~histone H3T6 kinase activity | 4 | 0.626539 | Q8R4K2, P83741, Q99KH8, O88351 | 1.24 | 0.94211823 |
| GOTERM_MF_DIRECT | GO:0004679~AMP-activated protein kinase activity | 4 | 0.634389 | Q8R4K2, P83741, Q99KH8, O88351 | 1.23 | 0.94211823 |
| INTERPRO | IPR008271:Ser/Thr_kinase_AS | 6 | 0.653585 | Q9ESL4, Q6PHZ2, Q63932, Q07832, P83741, O88351 | 1.08 | 0.98905908 |
| UP_KW_MOLECULAR_FUNCTION | KW-0723~Serine/threonine-protein kinase | 8 | 0.684941 | Q9ESL4, Q6PHZ2, Q8R4K2, Q63932, Q07832, P83741, Q99KH8, O88351 | 1.00 | 0.88311688 |
| UP_SEQ_FEATURE | DOMAIN:Protein kinase | 8 | 0.758222 | Q9ESL4, Q6PHZ2, Q8R4K2, Q63932, Q07832, P83741, Q99KH8, O88351 | 0.93 | 0.99251497 |
| INTERPRO | IPR000719:Prot_kinase_dom | 8 | 0.761019 | Q9ESL4, Q6PHZ2, Q8R4K2, Q63932, Q07832, P83741, Q99KH8, O88351 | 0.93 | 0.98905908 |
| INTERPRO | IPR017441:Protein_kinase_ATP_BS | 4 | 0.961292 | Q6PHZ2, Q63932, Q07832, Q99KH8 | 0.61 | 0.98905908 |

Annotation Cluster 80 — Enrichment Score: 0.32286690482420355

| Category | Term | Count | PValue | Genes | Fold Enrichment | FDR |
|---|---|---|---|---|---|---|
| UP_SEQ_FEATURE | LIPID:S-palmitoyl cysteine | 8 | 0.215888 | Q2TBE6, Q3TBT3, Q9CQW9, Q60634, Q99LX0, Q99J93, Q80VQ0, Q62351 | 1.64 | 0.99251497 |
| UP_KW_PTM | KW-0564~Palmitate | 10 | 0.693284 | Q2TBE6, Q3TBT3, Q9CQW9, Q60766, Q60634, Q99LX0, O35114, Q99J93, Q80VQ0, Q62351 | 0.98 | 0.96 |
| UP_KW_PTM | KW-0449~Lipoprotein | 22 | 0.718223 | Q2TBE6, Q3TBT3, P11928, Q8R1F1, Q9CQW9, P03975, Q60766, Q60634, P10810, Q99LX0, Q99J93, Q80VQ0, Q9Z0E6, Q62351, P63213, O35114, Q9CZE3, P61027, Q8R2Q8, P60766, Q3TGW2 | 0.94 | 0.96 |

Annotation Cluster 81 — Enrichment Score: 0.26488115863988465

| Category | Term | Count | PValue | Genes | Fold Enrichment | FDR |
|---|---|---|---|---|---|---|
| UP_SEQ_FEATURE | REPEAT:Solcar 1 | 3 | 0.192659 | Q791V5, Q9QXX4, Q8BH59 | 3.71 | 0.99251497 |
| UP_SEQ_FEATURE | REPEAT:Solcar 2 | 3 | 0.192659 | Q791V5, Q9QXX4, Q8BH59 | 3.71 | 0.99251497 |
| INTERPRO | IPR018108:Mitochondrial_sb/sol_carrier | 3 | 0.214704 | Q791V5, Q9QXX4, Q8BH59 | 3.45 | 0.98905908 |
| INTERPRO | IPR023395:Mt_carrier_dom_sf | 3 | 0.214704 | Q791V5, Q9QXX4, Q8BH59 | 3.45 | 0.98905908 |
| GOTERM_BP_DIRECT | GO:1902600~proton transmembrane transport | 3 | 0.479981 | Q3TBT3, Q9QXX4, Q8BH59 | 1.86 | 0.97339983 |
| GOTERM_BP_DIRECT | GO:0055085~transmembrane transport | 4 | 0.486627 | P36371, P21958, A1L314, Q9QXX4 | 1.53 | 0.97339983 |
| UP_SEQ_FEATURE | TRANSMEM:Helical; Name=8 | 3 | 0.527567 | P36371, P21958, Q78IQ7 | 1.70 | 0.99251497 |
| UP_SEQ_FEATURE | TRANSMEM:Helical; Name=3 | 7 | 0.983731 | Q3TBT3, P36371, P21958, Q78IQ7, Q791V5, Q9QXX4, Q8BH59 | 0.57 | 0.99251497 |
| UP_SEQ_FEATURE | TRANSMEM:Helical; Name=4 | 7 | 0.98459 | Q3TBT3, P36371, P21958, Q78IQ7, Q791V5, Q9QXX4, Q8BH59 | 0.57 | 0.99251497 |
| UP_SEQ_FEATURE | TRANSMEM:Helical; Name=1 | 7 | 0.986333 | Q3TBT3, P36371, P21958, Q78IQ7, Q791V5, Q9QXX4, Q8BH59 | 0.56 | 0.99251497 |
| UP_SEQ_FEATURE | TRANSMEM:Helical; Name=2 | 7 | 0.987063 | Q3TBT3, P36371, P21958, Q78IQ7, Q791V5, Q9QXX4, Q8BH59 | 0.56 | 0.99251497 |
| UP_SEQ_FEATURE | TRANSMEM:Helical; Name=6 | 6 | 0.992054 | P36371, P21958, Q78IQ7, Q791V5, Q9QXX4, Q8BH59 | 0.51 | 0.99251497 |
| UP_SEQ_FEATURE | TRANSMEM:Helical; Name=5 | 6 | 0.992602 | P36371, P21958, Q78IQ7, Q791V5, Q9QXX4, Q8BH59 | 0.51 | 0.99260153 |
| UP_SEQ_FEATURE | TRANSMEM:Helical; Name=7 | 3 | 0.999759 | P36371, P21958, Q78IQ7 | 0.28 | 0.99975891 |

Annotation Cluster 82 — Enrichment Score: 0.17698945109531977

| Category | Term | Count | PValue | Genes | Fold Enrichment | FDR |
|---|---|---|---|---|---|---|

| | | | | | | |
|---|---|---|---|---|---|---|
| GOTERM_MF_DIRECT | GO:0003677~DNA binding | 25 | 0.213714 | Q9CXY6, P09405, Q91VR5, Q9JKB3, Q8BVK9, Q9WVL2, Q62189, Q91XB0, Q99J87, P42225, Q6A028, Q52KI8, Q9Z2U2, A2BH40, Q3TGW2, E9QAM5, Q61881, P62960, O35892, P25976, Q5NC05, P17918, Q8BMQ2, E9PVX6, P97481 | 1.24 | 0.83211475 |
| UP_KW_MOLECULAR_FUNCTION | KW-0238~DNA-binding | 28 | 0.922109 | Q9CXY6, P09405, Q91VR5, Q9JKB3, Q8BVK9, Q9WVL2, P97789, P0DOV2, P42225, P49718, Q6A028, D3YXK2, Q52KI8, Q9Z2U2, A2BH40, Q9DBR1, Q8CGK3, Q9R002, E9QAM5, Q61881, P62960, O35892, P25976, Q5NC05, P17918, Q8BMQ2, E9PVX6, P97481 | 0.82 | 0.92210928 |
| UP_KW_BIOLOGICAL_PROCESS | KW-0804~Transcription | 31 | 0.996195 | Q9CXY6, Q922L6, Q91VR5, Q61656, Q9JKB3, Q8BVK9, Q501J6, Q9WVL2, P0DOV2, P42225, D3YXK2, Q9Z2U2, A2BH40, Q9DBR1, Q60809, Q91WQ5, Q60972, Q9WTM5, E9QAM5, Q8BRN9, Q9JIK5, P62960, O35892, P25976, Q8C166, Q5NC05, Q9QXG4, P63017, O35658, Q8BMQ2, P97481 | 0.69 | 0.99619473 |
| UP_KW_BIOLOGICAL_PROCESS | KW-0805~Transcription regulation | 29 | 0.99789 | Q9CXY6, Q922L6, Q91VR5, Q61656, Q9JKB3, Q8BVK9, Q501J6, Q9WVL2, P0DOV2, P42225, D3YXK2, Q9Z2U2, A2BH40, Q9DBR1, Q60809, Q91WQ5, Q60972, Q9WTM5, E9QAM5, Q8BRN9, P62960, O35892, P25976, Q8C166, Q5NC05, Q9QXG4, P63017, O35658, P97481 | 0.66 | 0.99789026 |
| Annotation Cluster 83 | Enrichment Score: 0.13736918562378367 | | | | | |
| Category | Term | Count | PValue | Genes | Fold Enrichment | FDR |
| UP_SEQ_FEATURE | DOMAIN:PDZ | 3 | 0.705985 | Q9QZQ1, P46062, Q9JLB0 | 1.22 | 0.99251497 |
| INTERPRO | IPR001478:PDZ | 3 | 0.72286 | Q9QZQ1, P46062, Q9JLB0 | 1.18 | 0.98905908 |
| INTERPRO | IPR036034:PDZ_sf | 3 | 0.732794 | Q9QZQ1, P46062, Q9JLB0 | 1.16 | 0.98905908 |
| SMART | SM00228:PDZ | 3 | 0.754556 | Q9QZQ1, P46062, Q9JLB0 | 1.11 | 1 |
| Annotation Cluster 84 | Enrichment Score: 0.13390069714518307 | | | | | |
| Category | Term | Count | PValue | Genes | Fold Enrichment | FDR |
| UP_SEQ_FEATURE | REPEAT:3 | 3 | 0.665491 | P09405, Q62433, P0DOV2 | 1.32 | 0.99251497 |
| UP_SEQ_FEATURE | REPEAT:1 | 3 | 0.769126 | P09405, Q62433, P0DOV2 | 1.08 | 0.99251497 |
| UP_SEQ_FEATURE | REPEAT:2 | 3 | 0.774744 | P09405, Q62433, P0DOV2 | 1.06 | 0.99251497 |
| Annotation Cluster 85 | Enrichment Score: 0.1127384295359617 | | | | | |
| Category | Term | Count | PValue | Genes | Fold Enrichment | FDR |
| INTERPRO | IPR011993:PH-like_dom_sf | 8 | 0.562478 | Q8R1F1, P26041, Q6A028, Q9D1J1, Q8CIH5, Q9JM90, P48193, Q8BVL3 | 1.13 | 0.98905908 |
| UP_SEQ_FEATURE | DOMAIN:PH | 4 | 0.815937 | Q8R1F1, Q6A028, Q8CIH5, Q9JM90 | 0.91 | 0.99251497 |
| INTERPRO | IPR001849:PH_domain | 4 | 0.81669 | Q8R1F1, Q6A028, Q8CIH5, Q9JM90 | 0.91 | 0.98905908 |
| SMART | SM00233:PH | 3 | 0.944554 | Q6A028, Q8CIH5, Q9JM90 | 0.66 | 1 |
| Annotation Cluster 86 | Enrichment Score: 0.10075908241253335 | | | | | |
| Category | Term | Count | PValue | Genes | Fold Enrichment | FDR |
| GOTERM_MF_DIRECT | GO:0020037~heme binding | 3 | 0.699465 | O55022, P22437, Q9CXV1 | 1.23 | 0.94211823 |
| UP_SEQ_FEATURE | BINDING:axial binding residue | 3 | 0.726303 | O55022, P22437, Q9CXV1 | 1.17 | 0.99251497 |
| UP_KW_LIGAND | KW-0408~Iron | 6 | 0.981385 | O55022, Q9CQA3, P22437, Q9CXV1, Q99KI0, Q8WTY4 | 0.57 | 1 |
| Annotation Cluster 87 | Enrichment Score: 0.07355478237336792 | | | | | |
| Category | Term | Count | PValue | Genes | Fold Enrichment | FDR |
| UP_KW_DOMAIN | KW-0245~EGF-like domain | 4 | 0.762729 | P21956, O88839, P22437, O70309 | 1.00 | 0.82604209 |
| UP_SEQ_FEATURE | DOMAIN:EGF-like | 3 | 0.859019 | P21956, O88839, P22437 | 0.87 | 0.99251497 |
| INTERPRO | IPR000742:EGF-like_dom | 3 | 0.918254 | P21956, O88839, P22437 | 0.73 | 0.98905908 |
| Annotation Cluster 88 | Enrichment Score: 0.03846543673059631 | | | | | |
| Category | Term | Count | PValue | Genes | Fold Enrichment | FDR |
| GOTERM_BP_DIRECT | GO:0006357~regulation of transcription by RNA polyme | 17 | 0.820802 | Q91WQ5, Q61656, Q8BVK9, Q501J6, Q9WTM5, Q9WVL2, Q9CPU0, Q8BRN9, P0DOV2, P42225, P45878, D3YXK2, Q9Z2U2, A2BH40, Q9JJ80, O88286, P97481 | 0.87 | 0.97339983 |
| GOTERM_MF_DIRECT | GO:0000981~DNA-binding transcription factor activity, | 9 | 0.95424 | P42225, Q8BVK9, O35892, Q9Z2U2, Q9WVL2, Q9JJ80, Q8BRN9, O88286, P97481 | 0.68 | 0.95423955 |
| GOTERM_MF_DIRECT | GO:0000978~RNA polymerase II cis-regulatory region s | 8 | 0.978831 | P42225, Q60972, D3YXK2, Q9WTM5, Q9WVL2, Q8BRN9, O88286, P97481 | 0.61 | 0.97883097 |
| Annotation Cluster 89 | Enrichment Score: 0.012379735021113886 | | | | | |
| Category | Term | Count | PValue | Genes | Fold Enrichment | FDR |
| UP_SEQ_FEATURE | CARBOHYD:N-linked (GlcNAc...) asparagine | 54 | 0.897033 | P30204, P06339, Q78IQ7, O70370, P97821, P18242, Q07797, A1L314, P98086, O35405, O89023, Q9Z0M5, P26151, P17439, Q8R180, P15379, P21855, P01902, O35114, P01901, Q9DCQ7, P32261, P01900, P29416, P10810, Q6R5N8, P20060, Q8R2E9, P10852, O88839, P22437, Q9WVJ3, Q9Z0F8, Q8K1T1, Q6ZQM8, Q8R2Q8, P05555, Q99P91, Q9EQH2, P03975, P11438, P97333, Q62351, P42082, P14426, Q9D8V0, O09159, P21956, Q8C129, Q8VIM6, O70309, Q9ERE7, O88188, Q8BG07 | 0.88 | 0.99251497 |
| UP_KW_DOMAIN | KW-0732~Signal | 58 | 0.994658 | P06339, Q78IQ7, O70370, P97821, P18242, Q07797, A1L314, P98086, O89023, Q9Z0M5, P26151, P17439, Q8R180, P15379, O55022, P01902, P01901, Q9DCQ7, P32261, P01900, P09103, P05064, P29416, Q9CWG1, Q6PHZ2, P10810, Q6R5N8, P20060, Q8R2E9, P45878, O88839, P36371, P22437, Q9WVJ3, Q9Z0F8, Q8K1T1, Q6ZQM8, Q8CGK3, P05555, P57759, Q99P91, Q9EQH2, P03975, P11438, P08905, P97333, P42082, P14426, O09159, P21956, Q8VIM6, O70309, P83741, Q9ERE7, Q9JK81, O88188, Q91YW3 | 0.78 | 0.99465755 |

| | | | | | | |
|---|---|---|---|---|---|---|
| UP_KW_PTM | KW-1015~Disulfide bond | 54 | 0.999998 | P30204, P06339, Q78IQ7, O70370, P97821, P18242, Q07797, P70227, Q9WTP6, P62075, A1L314, P98086, O35405, O89023, Q9DBC7, P26151, P17439, Q8R180, P15379, P21855, P01902, O35114, P01901, P32261, P01900, P09103, P29416, Q8K411, P10810, P20060, Q8R2E9, P10852, Q9CPU0, O88839, P22437, Q9Z0F8, Q8R2Q8, P05555, Q9EQH2, P03975, P11438, P08905, P97333, Q9WU84, Q62351, P42082, P14426, O09159, P21956, O70309, P38060, O88188, Q8BG07, Q91YW3 | 0.62 | 0.99999776 |
| UP_KW_PTM | KW-0325~Glycoprotein | 60 | 1 | P30204, P06339, Q78IQ7, O70370, P97821, P18242, Q07797, P12382, A1L314, P20152, P98086, O35405, O89023, Q9Z0M5, P26151, P17439, Q61881, Q8R180, P15379, P21855, O55022, P01902, O35114, P01901, Q9DCQ7, P32261, P01900, P29416, P10810, Q6R5N8, P20060, Q8R2E9, P10852, P48193, O88839, P22437, Q9WVJ3, Q9Z0F8, Q8K1T1, Q6ZQM8, Q8R2Q8, P05555, Q99P91, Q9EQH2, P03975, P11438, P97333, Q62351, P42082, P14426, Q9D8V0, O09159, P21956, Q8C129, Q8VIM6, O70309, Q9ERE7, O88188, Q9WUA3, Q8BG07 | 0.56 | 1 |
| Annotation Cluster 90 | Enrichment Score: 0.0037064470276916093 | | | | | |
| Category | Term | Count | PValue | Genes | Fold Enrichment | FDR |
| UP_SEQ_FEATURE | TOPO_DOM:Cytoplasmic | 45 | 0.960767 | P30204, P06339, Q80X80, Q9CQW9, Q6R5N8, Q8BYI6, Q9CZW5, Q78IQ7, P10852, Q3TZZ7, P70227, O88839, P36371, P63024, Q9Z0F8, Q9QUJ7, Q8K1T1, Q8R2Q8, O35405, P05555, Q3TBT3, Q99P91, Q9EQH2, P11438, P26151, P97333, Q99J93, Q791V5, Q6PA06, Q62351, P42082, P14426, Q99P72, Q9D8V0, P15379, Q8C129, P21855, O55022, O35114, P01902, P01901, P21958, P01900, O70309, O88983, Q8BG07 | 0.81 | 0.99251497 |
| UP_KW_DOMAIN | KW-1133~Transmembrane helix | 68 | 0.998507 | P30204, P06339, Q9CZW5, Q78IQ7, Q3TZZ7, P70227, Q9DBL1, A1L314, Q8BH59, O35405, Q9CXV1, P26151, O35682, Q99P72, Q8BVL3, P15379, Q58A65, P21855, O35435, O55022, P01902, O35114, P01901, P01900, Q3UMY5, Q9CWG1, Q9QXG4, Q62425, Q80X80, Q9CQW9, Q6R5N8, Q8BYI6, P10852, Q9QXX4, P48193, Q9CPQ1, P63024, O88839, P36371, Q9Z0F8, Q9QUJ7, Q8K1X4, Q8K1T1, Q6ZQM8, Q8R2Q8, Q9QYB1, Q9CZP5, P05555, Q3TBT3, Q99P91, Q9EQH2, P11438, P97333, Q99J93, Q791V5, Q6PA06, Q62351, P42082, P14426, Q9CQV7, Q9D8V0, O09159, Q8C129, P21958, Q6DVA0, O70309, Q78IK4, O88983, Q8BG07 | 0.77 | 0.99850729 |
| UP_SEQ_FEATURE | TOPO_DOM:Extracellular | 25 | 0.999012 | P30204, P06339, Q9CQW9, Q6R5N8, Q78IQ7, P10852, P70227, O88839, Q9Z0F8, Q8K1T1, Q8R2Q8, P05555, Q99P91, P26151, P97333, Q99J93, Q62351, P42082, P14426, P15379, Q8C129, P21855, P01902, P01901, P01900, O70309 | 0.61 | 0.99901228 |
| UP_KW_DOMAIN | KW-0812~Transmembrane | 71 | 0.999833 | P30204, P06339, Q9CZW5, Q78IQ7, Q3TZZ7, P70227, Q9DBL1, Q8BGH2, A1L314, Q8BH59, O35405, Q9CXV1, P26151, O35682, Q99P72, Q8BVL3, P15379, Q58A65, P21855, O35435, O55022, P01902, O35114, P01901, P01900, Q3UMY5, Q9CWG1, Q9QXG4, Q62425, Q80X80, Q9CQW9, Q6R5N8, Q8BYI6, P10852, Q9QXX4, P48193, Q9CPQ1, P63024, O88839, P36371, Q99388, Q9Z0F8, Q9QUJ7, Q8K1X4, Q8K1T1, Q6ZQM8, Q8R2Q8, Q9QYB1, Q9CZP5, P05555, Q3TBT3, Q99P91, Q9EQH2, Q60930, P11438, P97333, Q99J93, Q791V5, Q6PA06, Q62351, P42082, P14426, Q9CQV7, Q9D8V0, O09159, Q8C129, P21958, Q6DVA0, O70309, Q78IK4, O88983, Q8BG07 | 0.74 | 0.99983307 |
| UP_SEQ_FEATURE | TRANSMEM:Helical | 59 | 1 | P30204, P06339, Q9CZW5, Q78IQ7, Q3TZZ7, P70227, Q9DBL1, A1L314, O35405, Q9CXV1, P26151, O35682, Q99P72, Q8BVL3, P15379, Q58A65, P21855, O35435, O55022, P01902, O35114, P01901, P01900, Q3UMY5, Q9CWG1, Q9QXG4, Q62425, Q80X80, Q9CQW9, Q6R5N8, Q8BYI6, P10852, P48193, Q9CPQ1, O88839, P36371, Q99388, Q9Z0F8, Q8K1X4, Q8K1T1, Q6ZQM8, Q9CZP5, P05555, Q3TBT3, Q99P91, P11438, P97333, Q99J93, Q6PA06, Q62351, P42082, P14426, Q9CQV7, Q9D8V0, O09159, Q8C129, P21958, Q6DVA0, O70309, Q78IK4 | 0.54 | 1 |
| Annotation Cluster 91 | Enrichment Score: 1.1625924142684775E-4 | | | | | |
| Category | Term | Count | PValue | Genes | Fold Enrichment | FDR |
| UP_KW_BIOLOGICAL_PROCESS | KW-0221~Differentiation | 7 | 0.999225 | Q7TMB8, P97333, Q8C166, O08539, P42208, P60766, P97481 | 0.43 | 0.99922547 |
| UP_KW_MOLECULAR_FUNCTION | KW-0217~Developmental protein | 7 | 0.999986 | Q6P5H2, Q7TMB8, P97333, Q8K2Q9, O08539, P97481, P0DOV2 | 0.33 | 0.99998587 |
| UP_KW_MOLECULAR_FUNCTION | KW-9996~Developmental protein | 7 | 0.999986 | Q6P5H2, Q7TMB8, P97333, Q8K2Q9, O08539, P97481, P0DOV2 | 0.33 | 0.99998587 |
