## Supplemental Table 7 for "Effects of polystyrene and polylactide nanoparticles on macrophages under a repeated exposure mode"

Supplementary Table 7: Mitochondrial proteins modulated in response to PS or PLA particles
Color code: purple, proteins modulated in response to both particles,
blue, proteins modulated in response to PS particles only
green, proteins modulated in response to PLA particles only
black, unmodulated proteins related to the selected proteins

| accession | gene_name | description | U-PLA | U-PS | ratio PLA | ratio PS |
|---|---|---|---|---|---|---|
| O08528 | Hk2 | Hexokinase-2 | 0 | 3 | 1.25 | 0.94 |
| O08749 | Dld | Dihydrolipoyl dehydrogenase, mitochondrial | 10 | 1 | 0.96 | 0.79 |
| O09044 | Snap23 | Synaptosomal-associated protein 23 | 3 | 1 | 0.87 | 0.84 |
| O35435 | Dhodh | Dihydroorotate dehydrogenase (quinone), mitochondrial | 1 | 2 | 0.74 | 0.81 |
| O35658 | C1qbp | Complement component 1 Q subcomponent-binding protein, mitochondrial | 2 | 4 | 1.39 | 1.23 |
| O55022 | Pgrmc1 | Membrane-associated progesterone receptor component 1 | 0 | 3 | 1.13 | 1.12 |
| O88986 | Gcat | 2-amino-3-ketobutyrate coenzyme A ligase, mitochondrial | 6 | 0 | 1.21 | 1.41 |
| P10639 | Txn | Thioredoxin | 6 | 1 | 1.32 | 1.31 |
| P11352 | Gpx1 | Glutathione peroxidase 1 | 2 | 1 | 0.80 | 0.73 |
| P11928 | Oas1a | 2'-5'-oligoadenylate synthase 1A | 2 | 4 | 0.50 | 0.70 |
| P17710 | Hk1 | Hexokinase-1 | 0 | 5 | 1.17 | 1.03 |
| P29452 | Casp1 | Caspase-1 | 0 | 0 | 0.83 | 0.89 |
| P29758 | Oat | Ornithine aminotransferase, mitochondrial | 2 | 0 | 1.13 | 1.22 |
| P31750 | Akt1 | RAC-alpha serine/threonine-protein kinase | 7 | 1 | 0.96 | 1.71 |
| P36552 | Cpox | Oxygen-dependent coproporphyrinogen-III oxidase, mitochondrial | 1 | 1 | 1.79 | 1.91 |
| P38060 | Hmgcl | Hydroxymethylglutaryl-CoA lyase, mitochondrial | 2 | 3 | 1.22 | 1.20 |
| P38647 | Hspa9 | Stress-70 protein, mitochondrial | 2 | 4 | 1.08 | 1.05 |
| P45952 | Acadm | Medium-chain specific acyl-CoA dehydrogenase, mitochondrial | 2 | 7 | 0.89 | 0.98 |
| P47738 | Aldh2 | Aldehyde dehydrogenase, mitochondrial | 0 | 0 | 1.12 | 1.24 |
| P50171 | Hsd17b8 | Estradiol 17-beta-dehydrogenase 8 | 2 | 2 | 1.37 | 1.30 |
| P54987 | Acod1 | Cis-aconitate decarboxylase | 0 | 3 | 0.46 | 0.58 |
| P57716 | Ncstn | Nicastrin | 7 | 0 | 0.92 | 0.88 |
| P59017 | Bcl2l13 | Bcl-2-like protein 13 | 8 | 1 | 1.16 | 1.32 |
| P60603 | Romo1 | Reactive oxygen species modulator 1 | 5 | 0 | 1.46 | 2.00 |
| P62908 | Rps3 | 40S ribosomal protein S3 | 5 | 1 | 0.93 | 0.92 |
| P85094 | Isoc2a | Isochorismatase domain-containing protein 2A | 5 | 0 | 1.14 | 1.41 |
| P97333 | Nrp1 | Neuropilin-1 | 1 | 7 | 0.71 | 0.85 |
| P97494 | Gclc | Glutamate--cysteine ligase catalytic subunit | 7 | 0 | 1.17 | 1.35 |
| Q2TBE6 | Pi4k2a | Phosphatidylinositol 4-kinase type 2-alpha | 0 | 7 | 0.75 | 0.93 |
| Q3TBT3 | Tmem173 | Stimulator of interferon genes protein | 1 | 4 | 0.70 | 0.82 |
| Q3TRM8 | Hk3 | Hexokinase-3 | 2 | 0 | 0.91 | 0.88 |

| | | | | | | |
|---|---|---|---|---|---|---|
| Q60766 | Irgm1 | Immunity-related GTPase family M protein 1 | 0 | 0 | 0.60 | 0.61 |
| Q62465 | Vat1 | Synaptic vesicle membrane protein VAT-1 homolog | 0 | 0 | 1.61 | 1.58 |
| Q64521 | Gpd2 | Glycerol-3-phosphate dehydrogenase, mitochondrial | 4 | 0 | 1.12 | 1.17 |
| Q791T5 | Mtch1 | Mitochondrial carrier homolog 1 | 6 | 0 | 4.02 | 8.15 |
| Q791V5 | Mtch2 | Mitochondrial carrier homolog 2 | 0 | 1 | 1.15 | 1.21 |
| Q7TMY8 | Huwe1 | E3 ubiquitin-protein ligase HUWE1 | 6 | 1 | 1.29 | 1.33 |
| Q811U4 | Mfn1 | Mitofusin-1 | 4 | 1 | 1.34 | 1.43 |
| Q8BGH2 | Samm50 | Sorting and assembly machinery component 50 homolog | 2 | 5 | 1.08 | 1.05 |
| Q8BH04 | Pck2 | Phosphoenolpyruvate carboxykinase [GTP], mitochondrial | 1 | 10 | 1.11 | 1.00 |
| Q8BH59 | Slc25a12 | Calcium-binding mitochondrial carrier protein Aralar1 | 2 | 0 | 1.11 | 1.20 |
| Q8BH86 | Dglucy | D-glutamate cyclase, mitochondrial | 11 | 1 | 1.01 | 1.22 |
| Q8BHG1 | Nrdc | Nardilysin | 9 | 0 | 0.99 | 0.82 |
| Q8BMS1 | Hadha | Trifunctional enzyme subunit alpha, mitochondrial | 5 | 0 | 1.08 | 1.21 |
| Q8BTX9 | Hsdl1 | Inactive hydroxysteroid dehydrogenase-like protein 1 | 2 | 0 | 0.80 | 0.79 |
| Q8BVI4 | Qdpr | Dihydropteridine reductase | 8 | 1 | 1.02 | 1.19 |
| Q8BVN4 | Mterf4 | Transcription termination factor 4, mitochondrial 2 | 11 | 0 | 1.01 | 1.23 |
| Q8BY71 | Hat1 | Histone acetyltransferase type B catalytic subunit | 0 | 7 | 0.72 | 0.92 |
| Q8CG72 | Adprhl2 | Poly(ADP-ribose) glycohydrolase ARH3 | 0 | 9 | 1.24 | 1.03 |
| Q8CGK3 | Lonp1 | Lon protease homolog, mitochondrial | 0 | 1 | 1.25 | 1.14 |
| Q8JZU0 | Nudt13 | Nucleoside diphosphate-linked moiety X motif 13 | 11 | 1 | 0.73 | 1.14 |
| Q8K1J6 | Trnt1 | CCA tRNA nucleotidyltransferase 1, mitochondrial | 2 | 9 | 0.82 | 0.96 |
| Q8K411 | Pitrm1 | Presequence protease, mitochondrial | 2 | 4 | 1.13 | 1.10 |
| Q8QZT1 | Acat1 | Acetyl-CoA acetyltransferase, mitochondrial | 4 | 0 | 1.08 | 1.18 |
| Q8VCF0 | Mavs | Mitochondrial antiviral-signaling protein | 8 | 1 | 1.04 | 1.25 |
| Q8VCW8 | Acsf2 | Acyl-CoA synthetase family member 2, mitochondrial | 2 | 2 | 1.44 | 1.47 |
| Q8WTY4 | Ciapin1 | Anamorsin | 2 | 5 | 1.34 | 1.15 |
| Q91VJ1 | Aim2 | Interferon-inducible protein AIM2 | 12 | 0 | 1.00 | 1.15 |
| Q91VR5 | Ddx1 | ATP-dependent RNA helicase DDX1 | 2 | 0 | 1.10 | 1.10 |
| Q91YM4 | Tbrg4 | FAST kinase domain-containing protein 4 | 0 | 4 | 0.25 | 0.49 |
| Q921F2 | Tardbp | TAR DNA-binding protein 43 | 5 | 1 | 1.15 | 1.18 |
| Q922Q1 | Marc2 | Mitochondrial amidoxime reducing component 2 | 7 | 1 | 1.06 | 1.31 |
| Q99JR1 | Sfxn1 | Sideroflexin-1 | 7 | 1 | 0.94 | 1.13 |
| Q99JY0 | Hadhb | Trifunctional enzyme subunit beta, mitochondrial | 8 | 0 | 1.04 | 1.26 |
| Q99KR3 | Lactb2 | Endoribonuclease LACTB2 | 1 | 4 | 1.11 | 1.08 |
| Q99LX0 | Park7 | Protein/nucleic acid deglycase DJ-1 | 0 | 2 | 1.22 | 1.15 |
| Q9CZE3 | Rab32 | Ras-related protein Rab-32 | 2 | 2 | 0.50 | 0.57 |
| Q9CZP5 | Bcs1l | Mitochondrial chaperone BCS1 | 2 | 6 | 1.22 | 1.11 |

| | | | | | | |
|---|---|---|---|---|---|---|
| Q9D0G0 | Mrps30 | 28S ribosomal protein S30, mitochondrial | 4 | 0 | 1.86 | 1.97 |
| Q9D0K2 | Oxct1 | Succinyl-CoA:3-ketoacid coenzyme A transferase 1, mitochondrial | 5 | 0 | 1.13 | 1.28 |
| Q9D0L7 | Armc10 | Armadillo repeat-containing protein 10 | 9 | 0 | 1.09 | 1.35 |
| Q9DBF1 | Aldh7a1 | Alpha-aminoadipic semialdehyde dehydrogenase | 7 | 0 | 1.12 | 1.24 |
| Q9DBL1 | Acadsb | Short/branched chain specific acyl-CoA dehydrogenase, mitochondrial | 0 | 2 | 1.31 | 1.22 |
| Q9JIK5 | Ddx21 | Nucleolar RNA helicase 2 | 1 | 4 | 0.82 | 0.89 |
| Q9JIY5 | Htra2 | Serine protease HTRA2, mitochondrial | 11 | 0 | 1.07 | 2.23 |
| Q9JK81 | Myg1 | UPF0160 protein MYG1, mitochondrial | 0 | 0 | 0.80 | 0.77 |
| Q9JM90 | Stap1 | Signal-transducing adaptor protein 1 | 0 | 0 | 1.52 | 1.47 |
| Q9QXX4 | Slc25a13 | Calcium-binding mitochondrial carrier protein Aralar2 | 0 | 0 | 1.17 | 1.21 |
| Q9QYB1 | Clic4 | Chloride intracellular channel protein 4 | 0 | 0 | 1.28 | 1.22 |
| Q9WTP6 | Ak2 | Adenylate kinase 2, mitochondrial | 0 | 0 | 1.16 | 1.12 |
| Q99LC3 | Ndufa10 | NADH dehydrogenase [ubiquinone] 1 alpha subcomplex subunit 10, mitochondrial | 10 | 6 | 0.97 | 1.04 |
| Q9D8B4 | Ndufa11 | NADH dehydrogenase [ubiquinone] 1 alpha subcomplex subunit 11 | 12 | 2 | 1.03 | 1.17 |
| Q7TMF3 | Ndufa12 | NADH dehydrogenase [ubiquinone] 1 alpha subcomplex subunit 12 | 12 | 10 | 1.09 | 0.99 |
| Q9ERS2 | Ndufa13 | NADH dehydrogenase [ubiquinone] 1 alpha subcomplex subunit 13 | 8 | 9 | 1.10 | 1.03 |
| Q9CQ75 | Ndufa2 | NADH dehydrogenase [ubiquinone] 1 alpha subcomplex subunit 2 | 10 | 2 | 0.94 | 0.88 |
| Q62425 | Ndufa4 | Cytochrome c oxidase subunit NDUFA4 | 0 | 0 | 0.73 | 0.50 |
| Q9CPP6 | Ndufa5 | NADH dehydrogenase [ubiquinone] 1 alpha subcomplex subunit 5 | 7 | 4 | 1.87 | 1.94 |
| Q9DCJ5 | Ndufa8 | NADH dehydrogenase [ubiquinone] 1 alpha subcomplex subunit 8 | 5 | 9 | 0.90 | 0.96 |
| Q9DC69 | Ndufa9 | NADH dehydrogenase [ubiquinone] 1 alpha subcomplex subunit 9, mitochondrial | 12 | 8 | 0.89 | 0.97 |
| Q9DCS9 | Ndufb10 | NADH dehydrogenase [ubiquinone] 1 beta subcomplex subunit 10 | 5 | 9 | 0.70 | 0.96 |
| O09111 | Ndufb11 | NADH dehydrogenase [ubiquinone] 1 beta subcomplex subunit 11, mitochondrial | 10 | 0 | 2.05 | 10.25 |
| Q9CQZ6 | Ndufb3 | NADH dehydrogenase [ubiquinone] 1 beta subcomplex subunit 3 | 10 | 9 | 1.27 | 0.83 |
| Q9CQH3 | Ndufb5 | NADH dehydrogenase [ubiquinone] 1 beta subcomplex subunit 5, mitochondrial | 10 | 5 | 70423.52 | ### |
| Q9CQ54 | Ndufc2 | NADH dehydrogenase [ubiquinone] 1 subunit C2 | 10 | 10 | 0.94 | 0.97 |
| Q91VD9 | Ndufs1 | NADH-ubiquinone oxidoreductase 75 kDa subunit, mitochondrial | 8 | 7 | 1.02 | 1.02 |
| Q91WD5 | Ndufs2 | NADH dehydrogenase [ubiquinone] iron-sulfur protein 2, mitochondrial | 8 | 8 | 1.06 | 1.02 |
| Q9DCT2 | Ndufs3 | NADH dehydrogenase [ubiquinone] iron-sulfur protein 3, mitochondrial | 9 | 9 | 0.96 | 1.01 |

| | | | | | | |
|---|---|---|---|---|---|---|
| Q9DC70 | Ndufs7 | NADH dehydrogenase [ubiquinone] iron-sulfur protein 7, mitochondrial | 8 | 0 | 1.05 | 1.18 |
| Q8K3J1 | Ndufs8 | NADH dehydrogenase [ubiquinone] iron-sulfur protein 8, mitochondrial | 4 | 0 | 1.16 | 1.27 |
| Q91YT0 | Ndufv1 | NADH dehydrogenase [ubiquinone] flavoprotein 1, mitochondrial | 10 | 6 | 0.98 | 0.92 |
| Q9D6J6 | Ndufv2 | NADH dehydrogenase [ubiquinone] flavoprotein 2, mitochondrial | 4 | 6 | 1.16 | 1.06 |
| Q9CXV1 | Sdhd | Succinate dehydrogenase [ubiquinone] cytochrome b small subunit, mitochondrial | 2 | 0 | 0.61 | 0.63 |
| Q8K2B3 | Sdha | Succinate dehydrogenase [ubiquinone] flavoprotein subunit, mitochondrial | 3 | 6 | 1.19 | 1.07 |
| Q9CQA3 | Sdhb | Succinate dehydrogenase [ubiquinone] iron-sulfur subunit, mitochondrial | 0 | 0 | 1.19 | 1.11 |
| Q9CZB0 | Sdhc | Succinate dehydrogenase cytochrome b560 subunit, mitochondrial | 6 | 7 | 1.15 | 1.04 |
| Q9CWU6 | Uqcc1 | Ubiquinol-cytochrome-c reductase complex assembly factor 1 | 8 | 7 | 1.20 | 1.18 |
| Q8R1I1 | Uqcr10 | Cytochrome b-c1 complex subunit 9 | 10 | 8 | 0.86 | 0.86 |
| Q9CZ13 | Uqcrc1 | Cytochrome b-c1 complex subunit 1, mitochondrial | 5 | 2 | 1.07 | 1.15 |
| Q9DB77 | Uqcrc2 | Cytochrome b-c1 complex subunit 2, mitochondrial | 10 | 8 | 1.02 | 1.02 |
| Q9CR68 | Uqcrfs1 | Cytochrome b-c1 complex subunit Rieske, mitochondrial | 10 | 3 | 1.04 | 1.21 |
| P99028 | Uqcrh | Cytochrome b-c1 complex subunit 6, mitochondrial | 9 | 6 | 0.98 | 1.07 |
| Q7TQ16 | Uqcrq | Cytochrome b-c1 complex subunit 8 | 11.5 | 4 | 1.61 | 3.24 |
| Q8BJ03 | Cox15 | Cytochrome c oxidase assembly protein COX15 homolog | 9 | 8 | 1.14 | 1.37 |
| Q9D7J4 | Cox20 | Cytochrome c oxidase assembly protein COX20, mitochondrial | 4 | 5 | 0.69 | 0.89 |
| P56394 | Cox17 | Cytochrome c oxidase copper chaperone | 4 | 0 | 1.50 | 1.49 |
| P00405 | Mtco2 | Cytochrome c oxidase subunit 2 | 12 | 0 | 0.97 | 0.65 |
| P19783 | Cox4i1 | Cytochrome c oxidase subunit 4 isoform 1, mitochondrial | 5 | 0 | 0.93 | 0.82 |
| P11240 | Cox5a | Cytochrome c oxidase subunit 5A, mitochondrial | 2 | 0 | 0.83 | 0.77 |
| P19536 | Cox5b | Cytochrome c oxidase subunit 5B, mitochondrial | 1 | 0 | 0.81 | 0.65 |
| P56391 | Cox6b1 | Cytochrome c oxidase subunit 6B1 | 6 | 0 | 0.79 | 0.56 |
| Q9CPQ1 | Cox6c | Cytochrome c oxidase subunit 6C | 2 | 0 | 0.78 | 0.56 |
| P48771 | Cox7a2 | Cytochrome c oxidase subunit 7A2, mitochondrial | 6 | 0 | 0.76 | 0.31 |
| Q62425 | Ndufa4 | Cytochrome c oxidase subunit NDUFA4 | 0 | 0 | 0.73 | 0.50 |
| Q9CQQ7 | Atp5pb | ATP synthase F(0) complex subunit B1, mitochondrial | 8 | 5 | 0.93 | 0.94 |
| P56383 | Atp5mc2 | ATP synthase F(0) complex subunit C2, mitochondrial | 6 | 7 | 1.52 | 1.18 |
| Q91YY4 | Atpaf2 | ATP synthase mitochondrial F1 complex assembly factor 2 | 12 | 1 | 0.91 | 6.14 |

| | | | | | | |
|---|---|---|---|---|---|---|
| Q03265 | Atp5f1a | ATP synthase subunit alpha, mitochondrial | 10 | 9 | 0.99 | 1.00 |
| P15999 | Atp5f1a | ATP synthase subunit alpha, mitochondrial | 10 | 8 | 0.97 | 0.98 |
| P56480 | Atp5f1b | ATP synthase subunit beta, mitochondrial | 4 | 2 | 1.09 | 1.13 |
| Q9DCX2 | Atp5pd | ATP synthase subunit d, mitochondrial | 8 | 4 | 0.95 | 0.92 |
| Q9D3D9 | Atp5f1d | ATP synthase subunit delta, mitochondrial | 10 | 9 | 1.03 | 1.02 |
| Q06185 | Atp5me | ATP synthase subunit e, mitochondrial | 11 | 3 | 0.91 | 0.39 |
| P56135 | Atp5mf | ATP synthase subunit f, mitochondrial | 10 | 10 | 0.96 | 1.01 |
| Q91VR2 | Atp5f1c | ATP synthase subunit gamma, mitochondrial | 3 | 6 | 0.89 | 0.98 |
| Q9DB20 | Atp5po | ATP synthase subunit O, mitochondrial | 12 | 7 | 1.00 | 1.02 |
| P97450 | Atp5pf | ATP synthase-coupling factor 6, mitochondrial | 5 | 0 | 0.81 | 0.70 |
| O08734 | Bak1 | Bcl-2 homologous antagonist/killer | 12 | 4 | 1.04 | 0.92 |
| Q07813 | Bax | Apoptosis regulator BAX | 4 | 0 | 1.10 | 1.15 |
| Q8K019 | Bclaf1 | Bcl-2-associated transcription factor 1 | 11 | 4 | 0.98 | 0.94 |
| P53563 | Bcl2l1 | Bcl-2-like protein 1 | 11 | 6 | 1.01 | 0.92 |
| Q62760 | Tomm20 | Mitochondrial import receptor subunit TOM20 homolog | 9 | 8 | 1.28 | 1.07 |
| Q9CPQ3 | Tomm22 | Mitochondrial import receptor subunit TOM22 homolog | 6 | 1 | 1.16 | 1.15 |
| Q9CYG7 | Tomm34 | Mitochondrial import receptor subunit TOM34 | 0 | 0 | 1.20 | 1.20 |
| Q9QYA2 | Tomm40 | Mitochondrial import receptor subunit TOM40 homolog | 8 | 9 | 1.04 | 0.99 |
| B1AXP6 | Tomm5 | Mitochondrial import receptor subunit TOM5 homolog | 9 | 7 | 1.07 | 1.04 |
| Q9CZW5 | Tomm70 | Mitochondrial import receptor subunit TOM70 | 2 | 10 | 0.84 | 1.01 |
| P62075 | Timm13 | Mitochondrial import inner membrane translocase subunit Tim13 | 0 | 2 | 0.72 | 0.83 |
| Q9CQV7 | Dnajc19 | Mitochondrial import inner membrane translocase subunit TIM14 | 2 | 7 | 0.74 | 1.00 |
| O35092 | Timm17a | Mitochondrial import inner membrane translocase subunit Tim17-A | 11 | 4 | 0.70 | 0.85 |
| Q9Z0V7 | Timm17b | Mitochondrial import inner membrane translocase subunit Tim17-B | 7.5 | 7 | 0.00 | 8.01 |
| Q9WTQ8 | Timm23 | Mitochondrial import inner membrane translocase subunit Tim23 | 11 | 8 | 1.05 | 0.95 |
| O35857 | Timm44 | Mitochondrial import inner membrane translocase subunit TIM44 | 9 | 4 | 1.05 | 1.11 |
| Q9D880 | Timm50 | Mitochondrial import inner membrane translocase subunit TIM50 | 7 | 3 | 1.02 | 1.06 |

| | | | | | | |
|---|---|---|---|---|---|---|
| Q9Z2L0 | Vdac1 | Voltage-dependent anion-selective channel protein 1 | 2 | 2 | 1.12 | 1.09 |
| Q60930 | Vdac2 | Voltage-dependent anion-selective channel protein 2 | 2 | 7 | 1.11 | 1.05 |
| Q60931 | Vdac3 | Voltage-dependent anion-selective channel protein 3 | 4 | 6 | 1.19 | 1.12 |
| Q8R404 | Mic13 | MICOS complex subunit MIC13 | 7 | 10 | 1.13 | 0.98 |
| Q9CRB9 | Chchd3 | MICOS complex subunit Mic19 | 10 | 10 | 0.89 | 1.00 |
| Q91VN4 | Chchd6 | MICOS complex subunit Mic25 | 10 | 6 | 1.02 | 1.14 |
| Q9DCZ4 | Apoo | MICOS complex subunit Mic26 | 7 | 7 | 1.47 | 1.03 |
| Q78IK4 | Apool | MICOS complex subunit Mic27 | 2 | 2 | 1.23 | 1.14 |
| Q8CAQ8 | Immt | MICOS complex subunit Mic60 | 6 | 3 | 0.95 | 0.94 |
| P41216 | Acsl1 | Long-chain-fatty-acid--CoA ligase 1 | 6 | 4 | 0.94 | 0.90 |
| Q9CZW4 | Acsl3 | Long-chain-fatty-acid--CoA ligase 3 | 8 | 6 | 0.89 | 1.09 |
| Q9QUJ7 | Acsl4 | Long-chain-fatty-acid--CoA ligase 4 | 0 | 7 | 0.80 | 1.02 |
| Q8JZR0 | Acsl5 | Long-chain-fatty-acid--CoA ligase 5 | 4 | 9 | 0.91 | 1.01 |
| Q9CZU6 | Cs | Citrate synthase, mitochondrial | 9 | 7 | 0.97 | 1.02 |
| Q99KI0 | Aco2 | Aconitate hydratase, mitochondrial | 0 | 0 | 1.12 | 1.12 |
| P54071 | Idh2 | Isocitrate dehydrogenase [NADP], mitochondrial | 4 | 0 | 1.08 | 1.10 |
| Q9D6R2 | Idh3a | Isocitrate dehydrogenase [NAD] subunit alpha, mitochondrial | 9 | 8 | 1.03 | 0.98 |
| Q68FX0 | Idh3B | Isocitrate dehydrogenase [NAD] subunit beta, mitochondrial | 8 | 5 | 1.05 | 1.10 |
| P41565 | Idh3g | Isocitrate dehydrogenase [NAD] subunit gamma 1, mitochondrial | 6 | 0 | 1.11 | 1.23 |
| Q60597 | Ogdh | 2-oxoglutarate dehydrogenase, mitochondrial | 10 | 6 | 0.96 | 1.06 |
| Q9Z2I9 | Sucla2 | Succinate--CoA ligase [ADP-forming] subunit beta, mitochondrial | 2 | 0 | 1.22 | 1.24 |
| Q9WUM5 | Suclg1 | Succinate--CoA ligase [ADP/GDP-forming] subunit alpha, mitochondrial | 3 | 4 | 1.45 | 1.39 |
| Q9Z2I8 | Suclg2 | Succinate--CoA ligase [GDP-forming] subunit beta, mitochondrial | 1 | 0 | 1.18 | 1.20 |
| P08249 | Mdh2 | Malate dehydrogenase, mitochondrial | 11 | 8 | 1.00 | 0.99 |
| Q99KE1 | Me2 | NAD-dependent malic enzyme, mitochondrial | 12 | 9 | 1.00 | 1.00 |
