## Supplemental Table 8 for "Effects of polystyrene and polylactide nanoparticles on macrophages under a repeated exposure mode"

Supplementary Table 8: Proteins involved in central metabolism and modulated in response to PS or PLA particles

Color code: purple, proteins modulated in response to both particles,
blue, proteins modulated in response to PS particles only
green, proteins modulated in response to PLA particles only
black, unmodulated proteins related to the selected proteins

| accession | gene_name | description | U-PLA | U-PS | ratio PLA | ratio PS |
|---|---|---|---|---|---|---|
| P17710 | Hk1 | Hexokinase-1 | 0 | 5 | 1.17 | 1.03 |
| O08528 | Hk2 | Hexokinase-2 | 0 | 3 | 1.25 | 0.94 |
| Q3TRM8 | Hk3 | Hexokinase-3 | 2 | 0 | 0.91 | 0.88 |
| P06745 | Gpi | Glucose-6-phosphate isomerase | 5 | 3 | 1.11 | 1.08 |
| P12382 | Pfkl | ATP-dependent 6-phosphofructokinase, liver type | 2 | 8 | 1.17 | 1.06 |
| P47857 | Pfkm | ATP-dependent 6-phosphofructokinase, muscle type | 4 | 5 | 0.88 | 1.07 |
| Q9WUA3 | Pfkp | ATP-dependent 6-phosphofructokinase, platelet type | 0 | 3 | 1.32 | 1.07 |
| P05064 | Aldoa | Fructose-bisphosphate aldolase A | 1 | 8 | 1.19 | 1.04 |
| P05063 | Aldoc | Fructose-bisphosphate aldolase C | 0 | 6 | 1.20 | 1.05 |
| P17751 | Tpi1 | Triosephosphate isomerase | 3 | 7 | 1.15 | 0.94 |
| P16858 | Gapdh | Glyceraldehyde-3-phosphate dehydrogenase | 3 | 3 | 1.17 | 1.11 |
| P09411 | Pgk1 | Phosphoglycerate kinase 1 | 0 | 7 | 1.19 | 1.06 |
| P09041 | Pgk2 | Phosphoglycerate kinase 2 | 11 | 6 | 1.07 | 1.08 |
| P25113 | Pgam1 | Phosphoglycerate mutase 1 | 4 | 10 | 1.14 | 0.99 |
| P17182 | Eno1 | Alpha-enolase | 0 | 1 | 1.31 | 1.18 |
| P52480 | Pkm | Pyruvate kinase PKM | 4 | 7 | 1.12 | 0.99 |
| P06151 | Ldha | L-lactate dehydrogenase A chain | 3 | 10 | 1.11 | 1.01 |
| Q9D0F9 | Pgm1 | Phosphoglucomutase-1 | 0 | 10 | 1.19 | 1.02 |
| Q7TSV4 | Pgm2 | Phosphoglucomutase-2 | 7 | 1 | 0.62 | 0.58 |
| Q00612 | G6pdx | Glucose-6-phosphate 1-dehydrogenase X | 6 | 0 | 1.04 | 1.09 |
| Q9CQ60 | Pgls | 6-phosphogluconolactonase | 11 | 7 | 1.04 | 1.22 |
| Q9DCD0 | Pgd | 6-phosphogluconate dehydrogenase, decarboxylating | 1 | 0 | 1.15 | 1.23 |
| Q8VEE0 | Rpe | Ribulose-phosphate 3-epimerase | 10 | 10 | 0.90 | 0.88 |
| P47968 | Rpia | Ribose-5-phosphate isomerase | 8 | 8 | 0.90 | 0.97 |
| P40142 | Tkt | Transketolase | 4 | 1 | 1.06 | 1.09 |
| Q93092 | Taldo1 | Transaldolase | 9 | 10 | 0.96 | 0.99 |
| P14152 | Mdh1 | Malate dehydrogenase, cytoplasmic | 0 | 5 | 1.32 | 1.22 |
| P28271 | Aco1 | Cytoplasmic aconitate hydratase | 4 | 7 | 0.92 | 0.94 |

| | | | | | | |
|---|---|---|---|---|---|---|
| Q9QXG4 | Acss2 | Acetyl-coenzyme A synthetase, cytoplasmic | 2 | 6 | 1.14 | 1.07 |
