## Supplemental Table 9 for "Effects of polystyrene and polylactide nanoparticles on macrophages under a repeated exposure mode"

Supplementary Table 9: Lysosomal proteins modulated in response to PS or PLA particles
Color code: purple, proteins modulated in response to both particles,
blue, proteins modulated in response to PS particles only
green, proteins modulated in response to PLA particles only
black, unmodulated proteins related to the selected proteins

| accession | gene_name | description | U-PLA | U-PS | ratio PLA | ratio PS |
|---|---|---|---|---|---|---|
| O08585 | Clta | Clathrin light chain A | 0 | 0 | 0.80 | 0.82 |
| O09043 | Napsa | Napsin-A | 8 | 1 | 1.06 | 0.83 |
| O09159 | Man2b1 | Lysosomal alpha-mannosidase | 0 | 0 | 0.76 | 0.79 |
| O35114 | Scarb2 | Lysosome membrane protein 2 | 0 | 0 | 2.10 | 2.43 |
| O35405 | Pld3 | Phospholipase D3 | 0 | 9 | 0.59 | 0.99 |
| O88983 | Stx8 | Syntaxin-8 | 0 | 0 | 0.75 | 0.85 |
| O89023 | Tpp1 | Tripeptidyl-peptidase 1 | 2 | 4 | 1.21 | 1.15 |
| P01900 | H2-D1 | H-2 class I histocompatibility antigen, D-D alpha chain | 2 | 3 | 0.86 | 0.91 |
| P01901 | H2-K1 | H-2 class I histocompatibility antigen, K-B alpha chain | 0 | 2 | 0.84 | 0.91 |
| P01902 | H2-K1 | H-2 class I histocompatibility antigen, K-D alpha chain | 2 | 2 | 0.83 | 0.84 |
| P06339 | H2-T23 | H-2 class I histocompatibility antigen, D-37 alpha chain | 0 | 2 | 0.84 | 0.89 |
| P07356 | Anxa2 | Annexin A2 | 0 | 1 | 1.24 | 1.07 |
| P10852 | Slc3a2 | 4F2 cell-surface antigen heavy chain | 0 | 2 | 0.70 | 0.69 |
| P11438 | Lamp1 | Lysosome-associated membrane glycoprotein 1 | 2 | 8 | 1.19 | 1.03 |
| P14426 | H2-D1 | H-2 class I histocompatibility antigen, D-K alpha chain | 0 | 2 | 0.83 | 0.89 |
| P14824 | Anxa6 | Annexin A6 | 2 | 10 | 1.16 | 1.01 |
| P17439 | Gba | Glucosylceramidase | 0 | 9 | 0.77 | 0.98 |
| P29416 | Hexa | Beta-hexosaminidase subunit alpha | 2 | 6 | 0.85 | 0.92 |
| P20060 | Hexb | Beta-hexosaminidase subunit beta | 2 | 5 | 0.79 | 0.88 |
| P57716 | Ncstn | Nicastrin | 7 | 0 | 0.92 | 0.88 |
| P58681 | Tlr7 | Toll-like receptor 7 | 6 | 0 | 0.91 | 0.84 |
| P61027 | Rab10 | Ras-related protein Rab-10 | 0 | 0 | 1.10 | 1.10 |
| P63017 | Hspa8 | Heat shock cognate 71 kDa protein | 0 | 3 | 0.91 | 0.94 |
| P97821 | Ctsc | Dipeptidyl peptidase 1 | 0 | 0 | 0.69 | 0.76 |
| Q3TBT3 | Tmem173 | Stimulator of interferon genes prote | 1 | 4 | 0.70 | 0.82 |
| Q58A65 | Spag9 | C-Jun-amino-terminal kinase-interacting protein 4 | 1 | 1 | 1.49 | 1.42 |
| Q60766 | Irgm1 | Immunity-related GTPase family M protein 1 | 0 | 0 | 0.60 | 0.61 |
| Q68FD5 | Cltc | Clathrin heavy chain 1 | 4 | 1 | 0.94 | 0.95 |
| Q80TY0 | Fnbp1 | Formin-binding protein 1 | 2 | 6 | 1.12 | 0.92 |
| Q80V94 | Ap4e1 | AP-4 complex subunit epsilon-1 | 1 | 0 | 0.28 | 0.15 |
| Q8BFR4 | Gns | N-acetylglucosamine-6-sulfatase | 7 | 1 | 0.92 | 0.89 |
| Q8BG07 | Pld4 | Phospholipase D4 | 0 | 0 | 0.74 | 0.77 |
| Q8R5L3 | Vps39 | Vam6/Vps39-like protein | 2 | 8 | 1.34 | 1.16 |
| Q920I9 | Wdr7 | WD repeat-containing protein 7 | 0 | 4 | 0.70 | 0.91 |
| Q99J93 | Ifitm2 | Interferon-induced transmembrane protein 2 | 1 | 1 | 0.60 | 0.59 |
| Q9CQW9 | Ifitm3 | Interferon-induced transmembrane protein 3 | 1 | 1 | 0.59 | 0.55 |

| | | | | | | |
|---|---|---|---|---|---|---|
| Q9D1L9 | Lamtor5 | Ragulator complex protein LAMTOR5 | 0 | 8 | 0.78 | 0.95 |
| Q9JHJ3 | Glmp | Glycosylated lysosomal membrane protein | 12 | 0 | 1.25 | 1.95 |
| Q9WVJ3 | Cpq | Carboxypeptidase Q | 0 | 0 | 0.67 | 0.70 |
| Q9Z0M5 | Lipa | Lysosomal acid lipase/cholesteryl ester hydrolase | 0 | 0 | 2.04 | 3.16 |
| O54715 | Atp6ap1 | V-type proton ATPase subunit S1 | 0 | 3 | 0.75 | 0.79 |
| P50408 | Atp6v1f | V-type proton ATPase subunit F | 3 | 2 | 0.81 | 0.69 |
| P50516 | Atp6v1a | V-type proton ATPase catalytic subunit A | 9 | 6 | 1.02 | 1.05 |
| P50518 | Atp6v1e1 | V-type proton ATPase subunit E 1 | 3 | 9 | 0.73 | 0.95 |
| P51863 | Atp6v0d1 | V-type proton ATPase subunit d 1 | 9 | 6 | 1.03 | 1.03 |
| P57746 | Atp6v1d | V-type proton ATPase subunit D | 5 | 7 | 0.87 | 0.98 |
| P62814 | Atp6v1b2 | V-type proton ATPase subunit B, brain isoform | 12 | 6 | 1.00 | 0.99 |
| P63081 | Atp6v0c | V-type proton ATPase 16 kDa proteolipid subunit | 4 | 4 | 1.36 | 1.26 |
| Q5FVI6 | Atp6v1c1 | V-type proton ATPase subunit C 1 | 2 | 4 | 0.87 | 0.90 |
| Q8BVE3 | Atp6v1h | V-type proton ATPase subunit H | 2 | 1 | 1.21 | 1.20 |
| Q9CR51 | Atp6v1g1 | V-type proton ATPase subunit G 1 | 12 | 10 | 1.00 | 1.01 |
| Q9Z1G4 | Atp6v0a1 | V-type proton ATPase 116 kDa subunit a isoform 1 | 6 | 8 | 1.16 | 0.79 |
| P10605 | Ctsb | Cathepsin B | 7 | 5 | 0.94 | 0.93 |
| P18242 | Ctsd | Cathepsin D | 0 | 0 | 1.29 | 1.26 |
| P06797 | Ctsl | Cathepsin L1 | 6 | 6 | 1.16 | 1.08 |
| O70370 | Ctss | Cathepsin S | 0 | 1 | 0.73 | 0.71 |
| Q9WUU7 | Ctsz | Cathepsin Z | 12 | 10 | 0.99 | 0.99 |
| O35643 | Ap1b1 | AP-1 complex subunit beta-1 | 2 | 5 | 1.20 | 1.14 |
| P22892 | Ap1g1 | AP-1 complex subunit gamma-1 | 10 | 9 | 0.98 | 1.02 |
| O88512 | Ap1g2 | AP-1 complex subunit gamma-like 2 | 0 | 9 | 0.34 | 0.89 |
| P35585 | Ap1m1 | AP-1 complex subunit mu-1 | 7 | 5 | 1.23 | 1.30 |
| P61967 | Ap1s1 | AP-1 complex subunit sigma-1A | 12 | 8 | 1.00 | 1.05 |
| Q64514 | Tpp2 | Tripeptidyl-peptidase 2 | 8 | 5 | 1.06 | 1.13 |
| P17047 | Lamp2 | Lysosome-associated membrane glycoprotein 2 | 9 | 6 | 1.09 | 0.93 |
| Q9ET22 | Dpp7 | Dipeptidyl peptidase 2 | 11 | 4 | 1.04 | 1.16 |
| Q99KK7 | Dpp3 | Dipeptidyl peptidase 3 | 8 | 6 | 1.05 | 1.07 |
| Q8BVG4 | Dpp9 | Dipeptidyl peptidase 9 | 10 | 8 | 0.98 | 1.18 |
| Q6P791 | Lamtor1 | Ragulator complex protein LAMTOR1 | 2 | 2 | 0.69 | 0.78 |
| Q9JHS3 | Lamtor2 | Ragulator complex protein LAMTOR2 | 4 | 7 | 0.87 | 0.88 |
| O88653 | Lamtor3 | Ragulator complex protein LAMTOR3 | 6 | 4 | 2.00 | 1.83 |
