## Supplemental Table 10 for "Effects of polystyrene and polylactide nanoparticles on macrophages under a repeated exposure mode"

Supplementary Table 10: Proteins implicated in the immune response and modulated in response to PS or PLA particles

Color code: purple, proteins modulated in response to both particles,
blue, proteins modulated in response to PS particles only
green, proteins modulated in response to PLA particles only
black, unmodulated proteins related to the selected proteins

| accession | gene_name | description | U-PLA | U-PS | ratio PLA | ratio PS |
|---|---|---|---|---|---|---|
| P54987 | Acod1 | Cis-aconitate decarboxylase | 0 | 3 | 0.46 | 0.58 |
| Q91VJ1 | Aim2 | Interferon-inducible protein AIM2 | 12 | 0 | 1.00 | 1.15 |
| Q8R2Q8 | Bst2 | Bone marrow stromal antigen 2 | 1 | 2 | 0.64 | 0.68 |
| P98086 | C1qa | Complement C1q subcomponent subunit A | 0 | 3 | 0.54 | 0.67 |
| O35658 | C1qbp | Complement component 1 Q subcomponent-binding protein, mitochondrial | 2 | 4 | 1.39 | 1.23 |
| P10810 | Cd14 | Monocyte differentiation antigen CD14 | 0 | 0 | 0.80 | 0.71 |
| P42082 | Cd86 | T-lymphocyte activation antigen CD86 | 0 | 7 | 1.27 | 1.06 |
| Q91VR5 | Ddx1 | ATP-dependent RNA helicase DDX1 | 2 | 0 | 1.10 | 1.10 |
| Q501J6 | Ddx17 | Probable ATP-dependent RNA helicase DDX17 | 0 | 3 | 0.91 | 0.93 |
| Q9JIK5 | Ddx21 | Nucleolar RNA helicase 2 | 1 | 4 | 0.82 | 0.89 |
| Q62167 | Ddx3x | ATP-dependent RNA helicase DDX3X | 3 | 1 | 0.87 | 0.84 |
| Q99J87 | Dhx58 | Probable ATP-dependent RNA helicase DHX58 | 0 | 2 | 0.00 | 0.24 |
| Q9EQH2 | Erap1 | Endoplasmic reticulum aminopeptidase 1 | 1 | 4 | 0.83 | 0.89 |
| P26151 | Fcgr1 | High affinity immunoglobulin gamma Fc receptor I | 0 | 0 | 0.65 | 0.59 |
| Q9Z0E6 | Gbp2 | Guanylate-binding protein 2 | 1 | 1 | 0.21 | 0.37 |
| Q61107 | Gbp4 | Guanylate-binding protein 4 | 0 | 0 | 0.49 | 0.50 |
| P01900 | H2-D1 | H-2 class I histocompatibility antigen, D-D alpha chain | 2 | 3 | 0.86 | 0.91 |
| P14426 | H2-D1 | H-2 class I histocompatibility antigen, D-K alpha chain | 0 | 2 | 0.83 | 0.89 |
| P01901 | H2-K1 | H-2 class I histocompatibility antigen, K-B alpha chain | 0 | 2 | 0.84 | 0.91 |
| P01902 | H2-K1 | H-2 class I histocompatibility antigen, K-D alpha chain | 2 | 2 | 0.83 | 0.84 |
| P06339 | H2-T23 | H-2 class I histocompatibility antigen, D-37 alpha chain | 0 | 2 | 0.84 | 0.89 |
| P17710 | Hk1 | Hexokinase-1 | 0 | 5 | 1.17 | 1.03 |
| Q9R002 | Ifi202 | Interferon-activable protein 202 | 0 | 0 | 0.00 | 0.00 |
| P0DOV2 | Ifi204 | Interferon-activable protein 204 | 0 | 0 | 0.65 | 0.58 |
| Q99J93 | Ifitm2 | Interferon-induced transmembrane protein 2 | 1 | 1 | 0.60 | 0.59 |
| Q9CQW9 | Ifitm3 | Interferon-induced transmembrane protein 3 | 1 | 1 | 0.59 | 0.55 |
| Q8R4K2 | Irak4 | Interleukin-1 receptor-associated kinase 4 | 1 | 0 | 0.86 | 0.81 |
| Q60766 | Irgm1 | Immunity-related GTPase family M protein 1 | 0 | 0 | 0.60 | 0.61 |
| O88188 | Ly86 | Lymphocyte antigen 86 | 2 | 1 | 0.63 | 0.70 |
| Q8VCF0 | Mavs | Mitochondrial antiviral-signaling protein | 8 | 1 | 1.04 | 1.25 |

| | | | | | | |
|---|---|---|---|---|---|---|
| A1L314 | Mpeg1 | Macrophage-expressed gene 1 protein | 0 | 2 | 0.73 | 0.76 |
| P30204 | Msr1 | Macrophage scavenger receptor types I and II | 0 | 3 | 0.76 | 0.87 |
| Q9QZ08 | Nagk | N-acetyl-D-glucosamine kinase | 2 | 2 | 0.85 | 0.88 |
| O35309 | Nmi | N-myc-interactor | 0 | 0 | 0.80 | 0.80 |
| P11928 | Oas1a | 2'-5'-oligoadenylate synthase 1A | 2 | 4 | 0.50 | 0.70 |
| Q8VI94 | Oasl1 | 2'-5'-oligoadenylate synthase-like protein 1 | 4 | 0 | 0.25 | 0.07 |
| O35405 | Pld3 | Phospholipase D3 | 0 | 9 | 0.59 | 0.99 |
| Q8BG07 | Pld4 | Phospholipase D4 | 0 | 0 | 0.74 | 0.77 |
| P97814 | Pstpip1 | Proline-serine-threonine phosphatase-interacting protein 1 | 0 | 0 | 0.88 | 0.82 |
| E9Q555 | Rnf213 | E3 ubiquitin-protein ligase RNF213 | 2 | 4 | 0.81 | 0.89 |
| P97797 | Sirpa | Tyrosine-protein phosphatase non-receptor type substrate 1 | 6 | 1 | 0.92 | 0.91 |
| Q8BVK9 | Sp110 | Sp110 nuclear body protein | 0 | 0 | 0.82 | 0.71 |
| P21958 | Tap1 | Antigen peptide transporter 1 | 0 | 0 | 0.67 | 0.58 |
| P36371 | Tap2 | Antigen peptide transporter 2 | 0 | 0 | 0.57 | 0.60 |
| Q91YX0 | Themis2 | Protein THEMIS2 | 0 | 0 | 0.60 | 0.71 |
| Q6R5N8 | Tlr13 | Toll-like receptor 13 | 0 | 1 | 0.74 | 0.78 |
| Q9QUN7 | Tlr2 | Toll-like receptor 2 | 5 | 2 | 0.77 | 0.65 |
| Q99MB1 | Tlr3 | Toll-like receptor 3 | 10.5 | 7.5 | 1.69 | 0.49 |
| P58681 | Tlr7 | Toll-like receptor 7 | 6 | 0 | 0.91 | 0.84 |
| Q3TBT3 | Tmem173 | Stimulator of interferon genes protein | 1 | 4 | 0.70 | 0.82 |
| Q61510 | Trim25 | E3 ubiquitin/ISG15 ligase TRIM25 | 2 | 10 | 0.87 | 0.98 |
